# Human Cytomegalovirus Infection of Primary Human Oral Keratinocytes Induces Intermediate Keratinocyte Differentiation and an Altered Innate Immune Response

**DOI:** 10.1101/2025.09.16.676604

**Authors:** Olesea Cojohari, Yuming Cao, Askar Temirbek, Manuel Garber, Andres Colubri, Timothy Kowalik

**Author notes:** These authors contributed equally to this work (Olesea Cojohari, Yuming Cao).

## Abstract

The oral mucosa is central for human cytomegalovirus (HCMV) acquisition. However, the initial infection events in the oral cavity, including how HCMV modulates oral cells to establish infection and spread, are supported by limited experimental data and therefore remain poorly understood. We investigated HCMV infection in cultures of primary human oral keratinocytes (HOKs) as a model to study HCMV infection in the oral mucosa using the TB40E-Gfp and MOLD strains of HCMV. Both viral strains successfully infected and replicated in HOKs as demonstrated by the appearance of cytopathic effects, full expression of viral transcripts, and the shedding of infectious virus. To investigate how HCMV modulates the transcriptome of primary HOKs, we performed single cell RNA sequencing (scRNA-seq) which revealed 11 subtypes of cells. Furthermore, infection resulted in significant changes in host gene expression. HCMV upregulated genes involved in immune responses, cell cycle regulation, cancer-related pathways, neuronal and synaptic functions, metabolism, stress responses, molecular chaperones, and vesicular trafficking which might be critical for viral protein expression and assembly. HCMV also increased the expression of genes related to the extracellular matrix, cell adhesion, microtubules, signal transduction, kinases, and transcriptional regulation. Conversely, HCMV downregulated genes associated with inflammation and immune responses, cell cycle control and apoptosis, cell adhesion and migration, as well as signaling pathways, growth factors, ion channels, transporters, and transcription factors when compared to uninfected cells. These findings suggest that HCMV modulates a wide range of host cellular pathways to create a favorable environment for its replication. HCMV also induced changes in HOK differentiation genes downregulating basal state genes and upregulating intermediate genes and select terminal differentiation genes, indicating HCMV might be driving HOK differentiation towards a dysregulated intermediate phenotype while inhibiting their terminal differentiation to maintain a cell state capable of sustaining viral replication. Using SciViewer, our recently developed scRNA-seq data visualization tool, we determined that as the infection progressed from low to high viral transcript accumulation, HCMV infection upregulated E2F targeted genes, some antiviral and immune responses as well as autophagy and cellular stress responses, while downregulating interferon-stimulated genes, immune and antiviral response genes, pyroptosis, inflammatory cell death, and membrane remodeling and viral entry genes. Overall, our study shows that primary human HOKs can be used as a model to study HCMV infection in the oral mucosa and that HCMV drives dysregulated intermediate oral keratinocyte differentiation and altered immune responses which likely facilitate viral replication in the oral mucosa and spread into the rest of the host.

## Introduction

Human cytomegalovirus (HCMV) is a widely spread β-herpesvirus infecting 60-70% of the US population causing major morbidity and mortality in individuals with an underdeveloped or weakened immune system, such as neonates, solid organ and hematopoietic stem cell transplant recipients, AIDS patients, and those on chemo- or radiotherapy or other immunosuppressive regimens (1). HCMV is a major infectious cause of illness and death following organ transplantation, the primary infectious contributor to birth defects, and the most common cause of nonfamilial sensorineural hearing loss in children (2, 3).

Like other herpesviruses, HCMV causes lifelong infection manifesting as latent, persistent, and productive infection. Treatments for HCMV are limited and there is no licensed vaccine against this virus. Antiviral drugs, in particular ganciclovir, can be effective at decreasing HCMV replication and disease. However, ganciclovir and other antivirals can have severe side effects and may be teratogenic (4). Moreover, viral resistance to antivirals has been increasingly reported and creates a significant challenge to the management of HCMV infections (5, 6).

HCMV is usually spread between individuals through the oral mucosa. Previous work has shown that HCMV can be detected in saliva and in oral epithelial cells *in vivo* (7–11). Moreover, *in vitro* experiments have demonstrated that HCMV can infect and replicate in various oral cell lines such as telomerase-immortalized normal oral keratinocytes (NOKs), telomerase-immortalized gingival cells (hGETs) (12), various mucocutaneus cell lines (13), as well as a few specialized primary oral cell types such as primary tonsil epithelial cells (13), primary salivary gland epithelial cells (14), and a cultured gingival tissue model (15). These findings suggest that HCMV is capable of infecting various cell types within the oral mucosa and support the notion that the oral mucosa is an important site of viral transmission.

The oral epithelium consists of stratified layers of keratinocytes, beginning with the basal layer—the bottom-most layer that contains stem-like cells—followed by intermediately differentiated layers, and ending with the terminally differentiated or cornified layer, where cells lose their nuclei, harden, and undergo programmed cell death to form a protective barrier against the environment. Despite its importance to the HCMV lifecycle, it remains unclear which layer(s) of oral keratinocytes is targeted by HCMV and whether the virus modulates stratification or its function. Furthermore, the innate immune responses to HCMV infection in the oral cavity are also poorly understood.

To identify HCMV specific oral keratinocyte targets and how HCMV infection modulates its function, we carried out a systematic analysis of HCMV infection in primary human oral keratinocytes (HOKs) - the predominant cell type in the oral mucosa. We used cultures of primary human oral keratinocytes (HOKs) as a model to investigate how HCMV modulates the transcriptome of oral epithelial cells to promote viral replication. Using scRNA-seq in combination with our novel analysis platform, Single-cell Interactive Viewer (SciViewer), we were able to identify cells with phenotypes corresponding to distinct keratinocyte layers and assess how HCMV infection transcriptionally altered HOKs. HCMV infection upregulated genes involved in immune responses, cell cycle regulation, neuronal functions, metabolism, stress response, molecular chaperones, extracellular matrix, and signaling. Conversely, it downregulated genes associated with inflammation, cell cycle control and apoptosis, adhesion, migration, transcription factors, and signaling pathways when compared to uninfected cells. We found that HCMV drives infected HOKs toward a dysregulated, intermediate differentiation state. Furthermore, we observed that when focused on infected cells, increasing HCMV gene expression correlated with upregulation of E2F targets, G2/M checkpoint pathways, and hedgehog signaling, alongside downregulation of oxidative phosphorylation, adipogenesis, and fatty acid metabolism pathways. HCMV infection also significantly modulated innate immune responses in HOKs. Understanding which oral epithelial cell types are targeted by HCMV, and how the virus manipulates these cells to establish productive infection, provides crucial insights that may guide the development of novel therapeutics aimed at preventing primary infection or transmission through the oral mucosa.

## Results

### HCMV infects and replicates in primary human oral keratinocytes

To begin to understand primary HCMV transmission and dissemination through the oral cavity and the consequences of HCMV infection in oral cells, we used cultures of primary human oral epithelial cells, also known as human oral keratinocytes (HOKs), as an *in vitro* model of HCMV infection of the oral mucosa. To determine if HCMV infects primary cultures of HOKs, we mock- or HCMV-infected HOK monolayers at an MOI of 0.5 based on the viral titer in human retinal pigment epithelial (ARPE-19) cells and examined cells by microscopy at 1, 3, 5, and 8 days post infection (dpi) (**Fig. 1A**).

**Fig. 1:**
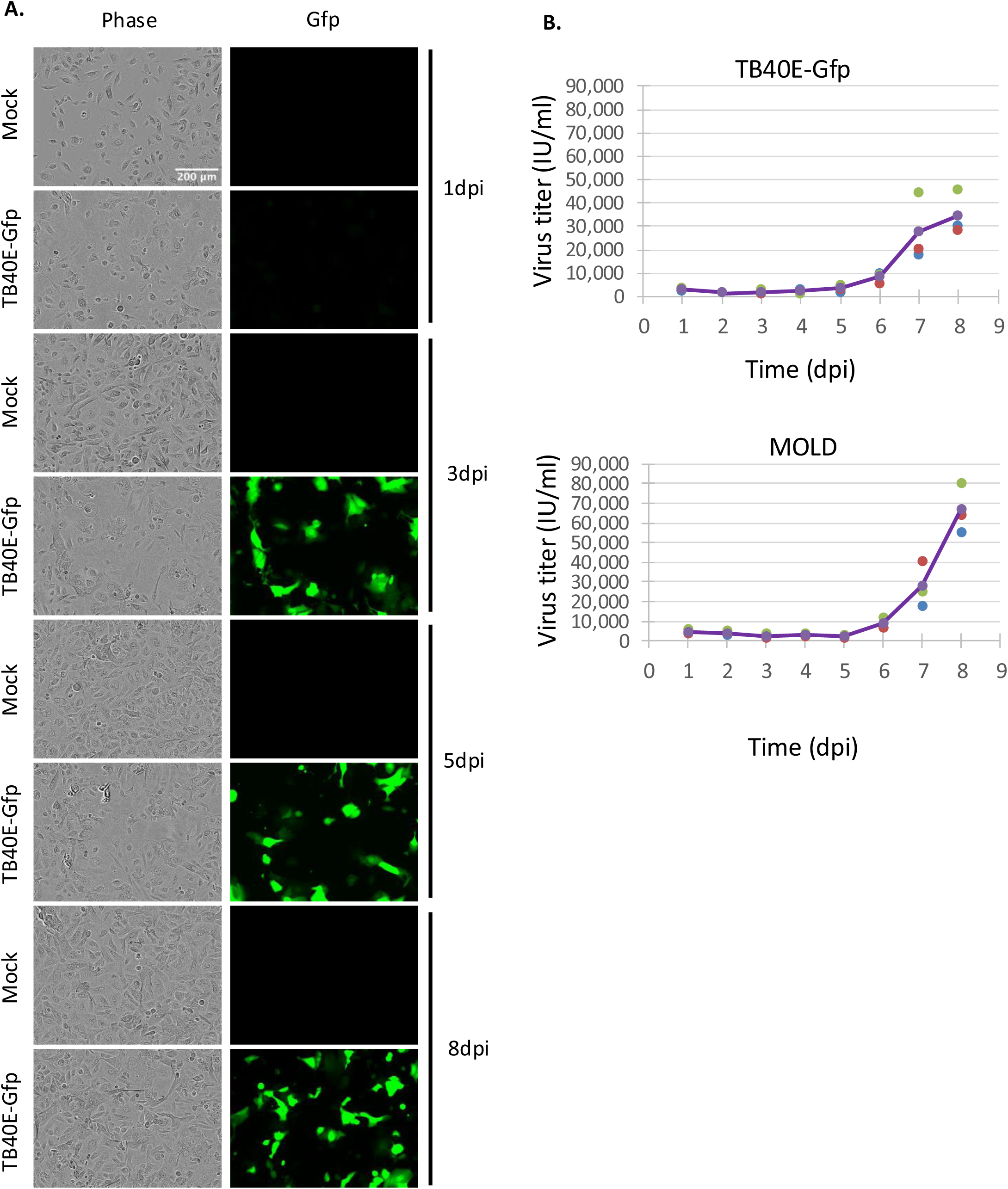
HCMV infects and replicates in primary human oral keratinocytes. **(A)** Primary human oral keratinocytes (HOKs) were either Mock- (media alone), or HCMV-infected with TB40E-Gfp or MOLD strains at MOI of 0.5 (based on HCMV titers in ARPE-19 cells) and imaged in phase and Gfp at 1, 3, 5, and 8 days post infection (dpi). **(B)** Primary HOKs were infected with TB40E-Gfp or MOLD at an MOI=2. Supernatants were collected at the indicated timepoints and infectious virus particles/infectious units (IU) from supernatants were quantified via a modified shell assay in HEL fibroblasts. The purple lines represent averages of three independent experiments shown in green, blue, and red dots.

We used the TB40E-Gfp strain of HCMV, which expresses enhanced green fluorescent protein (eGfp) from an SV40 promoter. We passaged the virus eight times in ARPE-19 cells to enhance its infectivity in epithelial cells (16, 17). We observed that HCMV was able to infect HOKs as evidenced by the cytopathic effects induced by the virus, particularly at 5 and 8 dpi, and by Gfp-positivity starting with 3 dpi. Gfp was visible as early as 1 dpi with auto exposure (data not shown). These data indicate that HOKs are permissive to HCMV infection.

We next aimed to determine whether HCMV replicates in HOKs. In addition to TB40E-Gfp, we utilized another HCMV strain, MOLD - a low-passage clinical isolate known for its strong tropism toward salivary gland epithelial cells (14, 18, 19). To assess viral replication, we performed single-step virus growth curve assays by infecting HOKs at a multiplicity of infection (MOI) of 2 and collecting supernatants daily for 8 days post-infection (dpi). Infectious virus particles in the supernatants were quantified at each time point using a modified shell assay on human embryonic lung (HEL) fibroblasts. We found that both viruses replicated in HOKs and resulted in the release of infectious progeny albeit at low levels (**Fig. 1B**), reaching 3.47 x 10^4^ infectious units (IU)/ml for TB40E-Gfp and 6.42 x 10^4^ IU/ml for MOLD by 8dpi. These findings are consistent with previous reports showing low-level HCMV replication in primary salivary gland epithelial cells (14). These data suggest that HOKs are fully susceptible to HCMV replication and support the use of primary HOKs as a relevant model system for studying HCMV infection in oral epithelial cells.

### Single cell RNA sequencing identified 11 subtypes of cells among mock- and HCMV-infected cultures of primary human oral keratinocytes (HOKs)

To investigate how HCMV changes the transcriptome of primary HOKs, we performed scRNA-seq on monolayer cultures of these cells either mock- or HCMV-infected with TB40E-Gfp or MOLD at an MOI of 0.5 and processed for scRNA-seq after 1 dpi and 3 dpi; time points where no evidence of viral shedding, and therefore, secondary infections would occur (**Fig. 2A**). The MOI of 0.5 was chosen as it resulted in 30-40% infection in HOKs based on Gfp positivity (**Fig. 1**).

**Fig. 2:**
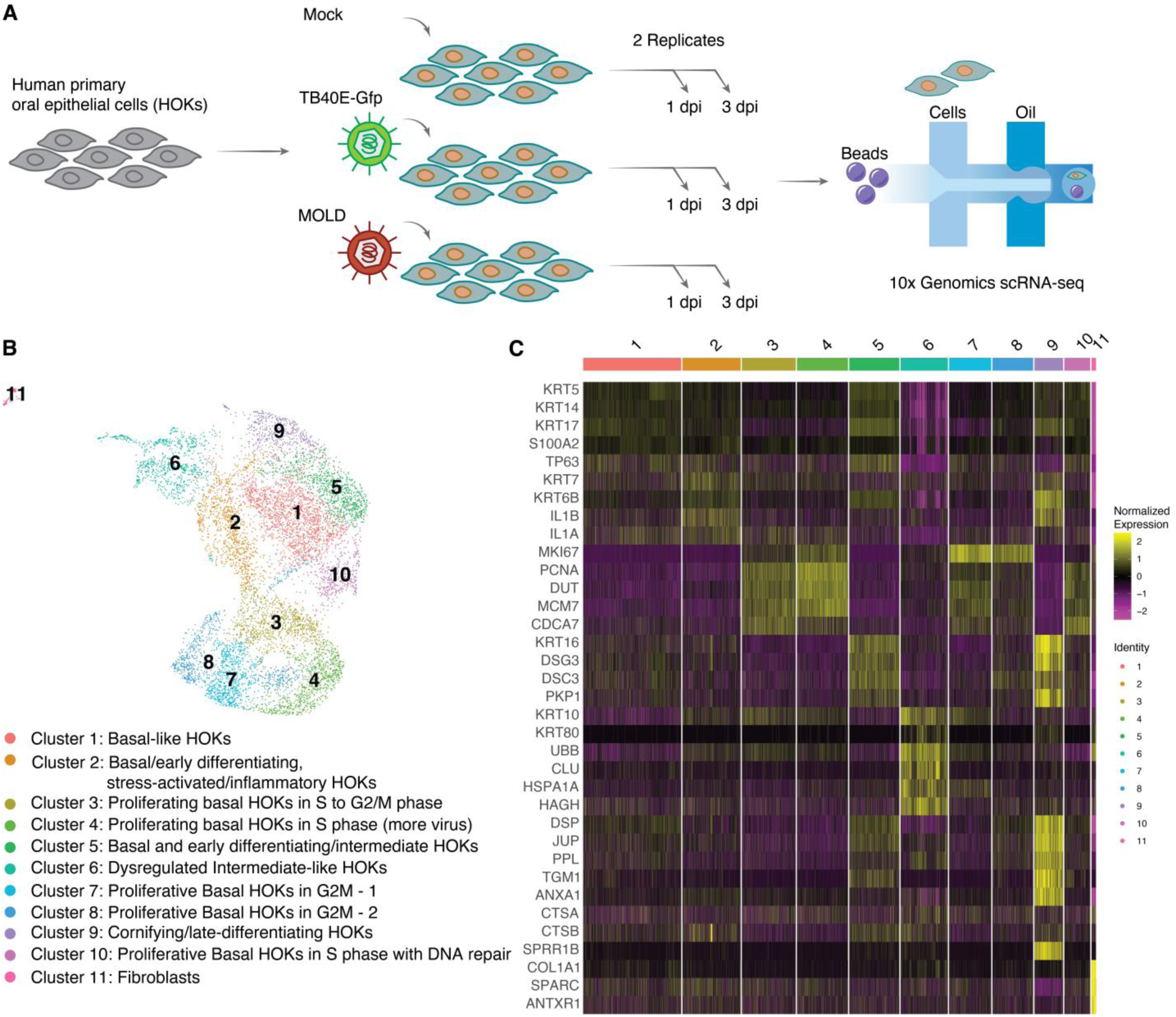
Primary human oral keratinocytes cluster into 11 subtypes based on single cell RNA sequencing. **(A)** We performed single cell RNA sequencing (scRNA-seq) on primary HOKs either mock- or HCMV-infected with TB40E-Gfp or MOLD at MOI of 0.5 (based on HCMV titers in ARPE-19 cells) and processed for scRNA-seq using the 10X Genomics platform after 1 dpi and 3 dpi. The experiment was performed in duplicate. **(B)** We performed uniform manifold approximation and projection (UMAP) mapping on cells from all samples processed for scRNA-seq and Louvain clustering observing that cells can be grouped into 11 clusters (cluster names were assigned based on terms associated with the top genes expressed in each cluster). **(C)** Single-cell expression heatmap of selected marker genes for each cluster. Markers were identified as differentially expressed host genes between the tested cluster against the rest of cells in the data. Markers were selected based on literature.

We performed uniform manifold approximation and projection (UMAP) mapping and Louvain clustering on 8623 cells pooled from all samples processed for scRNA-seq. We identified 11 clusters among mock- and HCMV-infected cells (**Fig. 2B**). Interestingly, non-infected cells were represented in 10/11 clusters, suggesting that primary HOKs are not homogenous. The top expressed genes in each cluster did not always include signature markers for HOK subtypes (**SI Appendix, Fig. S1**). Therefore, we examined the broader lists of significantly upregulated genes in each cluster to determine cell subtypes (**Fig. 2C and SI Appendix, Fig. S1E, Table S1**). Cluster 1 had higher expression levels of basal HOK genes: KRT5, KRT14, KRT17, S100A2, and TP63 (**Fig. 2C**). Cluster 2 showed higher levels of basal/early differentiating, stress-activated/inflammatory genes: KRT7, KRT6A, CAVIN3, IL1B, and IL1A. Clusters 3, 4, 7, 8, and 10 showed higher levels of genes associated with proliferation and cell cycle stages: MKI67, PCNA, DUT, MCM7, and CDCA7. Cluster 5 comprised a mixed population of basal and early suprabasal/intermediate/spinous-like HOKs characterized by higher levels of KRT16, KRT17, DSG3, DSC3, and PKP1. Cluster 6 contained dysregulated intermediate-like HOKs expressing KRT10, KRT80, UBB, CLU, HSPA1A, and HAGH. Cluster 9 was composed of cornifying/late-differentiating HOKs expressing DSP, JUP, PPL, TGM1, ANXA1, CTSA, CTSB, and SPRR1B. Finally, we identified cluster 11 as a small cluster of fibroblasts expressing COL1A1, SPARC, and ANTXR1. Overall, although HOK monolayer cultures were studied, we observed evidence of the three main layers or differentiation states of the oral mucosa - ranging from basal to intermediate to terminally differentiated oral keratinocytes – resembling those typically found in native oral tissues. This suggests that monolayers of cultured primary HOKs can serve as a model of HCMV infection in the oral mucosa as they broadly recapitulate the differentiation states observed in 3D tissues *in vivo*.

### HCMV-infected primary HOKs collectively express all temporal phases of HCMV genes

We next examined HCMV gene expression patterns in HOKs. As expected, no HCMV transcripts were detected in mock-infected samples (**SI Appendix, Table S2**) whereas HCMV transcripts were readily detected in HCMV-infected samples. The HCMV-infected samples included cells with no detected HCMV transcripts, which we considered to be uninfected (bystander) cells. Infected cells harboring HCMV transcripts had a clear bimodal gene expression distribution with one set of marginally infected cells having a mean of 0.0023% HCMV transcripts and a second set of infected cells having a mean of **1.656%** HCMV transcripts (**Fig. 3A and SI Appendix, Table S2**). The exact nature of marginally infected cells is unclear. They could represent cells in which HCMV replication is restricted by intrinsic immune barriers or abortively infected cells, or cells which may have taken up residual viral RNA from the supernatant. However, we detected low-level expression of genes from all viral classes (**Fig. S3**)(20).

**Fig. 3:**
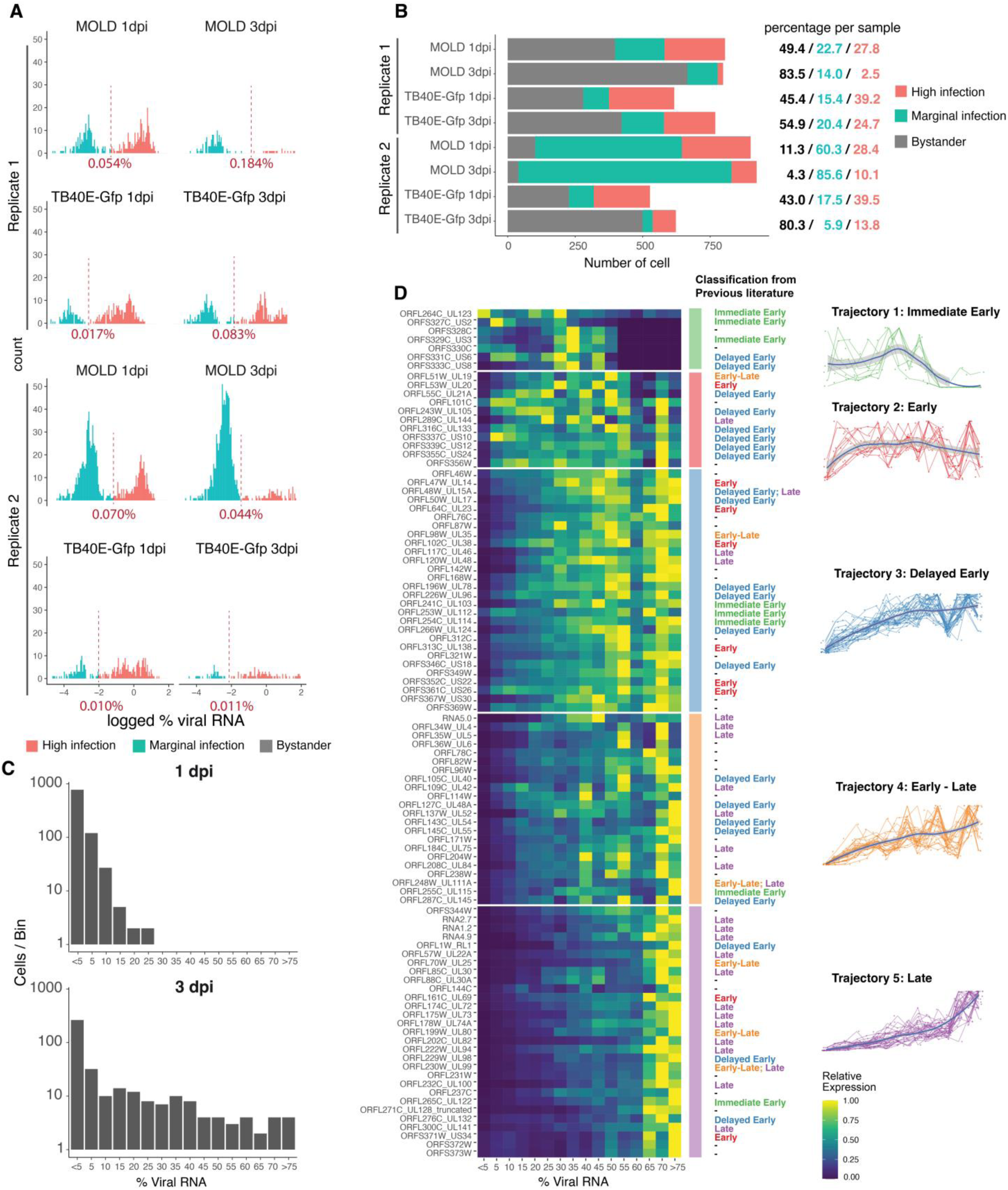
Primary HOKs infected with HCMV express all temporal phases of HCMV genes. **(A)** Histogram of viral transcript percentage out of total number of transcripts in virus positive cells for each HCMV infected sample. Viral transcript percentage is converted on log_10_ scale with a fudge factor added. The percentage of viral counts used to determine infection level is labeled for each sample. Cells with viral transcript percentage higher than the infection level are considered to be highly infected. The rest of virus positive cells are labeled as marginal infection. **(B)** Distributions of all cells in each infected sample. Bystander cells are cells without any viral transcript. Marginal infection and high infection cells are defined in A. **(C)** Histograms of the number of highly infected cells, split by days post infection. Cells were grouped in 5% bins based on increasing percentage of viral gene expression, going from cells expressing low levels of viral transcripts (left) to cells expressing high levels of viral transcripts (right). **(D)** Heatmap of the top 30% (97/344) expressed HCMV genes at 1dpi and 3dpi from both viral strains. We investigated how HCMV genes change in expression with increasing levels of viral transcript percentage only in highly infected cells, grouped. Represented are the means of the raw counts for each gene within a bin relative to the means of raw counts for that gene in the highest expressing bin. HCMV temporal gene classifications are based on Rozman et al. 2022 Cell Reports, Weekes et al. 2014 Cell, and Ye et al. 2020 Frontiers in Microbiology. HCMV genes were grouped based on their expression patterns into 5 trajectories using the k-means clustering (k=5).

The distributions were well separated allowing us to define a threshold distinguishing high from marginally infected cells. The thresholds were determined for each sample individually based on local minima in viral transcript distributions (**Fig. 3A and Methods**). Importantly, we observed a decrease in the number of cells containing viral transcripts between 1 and 3 dpi in both TB40E-Gfp replicates and in one of the MOLD replicates. Namely, in TB40E-Gfp-infected samples, 54.6% and 57% of cells in replicates 1 and 2, respectively, contained viral transcripts at 1dpi (marginally and highly infected cells combined) (**Fig. 3B and SI Appendix, Table S2**). By 3 dpi, these numbers declined to 45.1% (p<0.001, two-proportion z-test) and 19.7% (p<0.001, two-proportion z-test), respectively. In MOLD-infected samples, 50.5% and 88.7% of cells were infected at 1 dpi, with the percentages decreasing to 16.5% (p<0.001, two-proportion z-test) at 3 dpi for replicate 1, and increasing to 95.7% (p<0.001, two-proportion z-test) at 3 dpi for replicates 2 (**Fig. 3A-B and SI Appendix, Table S2**). Taken together these results suggest that viral replication is impeded in most HOKs during this period.

We then investigated the expression of the various temporal phases of HCMV genes in infected HOKs (**Fig. 3C-D**). We used the increasing amount of viral transcripts observed in highly infected cells as a proxy for infection time given the limited experimental time points (1 dpi and 3 dpi) in data collection. Indeed we found most of the cells with high amount of viral transcripts are cells from 3 dpi samples **(Fig. 3C)**. We ranked cells based on HCMV transcript levels (viral transcript load) from low (**Fig. 3D, left**) to high (**Fig. 3D, right**) and examined the expression patterns of the top 30% most highly expressed HCMV genes. Each gene was labeled with its temporal category - immediate early, early, delayed early, early-late, or late - based on previously established expression patterns (**Fig. 3D, middle**) (21–23). We identified five distinct gene expression trajectories across cells with increasing viral transcript load (**Fig. 3D**). Trajectory 1 genes (e.g., *UL123, US2, US6*) show high expression in cells with low viral transcript loads that decreases as the loads increase, suggesting these may be immediate early genes. Trajectory 2 genes (e.g., *UL19, UL20, UL105, UL144*) increase in expression starting with slow viral transcript loads, remain stable as the viral loads rise, and decrease at peak viral transcript loads, indicating they may be early genes. Trajectory 3 genes (e.g., *UL14, UL96, UL138*) exhibit a delayed increase in expression as viral transcript loads rise, plateauing at high levels, consistent with delayed early genes. Trajectory 4 genes (e.g., *UL4, UL48A, UL111A, UL115*) also show a gradual increase, but unlike trajectory 3 genes, they continue to rise as viral transcript loads peak, suggesting they are early-late genes. Finally, trajectory 5 genes (e.g., *UL22A, UL73, UL94, US34*) remain low until viral transcript loads reach peak levels, at which point they increase, indicating they may be late genes.

Although our analysis classified genes based on increasing viral transcript levels rather than timepoints, the overall expression dynamics align with previous studies that described temporal phases of HCMV gene expression in other cell types (24–26).

### HCMV infection in HOKs resulted in significant changes in host gene expression

We next investigated global host gene expression changes upon HCMV infection in HOKs (**Fig. 4**). We performed differential gene expression analysis between mock-infected cells and highly infected cells, marginally infected cells, and bystander cells (**Fig. 4A-C and SI Appendix, Table S3**). In the comparison of highly infected versus mock-infected cells, we identified 357 differentially expressed genes (DEGs), of which 271 were significantly upregulated and 86 were significantly downregulated (**Fig. 4A**). We observed that HCMV upregulated some immune response genes (ISG15, IFI6, MX1, OAS1, IFI27, ISG20, CXCL8, TNFAIP2, IL23A, CCL28, HLA-B, and HLA-F) along with a suppressor of cytokine signaling (SOCS1); cell cycle regulation genes (CCNE1, CCNA1, CCNE2, CDKN2D, CCDC69, and CCDC149); cell death and cancer-related genes (TP53INP2, TP53I11, BIRC3, and AGR2); neuronal and synaptic function genes (RBP7, ENO2, BEX2, BEX5, UCHL1, INA, SYNGR1, SNCG, CAMK2B, PRSS22, and MAP1B); molecular chaperones and metabolism, stress, and vesicular trafficking genes (METRNL, CLU, HMOX1, GCH1, GCHFR, RAB3A, RAB6B, DNAJA4, HSPA1A, HSP90B1); tight junctions and extracellular matrix and cell adhesion genes (NID1, CLDN4, CLDN23, TIMP2); microtubule-associated genes (MAP7D2, MAP1B, MAPRE3, and MAPRE2); kinases and signal transduction genes (AKAP12, RGS9, PKN1, and PDE4A); and transcription factors (FOXQ1, KLF2, IRF1, IRF7, HES6, E2F1, MAFG).

**Fig. 4:**
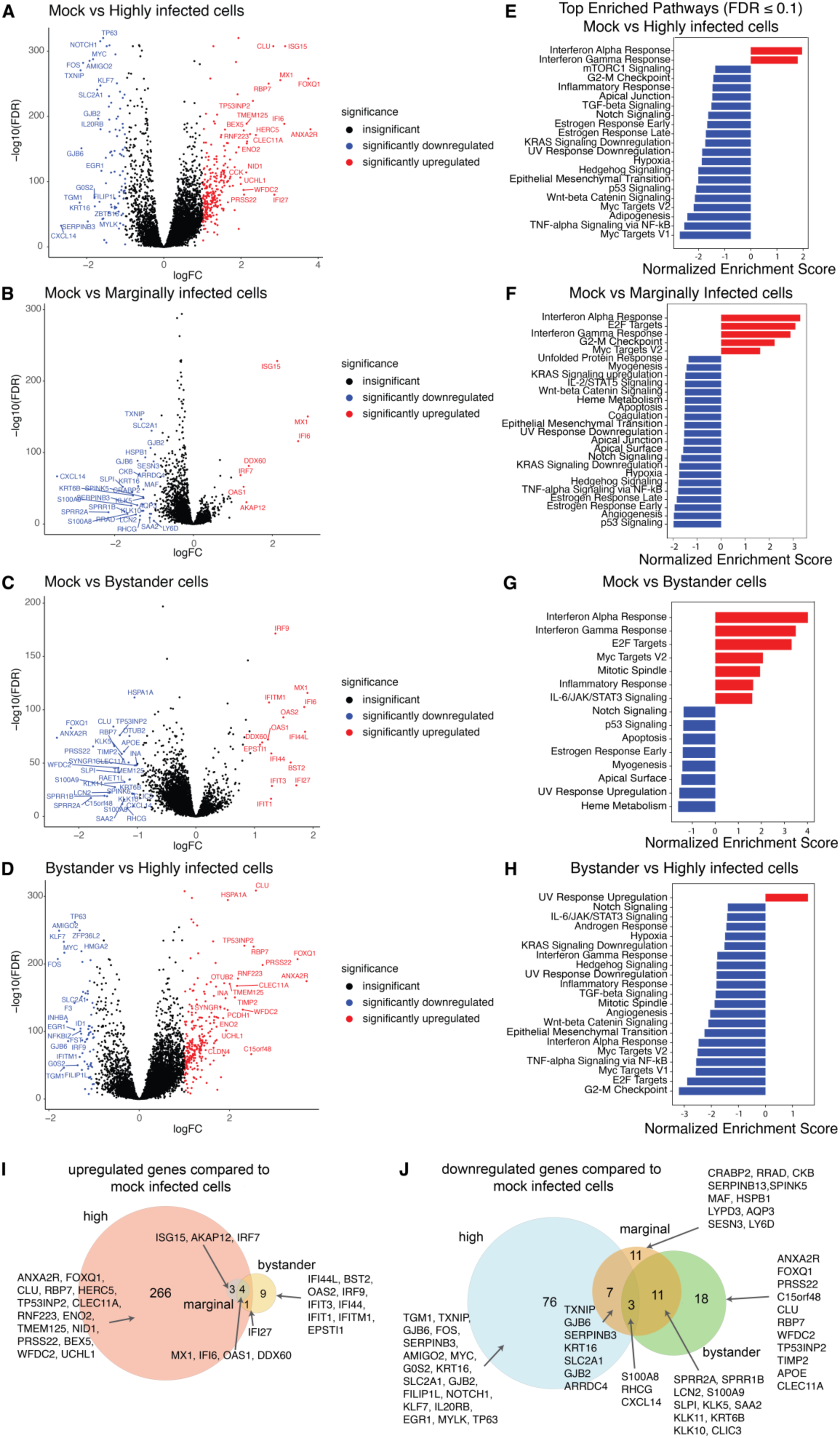
HCMV infection in HOKs resulted in significant changes in host gene expression. (A - B). Volcano plot of host gene differential expression (DE) analyses comparing mock vs highly infected cells (**A**), mock vs marginally infected cells (**B**), mock vs bystander cells (**C**), and bystander vs highly infected cells (**D**). DE genes are defined as genes have more than two-fold change and < 0.05 false discovery rate (FDR). The top 20 genes either upregulated or downregulated were labeled. X-axis represents gene expression log_2_ fold change (logFC). Y-axis is the –log10 of false discovery rate. (**E - H**) GSEA enriched pathways in mock vs highly infected cells comparison (**E**), mock vs marginally infected cells (**F**), mock vs bystander cells (**G**), and bystander vs highly infected cells (**H**). Pathways tested with FDR ≤ 0.1 are shown. (**I**) Venn Diagram of significantly upregulated genes in DE comparison in A - C. (**J**) Venn Diagram of significantly downregulated genes identified in DE comparison in A - C.

HCMV infection downregulated inflammation and immune response genes (CXCL14, IL33, S100A8, NFKBIZ, IL20RB, and ILRAP); stress, DNA damage response, and apoptosis genes (DUSP1, TP53AIP1, GADD45A, TP63, and TXNIP); junctions, cell adhesion, and cell migration genes (PKP1, DSC3, AJUBA, FAT2, FAT1, SH3PXD2A, TNS4, MMP28, SNAI2, AJUBA, and MYLK); cell signaling and growth factors (EGFR, FGFR3, IRS1, PTHLH); ion channels and transporters (KCNJ5, SLC7A11, SLC2A1, GJB2); and transcription factors (FOS, JUN, CEBPD, EGR1, MYC, KLF6, KLF7, ETS2) (**Fig. 4A and SI Appendix, Table S3**). These changes suggest that HCMV actively modulates host genes across multiple pathways, likely to create a cellular environment conducive to viral replication.

The differences in gene expression in mock-versus marginally infected cells were more modest (**Fig. 4B**). Specifically, we identified 39 differentially expressed genes with only 7 genes upregulated (MX1, IFI6, ISG15, DDX60, OAS1, IRF7, and AKAP12) and 31 being downregulated in marginally infected cells (**Fig. 4B and SI Appendix, Table S3**). Interestingly, all 7 upregulated genes are either interferon-stimulated or immune/antiviral response genes, suggesting that in these cells, the innate immune response may be effectively combating viral gene expression, at least at these time points, resulting in a marginal infection. On the other hand, other immune response and inflammation genes (S100A8, S100A9, SLPI, CXCL14, LCN2, SAA2) were downregulated in marginally infected cells, along with serine proteases and serine protease inhibitors involved in epithelial remodeling (KLK5, KLK10, KLK11, SERPINB3, SERPINB13, and SPINK5), transport genes (SLC2A1, AQP3, RHCG, CLIC3), metabolism and stress response (CKB, SESN3, TXNIP, HSPB1, ARRDC4), and keratinocyte barrier and differentiation (KRT16, KRT6B, SPRR1B, SPRR2A, LY6D, LYPD3, CRABP2) (**Fig. 4B and SI Appendix, Table S3**), suggesting that, despite low levels of viral gene expression, HCMV can modulate the host transcriptome likely to prepare the cell for subsequent increases in viral gene expression and replication.

In the comparison between bystander and mock-infected cells, we identified 46 differentially expressed genes, including 14 genes that were upregulated in bystander cells (**Fig. 4C**). All of these upregulated genes (MX1, IFI44L, IFI6, IFI27, BST2, OAS2, IRF9, IFIT3, IFI44, IFIT1, IFITM1, OAS1, DDX60, and EPSTI1) were associated with interferon and immune responses. Notably, though some of these were also upregulated in marginally and highly infected cells (MX1, IFI6, OAS1, and DDX60), the majority were unique to bystander cells (**Fig. 4G**). Among the genes downregulated in bystander cells, we observed immune and antimicrobial genes (S100A8, S100A9, CXCL14, RAET1L, SAA2, LCN2, CLEC11A, C15orf48, RBP7); keratinocyte differentiation and mucosal barrier genes (SPRR1B, SPRR2A, KRT6B); serine proteases and protease inhibitors involved in epithelial remodeling (KLK5, KLK10, KLK11, PRSS22, WFDC2, SLPI, SPINK6, TIMP2); molecular chaperones and stress response genes (HSPA1A, TP53INP2, CLU, OTUB2); neuronal genes (SYNGR1 and INA); and a transcription factor (FOXQ1) (**Fig. 4C and SI Appendix, Table S3**). Interestingly, while FOXQ1 was highly downregulated in bystander compared to mock infected cells, it was upregulated in highly infected versus mock-infected cells, suggesting FOXQ1 could potentially be a pro-viral factor actively downregulated in bystander cells and upregulated by the virus in infected cells with a potential role in viral gene expression and replication.

To distinguish gene expression changes directly caused by HCMV infection from those driven by the infection environment, we compared highly infected cells to bystander cells, identifying 316 differentially expressed genes. Of these, 252 were upregulated and 64 were downregulated in the highly infected cells. Highly infected cells upregulated genes involved in cell cycle and proliferation (CCNE1, CCNA1, CDKN2D, E2F1, DYRK1B, PPP2R5B, FAM110C, ID1); apoptosis regulation (DDIT3, TP53I3, TP53I11, NIBAN1, G0S2, and TNFAIP2); transcription factors and cell differentiation (FOXQ1, FOXL2, HES6, CEBPA, RORA, ELF3, MITF, TFAP2C, KLF2, FST); extracellular matrix, cell adhesion, and tight junctions (SAHS1, TIMP2, PCDH1, NID1, CLDN4, CLDN23, CGN, FLNC, FBLN2, SERPINB3, SERPINB5, MMP28, THBS1, COL7A1, COL8A1, DST, and AMIGO2); stress response, molecular chaperones, and protein folding (CHAC1, DUSP8, HSPA1A, HSPA1B, HSPA2, DNAJB9, DNAJA4, and SVIP); signal transduction and kinase activity (CAMK2B, PRKAA2, PKN1, MAP3K11, MRAS, RASGRP2, RND2, RHPN2, and RGS9); immune response and inflammation (CXCL8, CXCL14, CCL28, IL11, IL23A, NFKBIZ, TNFAIP2, RAET1L, ISG20, IFITM1, IFITM2, IFI44L, IRF9, and HERC5); metabolism, transport, and homeostasis (SLC2A1, SLC26A11, SLC41A2, SLC43A2, SLC25A4, SLC25A42, SLC46A3, SLC17A5, ENO2, PDK2, APOE, PLPPR2, MT1X, MT1F, MT1G, LPIN2, GCH1, and GCHFR); neuronal function and synaptic components (SYNGR1, INA, VSNL1, GABRE, SNCG, and NEFH); ubiquitination, deubiquitination, protein degradation, and protein quality control (UBB, FBXO32, HERC5, RNF208, MARCHF3, UCHL1, OTUB2, USP2, and KLHL24); cytoskeleton dynamics, intracellular transport, and vesicular trafficking (MAPRE3, MAPRE2, MAP1B, MAP1S, MAP7D2, KIF3C, KIF13B, RAB3A, RAB6B, RILP, SNX10, SNX16, COBL, FLNC, and DBNDD1); and epigenetic regulators and chromatin modifiers (HDAC9, ING2, LRIF1) (**Fig. 4D and SI Appendix, Table S3**).

Among the downregulated genes in highly infected compared to bystander cells, we identified immune response and inflammation genes (CXCL2, CXCL14, IL20RB, IL1RAP, IFI44L, IFITM1, IFITM2, IRF9, NFKBIZ, CEBPD, LGALS9B, LGALS9C); cell cycle regulation and proliferation genes (PLK1, PLK2, CDCA3, KIF20A, GADD45A, ID1); apoptosis regulators and stress response genes (DDIT4, PHLDA1, TXNIP, G0S2, MT1X); extracellular matrix, cell adhesion and motility genes (MMP28, THBS1, COL7A1, COL8A1, PCDH7, DST, AMIGO2, FILIP1L); growth factors, signaling pathways, and metabolism genes (PTHLH, IRS1, SLC2A1, FGFBP1, VEGFC, VSNL1, GABRE); cell fate and differentiation genes (DKK1, NOTCH1, JAG1, DLK2, INHBA), serine protease inhibitors (SERPINB3, SERPINB5); and transcription factors and gene expression regulators (FOS, FOSB, EGR1, MYC, KLF7, HMGA2, CEBPD, ETS2, TP63, SNAI2, ZFP36L1, ZFP36L2) (**Fig. 4D and SI Appendix, Table S3).**

Interestingly, we identified no significantly differentially expressed host genes in the comparison between marginally infected and bystander cells (data not shown) at least using our cutoffs (fold change > 2 and false discovery rate < 0.05). This suggests that marginally infected cells might be bystander cells which have ingested viral transcripts from the supernatant. Alternatively, these cells could be pseudo-latently infected with HCMV, or HCMV might have “deliberately” seeded them with viral transcripts from nearby infected cells to prepare them for an upcoming infection.

Gene Set Enrichment Analyses (GSEA) identified pathways and processes that were enriched among the differentially expressed genes (**Fig. 4E-H**). Interferon alpha and gamma were among the top pathways upregulated in highly infected, marginally infected, and bystander cells compared to mock-infected cells, whereas “UV response up” was the only pathway upregulated in highly infected compared to bystander cells (**Fig. 4H**). In contrast, the Notch signaling pathway was downregulated in highly infected, marginally infected, and bystander cells compared to mock-infected cells as well as in highly infected cells compared to bystander cells, whereas the p53 pathway and estrogen response early was downregulated in the first three comparisons only. Importantly, all pathways upregulated in bystander compared to mock-infected cells, namely Interferon alpha and gamma, E2F targets, Myc targets V2, Mitotic spindle, and Inflammatory response, IL-6/JAK/STAT3 signaling, were downregulated in highly infected cells compared to bystander cells (**Fig. 4G and H**). Similarly, “UV response up” was downregulated in bystander compared to mock-infected cells, but was upregulated in highly infected cells compared to bystander cells, indicating HCMV is specifically manipulating these pathways to counteract cellular responses to the infection.

Among the upregulated genes in the three comparisons against mock-infected cells (highly infected versus mock, marginally infected versus mock, and bystander versus mock) (**Fig. 4A-C**), only four genes were found in highly infected, marginally infected and bystander cells (**Fig. 4G**) -MX1, IFI6, OAS1 and DDX60 - which are interferon stimulated genes. IFI27 was also upregulated in highly infected and bystander cells. Bystander cells specifically upregulated 9 other interferon stimulated genes including IFI44L, BST2, OAS2, IRF9, IFIT1, IFIT3, IFI44, IFITM1 and EPSTI1 that were not found in HCMV infected cells. Two hundred sixty-six genes were uniquely upregulated in highly infected cells when compared to mock-infected cells, and 3 additional genes were upregulated in both highly infected and marginally infected cells (**Fig. 4G and SI Appendix, Table S3**).

Changes in infected cells compared to mock-infected cells indicate processes directly modified by the virus. Downregulated genes from the three comparisons against mock-infected cells had three genes in common: S100A8, RHCG, and CXCL14 (**Fig. 4I**). HCMV infection, regardless of the infection level, downregulated genes with a function in intercellular gap junctions (GJB2, GJB6), cell structure (KRT16) and glucose metabolism (TXNIP, SLC2A1, ARRDC4) (27). Highly infected cells downregulated genes involved in epidermis development (TGM1, KLF7, INHBA, NFKBIZ, FST, NOTCH1, JAG1, ZFP36L1, SNAI2, TP63), cellular stress response (FOS, MYC, EGR1, DDIT4, ZFP36L1), cell adhesion (AMIGO2, DSG3, TENM2, FLRT3, DSC3, FAT1, FAT2, AJUBA, LGALS7B), and cornified envelop and cell structure (IVL, KRT14, PKP1, EPPK1, MYLK) (**Fig. 4J**). Bystander cells and cells with marginal infection together downregulated a group of small proline-rich proteins (SPRR2A, SPRR1B) which are essential in keratinocyte cornified envelope formation, and kallikreins (KLK5, KLK10, KLK11) which are proteases that function in wound healing and immune response. Bystander cells specifically downregulated ANXA2R, FOXQ1, PRSS22, C15orf48, CLU, RBP7, WFDC2, TP53INP2, TIMP2, APOE, and CLEC11A that were otherwise upregulated in highly infected cells (**Fig. 4I-J, and Table S3**), which suggests these genes could be potentially pro-viral factors actively downregulated in bystander cells and upregulated by the virus in infected cells.

Our data showed that HCMV infection globally induced the upregulation of certain immune responses, oncogenesis and cell cycle regulation pathways while downregulating other immune genes and inter-/intra-cellular structure genes. Interferon response was highly induced in bystander cells yet less so in the infected cells.

### SciViewer Directional Analysis and GSEA reveal host gene pathways associated with increased HCMV gene expression in highly infected cells

We were next curious to explore the transcriptional changes which accompanied the increasing accumulation of viral RNA transcripts and sustained HCMV infection in HOKs, particularly in highly infected cells. Because HCMV viral transcript levels span a continuum from lower to higher expression levels even among highly infected cells (**Fig. 3A**), we used SciViewer - an interactive tool that enables directional analysis across UMAP embeddings - to capture this progression (28). Building upon our DE/GSEA results that delineated broad infection-associated programs (e.g., mock vs highly infected cells), directional analysis enabled us to examine continuous changes within similar cells along a viral-load continuum – thus refining and extending the findings with higher resolution. We focused on the UMAP of 3 dpi highly infected cells (Clusters 1, 2, 6, 8, 9, 10, and 11; **Fig. 5A**), which displayed a broad range of viral gene expression levels (0.01 - 82.55%; **Fig. 5B**). A direction drawn from basal-like HOKs with low viral transcripts (cluster 1) toward intermediate-like HOKs with highest transcripts (cluster 6) yielded gene lists ranked by Pearson correlation coefficients (PCC) and p-values (**SI Appendix, Table S4**).

**Fig. 5:**
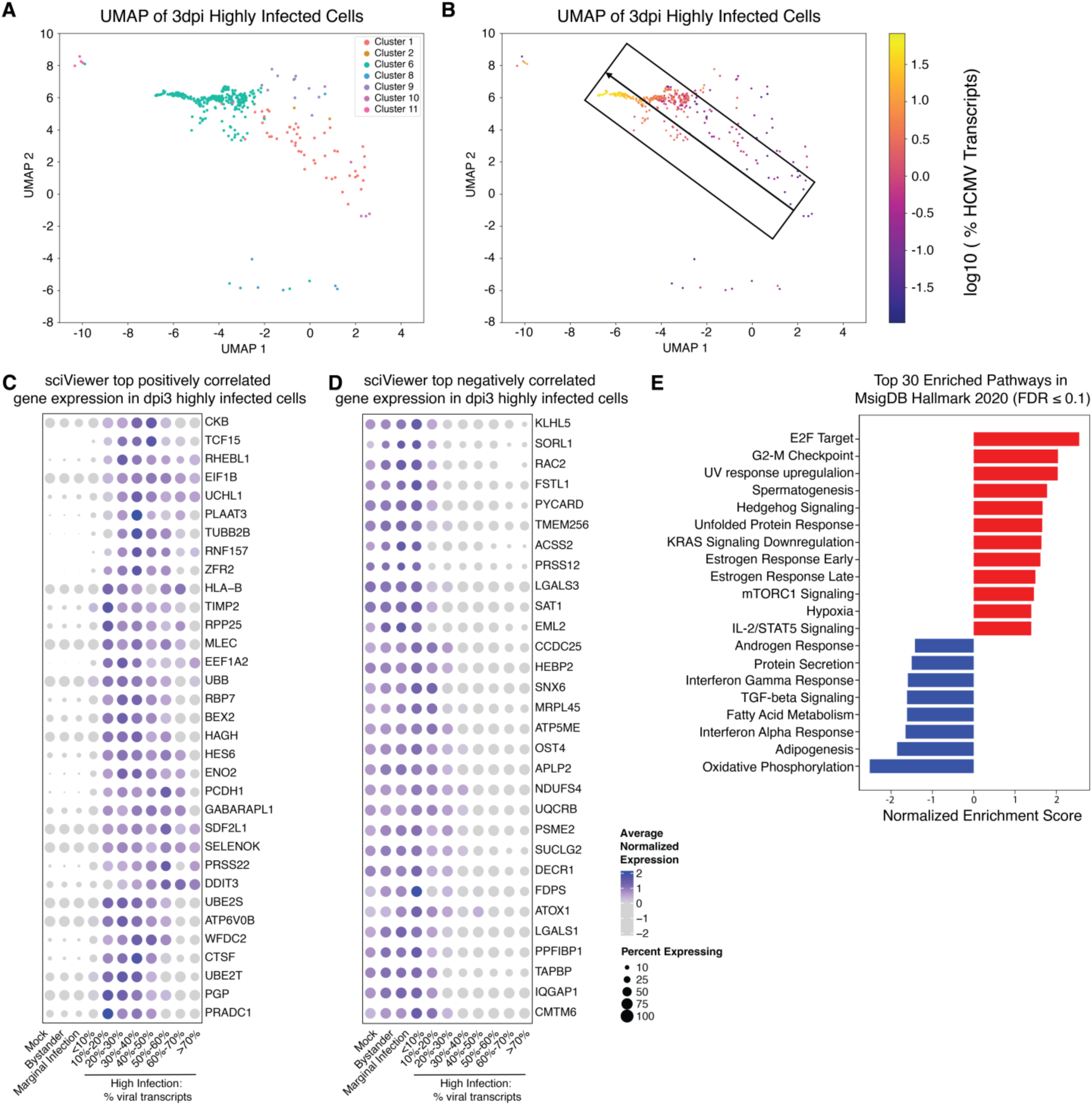
Increased HCMV replication is associated with decreased interferon and oxidative phosphorylation pathways as highlighted by SciViewer directional analysis and Gene Set Enrichment Analysis. **(A**) UMAP of 3dpi infected cells colored by Seurat cluster. This is a subset of the original UMAP containing all cells, but this subset contains only 3dpi infected cells . **(B)** UMAP of 3dpi infected cells colored by the log_10_ of viral transcript percentage. The arrow and box represents the directional analysis performed on SciViewer. The directional analysis goes from 3dpi infected cells with relatively low number of viral transcripts towards the 3dpi infected cells with the highest number of viral transcripts. Heatmaps of top positively **(C)** and negatively **(D)** correlated genes identified in the SciViewer directional analysis. 3dpi highly infected cells were grouped by their viral transcript percentage with 5% increment. Averaged expression was taken per bin for each gene. ( **E)** Gene set enrichment analysis result on the top enriched pathways ordered by correlation along SciViewer trajectory. Pathways that have FDR ≤ 0.1 are included.

Directional analysis highlighted additional genes whose expression changed progressively with viral load, complementing the DE results. The top positively correlated genes (**Fig. 5C**) included regulators of protein synthesis and metabolism (EIF1B, EEF1A2, TUBB2B), stress and autophagy factors (UCHL1, PLAAT3, GABARAPL1, ATP6V0B), immune modulators (HLA-B, RPP25), and transcriptional regulators (HES6, BEX2, RHEBL1), while the top negatively correlated genes (**Fig. 5D**) comprised innate immune and inflammatory mediators (PYCARD, FSTL1, RAC2), oxidative phosphorylation components (NDUFS4, UQCRB, SUCLG2, ATP5ME), and cell adhesion/structural components (PRSS12, EML2, LGALS3). Together, these patterns suggest that as viral load increases, HCMV promotes stress-adaptation and immune evasion while suppressing mitochondrial metabolism, pyroptosis, and membrane integrity. Beyond these top 30, additional genes from the full directional analysis (**SI Appendix**, **Table S4**) showed increasing expression of CD55, ATG101, DNASE2, HSPA13, and NFKBIB - implicating complement regulation and autophagy/stress pathways – and decreases in IFI27L2, KRT8, DSG2, and VAMP8, further supporting progressive dampening of inflammatory and membrane-remodeling programs (29). Notably, MAP1LC3A and IL32 overlapped with the DE analyses, providing independent confirmation of signals shared across both approaches.

GSEA on genes ranked by their correlation with viral transcript load revealed both overlapping and unique pathways compared to the earlier DE-based GSEA comparing mock, bystander, and infected cells (**Fig. 5E**). As in the earlier analyses, we observed enrichment of E2F targets and suppression of interferon alpha/gamma responses, confirming our previous conclusions.

Directional analysis further clarified that several metabolic programs - including oxidative phosphorylation, fatty acid metabolism - decline progressively with increasing viral load and revealed enrichment of IL-2/STAT5 pathway along the same trajectory (**Fig. 5E**, **SI Appendix**, **Table S5**). Together, these results indicate graded (continuous) transcriptional reprogramming, complementing the binary contrasts from DE by showing how host metabolism and immune signaling evolve stepwise as viral transcripts accumulate.

The results of the first directional analysis motivated a second, within-cluster analysis of the most infected population (cluster 6) (**SI Appendix**, **Fig. S4**) (29) - the dysregulated intermediate-like HOK population that harbors the highest viral transcript loads (**SI Appendix**, **Fig. S4A**).

Comparing cells with intermediate (0.01%) versus highest viral transcript levels through a second directional analysis (**SI Appendix**, **Fig. S4B and Table S6**) revealed a coherent, load-dependent program. At the gene level, SciViewer revealed unique upregulation of stress and apoptosis-related genes (ATF3, TRIB3, SDF2L1) and the complement regulator CD55, which were not detected in DE-based comparisons. These findings suggest that stress adaptation and complement evasion progressively intensify with higher viral loads. We also confirmed overlap with the DE analyses, which had highlighted induction of DDIT3, HERC5, TNFRSF18, NRIP3, and antiviral factors such as ISG15 and ISG20. On the other hand, SciViewer exposed a large set of downregulated genes absent from the DE results, including the inflammasome adaptor PYCARD, interferon signaling receptors (IFNGR1, IFNAR1), antigen presentation components (TAPBP, IL1RN, TOLLIP), epithelial barrier and differentiation genes (IRF6, CDH1, CD24, CD81, CD82, KRT23), and mitochondrial gene UQCRB. These findings extend beyond the DE-based suppression of IL32, ISG15, ISG20, FOXQ1, KRT6B, and S100A8/S100A9, showing that HCMV progressively disables a wider network of immune, epithelial, and metabolic regulators as viral loads increase. of IL32, ISG15, ISG20, FOXQ1, KRT6B, and S100A8/S100A9, showing that HCMV progressively disables a wider network of immune, epithelial, and metabolic regulators as viral loads increase.

A particularly novel observation was the pronounced downregulation of IRF6 and CDH1 (E-cadherin), which are central to keratinocyte differentiation and epithelial integrity, respectively.

Their suppression suggests that HCMV actively blocks epithelial maturation, perhaps to maintain an intermediate state that favors ongoing replication. We also noted extensive remodeling of cell surface proteome (CD9, CD24, CD47, CD58, CD81, CD82, CD99, CD109), a change not evident from DE results, which could facilitate immune evasion and viral spread (**SI Appendix**, **Fig. S4C-D**). Pathway-level analysis with GSEA supported these unique gene-level findings, showing significant negative enrichment of oxidative phosphorylation, complement, interferon alpha/gamma responses, and TNF-alpha signaling via NF-kB (**SI Appendix**, **Fig. S4E and Table S7**). Together, these results indicate that as infection intensifies within cluster 6, HCMV drives a coordinated program of stress response activation, immune suppression, impaired differentiation, and metabolic collapse – changes that became visible only when analyzed as a trajectory rather than as discrete infection states.

Primary HOKs has evolved a plethora of strategies to downregulate the host immune/antiviral response reviewed in (30, 31). Previous studies in fibroblasts and CD14+ monocytes have found interferon (IFN) stimulated genes (ISGs) together with other immune genes had lower expression in highly infected cells compared to bystander cells (32, 33). As we observed a differential upregulation of the IFN response in cells in different infection states (highly infected, marginally infected, and bystander cells), we further investigated the source of IFN and the stimulated immune responses towards HCMV infection in HOKs. Despite the high expression of ISGs, IFN genes were mostly not detected in the data, most likely due to the time of data collection in our experimental design (1 and 3 dpi) (**Fig. 6**. Sparse expression of type I and III IFN genes were observed in highly infected cells (whose transcriptomes contained <10% viral transcripts, **Fig. S4**). Expressions of type I and II IFN receptors (IFNAR1, IFNAR2, IFNGR1, IFNGR2) were found to be ubiquitous in cells without viral transcripts (mock and bystander) and cells with less than 20% of viral transcripts, but were downregulated when the cells acquired more viral transcripts (**Fig. 6B**). Expression of type III IFN receptor (IFNLR1) was specifically induced in infected cells with low levels of viral gene expression (<10% viral transcripts, **SI Appendix, Fig. S5**).

**Fig. 6:**
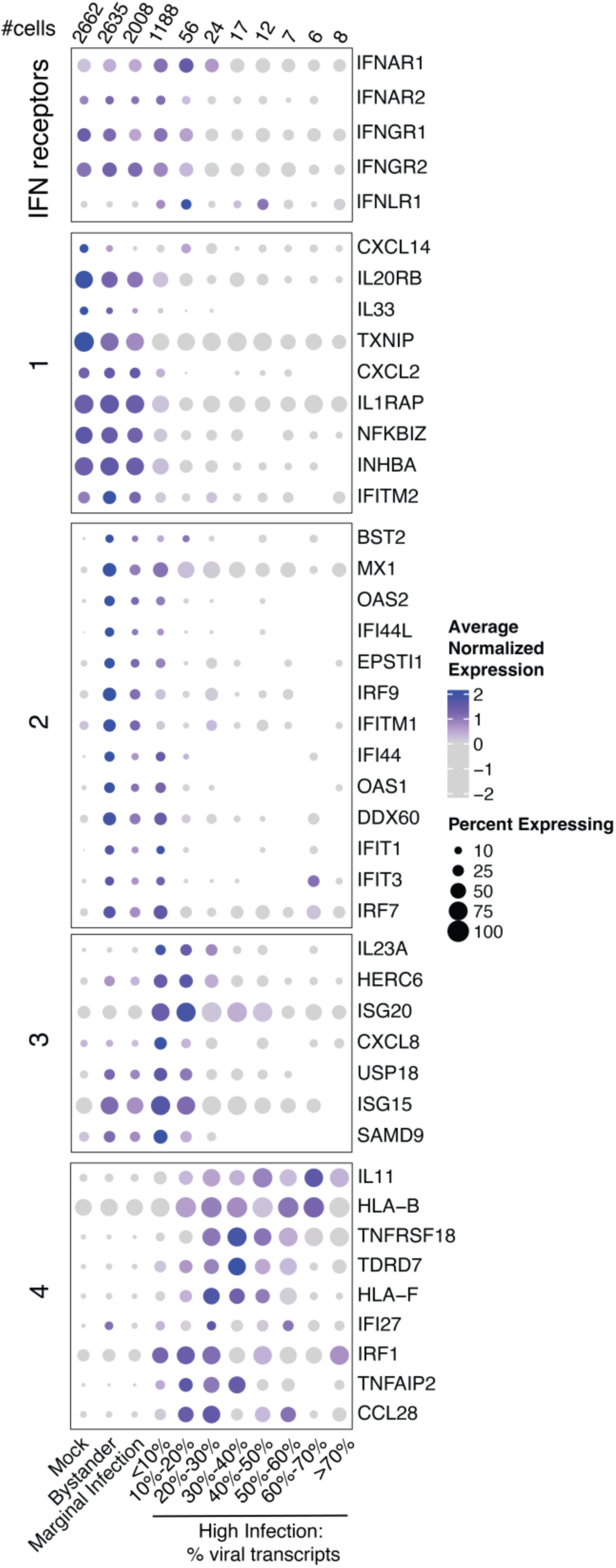
HCMV infection differentially induces interferon response in HOKs. Expression dotplot of significantly differentially expressed interferon responding genes and immune genes identified in comparisons in Fig. 5A**-C**.

HCMV infection differentially induced antiviral responses in the infected environment in HOKs. However, the expression patterns of these genes were different from those in previous scRNA-seq studies in fibroblasts and CD14+ monocytes (32, 33). We focused on ISGs and immune genes from the DEGs in comparisons presented in **Fig. 4**. These genes can be grouped into four groups based on their expression levels from cells at different infection stages (**Fig. 6C**). Group 1 genes, including CXCL2, CXCL14, NFKBIZ, were expressed highly in mock, bystander and marginal infection cells but were significantly downregulated in cells at high infection stage.

Group 2 genes including MX1, IRF7, IRF9, OAS1/2 and IFIT1,3 were specifically induced in the bystander cells but were downregulated when cells get infected. Expressions of group 3 and 4 genes were higher in highly infected cells compared to genes in group 1 and 2. Furthermore, group 4 genes including IRF1, IL11 and IFI27 maintained their expression as the cells became more heavily infected while group 3 genes which include ISG15, ISG20 and CXCL8 were downregulated in heavily infected cells. Our data suggest that HCMV infection successfully downregulated a group of ISGs and immune genes, but either failed to downregulate or selectively induced certain immune genes.

### HCMV infection drives intermediate HOK differentiation

Cellular differentiation has been shown to be critical for HCMV infection, replication, and/or reactivation in several contexts and HCMV has been shown to drive the differentiation process in multiple cell types, including monocyte-to-macrophage differentiation (34–36), neuronal differentiation (37, 38) as well as CD34^+^ hematopoietic progenitor cell differentiation (39). We therefore examined if HCMV infection of HOKs affected their differentiation state. In normal human oral mucosa - there are three general layers: the basal layer containing oral stem cells or basal HOKs from which the oral mucosa becomes regenerated; the intermediate layer composed of larger, stacked polyhedral shaped cells which have lost their ability to divide and which compromise the bulk of the oral epithelium; and the superficial layer which contains the terminal HOKs – which are highly cornified, flattened cells which undergo shedding as they age and die during tissue turnover (40). As cells progress from basal stem cells to terminally differentiated HOKs, there is a decrease in basal genes and an increase in intermediate genes, followed by an increase in terminal gene expression.

We observed that highly infected cells exhibited a marked downregulation of several epithelial subtypes present in mock-infected, bystander, and marginally infected cells (**SI Appendix, Fig. S6**). These included basal-like cells (clusters 1 and 2), proliferative basal HOKs at various stages of the cell cycle (clusters 3, 4, 7, 8, and 10), early differentiating/intermediate HOKs (cluster 5), and cornifying/late-differentiating HOKs (cluster 9). In contrast, a distinct population - dysregulated, intermediate-like HOKs (cluster 6) - emerged specifically in the highly infected cells and was nearly absent in the mock, bystander, and marginally infected groups (**Fig. 7A**). Indeed, when examining the expression of differentiation-related genes, we observed that HCMV downregulated basal keratinocyte genes - KRT5, KRT6A, KRT6B, KRT14, KRT16, KRT17, NOTCH1, and S100A2 (**Fig. 7B**). Concomitantly, HCMV infection induced an increase in select intermediate differentiation genes (KRT10, S100A4, and NOTCH3), and to a lesser extent some terminal differentiation genes (TGM2, SPRR2A, and SPRR1B), suggesting HCMV is driving HOK differentiation away from basal and towards a dysregulated intermediate state, while also slowing down terminal differentiation. Moreover, the top Gene Ontology (GO) terms downregulated in HCMV-infected cells were “cornified envelope”, “cell substrate junction”, and “apical part of cell” (**Fig. 7C**) - terms associated with terminal differentiation, indicating further that HCMV is diverting HOKs from the normal basal-to intermediate-to terminal differentiation towards dysregulated, intermediate-like HOKs.

**Fig. 7:**
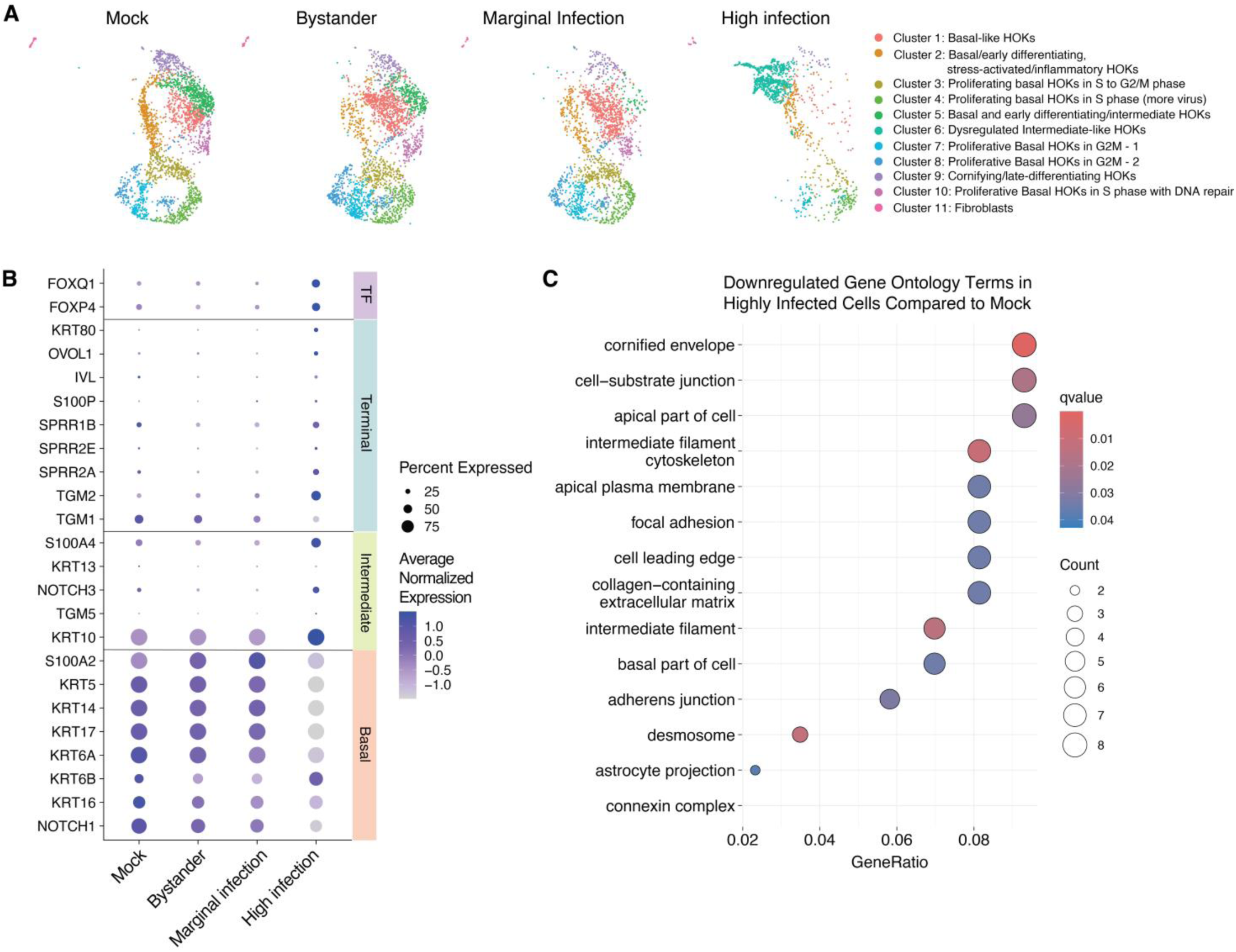
HCMV infection drives dysregulated intermediate HOK differentiation. **(A)** UMAP embedding of cells in different infection state, colored by identified cell population clusters. **(B)** Expressions of differentiation markers in HOKs. HCMV highly infected cells have higher expression of HOK intermediate differentiation markers. Intermediate markers KRT10, S100A4 and NOTCH3 are specifically upregulated in infected cells. S100P, SPRR1B, SPRR2A, and TGM1 are terminally differentiation markers. S100A4, KRT13, NOTCH3 and TGM5 are intermediate differentiation markers. S100A2, KRT5, KRT14 are basal epithelial cell markers. **(C)** Top gene ontology (GO) terms enriched in mock infected cells using DE analysis between mock and highly infected cells. Genes are associated with terminal differentiation terms such as cornified envelope.

Overall, our data indicate that upon infection in HOKs, HCMV drives a variety of host gene changes such as downregulation of basal (e.g. KRT5, KRT6A, KRT6B, and NOTCH1) genes, upregulations of intermediate differentiation genes (KRT10 and S100A4) and some terminal genes (TGM1) which together drives differentiation towards intermediate HOKs (**Fig. 8**). HCMV also induces the expression of genes encoding membrane proteins (e.g. FAM241B, TMEM125), molecular chaperones (e.g. CLU, AKAP12, and HSPA1A) which HCMV may be using for viral protein stability, virus assembly and egress; upregulates differentiation transcription factor FOXQ1 which likely plays a role in the differentiation process. We also identified HCMV downregulates transcription factors FOS, EGR-1, MYC, KLF-7, TP63, and ETS2, necessary for cell cycle and self-renewal of basal stem cells. Finally, we also determined HCMV induces an altered innate immune response downregulating a variety of immune genes (e.g. CXCL14, IL20RB, SERPINB3, and S100A8) while upregulating select immune genes (e.g. ISG15, IFI6, MX1, IFI27, and ISG20) potentially to create an inflammatory environment to aid its replication and/or recruit monocytes/myeloid cells to infect and spread in the body.

**Fig. 8:**
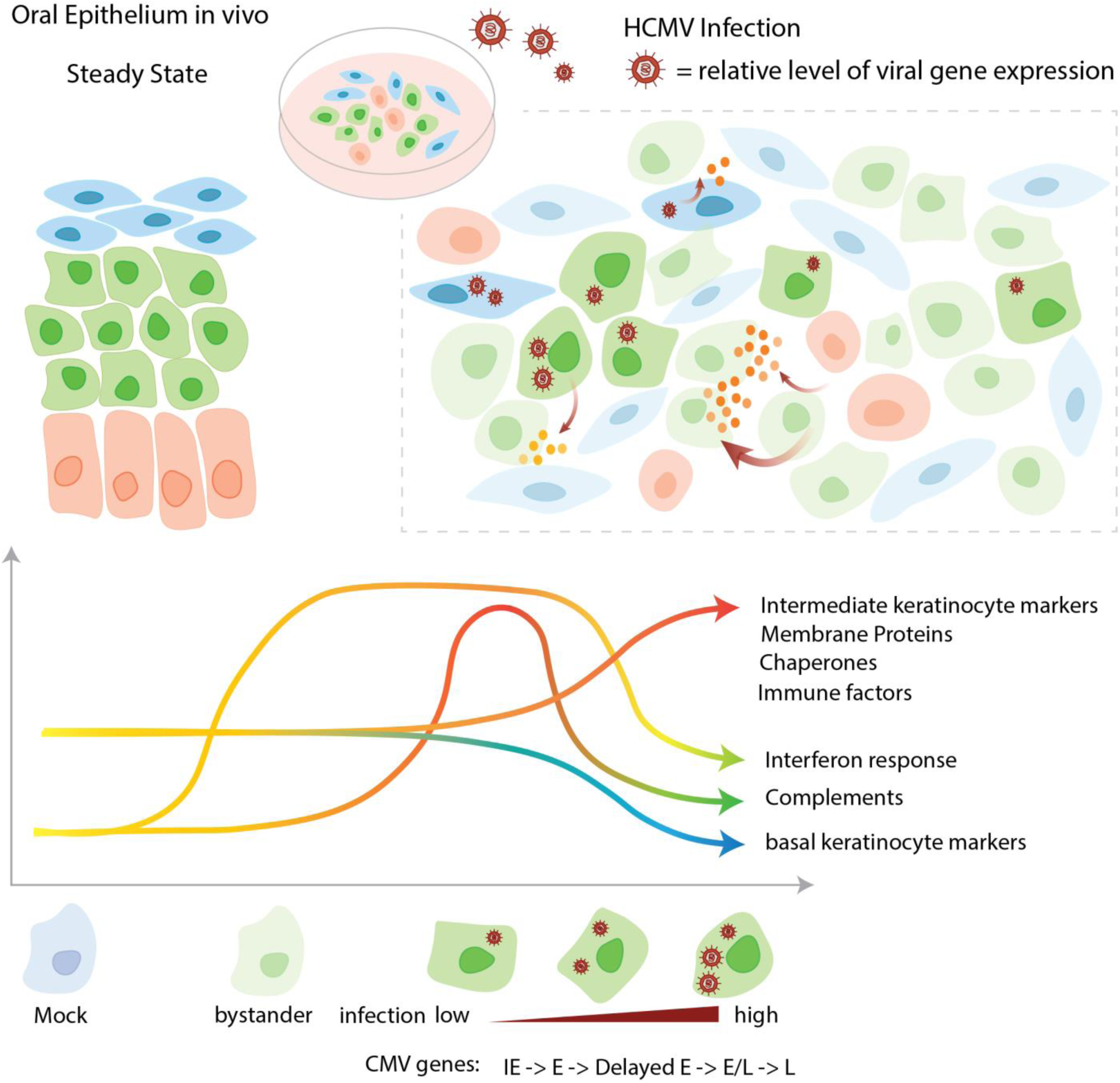
**Model of HCMV infection in HOKs**. Primary oral epithelium contains oral keratinocytes in basal, intermediate and terminal differentiation states. In primary HOK culture, we observed these three different states. HCMV infection directs the differentiation progress to intermediate state. In the HCMV infected HOKs, we observed upregulation of the expressions of intermediate markers, membrane proteins, chaperones and immune factors, while the expressions of basal markers, interferon responding genes, complement genes, and genes targeted by E2F and Myc are being downregulated. We also constructed HCMV temporal gene expression patterns that categorized genes into immediate early (IE), early (E), delayed early (Delayed E), early-late (E/L) and late (L).

## Discussion

Our study demonstrates that primary HOKs are permissive to HCMV infection and can serve as a robust model to explore early infection events in the oral mucosa. Through scRNA-seq and innovative visual analytics tools such as SciViewer, we characterized HCMV-induced transcriptomic alterations in HOKs, identifying significant modulation of cellular differentiation, innate immune responses, and cellular metabolism that likely facilitate viral replication and persistence.

The development of this model has broader implications for understanding the initial events of oral HCMV infection and its subsequent dissemination throughout the host. The oral mucosa represents a critical entry point for HCMV, yet specific interactions between the virus and oral epithelial cells remain poorly characterized. Our findings significantly contribute to the understanding of these early infection dynamics and may inform future therapeutic and preventative strategies targeting oral transmission.

Although we used a monolayer culture, we observed evidence of the cells being in the three main differentiation states of the layers in the oral mucosa - ranging from basal to intermediate to terminally differentiated oral keratinocytes – resembling those typically found in native oral tissues. This suggests that monolayers of primary HOKs can serve as a model of HCMV infection in the oral mucosa as they recapitulate the differentiation states observed in 3D tissues *in vivo*. Importantly, we observed similar HOKs gene expression patterns compared to other studies of oral mucosa with some differences. In Jung et al, the basal layer from 3D oral epithelial tissues expressed KRT5, KRT14, S100A2, intermediate layer expressed NOTCH3, S100A4, KRT13, TGM5, and the superficial layer was characterized by the expression of S100P, SPRR2A, SPRR2E (41).

Similarly, in our study, cluster 1 (basal-like cells) expressed KRT5, KRT14, KRT17, S100A2, and TP63. Our intermediate-like HOKs clusters, namely cluster 2 (basal/early differentiating, stress-activated/inflammatory HOKs), cluster 5 (mixed population of basal and early suprabasal/intermediate/spinous-like HOKs), and cluster 6 (dysregulated intermediate-like HOKs, largely present only in the infected samples) expressed some similar and some different gene signatures – namely KRT7, KRT6A, CAVIN3, IL1B, and IL1A for cluster 2; KRT16, KRT17, DSG3, DSC3, and PKP1, for cluster 5, and KRT10, KRT80, UBB, CLU, HSPA1A, and HAGH for cluster 6. Finally, Cluster 9 (cornifying/late-differentiating HOKs) expressed DSP, JUP, PPL, TGM1, S100A9, ANXA1, CTSA, CTSB, SPPR2A, SPRR2E, and SPRR1B. It is possible that the differences between Jung et al and this study arose from culture structure (two-dimensional for our study versus three-dimensional in Jung et al), or differences in the viruses used (HCMV vs. KSHV). Interestingly, Williams et al 2021 (42), who examined periodontic human oral biopsies by scRNA-seq and identified an epithelial cell population with an inflammatory signature (IL1B, SLPI, and S100A8/9) similar to our cluster 2. This suggests that the oral mucosa is an active tissue with subsets of cells showing enhanced proinflammatory or antimicrobial gene expression, consistent with its role as the first line of defense against pathogens entering the human body through the oral cavity. Overall, our study follows gene expression patterns in primary HOKs that align with various layers of differentiation indicating that this model can be used in future studies of HCMV and other viruses for which the oral mucosa is a key site of infection and transmission.

Somewhat surprisingly, the percentage of cells expressing viral transcripts decreased from 1 dpi to 3 dpi in infected samples as observed by scRNA-seq, indicating HCMV might not be very efficient in replicating in primary HOKs. Likewise, infectious virus yields were low and not detected until 6 dpi. This observation is in part consistent with a study by Morrison et al. which showed that HCMV fails to exhibit robust spread throughout salivary gland epithelial cells and persists in a low percentage of salivary cells (14). Interestingly, a study of HCMV infection in primary kidney and mammary gland epithelial cells determined that unlike fibroblasts, these cells are able to sequester HCMV at the plasma membrane and release it in the supernatant in the absence of viral replication at early timepoints, followed by viral replication at later timepoints (43). Our finding of low levels of HCMV replication in HOKs is also consistent with the fact that HCMV infections usually do not cause overt disease in the oral mucosa of healthy individuals though it can result in oral ulcers in immunocompromised patients. Another potential explanation for the decreased number of virus-positive cells between the two time-points is that a part of cells in infected samples could have died and may have been removed during the extensive washing and cell straining steps which are part of the scRNA-seq protocol.

In our UMAP clustering of cells, the presence of the fibroblast cluster was somewhat surprising, considering the cells had been specifically enriched for oral epithelial cells by the manufacturer. However, since these primary cultures are derived from an individual’s oral mucosa - an area naturally containing oral fibroblasts - their presence in small numbers is not entirely unexpected.

HCMV gene expression patterns in HOKs were largely similar to those observed in other productively infected cell types, such as fibroblasts and epithelial cells, although some differences were noted. For example, UL122, which is typically expressed with immediate early kinetics, clustered with the late genes in our dataset. This may indicate that UL122 is expressed with delayed kinetics in HOKs unlike other cell types, or that viral transcript levels fluctuate over the course of the three-day time course in primary HOKs, causing the trajectories we observed to not fully reflect the typical temporal phases of HCMV gene expression. Nonetheless, it is not entirely surprising that we identified some differences compared to established patterns of HCMV gene expression. Multiple previous studies have been conducted in fibroblasts or other cell lines, whereas our study focused on primary cultures of HOKs. Furthermore, even among cell lines, it has been shown before that differences exist in HCMV expression in fibroblasts, epithelial cells, and astrocytes (44). Nevertheless, our study contributes valuable insights to the current understanding of HCMV gene expression in productively infected primary cells.

HCMV infection induced extensive changes in the host transcriptome in primary HOKs. HCMV infection upregulated select immune genes while downregulating others, suggesting infection is modulating the immune response to avoid immune recognition while creating an inflammatory environment which has been shown to benefit the virus. HCMV infection also induced molecular chaperones and vesicular trafficking genes, which may be used for viral protein stability, virus assembly and egress. HCMV infection also upregulated a variety of differentiation transcription factors, with FOXQ1 being one of the top upregulated genes.

Interestingly, KSHV has been recently shown to highly upregulate FOXQ1 in primary human gingival epithelial (HGEP) cells and in that model, FOXQ1 was critical for sustaining KSHV lytic infection (45). It is possible that HCMV also uses FOXQ1 to induce viral replication in primary HOKs. Alternatively, FOXQ1 has been shown to be important in oral epithelial cell differentiation (46), thus FOXQ1 may also play a role in the HCMV-induced differentiation process.

HCMV infection downregulated a variety of transcription factors necessary for cell cycle progression and self-renewal of basal stem cells relative to uninfected cells, which is consistent with previous findings that HCMV alters the cell cycle, and with our observation that the virus drives HOK differentiation away from the basal state toward a dysregulated intermediate state. Among the downregulated transcription factors were JUN and FOS, which together form the Activator Protein-1 (AP-1) transcription factor complex involved in regulating cell proliferation, differentiation, apoptosis, and inflammation. One of their downstream targets, MYC, was also downregulated. Interestingly, early work showed that FOS, JUN, and MYC are rapidly induced by HCMV in human embryonic lung cells (47), while more recent studies reported that HCMV infection of ARPE-19 cells results in dampened AP-1 activity (48). Krishna et al. demonstrated that FOS expression and activity are attenuated by the HCMV US28 protein, facilitating latent infection in myeloid cells (49), and that AP-1 activity is critical for HCMV reactivation from latency in these cells (50). While it is not clear why HCMV downregulated JUN and FOS expression in primary HOKs which sustained a productive HCMV infection, FOS has been shown to play a role in both basal and suprabasal layers of skin keratinocytes (51), as well as in the late stages of epithelial differentiation just prior to cornification and cell death (52). Thus, it is possible that HCMV downregulates FOS in HOKs to drive cells away from the basal state but also prevent full terminal differentiation, thereby promoting a cell differentiation state that is optimal for viral replication. Importantly, disruption of FOS has been implicated in craniofacial anomalies (53), and HCMV is known to cause microcephaly and other neurodevelopmental disorders in congenital infections. It is therefore possible that HCMV contributes to these outcomes, at least in part, by downregulating FOS. Interestingly, although our study was conducted in primary HOKs rather than neurons, we observed upregulation of several genes involved in neuronal and synaptic regulation following HCMV infection. While it is not yet clear why HCMV may be upregulating these genes in oral epithelial cells, our findings may provide insight into how HCMV disrupts normal neuronal and craniofacial development.

HCMV also downregulated EGR1 in HOKs (**Fig. 4A and SI Appendix, Table S3**). Interestingly, HCMV miR-US22 has been shown to downregulate *EGR1* in CD34⁺ hematopoietic progenitor cells (HPCs), thereby blocking HPC self-renewal and proliferation and promoting a differentiation pathway necessary for HCMV reactivation (39). Therefore, it is possible that HCMV downregulates *EGR1* in HOKs to drive cells away from the basal stem cell state and toward the intermediate HOK phenotype that supports viral replication. It will be interesting to determine whether viral microRNAs are also involved in this process in HOKs.

We observed significant upregulation of autophagy-related genes (ATG101, MAP1LC3A, DNASE2, HSPA13, NFKBIB) in highly infected cells as viral transcripts increase (**Fig. 5**). HCMV is known to regulate autophagy, a vesicular pathway critical for the degradation and recycling of cellular components (54). Previous studies in fibroblasts have demonstrated HCMV manipulation of autophagy to optimize cytoplasmic envelopment of mature viral particles.

Some evidence suggests early HCMV infection inhibits autophagy, while later stages reactivate it to support viral egress. Thus, the observed upregulation of these autophagy-related genes in our directional analysis may indicate HCMV-induced activation of late-stage autophagy in HOKs, potentially facilitating viral trafficking, assembly, or release.

We identified marked downregulation of IRF6 as cells progressed from intermediate (0.01%) to highest viral transcript levels (82.55%) (**Fig. S4**). IRF6 has previously been implicated in promoting differentiation of oral keratinocytes in response to pathogens such as *P. gingivalis*, regulating critical differentiation markers (IVL and KRT13) and transcriptional regulators (GRHL3 and OVOL1) (55). Downregulation of IRF6 in highly infected HOKs aligns with our observations that HCMV drives cells away from the basal state toward an intermediate-like phenotype, likely creating a differentiation state conducive to viral replication. This finding underscores the precision with which HCMV targets host differentiation pathways to support its lifecycle.

We also observed extensive modulation of host immune responses. HCMV-mediated the downregulation of critical antiviral and inflammatory pathways, including interferon signaling (IFNGR1, LY6E, STING1), complement system (C3), and pyroptosis (PYCARD, CASP4), and significant upregulation of immune evasion genes such as CD55. C3 is a critical protein of the complement system which is a central protein across all three complement pathways. None of the three complement pathways can function without C3, because it is a key protein that is cleaved to generate active components. C3 plays a crucial role in opsonization, a process that marks pathogens and infected cells for destruction by phagocytes. C3 is critical to antibody-mediated complement lysis for elimination of virus-infected cells (56). CD55’s marked upregulation in highly infected cells underscores its role in protecting infected cells from complement-mediated destruction, aligning with prior studies demonstrating its critical function in viral immune evasion (57, 58). Additionally, CD55 has been previously implicated in facilitating viral entry into biliary epithelial cells through CD14, further emphasizing its importance in HCMV pathogenesis (59). Directional analysis 2 also identified extensive changes in cell surface proteins as we looked at gene expression changes from cells with intermediate number of viral transcripts (0.01%) to cells with highest number of viral transcripts (82.55%).

Several cell surface proteins critical for epithelial integrity, cell-cell adhesion, and immune cell interactions – including CD9, CD24, CD47, CD81, and CD82 – were markedly downregulated. These findings suggest active remodeling of the cell surface proteome by HCMV, potentially disrupting epithelial barriers, facilitating immune evasion, and promoting viral persistence and dissemination. Recently, it has been shown that virus entry efficiency is a major determinant of HCMV latency in monocytes and when a glycoprotein from fibroblasts was ectopically expressed in monocytes the virus would do a productive lytic infection (60). This demonstrated that cell surface proteins play a critical role in efficient viral entry, which can then dictate whether a lytic infection happens. Given the cell surface protein gene expression changes that we observe in HOKs, it would be a potential next step to explore whether the cell surface changes driven by virus in HOKs allow efficient entry and productive infection.

Moreover, our directional and gene set enrichment analyses revealed significant HCMV-driven metabolic reprogramming in highly infected cells. Notably, pathways associated with oxidative phosphorylation, fatty acid metabolism, and adipogenesis were strongly downregulated in cells with high viral transcript levels. The observed suppression of oxidative phosphorylation genes (such as DECR1, UQCRB, ATP5ME, NDUFS4, COX7C, and UQCRC1) suggests mitochondrial dysfunction, potentially impairing the energy production required for antiviral responses.

Similarly, decreased expression of fatty acid metabolism (DECR1, LGALS1, SUCLG2, SDHC) and adipogenesis-related genes (APLP2, NDUFA5, PDCD4, DBT) indicates a metabolic shift that likely favors the biosynthetic and structural requirements of viral replication over host cellular needs. Previous studies have shown that HCMV infection upregulates oxidative phosphorylation; in our study, we found that as 3 dpi infected cells have more viral transcripts, they have downregulation of oxidative phosphorylation (61). It is possible that oxidative phosphorylation is upregulated earlier on in infected cells before 3 dpi so that the energetic demands of virus replication are met or downregulation of oxidative phosphorylation in infected cells could be a factor that is either aiding or stifling the viral replication process. This is a research area that would need to be elucidated in future experiments. Aside from oxidative phosphorylation, the fatty acid and adipogenesis metabolic alterations align with prior studies documenting HCMV’s capacity to modulate host energy metabolism and redirect cellular resources toward viral assembly and maintenance, underscoring another critical dimension of viral-host interaction during infection (62).

SciViewer enabled directional analysis of UMAP trajectories, revealing subtle, viral-transcription-load-dependent changes that conventional DE/GSEA could not resolve. This interactive approach complemented existing pipelines by allowing user-guided exploration of continuous patterns, rather than relying solely on binary contrasts between infection states (63). In doing so, Sciviewer uncovered graded shifts in immunity, differentiation, and metabolism that became apparent only when cells were analyzed along a viral-load continuum. While pseudotime and trajectory inference methods are crucial for understanding dynamic biological processes (64,65), they typically generate rigid, algorithm-driven orderings that leave little room for biological intuition. SciViewer fills this gap by providing an intuitive framework for visually tracing progressive changes in embedding space and directly linking them to gene- or pathway-level outputs. This “directional search” approach highlighted additional candidate processes - including oxidative phosphorylation, innate immunity, epithelial maturation, and cell-surface remodeling - that merit functional validation in future work.

Our data presents a map of differentially downregulated immune genes in HCMV infected HOKs with different number of viral transcripts. We observed genes that are entirely downregulated in infected cells regardless of the amount of virus (**Fig. 6**), suggesting the strategy of HCMV in promoting its own survival from alerting the immune system. HCMV inhibits the production of inflammatory chemokines such as CXCL2 and CXCL14 and cytokines such as IL33, as well as the IL20 receptor IL20RB. HCMV utilizes the cellular Roquin to target genes including CXCL2 to evade host antiviral mechanisms (66). Though HCMV’s interference on CXCL14 production has not been indicated before, studies have shown that human papillomaviruses (HPV) are able to suppress the expression of CXCL14 to evade the host response (67), suggesting HCMV might utilize similar mechanisms. Regulatory T cells (T-regs), promoted by IL33, have been shown to confer protection from HCMV and MCMV infection (68). Downregulation of IL33 by HCMV has not been indicated previously yet has been found in cells infected by MCMV (69). HCMV also interferes with the TGFb family signaling pathway by inhibiting INHBA expression (70). We also observed HCMV in general downregulates interferon induced genes such as IRF7, IRF9, IFIT1, IFIT3, OAS1, OAS2, MX1, ISG15 and ISG20. On the contrary to the general trend of downregulation, a group of genes appeared with persistent expression when viral gene expression increased. Within these genes, IL11 maintained its expression in highly infected cells. IL11 was found to be induced in endothelial cells by HCMV infection, which may enhance survival of the infected host from apoptosis (71). It is possible that a similar mechanism is utilized by HCMV in epithelial cells, which needs to be further experimentally verified. Our immune profiling revealed extensive modulation of host responses, including upregulation of MHC molecules such as HLA-B and HLA-F in infected cells, suggesting potential interactions with cytotoxic T cells and immune regulation.

We observed that HCMV drives differentiation away from basal and towards a dysregulated intermediate phenotype, while stalling their progression towards terminal differentiation. It is possible that HCMV drives differentiation away from basal HOKs as stem-like properties have been shown to be unfavorable for HCMV replication (72–75). Additionally, it is plausible that HCMV delays the progression toward terminal differentiation to ensure the cells are conducive to its replication cycle. As human oral keratinocytes (HOKs) transition from intermediate to terminal stages, they lose their nuclei and undergo apoptosis, thereby forming a protective keratinized layer that is subsequently shed and replenished by newly cornified cells. If this differentiation pathway proceeds unimpeded, HCMV may not have sufficient time to complete its replication cycle before the host cells lose their nuclei and undergo apoptosis. Interestingly, HCMV downregulated most basal genes except KRT6B which was upregulated in a greater proportion of infected cells compared to mock-infected, bystander, and marginally infected cells. KRT6B plays an important role in wound-healing and cell migration and has been shown to be upregulated in some cancers, therefore HCMV might be upregulating it to promote migration of infected cells towards other non-infected cells to transmit the virus. Similarly, HCMV selectively upregulated certain intermediate and terminal differentiation genes, while others - such as the intermediate gene KRT13 and terminal genes TGM1, SPRR2E, and S100P - were not upregulated. This suggests that HCMV promotes specific aspects of intermediate and terminal differentiation that may favor its replication, while avoiding others that could be detrimental to its lifecycle.

Other viruses, such as human papillomavirus (HPV) and Kaposi’s sarcoma-associated herpesvirus (KSHV), have been shown to exploit epithelial cell differentiation to facilitate their replication. For example, productive HPV infection depends on the differentiation of basal cells into terminally differentiated cells in the cornified layer, while KSHV exhibits increased lytic gene expression in the more terminally differentiated layers of the epithelium (76–78). HCMV appears to follow a similar strategy; however, instead of promoting full or terminal differentiation of HOKs, it induces a dysregulated, intermediate-like differentiation state. We propose that this altered state - occurring just prior to terminal differentiation and cell death - creates a favorable environment for HCMV replication and spread.

In conclusion, our comprehensive study provides insights into the intricate strategies employed by HCMV to establish a productive infection in HOKs by manipulating their differentiation, immune responses, and metabolism. Our data suggests that HCMV uses these changes in order to create an environment suitable for viral replication in the oral mucosa and spread into the rest of the host. These insights offer promising directions for future research, including functional studies to validate the roles of identified genes in viral pathogenesis and developing targeted interventions to prevent oral transmission and systemic dissemination of HCMV. Also, we demonstrate how novel computational tools enabling rapid exploration of patterns in single-cell datasets can lead to biologically meaningful hypotheses-generating findings.

## Materials and Methods

### Cells and viruses

Primary human oral keratinocytes (HOKs) were isolated from human oral mucosa, cryopreserved at passage one, and delivered to us frozen (ScienCell Research Laboratories, Cat No 2610, Carlsbad, CA). Upon arrival, cells were stored in liquid nitrogen until needed for experiments. Upon thawing, HOKs can expand for an additional 10 population doublings. Cells were thawed and cultured in oral keratinocyte medium (OKM, Cat. No. 2611, ScienCell Research Laboratories, Carlsbad, CA) as per the product sheet instructions.

To increase infectivity in HOKs, we used the TB40E-Gfp strain of HCMV (generously provided by Dr. Gary Chan at SUNY Upstate Medical University) which expresses enhanced green fluorescent protein (eGfp) from the SV40 early promoter and is known to express the gH/gL/UL128-131 pentameric complex necessary for infection in epithelial cells (79)((80).

Considering that the pentameric complex can be quickly lost if HCMV is grown or passaged in fibroblasts (81), we adapted TB40E-Gfp to epithelial cells through eight passages in retinal pigment epithelial cells (ARPE-19, CRL-2302, American Type Culture Collection, Manassas, VA). ARPE-19 cells from a low passage number (passage 9 [P9] to P15) were subcultured in

Dulbecco’s Modified Eagle Medium Nutrient Mixture F12 (Ham) (DMEM/F12; 11330-032, Gibco, ThermoFisher, Grand Island, NY) with 10% fetal bovine serum (FBS; Cat 26140-079, Gibco). To expand viral stocks, ARPE-19 cultures were infected with TB40E-Gfp passage 7 in DMEM/F12 with 4% FBS. When the cultures were covered by cytopathic effect (CPE), supernatants were collected and spun at 4000rpm for 20min to remove cell debris. The virus was then further purified on a 20% sorbitol cushion by ultracentrifugation using SW28 rotors (Beckman Coulter) at 26,000 rpm resuspended in DMEM/F12 medium. The TB40E-Gfp stocks were titered in ARPE-19 cells by Gfp positivity at two days post infection.

We also used HCMV MOLD strain (generously provided by Dr. William Miller at the University of Cincinnati) - a primary low-passage HCMV isolate shown to be particularly tropic in salivary epithelial cells and endothelial cells (14, 19, 82). MOLD virus was expanded in human embryonic lung (HEL) 299 fibroblasts (CCL-137; American Type Culture Collection, Manassas, VA). HEL cells from a low passage number (passage 7 [P7] to P15) were subcultured in Dulbecco modified Eagle medium (DMEM; 11995-065, Gibco, ThermoFisher, Grand Island, NY) with 10% fetal bovine serum (FBS; Gibco). When the culture reached confluence, the cells were infected with MOLD in DMEM with 4% FBS. When the cultures were covered by CPE, virus was purified from the supernatant on a 20% sorbitol cushion by ultracentrifugation using SW28 rotors at 27,000 rpm and resuspended in DMEM/F12 medium (Gibco). The MOLD stocks were titered in ARPE-19 cells by plaque assay. A multiplicity of infection (MOI) of 0.5 or 2 was used depending on the experiment based on HCMV titers in ARPE-19 cells. Mock infection was performed by adding an equivalent volume of DMEM/F12 medium to HOKs.

### Microscopy

Primary cultures of HOKs were either mock- or HCMV-infected with TB40E-Gfp or MOLD at MOI of 0.5 (based on HCMV titers in ARPE-19 cells). Live cell imaging was performed in phase and Gfp in a Sartorius Incucyte SX5 microscope at 1, 3, 5, and 8 dpi using a 20x objective in the Sanderson Center for Optical Experimentation at UMass Chan Medical School. Gfp and phase intensities were normalized for all images using Fiji software.

### Viral Growth Curves

Primary human oral keratinocytes (HOKs) were HCMV-infected with the TB40E-Gfp strain or the MOLD strain at an MOI of 2 (based on HCMV titers in ARPE-19 cells) for an 8-day time course and supernatants were collected every day, and fresh media was added after supernatant collection. The experiment was performed in triplicate. The virus titer in supernatants was quantified on human embryonic lung (HEL) fibroblasts through a modified shell assay using the Quidel D^3^DFA Cytomegalovirus Immediate Early Antigen identification kit. Specifically, the assay was performed in 96 well plates instead of shell vials. HEL cells were seeded in 96 well plates in DMEM. To avoid edge effect often observed in 96 well plates (83), the plates were left at room temperature for 1h for the cells to settle prior to moving the plates into a humidified incubator set to 37C and 5% CO2 as previously described (84). The following day, the HEL cells were infected with supernatants from HOKs diluted at 1:100 in DMEM. The infections were performed in quintuplicates. Images were taken on an ImageXpress Micro XL system and IE positive nuclei were quantified using the MetaXpress software. Numbers from mock-infected wells (cells treated with DMEM media mixed with HOKM media at 1:100) were subtracted from numbers in infected wells.

### scRNA-seq experiments

Primary HOKs at population doubling ∼7-9 were either mock- or HCMV-infected with TB40E-Gfp or MOLD at MOI of 0.5 (based on HCMV titers in ARPE-19 cells) and processed for scRNA-seq after 1 dpi and 3 dpi aiming to sequence ∼1000 cells per sample. Libraries were prepared using the 10X Genomics Chromium Next GEM Single Cell 3ʹ Reagent Kits v3.1 (Dual Index) and sequenced on the Illumina NextSeq2000 using a high throughput flow cell as per manufacturer instructions. The experiment was performed in duplicate, and the libraries were sequenced on the same flow cell to avoid batch effect.

### Bioinformatics Analyses

#### scRNA-seq data processing and quality control

Pooled libraries were demultiplexed using bcl2fastq (v1.8.4) with default settings. Raw reads were processed with Cellranger v3.0.1 with the command *--include-introns = false* (85). A custom reference genome was created by combining GRCh38, an EGFP DNA sequence and a custom HCMV transcriptome assembly described in Hein and Weissman, 2021 (33).

We obtained ∼78 - 142 million (mln) reads per sample, yielding ∼84,000 – 230,000 median reads per cell with a ∼70-78% alignment rate (**SI Appendix, Table S2**). The number of cells sequenced per sample ranged from 572 to 1129 cells per sample.

SoupX (86) was applied to each sample to account for ambient RNA contamination with custom settings (*tfidfMin=0.8,soupQuantile=0.8)* except for two samples: Exp214_Mock_1dpi and Exp213_TB_3dpi. For these two samples, SoupX was applied using the custom setting: *tfidfMin=0.7,soupQuantile=0.7.* The contamination rates were estimated to be 0.02-0.08% in the samples. The aligned and SoupX corrected cell-by-gene matrices were merged across all samples. The downstream analysis was performed using Seurat v5 (87) in R v4.4.0 environment (88). Plots were generated using ggplot2 (89). Cells with high mitochondrial gene content (more than 25% of the transcriptome derived from mitochondrial genes) and less than 2500 expressed genes were removed. The final dataset contains 20388 genes and 8623 cells across 12 samples (524-922 cells per sample, **SI Appendix, Table S2**). On average, mock-infected samples had a smaller percentage of cells was removed (∼3-10% of cells/sample) whereas virus infected cells had a greater percentage of cells removed - 11-14% for TB40E-Gfp infected samples and 4-20% for MOLD infected samples. It is not entirely surprising that HCMV infected samples had a higher percentage of dead and dying cells as infected cells likely upregulate intrinsic antiviral defense pathways resulting in apoptosis as early as 1 dpi.

We identified several clusters of cells in various stages of cell division (clusters 3, 4, 7, 8, and 10). Importantly, even though the cell cycle has been shown to have a significant effect on gene expression (63, 90) and has been reported to be an important confounding factor of cell subpopulation identification (87)(86)(91, 92) we did not observe any major differences in clustering when cell cycle genes were excluded versus included prior to UMAP analysis (**SI Appendix, Fig. S2**). Therefore, we did not exclude cell cycle genes from our analyses.

The raw data were then normalized using log2 normalization. Dimension reduction was performed using Principal Component Analysis (PCA) on the top 2000 variable genes. The variable genes were selected using the *vst* method in Seurat. HCMV genes were excluded from the variable genes. 20 PCs were used to cluster cells using Seurat’s graph-based clustering method. Cells were embedded in 2-dimensional space using UMAP (93) and 11 clusters were found.

Cluster marker identification was performed using MAST algorithm through Seurat with default settings. Marker gene list for each cluster were ordered by increasing *p.adj*, decreasing *pct.1* and decreasing *ave_log2FC*. HCMV genes were removed from the final marker gene list.

Differential gene expression analyses for **Fig. 4** were performed using edgeR algorithm through custom code. HCMV genes were removed from the tested cells. Only genes that were expressed in at least 10% of the tested cells were kept in the analysis. Significantly differentially expressed genes were defined as those that had log2 fold change > 1 or < -1 and false discovery rate < 0.05.

HCMV gene expression trajectories were constructed only for HCMV genes that had more than 70th percentile mean and standard deviation. In the heatmap representation of gene expression trajectory along increasing viral load, cells were binned by 5% of log10 (% HCMV transcripts) increments. The 80-85% bin only contained one cell. This cell was included in the >75% bin. Raw counts of HCMV genes in each bin were averaged, and relative expressions to the bin containing the max average expression were taken. Genes were clustered using k-means with k = 5. The expression trajectories were aggregated for each cluster, and the smoothed curves were generated using local polynomial regression fitting from geom_smooth (method = stats::loess) in ggplot2.

#### SciViewer Analyses

An AnnData file version of the Seurat object was generated using SeuratDisk package and its *Convert()* function (94) in R environment. Following that, AnnData object was loaded into a Jupyter Notebook in Python environment. SciViewer was run by initializing a SciViewer object with the 2D UMAP embedding and the expression data and then running the *explore_data* method (28). The visualizer then appeared where the user can interactively analyze the UMAP embedding.

The AnnData object was filtered to remove viral genes and only looked at host genes. It was further filtered to keep only 3 dpi infected cells. A directional analysis mode was selected in the visualizer, and then an arrow was drawn across the embedding that started from cells with the lowest number of viral transcripts and ended up where cells had the highest number of viral transcripts. The rectangle area around the arrow was selected to specify which cells to include in analysis. The SciViewer object was exported back to the notebook and the visualizer was closed by clicking the EXPORT & CLOSE button in the visualizer. This SciViewer object retained the directional analysis results which contained the gene name, Pearson correlation coefficient and p-value for every gene transcript. By default, the list of genes was sorted from highest positive value to lowest negative value of Pearson correlation coefficient. This list of genes (**SI Appendix, Table S4 and Table S6**) was then used to run gene set enrichment analysis.

#### GSEA/GO Analyses

Gene set enrichment analysis (GSEA) was performed using the GSEApy (95) library in Python environment. For GSEA analysis comparing highly infected cells with increasing amount of viral transcripts, ranked gene lists derived from SciViewer (ranked by Pearson correlation coefficients from positive to negative) were used directly as input (**SI Appendix, Table S4 and S6**). For GSEA comparing cells in different infection stages, ranked gene lists were derived from DE analysis comparing cells in specific infection stages (**SI Appendix, Table S3**). The MSigDB Hallmark 2020 gene sets served as the reference database for both analyses, with enrichment analysis parameters set to a minimum gene set size of 3 and a maximum of 500. Significant gene set terms were defined by a False Discovery Rate (FDR) q-value <= 0.1.

Gene ontology analysis was performed using *enrichGO()* function from clusterProfiler (96) in R environment. Cellular component (CC) ontology terms were tested. Significant terms were considered to have q-value < 0.05.

#### Data, Materials, and Code Availability

### Data and Code Availability

Raw sequencing fastq files and the final count tables will be reposited in GEO repository soon. Codes for recreating the analyses and plots can be found in https://github.com/yumingcao/HCMV_HOK_scRNA-seq and https://github.com/askartemir/HCMV_HOK_scRNA-seq.

## Acknowledgements

We thank current and past members of the Kowalik lab - Anne Mirza, John Holik, and Xiaofei E, for support and useful discussions. We thank Dr. William Miller and his group at the University of Cincinnati for the MOLD strain and Dr. Gary Chan and his laboratory members at SUNY Upstate Medical University for the TB40E-Gfp strain. We also thank the following cores, labs, and individuals at UMass Chan Medical School for help with troubleshooting experiments: Christina Baer and Jill McConnel in the Sanderson Center for Optical Experimentation (SCOPE); Carol Schrader and Tammy Krumpoch in the flow cytometry core; the Molecular Biology Core Labs; Jennifer Wang lab; Doyle Ward lab, Kotcha Mangkalaphiban in Allan Jacobson lab, Chris Sassetti lab, Sam Behar lab, Sambra Reddick in the Greiner lab, and the Department of Microbiology. We also thank our funding sources - NIAID grants R21AI154168, P01AI129859, and R56AI159672 to TK; and NIH IMSD grant T32GM135751 to AC and AT.

## Author Contributions

OC and TK designed the research; OC performed the experiments; YC, AT, and AC performed the scRNA-seq bioinformatics analyses; AT and AC performed the SciViewer analyses; OC, YC, AT, AC, and TK wrote the paper; OC, YC, AT, AC, MG, and TK edited the paper.

## Supplemental Figures

**Fig. S1:**
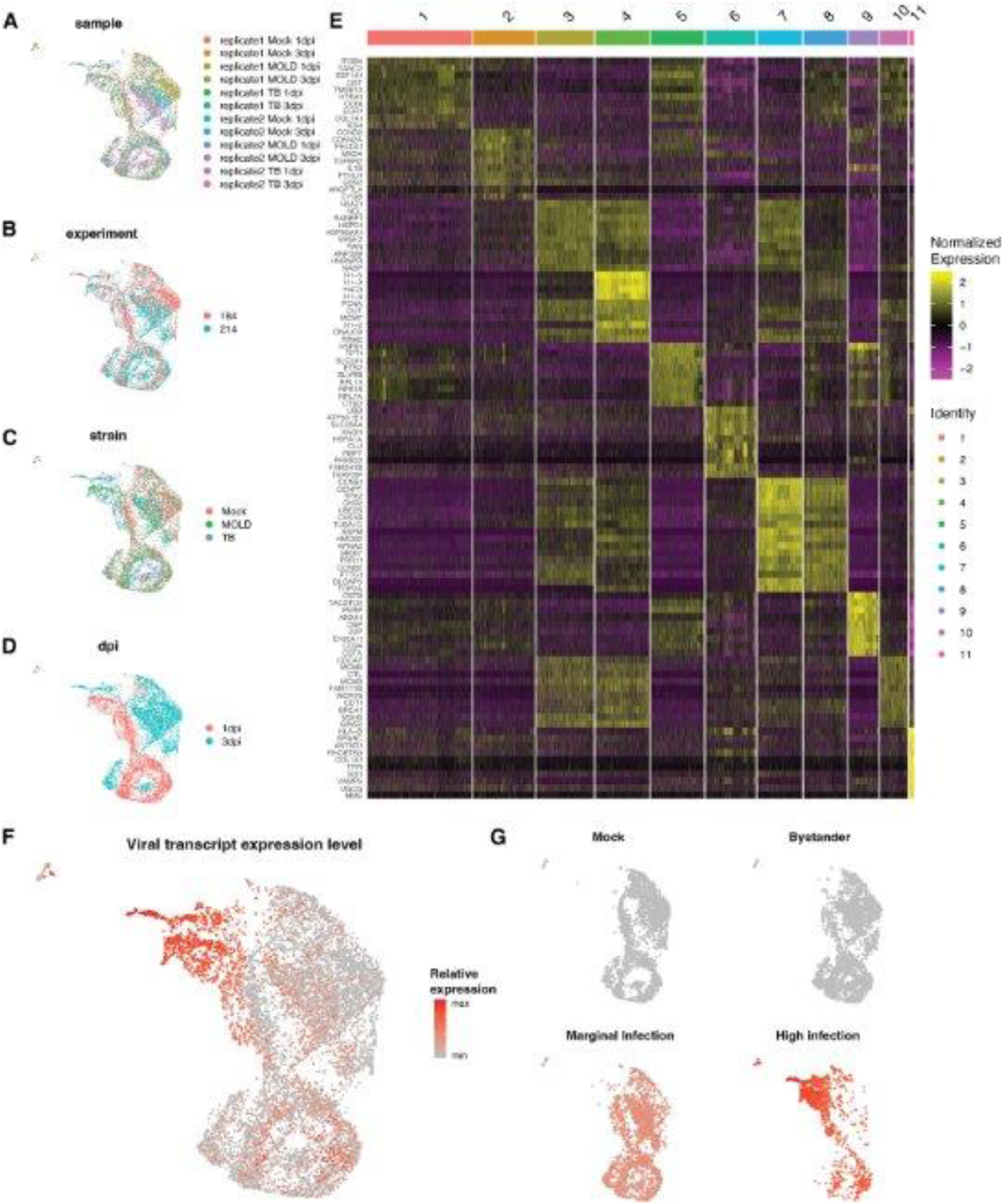
Clustering of cells and top marker genes of each cluster. **(A)** Cells were colored by collection sample, **(B)** experiment, **(C)** viral strain, and **(D)** days post infection (dpi). **(E)** Single-cell expression heatmap of top 10 markers for each cluster. Markers were identified as differentially expressed host genes between the tested cluster against the rest of cells in the data. **(F)** Cells were colored by viral transcript abundance. **(G)** UMAP embedding split by infection stages: mock, bystander, marginal infection and high infection.

**Fig. S2:**
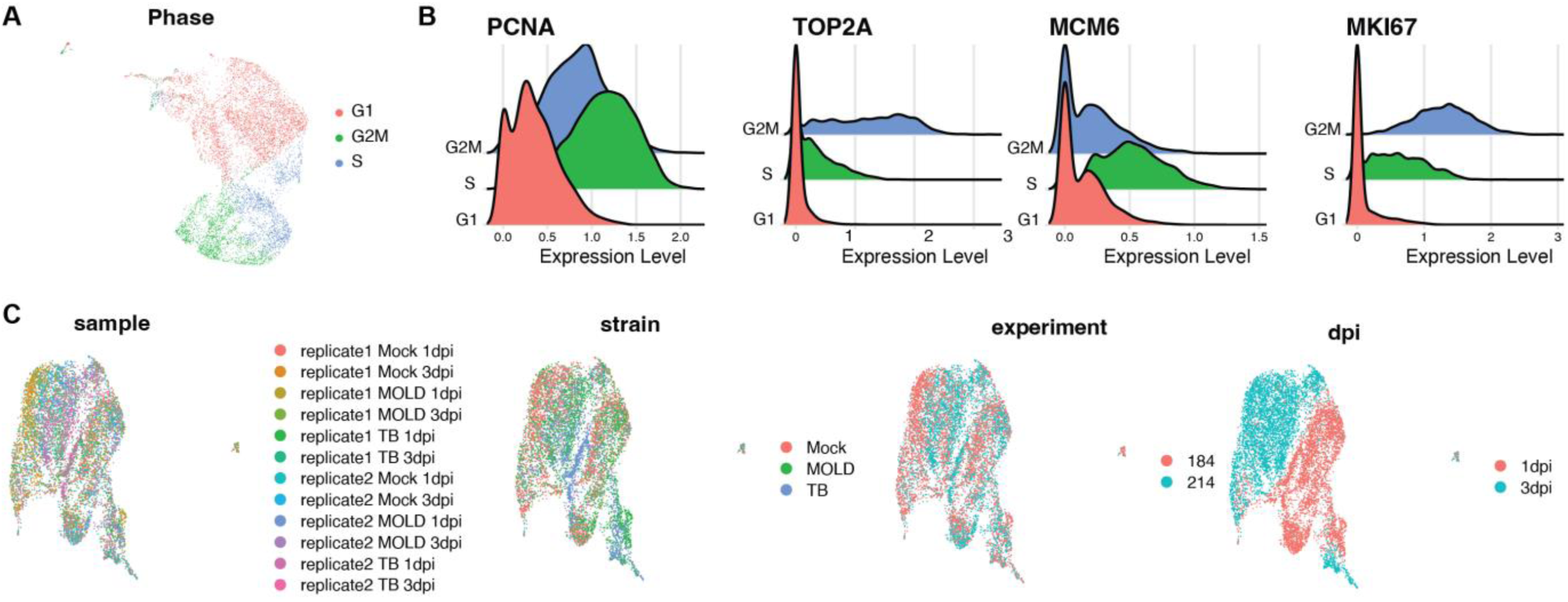
Cell cycle phase determination. **(A)** Cells were colored by the assigned cell cycle phase based on the expression of G2/M and S phase markers. **(B)** Expression of selected markers of cells in each cell cycle phase. **(C)** The dataset was rescaled by regressing out the cell cycle score. The new UMAP is colored by collection sample, viral strain, experiment, and days post infection (dpi).

**Fig. S3:**
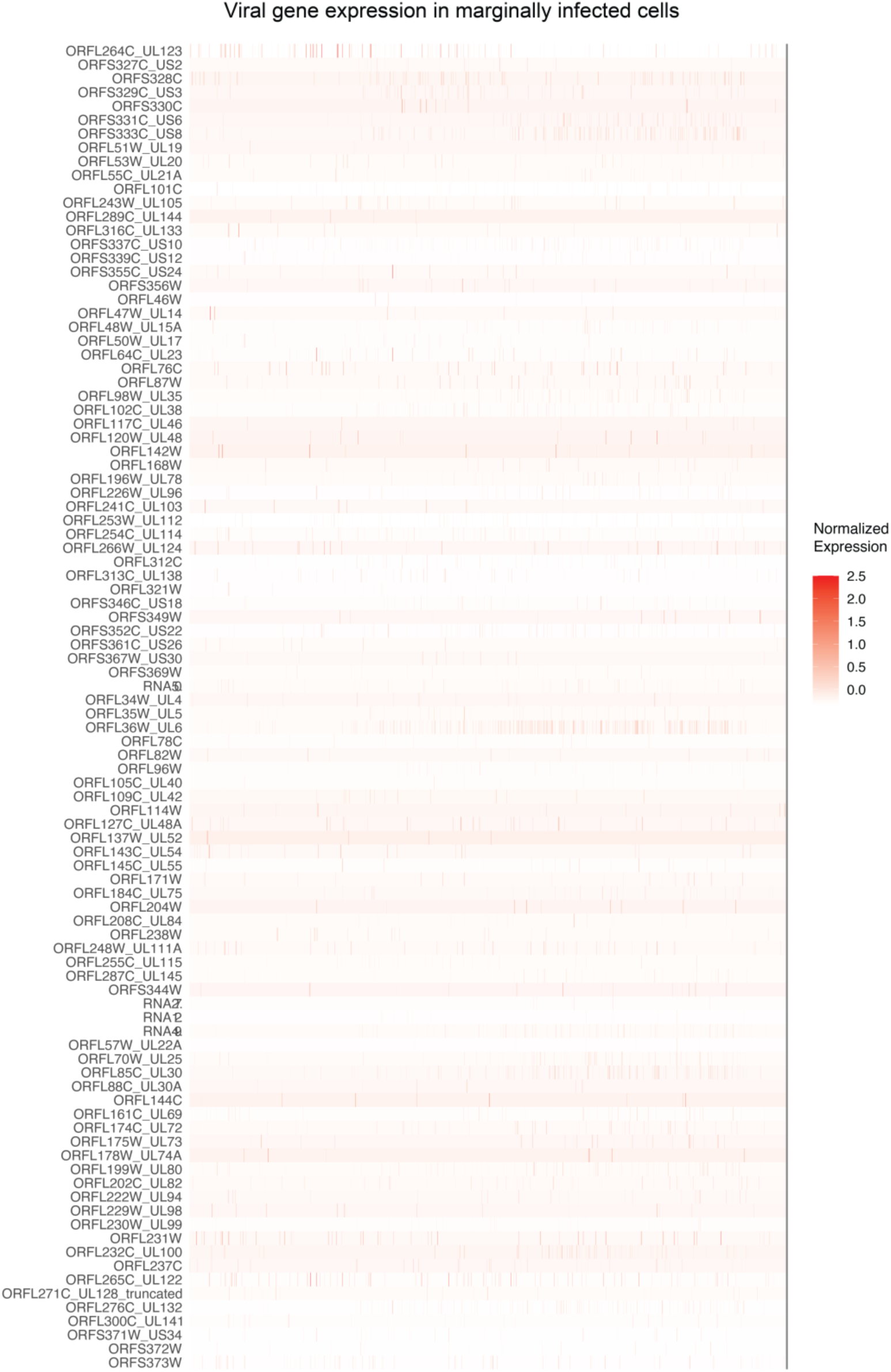
Expression of viral genes in marginally infected cells. Viral genes are selected and ordered based on Fig. 3D. Single cell normalized expression was plotted.

**Fig. S4:**
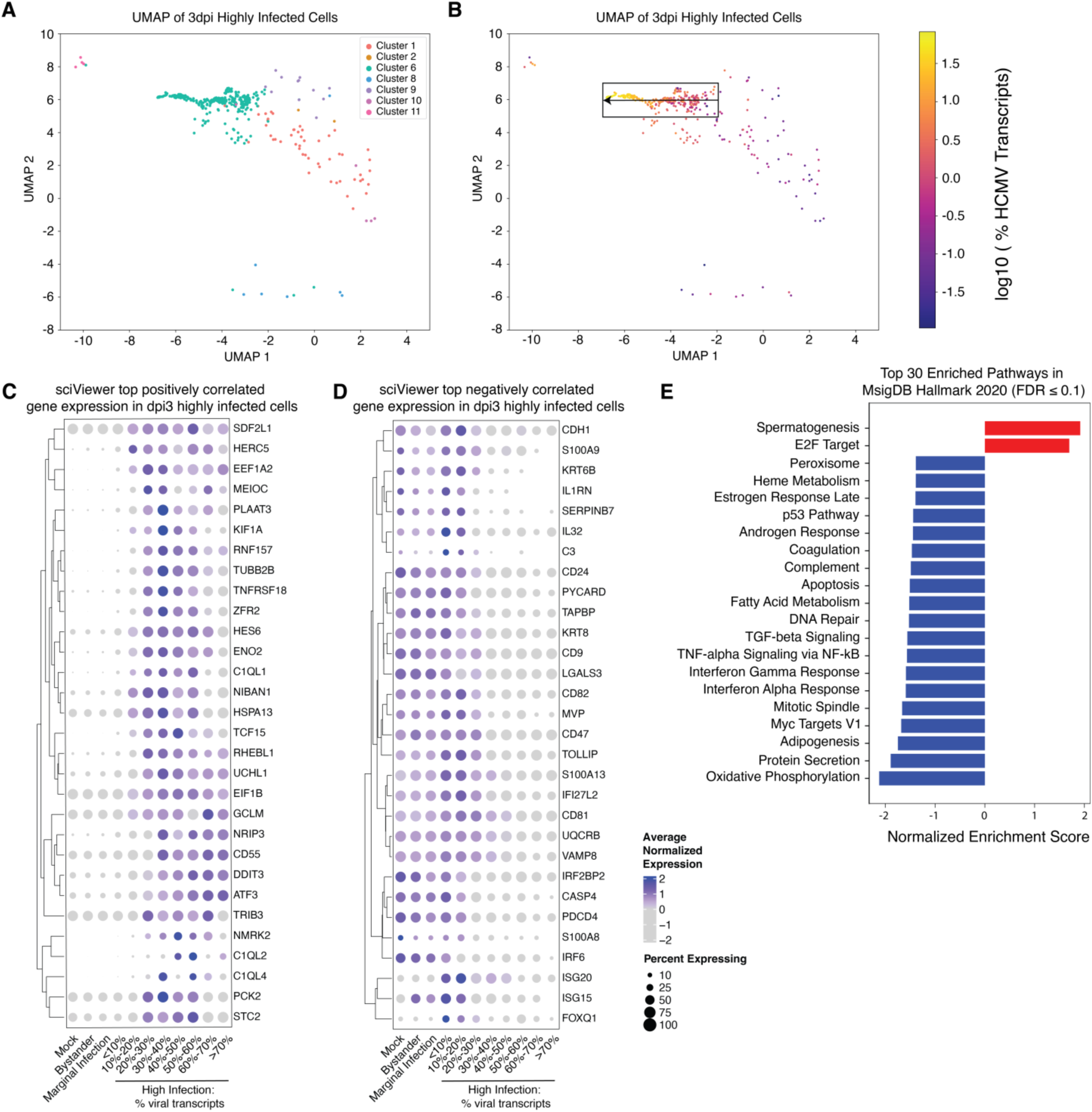
Interferon, complement, apoptotic and oxidative phosphorylation pathways associated with increased HCMV replication highlighted by SciViewer directional analysis and GSEA. **(A)** UMAP of 3dpi infected cells colored by Seurat cluster. This is a subset of the original UMAP containing all cells, but this subset contains only 3dpi infected cells. **(B)** UMAP of 3dpi Infected cells colored by the log_10_ of viral transcript percentage. The arrow and box represents the directional analysis performed on SciViewer. The directional analysis goes from 3dpi infected cells with relatively moderate number of viral transcripts towards the 3dpi infected cells with the highest number of viral transcripts. Heatmaps of top positively **(C)** and negatively **(D)** correlated genes identified in the SciViewer directional analysis. 3dpi highly infected cells were grouped by their viral transcript percentage with 5% increment. Averaged expression was taken per bin for each gene. **(E)** Gene set enrichment analysis result on the top enriched pathways on genes ordered by correlation along SciViewer trajectory. Pathways that have FDR ≤ 0.1 are included.

**Fig. S5:**
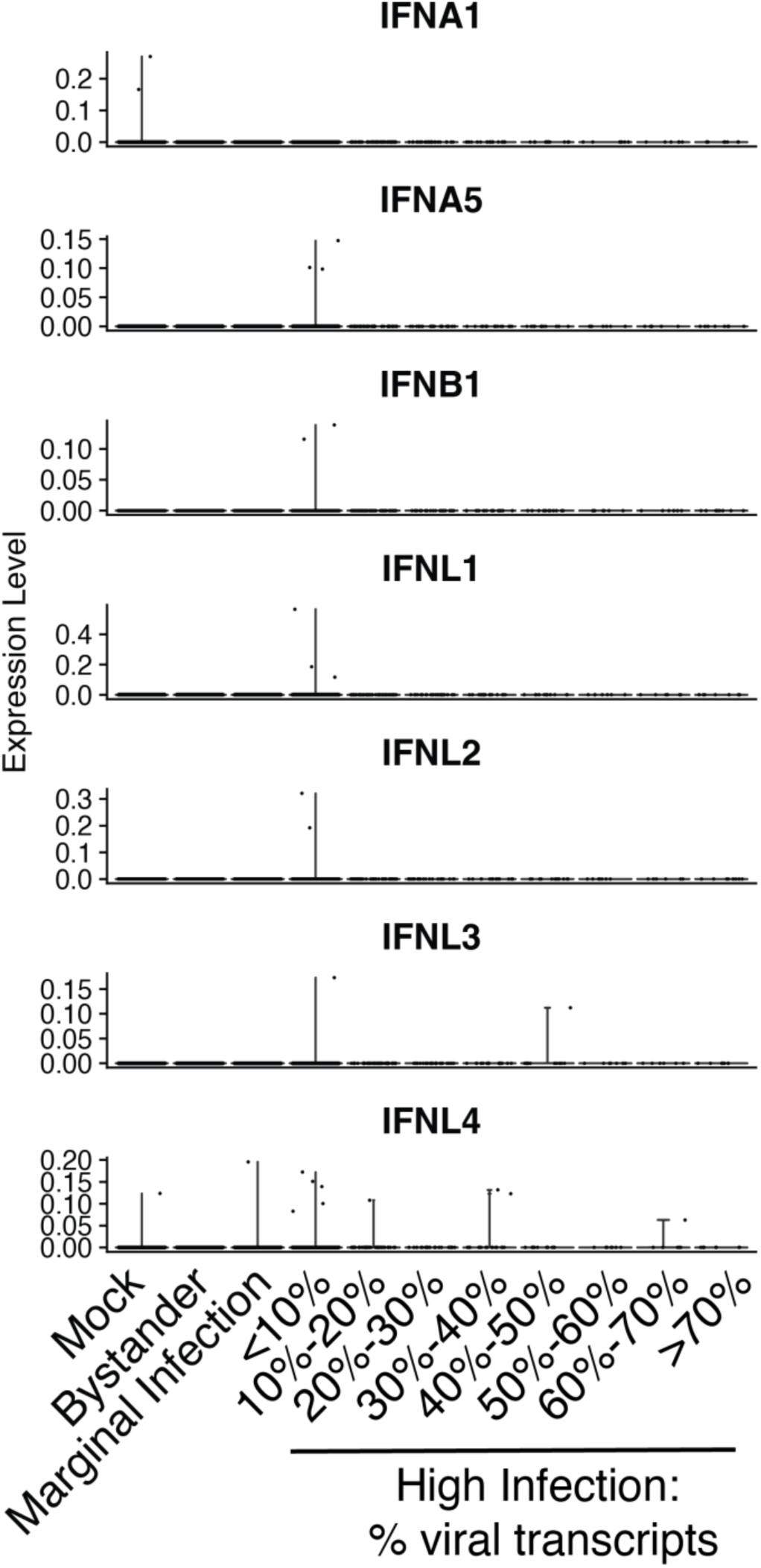
Expression of interferon genes. Violin plots of interferon genes expressed in the dataset. Cells are grouped by HCMV gene expression.

**Fig. S6:**
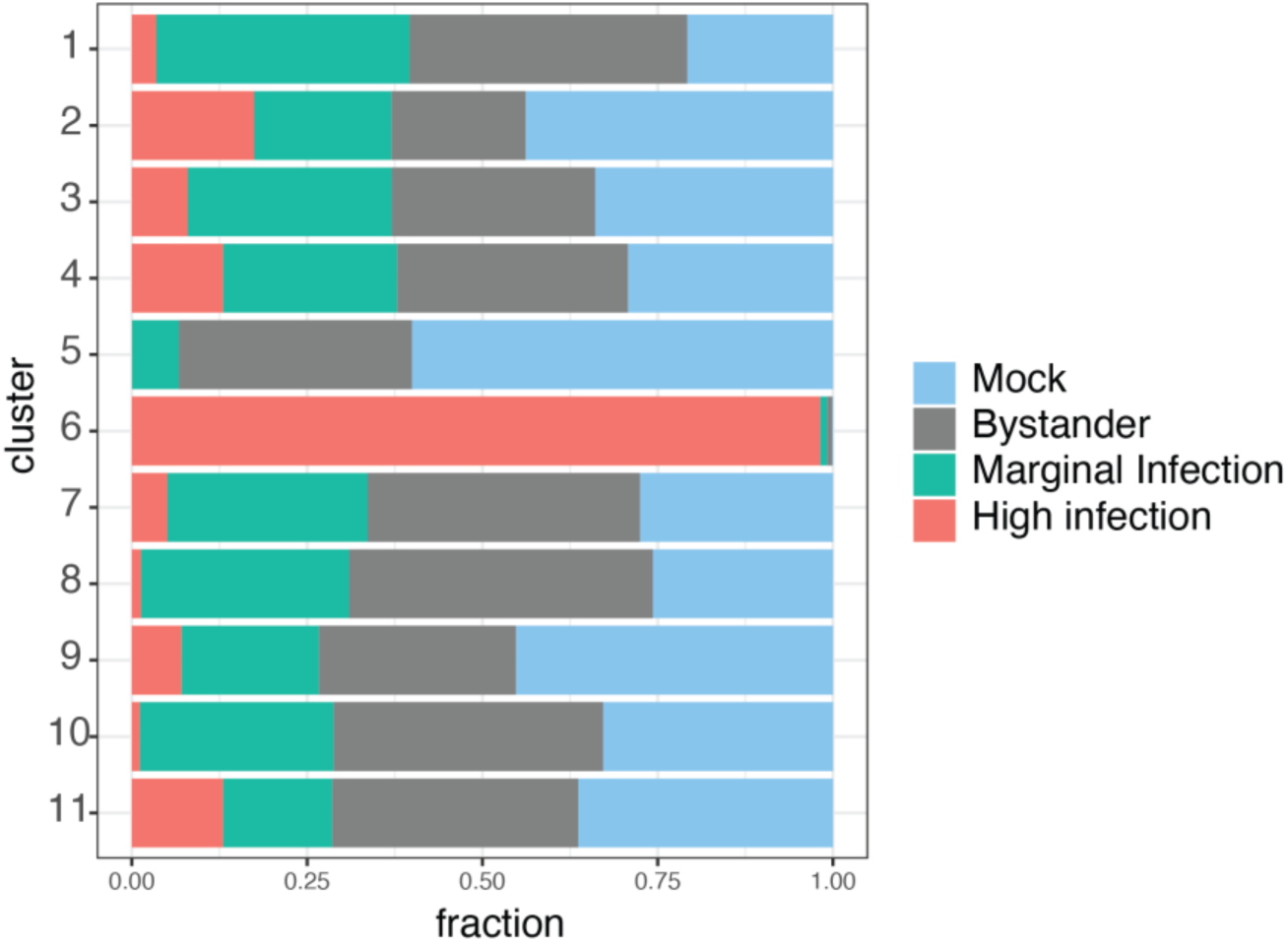
**Fraction of cells in each infection stage for each cluster.**

