## Supplemental Table 2 for "Human Cytomegalovirus Infection of Primary Human Oral Keratinocytes Induces Intermediate Keratinocyte Differentiation and an Altered Innate Immune Response"

| Sample | Total reads | Uniquely mapped | # total cell | Pre correction | |
|---|---|---|---|---|---|
| | | | | #virus positive cells | #GFP positive cells |
| **Mock 1dpi** | 104,619,141 | 75.4% | 858 | 0 | 0 |
| **Mock 3dpi** | 105,312,194 | 76.2% | 815 | 1 | 0 |
| **TB40E-Gfp 1dpi** | 89,394,174 | 73.1% | 701 | 474 | 261 |
| **TB40E-Gfp 3dpi** | 91,760,529 | 74.0% | 903 | 760 | 625 |
| **MOLD 1dpi** | 78,444,317 | 78.0% | 845 | 684 | 0 |
| **MOLD 3dpi** | 103,480,547 | 72.8% | 849 | 219 | 0 |
| **Mock 1dpi** | 84,317,141 | 75.8% | 572 | 0 | 0 |
| **Mock 3dpi** | 142,396,757 | 70.2% | 619 | 0 | 0 |
| **TB40E-Gfp 1dpi** | 122,774,458 | 74.2% | 600 | 517 | 321 |
| **TB40E-Gfp 3dpi** | 107,490,549 | 70.7% | 718 | 625 | 613 |
| **MOLD 1dpi** | 114,854,906 | 78.3% | 1129 | 888 | 4 |
| **MOLD 3dpi** | 86,609,890 | 74.1% | 1023 | 913 | 0 |

| | | | | Post correction | |
|---|---|---|---|---|---|
| # total cell | Median reads per cell | # virus positive cells | # GFP positive cells | # marginally infected cells | # highly infected cells |
| 765 | 121,934 | 0 | 0 | 0 | 0 |
| 786 | 129,817 | 1 | 0 | 0 | 0 |
| 617 | 127,524 | 337 | 179 | 95 | 242 |
| 769 | 101,617 | 347 | 160 | 157 | 190 |
| 805 | 92,834 | 407 | 0 | 183 | 224 |
| 798 | 121,885 | 132 | 0 | 112 | 20 |
| 524 | 147,408 | 0 | 0 | 0 | 0 |
| 587 | 230,043 | 0 | 0 | 0 | 0 |
| 527 | 204,624 | 300 | 123 | 92 | 208 |
| 623 | 149,708 | 123 | 41 | 37 | 86 |
| 900 | 101,732 | 798 | 0 | 543 | 255 |
| 922 | 84,663 | 882 | 0 | 789 | 93 |

| % marginally infected cells | % highly infected cells |
|---|---|
| 0 | 0 |
| 0 | 0 |
| 15.4% | 39.2% |
| 20.4% | 24.7% |
| 22.7% | 27.8% |
| 14.0% | 2.5% |
| 0 | 0 |
| 0 | 0 |
| 17.5% | 39.5% |
| 5.9% | 13.8% |
| 60.3% | 28.4% |
| 85.6% | 10.1% |
