## Supplemental Table 3 for "Human Cytomegalovirus Infection of Primary Human Oral Keratinocytes Induces Intermediate Keratinocyte Differentiation and an Altered Innate Immune Response"

| | logFC | logCPM | LR | PValue |
|---|---|---|---|---|
| ANXA2R | 3.800649983 | 5.809600611 | 833.3036574 | 3.10E-183 |
| FOXQ1 | 3.746897056 | 6.273025499 | 1192.832146 | 2.20E-261 |
| ISG15 | 3.146911245 | 7.31839218 | 2302.145644 | 0 |
| IFI6 | 3.123172604 | 5.814005856 | 871.012462 | 1.97E-191 |
| MX1 | 3.02117541 | 5.701980045 | 1181.553586 | 6.23E-259 |
| IFI27 | 2.865987436 | 5.534157736 | 367.9699087 | 5.18E-82 |
| CLU | 2.839307128 | 5.697592193 | 1676.875158 | 0 |
| RBP7 | 2.717511157 | 5.604709307 | 1157.304334 | 1.16E-253 |
| CXCL14 | -2.662479366 | 6.362579038 | 147.5547246 | 5.94E-34 |
| HERC5 | 2.386702728 | 5.306370351 | 792.79258 | 1.99E-174 |
| TP53INP2 | 2.30989154 | 5.357457098 | 1034.048897 | 7.13E-227 |
| CLEC11A | 2.234933113 | 5.347943718 | 798.6061904 | 1.08E-175 |
| TGM1 | -2.185529965 | 6.690900998 | 312.0435243 | 7.84E-70 |
| RNF223 | 2.160420418 | 5.269951808 | 746.6211746 | 2.18E-164 |
| TXNIP | -2.159230384 | 8.122046766 | 1251.583137 | 3.76E-274 |
| ENO2 | 2.152615973 | 5.277873633 | 731.7502409 | 3.73E-161 |
| TMEM125 | 2.145444248 | 5.338877447 | 872.3530262 | 1.00E-191 |
| GJB6 | -2.136648777 | 6.280772036 | 698.0786537 | 7.83E-154 |
| NID1 | 2.135775483 | 5.296350689 | 526.2702619 | 1.83E-116 |
| FOS | -2.091526843 | 7.606314206 | 1304.188518 | 1.39E-285 |
| PRSS22 | 2.077493136 | 5.59901283 | 368.3448857 | 4.29E-82 |
| BEX5 | 2.071845492 | 5.237567605 | 824.4022334 | 2.67E-181 |
| WFDC2 | 2.065854363 | 5.512346384 | 401.4170607 | 2.71E-89 |
| UCHL1 | 1.994414313 | 5.823028072 | 444.591848 | 1.08E-98 |
| CCK | 1.983684435 | 5.380583299 | 490.5526823 | 1.08E-108 |
| SERPINB3 | -1.969900562 | 6.135161218 | 177.9898169 | 1.33E-40 |
| INA | 1.944000881 | 5.304768129 | 705.6821002 | 1.74E-155 |
| C15orf48 | 1.942772236 | 5.817736671 | 125.0680455 | 4.92E-29 |
| FAM241B | 1.935027566 | 5.672070125 | 1480.662761 | 4.94065645841247e-324 |
| AMIGO2 | -1.926739441 | 6.710056396 | 1318.749066 | 9.52E-289 |
| ISG20 | 1.911053411 | 6.044223182 | 1080.857355 | 4.78E-237 |
| TLCD4 | 1.873156512 | 5.239311907 | 667.5907186 | 3.34E-147 |
| HSPA1A | 1.870571385 | 6.707431936 | 1310.65702 | 5.46E-287 |
| CLDN4 | 1.836775424 | 5.822894897 | 257.8790784 | 4.98E-58 |
| HMOX1 | 1.836749235 | 5.263092313 | 564.3492588 | 9.52E-125 |
| MYC | -1.835515454 | 8.069154355 | 1330.81025 | 2.28E-291 |
| G0S2 | -1.796883339 | 9.469352175 | 346.771609 | 2.14E-77 |
| KRT16 | -1.789966268 | 9.962321127 | 284.8748123 | 6.51E-64 |
| TIMP2 | 1.788991638 | 5.666808114 | 431.4450826 | 7.88E-96 |
| SLC2A1 | -1.727201651 | 7.377674174 | 1114.631724 | 2.18E-244 |
| SYNGR1 | 1.726344487 | 5.505954387 | 496.2776948 | 6.14E-110 |
| RETREG1 | 1.713008713 | 5.228940071 | 617.7429815 | 2.32E-136 |

| | | | | |
|---|---|---|---|---|
| TNNI3 | 1.711414467 | 5.275202024 | 482.8506556 | 5.12E-107 |
| DDX60 | 1.707467484 | 5.44257189 | 427.9883182 | 4.45E-95 |
| GJB2 | -1.703566959 | 8.119928455 | 907.8375021 | 1.94E-199 |
| SNCG | 1.668027526 | 5.751569951 | 466.4056213 | 1.94E-103 |
| FILIP1L | -1.666885058 | 5.761307528 | 317.1383697 | 6.08E-71 |
| SNX10 | 1.660619744 | 5.443642618 | 734.755998 | 8.28E-162 |
| PLAAT3 | 1.660124267 | 5.26150717 | 314.1426986 | 2.73E-70 |
| ACP5 | 1.655502765 | 5.372383567 | 562.4715539 | 2.44E-124 |
| RAB6B | 1.651920033 | 5.245234177 | 464.1590783 | 5.98E-103 |
| NOTCH1 | -1.647455904 | 6.620114242 | 1458.259265 | 4.59989938247576e-319 |
| KLF7 | -1.645805154 | 6.47110132 | 1143.28925 | 1.29E-250 |
| IL20RB | -1.641915102 | 6.399717431 | 832.9521105 | 3.70E-183 |
| RRAGD | 1.64027029 | 5.230305729 | 457.8568854 | 1.41E-101 |
| PDK2 | 1.634008768 | 5.534211324 | 735.4046478 | 5.98E-162 |
| EGR1 | -1.619026914 | 6.557650409 | 538.1148052 | 4.85E-119 |
| TSPYL5 | 1.596987414 | 5.332863266 | 497.117667 | 4.03E-110 |
| GCH1 | 1.593481833 | 5.328521852 | 588.8335876 | 4.49E-130 |
| MYLK | -1.59308274 | 5.610583456 | 194.6293328 | 3.10E-44 |
| GCHFR | 1.591327067 | 5.496120702 | 502.5715345 | 2.62E-111 |
| RPP25 | 1.589207821 | 5.575974012 | 683.2180321 | 1.33E-150 |
| BEX2 | 1.586632695 | 5.944463348 | 831.1980356 | 8.90E-183 |
| NRIP3 | 1.578492083 | 5.255458234 | 489.2519581 | 2.07E-108 |
| TP63 | -1.572265136 | 6.829114823 | 1481.353766 | 4.94065645841247e-324 |
| SMPD1 | 1.569473952 | 5.344226793 | 642.1671544 | 1.13E-141 |
| KLF2 | 1.569323616 | 5.224409678 | 433.0494612 | 3.53E-96 |
| ZBTB16 | -1.568423146 | 5.384675374 | 198.7807504 | 3.85E-45 |
| PARD6A | 1.565714465 | 5.330870755 | 540.4258685 | 1.52E-119 |
| GPRC5C | 1.563557682 | 5.29376094 | 527.0081723 | 1.26E-116 |
| MT1F | 1.559593132 | 5.527789119 | 714.4564329 | 2.15E-157 |
| NKAPL | 1.556976693 | 5.316521932 | 452.6570643 | 1.90E-100 |
| CGN | 1.551653506 | 5.327390477 | 466.0935464 | 2.27E-103 |
| CBR3 | 1.550348628 | 5.324412455 | 514.3085433 | 7.32E-114 |
| MITF | 1.54983779 | 5.392840209 | 576.6655096 | 1.99E-127 |
| TNFAIP2 | 1.54834364 | 5.271909225 | 267.5143095 | 3.95E-60 |
| AKAP12 | 1.54520936 | 5.663630081 | 256.4483849 | 1.02E-57 |
| S100A8 | -1.544304882 | 8.567178641 | 46.32071001 | 1.00E-11 |
| IRF7 | 1.539907213 | 5.383751058 | 535.5140193 | 1.78E-118 |
| TDRD7 | 1.525262774 | 5.257853801 | 541.1625994 | 1.05E-119 |
| TP53I11 | 1.517701766 | 5.402238044 | 421.7051767 | 1.04E-93 |
| RND2 | 1.511006363 | 5.225268179 | 386.1004948 | 5.85E-86 |
| ICAM1 | 1.50617831 | 5.400164802 | 269.9237043 | 1.18E-60 |
| NFKBIZ | -1.504905964 | 6.041722947 | 630.4977437 | 3.90E-139 |
| INHBA | -1.500697915 | 7.147527006 | 449.4266301 | 9.61E-100 |

| | | | | |
|---|---|---|---|---|
| GKAP1 | 1.500279973 | 5.223032843 | 470.9878392 | 1.95E-104 |
| DNAJB9 | 1.491910236 | 5.858774585 | 792.4818565 | 2.33E-174 |
| RGS9 | 1.488965153 | 5.376151422 | 527.9493762 | 7.89E-117 |
| IVL | -1.488790362 | 6.060241357 | 27.0637036 | 1.97E-07 |
| HERC6 | 1.487804906 | 5.3581606 | 513.2844037 | 1.22E-113 |
| OTUB2 | 1.484492358 | 5.357559496 | 499.2521506 | 1.38E-110 |
| ZFP36L2 | -1.481241325 | 8.723942633 | 1537.806607 | 0 |
| CCNE1 | 1.480932333 | 5.756617063 | 777.1490452 | 5.02E-171 |
| IL11 | 1.477548499 | 5.416566585 | 397.1338327 | 2.32E-88 |
| PTGES | 1.476098967 | 5.442140683 | 494.1367322 | 1.79E-109 |
| CCDC69 | 1.475649893 | 5.239485347 | 468.2743893 | 7.61E-104 |
| TNFRSF18 | 1.467427114 | 5.266492567 | 296.1582501 | 2.26E-66 |
| IL23A | 1.454138966 | 5.246842948 | 327.4657905 | 3.43E-73 |
| TGM2 | 1.450737143 | 5.586402119 | 366.3199956 | 1.18E-81 |
| SERTAD4 | 1.436131934 | 5.411627406 | 456.4060494 | 2.91E-101 |
| SYT8 | -1.430136588 | 5.851853511 | 243.6562567 | 6.27E-55 |
| DST | -1.423064597 | 10.4742298 | 993.4976194 | 4.65E-218 |
| ETS2 | -1.421811499 | 7.075815916 | 1067.511879 | 3.80E-234 |
| CTSF | 1.415555257 | 5.335609522 | 365.5875532 | 1.71E-81 |
| DDIT4 | -1.413835711 | 8.698748038 | 901.8996682 | 3.79E-198 |
| ZFP36L1 | -1.409143151 | 8.310786277 | 1429.346615 | 8.81848699475685e-313 |
| PHLDA1 | -1.40705703 | 7.899519543 | 814.7379334 | 3.37E-179 |
| RASGRP2 | 1.40428736 | 5.269913634 | 288.3967135 | 1.11E-64 |
| DMTN | 1.399424858 | 5.244257843 | 405.7944485 | 3.02E-90 |
| SAMHD1 | 1.392600291 | 5.428049616 | 553.3253055 | 2.38E-122 |
| MT1G | 1.392105363 | 5.324691107 | 151.2166464 | 9.40E-35 |
| DSG3 | -1.390996633 | 6.841631948 | 667.3137315 | 3.84E-147 |
| CXCL8 | 1.389181352 | 5.942779289 | 84.61002233 | 3.63E-20 |
| OAS1 | 1.386882891 | 5.26882421 | 279.4646926 | 9.82E-63 |
| NIBAN1 | 1.385108827 | 5.486079376 | 321.8410969 | 5.75E-72 |
| MAP7D2 | 1.379430238 | 5.26781447 | 342.3339449 | 1.98E-76 |
| CRB3 | 1.378960475 | 5.868153342 | 937.3013369 | 7.64E-206 |
| B3GNT3 | 1.377169339 | 5.409148992 | 504.2270275 | 1.14E-111 |
| GABARAPL1 | 1.374299771 | 5.627187361 | 601.7922699 | 6.82E-133 |
| DPF1 | 1.372069627 | 5.245539446 | 406.6714469 | 1.94E-90 |
| LGALS9C | -1.369656584 | 5.286716115 | 297.9894247 | 9.03E-67 |
| CAMK2B | 1.369559455 | 5.422499495 | 225.713935 | 5.13E-51 |
| PCDH1 | 1.368098816 | 5.309559824 | 191.3261995 | 1.63E-43 |
| VSNL1 | -1.367787107 | 5.779397318 | 525.3708713 | 2.87E-116 |
| JAG1 | -1.365259422 | 7.295392989 | 1021.482394 | 3.84E-224 |
| YPEL3 | 1.36489369 | 5.963554409 | 640.5774897 | 2.50E-141 |
| MAP1B | 1.364595834 | 6.492108254 | 483.9882721 | 2.90E-107 |
| TENM2 | -1.363536002 | 7.625957876 | 675.1626426 | 7.53E-149 |

| | | | | |
|---|---|---|---|---|
| PLAUR | 1.360777212 | 6.508642869 | 620.7308056 | 5.19E-137 |
| DNAJA4 | 1.358817143 | 5.326665694 | 431.3500213 | 8.26E-96 |
| TMEM121 | 1.357985777 | 5.325774142 | 314.7797744 | 1.99E-70 |
| PKN1 | 1.355916725 | 5.466716634 | 462.3677213 | 1.47E-102 |
| TP53AIP1 | -1.355528133 | 5.497419722 | 278.5726014 | 1.54E-62 |
| GPX2 | 1.353940754 | 5.241880048 | 286.1076391 | 3.50E-64 |
| PTAFR | 1.346847587 | 5.479165347 | 520.731303 | 2.93E-115 |
| IRS1 | -1.342939633 | 6.213124421 | 648.2668477 | 5.32E-143 |
| CNFN | 1.337866448 | 5.2873682 | 159.1873364 | 1.70E-36 |
| CLDN23 | 1.337810438 | 5.228661667 | 391.3041027 | 4.31E-87 |
| FUT3 | 1.337348079 | 5.264734005 | 296.996805 | 1.49E-66 |
| FAM43A | 1.334612442 | 5.24080848 | 331.5883687 | 4.33E-74 |
| ATP6V0E2 | 1.328136885 | 5.472688113 | 494.2622549 | 1.68E-109 |
| RASEF | 1.325573482 | 5.244585963 | 331.6629599 | 4.17E-74 |
| SDR16C5 | 1.325465447 | 5.295169195 | 311.4220211 | 1.07E-69 |
| SVIP | 1.324769278 | 5.923018213 | 873.5053769 | 5.64E-192 |
| CCL28 | 1.32470832 | 5.300066169 | 357.1701699 | 1.16E-79 |
| NACAD | 1.322956463 | 5.288151066 | 359.779274 | 3.15E-80 |
| ELF3 | 1.322836875 | 5.403658997 | 163.3475562 | 2.10E-37 |
| TRPV3 | 1.316845066 | 5.21548612 | 346.9899321 | 1.92E-77 |
| LPIN2 | 1.314869957 | 5.493099158 | 465.6057806 | 2.90E-103 |
| WFS1 | 1.309343384 | 5.539980944 | 580.8280756 | 2.48E-128 |
| KDELR3 | 1.307278145 | 5.739768131 | 621.6167271 | 3.33E-137 |
| CDA | 1.306548108 | 6.17023303 | 352.7644023 | 1.06E-78 |
| CST6 | 1.304482764 | 5.492520522 | 100.3547438 | 1.27E-23 |
| ABHD8 | 1.303880223 | 5.488366421 | 496.8026949 | 4.72E-110 |
| C1orf115 | 1.301989634 | 5.258860177 | 381.3064302 | 6.46E-85 |
| MARCHF3 | 1.300730535 | 5.257143772 | 287.2239724 | 2.00E-64 |
| MAGEH1 | 1.296775712 | 5.337924082 | 373.5148564 | 3.21E-83 |
| ITM2C | 1.296050441 | 6.051283352 | 889.1661688 | 2.22E-195 |
| FICD | 1.290471365 | 5.279917383 | 331.5315004 | 4.46E-74 |
| SLC25A4 | 1.286749903 | 6.780732382 | 1670.14624 | 0 |
| BIRC3 | 1.285924112 | 5.35985214 | 220.9888691 | 5.50E-50 |
| MMP28 | -1.277862508 | 5.767503601 | 341.1558364 | 3.57E-76 |
| USP2 | 1.277800179 | 5.20803753 | 342.2370176 | 2.08E-76 |
| RAB3A | 1.276707752 | 5.2179532 | 335.9242688 | 4.93E-75 |
| F3 | -1.275752661 | 9.358280638 | 531.3801172 | 1.41E-117 |
| PLK2 | -1.275411842 | 7.701569868 | 678.7828081 | 1.23E-149 |
| TMEM38A | 1.275107868 | 5.432252791 | 377.6266328 | 4.09E-84 |
| ERO1B | 1.274212033 | 5.361049601 | 392.5752452 | 2.28E-87 |
| LGALS9B | -1.274028171 | 5.308814844 | 282.8418478 | 1.80E-63 |
| FOSB | -1.271147119 | 6.137956005 | 270.266725 | 9.93E-61 |
| EPHX4 | 1.269298185 | 5.23943089 | 329.5318253 | 1.22E-73 |

| | | | | |
|---|---|---|---|---|
| HLA-B | 1.266749389 | 6.625000402 | 525.0437936 | 3.38E-116 |
| RORA | 1.266112451 | 5.295504653 | 318.9030608 | 2.51E-71 |
| HLA-F | 1.262231124 | 5.24003844 | 312.5604049 | 6.05E-70 |
| PDE4A | 1.256496525 | 5.211489924 | 311.2685929 | 1.16E-69 |
| SERPINB5 | -1.255592604 | 8.32183999 | 1364.861 | 9.08E-299 |
| PRR15 | 1.255016107 | 5.205218492 | 342.3639658 | 1.95E-76 |
| LBH | 1.254725921 | 5.315032863 | 239.1738972 | 5.95E-54 |
| KCNJ5 | -1.253635216 | 5.306531365 | 235.5458636 | 3.68E-53 |
| SLC41A2 | 1.253226285 | 5.201967264 | 349.9672698 | 4.31E-78 |
| APH1B | 1.249892575 | 5.387890309 | 410.7644188 | 2.50E-91 |
| SNAI2 | -1.248624913 | 6.792194973 | 830.730363 | 1.12E-182 |
| RHPN2 | 1.244564753 | 5.334826318 | 366.4402172 | 1.12E-81 |
| PCSK1N | 1.24409276 | 5.564319373 | 277.5416258 | 2.58E-62 |
| TMEM255B | 1.242613164 | 5.234552188 | 314.9202743 | 1.85E-70 |
| MAP1A | 1.232288562 | 5.280409057 | 174.9114237 | 6.26E-40 |
| GABRE | -1.229977269 | 5.357333362 | 277.0818775 | 3.25E-62 |
| NEU1 | 1.229255539 | 5.863310319 | 723.9522779 | 1.85E-159 |
| FBXO32 | 1.227528819 | 5.345401504 | 214.0731871 | 1.78E-48 |
| HMGA2 | -1.226080142 | 8.100000168 | 907.8324811 | 1.95E-199 |
| EMILIN2 | 1.223524868 | 5.220715777 | 323.2888297 | 2.78E-72 |
| SOCS1 | 1.22289564 | 5.373502931 | 299.9284817 | 3.41E-67 |
| METRNL | 1.222587675 | 5.682098707 | 209.0066436 | 2.26E-47 |
| KIF13B | 1.219936889 | 5.396183532 | 383.6872309 | 1.96E-85 |
| ZNF280B | 1.218690608 | 5.294549857 | 324.6516518 | 1.41E-72 |
| INSR | 1.217558936 | 5.237431979 | 262.3526845 | 5.27E-59 |
| HTRA1 | -1.217147338 | 7.995300894 | 1159.462957 | 3.94E-254 |
| PLPP2 | 1.217144769 | 6.00281287 | 497.3879804 | 3.52E-110 |
| RHEBL1 | 1.214397423 | 5.23930864 | 266.2406848 | 7.49E-60 |
| ATOSB | 1.213363221 | 5.557968565 | 440.607629 | 7.98E-98 |
| MAP1LC3A | 1.211808251 | 6.020522899 | 521.5186428 | 1.98E-115 |
| TANC2 | -1.210697429 | 6.780229991 | 784.2385544 | 1.44E-172 |
| HAGH | 1.20612867 | 6.59229023 | 1193.189662 | 1.84E-261 |
| ABHD3 | 1.204101869 | 5.464321789 | 479.629232 | 2.57E-106 |
| DIPK1B | 1.203381598 | 5.295372292 | 254.1711199 | 3.20E-57 |
| ULBP2 | 1.202908196 | 5.698201352 | 550.2568613 | 1.11E-121 |
| ADAP2 | 1.201228085 | 5.334968207 | 219.9330756 | 9.35E-50 |
| PLPPR2 | 1.200643625 | 5.464196665 | 439.1820187 | 1.63E-97 |
| CD83 | 1.198368047 | 5.303247955 | 301.0661398 | 1.93E-67 |
| TSPAN2 | 1.197325885 | 5.214538301 | 271.7767809 | 4.65E-61 |
| COBL | 1.194617063 | 5.230664776 | 224.3488928 | 1.02E-50 |
| SLC17A5 | 1.190369816 | 5.483547043 | 422.9565776 | 5.55E-94 |
| CCDC149 | 1.187868355 | 5.361841672 | 379.4386635 | 1.65E-84 |
| CEBPD | -1.187765054 | 7.890283015 | 923.8173398 | 6.52E-203 |

| | | | | |
|---|---|---|---|---|
| FLRT3 | -1.182827022 | 5.736392401 | 306.2034954 | 1.47E-68 |
| CDKN2D | 1.182248472 | 5.553460444 | 431.1172934 | 9.28E-96 |
| STX3 | 1.179836622 | 5.714647669 | 544.2740573 | 2.22E-120 |
| CCNA1 | 1.174193265 | 5.286755437 | 214.4630511 | 1.46E-48 |
| HSPA2 | 1.173537887 | 5.443933826 | 355.8225788 | 2.29E-79 |
| DLK2 | -1.171532952 | 5.564159362 | 247.944306 | 7.29E-56 |
| MAP1S | 1.165033886 | 5.65642595 | 638.298723 | 7.83E-141 |
| IL33 | -1.163099323 | 5.303526567 | 126.8550126 | 2.00E-29 |
| CHST12 | 1.157710288 | 5.628308981 | 614.3981807 | 1.24E-135 |
| FBLN2 | 1.157017805 | 5.291499213 | 216.6298505 | 4.92E-49 |
| FGFR3 | -1.155935933 | 5.974979372 | 386.0345455 | 6.04E-86 |
| FST | -1.152867747 | 9.064618445 | 357.9748344 | 7.77E-80 |
| XKR8 | 1.148935339 | 5.3448412 | 361.8949056 | 1.09E-80 |
| OCLN | 1.148705819 | 5.452558653 | 279.3891494 | 1.02E-62 |
| GADD45A | -1.147027534 | 8.474183315 | 825.4644905 | 1.57E-181 |
| KIF3C | 1.146342747 | 5.560372662 | 469.215801 | 4.75E-104 |
| MFSD9 | 1.146151244 | 5.381960227 | 374.3574386 | 2.11E-83 |
| SMCO4 | 1.14321858 | 5.461545208 | 360.5603579 | 2.13E-80 |
| PIERCE1 | 1.142025862 | 5.289186093 | 303.6191938 | 5.36E-68 |
| ARFGEF3 | 1.140403646 | 5.399732162 | 297.0057935 | 1.48E-66 |
| SELENOM | 1.137524615 | 5.932243114 | 384.818828 | 1.11E-85 |
| SLC38A2 | -1.137467224 | 8.009530742 | 1099.767304 | 3.71E-241 |
| PTHLH | -1.136538662 | 8.181633319 | 506.8301419 | 3.10E-112 |
| NCOA1 | 1.135514998 | 5.485078558 | 385.0144206 | 1.01E-85 |
| FAM171B | 1.135409705 | 5.263235783 | 271.2911378 | 5.94E-61 |
| DDIT3 | 1.135276102 | 5.624378008 | 195.1710093 | 2.36E-44 |
| FGFBP1 | -1.135166375 | 10.75832169 | 658.7492907 | 2.79E-145 |
| QPRT | 1.133740468 | 5.317292209 | 102.712783 | 3.87E-24 |
| KRT14 | -1.133541913 | 13.39787788 | 979.3382792 | 5.56E-215 |
| SIPA1L2 | 1.131995952 | 5.365219106 | 209.8453193 | 1.48E-47 |
| TNNI2 | -1.131222148 | 5.262057761 | 164.8669052 | 9.78E-38 |
| PLLP | 1.131218541 | 5.552117674 | 369.6272352 | 2.26E-82 |
| KCNN4 | 1.127646065 | 5.362269572 | 226.2961501 | 3.83E-51 |
| CASTOR2 | 1.125363827 | 5.356023762 | 309.991173 | 2.19E-69 |
| PPP2R5B | 1.123989525 | 5.493291169 | 422.3489259 | 7.52E-94 |
| HDHD3 | 1.123109498 | 5.551741397 | 458.851282 | 8.55E-102 |
| TFAP2C | 1.122457512 | 5.323612514 | 284.4420105 | 8.08E-64 |
| ANKRD9 | 1.121634755 | 5.783651819 | 570.0739776 | 5.41E-126 |
| C1orf210 | 1.118395098 | 5.331769921 | 321.1706661 | 8.05E-72 |
| FAM131C | 1.118286138 | 5.306282839 | 263.2170696 | 3.41E-59 |
| PRKAA2 | 1.117594133 | 5.284433584 | 240.2826054 | 3.41E-54 |
| PHTF1 | 1.116755029 | 5.463220251 | 416.7677055 | 1.23E-92 |
| HES6 | 1.11620885 | 5.569641996 | 228.0764336 | 1.57E-51 |

| | | | | |
|---|---|---|---|---|
| ARMCX1 | 1.11418034 | 5.864072536 | 664.7708209 | 1.37E-146 |
| KLK1 | 1.110900729 | 5.225181389 | 195.3523764 | 2.16E-44 |
| ATL1 | 1.110313416 | 5.256645109 | 262.1682385 | 5.78E-59 |
| IRF1 | 1.109560946 | 5.65867676 | 358.3927035 | 6.30E-80 |
| SLC25A42 | 1.108683957 | 5.256221221 | 257.1174071 | 7.29E-58 |
| RILP | 1.107367199 | 5.396585404 | 345.6541585 | 3.75E-77 |
| POMGNT2 | 1.106639523 | 5.449424518 | 383.2742853 | 2.41E-85 |
| OAS3 | 1.1053347 | 5.560594201 | 413.0024526 | 8.14E-92 |
| SLC7A11 | -1.103192093 | 6.544720697 | 400.8232938 | 3.65E-89 |
| A1BG | 1.102236981 | 5.4307947 | 261.3073715 | 8.90E-59 |
| ELOVL4 | 1.102110181 | 5.579397757 | 429.9585331 | 1.66E-95 |
| GULP1 | 1.101067219 | 5.329242824 | 262.3835542 | 5.19E-59 |
| H2BC11 | 1.099545735 | 5.307395905 | 239.0588768 | 6.31E-54 |
| HSPA1B | 1.099188167 | 5.868864723 | 417.8020526 | 7.34E-93 |
| USP18 | 1.098912816 | 5.335463264 | 304.3249939 | 3.76E-68 |
| NAP1L5 | 1.097958724 | 5.244992034 | 251.8094769 | 1.05E-56 |
| PCDH7 | -1.096738614 | 5.559559323 | 221.8777543 | 3.52E-50 |
| IL1RAP | -1.096472385 | 6.369484643 | 648.9586649 | 3.76E-143 |
| LRIF1 | 1.091666712 | 6.425883091 | 942.8389524 | 4.78E-207 |
| MAFG | 1.091161948 | 6.037159314 | 738.0334751 | 1.60E-162 |
| SAMD9 | 1.090375485 | 5.522472286 | 209.5082715 | 1.76E-47 |
| INPP5D | -1.089714085 | 5.909305909 | 426.2406575 | 1.07E-94 |
| TLN2 | 1.089246982 | 5.278969198 | 263.5402333 | 2.90E-59 |
| PTGS1 | 1.088893121 | 5.273031824 | 251.7747186 | 1.07E-56 |
| GARIN5A | 1.087407006 | 5.298640378 | 198.5627058 | 4.30E-45 |
| PINK1 | 1.087073225 | 6.117458456 | 635.0188724 | 4.05E-140 |
| SLC2A10 | 1.08656275 | 5.210751144 | 242.0176234 | 1.43E-54 |
| TRIM2 | 1.08562142 | 5.274238275 | 252.8293916 | 6.28E-57 |
| ZNF483 | 1.084221637 | 5.319751244 | 252.6701853 | 6.80E-57 |
| TSPAN33 | 1.078952668 | 5.307209436 | 277.1026768 | 3.21E-62 |
| FLNC | 1.078707371 | 5.338777517 | 176.493752 | 2.82E-40 |
| NIPA1 | 1.076929538 | 5.4441881 | 357.4398602 | 1.02E-79 |
| E2F1 | 1.0767666 | 5.454788457 | 240.1420337 | 3.66E-54 |
| SERPINI1 | 1.076353181 | 5.21331841 | 251.5128683 | 1.22E-56 |
| ANKRD22 | 1.076316332 | 5.295281962 | 125.6612081 | 3.65E-29 |
| KLHL24 | 1.075730079 | 5.574908244 | 300.1575173 | 3.04E-67 |
| PRSS16 | 1.07513886 | 5.381352809 | 302.835903 | 7.94E-68 |
| PRADC1 | 1.073056265 | 5.873153608 | 512.994384 | 1.41E-113 |
| DSC3 | -1.068563853 | 7.135272435 | 871.7753597 | 1.34E-191 |
| AIFM2 | 1.0672833 | 5.374994869 | 316.9972391 | 6.53E-71 |
| GSPT2 | 1.06627326 | 5.582117327 | 412.1834952 | 1.23E-91 |
| TSPAN15 | 1.065931135 | 5.324454575 | 279.5758553 | 9.29E-63 |
| KLF6 | -1.065343081 | 8.286068744 | 535.4459209 | 1.85E-118 |

| | | | | |
|---|---|---|---|---|
| BCAM | -1.065217705 | 6.607341807 | 688.7584881 | 8.32E-152 |
| MAPRE3 | 1.064257594 | 5.361402126 | 323.7277124 | 2.23E-72 |
| FAT1 | -1.063549231 | 8.030497362 | 1055.685683 | 1.41E-231 |
| RBM38 | 1.062432728 | 5.62202132 | 475.7549257 | 1.79E-105 |
| TRAPPC14 | 1.060650936 | 5.375394827 | 334.5575017 | 9.78E-75 |
| OSGIN1 | 1.060624109 | 5.278551457 | 226.3761126 | 3.68E-51 |
| AJUBA | -1.060563893 | 6.482675732 | 844.714139 | 1.03E-185 |
| PGM2L1 | 1.060393266 | 5.671536694 | 315.9311071 | 1.11E-70 |
| SLC46A3 | 1.060300536 | 5.24714769 | 221.5983145 | 4.05E-50 |
| ARRDC4 | -1.060121382 | 5.930435623 | 233.5495016 | 1.00E-52 |
| SHFL | 1.053125508 | 5.637208281 | 444.1774512 | 1.33E-98 |
| LGALS7B | -1.052526434 | 6.364734353 | 153.233763 | 3.41E-35 |
| TMEM61 | 1.045772692 | 5.251840538 | 221.8678195 | 3.54E-50 |
| TP53I3 | 1.045075241 | 6.644938701 | 556.9374077 | 3.90E-123 |
| DUSP1 | -1.043888069 | 6.888696297 | 339.359672 | 8.80E-76 |
| ZFAND2A | 1.043519709 | 5.714187695 | 435.1233767 | 1.25E-96 |
| JUN | -1.043022277 | 7.638632748 | 640.7277832 | 2.32E-141 |
| EPPK1 | -1.042815188 | 6.314034259 | 341.7789275 | 2.61E-76 |
| SIK1 | -1.042747931 | 6.142932329 | 391.9301794 | 3.15E-87 |
| P2RX4 | 1.039790523 | 5.344633906 | 286.6767171 | 2.63E-64 |
| ING2 | 1.038954181 | 6.219308118 | 738.3286767 | 1.38E-162 |
| MAPRE2 | 1.03837003 | 5.4050206 | 305.1138523 | 2.53E-68 |
| CCNE2 | 1.038220855 | 5.384454289 | 148.9591542 | 2.93E-34 |
| AGR2 | 1.037720179 | 5.59839364 | 123.0683524 | 1.35E-28 |
| PLEKHG6 | 1.037545765 | 5.255851228 | 202.6467049 | 5.52E-46 |
| SEPHS2 | 1.037321521 | 6.258927138 | 835.3073898 | 1.14E-183 |
| VCAN | -1.037062822 | 6.178592561 | 380.8044101 | 8.31E-85 |
| CFL2 | 1.036573837 | 5.804803062 | 549.4763826 | 1.64E-121 |
| GCA | 1.032386695 | 5.495669004 | 364.0510809 | 3.69E-81 |
| PLCB4 | 1.031671334 | 5.480898107 | 185.3860748 | 3.23E-42 |
| LYPD6B | 1.031552001 | 5.304174255 | 211.5396038 | 6.34E-48 |
| HMGN5 | 1.02723688 | 5.489897677 | 353.7350643 | 6.51E-79 |
| IFFO2 | -1.026259631 | 6.021799121 | 396.8685171 | 2.65E-88 |
| BPGM | 1.024178231 | 5.862271559 | 339.8673375 | 6.82E-76 |
| HDAC9 | 1.022592586 | 5.27324232 | 210.7635487 | 9.36E-48 |
| BICDL1 | 1.021584013 | 5.25373806 | 229.0691406 | 9.51E-52 |
| EGFR | -1.020126959 | 8.147678867 | 1079.289425 | 1.05E-236 |
| CPEB4 | 1.019444013 | 5.769833692 | 441.9102829 | 4.16E-98 |
| KRTAP2-3 | 1.018424935 | 5.887145356 | 43.11691358 | 5.16E-11 |
| MEIS3 | 1.017818924 | 5.219933604 | 179.7652783 | 5.45E-41 |
| MFAP3L | 1.015262356 | 5.208735048 | 219.4020334 | 1.22E-49 |
| CCDC68 | 1.015118091 | 5.479067617 | 342.7119506 | 1.64E-76 |
| MTSS1 | -1.014334352 | 6.096209761 | 396.9589137 | 2.53E-88 |

| | | | | |
|---|---|---|---|---|
| ADCK2 | 1.013051118 | 5.648088604 | 457.6971389 | 1.52E-101 |
| ICA1 | 1.012936567 | 5.523262666 | 309.0587693 | 3.50E-69 |
| TNS4 | -1.012781701 | 7.008888561 | 797.2308967 | 2.16E-175 |
| PGPEP1 | 1.011892251 | 5.316961152 | 236.3430678 | 2.47E-53 |
| NCF2 | 1.011841899 | 5.353087182 | 173.8446665 | 1.07E-39 |
| LIN7B | 1.011507528 | 5.323827103 | 260.5813504 | 1.28E-58 |
| TBX3 | 1.008723915 | 5.425594146 | 268.4230984 | 2.50E-60 |
| SH3PXD2A | -1.007756263 | 7.074848123 | 703.9558464 | 4.13E-155 |
| TPST1 | 1.007552297 | 5.533185364 | 344.9030013 | 5.46E-77 |
| HAPLN3 | 1.007348576 | 5.267914903 | 232.4536111 | 1.74E-52 |
| PAK3 | 1.005399282 | 5.248088396 | 195.4723974 | 2.03E-44 |
| PKP1 | -1.005163261 | 6.940328156 | 447.7296599 | 2.25E-99 |
| FAT2 | -1.002780309 | 6.195100512 | 480.7112843 | 1.50E-106 |
| RHCG | -1.002725766 | 6.580049739 | 15.22147413 | 9.56E-05 |

| FDR | sig | gene |
|---|---|---|
| 3.98E-181 | sig_up | ANXA2R |
| 9.46E-259 | sig_up | FOXQ1 |
| 0 | sig_up | ISG15 |
| 2.92E-189 | sig_up | IFI6 |
| 2.48E-256 | sig_up | MX1 |
| 1.07E-80 | sig_up | IFI27 |
| 0 | sig_up | CLU |
| 4.31E-251 | sig_up | RBP7 |
| 3.25E-33 | sig_down | CXCL14 |
| 2.10E-172 | sig_up | HERC5 |
| 1.85E-224 | sig_up | TP53INP2 |
| 1.16E-173 | sig_up | CLEC11A |
| 1.19E-68 | sig_down | TGM1 |
| 2.03E-162 | sig_up | RNF223 |
| 2.00E-271 | sig_down | TXNIP |
| 3.25E-159 | sig_up | ENO2 |
| 1.54E-189 | sig_up | TMEM125 |
| 6.06E-152 | sig_down | GJB6 |
| 7.95E-115 | sig_up | NID1 |
| 8.16E-283 | sig_down | FOS |
| 8.92E-81 | sig_up | PRSS22 |
| 3.17E-179 | sig_up | BEX5 |
| 6.67E-88 | sig_up | WFDC2 |
| 3.24E-97 | sig_up | UCHL1 |
| 4.05E-107 | sig_up | CCK |
| 9.03E-40 | sig_down | SERPINB3 |
| 1.41E-153 | sig_up | INA |
| 2.28E-28 | sig_up | C15orf48 |
| 6.89221575948539e-321 | sig_up | FAM241B |
| 6.64E-286 | sig_down | AMIGO2 |
| 1.44E-234 | sig_up | ISG20 |
| 2.36E-145 | sig_up | TLCD4 |
| 3.39E-284 | sig_up | HSPA1A |
| 5.64E-57 | sig_up | CLDN4 |
| 4.83E-123 | sig_up | HMOX1 |
| 1.70E-288 | sig_down | MYC |
| 3.98E-76 | sig_down | G0S2 |
| 8.53E-63 | sig_down | KRT16 |
| 2.20E-94 | sig_up | TIMP2 |
| 7.16E-242 | sig_down | SLC2A1 |
| 2.36E-108 | sig_up | SYNGR1 |
| 1.39E-134 | sig_up | RETREG1 |

| | | |
|---|---|---|
| 1.86E-105 | sig_up | TNNI3 |
| 1.22E-93 | sig_up | DDX60 |
| 3.39E-197 | sig_down | GJB2 |
| 6.43E-102 | sig_up | SNCG |
| 9.55E-70 | sig_down | FILIP1L |
| 7.28E-160 | sig_up | SNX10 |
| 4.20E-69 | sig_up | PLAAT3 |
| 1.23E-122 | sig_up | ACP5 |
| 1.96E-101 | sig_up | RAB6B |
| 5.70387523426994e-316 | sig_down | NOTCH1 |
| 4.64E-248 | sig_down | KLF7 |
| 4.64E-181 | sig_down | IL20RB |
| 4.45E-100 | sig_up | RRAGD |
| 5.30E-160 | sig_up | PDK2 |
| 2.24E-117 | sig_down | EGR1 |
| 1.56E-108 | sig_up | TSPYL5 |
| 2.50E-128 | sig_up | GCH1 |
| 2.38E-43 | sig_down | MYLK |
| 1.03E-109 | sig_up | GCHFR |
| 9.99E-149 | sig_up | RPP25 |
| 1.10E-180 | sig_up | BEX2 |
| 7.69E-107 | sig_up | NRIP3 |
| 6.89221575948539e-321 | sig_down | TP63 |
| 7.41E-140 | sig_up | SMPD1 |
| 9.91E-95 | sig_up | KLF2 |
| 3.02E-44 | sig_down | ZBTB16 |
| 7.08E-118 | sig_up | PARD6A |
| 5.51E-115 | sig_up | GPRC5C |
| 1.80E-155 | sig_up | MT1F |
| 5.89E-99 | sig_up | NKAPL |
| 7.49E-102 | sig_up | CGN |
| 3.03E-112 | sig_up | CBR3 |
| 1.06E-125 | sig_up | MITF |
| 4.75E-59 | sig_up | TNFAIP2 |
| 1.15E-56 | sig_up | AKAP12 |
| 2.32E-11 | sig_down | S100A8 |
| 8.00E-117 | sig_up | IRF7 |
| 4.94E-118 | sig_up | TDRD7 |
| 2.77E-92 | sig_up | TP53I11 |
| 1.34E-84 | sig_up | RND2 |
| 1.43E-59 | sig_up | ICAM1 |
| 2.40E-137 | sig_down | NFKBIZ |
| 2.94E-98 | sig_down | INHBA |

| | | |
|---|---|---|
| 6.65E-103 | sig_up | GKAP1 |
| 2.43E-172 | sig_up | DNAJB9 |
| 3.47E-115 | sig_up | RGS9 |
| 3.62E-07 | sig_down | IVL |
| 5.04E-112 | sig_up | HERC6 |
| 5.41E-109 | sig_up | OTUB2 |
| 0 | sig_down | ZFP36L2 |
| 4.95E-169 | sig_up | CCNE1 |
| 5.56E-87 | sig_up | IL11 |
| 6.85E-108 | sig_up | PTGES |
| 2.55E-102 | sig_up | CCDC69 |
| 3.12E-65 | sig_up | TNFRSF18 |
| 5.64E-72 | sig_up | IL23A |
| 2.44E-80 | sig_up | TGM2 |
| 9.10E-100 | sig_up | SERTAD4 |
| 6.59E-54 | sig_down | SYT8 |
| 1.02E-215 | sig_down | DST |
| 1.09E-231 | sig_down | ETS2 |
| 3.51E-80 | sig_up | CTSF |
| 6.51E-196 | sig_down | DDIT4 |
| 8.94675589649878e-310 | sig_down | ZFP36L1 |
| 3.88E-177 | sig_down | PHLDA1 |
| 1.49E-63 | sig_up | RASGRP2 |
| 7.60E-89 | sig_up | DMTN |
| 1.18E-120 | sig_up | SAMHD1 |
| 5.27E-34 | sig_up | MT1G |
| 2.69E-145 | sig_down | DSG3 |
| 1.21E-19 | sig_up | CXCL8 |
| 1.25E-61 | sig_up | OAS1 |
| 9.21E-71 | sig_up | NIBAN1 |
| 3.59E-75 | sig_up | MAP7D2 |
| 1.47E-203 | sig_up | CRB3 |
| 4.54E-110 | sig_up | B3GNT3 |
| 3.88E-131 | sig_up | GABARAPL1 |
| 4.92E-89 | sig_up | DPF1 |
| 1.27E-65 | sig_down | LGALS9C |
| 4.77E-50 | sig_up | CAMK2B |
| 1.22E-42 | sig_up | PCDH1 |
| 1.24E-114 | sig_down | VSNL1 |
| 9.33E-222 | sig_down | JAG1 |
| 1.60E-139 | sig_up | YPEL3 |
| 1.06E-105 | sig_up | MAP1B |
| 5.42E-147 | sig_down | TENM2 |

| | | |
|---|---|---|
| 3.13E-135 | sig_up | PLAUR |
| 2.29E-94 | sig_up | DNAJA4 |
| 3.07E-69 | sig_up | TMEM121 |
| 4.73E-101 | sig_up | PKN1 |
| 1.94E-61 | sig_down | TP53AIP1 |
| 4.64E-63 | sig_up | GPX2 |
| 1.24E-113 | sig_up | PTAFR |
| 3.56E-141 | sig_down | IRS1 |
| 1.00E-35 | sig_up | CNFN |
| 1.01E-85 | sig_up | CLDN23 |
| 2.07E-65 | sig_up | FUT3 |
| 7.29E-73 | sig_up | FAM43A |
| 6.46E-108 | sig_up | ATP6V0E2 |
| 7.04E-73 | sig_up | RASEF |
| 1.62E-68 | sig_up | SDR16C5 |
| 8.75E-190 | sig_up | SVIP |
| 2.25E-78 | sig_up | CCL28 |
| 6.27E-79 | sig_up | NACAD |
| 1.27E-36 | sig_up | ELF3 |
| 3.58E-76 | sig_up | TRPV3 |
| 9.51E-102 | sig_up | LPIN2 |
| 1.33E-126 | sig_up | WFS1 |
| 2.02E-135 | sig_up | KDELR3 |
| 2.02E-77 | sig_up | CDA |
| 4.89E-23 | sig_up | CST6 |
| 1.82E-108 | sig_up | ABHD8 |
| 1.43E-83 | sig_up | C1orf115 |
| 2.67E-63 | sig_up | MARCHF3 |
| 6.84E-82 | sig_up | MAGEH1 |
| 3.65E-193 | sig_up | ITM2C |
| 7.49E-73 | sig_up | FICD |
| 0 | sig_up | SLC25A4 |
| 4.95E-49 | sig_up | BIRC3 |
| 6.41E-75 | sig_down | MMP28 |
| 3.77E-75 | sig_up | USP2 |
| 8.54E-74 | sig_up | RAB3A |
| 6.27E-116 | sig_down | F3 |
| 8.97E-148 | sig_down | PLK2 |
| 8.91E-83 | sig_up | TMEM38A |
| 5.35E-86 | sig_up | ERO1B |
| 2.34E-62 | sig_down | LGALS9B |
| 1.21E-59 | sig_down | FOSB |
| 2.02E-72 | sig_up | EPHX4 |

| | | |
|---|---|---|
| 1.45E-114 | sig_up | HLA-B |
| 3.97E-70 | sig_up | RORA |
| 9.26E-69 | sig_up | HLA-F |
| 1.74E-68 | sig_up | PDE4A |
| 7.80E-296 | sig_down | SERPINB5 |
| 3.54E-75 | sig_up | PRR15 |
| 6.05E-53 | sig_up | LBH |
| 3.65E-52 | sig_down | KCNJ5 |
| 8.11E-77 | sig_up | SLC41A2 |
| 6.44E-90 | sig_up | APH1B |
| 1.38E-180 | sig_down | SNAI2 |
| 2.30E-80 | sig_up | RHPN2 |
| 3.25E-61 | sig_up | PCSK1N |
| 2.87E-69 | sig_up | TMEM255B |
| 4.13E-39 | sig_up | MAP1A |
| 4.08E-61 | sig_down | GABRE |
| 1.59E-157 | sig_up | NEU1 |
| 1.54E-47 | sig_up | FBXO32 |
| 3.39E-197 | sig_down | HMGA2 |
| 4.48E-71 | sig_up | EMILIN2 |
| 4.84E-66 | sig_up | SOCS1 |
| 1.90E-46 | sig_up | METRNL |
| 4.39E-84 | sig_up | KIF13B |
| 2.28E-71 | sig_up | ZNF280B |
| 6.14E-58 | sig_up | INSR |
| 1.52E-251 | sig_down | HTRA1 |
| 1.37E-108 | sig_up | PLPP2 |
| 8.96E-59 | sig_up | RHEBL1 |
| 2.33E-96 | sig_up | ATOSB |
| 8.39E-114 | sig_up | MAP1LC3A |
| 1.48E-170 | sig_down | TANC2 |
| 8.22E-259 | sig_up | HAGH |
| 9.14E-105 | sig_up | ABHD3 |
| 3.57E-56 | sig_up | DIPK1B |
| 5.42E-120 | sig_up | ULBP2 |
| 8.38E-49 | sig_up | ADAP2 |
| 4.70E-96 | sig_up | PLPPR2 |
| 2.75E-66 | sig_up | CD83 |
| 5.72E-60 | sig_up | TSPAN2 |
| 9.39E-50 | sig_up | COBL |
| 1.49E-92 | sig_up | SLC17A5 |
| 3.61E-83 | sig_up | CCDC149 |
| 1.21E-200 | sig_down | CEBPD |

| | | |
|---|---|---|
| 2.16E-67 | sig_down | FLRT3 |
| 2.57E-94 | sig_up | CDKN2D |
| 1.05E-118 | sig_up | STX3 |
| 1.27E-47 | sig_up | CCNA1 |
| 4.42E-78 | sig_up | HSPA2 |
| 7.82E-55 | sig_down | DLK2 |
| 4.97E-139 | sig_up | MAP1S |
| 9.37E-29 | sig_down | IL33 |
| 7.34E-134 | sig_up | CHST12 |
| 4.32E-48 | sig_up | FBLN2 |
| 1.38E-84 | sig_down | FGFR3 |
| 1.53E-78 | sig_down | FST |
| 2.21E-79 | sig_up | XKR8 |
| 1.30E-61 | sig_up | OCLN |
| 1.88E-179 | sig_down | GADD45A |
| 1.60E-102 | sig_up | KIF3C |
| 4.51E-82 | sig_up | MFSD9 |
| 4.27E-79 | sig_up | SMCO4 |
| 7.73E-67 | sig_up | PIERCE1 |
| 2.06E-65 | sig_up | ARFGEF3 |
| 2.52E-84 | sig_up | SELENOM |
| 1.15E-238 | sig_down | SLC38A2 |
| 1.25E-110 | sig_down | PTHLH |
| 2.29E-84 | sig_up | NCOA1 |
| 7.28E-60 | sig_up | FAM171B |
| 1.82E-43 | sig_up | DDIT3 |
| 1.93E-143 | sig_down | FGFBP1 |
| 1.51E-23 | sig_up | QPRT |
| 1.17E-212 | sig_down | KRT14 |
| 1.25E-46 | sig_up | SIPA1L2 |
| 5.99E-37 | sig_down | TNNI2 |
| 4.71E-81 | sig_up | PLLP |
| 3.58E-50 | sig_up | KCNN4 |
| 3.29E-68 | sig_up | CASTOR2 |
| 2.01E-92 | sig_up | PPP2R5B |
| 2.72E-100 | sig_up | HDHD3 |
| 1.05E-62 | sig_up | TFAP2C |
| 2.78E-124 | sig_up | ANKRD9 |
| 1.29E-70 | sig_up | C1orf210 |
| 4.00E-58 | sig_up | FAM131C |
| 3.50E-53 | sig_up | PRKAA2 |
| 3.23E-91 | sig_up | PHTF1 |
| 1.48E-50 | sig_up | HES6 |

| | | |
|---|---|---|
| 9.56E-145 | sig_up | ARMCX1 |
| 1.66E-43 | sig_up | KLK1 |
| 6.73E-58 | sig_up | ATL1 |
| 1.25E-78 | sig_up | IRF1 |
| 8.25E-57 | sig_up | SLC25A42 |
| 6.94E-76 | sig_up | RILP |
| 5.39E-84 | sig_up | POMGNT2 |
| 2.11E-90 | sig_up | OAS3 |
| 8.96E-88 | sig_down | SLC7A11 |
| 1.03E-57 | sig_up | A1BG |
| 4.56E-94 | sig_up | ELOVL4 |
| 6.06E-58 | sig_up | GULP1 |
| 6.41E-53 | sig_up | H2BC11 |
| 1.94E-91 | sig_up | HSPA1B |
| 5.47E-67 | sig_up | USP18 |
| 1.15E-55 | sig_up | NAP1L5 |
| 3.19E-49 | sig_down | PCDH7 |
| 2.53E-141 | sig_down | IL1RAP |
| 9.36E-205 | sig_up | LRIF1 |
| 1.43E-160 | sig_up | MAFG |
| 1.48E-46 | sig_up | SAMD9 |
| 2.90E-93 | sig_down | INPP5D |
| 3.42E-58 | sig_up | TLN2 |
| 1.17E-55 | sig_up | PTGS1 |
| 3.37E-44 | sig_up | GARIN5A |
| 2.55E-138 | sig_up | PINK1 |
| 1.48E-53 | sig_up | SLC2A10 |
| 6.95E-56 | sig_up | TRIM2 |
| 7.51E-56 | sig_up | ZNF483 |
| 4.04E-61 | sig_up | TSPAN33 |
| 1.89E-39 | sig_up | FLNC |
| 1.98E-78 | sig_up | NIPA1 |
| 3.75E-53 | sig_up | E2F1 |
| 1.33E-55 | sig_up | SERPINI1 |
| 1.70E-28 | sig_up | ANKRD22 |
| 4.32E-66 | sig_up | KLHL24 |
| 1.14E-66 | sig_up | PRSS16 |
| 5.81E-112 | sig_up | PRADC1 |
| 2.02E-189 | sig_down | DSC3 |
| 1.02E-69 | sig_up | AIFM2 |
| 3.17E-90 | sig_up | GSPT2 |
| 1.19E-61 | sig_up | TSPAN15 |
| 8.24E-117 | sig_down | KLF6 |

| | | |
|---|---|---|
| 6.36E-150 | sig_down | BCAM |
| 3.61E-71 | sig_up | MAPRE3 |
| 3.95E-229 | sig_down | FAT1 |
| 6.27E-104 | sig_up | RBM38 |
| 1.68E-73 | sig_up | TRAPPC14 |
| 3.44E-50 | sig_up | OSGIN1 |
| 1.41E-183 | sig_down | AJUBA |
| 1.73E-69 | sig_up | PGM2L1 |
| 3.66E-49 | sig_up | SLC46A3 |
| 9.84E-52 | sig_down | ARRDC4 |
| 3.97E-97 | sig_up | SHFL |
| 1.94E-34 | sig_down | LGALS7B |
| 3.21E-49 | sig_up | TMEM61 |
| 1.95E-121 | sig_up | TP53I3 |
| 1.56E-74 | sig_down | DUSP1 |
| 3.55E-95 | sig_up | ZFAND2A |
| 1.50E-139 | sig_down | JUN |
| 4.71E-75 | sig_down | EPPK1 |
| 7.36E-86 | sig_down | SIK1 |
| 3.50E-63 | sig_up | P2RX4 |
| 1.25E-160 | sig_up | ING2 |
| 3.71E-67 | sig_up | MAPRE2 |
| 1.62E-33 | sig_up | CCNE2 |
| 6.13E-28 | sig_up | AGR2 |
| 4.44E-45 | sig_up | PLEKHG6 |
| 1.48E-181 | sig_up | SEPHS2 |
| 1.84E-83 | sig_down | VCAN |
| 7.97E-120 | sig_up | CFL2 |
| 7.54E-80 | sig_up | GCA |
| 2.31E-41 | sig_up | PLCB4 |
| 5.42E-47 | sig_up | LYPD6B |
| 1.25E-77 | sig_up | HMGN5 |
| 6.32E-87 | sig_down | IFFO2 |
| 1.21E-74 | sig_up | BPGM |
| 7.94E-47 | sig_up | HDAC9 |
| 9.03E-51 | sig_up | BICDL1 |
| 3.08E-234 | sig_down | EGFR |
| 1.23E-96 | sig_up | CPEB4 |
| 1.15E-10 | sig_up | KRTAP2-3 |
| 3.75E-40 | sig_up | MEIS3 |
| 1.09E-48 | sig_up | MFAP3L |
| 2.98E-75 | sig_up | CCDC68 |
| 6.06E-87 | sig_down | MTSS1 |

| | | |
|---|---|---|
| 4.81E-100 | sig_up | ADCK2 |
| 5.22E-68 | sig_up | ICA1 |
| 2.29E-173 | sig_down | TNS4 |
| 2.46E-52 | sig_up | PGPEP1 |
| 7.03E-39 | sig_up | NCF2 |
| 1.47E-57 | sig_up | LIN7B |
| 3.01E-59 | sig_up | TBX3 |
| 3.27E-153 | sig_down | SH3PXD2A |
| 1.01E-75 | sig_up | TPST1 |
| 1.69E-51 | sig_up | HAPLN3 |
| 1.56E-43 | sig_up | PAK3 |
| 6.82E-98 | sig_down | PKP1 |
| 5.33E-105 | sig_down | FAT2 |
| 0.000148031 | sig_down | RHCG |

| | logFC | logCPM | LR | PValue | FDR | sig |
|---|---|---|---|---|---|---|
| CXCL14 | -3.459869126 | 6.353238461 | 309.7597405 | 2.46E-69 | 2.20E-67 | sig_down |
| MX1 | 2.899949519 | 5.927854159 | 698.2626204 | 7.14E-154 | 3.33E-151 | sig_up |
| IFI6 | 2.655397806 | 5.907387388 | 538.4126573 | 4.18E-119 | 1.15E-116 | sig_up |
| SPRR2A | -2.154749212 | 7.138670841 | 76.28256045 | 2.46E-18 | 2.90E-17 | sig_down |
| ISG15 | 2.124135529 | 6.949905404 | 1057.481964 | 5.76E-232 | 1.03E-228 | sig_up |
| SLPI | -1.795095362 | 8.039574068 | 241.9512203 | 1.48E-54 | 8.80E-53 | sig_down |
| SPRR1B | -1.753843759 | 7.840861558 | 81.95625053 | 1.39E-19 | 1.80E-18 | sig_down |
| S100A8 | -1.674320849 | 8.411063683 | 65.79743071 | 5.00E-16 | 5.05E-15 | sig_down |
| S100A9 | -1.575961958 | 10.28958057 | 119.0186879 | 1.04E-27 | 2.14E-26 | sig_down |
| CRABP2 | -1.557668642 | 6.719432285 | 206.2814556 | 8.90E-47 | 4.13E-45 | sig_down |
| KRT6B | -1.534078544 | 8.778970145 | 186.0562088 | 2.31E-42 | 9.10E-41 | sig_down |
| RRAD | -1.459986872 | 5.973777394 | 61.50432668 | 4.42E-15 | 4.15E-14 | sig_down |
| SERPINB3 | -1.425777795 | 6.237492444 | 122.3263333 | 1.96E-28 | 4.15E-27 | sig_down |
| GJB6 | -1.415630692 | 6.396488236 | 411.2788062 | 1.93E-91 | 3.09E-89 | sig_down |
| KRT16 | -1.40287263 | 9.889067274 | 245.7829569 | 2.16E-55 | 1.35E-53 | sig_down |
| DDX60 | 1.402750205 | 5.59188121 | 378.1715492 | 3.11E-84 | 4.07E-82 | sig_up |
| CKB | -1.389908391 | 6.935421419 | 321.1686865 | 8.06E-72 | 7.72E-70 | sig_down |
| RHCG | -1.361953679 | 6.563495411 | 27.0569735 | 1.98E-07 | 8.54E-07 | sig_down |
| LCN2 | -1.354669617 | 6.016073163 | 53.1684293 | 3.06E-13 | 2.48E-12 | sig_down |
| AKAP12 | 1.344263521 | 5.804589289 | 141.1108724 | 1.52E-32 | 3.96E-31 | sig_up |
| TXNIP | -1.327099737 | 8.184549816 | 680.5581858 | 5.05E-150 | 2.08E-147 | sig_down |
| KLK10 | -1.304459709 | 6.116031322 | 84.65912109 | 3.55E-20 | 4.71E-19 | sig_down |
| SERPINB13 | -1.283679121 | 5.639762619 | 129.4044392 | 5.53E-30 | 1.28E-28 | sig_down |
| SPINK5 | -1.278055535 | 6.17195949 | 175.8422702 | 3.92E-40 | 1.43E-38 | sig_down |
| OAS1 | 1.272280042 | 5.467673055 | 242.5602739 | 1.09E-54 | 6.63E-53 | sig_up |
| CLIC3 | -1.270905554 | 6.33619675 | 85.70570554 | 2.09E-20 | 2.84E-19 | sig_down |
| MAF | -1.26936867 | 5.597697392 | 224.7264057 | 8.42E-51 | 4.39E-49 | sig_down |
| KLK5 | -1.265672175 | 6.372393409 | 168.7830137 | 1.36E-38 | 4.62E-37 | sig_down |
| HSPB1 | -1.222378918 | 8.870700308 | 433.3103398 | 3.09E-96 | 5.82E-94 | sig_down |
| ARRDC4 | -1.212936269 | 6.014117339 | 346.905731 | 2.00E-77 | 2.14E-75 | sig_down |
| IRF7 | 1.193997127 | 5.540494853 | 317.2552094 | 5.74E-71 | 5.40E-69 | sig_up |
| SAA2 | -1.111543992 | 6.5387552 | 40.01455988 | 2.52E-10 | 1.53E-09 | sig_down |
| KLK11 | -1.105653811 | 5.806368921 | 66.42927542 | 3.63E-16 | 3.68E-15 | sig_down |
| GJB2 | -1.086510801 | 8.177396724 | 494.3969935 | 1.57E-109 | 3.84E-107 | sig_down |
| LYPD3 | -1.072793835 | 5.950282367 | 65.12369185 | 7.03E-16 | 7.01E-15 | sig_down |
| AQP3 | -1.070711288 | 6.784875031 | 136.3763533 | 1.65E-31 | 4.05E-30 | sig_down |
| SLC2A1 | -1.060135296 | 7.467692115 | 606.2245875 | 7.41E-134 | 2.65E-131 | sig_down |
| SESN3 | -1.011222456 | 6.465938434 | 407.0248898 | 1.63E-90 | 2.50E-88 | sig_down |
| LY6D | -1.004305125 | 6.265883698 | 19.59656171 | 9.56E-06 | 3.25E-05 | sig_down |

gene
CXCL14
MX1
IFI6
SPRR2A
ISG15
SLPI
SPRR1B
S100A8
S100A9
CRABP2
KRT6B
RRAD
SERPINB3
GJB6
KRT16
DDX60
CKB
RHCG
LCN2
AKAP12
TXNIP
KLK10
SERPINB13
SPINK5
OAS1
CLIC3
MAF
KLK5
HSPB1
ARRDC4
IRF7
SAA2
KLK11
GJB2
LYPD3
AQP3
SLC2A1
SESN3
LY6D

| | logFC | logCPM | LR | PValue | FDR |
|---|---|---|---|---|---|
| ANXA2R | -2.374178864 | 5.708623082 | 344.6347543 | 6.24E-77 | 1.72E-74 |
| FOXQ1 | -2.133772948 | 6.085607687 | 387.5029114 | 2.89E-86 | 1.51E-83 |
| MX1 | 1.904531485 | 6.262133673 | 541.6989191 | 8.05E-120 | 1.77E-116 |
| IFI44L | 1.862324631 | 5.492386113 | 371.5217782 | 8.73E-83 | 3.55E-80 |
| IFI6 | 1.849580077 | 6.37999722 | 479.575039 | 2.64E-106 | 2.91E-103 |
| SPRR2A | -1.792788312 | 7.197019804 | 79.03506504 | 6.10E-19 | 5.26E-18 |
| PRSS22 | -1.755884512 | 5.569182474 | 306.6968805 | 1.14E-68 | 2.21E-66 |
| IFI27 | 1.715303323 | 5.936345372 | 134.9206146 | 3.44E-31 | 8.10E-30 |
| BST2 | 1.618018102 | 5.465056839 | 236.2452869 | 2.59E-53 | 2.61E-51 |
| SPRR1B | -1.559618493 | 7.803361103 | 88.89941826 | 4.15E-21 | 4.29E-20 |
| C15orf48 | -1.516028337 | 5.751891922 | 86.68452399 | 1.27E-20 | 1.26E-19 |
| OAS2 | 1.492974733 | 5.476430981 | 434.6148726 | 1.61E-96 | 1.18E-93 |
| LCN2 | -1.470408262 | 6.071824293 | 103.8268351 | 2.21E-24 | 3.01E-23 |
| CLU | -1.408334786 | 5.663428431 | 394.4973007 | 8.69E-88 | 5.02E-85 |
| RBP7 | -1.395086089 | 5.589679731 | 308.0843679 | 5.71E-69 | 1.12E-66 |
| S100A9 | -1.383781393 | 10.28064737 | 127.4024534 | 1.52E-29 | 3.10E-28 |
| IRF9 | 1.35741327 | 5.539948018 | 799.9775366 | 5.46E-176 | 3.00E-172 |
| WFDC2 | -1.323950773 | 5.520001102 | 212.6996794 | 3.54E-48 | 2.72E-46 |
| SLPI | -1.309863956 | 8.161787785 | 192.0338493 | 1.14E-43 | 6.72E-42 |
| TP53INP2 | -1.301573475 | 5.402468859 | 355.6187045 | 2.53E-79 | 8.44E-77 |
| IFIT3 | 1.298214996 | 5.594000508 | 132.5771274 | 1.12E-30 | 2.52E-29 |
| IFI44 | 1.286795215 | 5.47629286 | 275.8690561 | 5.97E-62 | 8.75E-60 |
| IFIT1 | 1.281516643 | 5.78551788 | 75.93193254 | 2.94E-18 | 2.40E-17 |
| KLK5 | -1.26558049 | 6.411586588 | 273.5630723 | 1.90E-61 | 2.71E-59 |
| S100A8 | -1.257717599 | 8.260491676 | 51.94182104 | 5.72E-13 | 2.88E-12 |
| IFITM1 | 1.248513508 | 5.551850854 | 499.7823872 | 1.06E-110 | 1.46E-107 |
| TIMP2 | -1.244023633 | 5.646193818 | 250.7249261 | 1.80E-56 | 2.07E-54 |
| OAS1 | 1.230658761 | 5.493210491 | 336.7665437 | 3.23E-75 | 8.45E-73 |
| APOE | -1.227210776 | 7.245742067 | 283.9947817 | 1.01E-63 | 1.64E-61 |
| SAA2 | -1.222388067 | 6.48881308 | 72.27643033 | 1.87E-17 | 1.42E-16 |
| KLK11 | -1.221631014 | 5.846234547 | 149.8581309 | 1.86E-34 | 5.67E-33 |
| RHCG | -1.171671812 | 6.42576708 | 33.50387568 | 7.11E-09 | 2.34E-08 |
| CLEC11A | -1.149006109 | 5.398728961 | 236.2961868 | 2.53E-53 | 2.57E-51 |
| OTUB2 | -1.136261833 | 5.408331605 | 352.6458459 | 1.12E-78 | 3.43E-76 |
| KRT6B | -1.13585546 | 8.795401701 | 160.0874816 | 1.08E-36 | 3.78E-35 |
| DDX60 | 1.131179859 | 5.696370216 | 323.9091425 | 2.04E-72 | 4.87E-70 |
| RAET1L | -1.126586197 | 5.413204249 | 161.4802485 | 5.37E-37 | 1.95E-35 |
| EPSTI1 | 1.094280579 | 5.461749418 | 314.468172 | 2.32E-70 | 5.00E-68 |
| SPINK6 | -1.090131308 | 5.477730258 | 99.38098402 | 2.08E-23 | 2.62E-22 |
| KLK10 | -1.084756989 | 6.120440632 | 96.99648407 | 6.95E-23 | 8.25E-22 |
| CXCL14 | -1.071575196 | 6.256382252 | 42.20137632 | 8.23E-11 | 3.35E-10 |
| CLIC3 | -1.06263638 | 6.274336698 | 97.85957023 | 4.49E-23 | 5.42E-22 |

| | | | | | |
|---|---|---|---|---|---|
| HSPA1A | -1.048310385 | 6.562981221 | 521.4462496 | 2.05E-115 | 3.22E-112 |
| SYNGR1 | -1.042755896 | 5.528654999 | 223.5697264 | 1.51E-50 | 1.33E-48 |
| TMEM125 | -1.020164379 | 5.396961903 | 224.4035002 | 9.91E-51 | 8.85E-49 |
| INA | -1.006972048 | 5.371242502 | 226.2719613 | 3.88E-51 | 3.58E-49 |

| sig | gene |
|---|---|
| sig_down | ANXA2R |
| sig_down | FOXQ1 |
| sig_up | MX1 |
| sig_up | IFI44L |
| sig_up | IFI6 |
| sig_down | SPRR2A |
| sig_down | PRSS22 |
| sig_up | IFI27 |
| sig_up | BST2 |
| sig_down | SPRR1B |
| sig_down | C15orf48 |
| sig_up | OAS2 |
| sig_down | LCN2 |
| sig_down | CLU |
| sig_down | RBP7 |
| sig_down | S100A9 |
| sig_up | IRF9 |
| sig_down | WFDC2 |
| sig_down | SLPI |
| sig_down | TP53INP2 |
| sig_up | IFIT3 |
| sig_up | IFI44 |
| sig_up | IFIT1 |
| sig_down | KLK5 |
| sig_down | S100A8 |
| sig_up | IFITM1 |
| sig_down | TIMP2 |
| sig_up | OAS1 |
| sig_down | APOE |
| sig_down | SAA2 |
| sig_down | KLK11 |
| sig_down | RHCG |
| sig_down | CLEC11A |
| sig_down | OTUB2 |
| sig_down | KRT6B |
| sig_up | DDX60 |
| sig_down | RAET1L |
| sig_up | EPSTI1 |
| sig_down | SPINK6 |
| sig_down | KLK10 |
| sig_down | CXCL14 |
| sig_down | CLIC3 |

| | |
|---|---|
| sig_down | HSPA1A |
| sig_down | SYNGR1 |
| sig_down | TMEM125 |
| sig_down | INA |

| | logFC | logCPM | LR | PValue |
|---|---|---|---|---|
| ANXA2R | 3.706288045 | 5.967235313 | 807.1882273 | 1.48E-177 |
| FOXQ1 | 3.50711751 | 6.439089964 | 957.8137685 | 2.66E-210 |
| PRSS22 | 2.733349433 | 5.706810998 | 918.086143 | 1.15E-187 |
| CLU | 2.585867518 | 5.871780078 | 1430.450787 | 5.08E-299 |
| RBP7 | 2.533190064 | 5.774463266 | 1044.160883 | 4.52E-229 |
| C15orf48 | 2.482112882 | 5.931353551 | 305.4045797 | 2.19E-68 |
| TP53INP2 | 2.329333982 | 5.52405324 | 1050.574098 | 1.83E-230 |
| WFDC2 | 2.292877984 | 5.652394412 | 611.1414874 | 6.32E-135 |
| RNF223 | 2.187926578 | 5.440581279 | 824.9040222 | 2.08E-181 |
| CLEC11A | 2.167450461 | 5.518962431 | 774.7194149 | 1.69E-170 |
| TIMP2 | 2.119562983 | 5.780745789 | 695.3074242 | 3.13E-153 |
| TMEM125 | 2.004154796 | 5.51478117 | 788.9870389 | 1.34E-173 |
| HSPA1A | 1.963941729 | 6.80204517 | 1365.137824 | 7.91E-299 |
| PCDH1 | 1.960700043 | 5.442444083 | 609.455895 | 1.47E-134 |
| UCHL1 | 1.92616177 | 5.975008079 | 467.4821776 | 1.13E-103 |
| INA | 1.913496184 | 5.475301814 | 689.3862182 | 6.08E-152 |
| SYNGR1 | 1.887941955 | 5.64808253 | 618.4896529 | 1.59E-136 |
| OTUB2 | 1.881520341 | 5.497582504 | 792.6678989 | 2.12E-174 |
| CLDN4 | 1.872739296 | 5.952870869 | 296.0368193 | 2.41E-66 |
| ENO2 | 1.860893654 | 5.462157437 | 552.8131866 | 3.08E-122 |
| LCN2 | 1.812136699 | 6.03643604 | 146.8632714 | 8.41E-34 |
| FOS | -1.802771044 | 7.46311394 | 959.6041655 | 1.08E-210 |
| RAB6B | 1.798584277 | 5.410686335 | 627.8270916 | 1.48E-138 |
| AMIGO2 | -1.77313944 | 6.703955467 | 1154.228652 | 5.41E-253 |
| BEX5 | 1.748392706 | 5.423843162 | 554.805109 | 1.13E-122 |
| SPRR2A | 1.731165119 | 6.873446071 | 60.35755724 | 7.91E-15 |
| APOE | 1.713573321 | 7.267649114 | 402.5635502 | 1.52E-89 |
| HMOX1 | 1.712761285 | 5.440303311 | 493.9531375 | 1.97E-109 |
| TLCD4 | 1.710230649 | 5.419144495 | 560.2115576 | 7.56E-124 |
| PDK2 | 1.70143063 | 5.683323004 | 769.9463727 | 1.85E-169 |
| BEX2 | 1.691272223 | 6.062219372 | 932.0726649 | 1.05E-204 |
| TNFAIP2 | 1.68411685 | 5.433745509 | 394.3919174 | 9.16E-88 |
| KLF7 | -1.671170806 | 6.572162241 | 1077.76377 | 2.25E-236 |
| TGM1 | -1.667927237 | 6.458485157 | 212.1965266 | 4.56E-48 |
| TSPYL5 | 1.664773068 | 5.495137628 | 563.2164937 | 1.68E-124 |
| CCK | 1.659340545 | 5.567983601 | 380.046114 | 1.22E-84 |
| MYC | -1.656875743 | 8.007714257 | 1002.101556 | 6.27E-220 |
| PCSK1N | 1.654994101 | 5.669815444 | 520.7461307 | 2.91E-115 |
| TP53I11 | 1.654957293 | 5.553584514 | 538.821744 | 3.40E-119 |
| RPP25 | 1.652098354 | 5.722381741 | 705.8499428 | 1.60E-155 |
| NID1 | 1.650350763 | 5.491317015 | 261.0881753 | 9.94E-59 |
| PLAAT3 | 1.64579176 | 5.432940147 | 349.3062641 | 6.00E-78 |

| | | | | |
|---|---|---|---|---|
| FAM241B | 1.63861599 | 5.857137 | 1080.275843 | 6.40E-237 |
| CTSF | 1.607666499 | 5.488335391 | 477.8868905 | 6.16E-106 |
| PTGES | 1.602632068 | 5.591965542 | 590.5727193 | 1.88E-130 |
| INHBA | -1.600910054 | 7.279051345 | 528.9151716 | 4.86E-117 |
| YPEL3 | 1.584053817 | 6.056957018 | 800.6956696 | 3.81E-176 |
| GJB6 | -1.574588788 | 6.128731939 | 394.906975 | 7.07E-88 |
| GPRC5C | 1.572310478 | 5.461776614 | 517.2212187 | 1.70E-114 |
| KLK11 | 1.572049898 | 5.856080185 | 234.7637707 | 5.45E-53 |
| SNCG | 1.571698408 | 5.906250228 | 461.5423838 | 2.22E-102 |
| FBXO32 | 1.567860968 | 5.485783988 | 427.8818828 | 4.70E-95 |
| GCH1 | 1.553199844 | 5.497962952 | 560.9850203 | 5.13E-124 |
| SYNM | 1.534406509 | 5.401643471 | 451.5389267 | 3.34E-100 |
| METRNL | 1.505601237 | 5.784646571 | 380.1919514 | 1.13E-84 |
| CXCL8 | 1.495940512 | 6.049168461 | 107.6630233 | 3.19E-25 |
| MAP1LC3A | 1.495311873 | 6.094871367 | 785.2333415 | 8.76E-173 |
| TMEM38A | 1.483215764 | 5.573648952 | 534.3859615 | 3.14E-118 |
| NKAPL | 1.47836666 | 5.489221051 | 399.218933 | 8.15E-89 |
| RASGRP2 | 1.47226423 | 5.435033898 | 333.1520045 | 1.98E-74 |
| RAET1L | 1.471822723 | 5.45419288 | 274.0391773 | 1.49E-61 |
| GABARAPL1 | 1.468319549 | 5.764569591 | 668.5915231 | 2.02E-147 |
| KLF2 | 1.454652496 | 5.40276096 | 384.7500963 | 1.15E-85 |
| CCNE1 | 1.453881635 | 5.903926691 | 719.4976538 | 1.72E-158 |
| RETREG1 | 1.441499503 | 5.414614939 | 430.3936204 | 1.33E-95 |
| ADAP2 | 1.44026753 | 5.482894326 | 364.3440096 | 3.19E-81 |
| EGR1 | -1.439287504 | 6.539069171 | 475.2485558 | 2.31E-105 |
| ACP5 | 1.439282307 | 5.553115453 | 430.0917018 | 1.55E-95 |
| DNAJB9 | 1.435787078 | 6.005767661 | 735.0689635 | 7.08E-162 |
| F3 | -1.433422708 | 9.559414846 | 657.4580705 | 5.34E-145 |
| TP63 | -1.425095681 | 6.82465802 | 1214.191508 | 5.02E-266 |
| SIPA1L2 | 1.412461618 | 5.504802775 | 384.651613 | 1.21E-85 |
| GKAP1 | 1.412261366 | 5.400101025 | 424.4466769 | 2.63E-94 |
| FAM43A | 1.397326112 | 5.408989916 | 393.8101911 | 1.23E-87 |
| TNFRSF18 | 1.397283817 | 5.439901238 | 286.9973943 | 2.24E-64 |
| RORA | 1.396166053 | 5.454956625 | 401.2355364 | 2.96E-89 |
| KLHL24 | 1.392470384 | 5.684998418 | 476.4646827 | 1.26E-105 |
| ABHD8 | 1.391767577 | 5.637117081 | 555.3715947 | 8.54E-123 |
| PARD6A | 1.39077568 | 5.509916779 | 442.6005811 | 2.94E-98 |
| INSR | 1.389145102 | 5.399883908 | 371.4610775 | 9.00E-83 |
| ELF3 | 1.388193148 | 5.557048482 | 196.0750102 | 1.50E-44 |
| SMPD1 | 1.383307414 | 5.524226598 | 498.6418505 | 1.88E-110 |
| DDIT3 | 1.379648937 | 5.736308089 | 258.5677317 | 3.52E-58 |
| KLK5 | 1.377669242 | 6.323975749 | 274.4186883 | 1.24E-61 |
| MARCHF3 | 1.37710609 | 5.423365809 | 362.2806693 | 8.97E-81 |

| | | | | |
|---|---|---|---|---|
| NIBAN1 | 1.375639545 | 5.643358656 | 333.0735178 | 2.06E-74 |
| RASEF | 1.370575893 | 5.413734373 | 380.4326999 | 1.00E-84 |
| DMTN | 1.369346724 | 5.418011603 | 399.5365152 | 6.95E-89 |
| PLPPR2 | 1.365864893 | 5.606480716 | 556.8961012 | 3.98E-123 |
| TMEM121 | 1.362186807 | 5.491491278 | 341.088654 | 3.70E-76 |
| G0S2 | -1.361807065 | 9.188077511 | 229.1237361 | 9.26E-52 |
| DIPK1B | 1.360185119 | 5.452555835 | 330.4688673 | 7.60E-74 |
| HES6 | 1.360077618 | 5.690425559 | 354.6084152 | 4.20E-79 |
| CST6 | 1.359632443 | 5.643680136 | 147.7861353 | 5.28E-34 |
| RND2 | 1.351095782 | 5.405621 | 309.9102827 | 2.28E-69 |
| CCDC69 | 1.349551769 | 5.418062874 | 396.9008393 | 2.60E-88 |
| FLNC | 1.346047485 | 5.482603204 | 286.2966489 | 3.19E-64 |
| GCHFR | 1.34445505 | 5.678519318 | 411.5312488 | 1.70E-91 |
| SNX10 | 1.342280019 | 5.633846281 | 494.7555416 | 1.32E-109 |
| TNNI3 | 1.339540356 | 5.466937934 | 289.7147591 | 5.74E-65 |
| NRIP3 | 1.33704387 | 5.439411419 | 357.4471667 | 1.01E-79 |
| CASTOR2 | 1.332864376 | 5.503935214 | 425.5102423 | 1.54E-94 |
| MITF | 1.332055353 | 5.574102109 | 432.1002871 | 5.67E-96 |
| ATOSB | 1.331839713 | 5.697261876 | 556.9525863 | 3.87E-123 |
| TGM2 | 1.330712755 | 5.751777416 | 341.9655613 | 2.38E-76 |
| FUT3 | 1.329840026 | 5.435655611 | 332.9253497 | 2.22E-74 |
| BICDL2 | 1.329108127 | 5.532703445 | 319.2919921 | 2.07E-71 |
| ZFP36L2 | -1.317465789 | 8.651635846 | 1156.434947 | 1.79E-253 |
| IFITM1 | -1.316593162 | 5.680184521 | 286.3122013 | 3.16E-64 |
| TP53I3 | 1.307951458 | 6.675267966 | 998.6926061 | 3.45E-219 |
| RAB3A | 1.306526866 | 5.389777753 | 368.1422639 | 4.75E-82 |
| NFKBIZ | -1.30550654 | 6.074030546 | 462.5211042 | 1.36E-102 |
| WFS1 | 1.299063129 | 5.695167905 | 540.7788335 | 1.28E-119 |
| CRB3 | 1.298235476 | 6.019487415 | 827.7214819 | 5.07E-182 |
| ID1 | -1.297853844 | 7.080987467 | 481.1310823 | 1.21E-106 |
| ICAM1 | 1.29575632 | 5.578131511 | 201.4013214 | 1.03E-45 |
| CAMK2B | 1.295026256 | 5.588308139 | 198.2269489 | 5.09E-45 |
| S100A4 | 1.294874876 | 5.923962169 | 224.8839955 | 7.78E-51 |
| LRIF1 | 1.285375919 | 6.48831943 | 1190.257116 | 7.99E-261 |
| FST | -1.283566167 | 9.2475039 | 445.596262 | 6.55E-99 |
| DBNDD1 | 1.281780918 | 5.44314074 | 345.0346003 | 5.11E-77 |
| CXADR | 1.280186085 | 6.004866808 | 647.415384 | 8.15E-143 |
| HMGA2 | -1.277318213 | 8.205919075 | 1012.273205 | 3.86E-222 |
| OLFM2 | 1.274013778 | 5.445677146 | 243.5989161 | 6.46E-55 |
| MAGEH1 | 1.273593971 | 5.50498323 | 354.9301678 | 3.58E-79 |
| FAM131C | 1.273460453 | 5.462640103 | 338.6262714 | 1.27E-75 |
| SLC2A1 | -1.272157839 | 7.142604696 | 701.1245117 | 1.70E-154 |
| FILIP1L | -1.269368711 | 5.775199073 | 196.882755 | 1.00E-44 |

| | | | | |
|---|---|---|---|---|
| PRADC1 | 1.268535297 | 5.973394489 | 679.4107459 | 8.98E-150 |
| SLPI | 1.263948537 | 7.881155001 | 133.4328669 | 7.27E-31 |
| IRF9 | -1.263422622 | 5.681529018 | 400.2275607 | 4.91E-89 |
| DST | -1.257490015 | 10.39382702 | 938.979425 | 3.30E-206 |
| TXNIP | -1.255874063 | 7.504929844 | 312.0177561 | 7.94E-70 |
| SELENOM | 1.255299411 | 6.039465875 | 500.7486225 | 6.53E-111 |
| STX3 | 1.253582614 | 5.845816789 | 590.6066227 | 1.85E-130 |
| DNAJA4 | 1.250068261 | 5.501333869 | 367.6704848 | 6.02E-82 |
| PIERCE1 | 1.246305019 | 5.450331458 | 364.6725839 | 2.70E-81 |
| NOTCH1 | -1.242100986 | 6.493542576 | 734.0132577 | 1.20E-161 |
| MAP7D2 | 1.239481308 | 5.446056505 | 279.1789536 | 1.13E-62 |
| CGN | 1.237982917 | 5.516947718 | 299.1414841 | 5.07E-67 |
| RHPN2 | 1.23508554 | 5.500919751 | 360.8575821 | 1.83E-80 |
| CLDN23 | 1.231677477 | 5.406168035 | 333.3528407 | 1.79E-74 |
| DPF1 | 1.230272084 | 5.424588712 | 322.3125341 | 4.54E-72 |
| TRPV3 | 1.227706546 | 5.39309626 | 308.4429502 | 4.77E-69 |
| ANKRD22 | 1.224333849 | 5.450423901 | 206.9321389 | 6.42E-47 |
| QPRT | 1.222776911 | 5.474150233 | 141.0723313 | 1.55E-32 |
| RGS9 | 1.220883233 | 5.561694566 | 367.029274 | 8.30E-82 |
| RHOF | 1.220669339 | 5.76018154 | 485.5215534 | 1.34E-107 |
| PRR15 | 1.219641799 | 5.380826888 | 329.5254359 | 1.22E-73 |
| ITM2C | 1.219506741 | 6.195200825 | 784.9188951 | 1.03E-172 |
| TTC9 | 1.21530661 | 5.478097138 | 263.4657119 | 3.01E-59 |
| PLPP2 | 1.213337554 | 6.131255973 | 501.1243715 | 5.41E-111 |
| LGALS9C | -1.211052092 | 5.436019128 | 254.5079337 | 2.70E-57 |
| COBL | 1.209883808 | 5.401413319 | 259.1638511 | 2.61E-58 |
| IL11 | 1.207888013 | 5.602886941 | 279.2733837 | 1.08E-62 |
| LAYN | 1.20755684 | 5.860413768 | 332.2302423 | 3.14E-74 |
| FICD | 1.207181121 | 5.454098703 | 300.8744766 | 2.12E-67 |
| MT1G | 1.206115282 | 5.508704094 | 145.9565238 | 1.33E-33 |
| PGF | 1.204395214 | 6.425658199 | 742.9250045 | 1.39E-163 |
| CCL28 | 1.201339142 | 5.476339407 | 296.2782577 | 2.13E-66 |
| HAGH | 1.201030818 | 6.700890328 | 1211.201939 | 2.24E-265 |
| SERPINB3 | -1.198235104 | 5.923659942 | 65.45507186 | 5.95E-16 |
| VSNL1 | -1.197080619 | 5.854578923 | 389.1381456 | 1.27E-86 |
| CNFN | 1.194299693 | 5.464368087 | 113.4140179 | 1.75E-26 |
| NAP1L5 | 1.191681566 | 5.41051118 | 306.0484294 | 1.59E-68 |
| NATD1 | 1.188932328 | 5.395266434 | 301.9610559 | 1.23E-67 |
| JAG1 | -1.188834331 | 7.250018367 | 723.7720667 | 2.03E-159 |
| ZFAND2A | 1.187000979 | 5.834996695 | 533.5673311 | 4.73E-118 |
| GSPT2 | 1.184903662 | 5.716614121 | 474.5137893 | 3.34E-105 |
| SVIP | 1.184665908 | 6.084210483 | 740.8008893 | 4.01E-163 |
| ATP6V0E2 | 1.183622125 | 5.645525614 | 408.2513657 | 8.81E-91 |

| | | | | |
|---|---|---|---|---|
| CHAC1 | 1.179673946 | 5.74528442 | 200.3219089 | 1.78E-45 |
| MAPRE3 | 1.177535758 | 5.517191999 | 387.7481038 | 2.56E-86 |
| ANKRD9 | 1.174560044 | 5.913892339 | 619.1647482 | 1.14E-136 |
| GULP1 | 1.174488641 | 5.488618032 | 290.6729793 | 3.55E-65 |
| MAP1B | 1.172843257 | 6.648918218 | 399.4199392 | 7.37E-89 |
| CCDC149 | 1.172312294 | 5.527594577 | 373.8543791 | 2.71E-83 |
| CDA | 1.171353822 | 6.315305106 | 358.1631893 | 7.07E-80 |
| ERO1B | 1.171195415 | 5.53333992 | 334.8786091 | 8.32E-75 |
| SLC25A4 | 1.168713547 | 6.917902306 | 1379.727294 | 5.34E-302 |
| AIF1L | 1.168523223 | 5.525684612 | 284.2579971 | 8.87E-64 |
| ULBP2 | 1.16797381 | 5.850371698 | 523.7237732 | 6.55E-116 |
| CPEB4 | 1.167818157 | 5.88521244 | 574.6388199 | 5.50E-127 |
| KIF3C | 1.167555321 | 5.711024871 | 482.8165467 | 5.21E-107 |
| MLLT11 | 1.167547674 | 6.221093976 | 527.730431 | 8.80E-117 |
| GJB2 | -1.166341499 | 7.784049275 | 406.3917196 | 2.24E-90 |
| CHST12 | 1.16623693 | 5.776180682 | 591.3388856 | 1.28E-130 |
| ISG20 | 1.165672535 | 6.319594347 | 378.7058994 | 2.38E-84 |
| SLC43A2 | 1.165665601 | 5.517564951 | 341.5206343 | 2.98E-76 |
| KRTAP2-3 | 1.162118767 | 5.979783079 | 71.96137198 | 2.19E-17 |
| USP2 | 1.160731584 | 5.387120586 | 289.4158548 | 6.67E-65 |
| NEU1 | 1.160102801 | 6.011514257 | 635.0963661 | 3.89E-140 |
| SLC46A3 | 1.160021557 | 5.4121121 | 284.3832329 | 8.33E-64 |
| TAF7 | 1.156577151 | 6.882388337 | 1232.821566 | 4.49E-270 |
| FOXL2 | 1.154207146 | 5.426936419 | 246.38003 | 1.60E-55 |
| PDE4A | 1.153821207 | 5.38981111 | 269.4345286 | 1.51E-60 |
| FGFBP1 | -1.152751449 | 10.83912653 | 678.9777515 | 1.11E-149 |
| IFITM2 | -1.152738585 | 5.756406674 | 236.059548 | 2.84E-53 |
| COL8A1 | -1.148194248 | 6.511413699 | 410.5242989 | 2.82E-91 |
| SLC25A42 | 1.14699801 | 5.424114755 | 276.5865645 | 4.16E-62 |
| ATOSA | 1.13971475 | 5.622117271 | 388.2324534 | 2.01E-86 |
| ETS2 | -1.139583314 | 6.977398868 | 738.3386681 | 1.38E-162 |
| IL23A | 1.13863297 | 5.436633158 | 216.1098961 | 6.38E-49 |
| SEPHS2 | 1.13823594 | 6.350668041 | 964.6884516 | 8.51E-212 |
| IL1RAP | -1.137019437 | 6.484482201 | 721.0802824 | 7.79E-159 |
| SPRR1B | 1.135244367 | 7.345056089 | 35.66255826 | 2.35E-09 |
| C1orf115 | 1.133380351 | 5.43928372 | 294.2797287 | 5.81E-66 |
| APH1B | 1.130742215 | 5.561468373 | 340.00351 | 6.37E-76 |
| FOSB | -1.129065791 | 6.184325061 | 204.3078508 | 2.40E-46 |
| PGM2L1 | 1.126243913 | 5.805570247 | 397.7684334 | 1.69E-88 |
| VEGFC | -1.125226227 | 6.287435545 | 477.7834045 | 6.49E-106 |
| MAFG | 1.124455657 | 6.156378389 | 752.9345943 | 9.23E-166 |
| LGALS9B | -1.122568049 | 5.455524534 | 220.9134298 | 5.72E-50 |
| SIK1 | -1.121993151 | 6.276586055 | 437.3124556 | 4.16E-97 |

| | | | | |
|---|---|---|---|---|
| LPIN2 | 1.121988702 | 5.670312234 | 346.4904787 | 2.46E-77 |
| TMEM255B | 1.117541214 | 5.413522942 | 259.9809528 | 1.73E-58 |
| PINK1 | 1.117360455 | 6.23311828 | 646.2775978 | 1.44E-142 |
| TRAPPC14 | 1.117088182 | 5.533733646 | 364.4981561 | 2.95E-81 |
| CXCL14 | -1.116499731 | 5.838098806 | 31.54133688 | 1.95E-08 |
| COL7A1 | -1.115123928 | 6.623546302 | 499.068925 | 1.52E-110 |
| NEFH | 1.113736994 | 5.675706316 | 261.1755391 | 9.51E-59 |
| SLC41A2 | 1.112679754 | 5.382758083 | 280.1245602 | 7.05E-63 |
| DKK1 | -1.112109985 | 7.508450154 | 143.8843545 | 3.77E-33 |
| IRS1 | -1.11055791 | 6.216631863 | 449.9366774 | 7.45E-100 |
| NOL4L | 1.109457479 | 5.531986663 | 317.6140103 | 4.79E-71 |
| SLC17A5 | 1.106633245 | 5.649789254 | 374.4872288 | 1.97E-83 |
| A1BG | 1.106444281 | 5.588661671 | 272.4837132 | 3.26E-61 |
| HES4 | 1.105208126 | 6.317275775 | 261.3215383 | 8.84E-59 |
| EPHX4 | 1.104658589 | 5.420284395 | 246.2188439 | 1.73E-55 |
| ZNF280B | 1.103027995 | 5.470339371 | 269.4225996 | 1.52E-60 |
| TSPAN33 | 1.10260387 | 5.472224108 | 283.4812984 | 1.31E-63 |
| POMGNT2 | 1.099033482 | 5.608098918 | 358.8969777 | 4.90E-80 |
| ZNF483 | 1.098297324 | 5.484505195 | 252.1105993 | 9.00E-57 |
| CCN2 | -1.097141309 | 6.399166235 | 143.8691267 | 3.79E-33 |
| THBS1 | -1.095498415 | 9.699090381 | 498.4347562 | 2.08E-110 |
| ZFP36L1 | -1.094730922 | 8.137765707 | 808.4728134 | 7.76E-178 |
| HSPA1B | 1.092464084 | 6.006719394 | 381.7347499 | 5.22E-85 |
| TFAP2C | 1.091944729 | 5.490945415 | 269.9713915 | 1.15E-60 |
| TPRG1L | 1.090131416 | 5.946471136 | 597.5725996 | 5.65E-132 |
| IL20RB | -1.08920485 | 6.234314554 | 323.5920798 | 2.39E-72 |
| KIF13B | 1.084010946 | 5.570131619 | 315.7189128 | 1.24E-70 |
| PKN1 | 1.082624604 | 5.653881835 | 311.117453 | 1.25E-69 |
| SAA2 | 1.081523729 | 6.28637174 | 43.24156057 | 4.84E-11 |
| MFSD9 | 1.080903942 | 5.550084323 | 330.6008899 | 7.11E-74 |
| B3GNT3 | 1.079870314 | 5.600126757 | 327.1038442 | 4.11E-73 |
| CDCA3 | -1.07970202 | 5.916936224 | 148.5859495 | 3.53E-34 |
| GARIN5A | 1.079536024 | 5.46578131 | 202.7587357 | 5.22E-46 |
| SERPINB5 | -1.078944365 | 8.249248181 | 951.3023727 | 6.91E-209 |
| DLK2 | -1.07661584 | 5.686633565 | 232.7305864 | 1.51E-52 |
| PLK2 | -1.076316547 | 7.64083995 | 458.2607531 | 1.15E-101 |
| KLK10 | 1.072821117 | 6.039930705 | 72.67308795 | 1.53E-17 |
| NCOA1 | 1.072113535 | 5.648184684 | 346.0575053 | 3.06E-77 |
| GABRE | -1.070701124 | 5.495507502 | 225.2361007 | 6.52E-51 |
| ZFYVE28 | 1.069756344 | 5.449209863 | 269.7840252 | 1.26E-60 |
| PPP2R5B | 1.066244945 | 5.656185815 | 381.0208883 | 7.46E-85 |
| CCNA1 | 1.066047024 | 5.462871553 | 169.4543591 | 9.74E-39 |
| MMP28 | -1.064014023 | 5.828656049 | 243.9159501 | 5.51E-55 |

| | | | | |
|---|---|---|---|---|
| MRAS | 1.063480086 | 5.597519261 | 290.2785019 | 4.32E-65 |
| ULK1 | 1.06346371 | 5.569595358 | 289.1713401 | 7.54E-65 |
| SASH1 | 1.0633227 | 5.773237084 | 429.6764206 | 1.91E-95 |
| SNAI2 | -1.062960473 | 6.775163997 | 576.4830133 | 2.18E-127 |
| SLC26A11 | 1.060732862 | 5.404968954 | 257.1572021 | 7.15E-58 |
| TSPAN2 | 1.060202358 | 5.3942647 | 215.4963189 | 8.69E-49 |
| MT1F | 1.060030548 | 5.742148893 | 352.6622517 | 1.12E-78 |
| DUSP8 | 1.059709229 | 5.45241164 | 224.1795582 | 1.11E-50 |
| CDKN2D | 1.058800313 | 5.721753874 | 354.9472716 | 3.55E-79 |
| KIF20A | -1.057533859 | 5.620287744 | 134.067261 | 5.28E-31 |
| HERC5 | 1.056063712 | 5.567495071 | 99.39417349 | 2.07E-23 |
| IFI44L | -1.054805615 | 5.685940829 | 66.23664312 | 4.00E-16 |
| MFAP3L | 1.053471409 | 5.380132199 | 247.220225 | 1.05E-55 |
| RNF208 | 1.052709815 | 5.410260798 | 222.4149269 | 2.69E-50 |
| E2F1 | 1.047151976 | 5.615517708 | 239.8830848 | 4.17E-54 |
| MT1X | -1.04654506 | 9.002333261 | 488.9951664 | 2.36E-108 |
| FAM110C | 1.04623627 | 5.618903365 | 230.2711192 | 5.20E-52 |
| NIPA1 | 1.045505113 | 5.606487041 | 329.3668085 | 1.32E-73 |
| SDR16C5 | 1.045272777 | 5.483717071 | 212.8092684 | 3.35E-48 |
| SECISBP2L | 1.044884329 | 5.819659328 | 444.1381031 | 1.36E-98 |
| BEX4 | 1.043434089 | 6.496942234 | 528.6558108 | 5.54E-117 |
| PCDH7 | -1.042878841 | 5.691183533 | 216.290051 | 5.83E-49 |
| CEBPA | 1.041733757 | 5.492237176 | 208.2826934 | 3.25E-47 |
| ATL1 | 1.041213728 | 5.431457647 | 229.9050395 | 6.25E-52 |
| ADGRF4 | 1.039798475 | 5.525712683 | 236.9414664 | 1.83E-53 |
| FBLN2 | 1.035791291 | 5.467223823 | 178.3637371 | 1.10E-40 |
| CEBPD | -1.034375448 | 7.844765001 | 657.089745 | 6.42E-145 |
| MAP1S | 1.033873162 | 5.82365887 | 505.0226571 | 7.68E-112 |
| S100A9 | 1.032723206 | 9.763077196 | 56.48684665 | 5.66E-14 |
| CYSRT1 | 1.031193349 | 5.427094712 | 123.3057074 | 1.20E-28 |
| CXCL2 | -1.028298823 | 5.70144973 | 97.90637906 | 4.39E-23 |
| SERTAD4 | 1.027593605 | 5.611990389 | 248.626051 | 5.18E-56 |
| ARHGAP42 | 1.027210769 | 5.481285676 | 259.8838965 | 1.82E-58 |
| PHLDA1 | -1.026111513 | 7.703070414 | 444.2681015 | 1.28E-98 |
| GAL | 1.024212602 | 6.559651868 | 412.8730885 | 8.68E-92 |
| MAP3K11 | 1.023721781 | 5.932329972 | 511.238705 | 3.41E-113 |
| XKR8 | 1.022712742 | 5.520894119 | 283.7774363 | 1.13E-63 |
| ABHD3 | 1.022633858 | 5.641666644 | 355.4241382 | 2.79E-79 |
| DDIT4 | -1.020966091 | 8.464875819 | 416.1888681 | 1.65E-92 |
| LBH | 1.020606886 | 5.498290722 | 168.4183007 | 1.64E-38 |
| ING2 | 1.018735251 | 6.342493398 | 736.858277 | 2.89E-162 |
| MAP1A | 1.016631931 | 5.464066095 | 121.7557187 | 2.61E-28 |
| TTLL7 | 1.01565678 | 5.565533925 | 258.5455463 | 3.56E-58 |

| | | | | |
|---|---|---|---|---|
| TRIM61 | 1.014509079 | 5.423324991 | 162.2478077 | 3.65E-37 |
| FLRT2 | -1.013815522 | 5.94527066 | 336.4074355 | 3.87E-75 |
| HYAL3 | 1.01371773 | 5.541609326 | 277.4029697 | 2.76E-62 |
| PIP4K2C | 1.012196348 | 5.934145327 | 528.2200447 | 6.89E-117 |
| RILP | 1.010727997 | 5.568184204 | 291.4557866 | 2.40E-65 |
| DYRK1B | 1.008837403 | 5.528744463 | 256.8219415 | 8.46E-58 |
| HDAC9 | 1.00855798 | 5.443017287 | 210.0798771 | 1.32E-47 |
| PRKAA2 | 1.008265101 | 5.46034738 | 198.5924594 | 4.24E-45 |
| HSPA2 | 1.006556181 | 5.619887935 | 271.3767352 | 5.69E-61 |
| UBB | 1.005884666 | 9.068916753 | 1967.396332 | 0 |
| IL32 | 1.005040287 | 7.085503945 | 141.4682913 | 1.27E-32 |
| SNX16 | 1.004863059 | 5.596807698 | 312.7335043 | 5.54E-70 |
| PTHLH | -1.004094166 | 8.15967228 | 422.0375609 | 8.79E-94 |
| PLK1 | -1.003650529 | 6.24654715 | 99.05071692 | 2.46E-23 |
| OCLN | 1.002463219 | 5.625719436 | 227.4061937 | 2.19E-51 |
| GADD45A | -1.002182869 | 8.420566563 | 625.4848329 | 4.80E-138 |

| FDR | sig | gene |
|---|---|---|
| 3.27E-175 | sig_up | ANXA2R |
| 1.05E-207 | sig_up | FOXQ1 |
| 3.74E-199 | sig_up | PRSS22 |
| 0.00E+00 | sig_up | CLU |
| 2.51E-226 | sig_up | RBP7 |
| 3.71E-67 | sig_up | C15orf48 |
| 1.06E-227 | sig_up | TP53INP2 |
| 5.26E-133 | sig_up | WFDC2 |
| 5.00E-179 | sig_up | RNF223 |
| 3.03E-168 | sig_up | CLEC11A |
| 4.09E-151 | sig_up | TIMP2 |
| 2.65E-171 | sig_up | TMEM125 |
| 2.19E-295 | sig_up | HSPA1A |
| 1.21E-132 | sig_up | PCDH1 |
| 4.96E-102 | sig_up | UCHL1 |
| 7.57E-150 | sig_up | INA |
| 1.39E-134 | sig_up | SYNGR1 |
| 4.27E-172 | sig_up | OTUB2 |
| 3.85E-65 | sig_up | CLDN4 |
| 1.99E-120 | sig_up | ENO2 |
| 4.74E-33 | sig_up | LCN2 |
| 4.62E-208 | sig_down | FOS |
| 1.38E-136 | sig_up | RAB6B |
| 5.45E-250 | sig_down | AMIGO2 |
| 7.39E-121 | sig_up | BEX5 |
| 2.11E-14 | sig_up | SPRR2A |
| 4.82E-88 | sig_up | APOE |
| 9.77E-108 | sig_up | HMOX1 |
| 5.05E-122 | sig_up | TLCD4 |
| 3.25E-167 | sig_up | PDK2 |
| 3.51E-202 | sig_up | BEX2 |
| 2.73E-86 | sig_up | TNFAIP2 |
| 1.38E-233 | sig_down | KLF7 |
| 4.17E-47 | sig_down | TGM1 |
| 1.14E-122 | sig_up | TSPYL5 |
| 3.29E-83 | sig_up | CCK |
| 3.16E-217 | sig_down | MYC |
| 1.70E-113 | sig_up | PCSK1N |
| 2.14E-117 | sig_up | TP53I11 |
| 2.19E-153 | sig_up | RPP25 |
| 1.29E-57 | sig_up | NID1 |
| 1.36E-76 | sig_up | PLAAT3 |

| | | |
|---|---|---|
| 4.17E-234 | sig_up | FAM241B |
| 2.81E-104 | sig_up | CTSF |
| 1.42E-128 | sig_up | PTGES |
| 2.96E-115 | sig_down | INHBA |
| 8.28E-174 | sig_up | YPEL3 |
| 2.11E-86 | sig_down | GJB6 |
| 9.67E-113 | sig_up | GPRC5C |
| 5.95E-52 | sig_up | KLK11 |
| 9.49E-101 | sig_up | SNCG |
| 1.63E-93 | sig_up | FBXO32 |
| 3.45E-122 | sig_up | GCH1 |
| 1.38E-98 | sig_up | SYNM |
| 3.07E-83 | sig_up | METRNL |
| 1.33E-24 | sig_up | CXCL8 |
| 1.65E-170 | sig_up | MAP1LC3A |
| 1.93E-116 | sig_up | TMEM38A |
| 2.50E-87 | sig_up | NKAPL |
| 4.02E-73 | sig_up | RASGRP2 |
| 2.10E-60 | sig_up | RAET1L |
| 2.24E-145 | sig_up | GABARAPL1 |
| 3.20E-84 | sig_up | KLF2 |
| 2.41E-156 | sig_up | CCNE1 |
| 4.75E-94 | sig_up | RETREG1 |
| 7.92E-80 | sig_up | ADAP2 |
| 1.03E-103 | sig_down | EGR1 |
| 5.48E-94 | sig_up | ACP5 |
| 1.06E-159 | sig_up | DNAJB9 |
| 5.63E-143 | sig_down | F3 |
| 9.28E-263 | sig_down | TP63 |
| 3.36E-84 | sig_up | SIPA1L2 |
| 8.93E-93 | sig_up | GKAP1 |
| 3.63E-86 | sig_up | FAM43A |
| 3.41E-63 | sig_up | TNFRSF18 |
| 9.28E-88 | sig_up | RORA |
| 5.66E-104 | sig_up | KLHL24 |
| 5.60E-121 | sig_up | ABHD8 |
| 1.13E-96 | sig_up | PARD6A |
| 2.34E-81 | sig_up | INSR |
| 1.21E-43 | sig_up | ELF3 |
| 9.67E-109 | sig_up | SMPD1 |
| 4.46E-57 | sig_up | DDIT3 |
| 1.75E-60 | sig_up | KLK5 |
| 2.20E-79 | sig_up | MARCHF3 |

| | | |
|---|---|---|
| 4.18E-73 | sig_up | NIBAN1 |
| 2.73E-83 | sig_up | RASEF |
| 2.14E-87 | sig_up | DMTN |
| 2.62E-121 | sig_up | PLPPR2 |
| 7.85E-75 | sig_up | TMEM121 |
| 9.62E-51 | sig_down | G0S2 |
| 1.51E-72 | sig_up | DIPK1B |
| 9.76E-78 | sig_up | HES6 |
| 3.01E-33 | sig_up | CST6 |
| 4.02E-68 | sig_up | RND2 |
| 7.86E-87 | sig_up | CCDC69 |
| 4.81E-63 | sig_up | FLNC |
| 5.59E-90 | sig_up | GCHFR |
| 6.60E-108 | sig_up | SNX10 |
| 8.87E-64 | sig_up | TNNI3 |
| 2.40E-78 | sig_up | NRIP3 |
| 5.29E-93 | sig_up | CASTOR2 |
| 2.05E-94 | sig_up | MITF |
| 2.57E-121 | sig_up | ATOSB |
| 5.12E-75 | sig_up | TGM2 |
| 4.49E-73 | sig_up | FUT3 |
| 3.89E-70 | sig_up | BICDL2 |
| 1.99E-250 | sig_down | ZFP36L2 |
| 4.77E-63 | sig_down | IFITM1 |
| 1.66E-216 | sig_up | TP53I3 |
| 1.21E-80 | sig_up | RAB3A |
| 5.84E-101 | sig_down | NFKBIZ |
| 8.08E-118 | sig_up | WFS1 |
| 1.28E-179 | sig_up | CRB3 |
| 5.67E-105 | sig_down | ID1 |
| 8.73E-45 | sig_up | ICAM1 |
| 4.19E-44 | sig_up | CAMK2B |
| 7.81E-50 | sig_up | S100A4 |
| 1.11E-257 | sig_up | LRIF1 |
| 2.59E-97 | sig_down | FST |
| 1.12E-75 | sig_up | DBNDD1 |
| 8.06E-141 | sig_up | CXADR |
| 2.04E-219 | sig_down | HMGA2 |
| 7.51E-54 | sig_up | OLFM2 |
| 8.33E-78 | sig_up | MAGEH1 |
| 2.64E-74 | sig_up | FAM131C |
| 2.27E-152 | sig_down | SLC2A1 |
| 8.15E-44 | sig_down | FILIP1L |

| | | |
|---|---|---|
| 1.06E-147 | sig_up | PRADC1 |
| 3.70E-30 | sig_up | SLPI |
| 1.52E-87 | sig_down | IRF9 |
| 1.18E-203 | sig_down | DST |
| 1.42E-68 | sig_down | TXNIP |
| 3.48E-109 | sig_up | SELENOM |
| 1.40E-128 | sig_up | STX3 |
| 1.53E-80 | sig_up | DNAJA4 |
| 6.77E-80 | sig_up | PIERCE1 |
| 1.77E-159 | sig_down | NOTCH1 |
| 1.63E-61 | sig_up | MAP7D2 |
| 8.28E-66 | sig_up | CGN |
| 4.46E-79 | sig_up | RHPN2 |
| 3.64E-73 | sig_up | CLDN23 |
| 8.71E-71 | sig_up | DPF1 |
| 8.35E-68 | sig_up | TRPV3 |
| 5.66E-46 | sig_up | ANKRD22 |
| 8.41E-32 | sig_up | QPRT |
| 2.09E-80 | sig_up | RGS9 |
| 6.47E-106 | sig_up | RHOF |
| 2.41E-72 | sig_up | PRR15 |
| 1.89E-170 | sig_up | ITM2C |
| 3.97E-58 | sig_up | TTC9 |
| 2.90E-109 | sig_up | PLPP2 |
| 3.34E-56 | sig_down | LGALS9C |
| 3.32E-57 | sig_up | COBL |
| 1.56E-61 | sig_up | IL11 |
| 6.32E-73 | sig_up | LAYN |
| 3.51E-66 | sig_up | FICD |
| 7.42E-33 | sig_up | MT1G |
| 2.23E-161 | sig_up | PGF |
| 3.42E-65 | sig_up | CCL28 |
| 3.55E-262 | sig_up | HAGH |
| 1.68E-15 | sig_down | SERPINB3 |
| 3.67E-85 | sig_down | VSNL1 |
| 7.70E-26 | sig_up | CNFN |
| 2.71E-67 | sig_up | NAP1L5 |
| 2.04E-66 | sig_up | NATD1 |
| 2.91E-157 | sig_down | JAG1 |
| 2.90E-116 | sig_up | ZFAND2A |
| 1.49E-103 | sig_up | GSPT2 |
| 6.35E-161 | sig_up | SVIP |
| 2.84E-89 | sig_up | ATP6V0E2 |

| | | |
|---|---|---|
| 1.49E-44 | sig_up | CHAC1 |
| 7.33E-85 | sig_up | MAPRE3 |
| 9.99E-135 | sig_up | ANKRD9 |
| 5.55E-64 | sig_up | GULP1 |
| 2.27E-87 | sig_up | MAP1B |
| 7.15E-82 | sig_up | CCDC149 |
| 1.69E-78 | sig_up | CDA |
| 1.70E-73 | sig_up | ERO1B |
| 1.97E-298 | sig_up | SLC25A4 |
| 1.32E-62 | sig_up | AIF1L |
| 3.88E-114 | sig_up | ULBP2 |
| 3.96E-125 | sig_up | CPEB4 |
| 2.46E-105 | sig_up | KIF3C |
| 5.27E-115 | sig_up | MLLT11 |
| 7.16E-89 | sig_down | GJB2 |
| 9.86E-129 | sig_up | CHST12 |
| 6.40E-83 | sig_up | ISG20 |
| 6.33E-75 | sig_up | SLC43A2 |
| 6.59E-17 | sig_up | KRTAP2-3 |
| 1.03E-63 | sig_up | USP2 |
| 3.69E-138 | sig_up | NEU1 |
| 1.24E-62 | sig_up | SLC46A3 |
| 9.95E-267 | sig_up | TAF7 |
| 1.89E-54 | sig_up | FOXL2 |
| 2.04E-59 | sig_up | PDE4A |
| 1.29E-147 | sig_down | FGFBP1 |
| 3.14E-52 | sig_down | IFITM2 |
| 9.21E-90 | sig_down | COL8A1 |
| 5.93E-61 | sig_up | SLC25A42 |
| 5.76E-85 | sig_up | ATOSA |
| 2.12E-160 | sig_down | ETS2 |
| 5.96E-48 | sig_up | IL23A |
| 3.77E-209 | sig_up | SEPHS2 |
| 1.11E-156 | sig_down | IL1RAP |
| 4.76E-09 | sig_up | SPRR1B |
| 9.23E-65 | sig_up | C1orf115 |
| 1.35E-74 | sig_up | APH1B |
| 2.07E-45 | sig_down | FOSB |
| 5.12E-87 | sig_up | PGM2L1 |
| 2.95E-104 | sig_down | VEGFC |
| 1.60E-163 | sig_up | MAFG |
| 5.51E-49 | sig_down | LGALS9B |
| 1.57E-95 | sig_down | SIK1 |

| | | |
|---|---|---|
| 5.49E-76 | sig_up | LPIN2 |
| 2.23E-57 | sig_up | TMEM255B |
| 1.41E-140 | sig_up | PINK1 |
| 7.35E-80 | sig_up | TRAPPC14 |
| 3.77E-08 | sig_down | CXCL14 |
| 7.88E-109 | sig_down | COL7A1 |
| 1.23E-57 | sig_up | NEFH |
| 1.02E-61 | sig_up | SLC41A2 |
| 2.08E-32 | sig_down | DKK1 |
| 3.05E-98 | sig_down | IRS1 |
| 8.88E-70 | sig_up | NOL4L |
| 5.23E-82 | sig_up | SLC17A5 |
| 4.51E-60 | sig_up | A1BG |
| 1.15E-57 | sig_up | HES4 |
| 2.05E-54 | sig_up | EPHX4 |
| 2.05E-59 | sig_up | ZNF280B |
| 1.93E-62 | sig_up | TSPAN33 |
| 1.18E-78 | sig_up | POMGNT2 |
| 1.10E-55 | sig_up | ZNF483 |
| 2.09E-32 | sig_down | CCN2 |
| 1.06E-108 | sig_down | THBS1 |
| 1.75E-175 | sig_down | ZFP36L1 |
| 1.44E-83 | sig_up | HSPA1B |
| 1.57E-59 | sig_up | TFAP2C |
| 4.44E-130 | sig_up | TPRG1L |
| 4.64E-71 | sig_down | IL20RB |
| 2.27E-69 | sig_up | KIF13B |
| 2.21E-68 | sig_up | PKN1 |
| 1.07E-10 | sig_up | SAA2 |
| 1.41E-72 | sig_up | MFSD9 |
| 8.06E-72 | sig_up | B3GNT3 |
| 2.02E-33 | sig_down | CDCA3 |
| 4.45E-45 | sig_up | GARIN5A |
| 2.55E-206 | sig_down | SERPINB5 |
| 1.62E-51 | sig_down | DLK2 |
| 4.90E-100 | sig_down | PLK2 |
| 4.63E-17 | sig_up | KLK10 |
| 6.77E-76 | sig_up | NCOA1 |
| 6.57E-50 | sig_down | GABRE |
| 1.72E-59 | sig_up | ZFYVE28 |
| 2.04E-83 | sig_up | PPP2R5B |
| 6.56E-38 | sig_up | CCNA1 |
| 6.42E-54 | sig_down | MMP28 |

| | | |
|---|---|---|
| 6.73E-64 | sig_up | MRAS |
| 1.16E-63 | sig_up | ULK1 |
| 6.68E-94 | sig_up | SASH1 |
| 1.58E-125 | sig_down | SNAI2 |
| 9.01E-57 | sig_up | SLC26A11 |
| 8.09E-48 | sig_up | TSPAN2 |
| 2.57E-77 | sig_up | MT1F |
| 1.11E-49 | sig_up | DUSP8 |
| 8.27E-78 | sig_up | CDKN2D |
| 2.70E-30 | sig_down | KIF20A |
| 8.00E-23 | sig_up | HERC5 |
| 1.14E-15 | sig_down | IFI44L |
| 1.25E-54 | sig_up | MFAP3L |
| 2.63E-49 | sig_up | RNF208 |
| 4.72E-53 | sig_up | E2F1 |
| 1.15E-106 | sig_down | MT1X |
| 5.46E-51 | sig_up | FAM110C |
| 2.60E-72 | sig_up | NIPA1 |
| 3.07E-47 | sig_up | SDR16C5 |
| 5.33E-97 | sig_up | SECISBP2L |
| 3.35E-115 | sig_up | BEX4 |
| 5.46E-48 | sig_down | PCDH7 |
| 2.89E-46 | sig_up | CEBPA |
| 6.54E-51 | sig_up | ATL1 |
| 2.03E-52 | sig_up | ADGRF4 |
| 8.00E-40 | sig_up | FBLN2 |
| 6.71E-143 | sig_down | CEBPD |
| 4.21E-110 | sig_up | MAP1S |
| 1.45E-13 | sig_up | S100A9 |
| 5.62E-28 | sig_up | CYSRT1 |
| 1.67E-22 | sig_down | CXCL2 |
| 6.21E-55 | sig_up | SERTAD4 |
| 2.34E-57 | sig_up | ARHGAP42 |
| 5.01E-97 | sig_down | PHLDA1 |
| 2.86E-90 | sig_up | GAL |
| 1.91E-111 | sig_up | MAP3K11 |
| 1.67E-62 | sig_up | XKR8 |
| 6.53E-78 | sig_up | ABHD3 |
| 5.48E-91 | sig_down | DDIT4 |
| 1.09E-37 | sig_up | LBH |
| 4.39E-160 | sig_up | ING2 |
| 1.21E-27 | sig_up | MAP1A |
| 4.50E-57 | sig_up | TTLL7 |

| | | |
|---|---|---|
| 2.31E-36 | sig_up | TRIM61 |
| 7.98E-74 | sig_down | FLRT2 |
| 3.96E-61 | sig_up | HYAL3 |
| 4.15E-115 | sig_up | PIP4K2C |
| 3.78E-64 | sig_up | RILP |
| 1.06E-56 | sig_up | DYRK1B |
| 1.19E-46 | sig_up | HDAC9 |
| 3.51E-44 | sig_up | PRKAA2 |
| 7.81E-60 | sig_up | HSPA2 |
| 0 | sig_up | UBB |
| 6.91E-32 | sig_up | IL32 |
| 1.00E-68 | sig_up | SNX16 |
| 2.98E-92 | sig_down | PTHLH |
| 9.49E-23 | sig_down | PLK1 |
| 2.24E-50 | sig_up | OCLN |
| 4.39E-136 | sig_down | GADD45A |
