## Supplemental Table 4 for "Human Cytomegalovirus Infection of Primary Human Oral Keratinocytes Induces Intermediate Keratinocyte Differentiation and an Altered Innate Immune Response"

| Gene name | R | P | Correlation Strength |
|---|---|---|---|
| UBE2S | 0.564104 | 1.57E-28 | Strong Positive Correlation |
| SDF2L1 | 0.542983 | 3.64E-26 | Strong Positive Correlation |
| RBP7 | 0.541374 | 5.43E-26 | Strong Positive Correlation |
| TUBB2B | 0.483092 | 2.72E-20 | Moderate Positive Correlation |
| UCHL1 | 0.481833 | 3.52E-20 | Moderate Positive Correlation |
| HAGH | 0.481479 | 3.79E-20 | Moderate Positive Correlation |
| UBB | 0.475036 | 1.38E-19 | Moderate Positive Correlation |
| ENO2 | 0.472712 | 2.19E-19 | Moderate Positive Correlation |
| ATP6V0B | 0.472669 | 2.21E-19 | Moderate Positive Correlation |
| HES6 | 0.471303 | 2.89E-19 | Moderate Positive Correlation |
| SELENOK | 0.470639 | 3.3E-19 | Moderate Positive Correlation |
| EEF1A2 | 0.468116 | 5.4E-19 | Moderate Positive Correlation |
| RPP25 | 0.457283 | 4.28E-18 | Moderate Positive Correlation |
| TIMP2 | 0.452656 | 1.01E-17 | Moderate Positive Correlation |
| ZFR2 | 0.445812 | 3.54E-17 | Moderate Positive Correlation |
| CKB | 0.443419 | 5.44E-17 | Moderate Positive Correlation |
| RNF157 | 0.438222 | 1.37E-16 | Moderate Positive Correlation |
| BEX2 | 0.437428 | 1.58E-16 | Moderate Positive Correlation |
| PCDH1 | 0.433278 | 3.26E-16 | Moderate Positive Correlation |
| PLAAT3 | 0.428295 | 7.68E-16 | Moderate Positive Correlation |
| TCF15 | 0.425321 | 1.27E-15 | Moderate Positive Correlation |
| DDIT3 | 0.424038 | 1.58E-15 | Moderate Positive Correlation |
| HLA-B | 0.422048 | 2.2E-15 | Moderate Positive Correlation |
| PGP | 0.420031 | 3.09E-15 | Moderate Positive Correlation |
| WFDC2 | 0.41773 | 4.51E-15 | Moderate Positive Correlation |
| MLEC | 0.417282 | 4.86E-15 | Moderate Positive Correlation |
| CTSF | 0.410221 | 1.53E-14 | Moderate Positive Correlation |
| PRADC1 | 0.405889 | 3.06E-14 | Moderate Positive Correlation |
| UBE2T | 0.405705 | 3.15E-14 | Moderate Positive Correlation |
| RHEBL1 | 0.405014 | 3.51E-14 | Moderate Positive Correlation |
| EIF1B | 0.404446 | 3.84E-14 | Moderate Positive Correlation |
| PRSS22 | 0.403199 | 4.67E-14 | Moderate Positive Correlation |
| GABARAPL1 | 0.402156 | 5.5E-14 | Moderate Positive Correlation |
| KIF1A | 0.399738 | 8.02E-14 | Moderate Positive Correlation |
| NIBAN1 | 0.399035 | 8.94E-14 | Moderate Positive Correlation |
| MAPK8IP2 | 0.394509 | 1.79E-13 | Moderate Positive Correlation |
| E2F1 | 0.391338 | 2.9E-13 | Moderate Positive Correlation |
| NRIP3 | 0.391209 | 2.95E-13 | Moderate Positive Correlation |
| APOE | 0.389073 | 4.07E-13 | Moderate Positive Correlation |
| ZWINT | 0.388408 | 4.5E-13 | Moderate Positive Correlation |
| YBX2 | 0.384371 | 8.18E-13 | Moderate Positive Correlation |
| ATP6V1E1 | 0.382635 | 1.06E-12 | Moderate Positive Correlation |
| LMO4 | 0.379932 | 1.57E-12 | Moderate Positive Correlation |
| TMEM208 | 0.378578 | 1.9E-12 | Moderate Positive Correlation |
| CFL2 | 0.378453 | 1.94E-12 | Moderate Positive Correlation |
| ITM2C | 0.377319 | 2.28E-12 | Moderate Positive Correlation |
| TNFRSF18 | 0.375393 | 3.01E-12 | Moderate Positive Correlation |

| | | | |
|---|---|---|---|
| PDK2 | 0.375236 | 3.08E-12 | Moderate Positive Correlation |
| CLDN7 | 0.37509 | 3.14E-12 | Moderate Positive Correlation |
| DNAJB9 | 0.373583 | 3.89E-12 | Moderate Positive Correlation |
| METRNL | 0.372708 | 4.41E-12 | Moderate Positive Correlation |
| HERC5 | 0.369741 | 6.69E-12 | Moderate Positive Correlation |
| TLCD4 | 0.366577 | 1.04E-11 | Moderate Positive Correlation |
| MMP23B | 0.365877 | 1.15E-11 | Moderate Positive Correlation |
| TK1 | 0.365817 | 1.16E-11 | Moderate Positive Correlation |
| PRRX2 | 0.364851 | 1.32E-11 | Moderate Positive Correlation |
| ZFAND2A | 0.364675 | 1.35E-11 | Moderate Positive Correlation |
| ATAD2 | 0.364133 | 1.46E-11 | Moderate Positive Correlation |
| DNAJC3 | 0.363299 | 1.63E-11 | Moderate Positive Correlation |
| RASGRP2 | 0.36287 | 1.73E-11 | Moderate Positive Correlation |
| MED10 | 0.361456 | 2.1E-11 | Moderate Positive Correlation |
| PLPP2 | 0.361206 | 2.17E-11 | Moderate Positive Correlation |
| CHAC1 | 0.359546 | 2.72E-11 | Moderate Positive Correlation |
| RORA | 0.359532 | 2.73E-11 | Moderate Positive Correlation |
| PRSS8 | 0.358067 | 3.32E-11 | Moderate Positive Correlation |
| ATG101 | 0.357499 | 3.59E-11 | Moderate Positive Correlation |
| CD55 | 0.357419 | 3.63E-11 | Moderate Positive Correlation |
| FEN1 | 0.353447 | 6.16E-11 | Moderate Positive Correlation |
| CTSA | 0.351504 | 7.95E-11 | Moderate Positive Correlation |
| NEU1 | 0.34905 | 1.1E-10 | Moderate Positive Correlation |
| BSPRY | 0.348737 | 1.14E-10 | Moderate Positive Correlation |
| C1QL1 | 0.348642 | 1.16E-10 | Moderate Positive Correlation |
| SLC25A4 | 0.347783 | 1.29E-10 | Moderate Positive Correlation |
| CDC6 | 0.346853 | 1.46E-10 | Moderate Positive Correlation |
| GARIN5A | 0.34619 | 1.59E-10 | Moderate Positive Correlation |
| ASRGL1 | 0.345068 | 1.84E-10 | Moderate Positive Correlation |
| RRAGD | 0.345045 | 1.84E-10 | Moderate Positive Correlation |
| C1orf109 | 0.341828 | 2.78E-10 | Moderate Positive Correlation |
| SELENOM | 0.33754 | 4.77E-10 | Moderate Positive Correlation |
| CEBPB | 0.337378 | 4.87E-10 | Moderate Positive Correlation |
| NPTX2 | 0.335803 | 5.92E-10 | Moderate Positive Correlation |
| MAP1LC3B | 0.335266 | 6.33E-10 | Moderate Positive Correlation |
| DNASE2 | 0.335166 | 6.41E-10 | Moderate Positive Correlation |
| MAP1LC3A | 0.333863 | 7.53E-10 | Moderate Positive Correlation |
| SERTAD4 | 0.33257 | 8.83E-10 | Moderate Positive Correlation |
| MYH14 | 0.331493 | 1.01E-09 | Moderate Positive Correlation |
| MESP1 | 0.331161 | 1.05E-09 | Moderate Positive Correlation |
| HID1 | 0.329928 | 1.22E-09 | Moderate Positive Correlation |
| MAP1B | 0.329549 | 1.28E-09 | Moderate Positive Correlation |
| BARX1 | 0.329394 | 1.3E-09 | Moderate Positive Correlation |
| CRLF1 | 0.329378 | 1.3E-09 | Moderate Positive Correlation |
| SEC11C | 0.328855 | 1.39E-09 | Moderate Positive Correlation |
| RNF223 | 0.326895 | 1.76E-09 | Moderate Positive Correlation |
| TCF19 | 0.326374 | 1.88E-09 | Moderate Positive Correlation |
| NMRK2 | 0.326242 | 1.91E-09 | Moderate Positive Correlation |

| | | | |
|---|---|---|---|
| GMPR | 0.325043 | 2.2E-09 | Moderate Positive Correlation |
| BEX5 | 0.324428 | 2.37E-09 | Moderate Positive Correlation |
| TP53I11 | 0.323817 | 2.55E-09 | Moderate Positive Correlation |
| STX3 | 0.323196 | 2.74E-09 | Moderate Positive Correlation |
| MORC4 | 0.322754 | 2.89E-09 | Moderate Positive Correlation |
| HLA-C | 0.321887 | 3.2E-09 | Moderate Positive Correlation |
| VLDLR | 0.321536 | 3.34E-09 | Moderate Positive Correlation |
| BARD1 | 0.321263 | 3.45E-09 | Moderate Positive Correlation |
| SYNGR1 | 0.320967 | 3.57E-09 | Moderate Positive Correlation |
| GAL | 0.320648 | 3.71E-09 | Moderate Positive Correlation |
| HMX1 | 0.319349 | 4.32E-09 | Moderate Positive Correlation |
| C1QTNF4 | 0.318913 | 4.54E-09 | Moderate Positive Correlation |
| MESD | 0.318644 | 4.69E-09 | Moderate Positive Correlation |
| INHBB | 0.317555 | 5.32E-09 | Moderate Positive Correlation |
| YPEL3 | 0.316155 | 6.26E-09 | Moderate Positive Correlation |
| ORC6 | 0.316102 | 6.3E-09 | Moderate Positive Correlation |
| GCHFR | 0.315296 | 6.91E-09 | Moderate Positive Correlation |
| AIF1L | 0.314867 | 7.26E-09 | Moderate Positive Correlation |
| RPA2 | 0.313009 | 8.99E-09 | Moderate Positive Correlation |
| CDK2AP2 | 0.311677 | 1.05E-08 | Moderate Positive Correlation |
| NXPH4 | 0.3102 | 1.24E-08 | Moderate Positive Correlation |
| CLEC11A | 0.310069 | 1.26E-08 | Moderate Positive Correlation |
| RIC3 | 0.310024 | 1.26E-08 | Moderate Positive Correlation |
| POP7 | 0.309648 | 1.32E-08 | Moderate Positive Correlation |
| CCK | 0.30916 | 1.39E-08 | Moderate Positive Correlation |
| FICD | 0.30867 | 1.47E-08 | Moderate Positive Correlation |
| METTL4 | 0.308261 | 1.54E-08 | Moderate Positive Correlation |
| GAS2L3 | 0.308068 | 1.57E-08 | Moderate Positive Correlation |
| RHEB | 0.307868 | 1.61E-08 | Moderate Positive Correlation |
| MAFG | 0.306786 | 1.82E-08 | Moderate Positive Correlation |
| RAMP1 | 0.306685 | 1.84E-08 | Moderate Positive Correlation |
| ERBB3 | 0.306446 | 1.89E-08 | Moderate Positive Correlation |
| C1QL4 | 0.306311 | 1.92E-08 | Moderate Positive Correlation |
| FLOT1 | 0.306175 | 1.95E-08 | Moderate Positive Correlation |
| GLRX5 | 0.305447 | 2.11E-08 | Moderate Positive Correlation |
| INA | 0.304674 | 2.3E-08 | Moderate Positive Correlation |
| FAM241B | 0.304248 | 2.41E-08 | Moderate Positive Correlation |
| HES4 | 0.304161 | 2.43E-08 | Moderate Positive Correlation |
| PCSK1N | 0.303498 | 2.62E-08 | Moderate Positive Correlation |
| BEX4 | 0.303416 | 2.64E-08 | Moderate Positive Correlation |
| RND2 | 0.30327 | 2.69E-08 | Moderate Positive Correlation |
| EFHD1 | 0.302304 | 2.99E-08 | Moderate Positive Correlation |
| AURKA | 0.301146 | 3.39E-08 | Moderate Positive Correlation |
| APLP1 | 0.301078 | 3.42E-08 | Moderate Positive Correlation |
| HSPA13 | 0.300651 | 3.58E-08 | Moderate Positive Correlation |
| FCGRT | 0.299367 | 4.12E-08 | Other Positive Correlation |
| PKN1 | 0.299056 | 4.26E-08 | Other Positive Correlation |
| ERO1B | 0.298951 | 4.31E-08 | Other Positive Correlation |

| | | | |
|---|---|---|---|
| SHISA8 | 0.298419 | 4.56E-08 | Other Positive Correlation |
| CBX4 | 0.297732 | 4.92E-08 | Other Positive Correlation |
| TAC1 | 0.297177 | 5.22E-08 | Other Positive Correlation |
| TNRC6C | 0.295834 | 6.03E-08 | Other Positive Correlation |
| MTFR2 | 0.295323 | 6.37E-08 | Other Positive Correlation |
| C19orf25 | 0.295085 | 6.53E-08 | Other Positive Correlation |
| GADD45G | 0.294818 | 6.72E-08 | Other Positive Correlation |
| GKAP1 | 0.294521 | 6.94E-08 | Other Positive Correlation |
| TLE2 | 0.294293 | 7.11E-08 | Other Positive Correlation |
| MYBL1 | 0.294112 | 7.25E-08 | Other Positive Correlation |
| NFKBIB | 0.294074 | 7.28E-08 | Other Positive Correlation |
| FAM131C | 0.293936 | 7.39E-08 | Other Positive Correlation |
| DOC2B | 0.291816 | 9.25E-08 | Other Positive Correlation |
| TESC | 0.291796 | 9.27E-08 | Other Positive Correlation |
| ADM | 0.290332 | 1.08E-07 | Other Positive Correlation |
| RAB6B | 0.289999 | 1.12E-07 | Other Positive Correlation |
| PMAIP1 | 0.288554 | 1.3E-07 | Other Positive Correlation |
| RAB3IL1 | 0.286977 | 1.53E-07 | Other Positive Correlation |
| RELL2 | 0.286786 | 1.56E-07 | Other Positive Correlation |
| PPP1R1A | 0.286709 | 1.58E-07 | Other Positive Correlation |
| DIRAS1 | 0.286471 | 1.62E-07 | Other Positive Correlation |
| TCEAL2 | 0.286325 | 1.64E-07 | Other Positive Correlation |
| KLHL24 | 0.285772 | 1.74E-07 | Other Positive Correlation |
| RAD51 | 0.285023 | 1.88E-07 | Other Positive Correlation |
| SELENOS | 0.284936 | 1.89E-07 | Other Positive Correlation |
| SDHAF1 | 0.284694 | 1.94E-07 | Other Positive Correlation |
| BMP4 | 0.284109 | 2.06E-07 | Other Positive Correlation |
| ABHD3 | 0.283169 | 2.27E-07 | Other Positive Correlation |
| ERICH2 | 0.283139 | 2.28E-07 | Other Positive Correlation |
| MEIOC | 0.283104 | 2.28E-07 | Other Positive Correlation |
| SLC25A33 | 0.282299 | 2.48E-07 | Other Positive Correlation |
| ING2 | 0.28201 | 2.55E-07 | Other Positive Correlation |
| PDRG1 | 0.281562 | 2.67E-07 | Other Positive Correlation |
| DES | 0.280714 | 2.91E-07 | Other Positive Correlation |
| RHOQ | 0.280434 | 3E-07 | Other Positive Correlation |
| MACO1 | 0.280391 | 3.01E-07 | Other Positive Correlation |
| MCM5 | 0.280352 | 3.02E-07 | Other Positive Correlation |
| LYPD6B | 0.28024 | 3.06E-07 | Other Positive Correlation |
| TIMM10 | 0.279911 | 3.16E-07 | Other Positive Correlation |
| CYB561 | 0.279768 | 3.21E-07 | Other Positive Correlation |
| KLF15 | 0.279623 | 3.25E-07 | Other Positive Correlation |
| RNF212 | 0.279155 | 3.41E-07 | Other Positive Correlation |
| NEFH | 0.278762 | 3.55E-07 | Other Positive Correlation |
| GCLM | 0.278591 | 3.61E-07 | Other Positive Correlation |
| TMEM59L | 0.278393 | 3.68E-07 | Other Positive Correlation |
| PHF1 | 0.278269 | 3.73E-07 | Other Positive Correlation |
| ANKRD9 | 0.278031 | 3.82E-07 | Other Positive Correlation |
| FAM89A | 0.277687 | 3.95E-07 | Other Positive Correlation |

| | | | |
|---|---|---|---|
| ZNF85 | 0.27765 | 3.96E-07 | Other Positive Correlation |
| ARL6IP1 | 0.277442 | 4.05E-07 | Other Positive Correlation |
| SCN1B | 0.276666 | 4.37E-07 | Other Positive Correlation |
| STC2 | 0.276384 | 4.5E-07 | Other Positive Correlation |
| CALCB | 0.275474 | 4.92E-07 | Other Positive Correlation |
| ENPP5 | 0.27513 | 5.09E-07 | Other Positive Correlation |
| SCN5A | 0.275085 | 5.12E-07 | Other Positive Correlation |
| CYP27A1 | 0.27403 | 5.68E-07 | Other Positive Correlation |
| FSD1 | 0.272991 | 6.29E-07 | Other Positive Correlation |
| ENHO | 0.27297 | 6.3E-07 | Other Positive Correlation |
| COX19 | 0.272951 | 6.31E-07 | Other Positive Correlation |
| HS3ST6 | 0.272846 | 6.38E-07 | Other Positive Correlation |
| FBLL1 | 0.27282 | 6.39E-07 | Other Positive Correlation |
| TMEM106C | 0.272718 | 6.46E-07 | Other Positive Correlation |
| GATA2 | 0.272641 | 6.51E-07 | Other Positive Correlation |
| HSPH1 | 0.272457 | 6.62E-07 | Other Positive Correlation |
| RGS9 | 0.271117 | 7.55E-07 | Other Positive Correlation |
| PREB | 0.270999 | 7.64E-07 | Other Positive Correlation |
| CDKN2D | 0.27086 | 7.74E-07 | Other Positive Correlation |
| COL6A2 | 0.270654 | 7.9E-07 | Other Positive Correlation |
| STRBP | 0.269959 | 8.45E-07 | Other Positive Correlation |
| ADM2 | 0.269568 | 8.78E-07 | Other Positive Correlation |
| SEL1L | 0.269173 | 9.12E-07 | Other Positive Correlation |
| SVIP | 0.268434 | 9.79E-07 | Other Positive Correlation |
| MLF1 | 0.268207 | 1E-06 | Other Positive Correlation |
| RTN4RL2 | 0.268182 | 1E-06 | Other Positive Correlation |
| SMCO4 | 0.267248 | 1.1E-06 | Other Positive Correlation |
| SLC10A4 | 0.266753 | 1.15E-06 | Other Positive Correlation |
| MFGE8 | 0.266035 | 1.23E-06 | Other Positive Correlation |
| RNF26 | 0.2658 | 1.26E-06 | Other Positive Correlation |
| EVA1B | 0.265016 | 1.36E-06 | Other Positive Correlation |
| PRKAA2 | 0.264901 | 1.37E-06 | Other Positive Correlation |
| DSCC1 | 0.26431 | 1.45E-06 | Other Positive Correlation |
| SAT2 | 0.263897 | 1.51E-06 | Other Positive Correlation |
| B4GALNT1 | 0.263692 | 1.54E-06 | Other Positive Correlation |
| POLA2 | 0.263625 | 1.55E-06 | Other Positive Correlation |
| ESCO2 | 0.263595 | 1.55E-06 | Other Positive Correlation |
| WFS1 | 0.263553 | 1.56E-06 | Other Positive Correlation |
| OVOL2 | 0.263044 | 1.64E-06 | Other Positive Correlation |
| SAR1B | 0.262894 | 1.66E-06 | Other Positive Correlation |
| RNF175 | 0.262893 | 1.66E-06 | Other Positive Correlation |
| CAMK2N2 | 0.262606 | 1.71E-06 | Other Positive Correlation |
| MIEF1 | 0.262099 | 1.79E-06 | Other Positive Correlation |
| NRTN | 0.262034 | 1.8E-06 | Other Positive Correlation |
| GMPPB | 0.261932 | 1.82E-06 | Other Positive Correlation |
| ASF1B | 0.261524 | 1.89E-06 | Other Positive Correlation |
| SNAP25 | 0.261394 | 1.91E-06 | Other Positive Correlation |
| PALM | 0.261198 | 1.95E-06 | Other Positive Correlation |

| | | | |
|---|---|---|---|
| TXNRD1 | 0.260485 | 2.08E-06 | Other Positive Correlation |
| ETFDH | 0.260118 | 2.15E-06 | Other Positive Correlation |
| TOX2 | 0.259503 | 2.28E-06 | Other Positive Correlation |
| RASSF1 | 0.259498 | 2.28E-06 | Other Positive Correlation |
| KCNK12 | 0.25922 | 2.34E-06 | Other Positive Correlation |
| SDC2 | 0.258764 | 2.44E-06 | Other Positive Correlation |
| NES | 0.258181 | 2.58E-06 | Other Positive Correlation |
| TUG1 | 0.25813 | 2.59E-06 | Other Positive Correlation |
| TTC9 | 0.258118 | 2.59E-06 | Other Positive Correlation |
| PDE4A | 0.257835 | 2.66E-06 | Other Positive Correlation |
| PNPLA2 | 0.257647 | 2.71E-06 | Other Positive Correlation |
| BRCA1 | 0.257616 | 2.71E-06 | Other Positive Correlation |
| INSR | 0.257246 | 2.81E-06 | Other Positive Correlation |
| GFPT1 | 0.257122 | 2.84E-06 | Other Positive Correlation |
| ZNF483 | 0.256819 | 2.92E-06 | Other Positive Correlation |
| SEMA6B | 0.256107 | 3.12E-06 | Other Positive Correlation |
| MANF | 0.256067 | 3.13E-06 | Other Positive Correlation |
| ELF3 | 0.255953 | 3.16E-06 | Other Positive Correlation |
| KIAA2013 | 0.255743 | 3.22E-06 | Other Positive Correlation |
| TIMM44 | 0.255508 | 3.29E-06 | Other Positive Correlation |
| STK40 | 0.255499 | 3.3E-06 | Other Positive Correlation |
| C9orf40 | 0.255292 | 3.36E-06 | Other Positive Correlation |
| GABARAPL2 | 0.255269 | 3.37E-06 | Other Positive Correlation |
| KLK1 | 0.255044 | 3.43E-06 | Other Positive Correlation |
| C1QL2 | 0.254423 | 3.63E-06 | Other Positive Correlation |
| SKAP1 | 0.254406 | 3.64E-06 | Other Positive Correlation |
| TERF2IP | 0.254233 | 3.7E-06 | Other Positive Correlation |
| POLR3H | 0.254071 | 3.75E-06 | Other Positive Correlation |
| KCNK15 | 0.253711 | 3.88E-06 | Other Positive Correlation |
| HRK | 0.253669 | 3.89E-06 | Other Positive Correlation |
| KCNK1 | 0.253599 | 3.92E-06 | Other Positive Correlation |
| IGF2 | 0.253273 | 4.03E-06 | Other Positive Correlation |
| KCNH2 | 0.253115 | 4.09E-06 | Other Positive Correlation |
| INHBE | 0.253005 | 4.13E-06 | Other Positive Correlation |
| ATF3 | 0.253004 | 4.13E-06 | Other Positive Correlation |
| CLPTM1 | 0.252824 | 4.2E-06 | Other Positive Correlation |
| SMC3 | 0.252805 | 4.21E-06 | Other Positive Correlation |
| ZBTB18 | 0.25261 | 4.28E-06 | Other Positive Correlation |
| TMEM151A | 0.252533 | 4.31E-06 | Other Positive Correlation |
| FAM184A | 0.252272 | 4.42E-06 | Other Positive Correlation |
| SASS6 | 0.252015 | 4.52E-06 | Other Positive Correlation |
| MCOLN1 | 0.251972 | 4.54E-06 | Other Positive Correlation |
| CXXC5 | 0.251932 | 4.55E-06 | Other Positive Correlation |
| GPRC5C | 0.251884 | 4.57E-06 | Other Positive Correlation |
| ITGA5 | 0.251029 | 4.94E-06 | Other Positive Correlation |
| TAF5 | 0.250595 | 5.13E-06 | Other Positive Correlation |
| DMTN | 0.250591 | 5.14E-06 | Other Positive Correlation |
| MGAT4A | 0.250459 | 5.2E-06 | Other Positive Correlation |

| | | | |
|---|---|---|---|
| AKTIP | 0.250316 | 5.26E-06 | Other Positive Correlation |
| RAB39B | 0.250002 | 5.41E-06 | Other Positive Correlation |
| ZNF280B | 0.249641 | 5.59E-06 | Other Positive Correlation |
| NUP210 | 0.249617 | 5.6E-06 | Other Positive Correlation |
| RAB17 | 0.249593 | 5.61E-06 | Other Positive Correlation |
| PCK2 | 0.249031 | 5.9E-06 | Other Positive Correlation |
| EMC6 | 0.248901 | 5.97E-06 | Other Positive Correlation |
| NFIL3 | 0.248431 | 6.23E-06 | Other Positive Correlation |
| FAHD1 | 0.248408 | 6.24E-06 | Other Positive Correlation |
| MICA | 0.247914 | 6.52E-06 | Other Positive Correlation |
| CLDN6 | 0.247524 | 6.75E-06 | Other Positive Correlation |
| STAC2 | 0.247406 | 6.82E-06 | Other Positive Correlation |
| MNX1 | 0.247272 | 6.9E-06 | Other Positive Correlation |
| CADM4 | 0.247269 | 6.9E-06 | Other Positive Correlation |
| SHE | 0.247194 | 6.95E-06 | Other Positive Correlation |
| TESK1 | 0.247019 | 7.05E-06 | Other Positive Correlation |
| TLX3 | 0.246777 | 7.21E-06 | Other Positive Correlation |
| CRB3 | 0.246485 | 7.39E-06 | Other Positive Correlation |
| RIMKLA | 0.246274 | 7.53E-06 | Other Positive Correlation |
| MAFA | 0.24589 | 7.79E-06 | Other Positive Correlation |
| UBE2O | 0.245848 | 7.82E-06 | Other Positive Correlation |
| NDRG2 | 0.245432 | 8.11E-06 | Other Positive Correlation |
| KIF22 | 0.245112 | 8.34E-06 | Other Positive Correlation |
| PPP1R9A | 0.244773 | 8.59E-06 | Other Positive Correlation |
| RYBP | 0.244696 | 8.65E-06 | Other Positive Correlation |
| TMEM121 | 0.244293 | 8.96E-06 | Other Positive Correlation |
| ZAR1 | 0.244202 | 9.03E-06 | Other Positive Correlation |
| CAMK1D | 0.244028 | 9.17E-06 | Other Positive Correlation |
| GNS | 0.243896 | 9.27E-06 | Other Positive Correlation |
| HAX1 | 0.243833 | 9.32E-06 | Other Positive Correlation |
| CISD2 | 0.243832 | 9.32E-06 | Other Positive Correlation |
| NKAIN4 | 0.243712 | 9.42E-06 | Other Positive Correlation |
| LTBP3 | 0.243578 | 9.53E-06 | Other Positive Correlation |
| FGF21 | 0.243411 | 9.67E-06 | Other Positive Correlation |
| SATL1 | 0.243022 | 1E-05 | Other Positive Correlation |
| SIGIRR | 0.242848 | 1.02E-05 | Other Positive Correlation |
| SCAMP5 | 0.242494 | 1.05E-05 | Other Positive Correlation |
| RPA1 | 0.242353 | 1.06E-05 | Other Positive Correlation |
| COL6A1 | 0.242236 | 1.07E-05 | Other Positive Correlation |
| NRGN | 0.242076 | 1.09E-05 | Other Positive Correlation |
| PALM3 | 0.241365 | 1.15E-05 | Other Positive Correlation |
| NPTXR | 0.240776 | 1.21E-05 | Other Positive Correlation |
| USP1 | 0.240212 | 1.27E-05 | Other Positive Correlation |
| KAZALD1 | 0.240073 | 1.29E-05 | Other Positive Correlation |
| ART5 | 0.239746 | 1.33E-05 | Other Positive Correlation |
| SAP30 | 0.239744 | 1.33E-05 | Other Positive Correlation |
| SLC39A7 | 0.23937 | 1.37E-05 | Other Positive Correlation |
| MTHFD2 | 0.238944 | 1.42E-05 | Other Positive Correlation |

| | | | |
|---|---|---|---|
| DEGS2 | 0.238895 | 1.43E-05 | Other Positive Correlation |
| RECQL4 | 0.238778 | 1.44E-05 | Other Positive Correlation |
| RCAN2 | 0.238724 | 1.45E-05 | Other Positive Correlation |
| TMEM259 | 0.238674 | 1.45E-05 | Other Positive Correlation |
| APBB1 | 0.23847 | 1.48E-05 | Other Positive Correlation |
| MYDGF | 0.238463 | 1.48E-05 | Other Positive Correlation |
| BHLHE41 | 0.238202 | 1.51E-05 | Other Positive Correlation |
| ARHGAP44 | 0.237276 | 1.64E-05 | Other Positive Correlation |
| SLC16A11 | 0.237158 | 1.65E-05 | Other Positive Correlation |
| GPAT3 | 0.236975 | 1.68E-05 | Other Positive Correlation |
| EPCAM | 0.236973 | 1.68E-05 | Other Positive Correlation |
| ONECUT2 | 0.23663 | 1.73E-05 | Other Positive Correlation |
| CDT1 | 0.2364 | 1.76E-05 | Other Positive Correlation |
| SMIM13 | 0.23611 | 1.8E-05 | Other Positive Correlation |
| ATP6V0E2 | 0.236065 | 1.81E-05 | Other Positive Correlation |
| SOCS7 | 0.23587 | 1.84E-05 | Other Positive Correlation |
| RGL3 | 0.235849 | 1.85E-05 | Other Positive Correlation |
| DPP7 | 0.235727 | 1.86E-05 | Other Positive Correlation |
| DOHH | 0.235014 | 1.98E-05 | Other Positive Correlation |
| NTNG1 | 0.234807 | 2.01E-05 | Other Positive Correlation |
| DTNBP1 | 0.234242 | 2.11E-05 | Other Positive Correlation |
| IFT27 | 0.234093 | 2.14E-05 | Other Positive Correlation |
| ATP6V0E1 | 0.234076 | 2.14E-05 | Other Positive Correlation |
| CLDN3 | 0.234023 | 2.15E-05 | Other Positive Correlation |
| HAUS6 | 0.233897 | 2.17E-05 | Other Positive Correlation |
| SEPTIN3 | 0.233866 | 2.18E-05 | Other Positive Correlation |
| TNNI3 | 0.233833 | 2.18E-05 | Other Positive Correlation |
| SYT13 | 0.233375 | 2.27E-05 | Other Positive Correlation |
| CALCOCO2 | 0.233345 | 2.27E-05 | Other Positive Correlation |
| LAPTM4A | 0.233007 | 2.34E-05 | Other Positive Correlation |
| NCBP2AS2 | 0.232945 | 2.35E-05 | Other Positive Correlation |
| EIF5 | 0.232858 | 2.37E-05 | Other Positive Correlation |
| ARHGAP9 | 0.232726 | 2.39E-05 | Other Positive Correlation |
| ACYP1 | 0.232463 | 2.45E-05 | Other Positive Correlation |
| MAP7D2 | 0.232239 | 2.49E-05 | Other Positive Correlation |
| CEP128 | 0.232186 | 2.5E-05 | Other Positive Correlation |
| WT1 | 0.232067 | 2.53E-05 | Other Positive Correlation |
| LINGO1 | 0.231725 | 2.6E-05 | Other Positive Correlation |
| RFC4 | 0.231639 | 2.62E-05 | Other Positive Correlation |
| PIEZO2 | 0.231576 | 2.63E-05 | Other Positive Correlation |
| CTH | 0.231559 | 2.64E-05 | Other Positive Correlation |
| NEMP1 | 0.231531 | 2.64E-05 | Other Positive Correlation |
| TIMP4 | 0.231248 | 2.71E-05 | Other Positive Correlation |
| CRACD | 0.23096 | 2.77E-05 | Other Positive Correlation |
| AGT | 0.230891 | 2.79E-05 | Other Positive Correlation |
| LPL | 0.230867 | 2.79E-05 | Other Positive Correlation |
| DOK5 | 0.230788 | 2.81E-05 | Other Positive Correlation |
| PKD1 | 0.230587 | 2.86E-05 | Other Positive Correlation |

| | | | |
|---|---|---|---|
| DUSP15 | 0.230441 | 2.89E-05 | Other Positive Correlation |
| BATF3 | 0.230231 | 2.94E-05 | Other Positive Correlation |
| CCSAP | 0.230117 | 2.97E-05 | Other Positive Correlation |
| KIFC2 | 0.229928 | 3.01E-05 | Other Positive Correlation |
| NAGK | 0.229925 | 3.02E-05 | Other Positive Correlation |
| ULK1 | 0.229903 | 3.02E-05 | Other Positive Correlation |
| CENPU | 0.229662 | 3.08E-05 | Other Positive Correlation |
| PTP4A2 | 0.229443 | 3.14E-05 | Other Positive Correlation |
| SORT1 | 0.229051 | 3.24E-05 | Other Positive Correlation |
| ABHD8 | 0.228955 | 3.26E-05 | Other Positive Correlation |
| CLIC6 | 0.228911 | 3.28E-05 | Other Positive Correlation |
| DUSP26 | 0.228841 | 3.29E-05 | Other Positive Correlation |
| FBXO48 | 0.228826 | 3.3E-05 | Other Positive Correlation |
| CHAF1A | 0.228673 | 3.34E-05 | Other Positive Correlation |
| SEPTIN4 | 0.228568 | 3.37E-05 | Other Positive Correlation |
| C16orf91 | 0.228451 | 3.4E-05 | Other Positive Correlation |
| NXT1 | 0.22833 | 3.43E-05 | Other Positive Correlation |
| SLC38A3 | 0.22787 | 3.57E-05 | Other Positive Correlation |
| SCGB3A1 | 0.227565 | 3.65E-05 | Other Positive Correlation |
| KIAA0232 | 0.227475 | 3.68E-05 | Other Positive Correlation |
| SAPCD2 | 0.227373 | 3.71E-05 | Other Positive Correlation |
| SPOCK2 | 0.227361 | 3.72E-05 | Other Positive Correlation |
| TSPAN33 | 0.227094 | 3.8E-05 | Other Positive Correlation |
| ARFGEF3 | 0.22673 | 3.91E-05 | Other Positive Correlation |
| TLX2 | 0.22672 | 3.91E-05 | Other Positive Correlation |
| SLC1A4 | 0.226636 | 3.94E-05 | Other Positive Correlation |
| MSX1 | 0.226588 | 3.96E-05 | Other Positive Correlation |
| FGF9 | 0.226523 | 3.98E-05 | Other Positive Correlation |
| PAIP2B | 0.226488 | 3.99E-05 | Other Positive Correlation |
| MARCHF3 | 0.226345 | 4.03E-05 | Other Positive Correlation |
| COBL | 0.226266 | 4.06E-05 | Other Positive Correlation |
| GNA13 | 0.225878 | 4.19E-05 | Other Positive Correlation |
| CEBPG | 0.22581 | 4.21E-05 | Other Positive Correlation |
| TMEM167B | 0.22581 | 4.21E-05 | Other Positive Correlation |
| TMEM108 | 0.225697 | 4.25E-05 | Other Positive Correlation |
| PLEKHO1 | 0.225576 | 4.29E-05 | Other Positive Correlation |
| TEDC2 | 0.225548 | 4.3E-05 | Other Positive Correlation |
| HSPA9 | 0.22544 | 4.34E-05 | Other Positive Correlation |
| STOX1 | 0.225244 | 4.41E-05 | Other Positive Correlation |
| MIS18A | 0.224925 | 4.52E-05 | Other Positive Correlation |
| SCML2 | 0.224882 | 4.54E-05 | Other Positive Correlation |
| RNF5 | 0.224784 | 4.57E-05 | Other Positive Correlation |
| LRP1 | 0.224667 | 4.61E-05 | Other Positive Correlation |
| FN3K | 0.224476 | 4.69E-05 | Other Positive Correlation |
| HSPB8 | 0.224232 | 4.78E-05 | Other Positive Correlation |
| SIPA1L2 | 0.223529 | 5.05E-05 | Other Positive Correlation |
| PDXDC1 | 0.22348 | 5.07E-05 | Other Positive Correlation |
| ARHGAP45 | 0.222664 | 5.41E-05 | Other Positive Correlation |

| | | | |
|---|---|---|---|
| SYCP2 | 0.222562 | 5.46E-05 | Other Positive Correlation |
| FLAD1 | 0.222434 | 5.51E-05 | Other Positive Correlation |
| FHIT | 0.222186 | 5.62E-05 | Other Positive Correlation |
| PLAGL2 | 0.222052 | 5.68E-05 | Other Positive Correlation |
| TRAK2 | 0.221987 | 5.71E-05 | Other Positive Correlation |
| OLFM2 | 0.221756 | 5.82E-05 | Other Positive Correlation |
| CRYBA2 | 0.221492 | 5.94E-05 | Other Positive Correlation |
| DAPK1 | 0.221311 | 6.02E-05 | Other Positive Correlation |
| REEP1 | 0.22124 | 6.06E-05 | Other Positive Correlation |
| SDR16C5 | 0.221189 | 6.08E-05 | Other Positive Correlation |
| SYT17 | 0.220908 | 6.22E-05 | Other Positive Correlation |
| TIMM29 | 0.220831 | 6.26E-05 | Other Positive Correlation |
| REEP4 | 0.220765 | 6.29E-05 | Other Positive Correlation |
| SEPHS2 | 0.220694 | 6.32E-05 | Other Positive Correlation |
| SYT7 | 0.220477 | 6.43E-05 | Other Positive Correlation |
| ASCL2 | 0.22023 | 6.56E-05 | Other Positive Correlation |
| RASEF | 0.219905 | 6.73E-05 | Other Positive Correlation |
| TYRO3 | 0.219545 | 6.92E-05 | Other Positive Correlation |
| NKIRAS1 | 0.219333 | 7.03E-05 | Other Positive Correlation |
| MAST1 | 0.219201 | 7.11E-05 | Other Positive Correlation |
| CIB2 | 0.219117 | 7.15E-05 | Other Positive Correlation |
| RBPMS2 | 0.219046 | 7.19E-05 | Other Positive Correlation |
| LMCD1 | 0.218859 | 7.3E-05 | Other Positive Correlation |
| C14orf132 | 0.218807 | 7.33E-05 | Other Positive Correlation |
| TJP3 | 0.218641 | 7.42E-05 | Other Positive Correlation |
| SEMA4D | 0.218455 | 7.53E-05 | Other Positive Correlation |
| CENPM | 0.218108 | 7.74E-05 | Other Positive Correlation |
| TPRA1 | 0.217984 | 7.81E-05 | Other Positive Correlation |
| CNNM1 | 0.217864 | 7.89E-05 | Other Positive Correlation |
| CDX1 | 0.217418 | 8.16E-05 | Other Positive Correlation |
| KCND3 | 0.217248 | 8.27E-05 | Other Positive Correlation |
| LONRF3 | 0.217164 | 8.32E-05 | Other Positive Correlation |
| SLC7A1 | 0.217013 | 8.42E-05 | Other Positive Correlation |
| RPS19BP1 | 0.216719 | 8.62E-05 | Other Positive Correlation |
| ST7 | 0.216706 | 8.62E-05 | Other Positive Correlation |
| USP44 | 0.216703 | 8.63E-05 | Other Positive Correlation |
| ADCK2 | 0.216597 | 8.7E-05 | Other Positive Correlation |
| MAL | 0.216403 | 8.83E-05 | Other Positive Correlation |
| TMEM255B | 0.216393 | 8.84E-05 | Other Positive Correlation |
| MOXD1 | 0.216351 | 8.86E-05 | Other Positive Correlation |
| GALK1 | 0.216344 | 8.87E-05 | Other Positive Correlation |
| HHEX | 0.216308 | 8.89E-05 | Other Positive Correlation |
| SPTBN4 | 0.216295 | 8.9E-05 | Other Positive Correlation |
| RFC2 | 0.216149 | 9E-05 | Other Positive Correlation |
| ALPK3 | 0.216108 | 9.03E-05 | Other Positive Correlation |
| ITPKA | 0.215969 | 9.13E-05 | Other Positive Correlation |
| TMEM38A | 0.215903 | 9.18E-05 | Other Positive Correlation |
| GPRIN2 | 0.215821 | 9.23E-05 | Other Positive Correlation |

| | | | |
|---|---|---|---|
| BORCS8 | 0.215728 | 9.3E-05 | Other Positive Correlation |
| C15orf48 | 0.215654 | 9.35E-05 | Other Positive Correlation |
| PPP2R2C | 0.21562 | 9.38E-05 | Other Positive Correlation |
| EFCAB11 | 0.215448 | 9.5E-05 | Other Positive Correlation |
| SPINT1 | 0.215129 | 9.74E-05 | Other Positive Correlation |
| GUCY1B1 | 0.214825 | 9.97E-05 | Other Positive Correlation |
| GTPBP2 | 0.2148 | 9.98E-05 | Other Positive Correlation |
| OSGIN2 | 0.214745 | 0.0001 | Other Positive Correlation |
| ZNF84 | 0.214698 | 0.000101 | Other Positive Correlation |
| ZBTB21 | 0.214395 | 0.000103 | Other Positive Correlation |
| ZNF367 | 0.214318 | 0.000104 | Other Positive Correlation |
| ALKAL2 | 0.214078 | 0.000105 | Other Positive Correlation |
| CHST1 | 0.213563 | 0.00011 | Other Positive Correlation |
| SERTAD2 | 0.213271 | 0.000112 | Other Positive Correlation |
| ZNF331 | 0.21324 | 0.000112 | Other Positive Correlation |
| THAP7 | 0.21261 | 0.000118 | Other Positive Correlation |
| HERPUD1 | 0.212463 | 0.000119 | Other Positive Correlation |
| FRY | 0.212455 | 0.000119 | Other Positive Correlation |
| PIP4P1 | 0.212376 | 0.00012 | Other Positive Correlation |
| AMOT | 0.212173 | 0.000122 | Other Positive Correlation |
| C5orf34 | 0.212155 | 0.000122 | Other Positive Correlation |
| ADRA2C | 0.212148 | 0.000122 | Other Positive Correlation |
| TRIM69 | 0.211797 | 0.000125 | Other Positive Correlation |
| STRN4 | 0.21169 | 0.000126 | Other Positive Correlation |
| COX7A1 | 0.211681 | 0.000127 | Other Positive Correlation |
| NR1D2 | 0.211676 | 0.000127 | Other Positive Correlation |
| PRSS36 | 0.211566 | 0.000128 | Other Positive Correlation |
| POC1A | 0.211492 | 0.000128 | Other Positive Correlation |
| ZBTB10 | 0.211162 | 0.000132 | Other Positive Correlation |
| RPL39L | 0.21116 | 0.000132 | Other Positive Correlation |
| HOXD8 | 0.211138 | 0.000132 | Other Positive Correlation |
| GSTM3 | 0.210898 | 0.000134 | Other Positive Correlation |
| PSAP | 0.210744 | 0.000136 | Other Positive Correlation |
| TTC39A | 0.210728 | 0.000136 | Other Positive Correlation |
| CLCN5 | 0.210641 | 0.000137 | Other Positive Correlation |
| NOTUM | 0.210371 | 0.00014 | Other Positive Correlation |
| MKX | 0.210129 | 0.000142 | Other Positive Correlation |
| RRAGC | 0.210039 | 0.000143 | Other Positive Correlation |
| SLC16A10 | 0.209995 | 0.000144 | Other Positive Correlation |
| HYLS1 | 0.209947 | 0.000144 | Other Positive Correlation |
| TSPAN18 | 0.209899 | 0.000145 | Other Positive Correlation |
| EBAG9 | 0.209897 | 0.000145 | Other Positive Correlation |
| NOVA1 | 0.209623 | 0.000148 | Other Positive Correlation |
| HSPA1A | 0.209569 | 0.000148 | Other Positive Correlation |
| FLNC | 0.209318 | 0.000151 | Other Positive Correlation |
| LY6H | 0.209155 | 0.000153 | Other Positive Correlation |
| COX14 | 0.208662 | 0.000159 | Other Positive Correlation |
| TRIB3 | 0.208602 | 0.000159 | Other Positive Correlation |

| | | | |
|---|---|---|---|
| TMEM70 | 0.208574 | 0.00016 | Other Positive Correlation |
| PITHD1 | 0.208487 | 0.000161 | Other Positive Correlation |
| SYNM | 0.208313 | 0.000163 | Other Positive Correlation |
| HMOX1 | 0.208196 | 0.000164 | Other Positive Correlation |
| LPIN2 | 0.20807 | 0.000166 | Other Positive Correlation |
| RGS16 | 0.208044 | 0.000166 | Other Positive Correlation |
| GPRC5B | 0.207971 | 0.000167 | Other Positive Correlation |
| NFATC2 | 0.207892 | 0.000168 | Other Positive Correlation |
| CPEB3 | 0.207857 | 0.000168 | Other Positive Correlation |
| MRPL18 | 0.207817 | 0.000169 | Other Positive Correlation |
| YTHDF2 | 0.207779 | 0.000169 | Other Positive Correlation |
| NOTCH3 | 0.207647 | 0.000171 | Other Positive Correlation |
| HLTF | 0.207612 | 0.000171 | Other Positive Correlation |
| CDKN2C | 0.207462 | 0.000173 | Other Positive Correlation |
| NCOA1 | 0.207427 | 0.000174 | Other Positive Correlation |
| TBC1D20 | 0.207245 | 0.000176 | Other Positive Correlation |
| SAMD11 | 0.207218 | 0.000176 | Other Positive Correlation |
| STARD3 | 0.207121 | 0.000178 | Other Positive Correlation |
| TMPRSS13 | 0.206921 | 0.00018 | Other Positive Correlation |
| HMX3 | 0.206876 | 0.000181 | Other Positive Correlation |
| FA2H | 0.206691 | 0.000183 | Other Positive Correlation |
| DERL3 | 0.206624 | 0.000184 | Other Positive Correlation |
| WNT6 | 0.206618 | 0.000184 | Other Positive Correlation |
| NRK | 0.206552 | 0.000185 | Other Positive Correlation |
| MAGI3 | 0.206362 | 0.000188 | Other Positive Correlation |
| STXBP6 | 0.206346 | 0.000188 | Other Positive Correlation |
| DNAJC1 | 0.206161 | 0.000191 | Other Positive Correlation |
| SQSTM1 | 0.206131 | 0.000191 | Other Positive Correlation |
| PELI2 | 0.206067 | 0.000192 | Other Positive Correlation |
| TMEM50B | 0.205994 | 0.000193 | Other Positive Correlation |
| CCPG1 | 0.205882 | 0.000195 | Other Positive Correlation |
| GGH | 0.205653 | 0.000198 | Other Positive Correlation |
| WDR24 | 0.205636 | 0.000198 | Other Positive Correlation |
| NIPA2 | 0.205014 | 0.000207 | Other Positive Correlation |
| SRSF12 | 0.204802 | 0.000211 | Other Positive Correlation |
| PEPD | 0.204549 | 0.000215 | Other Positive Correlation |
| YPEL5 | 0.20438 | 0.000217 | Other Positive Correlation |
| GULP1 | 0.204364 | 0.000218 | Other Positive Correlation |
| SLTM | 0.204229 | 0.00022 | Other Positive Correlation |
| CCDC167 | 0.2041 | 0.000222 | Other Positive Correlation |
| STMND1 | 0.203955 | 0.000224 | Other Positive Correlation |
| BTF3L4 | 0.203807 | 0.000227 | Other Positive Correlation |
| DSN1 | 0.20362 | 0.00023 | Other Positive Correlation |
| CRABP2 | 0.20362 | 0.00023 | Other Positive Correlation |
| PPP6R1 | 0.203524 | 0.000231 | Other Positive Correlation |
| DIPK1B | 0.203391 | 0.000233 | Other Positive Correlation |
| CDH2 | 0.203361 | 0.000234 | Other Positive Correlation |
| H2AJ | 0.203151 | 0.000238 | Other Positive Correlation |

| | | | |
|---|---|---|---|
| CNNM2 | 0.203096 | 0.000238 | Other Positive Correlation |
| BMP7 | 0.203055 | 0.000239 | Other Positive Correlation |
| GPS1 | 0.20291 | 0.000242 | Other Positive Correlation |
| OTULINL | 0.202752 | 0.000245 | Other Positive Correlation |
| TRMU | 0.202689 | 0.000246 | Other Positive Correlation |
| FOXA1 | 0.202634 | 0.000247 | Other Positive Correlation |
| AUNIP | 0.202587 | 0.000247 | Other Positive Correlation |
| FAM171B | 0.202543 | 0.000248 | Other Positive Correlation |
| GPR160 | 0.202491 | 0.000249 | Other Positive Correlation |
| SMOX | 0.202487 | 0.000249 | Other Positive Correlation |
| TP53I13 | 0.202359 | 0.000252 | Other Positive Correlation |
| LSR | 0.202287 | 0.000253 | Other Positive Correlation |
| LRRN1 | 0.202038 | 0.000257 | Other Positive Correlation |
| ENPP4 | 0.201636 | 0.000265 | Other Positive Correlation |
| SLC31A1 | 0.201515 | 0.000267 | Other Positive Correlation |
| NR4A1 | 0.201419 | 0.000269 | Other Positive Correlation |
| NAA30 | 0.201328 | 0.000271 | Other Positive Correlation |
| EPHB6 | 0.201321 | 0.000271 | Other Positive Correlation |
| CDKN1C | 0.201081 | 0.000276 | Other Positive Correlation |
| TMEM37 | 0.200995 | 0.000277 | Other Positive Correlation |
| PPP2R3B | 0.200789 | 0.000282 | Other Positive Correlation |
| SNTB1 | 0.200529 | 0.000287 | Other Positive Correlation |
| ARHGDIG | 0.200179 | 0.000294 | Other Positive Correlation |
| COLEC12 | 0.200129 | 0.000295 | Other Positive Correlation |
| ID4 | 0.200103 | 0.000296 | Other Positive Correlation |
| CFAP43 | 0.200025 | 0.000297 | Other Positive Correlation |
| SRRM3 | 0.199945 | 0.000299 | Other Positive Correlation |
| MSI1 | 0.199699 | 0.000304 | Other Positive Correlation |
| ASB8 | 0.199658 | 0.000305 | Other Positive Correlation |
| WDR62 | 0.199546 | 0.000308 | Other Positive Correlation |
| DOLPP1 | 0.199481 | 0.000309 | Other Positive Correlation |
| BCL2L12 | 0.199438 | 0.00031 | Other Positive Correlation |
| ANKRD53 | 0.199303 | 0.000313 | Other Positive Correlation |
| CENPQ | 0.199039 | 0.000319 | Other Positive Correlation |
| SH3KBP1 | 0.198967 | 0.000321 | Other Positive Correlation |
| PDLIM3 | 0.198967 | 0.000321 | Other Positive Correlation |
| C21orf58 | 0.198771 | 0.000325 | Other Positive Correlation |
| CGN | 0.198552 | 0.00033 | Other Positive Correlation |
| RIMOC1 | 0.198407 | 0.000334 | Other Positive Correlation |
| EBF3 | 0.198391 | 0.000334 | Other Positive Correlation |
| RNF208 | 0.19834 | 0.000335 | Other Positive Correlation |
| STIL | 0.198279 | 0.000337 | Other Positive Correlation |
| MLXIPL | 0.198106 | 0.000341 | Other Positive Correlation |
| SWI5 | 0.198046 | 0.000342 | Other Positive Correlation |
| PSIP1 | 0.197805 | 0.000348 | Other Positive Correlation |
| NDEL1 | 0.197579 | 0.000354 | Other Positive Correlation |
| TMSB15A | 0.197576 | 0.000354 | Other Positive Correlation |
| GPR137 | 0.197071 | 0.000367 | Other Positive Correlation |

| | | | |
|---|---|---|---|
| RMC1 | 0.196888 | 0.000371 | Other Positive Correlation |
| BCAS2 | 0.196883 | 0.000371 | Other Positive Correlation |
| USP2 | 0.196804 | 0.000373 | Other Positive Correlation |
| DCAF7 | 0.196717 | 0.000376 | Other Positive Correlation |
| RAB5IF | 0.196622 | 0.000378 | Other Positive Correlation |
| SP4 | 0.196612 | 0.000379 | Other Positive Correlation |
| BRSK2 | 0.196242 | 0.000388 | Other Positive Correlation |
| KLHDC3 | 0.196211 | 0.000389 | Other Positive Correlation |
| SYVN1 | 0.196125 | 0.000392 | Other Positive Correlation |
| TMEM145 | 0.196122 | 0.000392 | Other Positive Correlation |
| ELOVL7 | 0.195767 | 0.000402 | Other Positive Correlation |
| HHIPL1 | 0.19547 | 0.00041 | Other Positive Correlation |
| SCAPER | 0.195456 | 0.00041 | Other Positive Correlation |
| MCM3 | 0.195405 | 0.000412 | Other Positive Correlation |
| UBN1 | 0.195025 | 0.000423 | Other Positive Correlation |
| NAGPA | 0.194906 | 0.000426 | Other Positive Correlation |
| PAQR5 | 0.194656 | 0.000434 | Other Positive Correlation |
| JPH4 | 0.194222 | 0.000447 | Other Positive Correlation |
| ECEL1 | 0.194189 | 0.000448 | Other Positive Correlation |
| EP300 | 0.194119 | 0.00045 | Other Positive Correlation |
| NAT8L | 0.194085 | 0.000451 | Other Positive Correlation |
| HOOK1 | 0.194072 | 0.000452 | Other Positive Correlation |
| PEG3 | 0.193989 | 0.000454 | Other Positive Correlation |
| UBE2Q1 | 0.1937 | 0.000464 | Other Positive Correlation |
| TMED5 | 0.193337 | 0.000475 | Other Positive Correlation |
| MANEAL | 0.193305 | 0.000476 | Other Positive Correlation |
| SFMBT2 | 0.193113 | 0.000483 | Other Positive Correlation |
| C1QTNF2 | 0.19285 | 0.000492 | Other Positive Correlation |
| IGFBPL1 | 0.19269 | 0.000497 | Other Positive Correlation |
| HPCAL4 | 0.192602 | 0.0005 | Other Positive Correlation |
| YOD1 | 0.192397 | 0.000507 | Other Positive Correlation |
| HTRA3 | 0.192311 | 0.00051 | Other Positive Correlation |
| SLC30A3 | 0.19227 | 0.000512 | Other Positive Correlation |
| ZNF93 | 0.192176 | 0.000515 | Other Positive Correlation |
| DYNC1I1 | 0.192133 | 0.000516 | Other Positive Correlation |
| DNM1 | 0.192066 | 0.000519 | Other Positive Correlation |
| TP53BP2 | 0.191949 | 0.000523 | Other Positive Correlation |
| CPT1C | 0.191899 | 0.000525 | Other Positive Correlation |
| SLBP | 0.191774 | 0.000529 | Other Positive Correlation |
| CDC25A | 0.191763 | 0.00053 | Other Positive Correlation |
| CENPB | 0.191762 | 0.00053 | Other Positive Correlation |
| FAM162B | 0.19168 | 0.000533 | Other Positive Correlation |
| MB | 0.191475 | 0.00054 | Other Positive Correlation |
| MFHAS1 | 0.191319 | 0.000546 | Other Positive Correlation |
| TMEM125 | 0.191115 | 0.000554 | Other Positive Correlation |
| AQP5 | 0.190961 | 0.00056 | Other Positive Correlation |
| RAB5B | 0.190759 | 0.000567 | Other Positive Correlation |
| CHN1 | 0.190572 | 0.000575 | Other Positive Correlation |

| | | | |
|---|---|---|---|
| PLIN2 | 0.190157 | 0.000591 | Other Positive Correlation |
| PNMA2 | 0.190011 | 0.000597 | Other Positive Correlation |
| HCN2 | 0.189972 | 0.000599 | Other Positive Correlation |
| LIN9 | 0.189858 | 0.000603 | Other Positive Correlation |
| WIPI1 | 0.189806 | 0.000605 | Other Positive Correlation |
| PRKG1 | 0.1897 | 0.00061 | Other Positive Correlation |
| LONP1 | 0.189626 | 0.000613 | Other Positive Correlation |
| MISP3 | 0.189416 | 0.000622 | Other Positive Correlation |
| LRRC4B | 0.189412 | 0.000622 | Other Positive Correlation |
| CDK5R2 | 0.189005 | 0.000639 | Other Positive Correlation |
| S1PR1 | 0.188954 | 0.000641 | Other Positive Correlation |
| BICDL2 | 0.188869 | 0.000645 | Other Positive Correlation |
| CENPT | 0.188719 | 0.000651 | Other Positive Correlation |
| BGN | 0.188653 | 0.000654 | Other Positive Correlation |
| ENPP1 | 0.188632 | 0.000655 | Other Positive Correlation |
| RNF168 | 0.188504 | 0.000661 | Other Positive Correlation |
| CNOT1 | 0.188431 | 0.000664 | Other Positive Correlation |
| INPP5F | 0.188381 | 0.000666 | Other Positive Correlation |
| DLX4 | 0.188371 | 0.000667 | Other Positive Correlation |
| ANK1 | 0.18831 | 0.00067 | Other Positive Correlation |
| C6orf120 | 0.188293 | 0.00067 | Other Positive Correlation |
| GNAZ | 0.188253 | 0.000672 | Other Positive Correlation |
| HOXD1 | 0.188122 | 0.000678 | Other Positive Correlation |
| HR | 0.18811 | 0.000679 | Other Positive Correlation |
| EGFL7 | 0.187868 | 0.00069 | Other Positive Correlation |
| ULBP1 | 0.187808 | 0.000693 | Other Positive Correlation |
| HOXB7 | 0.187748 | 0.000695 | Other Positive Correlation |
| NDUFA2 | 0.187605 | 0.000702 | Other Positive Correlation |
| C3orf70 | 0.187556 | 0.000704 | Other Positive Correlation |
| BAMBI | 0.187504 | 0.000707 | Other Positive Correlation |
| ZNF530 | 0.187436 | 0.00071 | Other Positive Correlation |
| PPP2R5B | 0.187284 | 0.000717 | Other Positive Correlation |
| RARRES2 | 0.187215 | 0.000721 | Other Positive Correlation |
| ATRNL1 | 0.187092 | 0.000727 | Other Positive Correlation |
| RMI1 | 0.187081 | 0.000727 | Other Positive Correlation |
| SCN8A | 0.187076 | 0.000727 | Other Positive Correlation |
| DDX24 | 0.18707 | 0.000728 | Other Positive Correlation |
| SYNJ2 | 0.187001 | 0.000731 | Other Positive Correlation |
| SLC30A2 | 0.186993 | 0.000731 | Other Positive Correlation |
| SNN | 0.186946 | 0.000734 | Other Positive Correlation |
| MAGEF1 | 0.186831 | 0.00074 | Other Positive Correlation |
| TUSC2 | 0.186613 | 0.00075 | Other Positive Correlation |
| ALDH1L2 | 0.186562 | 0.000753 | Other Positive Correlation |
| ATXN1L | 0.186499 | 0.000756 | Other Positive Correlation |
| NEBL | 0.18647 | 0.000758 | Other Positive Correlation |
| OTUB2 | 0.186324 | 0.000765 | Other Positive Correlation |
| LIN54 | 0.185968 | 0.000783 | Other Positive Correlation |
| YIPF5 | 0.185911 | 0.000786 | Other Positive Correlation |

| | | | |
|---|---|---|---|
| ADAP2 | 0.185695 | 0.000798 | Other Positive Correlation |
| TIMELESS | 0.185538 | 0.000806 | Other Positive Correlation |
| PTGER2 | 0.185468 | 0.00081 | Other Positive Correlation |
| SLC44A4 | 0.185101 | 0.00083 | Other Positive Correlation |
| EML1 | 0.184932 | 0.000839 | Other Positive Correlation |
| ZNF778 | 0.184912 | 0.00084 | Other Positive Correlation |
| FLYWCH2 | 0.184834 | 0.000844 | Other Positive Correlation |
| TTLL7 | 0.184828 | 0.000845 | Other Positive Correlation |
| C19orf81 | 0.184755 | 0.000849 | Other Positive Correlation |
| SBK1 | 0.18471 | 0.000851 | Other Positive Correlation |
| SLC25A42 | 0.18469 | 0.000853 | Other Positive Correlation |
| RBM11 | 0.184498 | 0.000863 | Other Positive Correlation |
| RUNDC3B | 0.184311 | 0.000874 | Other Positive Correlation |
| SGCG | 0.184253 | 0.000877 | Other Positive Correlation |
| AMN | 0.18425 | 0.000878 | Other Positive Correlation |
| CDC42EP1 | 0.184143 | 0.000884 | Other Positive Correlation |
| ATP1B2 | 0.184046 | 0.00089 | Other Positive Correlation |
| CDYL | 0.18392 | 0.000897 | Other Positive Correlation |
| ZMAT1 | 0.183756 | 0.000907 | Other Positive Correlation |
| BPGM | 0.183637 | 0.000914 | Other Positive Correlation |
| FOXA3 | 0.183612 | 0.000915 | Other Positive Correlation |
| GAREM2 | 0.183538 | 0.00092 | Other Positive Correlation |
| SNTA1 | 0.183468 | 0.000924 | Other Positive Correlation |
| FBXO30 | 0.183388 | 0.000929 | Other Positive Correlation |
| S100A4 | 0.183347 | 0.000931 | Other Positive Correlation |
| THAP10 | 0.183276 | 0.000936 | Other Positive Correlation |
| THAP11 | 0.183162 | 0.000943 | Other Positive Correlation |
| KCTD13 | 0.183096 | 0.000947 | Other Positive Correlation |
| LTK | 0.183028 | 0.000951 | Other Positive Correlation |
| VPS29 | 0.183014 | 0.000952 | Other Positive Correlation |
| CCDC74A | 0.183001 | 0.000953 | Other Positive Correlation |
| RPUSD2 | 0.182919 | 0.000958 | Other Positive Correlation |
| UPF2 | 0.182762 | 0.000968 | Other Positive Correlation |
| ADAMTS2 | 0.182659 | 0.000974 | Other Positive Correlation |
| MAP6 | 0.18254 | 0.000982 | Other Positive Correlation |
| CERS1 | 0.18227 | 0.000999 | Other Positive Correlation |
| ZFP91 | 0.182251 | 0.001001 | Other Positive Correlation |
| ARMH4 | 0.182216 | 0.001003 | Other Positive Correlation |
| VAMP2 | 0.182008 | 0.001017 | Other Positive Correlation |
| MRPL34 | 0.181995 | 0.001017 | Other Positive Correlation |
| KLK11 | 0.181699 | 0.001037 | Other Positive Correlation |
| PSENEN | 0.181618 | 0.001043 | Other Positive Correlation |
| DHRS3 | 0.181544 | 0.001048 | Other Positive Correlation |
| B4GALT5 | 0.181498 | 0.001051 | Other Positive Correlation |
| ATG16L1 | 0.181441 | 0.001055 | Other Positive Correlation |
| PREX1 | 0.181409 | 0.001057 | Other Positive Correlation |
| MRE11 | 0.181247 | 0.001068 | Other Positive Correlation |
| COMMD5 | 0.180922 | 0.001091 | Other Positive Correlation |

| | | | |
|---|---|---|---|
| SAYSD1 | 0.180921 | 0.001091 | Other Positive Correlation |
| DCP2 | 0.180918 | 0.001091 | Other Positive Correlation |
| SLC22A17 | 0.180713 | 0.001106 | Other Positive Correlation |
| C2CD4C | 0.180699 | 0.001107 | Other Positive Correlation |
| SULT4A1 | 0.180638 | 0.001111 | Other Positive Correlation |
| PTPRS | 0.180484 | 0.001122 | Other Positive Correlation |
| ERCC3 | 0.180409 | 0.001128 | Other Positive Correlation |
| RNASEH2A | 0.180208 | 0.001143 | Other Positive Correlation |
| TTC32 | 0.18007 | 0.001153 | Other Positive Correlation |
| KISS1R | 0.180014 | 0.001157 | Other Positive Correlation |
| NFE2L1 | 0.179997 | 0.001158 | Other Positive Correlation |
| RTKN2 | 0.17997 | 0.00116 | Other Positive Correlation |
| DCHS1 | 0.179836 | 0.00117 | Other Positive Correlation |
| PAX2 | 0.179679 | 0.001182 | Other Positive Correlation |
| POMGNT2 | 0.179646 | 0.001185 | Other Positive Correlation |
| CLDN4 | 0.179635 | 0.001186 | Other Positive Correlation |
| VEGFB | 0.179514 | 0.001195 | Other Positive Correlation |
| ICA1 | 0.179494 | 0.001196 | Other Positive Correlation |
| GTF2A1 | 0.179239 | 0.001216 | Other Positive Correlation |
| PANX2 | 0.179155 | 0.001223 | Other Positive Correlation |
| CRAT | 0.179154 | 0.001223 | Other Positive Correlation |
| PHOX2A | 0.178988 | 0.001236 | Other Positive Correlation |
| CDR2 | 0.17887 | 0.001245 | Other Positive Correlation |
| POU4F1 | 0.178838 | 0.001248 | Other Positive Correlation |
| FAM219A | 0.178369 | 0.001286 | Other Positive Correlation |
| AGO2 | 0.178329 | 0.001289 | Other Positive Correlation |
| PURB | 0.178177 | 0.001302 | Other Positive Correlation |
| LHX2 | 0.178156 | 0.001304 | Other Positive Correlation |
| ZMYND19 | 0.177944 | 0.001321 | Other Positive Correlation |
| TFR2 | 0.17793 | 0.001322 | Other Positive Correlation |
| YPEL2 | 0.177914 | 0.001324 | Other Positive Correlation |
| EPB41L1 | 0.177822 | 0.001332 | Other Positive Correlation |
| OSBP | 0.177796 | 0.001334 | Other Positive Correlation |
| INPP5J | 0.177708 | 0.001341 | Other Positive Correlation |
| PARD3B | 0.177604 | 0.00135 | Other Positive Correlation |
| TMEM11 | 0.177516 | 0.001358 | Other Positive Correlation |
| WNK2 | 0.177448 | 0.001364 | Other Positive Correlation |
| FOXE3 | 0.177397 | 0.001368 | Other Positive Correlation |
| MAN1A1 | 0.177296 | 0.001377 | Other Positive Correlation |
| DTX1 | 0.177226 | 0.001383 | Other Positive Correlation |
| SLCO4A1 | 0.177164 | 0.001389 | Other Positive Correlation |
| CBS | 0.177151 | 0.00139 | Other Positive Correlation |
| ACSL1 | 0.17713 | 0.001392 | Other Positive Correlation |
| YPEL1 | 0.176961 | 0.001407 | Other Positive Correlation |
| SMPD1 | 0.176887 | 0.001413 | Other Positive Correlation |
| SRPRA | 0.176861 | 0.001416 | Other Positive Correlation |
| C10orf82 | 0.176847 | 0.001417 | Other Positive Correlation |
| BHMT2 | 0.176739 | 0.001427 | Other Positive Correlation |

| | | | |
|---|---|---|---|
| CECR2 | 0.176713 | 0.001429 | Other Positive Correlation |
| BOLA1 | 0.176558 | 0.001443 | Other Positive Correlation |
| GAN | 0.176463 | 0.001452 | Other Positive Correlation |
| TNFRSF1B | 0.176397 | 0.001458 | Other Positive Correlation |
| CYRIA | 0.176307 | 0.001466 | Other Positive Correlation |
| ACE | 0.176216 | 0.001475 | Other Positive Correlation |
| DOK6 | 0.175931 | 0.001501 | Other Positive Correlation |
| CAND2 | 0.175844 | 0.00151 | Other Positive Correlation |
| SLC25A25 | 0.175666 | 0.001527 | Other Positive Correlation |
| C19orf73 | 0.175584 | 0.001535 | Other Positive Correlation |
| RAB11FIP1 | 0.175576 | 0.001535 | Other Positive Correlation |
| HCN4 | 0.175095 | 0.001583 | Other Positive Correlation |
| DNAJB2 | 0.175061 | 0.001586 | Other Positive Correlation |
| PID1 | 0.174887 | 0.001603 | Other Positive Correlation |
| TADA2B | 0.17484 | 0.001608 | Other Positive Correlation |
| DEUP1 | 0.174821 | 0.00161 | Other Positive Correlation |
| MCOLN3 | 0.17476 | 0.001616 | Other Positive Correlation |
| IER3IP1 | 0.174704 | 0.001622 | Other Positive Correlation |
| SRGAP2 | 0.174619 | 0.001631 | Other Positive Correlation |
| GOT1 | 0.174604 | 0.001632 | Other Positive Correlation |
| TMEM251 | 0.174543 | 0.001638 | Other Positive Correlation |
| COL9A2 | 0.174476 | 0.001645 | Other Positive Correlation |
| SLC39A14 | 0.174458 | 0.001647 | Other Positive Correlation |
| ISOC2 | 0.17442 | 0.001651 | Other Positive Correlation |
| SNCB | 0.174395 | 0.001654 | Other Positive Correlation |
| FAM43A | 0.174394 | 0.001654 | Other Positive Correlation |
| MARS1 | 0.174256 | 0.001668 | Other Positive Correlation |
| CCDC117 | 0.174207 | 0.001673 | Other Positive Correlation |
| GABRQ | 0.174169 | 0.001677 | Other Positive Correlation |
| PLPPR3 | 0.174065 | 0.001688 | Other Positive Correlation |
| SPNS2 | 0.173987 | 0.001697 | Other Positive Correlation |
| AMZ1 | 0.173969 | 0.001698 | Other Positive Correlation |
| CLGN | 0.173943 | 0.001701 | Other Positive Correlation |
| ALG2 | 0.173793 | 0.001717 | Other Positive Correlation |
| MAST2 | 0.173655 | 0.001732 | Other Positive Correlation |
| FOXL2 | 0.173572 | 0.001741 | Other Positive Correlation |
| DAG1 | 0.17357 | 0.001741 | Other Positive Correlation |
| MGAT3 | 0.173492 | 0.00175 | Other Positive Correlation |
| PRR18 | 0.1734 | 0.00176 | Other Positive Correlation |
| DCK | 0.172988 | 0.001806 | Other Positive Correlation |
| MUS81 | 0.172935 | 0.001812 | Other Positive Correlation |
| ZNF439 | 0.172868 | 0.001819 | Other Positive Correlation |
| AGPAT2 | 0.172846 | 0.001822 | Other Positive Correlation |
| AMDHD2 | 0.172817 | 0.001825 | Other Positive Correlation |
| TNFAIP8L3 | 0.172807 | 0.001826 | Other Positive Correlation |
| SLAIN1 | 0.172798 | 0.001827 | Other Positive Correlation |
| SLC27A6 | 0.172545 | 0.001856 | Other Positive Correlation |
| KIAA1958 | 0.17248 | 0.001863 | Other Positive Correlation |

| | | | |
|---|---|---|---|
| CNN1 | 0.172473 | 0.001864 | Other Positive Correlation |
| UBE2A | 0.172446 | 0.001867 | Other Positive Correlation |
| FBXO2 | 0.172287 | 0.001886 | Other Positive Correlation |
| TBX2 | 0.172274 | 0.001887 | Other Positive Correlation |
| PHTF2 | 0.17224 | 0.001891 | Other Positive Correlation |
| CIAO2B | 0.172142 | 0.001903 | Other Positive Correlation |
| CNPPD1 | 0.172074 | 0.001911 | Other Positive Correlation |
| CDIPT | 0.171973 | 0.001923 | Other Positive Correlation |
| FUZ | 0.171826 | 0.00194 | Other Positive Correlation |
| ZNF560 | 0.171804 | 0.001943 | Other Positive Correlation |
| PAX6 | 0.171662 | 0.00196 | Other Positive Correlation |
| TFE3 | 0.171494 | 0.001981 | Other Positive Correlation |
| ALPL | 0.171443 | 0.001987 | Other Positive Correlation |
| GPC2 | 0.171384 | 0.001994 | Other Positive Correlation |
| ABCA3 | 0.171332 | 0.002001 | Other Positive Correlation |
| MT1G | 0.171322 | 0.002002 | Other Positive Correlation |
| FAM107B | 0.17129 | 0.002006 | Other Positive Correlation |
| NPNT | 0.171109 | 0.002028 | Other Positive Correlation |
| GCH1 | 0.171091 | 0.00203 | Other Positive Correlation |
| PRPH | 0.170993 | 0.002043 | Other Positive Correlation |
| SLC7A3 | 0.170906 | 0.002054 | Other Positive Correlation |
| HUNK | 0.170881 | 0.002057 | Other Positive Correlation |
| MED8 | 0.1708 | 0.002067 | Other Positive Correlation |
| PIANP | 0.170653 | 0.002086 | Other Positive Correlation |
| PHIP | 0.170496 | 0.002106 | Other Positive Correlation |
| FBLN5 | 0.170228 | 0.002141 | Other Positive Correlation |
| PTEN | 0.170117 | 0.002156 | Other Positive Correlation |
| C17orf107 | 0.169965 | 0.002176 | Other Positive Correlation |
| LSM10 | 0.169867 | 0.002189 | Other Positive Correlation |
| CPEB1 | 0.169709 | 0.00221 | Other Positive Correlation |
| FXYD6 | 0.169697 | 0.002212 | Other Positive Correlation |
| RAB3A | 0.169537 | 0.002233 | Other Positive Correlation |
| KLHL35 | 0.169388 | 0.002254 | Other Positive Correlation |
| TPRN | 0.169265 | 0.002271 | Other Positive Correlation |
| EEF1AKMT4 | 0.169232 | 0.002275 | Other Positive Correlation |
| RAB29 | 0.169127 | 0.00229 | Other Positive Correlation |
| TUBB4A | 0.168983 | 0.00231 | Other Positive Correlation |
| NCAPD3 | 0.168847 | 0.002329 | Other Positive Correlation |
| SUMO1 | 0.168807 | 0.002335 | Other Positive Correlation |
| OLFM1 | 0.16876 | 0.002342 | Other Positive Correlation |
| CRACDL | 0.168637 | 0.002359 | Other Positive Correlation |
| SLC9A3 | 0.168463 | 0.002384 | Other Positive Correlation |
| CBX7 | 0.168459 | 0.002385 | Other Positive Correlation |
| FAM149A | 0.168386 | 0.002395 | Other Positive Correlation |
| NCAM1 | 0.168352 | 0.0024 | Other Positive Correlation |
| RADIL | 0.168167 | 0.002428 | Other Positive Correlation |
| RBP4 | 0.168128 | 0.002433 | Other Positive Correlation |
| MAFK | 0.167882 | 0.00247 | Other Positive Correlation |

| | | | |
|---|---|---|---|
| TSPYL5 | 0.167845 | 0.002475 | Other Positive Correlation |
| PDE6D | 0.167748 | 0.00249 | Other Positive Correlation |
| MPLKIP | 0.16773 | 0.002493 | Other Positive Correlation |
| KCNN1 | 0.167653 | 0.002504 | Other Positive Correlation |
| TMEM200C | 0.167498 | 0.002528 | Other Positive Correlation |
| SESTD1 | 0.16749 | 0.002529 | Other Positive Correlation |
| EN2 | 0.167373 | 0.002547 | Other Positive Correlation |
| FERMT2 | 0.167276 | 0.002562 | Other Positive Correlation |
| HSPA12B | 0.167262 | 0.002564 | Other Positive Correlation |
| CNIH2 | 0.167184 | 0.002576 | Other Positive Correlation |
| KCNJ12 | 0.167041 | 0.002598 | Other Positive Correlation |
| BEX1 | 0.166882 | 0.002623 | Other Positive Correlation |
| PPP2R3C | 0.166869 | 0.002625 | Other Positive Correlation |
| NUSAP1 | 0.166754 | 0.002644 | Other Positive Correlation |
| LDHC | 0.166654 | 0.00266 | Other Positive Correlation |
| IQGAP2 | 0.166627 | 0.002664 | Other Positive Correlation |
| UBE2B | 0.166491 | 0.002686 | Other Positive Correlation |
| KIRREL2 | 0.166485 | 0.002687 | Other Positive Correlation |
| TLNRD1 | 0.166455 | 0.002692 | Other Positive Correlation |
| SPNS1 | 0.16641 | 0.002699 | Other Positive Correlation |
| SERPING1 | 0.166275 | 0.002721 | Other Positive Correlation |
| HOXB9 | 0.165983 | 0.002769 | Other Positive Correlation |
| FGF13 | 0.165931 | 0.002778 | Other Positive Correlation |
| BNC2 | 0.165762 | 0.002806 | Other Positive Correlation |
| CDC25B | 0.165754 | 0.002807 | Other Positive Correlation |
| PTPRN | 0.165685 | 0.002819 | Other Positive Correlation |
| NOL4L | 0.165665 | 0.002822 | Other Positive Correlation |
| PLEKHG4B | 0.16552 | 0.002847 | Other Positive Correlation |
| PLPPR2 | 0.165484 | 0.002853 | Other Positive Correlation |
| FAM110B | 0.165444 | 0.00286 | Other Positive Correlation |
| TMEM47 | 0.165441 | 0.00286 | Other Positive Correlation |
| BMP8B | 0.165353 | 0.002875 | Other Positive Correlation |
| CA8 | 0.165314 | 0.002882 | Other Positive Correlation |
| SLC41A2 | 0.165194 | 0.002902 | Other Positive Correlation |
| MAP3K9 | 0.165175 | 0.002906 | Other Positive Correlation |
| MPC1 | 0.165109 | 0.002917 | Other Positive Correlation |
| PHKG2 | 0.165036 | 0.00293 | Other Positive Correlation |
| CRY1 | 0.164898 | 0.002954 | Other Positive Correlation |
| RETREG1 | 0.164875 | 0.002958 | Other Positive Correlation |
| ARHGEF39 | 0.164848 | 0.002963 | Other Positive Correlation |
| ADO | 0.16484 | 0.002964 | Other Positive Correlation |
| LGI2 | 0.164817 | 0.002968 | Other Positive Correlation |
| GPANK1 | 0.164784 | 0.002974 | Other Positive Correlation |
| ENSG00000286132 | 0.164738 | 0.002982 | Other Positive Correlation |
| WDR83 | 0.164574 | 0.003012 | Other Positive Correlation |
| FBXL15 | 0.1645 | 0.003025 | Other Positive Correlation |
| PLCB1 | 0.164492 | 0.003026 | Other Positive Correlation |
| ELOVL2 | 0.164373 | 0.003048 | Other Positive Correlation |

| | | | |
|---|---|---|---|
| PLEKHD1 | 0.164277 | 0.003065 | Other Positive Correlation |
| KALRN | 0.164136 | 0.003091 | Other Positive Correlation |
| NOS1AP | 0.164059 | 0.003105 | Other Positive Correlation |
| PNMA8A | 0.164012 | 0.003114 | Other Positive Correlation |
| PRR15 | 0.164005 | 0.003115 | Other Positive Correlation |
| WEE1 | 0.163877 | 0.003139 | Other Positive Correlation |
| RAI2 | 0.163811 | 0.003151 | Other Positive Correlation |
| SDS | 0.163721 | 0.003168 | Other Positive Correlation |
| RAB27A | 0.163643 | 0.003183 | Other Positive Correlation |
| DACT3 | 0.163626 | 0.003186 | Other Positive Correlation |
| TET3 | 0.163586 | 0.003193 | Other Positive Correlation |
| RASAL1 | 0.16353 | 0.003204 | Other Positive Correlation |
| SLC27A2 | 0.163189 | 0.003269 | Other Positive Correlation |
| RNPEPL1 | 0.163145 | 0.003277 | Other Positive Correlation |
| DUSP8 | 0.163103 | 0.003286 | Other Positive Correlation |
| GYPC | 0.163091 | 0.003288 | Other Positive Correlation |
| TUBE1 | 0.16278 | 0.003349 | Other Positive Correlation |
| GOLT1B | 0.162771 | 0.003351 | Other Positive Correlation |
| GALM | 0.162762 | 0.003352 | Other Positive Correlation |
| CKLF | 0.162731 | 0.003358 | Other Positive Correlation |
| PHGDH | 0.16259 | 0.003386 | Other Positive Correlation |
| NSMCE3 | 0.162519 | 0.003401 | Other Positive Correlation |
| HLA-F | 0.162487 | 0.003407 | Other Positive Correlation |
| LIN7B | 0.162382 | 0.003428 | Other Positive Correlation |
| NR1H2 | 0.16231 | 0.003443 | Other Positive Correlation |
| AVP | 0.162273 | 0.00345 | Other Positive Correlation |
| ZNF598 | 0.162204 | 0.003464 | Other Positive Correlation |
| UNC5B | 0.162134 | 0.003478 | Other Positive Correlation |
| ENTPD8 | 0.162047 | 0.003496 | Other Positive Correlation |
| CACNA2D2 | 0.161981 | 0.00351 | Other Positive Correlation |
| ATP2C2 | 0.1619 | 0.003526 | Other Positive Correlation |
| TRIOBP | 0.161754 | 0.003557 | Other Positive Correlation |
| ISYNA1 | 0.161664 | 0.003575 | Other Positive Correlation |
| PELI3 | 0.161606 | 0.003588 | Other Positive Correlation |
| CYB5A | 0.161348 | 0.003642 | Other Positive Correlation |
| CABLES1 | 0.161273 | 0.003658 | Other Positive Correlation |
| MPP2 | 0.161234 | 0.003666 | Other Positive Correlation |
| TMEM245 | 0.161217 | 0.00367 | Other Positive Correlation |
| SHC2 | 0.161198 | 0.003674 | Other Positive Correlation |
| IFI27L1 | 0.161141 | 0.003686 | Other Positive Correlation |
| RHOU | 0.16092 | 0.003734 | Other Positive Correlation |
| ZNF711 | 0.160885 | 0.003742 | Other Positive Correlation |
| NAPA | 0.160584 | 0.003808 | Other Positive Correlation |
| GPN3 | 0.160578 | 0.003809 | Other Positive Correlation |
| RAPH1 | 0.160452 | 0.003837 | Other Positive Correlation |
| RCN3 | 0.160446 | 0.003838 | Other Positive Correlation |
| TMX2 | 0.160414 | 0.003846 | Other Positive Correlation |
| ELANE | 0.160309 | 0.003869 | Other Positive Correlation |

| | | | |
|---|---|---|---|
| NXNL2 | 0.160258 | 0.003881 | Other Positive Correlation |
| VSIG10 | 0.160255 | 0.003881 | Other Positive Correlation |
| AKR7A3 | 0.160118 | 0.003912 | Other Positive Correlation |
| E2F2 | 0.160087 | 0.003919 | Other Positive Correlation |
| CKAP2 | 0.159949 | 0.003951 | Other Positive Correlation |
| CACNB4 | 0.159852 | 0.003973 | Other Positive Correlation |
| ZBTB5 | 0.159846 | 0.003974 | Other Positive Correlation |
| ZNF138 | 0.159818 | 0.003981 | Other Positive Correlation |
| SNX16 | 0.159679 | 0.004013 | Other Positive Correlation |
| OCLN | 0.15965 | 0.00402 | Other Positive Correlation |
| TMEM74B | 0.159621 | 0.004026 | Other Positive Correlation |
| MFSD6 | 0.159532 | 0.004047 | Other Positive Correlation |
| DPYSL5 | 0.159515 | 0.004051 | Other Positive Correlation |
| ADNP2 | 0.159512 | 0.004052 | Other Positive Correlation |
| FEM1A | 0.159507 | 0.004053 | Other Positive Correlation |
| PLEKHH2 | 0.159421 | 0.004073 | Other Positive Correlation |
| HMGN5 | 0.159415 | 0.004075 | Other Positive Correlation |
| MFSD9 | 0.159375 | 0.004084 | Other Positive Correlation |
| FAM171A2 | 0.159368 | 0.004086 | Other Positive Correlation |
| CABLES2 | 0.159343 | 0.004092 | Other Positive Correlation |
| JADE1 | 0.159178 | 0.004131 | Other Positive Correlation |
| JSRP1 | 0.159122 | 0.004144 | Other Positive Correlation |
| CNTFR | 0.159065 | 0.004158 | Other Positive Correlation |
| PPP1R3E | 0.158914 | 0.004194 | Other Positive Correlation |
| ARL8A | 0.158802 | 0.004221 | Other Positive Correlation |
| AP1AR | 0.158711 | 0.004243 | Other Positive Correlation |
| OXT | 0.158626 | 0.004264 | Other Positive Correlation |
| DCDC2 | 0.15857 | 0.004278 | Other Positive Correlation |
| LONRF2 | 0.158333 | 0.004337 | Other Positive Correlation |
| EMC9 | 0.158256 | 0.004356 | Other Positive Correlation |
| SMAP1 | 0.158242 | 0.004359 | Other Positive Correlation |
| TMOD1 | 0.158226 | 0.004363 | Other Positive Correlation |
| KIF3B | 0.158195 | 0.004371 | Other Positive Correlation |
| NMUR1 | 0.15809 | 0.004398 | Other Positive Correlation |
| CMTM8 | 0.158049 | 0.004408 | Other Positive Correlation |
| DERL1 | 0.158001 | 0.00442 | Other Positive Correlation |
| KHDRBS3 | 0.157969 | 0.004428 | Other Positive Correlation |
| TMEM163 | 0.157959 | 0.004431 | Other Positive Correlation |
| FAM174B | 0.157929 | 0.004438 | Other Positive Correlation |
| CGNL1 | 0.157882 | 0.00445 | Other Positive Correlation |
| ZSCAN26 | 0.157811 | 0.004468 | Other Positive Correlation |
| KCNG3 | 0.157772 | 0.004478 | Other Positive Correlation |
| CCL28 | 0.157727 | 0.00449 | Other Positive Correlation |
| ABCC6 | 0.157695 | 0.004498 | Other Positive Correlation |
| SLC16A8 | 0.157694 | 0.004498 | Other Positive Correlation |
| LSM8 | 0.157634 | 0.004514 | Other Positive Correlation |
| FCRLB | 0.157619 | 0.004518 | Other Positive Correlation |
| B3GNT9 | 0.15741 | 0.004572 | Other Positive Correlation |

| | | | |
|---|---|---|---|
| SNAP91 | 0.157283 | 0.004605 | Other Positive Correlation |
| CPEB4 | 0.157223 | 0.004621 | Other Positive Correlation |
| YEATS4 | 0.157162 | 0.004637 | Other Positive Correlation |
| NDRG4 | 0.157131 | 0.004645 | Other Positive Correlation |
| HMX2 | 0.157086 | 0.004657 | Other Positive Correlation |
| CASTOR2 | 0.156943 | 0.004695 | Other Positive Correlation |
| IDUA | 0.156918 | 0.004702 | Other Positive Correlation |
| RAD54L | 0.156869 | 0.004715 | Other Positive Correlation |
| MCF2L | 0.156806 | 0.004732 | Other Positive Correlation |
| RCE1 | 0.156777 | 0.00474 | Other Positive Correlation |
| HYCC2 | 0.156759 | 0.004745 | Other Positive Correlation |
| TMEM203 | 0.156728 | 0.004753 | Other Positive Correlation |
| MED1 | 0.156685 | 0.004765 | Other Positive Correlation |
| TINCR | 0.156671 | 0.004769 | Other Positive Correlation |
| ZRANB1 | 0.156653 | 0.004774 | Other Positive Correlation |
| DCTN3 | 0.156616 | 0.004784 | Other Positive Correlation |
| THRA | 0.156595 | 0.004789 | Other Positive Correlation |
| AK1 | 0.156538 | 0.004805 | Other Positive Correlation |
| CPE | 0.156537 | 0.004805 | Other Positive Correlation |
| FRS2 | 0.156506 | 0.004814 | Other Positive Correlation |
| MADCAM1 | 0.156478 | 0.004821 | Other Positive Correlation |
| RCN2 | 0.156424 | 0.004836 | Other Positive Correlation |
| NEXMIF | 0.156375 | 0.00485 | Other Positive Correlation |
| TSPAN2 | 0.156324 | 0.004864 | Other Positive Correlation |
| CCDC149 | 0.156323 | 0.004864 | Other Positive Correlation |
| XBP1 | 0.156315 | 0.004866 | Other Positive Correlation |
| SLC6A17 | 0.156314 | 0.004866 | Other Positive Correlation |
| LMX1B | 0.156128 | 0.004918 | Other Positive Correlation |
| KRT80 | 0.156126 | 0.004919 | Other Positive Correlation |
| NUDT16 | 0.156093 | 0.004928 | Other Positive Correlation |
| ENSG00000284337 | 0.156027 | 0.004946 | Other Positive Correlation |
| SYT3 | 0.155891 | 0.004985 | Other Positive Correlation |
| ZMAT5 | 0.155686 | 0.005043 | Other Positive Correlation |
| PLXND1 | 0.155668 | 0.005048 | Other Positive Correlation |
| SCAND1 | 0.155574 | 0.005075 | Other Positive Correlation |
| XYLT1 | 0.155556 | 0.00508 | Other Positive Correlation |
| DPM2 | 0.155442 | 0.005113 | Other Positive Correlation |
| ARHGAP31 | 0.155325 | 0.005147 | Other Positive Correlation |
| CHP2 | 0.155256 | 0.005167 | Other Positive Correlation |
| TAFA4 | 0.155118 | 0.005207 | Other Positive Correlation |
| C19orf38 | 0.155056 | 0.005225 | Other Positive Correlation |
| GIPC3 | 0.154899 | 0.005272 | Other Positive Correlation |
| SLC35A4 | 0.154881 | 0.005277 | Other Positive Correlation |
| PCBP3 | 0.154625 | 0.005354 | Other Positive Correlation |
| PRDM16 | 0.154554 | 0.005375 | Other Positive Correlation |
| ISCA2 | 0.154542 | 0.005379 | Other Positive Correlation |
| GXYLT1 | 0.154455 | 0.005405 | Other Positive Correlation |
| MFAP3L | 0.154436 | 0.005411 | Other Positive Correlation |

| | | | |
|---|---|---|---|
| SYN2 | 0.154306 | 0.005451 | Other Positive Correlation |
| ANO5 | 0.154163 | 0.005494 | Other Positive Correlation |
| LOXL4 | 0.154156 | 0.005497 | Other Positive Correlation |
| PPM1B | 0.154096 | 0.005515 | Other Positive Correlation |
| UTF1 | 0.154008 | 0.005542 | Other Positive Correlation |
| PIGQ | 0.153987 | 0.005549 | Other Positive Correlation |
| TRPV3 | 0.153894 | 0.005578 | Other Positive Correlation |
| ATP6V0A2 | 0.153808 | 0.005605 | Other Positive Correlation |
| NGF | 0.153724 | 0.005631 | Other Positive Correlation |
| FMC1 | 0.153563 | 0.005682 | Other Positive Correlation |
| ATP6V0D1 | 0.153508 | 0.005699 | Other Positive Correlation |
| PEA15 | 0.153502 | 0.005701 | Other Positive Correlation |
| FBP1 | 0.153315 | 0.005761 | Other Positive Correlation |
| HLA-A | 0.153297 | 0.005767 | Other Positive Correlation |
| HAT1 | 0.153191 | 0.005801 | Other Positive Correlation |
| SESN2 | 0.152739 | 0.005949 | Other Positive Correlation |
| AGAP1 | 0.152657 | 0.005976 | Other Positive Correlation |
| ADCY9 | 0.152618 | 0.005989 | Other Positive Correlation |
| TMEM38B | 0.152613 | 0.005991 | Other Positive Correlation |
| EMILIN2 | 0.152544 | 0.006014 | Other Positive Correlation |
| SLC35B1 | 0.152463 | 0.006041 | Other Positive Correlation |
| STAC | 0.152375 | 0.006071 | Other Positive Correlation |
| EPOP | 0.152373 | 0.006071 | Other Positive Correlation |
| DPF1 | 0.152363 | 0.006075 | Other Positive Correlation |
| ZNF726 | 0.152331 | 0.006086 | Other Positive Correlation |
| CGRRF1 | 0.152288 | 0.0061 | Other Positive Correlation |
| TMEM225B | 0.152237 | 0.006117 | Other Positive Correlation |
| MSANTD4 | 0.152123 | 0.006156 | Other Positive Correlation |
| TCEA3 | 0.152094 | 0.006166 | Other Positive Correlation |
| ILDR2 | 0.152078 | 0.006172 | Other Positive Correlation |
| PIK3CB | 0.15199 | 0.006202 | Other Positive Correlation |
| ACSS1 | 0.151777 | 0.006275 | Other Positive Correlation |
| MPC2 | 0.151719 | 0.006295 | Other Positive Correlation |
| GNA11 | 0.151715 | 0.006297 | Other Positive Correlation |
| MRPL41 | 0.151587 | 0.006342 | Other Positive Correlation |
| KIF5C | 0.151532 | 0.006361 | Other Positive Correlation |
| LRRC10B | 0.151512 | 0.006368 | Other Positive Correlation |
| ADAMTS17 | 0.151409 | 0.006404 | Other Positive Correlation |
| PNMT | 0.15134 | 0.006429 | Other Positive Correlation |
| XKR8 | 0.151262 | 0.006456 | Other Positive Correlation |
| GFRA1 | 0.151226 | 0.006469 | Other Positive Correlation |
| COLGALT2 | 0.151149 | 0.006497 | Other Positive Correlation |
| CDK11A | 0.15107 | 0.006525 | Other Positive Correlation |
| RBPJ | 0.151017 | 0.006544 | Other Positive Correlation |
| RAB39A | 0.151 | 0.00655 | Other Positive Correlation |
| KDM2A | 0.15095 | 0.006568 | Other Positive Correlation |
| MRPL49 | 0.150938 | 0.006573 | Other Positive Correlation |
| FOXJ2 | 0.150867 | 0.006598 | Other Positive Correlation |

| | | | |
|---|---|---|---|
| TMEM59 | 0.150857 | 0.006602 | Other Positive Correlation |
| CHRNA3 | 0.150846 | 0.006606 | Other Positive Correlation |
| PINK1 | 0.15084 | 0.006608 | Other Positive Correlation |
| FGF19 | 0.150752 | 0.00664 | Other Positive Correlation |
| FAM167A | 0.150729 | 0.006649 | Other Positive Correlation |
| EIF4E3 | 0.150719 | 0.006652 | Other Positive Correlation |
| SDK1 | 0.150681 | 0.006666 | Other Positive Correlation |
| L1CAM | 0.150599 | 0.006696 | Other Positive Correlation |
| SFMBT1 | 0.150551 | 0.006714 | Other Positive Correlation |
| GRB7 | 0.150451 | 0.006751 | Other Positive Correlation |
| CENPP | 0.15045 | 0.006751 | Other Positive Correlation |
| SMC1B | 0.150389 | 0.006774 | Other Positive Correlation |
| TDRD9 | 0.150361 | 0.006784 | Other Positive Correlation |
| TAL1 | 0.150359 | 0.006785 | Other Positive Correlation |
| DMBX1 | 0.150262 | 0.006821 | Other Positive Correlation |
| CCDC110 | 0.150199 | 0.006845 | Other Positive Correlation |
| FBXL3 | 0.150164 | 0.006858 | Other Positive Correlation |
| ORAI1 | 0.150127 | 0.006872 | Other Positive Correlation |
| SEZ6L2 | 0.150107 | 0.00688 | Other Positive Correlation |
| C11orf96 | 0.150085 | 0.006888 | Other Positive Correlation |
| NKIRAS2 | 0.150015 | 0.006914 | Other Positive Correlation |
| GAB2 | 0.149997 | 0.006921 | Other Positive Correlation |
| NKX2-3 | 0.149922 | 0.00695 | Other Positive Correlation |
| OTUD7B | 0.149916 | 0.006952 | Other Positive Correlation |
| HOXA10 | 0.149868 | 0.00697 | Other Positive Correlation |
| SKOR1 | 0.149671 | 0.007045 | Other Positive Correlation |
| CCDC69 | 0.149647 | 0.007055 | Other Positive Correlation |
| SLC52A2 | 0.14964 | 0.007057 | Other Positive Correlation |
| SPTLC2 | 0.149623 | 0.007064 | Other Positive Correlation |
| PIP5K1B | 0.149414 | 0.007145 | Other Positive Correlation |
| UCK1 | 0.149376 | 0.00716 | Other Positive Correlation |
| CDC25C | 0.149372 | 0.007162 | Other Positive Correlation |
| TMEM268 | 0.149296 | 0.007191 | Other Positive Correlation |
| FBXO45 | 0.149286 | 0.007195 | Other Positive Correlation |
| SYTL2 | 0.149262 | 0.007204 | Other Positive Correlation |
| LCORL | 0.149253 | 0.007208 | Other Positive Correlation |
| SLC39A3 | 0.149247 | 0.007211 | Other Positive Correlation |
| MED12L | 0.149152 | 0.007248 | Other Positive Correlation |
| SCYL3 | 0.149042 | 0.007291 | Other Positive Correlation |
| UBE2J1 | 0.148939 | 0.007333 | Other Positive Correlation |
| TDRD5 | 0.14893 | 0.007336 | Other Positive Correlation |
| DOC2A | 0.148929 | 0.007336 | Other Positive Correlation |
| TMEM176B | 0.148921 | 0.00734 | Other Positive Correlation |
| PGF | 0.148879 | 0.007356 | Other Positive Correlation |
| RPAIN | 0.148872 | 0.007359 | Other Positive Correlation |
| MPZL3 | 0.148827 | 0.007377 | Other Positive Correlation |
| CALY | 0.148813 | 0.007383 | Other Positive Correlation |
| NEO1 | 0.148787 | 0.007393 | Other Positive Correlation |

| | | | |
|---|---|---|---|
| ADCYAP1 | 0.148756 | 0.007406 | Other Positive Correlation |
| TCEA2 | 0.148746 | 0.00741 | Other Positive Correlation |
| CYP2R1 | 0.148724 | 0.007419 | Other Positive Correlation |
| NEURL1 | 0.148677 | 0.007438 | Other Positive Correlation |
| FREM2 | 0.148672 | 0.00744 | Other Positive Correlation |
| ATP2A3 | 0.148601 | 0.007469 | Other Positive Correlation |
| PTH2 | 0.14856 | 0.007485 | Other Positive Correlation |
| FASTKD3 | 0.148535 | 0.007495 | Other Positive Correlation |
| GCC1 | 0.148497 | 0.007511 | Other Positive Correlation |
| SNX24 | 0.148475 | 0.00752 | Other Positive Correlation |
| N4BP2L1 | 0.148465 | 0.007524 | Other Positive Correlation |
| SUZ12 | 0.148417 | 0.007543 | Other Positive Correlation |
| NPY | 0.148413 | 0.007545 | Other Positive Correlation |
| CRIP1 | 0.148286 | 0.007597 | Other Positive Correlation |
| C1orf115 | 0.148286 | 0.007597 | Other Positive Correlation |
| RHNO1 | 0.148254 | 0.00761 | Other Positive Correlation |
| SHISAL1 | 0.148069 | 0.007687 | Other Positive Correlation |
| CLIC5 | 0.148042 | 0.007698 | Other Positive Correlation |
| WWOX | 0.148023 | 0.007706 | Other Positive Correlation |
| INCENP | 0.147935 | 0.007743 | Other Positive Correlation |
| POLD3 | 0.147927 | 0.007746 | Other Positive Correlation |
| ELMOD1 | 0.147779 | 0.007808 | Other Positive Correlation |
| ZNF583 | 0.147735 | 0.007827 | Other Positive Correlation |
| ENTPD4 | 0.147634 | 0.00787 | Other Positive Correlation |
| SLFN13 | 0.14751 | 0.007923 | Other Positive Correlation |
| DSCAML1 | 0.147507 | 0.007924 | Other Positive Correlation |
| CACNA1H | 0.147501 | 0.007927 | Other Positive Correlation |
| RFLNB | 0.1475 | 0.007927 | Other Positive Correlation |
| DCAF5 | 0.147411 | 0.007965 | Other Positive Correlation |
| RNFT2 | 0.147379 | 0.007979 | Other Positive Correlation |
| SH3BP2 | 0.147295 | 0.008015 | Other Positive Correlation |
| NATD1 | 0.147179 | 0.008065 | Other Positive Correlation |
| TOPBP1 | 0.147123 | 0.00809 | Other Positive Correlation |
| COA5 | 0.147122 | 0.00809 | Other Positive Correlation |
| EID2 | 0.146993 | 0.008147 | Other Positive Correlation |
| MAPK8IP1 | 0.146986 | 0.008149 | Other Positive Correlation |
| UBE2G1 | 0.146934 | 0.008173 | Other Positive Correlation |
| PGM2L1 | 0.146821 | 0.008222 | Other Positive Correlation |
| ITGAE | 0.146817 | 0.008224 | Other Positive Correlation |
| SCNN1A | 0.146712 | 0.00827 | Other Positive Correlation |
| CMC2 | 0.146692 | 0.008279 | Other Positive Correlation |
| SNAP29 | 0.146653 | 0.008297 | Other Positive Correlation |
| IGHMBP2 | 0.146609 | 0.008316 | Other Positive Correlation |
| DNAJB1 | 0.146597 | 0.008322 | Other Positive Correlation |
| RGCC | 0.14651 | 0.008361 | Other Positive Correlation |
| TMX4 | 0.146487 | 0.008371 | Other Positive Correlation |
| SCUBE2 | 0.146431 | 0.008396 | Other Positive Correlation |
| FAM102A | 0.146394 | 0.008413 | Other Positive Correlation |

| | | | |
|---|---|---|---|
| WSCD1 | 0.146357 | 0.008429 | Other Positive Correlation |
| GLIPR2 | 0.146291 | 0.008459 | Other Positive Correlation |
| TMEM240 | 0.146278 | 0.008466 | Other Positive Correlation |
| ANXA6 | 0.146263 | 0.008472 | Other Positive Correlation |
| LIN28B | 0.146233 | 0.008486 | Other Positive Correlation |
| C1orf52 | 0.146215 | 0.008494 | Other Positive Correlation |
| NAGA | 0.146077 | 0.008557 | Other Positive Correlation |
| DBNDD1 | 0.146025 | 0.008581 | Other Positive Correlation |
| CLU | 0.145962 | 0.00861 | Other Positive Correlation |
| AMIGO1 | 0.145946 | 0.008617 | Other Positive Correlation |
| ZNF76 | 0.145929 | 0.008625 | Other Positive Correlation |
| PHF19 | 0.145894 | 0.008641 | Other Positive Correlation |
| CMTM4 | 0.145862 | 0.008656 | Other Positive Correlation |
| CHRD | 0.145795 | 0.008687 | Other Positive Correlation |
| CHRM3 | 0.14579 | 0.008689 | Other Positive Correlation |
| PGAP4 | 0.145764 | 0.008701 | Other Positive Correlation |
| KCNJ4 | 0.145731 | 0.008717 | Other Positive Correlation |
| GRPEL2 | 0.145672 | 0.008744 | Other Positive Correlation |
| GLRB | 0.145527 | 0.008812 | Other Positive Correlation |
| PPP1R9B | 0.145492 | 0.008828 | Other Positive Correlation |
| NMNAT3 | 0.145442 | 0.008852 | Other Positive Correlation |
| CCN3 | 0.145424 | 0.008861 | Other Positive Correlation |
| MAPT | 0.14542 | 0.008862 | Other Positive Correlation |
| SUV39H2 | 0.145388 | 0.008878 | Other Positive Correlation |
| SLC25A35 | 0.145354 | 0.008894 | Other Positive Correlation |
| C2orf69 | 0.14535 | 0.008896 | Other Positive Correlation |
| FOXRED1 | 0.145295 | 0.008922 | Other Positive Correlation |
| GABRB3 | 0.145283 | 0.008927 | Other Positive Correlation |
| CCDC106 | 0.145259 | 0.008939 | Other Positive Correlation |
| SERP1 | 0.145217 | 0.008959 | Other Positive Correlation |
| FIBIN | 0.145213 | 0.008961 | Other Positive Correlation |
| ATP8A2 | 0.145191 | 0.008971 | Other Positive Correlation |
| CPTP | 0.145148 | 0.008992 | Other Positive Correlation |
| TACC1 | 0.145125 | 0.009003 | Other Positive Correlation |
| KBTBD2 | 0.145077 | 0.009026 | Other Positive Correlation |
| FADS6 | 0.144773 | 0.009173 | Other Positive Correlation |
| PTCHD1 | 0.144749 | 0.009184 | Other Positive Correlation |
| ELOVL3 | 0.14473 | 0.009194 | Other Positive Correlation |
| CTNS | 0.144726 | 0.009196 | Other Positive Correlation |
| ZNRF3 | 0.144667 | 0.009225 | Other Positive Correlation |
| PLAAT1 | 0.144548 | 0.009283 | Other Positive Correlation |
| MIS18BP1 | 0.144435 | 0.009339 | Other Positive Correlation |
| LIN52 | 0.144385 | 0.009364 | Other Positive Correlation |
| PDZD7 | 0.14434 | 0.009386 | Other Positive Correlation |
| CRMP1 | 0.144321 | 0.009395 | Other Positive Correlation |
| YY1AP1 | 0.144222 | 0.009445 | Other Positive Correlation |
| PRR14 | 0.144178 | 0.009467 | Other Positive Correlation |
| DYNC1LI1 | 0.144146 | 0.009483 | Other Positive Correlation |

| | | | |
|---|---|---|---|
| FGF22 | 0.144121 | 0.009496 | Other Positive Correlation |
| GLMP | 0.14405 | 0.009531 | Other Positive Correlation |
| DMAP1 | 0.144012 | 0.00955 | Other Positive Correlation |
| REPIN1 | 0.143971 | 0.009571 | Other Positive Correlation |
| WIZ | 0.143918 | 0.009598 | Other Positive Correlation |
| CXADR | 0.143888 | 0.009613 | Other Positive Correlation |
| ZBTB43 | 0.143887 | 0.009614 | Other Positive Correlation |
| PLA2G10 | 0.143877 | 0.009619 | Other Positive Correlation |
| TSC22D3 | 0.143875 | 0.00962 | Other Positive Correlation |
| SLC32A1 | 0.143799 | 0.009658 | Other Positive Correlation |
| ERC2 | 0.143787 | 0.009665 | Other Positive Correlation |
| ATXN1 | 0.143779 | 0.009668 | Other Positive Correlation |
| DDIT4L | 0.14377 | 0.009673 | Other Positive Correlation |
| PLD3 | 0.14376 | 0.009678 | Other Positive Correlation |
| ZNF805 | 0.143737 | 0.00969 | Other Positive Correlation |
| DPP10 | 0.143659 | 0.00973 | Other Positive Correlation |
| CDKN1B | 0.143639 | 0.00974 | Other Positive Correlation |
| CLDN10 | 0.143626 | 0.009747 | Other Positive Correlation |
| FAM161A | 0.143589 | 0.009766 | Other Positive Correlation |
| GNPTG | 0.143487 | 0.009818 | Other Positive Correlation |
| SLC2A8 | 0.143452 | 0.009836 | Other Positive Correlation |
| CCSER1 | 0.143428 | 0.009849 | Other Positive Correlation |
| PARM1 | 0.143399 | 0.009864 | Other Positive Correlation |
| FAM174C | 0.143234 | 0.00995 | Other Positive Correlation |
| TEX261 | 0.14305 | 0.010047 | Other Positive Correlation |
| FASTK | 0.143038 | 0.010053 | Other Positive Correlation |
| DNAJA4 | 0.142993 | 0.010077 | Other Positive Correlation |
| DHX29 | 0.142986 | 0.010081 | Other Positive Correlation |
| SLC25A44 | 0.142974 | 0.010087 | Other Positive Correlation |
| ENSG00000285920 | 0.142973 | 0.010088 | Other Positive Correlation |
| DOCK11 | 0.142941 | 0.010105 | Other Positive Correlation |
| PTPN12 | 0.14291 | 0.010121 | Other Positive Correlation |
| ATP8B2 | 0.142909 | 0.010122 | Other Positive Correlation |
| AR | 0.142866 | 0.010145 | Other Positive Correlation |
| GAS7 | 0.142829 | 0.010164 | Other Positive Correlation |
| ARHGAP20 | 0.142775 | 0.010193 | Other Positive Correlation |
| TP53INP2 | 0.142755 | 0.010204 | Other Positive Correlation |
| TOX3 | 0.142744 | 0.01021 | Other Positive Correlation |
| KCTD6 | 0.142646 | 0.010262 | Other Positive Correlation |
| CDON | 0.142611 | 0.010281 | Other Positive Correlation |
| ZNF576 | 0.142542 | 0.010318 | Other Positive Correlation |
| TTC28 | 0.142446 | 0.01037 | Other Positive Correlation |
| GCK | 0.142279 | 0.010461 | Other Positive Correlation |
| MYLIP | 0.142212 | 0.010498 | Other Positive Correlation |
| VPS18 | 0.142196 | 0.010507 | Other Positive Correlation |
| TMEM179 | 0.142148 | 0.010533 | Other Positive Correlation |
| TMEM176A | 0.142136 | 0.01054 | Other Positive Correlation |
| BDH1 | 0.142065 | 0.010579 | Other Positive Correlation |

| | | | |
|---|---|---|---|
| FAAP24 | 0.141987 | 0.010622 | Other Positive Correlation |
| C16orf87 | 0.141964 | 0.010635 | Other Positive Correlation |
| BLTP3A | 0.141898 | 0.010672 | Other Positive Correlation |
| GALR3 | 0.141879 | 0.010682 | Other Positive Correlation |
| DPYD | 0.141746 | 0.010757 | Other Positive Correlation |
| ROR2 | 0.141737 | 0.010762 | Other Positive Correlation |
| FAM32A | 0.141701 | 0.010782 | Other Positive Correlation |
| DAZAP2 | 0.141661 | 0.010804 | Other Positive Correlation |
| SLC15A1 | 0.141529 | 0.010879 | Other Positive Correlation |
| KLHDC9 | 0.141464 | 0.010916 | Other Positive Correlation |
| ZNF714 | 0.141454 | 0.010922 | Other Positive Correlation |
| LMF2 | 0.141452 | 0.010923 | Other Positive Correlation |
| SLC7A8 | 0.141352 | 0.01098 | Other Positive Correlation |
| FXYD7 | 0.141262 | 0.011032 | Other Positive Correlation |
| CDC42BPA | 0.141231 | 0.011049 | Other Positive Correlation |
| WBP2 | 0.141164 | 0.011088 | Other Positive Correlation |
| EZH2 | 0.141055 | 0.01115 | Other Positive Correlation |
| SMG5 | 0.141014 | 0.011175 | Other Positive Correlation |
| ERAS | 0.141003 | 0.011181 | Other Positive Correlation |
| CCZ1B | 0.141002 | 0.011182 | Other Positive Correlation |
| SAP30L | 0.140972 | 0.011199 | Other Positive Correlation |
| COL11A2 | 0.140881 | 0.011252 | Other Positive Correlation |
| SCRN1 | 0.140699 | 0.011359 | Other Positive Correlation |
| INTS5 | 0.140678 | 0.011371 | Other Positive Correlation |
| PENK | 0.14062 | 0.011405 | Other Positive Correlation |
| ZNF704 | 0.140421 | 0.011523 | Other Positive Correlation |
| MUC1 | 0.14042 | 0.011524 | Other Positive Correlation |
| KIAA1549 | 0.140364 | 0.011557 | Other Positive Correlation |
| FECH | 0.140309 | 0.01159 | Other Positive Correlation |
| CRTAC1 | 0.140076 | 0.01173 | Other Positive Correlation |
| CDK5RAP2 | 0.139953 | 0.011805 | Other Positive Correlation |
| DYNLL2 | 0.13994 | 0.011813 | Other Positive Correlation |
| FAM219B | 0.139849 | 0.011869 | Other Positive Correlation |
| PIERCE1 | 0.139843 | 0.011873 | Other Positive Correlation |
| BMP6 | 0.139835 | 0.011877 | Other Positive Correlation |
| CCDC174 | 0.139745 | 0.011933 | Other Positive Correlation |
| CHMP4C | 0.139727 | 0.011943 | Other Positive Correlation |
| MRPS12 | 0.13971 | 0.011954 | Other Positive Correlation |
| SHOX2 | 0.13969 | 0.011966 | Other Positive Correlation |
| CCDC160 | 0.139581 | 0.012034 | Other Positive Correlation |
| PAK4 | 0.139518 | 0.012072 | Other Positive Correlation |
| CFAP119 | 0.139502 | 0.012083 | Other Positive Correlation |
| CLCC1 | 0.139491 | 0.012089 | Other Positive Correlation |
| ACOT4 | 0.139467 | 0.012105 | Other Positive Correlation |
| COPZ2 | 0.139381 | 0.012158 | Other Positive Correlation |
| F12 | 0.139345 | 0.012181 | Other Positive Correlation |
| FBXO34 | 0.139257 | 0.012236 | Other Positive Correlation |
| ZFP41 | 0.139254 | 0.012237 | Other Positive Correlation |

| | | | |
|---|---|---|---|
| FBXO28 | 0.139249 | 0.012241 | Other Positive Correlation |
| FAM3B | 0.139232 | 0.012251 | Other Positive Correlation |
| CCNA1 | 0.139053 | 0.012364 | Other Positive Correlation |
| GSTM4 | 0.139053 | 0.012364 | Other Positive Correlation |
| ALDH1A2 | 0.139046 | 0.012369 | Other Positive Correlation |
| TNRC6A | 0.139025 | 0.012382 | Other Positive Correlation |
| RB1CC1 | 0.138935 | 0.01244 | Other Positive Correlation |
| LRRC2 | 0.138935 | 0.01244 | Other Positive Correlation |
| EPB41L3 | 0.138878 | 0.012476 | Other Positive Correlation |
| ANKRD39 | 0.13885 | 0.012494 | Other Positive Correlation |
| MYL9 | 0.138794 | 0.01253 | Other Positive Correlation |
| ZNF695 | 0.138621 | 0.012642 | Other Positive Correlation |
| KCNH3 | 0.138579 | 0.012669 | Other Positive Correlation |
| PPM1N | 0.138527 | 0.012702 | Other Positive Correlation |
| GFPT2 | 0.138511 | 0.012713 | Other Positive Correlation |
| RTBDN | 0.138492 | 0.012725 | Other Positive Correlation |
| EPHX3 | 0.138466 | 0.012742 | Other Positive Correlation |
| PXDC1 | 0.138382 | 0.012797 | Other Positive Correlation |
| KBTBD11 | 0.138362 | 0.01281 | Other Positive Correlation |
| ZNF391 | 0.138362 | 0.01281 | Other Positive Correlation |
| GLA | 0.138322 | 0.012836 | Other Positive Correlation |
| GARS1 | 0.138285 | 0.01286 | Other Positive Correlation |
| WLS | 0.138253 | 0.012881 | Other Positive Correlation |
| BEGAIN | 0.138246 | 0.012886 | Other Positive Correlation |
| YJEFN3 | 0.138204 | 0.012913 | Other Positive Correlation |
| ZNF492 | 0.138141 | 0.012955 | Other Positive Correlation |
| RNF24 | 0.138052 | 0.013014 | Other Positive Correlation |
| PHOSPHO1 | 0.138031 | 0.013028 | Other Positive Correlation |
| DMRT2 | 0.138017 | 0.013037 | Other Positive Correlation |
| C3orf38 | 0.137849 | 0.013149 | Other Positive Correlation |
| ZSCAN16 | 0.137671 | 0.013269 | Other Positive Correlation |
| PRTN3 | 0.137599 | 0.013318 | Other Positive Correlation |
| NKX2-5 | 0.137576 | 0.013333 | Other Positive Correlation |
| TAB2 | 0.137568 | 0.013339 | Other Positive Correlation |
| MMP11 | 0.137558 | 0.013345 | Other Positive Correlation |
| NUP50 | 0.137532 | 0.013363 | Other Positive Correlation |
| WDSUB1 | 0.137494 | 0.013389 | Other Positive Correlation |
| COL9A3 | 0.137484 | 0.013396 | Other Positive Correlation |
| CTSD | 0.137482 | 0.013398 | Other Positive Correlation |
| ATG13 | 0.137355 | 0.013484 | Other Positive Correlation |
| VMAC | 0.137287 | 0.013531 | Other Positive Correlation |
| LRFN1 | 0.13728 | 0.013535 | Other Positive Correlation |
| UBALD2 | 0.137257 | 0.013552 | Other Positive Correlation |
| DOCK3 | 0.137248 | 0.013557 | Other Positive Correlation |
| DCLRE1B | 0.137212 | 0.013582 | Other Positive Correlation |
| POPDC3 | 0.137175 | 0.013608 | Other Positive Correlation |
| SLC43A3 | 0.137152 | 0.013623 | Other Positive Correlation |
| LYSMD1 | 0.137093 | 0.013665 | Other Positive Correlation |

| | | | |
|---|---|---|---|
| GATA4 | 0.137087 | 0.013669 | Other Positive Correlation |
| RETREG2 | 0.137075 | 0.013677 | Other Positive Correlation |
| HEY2 | 0.13707 | 0.01368 | Other Positive Correlation |
| CGGBP1 | 0.137055 | 0.013691 | Other Positive Correlation |
| ZIK1 | 0.136987 | 0.013738 | Other Positive Correlation |
| PARD6B | 0.13697 | 0.01375 | Other Positive Correlation |
| ZNF318 | 0.136918 | 0.013786 | Other Positive Correlation |
| MID2 | 0.136861 | 0.013826 | Other Positive Correlation |
| C18orf25 | 0.136839 | 0.013841 | Other Positive Correlation |
| FAM120AOS | 0.136742 | 0.01391 | Other Positive Correlation |
| KIF3C | 0.136726 | 0.013921 | Other Positive Correlation |
| LARGE1 | 0.136676 | 0.013956 | Other Positive Correlation |
| SLC4A2 | 0.13663 | 0.013988 | Other Positive Correlation |
| SLC22A5 | 0.136563 | 0.014036 | Other Positive Correlation |
| ZNFX1 | 0.136554 | 0.014042 | Other Positive Correlation |
| WASH6P | 0.136512 | 0.014072 | Other Positive Correlation |
| PKIB | 0.136466 | 0.014105 | Other Positive Correlation |
| PBX4 | 0.136463 | 0.014107 | Other Positive Correlation |
| SLC7A4 | 0.136461 | 0.014109 | Other Positive Correlation |
| TNIK | 0.136396 | 0.014155 | Other Positive Correlation |
| MT1F | 0.136359 | 0.014181 | Other Positive Correlation |
| DONSON | 0.136331 | 0.014201 | Other Positive Correlation |
| KRBOX5 | 0.136304 | 0.014221 | Other Positive Correlation |
| FOXI2 | 0.136247 | 0.014262 | Other Positive Correlation |
| ANKRD44 | 0.136212 | 0.014287 | Other Positive Correlation |
| TRPS1 | 0.136187 | 0.014305 | Other Positive Correlation |
| CHST8 | 0.136165 | 0.014321 | Other Positive Correlation |
| SH3GL2 | 0.136141 | 0.014338 | Other Positive Correlation |
| PMM2 | 0.136125 | 0.014349 | Other Positive Correlation |
| LY6G5C | 0.136121 | 0.014353 | Other Positive Correlation |
| MBTPS1 | 0.135988 | 0.014449 | Other Positive Correlation |
| BCL2L11 | 0.135919 | 0.014499 | Other Positive Correlation |
| CELF4 | 0.135865 | 0.014539 | Other Positive Correlation |
| ST8SIA2 | 0.135824 | 0.014569 | Other Positive Correlation |
| RBMXL2 | 0.135817 | 0.014574 | Other Positive Correlation |
| RHPN2 | 0.135789 | 0.014595 | Other Positive Correlation |
| FANCE | 0.135741 | 0.014629 | Other Positive Correlation |
| SOWAHA | 0.135741 | 0.01463 | Other Positive Correlation |
| CTSL | 0.135713 | 0.01465 | Other Positive Correlation |
| FNDC5 | 0.135606 | 0.014729 | Other Positive Correlation |
| SHMT1 | 0.135545 | 0.014775 | Other Positive Correlation |
| BRPF3 | 0.135543 | 0.014776 | Other Positive Correlation |
| RBFOX3 | 0.13553 | 0.014786 | Other Positive Correlation |
| PODXL | 0.135495 | 0.014812 | Other Positive Correlation |
| ZNF273 | 0.135457 | 0.01484 | Other Positive Correlation |
| HSPA2 | 0.135319 | 0.014943 | Other Positive Correlation |
| TMEM200B | 0.135302 | 0.014956 | Other Positive Correlation |
| PIGM | 0.135141 | 0.015077 | Other Positive Correlation |

| | | | |
|---|---|---|---|
| BLOC1S3 | 0.135092 | 0.015114 | Other Positive Correlation |
| KRT10 | 0.135027 | 0.015163 | Other Positive Correlation |
| PRTFDC1 | 0.134887 | 0.01527 | Other Positive Correlation |
| GPR137C | 0.134874 | 0.01528 | Other Positive Correlation |
| DACT1 | 0.134831 | 0.015313 | Other Positive Correlation |
| SELENOV | 0.134829 | 0.015314 | Other Positive Correlation |
| JPH3 | 0.134766 | 0.015362 | Other Positive Correlation |
| SEC61A2 | 0.134754 | 0.015372 | Other Positive Correlation |
| DLX3 | 0.134749 | 0.015376 | Other Positive Correlation |
| EDC4 | 0.134693 | 0.015419 | Other Positive Correlation |
| TBPL1 | 0.134652 | 0.015451 | Other Positive Correlation |
| KLF2 | 0.134608 | 0.015485 | Other Positive Correlation |
| TEAD3 | 0.13453 | 0.015544 | Other Positive Correlation |
| PDZD4 | 0.134522 | 0.015551 | Other Positive Correlation |
| LCN2 | 0.134486 | 0.015579 | Other Positive Correlation |
| EMX2 | 0.134447 | 0.015609 | Other Positive Correlation |
| RBBP5 | 0.134447 | 0.015609 | Other Positive Correlation |
| SMIM10 | 0.134309 | 0.015717 | Other Positive Correlation |
| PTF1A | 0.134255 | 0.01576 | Other Positive Correlation |
| SLC66A1 | 0.134042 | 0.015927 | Other Positive Correlation |
| FBXW11 | 0.134015 | 0.015949 | Other Positive Correlation |
| TUB | 0.133861 | 0.016071 | Other Positive Correlation |
| KCNIP3 | 0.133821 | 0.016103 | Other Positive Correlation |
| CHRM4 | 0.133798 | 0.016122 | Other Positive Correlation |
| SIRT7 | 0.133787 | 0.016131 | Other Positive Correlation |
| U2AF1L4 | 0.13377 | 0.016144 | Other Positive Correlation |
| LYRM1 | 0.133691 | 0.016207 | Other Positive Correlation |
| CHD5 | 0.13366 | 0.016233 | Other Positive Correlation |
| GAD1 | 0.133642 | 0.016247 | Other Positive Correlation |
| GOLT1A | 0.133607 | 0.016276 | Other Positive Correlation |
| MMP15 | 0.133578 | 0.016299 | Other Positive Correlation |
| SCN3B | 0.133567 | 0.016308 | Other Positive Correlation |
| FMNL3 | 0.133476 | 0.016381 | Other Positive Correlation |
| SLC25A48 | 0.133402 | 0.016441 | Other Positive Correlation |
| BBC3 | 0.133322 | 0.016506 | Other Positive Correlation |
| OSBPL6 | 0.133283 | 0.016539 | Other Positive Correlation |
| PNMA6A | 0.133273 | 0.016547 | Other Positive Correlation |
| LGI3 | 0.13327 | 0.01655 | Other Positive Correlation |
| ATP6AP1 | 0.133255 | 0.016561 | Other Positive Correlation |
| PLEKHA3 | 0.133247 | 0.016568 | Other Positive Correlation |
| S100PBP | 0.133237 | 0.016576 | Other Positive Correlation |
| RAB11FIP4 | 0.133182 | 0.016622 | Other Positive Correlation |
| SRP54 | 0.133152 | 0.016647 | Other Positive Correlation |
| CHPF | 0.133112 | 0.016679 | Other Positive Correlation |
| ERFE | 0.133091 | 0.016696 | Other Positive Correlation |
| ECSIT | 0.133051 | 0.01673 | Other Positive Correlation |
| FBLN2 | 0.133032 | 0.016745 | Other Positive Correlation |
| NBEA | 0.132959 | 0.016806 | Other Positive Correlation |

| | | | |
|---|---|---|---|
| EGLN3 | 0.132939 | 0.016823 | Other Positive Correlation |
| TUBD1 | 0.132912 | 0.016845 | Other Positive Correlation |
| GATM | 0.132895 | 0.016859 | Other Positive Correlation |
| GPX4 | 0.132874 | 0.016877 | Other Positive Correlation |
| GCA | 0.132803 | 0.016936 | Other Positive Correlation |
| CHRDL2 | 0.13278 | 0.016955 | Other Positive Correlation |
| SPR | 0.132707 | 0.017016 | Other Positive Correlation |
| ERBB4 | 0.132704 | 0.017019 | Other Positive Correlation |
| OTUD3 | 0.132675 | 0.017043 | Other Positive Correlation |
| C2orf50 | 0.132616 | 0.017093 | Other Positive Correlation |
| SLC35F1 | 0.132595 | 0.01711 | Other Positive Correlation |
| FRRS1L | 0.132567 | 0.017134 | Other Positive Correlation |
| HLA-G | 0.13256 | 0.01714 | Other Positive Correlation |
| INPP5K | 0.132515 | 0.017178 | Other Positive Correlation |
| KCNS3 | 0.13251 | 0.017183 | Other Positive Correlation |
| EBF4 | 0.132505 | 0.017187 | Other Positive Correlation |
| GSE1 | 0.132497 | 0.017193 | Other Positive Correlation |
| RTN3 | 0.132467 | 0.017218 | Other Positive Correlation |
| ARX | 0.132458 | 0.017226 | Other Positive Correlation |
| NECAB2 | 0.132445 | 0.017238 | Other Positive Correlation |
| SIX3 | 0.132313 | 0.01735 | Other Positive Correlation |
| MARCHF9 | 0.132311 | 0.017352 | Other Positive Correlation |
| PTAR1 | 0.132286 | 0.017373 | Other Positive Correlation |
| HOMER2 | 0.132247 | 0.017407 | Other Positive Correlation |
| PRKAR2B | 0.132244 | 0.017409 | Other Positive Correlation |
| ZNF774 | 0.132221 | 0.017429 | Other Positive Correlation |
| MOAP1 | 0.132218 | 0.017431 | Other Positive Correlation |
| MATN3 | 0.132132 | 0.017505 | Other Positive Correlation |
| CCDC136 | 0.132109 | 0.017525 | Other Positive Correlation |
| TNNC1 | 0.132056 | 0.017571 | Other Positive Correlation |
| PPCDC | 0.132045 | 0.01758 | Other Positive Correlation |
| SPINK2 | 0.132016 | 0.017606 | Other Positive Correlation |
| MARVELD3 | 0.131934 | 0.017676 | Other Positive Correlation |
| TRIM71 | 0.131922 | 0.017687 | Other Positive Correlation |
| PWWP2A | 0.131917 | 0.017691 | Other Positive Correlation |
| WDR53 | 0.13173 | 0.017854 | Other Positive Correlation |
| ZNF593 | 0.131676 | 0.017901 | Other Positive Correlation |
| ARRB1 | 0.131669 | 0.017908 | Other Positive Correlation |
| BTNL9 | 0.131595 | 0.017972 | Other Positive Correlation |
| RAD9A | 0.131548 | 0.018014 | Other Positive Correlation |
| EXO5 | 0.131399 | 0.018146 | Other Positive Correlation |
| EML5 | 0.131329 | 0.018208 | Other Positive Correlation |
| LOXL2 | 0.131327 | 0.01821 | Other Positive Correlation |
| ZBTB7C | 0.131237 | 0.018291 | Other Positive Correlation |
| SLC46A3 | 0.131187 | 0.018335 | Other Positive Correlation |
| BCOR | 0.131162 | 0.018357 | Other Positive Correlation |
| PI4KA | 0.131097 | 0.018416 | Other Positive Correlation |
| EMX1 | 0.131067 | 0.018443 | Other Positive Correlation |

| | | | |
|---|---|---|---|
| SLC37A4 | 0.131032 | 0.018475 | Other Positive Correlation |
| DBF4B | 0.131031 | 0.018475 | Other Positive Correlation |
| GNG7 | 0.130968 | 0.018532 | Other Positive Correlation |
| RAB33A | 0.130963 | 0.018537 | Other Positive Correlation |
| EFCAB14 | 0.130955 | 0.018544 | Other Positive Correlation |
| LYPD1 | 0.13091 | 0.018585 | Other Positive Correlation |
| PTP4A3 | 0.130898 | 0.018596 | Other Positive Correlation |
| PPP1R32 | 0.130876 | 0.018615 | Other Positive Correlation |
| DHRS11 | 0.130866 | 0.018625 | Other Positive Correlation |
| CRYM | 0.130811 | 0.018675 | Other Positive Correlation |
| WNT7B | 0.130801 | 0.018684 | Other Positive Correlation |
| MERTK | 0.130787 | 0.018696 | Other Positive Correlation |
| JOSD1 | 0.130726 | 0.018752 | Other Positive Correlation |
| SLC7A10 | 0.13072 | 0.018758 | Other Positive Correlation |
| HSPA1B | 0.130578 | 0.018888 | Other Positive Correlation |
| LHFPL2 | 0.130578 | 0.018888 | Other Positive Correlation |
| N4BP2 | 0.130505 | 0.018955 | Other Positive Correlation |
| DLC1 | 0.130464 | 0.018993 | Other Positive Correlation |
| RNF182 | 0.130457 | 0.019 | Other Positive Correlation |
| RER1 | 0.130398 | 0.019054 | Other Positive Correlation |
| EFS | 0.130348 | 0.0191 | Other Positive Correlation |
| MRM1 | 0.130291 | 0.019153 | Other Positive Correlation |
| BUD31 | 0.130254 | 0.019188 | Other Positive Correlation |
| TLE3 | 0.130219 | 0.019221 | Other Positive Correlation |
| SEC61A1 | 0.130213 | 0.019226 | Other Positive Correlation |
| RFK | 0.130119 | 0.019314 | Other Positive Correlation |
| IPMK | 0.130117 | 0.019316 | Other Positive Correlation |
| SEC14L6 | 0.13008 | 0.019351 | Other Positive Correlation |
| FZD9 | 0.130031 | 0.019397 | Other Positive Correlation |
| SOX8 | 0.129955 | 0.019468 | Other Positive Correlation |
| RIMS1 | 0.129943 | 0.01948 | Other Positive Correlation |
| MELTF | 0.129829 | 0.019588 | Other Positive Correlation |
| ILRUN | 0.129802 | 0.019613 | Other Positive Correlation |
| PGM5 | 0.129648 | 0.01976 | Other Positive Correlation |
| HVCN1 | 0.129613 | 0.019794 | Other Positive Correlation |
| OGFOD2 | 0.129554 | 0.01985 | Other Positive Correlation |
| BRME1 | 0.129422 | 0.019978 | Other Positive Correlation |
| UPF1 | 0.129405 | 0.019993 | Other Positive Correlation |
| GNMT | 0.129404 | 0.019995 | Other Positive Correlation |
| BAZ2A | 0.129397 | 0.020001 | Other Positive Correlation |
| ME1 | 0.129335 | 0.020061 | Other Positive Correlation |
| PIH1D1 | 0.129213 | 0.02018 | Other Positive Correlation |
| IL17RA | 0.129193 | 0.020199 | Other Positive Correlation |
| TLX1 | 0.129183 | 0.020209 | Other Positive Correlation |
| VSTM2B | 0.129147 | 0.020245 | Other Positive Correlation |
| DENND2D | 0.129088 | 0.020302 | Other Positive Correlation |
| ZDHHC13 | 0.129061 | 0.020328 | Other Positive Correlation |
| RUNX3 | 0.129033 | 0.020356 | Other Positive Correlation |

| | | | |
|---|---|---|---|
| ATP1A3 | 0.129025 | 0.020364 | Other Positive Correlation |
| CCHCR1 | 0.129013 | 0.020375 | Other Positive Correlation |
| TNPO2 | 0.129005 | 0.020384 | Other Positive Correlation |
| HTR1D | 0.128979 | 0.020409 | Other Positive Correlation |
| ZNF836 | 0.128861 | 0.020526 | Other Positive Correlation |
| SMIM29 | 0.128836 | 0.02055 | Other Positive Correlation |
| SCAMP4 | 0.128828 | 0.020558 | Other Positive Correlation |
| C15orf39 | 0.128814 | 0.020572 | Other Positive Correlation |
| KBTBD8 | 0.128737 | 0.020648 | Other Positive Correlation |
| ZFP3 | 0.128627 | 0.020758 | Other Positive Correlation |
| HS1BP3 | 0.128606 | 0.020779 | Other Positive Correlation |
| CREB3L1 | 0.128602 | 0.020782 | Other Positive Correlation |
| LZIC | 0.128601 | 0.020784 | Other Positive Correlation |
| CTDSP2 | 0.12854 | 0.020845 | Other Positive Correlation |
| ADD2 | 0.128519 | 0.020865 | Other Positive Correlation |
| FGF16 | 0.128509 | 0.020876 | Other Positive Correlation |
| TBX15 | 0.128465 | 0.02092 | Other Positive Correlation |
| CDH15 | 0.128448 | 0.020938 | Other Positive Correlation |
| FAM104A | 0.128447 | 0.020938 | Other Positive Correlation |
| PLA2G3 | 0.128398 | 0.020988 | Other Positive Correlation |
| DNAJC30 | 0.128391 | 0.020995 | Other Positive Correlation |
| FBXL16 | 0.128312 | 0.021074 | Other Positive Correlation |
| DLX6 | 0.128287 | 0.0211 | Other Positive Correlation |
| AKNA | 0.128259 | 0.021128 | Other Positive Correlation |
| VKORC1L1 | 0.128191 | 0.021197 | Other Positive Correlation |
| RASL10A | 0.128105 | 0.021285 | Other Positive Correlation |
| CHKA | 0.128091 | 0.021299 | Other Positive Correlation |
| ASGR1 | 0.12808 | 0.02131 | Other Positive Correlation |
| CELSR3 | 0.128076 | 0.021314 | Other Positive Correlation |
| NOVA2 | 0.127984 | 0.021408 | Other Positive Correlation |
| SH3BP5L | 0.12787 | 0.021525 | Other Positive Correlation |
| GNG4 | 0.127858 | 0.021538 | Other Positive Correlation |
| NCR3LG1 | 0.127812 | 0.021585 | Other Positive Correlation |
| TBC1D10B | 0.127761 | 0.021638 | Other Positive Correlation |
| SYNE2 | 0.12776 | 0.021639 | Other Positive Correlation |
| NMU | 0.127704 | 0.021697 | Other Positive Correlation |
| TCEAL7 | 0.127595 | 0.021811 | Other Positive Correlation |
| CXCR4 | 0.127529 | 0.021879 | Other Positive Correlation |
| PARD6A | 0.127527 | 0.021881 | Other Positive Correlation |
| TMEM170B | 0.127367 | 0.022049 | Other Positive Correlation |
| KCNJ10 | 0.127348 | 0.022069 | Other Positive Correlation |
| ETV5 | 0.127219 | 0.022206 | Other Positive Correlation |
| NECTIN3 | 0.127215 | 0.022209 | Other Positive Correlation |
| KCTD2 | 0.127209 | 0.022216 | Other Positive Correlation |
| EPOR | 0.127183 | 0.022244 | Other Positive Correlation |
| MIA2 | 0.127046 | 0.022389 | Other Positive Correlation |
| HOXD9 | 0.126975 | 0.022465 | Other Positive Correlation |
| THAP3 | 0.126964 | 0.022477 | Other Positive Correlation |

| | | | |
|---|---|---|---|
| PALB2 | 0.126963 | 0.022477 | Other Positive Correlation |
| C3orf18 | 0.126933 | 0.022509 | Other Positive Correlation |
| C2orf68 | 0.126816 | 0.022635 | Other Positive Correlation |
| NKD2 | 0.12679 | 0.022663 | Other Positive Correlation |
| THAP2 | 0.126767 | 0.022688 | Other Positive Correlation |
| FAM124A | 0.126756 | 0.022699 | Other Positive Correlation |
| CCNE2 | 0.12662 | 0.022846 | Other Positive Correlation |
| DNAJC14 | 0.126574 | 0.022897 | Other Positive Correlation |
| KLHL15 | 0.126468 | 0.023012 | Other Positive Correlation |
| TERF2 | 0.126395 | 0.023092 | Other Positive Correlation |
| SAP130 | 0.126326 | 0.023168 | Other Positive Correlation |
| C1orf226 | 0.126289 | 0.023208 | Other Positive Correlation |
| B3GLCT | 0.126219 | 0.023286 | Other Positive Correlation |
| SHISA4 | 0.126187 | 0.023321 | Other Positive Correlation |
| SCML1 | 0.126184 | 0.023324 | Other Positive Correlation |
| UCP2 | 0.12618 | 0.023329 | Other Positive Correlation |
| FAM234B | 0.126156 | 0.023355 | Other Positive Correlation |
| CYP2E1 | 0.126086 | 0.023433 | Other Positive Correlation |
| RBBP6 | 0.12607 | 0.02345 | Other Positive Correlation |
| PITX3 | 0.126053 | 0.023469 | Other Positive Correlation |
| SYNJ1 | 0.126032 | 0.023492 | Other Positive Correlation |
| CHRM1 | 0.125985 | 0.023545 | Other Positive Correlation |
| C6orf47 | 0.125983 | 0.023547 | Other Positive Correlation |
| PRKD1 | 0.125919 | 0.023618 | Other Positive Correlation |
| TGFBR3 | 0.125647 | 0.023923 | Other Positive Correlation |
| SLC29A4 | 0.125565 | 0.024017 | Other Positive Correlation |
| FZD5 | 0.125515 | 0.024073 | Other Positive Correlation |
| JAM2 | 0.125512 | 0.024076 | Other Positive Correlation |
| H2BC11 | 0.125388 | 0.024217 | Other Positive Correlation |
| NR3C2 | 0.125342 | 0.02427 | Other Positive Correlation |
| THY1 | 0.125267 | 0.024356 | Other Positive Correlation |
| GAL3ST3 | 0.125245 | 0.024381 | Other Positive Correlation |
| CNBD2 | 0.125238 | 0.024389 | Other Positive Correlation |
| NR2F1 | 0.125234 | 0.024394 | Other Positive Correlation |
| ATP6V1F | 0.12523 | 0.024398 | Other Positive Correlation |
| CERS4 | 0.1252 | 0.024433 | Other Positive Correlation |
| ARIH2 | 0.125177 | 0.02446 | Other Positive Correlation |
| CEP78 | 0.125169 | 0.024468 | Other Positive Correlation |
| MAMLD1 | 0.125144 | 0.024498 | Other Positive Correlation |
| CERT1 | 0.12509 | 0.02456 | Other Positive Correlation |
| ACOT8 | 0.125075 | 0.024576 | Other Positive Correlation |
| UNC13A | 0.125009 | 0.024653 | Other Positive Correlation |
| SNAPC5 | 0.124987 | 0.024678 | Other Positive Correlation |
| VAT1 | 0.124977 | 0.024691 | Other Positive Correlation |
| GPR108 | 0.124781 | 0.024918 | Other Positive Correlation |
| SSX2IP | 0.124732 | 0.024975 | Other Positive Correlation |
| NCCRP1 | 0.124725 | 0.024983 | Other Positive Correlation |
| ARMC7 | 0.124714 | 0.024997 | Other Positive Correlation |

| | | | |
|---|---|---|---|
| LACTB2 | 0.124696 | 0.025017 | Other Positive Correlation |
| SDR42E1 | 0.124646 | 0.025077 | Other Positive Correlation |
| AHSA1 | 0.124606 | 0.025124 | Other Positive Correlation |
| GGA2 | 0.124505 | 0.025243 | Other Positive Correlation |
| KCNMB4 | 0.124498 | 0.025251 | Other Positive Correlation |
| NARF | 0.124463 | 0.025292 | Other Positive Correlation |
| EPHX4 | 0.124456 | 0.025301 | Other Positive Correlation |
| PRR19 | 0.124298 | 0.025489 | Other Positive Correlation |
| RALGPS1 | 0.124286 | 0.025503 | Other Positive Correlation |
| RAB9A | 0.124185 | 0.025624 | Other Positive Correlation |
| GATA5 | 0.12418 | 0.02563 | Other Positive Correlation |
| LMO2 | 0.124167 | 0.025645 | Other Positive Correlation |
| ZNF550 | 0.124112 | 0.025711 | Other Positive Correlation |
| FAM78A | 0.124065 | 0.025767 | Other Positive Correlation |
| HLA-DPA1 | 0.124023 | 0.025818 | Other Positive Correlation |
| ARID5A | 0.123955 | 0.0259 | Other Positive Correlation |
| TMPRSS2 | 0.123933 | 0.025927 | Other Positive Correlation |
| RNF145 | 0.12391 | 0.025954 | Other Positive Correlation |
| PRDM12 | 0.123816 | 0.026069 | Other Positive Correlation |
| MAGEH1 | 0.123811 | 0.026075 | Other Positive Correlation |
| PXMP4 | 0.123713 | 0.026194 | Other Positive Correlation |
| DTX3 | 0.123699 | 0.026211 | Other Positive Correlation |
| STOX2 | 0.123674 | 0.026242 | Other Positive Correlation |
| OSGIN1 | 0.123638 | 0.026286 | Other Positive Correlation |
| ACADL | 0.123633 | 0.026292 | Other Positive Correlation |
| GLI2 | 0.123625 | 0.026301 | Other Positive Correlation |
| FLT4 | 0.123609 | 0.026321 | Other Positive Correlation |
| GPAM | 0.123444 | 0.026524 | Other Positive Correlation |
| CNNM3 | 0.123431 | 0.02654 | Other Positive Correlation |
| FDX1 | 0.12334 | 0.026653 | Other Positive Correlation |
| PPFIBP2 | 0.123329 | 0.026666 | Other Positive Correlation |
| ZBTB8B | 0.123305 | 0.026695 | Other Positive Correlation |
| SLC26A5 | 0.123187 | 0.026842 | Other Positive Correlation |
| AKIRIN2 | 0.123176 | 0.026857 | Other Positive Correlation |
| AARD | 0.123175 | 0.026858 | Other Positive Correlation |
| TRPV6 | 0.123108 | 0.026942 | Other Positive Correlation |
| TEX30 | 0.123088 | 0.026966 | Other Positive Correlation |
| MVB12B | 0.123058 | 0.027004 | Other Positive Correlation |
| PPM1E | 0.123023 | 0.027047 | Other Positive Correlation |
| SMIM10L1 | 0.122939 | 0.027154 | Other Positive Correlation |
| TNNC2 | 0.122927 | 0.027169 | Other Positive Correlation |
| ZNF850 | 0.122918 | 0.027179 | Other Positive Correlation |
| ZNF467 | 0.122895 | 0.027209 | Other Positive Correlation |
| DPH3 | 0.122819 | 0.027304 | Other Positive Correlation |
| DRD2 | 0.122698 | 0.027458 | Other Positive Correlation |
| PHYHIPL | 0.12269 | 0.027469 | Other Positive Correlation |
| GINS3 | 0.122572 | 0.027619 | Other Positive Correlation |
| ZDHHC18 | 0.122446 | 0.02778 | Other Positive Correlation |

| | | | |
|---|---|---|---|
| SPATC1L | 0.122413 | 0.027823 | Other Positive Correlation |
| TESK2 | 0.122379 | 0.027866 | Other Positive Correlation |
| LRRC37B | 0.12225 | 0.028032 | Other Positive Correlation |
| HSD17B14 | 0.122244 | 0.028041 | Other Positive Correlation |
| RCHY1 | 0.122194 | 0.028105 | Other Positive Correlation |
| TDRD10 | 0.122193 | 0.028106 | Other Positive Correlation |
| GCGR | 0.122168 | 0.028139 | Other Positive Correlation |
| CLDN23 | 0.122165 | 0.028143 | Other Positive Correlation |
| PFN3 | 0.12214 | 0.028175 | Other Positive Correlation |
| TEX15 | 0.122123 | 0.028197 | Other Positive Correlation |
| PLEKHG1 | 0.122037 | 0.02831 | Other Positive Correlation |
| SP5 | 0.121868 | 0.02853 | Other Positive Correlation |
| KIF12 | 0.121835 | 0.028574 | Other Positive Correlation |
| SHCBP1L | 0.121806 | 0.028611 | Other Positive Correlation |
| ZNF200 | 0.121766 | 0.028665 | Other Positive Correlation |
| KLHL30 | 0.12176 | 0.028672 | Other Positive Correlation |
| BEND7 | 0.121753 | 0.028681 | Other Positive Correlation |
| NOTO | 0.121723 | 0.028721 | Other Positive Correlation |
| SIAH2 | 0.12171 | 0.028739 | Other Positive Correlation |
| FEM1C | 0.121609 | 0.028873 | Other Positive Correlation |
| NPBWR1 | 0.121596 | 0.028889 | Other Positive Correlation |
| UBAP1 | 0.121401 | 0.02915 | Other Positive Correlation |
| SSBP2 | 0.121309 | 0.029273 | Other Positive Correlation |
| NPRL2 | 0.121274 | 0.02932 | Other Positive Correlation |
| SENP3 | 0.121213 | 0.029402 | Other Positive Correlation |
| ADRA1B | 0.121175 | 0.029453 | Other Positive Correlation |
| CDK5RAP1 | 0.121154 | 0.029482 | Other Positive Correlation |
| P2RX5 | 0.121057 | 0.029612 | Other Positive Correlation |
| PLEKHB1 | 0.121051 | 0.029621 | Other Positive Correlation |
| GPR4 | 0.12105 | 0.029622 | Other Positive Correlation |
| SLC30A4 | 0.120969 | 0.029732 | Other Positive Correlation |
| SMIM14 | 0.120966 | 0.029737 | Other Positive Correlation |
| WIPF1 | 0.120953 | 0.029754 | Other Positive Correlation |
| TLN2 | 0.120918 | 0.029801 | Other Positive Correlation |
| CHST9 | 0.120788 | 0.029979 | Other Positive Correlation |
| SORBS1 | 0.120783 | 0.029986 | Other Positive Correlation |
| CDK19 | 0.120724 | 0.030067 | Other Positive Correlation |
| ZNF473 | 0.120642 | 0.03018 | Other Positive Correlation |
| WNT9A | 0.120622 | 0.030207 | Other Positive Correlation |
| TENT5C | 0.120535 | 0.030327 | Other Positive Correlation |
| FXYD1 | 0.1205 | 0.030376 | Other Positive Correlation |
| MAN1C1 | 0.120495 | 0.030383 | Other Positive Correlation |
| CLXN | 0.120472 | 0.030415 | Other Positive Correlation |
| DOLK | 0.12047 | 0.030418 | Other Positive Correlation |
| EVI5L | 0.120459 | 0.030433 | Other Positive Correlation |
| ZNF574 | 0.12045 | 0.030446 | Other Positive Correlation |
| STX11 | 0.12042 | 0.030488 | Other Positive Correlation |
| SERPINF1 | 0.12037 | 0.030557 | Other Positive Correlation |

| | | | |
|---|---|---|---|
| AAK1 | 0.12024 | 0.030739 | Other Positive Correlation |
| ALAS1 | 0.120239 | 0.030739 | Other Positive Correlation |
| ABHD12 | 0.120207 | 0.030785 | Other Positive Correlation |
| TAMALIN | 0.120175 | 0.03083 | Other Positive Correlation |
| KATNAL2 | 0.120172 | 0.030833 | Other Positive Correlation |
| C22orf39 | 0.120078 | 0.030965 | Other Positive Correlation |
| RGS9BP | 0.120057 | 0.030996 | Other Positive Correlation |
| YKT6 | 0.120003 | 0.031072 | Other Positive Correlation |
| DNAJC22 | 0.119977 | 0.031108 | Other Positive Correlation |
| ATP9A | 0.119947 | 0.031151 | Other Positive Correlation |
| IKZF1 | 0.119934 | 0.031169 | Other Positive Correlation |
| DDX20 | 0.119912 | 0.0312 | Other Positive Correlation |
| EPHX1 | 0.119815 | 0.031338 | Other Positive Correlation |
| TFAP2C | 0.119771 | 0.0314 | Other Positive Correlation |
| NECAP1 | 0.11975 | 0.031432 | Other Positive Correlation |
| SEMA6D | 0.119617 | 0.031621 | Other Positive Correlation |
| RANBP6 | 0.1196 | 0.031646 | Other Positive Correlation |
| GLIS2 | 0.119538 | 0.031734 | Other Positive Correlation |
| FLVCR2 | 0.119538 | 0.031734 | Other Positive Correlation |
| CACTIN | 0.11949 | 0.031803 | Other Positive Correlation |
| SDE2 | 0.119487 | 0.031807 | Other Positive Correlation |
| KLHL28 | 0.119466 | 0.031838 | Other Positive Correlation |
| FXR2 | 0.119422 | 0.031902 | Other Positive Correlation |
| OIP5 | 0.119352 | 0.032003 | Other Positive Correlation |
| OSER1 | 0.119349 | 0.032008 | Other Positive Correlation |
| CNOT8 | 0.119346 | 0.032012 | Other Positive Correlation |
| NFATC2IP | 0.119301 | 0.032077 | Other Positive Correlation |
| CENPS | 0.11928 | 0.032107 | Other Positive Correlation |
| FIBCD1 | 0.11926 | 0.032137 | Other Positive Correlation |
| CD14 | 0.119234 | 0.032174 | Other Positive Correlation |
| MADD | 0.119208 | 0.032212 | Other Positive Correlation |
| SLC25A14 | 0.119196 | 0.03223 | Other Positive Correlation |
| ZNF568 | 0.119096 | 0.032375 | Other Positive Correlation |
| ZNF92 | 0.119042 | 0.032455 | Other Positive Correlation |
| YIPF7 | 0.119009 | 0.032503 | Other Positive Correlation |
| MTMR4 | 0.118943 | 0.0326 | Other Positive Correlation |
| IQSEC3 | 0.118936 | 0.032611 | Other Positive Correlation |
| CLDN9 | 0.118918 | 0.032637 | Other Positive Correlation |
| SHH | 0.11891 | 0.032649 | Other Positive Correlation |
| DENND2A | 0.118905 | 0.032657 | Other Positive Correlation |
| PAIP2 | 0.11889 | 0.032678 | Other Positive Correlation |
| PODN | 0.11878 | 0.032841 | Other Positive Correlation |
| PHLPP1 | 0.118771 | 0.032855 | Other Positive Correlation |
| HAP1 | 0.118761 | 0.032869 | Other Positive Correlation |
| EPS8 | 0.118761 | 0.03287 | Other Positive Correlation |
| SPOPL | 0.118757 | 0.032875 | Other Positive Correlation |
| LPO | 0.118737 | 0.032904 | Other Positive Correlation |
| FBXO32 | 0.118706 | 0.032951 | Other Positive Correlation |

| | | | |
|---|---|---|---|
| SEC24A | 0.118689 | 0.032976 | Other Positive Correlation |
| EIF4EBP2 | 0.118685 | 0.032981 | Other Positive Correlation |
| ALKAL1 | 0.118658 | 0.033023 | Other Positive Correlation |
| SLC6A9 | 0.118631 | 0.033062 | Other Positive Correlation |
| TPRG1L | 0.118618 | 0.033082 | Other Positive Correlation |
| PARS2 | 0.118603 | 0.033104 | Other Positive Correlation |
| UPK3A | 0.118479 | 0.03329 | Other Positive Correlation |
| ZDHHC23 | 0.11841 | 0.033393 | Other Positive Correlation |
| PPM1H | 0.118367 | 0.033458 | Other Positive Correlation |
| PRRT4 | 0.11834 | 0.033499 | Other Positive Correlation |
| SLC2A4 | 0.118241 | 0.033648 | Other Positive Correlation |
| BCL2L10 | 0.118218 | 0.033683 | Other Positive Correlation |
| C6orf89 | 0.1182 | 0.033709 | Other Positive Correlation |
| ECHDC2 | 0.118018 | 0.033986 | Other Positive Correlation |
| CSAD | 0.118003 | 0.034008 | Other Positive Correlation |
| KATNB1 | 0.118002 | 0.03401 | Other Positive Correlation |
| RPS6KL1 | 0.117979 | 0.034046 | Other Positive Correlation |
| UBR7 | 0.11793 | 0.03412 | Other Positive Correlation |
| SMKR1 | 0.117916 | 0.034142 | Other Positive Correlation |
| SYNGR3 | 0.117765 | 0.034374 | Other Positive Correlation |
| PRR36 | 0.117748 | 0.0344 | Other Positive Correlation |
| ZFYVE1 | 0.117739 | 0.034414 | Other Positive Correlation |
| ARRDC2 | 0.117736 | 0.034418 | Other Positive Correlation |
| ACADS | 0.11768 | 0.034506 | Other Positive Correlation |
| MXRA8 | 0.117652 | 0.034549 | Other Positive Correlation |
| POMC | 0.117621 | 0.034596 | Other Positive Correlation |
| PACSIN1 | 0.117611 | 0.034612 | Other Positive Correlation |
| PLPP7 | 0.1176 | 0.034629 | Other Positive Correlation |
| ARID3A | 0.117598 | 0.034631 | Other Positive Correlation |
| WIPF3 | 0.11754 | 0.034722 | Other Positive Correlation |
| TOP3B | 0.117515 | 0.03476 | Other Positive Correlation |
| ARHGEF33 | 0.117503 | 0.034779 | Other Positive Correlation |
| ICOSLG | 0.117485 | 0.034808 | Other Positive Correlation |
| SLC30A5 | 0.117385 | 0.034963 | Other Positive Correlation |
| TRIM23 | 0.117347 | 0.035023 | Other Positive Correlation |
| RAB23 | 0.117235 | 0.035198 | Other Positive Correlation |
| CCNE1 | 0.117161 | 0.035315 | Other Positive Correlation |
| NSD3 | 0.11714 | 0.035348 | Other Positive Correlation |
| EMP2 | 0.117133 | 0.03536 | Other Positive Correlation |
| PGRMC2 | 0.117112 | 0.035392 | Other Positive Correlation |
| FARSA | 0.117077 | 0.035447 | Other Positive Correlation |
| PNMA3 | 0.117071 | 0.035456 | Other Positive Correlation |
| ATP6V1B2 | 0.117027 | 0.035526 | Other Positive Correlation |
| MRPL57 | 0.116981 | 0.035599 | Other Positive Correlation |
| FIGNL1 | 0.116979 | 0.035602 | Other Positive Correlation |
| DUSP23 | 0.11697 | 0.035617 | Other Positive Correlation |
| ARHGEF6 | 0.116942 | 0.035661 | Other Positive Correlation |
| C11orf24 | 0.116931 | 0.035679 | Other Positive Correlation |

| | | | |
|---|---|---|---|
| LLGL2 | 0.116885 | 0.035751 | Other Positive Correlation |
| VIPR2 | 0.116869 | 0.035778 | Other Positive Correlation |
| ABHD12B | 0.116836 | 0.03583 | Other Positive Correlation |
| EFR3B | 0.116788 | 0.035907 | Other Positive Correlation |
| TRPC6 | 0.116736 | 0.035989 | Other Positive Correlation |
| ISM1 | 0.116724 | 0.036009 | Other Positive Correlation |
| KLHDC8B | 0.116621 | 0.036174 | Other Positive Correlation |
| ZCCHC14 | 0.116477 | 0.036406 | Other Positive Correlation |
| GRHL1 | 0.116462 | 0.036431 | Other Positive Correlation |
| TMEM63C | 0.116403 | 0.036527 | Other Positive Correlation |
| HS3ST3A1 | 0.11634 | 0.036629 | Other Positive Correlation |
| BIN1 | 0.116171 | 0.036904 | Other Positive Correlation |
| LMBR1L | 0.116139 | 0.036957 | Other Positive Correlation |
| NQO2 | 0.116137 | 0.03696 | Other Positive Correlation |
| RUBCN | 0.116109 | 0.037006 | Other Positive Correlation |
| GTF3C2 | 0.11609 | 0.037036 | Other Positive Correlation |
| FXYD4 | 0.116078 | 0.037057 | Other Positive Correlation |
| SGSH | 0.116042 | 0.037115 | Other Positive Correlation |
| CH25H | 0.11602 | 0.037151 | Other Positive Correlation |
| CDHR3 | 0.116014 | 0.037162 | Other Positive Correlation |
| CHL1 | 0.11599 | 0.037201 | Other Positive Correlation |
| RIMS4 | 0.115887 | 0.037372 | Other Positive Correlation |
| DCUN1D2 | 0.115748 | 0.037601 | Other Positive Correlation |
| ZNF738 | 0.115704 | 0.037675 | Other Positive Correlation |
| CCNJ | 0.11566 | 0.037748 | Other Positive Correlation |
| TMEM86A | 0.115613 | 0.037827 | Other Positive Correlation |
| EPS15L1 | 0.115506 | 0.038005 | Other Positive Correlation |
| PLCL1 | 0.115373 | 0.038229 | Other Positive Correlation |
| WNT2B | 0.115372 | 0.03823 | Other Positive Correlation |
| PANK4 | 0.115351 | 0.038266 | Other Positive Correlation |
| ING1 | 0.115328 | 0.038305 | Other Positive Correlation |
| LIFR | 0.115305 | 0.038343 | Other Positive Correlation |
| MRPL36 | 0.115255 | 0.038428 | Other Positive Correlation |
| BLOC1S4 | 0.115235 | 0.038461 | Other Positive Correlation |
| USP11 | 0.115235 | 0.038462 | Other Positive Correlation |
| PDE8B | 0.115184 | 0.038548 | Other Positive Correlation |
| CEP162 | 0.115154 | 0.038599 | Other Positive Correlation |
| USP30 | 0.11511 | 0.038675 | Other Positive Correlation |
| ZNF674 | 0.115093 | 0.038704 | Other Positive Correlation |
| MAGI2 | 0.115051 | 0.038774 | Other Positive Correlation |
| MGARP | 0.114963 | 0.038925 | Other Positive Correlation |
| ZNF226 | 0.114879 | 0.039069 | Other Positive Correlation |
| SLC1A1 | 0.114874 | 0.039078 | Other Positive Correlation |
| FZD2 | 0.114868 | 0.039089 | Other Positive Correlation |
| ADCY1 | 0.114847 | 0.039123 | Other Positive Correlation |
| GMPPA | 0.114807 | 0.039193 | Other Positive Correlation |
| RDH13 | 0.114781 | 0.039237 | Other Positive Correlation |
| CCDC126 | 0.114781 | 0.039237 | Other Positive Correlation |

| | | | |
|---|---|---|---|
| CCDC85A | 0.114768 | 0.03926 | Other Positive Correlation |
| C17orf67 | 0.114767 | 0.039262 | Other Positive Correlation |
| FABP3 | 0.114737 | 0.039313 | Other Positive Correlation |
| SRPK3 | 0.114707 | 0.039365 | Other Positive Correlation |
| KL | 0.1147 | 0.039377 | Other Positive Correlation |
| PRIM2 | 0.114672 | 0.039425 | Other Positive Correlation |
| MTMR14 | 0.114638 | 0.039485 | Other Positive Correlation |
| GDPD5 | 0.114612 | 0.03953 | Other Positive Correlation |
| HIC2 | 0.114609 | 0.039535 | Other Positive Correlation |
| AHCYL2 | 0.114513 | 0.039702 | Other Positive Correlation |
| ZDHHC2 | 0.114484 | 0.039753 | Other Positive Correlation |
| SDF4 | 0.114445 | 0.039819 | Other Positive Correlation |
| RAD54L2 | 0.11444 | 0.039829 | Other Positive Correlation |
| CHPT1 | 0.114411 | 0.039878 | Other Positive Correlation |
| DLX1 | 0.114402 | 0.039894 | Other Positive Correlation |
| CREBRF | 0.114373 | 0.039945 | Other Positive Correlation |
| GOSR2 | 0.114292 | 0.040088 | Other Positive Correlation |
| FAM221A | 0.11429 | 0.040092 | Other Positive Correlation |
| RFLNA | 0.114264 | 0.040137 | Other Positive Correlation |
| STAG3 | 0.114228 | 0.040201 | Other Positive Correlation |
| RASSF8 | 0.114194 | 0.040259 | Other Positive Correlation |
| WASL | 0.114191 | 0.040265 | Other Positive Correlation |
| PAG1 | 0.114169 | 0.040304 | Other Positive Correlation |
| CACNA1G | 0.114157 | 0.040325 | Other Positive Correlation |
| KCNC3 | 0.11408 | 0.040461 | Other Positive Correlation |
| RASD1 | 0.113991 | 0.040618 | Other Positive Correlation |
| ITGA7 | 0.113949 | 0.040693 | Other Positive Correlation |
| SIK3 | 0.113929 | 0.040729 | Other Positive Correlation |
| TOR2A | 0.113912 | 0.040759 | Other Positive Correlation |
| ZNF232 | 0.113895 | 0.040789 | Other Positive Correlation |
| KIF26A | 0.113889 | 0.040799 | Other Positive Correlation |
| TPPP3 | 0.113843 | 0.040882 | Other Positive Correlation |
| POP5 | 0.113793 | 0.040971 | Other Positive Correlation |
| DCAF12 | 0.113762 | 0.041026 | Other Positive Correlation |
| PKDCC | 0.113741 | 0.041063 | Other Positive Correlation |
| CHCHD3 | 0.113727 | 0.04109 | Other Positive Correlation |
| CAMK2N1 | 0.113723 | 0.041095 | Other Positive Correlation |
| GRIN2B | 0.113717 | 0.041107 | Other Positive Correlation |
| ERVK3-1 | 0.113493 | 0.041509 | Other Positive Correlation |
| KDM6B | 0.113467 | 0.041557 | Other Positive Correlation |
| FUT4 | 0.11346 | 0.04157 | Other Positive Correlation |
| PTH1R | 0.113414 | 0.041653 | Other Positive Correlation |
| MAP3K2 | 0.113384 | 0.041707 | Other Positive Correlation |
| LEFTY2 | 0.113364 | 0.041744 | Other Positive Correlation |
| LKAAEAR1 | 0.113336 | 0.041794 | Other Positive Correlation |
| MRAS | 0.1133 | 0.04186 | Other Positive Correlation |
| TAX1BP1 | 0.113248 | 0.041953 | Other Positive Correlation |
| RMDN3 | 0.113148 | 0.042137 | Other Positive Correlation |

| | | | |
|---|---|---|---|
| UAP1L1 | 0.113115 | 0.042197 | Other Positive Correlation |
| TOR1A | 0.113115 | 0.042198 | Other Positive Correlation |
| EPHB3 | 0.113077 | 0.042267 | Other Positive Correlation |
| FNDC3A | 0.113063 | 0.042292 | Other Positive Correlation |
| DLGAP3 | 0.113045 | 0.042326 | Other Positive Correlation |
| TRIM11 | 0.112994 | 0.042419 | Other Positive Correlation |
| AP3D1 | 0.112967 | 0.042469 | Other Positive Correlation |
| GPM6B | 0.112961 | 0.04248 | Other Positive Correlation |
| TSC2 | 0.112959 | 0.042484 | Other Positive Correlation |
| L3MBTL2 | 0.112895 | 0.042602 | Other Positive Correlation |
| BAIAP3 | 0.112885 | 0.04262 | Other Positive Correlation |
| TICRR | 0.112871 | 0.042646 | Other Positive Correlation |
| MAX | 0.112838 | 0.042707 | Other Positive Correlation |
| ZNF239 | 0.112744 | 0.042881 | Other Positive Correlation |
| MAP1S | 0.112739 | 0.042892 | Other Positive Correlation |
| HBP1 | 0.112675 | 0.04301 | Other Positive Correlation |
| PODXL2 | 0.112668 | 0.043022 | Other Positive Correlation |
| HRH2 | 0.112636 | 0.043083 | Other Positive Correlation |
| NPIPA1 | 0.112588 | 0.043171 | Other Positive Correlation |
| MYOD1 | 0.112588 | 0.043172 | Other Positive Correlation |
| CEP350 | 0.112576 | 0.043194 | Other Positive Correlation |
| HCK | 0.11252 | 0.043299 | Other Positive Correlation |
| PNKD | 0.112508 | 0.043322 | Other Positive Correlation |
| ZNF48 | 0.112456 | 0.043419 | Other Positive Correlation |
| MOSPD1 | 0.112455 | 0.043421 | Other Positive Correlation |
| SUPT7L | 0.112398 | 0.043528 | Other Positive Correlation |
| STAU1 | 0.112368 | 0.043583 | Other Positive Correlation |
| ADRA2B | 0.112345 | 0.043628 | Other Positive Correlation |
| PTGIS | 0.112285 | 0.043741 | Other Positive Correlation |
| MFSD6L | 0.112147 | 0.044001 | Other Positive Correlation |
| IQCD | 0.112032 | 0.044221 | Other Positive Correlation |
| MOB2 | 0.112022 | 0.04424 | Other Positive Correlation |
| WDR86 | 0.111981 | 0.044317 | Other Positive Correlation |
| RDM1 | 0.111955 | 0.044367 | Other Positive Correlation |
| TCAP | 0.1119 | 0.044474 | Other Positive Correlation |
| ADAL | 0.111898 | 0.044477 | Other Positive Correlation |
| SLC26A11 | 0.111897 | 0.044479 | Other Positive Correlation |
| MCCC1 | 0.111827 | 0.044613 | Other Positive Correlation |
| DPYSL4 | 0.111823 | 0.044621 | Other Positive Correlation |
| ZNF606 | 0.111821 | 0.044624 | Other Positive Correlation |
| FNIP2 | 0.111802 | 0.044661 | Other Positive Correlation |
| ARHGAP24 | 0.111798 | 0.044669 | Other Positive Correlation |
| BMAL1 | 0.11179 | 0.044684 | Other Positive Correlation |
| HELT | 0.111729 | 0.044801 | Other Positive Correlation |
| NGB | 0.111724 | 0.044811 | Other Positive Correlation |
| RPN2 | 0.111722 | 0.044815 | Other Positive Correlation |
| GRAMD1B | 0.111698 | 0.044861 | Other Positive Correlation |
| MAP3K3 | 0.111655 | 0.044943 | Other Positive Correlation |

| | | | |
|---|---|---|---|
| LYNX1 | 0.111619 | 0.045013 | Other Positive Correlation |
| SPATA13 | 0.111616 | 0.045019 | Other Positive Correlation |
| CREBBP | 0.111593 | 0.045065 | Other Positive Correlation |
| PLEKHF1 | 0.111588 | 0.045073 | Other Positive Correlation |
| PPM1L | 0.11157 | 0.045109 | Other Positive Correlation |
| NAXD | 0.111567 | 0.045114 | Other Positive Correlation |
| PSD | 0.11155 | 0.045146 | Other Positive Correlation |
| DPYSL2 | 0.111516 | 0.045213 | Other Positive Correlation |
| CWH43 | 0.111444 | 0.045352 | Other Positive Correlation |
| ELAVL3 | 0.111395 | 0.045449 | Other Positive Correlation |
| ANKRD34B | 0.111391 | 0.045455 | Other Positive Correlation |
| SGPP2 | 0.111281 | 0.045671 | Other Positive Correlation |
| MIIP | 0.111213 | 0.045804 | Other Positive Correlation |
| SSBP3 | 0.111206 | 0.045819 | Other Positive Correlation |
| MCAM | 0.1112 | 0.04583 | Other Positive Correlation |
| SEC22B | 0.111174 | 0.045881 | Other Positive Correlation |
| TENT5A | 0.111122 | 0.045984 | Other Positive Correlation |
| SLC34A2 | 0.111018 | 0.046188 | Other Positive Correlation |
| CRTC3 | 0.11098 | 0.046264 | Other Positive Correlation |
| EIF2AK3 | 0.110951 | 0.046321 | Other Positive Correlation |
| RIPPLY3 | 0.110931 | 0.046361 | Other Positive Correlation |
| HAPLN3 | 0.110913 | 0.046396 | Other Positive Correlation |
| GFOD1 | 0.110888 | 0.046446 | Other Positive Correlation |
| POLR2C | 0.110882 | 0.046458 | Other Positive Correlation |
| MYEF2 | 0.110797 | 0.046626 | Other Positive Correlation |
| RBM15 | 0.110748 | 0.046725 | Other Positive Correlation |
| SLC5A10 | 0.110648 | 0.046923 | Other Positive Correlation |
| SLC44A3 | 0.110642 | 0.046936 | Other Positive Correlation |
| CCDC93 | 0.110611 | 0.046997 | Other Positive Correlation |
| CFAP74 | 0.110611 | 0.046999 | Other Positive Correlation |
| NKAIN1 | 0.110582 | 0.047056 | Other Positive Correlation |
| OXLD1 | 0.110545 | 0.047132 | Other Positive Correlation |
| FNBP1L | 0.110544 | 0.047133 | Other Positive Correlation |
| PDE10A | 0.110543 | 0.047134 | Other Positive Correlation |
| KCNK5 | 0.110487 | 0.047247 | Other Positive Correlation |
| KEAP1 | 0.110417 | 0.047389 | Other Positive Correlation |
| WNT11 | 0.11032 | 0.047584 | Other Positive Correlation |
| SYTL4 | 0.110292 | 0.047641 | Other Positive Correlation |
| CYP39A1 | 0.11024 | 0.047747 | Other Positive Correlation |
| DNAL1 | 0.110207 | 0.047814 | Other Positive Correlation |
| TAPT1 | 0.110198 | 0.047834 | Other Positive Correlation |
| GDF15 | 0.110092 | 0.048049 | Other Positive Correlation |
| STK32A | 0.110063 | 0.048108 | Other Positive Correlation |
| KCNK13 | 0.110016 | 0.048203 | Other Positive Correlation |
| ZNF777 | 0.109979 | 0.04828 | Other Positive Correlation |
| C19orf18 | 0.109957 | 0.048324 | Other Positive Correlation |
| TDRD7 | 0.109949 | 0.048342 | Other Positive Correlation |
| FDX2 | 0.109932 | 0.048377 | Other Positive Correlation |

| | | | |
|---|---|---|---|
| CENPC | 0.109914 | 0.048412 | Other Positive Correlation |
| TIFA | 0.109901 | 0.048441 | Other Positive Correlation |
| MTERF4 | 0.109851 | 0.048543 | Other Positive Correlation |
| PGPEP1 | 0.109738 | 0.048775 | Other Positive Correlation |
| SLC18A3 | 0.10969 | 0.048875 | Other Positive Correlation |
| HSPB3 | 0.109664 | 0.048929 | Other Positive Correlation |
| TRIM58 | 0.109581 | 0.049102 | Other Positive Correlation |
| AARS1 | 0.109575 | 0.049115 | Other Positive Correlation |
| ZNF768 | 0.109563 | 0.04914 | Other Positive Correlation |
| AOC2 | 0.109413 | 0.049452 | Other Positive Correlation |
| MYRIP | 0.109384 | 0.049513 | Other Positive Correlation |
| GPX8 | 0.109284 | 0.049722 | Other Positive Correlation |
| VAV1 | 0.109278 | 0.049735 | Other Positive Correlation |
| CDC14B | 0.109259 | 0.049776 | Other Positive Correlation |
| MAIP1 | 0.109121 | 0.050067 | Other Positive Correlation |
| BEND6 | 0.109071 | 0.050172 | Other Positive Correlation |
| EFCC1 | 0.109061 | 0.050193 | Other Positive Correlation |
| TAOK2 | 0.109055 | 0.050206 | Other Positive Correlation |
| TMEM158 | 0.108984 | 0.050357 | Other Positive Correlation |
| NKX1-1 | 0.108983 | 0.050359 | Other Positive Correlation |
| MBLAC1 | 0.108962 | 0.050403 | Other Positive Correlation |
| MSH2 | 0.108891 | 0.050554 | Other Positive Correlation |
| TMEM62 | 0.108887 | 0.050562 | Other Positive Correlation |
| PAF1 | 0.108806 | 0.050737 | Other Positive Correlation |
| APBA1 | 0.108797 | 0.050756 | Other Positive Correlation |
| C4orf19 | 0.108776 | 0.050799 | Other Positive Correlation |
| PCOLCE2 | 0.108762 | 0.05083 | Other Positive Correlation |
| APCDD1 | 0.108731 | 0.050897 | Other Positive Correlation |
| AURKC | 0.10867 | 0.051027 | Other Positive Correlation |
| ESYT3 | 0.108631 | 0.051112 | Other Positive Correlation |
| PIK3C2A | 0.108601 | 0.051176 | Other Positive Correlation |
| VASH1 | 0.108517 | 0.051357 | Other Positive Correlation |
| ZFHX3 | 0.108516 | 0.051359 | Other Positive Correlation |
| PEAK1 | 0.108484 | 0.051429 | Other Positive Correlation |
| GMEB2 | 0.108419 | 0.051568 | Other Positive Correlation |
| RNF10 | 0.108393 | 0.051626 | Other Positive Correlation |
| AP3M2 | 0.108373 | 0.051669 | Other Positive Correlation |
| TMEM45B | 0.108349 | 0.051721 | Other Positive Correlation |
| RALGPS2 | 0.10829 | 0.051848 | Other Positive Correlation |
| WNT1 | 0.108275 | 0.051881 | Other Positive Correlation |
| CCDC77 | 0.108269 | 0.051895 | Other Positive Correlation |
| FAM229A | 0.108233 | 0.051973 | Other Positive Correlation |
| PLXDC1 | 0.108162 | 0.052128 | Other Positive Correlation |
| MCEE | 0.10814 | 0.052175 | Other Positive Correlation |
| ITSN2 | 0.108139 | 0.052178 | Other Positive Correlation |
| SLC2A13 | 0.108067 | 0.052337 | Other Positive Correlation |
| DISP1 | 0.108037 | 0.052401 | Other Positive Correlation |
| ZNF148 | 0.108001 | 0.052482 | Other Positive Correlation |

| | | | |
|---|---|---|---|
| TSSC4 | 0.107983 | 0.052521 | Other Positive Correlation |
| NTF3 | 0.107863 | 0.052785 | Other Positive Correlation |
| CCDC74B | 0.107843 | 0.052829 | Other Positive Correlation |
| DGKE | 0.107788 | 0.05295 | Other Positive Correlation |
| ZNF117 | 0.107745 | 0.053046 | Other Positive Correlation |
| TOM1 | 0.107697 | 0.053153 | Other Positive Correlation |
| FOXC2 | 0.107679 | 0.053193 | Other Positive Correlation |
| NUTM2E | 0.107665 | 0.053223 | Other Positive Correlation |
| KMT2B | 0.107662 | 0.053231 | Other Positive Correlation |
| MRGPRF | 0.107659 | 0.053238 | Other Positive Correlation |
| SLC43A1 | 0.107646 | 0.053266 | Other Positive Correlation |
| ALKBH4 | 0.107626 | 0.05331 | Other Positive Correlation |
| TRIM61 | 0.107624 | 0.053315 | Other Positive Correlation |
| CPAMD8 | 0.10761 | 0.053348 | Other Positive Correlation |
| METTL3 | 0.107588 | 0.053395 | Other Positive Correlation |
| PPFIA3 | 0.107584 | 0.053404 | Other Positive Correlation |
| PTOV1 | 0.10758 | 0.053415 | Other Positive Correlation |
| HOXC9 | 0.107554 | 0.053472 | Other Positive Correlation |
| EMID1 | 0.107534 | 0.053517 | Other Positive Correlation |
| RP9 | 0.107516 | 0.053556 | Other Positive Correlation |
| LHX3 | 0.107497 | 0.053599 | Other Positive Correlation |
| PDE6B | 0.107496 | 0.053602 | Other Positive Correlation |
| ADPRH | 0.107441 | 0.053724 | Other Positive Correlation |
| BEAN1 | 0.10733 | 0.053973 | Other Positive Correlation |
| SLC25A40 | 0.107329 | 0.053976 | Other Positive Correlation |
| UNC5D | 0.107302 | 0.054036 | Other Positive Correlation |
| LRRC71 | 0.107268 | 0.054113 | Other Positive Correlation |
| FES | 0.107247 | 0.054162 | Other Positive Correlation |
| PET100 | 0.107246 | 0.054163 | Other Positive Correlation |
| PLBD2 | 0.107163 | 0.05435 | Other Positive Correlation |
| RELA | 0.107122 | 0.054443 | Other Positive Correlation |
| ATP11B | 0.107111 | 0.054467 | Other Positive Correlation |
| CRACR2B | 0.107078 | 0.054544 | Other Positive Correlation |
| RPP38 | 0.107054 | 0.054597 | Other Positive Correlation |
| SH2D4A | 0.106963 | 0.054805 | Other Positive Correlation |
| FAM50A | 0.106949 | 0.054837 | Other Positive Correlation |
| COL4A4 | 0.106937 | 0.054864 | Other Positive Correlation |
| DPY19L3 | 0.106878 | 0.054998 | Other Positive Correlation |
| RHBG | 0.106853 | 0.055056 | Other Positive Correlation |
| CBX2 | 0.106837 | 0.055093 | Other Positive Correlation |
| VCPKMT | 0.106824 | 0.055123 | Other Positive Correlation |
| ATF6 | 0.106822 | 0.055128 | Other Positive Correlation |
| SASH1 | 0.106814 | 0.055146 | Other Positive Correlation |
| VCPIP1 | 0.106775 | 0.055235 | Other Positive Correlation |
| KIAA1614 | 0.106765 | 0.055258 | Other Positive Correlation |
| BANK1 | 0.106631 | 0.055566 | Other Positive Correlation |
| PCDH8 | 0.106602 | 0.055632 | Other Positive Correlation |
| ENSG00000286022 | 0.106586 | 0.055671 | Other Positive Correlation |

| | | | |
|---|---|---|---|
| OGDHL | 0.106574 | 0.055696 | Other Positive Correlation |
| PCGF5 | 0.106551 | 0.055751 | Other Positive Correlation |
| DDX23 | 0.106509 | 0.055848 | Other Positive Correlation |
| CSPG5 | 0.106471 | 0.055935 | Other Positive Correlation |
| ABTB2 | 0.106464 | 0.055952 | Other Positive Correlation |
| DLL4 | 0.106462 | 0.055956 | Other Positive Correlation |
| LINGO2 | 0.106427 | 0.056038 | Other Positive Correlation |
| GK | 0.106408 | 0.056083 | Other Positive Correlation |
| PICK1 | 0.106357 | 0.056199 | Other Positive Correlation |
| CA2 | 0.106354 | 0.056207 | Other Positive Correlation |
| USP32 | 0.106354 | 0.056208 | Other Positive Correlation |
| TNFSF12 | 0.106351 | 0.056215 | Other Positive Correlation |
| ZSCAN1 | 0.106289 | 0.056359 | Other Positive Correlation |
| ICA1L | 0.106193 | 0.056582 | Other Positive Correlation |
| PRKCB | 0.106165 | 0.056649 | Other Positive Correlation |
| KLHDC8A | 0.106117 | 0.056762 | Other Positive Correlation |
| PDE4D | 0.106016 | 0.056999 | Other Positive Correlation |
| ATP6V1C2 | 0.105984 | 0.057075 | Other Positive Correlation |
| PPP1R11 | 0.10597 | 0.057107 | Other Positive Correlation |
| MDGA1 | 0.105949 | 0.057156 | Other Positive Correlation |
| CTSH | 0.105931 | 0.0572 | Other Positive Correlation |
| TP53RK | 0.105894 | 0.057288 | Other Positive Correlation |
| NKX6-2 | 0.105804 | 0.0575 | Other Positive Correlation |
| HIF1AN | 0.105766 | 0.05759 | Other Positive Correlation |
| DNM2 | 0.105715 | 0.057711 | Other Positive Correlation |
| ZNF454 | 0.105683 | 0.057788 | Other Positive Correlation |
| MTCH1 | 0.105652 | 0.057861 | Other Positive Correlation |
| EMSY | 0.105575 | 0.058045 | Other Positive Correlation |
| CAPN5 | 0.105552 | 0.0581 | Other Positive Correlation |
| FOXP4 | 0.105547 | 0.058113 | Other Positive Correlation |
| MTFR1L | 0.105518 | 0.058181 | Other Positive Correlation |
| NPLOC4 | 0.105491 | 0.058245 | Other Positive Correlation |
| CABIN1 | 0.105484 | 0.058263 | Other Positive Correlation |
| UCP1 | 0.105477 | 0.058279 | Other Positive Correlation |
| CFAP92 | 0.105472 | 0.05829 | Other Positive Correlation |
| VSTM4 | 0.105432 | 0.058388 | Other Positive Correlation |
| CA11 | 0.105403 | 0.058456 | Other Positive Correlation |
| PIK3AP1 | 0.105382 | 0.058508 | Other Positive Correlation |
| DMRT3 | 0.105305 | 0.058693 | Other Positive Correlation |
| AMPD2 | 0.105302 | 0.058699 | Other Positive Correlation |
| STX12 | 0.105253 | 0.058818 | Other Positive Correlation |
| CALCA | 0.10523 | 0.058874 | Other Positive Correlation |
| TMEM147 | 0.105207 | 0.058931 | Other Positive Correlation |
| CRY2 | 0.105151 | 0.059066 | Other Positive Correlation |
| ESS2 | 0.105119 | 0.059143 | Other Positive Correlation |
| RBM18 | 0.105083 | 0.059231 | Other Positive Correlation |
| CCL3L1 | 0.104992 | 0.059452 | Other Positive Correlation |
| KLHL22 | 0.104958 | 0.059535 | Other Positive Correlation |

| | | | |
|---|---|---|---|
| MED26 | 0.104845 | 0.059811 | Other Positive Correlation |
| QPRT | 0.104815 | 0.059886 | Other Positive Correlation |
| C2orf72 | 0.104813 | 0.059891 | Other Positive Correlation |
| SAMD14 | 0.104811 | 0.059896 | Other Positive Correlation |
| UNC119 | 0.10481 | 0.059897 | Other Positive Correlation |
| C4orf48 | 0.104793 | 0.059939 | Other Positive Correlation |
| CRACR2A | 0.104787 | 0.059953 | Other Positive Correlation |
| MIER2 | 0.104783 | 0.059963 | Other Positive Correlation |
| NUDT8 | 0.104774 | 0.059985 | Other Positive Correlation |
| GPR161 | 0.104663 | 0.060258 | Other Positive Correlation |
| TULP4 | 0.104649 | 0.060294 | Other Positive Correlation |
| ATOH7 | 0.104641 | 0.060312 | Other Positive Correlation |
| VEZF1 | 0.104594 | 0.060429 | Other Positive Correlation |
| GLI1 | 0.104579 | 0.060467 | Other Positive Correlation |
| HNRNPUL2 | 0.104577 | 0.06047 | Other Positive Correlation |
| CPLX1 | 0.104548 | 0.060542 | Other Positive Correlation |
| SMIM3 | 0.104547 | 0.060546 | Other Positive Correlation |
| BICDL1 | 0.104517 | 0.06062 | Other Positive Correlation |
| LAYN | 0.104483 | 0.060703 | Other Positive Correlation |
| GNAO1 | 0.104474 | 0.060726 | Other Positive Correlation |
| RSRC1 | 0.104462 | 0.060757 | Other Positive Correlation |
| PNMA8B | 0.104461 | 0.060758 | Other Positive Correlation |
| APC2 | 0.104458 | 0.060765 | Other Positive Correlation |
| IPO13 | 0.104458 | 0.060766 | Other Positive Correlation |
| EYA4 | 0.104437 | 0.060818 | Other Positive Correlation |
| STK33 | 0.104424 | 0.060852 | Other Positive Correlation |
| ATOSA | 0.104349 | 0.061038 | Other Positive Correlation |
| ZFP92 | 0.104328 | 0.061089 | Other Positive Correlation |
| ZNF304 | 0.104325 | 0.061096 | Other Positive Correlation |
| MTOR | 0.104232 | 0.061329 | Other Positive Correlation |
| DNAJB11 | 0.10419 | 0.061434 | Other Positive Correlation |
| ARMC5 | 0.104184 | 0.061449 | Other Positive Correlation |
| MSRB1 | 0.104107 | 0.061643 | Other Positive Correlation |
| KCTD16 | 0.104055 | 0.061772 | Other Positive Correlation |
| STAM | 0.104005 | 0.061898 | Other Positive Correlation |
| DNMT3B | 0.10399 | 0.061936 | Other Positive Correlation |
| VIPR1 | 0.103946 | 0.062048 | Other Positive Correlation |
| LRRC20 | 0.103928 | 0.062094 | Other Positive Correlation |
| HLA-DPB1 | 0.103908 | 0.062144 | Other Positive Correlation |
| PROSER1 | 0.103806 | 0.062402 | Other Positive Correlation |
| FAM81A | 0.103805 | 0.062404 | Other Positive Correlation |
| TXNDC16 | 0.103793 | 0.062435 | Other Positive Correlation |
| SRP19 | 0.103788 | 0.062447 | Other Positive Correlation |
| FAM83F | 0.103782 | 0.062464 | Other Positive Correlation |
| PLEKHG6 | 0.103707 | 0.062653 | Other Positive Correlation |
| BEST1 | 0.103686 | 0.062709 | Other Positive Correlation |
| ANKRD10 | 0.103677 | 0.062729 | Other Positive Correlation |
| TMTC2 | 0.103667 | 0.062757 | Other Positive Correlation |

| | | | |
|---|---|---|---|
| MSX2 | 0.103598 | 0.062931 | Other Positive Correlation |
| MBD1 | 0.103572 | 0.062999 | Other Positive Correlation |
| DIS3L | 0.103566 | 0.063014 | Other Positive Correlation |
| ELOVL4 | 0.103521 | 0.063129 | Other Positive Correlation |
| NANOS1 | 0.103463 | 0.063279 | Other Positive Correlation |
| MDK | 0.103453 | 0.063302 | Other Positive Correlation |
| APOLD1 | 0.103452 | 0.063305 | Other Positive Correlation |
| DNAJB6 | 0.103385 | 0.063479 | Other Positive Correlation |
| MME | 0.103365 | 0.06353 | Other Positive Correlation |
| HRH3 | 0.103352 | 0.063563 | Other Positive Correlation |
| CYP11A1 | 0.103343 | 0.063586 | Other Positive Correlation |
| SOAT1 | 0.103313 | 0.063665 | Other Positive Correlation |
| ZNF577 | 0.1033 | 0.063696 | Other Positive Correlation |
| BTBD9 | 0.103287 | 0.063731 | Other Positive Correlation |
| EDA | 0.103275 | 0.063762 | Other Positive Correlation |
| MMD | 0.103226 | 0.063887 | Other Positive Correlation |
| GBX2 | 0.103112 | 0.064185 | Other Positive Correlation |
| ESPN | 0.10308 | 0.064267 | Other Positive Correlation |
| PNRC2 | 0.103061 | 0.064317 | Other Positive Correlation |
| SPHK2 | 0.103002 | 0.064469 | Other Positive Correlation |
| OR1R1P | 0.102973 | 0.064544 | Other Positive Correlation |
| KNDC1 | 0.102922 | 0.06468 | Other Positive Correlation |
| SDF2 | 0.102847 | 0.064876 | Other Positive Correlation |
| GRB2 | 0.102837 | 0.064902 | Other Positive Correlation |
| ZNF230 | 0.102769 | 0.06508 | Other Positive Correlation |
| ZNF684 | 0.102755 | 0.065117 | Other Positive Correlation |
| FAM117A | 0.102601 | 0.065523 | Other Positive Correlation |
| GMCL1 | 0.102589 | 0.065555 | Other Positive Correlation |
| CCSER2 | 0.102579 | 0.06558 | Other Positive Correlation |
| KCNB1 | 0.102573 | 0.065596 | Other Positive Correlation |
| SYTL3 | 0.102552 | 0.065652 | Other Positive Correlation |
| CADM2 | 0.102536 | 0.065693 | Other Positive Correlation |
| SPIN2B | 0.102528 | 0.065714 | Other Positive Correlation |
| PPM1K | 0.102501 | 0.065787 | Other Positive Correlation |
| ENTREP2 | 0.10246 | 0.065896 | Other Positive Correlation |
| LBH | 0.102389 | 0.066085 | Other Positive Correlation |
| TMIE | 0.102384 | 0.066096 | Other Positive Correlation |
| CEP192 | 0.102373 | 0.066127 | Other Positive Correlation |
| AZIN2 | 0.10237 | 0.066133 | Other Positive Correlation |
| SMIM1 | 0.10233 | 0.066241 | Other Positive Correlation |
| EXOC3L4 | 0.102265 | 0.066415 | Other Positive Correlation |
| HS6ST3 | 0.102238 | 0.066487 | Other Positive Correlation |
| HPGD | 0.102216 | 0.066544 | Other Positive Correlation |
| IDH3A | 0.102142 | 0.066743 | Other Positive Correlation |
| FZR1 | 0.102122 | 0.066796 | Other Positive Correlation |
| UBE2J2 | 0.102122 | 0.066798 | Other Positive Correlation |
| DESI2 | 0.10212 | 0.066802 | Other Positive Correlation |
| ZNF140 | 0.102108 | 0.066835 | Other Positive Correlation |

| | | | |
|---|---|---|---|
| NPHS1 | 0.102077 | 0.066918 | Other Positive Correlation |
| SHD | 0.102038 | 0.067024 | Other Positive Correlation |
| CREG1 | 0.102027 | 0.067053 | Other Positive Correlation |
| GPR37 | 0.101921 | 0.067339 | Other Positive Correlation |
| CCDC40 | 0.101856 | 0.067513 | Other Positive Correlation |
| CNOT3 | 0.101817 | 0.06762 | Other Positive Correlation |
| ADGRB1 | 0.101803 | 0.067659 | Other Positive Correlation |
| PICALM | 0.101799 | 0.067668 | Other Positive Correlation |
| PLCE1 | 0.101779 | 0.067721 | Other Positive Correlation |
| LYSMD4 | 0.101761 | 0.067772 | Other Positive Correlation |
| SEMA4C | 0.101744 | 0.067817 | Other Positive Correlation |
| HBA1 | 0.101711 | 0.067907 | Other Positive Correlation |
| PRRX1 | 0.101705 | 0.067925 | Other Positive Correlation |
| PROX1 | 0.1017 | 0.067939 | Other Positive Correlation |
| MOCS2 | 0.101652 | 0.068068 | Other Positive Correlation |
| B3GALT6 | 0.101622 | 0.068151 | Other Positive Correlation |
| TMEM132E | 0.101596 | 0.068221 | Other Positive Correlation |
| FBXO8 | 0.101579 | 0.068267 | Other Positive Correlation |
| ZFP14 | 0.101566 | 0.068302 | Other Positive Correlation |
| GAK | 0.101531 | 0.068397 | Other Positive Correlation |
| NDRG3 | 0.101513 | 0.068447 | Other Positive Correlation |
| RBM42 | 0.101432 | 0.06867 | Other Positive Correlation |
| PFN4 | 0.101357 | 0.068876 | Other Positive Correlation |
| ZNF98 | 0.101341 | 0.068921 | Other Positive Correlation |
| DZIP1L | 0.10128 | 0.069087 | Other Positive Correlation |
| ATF5 | 0.101274 | 0.069105 | Other Positive Correlation |
| GTPBP1 | 0.101237 | 0.069206 | Other Positive Correlation |
| ZBTB7B | 0.101208 | 0.069288 | Other Positive Correlation |
| B2M | 0.101117 | 0.06954 | Other Positive Correlation |
| KCNA1 | 0.10107 | 0.069669 | Other Positive Correlation |
| ZNF468 | 0.101033 | 0.069771 | Other Positive Correlation |
| MYBPC2 | 0.100952 | 0.069998 | Other Positive Correlation |
| ZNF599 | 0.100949 | 0.070006 | Other Positive Correlation |
| ODAD3 | 0.100897 | 0.070151 | Other Positive Correlation |
| BHLHA9 | 0.100868 | 0.070233 | Other Positive Correlation |
| ARHGEF11 | 0.100779 | 0.070481 | Other Positive Correlation |
| RBM4B | 0.100748 | 0.070569 | Other Positive Correlation |
| NUP62CL | 0.100738 | 0.070597 | Other Positive Correlation |
| SMIM24 | 0.100728 | 0.070625 | Other Positive Correlation |
| TP53TG5 | 0.100713 | 0.070665 | Other Positive Correlation |
| EIF2AK1 | 0.100696 | 0.070713 | Other Positive Correlation |
| GLIS3 | 0.100675 | 0.070774 | Other Positive Correlation |
| MBD6 | 0.100635 | 0.070885 | Other Positive Correlation |
| ZFYVE19 | 0.100518 | 0.071215 | Other Positive Correlation |
| LRRC46 | 0.100485 | 0.071309 | Other Positive Correlation |
| CIAPIN1 | 0.100482 | 0.071319 | Other Positive Correlation |
| ELAVL4 | 0.100471 | 0.071349 | Other Positive Correlation |
| SPECC1L | 0.100446 | 0.071419 | Other Positive Correlation |

| | | | |
|---|---|---|---|
| LZTS1 | 0.100411 | 0.07152 | Other Positive Correlation |
| TWIST1 | 0.100404 | 0.07154 | Other Positive Correlation |
| MTF1 | 0.100394 | 0.071567 | Other Positive Correlation |
| TCF24 | 0.100288 | 0.071868 | Other Positive Correlation |
| KLHDC7B | 0.100259 | 0.071953 | Other Positive Correlation |
| SOX17 | 0.100253 | 0.071969 | Other Positive Correlation |
| ZNF639 | 0.10025 | 0.071977 | Other Positive Correlation |
| NRXN2 | 0.100229 | 0.072038 | Other Positive Correlation |
| CCDC33 | 0.100126 | 0.07233 | Other Positive Correlation |
| ARL8B | 0.100086 | 0.072446 | Other Positive Correlation |
| PDXP | 0.100084 | 0.072451 | Other Positive Correlation |
| CNOT9 | 0.10007 | 0.072492 | Other Positive Correlation |
| VAC14 | 0.099953 | 0.072827 | Other Positive Correlation |
| CCDC148 | 0.099886 | 0.073022 | Other Positive Correlation |
| GPR180 | 0.099877 | 0.073048 | Other Positive Correlation |
| PIEZO1 | 0.099868 | 0.073073 | Other Positive Correlation |
| TDP1 | 0.099812 | 0.073234 | Other Positive Correlation |
| SNX10 | 0.099801 | 0.073268 | Other Positive Correlation |
| TEC | 0.099766 | 0.073368 | Other Positive Correlation |
| FOXO6 | 0.099712 | 0.073524 | Other Positive Correlation |
| MBOAT7 | 0.099693 | 0.07358 | Other Positive Correlation |
| RHBDD3 | 0.09968 | 0.073617 | Other Positive Correlation |
| TMEM116 | 0.09965 | 0.073706 | Other Positive Correlation |
| PYGO2 | 0.099632 | 0.073757 | Other Positive Correlation |
| C8orf88 | 0.099613 | 0.073812 | Other Positive Correlation |
| RENBP | 0.099608 | 0.073828 | Other Positive Correlation |
| NANP | 0.0996 | 0.073849 | Other Positive Correlation |
| SLC38A8 | 0.099589 | 0.073884 | Other Positive Correlation |
| BMP2 | 0.099579 | 0.073913 | Other Positive Correlation |
| SLC25A23 | 0.099536 | 0.074036 | Other Positive Correlation |
| GPRIN1 | 0.099526 | 0.074065 | Other Positive Correlation |
| ZNF414 | 0.099508 | 0.074119 | Other Positive Correlation |
| SPSB3 | 0.099493 | 0.074162 | Other Positive Correlation |
| CA3 | 0.099443 | 0.074308 | Other Positive Correlation |
| KCNJ2 | 0.099371 | 0.074519 | Other Positive Correlation |
| C16orf86 | 0.099351 | 0.074579 | Other Positive Correlation |
| SCCPDH | 0.099349 | 0.074586 | Other Positive Correlation |
| DHX16 | 0.099342 | 0.074604 | Other Positive Correlation |
| SPSB1 | 0.099329 | 0.074644 | Other Positive Correlation |
| RFWD3 | 0.099271 | 0.074814 | Other Positive Correlation |
| GLCE | 0.099263 | 0.074839 | Other Positive Correlation |
| SEMA6C | 0.09926 | 0.074845 | Other Positive Correlation |
| SIRPA | 0.099234 | 0.074923 | Other Positive Correlation |
| TDRP | 0.099201 | 0.075019 | Other Positive Correlation |
| ZNF536 | 0.099193 | 0.075045 | Other Positive Correlation |
| RAB3IP | 0.099191 | 0.075048 | Other Positive Correlation |
| PASK | 0.099044 | 0.075486 | Other Positive Correlation |
| HRC | 0.099011 | 0.075583 | Other Positive Correlation |

| | | | |
|---|---|---|---|
| COL2A1 | 0.098977 | 0.075682 | Other Positive Correlation |
| CCDC82 | 0.098964 | 0.075722 | Other Positive Correlation |
| SLC15A3 | 0.098959 | 0.075737 | Other Positive Correlation |
| PCYT1B | 0.098945 | 0.075777 | Other Positive Correlation |
| RPP25L | 0.098927 | 0.075831 | Other Positive Correlation |
| INPP5B | 0.098883 | 0.075962 | Other Positive Correlation |
| FAM53C | 0.09888 | 0.075972 | Other Positive Correlation |
| PRTG | 0.09887 | 0.076001 | Other Positive Correlation |
| NFKBIL1 | 0.098858 | 0.076038 | Other Positive Correlation |
| SLC6A6 | 0.098831 | 0.076118 | Other Positive Correlation |
| EFNB3 | 0.098779 | 0.076272 | Other Positive Correlation |
| SPSB4 | 0.098739 | 0.076393 | Other Positive Correlation |
| SCN9A | 0.098735 | 0.076406 | Other Positive Correlation |
| ANKRD52 | 0.098673 | 0.076592 | Other Positive Correlation |
| ENSG00000267385 | 0.098652 | 0.076655 | Other Positive Correlation |
| SPON2 | 0.098643 | 0.076682 | Other Positive Correlation |
| CDC34 | 0.098613 | 0.076771 | Other Positive Correlation |
| AKR1B1 | 0.098602 | 0.076804 | Other Positive Correlation |
| DLEU7 | 0.098579 | 0.076872 | Other Positive Correlation |
| ZNF549 | 0.098559 | 0.076934 | Other Positive Correlation |
| SULT2B1 | 0.098536 | 0.077001 | Other Positive Correlation |
| FAM50B | 0.098518 | 0.077056 | Other Positive Correlation |
| CNTN4 | 0.098513 | 0.077071 | Other Positive Correlation |
| ACVRL1 | 0.098513 | 0.077071 | Other Positive Correlation |
| CTNND2 | 0.098513 | 0.077072 | Other Positive Correlation |
| EPM2AIP1 | 0.098498 | 0.077118 | Other Positive Correlation |
| MAGIX | 0.09843 | 0.077323 | Other Positive Correlation |
| SLC16A6 | 0.098425 | 0.077337 | Other Positive Correlation |
| ZCCHC24 | 0.0984 | 0.077414 | Other Positive Correlation |
| FBXO10 | 0.098366 | 0.077515 | Other Positive Correlation |
| CNEP1R1 | 0.098311 | 0.077682 | Other Positive Correlation |
| MXI1 | 0.098275 | 0.077793 | Other Positive Correlation |
| BVES | 0.098272 | 0.0778 | Other Positive Correlation |
| BHLHA15 | 0.098258 | 0.077844 | Other Positive Correlation |
| LEMD2 | 0.098257 | 0.077846 | Other Positive Correlation |
| PLCH1 | 0.098246 | 0.07788 | Other Positive Correlation |
| TFAP2E | 0.098187 | 0.078059 | Other Positive Correlation |
| MRAP2 | 0.098151 | 0.07817 | Other Positive Correlation |
| DNAJA3 | 0.098109 | 0.078299 | Other Positive Correlation |
| GPR27 | 0.098082 | 0.078381 | Other Positive Correlation |
| C2orf81 | 0.098078 | 0.078394 | Other Positive Correlation |
| SLC19A2 | 0.098076 | 0.078397 | Other Positive Correlation |
| CA13 | 0.098076 | 0.0784 | Other Positive Correlation |
| C2CD4D | 0.098074 | 0.078404 | Other Positive Correlation |
| PDHX | 0.098054 | 0.078467 | Other Positive Correlation |
| TNKS2 | 0.097988 | 0.078669 | Other Positive Correlation |
| LRIF1 | 0.097982 | 0.078685 | Other Positive Correlation |
| CWF19L2 | 0.097974 | 0.078712 | Other Positive Correlation |

| | | | |
|---|---|---|---|
| FAM118B | 0.097886 | 0.078981 | Other Positive Correlation |
| SOX5 | 0.097848 | 0.079099 | Other Positive Correlation |
| ESAM | 0.097846 | 0.079106 | Other Positive Correlation |
| DDX25 | 0.097829 | 0.079156 | Other Positive Correlation |
| CKAP5 | 0.097817 | 0.079193 | Other Positive Correlation |
| HPCA | 0.097802 | 0.079241 | Other Positive Correlation |
| CAMK2A | 0.097759 | 0.079371 | Other Positive Correlation |
| CCDC187 | 0.09774 | 0.079433 | Other Positive Correlation |
| CYS1 | 0.097738 | 0.079436 | Other Positive Correlation |
| LIG4 | 0.097697 | 0.079564 | Other Positive Correlation |
| CCKBR | 0.09768 | 0.079617 | Other Positive Correlation |
| OCRL | 0.097667 | 0.079658 | Other Positive Correlation |
| ATG9B | 0.097638 | 0.079747 | Other Positive Correlation |
| FBXO25 | 0.097635 | 0.079756 | Other Positive Correlation |
| C1orf210 | 0.097625 | 0.079787 | Other Positive Correlation |
| LHX1 | 0.097617 | 0.079813 | Other Positive Correlation |
| MARCHF8 | 0.097606 | 0.079845 | Other Positive Correlation |
| RASGRF2 | 0.097582 | 0.079922 | Other Positive Correlation |
| DDAH1 | 0.097572 | 0.079952 | Other Positive Correlation |
| KRAS | 0.097532 | 0.080078 | Other Positive Correlation |
| MAT1A | 0.097467 | 0.080279 | Other Positive Correlation |
| RASGRP1 | 0.097436 | 0.080376 | Other Positive Correlation |
| PTDSS2 | 0.097426 | 0.080405 | Other Positive Correlation |
| SNRK | 0.097412 | 0.08045 | Other Positive Correlation |
| SKIC2 | 0.097353 | 0.080633 | Other Positive Correlation |
| IL1RL2 | 0.097324 | 0.080724 | Other Positive Correlation |
| ADRB1 | 0.097324 | 0.080725 | Other Positive Correlation |
| SECTM1 | 0.097299 | 0.080805 | Other Positive Correlation |
| ARHGAP12 | 0.09729 | 0.080833 | Other Positive Correlation |
| UBE2E2 | 0.097282 | 0.080856 | Other Positive Correlation |
| CCNP | 0.097229 | 0.081024 | Other Positive Correlation |
| FOXD2 | 0.097155 | 0.081257 | Other Positive Correlation |
| RAB9B | 0.097147 | 0.08128 | Other Positive Correlation |
| MLYCD | 0.097119 | 0.08137 | Other Positive Correlation |
| TMUB2 | 0.097113 | 0.081388 | Other Positive Correlation |
| SIM1 | 0.097112 | 0.081391 | Other Positive Correlation |
| AGTR1 | 0.097095 | 0.081445 | Other Positive Correlation |
| PARP2 | 0.097075 | 0.081507 | Other Positive Correlation |
| GAST | 0.09704 | 0.081619 | Other Positive Correlation |
| TMEM127 | 0.097007 | 0.081723 | Other Positive Correlation |
| CDKL5 | 0.096988 | 0.081784 | Other Positive Correlation |
| APOBEC3B | 0.096976 | 0.081821 | Other Positive Correlation |
| PEX6 | 0.096962 | 0.081864 | Other Positive Correlation |
| PDIK1L | 0.096892 | 0.082088 | Other Positive Correlation |
| ZNF365 | 0.096866 | 0.082168 | Other Positive Correlation |
| LIMK1 | 0.09685 | 0.082219 | Other Positive Correlation |
| OTUD1 | 0.096792 | 0.082404 | Other Positive Correlation |
| FRAT1 | 0.09679 | 0.082412 | Other Positive Correlation |

| | | | |
|---|---|---|---|
| CD40 | 0.096762 | 0.082499 | Other Positive Correlation |
| FAM234A | 0.096758 | 0.082514 | Other Positive Correlation |
| GUCY1A2 | 0.096704 | 0.082686 | Other Positive Correlation |
| MIOS | 0.096701 | 0.082693 | Other Positive Correlation |
| TMEFF1 | 0.096634 | 0.08291 | Other Positive Correlation |
| UTP25 | 0.096594 | 0.083035 | Other Positive Correlation |
| EIF4E1B | 0.09653 | 0.083241 | Other Positive Correlation |
| LRWD1 | 0.096481 | 0.0834 | Other Positive Correlation |
| SNW1 | 0.09647 | 0.083436 | Other Positive Correlation |
| BBS9 | 0.096465 | 0.083451 | Other Positive Correlation |
| MICB | 0.096458 | 0.083473 | Other Positive Correlation |
| LEFTY1 | 0.096445 | 0.083516 | Other Positive Correlation |
| TM6SF2 | 0.09641 | 0.083629 | Other Positive Correlation |
| ZIC1 | 0.096355 | 0.083804 | Other Positive Correlation |
| SMIM34 | 0.096347 | 0.08383 | Other Positive Correlation |
| TSPAN15 | 0.096339 | 0.083857 | Other Positive Correlation |
| AMN1 | 0.096316 | 0.083931 | Other Positive Correlation |
| COL1A1 | 0.096314 | 0.083939 | Other Positive Correlation |
| SYPL2 | 0.096266 | 0.084091 | Other Positive Correlation |
| ADAM19 | 0.096238 | 0.084182 | Other Positive Correlation |
| DSEL | 0.096238 | 0.084182 | Other Positive Correlation |
| CFD | 0.096221 | 0.084239 | Other Positive Correlation |
| PAQR9 | 0.096182 | 0.084365 | Other Positive Correlation |
| MTMR9 | 0.096139 | 0.084505 | Other Positive Correlation |
| CREM | 0.096121 | 0.084562 | Other Positive Correlation |
| C17orf100 | 0.096096 | 0.084642 | Other Positive Correlation |
| LRRC23 | 0.096026 | 0.084872 | Other Positive Correlation |
| AEN | 0.096022 | 0.084886 | Other Positive Correlation |
| RNF150 | 0.09597 | 0.085055 | Other Positive Correlation |
| LHX9 | 0.095927 | 0.085196 | Other Positive Correlation |
| MARCHF2 | 0.095923 | 0.085208 | Other Positive Correlation |
| VWA5B2 | 0.095921 | 0.085215 | Other Positive Correlation |
| FSTL4 | 0.095918 | 0.085224 | Other Positive Correlation |
| LNX1 | 0.095898 | 0.085289 | Other Positive Correlation |
| GALR2 | 0.095894 | 0.085305 | Other Positive Correlation |
| ENDOU | 0.095865 | 0.085399 | Other Positive Correlation |
| PWWP2B | 0.095849 | 0.085451 | Other Positive Correlation |
| LBX1 | 0.095816 | 0.085558 | Other Positive Correlation |
| EXOC4 | 0.095808 | 0.085584 | Other Positive Correlation |
| MCM8 | 0.095798 | 0.085619 | Other Positive Correlation |
| SERAC1 | 0.095752 | 0.085769 | Other Positive Correlation |
| ZNF75D | 0.09569 | 0.085971 | Other Positive Correlation |
| E2F8 | 0.095675 | 0.086023 | Other Positive Correlation |
| SRGAP2C | 0.095655 | 0.086087 | Other Positive Correlation |
| TMEM60 | 0.095595 | 0.086286 | Other Positive Correlation |
| ASH2L | 0.095571 | 0.086365 | Other Positive Correlation |
| HOXD4 | 0.095543 | 0.086457 | Other Positive Correlation |
| ZNF724 | 0.095499 | 0.086605 | Other Positive Correlation |

| | | | |
|---|---|---|---|
| LRRC8B | 0.09546 | 0.086733 | Other Positive Correlation |
| HINFP | 0.09546 | 0.086733 | Other Positive Correlation |
| CACNG5 | 0.095446 | 0.08678 | Other Positive Correlation |
| SYT14 | 0.095432 | 0.086827 | Other Positive Correlation |
| IRX3 | 0.095391 | 0.086961 | Other Positive Correlation |
| CASKIN1 | 0.09539 | 0.086966 | Other Positive Correlation |
| SLC41A1 | 0.095383 | 0.086988 | Other Positive Correlation |
| CSGALNACT2 | 0.095369 | 0.087034 | Other Positive Correlation |
| PLIN5 | 0.095344 | 0.087119 | Other Positive Correlation |
| LGMN | 0.09531 | 0.087232 | Other Positive Correlation |
| KCNS1 | 0.095286 | 0.087311 | Other Positive Correlation |
| ATG9A | 0.095269 | 0.087368 | Other Positive Correlation |
| GBA1 | 0.095249 | 0.087437 | Other Positive Correlation |
| RNF4 | 0.095179 | 0.08767 | Other Positive Correlation |
| IGSF9B | 0.09514 | 0.087799 | Other Positive Correlation |
| HEYL | 0.095123 | 0.087856 | Other Positive Correlation |
| IGLON5 | 0.095103 | 0.087923 | Other Positive Correlation |
| SNX11 | 0.095072 | 0.088028 | Other Positive Correlation |
| LTBR | 0.094997 | 0.08828 | Other Positive Correlation |
| ENSG00000285085 | 0.094979 | 0.088339 | Other Positive Correlation |
| SPATA24 | 0.094961 | 0.088399 | Other Positive Correlation |
| SLC35D3 | 0.094942 | 0.088466 | Other Positive Correlation |
| SMARCC2 | 0.094893 | 0.08863 | Other Positive Correlation |
| PSMG3 | 0.094886 | 0.088654 | Other Positive Correlation |
| CDH22 | 0.094886 | 0.088654 | Other Positive Correlation |
| CDK5 | 0.09486 | 0.08874 | Other Positive Correlation |
| MMP25 | 0.094844 | 0.088797 | Other Positive Correlation |
| HSPA12A | 0.094783 | 0.089001 | Other Positive Correlation |
| PLBD1 | 0.09477 | 0.089045 | Other Positive Correlation |
| TTI1 | 0.094768 | 0.089053 | Other Positive Correlation |
| NKX6-1 | 0.094698 | 0.08929 | Other Positive Correlation |
| KIAA0319 | 0.094604 | 0.08961 | Other Positive Correlation |
| BEND3 | 0.094553 | 0.089784 | Other Positive Correlation |
| TMEM205 | 0.094552 | 0.089786 | Other Positive Correlation |
| LMO1 | 0.094547 | 0.089803 | Other Positive Correlation |
| SLC5A5 | 0.09452 | 0.089896 | Other Positive Correlation |
| KHDC4 | 0.094511 | 0.089927 | Other Positive Correlation |
| ANGPTL2 | 0.094477 | 0.090042 | Other Positive Correlation |
| ZNF253 | 0.094436 | 0.090184 | Other Positive Correlation |
| CYREN | 0.094416 | 0.090252 | Other Positive Correlation |
| WDR72 | 0.094405 | 0.090288 | Other Positive Correlation |
| ITGA1 | 0.094368 | 0.090415 | Other Positive Correlation |
| PAPPA | 0.094282 | 0.090711 | Other Positive Correlation |
| SMARCA2 | 0.094281 | 0.090715 | Other Positive Correlation |
| PDGFRB | 0.094263 | 0.090777 | Other Positive Correlation |
| ZNF680 | 0.094261 | 0.090783 | Other Positive Correlation |
| BRPF1 | 0.094234 | 0.090875 | Other Positive Correlation |
| WNT3 | 0.094219 | 0.09093 | Other Positive Correlation |

| | | | |
|---|---|---|---|
| CHD1 | 0.094202 | 0.090987 | Other Positive Correlation |
| NKX2-6 | 0.09418 | 0.091064 | Other Positive Correlation |
| ANKRD49 | 0.094176 | 0.091076 | Other Positive Correlation |
| SLC25A29 | 0.094132 | 0.091229 | Other Positive Correlation |
| FZD7 | 0.094113 | 0.091294 | Other Positive Correlation |
| FCSK | 0.094105 | 0.091322 | Other Positive Correlation |
| LMNTD2 | 0.09403 | 0.091581 | Other Positive Correlation |
| VIP | 0.094024 | 0.091602 | Other Positive Correlation |
| EAF2 | 0.093993 | 0.09171 | Other Positive Correlation |
| FOXI3 | 0.093932 | 0.09192 | Other Positive Correlation |
| CYP26C1 | 0.093931 | 0.091925 | Other Positive Correlation |
| TREML2 | 0.093861 | 0.09217 | Other Positive Correlation |
| FAF1 | 0.093854 | 0.092194 | Other Positive Correlation |
| CCDC184 | 0.093835 | 0.092259 | Other Positive Correlation |
| PELO | 0.093785 | 0.092434 | Other Positive Correlation |
| ALPK2 | 0.093766 | 0.092499 | Other Positive Correlation |
| PET117 | 0.093735 | 0.092608 | Other Positive Correlation |
| SALL4 | 0.09373 | 0.092624 | Other Positive Correlation |
| EZH1 | 0.093719 | 0.092665 | Other Positive Correlation |
| PTGES | 0.093626 | 0.09299 | Other Positive Correlation |
| DCLK2 | 0.093602 | 0.093073 | Other Positive Correlation |
| LRFN3 | 0.093529 | 0.093332 | Other Positive Correlation |
| ABHD13 | 0.09348 | 0.093503 | Other Positive Correlation |
| ANOS1 | 0.093463 | 0.093562 | Other Positive Correlation |
| PEX13 | 0.093455 | 0.09359 | Other Positive Correlation |
| TIMM8A | 0.093445 | 0.093628 | Other Positive Correlation |
| NREP | 0.093439 | 0.093648 | Other Positive Correlation |
| HS6ST1 | 0.093407 | 0.093762 | Other Positive Correlation |
| UPRT | 0.093404 | 0.09377 | Other Positive Correlation |
| SMIM4 | 0.093353 | 0.09395 | Other Positive Correlation |
| SMUG1 | 0.093344 | 0.093984 | Other Positive Correlation |
| KCNT2 | 0.093261 | 0.094277 | Other Positive Correlation |
| RASL10B | 0.093208 | 0.094465 | Other Positive Correlation |
| LBX2 | 0.093188 | 0.094537 | Other Positive Correlation |
| CSGALNACT1 | 0.093177 | 0.094576 | Other Positive Correlation |
| GAL3ST1 | 0.093168 | 0.094608 | Other Positive Correlation |
| SARM1 | 0.093154 | 0.094658 | Other Positive Correlation |
| LDLRAD4 | 0.093057 | 0.095002 | Other Positive Correlation |
| UNC5A | 0.093041 | 0.095061 | Other Positive Correlation |
| ZNF470 | 0.092978 | 0.095284 | Other Positive Correlation |
| GPAT2 | 0.092968 | 0.095319 | Other Positive Correlation |
| ADRA1D | 0.092964 | 0.095335 | Other Positive Correlation |
| ZNF528 | 0.092955 | 0.095368 | Other Positive Correlation |
| DEF8 | 0.092924 | 0.095477 | Other Positive Correlation |
| KIFBP | 0.092905 | 0.095546 | Other Positive Correlation |
| CASP9 | 0.09287 | 0.095671 | Other Positive Correlation |
| NLGN1 | 0.092814 | 0.095871 | Other Positive Correlation |
| SPRY3 | 0.092809 | 0.095889 | Other Positive Correlation |

| | | | |
|---|---|---|---|
| HCFC1 | 0.092805 | 0.095904 | Other Positive Correlation |
| KIF26B | 0.0928 | 0.095924 | Other Positive Correlation |
| MAP3K15 | 0.092791 | 0.095955 | Other Positive Correlation |
| LYL1 | 0.092785 | 0.095976 | Other Positive Correlation |
| SEMA6A | 0.092767 | 0.096042 | Other Positive Correlation |
| LGR6 | 0.092738 | 0.096145 | Other Positive Correlation |
| ZG16 | 0.092694 | 0.096304 | Other Positive Correlation |
| SLF2 | 0.09267 | 0.096391 | Other Positive Correlation |
| CTSV | 0.092647 | 0.096473 | Other Positive Correlation |
| HAPLN2 | 0.09264 | 0.0965 | Other Positive Correlation |
| CPQ | 0.092601 | 0.096639 | Other Positive Correlation |
| UBAC2 | 0.092585 | 0.096699 | Other Positive Correlation |
| IFNL4 | 0.092571 | 0.096749 | Other Positive Correlation |
| GIGYF1 | 0.092569 | 0.096756 | Other Positive Correlation |
| PPP2R5A | 0.092535 | 0.09688 | Other Positive Correlation |
| ADGRG2 | 0.092512 | 0.096964 | Other Positive Correlation |
| ASB6 | 0.092503 | 0.096994 | Other Positive Correlation |
| SH3GL3 | 0.09247 | 0.097115 | Other Positive Correlation |
| MPHOSPH10 | 0.092461 | 0.097149 | Other Positive Correlation |
| RADX | 0.09246 | 0.097152 | Other Positive Correlation |
| CIAO1 | 0.09245 | 0.097186 | Other Positive Correlation |
| SLC25A11 | 0.092349 | 0.097555 | Other Positive Correlation |
| SMIM45 | 0.092332 | 0.097618 | Other Positive Correlation |
| SNX21 | 0.092321 | 0.097658 | Other Positive Correlation |
| CACFD1 | 0.092321 | 0.097659 | Other Positive Correlation |
| XPO7 | 0.092308 | 0.097704 | Other Positive Correlation |
| RALGAPA1 | 0.092276 | 0.09782 | Other Positive Correlation |
| MUC20 | 0.092224 | 0.098011 | Other Positive Correlation |
| SLC4A4 | 0.092222 | 0.098019 | Other Positive Correlation |
| MT-CYB | 0.092157 | 0.098259 | Other Positive Correlation |
| PWWP3B | 0.092111 | 0.098424 | Other Positive Correlation |
| MINPP1 | 0.092063 | 0.098603 | Other Positive Correlation |
| RASSF3 | 0.092034 | 0.098708 | Other Positive Correlation |
| ROR1 | 0.092015 | 0.09878 | Other Positive Correlation |
| CPT1A | 0.091982 | 0.0989 | Other Positive Correlation |
| CYB5RL | 0.091807 | 0.099546 | Other Positive Correlation |
| ZNF10 | 0.091801 | 0.099568 | Other Positive Correlation |
| B3GNT8 | 0.091771 | 0.099679 | Other Positive Correlation |
| TSEN15 | 0.091698 | 0.099951 | Other Positive Correlation |
| CLSTN2 | 0.091696 | 0.099958 | Other Positive Correlation |
| SMIM10L2B | 0.091651 | 0.100126 | Other Positive Correlation |
| TMEM117 | 0.091614 | 0.100263 | Other Positive Correlation |
| DCAF12L1 | 0.09161 | 0.100277 | Other Positive Correlation |
| FOXL2NB | 0.091569 | 0.100429 | Other Positive Correlation |
| C15orf61 | 0.091537 | 0.100552 | Other Positive Correlation |
| AKAP17A | 0.09144 | 0.100913 | Other Positive Correlation |
| PSTK | 0.09139 | 0.101101 | Other Positive Correlation |
| TBCE.1 | 0.091366 | 0.10119 | Other Positive Correlation |

| | | | |
|---|---|---|---|
| H3C10 | 0.091295 | 0.101458 | Other Positive Correlation |
| EN1 | 0.091237 | 0.101676 | Other Positive Correlation |
| HPN | 0.091212 | 0.101769 | Other Positive Correlation |
| ENSG00000285526 | 0.091202 | 0.101805 | Other Positive Correlation |
| TMUB1 | 0.091191 | 0.101849 | Other Positive Correlation |
| RITA1 | 0.091177 | 0.101901 | Other Positive Correlation |
| ANKS1B | 0.091166 | 0.101944 | Other Positive Correlation |
| NEGR1 | 0.091148 | 0.10201 | Other Positive Correlation |
| TRAF3IP1 | 0.091131 | 0.102075 | Other Positive Correlation |
| SLC12A7 | 0.091112 | 0.102146 | Other Positive Correlation |
| PYDC1 | 0.091075 | 0.102287 | Other Positive Correlation |
| ZNF728 | 0.09107 | 0.102304 | Other Positive Correlation |
| SOX10 | 0.09104 | 0.102419 | Other Positive Correlation |
| HAND1 | 0.09104 | 0.102419 | Other Positive Correlation |
| AURKAIP1 | 0.091025 | 0.102474 | Other Positive Correlation |
| ZFHX2 | 0.091015 | 0.102514 | Other Positive Correlation |
| ZNF394 | 0.091001 | 0.102565 | Other Positive Correlation |
| ZNF418 | 0.090988 | 0.102617 | Other Positive Correlation |
| MEGF9 | 0.090983 | 0.102634 | Other Positive Correlation |
| NCDN | 0.090931 | 0.102831 | Other Positive Correlation |
| CHST13 | 0.090883 | 0.103015 | Other Positive Correlation |
| SLC6A2 | 0.090876 | 0.10304 | Other Positive Correlation |
| ATG4A | 0.090857 | 0.103115 | Other Positive Correlation |
| TMEM51 | 0.090853 | 0.103127 | Other Positive Correlation |
| RGS14 | 0.090818 | 0.103262 | Other Positive Correlation |
| CCDC157 | 0.090812 | 0.103284 | Other Positive Correlation |
| PRIMA1 | 0.090794 | 0.103354 | Other Positive Correlation |
| ARRB2 | 0.090724 | 0.10362 | Other Positive Correlation |
| LRRN2 | 0.090714 | 0.103659 | Other Positive Correlation |
| RBM48 | 0.090707 | 0.103687 | Other Positive Correlation |
| PURA | 0.090704 | 0.103699 | Other Positive Correlation |
| GATC | 0.090669 | 0.103831 | Other Positive Correlation |
| UST | 0.090651 | 0.103899 | Other Positive Correlation |
| MPPED2 | 0.090553 | 0.104276 | Other Positive Correlation |
| HSD17B6 | 0.090547 | 0.1043 | Other Positive Correlation |
| FGF18 | 0.090536 | 0.104341 | Other Positive Correlation |
| LSM11 | 0.090524 | 0.104388 | Other Positive Correlation |
| NPR1 | 0.090522 | 0.104395 | Other Positive Correlation |
| GDNF | 0.090493 | 0.104508 | Other Positive Correlation |
| CPXM2 | 0.090471 | 0.104593 | Other Positive Correlation |
| MXD3 | 0.0904 | 0.104867 | Other Positive Correlation |
| ST6GAL2 | 0.090281 | 0.105325 | Other Positive Correlation |
| LRRC61 | 0.090266 | 0.105384 | Other Positive Correlation |
| PRUNE2 | 0.090234 | 0.105506 | Other Positive Correlation |
| PABIR3 | 0.090141 | 0.105867 | Other Positive Correlation |
| NPC2 | 0.09014 | 0.105872 | Other Positive Correlation |
| ABHD10 | 0.090104 | 0.106014 | Other Positive Correlation |
| MAP3K8 | 0.090066 | 0.106159 | Other Positive Correlation |

| | | | |
|---|---|---|---|
| FGF17 | 0.090043 | 0.106251 | Other Positive Correlation |
| LRRK2 | 0.08999 | 0.106456 | Other Positive Correlation |
| HPF1 | 0.089989 | 0.106463 | Other Positive Correlation |
| WNT5B | 0.089925 | 0.10671 | Other Positive Correlation |
| EOLA2 | 0.089879 | 0.106891 | Other Positive Correlation |
| ST6GALNAC3 | 0.089864 | 0.10695 | Other Positive Correlation |
| E2F7 | 0.089863 | 0.106952 | Other Positive Correlation |
| ZNF670 | 0.089855 | 0.106984 | Other Positive Correlation |
| SRCIN1 | 0.089815 | 0.107143 | Other Positive Correlation |
| PPP1R15B | 0.089777 | 0.107291 | Other Positive Correlation |
| SDR42E2 | 0.089765 | 0.107339 | Other Positive Correlation |
| KBTBD12 | 0.089744 | 0.107419 | Other Positive Correlation |
| EFNA3 | 0.089727 | 0.107488 | Other Positive Correlation |
| GLCCI1 | 0.089705 | 0.107574 | Other Positive Correlation |
| ZBTB17 | 0.089634 | 0.107854 | Other Positive Correlation |
| COA7 | 0.089604 | 0.107972 | Other Positive Correlation |
| PRRT1B | 0.089599 | 0.107994 | Other Positive Correlation |
| GLT1D1 | 0.089587 | 0.108039 | Other Positive Correlation |
| CHRNB4 | 0.089501 | 0.10838 | Other Positive Correlation |
| UPF3B | 0.089478 | 0.108471 | Other Positive Correlation |
| DUSP28 | 0.089438 | 0.10863 | Other Positive Correlation |
| IRX6 | 0.08938 | 0.108861 | Other Positive Correlation |
| TBC1D15 | 0.08936 | 0.108941 | Other Positive Correlation |
| TRIM31 | 0.089352 | 0.108972 | Other Positive Correlation |
| UBXN11 | 0.089345 | 0.108997 | Other Positive Correlation |
| BARHL2 | 0.089339 | 0.109024 | Other Positive Correlation |
| HLA-DRB5 | 0.089323 | 0.109085 | Other Positive Correlation |
| EIF1AD | 0.089296 | 0.109193 | Other Positive Correlation |
| PLAGL1 | 0.089265 | 0.109318 | Other Positive Correlation |
| MFSD12 | 0.089253 | 0.109364 | Other Positive Correlation |
| ZBTB26 | 0.089248 | 0.109384 | Other Positive Correlation |
| AIFM2 | 0.089244 | 0.1094 | Other Positive Correlation |
| PLCG1 | 0.089216 | 0.109514 | Other Positive Correlation |
| SYNE3 | 0.089156 | 0.109753 | Other Positive Correlation |
| DBP | 0.089154 | 0.10976 | Other Positive Correlation |
| FN3KRP | 0.089109 | 0.109943 | Other Positive Correlation |
| FAM222A | 0.089088 | 0.110025 | Other Positive Correlation |
| ENSG00000288695 | 0.089088 | 0.110026 | Other Positive Correlation |
| RNLS | 0.089086 | 0.110035 | Other Positive Correlation |
| SAFB2 | 0.089049 | 0.110183 | Other Positive Correlation |
| PTGES3L | 0.089046 | 0.110192 | Other Positive Correlation |
| KCTD8 | 0.089034 | 0.110243 | Other Positive Correlation |
| TMEM61 | 0.089025 | 0.110277 | Other Positive Correlation |
| PRKCE | 0.089006 | 0.110355 | Other Positive Correlation |
| HES5 | 0.088982 | 0.110449 | Other Positive Correlation |
| ANKRD30BL | 0.088967 | 0.110513 | Other Positive Correlation |
| ZBTB46 | 0.088965 | 0.110519 | Other Positive Correlation |
| ZNF697 | 0.088941 | 0.110617 | Other Positive Correlation |

| | | | |
|---|---|---|---|
| SOS1 | 0.088926 | 0.110675 | Other Positive Correlation |
| FAAH | 0.088922 | 0.110694 | Other Positive Correlation |
| NR5A1 | 0.088873 | 0.11089 | Other Positive Correlation |
| KCNQ4 | 0.088856 | 0.110957 | Other Positive Correlation |
| CNTNAP2 | 0.088824 | 0.111087 | Other Positive Correlation |
| POU2F3 | 0.088817 | 0.111116 | Other Positive Correlation |
| ICAM3 | 0.088813 | 0.111133 | Other Positive Correlation |
| SLC25A21 | 0.088807 | 0.111154 | Other Positive Correlation |
| C1orf122 | 0.088781 | 0.11126 | Other Positive Correlation |
| MIS12 | 0.088766 | 0.111324 | Other Positive Correlation |
| CLCN7 | 0.088762 | 0.111337 | Other Positive Correlation |
| TMCO3 | 0.088735 | 0.111445 | Other Positive Correlation |
| RGS17 | 0.088682 | 0.111661 | Other Positive Correlation |
| GPX3 | 0.088658 | 0.11176 | Other Positive Correlation |
| GRIK5 | 0.08861 | 0.111953 | Other Positive Correlation |
| KLF9 | 0.088599 | 0.111999 | Other Positive Correlation |
| MKS1 | 0.08854 | 0.112239 | Other Positive Correlation |
| SLC35E4 | 0.088505 | 0.112382 | Other Positive Correlation |
| ISLR2 | 0.088485 | 0.112463 | Other Positive Correlation |
| METTL24 | 0.08847 | 0.112524 | Other Positive Correlation |
| MAP2 | 0.088427 | 0.1127 | Other Positive Correlation |
| ALDH5A1 | 0.088392 | 0.112845 | Other Positive Correlation |
| WDR33 | 0.088359 | 0.112979 | Other Positive Correlation |
| PADI2 | 0.088339 | 0.11306 | Other Positive Correlation |
| DLGAP1 | 0.088324 | 0.113122 | Other Positive Correlation |
| JKAMP | 0.088287 | 0.113273 | Other Positive Correlation |
| CMTR2 | 0.088287 | 0.113274 | Other Positive Correlation |
| SYT6 | 0.088256 | 0.113399 | Other Positive Correlation |
| SORCS1 | 0.088251 | 0.113421 | Other Positive Correlation |
| STAT5B | 0.088187 | 0.113685 | Other Positive Correlation |
| UBL3 | 0.088125 | 0.113936 | Other Positive Correlation |
| CYP46A1 | 0.088122 | 0.11395 | Other Positive Correlation |
| RAPGEF5 | 0.0881 | 0.114042 | Other Positive Correlation |
| P2RX7 | 0.088083 | 0.11411 | Other Positive Correlation |
| TIGD2 | 0.088073 | 0.114151 | Other Positive Correlation |
| DNAAF2 | 0.088043 | 0.114278 | Other Positive Correlation |
| NGDN | 0.088014 | 0.114397 | Other Positive Correlation |
| ZNF773 | 0.087997 | 0.114468 | Other Positive Correlation |
| NGFR | 0.087995 | 0.114477 | Other Positive Correlation |
| CA7 | 0.08799 | 0.114493 | Other Positive Correlation |
| MTTP | 0.087863 | 0.115023 | Other Positive Correlation |
| PIP5K1C | 0.087862 | 0.115026 | Other Positive Correlation |
| MACROD2 | 0.087831 | 0.115156 | Other Positive Correlation |
| ACBD3 | 0.087811 | 0.115236 | Other Positive Correlation |
| HOXD11 | 0.087784 | 0.115348 | Other Positive Correlation |
| FRMD3 | 0.087763 | 0.115435 | Other Positive Correlation |
| ACER3 | 0.087762 | 0.115442 | Other Positive Correlation |
| SUSD4 | 0.087717 | 0.115627 | Other Positive Correlation |

| | | | |
|---|---|---|---|
| GCKR | 0.087695 | 0.115719 | Other Positive Correlation |
| UBXN7 | 0.08766 | 0.115866 | Other Positive Correlation |
| ATOSB | 0.087575 | 0.116221 | Other Positive Correlation |
| ENSG00000288637 | 0.087554 | 0.116308 | Other Positive Correlation |
| PRLR | 0.087429 | 0.116831 | Other Positive Correlation |
| NIPA1 | 0.087406 | 0.116931 | Other Positive Correlation |
| RND1 | 0.087373 | 0.117067 | Other Positive Correlation |
| TRAPPC4 | 0.087314 | 0.117317 | Other Positive Correlation |
| AGFG2 | 0.087314 | 0.117319 | Other Positive Correlation |
| ST3GAL2 | 0.087248 | 0.117597 | Other Positive Correlation |
| CALU | 0.087223 | 0.1177 | Other Positive Correlation |
| RAB26 | 0.087211 | 0.117754 | Other Positive Correlation |
| AKT3 | 0.087179 | 0.117886 | Other Positive Correlation |
| ARL16 | 0.087167 | 0.117939 | Other Positive Correlation |
| CAPN7 | 0.087164 | 0.117949 | Other Positive Correlation |
| AZIN1 | 0.08715 | 0.118011 | Other Positive Correlation |
| ZNF816 | 0.087139 | 0.118055 | Other Positive Correlation |
| DERL2 | 0.087102 | 0.118214 | Other Positive Correlation |
| MICU1 | 0.087101 | 0.118216 | Other Positive Correlation |
| MTHFD2L | 0.087094 | 0.118247 | Other Positive Correlation |
| SMIM12 | 0.08709 | 0.118263 | Other Positive Correlation |
| FRS3 | 0.087089 | 0.118269 | Other Positive Correlation |
| CYBC1 | 0.087083 | 0.118295 | Other Positive Correlation |
| P4HTM | 0.086975 | 0.118753 | Other Positive Correlation |
| SOCS6 | 0.086966 | 0.118789 | Other Positive Correlation |
| BEND4 | 0.086954 | 0.118841 | Other Positive Correlation |
| TERF1 | 0.086892 | 0.119108 | Other Positive Correlation |
| FBXO42 | 0.086835 | 0.11935 | Other Positive Correlation |
| STK32C | 0.086819 | 0.11942 | Other Positive Correlation |
| SLC8A2 | 0.086775 | 0.119606 | Other Positive Correlation |
| RASGEF1B | 0.08675 | 0.119714 | Other Positive Correlation |
| SPEN | 0.086728 | 0.119809 | Other Positive Correlation |
| MED9 | 0.086721 | 0.119839 | Other Positive Correlation |
| CABP1 | 0.086697 | 0.11994 | Other Positive Correlation |
| DNAH6 | 0.086654 | 0.120123 | Other Positive Correlation |
| AFF3 | 0.086635 | 0.120206 | Other Positive Correlation |
| OAZ2 | 0.086612 | 0.120306 | Other Positive Correlation |
| GOLPH3 | 0.086587 | 0.120413 | Other Positive Correlation |
| USP38 | 0.086587 | 0.120415 | Other Positive Correlation |
| BCAR1 | 0.086524 | 0.120685 | Other Positive Correlation |
| A4GALT | 0.086523 | 0.120687 | Other Positive Correlation |
| PSPH | 0.08648 | 0.120875 | Other Positive Correlation |
| CDO1 | 0.086479 | 0.120879 | Other Positive Correlation |
| GALT | 0.086466 | 0.120936 | Other Positive Correlation |
| LRP6 | 0.086438 | 0.121055 | Other Positive Correlation |
| HARBI1 | 0.086423 | 0.121119 | Other Positive Correlation |
| ASTL | 0.086422 | 0.121123 | Other Positive Correlation |
| MOV10L1 | 0.086419 | 0.121135 | Other Positive Correlation |

| | | | |
|---|---|---|---|
| IGSF10 | 0.086403 | 0.121207 | Other Positive Correlation |
| CXXC1 | 0.086386 | 0.12128 | Other Positive Correlation |
| RNF113A | 0.086376 | 0.121324 | Other Positive Correlation |
| TFF2 | 0.08636 | 0.12139 | Other Positive Correlation |
| PTCH1 | 0.086344 | 0.12146 | Other Positive Correlation |
| SPART | 0.086338 | 0.121486 | Other Positive Correlation |
| KCNK9 | 0.08633 | 0.121522 | Other Positive Correlation |
| ADAM11 | 0.086285 | 0.121719 | Other Positive Correlation |
| CLCN3 | 0.086277 | 0.12175 | Other Positive Correlation |
| HECW2 | 0.08626 | 0.121826 | Other Positive Correlation |
| SLC17A7 | 0.086259 | 0.121828 | Other Positive Correlation |
| NELL2 | 0.086248 | 0.121876 | Other Positive Correlation |
| OR51M1 | 0.086196 | 0.122102 | Other Positive Correlation |
| SEC14L5 | 0.086152 | 0.122294 | Other Positive Correlation |
| ALKBH5 | 0.0861 | 0.12252 | Other Positive Correlation |
| SAMD12 | 0.086074 | 0.122634 | Other Positive Correlation |
| ZNF461 | 0.086073 | 0.122637 | Other Positive Correlation |
| GNAL | 0.086053 | 0.122729 | Other Positive Correlation |
| DCP1A | 0.086009 | 0.122917 | Other Positive Correlation |
| RAB37 | 0.085997 | 0.122973 | Other Positive Correlation |
| ZNHIT2 | 0.085973 | 0.123077 | Other Positive Correlation |
| GSK3A | 0.08597 | 0.123088 | Other Positive Correlation |
| ENKD1 | 0.085945 | 0.123198 | Other Positive Correlation |
| TMEM104 | 0.085888 | 0.123449 | Other Positive Correlation |
| MUC15 | 0.08572 | 0.124185 | Other Positive Correlation |
| EFCAB13 | 0.085709 | 0.124237 | Other Positive Correlation |
| TMTC1 | 0.085703 | 0.124261 | Other Positive Correlation |
| HAPLN4 | 0.085703 | 0.124263 | Other Positive Correlation |
| B4GALT3 | 0.085688 | 0.124326 | Other Positive Correlation |
| CYTH2 | 0.085667 | 0.124422 | Other Positive Correlation |
| TRIM62 | 0.085648 | 0.124504 | Other Positive Correlation |
| PPARGC1A | 0.085608 | 0.124683 | Other Positive Correlation |
| SNX18 | 0.085589 | 0.124764 | Other Positive Correlation |
| SLC6A11 | 0.085568 | 0.124858 | Other Positive Correlation |
| CACNB2 | 0.085567 | 0.124861 | Other Positive Correlation |
| DNAJB12 | 0.085555 | 0.124916 | Other Positive Correlation |
| GAS2L1 | 0.085504 | 0.125142 | Other Positive Correlation |
| ASAP3 | 0.085502 | 0.125152 | Other Positive Correlation |
| SPATA2 | 0.085492 | 0.125196 | Other Positive Correlation |
| LURAP1L | 0.085485 | 0.125225 | Other Positive Correlation |
| TSR2 | 0.085443 | 0.125412 | Other Positive Correlation |
| ACSL6 | 0.085438 | 0.125432 | Other Positive Correlation |
| CANT1 | 0.085416 | 0.125531 | Other Positive Correlation |
| KAT6B | 0.08541 | 0.125558 | Other Positive Correlation |
| WWC2 | 0.085382 | 0.125683 | Other Positive Correlation |
| CACNA2D1 | 0.08538 | 0.125693 | Other Positive Correlation |
| TRH | 0.085373 | 0.125725 | Other Positive Correlation |
| PRR35 | 0.085353 | 0.125814 | Other Positive Correlation |

| | | | |
|---|---|---|---|
| ARFGEF2 | 0.085327 | 0.125926 | Other Positive Correlation |
| TBKBP1 | 0.085291 | 0.126087 | Other Positive Correlation |
| FBH1 | 0.085271 | 0.126179 | Other Positive Correlation |
| ARHGAP28 | 0.085261 | 0.126221 | Other Positive Correlation |
| ZNF329 | 0.085235 | 0.126338 | Other Positive Correlation |
| GOLPH3L | 0.085205 | 0.126473 | Other Positive Correlation |
| H3C6 | 0.085109 | 0.126902 | Other Positive Correlation |
| CAMSAP3 | 0.085036 | 0.12723 | Other Positive Correlation |
| ARHGEF26 | 0.085025 | 0.127277 | Other Positive Correlation |
| CYP2U1 | 0.085024 | 0.127283 | Other Positive Correlation |
| IP6K1 | 0.084979 | 0.127484 | Other Positive Correlation |
| CLEC2L | 0.084957 | 0.127583 | Other Positive Correlation |
| ACBD7 | 0.084925 | 0.127728 | Other Positive Correlation |
| MRPL37 | 0.084919 | 0.127754 | Other Positive Correlation |
| BRSK1 | 0.084882 | 0.127922 | Other Positive Correlation |
| CACNG8 | 0.084824 | 0.128184 | Other Positive Correlation |
| FAM163A | 0.084821 | 0.128197 | Other Positive Correlation |
| TMC5 | 0.084818 | 0.128208 | Other Positive Correlation |
| KHDC1 | 0.084802 | 0.12828 | Other Positive Correlation |
| ARHGAP26 | 0.084773 | 0.128411 | Other Positive Correlation |
| CACNG4 | 0.084753 | 0.128502 | Other Positive Correlation |
| RELT | 0.084753 | 0.128505 | Other Positive Correlation |
| MATN4 | 0.08469 | 0.128791 | Other Positive Correlation |
| ECRG4 | 0.084641 | 0.129011 | Other Positive Correlation |
| MTSS2 | 0.084558 | 0.12939 | Other Positive Correlation |
| SPAG6 | 0.08455 | 0.129423 | Other Positive Correlation |
| ZNF32 | 0.084525 | 0.12954 | Other Positive Correlation |
| RNASEH2C | 0.084511 | 0.129603 | Other Positive Correlation |
| NFE2L3 | 0.084447 | 0.129894 | Other Positive Correlation |
| URGCP | 0.084441 | 0.12992 | Other Positive Correlation |
| WBP2NL | 0.084441 | 0.129921 | Other Positive Correlation |
| ZDHHC11B | 0.084392 | 0.130143 | Other Positive Correlation |
| ADGRA2 | 0.084385 | 0.130176 | Other Positive Correlation |
| PSME4 | 0.084376 | 0.130218 | Other Positive Correlation |
| RBM20 | 0.084349 | 0.130339 | Other Positive Correlation |
| USP37 | 0.084344 | 0.130366 | Other Positive Correlation |
| MLPH | 0.084203 | 0.131009 | Other Positive Correlation |
| MIGA2 | 0.084202 | 0.131016 | Other Positive Correlation |
| GET4 | 0.084178 | 0.131127 | Other Positive Correlation |
| ELMOD3 | 0.084175 | 0.13114 | Other Positive Correlation |
| ABHD4 | 0.08417 | 0.131161 | Other Positive Correlation |
| DDTL | 0.084168 | 0.13117 | Other Positive Correlation |
| CCDC59 | 0.084167 | 0.131176 | Other Positive Correlation |
| EFNA2 | 0.08416 | 0.13121 | Other Positive Correlation |
| WDTC1 | 0.084119 | 0.131396 | Other Positive Correlation |
| MAJIN | 0.084108 | 0.131448 | Other Positive Correlation |
| SLPI | 0.084085 | 0.131555 | Other Positive Correlation |
| KLHL36 | 0.084075 | 0.131598 | Other Positive Correlation |

| | | | |
|---|---|---|---|
| SDR39U1 | 0.084063 | 0.131656 | Other Positive Correlation |
| PRMT6 | 0.084062 | 0.131657 | Other Positive Correlation |
| HYKK | 0.084033 | 0.131791 | Other Positive Correlation |
| PIK3R3 | 0.084009 | 0.131904 | Other Positive Correlation |
| POLR2K | 0.084001 | 0.131941 | Other Positive Correlation |
| DCAF10 | 0.083975 | 0.13206 | Other Positive Correlation |
| ACHE | 0.083962 | 0.132121 | Other Positive Correlation |
| DR1 | 0.083843 | 0.132671 | Other Positive Correlation |
| HERC1 | 0.083837 | 0.132701 | Other Positive Correlation |
| COQ7 | 0.083807 | 0.132839 | Other Positive Correlation |
| EMC3 | 0.08375 | 0.133101 | Other Positive Correlation |
| PIGC | 0.083726 | 0.133213 | Other Positive Correlation |
| CARMIL2 | 0.083698 | 0.133344 | Other Positive Correlation |
| SLC8A3 | 0.083654 | 0.133549 | Other Positive Correlation |
| ABO | 0.08362 | 0.13371 | Other Positive Correlation |
| SLC5A3 | 0.083617 | 0.133723 | Other Positive Correlation |
| CHRNA5 | 0.08357 | 0.13394 | Other Positive Correlation |
| ZNF251 | 0.083477 | 0.134379 | Other Positive Correlation |
| PLXDC2 | 0.083469 | 0.134415 | Other Positive Correlation |
| MT1M | 0.083455 | 0.134481 | Other Positive Correlation |
| SENP5 | 0.08338 | 0.13483 | Other Positive Correlation |
| KLB | 0.083334 | 0.135048 | Other Positive Correlation |
| IRF2BP1 | 0.083331 | 0.13506 | Other Positive Correlation |
| KLK7 | 0.083326 | 0.135084 | Other Positive Correlation |
| EIF4EBP3 | 0.083298 | 0.135219 | Other Positive Correlation |
| ZNF433 | 0.083198 | 0.13569 | Other Positive Correlation |
| ZNF213 | 0.083196 | 0.135697 | Other Positive Correlation |
| MAB21L4 | 0.08319 | 0.135725 | Other Positive Correlation |
| ANKS3 | 0.083061 | 0.136336 | Other Positive Correlation |
| FNDC10 | 0.083049 | 0.136392 | Other Positive Correlation |
| ELF5 | 0.083021 | 0.136527 | Other Positive Correlation |
| ZNF519 | 0.083018 | 0.136539 | Other Positive Correlation |
| RAB6D | 0.083009 | 0.136581 | Other Positive Correlation |
| IGSF22 | 0.082993 | 0.136657 | Other Positive Correlation |
| LRAT | 0.082991 | 0.136669 | Other Positive Correlation |
| SNED1 | 0.082896 | 0.137119 | Other Positive Correlation |
| ELFN1 | 0.082852 | 0.137326 | Other Positive Correlation |
| ANKMY1 | 0.082842 | 0.137373 | Other Positive Correlation |
| CARNS1 | 0.082842 | 0.137376 | Other Positive Correlation |
| CNTLN | 0.082822 | 0.137469 | Other Positive Correlation |
| PLRG1 | 0.082756 | 0.137784 | Other Positive Correlation |
| PRR22 | 0.082737 | 0.137874 | Other Positive Correlation |
| NAALAD2 | 0.082672 | 0.138187 | Other Positive Correlation |
| PDGFRA | 0.082629 | 0.138393 | Other Positive Correlation |
| APBB1IP | 0.082611 | 0.138479 | Other Positive Correlation |
| EPHA7 | 0.082601 | 0.138526 | Other Positive Correlation |
| SYN1 | 0.082593 | 0.138567 | Other Positive Correlation |
| CDYL2 | 0.082579 | 0.138634 | Other Positive Correlation |

| | | | |
|---|---|---|---|
| GREB1 | 0.082572 | 0.138666 | Other Positive Correlation |
| USP8 | 0.082565 | 0.138701 | Other Positive Correlation |
| ZNF610 | 0.082516 | 0.138933 | Other Positive Correlation |
| FLCN | 0.082469 | 0.139159 | Other Positive Correlation |
| PROC | 0.082441 | 0.139293 | Other Positive Correlation |
| HOXC8 | 0.082412 | 0.139433 | Other Positive Correlation |
| MT3 | 0.082411 | 0.139441 | Other Positive Correlation |
| N4BP3 | 0.082398 | 0.139501 | Other Positive Correlation |
| MOCS1 | 0.082397 | 0.139505 | Other Positive Correlation |
| PDK3 | 0.082388 | 0.139549 | Other Positive Correlation |
| ATP8A1 | 0.082299 | 0.139978 | Other Positive Correlation |
| VGLL2 | 0.082281 | 0.140068 | Other Positive Correlation |
| TAF4 | 0.082268 | 0.140132 | Other Positive Correlation |
| RPS6KB1 | 0.082263 | 0.140156 | Other Positive Correlation |
| EPDR1 | 0.082222 | 0.140353 | Other Positive Correlation |
| NPPC | 0.082217 | 0.140375 | Other Positive Correlation |
| DMGDH | 0.082193 | 0.140492 | Other Positive Correlation |
| RTL8A | 0.082173 | 0.140589 | Other Positive Correlation |
| RIPOR1 | 0.082161 | 0.140649 | Other Positive Correlation |
| RAD1 | 0.082065 | 0.141116 | Other Positive Correlation |
| B3GNT7 | 0.082061 | 0.141135 | Other Positive Correlation |
| COQ6 | 0.082039 | 0.141242 | Other Positive Correlation |
| CADPS2 | 0.081999 | 0.141438 | Other Positive Correlation |
| ODAD2 | 0.081987 | 0.141492 | Other Positive Correlation |
| RIPK2 | 0.081941 | 0.14172 | Other Positive Correlation |
| VAX1 | 0.081925 | 0.141795 | Other Positive Correlation |
| PITPNM2 | 0.081911 | 0.141863 | Other Positive Correlation |
| SYT9 | 0.081911 | 0.141864 | Other Positive Correlation |
| SIPA1 | 0.081881 | 0.14201 | Other Positive Correlation |
| SIAH3 | 0.081865 | 0.142092 | Other Positive Correlation |
| OTUD7A | 0.081849 | 0.142167 | Other Positive Correlation |
| RNF180 | 0.081821 | 0.142306 | Other Positive Correlation |
| GALNT12 | 0.081777 | 0.142519 | Other Positive Correlation |
| PDK4 | 0.08173 | 0.142748 | Other Positive Correlation |
| ZSCAN2 | 0.081728 | 0.14276 | Other Positive Correlation |
| TSC22D2 | 0.081707 | 0.142863 | Other Positive Correlation |
| SPICE1 | 0.081658 | 0.143101 | Other Positive Correlation |
| SERP2 | 0.081656 | 0.143111 | Other Positive Correlation |
| PLEKHA7 | 0.081645 | 0.143167 | Other Positive Correlation |
| SLC40A1 | 0.081534 | 0.143713 | Other Positive Correlation |
| GPT | 0.08153 | 0.143736 | Other Positive Correlation |
| STK11 | 0.081529 | 0.143741 | Other Positive Correlation |
| TMEM221 | 0.081501 | 0.143877 | Other Positive Correlation |
| TTPA | 0.0815 | 0.143883 | Other Positive Correlation |
| NSUN7 | 0.081495 | 0.143906 | Other Positive Correlation |
| IHH | 0.081448 | 0.144139 | Other Positive Correlation |
| SCAMP2 | 0.081415 | 0.144304 | Other Positive Correlation |
| NCMAP | 0.081413 | 0.144312 | Other Positive Correlation |

| | | | |
|---|---|---|---|
| TRAPPC2B | 0.081412 | 0.144315 | Other Positive Correlation |
| RIMBP2 | 0.081411 | 0.144323 | Other Positive Correlation |
| SMAD9 | 0.081351 | 0.144619 | Other Positive Correlation |
| PLCXD3 | 0.081277 | 0.144984 | Other Positive Correlation |
| TAF7L | 0.08126 | 0.14507 | Other Positive Correlation |
| DOT1L | 0.081208 | 0.14533 | Other Positive Correlation |
| EGFL6 | 0.081182 | 0.145459 | Other Positive Correlation |
| GDF6 | 0.081163 | 0.14555 | Other Positive Correlation |
| CCZ1 | 0.081152 | 0.145608 | Other Positive Correlation |
| CYP1A1 | 0.081096 | 0.145883 | Other Positive Correlation |
| PLA2G4A | 0.081086 | 0.145934 | Other Positive Correlation |
| ZCCHC3 | 0.08108 | 0.145965 | Other Positive Correlation |
| EMC4 | 0.081065 | 0.146041 | Other Positive Correlation |
| NKX3-2 | 0.081046 | 0.146137 | Other Positive Correlation |
| SNCAIP | 0.08104 | 0.146167 | Other Positive Correlation |
| GABRG3 | 0.081024 | 0.146244 | Other Positive Correlation |
| RHBDL1 | 0.081013 | 0.1463 | Other Positive Correlation |
| KAT2B | 0.080991 | 0.146409 | Other Positive Correlation |
| L3MBTL4 | 0.080949 | 0.146622 | Other Positive Correlation |
| ADCY5 | 0.080912 | 0.146806 | Other Positive Correlation |
| ITPRIPL1 | 0.08086 | 0.147066 | Other Positive Correlation |
| RUNX1T1 | 0.080847 | 0.147129 | Other Positive Correlation |
| PLEKHJ1 | 0.080843 | 0.147152 | Other Positive Correlation |
| HOXA7 | 0.080841 | 0.147162 | Other Positive Correlation |
| TNFRSF9 | 0.080814 | 0.147296 | Other Positive Correlation |
| ALKBH6 | 0.080806 | 0.147336 | Other Positive Correlation |
| MOSMO | 0.080793 | 0.147403 | Other Positive Correlation |
| GRB10 | 0.080759 | 0.147574 | Other Positive Correlation |
| PPARG | 0.080755 | 0.147594 | Other Positive Correlation |
| GFER | 0.080751 | 0.147611 | Other Positive Correlation |
| ABCG4 | 0.080749 | 0.147624 | Other Positive Correlation |
| POR | 0.080683 | 0.147956 | Other Positive Correlation |
| MAP3K5 | 0.080674 | 0.148001 | Other Positive Correlation |
| CNIH3 | 0.080673 | 0.148008 | Other Positive Correlation |
| PLAC9 | 0.080587 | 0.148442 | Other Positive Correlation |
| ZNF821 | 0.080579 | 0.14848 | Other Positive Correlation |
| LAMC3 | 0.080555 | 0.1486 | Other Positive Correlation |
| CLPX | 0.080525 | 0.148754 | Other Positive Correlation |
| LDLRAD1 | 0.08051 | 0.148829 | Other Positive Correlation |
| NT5DC4 | 0.080496 | 0.148899 | Other Positive Correlation |
| NDNF | 0.080496 | 0.148903 | Other Positive Correlation |
| LPCAT1 | 0.080495 | 0.148906 | Other Positive Correlation |
| COG6 | 0.080477 | 0.148998 | Other Positive Correlation |
| ZNF808 | 0.080472 | 0.149024 | Other Positive Correlation |
| MINDY3 | 0.080432 | 0.149225 | Other Positive Correlation |
| ETV1 | 0.080404 | 0.149369 | Other Positive Correlation |
| SLC9A6 | 0.080357 | 0.149604 | Other Positive Correlation |
| ANKH | 0.080286 | 0.149969 | Other Positive Correlation |

| | | | |
|---|---|---|---|
| ZFAND3 | 0.080278 | 0.150009 | Other Positive Correlation |
| JAM3 | 0.080262 | 0.150088 | Other Positive Correlation |
| CDADC1 | 0.080228 | 0.150261 | Other Positive Correlation |
| WNT3A | 0.080186 | 0.150476 | Other Positive Correlation |
| GPATCH11 | 0.08018 | 0.150508 | Other Positive Correlation |
| ST8SIA5 | 0.080167 | 0.150576 | Other Positive Correlation |
| ARHGEF2 | 0.080162 | 0.150598 | Other Positive Correlation |
| CDK9 | 0.080159 | 0.150616 | Other Positive Correlation |
| SLC17A9 | 0.080156 | 0.150628 | Other Positive Correlation |
| TLCD3A | 0.080152 | 0.150653 | Other Positive Correlation |
| RNF6 | 0.080141 | 0.150707 | Other Positive Correlation |
| SLC12A8 | 0.080046 | 0.151193 | Other Positive Correlation |
| GRM4 | 0.080022 | 0.151317 | Other Positive Correlation |
| STX5 | 0.080013 | 0.151361 | Other Positive Correlation |
| EOLA1 | 0.079994 | 0.151459 | Other Positive Correlation |
| RECK | 0.079993 | 0.151465 | Other Positive Correlation |
| PPARA | 0.079993 | 0.151467 | Other Positive Correlation |
| TSPAN13 | 0.079989 | 0.151484 | Other Positive Correlation |
| STX18 | 0.079952 | 0.151674 | Other Positive Correlation |
| IGFBP5 | 0.07995 | 0.151686 | Other Positive Correlation |
| PITPNM1 | 0.079945 | 0.15171 | Other Positive Correlation |
| EDNRB | 0.079942 | 0.151729 | Other Positive Correlation |
| CILP2 | 0.079888 | 0.152003 | Other Positive Correlation |
| OXR1 | 0.079872 | 0.152086 | Other Positive Correlation |
| MPP1 | 0.079858 | 0.152159 | Other Positive Correlation |
| JAKMIP2 | 0.07981 | 0.152406 | Other Positive Correlation |
| CDR2L | 0.079793 | 0.152493 | Other Positive Correlation |
| TAF11 | 0.079784 | 0.152542 | Other Positive Correlation |
| WAPL | 0.079779 | 0.152564 | Other Positive Correlation |
| C12orf56 | 0.079741 | 0.152762 | Other Positive Correlation |
| TVP23A | 0.079731 | 0.152813 | Other Positive Correlation |
| PRKD3 | 0.079727 | 0.152832 | Other Positive Correlation |
| DENND11 | 0.079697 | 0.15299 | Other Positive Correlation |
| DCLRE1A | 0.079671 | 0.153125 | Other Positive Correlation |
| LBHD2 | 0.079653 | 0.153216 | Other Positive Correlation |
| ZMYM5 | 0.079598 | 0.153503 | Other Positive Correlation |
| ZEB1 | 0.079592 | 0.153535 | Other Positive Correlation |
| MPV17L | 0.079558 | 0.15371 | Other Positive Correlation |
| ENSG00000286098 | 0.079548 | 0.15376 | Other Positive Correlation |
| ZNF761 | 0.079522 | 0.153894 | Other Positive Correlation |
| PLPP6 | 0.079498 | 0.154023 | Other Positive Correlation |
| ZNF629 | 0.079486 | 0.154081 | Other Positive Correlation |
| SERPINB6 | 0.079418 | 0.154436 | Other Positive Correlation |
| MDGA2 | 0.079417 | 0.15444 | Other Positive Correlation |
| SLC2A10 | 0.079375 | 0.154663 | Other Positive Correlation |
| ZNF841 | 0.079366 | 0.154708 | Other Positive Correlation |
| PLCL2 | 0.079315 | 0.154973 | Other Positive Correlation |
| NUP88 | 0.079285 | 0.155129 | Other Positive Correlation |

| | | | |
|---|---|---|---|
| GOLGA7B | 0.079178 | 0.155688 | Other Positive Correlation |
| AKAP5 | 0.079174 | 0.155712 | Other Positive Correlation |
| ACP2 | 0.079173 | 0.155716 | Other Positive Correlation |
| LMAN2L | 0.079162 | 0.155774 | Other Positive Correlation |
| PTGER3 | 0.079147 | 0.155854 | Other Positive Correlation |
| PPIP5K1 | 0.079135 | 0.155915 | Other Positive Correlation |
| CC2D1A | 0.079119 | 0.155997 | Other Positive Correlation |
| KIF5A | 0.079089 | 0.156157 | Other Positive Correlation |
| AXIN2 | 0.079081 | 0.156197 | Other Positive Correlation |
| CLCN4 | 0.079055 | 0.156337 | Other Positive Correlation |
| DMRTA1 | 0.079053 | 0.156347 | Other Positive Correlation |
| ZMYND12 | 0.079024 | 0.156499 | Other Positive Correlation |
| TMEM248 | 0.078987 | 0.156696 | Other Positive Correlation |
| DNASE1L2 | 0.078973 | 0.156768 | Other Positive Correlation |
| ENSG00000272822 | 0.07893 | 0.156995 | Other Positive Correlation |
| CDC42BPG | 0.078914 | 0.157079 | Other Positive Correlation |
| ZFAND5 | 0.078892 | 0.157195 | Other Positive Correlation |
| CREB3L4 | 0.078865 | 0.157339 | Other Positive Correlation |
| RNF19A | 0.07886 | 0.157362 | Other Positive Correlation |
| PRKRIP1 | 0.078826 | 0.157542 | Other Positive Correlation |
| RIOX1 | 0.078822 | 0.157562 | Other Positive Correlation |
| RYR1 | 0.078813 | 0.15761 | Other Positive Correlation |
| SHANK1 | 0.078798 | 0.157693 | Other Positive Correlation |
| TLE4 | 0.078764 | 0.157871 | Other Positive Correlation |
| TNFRSF11A | 0.078747 | 0.15796 | Other Positive Correlation |
| AQP11 | 0.078739 | 0.158002 | Other Positive Correlation |
| SOX3 | 0.078725 | 0.158075 | Other Positive Correlation |
| CELF1 | 0.078701 | 0.158202 | Other Positive Correlation |
| MT1H | 0.078688 | 0.158275 | Other Positive Correlation |
| ALG10B | 0.078667 | 0.158387 | Other Positive Correlation |
| THSD7A | 0.078657 | 0.158436 | Other Positive Correlation |
| NUCB2 | 0.078657 | 0.158437 | Other Positive Correlation |
| SHLD1 | 0.078648 | 0.158483 | Other Positive Correlation |
| SOX18 | 0.078636 | 0.158549 | Other Positive Correlation |
| REM1 | 0.078599 | 0.158743 | Other Positive Correlation |
| TSPOAP1 | 0.078584 | 0.158828 | Other Positive Correlation |
| KLHL23 | 0.078517 | 0.159182 | Other Positive Correlation |
| COL5A3 | 0.078473 | 0.159416 | Other Positive Correlation |
| GABRA5 | 0.078472 | 0.159422 | Other Positive Correlation |
| BDNF | 0.078464 | 0.159463 | Other Positive Correlation |
| XKR9 | 0.078458 | 0.159495 | Other Positive Correlation |
| EPHB1 | 0.078446 | 0.159559 | Other Positive Correlation |
| DAPK2 | 0.078444 | 0.159573 | Other Positive Correlation |
| PEX11A | 0.078408 | 0.159766 | Other Positive Correlation |
| RGMA | 0.078378 | 0.159922 | Other Positive Correlation |
| ADGRL1 | 0.078378 | 0.159925 | Other Positive Correlation |
| BMF | 0.078373 | 0.159949 | Other Positive Correlation |
| IQCB1 | 0.078328 | 0.160192 | Other Positive Correlation |

| | | | |
|---|---|---|---|
| RANBP3L | 0.078313 | 0.160273 | Other Positive Correlation |
| TRIM2 | 0.078235 | 0.160688 | Other Positive Correlation |
| ARFGAP3 | 0.078233 | 0.160699 | Other Positive Correlation |
| PLEKHM2 | 0.078226 | 0.160738 | Other Positive Correlation |
| KIF19 | 0.078214 | 0.1608 | Other Positive Correlation |
| GREB1L | 0.078144 | 0.161181 | Other Positive Correlation |
| SDHAF2 | 0.078114 | 0.161338 | Other Positive Correlation |
| VMO1 | 0.078094 | 0.161447 | Other Positive Correlation |
| ACAP2 | 0.078072 | 0.161564 | Other Positive Correlation |
| TBC1D28 | 0.078048 | 0.161697 | Other Positive Correlation |
| EAF1 | 0.078021 | 0.161841 | Other Positive Correlation |
| ZNF689 | 0.078014 | 0.161879 | Other Positive Correlation |
| SLC22A31 | 0.078005 | 0.161929 | Other Positive Correlation |
| CMPK2 | 0.077959 | 0.162178 | Other Positive Correlation |
| B3GNT2 | 0.07793 | 0.162332 | Other Positive Correlation |
| ADAMTS14 | 0.077792 | 0.163078 | Other Positive Correlation |
| ZNF615 | 0.07779 | 0.163092 | Other Positive Correlation |
| DDX19A | 0.077786 | 0.163113 | Other Positive Correlation |
| CIART | 0.077778 | 0.163155 | Other Positive Correlation |
| SORCS3 | 0.077761 | 0.163248 | Other Positive Correlation |
| IL17D | 0.077738 | 0.163376 | Other Positive Correlation |
| PTPRH | 0.077715 | 0.163499 | Other Positive Correlation |
| INAFM1 | 0.077647 | 0.163867 | Other Positive Correlation |
| SETDB2 | 0.077647 | 0.16387 | Other Positive Correlation |
| NRSN2 | 0.077639 | 0.163914 | Other Positive Correlation |
| KIAA2026 | 0.077583 | 0.164219 | Other Positive Correlation |
| TXLNA | 0.077575 | 0.164263 | Other Positive Correlation |
| ACVR1B | 0.077534 | 0.164483 | Other Positive Correlation |
| ATP23 | 0.077517 | 0.164578 | Other Positive Correlation |
| PIGA | 0.077475 | 0.164809 | Other Positive Correlation |
| DHX32 | 0.07747 | 0.164835 | Other Positive Correlation |
| PTPRJ | 0.077417 | 0.165124 | Other Positive Correlation |
| RGS3 | 0.077403 | 0.165201 | Other Positive Correlation |
| FAM178B | 0.077356 | 0.165459 | Other Positive Correlation |
| FAM118A | 0.07731 | 0.16571 | Other Positive Correlation |
| ZNF333 | 0.077303 | 0.165748 | Other Positive Correlation |
| AKAP12 | 0.077299 | 0.165772 | Other Positive Correlation |
| LARP6 | 0.077275 | 0.1659 | Other Positive Correlation |
| ZNF727 | 0.077274 | 0.165909 | Other Positive Correlation |
| ACO2 | 0.07724 | 0.166094 | Other Positive Correlation |
| GPKOW | 0.077224 | 0.166186 | Other Positive Correlation |
| TOMM40L | 0.077158 | 0.166545 | Other Positive Correlation |
| FBLN1 | 0.077128 | 0.166712 | Other Positive Correlation |
| BLACAT1 | 0.077091 | 0.166916 | Other Positive Correlation |
| CNNM4 | 0.077067 | 0.167051 | Other Positive Correlation |
| PFDN6 | 0.077063 | 0.167072 | Other Positive Correlation |
| RASA4 | 0.077054 | 0.16712 | Other Positive Correlation |
| NOCT | 0.077054 | 0.167122 | Other Positive Correlation |

| | | | |
|---|---|---|---|
| ZNF620 | 0.077037 | 0.167217 | Other Positive Correlation |
| ENSG00000285641 | 0.077031 | 0.167246 | Other Positive Correlation |
| TMED4 | 0.07703 | 0.167253 | Other Positive Correlation |
| SV2A | 0.077029 | 0.167258 | Other Positive Correlation |
| SCT | 0.077023 | 0.167292 | Other Positive Correlation |
| ITIH5 | 0.077007 | 0.167382 | Other Positive Correlation |
| MESP2 | 0.076968 | 0.167597 | Other Positive Correlation |
| DIP2C | 0.076967 | 0.167602 | Other Positive Correlation |
| MED21 | 0.076955 | 0.167668 | Other Positive Correlation |
| RAPGEF1 | 0.07694 | 0.167752 | Other Positive Correlation |
| NALF2 | 0.076934 | 0.167784 | Other Positive Correlation |
| ZNF765 | 0.076927 | 0.167825 | Other Positive Correlation |
| FGFRL1 | 0.076915 | 0.16789 | Other Positive Correlation |
| ANO4 | 0.076857 | 0.168212 | Other Positive Correlation |
| PPP1R3G | 0.076814 | 0.168453 | Other Positive Correlation |
| MFSD5 | 0.076801 | 0.168523 | Other Positive Correlation |
| HYOU1 | 0.076759 | 0.168758 | Other Positive Correlation |
| CEBPA | 0.076757 | 0.168766 | Other Positive Correlation |
| CUL3 | 0.076746 | 0.168828 | Other Positive Correlation |
| MED31 | 0.076668 | 0.169266 | Other Positive Correlation |
| RAVER2 | 0.076645 | 0.169393 | Other Positive Correlation |
| GABRB2 | 0.076635 | 0.169446 | Other Positive Correlation |
| TMEM107 | 0.076599 | 0.169646 | Other Positive Correlation |
| GET3 | 0.076583 | 0.169736 | Other Positive Correlation |
| CFTR | 0.076573 | 0.169797 | Other Positive Correlation |
| HIPK1 | 0.076535 | 0.170004 | Other Positive Correlation |
| VPS37D | 0.076483 | 0.170298 | Other Positive Correlation |
| COX6A1 | 0.07648 | 0.170316 | Other Positive Correlation |
| CDK14 | 0.076468 | 0.170381 | Other Positive Correlation |
| B3GALT4 | 0.076423 | 0.170636 | Other Positive Correlation |
| MFSD13A | 0.076393 | 0.170805 | Other Positive Correlation |
| FAM153A | 0.076382 | 0.170862 | Other Positive Correlation |
| S100P | 0.076371 | 0.170924 | Other Positive Correlation |
| NPEPL1 | 0.076366 | 0.170956 | Other Positive Correlation |
| C1orf198 | 0.076352 | 0.171033 | Other Positive Correlation |
| IDH2 | 0.076307 | 0.171286 | Other Positive Correlation |
| RBSN | 0.076296 | 0.171346 | Other Positive Correlation |
| ADAMTS9 | 0.076284 | 0.171415 | Other Positive Correlation |
| PLPP3 | 0.076279 | 0.171444 | Other Positive Correlation |
| SCAI | 0.076258 | 0.171564 | Other Positive Correlation |
| TRIM63 | 0.076241 | 0.171656 | Other Positive Correlation |
| WDR20 | 0.076203 | 0.17187 | Other Positive Correlation |
| C6orf163 | 0.076121 | 0.172334 | Other Positive Correlation |
| RUSC1 | 0.07609 | 0.172513 | Other Positive Correlation |
| SLC44A2 | 0.076079 | 0.17257 | Other Positive Correlation |
| MYO16 | 0.076065 | 0.172651 | Other Positive Correlation |
| ZNF175 | 0.076055 | 0.172709 | Other Positive Correlation |
| ZNF506 | 0.075991 | 0.173069 | Other Positive Correlation |

| | | | |
|---|---|---|---|
| LPAR1 | 0.075975 | 0.17316 | Other Positive Correlation |
| CLEC4A | 0.075968 | 0.1732 | Other Positive Correlation |
| TIAM1 | 0.075955 | 0.173277 | Other Positive Correlation |
| SLC26A1 | 0.075952 | 0.173291 | Other Positive Correlation |
| PKHD1L1 | 0.075904 | 0.173565 | Other Positive Correlation |
| ACER1 | 0.0759 | 0.173586 | Other Positive Correlation |
| FBRS | 0.075825 | 0.174013 | Other Positive Correlation |
| ANKRD35 | 0.075803 | 0.174142 | Other Positive Correlation |
| PRDM11 | 0.075781 | 0.174263 | Other Positive Correlation |
| ZNF316 | 0.075736 | 0.174523 | Other Positive Correlation |
| GSKIP | 0.075726 | 0.174581 | Other Positive Correlation |
| ANKRD63 | 0.075626 | 0.175152 | Other Positive Correlation |
| CFAP298 | 0.075622 | 0.175172 | Other Positive Correlation |
| CEP295NL | 0.0756 | 0.175297 | Other Positive Correlation |
| SPEF2 | 0.075523 | 0.175738 | Other Positive Correlation |
| SLC25A10 | 0.075514 | 0.175791 | Other Positive Correlation |
| DPP4 | 0.075504 | 0.17585 | Other Positive Correlation |
| GATAD1 | 0.075497 | 0.175892 | Other Positive Correlation |
| BRF1 | 0.075459 | 0.176106 | Other Positive Correlation |
| RPRD2 | 0.075429 | 0.176281 | Other Positive Correlation |
| SLC22A16 | 0.075406 | 0.176412 | Other Positive Correlation |
| TULP1 | 0.075363 | 0.176659 | Other Positive Correlation |
| NAT16 | 0.075325 | 0.17688 | Other Positive Correlation |
| STRADB | 0.075293 | 0.177066 | Other Positive Correlation |
| ACD | 0.075271 | 0.177191 | Other Positive Correlation |
| RASGRP3 | 0.075237 | 0.177387 | Other Positive Correlation |
| GPD1 | 0.075194 | 0.177636 | Other Positive Correlation |
| PROM1 | 0.07515 | 0.17789 | Other Positive Correlation |
| KCNA7 | 0.075126 | 0.178031 | Other Positive Correlation |
| C6orf136 | 0.0751 | 0.178179 | Other Positive Correlation |
| HDAC9 | 0.075068 | 0.178365 | Other Positive Correlation |
| SERPINI1 | 0.075049 | 0.178473 | Other Positive Correlation |
| DMRT1 | 0.075047 | 0.178485 | Other Positive Correlation |
| IKZF3 | 0.075 | 0.178762 | Other Positive Correlation |
| ZNF415 | 0.074997 | 0.178776 | Other Positive Correlation |
| KISS1 | 0.074908 | 0.179295 | Other Positive Correlation |
| VSIR | 0.074895 | 0.179373 | Other Positive Correlation |
| DEPDC5 | 0.074883 | 0.179443 | Other Positive Correlation |
| ZNF730 | 0.074863 | 0.179555 | Other Positive Correlation |
| PKN3 | 0.074863 | 0.179559 | Other Positive Correlation |
| PRDX2 | 0.074854 | 0.179606 | Other Positive Correlation |
| SPEF1 | 0.074851 | 0.179626 | Other Positive Correlation |
| OXTR | 0.074848 | 0.179643 | Other Positive Correlation |
| KCNJ11 | 0.074809 | 0.179869 | Other Positive Correlation |
| KLF12 | 0.07478 | 0.180044 | Other Positive Correlation |
| GVQW3 | 0.074765 | 0.180129 | Other Positive Correlation |
| INSYN1 | 0.074764 | 0.180137 | Other Positive Correlation |
| FIGN | 0.074762 | 0.180149 | Other Positive Correlation |

| | | | |
|---|---|---|---|
| GNPDA2 | 0.074744 | 0.180251 | Other Positive Correlation |
| MPND | 0.074725 | 0.180362 | Other Positive Correlation |
| PLEKHB2 | 0.074716 | 0.180413 | Other Positive Correlation |
| PRR12 | 0.07471 | 0.18045 | Other Positive Correlation |
| NXPH3 | 0.074707 | 0.180467 | Other Positive Correlation |
| ABHD6 | 0.074651 | 0.180798 | Other Positive Correlation |
| ZNF211 | 0.074638 | 0.180871 | Other Positive Correlation |
| ABLIM2 | 0.07461 | 0.181038 | Other Positive Correlation |
| C12orf43 | 0.074565 | 0.181299 | Other Positive Correlation |
| SHOC2 | 0.074561 | 0.181322 | Other Positive Correlation |
| BID | 0.074552 | 0.181378 | Other Positive Correlation |
| SAMD10 | 0.074539 | 0.18145 | Other Positive Correlation |
| ARMCX6 | 0.074533 | 0.18149 | Other Positive Correlation |
| LYPD6 | 0.074515 | 0.181593 | Other Positive Correlation |
| HSF4 | 0.074495 | 0.181712 | Other Positive Correlation |
| TMEM132B | 0.074494 | 0.181716 | Other Positive Correlation |
| NARS1 | 0.074443 | 0.182016 | Other Positive Correlation |
| HIP1 | 0.074385 | 0.182359 | Other Positive Correlation |
| SYDE1 | 0.074376 | 0.18241 | Other Positive Correlation |
| AUH | 0.074343 | 0.182605 | Other Positive Correlation |
| KLK5 | 0.074316 | 0.182767 | Other Positive Correlation |
| KLLN | 0.074308 | 0.182812 | Other Positive Correlation |
| PAK6 | 0.074285 | 0.18295 | Other Positive Correlation |
| SHISA2 | 0.074269 | 0.183046 | Other Positive Correlation |
| RCC1L | 0.074233 | 0.183258 | Other Positive Correlation |
| HENMT1 | 0.074186 | 0.183537 | Other Positive Correlation |
| ESX1 | 0.074155 | 0.183719 | Other Positive Correlation |
| JMY | 0.07409 | 0.184102 | Other Positive Correlation |
| CBX8 | 0.074088 | 0.184118 | Other Positive Correlation |
| SNX4 | 0.074082 | 0.184154 | Other Positive Correlation |
| IGFL1 | 0.07408 | 0.184165 | Other Positive Correlation |
| DNAJC8 | 0.074044 | 0.184376 | Other Positive Correlation |
| C2CD4B | 0.074036 | 0.184425 | Other Positive Correlation |
| MPPED1 | 0.07402 | 0.184522 | Other Positive Correlation |
| SLC24A2 | 0.07402 | 0.184523 | Other Positive Correlation |
| GPRC5A | 0.074018 | 0.184533 | Other Positive Correlation |
| ZACN | 0.074002 | 0.184627 | Other Positive Correlation |
| NOS3 | 0.073997 | 0.184655 | Other Positive Correlation |
| MON1A | 0.073997 | 0.184656 | Other Positive Correlation |
| ENSG00000230707 | 0.073986 | 0.184722 | Other Positive Correlation |
| SNX19 | 0.073968 | 0.18483 | Other Positive Correlation |
| DHRS13 | 0.073957 | 0.184895 | Other Positive Correlation |
| RALA | 0.073956 | 0.1849 | Other Positive Correlation |
| LRP3 | 0.073955 | 0.184907 | Other Positive Correlation |
| ZSCAN22 | 0.073925 | 0.185088 | Other Positive Correlation |
| RUBCNL | 0.073904 | 0.185208 | Other Positive Correlation |
| CTTNBP2 | 0.073884 | 0.18533 | Other Positive Correlation |
| SLC35B2 | 0.073883 | 0.185334 | Other Positive Correlation |

| | | | |
|---|---|---|---|
| PNPLA5 | 0.073873 | 0.185397 | Other Positive Correlation |
| SMCR8 | 0.073872 | 0.185403 | Other Positive Correlation |
| PDE1C | 0.073865 | 0.185444 | Other Positive Correlation |
| SOX13 | 0.073859 | 0.185479 | Other Positive Correlation |
| ARMCX3 | 0.073846 | 0.185554 | Other Positive Correlation |
| MRPL46 | 0.073795 | 0.185859 | Other Positive Correlation |
| INO80B | 0.073718 | 0.186322 | Other Positive Correlation |
| MFAP4 | 0.073707 | 0.186385 | Other Positive Correlation |
| RNF103 | 0.073669 | 0.186617 | Other Positive Correlation |
| CACNA1S | 0.073668 | 0.186621 | Other Positive Correlation |
| DEFB134 | 0.073668 | 0.186621 | Other Positive Correlation |
| MYO3A | 0.073572 | 0.187198 | Other Positive Correlation |
| TGFBR3L | 0.073553 | 0.18731 | Other Positive Correlation |
| CEP112 | 0.073525 | 0.187482 | Other Positive Correlation |
| IQSEC1 | 0.073486 | 0.187717 | Other Positive Correlation |
| SLC49A3 | 0.073484 | 0.187726 | Other Positive Correlation |
| PPP1R3F | 0.073453 | 0.187913 | Other Positive Correlation |
| ARVCF | 0.073446 | 0.187957 | Other Positive Correlation |
| FGF12 | 0.073437 | 0.18801 | Other Positive Correlation |
| ZNF736 | 0.07343 | 0.188054 | Other Positive Correlation |
| RNF146 | 0.073429 | 0.188057 | Other Positive Correlation |
| USP48 | 0.073424 | 0.188091 | Other Positive Correlation |
| GFRA4 | 0.073405 | 0.188205 | Other Positive Correlation |
| ENTPD2 | 0.073336 | 0.188618 | Other Positive Correlation |
| DGKD | 0.073331 | 0.18865 | Other Positive Correlation |
| THRB | 0.07326 | 0.189081 | Other Positive Correlation |
| TMPRSS4 | 0.073247 | 0.189161 | Other Positive Correlation |
| ENSG00000257355 | 0.073178 | 0.189576 | Other Positive Correlation |
| PAM | 0.073174 | 0.189599 | Other Positive Correlation |
| IL18RAP | 0.073167 | 0.189644 | Other Positive Correlation |
| FCAMR | 0.073167 | 0.189644 | Other Positive Correlation |
| CSN1S1 | 0.073167 | 0.189644 | Other Positive Correlation |
| ZC3HC1 | 0.073147 | 0.189767 | Other Positive Correlation |
| THPO | 0.07314 | 0.189808 | Other Positive Correlation |
| DCDC2C | 0.073099 | 0.190054 | Other Positive Correlation |
| GFM1 | 0.07306 | 0.190292 | Other Positive Correlation |
| ARID1A | 0.073039 | 0.190425 | Other Positive Correlation |
| PUS10 | 0.073021 | 0.190534 | Other Positive Correlation |
| PPT2 | 0.073011 | 0.190592 | Other Positive Correlation |
| SHANK2 | 0.072993 | 0.1907 | Other Positive Correlation |
| TPPP | 0.072937 | 0.191041 | Other Positive Correlation |
| ISLR | 0.072798 | 0.191892 | Other Positive Correlation |
| MIOX | 0.072798 | 0.191895 | Other Positive Correlation |
| XAGE2 | 0.072774 | 0.192039 | Other Positive Correlation |
| CD300C | 0.072774 | 0.192039 | Other Positive Correlation |
| GPR158 | 0.072774 | 0.192039 | Other Positive Correlation |
| C1orf87 | 0.072774 | 0.192039 | Other Positive Correlation |
| CACNA1A | 0.07276 | 0.192124 | Other Positive Correlation |

| | | | |
|---|---|---|---|
| UTS2R | 0.07274 | 0.192247 | Other Positive Correlation |
| PSMB4 | 0.072739 | 0.192252 | Other Positive Correlation |
| CFAP107 | 0.072696 | 0.192518 | Other Positive Correlation |
| FEV | 0.072696 | 0.192518 | Other Positive Correlation |
| WNT10B | 0.072681 | 0.192609 | Other Positive Correlation |
| SHQ1 | 0.07268 | 0.192617 | Other Positive Correlation |
| SMG8 | 0.072655 | 0.192769 | Other Positive Correlation |
| ARMC1 | 0.072655 | 0.192771 | Other Positive Correlation |
| SNTG2 | 0.072635 | 0.192891 | Other Positive Correlation |
| ZFPM1 | 0.072611 | 0.193038 | Other Positive Correlation |
| ATP13A2 | 0.072588 | 0.19318 | Other Positive Correlation |
| SLC25A30 | 0.072546 | 0.193439 | Other Positive Correlation |
| TRIM52 | 0.072511 | 0.193654 | Other Positive Correlation |
| MALT1 | 0.072508 | 0.193672 | Other Positive Correlation |
| PBOV1 | 0.072504 | 0.193698 | Other Positive Correlation |
| CHST5 | 0.072504 | 0.193698 | Other Positive Correlation |
| IGSF1 | 0.072497 | 0.193744 | Other Positive Correlation |
| NPTX1 | 0.072488 | 0.1938 | Other Positive Correlation |
| NFATC3 | 0.072475 | 0.193875 | Other Positive Correlation |
| BCL2 | 0.072475 | 0.193878 | Other Positive Correlation |
| KLHL34 | 0.072473 | 0.19389 | Other Positive Correlation |
| LCA5 | 0.072435 | 0.194123 | Other Positive Correlation |
| ZHX1-C8orf76 | 0.072428 | 0.19417 | Other Positive Correlation |
| HOXC13 | 0.072405 | 0.194312 | Other Positive Correlation |
| PYCR2 | 0.072357 | 0.194608 | Other Positive Correlation |
| KRT76 | 0.072353 | 0.194629 | Other Positive Correlation |
| FIGNL2 | 0.072324 | 0.194812 | Other Positive Correlation |
| MYO5B | 0.072248 | 0.19528 | Other Positive Correlation |
| ALAD | 0.072233 | 0.195375 | Other Positive Correlation |
| MAGED4B | 0.072219 | 0.195459 | Other Positive Correlation |
| BMP3 | 0.072219 | 0.195459 | Other Positive Correlation |
| ENSG00000261915 | 0.072217 | 0.19547 | Other Positive Correlation |
| DBR1 | 0.07219 | 0.195638 | Other Positive Correlation |
| PRPF31 | 0.072187 | 0.195657 | Other Positive Correlation |
| DEFB124 | 0.072171 | 0.195759 | Other Positive Correlation |
| CLCNKA | 0.072171 | 0.195759 | Other Positive Correlation |
| DUSP9 | 0.072158 | 0.195839 | Other Positive Correlation |
| ZXDB | 0.072102 | 0.196186 | Other Positive Correlation |
| ATF4 | 0.072095 | 0.196231 | Other Positive Correlation |
| TAF6L | 0.072089 | 0.196265 | Other Positive Correlation |
| TICAM1 | 0.072054 | 0.196485 | Other Positive Correlation |
| NDUFB1 | 0.072041 | 0.196566 | Other Positive Correlation |
| PRR5 | 0.072032 | 0.196621 | Other Positive Correlation |
| SLC16A7 | 0.072027 | 0.196655 | Other Positive Correlation |
| INSC | 0.072018 | 0.196713 | Other Positive Correlation |
| SP8 | 0.071996 | 0.196846 | Other Positive Correlation |
| LIMS2 | 0.071985 | 0.196918 | Other Positive Correlation |
| LOX | 0.071954 | 0.19711 | Other Positive Correlation |

| | | | |
|---|---|---|---|
| TMEM214 | 0.071941 | 0.197189 | Other Positive Correlation |
| THAP5 | 0.071924 | 0.197297 | Other Positive Correlation |
| ADGRD1 | 0.071903 | 0.197427 | Other Positive Correlation |
| NBPF11 | 0.071868 | 0.197647 | Other Positive Correlation |
| PIP4K2C | 0.071867 | 0.19765 | Other Positive Correlation |
| GATA3 | 0.071867 | 0.197653 | Other Positive Correlation |
| PPP1R26 | 0.07185 | 0.197756 | Other Positive Correlation |
| AIFM3 | 0.071786 | 0.198159 | Other Positive Correlation |
| UBQLN1 | 0.071747 | 0.198402 | Other Positive Correlation |
| MEGF8 | 0.071746 | 0.198408 | Other Positive Correlation |
| DSTYK | 0.071735 | 0.198477 | Other Positive Correlation |
| HGD | 0.071701 | 0.198693 | Other Positive Correlation |
| COPRS | 0.07168 | 0.198828 | Other Positive Correlation |
| MBOAT1 | 0.071651 | 0.199009 | Other Positive Correlation |
| MYOZ3 | 0.071643 | 0.199056 | Other Positive Correlation |
| AGPAT4 | 0.071631 | 0.199133 | Other Positive Correlation |
| ZNF594 | 0.071588 | 0.199406 | Other Positive Correlation |
| HSPA1L | 0.071518 | 0.199842 | Other Positive Correlation |
| TRIP11 | 0.071462 | 0.200195 | Other Positive Correlation |
| STIMATE | 0.071459 | 0.200216 | Other Positive Correlation |
| LYSMD3 | 0.071428 | 0.200413 | Other Positive Correlation |
| RAB40C | 0.071399 | 0.200594 | Other Positive Correlation |
| KCNK4 | 0.071395 | 0.20062 | Other Positive Correlation |
| COL8A2 | 0.071387 | 0.200671 | Other Positive Correlation |
| ZNF396 | 0.071373 | 0.200763 | Other Positive Correlation |
| GRIN2C | 0.07134 | 0.200967 | Other Positive Correlation |
| FAM111A | 0.07134 | 0.200969 | Other Positive Correlation |
| NOS2 | 0.071335 | 0.200998 | Other Positive Correlation |
| DCAF12L2 | 0.071324 | 0.201071 | Other Positive Correlation |
| MRPL50 | 0.071312 | 0.201147 | Other Positive Correlation |
| RNASEH1 | 0.071301 | 0.201214 | Other Positive Correlation |
| POMT1 | 0.071294 | 0.201261 | Other Positive Correlation |
| RSPO1 | 0.071283 | 0.201328 | Other Positive Correlation |
| NYAP1 | 0.071275 | 0.201384 | Other Positive Correlation |
| CBARP | 0.071212 | 0.201783 | Other Positive Correlation |
| SLC6A16 | 0.071196 | 0.201884 | Other Positive Correlation |
| TMEM186 | 0.071183 | 0.201965 | Other Positive Correlation |
| SNAPC3 | 0.071135 | 0.202271 | Other Positive Correlation |
| GUCD1 | 0.071052 | 0.202798 | Other Positive Correlation |
| FGFR4 | 0.071043 | 0.202854 | Other Positive Correlation |
| TCP11 | 0.071043 | 0.202855 | Other Positive Correlation |
| FZD8 | 0.071039 | 0.202879 | Other Positive Correlation |
| NPAT | 0.071035 | 0.202905 | Other Positive Correlation |
| UQCC3 | 0.071019 | 0.203008 | Other Positive Correlation |
| CSNK1D | 0.071001 | 0.203123 | Other Positive Correlation |
| SSTR2 | 0.070973 | 0.203305 | Other Positive Correlation |
| KCNMB3 | 0.070953 | 0.203427 | Other Positive Correlation |
| NPB | 0.07092 | 0.203638 | Other Positive Correlation |

| | | | |
|---|---|---|---|
| CBFA2T3 | 0.070912 | 0.203692 | Other Positive Correlation |
| IL19 | 0.0709 | 0.203767 | Other Positive Correlation |
| ZNRF2 | 0.07089 | 0.203835 | Other Positive Correlation |
| SYNPO2 | 0.070852 | 0.204074 | Other Positive Correlation |
| IPO8 | 0.070846 | 0.204114 | Other Positive Correlation |
| BAP1 | 0.070843 | 0.204133 | Other Positive Correlation |
| CHST12 | 0.070833 | 0.204194 | Other Positive Correlation |
| DND1 | 0.070799 | 0.204417 | Other Positive Correlation |
| TAS2R3 | 0.070775 | 0.204571 | Other Positive Correlation |
| KCNA4 | 0.070775 | 0.204571 | Other Positive Correlation |
| TUBB1 | 0.070764 | 0.204641 | Other Positive Correlation |
| TCHP | 0.070741 | 0.204787 | Other Positive Correlation |
| ZNRF1 | 0.070713 | 0.204966 | Other Positive Correlation |
| CSTL1 | 0.070698 | 0.20506 | Other Positive Correlation |
| HILPDA | 0.070676 | 0.205204 | Other Positive Correlation |
| LRRC63 | 0.070671 | 0.205238 | Other Positive Correlation |
| APLF | 0.070622 | 0.205551 | Other Positive Correlation |
| ARHGAP36 | 0.070613 | 0.205607 | Other Positive Correlation |
| PSMD11 | 0.070607 | 0.205646 | Other Positive Correlation |
| CALB2 | 0.070594 | 0.205734 | Other Positive Correlation |
| JAK1 | 0.070578 | 0.205835 | Other Positive Correlation |
| POU3F1 | 0.070549 | 0.206023 | Other Positive Correlation |
| POLR2J2 | 0.070545 | 0.206047 | Other Positive Correlation |
| HELB | 0.070522 | 0.206193 | Other Positive Correlation |
| TIGD3 | 0.070521 | 0.206202 | Other Positive Correlation |
| TYW5 | 0.07051 | 0.206275 | Other Positive Correlation |
| FZD10 | 0.070502 | 0.206326 | Other Positive Correlation |
| NME5 | 0.070497 | 0.206354 | Other Positive Correlation |
| PIP4K2A | 0.070454 | 0.206634 | Other Positive Correlation |
| ENOX1 | 0.070411 | 0.206912 | Other Positive Correlation |
| AMPH | 0.070381 | 0.207108 | Other Positive Correlation |
| ZNF682 | 0.070378 | 0.207125 | Other Positive Correlation |
| TNFRSF8 | 0.070373 | 0.207156 | Other Positive Correlation |
| ZNF71 | 0.070355 | 0.207275 | Other Positive Correlation |
| CLDN5 | 0.070328 | 0.207448 | Other Positive Correlation |
| KIF21B | 0.070269 | 0.207834 | Other Positive Correlation |
| PI4KB | 0.07025 | 0.207952 | Other Positive Correlation |
| NUCB1 | 0.070218 | 0.20816 | Other Positive Correlation |
| SUCLG1 | 0.07021 | 0.208212 | Other Positive Correlation |
| DDX28 | 0.070189 | 0.208349 | Other Positive Correlation |
| ADA2 | 0.070181 | 0.208403 | Other Positive Correlation |
| BAZ1B | 0.070177 | 0.208432 | Other Positive Correlation |
| RNF166 | 0.070159 | 0.208545 | Other Positive Correlation |
| ADAM33 | 0.070143 | 0.208648 | Other Positive Correlation |
| DAGLA | 0.070118 | 0.208814 | Other Positive Correlation |
| TLK2 | 0.070117 | 0.20882 | Other Positive Correlation |
| CD200 | 0.070072 | 0.209109 | Other Positive Correlation |
| HSPA14 | 0.070068 | 0.209138 | Other Positive Correlation |

| | | | |
|---|---|---|---|
| AQP1 | 0.07006 | 0.209189 | Other Positive Correlation |
| MED7 | 0.070037 | 0.209343 | Other Positive Correlation |
| SBSN | 0.070014 | 0.209487 | Other Positive Correlation |
| RANBP3 | 0.070009 | 0.20952 | Other Positive Correlation |
| PCDH19 | 0.069979 | 0.209718 | Other Positive Correlation |
| PCED1A | 0.069967 | 0.209795 | Other Positive Correlation |
| TRIM39 | 0.069945 | 0.209942 | Other Positive Correlation |
| CCDC144A | 0.069904 | 0.210211 | Other Positive Correlation |
| MDM1 | 0.069897 | 0.210255 | Other Positive Correlation |
| DOCK8 | 0.069894 | 0.210271 | Other Positive Correlation |
| ALS2 | 0.069884 | 0.210337 | Other Positive Correlation |
| PHRF1 | 0.069874 | 0.210403 | Other Positive Correlation |
| MYO15A | 0.069868 | 0.210442 | Other Positive Correlation |
| CYLD | 0.069849 | 0.21057 | Other Positive Correlation |
| CELF5 | 0.069842 | 0.210616 | Other Positive Correlation |
| NGEF | 0.069815 | 0.21079 | Other Positive Correlation |
| FBXO9 | 0.06981 | 0.210823 | Other Positive Correlation |
| LRRC58 | 0.069718 | 0.211428 | Other Positive Correlation |
| PCOLCE | 0.069691 | 0.211602 | Other Positive Correlation |
| BCL7B | 0.069658 | 0.211824 | Other Positive Correlation |
| CCDC97 | 0.069646 | 0.211902 | Other Positive Correlation |
| TAS1R1 | 0.069643 | 0.21192 | Other Positive Correlation |
| PUM2 | 0.06963 | 0.212004 | Other Positive Correlation |
| MEIOB | 0.069612 | 0.212126 | Other Positive Correlation |
| DNAJC28 | 0.069611 | 0.212132 | Other Positive Correlation |
| XK | 0.069604 | 0.212179 | Other Positive Correlation |
| CA4 | 0.069566 | 0.21243 | Other Positive Correlation |
| PATE2 | 0.069542 | 0.212588 | Other Positive Correlation |
| BRAP | 0.069534 | 0.212638 | Other Positive Correlation |
| NALCN | 0.06952 | 0.212728 | Other Positive Correlation |
| GPC3 | 0.069516 | 0.212756 | Other Positive Correlation |
| SAMHD1 | 0.069486 | 0.212957 | Other Positive Correlation |
| PYGO1 | 0.06948 | 0.212996 | Other Positive Correlation |
| F7 | 0.069456 | 0.213152 | Other Positive Correlation |
| SIX4 | 0.069454 | 0.213169 | Other Positive Correlation |
| ZNF584 | 0.069442 | 0.213245 | Other Positive Correlation |
| PRRT1 | 0.06939 | 0.213589 | Other Positive Correlation |
| FGFR1 | 0.069362 | 0.213775 | Other Positive Correlation |
| ADGRV1 | 0.069347 | 0.213874 | Other Positive Correlation |
| DHX37 | 0.069334 | 0.213962 | Other Positive Correlation |
| SNIP1 | 0.069306 | 0.214147 | Other Positive Correlation |
| CRAMP1 | 0.069306 | 0.214147 | Other Positive Correlation |
| TTYH2 | 0.069283 | 0.214296 | Other Positive Correlation |
| TIGD7 | 0.069283 | 0.214299 | Other Positive Correlation |
| CLIP3 | 0.069264 | 0.214426 | Other Positive Correlation |
| MORC2 | 0.069252 | 0.214501 | Other Positive Correlation |
| SETD1B | 0.069248 | 0.214531 | Other Positive Correlation |
| GFI1 | 0.06922 | 0.214713 | Other Positive Correlation |

| | | | |
|---|---|---|---|
| ZNF248 | 0.069192 | 0.214904 | Other Positive Correlation |
| CSRP2 | 0.069188 | 0.214925 | Other Positive Correlation |
| SGK3 | 0.069174 | 0.215021 | Other Positive Correlation |
| RAF1 | 0.069172 | 0.215038 | Other Positive Correlation |
| LAMP3 | 0.069155 | 0.215151 | Other Positive Correlation |
| SLC29A3 | 0.069144 | 0.215219 | Other Positive Correlation |
| OVOL1 | 0.069142 | 0.215234 | Other Positive Correlation |
| CMTM1 | 0.069099 | 0.215518 | Other Positive Correlation |
| CGAS | 0.069093 | 0.215561 | Other Positive Correlation |
| ZSWIM8 | 0.06909 | 0.215579 | Other Positive Correlation |
| ZNF157 | 0.069083 | 0.215625 | Other Positive Correlation |
| NKAPL | 0.068975 | 0.216347 | Other Positive Correlation |
| CFAP77 | 0.068975 | 0.21635 | Other Positive Correlation |
| ESRP1 | 0.068939 | 0.216587 | Other Positive Correlation |
| SLC22A4 | 0.068936 | 0.216605 | Other Positive Correlation |
| SECISBP2L | 0.0689 | 0.216852 | Other Positive Correlation |
| SH3RF3 | 0.068874 | 0.217024 | Other Positive Correlation |
| ZNF45 | 0.068865 | 0.21708 | Other Positive Correlation |
| ASTN2 | 0.068854 | 0.217155 | Other Positive Correlation |
| ZFYVE28 | 0.068853 | 0.217163 | Other Positive Correlation |
| BCL6 | 0.06885 | 0.217183 | Other Positive Correlation |
| TSPAN12 | 0.068847 | 0.217205 | Other Positive Correlation |
| EPRS1 | 0.068844 | 0.217222 | Other Positive Correlation |
| MUC12 | 0.068826 | 0.217346 | Other Positive Correlation |
| ZC4H2 | 0.068805 | 0.217488 | Other Positive Correlation |
| BMERB1 | 0.068775 | 0.217687 | Other Positive Correlation |
| CNP | 0.068724 | 0.218027 | Other Positive Correlation |
| PRRG3 | 0.068644 | 0.218566 | Other Positive Correlation |
| NKX2-4 | 0.0686 | 0.218863 | Other Positive Correlation |
| ZNF804A | 0.0686 | 0.218863 | Other Positive Correlation |
| LCK | 0.068579 | 0.219 | Other Positive Correlation |
| RNF103-CHMP3 | 0.068524 | 0.219372 | Other Positive Correlation |
| PGLYRP2 | 0.068524 | 0.219372 | Other Positive Correlation |
| SLC22A10 | 0.068524 | 0.219372 | Other Positive Correlation |
| FAM133A | 0.068524 | 0.219372 | Other Positive Correlation |
| KLHL14 | 0.068524 | 0.219372 | Other Positive Correlation |
| EYA1 | 0.068458 | 0.219819 | Other Positive Correlation |
| INCA1 | 0.068457 | 0.219826 | Other Positive Correlation |
| CDC42EP4 | 0.068434 | 0.219977 | Other Positive Correlation |
| MANSC1 | 0.068429 | 0.220014 | Other Positive Correlation |
| IL20RA | 0.068417 | 0.220094 | Other Positive Correlation |
| TMEM192 | 0.068417 | 0.220097 | Other Positive Correlation |
| CLMN | 0.068412 | 0.220131 | Other Positive Correlation |
| TSKU | 0.068397 | 0.220233 | Other Positive Correlation |
| SLC1A6 | 0.068357 | 0.220498 | Other Positive Correlation |
| CACNB1 | 0.06835 | 0.220549 | Other Positive Correlation |
| ZNF844 | 0.068331 | 0.220677 | Other Positive Correlation |
| RMI2 | 0.068323 | 0.220729 | Other Positive Correlation |

| | | | |
|---|---|---|---|
| ATOH8 | 0.068284 | 0.220999 | Other Positive Correlation |
| CEP19 | 0.068279 | 0.22103 | Other Positive Correlation |
| WFIKKN2 | 0.068276 | 0.221053 | Other Positive Correlation |
| GALNTL6 | 0.068276 | 0.221053 | Other Positive Correlation |
| GALNT17 | 0.068276 | 0.221053 | Other Positive Correlation |
| STH | 0.068276 | 0.221053 | Other Positive Correlation |
| SFRP4 | 0.068276 | 0.221053 | Other Positive Correlation |
| TMEM215 | 0.068276 | 0.221053 | Other Positive Correlation |
| LRRIQ4 | 0.068276 | 0.221053 | Other Positive Correlation |
| SRRM4 | 0.068276 | 0.221053 | Other Positive Correlation |
| RAB11FIP5 | 0.068258 | 0.221174 | Other Positive Correlation |
| EMILIN3 | 0.068251 | 0.221221 | Other Positive Correlation |
| PYGM | 0.068195 | 0.221599 | Other Positive Correlation |
| LAMTOR3 | 0.068175 | 0.221738 | Other Positive Correlation |
| PGBD5 | 0.068152 | 0.221892 | Other Positive Correlation |
| LZTS3 | 0.068003 | 0.222907 | Other Positive Correlation |
| ABHD15 | 0.068001 | 0.222923 | Other Positive Correlation |
| ZSWIM4 | 0.067969 | 0.223141 | Other Positive Correlation |
| CTDP1 | 0.067938 | 0.223347 | Other Positive Correlation |
| FCHSD2 | 0.067917 | 0.223493 | Other Positive Correlation |
| SKOR2 | 0.067891 | 0.223674 | Other Positive Correlation |
| HSPB7 | 0.067891 | 0.223674 | Other Positive Correlation |
| CYP4F3 | 0.067876 | 0.223775 | Other Positive Correlation |
| CBLN1 | 0.067805 | 0.224257 | Other Positive Correlation |
| COX6B2 | 0.067792 | 0.224352 | Other Positive Correlation |
| GBGT1 | 0.067783 | 0.224409 | Other Positive Correlation |
| PCP4L1 | 0.067737 | 0.224726 | Other Positive Correlation |
| VAMP4 | 0.067713 | 0.224891 | Other Positive Correlation |
| ZNF701 | 0.067693 | 0.22503 | Other Positive Correlation |
| ZFPM2 | 0.067675 | 0.22515 | Other Positive Correlation |
| TBC1D9 | 0.067605 | 0.225635 | Other Positive Correlation |
| MEST | 0.067575 | 0.22584 | Other Positive Correlation |
| SPACDR | 0.067548 | 0.226022 | Other Positive Correlation |
| MEIG1 | 0.067513 | 0.226267 | Other Positive Correlation |
| PDCD1 | 0.067468 | 0.226576 | Other Positive Correlation |
| GPR37L1 | 0.067463 | 0.226614 | Other Positive Correlation |
| ELMO1 | 0.06743 | 0.226835 | Other Positive Correlation |
| TRO | 0.06743 | 0.226837 | Other Positive Correlation |
| ENSG00000288658 | 0.067364 | 0.227297 | Other Positive Correlation |
| PDSS2 | 0.067344 | 0.227434 | Other Positive Correlation |
| HEATR5A | 0.06733 | 0.227529 | Other Positive Correlation |
| ZC3H10 | 0.06732 | 0.227598 | Other Positive Correlation |
| MSL2 | 0.067298 | 0.227748 | Other Positive Correlation |
| PLEKHM3 | 0.067279 | 0.227883 | Other Positive Correlation |
| GPT2 | 0.067262 | 0.227997 | Other Positive Correlation |
| RTN4RL1 | 0.067229 | 0.228226 | Other Positive Correlation |
| ING4 | 0.067058 | 0.229415 | Other Positive Correlation |
| RAB28 | 0.067052 | 0.22946 | Other Positive Correlation |

| | | | |
|---|---|---|---|
| LCMT2 | 0.067011 | 0.229744 | Other Positive Correlation |
| KDF1 | 0.067004 | 0.229793 | Other Positive Correlation |
| HBZ | 0.066958 | 0.230111 | Other Positive Correlation |
| DIRAS3 | 0.066932 | 0.230299 | Other Positive Correlation |
| SLC52A1 | 0.066926 | 0.23034 | Other Positive Correlation |
| SPTSSA | 0.066882 | 0.230647 | Other Positive Correlation |
| NEURL1B | 0.066866 | 0.230758 | Other Positive Correlation |
| DLX2 | 0.066861 | 0.230793 | Other Positive Correlation |
| RGS7BP | 0.066855 | 0.230834 | Other Positive Correlation |
| TTF2 | 0.066814 | 0.231121 | Other Positive Correlation |
| HPS1 | 0.066808 | 0.231163 | Other Positive Correlation |
| ZNF81 | 0.066771 | 0.231422 | Other Positive Correlation |
| HECTD3 | 0.066745 | 0.231602 | Other Positive Correlation |
| RNF212B | 0.066741 | 0.231631 | Other Positive Correlation |
| C11orf71 | 0.066717 | 0.231798 | Other Positive Correlation |
| OXNAD1 | 0.066695 | 0.231951 | Other Positive Correlation |
| SBK2 | 0.06669 | 0.23199 | Other Positive Correlation |
| NRL | 0.066689 | 0.231995 | Other Positive Correlation |
| INTS1 | 0.066637 | 0.232361 | Other Positive Correlation |
| RNF181 | 0.066632 | 0.232394 | Other Positive Correlation |
| RNF41 | 0.066609 | 0.232555 | Other Positive Correlation |
| PCDHGA5 | 0.066572 | 0.232817 | Other Positive Correlation |
| SEPTIN1 | 0.066562 | 0.23289 | Other Positive Correlation |
| GPR63 | 0.066558 | 0.232916 | Other Positive Correlation |
| LRRC45 | 0.066548 | 0.23299 | Other Positive Correlation |
| UNK | 0.066512 | 0.233244 | Other Positive Correlation |
| GSN | 0.066509 | 0.23326 | Other Positive Correlation |
| TLCD2 | 0.066487 | 0.233414 | Other Positive Correlation |
| SLC6A20 | 0.066476 | 0.233496 | Other Positive Correlation |
| E2F3 | 0.066469 | 0.233543 | Other Positive Correlation |
| FGFBP3 | 0.066438 | 0.233764 | Other Positive Correlation |
| RGS19 | 0.066431 | 0.233815 | Other Positive Correlation |
| ZNF585A | 0.066427 | 0.233838 | Other Positive Correlation |
| ANKLE2 | 0.066366 | 0.234272 | Other Positive Correlation |
| TRAPPC9 | 0.066353 | 0.234359 | Other Positive Correlation |
| TEFM | 0.066337 | 0.234477 | Other Positive Correlation |
| KCNV1 | 0.066281 | 0.234869 | Other Positive Correlation |
| TRPC4AP | 0.066259 | 0.23503 | Other Positive Correlation |
| MAFB | 0.066254 | 0.235063 | Other Positive Correlation |
| COCH | 0.066239 | 0.235171 | Other Positive Correlation |
| MITF | 0.066189 | 0.235525 | Other Positive Correlation |
| MOBP | 0.066178 | 0.235599 | Other Positive Correlation |
| SEMA3D | 0.066177 | 0.235609 | Other Positive Correlation |
| CHST2 | 0.066168 | 0.235671 | Other Positive Correlation |
| CPNE5 | 0.066164 | 0.235702 | Other Positive Correlation |
| FAM199X | 0.066154 | 0.235773 | Other Positive Correlation |
| MAN2A2 | 0.066146 | 0.235832 | Other Positive Correlation |
| RHOF | 0.066141 | 0.235866 | Other Positive Correlation |

| | | | |
|---|---|---|---|
| TMEM130 | 0.066107 | 0.236103 | Other Positive Correlation |
| MAPK12 | 0.066088 | 0.236236 | Other Positive Correlation |
| SEZ6 | 0.066059 | 0.236447 | Other Positive Correlation |
| CCL22 | 0.06598 | 0.23701 | Other Positive Correlation |
| ATMIN | 0.065963 | 0.237132 | Other Positive Correlation |
| TFPT | 0.065909 | 0.237512 | Other Positive Correlation |
| NRG2 | 0.065851 | 0.237925 | Other Positive Correlation |
| CLCN1 | 0.065835 | 0.238041 | Other Positive Correlation |
| BEST2 | 0.065823 | 0.238131 | Other Positive Correlation |
| ACTRT3 | 0.065819 | 0.238156 | Other Positive Correlation |
| PACSIN2 | 0.06581 | 0.238225 | Other Positive Correlation |
| ASXL2 | 0.065809 | 0.238226 | Other Positive Correlation |
| RDH12 | 0.065805 | 0.238258 | Other Positive Correlation |
| ZNF835 | 0.065768 | 0.238523 | Other Positive Correlation |
| CDS2 | 0.065754 | 0.238619 | Other Positive Correlation |
| CLCN2 | 0.065644 | 0.239408 | Other Positive Correlation |
| ABCD1 | 0.065641 | 0.239428 | Other Positive Correlation |
| ZFAT | 0.065608 | 0.239665 | Other Positive Correlation |
| COASY | 0.065607 | 0.239677 | Other Positive Correlation |
| CLNK | 0.065552 | 0.240069 | Other Positive Correlation |
| TRIM6 | 0.065542 | 0.240146 | Other Positive Correlation |
| SPRTN | 0.065533 | 0.240204 | Other Positive Correlation |
| SAC3D1 | 0.065532 | 0.240217 | Other Positive Correlation |
| CEP63 | 0.065517 | 0.240321 | Other Positive Correlation |
| TM4SF19 | 0.065508 | 0.240389 | Other Positive Correlation |
| PPP1R16B | 0.06549 | 0.240519 | Other Positive Correlation |
| ZNF398 | 0.065473 | 0.240637 | Other Positive Correlation |
| GNG2 | 0.065455 | 0.240766 | Other Positive Correlation |
| TGIF2 | 0.06543 | 0.24095 | Other Positive Correlation |
| ZNF672 | 0.065402 | 0.241154 | Other Positive Correlation |
| ENSG00000257390 | 0.065367 | 0.241404 | Other Positive Correlation |
| ZNF324B | 0.065313 | 0.241793 | Other Positive Correlation |
| TNFAIP2 | 0.065291 | 0.241951 | Other Positive Correlation |
| ZNF501 | 0.065275 | 0.242066 | Other Positive Correlation |
| SLC25A27 | 0.065274 | 0.242075 | Other Positive Correlation |
| ZNF524 | 0.065252 | 0.242231 | Other Positive Correlation |
| RLF | 0.065213 | 0.242516 | Other Positive Correlation |
| EPHA6 | 0.065193 | 0.242663 | Other Positive Correlation |
| C2CD2L | 0.065185 | 0.242718 | Other Positive Correlation |
| CYB5R4 | 0.065165 | 0.242862 | Other Positive Correlation |
| STXBP1 | 0.065121 | 0.24318 | Other Positive Correlation |
| AFTPH | 0.065082 | 0.243462 | Other Positive Correlation |
| SLC7A7 | 0.06505 | 0.243699 | Other Positive Correlation |
| ZFAND4 | 0.065001 | 0.24405 | Other Positive Correlation |
| LZTS2 | 0.064997 | 0.244086 | Other Positive Correlation |
| ACOXL | 0.064991 | 0.244127 | Other Positive Correlation |
| GSG1L | 0.06489 | 0.24486 | Other Positive Correlation |
| RNF2 | 0.064856 | 0.245106 | Other Positive Correlation |

| | | | |
|---|---|---|---|
| ST14 | 0.064777 | 0.245686 | Other Positive Correlation |
| PNMA1 | 0.064774 | 0.24571 | Other Positive Correlation |
| PRR15L | 0.064771 | 0.245729 | Other Positive Correlation |
| CSRNP1 | 0.064732 | 0.246013 | Other Positive Correlation |
| VASN | 0.064719 | 0.246111 | Other Positive Correlation |
| F8 | 0.064718 | 0.246119 | Other Positive Correlation |
| RGS11 | 0.064715 | 0.24614 | Other Positive Correlation |
| VRK3 | 0.064705 | 0.246214 | Other Positive Correlation |
| PURG | 0.064703 | 0.24623 | Other Positive Correlation |
| B3GAT1 | 0.06468 | 0.246399 | Other Positive Correlation |
| ZBTB47 | 0.064673 | 0.246447 | Other Positive Correlation |
| TMED3 | 0.06466 | 0.24654 | Other Positive Correlation |
| USPL1 | 0.064609 | 0.246916 | Other Positive Correlation |
| ZNF423 | 0.064607 | 0.24693 | Other Positive Correlation |
| OR2C3 | 0.064606 | 0.246938 | Other Positive Correlation |
| CD8A | 0.064606 | 0.24694 | Other Positive Correlation |
| SWSAP1 | 0.064541 | 0.247415 | Other Positive Correlation |
| ABTB3 | 0.064531 | 0.247486 | Other Positive Correlation |
| ADPRS | 0.064519 | 0.247574 | Other Positive Correlation |
| STRA6 | 0.064517 | 0.247591 | Other Positive Correlation |
| H2AC25 | 0.064406 | 0.248406 | Other Positive Correlation |
| AZGP1 | 0.064397 | 0.248473 | Other Positive Correlation |
| MARCHF5 | 0.064369 | 0.248677 | Other Positive Correlation |
| HSD11B1L | 0.064348 | 0.248833 | Other Positive Correlation |
| SYNGAP1 | 0.064333 | 0.248945 | Other Positive Correlation |
| CFAP45 | 0.064285 | 0.249297 | Other Positive Correlation |
| SLC4A9 | 0.064284 | 0.249306 | Other Positive Correlation |
| SLC1A2 | 0.06425 | 0.249555 | Other Positive Correlation |
| BLTP3B | 0.064238 | 0.249648 | Other Positive Correlation |
| LTO1 | 0.064189 | 0.250009 | Other Positive Correlation |
| PPP1R1B | 0.06417 | 0.250148 | Other Positive Correlation |
| GABRR2 | 0.064126 | 0.250473 | Other Positive Correlation |
| ENSG00000287908 | 0.064103 | 0.250641 | Other Positive Correlation |
| TSFM | 0.064103 | 0.250647 | Other Positive Correlation |
| SEC23B | 0.064072 | 0.250876 | Other Positive Correlation |
| RAD21 | 0.064039 | 0.25112 | Other Positive Correlation |
| SCRG1 | 0.064034 | 0.251153 | Other Positive Correlation |
| APH1B | 0.064028 | 0.251197 | Other Positive Correlation |
| TEX19 | 0.064024 | 0.251229 | Other Positive Correlation |
| ERICH5 | 0.063949 | 0.251786 | Other Positive Correlation |
| ZNF114 | 0.063941 | 0.251848 | Other Positive Correlation |
| PDCL | 0.063928 | 0.251943 | Other Positive Correlation |
| TMEM9 | 0.063912 | 0.25206 | Other Positive Correlation |
| MAFF | 0.063885 | 0.252263 | Other Positive Correlation |
| TMEM198 | 0.063862 | 0.252436 | Other Positive Correlation |
| SMCO1 | 0.063817 | 0.252769 | Other Positive Correlation |
| CUL2 | 0.0638 | 0.252898 | Other Positive Correlation |
| BRD8 | 0.063784 | 0.253013 | Other Positive Correlation |

| | | | |
|---|---|---|---|
| IRX5 | 0.063735 | 0.253382 | Other Positive Correlation |
| HSF2 | 0.063676 | 0.253822 | Other Positive Correlation |
| FAAH2 | 0.063644 | 0.25406 | Other Positive Correlation |
| ZBED8 | 0.063631 | 0.254155 | Other Positive Correlation |
| NRN1L | 0.063621 | 0.254231 | Other Positive Correlation |
| DUOX2 | 0.063608 | 0.254328 | Other Positive Correlation |
| FBXO46 | 0.063591 | 0.254456 | Other Positive Correlation |
| CACHD1 | 0.063577 | 0.254562 | Other Positive Correlation |
| C6orf15 | 0.063569 | 0.254624 | Other Positive Correlation |
| TMEM120B | 0.063551 | 0.254759 | Other Positive Correlation |
| DAB2 | 0.06354 | 0.254837 | Other Positive Correlation |
| ATP6V1C1 | 0.063405 | 0.255848 | Other Positive Correlation |
| ZNF668 | 0.063401 | 0.255884 | Other Positive Correlation |
| CAPN11 | 0.063362 | 0.256172 | Other Positive Correlation |
| APOB | 0.06336 | 0.256186 | Other Positive Correlation |
| CCDC92B | 0.063341 | 0.256333 | Other Positive Correlation |
| MAP3K4 | 0.063314 | 0.256535 | Other Positive Correlation |
| PALD1 | 0.063301 | 0.256633 | Other Positive Correlation |
| KCNQ2 | 0.063266 | 0.256894 | Other Positive Correlation |
| COL4A3 | 0.063264 | 0.256915 | Other Positive Correlation |
| ILDR1 | 0.063256 | 0.256972 | Other Positive Correlation |
| RUNDC1 | 0.063245 | 0.257058 | Other Positive Correlation |
| CYP4X1 | 0.063199 | 0.2574 | Other Positive Correlation |
| IWS1 | 0.063186 | 0.257499 | Other Positive Correlation |
| PCDHAC2 | 0.063163 | 0.257674 | Other Positive Correlation |
| TTC9C | 0.063147 | 0.257794 | Other Positive Correlation |
| VSIG10L | 0.063124 | 0.257967 | Other Positive Correlation |
| SATB2 | 0.063119 | 0.258007 | Other Positive Correlation |
| SIM2 | 0.063098 | 0.258166 | Other Positive Correlation |
| NEUROD2 | 0.063088 | 0.258242 | Other Positive Correlation |
| ATG12 | 0.063085 | 0.258261 | Other Positive Correlation |
| AP1G1 | 0.063055 | 0.258488 | Other Positive Correlation |
| ODAD1 | 0.063037 | 0.258625 | Other Positive Correlation |
| STAT5A | 0.063023 | 0.258729 | Other Positive Correlation |
| ZC3H12B | 0.063 | 0.258903 | Other Positive Correlation |
| LRRC32 | 0.062999 | 0.258915 | Other Positive Correlation |
| MED22 | 0.062983 | 0.259033 | Other Positive Correlation |
| PABPC5 | 0.062982 | 0.25904 | Other Positive Correlation |
| HCCS | 0.062893 | 0.259718 | Other Positive Correlation |
| TMPRSS3 | 0.06289 | 0.259739 | Other Positive Correlation |
| ACOT2 | 0.062868 | 0.259906 | Other Positive Correlation |
| NEURL2 | 0.062834 | 0.260168 | Other Positive Correlation |
| TMLHE | 0.062824 | 0.260244 | Other Positive Correlation |
| STK32B | 0.062816 | 0.260303 | Other Positive Correlation |
| ADAM22 | 0.062805 | 0.260388 | Other Positive Correlation |
| UBR3 | 0.062788 | 0.260514 | Other Positive Correlation |
| ZKSCAN5 | 0.062784 | 0.260544 | Other Positive Correlation |
| PLD6 | 0.062762 | 0.260716 | Other Positive Correlation |

| | | | |
|---|---|---|---|
| HIPK4 | 0.062749 | 0.260814 | Other Positive Correlation |
| GRHL3 | 0.062653 | 0.261544 | Other Positive Correlation |
| ARID3C | 0.062639 | 0.261649 | Other Positive Correlation |
| PIM1 | 0.062623 | 0.261774 | Other Positive Correlation |
| CACNA1F | 0.062615 | 0.261835 | Other Positive Correlation |
| INPP5E | 0.062586 | 0.262052 | Other Positive Correlation |
| HLA-DQB1 | 0.062564 | 0.262221 | Other Positive Correlation |
| ZNF529 | 0.06249 | 0.262792 | Other Positive Correlation |
| EIF2B5 | 0.062473 | 0.26292 | Other Positive Correlation |
| FUNDC2 | 0.062446 | 0.263126 | Other Positive Correlation |
| TBXA2R | 0.062446 | 0.263127 | Other Positive Correlation |
| TMEM52 | 0.062439 | 0.263176 | Other Positive Correlation |
| E2F6 | 0.06241 | 0.263401 | Other Positive Correlation |
| SPPL2B | 0.062371 | 0.2637 | Other Positive Correlation |
| MSTO1 | 0.062345 | 0.263898 | Other Positive Correlation |
| ZNF700 | 0.06234 | 0.263935 | Other Positive Correlation |
| CCS | 0.062333 | 0.263996 | Other Positive Correlation |
| OOSP4A | 0.062326 | 0.264045 | Other Positive Correlation |
| SOCS1 | 0.062307 | 0.264195 | Other Positive Correlation |
| SH2B1 | 0.062306 | 0.264199 | Other Positive Correlation |
| TRMT12 | 0.062272 | 0.264465 | Other Positive Correlation |
| LRP1B | 0.062211 | 0.264929 | Other Positive Correlation |
| RAB21 | 0.062207 | 0.264962 | Other Positive Correlation |
| AMPD3 | 0.062187 | 0.265112 | Other Positive Correlation |
| FOXH1 | 0.062176 | 0.265202 | Other Positive Correlation |
| NANS | 0.062144 | 0.26545 | Other Positive Correlation |
| EDN2 | 0.062103 | 0.265766 | Other Positive Correlation |
| NLRC5 | 0.062098 | 0.2658 | Other Positive Correlation |
| BRD4 | 0.06206 | 0.266093 | Other Positive Correlation |
| IRF1 | 0.062052 | 0.266152 | Other Positive Correlation |
| EVPLL | 0.062022 | 0.266385 | Other Positive Correlation |
| ATG14 | 0.062022 | 0.266387 | Other Positive Correlation |
| PLA2G15 | 0.062017 | 0.266426 | Other Positive Correlation |
| ZNF275 | 0.061992 | 0.266618 | Other Positive Correlation |
| DENND1A | 0.061954 | 0.266912 | Other Positive Correlation |
| TCTN3 | 0.061947 | 0.266964 | Other Positive Correlation |
| AMH | 0.061902 | 0.267316 | Other Positive Correlation |
| RAB15 | 0.061883 | 0.267459 | Other Positive Correlation |
| DNAJC25 | 0.06187 | 0.267563 | Other Positive Correlation |
| MATK | 0.06187 | 0.267563 | Other Positive Correlation |
| USP36 | 0.061845 | 0.267756 | Other Positive Correlation |
| HSPA6 | 0.06184 | 0.267799 | Other Positive Correlation |
| TBC1D12 | 0.061824 | 0.267923 | Other Positive Correlation |
| FOLR1 | 0.061804 | 0.268077 | Other Positive Correlation |
| C17orf75 | 0.061793 | 0.268161 | Other Positive Correlation |
| LARP4 | 0.061774 | 0.268306 | Other Positive Correlation |
| AK9 | 0.061723 | 0.268701 | Other Positive Correlation |
| FKRP | 0.061722 | 0.268709 | Other Positive Correlation |

| | | | |
|---|---|---|---|
| TMCO5A | 0.061705 | 0.268846 | Other Positive Correlation |
| MTUS2 | 0.061677 | 0.269063 | Other Positive Correlation |
| SSTR5 | 0.061649 | 0.26928 | Other Positive Correlation |
| EXOSC1 | 0.061634 | 0.269399 | Other Positive Correlation |
| AMBRA1 | 0.061619 | 0.269515 | Other Positive Correlation |
| BCDIN3D | 0.06159 | 0.269739 | Other Positive Correlation |
| WDR45 | 0.061583 | 0.269795 | Other Positive Correlation |
| RAP1GAP | 0.061583 | 0.269796 | Other Positive Correlation |
| ZNF428 | 0.061576 | 0.269846 | Other Positive Correlation |
| MED30 | 0.061517 | 0.270306 | Other Positive Correlation |
| CXCL12 | 0.061515 | 0.270323 | Other Positive Correlation |
| POP4 | 0.061514 | 0.270334 | Other Positive Correlation |
| CALML5 | 0.061495 | 0.270478 | Other Positive Correlation |
| ZFP90 | 0.061467 | 0.2707 | Other Positive Correlation |
| IKZF5 | 0.061449 | 0.270836 | Other Positive Correlation |
| HLA-DMB | 0.061447 | 0.270854 | Other Positive Correlation |
| TFAP2B | 0.061441 | 0.270902 | Other Positive Correlation |
| CDC23 | 0.061437 | 0.270935 | Other Positive Correlation |
| SLFN12L | 0.061425 | 0.271024 | Other Positive Correlation |
| IL12RB1 | 0.061411 | 0.271139 | Other Positive Correlation |
| ANKRD45 | 0.061409 | 0.271151 | Other Positive Correlation |
| ZNF469 | 0.0614 | 0.271219 | Other Positive Correlation |
| FYCO1 | 0.061358 | 0.271554 | Other Positive Correlation |
| GGCX | 0.06133 | 0.271766 | Other Positive Correlation |
| NRBP1 | 0.061307 | 0.27195 | Other Positive Correlation |
| DEPTOR | 0.061296 | 0.272036 | Other Positive Correlation |
| GARIN1B | 0.061294 | 0.272055 | Other Positive Correlation |
| ZNF107 | 0.061294 | 0.272055 | Other Positive Correlation |
| MEI4 | 0.061279 | 0.272167 | Other Positive Correlation |
| PARG | 0.061245 | 0.272439 | Other Positive Correlation |
| SPNS3 | 0.061243 | 0.272454 | Other Positive Correlation |
| FAM102B | 0.06123 | 0.272553 | Other Positive Correlation |
| MYCL | 0.061211 | 0.272699 | Other Positive Correlation |
| ANKRD42 | 0.061205 | 0.272749 | Other Positive Correlation |
| ZNF417 | 0.061203 | 0.272767 | Other Positive Correlation |
| CRISPLD2 | 0.061194 | 0.272832 | Other Positive Correlation |
| DENND5B | 0.061162 | 0.27309 | Other Positive Correlation |
| IL17RB | 0.061156 | 0.273136 | Other Positive Correlation |
| ZNF431 | 0.061153 | 0.273159 | Other Positive Correlation |
| TTC9B | 0.061114 | 0.273466 | Other Positive Correlation |
| UQCRFS1 | 0.061109 | 0.273506 | Other Positive Correlation |
| RASD2 | 0.061045 | 0.274008 | Other Positive Correlation |
| ACSS3 | 0.061043 | 0.274023 | Other Positive Correlation |
| PRXL2A | 0.061031 | 0.274117 | Other Positive Correlation |
| NACC2 | 0.060986 | 0.274473 | Other Positive Correlation |
| NUPR2 | 0.060961 | 0.274672 | Other Positive Correlation |
| EIF5A2 | 0.060922 | 0.274976 | Other Positive Correlation |
| LHX6 | 0.06092 | 0.274992 | Other Positive Correlation |

| | | | |
|---|---|---|---|
| DPEP3 | 0.060917 | 0.275018 | Other Positive Correlation |
| PROCA1 | 0.060907 | 0.275096 | Other Positive Correlation |
| CLEC20A | 0.060871 | 0.275383 | Other Positive Correlation |
| RSPO4 | 0.060833 | 0.275684 | Other Positive Correlation |
| AEBP1 | 0.060806 | 0.275897 | Other Positive Correlation |
| SYNDIG1 | 0.060778 | 0.276119 | Other Positive Correlation |
| CRABP1 | 0.06076 | 0.27626 | Other Positive Correlation |
| CD247 | 0.06075 | 0.276338 | Other Positive Correlation |
| DLG3 | 0.060746 | 0.276367 | Other Positive Correlation |
| ABCA2 | 0.060685 | 0.27685 | Other Positive Correlation |
| FGF14 | 0.060684 | 0.276864 | Other Positive Correlation |
| ZNF66 | 0.060627 | 0.277315 | Other Positive Correlation |
| HKDC1 | 0.060623 | 0.277347 | Other Positive Correlation |
| IRX4 | 0.060606 | 0.277482 | Other Positive Correlation |
| CALHM3 | 0.060602 | 0.277512 | Other Positive Correlation |
| ATP6V1FNB | 0.060549 | 0.277929 | Other Positive Correlation |
| ATF7IP2 | 0.060528 | 0.278101 | Other Positive Correlation |
| DOK1 | 0.06051 | 0.278239 | Other Positive Correlation |
| FRAT2 | 0.060475 | 0.278522 | Other Positive Correlation |
| NFATC4 | 0.060441 | 0.278795 | Other Positive Correlation |
| EFHB | 0.060399 | 0.279129 | Other Positive Correlation |
| ZNF208 | 0.060374 | 0.279323 | Other Positive Correlation |
| CTDSPL2 | 0.060364 | 0.279404 | Other Positive Correlation |
| RTCB | 0.060356 | 0.279467 | Other Positive Correlation |
| CDAN1 | 0.060347 | 0.279542 | Other Positive Correlation |
| PLIN1 | 0.060334 | 0.279647 | Other Positive Correlation |
| ME3 | 0.060333 | 0.279651 | Other Positive Correlation |
| TMDD1 | 0.060333 | 0.279656 | Other Positive Correlation |
| PABIR1 | 0.060326 | 0.279708 | Other Positive Correlation |
| C2CD6 | 0.060303 | 0.279892 | Other Positive Correlation |
| BTC | 0.06029 | 0.279997 | Other Positive Correlation |
| RLN1 | 0.060268 | 0.280174 | Other Positive Correlation |
| ATG4D | 0.060129 | 0.281284 | Other Positive Correlation |
| ZNF189 | 0.060124 | 0.281325 | Other Positive Correlation |
| EDEM2 | 0.060115 | 0.281395 | Other Positive Correlation |
| SMIM43 | 0.0601 | 0.281519 | Other Positive Correlation |
| IKZF4 | 0.060082 | 0.281665 | Other Positive Correlation |
| CGB3 | 0.060075 | 0.281721 | Other Positive Correlation |
| CCR2 | 0.060075 | 0.281721 | Other Positive Correlation |
| PI16 | 0.060075 | 0.281721 | Other Positive Correlation |
| PNLIPRP2 | 0.060075 | 0.281721 | Other Positive Correlation |
| IL18R1 | 0.060061 | 0.281835 | Other Positive Correlation |
| CHDH | 0.059908 | 0.283061 | Other Positive Correlation |
| DUSP3 | 0.059886 | 0.283239 | Other Positive Correlation |
| TLE6 | 0.059885 | 0.283243 | Other Positive Correlation |
| CEP120 | 0.059882 | 0.283267 | Other Positive Correlation |
| GEMIN7 | 0.059862 | 0.283431 | Other Positive Correlation |
| MOB4 | 0.059851 | 0.283522 | Other Positive Correlation |

| | | | |
|---|---|---|---|
| PCGF3 | 0.059848 | 0.283543 | Other Positive Correlation |
| GLT8D2 | 0.059798 | 0.28395 | Other Positive Correlation |
| GIMAP5 | 0.059776 | 0.284123 | Other Positive Correlation |
| SLC27A4 | 0.059755 | 0.284295 | Other Positive Correlation |
| TNFSF13B | 0.059749 | 0.284344 | Other Positive Correlation |
| PPP6R3 | 0.05974 | 0.284418 | Other Positive Correlation |
| C19orf44 | 0.059732 | 0.284478 | Other Positive Correlation |
| NOTCH2NLC | 0.05971 | 0.284658 | Other Positive Correlation |
| GABBR2 | 0.059708 | 0.284671 | Other Positive Correlation |
| CTXN2 | 0.059677 | 0.284927 | Other Positive Correlation |
| EML6 | 0.059671 | 0.284976 | Other Positive Correlation |
| SLC39A9 | 0.059644 | 0.285195 | Other Positive Correlation |
| KLK12 | 0.059603 | 0.285525 | Other Positive Correlation |
| ZNF585B | 0.059594 | 0.285594 | Other Positive Correlation |
| MYB | 0.059573 | 0.285765 | Other Positive Correlation |
| PLA2G7 | 0.05956 | 0.285873 | Other Positive Correlation |
| GPR137B | 0.059531 | 0.286104 | Other Positive Correlation |
| MED13 | 0.059524 | 0.286161 | Other Positive Correlation |
| PSKH1 | 0.059509 | 0.286283 | Other Positive Correlation |
| FOXF2 | 0.059489 | 0.286447 | Other Positive Correlation |
| CYP3A4 | 0.059477 | 0.286546 | Other Positive Correlation |
| NFE2 | 0.059467 | 0.286622 | Other Positive Correlation |
| NLRP14 | 0.059459 | 0.286692 | Other Positive Correlation |
| KCNK10 | 0.059423 | 0.286983 | Other Positive Correlation |
| C1orf167 | 0.059407 | 0.28711 | Other Positive Correlation |
| TMEM171 | 0.059357 | 0.287518 | Other Positive Correlation |
| OGT | 0.059333 | 0.287717 | Other Positive Correlation |
| MUC3A | 0.059327 | 0.287763 | Other Positive Correlation |
| TMEM86B | 0.059277 | 0.28817 | Other Positive Correlation |
| SLC35B4 | 0.059271 | 0.288221 | Other Positive Correlation |
| VPS33A | 0.059261 | 0.2883 | Other Positive Correlation |
| FUT1 | 0.059252 | 0.288373 | Other Positive Correlation |
| CHAF1B | 0.059246 | 0.28842 | Other Positive Correlation |
| DLK1 | 0.059238 | 0.288486 | Other Positive Correlation |
| UBE2QL1 | 0.059238 | 0.288486 | Other Positive Correlation |
| NMBR | 0.059238 | 0.288486 | Other Positive Correlation |
| ENSG00000256206 | 0.059238 | 0.288486 | Other Positive Correlation |
| DPPA5 | 0.059238 | 0.288486 | Other Positive Correlation |
| PTPRC | 0.059238 | 0.288486 | Other Positive Correlation |
| ARHGAP33 | 0.059193 | 0.288853 | Other Positive Correlation |
| ZYG11A | 0.059169 | 0.289054 | Other Positive Correlation |
| NDUFA9 | 0.059169 | 0.289054 | Other Positive Correlation |
| ENDOG | 0.059168 | 0.28906 | Other Positive Correlation |
| TMEM184C | 0.059151 | 0.289202 | Other Positive Correlation |
| SEMA3E | 0.05915 | 0.289209 | Other Positive Correlation |
| RTL5 | 0.059148 | 0.289227 | Other Positive Correlation |
| PACSIN3 | 0.059146 | 0.289244 | Other Positive Correlation |
| CCDC183 | 0.059145 | 0.28925 | Other Positive Correlation |

| | | | |
|---|---|---|---|
| LSG1 | 0.059139 | 0.2893 | Other Positive Correlation |
| ANKLE1 | 0.059088 | 0.289713 | Other Positive Correlation |
| YJU2B | 0.059078 | 0.2898 | Other Positive Correlation |
| FAM25A | 0.05907 | 0.289858 | Other Positive Correlation |
| ODF3L2 | 0.059062 | 0.289926 | Other Positive Correlation |
| GSTM2 | 0.05906 | 0.289941 | Other Positive Correlation |
| TC2N | 0.059044 | 0.290072 | Other Positive Correlation |
| BUD13 | 0.058986 | 0.290545 | Other Positive Correlation |
| MEX3D | 0.058963 | 0.290737 | Other Positive Correlation |
| DEPDC4 | 0.058932 | 0.290991 | Other Positive Correlation |
| SP9 | 0.058891 | 0.291325 | Other Positive Correlation |
| UVSSA | 0.058885 | 0.291381 | Other Positive Correlation |
| VSTM5 | 0.058881 | 0.291407 | Other Positive Correlation |
| PBLD | 0.05887 | 0.291497 | Other Positive Correlation |
| LEPROTL1 | 0.058868 | 0.291519 | Other Positive Correlation |
| ACVR1C | 0.058842 | 0.29173 | Other Positive Correlation |
| C2CD4A | 0.058829 | 0.29184 | Other Positive Correlation |
| ENSG00000286165 | 0.058804 | 0.292042 | Other Positive Correlation |
| PRCD | 0.05879 | 0.292157 | Other Positive Correlation |
| RAP2A | 0.058772 | 0.292306 | Other Positive Correlation |
| IL5 | 0.058767 | 0.292348 | Other Positive Correlation |
| TBCC | 0.058752 | 0.292471 | Other Positive Correlation |
| TCEAL4 | 0.058747 | 0.292513 | Other Positive Correlation |
| ZDHHC21 | 0.058742 | 0.292548 | Other Positive Correlation |
| GSG1 | 0.058738 | 0.292583 | Other Positive Correlation |
| FASTKD5 | 0.058731 | 0.292642 | Other Positive Correlation |
| P2RX4 | 0.058703 | 0.292873 | Other Positive Correlation |
| CCDC9 | 0.058662 | 0.29321 | Other Positive Correlation |
| GPR19 | 0.058644 | 0.293361 | Other Positive Correlation |
| KMT5C | 0.058603 | 0.293696 | Other Positive Correlation |
| CCDC102B | 0.058593 | 0.293782 | Other Positive Correlation |
| KHDC3L | 0.058575 | 0.293932 | Other Positive Correlation |
| PTK2B | 0.058489 | 0.294636 | Other Positive Correlation |
| ABCB9 | 0.058477 | 0.29474 | Other Positive Correlation |
| FGD5 | 0.058449 | 0.294971 | Other Positive Correlation |
| C11orf68 | 0.058448 | 0.294975 | Other Positive Correlation |
| AMMECR1L | 0.0584 | 0.295379 | Other Positive Correlation |
| XKR4 | 0.058395 | 0.295416 | Other Positive Correlation |
| ZNF625 | 0.058388 | 0.295479 | Other Positive Correlation |
| ZNF676 | 0.058385 | 0.295504 | Other Positive Correlation |
| PSMC4 | 0.05837 | 0.295623 | Other Positive Correlation |
| TMEM233 | 0.058365 | 0.295666 | Other Positive Correlation |
| ANO8 | 0.05835 | 0.295788 | Other Positive Correlation |
| NFATC1 | 0.058308 | 0.296138 | Other Positive Correlation |
| C8orf76 | 0.058307 | 0.296143 | Other Positive Correlation |
| PCDH9 | 0.058254 | 0.29659 | Other Positive Correlation |
| GIPC2 | 0.058234 | 0.29675 | Other Positive Correlation |
| H2BC15 | 0.05823 | 0.29679 | Other Positive Correlation |

| | | | |
|---|---|---|---|
| IRF4 | 0.058208 | 0.296969 | Other Positive Correlation |
| FKBPL | 0.058197 | 0.297064 | Other Positive Correlation |
| KANK2 | 0.058195 | 0.297081 | Other Positive Correlation |
| VGF | 0.058193 | 0.297096 | Other Positive Correlation |
| TSPAN17 | 0.058185 | 0.297158 | Other Positive Correlation |
| KCTD5 | 0.058182 | 0.297184 | Other Positive Correlation |
| ZNF263 | 0.058163 | 0.297345 | Other Positive Correlation |
| HOXA9 | 0.058162 | 0.297351 | Other Positive Correlation |
| LGI4 | 0.058127 | 0.297639 | Other Positive Correlation |
| ZNF678 | 0.058126 | 0.297653 | Other Positive Correlation |
| CHCT1 | 0.058125 | 0.297658 | Other Positive Correlation |
| KLF1 | 0.058114 | 0.297755 | Other Positive Correlation |
| FAXC | 0.058109 | 0.297796 | Other Positive Correlation |
| C2CD2 | 0.058084 | 0.298004 | Other Positive Correlation |
| TMEM175 | 0.058023 | 0.298509 | Other Positive Correlation |
| LSM1 | 0.058017 | 0.298559 | Other Positive Correlation |
| GPATCH3 | 0.05798 | 0.298872 | Other Positive Correlation |
| FOXJ1 | 0.057979 | 0.298876 | Other Positive Correlation |
| GOLGA5 | 0.057938 | 0.299221 | Other Positive Correlation |
| SV2B | 0.057883 | 0.29968 | Other Positive Correlation |
| GPC5 | 0.057873 | 0.29976 | Other Positive Correlation |
| DLX5 | 0.057863 | 0.29985 | Other Positive Correlation |
| ZNF347 | 0.057862 | 0.299853 | Other Positive Correlation |
| FAM174A | 0.057842 | 0.300024 | Other Positive Correlation |
| FAM172A | 0.05783 | 0.300123 | Other Positive Correlation |
| SOBP | 0.057817 | 0.30023 | Other Positive Correlation |
| TIMM10B | 0.057807 | 0.300317 | Other Positive Correlation |
| CLN3 | 0.057774 | 0.300595 | Other Positive Correlation |
| ANKZF1 | 0.057748 | 0.300808 | Other Positive Correlation |
| RDH10 | 0.057744 | 0.300842 | Other Positive Correlation |
| ZNF740 | 0.057739 | 0.300889 | Other Positive Correlation |
| ROPN1B | 0.057706 | 0.301161 | Other Positive Correlation |
| DENND4A | 0.057701 | 0.301208 | Other Positive Correlation |
| TRAF5 | 0.057697 | 0.30124 | Other Positive Correlation |
| SOX2 | 0.057693 | 0.301275 | Other Positive Correlation |
| IFNL3 | 0.057636 | 0.301752 | Other Positive Correlation |
| FRMPD3 | 0.057636 | 0.301752 | Other Positive Correlation |
| PCP4 | 0.057636 | 0.301752 | Other Positive Correlation |
| TMEM132A | 0.057622 | 0.301866 | Other Positive Correlation |
| NID2 | 0.057601 | 0.302044 | Other Positive Correlation |
| NTN1 | 0.057589 | 0.302146 | Other Positive Correlation |
| ZMPSTE24 | 0.057551 | 0.302468 | Other Positive Correlation |
| MYO5C | 0.057547 | 0.3025 | Other Positive Correlation |
| ENSG00000256950 | 0.057546 | 0.302505 | Other Positive Correlation |
| SOWAHB | 0.057538 | 0.302571 | Other Positive Correlation |
| DENND5A | 0.057533 | 0.302619 | Other Positive Correlation |
| SLC18A2 | 0.057505 | 0.302853 | Other Positive Correlation |
| IL4I1 | 0.057503 | 0.302872 | Other Positive Correlation |

| | | | |
|---|---|---|---|
| PRRT2 | 0.057479 | 0.303067 | Other Positive Correlation |
| PRKCG | 0.057476 | 0.303098 | Other Positive Correlation |
| HDHD3 | 0.057473 | 0.303124 | Other Positive Correlation |
| FAM240C | 0.057465 | 0.303185 | Other Positive Correlation |
| VWF | 0.05744 | 0.303395 | Other Positive Correlation |
| TMEM255A | 0.057438 | 0.303417 | Other Positive Correlation |
| AVPR1B | 0.057422 | 0.303547 | Other Positive Correlation |
| ZNF648 | 0.057422 | 0.303547 | Other Positive Correlation |
| PSMB11 | 0.057422 | 0.303547 | Other Positive Correlation |
| SHPK | 0.05739 | 0.303824 | Other Positive Correlation |
| CPNE2 | 0.05736 | 0.304075 | Other Positive Correlation |
| SYNRG | 0.057344 | 0.304209 | Other Positive Correlation |
| RASSF2 | 0.057336 | 0.30428 | Other Positive Correlation |
| ZNF444 | 0.05733 | 0.304325 | Other Positive Correlation |
| C2 | 0.057274 | 0.304802 | Other Positive Correlation |
| DYRK2 | 0.057268 | 0.304852 | Other Positive Correlation |
| MPO | 0.057249 | 0.305009 | Other Positive Correlation |
| ASPG | 0.057225 | 0.305215 | Other Positive Correlation |
| AVPI1 | 0.057209 | 0.305351 | Other Positive Correlation |
| CHFR | 0.057203 | 0.305403 | Other Positive Correlation |
| CHRNA4 | 0.057199 | 0.305433 | Other Positive Correlation |
| SMPDL3B | 0.05718 | 0.305596 | Other Positive Correlation |
| UNC93B1 | 0.057171 | 0.305676 | Other Positive Correlation |
| RNASEK-C17orf49 | 0.057147 | 0.305879 | Other Positive Correlation |
| ACRBP | 0.057141 | 0.305931 | Other Positive Correlation |
| PARP11 | 0.057119 | 0.306114 | Other Positive Correlation |
| SH2B2 | 0.057116 | 0.306137 | Other Positive Correlation |
| NEURL3 | 0.057115 | 0.306149 | Other Positive Correlation |
| ERLEC1 | 0.057111 | 0.306183 | Other Positive Correlation |
| ATL1 | 0.057105 | 0.306231 | Other Positive Correlation |
| EMC8 | 0.057079 | 0.306456 | Other Positive Correlation |
| MINAR1 | 0.057022 | 0.306941 | Other Positive Correlation |
| SNRNP27 | 0.056981 | 0.307285 | Other Positive Correlation |
| TMEM91 | 0.056981 | 0.307288 | Other Positive Correlation |
| ZNF154 | 0.05698 | 0.307291 | Other Positive Correlation |
| CCDC158 | 0.05698 | 0.307299 | Other Positive Correlation |
| RAMP2 | 0.056971 | 0.307368 | Other Positive Correlation |
| PIAS4 | 0.056954 | 0.307513 | Other Positive Correlation |
| TRABD2A | 0.056947 | 0.307571 | Other Positive Correlation |
| KLC2 | 0.056881 | 0.308137 | Other Positive Correlation |
| TPST1 | 0.056864 | 0.30828 | Other Positive Correlation |
| SOX1 | 0.056849 | 0.30841 | Other Positive Correlation |
| INTS8 | 0.056816 | 0.308693 | Other Positive Correlation |
| RNF148 | 0.056806 | 0.30878 | Other Positive Correlation |
| LTBP1 | 0.056738 | 0.309353 | Other Positive Correlation |
| CDC7 | 0.056729 | 0.30943 | Other Positive Correlation |
| USP6NL | 0.056685 | 0.30981 | Other Positive Correlation |
| ZNF883 | 0.056677 | 0.30988 | Other Positive Correlation |

| | | | |
|---|---|---|---|
| VSIG8 | 0.056675 | 0.309893 | Other Positive Correlation |
| TMEM54 | 0.056673 | 0.309912 | Other Positive Correlation |
| SCIN | 0.056672 | 0.309923 | Other Positive Correlation |
| PCDHGB4 | 0.056662 | 0.310004 | Other Positive Correlation |
| EPHA10 | 0.056636 | 0.310225 | Other Positive Correlation |
| ABTB1 | 0.056629 | 0.31029 | Other Positive Correlation |
| MLLT3 | 0.056602 | 0.310518 | Other Positive Correlation |
| ZNF518B | 0.05659 | 0.310618 | Other Positive Correlation |
| RC3H1 | 0.056575 | 0.310747 | Other Positive Correlation |
| NPDC1 | 0.05655 | 0.310965 | Other Positive Correlation |
| SCYL1 | 0.056474 | 0.311615 | Other Positive Correlation |
| PLA2G2A | 0.056445 | 0.31187 | Other Positive Correlation |
| NIBAN3 | 0.056442 | 0.311889 | Other Positive Correlation |
| ZFP69 | 0.056413 | 0.312141 | Other Positive Correlation |
| ONECUT3 | 0.056406 | 0.312199 | Other Positive Correlation |
| UBFD1 | 0.056397 | 0.312277 | Other Positive Correlation |
| GOLM2 | 0.056356 | 0.312631 | Other Positive Correlation |
| ICMT | 0.056352 | 0.312667 | Other Positive Correlation |
| SOSTDC1 | 0.056351 | 0.312673 | Other Positive Correlation |
| CACNA1C | 0.056315 | 0.31298 | Other Positive Correlation |
| YBEY | 0.056265 | 0.313413 | Other Positive Correlation |
| NR5A2 | 0.056222 | 0.313783 | Other Positive Correlation |
| PLK5 | 0.056222 | 0.313783 | Other Positive Correlation |
| HOXC12 | 0.056222 | 0.313783 | Other Positive Correlation |
| ENSG00000269533 | 0.056222 | 0.313783 | Other Positive Correlation |
| DNAAF6 | 0.056222 | 0.313783 | Other Positive Correlation |
| PPP4R4 | 0.056221 | 0.313797 | Other Positive Correlation |
| ST20-MTHFS | 0.0562 | 0.313971 | Other Positive Correlation |
| OTX1 | 0.056195 | 0.31402 | Other Positive Correlation |
| ZNF286A | 0.056194 | 0.314026 | Other Positive Correlation |
| ENO4 | 0.05614 | 0.314495 | Other Positive Correlation |
| VENTX | 0.056066 | 0.315135 | Other Positive Correlation |
| GPRASP1 | 0.056065 | 0.315138 | Other Positive Correlation |
| TRMT9B | 0.056046 | 0.315306 | Other Positive Correlation |
| RABL2A | 0.056041 | 0.315347 | Other Positive Correlation |
| CDC42SE2 | 0.05597 | 0.315963 | Other Positive Correlation |
| NPY1R | 0.055961 | 0.316042 | Other Positive Correlation |
| PRM3 | 0.055957 | 0.316075 | Other Positive Correlation |
| SCRN3 | 0.055953 | 0.316107 | Other Positive Correlation |
| TMEM39A | 0.055941 | 0.316211 | Other Positive Correlation |
| PBX3 | 0.055939 | 0.316228 | Other Positive Correlation |
| ITGAX | 0.055939 | 0.316228 | Other Positive Correlation |
| RNF222 | 0.055919 | 0.316408 | Other Positive Correlation |
| TRIM49B | 0.0559 | 0.31657 | Other Positive Correlation |
| RASAL3 | 0.055866 | 0.316864 | Other Positive Correlation |
| CAVIN4 | 0.055848 | 0.317021 | Other Positive Correlation |
| MRPL43 | 0.055835 | 0.317133 | Other Positive Correlation |
| RNASEK | 0.055818 | 0.317284 | Other Positive Correlation |

| | | | |
|---|---|---|---|
| USP39 | 0.055802 | 0.31742 | Other Positive Correlation |
| SFRP2 | 0.055798 | 0.317455 | Other Positive Correlation |
| CD8B | 0.055798 | 0.317455 | Other Positive Correlation |
| ADRA1A | 0.055798 | 0.317455 | Other Positive Correlation |
| ITIH3 | 0.055798 | 0.317455 | Other Positive Correlation |
| JAKMIP1 | 0.055798 | 0.317455 | Other Positive Correlation |
| ENSG00000285329 | 0.055797 | 0.317462 | Other Positive Correlation |
| MBD3L2B | 0.055797 | 0.317462 | Other Positive Correlation |
| MUC4 | 0.055783 | 0.317587 | Other Positive Correlation |
| CCDC71L | 0.055751 | 0.317859 | Other Positive Correlation |
| ABCC4 | 0.055747 | 0.317901 | Other Positive Correlation |
| SKI | 0.055691 | 0.318388 | Other Positive Correlation |
| ZNF165 | 0.055683 | 0.318456 | Other Positive Correlation |
| OSCAR | 0.055681 | 0.318475 | Other Positive Correlation |
| ENSG00000187186 | 0.055665 | 0.318609 | Other Positive Correlation |
| TTC38 | 0.055645 | 0.318782 | Other Positive Correlation |
| FOXF1 | 0.055639 | 0.318837 | Other Positive Correlation |
| TSPO2 | 0.055615 | 0.319042 | Other Positive Correlation |
| DNAAF3 | 0.055606 | 0.319125 | Other Positive Correlation |
| AKR1A1 | 0.055546 | 0.319644 | Other Positive Correlation |
| DRAM2 | 0.05552 | 0.319872 | Other Positive Correlation |
| SEC16A | 0.055508 | 0.319976 | Other Positive Correlation |
| UVRAG | 0.055471 | 0.3203 | Other Positive Correlation |
| ZNF100 | 0.055465 | 0.320359 | Other Positive Correlation |
| GRAMD1A | 0.055459 | 0.320404 | Other Positive Correlation |
| ALX1 | 0.055408 | 0.320854 | Other Positive Correlation |
| ENO3 | 0.055399 | 0.320936 | Other Positive Correlation |
| TMEM129 | 0.055385 | 0.321052 | Other Positive Correlation |
| CNTN5 | 0.055364 | 0.321241 | Other Positive Correlation |
| ADAMTS8 | 0.055364 | 0.321241 | Other Positive Correlation |
| GFY | 0.055364 | 0.321241 | Other Positive Correlation |
| GDAP2 | 0.055344 | 0.321414 | Other Positive Correlation |
| PAX5 | 0.055309 | 0.321724 | Other Positive Correlation |
| TEN1 | 0.055307 | 0.321742 | Other Positive Correlation |
| SYT5 | 0.05528 | 0.321978 | Other Positive Correlation |
| GRID1 | 0.05524 | 0.322327 | Other Positive Correlation |
| GPM6A | 0.055232 | 0.322395 | Other Positive Correlation |
| ASB4 | 0.055229 | 0.32242 | Other Positive Correlation |
| PPP1R12C | 0.055184 | 0.322813 | Other Positive Correlation |
| STX8 | 0.055184 | 0.32282 | Other Positive Correlation |
| NAT14 | 0.055176 | 0.322885 | Other Positive Correlation |
| KIT | 0.055139 | 0.32321 | Other Positive Correlation |
| PTGDS | 0.055115 | 0.323423 | Other Positive Correlation |
| CHMP2B | 0.055109 | 0.323474 | Other Positive Correlation |
| DEDD2 | 0.055097 | 0.323578 | Other Positive Correlation |
| RDH16 | 0.055084 | 0.323693 | Other Positive Correlation |
| CACNA1B | 0.055084 | 0.323698 | Other Positive Correlation |
| GTF2E1 | 0.055083 | 0.323702 | Other Positive Correlation |

| | | | |
|---|---|---|---|
| SHC3 | 0.055082 | 0.323709 | Other Positive Correlation |
| GAB3 | 0.05503 | 0.324174 | Other Positive Correlation |
| CRYBG2 | 0.055025 | 0.324213 | Other Positive Correlation |
| RIMS3 | 0.055013 | 0.324317 | Other Positive Correlation |
| ORMDL3 | 0.054977 | 0.324635 | Other Positive Correlation |
| PCTP | 0.054972 | 0.324681 | Other Positive Correlation |
| ESR2 | 0.05496 | 0.324792 | Other Positive Correlation |
| ECM1 | 0.054946 | 0.324907 | Other Positive Correlation |
| CNKSR1 | 0.054939 | 0.324973 | Other Positive Correlation |
| PPFIA4 | 0.054935 | 0.325006 | Other Positive Correlation |
| SPESP1 | 0.054932 | 0.325031 | Other Positive Correlation |
| KCNK3 | 0.054918 | 0.325162 | Other Positive Correlation |
| RASL12 | 0.054899 | 0.32533 | Other Positive Correlation |
| TMEM45A | 0.054894 | 0.325371 | Other Positive Correlation |
| CASP8AP2 | 0.054891 | 0.325395 | Other Positive Correlation |
| SWT1 | 0.054879 | 0.325502 | Other Positive Correlation |
| ILVBL | 0.05486 | 0.325667 | Other Positive Correlation |
| TVP23C-CDRT4 | 0.05485 | 0.325759 | Other Positive Correlation |
| LEMD3 | 0.054841 | 0.325837 | Other Positive Correlation |
| TRMT2B | 0.054827 | 0.325962 | Other Positive Correlation |
| FNDC1 | 0.054811 | 0.326107 | Other Positive Correlation |
| TMEM8B | 0.054807 | 0.326134 | Other Positive Correlation |
| TERB1 | 0.054776 | 0.326414 | Other Positive Correlation |
| ARF4 | 0.054773 | 0.326436 | Other Positive Correlation |
| STEAP4 | 0.054765 | 0.326511 | Other Positive Correlation |
| MTRFR | 0.054758 | 0.326568 | Other Positive Correlation |
| IDNK | 0.054733 | 0.326792 | Other Positive Correlation |
| PON3 | 0.054683 | 0.327238 | Other Positive Correlation |
| GPR107 | 0.054682 | 0.327241 | Other Positive Correlation |
| CR2 | 0.054646 | 0.327564 | Other Positive Correlation |
| TMEM150C | 0.054643 | 0.327593 | Other Positive Correlation |
| PHETA2 | 0.054633 | 0.327682 | Other Positive Correlation |
| ZNF670-ZNF695 | 0.054619 | 0.327807 | Other Positive Correlation |
| CCN5 | 0.054616 | 0.327829 | Other Positive Correlation |
| SLC39A8 | 0.054608 | 0.327903 | Other Positive Correlation |
| GID4 | 0.054556 | 0.328366 | Other Positive Correlation |
| CSTPP1 | 0.054533 | 0.32857 | Other Positive Correlation |
| FAM228B | 0.054521 | 0.328676 | Other Positive Correlation |
| CEACAM1 | 0.054518 | 0.328704 | Other Positive Correlation |
| GPR156 | 0.054499 | 0.328871 | Other Positive Correlation |
| LRRC73 | 0.054488 | 0.328967 | Other Positive Correlation |
| USP12 | 0.054483 | 0.329012 | Other Positive Correlation |
| GLDC | 0.054476 | 0.329072 | Other Positive Correlation |
| GPLD1 | 0.054473 | 0.329099 | Other Positive Correlation |
| UGT3A2 | 0.054464 | 0.329176 | Other Positive Correlation |
| ABRAXAS2 | 0.054459 | 0.329223 | Other Positive Correlation |
| PTPRR | 0.054458 | 0.329236 | Other Positive Correlation |
| KLK10 | 0.05439 | 0.329836 | Other Positive Correlation |

| | | | |
|---|---|---|---|
| NRBF2 | 0.054362 | 0.330092 | Other Positive Correlation |
| PPP3CC | 0.05432 | 0.330464 | Other Positive Correlation |
| CCDC28A | 0.054315 | 0.330505 | Other Positive Correlation |
| PABPN1L | 0.0543 | 0.330639 | Other Positive Correlation |
| CAPSL | 0.0543 | 0.330639 | Other Positive Correlation |
| ZNF35 | 0.05429 | 0.330733 | Other Positive Correlation |
| ZNF611 | 0.054276 | 0.33085 | Other Positive Correlation |
| SLC36A4 | 0.054266 | 0.330944 | Other Positive Correlation |
| LRRC38 | 0.054248 | 0.331107 | Other Positive Correlation |
| GSC2 | 0.054248 | 0.331107 | Other Positive Correlation |
| ETNPPL | 0.054248 | 0.331107 | Other Positive Correlation |
| ASB18 | 0.054248 | 0.331107 | Other Positive Correlation |
| CHAD | 0.054248 | 0.331107 | Other Positive Correlation |
| LENG1 | 0.054241 | 0.331166 | Other Positive Correlation |
| C4orf46 | 0.054209 | 0.331454 | Other Positive Correlation |
| TMEM223 | 0.054204 | 0.331501 | Other Positive Correlation |
| HMGCL | 0.054192 | 0.331609 | Other Positive Correlation |
| CRYGD | 0.054181 | 0.331703 | Other Positive Correlation |
| RAMP3 | 0.054181 | 0.331703 | Other Positive Correlation |
| SLC5A8 | 0.054181 | 0.331703 | Other Positive Correlation |
| WSCD2 | 0.054181 | 0.331703 | Other Positive Correlation |
| ZNF559-ZNF177 | 0.054181 | 0.331703 | Other Positive Correlation |
| CCR9 | 0.054181 | 0.331703 | Other Positive Correlation |
| TMEM220 | 0.05417 | 0.331798 | Other Positive Correlation |
| MAF | 0.054136 | 0.332106 | Other Positive Correlation |
| EMC7 | 0.054074 | 0.332663 | Other Positive Correlation |
| USP35 | 0.054069 | 0.332706 | Other Positive Correlation |
| CDKL3 | 0.05402 | 0.333147 | Other Positive Correlation |
| SNX9 | 0.053988 | 0.33343 | Other Positive Correlation |
| TMEM270 | 0.05398 | 0.333505 | Other Positive Correlation |
| SIX5 | 0.05396 | 0.333681 | Other Positive Correlation |
| LCTL | 0.053947 | 0.333798 | Other Positive Correlation |
| PHF10 | 0.053907 | 0.334159 | Other Positive Correlation |
| UBTD2 | 0.053902 | 0.334199 | Other Positive Correlation |
| CD6 | 0.053897 | 0.334249 | Other Positive Correlation |
| CHCHD4 | 0.053894 | 0.334271 | Other Positive Correlation |
| DDN | 0.053883 | 0.334372 | Other Positive Correlation |
| ACACB | 0.053825 | 0.334895 | Other Positive Correlation |
| USP27X | 0.053808 | 0.335045 | Other Positive Correlation |
| C15orf62 | 0.0538 | 0.335117 | Other Positive Correlation |
| UPK2 | 0.053797 | 0.335144 | Other Positive Correlation |
| DNAI4 | 0.053793 | 0.335182 | Other Positive Correlation |
| HECTD2 | 0.053755 | 0.335521 | Other Positive Correlation |
| COPE | 0.05374 | 0.33566 | Other Positive Correlation |
| TRIM49C | 0.053735 | 0.335705 | Other Positive Correlation |
| TRIM43 | 0.053735 | 0.335705 | Other Positive Correlation |
| ARHGEF35 | 0.053734 | 0.335718 | Other Positive Correlation |
| IRAK3 | 0.053688 | 0.336127 | Other Positive Correlation |

| | | | |
|---|---|---|---|
| NTRK1 | 0.053681 | 0.336187 | Other Positive Correlation |
| ENSG00000286221 | 0.053671 | 0.336284 | Other Positive Correlation |
| S1PR3 | 0.053613 | 0.336804 | Other Positive Correlation |
| ZIC2 | 0.05361 | 0.336832 | Other Positive Correlation |
| SHISA3 | 0.053593 | 0.336988 | Other Positive Correlation |
| PLOD1 | 0.053592 | 0.33699 | Other Positive Correlation |
| CSMD2 | 0.053523 | 0.337618 | Other Positive Correlation |
| RLN2 | 0.053515 | 0.337692 | Other Positive Correlation |
| ZNF69 | 0.053508 | 0.337753 | Other Positive Correlation |
| CACNA1D | 0.053488 | 0.337929 | Other Positive Correlation |
| STK38 | 0.053459 | 0.338192 | Other Positive Correlation |
| GRIN3A | 0.053458 | 0.338208 | Other Positive Correlation |
| TCHH | 0.053399 | 0.338743 | Other Positive Correlation |
| OR2AG1 | 0.05336 | 0.339096 | Other Positive Correlation |
| SLC38A9 | 0.053333 | 0.339337 | Other Positive Correlation |
| CCDC159 | 0.05331 | 0.339546 | Other Positive Correlation |
| CNOT6L | 0.05328 | 0.339821 | Other Positive Correlation |
| ADGRB2 | 0.053224 | 0.340325 | Other Positive Correlation |
| FTCDNL1 | 0.053188 | 0.340657 | Other Positive Correlation |
| PPL | 0.053183 | 0.340699 | Other Positive Correlation |
| GIMAP2 | 0.053179 | 0.34074 | Other Positive Correlation |
| DCAF16 | 0.05316 | 0.34091 | Other Positive Correlation |
| ZNF729 | 0.053143 | 0.341066 | Other Positive Correlation |
| SLC4A1AP | 0.053131 | 0.341175 | Other Positive Correlation |
| FCGBP | 0.05308 | 0.341639 | Other Positive Correlation |
| TACC2 | 0.053065 | 0.34177 | Other Positive Correlation |
| MIA3 | 0.052982 | 0.342528 | Other Positive Correlation |
| NAA60 | 0.052961 | 0.342719 | Other Positive Correlation |
| PHF13 | 0.052942 | 0.342897 | Other Positive Correlation |
| PCP2 | 0.052935 | 0.342961 | Other Positive Correlation |
| RBM46 | 0.052893 | 0.343345 | Other Positive Correlation |
| RTN4R | 0.052889 | 0.343383 | Other Positive Correlation |
| INIP | 0.052881 | 0.343453 | Other Positive Correlation |
| ALG9 | 0.052875 | 0.343503 | Other Positive Correlation |
| ENSG00000283782 | 0.052858 | 0.343666 | Other Positive Correlation |
| ADCYAP1R1 | 0.052846 | 0.343768 | Other Positive Correlation |
| CACNG7 | 0.052846 | 0.343768 | Other Positive Correlation |
| C4orf51 | 0.052846 | 0.343768 | Other Positive Correlation |
| HS3ST5 | 0.052846 | 0.343768 | Other Positive Correlation |
| TLR2 | 0.052827 | 0.343945 | Other Positive Correlation |
| SMG9 | 0.052821 | 0.344001 | Other Positive Correlation |
| RALGAPB | 0.052821 | 0.344002 | Other Positive Correlation |
| HNF1B | 0.052819 | 0.344018 | Other Positive Correlation |
| PRR14L | 0.052782 | 0.344359 | Other Positive Correlation |
| RHBDL3 | 0.052781 | 0.344364 | Other Positive Correlation |
| MOG | 0.052772 | 0.344446 | Other Positive Correlation |
| BRINP1 | 0.052763 | 0.344528 | Other Positive Correlation |
| LDHD | 0.052755 | 0.344608 | Other Positive Correlation |

| | | | |
|---|---|---|---|
| FAIM2 | 0.052732 | 0.344819 | Other Positive Correlation |
| SELENBP1 | 0.052724 | 0.344886 | Other Positive Correlation |
| SETD9 | 0.052711 | 0.345009 | Other Positive Correlation |
| NINJ1 | 0.052689 | 0.345209 | Other Positive Correlation |
| WHRN | 0.052689 | 0.345212 | Other Positive Correlation |
| FMO5 | 0.052676 | 0.345327 | Other Positive Correlation |
| RECQL5 | 0.052673 | 0.345353 | Other Positive Correlation |
| FAM184B | 0.052648 | 0.345586 | Other Positive Correlation |
| CALHM6 | 0.052636 | 0.345697 | Other Positive Correlation |
| ANKRD54 | 0.052628 | 0.345769 | Other Positive Correlation |
| ADCK1 | 0.052621 | 0.34583 | Other Positive Correlation |
| ZIC5 | 0.052604 | 0.345992 | Other Positive Correlation |
| LIN28A | 0.052604 | 0.345992 | Other Positive Correlation |
| HTR3A | 0.052576 | 0.34625 | Other Positive Correlation |
| SLURP1 | 0.052576 | 0.34625 | Other Positive Correlation |
| FTCD | 0.052576 | 0.34625 | Other Positive Correlation |
| ACAN | 0.052576 | 0.34625 | Other Positive Correlation |
| GAP43 | 0.052576 | 0.34625 | Other Positive Correlation |
| PRSS56 | 0.052576 | 0.34625 | Other Positive Correlation |
| LRFN2 | 0.052576 | 0.34625 | Other Positive Correlation |
| SMCHD1 | 0.052565 | 0.346344 | Other Positive Correlation |
| FGF3 | 0.052546 | 0.346526 | Other Positive Correlation |
| SLC25A51 | 0.052529 | 0.346677 | Other Positive Correlation |
| HSPB11 | 0.05252 | 0.346763 | Other Positive Correlation |
| FAM76B | 0.052464 | 0.347274 | Other Positive Correlation |
| ZNF763 | 0.052427 | 0.347618 | Other Positive Correlation |
| CNFN | 0.05241 | 0.347772 | Other Positive Correlation |
| GPC4 | 0.052389 | 0.347963 | Other Positive Correlation |
| ZNF566 | 0.052266 | 0.349101 | Other Positive Correlation |
| ALDH3B2 | 0.052261 | 0.34915 | Other Positive Correlation |
| DRD5 | 0.052257 | 0.349183 | Other Positive Correlation |
| ARMC8 | 0.052232 | 0.349417 | Other Positive Correlation |
| RASSF4 | 0.052229 | 0.349439 | Other Positive Correlation |
| CHRNA7 | 0.052209 | 0.349625 | Other Positive Correlation |
| R3HCC1 | 0.052182 | 0.349875 | Other Positive Correlation |
| ISL2 | 0.052173 | 0.349961 | Other Positive Correlation |
| USP3 | 0.052148 | 0.350196 | Other Positive Correlation |
| PCK1 | 0.052145 | 0.350223 | Other Positive Correlation |
| TERT | 0.052145 | 0.350223 | Other Positive Correlation |
| KCTD9 | 0.052137 | 0.350292 | Other Positive Correlation |
| OLFML2B | 0.052122 | 0.350431 | Other Positive Correlation |
| PTGR1 | 0.052113 | 0.35052 | Other Positive Correlation |
| ZNF559 | 0.052093 | 0.350706 | Other Positive Correlation |
| CNRIP1 | 0.052086 | 0.350769 | Other Positive Correlation |
| ZNF879 | 0.052083 | 0.350797 | Other Positive Correlation |
| PXK | 0.052078 | 0.350841 | Other Positive Correlation |
| ARNT2 | 0.052074 | 0.350876 | Other Positive Correlation |
| LCLAT1 | 0.052054 | 0.351063 | Other Positive Correlation |

| | | | |
|---|---|---|---|
| CDH7 | 0.052049 | 0.351107 | Other Positive Correlation |
| SLC6A1 | 0.052049 | 0.351107 | Other Positive Correlation |
| RAB19 | 0.052003 | 0.351531 | Other Positive Correlation |
| STARD8 | 0.052003 | 0.351531 | Other Positive Correlation |
| CERS2 | 0.051978 | 0.351768 | Other Positive Correlation |
| WNT5A | 0.051929 | 0.352225 | Other Positive Correlation |
| CRB2 | 0.05192 | 0.352308 | Other Positive Correlation |
| CHGB | 0.051915 | 0.352348 | Other Positive Correlation |
| CHIC1 | 0.051911 | 0.352393 | Other Positive Correlation |
| RPS6KC1 | 0.051893 | 0.352553 | Other Positive Correlation |
| FNTB | 0.051871 | 0.352758 | Other Positive Correlation |
| SPDYA | 0.051867 | 0.352793 | Other Positive Correlation |
| ZNF2 | 0.05186 | 0.352862 | Other Positive Correlation |
| HOXC6 | 0.051855 | 0.35291 | Other Positive Correlation |
| VPREB3 | 0.05185 | 0.352958 | Other Positive Correlation |
| SRD5A3 | 0.051839 | 0.353054 | Other Positive Correlation |
| COPS5 | 0.051834 | 0.3531 | Other Positive Correlation |
| BBOF1 | 0.051811 | 0.35332 | Other Positive Correlation |
| FAM161B | 0.051806 | 0.353363 | Other Positive Correlation |
| ZFP2 | 0.051759 | 0.353807 | Other Positive Correlation |
| EEPD1 | 0.051757 | 0.353818 | Other Positive Correlation |
| IFT22 | 0.051756 | 0.353831 | Other Positive Correlation |
| TNNT3 | 0.051746 | 0.35392 | Other Positive Correlation |
| CYBA | 0.051707 | 0.354286 | Other Positive Correlation |
| DPY19L1 | 0.051695 | 0.354398 | Other Positive Correlation |
| ZCWPW1 | 0.051687 | 0.354476 | Other Positive Correlation |
| SCGB2B2 | 0.05165 | 0.354821 | Other Positive Correlation |
| BTN2A1 | 0.051644 | 0.354876 | Other Positive Correlation |
| SHB | 0.051642 | 0.354892 | Other Positive Correlation |
| PTPRB | 0.051618 | 0.355113 | Other Positive Correlation |
| KIF27 | 0.051599 | 0.35529 | Other Positive Correlation |
| RHBDD2 | 0.051558 | 0.355676 | Other Positive Correlation |
| NUDT2 | 0.051528 | 0.355959 | Other Positive Correlation |
| SUPT20H | 0.051526 | 0.355973 | Other Positive Correlation |
| FKBP2 | 0.051504 | 0.356185 | Other Positive Correlation |
| TEX38 | 0.051499 | 0.356227 | Other Positive Correlation |
| DNAI1 | 0.051463 | 0.356566 | Other Positive Correlation |
| CSF1R | 0.051437 | 0.356809 | Other Positive Correlation |
| ADAT3 | 0.051355 | 0.357575 | Other Positive Correlation |
| SNX2 | 0.051264 | 0.358434 | Other Positive Correlation |
| BRMS1L | 0.051262 | 0.358451 | Other Positive Correlation |
| ENSG00000258691 | 0.051234 | 0.358717 | Other Positive Correlation |
| ADORA3 | 0.051234 | 0.358717 | Other Positive Correlation |
| ENPP6 | 0.051234 | 0.358717 | Other Positive Correlation |
| SIRT6 | 0.051225 | 0.358801 | Other Positive Correlation |
| CST6 | 0.051206 | 0.358975 | Other Positive Correlation |
| EPHB2 | 0.051171 | 0.359304 | Other Positive Correlation |
| TMEM74 | 0.051142 | 0.359577 | Other Positive Correlation |

| | | | |
|---|---|---|---|
| PEX2 | 0.051134 | 0.359655 | Other Positive Correlation |
| PVALB | 0.051104 | 0.359932 | Other Positive Correlation |
| SPATA1 | 0.0511 | 0.359969 | Other Positive Correlation |
| SPATA6 | 0.051077 | 0.360192 | Other Positive Correlation |
| CAMK2G | 0.051075 | 0.36021 | Other Positive Correlation |
| LCN12 | 0.051071 | 0.36025 | Other Positive Correlation |
| MZB1 | 0.051049 | 0.360449 | Other Positive Correlation |
| EPG5 | 0.051045 | 0.360495 | Other Positive Correlation |
| CDKL2 | 0.051035 | 0.360588 | Other Positive Correlation |
| RYR2 | 0.051005 | 0.360865 | Other Positive Correlation |
| HOXC11 | 0.050999 | 0.360923 | Other Positive Correlation |
| PACC1 | 0.050985 | 0.361055 | Other Positive Correlation |
| CASKIN2 | 0.050979 | 0.361116 | Other Positive Correlation |
| KMT2D | 0.050978 | 0.361124 | Other Positive Correlation |
| LRCH4 | 0.050966 | 0.36124 | Other Positive Correlation |
| BCL2L15 | 0.050939 | 0.361494 | Other Positive Correlation |
| ELK1 | 0.050893 | 0.361927 | Other Positive Correlation |
| NOD2 | 0.05088 | 0.36205 | Other Positive Correlation |
| ABT1 | 0.050867 | 0.362175 | Other Positive Correlation |
| VPS8 | 0.050867 | 0.362175 | Other Positive Correlation |
| ENSG00000170846 | 0.050863 | 0.362204 | Other Positive Correlation |
| ITGB3 | 0.050837 | 0.362454 | Other Positive Correlation |
| EXOC3L2 | 0.050829 | 0.362534 | Other Positive Correlation |
| KPTN | 0.050798 | 0.362824 | Other Positive Correlation |
| SMG6 | 0.050797 | 0.362827 | Other Positive Correlation |
| ZUP1 | 0.050796 | 0.36284 | Other Positive Correlation |
| STX4 | 0.050766 | 0.363121 | Other Positive Correlation |
| NELFA | 0.050762 | 0.363162 | Other Positive Correlation |
| NUP42 | 0.050699 | 0.363757 | Other Positive Correlation |
| XKR7 | 0.050699 | 0.363758 | Other Positive Correlation |
| CADPS | 0.050699 | 0.363758 | Other Positive Correlation |
| LMAN1L | 0.050699 | 0.363758 | Other Positive Correlation |
| GOLGA8H | 0.050699 | 0.363758 | Other Positive Correlation |
| TM4SF5 | 0.050699 | 0.363758 | Other Positive Correlation |
| SIRT1 | 0.050696 | 0.363786 | Other Positive Correlation |
| VASH2 | 0.050695 | 0.363795 | Other Positive Correlation |
| DYSF | 0.050657 | 0.36416 | Other Positive Correlation |
| TMEM87A | 0.050624 | 0.364466 | Other Positive Correlation |
| ZNF703 | 0.050621 | 0.364496 | Other Positive Correlation |
| FAM78B | 0.050604 | 0.364664 | Other Positive Correlation |
| FAF2 | 0.050594 | 0.364756 | Other Positive Correlation |
| GDPGP1 | 0.050591 | 0.364787 | Other Positive Correlation |
| RGS6 | 0.050567 | 0.365011 | Other Positive Correlation |
| ITPR2 | 0.050566 | 0.365022 | Other Positive Correlation |
| STX1A | 0.050549 | 0.365179 | Other Positive Correlation |
| OAZ3 | 0.050517 | 0.365487 | Other Positive Correlation |
| GABRD | 0.050506 | 0.365595 | Other Positive Correlation |
| SMAD6 | 0.050499 | 0.365662 | Other Positive Correlation |

| | | | |
|---|---|---|---|
| TELO2 | 0.050497 | 0.365675 | Other Positive Correlation |
| ZCCHC7 | 0.050495 | 0.365698 | Other Positive Correlation |
| OBI1 | 0.05044 | 0.366221 | Other Positive Correlation |
| TEX264 | 0.050437 | 0.366244 | Other Positive Correlation |
| PCDH10 | 0.0504 | 0.366601 | Other Positive Correlation |
| CDK8 | 0.050392 | 0.366674 | Other Positive Correlation |
| BLVRB | 0.050377 | 0.366816 | Other Positive Correlation |
| CRYGC | 0.050343 | 0.367143 | Other Positive Correlation |
| DGCR6 | 0.050306 | 0.367494 | Other Positive Correlation |
| CAMKMT | 0.05028 | 0.367745 | Other Positive Correlation |
| WDR26 | 0.050274 | 0.367796 | Other Positive Correlation |
| GPSM1 | 0.050274 | 0.367801 | Other Positive Correlation |
| DUOXA1 | 0.050262 | 0.367917 | Other Positive Correlation |
| SLC25A53 | 0.050251 | 0.368016 | Other Positive Correlation |
| GRIK3 | 0.050222 | 0.3683 | Other Positive Correlation |
| CNMD | 0.050222 | 0.3683 | Other Positive Correlation |
| CRHR2 | 0.050222 | 0.3683 | Other Positive Correlation |
| NCBP2L | 0.050222 | 0.3683 | Other Positive Correlation |
| SIRT4 | 0.050213 | 0.368381 | Other Positive Correlation |
| METTL21A | 0.050174 | 0.368754 | Other Positive Correlation |
| ZNF597 | 0.050157 | 0.368921 | Other Positive Correlation |
| GDPD3 | 0.050131 | 0.369171 | Other Positive Correlation |
| ABCB8 | 0.050129 | 0.369189 | Other Positive Correlation |
| TBC1D17 | 0.050095 | 0.369508 | Other Positive Correlation |
| DEPP1 | 0.050085 | 0.369607 | Other Positive Correlation |
| KLK4 | 0.050065 | 0.369794 | Other Positive Correlation |
| MT4 | 0.050061 | 0.369838 | Other Positive Correlation |
| UNG | 0.050045 | 0.369988 | Other Positive Correlation |
| KLHL40 | 0.050036 | 0.370075 | Other Positive Correlation |
| PLPP1 | 0.049993 | 0.370488 | Other Positive Correlation |
| RORB | 0.049988 | 0.370538 | Other Positive Correlation |
| SNX30 | 0.049972 | 0.370693 | Other Positive Correlation |
| ECHDC3 | 0.049969 | 0.370714 | Other Positive Correlation |
| RNF152 | 0.049953 | 0.370871 | Other Positive Correlation |
| TARS3 | 0.049908 | 0.371299 | Other Positive Correlation |
| PRDM4 | 0.049905 | 0.371332 | Other Positive Correlation |
| DGKQ | 0.04989 | 0.371475 | Other Positive Correlation |
| RCCD1 | 0.04987 | 0.371669 | Other Positive Correlation |
| ANGPT2 | 0.049857 | 0.37179 | Other Positive Correlation |
| ZBTB40 | 0.049822 | 0.372127 | Other Positive Correlation |
| GGCT | 0.049811 | 0.372237 | Other Positive Correlation |
| FUT3 | 0.049801 | 0.372325 | Other Positive Correlation |
| GSTT2 | 0.049729 | 0.373026 | Other Positive Correlation |
| ADAMTS3 | 0.049726 | 0.373053 | Other Positive Correlation |
| IPCEF1 | 0.049722 | 0.373089 | Other Positive Correlation |
| ARC | 0.049721 | 0.373099 | Other Positive Correlation |
| CLN5 | 0.049718 | 0.373131 | Other Positive Correlation |
| FILIP1 | 0.049698 | 0.373318 | Other Positive Correlation |

| | | | |
|---|---|---|---|
| ZCCHC12 | 0.049693 | 0.373374 | Other Positive Correlation |
| OSTM1 | 0.049687 | 0.37343 | Other Positive Correlation |
| NKX2-2 | 0.049679 | 0.373506 | Other Positive Correlation |
| CLCNKB | 0.049668 | 0.373611 | Other Positive Correlation |
| UNKL | 0.049661 | 0.373677 | Other Positive Correlation |
| MST1 | 0.049645 | 0.373836 | Other Positive Correlation |
| ZSCAN29 | 0.049586 | 0.374398 | Other Positive Correlation |
| HTATSF1 | 0.049563 | 0.374621 | Other Positive Correlation |
| HOXA5 | 0.049534 | 0.374906 | Other Positive Correlation |
| SEC14L1 | 0.049503 | 0.375197 | Other Positive Correlation |
| POU3F3 | 0.049503 | 0.375206 | Other Positive Correlation |
| ZNF502 | 0.0495 | 0.375231 | Other Positive Correlation |
| RSPH10B2 | 0.049468 | 0.37554 | Other Positive Correlation |
| C1QL3 | 0.049468 | 0.37554 | Other Positive Correlation |
| CCDC30 | 0.049464 | 0.375583 | Other Positive Correlation |
| PPP3R1 | 0.049449 | 0.375727 | Other Positive Correlation |
| POU2AF3 | 0.049434 | 0.375868 | Other Positive Correlation |
| STYK1 | 0.049429 | 0.37592 | Other Positive Correlation |
| CARF | 0.049428 | 0.375929 | Other Positive Correlation |
| INKA2 | 0.049396 | 0.376232 | Other Positive Correlation |
| TRIM34 | 0.049391 | 0.376286 | Other Positive Correlation |
| PHF21B | 0.049371 | 0.376476 | Other Positive Correlation |
| CEP44 | 0.049364 | 0.376545 | Other Positive Correlation |
| FOXRED2 | 0.049267 | 0.37748 | Other Positive Correlation |
| TM2D2 | 0.049256 | 0.377592 | Other Positive Correlation |
| RNF149 | 0.049251 | 0.377637 | Other Positive Correlation |
| ZNF546 | 0.049233 | 0.377817 | Other Positive Correlation |
| SLITRK5 | 0.049232 | 0.377822 | Other Positive Correlation |
| PRAP1 | 0.04922 | 0.377938 | Other Positive Correlation |
| TMEM63A | 0.049195 | 0.378179 | Other Positive Correlation |
| SNAP47 | 0.04919 | 0.378236 | Other Positive Correlation |
| ADGRL3 | 0.049169 | 0.378431 | Other Positive Correlation |
| BRD1 | 0.049165 | 0.378472 | Other Positive Correlation |
| PRPF19 | 0.049146 | 0.378656 | Other Positive Correlation |
| MTMR10 | 0.049134 | 0.378777 | Other Positive Correlation |
| ZNF491 | 0.049117 | 0.378941 | Other Positive Correlation |
| VEPH1 | 0.049101 | 0.3791 | Other Positive Correlation |
| ZNF564 | 0.049095 | 0.379157 | Other Positive Correlation |
| ZNF136 | 0.049083 | 0.379269 | Other Positive Correlation |
| DRD4 | 0.049068 | 0.379416 | Other Positive Correlation |
| NAGS | 0.04905 | 0.379594 | Other Positive Correlation |
| ZNF43 | 0.049034 | 0.379746 | Other Positive Correlation |
| IL17C | 0.049014 | 0.379939 | Other Positive Correlation |
| ABHD1 | 0.049006 | 0.380022 | Other Positive Correlation |
| KSR2 | 0.048949 | 0.380577 | Other Positive Correlation |
| IL23R | 0.048927 | 0.380787 | Other Positive Correlation |
| ZMYND15 | 0.048848 | 0.381559 | Other Positive Correlation |
| GLB1 | 0.048845 | 0.381585 | Other Positive Correlation |

| | | | |
|---|---|---|---|
| PDGFD | 0.048837 | 0.381671 | Other Positive Correlation |
| DPEP1 | 0.048816 | 0.381871 | Other Positive Correlation |
| KLHL29 | 0.048812 | 0.381908 | Other Positive Correlation |
| PRKD2 | 0.048812 | 0.381911 | Other Positive Correlation |
| CD79B | 0.048803 | 0.381997 | Other Positive Correlation |
| CBX6 | 0.048785 | 0.382175 | Other Positive Correlation |
| ZNF571 | 0.048763 | 0.382386 | Other Positive Correlation |
| NECTIN4 | 0.048763 | 0.382388 | Other Positive Correlation |
| ZBTB34 | 0.048744 | 0.38258 | Other Positive Correlation |
| PPP1R21 | 0.048683 | 0.38317 | Other Positive Correlation |
| AKR1B15 | 0.048638 | 0.383618 | Other Positive Correlation |
| S100A7 | 0.048607 | 0.383918 | Other Positive Correlation |
| KATNIP | 0.0486 | 0.383988 | Other Positive Correlation |
| CFAP73 | 0.048576 | 0.384218 | Other Positive Correlation |
| GON7 | 0.048561 | 0.384367 | Other Positive Correlation |
| TXNL4B | 0.048555 | 0.384431 | Other Positive Correlation |
| KHK | 0.048551 | 0.384468 | Other Positive Correlation |
| AVEN | 0.048536 | 0.384617 | Other Positive Correlation |
| NCAN | 0.048533 | 0.384638 | Other Positive Correlation |
| CPPED1 | 0.048521 | 0.384755 | Other Positive Correlation |
| HCFC2 | 0.048516 | 0.384811 | Other Positive Correlation |
| KCNQ3 | 0.048455 | 0.385404 | Other Positive Correlation |
| TAF7 | 0.048432 | 0.385633 | Other Positive Correlation |
| ACSBG1 | 0.048425 | 0.385698 | Other Positive Correlation |
| TSPYL2 | 0.048424 | 0.385712 | Other Positive Correlation |
| SLC25A18 | 0.048336 | 0.386575 | Other Positive Correlation |
| ZNF7 | 0.048323 | 0.386703 | Other Positive Correlation |
| POLR3D | 0.048285 | 0.387078 | Other Positive Correlation |
| BATF | 0.048274 | 0.387183 | Other Positive Correlation |
| KLHL11 | 0.048247 | 0.387453 | Other Positive Correlation |
| RASA2 | 0.048236 | 0.387557 | Other Positive Correlation |
| BET1L | 0.048194 | 0.387972 | Other Positive Correlation |
| PGBD4 | 0.048187 | 0.388041 | Other Positive Correlation |
| CNTD1 | 0.048178 | 0.38813 | Other Positive Correlation |
| SPATA17 | 0.048174 | 0.38817 | Other Positive Correlation |
| FYN | 0.048144 | 0.388467 | Other Positive Correlation |
| WBP1 | 0.048139 | 0.388515 | Other Positive Correlation |
| TXNDC11 | 0.048115 | 0.388754 | Other Positive Correlation |
| LRRC14 | 0.048084 | 0.389059 | Other Positive Correlation |
| LAMB2 | 0.048058 | 0.389315 | Other Positive Correlation |
| ZNF177 | 0.048058 | 0.38932 | Other Positive Correlation |
| ALOX5 | 0.048057 | 0.38933 | Other Positive Correlation |
| AGTPBP1 | 0.048043 | 0.389468 | Other Positive Correlation |
| MAD2L1BP | 0.048034 | 0.38955 | Other Positive Correlation |
| ZNF555 | 0.048029 | 0.389605 | Other Positive Correlation |
| JCAD | 0.048023 | 0.389665 | Other Positive Correlation |
| RALYL | 0.048012 | 0.389776 | Other Positive Correlation |
| CDC14A | 0.047995 | 0.389941 | Other Positive Correlation |

| | | | |
|---|---|---|---|
| C3orf80 | 0.047931 | 0.390571 | Other Positive Correlation |
| MTCP1 | 0.047882 | 0.391062 | Other Positive Correlation |
| SHISAL2A | 0.047881 | 0.391066 | Other Positive Correlation |
| EEFSEC | 0.047865 | 0.39123 | Other Positive Correlation |
| LCNL1 | 0.047851 | 0.39136 | Other Positive Correlation |
| TCEAL5 | 0.047851 | 0.39136 | Other Positive Correlation |
| RPH3A | 0.047851 | 0.39136 | Other Positive Correlation |
| LARP1B | 0.047836 | 0.39151 | Other Positive Correlation |
| ZBTB14 | 0.047807 | 0.391805 | Other Positive Correlation |
| CHST10 | 0.047806 | 0.391813 | Other Positive Correlation |
| LIPE | 0.047789 | 0.391979 | Other Positive Correlation |
| TMEM187 | 0.047779 | 0.392075 | Other Positive Correlation |
| CTNNA2 | 0.047756 | 0.392306 | Other Positive Correlation |
| HOXA6 | 0.047743 | 0.392435 | Other Positive Correlation |
| TBC1D1 | 0.047738 | 0.39249 | Other Positive Correlation |
| PABPC1L2B | 0.04772 | 0.392661 | Other Positive Correlation |
| PDPK1 | 0.047715 | 0.392714 | Other Positive Correlation |
| RUNDC3A | 0.047714 | 0.392723 | Other Positive Correlation |
| NME1-NME2 | 0.047702 | 0.392848 | Other Positive Correlation |
| NKAP | 0.0477 | 0.392863 | Other Positive Correlation |
| NCOA6 | 0.047684 | 0.393019 | Other Positive Correlation |
| AGPAT1 | 0.047673 | 0.393133 | Other Positive Correlation |
| DYNC1LI2 | 0.047659 | 0.393265 | Other Positive Correlation |
| DCAF15 | 0.047641 | 0.393448 | Other Positive Correlation |
| PPP1R37 | 0.047617 | 0.393685 | Other Positive Correlation |
| SLC5A1 | 0.047577 | 0.394086 | Other Positive Correlation |
| FAM3C | 0.047556 | 0.394298 | Other Positive Correlation |
| KCTD1 | 0.047543 | 0.394427 | Other Positive Correlation |
| NUBP1 | 0.047499 | 0.394859 | Other Positive Correlation |
| GAREM1 | 0.047449 | 0.395361 | Other Positive Correlation |
| OLIG2 | 0.047438 | 0.395471 | Other Positive Correlation |
| C1orf141 | 0.047438 | 0.395471 | Other Positive Correlation |
| SMIM26 | 0.047426 | 0.395594 | Other Positive Correlation |
| RHOB | 0.04741 | 0.395745 | Other Positive Correlation |
| RNF38 | 0.047374 | 0.396104 | Other Positive Correlation |
| ZNF862 | 0.047351 | 0.396343 | Other Positive Correlation |
| STK17B | 0.047339 | 0.396454 | Other Positive Correlation |
| PTPDC1 | 0.047324 | 0.396604 | Other Positive Correlation |
| MGAT2 | 0.04731 | 0.396751 | Other Positive Correlation |
| WNT10A | 0.0473 | 0.396853 | Other Positive Correlation |
| WDR5B | 0.047257 | 0.397279 | Other Positive Correlation |
| EXOC3L1 | 0.047246 | 0.397387 | Other Positive Correlation |
| A1BG | 0.047244 | 0.397408 | Other Positive Correlation |
| ZNF404 | 0.047212 | 0.397729 | Other Positive Correlation |
| PDZRN3 | 0.047197 | 0.397878 | Other Positive Correlation |
| SUPV3L1 | 0.047189 | 0.397959 | Other Positive Correlation |
| LRRC3 | 0.047171 | 0.39814 | Other Positive Correlation |
| PHF24 | 0.04712 | 0.398647 | Other Positive Correlation |

| | | | |
|---|---|---|---|
| OR3A3 | 0.04711 | 0.398749 | Other Positive Correlation |
| FBXO15 | 0.047104 | 0.398805 | Other Positive Correlation |
| PHYH | 0.047101 | 0.398837 | Other Positive Correlation |
| SERTAD3 | 0.047094 | 0.398909 | Other Positive Correlation |
| PTCD1 | 0.047059 | 0.399264 | Other Positive Correlation |
| LHX8 | 0.046994 | 0.39991 | Other Positive Correlation |
| KCNJ9 | 0.046994 | 0.39991 | Other Positive Correlation |
| CRYGN | 0.046994 | 0.39991 | Other Positive Correlation |
| INSM1 | 0.046994 | 0.39991 | Other Positive Correlation |
| UNC13D | 0.046987 | 0.399985 | Other Positive Correlation |
| ZNF75A | 0.046976 | 0.400091 | Other Positive Correlation |
| DYNLT2 | 0.046952 | 0.400333 | Other Positive Correlation |
| LINS1 | 0.046923 | 0.400629 | Other Positive Correlation |
| HECA | 0.046887 | 0.400987 | Other Positive Correlation |
| DHDDS | 0.046887 | 0.400992 | Other Positive Correlation |
| CACNA1I | 0.046877 | 0.401091 | Other Positive Correlation |
| CHRNB1 | 0.046871 | 0.401149 | Other Positive Correlation |
| ACOT1 | 0.046855 | 0.401307 | Other Positive Correlation |
| YY2 | 0.046831 | 0.40155 | Other Positive Correlation |
| STAB1 | 0.046831 | 0.401557 | Other Positive Correlation |
| SLC51A | 0.046822 | 0.40164 | Other Positive Correlation |
| HSPA5 | 0.046803 | 0.401833 | Other Positive Correlation |
| UBXN10 | 0.046802 | 0.401842 | Other Positive Correlation |
| CSTF1 | 0.04679 | 0.401967 | Other Positive Correlation |
| SNAI1 | 0.046782 | 0.402047 | Other Positive Correlation |
| ASF1A | 0.046767 | 0.402191 | Other Positive Correlation |
| GPR89B | 0.046705 | 0.402816 | Other Positive Correlation |
| DYNC2I2 | 0.046685 | 0.403027 | Other Positive Correlation |
| LRCH2 | 0.046656 | 0.403316 | Other Positive Correlation |
| NKAIN3 | 0.046641 | 0.403469 | Other Positive Correlation |
| YAE1 | 0.046626 | 0.403618 | Other Positive Correlation |
| COQ10A | 0.046606 | 0.403816 | Other Positive Correlation |
| CCNI2 | 0.046596 | 0.403918 | Other Positive Correlation |
| BCORL1 | 0.046585 | 0.40403 | Other Positive Correlation |
| SIAH1 | 0.046576 | 0.40412 | Other Positive Correlation |
| ASCL5 | 0.046565 | 0.404231 | Other Positive Correlation |
| CCDC17 | 0.046547 | 0.404415 | Other Positive Correlation |
| SLC46A2 | 0.046518 | 0.40471 | Other Positive Correlation |
| GET1-SH3BGR | 0.046518 | 0.40471 | Other Positive Correlation |
| ABCB1 | 0.046518 | 0.40471 | Other Positive Correlation |
| ZNF865 | 0.04651 | 0.404792 | Other Positive Correlation |
| TSHR | 0.046498 | 0.404914 | Other Positive Correlation |
| ZCCHC18 | 0.046494 | 0.404956 | Other Positive Correlation |
| KIAA0513 | 0.046483 | 0.405063 | Other Positive Correlation |
| COX4I2 | 0.046462 | 0.40528 | Other Positive Correlation |
| ABCD4 | 0.046434 | 0.405555 | Other Positive Correlation |
| LNPK | 0.046374 | 0.406169 | Other Positive Correlation |
| CPO | 0.046354 | 0.406366 | Other Positive Correlation |

| | | | |
|---|---|---|---|
| PLEKHF2 | 0.046339 | 0.406518 | Other Positive Correlation |
| NUP43 | 0.046334 | 0.406575 | Other Positive Correlation |
| HIRA | 0.046322 | 0.406697 | Other Positive Correlation |
| NRXN3 | 0.046318 | 0.406733 | Other Positive Correlation |
| ATP6V0A4 | 0.046299 | 0.40693 | Other Positive Correlation |
| CLP1 | 0.046288 | 0.407043 | Other Positive Correlation |
| CATSPERZ | 0.046286 | 0.407057 | Other Positive Correlation |
| TNPO1 | 0.046283 | 0.407085 | Other Positive Correlation |
| ATPAF1 | 0.046276 | 0.407159 | Other Positive Correlation |
| MXD1 | 0.046262 | 0.407304 | Other Positive Correlation |
| PKD1L2 | 0.046254 | 0.407382 | Other Positive Correlation |
| PRSS16 | 0.046202 | 0.407909 | Other Positive Correlation |
| REV3L | 0.046185 | 0.408084 | Other Positive Correlation |
| DLAT | 0.046172 | 0.408216 | Other Positive Correlation |
| HACD2 | 0.046167 | 0.408265 | Other Positive Correlation |
| AK8 | 0.046142 | 0.408526 | Other Positive Correlation |
| SAP30BP | 0.046138 | 0.408566 | Other Positive Correlation |
| TENT4A | 0.046134 | 0.408602 | Other Positive Correlation |
| HLA-DRB1 | 0.046108 | 0.40887 | Other Positive Correlation |
| CEP85L | 0.046106 | 0.408883 | Other Positive Correlation |
| KCNJ3 | 0.046105 | 0.408893 | Other Positive Correlation |
| POLR3E | 0.046086 | 0.409089 | Other Positive Correlation |
| ZNF214 | 0.046078 | 0.40917 | Other Positive Correlation |
| ZSCAN23 | 0.046048 | 0.409474 | Other Positive Correlation |
| KRT32 | 0.046038 | 0.409575 | Other Positive Correlation |
| MYRFL | 0.046007 | 0.409891 | Other Positive Correlation |
| ULK4 | 0.046004 | 0.409927 | Other Positive Correlation |
| PGM3 | 0.046002 | 0.409943 | Other Positive Correlation |
| SPX | 0.046002 | 0.409949 | Other Positive Correlation |
| C20orf204 | 0.045967 | 0.410307 | Other Positive Correlation |
| FAM120C | 0.045958 | 0.410393 | Other Positive Correlation |
| FAM47E | 0.045956 | 0.41042 | Other Positive Correlation |
| TRAPPC11 | 0.045921 | 0.410772 | Other Positive Correlation |
| KCNF1 | 0.045878 | 0.41121 | Other Positive Correlation |
| GPR62 | 0.045867 | 0.411322 | Other Positive Correlation |
| IL10RA | 0.045853 | 0.411462 | Other Positive Correlation |
| CRYL1 | 0.045851 | 0.411487 | Other Positive Correlation |
| PDGFRL | 0.045851 | 0.411488 | Other Positive Correlation |
| PRPSAP1 | 0.045847 | 0.411524 | Other Positive Correlation |
| ZNF266 | 0.045839 | 0.411613 | Other Positive Correlation |
| SOWAHD | 0.045823 | 0.411768 | Other Positive Correlation |
| GEM | 0.045802 | 0.411987 | Other Positive Correlation |
| MAP3K11 | 0.045745 | 0.412566 | Other Positive Correlation |
| ZNF816-ZNF321P | 0.045741 | 0.412615 | Other Positive Correlation |
| SYT11 | 0.045737 | 0.412647 | Other Positive Correlation |
| MBIP | 0.045736 | 0.41266 | Other Positive Correlation |
| C2orf88 | 0.045703 | 0.413003 | Other Positive Correlation |
| IQCG | 0.045697 | 0.413063 | Other Positive Correlation |

| | | | |
|---|---|---|---|
| FAM9B | 0.045686 | 0.413178 | Other Positive Correlation |
| PKDREJ | 0.045686 | 0.413178 | Other Positive Correlation |
| BBS5 | 0.045677 | 0.413263 | Other Positive Correlation |
| SCUBE1 | 0.045674 | 0.4133 | Other Positive Correlation |
| ELP6 | 0.04565 | 0.413542 | Other Positive Correlation |
| MTM1 | 0.045597 | 0.414086 | Other Positive Correlation |
| TECPR1 | 0.045558 | 0.414488 | Other Positive Correlation |
| PRRG1 | 0.045556 | 0.414503 | Other Positive Correlation |
| AK7 | 0.045553 | 0.414538 | Other Positive Correlation |
| SVOPL | 0.045548 | 0.414588 | Other Positive Correlation |
| LY6L | 0.045539 | 0.414679 | Other Positive Correlation |
| SRGN | 0.04553 | 0.414776 | Other Positive Correlation |
| MIXL1 | 0.045528 | 0.414796 | Other Positive Correlation |
| COQ10B | 0.045427 | 0.41583 | Other Positive Correlation |
| NEUROG3 | 0.045407 | 0.416034 | Other Positive Correlation |
| SLC6A3 | 0.045407 | 0.416034 | Other Positive Correlation |
| DEFB126 | 0.045407 | 0.416034 | Other Positive Correlation |
| HNF4G | 0.045407 | 0.416034 | Other Positive Correlation |
| CCM2L | 0.045407 | 0.416034 | Other Positive Correlation |
| DNAJC6 | 0.045395 | 0.416157 | Other Positive Correlation |
| NSF | 0.045383 | 0.416279 | Other Positive Correlation |
| CCDC96 | 0.045376 | 0.416354 | Other Positive Correlation |
| ENSG00000285932 | 0.04532 | 0.416928 | Other Positive Correlation |
| NDST3 | 0.04532 | 0.416933 | Other Positive Correlation |
| GAS2 | 0.04531 | 0.417033 | Other Positive Correlation |
| PDX1 | 0.045286 | 0.417281 | Other Positive Correlation |
| ANO2 | 0.045282 | 0.417324 | Other Positive Correlation |
| CCDC181 | 0.045271 | 0.417429 | Other Positive Correlation |
| TMCC1 | 0.045257 | 0.417575 | Other Positive Correlation |
| MARF1 | 0.045256 | 0.417591 | Other Positive Correlation |
| MAPK15 | 0.045238 | 0.41777 | Other Positive Correlation |
| WDR17 | 0.045194 | 0.418232 | Other Positive Correlation |
| RMND5B | 0.045161 | 0.418563 | Other Positive Correlation |
| AKR1D1 | 0.04512 | 0.41899 | Other Positive Correlation |
| STRA8 | 0.04509 | 0.419301 | Other Positive Correlation |
| SENP1 | 0.045079 | 0.419418 | Other Positive Correlation |
| SLC6A8 | 0.045075 | 0.419456 | Other Positive Correlation |
| NIM1K | 0.045063 | 0.419576 | Other Positive Correlation |
| PPM1M | 0.045057 | 0.419644 | Other Positive Correlation |
| TAF3 | 0.045011 | 0.420115 | Other Positive Correlation |
| TRPM1 | 0.044984 | 0.420391 | Other Positive Correlation |
| NMB | 0.044957 | 0.420673 | Other Positive Correlation |
| PTER | 0.044954 | 0.420704 | Other Positive Correlation |
| GTF3C1 | 0.044926 | 0.420991 | Other Positive Correlation |
| HCRTR1 | 0.044862 | 0.421661 | Other Positive Correlation |
| CCDC91 | 0.044833 | 0.421959 | Other Positive Correlation |
| RBM24 | 0.04479 | 0.422408 | Other Positive Correlation |
| MYOM2 | 0.04474 | 0.422925 | Other Positive Correlation |

| | | | |
|---|---|---|---|
| WNK1 | 0.044722 | 0.423106 | Other Positive Correlation |
| KPNA6 | 0.04469 | 0.423445 | Other Positive Correlation |
| NPR3 | 0.044688 | 0.423468 | Other Positive Correlation |
| VPS37C | 0.044651 | 0.423842 | Other Positive Correlation |
| UBN2 | 0.044639 | 0.423971 | Other Positive Correlation |
| ABHD11 | 0.044636 | 0.424006 | Other Positive Correlation |
| ZNF600 | 0.044599 | 0.424383 | Other Positive Correlation |
| SOCS2 | 0.044587 | 0.424512 | Other Positive Correlation |
| RPRM | 0.044578 | 0.424607 | Other Positive Correlation |
| NPFFR2 | 0.044554 | 0.424854 | Other Positive Correlation |
| TEPSIN | 0.044508 | 0.425333 | Other Positive Correlation |
| DLEC1 | 0.044479 | 0.425637 | Other Positive Correlation |
| CPA5 | 0.04446 | 0.425839 | Other Positive Correlation |
| SFTPC | 0.044442 | 0.426023 | Other Positive Correlation |
| PPP1R3D | 0.04444 | 0.426042 | Other Positive Correlation |
| CXCL17 | 0.044433 | 0.426117 | Other Positive Correlation |
| TRPV4 | 0.044421 | 0.426245 | Other Positive Correlation |
| MRPS6 | 0.044419 | 0.426262 | Other Positive Correlation |
| ERN1 | 0.044395 | 0.426512 | Other Positive Correlation |
| CPXM1 | 0.044339 | 0.427098 | Other Positive Correlation |
| EPS8L1 | 0.044311 | 0.427388 | Other Positive Correlation |
| BMI1 | 0.044271 | 0.427806 | Other Positive Correlation |
| TMPRSS5 | 0.044237 | 0.428161 | Other Positive Correlation |
| TRIM48 | 0.044226 | 0.42828 | Other Positive Correlation |
| NAIF1 | 0.044223 | 0.428309 | Other Positive Correlation |
| CPZ | 0.04422 | 0.428344 | Other Positive Correlation |
| ENSG00000289720 | 0.044218 | 0.42836 | Other Positive Correlation |
| CRIP3 | 0.044203 | 0.428512 | Other Positive Correlation |
| SLC26A8 | 0.044188 | 0.428677 | Other Positive Correlation |
| MPV17L2 | 0.044178 | 0.428778 | Other Positive Correlation |
| ZNF382 | 0.044167 | 0.428892 | Other Positive Correlation |
| RANBP17 | 0.044151 | 0.429058 | Other Positive Correlation |
| RASL11B | 0.044109 | 0.429498 | Other Positive Correlation |
| NOMO2 | 0.044093 | 0.429671 | Other Positive Correlation |
| CAMTA2 | 0.04408 | 0.4298 | Other Positive Correlation |
| TEF | 0.044077 | 0.429836 | Other Positive Correlation |
| HAS1 | 0.044076 | 0.429848 | Other Positive Correlation |
| ATP1A2 | 0.044071 | 0.429903 | Other Positive Correlation |
| GOLGA2 | 0.044045 | 0.430173 | Other Positive Correlation |
| CFAP298-TCP10L | 0.044026 | 0.430365 | Other Positive Correlation |
| MTX2 | 0.044021 | 0.430423 | Other Positive Correlation |
| MATCAP2 | 0.043986 | 0.430785 | Other Positive Correlation |
| SLCO5A1 | 0.043981 | 0.430846 | Other Positive Correlation |
| USP6 | 0.04397 | 0.430954 | Other Positive Correlation |
| TMEM132C | 0.043952 | 0.431149 | Other Positive Correlation |
| PPOX | 0.043947 | 0.431202 | Other Positive Correlation |
| ERG | 0.043941 | 0.43126 | Other Positive Correlation |
| PIGU | 0.043902 | 0.431666 | Other Positive Correlation |

| | | | |
|---|---|---|---|
| DMXL2 | 0.043824 | 0.432487 | Other Positive Correlation |
| TSKS | 0.043814 | 0.43259 | Other Positive Correlation |
| FOXP2 | 0.043811 | 0.432625 | Other Positive Correlation |
| TOX | 0.043804 | 0.432701 | Other Positive Correlation |
| PHYHIP | 0.043801 | 0.432728 | Other Positive Correlation |
| SCNN1G | 0.043798 | 0.432765 | Other Positive Correlation |
| SCN4A | 0.043795 | 0.432794 | Other Positive Correlation |
| TOP3A | 0.043763 | 0.433134 | Other Positive Correlation |
| FOXP3 | 0.043737 | 0.433407 | Other Positive Correlation |
| PLXNA3 | 0.04372 | 0.433588 | Other Positive Correlation |
| SMAP2 | 0.043707 | 0.433726 | Other Positive Correlation |
| RIPK4 | 0.043692 | 0.433878 | Other Positive Correlation |
| ABCA7 | 0.043675 | 0.434055 | Other Positive Correlation |
| NECTIN2 | 0.04367 | 0.434111 | Other Positive Correlation |
| TRIM24 | 0.043668 | 0.434137 | Other Positive Correlation |
| EPM2A | 0.043664 | 0.434172 | Other Positive Correlation |
| MON1B | 0.043643 | 0.434395 | Other Positive Correlation |
| DICER1 | 0.043625 | 0.434588 | Other Positive Correlation |
| BCKDHA | 0.043605 | 0.434793 | Other Positive Correlation |
| TAL2 | 0.043602 | 0.434832 | Other Positive Correlation |
| GORASP1 | 0.043572 | 0.435143 | Other Positive Correlation |
| KSR1 | 0.043527 | 0.435621 | Other Positive Correlation |
| TCTE1 | 0.043507 | 0.435835 | Other Positive Correlation |
| PTCHD3 | 0.043496 | 0.435949 | Other Positive Correlation |
| COMMD9 | 0.043393 | 0.437037 | Other Positive Correlation |
| PPP1R2 | 0.043356 | 0.437426 | Other Positive Correlation |
| PHF7 | 0.04335 | 0.437494 | Other Positive Correlation |
| PRCP | 0.043325 | 0.437754 | Other Positive Correlation |
| ZZEF1 | 0.043267 | 0.438366 | Other Positive Correlation |
| AMER2 | 0.043225 | 0.438811 | Other Positive Correlation |
| HNF4A | 0.043212 | 0.438949 | Other Positive Correlation |
| ITGA9 | 0.043212 | 0.438949 | Other Positive Correlation |
| GATA1 | 0.043212 | 0.438949 | Other Positive Correlation |
| CACNG1 | 0.043212 | 0.438949 | Other Positive Correlation |
| OR2L2 | 0.043212 | 0.438949 | Other Positive Correlation |
| GJA4 | 0.043188 | 0.439208 | Other Positive Correlation |
| KRT33B | 0.043155 | 0.439552 | Other Positive Correlation |
| ACP5 | 0.043127 | 0.439848 | Other Positive Correlation |
| PCDHB14 | 0.043125 | 0.439871 | Other Positive Correlation |
| FSCN2 | 0.043088 | 0.440266 | Other Positive Correlation |
| SUPT6H | 0.043065 | 0.440513 | Other Positive Correlation |
| TSLP | 0.043064 | 0.44052 | Other Positive Correlation |
| YPEL4 | 0.043054 | 0.440627 | Other Positive Correlation |
| HIF3A | 0.043048 | 0.440697 | Other Positive Correlation |
| MMP16 | 0.043039 | 0.440787 | Other Positive Correlation |
| FOXD4L3 | 0.043021 | 0.440979 | Other Positive Correlation |
| HCST | 0.042984 | 0.441374 | Other Positive Correlation |
| C1GALT1C1L | 0.04292 | 0.442052 | Other Positive Correlation |

| | | | |
|---|---|---|---|
| ARMCX4 | 0.042917 | 0.442087 | Other Positive Correlation |
| ERBB2 | 0.042911 | 0.442148 | Other Positive Correlation |
| RAB14 | 0.042906 | 0.442207 | Other Positive Correlation |
| GTF2IRD2 | 0.042902 | 0.442245 | Other Positive Correlation |
| SLC25A13 | 0.042882 | 0.442455 | Other Positive Correlation |
| GPR12 | 0.042846 | 0.442846 | Other Positive Correlation |
| C1GALT1C1 | 0.042837 | 0.442932 | Other Positive Correlation |
| KAZN | 0.042802 | 0.443312 | Other Positive Correlation |
| CHST6 | 0.042796 | 0.443371 | Other Positive Correlation |
| MYL3 | 0.042782 | 0.443527 | Other Positive Correlation |
| NPC1 | 0.04278 | 0.443545 | Other Positive Correlation |
| ASS1 | 0.042745 | 0.443923 | Other Positive Correlation |
| R3HCC1L | 0.042727 | 0.44411 | Other Positive Correlation |
| SMARCD3 | 0.042694 | 0.444458 | Other Positive Correlation |
| GLS2 | 0.042691 | 0.444499 | Other Positive Correlation |
| ELAPOR2 | 0.042668 | 0.444743 | Other Positive Correlation |
| AXDND1 | 0.042667 | 0.444756 | Other Positive Correlation |
| RAD51B | 0.04265 | 0.444931 | Other Positive Correlation |
| FNDC11 | 0.04265 | 0.444932 | Other Positive Correlation |
| SLC22A14 | 0.042631 | 0.445135 | Other Positive Correlation |
| NSUN3 | 0.042626 | 0.445185 | Other Positive Correlation |
| CACNA2D3 | 0.042605 | 0.445414 | Other Positive Correlation |
| RELN | 0.042596 | 0.445511 | Other Positive Correlation |
| TSGA10 | 0.042592 | 0.44555 | Other Positive Correlation |
| RNF139 | 0.042587 | 0.44561 | Other Positive Correlation |
| S100A5 | 0.042573 | 0.445758 | Other Positive Correlation |
| VPS37A | 0.042573 | 0.445759 | Other Positive Correlation |
| ADAM21 | 0.042572 | 0.445768 | Other Positive Correlation |
| USH1C | 0.042568 | 0.445806 | Other Positive Correlation |
| RNASE4 | 0.042541 | 0.446101 | Other Positive Correlation |
| EXOSC4 | 0.042512 | 0.446411 | Other Positive Correlation |
| KLRG2 | 0.042509 | 0.446443 | Other Positive Correlation |
| CD34 | 0.04248 | 0.446751 | Other Positive Correlation |
| LSM14B | 0.042457 | 0.446995 | Other Positive Correlation |
| TBP | 0.042423 | 0.447357 | Other Positive Correlation |
| ASB2 | 0.042421 | 0.447388 | Other Positive Correlation |
| PODNL1 | 0.0424 | 0.447613 | Other Positive Correlation |
| TIGD6 | 0.042386 | 0.447759 | Other Positive Correlation |
| BRICD5 | 0.042385 | 0.447773 | Other Positive Correlation |
| GPX7 | 0.042355 | 0.448089 | Other Positive Correlation |
| GRAP | 0.042348 | 0.448163 | Other Positive Correlation |
| KCNAB3 | 0.042283 | 0.448863 | Other Positive Correlation |
| C9orf64 | 0.042282 | 0.448874 | Other Positive Correlation |
| RNF111 | 0.042184 | 0.449928 | Other Positive Correlation |
| NTRK2 | 0.042174 | 0.450034 | Other Positive Correlation |
| NAT9 | 0.042162 | 0.450161 | Other Positive Correlation |
| RIBC1 | 0.042161 | 0.450174 | Other Positive Correlation |
| EID2B | 0.042117 | 0.45065 | Other Positive Correlation |

| | | | |
|---|---|---|---|
| FHIP1B | 0.04211 | 0.450728 | Other Positive Correlation |
| TSC22D4 | 0.042109 | 0.450739 | Other Positive Correlation |
| SELENOH | 0.042106 | 0.450766 | Other Positive Correlation |
| PRKAG2 | 0.042098 | 0.450853 | Other Positive Correlation |
| ZCCHC2 | 0.042077 | 0.451084 | Other Positive Correlation |
| PDE4DIP | 0.042033 | 0.451553 | Other Positive Correlation |
| BTBD3 | 0.042028 | 0.451609 | Other Positive Correlation |
| IL16 | 0.041999 | 0.451918 | Other Positive Correlation |
| SKIDA1 | 0.041985 | 0.452072 | Other Positive Correlation |
| ADIPOR1 | 0.041967 | 0.452266 | Other Positive Correlation |
| NADSYN1 | 0.041957 | 0.452371 | Other Positive Correlation |
| SHROOM3 | 0.041936 | 0.452598 | Other Positive Correlation |
| ARHGAP39 | 0.041929 | 0.452674 | Other Positive Correlation |
| GIPR | 0.041915 | 0.452823 | Other Positive Correlation |
| PCDHB5 | 0.041904 | 0.452943 | Other Positive Correlation |
| SLC6A4 | 0.041852 | 0.453503 | Other Positive Correlation |
| IL37 | 0.041838 | 0.453657 | Other Positive Correlation |
| MYNN | 0.041829 | 0.453758 | Other Positive Correlation |
| DLGAP2 | 0.041828 | 0.453764 | Other Positive Correlation |
| TEKT4 | 0.041794 | 0.454136 | Other Positive Correlation |
| ADORA1 | 0.041768 | 0.454414 | Other Positive Correlation |
| RBM47 | 0.041762 | 0.454473 | Other Positive Correlation |
| SPRING1 | 0.041734 | 0.454785 | Other Positive Correlation |
| SLC9A8 | 0.041719 | 0.454946 | Other Positive Correlation |
| SCGB3A2 | 0.041718 | 0.454951 | Other Positive Correlation |
| CCDC134 | 0.041715 | 0.454983 | Other Positive Correlation |
| POLR3F | 0.041681 | 0.45535 | Other Positive Correlation |
| LUM | 0.041663 | 0.455553 | Other Positive Correlation |
| GRIP1 | 0.041662 | 0.455556 | Other Positive Correlation |
| ZNF592 | 0.041648 | 0.45571 | Other Positive Correlation |
| EEF1AKMT3 | 0.04162 | 0.456014 | Other Positive Correlation |
| NLRP2 | 0.041603 | 0.456196 | Other Positive Correlation |
| PSORS1C2 | 0.041545 | 0.456827 | Other Positive Correlation |
| FNBP1 | 0.041543 | 0.456846 | Other Positive Correlation |
| POM121 | 0.041535 | 0.456935 | Other Positive Correlation |
| ESRRG | 0.041534 | 0.456947 | Other Positive Correlation |
| ZNF350 | 0.041532 | 0.456969 | Other Positive Correlation |
| TST | 0.041531 | 0.456975 | Other Positive Correlation |
| YTHDF3 | 0.041524 | 0.457051 | Other Positive Correlation |
| LRP11 | 0.041511 | 0.457198 | Other Positive Correlation |
| KIAA1143 | 0.041477 | 0.457561 | Other Positive Correlation |
| LINGO3 | 0.041475 | 0.457582 | Other Positive Correlation |
| TULP2 | 0.041475 | 0.457582 | Other Positive Correlation |
| FOXB1 | 0.041475 | 0.457582 | Other Positive Correlation |
| AQP4 | 0.041475 | 0.457582 | Other Positive Correlation |
| RPS6KA5 | 0.041475 | 0.457592 | Other Positive Correlation |
| PLA2G2C | 0.041469 | 0.457647 | Other Positive Correlation |
| ARHGEF5 | 0.041466 | 0.457685 | Other Positive Correlation |

| | | | |
|---|---|---|---|
| SPI1 | 0.041453 | 0.457823 | Other Positive Correlation |
| SRGAP2B | 0.041439 | 0.457982 | Other Positive Correlation |
| DAB2IP | 0.04143 | 0.458074 | Other Positive Correlation |
| SEMA5B | 0.04143 | 0.458075 | Other Positive Correlation |
| LNP1 | 0.041372 | 0.4587 | Other Positive Correlation |
| RAB42 | 0.04133 | 0.459158 | Other Positive Correlation |
| WDR25 | 0.041326 | 0.459204 | Other Positive Correlation |
| POC5 | 0.041276 | 0.459754 | Other Positive Correlation |
| PCBD1 | 0.041256 | 0.459964 | Other Positive Correlation |
| RABIF | 0.041246 | 0.460075 | Other Positive Correlation |
| HOXC5 | 0.041244 | 0.460099 | Other Positive Correlation |
| ETV7 | 0.041229 | 0.460258 | Other Positive Correlation |
| SOCS4 | 0.04122 | 0.460358 | Other Positive Correlation |
| SPATA4 | 0.041217 | 0.460393 | Other Positive Correlation |
| DNER | 0.04121 | 0.460471 | Other Positive Correlation |
| SPRN | 0.041205 | 0.46052 | Other Positive Correlation |
| MFN1 | 0.041193 | 0.460648 | Other Positive Correlation |
| XKR6 | 0.04119 | 0.460689 | Other Positive Correlation |
| DVL2 | 0.041172 | 0.460887 | Other Positive Correlation |
| ANKRD33B | 0.041129 | 0.461354 | Other Positive Correlation |
| MYO7A | 0.041114 | 0.461515 | Other Positive Correlation |
| TENM3 | 0.041111 | 0.461547 | Other Positive Correlation |
| CFDP1 | 0.041109 | 0.461572 | Other Positive Correlation |
| KTI12 | 0.041102 | 0.461647 | Other Positive Correlation |
| SPOCK3 | 0.041102 | 0.461648 | Other Positive Correlation |
| ZNF91 | 0.041058 | 0.462125 | Other Positive Correlation |
| TBX3 | 0.041058 | 0.462127 | Other Positive Correlation |
| NKX2-8 | 0.04105 | 0.462209 | Other Positive Correlation |
| PCSK4 | 0.04105 | 0.462211 | Other Positive Correlation |
| BUD23 | 0.04104 | 0.462325 | Other Positive Correlation |
| ENSG00000282936 | 0.040987 | 0.462905 | Other Positive Correlation |
| DKK4 | 0.040931 | 0.463511 | Other Positive Correlation |
| NOTCH2NLB | 0.040923 | 0.463596 | Other Positive Correlation |
| CHRFAM7A | 0.04092 | 0.463636 | Other Positive Correlation |
| SNX25 | 0.040902 | 0.463834 | Other Positive Correlation |
| GMNC | 0.0409 | 0.463851 | Other Positive Correlation |
| PCM1 | 0.040878 | 0.464088 | Other Positive Correlation |
| ZFP28 | 0.040872 | 0.464156 | Other Positive Correlation |
| EOMES | 0.040868 | 0.464206 | Other Positive Correlation |
| HSD3B7 | 0.040853 | 0.464368 | Other Positive Correlation |
| FZD1 | 0.040844 | 0.464465 | Other Positive Correlation |
| ANKRD62 | 0.040827 | 0.464651 | Other Positive Correlation |
| CDHR1 | 0.040803 | 0.464914 | Other Positive Correlation |
| ZNF184 | 0.040779 | 0.465178 | Other Positive Correlation |
| GRM5 | 0.040777 | 0.465193 | Other Positive Correlation |
| LRRN4 | 0.04077 | 0.465278 | Other Positive Correlation |
| KIF13B | 0.040767 | 0.465305 | Other Positive Correlation |
| RAB44 | 0.040765 | 0.465334 | Other Positive Correlation |

| | | | |
|---|---|---|---|
| HIP1R | 0.040763 | 0.465353 | Other Positive Correlation |
| TXLNB | 0.040754 | 0.465447 | Other Positive Correlation |
| RASIP1 | 0.040744 | 0.465558 | Other Positive Correlation |
| CRISP2 | 0.0407 | 0.466039 | Other Positive Correlation |
| CNOT11 | 0.040698 | 0.466067 | Other Positive Correlation |
| SUSD1 | 0.040695 | 0.466094 | Other Positive Correlation |
| GNRHR | 0.040655 | 0.466535 | Other Positive Correlation |
| OR1F1 | 0.040609 | 0.467045 | Other Positive Correlation |
| FBN1 | 0.040585 | 0.467309 | Other Positive Correlation |
| CLSTN3 | 0.040581 | 0.467343 | Other Positive Correlation |
| RNF121 | 0.040563 | 0.467546 | Other Positive Correlation |
| CORO2B | 0.040518 | 0.46804 | Other Positive Correlation |
| FAAP100 | 0.0405 | 0.468241 | Other Positive Correlation |
| B3GNT3 | 0.040494 | 0.468304 | Other Positive Correlation |
| DENND4B | 0.040475 | 0.468512 | Other Positive Correlation |
| PSEN2 | 0.040466 | 0.46861 | Other Positive Correlation |
| DCAF1 | 0.040455 | 0.468727 | Other Positive Correlation |
| KDM3A | 0.04041 | 0.469228 | Other Positive Correlation |
| SLC12A2 | 0.040393 | 0.469413 | Other Positive Correlation |
| ZNF471 | 0.040391 | 0.469438 | Other Positive Correlation |
| STEAP2 | 0.040375 | 0.469616 | Other Positive Correlation |
| PLD5P1 | 0.040359 | 0.469792 | Other Positive Correlation |
| C8G | 0.040351 | 0.46988 | Other Positive Correlation |
| TCERG1L | 0.040348 | 0.469909 | Other Positive Correlation |
| BSN | 0.040326 | 0.470153 | Other Positive Correlation |
| INTS15 | 0.040234 | 0.471172 | Other Positive Correlation |
| BTG1 | 0.040232 | 0.471187 | Other Positive Correlation |
| ENSG00000284956 | 0.040223 | 0.471289 | Other Positive Correlation |
| ZDHHC14 | 0.040222 | 0.471295 | Other Positive Correlation |
| ZER1 | 0.040219 | 0.471336 | Other Positive Correlation |
| ASNSD1 | 0.040218 | 0.471341 | Other Positive Correlation |
| SLC48A1 | 0.04019 | 0.471657 | Other Positive Correlation |
| PRXL2B | 0.040165 | 0.471929 | Other Positive Correlation |
| JMJD7-PLA2G4B | 0.040087 | 0.472791 | Other Positive Correlation |
| NPR2 | 0.040037 | 0.473344 | Other Positive Correlation |
| FAM110A | 0.040035 | 0.473361 | Other Positive Correlation |
| HECTD4 | 0.040033 | 0.473386 | Other Positive Correlation |
| HOMEZ | 0.040019 | 0.473544 | Other Positive Correlation |
| ENSG00000272104 | 0.039989 | 0.473872 | Other Positive Correlation |
| ZNF853 | 0.039986 | 0.473904 | Other Positive Correlation |
| BAIAP2L2 | 0.039978 | 0.473994 | Other Positive Correlation |
| AGAP2 | 0.039966 | 0.474126 | Other Positive Correlation |
| MRFAP1 | 0.039951 | 0.474297 | Other Positive Correlation |
| C17orf50 | 0.039938 | 0.474441 | Other Positive Correlation |
| SLCO4C1 | 0.039935 | 0.47447 | Other Positive Correlation |
| UBE3D | 0.039918 | 0.474657 | Other Positive Correlation |
| KLF14 | 0.039887 | 0.475 | Other Positive Correlation |
| PIP5K1A | 0.03986 | 0.475307 | Other Positive Correlation |

| | | | |
|---|---|---|---|
| MGST3 | 0.039804 | 0.475926 | Other Positive Correlation |
| LY75 | 0.039773 | 0.476271 | Other Positive Correlation |
| C7orf57 | 0.039743 | 0.476603 | Other Positive Correlation |
| ZNF254 | 0.039733 | 0.476712 | Other Positive Correlation |
| OGFOD1 | 0.039711 | 0.476953 | Other Positive Correlation |
| MRTFA | 0.039696 | 0.477123 | Other Positive Correlation |
| TM4SF19-DYNLT2B | 0.039695 | 0.477131 | Other Positive Correlation |
| FBXW4 | 0.039687 | 0.477225 | Other Positive Correlation |
| ZNF260 | 0.039682 | 0.477275 | Other Positive Correlation |
| DACH1 | 0.039583 | 0.478377 | Other Positive Correlation |
| WDR97 | 0.039579 | 0.478418 | Other Positive Correlation |
| CD248 | 0.039572 | 0.478498 | Other Positive Correlation |
| TRIM49 | 0.039569 | 0.478535 | Other Positive Correlation |
| NDUFA8 | 0.039555 | 0.47869 | Other Positive Correlation |
| WNK3 | 0.039554 | 0.478698 | Other Positive Correlation |
| TRAPPC5 | 0.039544 | 0.478809 | Other Positive Correlation |
| RNF14 | 0.039525 | 0.479018 | Other Positive Correlation |
| IP6K3 | 0.039525 | 0.479025 | Other Positive Correlation |
| PGAM4 | 0.039511 | 0.479178 | Other Positive Correlation |
| ATF6B | 0.039503 | 0.479267 | Other Positive Correlation |
| SCNN1D | 0.039432 | 0.480061 | Other Positive Correlation |
| TRIM36 | 0.039426 | 0.480128 | Other Positive Correlation |
| KCNG1 | 0.039423 | 0.480162 | Other Positive Correlation |
| MYL2 | 0.039409 | 0.48032 | Other Positive Correlation |
| WDR7 | 0.039404 | 0.480369 | Other Positive Correlation |
| PAQR4 | 0.039388 | 0.480551 | Other Positive Correlation |
| MYT1 | 0.039357 | 0.480894 | Other Positive Correlation |
| MATN1 | 0.039353 | 0.480938 | Other Positive Correlation |
| NHLRC2 | 0.039351 | 0.480966 | Other Positive Correlation |
| MUC5B | 0.039314 | 0.48138 | Other Positive Correlation |
| ADAMTS13 | 0.039304 | 0.481483 | Other Positive Correlation |
| RNF20 | 0.039273 | 0.481831 | Other Positive Correlation |
| CYP7B1 | 0.039271 | 0.48186 | Other Positive Correlation |
| RIIAD1 | 0.039259 | 0.481995 | Other Positive Correlation |
| ARL6 | 0.03925 | 0.482095 | Other Positive Correlation |
| SLC10A3 | 0.03923 | 0.48231 | Other Positive Correlation |
| C2orf76 | 0.039217 | 0.482459 | Other Positive Correlation |
| ZNF569 | 0.039177 | 0.482902 | Other Positive Correlation |
| VTI1A | 0.039177 | 0.482909 | Other Positive Correlation |
| LDLRAD3 | 0.039157 | 0.483133 | Other Positive Correlation |
| CACNB3 | 0.039119 | 0.483554 | Other Positive Correlation |
| RDH14 | 0.039056 | 0.484255 | Other Positive Correlation |
| BTBD17 | 0.039048 | 0.48435 | Other Positive Correlation |
| APOL4 | 0.038999 | 0.484901 | Other Positive Correlation |
| ASPRV1 | 0.038968 | 0.48525 | Other Positive Correlation |
| PDE8A | 0.038967 | 0.485262 | Other Positive Correlation |
| SENP7 | 0.038966 | 0.485271 | Other Positive Correlation |
| ZNF578 | 0.038929 | 0.485681 | Other Positive Correlation |

| | | | |
|---|---|---|---|
| RIMBP3C | 0.038925 | 0.485724 | Other Positive Correlation |
| LEMD1 | 0.038918 | 0.485803 | Other Positive Correlation |
| TRIM56 | 0.038915 | 0.485835 | Other Positive Correlation |
| DNAAF10 | 0.038823 | 0.486876 | Other Positive Correlation |
| FAM89B | 0.038816 | 0.486952 | Other Positive Correlation |
| TIAM2 | 0.038784 | 0.487312 | Other Positive Correlation |
| ZNF609 | 0.038778 | 0.487382 | Other Positive Correlation |
| SLC28A3 | 0.038742 | 0.487778 | Other Positive Correlation |
| ZNF793 | 0.038735 | 0.487865 | Other Positive Correlation |
| IZUMO4 | 0.038711 | 0.488132 | Other Positive Correlation |
| PIPOX | 0.038702 | 0.488227 | Other Positive Correlation |
| SENP6 | 0.038687 | 0.488399 | Other Positive Correlation |
| TIGD1 | 0.038669 | 0.488604 | Other Positive Correlation |
| STX16-NPEPL1 | 0.038598 | 0.489407 | Other Positive Correlation |
| CYBRD1 | 0.038598 | 0.489408 | Other Positive Correlation |
| NLRP9 | 0.038588 | 0.489517 | Other Positive Correlation |
| CSRNP2 | 0.038569 | 0.489731 | Other Positive Correlation |
| KCNB2 | 0.038565 | 0.489773 | Other Positive Correlation |
| PEX14 | 0.038541 | 0.490042 | Other Positive Correlation |
| ZNF541 | 0.038535 | 0.490108 | Other Positive Correlation |
| IZUMO2 | 0.038533 | 0.490131 | Other Positive Correlation |
| CKMT2 | 0.038514 | 0.490348 | Other Positive Correlation |
| HPDL | 0.038508 | 0.490415 | Other Positive Correlation |
| TMEM217 | 0.038493 | 0.490586 | Other Positive Correlation |
| CPNE7 | 0.038483 | 0.490703 | Other Positive Correlation |
| GLOD4 | 0.038449 | 0.491082 | Other Positive Correlation |
| CLIC2 | 0.038448 | 0.491089 | Other Positive Correlation |
| SNX27 | 0.038448 | 0.491091 | Other Positive Correlation |
| ZNF792 | 0.038437 | 0.491215 | Other Positive Correlation |
| UPK1A | 0.038396 | 0.491684 | Other Positive Correlation |
| TGM5 | 0.038382 | 0.491836 | Other Positive Correlation |
| GRK3 | 0.038368 | 0.492 | Other Positive Correlation |
| SH2D6 | 0.038329 | 0.492432 | Other Positive Correlation |
| SDC3 | 0.038321 | 0.492532 | Other Positive Correlation |
| SAMD4A | 0.038317 | 0.492577 | Other Positive Correlation |
| KLF11 | 0.038311 | 0.492642 | Other Positive Correlation |
| FUT2 | 0.038284 | 0.49294 | Other Positive Correlation |
| SLC38A7 | 0.038228 | 0.493577 | Other Positive Correlation |
| CNKSR3 | 0.038223 | 0.493639 | Other Positive Correlation |
| SPRR2D | 0.038206 | 0.493831 | Other Positive Correlation |
| DEDD | 0.038204 | 0.493853 | Other Positive Correlation |
| FNIP1 | 0.03818 | 0.494125 | Other Positive Correlation |
| MAP3K10 | 0.038154 | 0.49442 | Other Positive Correlation |
| ARCN1 | 0.038112 | 0.494891 | Other Positive Correlation |
| ZNF221 | 0.0381 | 0.495024 | Other Positive Correlation |
| EDARADD | 0.038098 | 0.495048 | Other Positive Correlation |
| PGBD1 | 0.038095 | 0.495087 | Other Positive Correlation |
| SCUBE3 | 0.038074 | 0.49532 | Other Positive Correlation |

| | | | |
|---|---|---|---|
| FOXD3 | 0.038046 | 0.495639 | Other Positive Correlation |
| TCEAL3 | 0.038025 | 0.495872 | Other Positive Correlation |
| ZBTB7A | 0.038019 | 0.495949 | Other Positive Correlation |
| SSUH2 | 0.038012 | 0.496026 | Other Positive Correlation |
| RHOBTB1 | 0.037994 | 0.496229 | Other Positive Correlation |
| FAM110D | 0.037988 | 0.496296 | Other Positive Correlation |
| RFXAP | 0.037988 | 0.496297 | Other Positive Correlation |
| CHIC2 | 0.037967 | 0.496533 | Other Positive Correlation |
| LRRC24 | 0.037957 | 0.496644 | Other Positive Correlation |
| PRDM8 | 0.037955 | 0.496677 | Other Positive Correlation |
| P2RY6 | 0.03795 | 0.496723 | Other Positive Correlation |
| MTMR8 | 0.037892 | 0.497389 | Other Positive Correlation |
| TM9SF1 | 0.037891 | 0.497397 | Other Positive Correlation |
| SH2B3 | 0.037888 | 0.497437 | Other Positive Correlation |
| SLC39A11 | 0.037888 | 0.497437 | Other Positive Correlation |
| ARMCX5 | 0.037872 | 0.49761 | Other Positive Correlation |
| CHODL | 0.03787 | 0.497634 | Other Positive Correlation |
| LRRC26 | 0.037863 | 0.497715 | Other Positive Correlation |
| ATL2 | 0.037856 | 0.497802 | Other Positive Correlation |
| TSPYL4 | 0.037855 | 0.497805 | Other Positive Correlation |
| PEX3 | 0.037829 | 0.498102 | Other Positive Correlation |
| MAML1 | 0.037811 | 0.498307 | Other Positive Correlation |
| FRMD5 | 0.037768 | 0.498796 | Other Positive Correlation |
| DMRTA2 | 0.037701 | 0.499557 | Other Positive Correlation |
| MORN3 | 0.037689 | 0.499693 | Other Positive Correlation |
| ZSWIM3 | 0.037687 | 0.499714 | Other Positive Correlation |
| BTN2A2 | 0.037673 | 0.499883 | Other Positive Correlation |
| CCDC171 | 0.037665 | 0.499964 | Other Positive Correlation |
| NAPEPLD | 0.037623 | 0.500448 | Other Positive Correlation |
| OR52A1 | 0.037582 | 0.500917 | Other Positive Correlation |
| SLC35A5 | 0.037581 | 0.500932 | Other Positive Correlation |
| SFTPD | 0.037577 | 0.500972 | Other Positive Correlation |
| FAM166C | 0.037558 | 0.501189 | Other Positive Correlation |
| PPIC | 0.03755 | 0.501275 | Other Positive Correlation |
| C5orf22 | 0.03755 | 0.501277 | Other Positive Correlation |
| SLFN11 | 0.037532 | 0.501482 | Other Positive Correlation |
| PRRG2 | 0.037527 | 0.501547 | Other Positive Correlation |
| ALOX15 | 0.037525 | 0.501564 | Other Positive Correlation |
| CSDC2 | 0.037502 | 0.501833 | Other Positive Correlation |
| CA10 | 0.037493 | 0.501933 | Other Positive Correlation |
| SOHLH2 | 0.037482 | 0.50206 | Other Positive Correlation |
| PPP4R3B | 0.037379 | 0.503229 | Other Positive Correlation |
| EHMT2 | 0.037373 | 0.503296 | Other Positive Correlation |
| SLC7A6OS | 0.037348 | 0.503592 | Other Positive Correlation |
| CHGA | 0.037332 | 0.503764 | Other Positive Correlation |
| C1orf232 | 0.037324 | 0.503867 | Other Positive Correlation |
| PCDHB10 | 0.037311 | 0.504014 | Other Positive Correlation |
| ANGEL1 | 0.037307 | 0.504053 | Other Positive Correlation |

| | | | |
|---|---|---|---|
| TAF11L12 | 0.037285 | 0.504311 | Other Positive Correlation |
| FGF20 | 0.037282 | 0.504344 | Other Positive Correlation |
| ZNF26 | 0.037258 | 0.50462 | Other Positive Correlation |
| MTPAP | 0.037242 | 0.504801 | Other Positive Correlation |
| INPP1 | 0.037236 | 0.50487 | Other Positive Correlation |
| SHF | 0.037183 | 0.505476 | Other Positive Correlation |
| REXO1 | 0.037174 | 0.505581 | Other Positive Correlation |
| AP3B2 | 0.03716 | 0.505734 | Other Positive Correlation |
| AKR1E2 | 0.037153 | 0.505818 | Other Positive Correlation |
| MCFD2 | 0.03713 | 0.506081 | Other Positive Correlation |
| C1orf54 | 0.037076 | 0.506696 | Other Positive Correlation |
| ZNF671 | 0.037028 | 0.507253 | Other Positive Correlation |
| SLC2A12 | 0.037015 | 0.5074 | Other Positive Correlation |
| CLOCK | 0.036987 | 0.507723 | Other Positive Correlation |
| MFSD4A | 0.03693 | 0.508382 | Other Positive Correlation |
| IFRD1 | 0.03691 | 0.508605 | Other Positive Correlation |
| ZYG11B | 0.036904 | 0.508679 | Other Positive Correlation |
| FLT1 | 0.036886 | 0.508878 | Other Positive Correlation |
| ADAMTS19 | 0.036881 | 0.50894 | Other Positive Correlation |
| C5orf46 | 0.036832 | 0.509499 | Other Positive Correlation |
| CNTROB | 0.036822 | 0.509619 | Other Positive Correlation |
| CAPN6 | 0.036813 | 0.509719 | Other Positive Correlation |
| PPM1D | 0.036812 | 0.509733 | Other Positive Correlation |
| CRIPT | 0.036801 | 0.509856 | Other Positive Correlation |
| TSG101 | 0.036775 | 0.510155 | Other Positive Correlation |
| MYPOP | 0.036766 | 0.51026 | Other Positive Correlation |
| LIN37 | 0.036756 | 0.51038 | Other Positive Correlation |
| GBA2 | 0.036724 | 0.510745 | Other Positive Correlation |
| METTL13 | 0.036698 | 0.51105 | Other Positive Correlation |
| AQP7 | 0.036692 | 0.51111 | Other Positive Correlation |
| SERINC4 | 0.036676 | 0.511299 | Other Positive Correlation |
| OR2B6 | 0.036667 | 0.511408 | Other Positive Correlation |
| MSMB | 0.036609 | 0.512075 | Other Positive Correlation |
| FARS2 | 0.036573 | 0.512483 | Other Positive Correlation |
| GCNT3 | 0.036562 | 0.51261 | Other Positive Correlation |
| GOLM1 | 0.036556 | 0.512685 | Other Positive Correlation |
| RGS7 | 0.036553 | 0.512723 | Other Positive Correlation |
| CDKN2AIPNL | 0.036548 | 0.512772 | Other Positive Correlation |
| MFSD8 | 0.036516 | 0.513149 | Other Positive Correlation |
| WAS | 0.036511 | 0.513202 | Other Positive Correlation |
| ZNF229 | 0.036509 | 0.513224 | Other Positive Correlation |
| SLCO2A1 | 0.036503 | 0.513294 | Other Positive Correlation |
| NLGN2 | 0.036495 | 0.513388 | Other Positive Correlation |
| LYPD5 | 0.036484 | 0.513518 | Other Positive Correlation |
| TSNAXIP1 | 0.036396 | 0.514528 | Other Positive Correlation |
| ZNF131 | 0.036382 | 0.51469 | Other Positive Correlation |
| VWA1 | 0.036376 | 0.514761 | Other Positive Correlation |
| ZNF616 | 0.036372 | 0.514808 | Other Positive Correlation |

| | | | |
|---|---|---|---|
| RHOV | 0.036334 | 0.515248 | Other Positive Correlation |
| TRAPPC6B | 0.036329 | 0.515304 | Other Positive Correlation |
| ADAT1 | 0.036321 | 0.515397 | Other Positive Correlation |
| SLC26A2 | 0.03632 | 0.515415 | Other Positive Correlation |
| HMGB1P1 | 0.036292 | 0.515736 | Other Positive Correlation |
| ALKBH3 | 0.036267 | 0.516021 | Other Positive Correlation |
| SLC36A1 | 0.036266 | 0.516042 | Other Positive Correlation |
| AUTS2 | 0.03624 | 0.516339 | Other Positive Correlation |
| ESR1 | 0.036232 | 0.516434 | Other Positive Correlation |
| MCOLN2 | 0.036231 | 0.516445 | Other Positive Correlation |
| LMO3 | 0.036222 | 0.516551 | Other Positive Correlation |
| CEP41 | 0.036196 | 0.516848 | Other Positive Correlation |
| CCDC87 | 0.036166 | 0.517195 | Other Positive Correlation |
| PNPLA8 | 0.036143 | 0.517464 | Other Positive Correlation |
| APBA2 | 0.036134 | 0.517567 | Other Positive Correlation |
| HM13 | 0.036134 | 0.51757 | Other Positive Correlation |
| CARS1 | 0.036119 | 0.517745 | Other Positive Correlation |
| BTAF1 | 0.036115 | 0.51779 | Other Positive Correlation |
| ENSG00000285000 | 0.036102 | 0.51794 | Other Positive Correlation |
| PROZ | 0.036102 | 0.51794 | Other Positive Correlation |
| RLBP1 | 0.036099 | 0.517968 | Other Positive Correlation |
| SYCP1 | 0.036098 | 0.517985 | Other Positive Correlation |
| RET | 0.036034 | 0.518726 | Other Positive Correlation |
| GNB5 | 0.035992 | 0.519219 | Other Positive Correlation |
| H2AC8 | 0.03596 | 0.51959 | Other Positive Correlation |
| TRNT1 | 0.035942 | 0.519791 | Other Positive Correlation |
| ZNF664 | 0.035942 | 0.519795 | Other Positive Correlation |
| AARS2 | 0.03588 | 0.520517 | Other Positive Correlation |
| CEP76 | 0.035867 | 0.520665 | Other Positive Correlation |
| COLEC10 | 0.035845 | 0.520926 | Other Positive Correlation |
| GJA5 | 0.035829 | 0.521104 | Other Positive Correlation |
| NIF3L1 | 0.035788 | 0.521592 | Other Positive Correlation |
| NEDD1 | 0.035772 | 0.521778 | Other Positive Correlation |
| SV2C | 0.035749 | 0.52204 | Other Positive Correlation |
| C5orf49 | 0.035742 | 0.522126 | Other Positive Correlation |
| RBBP8NL | 0.035734 | 0.522219 | Other Positive Correlation |
| TBL1X | 0.035724 | 0.522329 | Other Positive Correlation |
| NFKBID | 0.035695 | 0.522667 | Other Positive Correlation |
| XPO6 | 0.035668 | 0.522984 | Other Positive Correlation |
| CFAP65 | 0.035661 | 0.523064 | Other Positive Correlation |
| TUBB8B | 0.035639 | 0.523319 | Other Positive Correlation |
| SCNN1B | 0.035612 | 0.523637 | Other Positive Correlation |
| NRN1 | 0.035592 | 0.523867 | Other Positive Correlation |
| SLC11A2 | 0.035579 | 0.524018 | Other Positive Correlation |
| DNAJC27 | 0.035555 | 0.524305 | Other Positive Correlation |
| ROPN1L | 0.035544 | 0.52443 | Other Positive Correlation |
| FBXO24 | 0.035522 | 0.52469 | Other Positive Correlation |
| CBLL1 | 0.035517 | 0.524751 | Other Positive Correlation |

| | | | |
|---|---|---|---|
| ZNF317 | 0.035442 | 0.525628 | Other Positive Correlation |
| CNKSR2 | 0.035433 | 0.525723 | Other Positive Correlation |
| ARMH3 | 0.035433 | 0.525728 | Other Positive Correlation |
| TLR5 | 0.035404 | 0.526066 | Other Positive Correlation |
| PIM3 | 0.035402 | 0.526094 | Other Positive Correlation |
| AP5M1 | 0.035381 | 0.526333 | Other Positive Correlation |
| P3H1 | 0.035381 | 0.526335 | Other Positive Correlation |
| FAM222B | 0.035374 | 0.526424 | Other Positive Correlation |
| EFCAB5 | 0.035371 | 0.526449 | Other Positive Correlation |
| HTRA2 | 0.035317 | 0.527083 | Other Positive Correlation |
| MSANTD1 | 0.035313 | 0.527134 | Other Positive Correlation |
| ENSG00000256029 | 0.035303 | 0.527245 | Other Positive Correlation |
| ANKRD6 | 0.035229 | 0.528119 | Other Positive Correlation |
| ANKHD1 | 0.03506 | 0.530101 | Other Positive Correlation |
| PAFAH2 | 0.035058 | 0.530127 | Other Positive Correlation |
| GMIP | 0.03504 | 0.53033 | Other Positive Correlation |
| EGLN1 | 0.035003 | 0.530772 | Other Positive Correlation |
| STBD1 | 0.034999 | 0.530819 | Other Positive Correlation |
| ZNF607 | 0.034985 | 0.530978 | Other Positive Correlation |
| RTN4IP1 | 0.034976 | 0.531085 | Other Positive Correlation |
| SLX1A | 0.034942 | 0.531479 | Other Positive Correlation |
| TRAPPC10 | 0.0349 | 0.531975 | Other Positive Correlation |
| LYPD3 | 0.034891 | 0.532089 | Other Positive Correlation |
| CIITA | 0.034878 | 0.532234 | Other Positive Correlation |
| HCN3 | 0.034834 | 0.532755 | Other Positive Correlation |
| TMPRSS6 | 0.034833 | 0.532767 | Other Positive Correlation |
| PTCD3 | 0.03481 | 0.53304 | Other Positive Correlation |
| SLC49A4 | 0.034797 | 0.533193 | Other Positive Correlation |
| IL11RA | 0.034792 | 0.533247 | Other Positive Correlation |
| PRDM15 | 0.03469 | 0.534457 | Other Positive Correlation |
| NT5C1A | 0.034684 | 0.534518 | Other Positive Correlation |
| SMPDL3A | 0.03461 | 0.535392 | Other Positive Correlation |
| ELOC | 0.034609 | 0.535405 | Other Positive Correlation |
| ZBTB8OS | 0.034595 | 0.53557 | Other Positive Correlation |
| LTN1 | 0.034586 | 0.535681 | Other Positive Correlation |
| ATXN7L3B | 0.034564 | 0.535939 | Other Positive Correlation |
| CLPSL1 | 0.034534 | 0.536295 | Other Positive Correlation |
| TRIM17 | 0.034509 | 0.536582 | Other Positive Correlation |
| ADCY3 | 0.034505 | 0.536633 | Other Positive Correlation |
| KANSL2 | 0.034491 | 0.536801 | Other Positive Correlation |
| ZNF567 | 0.034474 | 0.537005 | Other Positive Correlation |
| PCGF6 | 0.034455 | 0.537222 | Other Positive Correlation |
| UFSP1 | 0.034429 | 0.53753 | Other Positive Correlation |
| NAP1L2 | 0.034427 | 0.537555 | Other Positive Correlation |
| KRT34 | 0.034426 | 0.537564 | Other Positive Correlation |
| EXD1 | 0.034415 | 0.537694 | Other Positive Correlation |
| GTF2IRD1 | 0.034403 | 0.537844 | Other Positive Correlation |
| SRP68 | 0.034398 | 0.537902 | Other Positive Correlation |

| | | | |
|---|---|---|---|
| TRIM54 | 0.034391 | 0.53798 | Other Positive Correlation |
| ZNF74 | 0.034376 | 0.53816 | Other Positive Correlation |
| NAP1L5 | 0.034373 | 0.538197 | Other Positive Correlation |
| CIB3 | 0.034366 | 0.538278 | Other Positive Correlation |
| CGREF1 | 0.034351 | 0.538461 | Other Positive Correlation |
| SLC8B1 | 0.034337 | 0.538621 | Other Positive Correlation |
| ARSH | 0.034326 | 0.538751 | Other Positive Correlation |
| PDGFB | 0.034234 | 0.539845 | Other Positive Correlation |
| ZBTB2 | 0.034228 | 0.539913 | Other Positive Correlation |
| ROBO2 | 0.034218 | 0.540025 | Other Positive Correlation |
| PBX1 | 0.034206 | 0.540175 | Other Positive Correlation |
| ZNF527 | 0.034179 | 0.540491 | Other Positive Correlation |
| CDH16 | 0.03415 | 0.540839 | Other Positive Correlation |
| DGCR2 | 0.034141 | 0.540944 | Other Positive Correlation |
| RNF224 | 0.034119 | 0.5412 | Other Positive Correlation |
| PANK3 | 0.034108 | 0.541335 | Other Positive Correlation |
| ANO9 | 0.034089 | 0.541562 | Other Positive Correlation |
| ENSG00000283128 | 0.034059 | 0.541915 | Other Positive Correlation |
| MYO1F | 0.03405 | 0.542017 | Other Positive Correlation |
| CEP57 | 0.034048 | 0.542045 | Other Positive Correlation |
| CAPN14 | 0.034046 | 0.542073 | Other Positive Correlation |
| PPM1A | 0.033995 | 0.54268 | Other Positive Correlation |
| TMCC3 | 0.033983 | 0.542814 | Other Positive Correlation |
| RESF1 | 0.033979 | 0.54287 | Other Positive Correlation |
| SLC45A1 | 0.033946 | 0.543261 | Other Positive Correlation |
| SGSM1 | 0.033942 | 0.543307 | Other Positive Correlation |
| CAPN8 | 0.03394 | 0.543329 | Other Positive Correlation |
| HSD11B1 | 0.033918 | 0.543595 | Other Positive Correlation |
| ZNF675 | 0.033863 | 0.54425 | Other Positive Correlation |
| SYN3 | 0.033858 | 0.544309 | Other Positive Correlation |
| SYNDIG1L | 0.033839 | 0.544529 | Other Positive Correlation |
| APPBP2 | 0.033839 | 0.544531 | Other Positive Correlation |
| NAALADL2 | 0.033838 | 0.544538 | Other Positive Correlation |
| RAB36 | 0.033805 | 0.544936 | Other Positive Correlation |
| UBE4A | 0.0338 | 0.544996 | Other Positive Correlation |
| SLC25A45 | 0.033797 | 0.545032 | Other Positive Correlation |
| MEDAG | 0.033796 | 0.545038 | Other Positive Correlation |
| MAGEE1 | 0.033795 | 0.545057 | Other Positive Correlation |
| H3Y1 | 0.033769 | 0.545363 | Other Positive Correlation |
| ZNF624 | 0.033763 | 0.54543 | Other Positive Correlation |
| RAPGEF2 | 0.033763 | 0.545436 | Other Positive Correlation |
| MROH8 | 0.03374 | 0.545714 | Other Positive Correlation |
| TRIM55 | 0.03373 | 0.545824 | Other Positive Correlation |
| TBX19 | 0.03373 | 0.545829 | Other Positive Correlation |
| GPER1 | 0.03372 | 0.545952 | Other Positive Correlation |
| ABHD17B | 0.03368 | 0.546421 | Other Positive Correlation |
| MYCN | 0.033678 | 0.546452 | Other Positive Correlation |
| TMED8 | 0.033647 | 0.546817 | Other Positive Correlation |

| | | | |
|---|---|---|---|
| KCNQ1 | 0.033634 | 0.546974 | Other Positive Correlation |
| SLC22A1 | 0.033573 | 0.547702 | Other Positive Correlation |
| UNC13B | 0.033542 | 0.548071 | Other Positive Correlation |
| CDH5 | 0.033527 | 0.548254 | Other Positive Correlation |
| ZNF653 | 0.033524 | 0.548282 | Other Positive Correlation |
| ADCY8 | 0.033492 | 0.548666 | Other Positive Correlation |
| PRSS27 | 0.033454 | 0.549122 | Other Positive Correlation |
| NPAS3 | 0.03344 | 0.549283 | Other Positive Correlation |
| AKT1 | 0.033437 | 0.549322 | Other Positive Correlation |
| BMP2K | 0.033414 | 0.5496 | Other Positive Correlation |
| ZCRB1 | 0.033386 | 0.549939 | Other Positive Correlation |
| VAMP7 | 0.033383 | 0.549966 | Other Positive Correlation |
| TECTA | 0.033372 | 0.550104 | Other Positive Correlation |
| NBAS | 0.033321 | 0.550714 | Other Positive Correlation |
| GGTA1 | 0.03332 | 0.550728 | Other Positive Correlation |
| SPTB | 0.033287 | 0.551113 | Other Positive Correlation |
| TBC1D10A | 0.033266 | 0.551368 | Other Positive Correlation |
| BMPER | 0.03321 | 0.552035 | Other Positive Correlation |
| CRELD1 | 0.033205 | 0.552099 | Other Positive Correlation |
| B3GALT9 | 0.033198 | 0.552187 | Other Positive Correlation |
| OR3A1 | 0.03317 | 0.552516 | Other Positive Correlation |
| TTC14 | 0.033168 | 0.552548 | Other Positive Correlation |
| CCIN | 0.033145 | 0.552814 | Other Positive Correlation |
| DHRS7B | 0.033137 | 0.552919 | Other Positive Correlation |
| LYPD2 | 0.033101 | 0.553345 | Other Positive Correlation |
| THUMPD3 | 0.033087 | 0.553508 | Other Positive Correlation |
| TBC1D4 | 0.033079 | 0.553605 | Other Positive Correlation |
| DUS2 | 0.03303 | 0.554197 | Other Positive Correlation |
| EXOC2 | 0.032998 | 0.554576 | Other Positive Correlation |
| RAB33B | 0.032995 | 0.554615 | Other Positive Correlation |
| PTPRN2 | 0.032979 | 0.554813 | Other Positive Correlation |
| RNF123 | 0.032967 | 0.554958 | Other Positive Correlation |
| ZNF780A | 0.032939 | 0.555293 | Other Positive Correlation |
| SLIT2 | 0.032936 | 0.55532 | Other Positive Correlation |
| ANKRD29 | 0.03287 | 0.556124 | Other Positive Correlation |
| XYLT2 | 0.032862 | 0.55622 | Other Positive Correlation |
| TFF3 | 0.032853 | 0.556322 | Other Positive Correlation |
| FOXA2 | 0.032837 | 0.556511 | Other Positive Correlation |
| C4orf33 | 0.032792 | 0.557053 | Other Positive Correlation |
| OR7C1 | 0.032772 | 0.557298 | Other Positive Correlation |
| SPIB | 0.032732 | 0.557782 | Other Positive Correlation |
| GRIP2 | 0.032729 | 0.557818 | Other Positive Correlation |
| HMGN3 | 0.032676 | 0.558449 | Other Positive Correlation |
| BET1 | 0.03264 | 0.558889 | Other Positive Correlation |
| GPNMB | 0.032622 | 0.559107 | Other Positive Correlation |
| GJC3 | 0.032609 | 0.559266 | Other Positive Correlation |
| ZNF419 | 0.032606 | 0.559294 | Other Positive Correlation |
| RNF8 | 0.032545 | 0.560034 | Other Positive Correlation |

| | | | |
|---|---|---|---|
| TTYH1 | 0.032513 | 0.560425 | Other Positive Correlation |
| ENDOD1 | 0.032495 | 0.56064 | Other Positive Correlation |
| PRR16 | 0.032476 | 0.560865 | Other Positive Correlation |
| CUL7 | 0.032451 | 0.561168 | Other Positive Correlation |
| LRRC14B | 0.032442 | 0.561279 | Other Positive Correlation |
| RANGRF | 0.032438 | 0.561322 | Other Positive Correlation |
| FAM72A | 0.032427 | 0.561458 | Other Positive Correlation |
| GUCA1A | 0.032397 | 0.561822 | Other Positive Correlation |
| APOBEC3D | 0.032389 | 0.561915 | Other Positive Correlation |
| DNAJC12 | 0.032371 | 0.562134 | Other Positive Correlation |
| PANX1 | 0.032369 | 0.562162 | Other Positive Correlation |
| PIKFYVE | 0.032366 | 0.562194 | Other Positive Correlation |
| ZNF496 | 0.032344 | 0.562464 | Other Positive Correlation |
| NRROS | 0.032343 | 0.562474 | Other Positive Correlation |
| CLPSL2 | 0.03234 | 0.562513 | Other Positive Correlation |
| CRISPLD1 | 0.032331 | 0.562616 | Other Positive Correlation |
| GNPDA1 | 0.032274 | 0.563303 | Other Positive Correlation |
| KDM4B | 0.03223 | 0.563845 | Other Positive Correlation |
| YIPF3 | 0.032224 | 0.563906 | Other Positive Correlation |
| COA1 | 0.032206 | 0.564135 | Other Positive Correlation |
| VPS26C | 0.032199 | 0.564209 | Other Positive Correlation |
| GON4L | 0.032196 | 0.564245 | Other Positive Correlation |
| DYRK1B | 0.032185 | 0.56439 | Other Positive Correlation |
| SLC19A3 | 0.032137 | 0.564962 | Other Positive Correlation |
| ZBTB39 | 0.032136 | 0.564972 | Other Positive Correlation |
| CTAGE15 | 0.032105 | 0.565354 | Other Positive Correlation |
| WTIP | 0.03206 | 0.565901 | Other Positive Correlation |
| PDE9A | 0.032041 | 0.566126 | Other Positive Correlation |
| INO80D | 0.032034 | 0.566209 | Other Positive Correlation |
| ACVR2A | 0.032007 | 0.566536 | Other Positive Correlation |
| TRMT44 | 0.032006 | 0.566556 | Other Positive Correlation |
| DPM3 | 0.032004 | 0.566573 | Other Positive Correlation |
| FAN1 | 0.031974 | 0.566944 | Other Positive Correlation |
| OPRL1 | 0.031967 | 0.567022 | Other Positive Correlation |
| KLHL25 | 0.031948 | 0.567257 | Other Positive Correlation |
| ETFRF1 | 0.031926 | 0.567519 | Other Positive Correlation |
| SLC30A8 | 0.031919 | 0.567606 | Other Positive Correlation |
| PRUNE1 | 0.031902 | 0.567817 | Other Positive Correlation |
| XPOT | 0.0319 | 0.567844 | Other Positive Correlation |
| RP1L1 | 0.031874 | 0.568158 | Other Positive Correlation |
| PIGBOS1 | 0.03187 | 0.568202 | Other Positive Correlation |
| TGDS | 0.031851 | 0.568439 | Other Positive Correlation |
| RCSD1 | 0.031844 | 0.568523 | Other Positive Correlation |
| PPP1R7 | 0.031808 | 0.56896 | Other Positive Correlation |
| GIP | 0.031766 | 0.569464 | Other Positive Correlation |
| KMT2A | 0.031758 | 0.569562 | Other Positive Correlation |
| SEC23IP | 0.031753 | 0.569624 | Other Positive Correlation |
| IGSF5 | 0.031743 | 0.569751 | Other Positive Correlation |

| | | | |
|---|---|---|---|
| TNFRSF1A | 0.031739 | 0.569801 | Other Positive Correlation |
| ZNF563 | 0.031732 | 0.569878 | Other Positive Correlation |
| VPS9D1 | 0.031695 | 0.570332 | Other Positive Correlation |
| PIH1D2 | 0.031686 | 0.570437 | Other Positive Correlation |
| NTHL1 | 0.031667 | 0.570676 | Other Positive Correlation |
| CSTF3 | 0.031642 | 0.570976 | Other Positive Correlation |
| TLR4 | 0.031632 | 0.571099 | Other Positive Correlation |
| DNAH7 | 0.031611 | 0.571349 | Other Positive Correlation |
| FHAD1 | 0.031598 | 0.571516 | Other Positive Correlation |
| KLRF1 | 0.031583 | 0.571701 | Other Positive Correlation |
| GNGT2 | 0.031582 | 0.571702 | Other Positive Correlation |
| RASSF7 | 0.031519 | 0.572474 | Other Positive Correlation |
| ADAMTSL3 | 0.0315 | 0.572701 | Other Positive Correlation |
| ZNF665 | 0.0315 | 0.572705 | Other Positive Correlation |
| C17orf113 | 0.031495 | 0.572762 | Other Positive Correlation |
| TRPC1 | 0.031483 | 0.572916 | Other Positive Correlation |
| CLUH | 0.031398 | 0.573951 | Other Positive Correlation |
| CCL5 | 0.031398 | 0.573954 | Other Positive Correlation |
| CADM3 | 0.031351 | 0.57453 | Other Positive Correlation |
| CYB561D2 | 0.031311 | 0.575018 | Other Positive Correlation |
| CD22 | 0.031309 | 0.575038 | Other Positive Correlation |
| SCIMP | 0.031298 | 0.575169 | Other Positive Correlation |
| NAA40 | 0.031297 | 0.575185 | Other Positive Correlation |
| CCT6B | 0.031286 | 0.575316 | Other Positive Correlation |
| CXCL6 | 0.031283 | 0.575357 | Other Positive Correlation |
| GGA1 | 0.031267 | 0.575547 | Other Positive Correlation |
| RASA4B | 0.031245 | 0.575821 | Other Positive Correlation |
| SMPD3 | 0.031242 | 0.575858 | Other Positive Correlation |
| NTN5 | 0.031221 | 0.576117 | Other Positive Correlation |
| C19orf84 | 0.031207 | 0.576289 | Other Positive Correlation |
| CCM2 | 0.031204 | 0.576326 | Other Positive Correlation |
| GPBP1 | 0.031174 | 0.576692 | Other Positive Correlation |
| PXYLP1 | 0.031161 | 0.576845 | Other Positive Correlation |
| SLC25A28 | 0.031141 | 0.577087 | Other Positive Correlation |
| ARID2 | 0.03114 | 0.577102 | Other Positive Correlation |
| NR2C2 | 0.031131 | 0.577216 | Other Positive Correlation |
| FAM200B | 0.031103 | 0.577552 | Other Positive Correlation |
| CARD19 | 0.031096 | 0.577641 | Other Positive Correlation |
| PCDHB2 | 0.031077 | 0.577878 | Other Positive Correlation |
| ELMO2 | 0.031052 | 0.578177 | Other Positive Correlation |
| RNFT1 | 0.031015 | 0.578635 | Other Positive Correlation |
| CFAP221 | 0.030988 | 0.578962 | Other Positive Correlation |
| C8orf34 | 0.030986 | 0.57899 | Other Positive Correlation |
| KPNA5 | 0.030971 | 0.579168 | Other Positive Correlation |
| PLEKHH3 | 0.030942 | 0.579527 | Other Positive Correlation |
| SGK2 | 0.030926 | 0.57973 | Other Positive Correlation |
| FGF2 | 0.030906 | 0.579968 | Other Positive Correlation |
| RBP5 | 0.030886 | 0.58022 | Other Positive Correlation |

| | | | |
|---|---|---|---|
| RHOG | 0.030884 | 0.580235 | Other Positive Correlation |
| NWD2 | 0.030855 | 0.580597 | Other Positive Correlation |
| RNF144A | 0.030848 | 0.580688 | Other Positive Correlation |
| SMIM5 | 0.030835 | 0.580836 | Other Positive Correlation |
| HS3ST3B1 | 0.030826 | 0.580958 | Other Positive Correlation |
| CENATAC | 0.030792 | 0.581372 | Other Positive Correlation |
| XKR5 | 0.030778 | 0.581539 | Other Positive Correlation |
| KDM5A | 0.030742 | 0.581987 | Other Positive Correlation |
| DPF2 | 0.030737 | 0.582049 | Other Positive Correlation |
| ENGASE | 0.030727 | 0.582168 | Other Positive Correlation |
| SPATA7 | 0.030721 | 0.582241 | Other Positive Correlation |
| ATP13A4 | 0.030699 | 0.582507 | Other Positive Correlation |
| CTXN1 | 0.030696 | 0.58255 | Other Positive Correlation |
| SHISA6 | 0.030677 | 0.582782 | Other Positive Correlation |
| PACRG | 0.030662 | 0.582968 | Other Positive Correlation |
| IFT20 | 0.030659 | 0.583 | Other Positive Correlation |
| USH1G | 0.030641 | 0.58323 | Other Positive Correlation |
| KIN | 0.030628 | 0.583387 | Other Positive Correlation |
| MSRB3 | 0.030613 | 0.583565 | Other Positive Correlation |
| GATA6 | 0.030606 | 0.583656 | Other Positive Correlation |
| CLEC7A | 0.030564 | 0.584175 | Other Positive Correlation |
| ARL11 | 0.030562 | 0.584193 | Other Positive Correlation |
| TMEM35A | 0.030534 | 0.584538 | Other Positive Correlation |
| CDKAL1 | 0.030533 | 0.584549 | Other Positive Correlation |
| GRAMD4 | 0.030517 | 0.584753 | Other Positive Correlation |
| TMEM120A | 0.030479 | 0.585219 | Other Positive Correlation |
| GALNT13 | 0.030456 | 0.585501 | Other Positive Correlation |
| FZD4 | 0.030453 | 0.585542 | Other Positive Correlation |
| TTC34 | 0.030444 | 0.585653 | Other Positive Correlation |
| GOSR1 | 0.030443 | 0.585658 | Other Positive Correlation |
| IL11 | 0.030442 | 0.585677 | Other Positive Correlation |
| GSC | 0.030403 | 0.586149 | Other Positive Correlation |
| ENSG00000284188 | 0.030361 | 0.586667 | Other Positive Correlation |
| ZNF174 | 0.030357 | 0.586718 | Other Positive Correlation |
| AAMP | 0.030351 | 0.586796 | Other Positive Correlation |
| KIF9 | 0.03035 | 0.586803 | Other Positive Correlation |
| PPP1R13B | 0.030318 | 0.587208 | Other Positive Correlation |
| RABAC1 | 0.03031 | 0.587302 | Other Positive Correlation |
| ZNF878 | 0.030288 | 0.587574 | Other Positive Correlation |
| DACT2 | 0.030268 | 0.587814 | Other Positive Correlation |
| MAMSTR | 0.030256 | 0.587971 | Other Positive Correlation |
| MYH3 | 0.030255 | 0.587984 | Other Positive Correlation |
| OR10A2 | 0.030241 | 0.588152 | Other Positive Correlation |
| RNF214 | 0.030196 | 0.588707 | Other Positive Correlation |
| MYL11 | 0.030193 | 0.58875 | Other Positive Correlation |
| ADAMTS10 | 0.03019 | 0.588779 | Other Positive Correlation |
| NPIPA2 | 0.030184 | 0.588857 | Other Positive Correlation |
| ENSG00000249141 | 0.030168 | 0.58905 | Other Positive Correlation |

| | | | |
|---|---|---|---|
| TLCD3B | 0.030138 | 0.589423 | Other Positive Correlation |
| KCNG2 | 0.030138 | 0.589423 | Other Positive Correlation |
| UBAP1L | 0.030115 | 0.589711 | Other Positive Correlation |
| ADAD2 | 0.03009 | 0.590018 | Other Positive Correlation |
| EXD3 | 0.030087 | 0.590054 | Other Positive Correlation |
| MAGEL2 | 0.030087 | 0.590054 | Other Positive Correlation |
| FHOD3 | 0.030058 | 0.590418 | Other Positive Correlation |
| ACSL5 | 0.030052 | 0.590487 | Other Positive Correlation |
| PDPR | 0.030045 | 0.590576 | Other Positive Correlation |
| ALDH3B1 | 0.029984 | 0.591335 | Other Positive Correlation |
| CDK2 | 0.029976 | 0.591431 | Other Positive Correlation |
| MEF2B | 0.0299 | 0.592375 | Other Positive Correlation |
| ANXA9 | 0.029898 | 0.5924 | Other Positive Correlation |
| GFRA3 | 0.029888 | 0.592515 | Other Positive Correlation |
| CHD8 | 0.029885 | 0.592549 | Other Positive Correlation |
| ACVR2B | 0.029819 | 0.593376 | Other Positive Correlation |
| CPLANE2 | 0.029815 | 0.593422 | Other Positive Correlation |
| H3Y2 | 0.029792 | 0.593713 | Other Positive Correlation |
| TRIM51 | 0.029792 | 0.593713 | Other Positive Correlation |
| PMP22 | 0.029788 | 0.59376 | Other Positive Correlation |
| ENSG00000285253 | 0.029777 | 0.593899 | Other Positive Correlation |
| KRT13 | 0.029768 | 0.594006 | Other Positive Correlation |
| RSPRY1 | 0.029752 | 0.594198 | Other Positive Correlation |
| SLC47A1 | 0.029748 | 0.594259 | Other Positive Correlation |
| SLCO3A1 | 0.029718 | 0.59463 | Other Positive Correlation |
| WWC1 | 0.029679 | 0.595111 | Other Positive Correlation |
| REPS2 | 0.029677 | 0.595132 | Other Positive Correlation |
| PRAC1 | 0.029672 | 0.595194 | Other Positive Correlation |
| ANKRD28 | 0.029672 | 0.595199 | Other Positive Correlation |
| UBAC1 | 0.029652 | 0.595451 | Other Positive Correlation |
| TIRAP | 0.029645 | 0.595533 | Other Positive Correlation |
| PKLR | 0.029572 | 0.596442 | Other Positive Correlation |
| AMY2B | 0.029565 | 0.596521 | Other Positive Correlation |
| AIFM1 | 0.029563 | 0.596556 | Other Positive Correlation |
| PRX | 0.029548 | 0.596742 | Other Positive Correlation |
| NAPSA | 0.029539 | 0.596844 | Other Positive Correlation |
| SLC17A5 | 0.029509 | 0.597217 | Other Positive Correlation |
| SLC6A12 | 0.029463 | 0.597799 | Other Positive Correlation |
| ENSG00000285953 | 0.029446 | 0.59801 | Other Positive Correlation |
| NACAD | 0.02943 | 0.598203 | Other Positive Correlation |
| CKLF-CMTM1 | 0.029413 | 0.598416 | Other Positive Correlation |
| DCUN1D4 | 0.029409 | 0.598464 | Other Positive Correlation |
| MAPK3 | 0.029405 | 0.598513 | Other Positive Correlation |
| PRDM2 | 0.02936 | 0.59907 | Other Positive Correlation |
| NKD1 | 0.02934 | 0.599319 | Other Positive Correlation |
| RNF185 | 0.029316 | 0.599627 | Other Positive Correlation |
| PGS1 | 0.029299 | 0.599834 | Other Positive Correlation |
| PAK5 | 0.029285 | 0.600009 | Other Positive Correlation |

| | | | |
|---|---|---|---|
| KLHL38 | 0.029274 | 0.600147 | Other Positive Correlation |
| ERAL1 | 0.029259 | 0.600333 | Other Positive Correlation |
| TMEM121B | 0.029244 | 0.600525 | Other Positive Correlation |
| CFAP61 | 0.029234 | 0.600647 | Other Positive Correlation |
| SPPL3 | 0.029206 | 0.600991 | Other Positive Correlation |
| GTDC1 | 0.0292 | 0.601068 | Other Positive Correlation |
| ATPSCKMT | 0.029196 | 0.601123 | Other Positive Correlation |
| CILK1 | 0.029188 | 0.601215 | Other Positive Correlation |
| SBF1 | 0.029187 | 0.601226 | Other Positive Correlation |
| SAMD5 | 0.029149 | 0.601705 | Other Positive Correlation |
| ANKAR | 0.029142 | 0.601789 | Other Positive Correlation |
| MAU2 | 0.029131 | 0.601924 | Other Positive Correlation |
| SH3BGRL2 | 0.029119 | 0.602083 | Other Positive Correlation |
| TRPC3 | 0.029053 | 0.602901 | Other Positive Correlation |
| TXK | 0.029053 | 0.602901 | Other Positive Correlation |
| MUCL1 | 0.029036 | 0.603115 | Other Positive Correlation |
| PCDHGA11 | 0.028991 | 0.603673 | Other Positive Correlation |
| AOC3 | 0.028986 | 0.603733 | Other Positive Correlation |
| EPO | 0.028947 | 0.604221 | Other Positive Correlation |
| FHIP1A | 0.028938 | 0.604344 | Other Positive Correlation |
| IL1R2 | 0.028934 | 0.604394 | Other Positive Correlation |
| SCARA3 | 0.028932 | 0.604415 | Other Positive Correlation |
| C12orf29 | 0.028918 | 0.604583 | Other Positive Correlation |
| DNAAF8 | 0.028911 | 0.604672 | Other Positive Correlation |
| LZTR1 | 0.028909 | 0.604704 | Other Positive Correlation |
| SLC35C2 | 0.028893 | 0.604902 | Other Positive Correlation |
| CREB3L3 | 0.028863 | 0.605274 | Other Positive Correlation |
| POU2AF1 | 0.028846 | 0.605483 | Other Positive Correlation |
| HSCB | 0.02881 | 0.605943 | Other Positive Correlation |
| BNIPL | 0.028809 | 0.605946 | Other Positive Correlation |
| CATSPERE | 0.028788 | 0.606216 | Other Positive Correlation |
| ANKRD46 | 0.028775 | 0.606375 | Other Positive Correlation |
| EFCAB7 | 0.028768 | 0.60646 | Other Positive Correlation |
| ZBTB48 | 0.028747 | 0.60673 | Other Positive Correlation |
| ANKRD24 | 0.028712 | 0.607161 | Other Positive Correlation |
| RAB2B | 0.028694 | 0.607389 | Other Positive Correlation |
| ITIH6 | 0.028643 | 0.60803 | Other Positive Correlation |
| TSNAX | 0.028641 | 0.60806 | Other Positive Correlation |
| CRYBG3 | 0.028592 | 0.608669 | Other Positive Correlation |
| SLC7A2 | 0.028572 | 0.608925 | Other Positive Correlation |
| SLC11A1 | 0.028555 | 0.609129 | Other Positive Correlation |
| VPS54 | 0.028547 | 0.609228 | Other Positive Correlation |
| H2AL3 | 0.028544 | 0.609272 | Other Positive Correlation |
| GPR155 | 0.028466 | 0.610246 | Other Positive Correlation |
| MTMR7 | 0.028456 | 0.610376 | Other Positive Correlation |
| BCAS3 | 0.02843 | 0.610705 | Other Positive Correlation |
| CUEDC1 | 0.028417 | 0.61087 | Other Positive Correlation |
| DDI2 | 0.028403 | 0.611044 | Other Positive Correlation |

| | | | |
|---|---|---|---|
| DUSP19 | 0.028328 | 0.611983 | Other Positive Correlation |
| GAS1 | 0.028315 | 0.612149 | Other Positive Correlation |
| PF4V1 | 0.028288 | 0.612487 | Other Positive Correlation |
| AZU1 | 0.028267 | 0.612751 | Other Positive Correlation |
| PRAF2 | 0.028258 | 0.612868 | Other Positive Correlation |
| MAPKAPK5 | 0.028216 | 0.613395 | Other Positive Correlation |
| PLA2G2F | 0.028116 | 0.614651 | Other Positive Correlation |
| SPATA5 | 0.02811 | 0.61473 | Other Positive Correlation |
| CLDN14 | 0.028063 | 0.615318 | Other Positive Correlation |
| MGAT5 | 0.028035 | 0.615676 | Other Positive Correlation |
| ZNF681 | 0.028019 | 0.615878 | Other Positive Correlation |
| SGPP1 | 0.028012 | 0.615958 | Other Positive Correlation |
| STK35 | 0.027982 | 0.616342 | Other Positive Correlation |
| MYPN | 0.027961 | 0.616604 | Other Positive Correlation |
| NOP9 | 0.027953 | 0.616698 | Other Positive Correlation |
| TRAM1L1 | 0.027897 | 0.617414 | Other Positive Correlation |
| ABCD3 | 0.027888 | 0.617525 | Other Positive Correlation |
| SRXN1 | 0.027873 | 0.61771 | Other Positive Correlation |
| RAPGEF4 | 0.027848 | 0.61803 | Other Positive Correlation |
| PLPPR5 | 0.027808 | 0.618532 | Other Positive Correlation |
| MRGBP | 0.027802 | 0.61861 | Other Positive Correlation |
| SH2D3C | 0.027801 | 0.618618 | Other Positive Correlation |
| ZNF341 | 0.027762 | 0.619116 | Other Positive Correlation |
| MKRN1 | 0.027739 | 0.619407 | Other Positive Correlation |
| RPL17-C18orf32 | 0.027735 | 0.619456 | Other Positive Correlation |
| CLPTM1L | 0.027702 | 0.619875 | Other Positive Correlation |
| MBTPS2 | 0.027694 | 0.619974 | Other Positive Correlation |
| ZFTRAF1 | 0.02768 | 0.620157 | Other Positive Correlation |
| TMEM41A | 0.027665 | 0.620347 | Other Positive Correlation |
| NCALD | 0.027658 | 0.620423 | Other Positive Correlation |
| BSDC1 | 0.027651 | 0.620514 | Other Positive Correlation |
| DNHD1 | 0.027636 | 0.620703 | Other Positive Correlation |
| COL5A1 | 0.027578 | 0.621435 | Other Positive Correlation |
| MRM3 | 0.027566 | 0.621593 | Other Positive Correlation |
| SMG7 | 0.02756 | 0.621666 | Other Positive Correlation |
| CLEC2B | 0.027539 | 0.621936 | Other Positive Correlation |
| ZNF436 | 0.027463 | 0.6229 | Other Positive Correlation |
| GBP5 | 0.027427 | 0.623347 | Other Positive Correlation |
| TBC1D16 | 0.027381 | 0.623936 | Other Positive Correlation |
| HOXB2 | 0.027331 | 0.624571 | Other Positive Correlation |
| SMAD4 | 0.027316 | 0.624761 | Other Positive Correlation |
| ZNF432 | 0.027311 | 0.62482 | Other Positive Correlation |
| CDK5R1 | 0.027306 | 0.624887 | Other Positive Correlation |
| ENSG00000260914 | 0.027299 | 0.624981 | Other Positive Correlation |
| RALGDS | 0.027292 | 0.625059 | Other Positive Correlation |
| NME6 | 0.027283 | 0.625173 | Other Positive Correlation |
| SIGLEC11 | 0.027265 | 0.625409 | Other Positive Correlation |
| ZXDA | 0.02726 | 0.625475 | Other Positive Correlation |

| | | | |
|---|---|---|---|
| MOCS3 | 0.027246 | 0.625646 | Other Positive Correlation |
| TOX4 | 0.027219 | 0.625991 | Other Positive Correlation |
| ARV1 | 0.027214 | 0.626056 | Other Positive Correlation |
| TNS1 | 0.027212 | 0.626077 | Other Positive Correlation |
| TFIP11 | 0.027206 | 0.626154 | Other Positive Correlation |
| CD3D | 0.027196 | 0.626282 | Other Positive Correlation |
| ENSG00000270249 | 0.027187 | 0.626398 | Other Positive Correlation |
| NEDD8-MDP1 | 0.027187 | 0.626398 | Other Positive Correlation |
| OPRK1 | 0.027187 | 0.626398 | Other Positive Correlation |
| OTOP3 | 0.027175 | 0.626543 | Other Positive Correlation |
| CYP2W1 | 0.02716 | 0.626741 | Other Positive Correlation |
| LDB3 | 0.02715 | 0.626871 | Other Positive Correlation |
| CD63 | 0.027136 | 0.627044 | Other Positive Correlation |
| DIABLO | 0.027119 | 0.627265 | Other Positive Correlation |
| PTPN20 | 0.027101 | 0.627483 | Other Positive Correlation |
| PLS1 | 0.027016 | 0.628574 | Other Positive Correlation |
| SULT1A3 | 0.026998 | 0.628794 | Other Positive Correlation |
| ASTE1 | 0.026993 | 0.628863 | Other Positive Correlation |
| ICAM2 | 0.026976 | 0.629074 | Other Positive Correlation |
| BMPR1A | 0.026962 | 0.629251 | Other Positive Correlation |
| RAI1 | 0.02696 | 0.629277 | Other Positive Correlation |
| RIPK3 | 0.026908 | 0.629941 | Other Positive Correlation |
| ZNF383 | 0.026899 | 0.630051 | Other Positive Correlation |
| PDE1B | 0.026858 | 0.630583 | Other Positive Correlation |
| CCDC170 | 0.026854 | 0.630634 | Other Positive Correlation |
| SLC25A36 | 0.026828 | 0.630955 | Other Positive Correlation |
| SPRY1 | 0.026827 | 0.630977 | Other Positive Correlation |
| FYB2 | 0.026824 | 0.631015 | Other Positive Correlation |
| TMCO4 | 0.026821 | 0.631054 | Other Positive Correlation |
| C6orf58 | 0.026818 | 0.631093 | Other Positive Correlation |
| RAG1 | 0.026789 | 0.631458 | Other Positive Correlation |
| INPP5A | 0.026785 | 0.631506 | Other Positive Correlation |
| MXRA5 | 0.026774 | 0.631649 | Other Positive Correlation |
| COMMD1 | 0.026771 | 0.631689 | Other Positive Correlation |
| CTBP1 | 0.026749 | 0.631972 | Other Positive Correlation |
| MPDZ | 0.026741 | 0.632068 | Other Positive Correlation |
| AHDC1 | 0.026736 | 0.632136 | Other Positive Correlation |
| PDE4B | 0.026736 | 0.632137 | Other Positive Correlation |
| MINDY4 | 0.026728 | 0.632233 | Other Positive Correlation |
| ZFP82 | 0.026724 | 0.632283 | Other Positive Correlation |
| GRPR | 0.026716 | 0.632382 | Other Positive Correlation |
| COX10 | 0.026699 | 0.632599 | Other Positive Correlation |
| CCDC166 | 0.026692 | 0.632695 | Other Positive Correlation |
| MUC16 | 0.026639 | 0.633373 | Other Positive Correlation |
| ITGA2B | 0.026607 | 0.633778 | Other Positive Correlation |
| ID2 | 0.026565 | 0.634308 | Other Positive Correlation |
| ZSCAN12 | 0.026536 | 0.634687 | Other Positive Correlation |
| MEGF6 | 0.026519 | 0.634898 | Other Positive Correlation |

| | | | |
|---|---|---|---|
| ETV3 | 0.026491 | 0.635252 | Other Positive Correlation |
| UNC45A | 0.026491 | 0.63526 | Other Positive Correlation |
| SLMAP | 0.026458 | 0.635684 | Other Positive Correlation |
| ZNF37A | 0.026447 | 0.635825 | Other Positive Correlation |
| SCMH1 | 0.026422 | 0.636146 | Other Positive Correlation |
| NDUFAF5 | 0.026419 | 0.636172 | Other Positive Correlation |
| TIMM23B-AGAP6 | 0.026413 | 0.63626 | Other Positive Correlation |
| EGFLAM | 0.026411 | 0.636278 | Other Positive Correlation |
| NT5C2 | 0.026371 | 0.636789 | Other Positive Correlation |
| ALX3 | 0.026369 | 0.636819 | Other Positive Correlation |
| ZNF771 | 0.026368 | 0.636827 | Other Positive Correlation |
| WNT4 | 0.026356 | 0.636983 | Other Positive Correlation |
| CHSY1 | 0.026329 | 0.637334 | Other Positive Correlation |
| RFX4 | 0.026321 | 0.637435 | Other Positive Correlation |
| PLGLB1 | 0.026304 | 0.637654 | Other Positive Correlation |
| FURIN | 0.026249 | 0.638348 | Other Positive Correlation |
| CD47 | 0.026223 | 0.63868 | Other Positive Correlation |
| GPR173 | 0.02621 | 0.638847 | Other Positive Correlation |
| VPS11 | 0.026183 | 0.639197 | Other Positive Correlation |
| PPP1R14D | 0.026179 | 0.639251 | Other Positive Correlation |
| SLC16A14 | 0.026158 | 0.639514 | Other Positive Correlation |
| SLC30A6 | 0.026132 | 0.63985 | Other Positive Correlation |
| TMEM88 | 0.026113 | 0.640089 | Other Positive Correlation |
| COL21A1 | 0.026053 | 0.640863 | Other Positive Correlation |
| FRMD1 | 0.02604 | 0.64103 | Other Positive Correlation |
| RNF31 | 0.026002 | 0.641512 | Other Positive Correlation |
| CIZ1 | 0.025967 | 0.641967 | Other Positive Correlation |
| CFAP300 | 0.02596 | 0.642056 | Other Positive Correlation |
| LEKR1 | 0.025924 | 0.642522 | Other Positive Correlation |
| HNMT | 0.025911 | 0.642685 | Other Positive Correlation |
| FGF5 | 0.025902 | 0.642803 | Other Positive Correlation |
| SLIT1 | 0.025858 | 0.643358 | Other Positive Correlation |
| FAM90A27P | 0.025846 | 0.643523 | Other Positive Correlation |
| PDE6G | 0.02584 | 0.643593 | Other Positive Correlation |
| RABEP2 | 0.025785 | 0.644303 | Other Positive Correlation |
| PNPLA7 | 0.025747 | 0.644794 | Other Positive Correlation |
| ZNF613 | 0.025732 | 0.644982 | Other Positive Correlation |
| WFDC10B | 0.025723 | 0.645101 | Other Positive Correlation |
| CALB1 | 0.025705 | 0.645334 | Other Positive Correlation |
| NPIPB3 | 0.025689 | 0.64553 | Other Positive Correlation |
| IGIP | 0.025673 | 0.645745 | Other Positive Correlation |
| ZSCAN18 | 0.025669 | 0.645797 | Other Positive Correlation |
| AVPR2 | 0.025636 | 0.646217 | Other Positive Correlation |
| RHOBTB3 | 0.025623 | 0.646385 | Other Positive Correlation |
| LTF | 0.025611 | 0.646543 | Other Positive Correlation |
| ZNF582 | 0.025593 | 0.646768 | Other Positive Correlation |
| KHDC1L | 0.025515 | 0.647772 | Other Positive Correlation |
| TAGAP | 0.025502 | 0.647947 | Other Positive Correlation |

| | | | |
|---|---|---|---|
| GABRB1 | 0.025484 | 0.648167 | Other Positive Correlation |
| CC2D2B | 0.025404 | 0.649199 | Other Positive Correlation |
| PTCRA | 0.025397 | 0.649291 | Other Positive Correlation |
| GRM7 | 0.025397 | 0.649291 | Other Positive Correlation |
| CITED1 | 0.025397 | 0.649291 | Other Positive Correlation |
| TUBB8 | 0.025386 | 0.649438 | Other Positive Correlation |
| LGALS2 | 0.025386 | 0.649438 | Other Positive Correlation |
| SPAAR | 0.025345 | 0.649967 | Other Positive Correlation |
| SCG2 | 0.025319 | 0.650295 | Other Positive Correlation |
| TENT5B | 0.025291 | 0.65066 | Other Positive Correlation |
| TSSK4 | 0.02529 | 0.65067 | Other Positive Correlation |
| MCIDAS | 0.025275 | 0.65086 | Other Positive Correlation |
| TSTD2 | 0.025272 | 0.6509 | Other Positive Correlation |
| SALL3 | 0.025252 | 0.651164 | Other Positive Correlation |
| SPCS2 | 0.025229 | 0.651462 | Other Positive Correlation |
| ENSG00000261582 | 0.025227 | 0.651485 | Other Positive Correlation |
| VTCN1 | 0.025204 | 0.651785 | Other Positive Correlation |
| NPW | 0.025204 | 0.651789 | Other Positive Correlation |
| TRAF1 | 0.025201 | 0.651825 | Other Positive Correlation |
| ENSG00000284797 | 0.025164 | 0.6523 | Other Positive Correlation |
| TRAPPC14 | 0.025112 | 0.652974 | Other Positive Correlation |
| RALGAPA2 | 0.025089 | 0.653271 | Other Positive Correlation |
| PIAS2 | 0.025085 | 0.653325 | Other Positive Correlation |
| BCL6B | 0.025069 | 0.653522 | Other Positive Correlation |
| OR4F17 | 0.025069 | 0.653526 | Other Positive Correlation |
| MAGEE2 | 0.025069 | 0.653526 | Other Positive Correlation |
| NALF1 | 0.025069 | 0.653526 | Other Positive Correlation |
| FOXJ3 | 0.025031 | 0.654023 | Other Positive Correlation |
| FAM193B | 0.025026 | 0.654086 | Other Positive Correlation |
| ZSCAN32 | 0.025008 | 0.654318 | Other Positive Correlation |
| TLDC2 | 0.024981 | 0.654662 | Other Positive Correlation |
| FCMR | 0.024962 | 0.654904 | Other Positive Correlation |
| CARMIL3 | 0.02492 | 0.655447 | Other Positive Correlation |
| HEPN1 | 0.024903 | 0.655668 | Other Positive Correlation |
| PIP4P2 | 0.024894 | 0.655788 | Other Positive Correlation |
| RNF114 | 0.024872 | 0.656078 | Other Positive Correlation |
| DMAC2 | 0.024857 | 0.656268 | Other Positive Correlation |
| PLSCR4 | 0.024847 | 0.656398 | Other Positive Correlation |
| CAPN10 | 0.024838 | 0.656516 | Other Positive Correlation |
| WFDC12 | 0.024814 | 0.65682 | Other Positive Correlation |
| SELENOT | 0.024806 | 0.656929 | Other Positive Correlation |
| TM6SF1 | 0.024804 | 0.656949 | Other Positive Correlation |
| BRI3 | 0.024801 | 0.656991 | Other Positive Correlation |
| PRR33 | 0.0248 | 0.657 | Other Positive Correlation |
| OTOS | 0.024786 | 0.657191 | Other Positive Correlation |
| TRPC4 | 0.024753 | 0.657613 | Other Positive Correlation |
| TRIM47 | 0.024751 | 0.657639 | Other Positive Correlation |
| LY6G6C | 0.024743 | 0.657749 | Other Positive Correlation |

| | | | |
|---|---|---|---|
| TRIL | 0.02474 | 0.657781 | Other Positive Correlation |
| ANKRD40 | 0.024725 | 0.65798 | Other Positive Correlation |
| CBL | 0.024697 | 0.658335 | Other Positive Correlation |
| TMCO1 | 0.024688 | 0.658457 | Other Positive Correlation |
| TMEM31 | 0.024686 | 0.658486 | Other Positive Correlation |
| TPO | 0.024615 | 0.659404 | Other Positive Correlation |
| DDX42 | 0.024604 | 0.65955 | Other Positive Correlation |
| GGN | 0.024597 | 0.659634 | Other Positive Correlation |
| CASP6 | 0.024589 | 0.659746 | Other Positive Correlation |
| LMTK3 | 0.024566 | 0.660033 | Other Positive Correlation |
| B4GALT7 | 0.024541 | 0.660369 | Other Positive Correlation |
| ATG3 | 0.02453 | 0.660512 | Other Positive Correlation |
| PRR3 | 0.024469 | 0.661293 | Other Positive Correlation |
| BANP | 0.024437 | 0.661712 | Other Positive Correlation |
| THEMIS2 | 0.024411 | 0.662046 | Other Positive Correlation |
| KRT3 | 0.024383 | 0.662413 | Other Positive Correlation |
| ABCD2 | 0.024383 | 0.662413 | Other Positive Correlation |
| RO60 | 0.024315 | 0.663292 | Other Positive Correlation |
| SLC2A11 | 0.024295 | 0.663558 | Other Positive Correlation |
| TEP1 | 0.02428 | 0.663747 | Other Positive Correlation |
| FAM43B | 0.024278 | 0.663784 | Other Positive Correlation |
| AAMDC | 0.02423 | 0.664398 | Other Positive Correlation |
| PTPRD | 0.024215 | 0.6646 | Other Positive Correlation |
| MCRIP2 | 0.024178 | 0.665085 | Other Positive Correlation |
| ZNF292 | 0.024172 | 0.665156 | Other Positive Correlation |
| ST8SIA1 | 0.02412 | 0.665838 | Other Positive Correlation |
| CD70 | 0.02403 | 0.667005 | Other Positive Correlation |
| STAM2 | 0.024014 | 0.667213 | Other Positive Correlation |
| FRMPD1 | 0.024014 | 0.667214 | Other Positive Correlation |
| CERK | 0.024005 | 0.667329 | Other Positive Correlation |
| BORCS6 | 0.02399 | 0.667523 | Other Positive Correlation |
| DOCK9 | 0.023986 | 0.667579 | Other Positive Correlation |
| GGA3 | 0.023981 | 0.667646 | Other Positive Correlation |
| P2RY1 | 0.023977 | 0.667703 | Other Positive Correlation |
| RSPH14 | 0.023935 | 0.668246 | Other Positive Correlation |
| FCGR2A | 0.023915 | 0.668504 | Other Positive Correlation |
| INHA | 0.023907 | 0.66861 | Other Positive Correlation |
| OLAH | 0.023886 | 0.668882 | Other Positive Correlation |
| UPK1B | 0.023878 | 0.668994 | Other Positive Correlation |
| RNF216 | 0.023865 | 0.66916 | Other Positive Correlation |
| HOGA1 | 0.023845 | 0.669423 | Other Positive Correlation |
| FOXO3B | 0.023781 | 0.670255 | Other Positive Correlation |
| PROB1 | 0.023727 | 0.670959 | Other Positive Correlation |
| SLC28A1 | 0.023721 | 0.671035 | Other Positive Correlation |
| PRKN | 0.023707 | 0.671214 | Other Positive Correlation |
| GSTCD | 0.023683 | 0.67153 | Other Positive Correlation |
| RCBTB1 | 0.023675 | 0.671638 | Other Positive Correlation |
| TBX1 | 0.023627 | 0.672269 | Other Positive Correlation |

| | | | |
|---|---|---|---|
| ZDHHC3 | 0.023586 | 0.6728 | Other Positive Correlation |
| CYP1A2 | 0.023568 | 0.673031 | Other Positive Correlation |
| HMGXB3 | 0.023517 | 0.673706 | Other Positive Correlation |
| CEND1 | 0.023512 | 0.673771 | Other Positive Correlation |
| NLRC3 | 0.023471 | 0.674308 | Other Positive Correlation |
| ZNF354B | 0.023447 | 0.674617 | Other Positive Correlation |
| SCD5 | 0.023435 | 0.67477 | Other Positive Correlation |
| C6orf52 | 0.023419 | 0.674983 | Other Positive Correlation |
| LPAR2 | 0.023413 | 0.675059 | Other Positive Correlation |
| NUDT10 | 0.023389 | 0.67538 | Other Positive Correlation |
| HAUS5 | 0.023366 | 0.675679 | Other Positive Correlation |
| SDSL | 0.023353 | 0.675854 | Other Positive Correlation |
| GRM1 | 0.023349 | 0.6759 | Other Positive Correlation |
| NMUR2 | 0.023349 | 0.6759 | Other Positive Correlation |
| SCGB1C2 | 0.023349 | 0.6759 | Other Positive Correlation |
| NTRK3 | 0.023348 | 0.675916 | Other Positive Correlation |
| CST2 | 0.023348 | 0.675916 | Other Positive Correlation |
| TMEM210 | 0.023348 | 0.675916 | Other Positive Correlation |
| IGSF21 | 0.023348 | 0.675916 | Other Positive Correlation |
| TACR1 | 0.023348 | 0.675916 | Other Positive Correlation |
| ULBP2 | 0.023343 | 0.675982 | Other Positive Correlation |
| ZNF705A | 0.023307 | 0.676446 | Other Positive Correlation |
| TRIP10 | 0.023288 | 0.676703 | Other Positive Correlation |
| PCDHB8 | 0.023286 | 0.676721 | Other Positive Correlation |
| BBS1 | 0.02327 | 0.67694 | Other Positive Correlation |
| PROS1 | 0.023266 | 0.676992 | Other Positive Correlation |
| KRT37 | 0.02325 | 0.677198 | Other Positive Correlation |
| ICAM5 | 0.023228 | 0.677481 | Other Positive Correlation |
| MYLK3 | 0.023213 | 0.677686 | Other Positive Correlation |
| DLG4 | 0.023194 | 0.67793 | Other Positive Correlation |
| MYO1G | 0.023192 | 0.677959 | Other Positive Correlation |
| RAD21L1 | 0.023186 | 0.678035 | Other Positive Correlation |
| SCGB1A1 | 0.023162 | 0.678354 | Other Positive Correlation |
| SYS1 | 0.023147 | 0.678544 | Other Positive Correlation |
| GPR84 | 0.023127 | 0.678811 | Other Positive Correlation |
| NELFB | 0.023086 | 0.67935 | Other Positive Correlation |
| UBQLN4 | 0.023049 | 0.679834 | Other Positive Correlation |
| ATG2A | 0.023049 | 0.679839 | Other Positive Correlation |
| CDH23 | 0.023036 | 0.680003 | Other Positive Correlation |
| TM4SF18 | 0.023002 | 0.680453 | Other Positive Correlation |
| UNCX | 0.022994 | 0.680552 | Other Positive Correlation |
| GPR132 | 0.022993 | 0.680573 | Other Positive Correlation |
| TEX37 | 0.022983 | 0.680701 | Other Positive Correlation |
| NPIPA8 | 0.022972 | 0.68084 | Other Positive Correlation |
| GOLGA8M | 0.022952 | 0.681107 | Other Positive Correlation |
| MTF2 | 0.022945 | 0.681194 | Other Positive Correlation |
| HINT2 | 0.022923 | 0.681482 | Other Positive Correlation |
| TOM1L2 | 0.022876 | 0.6821 | Other Positive Correlation |

| | | | |
|---|---|---|---|
| CCDC78 | 0.022863 | 0.682276 | Other Positive Correlation |
| LRGUK | 0.022856 | 0.682369 | Other Positive Correlation |
| FAM107A | 0.022806 | 0.68302 | Other Positive Correlation |
| ERN2 | 0.022784 | 0.683309 | Other Positive Correlation |
| DRAXIN | 0.022776 | 0.68342 | Other Positive Correlation |
| CDSN | 0.022774 | 0.683442 | Other Positive Correlation |
| USP34 | 0.022767 | 0.683536 | Other Positive Correlation |
| FAM168A | 0.022751 | 0.683748 | Other Positive Correlation |
| TTC30A | 0.022729 | 0.684035 | Other Positive Correlation |
| TP53TG3D | 0.022701 | 0.684404 | Other Positive Correlation |
| MTMR1 | 0.022672 | 0.684789 | Other Positive Correlation |
| H3-5 | 0.022671 | 0.684803 | Other Positive Correlation |
| IQSEC2 | 0.022667 | 0.684855 | Other Positive Correlation |
| ANKDD1B | 0.022664 | 0.684894 | Other Positive Correlation |
| ANKRD13D | 0.022653 | 0.685041 | Other Positive Correlation |
| JRKL | 0.022649 | 0.685086 | Other Positive Correlation |
| NKX3-1 | 0.022641 | 0.685195 | Other Positive Correlation |
| PRKACA | 0.022632 | 0.685318 | Other Positive Correlation |
| CCR1 | 0.022589 | 0.685875 | Other Positive Correlation |
| CD19 | 0.022563 | 0.686219 | Other Positive Correlation |
| TBRG1 | 0.022561 | 0.686254 | Other Positive Correlation |
| GPR3 | 0.02256 | 0.686262 | Other Positive Correlation |
| C2orf42 | 0.022544 | 0.686477 | Other Positive Correlation |
| MKNK2 | 0.022511 | 0.686902 | Other Positive Correlation |
| TMC1 | 0.02243 | 0.687979 | Other Positive Correlation |
| IL2RB | 0.022422 | 0.688082 | Other Positive Correlation |
| ZNF44 | 0.022419 | 0.688124 | Other Positive Correlation |
| MT1B | 0.022408 | 0.688261 | Other Positive Correlation |
| MIEF2 | 0.022378 | 0.688658 | Other Positive Correlation |
| TMEM25 | 0.022347 | 0.689072 | Other Positive Correlation |
| NUDCD3 | 0.022344 | 0.689105 | Other Positive Correlation |
| SHROOM1 | 0.022336 | 0.689213 | Other Positive Correlation |
| ERV3-1 | 0.02229 | 0.689819 | Other Positive Correlation |
| ARHGEF19 | 0.022258 | 0.690243 | Other Positive Correlation |
| VGLL3 | 0.022235 | 0.690545 | Other Positive Correlation |
| LCMT1 | 0.022226 | 0.690662 | Other Positive Correlation |
| ZHX2 | 0.022221 | 0.690727 | Other Positive Correlation |
| ZNF662 | 0.022194 | 0.691085 | Other Positive Correlation |
| CCDC73 | 0.02217 | 0.691409 | Other Positive Correlation |
| KLF16 | 0.022167 | 0.691443 | Other Positive Correlation |
| ADI1 | 0.022149 | 0.691685 | Other Positive Correlation |
| DGKI | 0.022143 | 0.691766 | Other Positive Correlation |
| DCUN1D1 | 0.022135 | 0.691871 | Other Positive Correlation |
| ZNF416 | 0.022116 | 0.692117 | Other Positive Correlation |
| NID1 | 0.022102 | 0.692304 | Other Positive Correlation |
| PAMR1 | 0.022093 | 0.692421 | Other Positive Correlation |
| TRIM33 | 0.022083 | 0.69256 | Other Positive Correlation |
| CMKLR2 | 0.02208 | 0.692601 | Other Positive Correlation |

| | | | |
|---|---|---|---|
| CTC1 | 0.022056 | 0.692908 | Other Positive Correlation |
| GJA3 | 0.022053 | 0.69295 | Other Positive Correlation |
| ANKMY2 | 0.022042 | 0.693101 | Other Positive Correlation |
| CHRNA6 | 0.022024 | 0.693339 | Other Positive Correlation |
| IFT74 | 0.022017 | 0.69343 | Other Positive Correlation |
| PCDHB4 | 0.022014 | 0.693469 | Other Positive Correlation |
| SHOX | 0.022004 | 0.693598 | Other Positive Correlation |
| TMEM238L | 0.021992 | 0.693755 | Other Positive Correlation |
| KDM4E | 0.021989 | 0.693801 | Other Positive Correlation |
| SSU72P8 | 0.021989 | 0.693801 | Other Positive Correlation |
| BICD1 | 0.021976 | 0.693975 | Other Positive Correlation |
| NCF4 | 0.021961 | 0.694165 | Other Positive Correlation |
| MYO3B | 0.021952 | 0.694288 | Other Positive Correlation |
| PHF8 | 0.021949 | 0.694333 | Other Positive Correlation |
| TMEM243 | 0.021932 | 0.694556 | Other Positive Correlation |
| TEK | 0.021926 | 0.694634 | Other Positive Correlation |
| MYL4 | 0.021925 | 0.69464 | Other Positive Correlation |
| SLIT3 | 0.02192 | 0.694709 | Other Positive Correlation |
| WASHC2A | 0.0219 | 0.69498 | Other Positive Correlation |
| NIPSNAP3B | 0.021893 | 0.695062 | Other Positive Correlation |
| MAPK11 | 0.021759 | 0.69684 | Other Positive Correlation |
| ZNF717 | 0.021759 | 0.696848 | Other Positive Correlation |
| TMEM178B | 0.021757 | 0.696874 | Other Positive Correlation |
| GFOD2 | 0.021729 | 0.697233 | Other Positive Correlation |
| SPRED2 | 0.021709 | 0.697501 | Other Positive Correlation |
| ZNF3 | 0.021693 | 0.697718 | Other Positive Correlation |
| TNPO3 | 0.021676 | 0.697948 | Other Positive Correlation |
| IL15 | 0.021635 | 0.698487 | Other Positive Correlation |
| EPHX2 | 0.021598 | 0.698972 | Other Positive Correlation |
| CCL26 | 0.021596 | 0.698999 | Other Positive Correlation |
| RAB5A | 0.021575 | 0.699286 | Other Positive Correlation |
| UFM1 | 0.021572 | 0.699317 | Other Positive Correlation |
| DAPK3 | 0.021542 | 0.699717 | Other Positive Correlation |
| PIK3C2G | 0.021527 | 0.699915 | Other Positive Correlation |
| GPR20 | 0.021503 | 0.700229 | Other Positive Correlation |
| DPY19L2 | 0.021474 | 0.700622 | Other Positive Correlation |
| ENPP2 | 0.021467 | 0.700718 | Other Positive Correlation |
| ZNF296 | 0.021403 | 0.701563 | Other Positive Correlation |
| PREPL | 0.021384 | 0.70181 | Other Positive Correlation |
| TMEM115 | 0.021371 | 0.701984 | Other Positive Correlation |
| EPN2 | 0.021365 | 0.702063 | Other Positive Correlation |
| KIAA1755 | 0.021346 | 0.702318 | Other Positive Correlation |
| SLC45A2 | 0.02131 | 0.702805 | Other Positive Correlation |
| SLC46A1 | 0.021301 | 0.702918 | Other Positive Correlation |
| CEP152 | 0.021298 | 0.702963 | Other Positive Correlation |
| ABI3 | 0.021291 | 0.703057 | Other Positive Correlation |
| TRADD | 0.021282 | 0.70317 | Other Positive Correlation |
| COG2 | 0.021251 | 0.703581 | Other Positive Correlation |

| | | | |
|---|---|---|---|
| BCAS1 | 0.021228 | 0.703885 | Other Positive Correlation |
| BAHD1 | 0.021198 | 0.704286 | Other Positive Correlation |
| ACSM3 | 0.021198 | 0.704286 | Other Positive Correlation |
| IKBKG | 0.021165 | 0.704732 | Other Positive Correlation |
| PTRHD1 | 0.021147 | 0.704964 | Other Positive Correlation |
| UHMK1 | 0.021126 | 0.705248 | Other Positive Correlation |
| ABCC10 | 0.021097 | 0.705627 | Other Positive Correlation |
| VWA7 | 0.021091 | 0.705708 | Other Positive Correlation |
| WASHC5 | 0.021089 | 0.705738 | Other Positive Correlation |
| LCE5A | 0.021082 | 0.705824 | Other Positive Correlation |
| TEKT2 | 0.021078 | 0.705882 | Other Positive Correlation |
| MRPS18C | 0.021031 | 0.706504 | Other Positive Correlation |
| VPS4A | 0.021021 | 0.706637 | Other Positive Correlation |
| MTG2 | 0.021002 | 0.706895 | Other Positive Correlation |
| ASB3 | 0.020982 | 0.707162 | Other Positive Correlation |
| RRAD | 0.020969 | 0.707333 | Other Positive Correlation |
| TOGARAM2 | 0.020955 | 0.707522 | Other Positive Correlation |
| CYP24A1 | 0.020938 | 0.70774 | Other Positive Correlation |
| RNF40 | 0.020925 | 0.707916 | Other Positive Correlation |
| ISM2 | 0.020911 | 0.708109 | Other Positive Correlation |
| ZNF622 | 0.020906 | 0.708176 | Other Positive Correlation |
| ZNF212 | 0.020906 | 0.708179 | Other Positive Correlation |
| CENPX | 0.020888 | 0.70841 | Other Positive Correlation |
| MEIOSIN | 0.020886 | 0.708445 | Other Positive Correlation |
| TMEM204 | 0.020835 | 0.70912 | Other Positive Correlation |
| DHTKD1 | 0.020821 | 0.7093 | Other Positive Correlation |
| PNMA8C | 0.020775 | 0.70992 | Other Positive Correlation |
| PDE7B | 0.02076 | 0.710118 | Other Positive Correlation |
| PTPRA | 0.020757 | 0.710161 | Other Positive Correlation |
| MOGAT3 | 0.020754 | 0.710203 | Other Positive Correlation |
| NPFFR1 | 0.020754 | 0.710203 | Other Positive Correlation |
| BIN2 | 0.020754 | 0.710203 | Other Positive Correlation |
| NAP1L4 | 0.020752 | 0.71023 | Other Positive Correlation |
| RINT1 | 0.020751 | 0.71024 | Other Positive Correlation |
| XKRX | 0.020738 | 0.71041 | Other Positive Correlation |
| IKZF2 | 0.020724 | 0.710604 | Other Positive Correlation |
| ZNF112 | 0.020717 | 0.710692 | Other Positive Correlation |
| LAG3 | 0.020712 | 0.710762 | Other Positive Correlation |
| KCNH8 | 0.020678 | 0.711209 | Other Positive Correlation |
| VTA1 | 0.020666 | 0.711368 | Other Positive Correlation |
| RNF25 | 0.020652 | 0.711556 | Other Positive Correlation |
| ERFL | 0.020607 | 0.712165 | Other Positive Correlation |
| SYNJ2BP | 0.020597 | 0.712288 | Other Positive Correlation |
| CLTRN | 0.020574 | 0.7126 | Other Positive Correlation |
| TSNARE1 | 0.020556 | 0.71284 | Other Positive Correlation |
| PIFO | 0.020547 | 0.712964 | Other Positive Correlation |
| EAPP | 0.02053 | 0.713184 | Other Positive Correlation |
| NCOA2 | 0.020523 | 0.713278 | Other Positive Correlation |

| | | | |
|---|---|---|---|
| MYOM1 | 0.020498 | 0.713619 | Other Positive Correlation |
| ZNF442 | 0.020494 | 0.713667 | Other Positive Correlation |
| FABP6 | 0.020479 | 0.713872 | Other Positive Correlation |
| C2orf73 | 0.020449 | 0.714272 | Other Positive Correlation |
| MAP4K1 | 0.020439 | 0.714403 | Other Positive Correlation |
| TACR2 | 0.020435 | 0.714455 | Other Positive Correlation |
| TNNT1 | 0.020418 | 0.71469 | Other Positive Correlation |
| ENSG00000261832 | 0.020405 | 0.714852 | Other Positive Correlation |
| ELL3 | 0.020387 | 0.715098 | Other Positive Correlation |
| CWC27 | 0.020362 | 0.715428 | Other Positive Correlation |
| C1QTNF3 | 0.020338 | 0.715748 | Other Positive Correlation |
| GRIN3B | 0.02033 | 0.715862 | Other Positive Correlation |
| RIPOR2 | 0.020305 | 0.716195 | Other Positive Correlation |
| CCR7 | 0.020299 | 0.716282 | Other Positive Correlation |
| IRF2BPL | 0.020292 | 0.716369 | Other Positive Correlation |
| RORC | 0.020257 | 0.716834 | Other Positive Correlation |
| MOSPD3 | 0.020208 | 0.717491 | Other Positive Correlation |
| PPBP | 0.020202 | 0.717569 | Other Positive Correlation |
| TUBB2A | 0.020194 | 0.717683 | Other Positive Correlation |
| PIP5KL1 | 0.020187 | 0.717772 | Other Positive Correlation |
| C4orf47 | 0.020185 | 0.717797 | Other Positive Correlation |
| CCDC116 | 0.020178 | 0.717895 | Other Positive Correlation |
| FRMD8 | 0.020124 | 0.718615 | Other Positive Correlation |
| PXDNL | 0.020113 | 0.718772 | Other Positive Correlation |
| TRIM43B | 0.020078 | 0.71924 | Other Positive Correlation |
| TAF11L14 | 0.020078 | 0.71924 | Other Positive Correlation |
| BRINP2 | 0.020078 | 0.71924 | Other Positive Correlation |
| ENSG00000283321 | 0.020078 | 0.71924 | Other Positive Correlation |
| TAF11L13 | 0.020078 | 0.71924 | Other Positive Correlation |
| CD8B2 | 0.020078 | 0.71924 | Other Positive Correlation |
| HMSD | 0.020064 | 0.719416 | Other Positive Correlation |
| GNG11 | 0.020057 | 0.719509 | Other Positive Correlation |
| APOBR | 0.020057 | 0.719516 | Other Positive Correlation |
| ZNF829 | 0.020043 | 0.719697 | Other Positive Correlation |
| FBXL12 | 0.020013 | 0.720104 | Other Positive Correlation |
| RAET1L | 0.020008 | 0.720177 | Other Positive Correlation |
| GUCA2B | 0.02 | 0.720277 | Other Positive Correlation |
| ANAPC2 | 0.019991 | 0.720398 | Other Positive Correlation |
| OTOF | 0.019964 | 0.720759 | Other Positive Correlation |
| MAPRE2 | 0.019957 | 0.720859 | Other Positive Correlation |
| TTC7B | 0.019956 | 0.720866 | Other Positive Correlation |
| C8orf44-SGK3 | 0.019954 | 0.720889 | Other Positive Correlation |
| C11orf86 | 0.019922 | 0.721319 | Other Positive Correlation |
| ENSG00000268279 | 0.019907 | 0.721524 | Other Positive Correlation |
| PRSS33 | 0.019906 | 0.721544 | Other Positive Correlation |
| C5 | 0.019891 | 0.721735 | Other Positive Correlation |
| PRAM1 | 0.019869 | 0.722041 | Other Positive Correlation |
| PTPN23 | 0.019862 | 0.722124 | Other Positive Correlation |

| | | | |
|---|---|---|---|
| ENSG00000288721 | 0.019825 | 0.722621 | Other Positive Correlation |
| ALPP | 0.019799 | 0.722976 | Other Positive Correlation |
| KIF1B | 0.019763 | 0.72346 | Other Positive Correlation |
| DCTN5 | 0.019758 | 0.723529 | Other Positive Correlation |
| NCKAP1L | 0.019747 | 0.723667 | Other Positive Correlation |
| RCAN1 | 0.019741 | 0.723754 | Other Positive Correlation |
| ZBTB41 | 0.019696 | 0.724361 | Other Positive Correlation |
| PHOSPHO2 | 0.019668 | 0.72473 | Other Positive Correlation |
| GSTA1 | 0.019639 | 0.72512 | Other Positive Correlation |
| CDH18 | 0.019639 | 0.72512 | Other Positive Correlation |
| GRIK4 | 0.019628 | 0.725267 | Other Positive Correlation |
| CALCR | 0.019626 | 0.725297 | Other Positive Correlation |
| CTXND1 | 0.019611 | 0.725504 | Other Positive Correlation |
| TAC3 | 0.019611 | 0.725504 | Other Positive Correlation |
| TRMT1L | 0.019605 | 0.725573 | Other Positive Correlation |
| GPHA2 | 0.019573 | 0.726009 | Other Positive Correlation |
| H2BC26 | 0.019536 | 0.726501 | Other Positive Correlation |
| VAMP5 | 0.019536 | 0.726503 | Other Positive Correlation |
| GLIS1 | 0.019535 | 0.726513 | Other Positive Correlation |
| PMM1 | 0.019534 | 0.726539 | Other Positive Correlation |
| FAM117B | 0.019509 | 0.72687 | Other Positive Correlation |
| CYTH4 | 0.019451 | 0.727645 | Other Positive Correlation |
| DUSP10 | 0.019444 | 0.727741 | Other Positive Correlation |
| CLRN3 | 0.019411 | 0.728184 | Other Positive Correlation |
| CHST11 | 0.01937 | 0.728742 | Other Positive Correlation |
| NLK | 0.019363 | 0.728833 | Other Positive Correlation |
| RNMT | 0.019308 | 0.729579 | Other Positive Correlation |
| LMBRD2 | 0.019299 | 0.729689 | Other Positive Correlation |
| ZNF525 | 0.019262 | 0.730187 | Other Positive Correlation |
| TMPRSS11E | 0.019262 | 0.730196 | Other Positive Correlation |
| SNX15 | 0.019242 | 0.730465 | Other Positive Correlation |
| CEP89 | 0.019241 | 0.730472 | Other Positive Correlation |
| SP7 | 0.019208 | 0.730923 | Other Positive Correlation |
| DPP8 | 0.019204 | 0.730967 | Other Positive Correlation |
| FMO1 | 0.019155 | 0.731626 | Other Positive Correlation |
| RNGTT | 0.019153 | 0.731655 | Other Positive Correlation |
| PARVG | 0.019134 | 0.73191 | Other Positive Correlation |
| TRPM2 | 0.019133 | 0.731935 | Other Positive Correlation |
| NOTCH2 | 0.019109 | 0.732246 | Other Positive Correlation |
| PAOX | 0.019086 | 0.732567 | Other Positive Correlation |
| TSSK3 | 0.019083 | 0.732604 | Other Positive Correlation |
| NAT2 | 0.019055 | 0.732983 | Other Positive Correlation |
| PERM1 | 0.019023 | 0.733414 | Other Positive Correlation |
| EIF1AY | 0.019015 | 0.733522 | Other Positive Correlation |
| ZC3H12C | 0.019013 | 0.733542 | Other Positive Correlation |
| MAML3 | 0.018962 | 0.734234 | Other Positive Correlation |
| WDR88 | 0.018916 | 0.734854 | Other Positive Correlation |
| DEFB1 | 0.018902 | 0.735047 | Other Positive Correlation |

| | | | |
|---|---|---|---|
| ITCH | 0.018874 | 0.735416 | Other Positive Correlation |
| CSK | 0.018853 | 0.735703 | Other Positive Correlation |
| PEG10 | 0.018826 | 0.736071 | Other Positive Correlation |
| STAT1 | 0.018816 | 0.736204 | Other Positive Correlation |
| MUC5AC | 0.018807 | 0.736324 | Other Positive Correlation |
| RNF17 | 0.018789 | 0.736567 | Other Positive Correlation |
| ANO3 | 0.018769 | 0.736834 | Other Positive Correlation |
| CHRNE | 0.018765 | 0.736891 | Other Positive Correlation |
| ITGB7 | 0.018765 | 0.736894 | Other Positive Correlation |
| PRSS35 | 0.018753 | 0.73705 | Other Positive Correlation |
| ABHD2 | 0.018745 | 0.73716 | Other Positive Correlation |
| SCN11A | 0.018707 | 0.737674 | Other Positive Correlation |
| MBD4 | 0.018704 | 0.737723 | Other Positive Correlation |
| USP54 | 0.01869 | 0.737905 | Other Positive Correlation |
| MAD1L1 | 0.018688 | 0.737935 | Other Positive Correlation |
| CDH19 | 0.018668 | 0.7382 | Other Positive Correlation |
| CCL24 | 0.018659 | 0.738321 | Other Positive Correlation |
| COMMD3-BMI1 | 0.018658 | 0.738334 | Other Positive Correlation |
| HGFAC | 0.018619 | 0.738864 | Other Positive Correlation |
| FAM181A | 0.018615 | 0.738916 | Other Positive Correlation |
| CFAP52 | 0.018603 | 0.739078 | Other Positive Correlation |
| BHMT | 0.018594 | 0.7392 | Other Positive Correlation |
| FOXO1 | 0.018581 | 0.739383 | Other Positive Correlation |
| MEOX1 | 0.018563 | 0.739625 | Other Positive Correlation |
| ENSG00000258989 | 0.018561 | 0.739647 | Other Positive Correlation |
| PFKFB3 | 0.018549 | 0.739815 | Other Positive Correlation |
| MEF2C | 0.018534 | 0.74001 | Other Positive Correlation |
| KCNA2 | 0.0185 | 0.740472 | Other Positive Correlation |
| KCND2 | 0.018471 | 0.740869 | Other Positive Correlation |
| DNAH14 | 0.018468 | 0.740908 | Other Positive Correlation |
| DYNLRB1 | 0.018467 | 0.740927 | Other Positive Correlation |
| ADAM30 | 0.018446 | 0.74121 | Other Positive Correlation |
| SYNPO2L | 0.018437 | 0.741332 | Other Positive Correlation |
| H4C14 | 0.018417 | 0.741599 | Other Positive Correlation |
| P4HA3 | 0.018414 | 0.74164 | Other Positive Correlation |
| GALNT15 | 0.018407 | 0.741732 | Other Positive Correlation |
| C11orf52 | 0.01839 | 0.741959 | Other Positive Correlation |
| GRIK2 | 0.018389 | 0.741982 | Other Positive Correlation |
| ZNF557 | 0.018387 | 0.742006 | Other Positive Correlation |
| PLAAT2 | 0.018373 | 0.742198 | Other Positive Correlation |
| HCG22 | 0.01836 | 0.742368 | Other Positive Correlation |
| CRNN | 0.018318 | 0.742934 | Other Positive Correlation |
| TMEM52B | 0.018318 | 0.742935 | Other Positive Correlation |
| IRS4 | 0.018316 | 0.742967 | Other Positive Correlation |
| MAN2B2 | 0.018274 | 0.743528 | Other Positive Correlation |
| POMT2 | 0.018264 | 0.743673 | Other Positive Correlation |
| APC | 0.018263 | 0.743687 | Other Positive Correlation |
| MTIF3 | 0.018256 | 0.74378 | Other Positive Correlation |

| | | | |
|---|---|---|---|
| MCF2L2 | 0.018222 | 0.744235 | Other Positive Correlation |
| MT-ND4L | 0.01822 | 0.744262 | Other Positive Correlation |
| LCE3D | 0.018192 | 0.74465 | Other Positive Correlation |
| TSC1 | 0.018181 | 0.744799 | Other Positive Correlation |
| PATL1 | 0.018173 | 0.744901 | Other Positive Correlation |
| SERPINA6 | 0.018162 | 0.745049 | Other Positive Correlation |
| DENND1B | 0.018153 | 0.74517 | Other Positive Correlation |
| ARL15 | 0.018152 | 0.745182 | Other Positive Correlation |
| MFAP2 | 0.018124 | 0.745572 | Other Positive Correlation |
| OOEP | 0.018117 | 0.745658 | Other Positive Correlation |
| CPT2 | 0.018109 | 0.745769 | Other Positive Correlation |
| M1AP | 0.018087 | 0.746064 | Other Positive Correlation |
| MRGPRE | 0.018074 | 0.74624 | Other Positive Correlation |
| EPYC | 0.018074 | 0.74624 | Other Positive Correlation |
| STC1 | 0.018065 | 0.746365 | Other Positive Correlation |
| DBX2 | 0.018003 | 0.747212 | Other Positive Correlation |
| RLN3 | 0.018003 | 0.747212 | Other Positive Correlation |
| HOXD13 | 0.018003 | 0.747212 | Other Positive Correlation |
| ZNF679 | 0.018003 | 0.747212 | Other Positive Correlation |
| HOXC10 | 0.018003 | 0.747212 | Other Positive Correlation |
| NRSN1 | 0.017991 | 0.747369 | Other Positive Correlation |
| STUM | 0.017991 | 0.747369 | Other Positive Correlation |
| GOLGA6L9 | 0.017964 | 0.747737 | Other Positive Correlation |
| GRK5 | 0.017943 | 0.748023 | Other Positive Correlation |
| PEDS1 | 0.017928 | 0.748221 | Other Positive Correlation |
| FAM131B | 0.017922 | 0.748311 | Other Positive Correlation |
| TTR | 0.017904 | 0.748555 | Other Positive Correlation |
| POGLUT3 | 0.017899 | 0.748623 | Other Positive Correlation |
| PLA2G4D | 0.017889 | 0.748758 | Other Positive Correlation |
| HLA-DMA | 0.017859 | 0.749158 | Other Positive Correlation |
| CSF2 | 0.017859 | 0.749163 | Other Positive Correlation |
| NCSTN | 0.017842 | 0.749385 | Other Positive Correlation |
| RBM23 | 0.017823 | 0.749648 | Other Positive Correlation |
| HFM1 | 0.017821 | 0.749682 | Other Positive Correlation |
| LMF1 | 0.017792 | 0.750067 | Other Positive Correlation |
| ZNF426 | 0.017784 | 0.750179 | Other Positive Correlation |
| DHPS | 0.017767 | 0.750413 | Other Positive Correlation |
| S1PR4 | 0.017746 | 0.750699 | Other Positive Correlation |
| CWC15 | 0.017671 | 0.751715 | Other Positive Correlation |
| SLC22A23 | 0.017649 | 0.752007 | Other Positive Correlation |
| BTRC | 0.01764 | 0.752135 | Other Positive Correlation |
| TEX35 | 0.017629 | 0.752286 | Other Positive Correlation |
| S100B | 0.017617 | 0.752442 | Other Positive Correlation |
| MARK2 | 0.017614 | 0.752494 | Other Positive Correlation |
| H1-0 | 0.017588 | 0.752836 | Other Positive Correlation |
| PGGT1B | 0.017575 | 0.75302 | Other Positive Correlation |
| HMGCLL1 | 0.017569 | 0.753105 | Other Positive Correlation |
| STMN4 | 0.017549 | 0.753369 | Other Positive Correlation |

| | | | |
|---|---|---|---|
| OLR1 | 0.017536 | 0.753548 | Other Positive Correlation |
| SF3B5 | 0.017501 | 0.754025 | Other Positive Correlation |
| ALDH9A1 | 0.017461 | 0.754567 | Other Positive Correlation |
| HEXIM2 | 0.017457 | 0.754619 | Other Positive Correlation |
| GREP1 | 0.017449 | 0.754736 | Other Positive Correlation |
| ZNF256 | 0.017432 | 0.754969 | Other Positive Correlation |
| HUWE1 | 0.017412 | 0.755232 | Other Positive Correlation |
| PAPOLB | 0.017411 | 0.755245 | Other Positive Correlation |
| ANO1 | 0.017376 | 0.755723 | Other Positive Correlation |
| VAV3 | 0.017372 | 0.75578 | Other Positive Correlation |
| ENSG00000276418 | 0.017371 | 0.755791 | Other Positive Correlation |
| EDRF1 | 0.017359 | 0.755963 | Other Positive Correlation |
| KHDRBS2 | 0.017355 | 0.756007 | Other Positive Correlation |
| HEPACAM | 0.017355 | 0.756007 | Other Positive Correlation |
| F2RL2 | 0.017352 | 0.756056 | Other Positive Correlation |
| EXOC6 | 0.01734 | 0.75622 | Other Positive Correlation |
| GRIN2D | 0.017332 | 0.756322 | Other Positive Correlation |
| CD83 | 0.017287 | 0.756939 | Other Positive Correlation |
| SLC18B1 | 0.017284 | 0.756977 | Other Positive Correlation |
| PTGFR | 0.017278 | 0.757055 | Other Positive Correlation |
| F2RL3 | 0.017266 | 0.757229 | Other Positive Correlation |
| PPP4C | 0.017247 | 0.757482 | Other Positive Correlation |
| SMAD7 | 0.017233 | 0.75767 | Other Positive Correlation |
| AP5Z1 | 0.017231 | 0.757708 | Other Positive Correlation |
| TRPV2 | 0.017225 | 0.757777 | Other Positive Correlation |
| NUMBL | 0.017198 | 0.758148 | Other Positive Correlation |
| SCAMP3 | 0.017169 | 0.758549 | Other Positive Correlation |
| LDAH | 0.01713 | 0.75908 | Other Positive Correlation |
| ZNF843 | 0.017126 | 0.759136 | Other Positive Correlation |
| PAGE2B | 0.017113 | 0.759315 | Other Positive Correlation |
| RILP | 0.017094 | 0.759564 | Other Positive Correlation |
| PJVK | 0.017067 | 0.759932 | Other Positive Correlation |
| PIGZ | 0.017036 | 0.760357 | Other Positive Correlation |
| SFT2D3 | 0.017032 | 0.760408 | Other Positive Correlation |
| ENSG00000267706 | 0.017021 | 0.76057 | Other Positive Correlation |
| ENSG00000268400 | 0.017015 | 0.760639 | Other Positive Correlation |
| C11orf91 | 0.017001 | 0.760835 | Other Positive Correlation |
| RPRML | 0.016998 | 0.76088 | Other Positive Correlation |
| RAB6C | 0.016998 | 0.76088 | Other Positive Correlation |
| CHRM2 | 0.016998 | 0.76088 | Other Positive Correlation |
| HOXB13 | 0.016998 | 0.76088 | Other Positive Correlation |
| HAND2 | 0.016998 | 0.76088 | Other Positive Correlation |
| NEU4 | 0.016998 | 0.76088 | Other Positive Correlation |
| ZNF688 | 0.016974 | 0.761208 | Other Positive Correlation |
| GOLGA7 | 0.016973 | 0.76122 | Other Positive Correlation |
| AMDHD1 | 0.016971 | 0.761243 | Other Positive Correlation |
| CCDC112 | 0.016966 | 0.761313 | Other Positive Correlation |
| CLDN12 | 0.016931 | 0.761786 | Other Positive Correlation |

| | | | |
|---|---|---|---|
| ABCG2 | 0.016875 | 0.762552 | Other Positive Correlation |
| MPHOSPH8 | 0.016836 | 0.763088 | Other Positive Correlation |
| CATSPERG | 0.016823 | 0.76326 | Other Positive Correlation |
| UPF3A | 0.016817 | 0.763341 | Other Positive Correlation |
| SPSB2 | 0.016816 | 0.763355 | Other Positive Correlation |
| CASTOR1 | 0.016801 | 0.763564 | Other Positive Correlation |
| THEG | 0.0168 | 0.763582 | Other Positive Correlation |
| LRATD1 | 0.016784 | 0.763802 | Other Positive Correlation |
| ENSG00000288698 | 0.016773 | 0.763949 | Other Positive Correlation |
| TMPRSS12 | 0.01677 | 0.763992 | Other Positive Correlation |
| NETO2 | 0.016715 | 0.764742 | Other Positive Correlation |
| EGR3 | 0.016706 | 0.764857 | Other Positive Correlation |
| EPB41 | 0.016701 | 0.764931 | Other Positive Correlation |
| PRR20G | 0.0167 | 0.764945 | Other Positive Correlation |
| TTC33 | 0.016674 | 0.765297 | Other Positive Correlation |
| ADGRG3 | 0.016666 | 0.765416 | Other Positive Correlation |
| SUCO | 0.01663 | 0.765904 | Other Positive Correlation |
| CARD14 | 0.016608 | 0.766205 | Other Positive Correlation |
| FKBP6 | 0.016599 | 0.766323 | Other Positive Correlation |
| FBXO41 | 0.016572 | 0.766698 | Other Positive Correlation |
| NPAS1 | 0.016566 | 0.766782 | Other Positive Correlation |
| HHIPL2 | 0.016557 | 0.766904 | Other Positive Correlation |
| FAM110C | 0.016543 | 0.767096 | Other Positive Correlation |
| NLGN4X | 0.016459 | 0.768243 | Other Positive Correlation |
| ENSG00000284773 | 0.016445 | 0.768425 | Other Positive Correlation |
| IL21R | 0.016438 | 0.768531 | Other Positive Correlation |
| NSD1 | 0.016421 | 0.768763 | Other Positive Correlation |
| PLAAT5 | 0.016417 | 0.768816 | Other Positive Correlation |
| TAS2R7 | 0.016417 | 0.768816 | Other Positive Correlation |
| CCDC127 | 0.016401 | 0.76904 | Other Positive Correlation |
| ELF2 | 0.016397 | 0.76909 | Other Positive Correlation |
| LRTM1 | 0.016388 | 0.769218 | Other Positive Correlation |
| LRP10 | 0.016383 | 0.769281 | Other Positive Correlation |
| CPA1 | 0.016378 | 0.769344 | Other Positive Correlation |
| MIER3 | 0.016378 | 0.769352 | Other Positive Correlation |
| ZNF385C | 0.016365 | 0.769523 | Other Positive Correlation |
| XNDC1N | 0.016364 | 0.769534 | Other Positive Correlation |
| UBASH3B | 0.016354 | 0.769679 | Other Positive Correlation |
| FKBP1B | 0.016346 | 0.769788 | Other Positive Correlation |
| PAX1 | 0.01633 | 0.770002 | Other Positive Correlation |
| NOSIP | 0.01632 | 0.770138 | Other Positive Correlation |
| FGD1 | 0.016319 | 0.770157 | Other Positive Correlation |
| SLC2A14 | 0.016307 | 0.770322 | Other Positive Correlation |
| CACNA1E | 0.01629 | 0.77055 | Other Positive Correlation |
| TEX14 | 0.016285 | 0.770626 | Other Positive Correlation |
| ZNF233 | 0.016218 | 0.771543 | Other Positive Correlation |
| EFNA4 | 0.016217 | 0.771559 | Other Positive Correlation |
| DNAL4 | 0.016212 | 0.771619 | Other Positive Correlation |

| | | | |
|---|---|---|---|
| THBS4 | 0.016188 | 0.771956 | Other Positive Correlation |
| SPINDOC | 0.016186 | 0.771972 | Other Positive Correlation |
| CYP2C18 | 0.016182 | 0.772035 | Other Positive Correlation |
| CLN8 | 0.016152 | 0.772437 | Other Positive Correlation |
| FFAR2 | 0.016122 | 0.772858 | Other Positive Correlation |
| PCDHA13 | 0.016121 | 0.772875 | Other Positive Correlation |
| ENSG00000269476 | 0.016121 | 0.772875 | Other Positive Correlation |
| ASIP | 0.016121 | 0.772875 | Other Positive Correlation |
| MLLT1 | 0.016118 | 0.772912 | Other Positive Correlation |
| LRRN3 | 0.016117 | 0.772929 | Other Positive Correlation |
| ATP10B | 0.016087 | 0.773336 | Other Positive Correlation |
| ADCY6 | 0.016081 | 0.773419 | Other Positive Correlation |
| SPTSSB | 0.016061 | 0.773688 | Other Positive Correlation |
| RANBP10 | 0.01603 | 0.774119 | Other Positive Correlation |
| MEX3A | 0.016004 | 0.774477 | Other Positive Correlation |
| ZNF605 | 0.015992 | 0.774635 | Other Positive Correlation |
| GABARAP | 0.015974 | 0.774883 | Other Positive Correlation |
| ATF7 | 0.015941 | 0.775337 | Other Positive Correlation |
| ITM2A | 0.015896 | 0.775952 | Other Positive Correlation |
| HDAC5 | 0.01586 | 0.776444 | Other Positive Correlation |
| OVGP1 | 0.015851 | 0.776577 | Other Positive Correlation |
| TGFB3 | 0.015826 | 0.776913 | Other Positive Correlation |
| TMSB15B | 0.015825 | 0.776922 | Other Positive Correlation |
| DEXI | 0.015817 | 0.77704 | Other Positive Correlation |
| SEPTIN12 | 0.015805 | 0.7772 | Other Positive Correlation |
| CCDC190 | 0.015805 | 0.7772 | Other Positive Correlation |
| SSMEM1 | 0.015805 | 0.7772 | Other Positive Correlation |
| HSPB9 | 0.01579 | 0.777408 | Other Positive Correlation |
| TCN1 | 0.015765 | 0.777748 | Other Positive Correlation |
| NEFM | 0.015732 | 0.778204 | Other Positive Correlation |
| TTC36 | 0.015707 | 0.778545 | Other Positive Correlation |
| AOAH | 0.015698 | 0.778669 | Other Positive Correlation |
| RIC8B | 0.015694 | 0.778726 | Other Positive Correlation |
| HIVEP2 | 0.015683 | 0.778884 | Other Positive Correlation |
| CCP110 | 0.015625 | 0.779672 | Other Positive Correlation |
| OR3A2 | 0.015595 | 0.780091 | Other Positive Correlation |
| ZNF554 | 0.015575 | 0.780363 | Other Positive Correlation |
| FAM210A | 0.015539 | 0.780855 | Other Positive Correlation |
| SULT1C2 | 0.015505 | 0.781326 | Other Positive Correlation |
| FAM90A1 | 0.015477 | 0.781706 | Other Positive Correlation |
| DYNLT1 | 0.015458 | 0.781971 | Other Positive Correlation |
| CYP26B1 | 0.015435 | 0.78229 | Other Positive Correlation |
| PDE3B | 0.015434 | 0.782304 | Other Positive Correlation |
| JADE2 | 0.01543 | 0.782359 | Other Positive Correlation |
| ZNF750 | 0.015351 | 0.783439 | Other Positive Correlation |
| APOM | 0.015344 | 0.783537 | Other Positive Correlation |
| DPEP2NB | 0.015314 | 0.783952 | Other Positive Correlation |
| GRID2 | 0.015288 | 0.784308 | Other Positive Correlation |

| | | | |
|---|---|---|---|
| ADGRG7 | 0.015288 | 0.784308 | Other Positive Correlation |
| SPANXC | 0.015288 | 0.784308 | Other Positive Correlation |
| DMP1 | 0.015288 | 0.784308 | Other Positive Correlation |
| C4orf54 | 0.015288 | 0.784308 | Other Positive Correlation |
| SPANXA1 | 0.015288 | 0.784308 | Other Positive Correlation |
| PRDM9 | 0.015288 | 0.784308 | Other Positive Correlation |
| ARHGEF38 | 0.015288 | 0.784308 | Other Positive Correlation |
| CDH20 | 0.015288 | 0.784308 | Other Positive Correlation |
| AMBN | 0.015288 | 0.784308 | Other Positive Correlation |
| GIMD1 | 0.015288 | 0.784308 | Other Positive Correlation |
| KEL | 0.015288 | 0.784308 | Other Positive Correlation |
| EMCN | 0.015288 | 0.784308 | Other Positive Correlation |
| SPANXB1 | 0.015288 | 0.784308 | Other Positive Correlation |
| TNKS | 0.015277 | 0.784454 | Other Positive Correlation |
| PCDHAC1 | 0.015272 | 0.784523 | Other Positive Correlation |
| LRCH1 | 0.01527 | 0.784557 | Other Positive Correlation |
| VRTN | 0.01526 | 0.784693 | Other Positive Correlation |
| JAML | 0.01526 | 0.784694 | Other Positive Correlation |
| GNRH2 | 0.01526 | 0.784694 | Other Positive Correlation |
| PRAMEF14 | 0.015247 | 0.784868 | Other Positive Correlation |
| CAPN9 | 0.015235 | 0.78504 | Other Positive Correlation |
| CASC3 | 0.01522 | 0.785246 | Other Positive Correlation |
| RNF151 | 0.015206 | 0.785432 | Other Positive Correlation |
| C20orf96 | 0.0152 | 0.785525 | Other Positive Correlation |
| FBXO7 | 0.015193 | 0.785621 | Other Positive Correlation |
| CA6 | 0.01519 | 0.785654 | Other Positive Correlation |
| TNFAIP8L2 | 0.01519 | 0.785654 | Other Positive Correlation |
| OR2A5 | 0.01519 | 0.785654 | Other Positive Correlation |
| GDA | 0.015189 | 0.785669 | Other Positive Correlation |
| CST9L | 0.015179 | 0.785811 | Other Positive Correlation |
| OR7D2 | 0.015179 | 0.785811 | Other Positive Correlation |
| COL20A1 | 0.015179 | 0.785811 | Other Positive Correlation |
| ADAMTS18 | 0.015179 | 0.785811 | Other Positive Correlation |
| FGFBP2 | 0.015179 | 0.785811 | Other Positive Correlation |
| MAOB | 0.015172 | 0.785906 | Other Positive Correlation |
| SYT15 | 0.01516 | 0.786065 | Other Positive Correlation |
| ERP27 | 0.015159 | 0.786077 | Other Positive Correlation |
| ABCA8 | 0.015149 | 0.786222 | Other Positive Correlation |
| LILRB1 | 0.015144 | 0.786295 | Other Positive Correlation |
| HGS | 0.01512 | 0.786616 | Other Positive Correlation |
| WNK4 | 0.015081 | 0.787163 | Other Positive Correlation |
| SRBD1 | 0.015054 | 0.787525 | Other Positive Correlation |
| CDS1 | 0.015011 | 0.788122 | Other Positive Correlation |
| NR1D1 | 0.014934 | 0.789183 | Other Positive Correlation |
| ARHGAP10 | 0.01493 | 0.789238 | Other Positive Correlation |
| C2orf49 | 0.014916 | 0.78943 | Other Positive Correlation |
| PTGER1 | 0.014903 | 0.789612 | Other Positive Correlation |
| BAG4 | 0.0149 | 0.789658 | Other Positive Correlation |

| | | | |
|---|---|---|---|
| PRSS1 | 0.01488 | 0.789929 | Other Positive Correlation |
| RDH8 | 0.01488 | 0.789929 | Other Positive Correlation |
| HEY1 | 0.014875 | 0.789993 | Other Positive Correlation |
| SPDYC | 0.014872 | 0.790033 | Other Positive Correlation |
| SMIM18 | 0.014872 | 0.790033 | Other Positive Correlation |
| CLDN17 | 0.014872 | 0.790033 | Other Positive Correlation |
| TSPAN32 | 0.014872 | 0.790033 | Other Positive Correlation |
| GPR17 | 0.014872 | 0.790033 | Other Positive Correlation |
| RXFP4 | 0.014872 | 0.790033 | Other Positive Correlation |
| CACNG2 | 0.014872 | 0.790033 | Other Positive Correlation |
| ENSG00000249319 | 0.014872 | 0.790033 | Other Positive Correlation |
| TNFRSF19 | 0.014868 | 0.790088 | Other Positive Correlation |
| ATP6V0D2 | 0.014842 | 0.790455 | Other Positive Correlation |
| SIGLEC6 | 0.014838 | 0.790501 | Other Positive Correlation |
| IRAG2 | 0.014838 | 0.790501 | Other Positive Correlation |
| POU2AF2 | 0.014838 | 0.790501 | Other Positive Correlation |
| KRTAP1-4 | 0.014838 | 0.790501 | Other Positive Correlation |
| CLEC4D | 0.014838 | 0.790501 | Other Positive Correlation |
| KRTAP3-3 | 0.014838 | 0.790501 | Other Positive Correlation |
| SAMD7 | 0.014838 | 0.790501 | Other Positive Correlation |
| TBPL2 | 0.014838 | 0.790501 | Other Positive Correlation |
| KRTAP2-1 | 0.014838 | 0.790501 | Other Positive Correlation |
| KRT38 | 0.014838 | 0.790501 | Other Positive Correlation |
| DIO3 | 0.014838 | 0.790501 | Other Positive Correlation |
| CD69 | 0.014838 | 0.790501 | Other Positive Correlation |
| CLEC12A | 0.014838 | 0.790501 | Other Positive Correlation |
| CLEC12B | 0.014838 | 0.790501 | Other Positive Correlation |
| STOML3 | 0.014838 | 0.790501 | Other Positive Correlation |
| TLR7 | 0.014838 | 0.790501 | Other Positive Correlation |
| EVX1 | 0.014838 | 0.790501 | Other Positive Correlation |
| KRTAP2-4 | 0.014838 | 0.790501 | Other Positive Correlation |
| KCNH7 | 0.014838 | 0.790501 | Other Positive Correlation |
| CASP12 | 0.014838 | 0.790501 | Other Positive Correlation |
| CLEC2A | 0.014838 | 0.790501 | Other Positive Correlation |
| PPIAL4H | 0.014838 | 0.790501 | Other Positive Correlation |
| LYZ | 0.014838 | 0.790501 | Other Positive Correlation |
| PKD2L1 | 0.014838 | 0.790501 | Other Positive Correlation |
| HEMGN | 0.014838 | 0.790501 | Other Positive Correlation |
| HMGCS2 | 0.014838 | 0.790501 | Other Positive Correlation |
| KERA | 0.014838 | 0.790501 | Other Positive Correlation |
| GTSF1 | 0.014838 | 0.790501 | Other Positive Correlation |
| HOXA11 | 0.014838 | 0.790501 | Other Positive Correlation |
| KRTAP3-2 | 0.014838 | 0.790501 | Other Positive Correlation |
| ANAPC1 | 0.014831 | 0.790603 | Other Positive Correlation |
| SEPSECS | 0.014818 | 0.790777 | Other Positive Correlation |
| EPHA8 | 0.01475 | 0.791723 | Other Positive Correlation |
| TMEM41B | 0.014743 | 0.791819 | Other Positive Correlation |
| GOLGA6L1 | 0.014737 | 0.791907 | Other Positive Correlation |

| | | | |
|---|---|---|---|
| CCDC178 | 0.014724 | 0.79208 | Other Positive Correlation |
| ADGB | 0.014715 | 0.792204 | Other Positive Correlation |
| H1-8 | 0.014715 | 0.792204 | Other Positive Correlation |
| KLK15 | 0.014715 | 0.792204 | Other Positive Correlation |
| DLL3 | 0.014679 | 0.792703 | Other Positive Correlation |
| KLF17 | 0.014677 | 0.792732 | Other Positive Correlation |
| PCDHA9 | 0.014665 | 0.792895 | Other Positive Correlation |
| MMP24 | 0.014664 | 0.792911 | Other Positive Correlation |
| MCM3AP | 0.014659 | 0.792976 | Other Positive Correlation |
| KRT40 | 0.014659 | 0.792977 | Other Positive Correlation |
| CDK11B | 0.014651 | 0.79309 | Other Positive Correlation |
| TAF6 | 0.014648 | 0.793125 | Other Positive Correlation |
| ATP6V0A1 | 0.014606 | 0.793706 | Other Positive Correlation |
| LEP | 0.014591 | 0.793922 | Other Positive Correlation |
| TNFRSF4 | 0.014563 | 0.794296 | Other Positive Correlation |
| PLEKHM1 | 0.014554 | 0.794429 | Other Positive Correlation |
| CCDC3 | 0.014526 | 0.79481 | Other Positive Correlation |
| RHCE | 0.014525 | 0.794822 | Other Positive Correlation |
| HSPB2 | 0.014511 | 0.795016 | Other Positive Correlation |
| SARDH | 0.014497 | 0.795212 | Other Positive Correlation |
| METTL25B | 0.014481 | 0.795429 | Other Positive Correlation |
| GTF3C3 | 0.014464 | 0.795668 | Other Positive Correlation |
| ABCC2 | 0.014452 | 0.79584 | Other Positive Correlation |
| ZNF142 | 0.014444 | 0.795942 | Other Positive Correlation |
| OR6A2 | 0.014444 | 0.795947 | Other Positive Correlation |
| EMC10 | 0.014434 | 0.796084 | Other Positive Correlation |
| PIGK | 0.014433 | 0.796104 | Other Positive Correlation |
| MFRP | 0.014432 | 0.79611 | Other Positive Correlation |
| SPACA4 | 0.014432 | 0.79611 | Other Positive Correlation |
| CYP4F11 | 0.014425 | 0.796211 | Other Positive Correlation |
| C18orf21 | 0.014401 | 0.796539 | Other Positive Correlation |
| LYG1 | 0.014398 | 0.79658 | Other Positive Correlation |
| TTC7A | 0.014395 | 0.796628 | Other Positive Correlation |
| TMEM222 | 0.014391 | 0.796677 | Other Positive Correlation |
| SERPIND1 | 0.014387 | 0.796733 | Other Positive Correlation |
| NPEPPS | 0.014367 | 0.797014 | Other Positive Correlation |
| ZNF132 | 0.014359 | 0.797117 | Other Positive Correlation |
| ANKRD13C | 0.014324 | 0.797603 | Other Positive Correlation |
| HABP4 | 0.014316 | 0.797713 | Other Positive Correlation |
| CALHM4 | 0.014316 | 0.79772 | Other Positive Correlation |
| DOK4 | 0.014313 | 0.797757 | Other Positive Correlation |
| ACP7 | 0.014305 | 0.797866 | Other Positive Correlation |
| MASP1 | 0.014297 | 0.797973 | Other Positive Correlation |
| NTS | 0.01429 | 0.798069 | Other Positive Correlation |
| ZC3H11B | 0.014278 | 0.798234 | Other Positive Correlation |
| ATCAY | 0.014276 | 0.798267 | Other Positive Correlation |
| CFAP99 | 0.014241 | 0.798756 | Other Positive Correlation |
| LEF1 | 0.014229 | 0.798916 | Other Positive Correlation |

| | | | |
|---|---|---|---|
| PVR | 0.014204 | 0.799259 | Other Positive Correlation |
| COL26A1 | 0.014204 | 0.79927 | Other Positive Correlation |
| SNURF | 0.014202 | 0.799293 | Other Positive Correlation |
| ANKHD1-EIF4EBP3 | 0.014172 | 0.799701 | Other Positive Correlation |
| FAM124B | 0.014171 | 0.799727 | Other Positive Correlation |
| PRSS2 | 0.014171 | 0.799727 | Other Positive Correlation |
| CEMP1 | 0.014171 | 0.799727 | Other Positive Correlation |
| TFCP2 | 0.014165 | 0.799798 | Other Positive Correlation |
| GLRX2 | 0.014153 | 0.799967 | Other Positive Correlation |
| SLC35F3 | 0.014151 | 0.799992 | Other Positive Correlation |
| UNC119B | 0.014143 | 0.800112 | Other Positive Correlation |
| SH3GL1 | 0.014143 | 0.800112 | Other Positive Correlation |
| FAM83G | 0.014137 | 0.800195 | Other Positive Correlation |
| B4GALT4 | 0.014116 | 0.800488 | Other Positive Correlation |
| GALNT16 | 0.014102 | 0.800669 | Other Positive Correlation |
| POLN | 0.0141 | 0.800703 | Other Positive Correlation |
| DCTN4 | 0.014085 | 0.800909 | Other Positive Correlation |
| OR2AT4 | 0.01407 | 0.801112 | Other Positive Correlation |
| C14orf180 | 0.01407 | 0.801112 | Other Positive Correlation |
| CD53 | 0.014013 | 0.80191 | Other Positive Correlation |
| ZNF429 | 0.013995 | 0.802161 | Other Positive Correlation |
| ABCA9 | 0.013992 | 0.802191 | Other Positive Correlation |
| ARSL | 0.013946 | 0.802836 | Other Positive Correlation |
| BTN3A1 | 0.013937 | 0.802954 | Other Positive Correlation |
| STATH | 0.013931 | 0.803041 | Other Positive Correlation |
| KRT28 | 0.013931 | 0.803041 | Other Positive Correlation |
| C1orf56 | 0.013904 | 0.80341 | Other Positive Correlation |
| BRD7 | 0.013901 | 0.803459 | Other Positive Correlation |
| IL18BP | 0.013897 | 0.803507 | Other Positive Correlation |
| DNAAF1 | 0.013885 | 0.803674 | Other Positive Correlation |
| ZKSCAN2 | 0.013883 | 0.803713 | Other Positive Correlation |
| EPB41L4A | 0.013882 | 0.803716 | Other Positive Correlation |
| MBD3L2 | 0.013867 | 0.803925 | Other Positive Correlation |
| PHGR1 | 0.01386 | 0.80402 | Other Positive Correlation |
| ACAD10 | 0.013827 | 0.804481 | Other Positive Correlation |
| TESPA1 | 0.013823 | 0.804533 | Other Positive Correlation |
| PPFIA1 | 0.013785 | 0.80507 | Other Positive Correlation |
| RCOR2 | 0.01378 | 0.805139 | Other Positive Correlation |
| SETD1A | 0.013779 | 0.805145 | Other Positive Correlation |
| SMURF1 | 0.013778 | 0.805161 | Other Positive Correlation |
| TCIRG1 | 0.013751 | 0.805542 | Other Positive Correlation |
| FIG4 | 0.013693 | 0.806346 | Other Positive Correlation |
| USHBP1 | 0.013649 | 0.806943 | Other Positive Correlation |
| CUL5 | 0.013649 | 0.806949 | Other Positive Correlation |
| KRT12 | 0.013613 | 0.807453 | Other Positive Correlation |
| KRTAP1-3 | 0.013613 | 0.807453 | Other Positive Correlation |
| KRTAP1-5 | 0.013613 | 0.807453 | Other Positive Correlation |
| KRT20 | 0.013613 | 0.807453 | Other Positive Correlation |

| | | | |
|---|---|---|---|
| KRT24 | 0.013613 | 0.807453 | Other Positive Correlation |
| ATP13A5 | 0.013613 | 0.807453 | Other Positive Correlation |
| KRTAP2-2 | 0.013613 | 0.807453 | Other Positive Correlation |
| KRTAP1-1 | 0.013613 | 0.807453 | Other Positive Correlation |
| KRTAP3-1 | 0.013613 | 0.807453 | Other Positive Correlation |
| KRT25 | 0.013613 | 0.807453 | Other Positive Correlation |
| DOK3 | 0.013593 | 0.807733 | Other Positive Correlation |
| COL14A1 | 0.013585 | 0.80784 | Other Positive Correlation |
| GRTP1 | 0.013572 | 0.808017 | Other Positive Correlation |
| TM9SF2 | 0.013571 | 0.808028 | Other Positive Correlation |
| PLEKHA6 | 0.013552 | 0.808298 | Other Positive Correlation |
| AFAP1L1 | 0.013552 | 0.808301 | Other Positive Correlation |
| PRR5-ARHGAP8 | 0.013544 | 0.808406 | Other Positive Correlation |
| DNALI1 | 0.013542 | 0.80844 | Other Positive Correlation |
| ABCF1 | 0.01353 | 0.808601 | Other Positive Correlation |
| ENSG00000260342 | 0.013522 | 0.808708 | Other Positive Correlation |
| AOC1 | 0.013522 | 0.808708 | Other Positive Correlation |
| TIGD5 | 0.013509 | 0.808897 | Other Positive Correlation |
| SEPTIN5 | 0.013502 | 0.808992 | Other Positive Correlation |
| CARMIL1 | 0.013487 | 0.809203 | Other Positive Correlation |
| ADAMTS7 | 0.013469 | 0.809445 | Other Positive Correlation |
| SLC15A2 | 0.013458 | 0.809602 | Other Positive Correlation |
| INTS9 | 0.013448 | 0.809742 | Other Positive Correlation |
| TRPM8 | 0.01343 | 0.809984 | Other Positive Correlation |
| TBC1D22B | 0.013421 | 0.810119 | Other Positive Correlation |
| ENSG00000283201 | 0.013417 | 0.810174 | Other Positive Correlation |
| SPP1 | 0.013372 | 0.810798 | Other Positive Correlation |
| PCDHB9 | 0.013339 | 0.811258 | Other Positive Correlation |
| C19orf67 | 0.013258 | 0.812371 | Other Positive Correlation |
| MYOZ2 | 0.013258 | 0.812371 | Other Positive Correlation |
| PLP1 | 0.013258 | 0.812371 | Other Positive Correlation |
| TYROBP | 0.013258 | 0.812371 | Other Positive Correlation |
| POC1B-GALNT4 | 0.013258 | 0.812371 | Other Positive Correlation |
| MOGAT2 | 0.013258 | 0.812371 | Other Positive Correlation |
| EPS8L3 | 0.013258 | 0.812371 | Other Positive Correlation |
| RNASE1 | 0.013258 | 0.812371 | Other Positive Correlation |
| LILRA6 | 0.013258 | 0.812371 | Other Positive Correlation |
| KRTAP5-10 | 0.013258 | 0.812371 | Other Positive Correlation |
| OR1L8 | 0.013258 | 0.812371 | Other Positive Correlation |
| TRPM5 | 0.013258 | 0.812371 | Other Positive Correlation |
| GARIN5B | 0.013258 | 0.812371 | Other Positive Correlation |
| TIE1 | 0.013258 | 0.812371 | Other Positive Correlation |
| ZDHHC19 | 0.013258 | 0.812371 | Other Positive Correlation |
| PRF1 | 0.013258 | 0.812371 | Other Positive Correlation |
| MUC21 | 0.013258 | 0.812371 | Other Positive Correlation |
| TRIML1 | 0.013258 | 0.812371 | Other Positive Correlation |
| MEFV | 0.013258 | 0.812371 | Other Positive Correlation |
| SIGLEC9 | 0.013258 | 0.812371 | Other Positive Correlation |

| | | | |
|---|---|---|---|
| GPRC5D | 0.013258 | 0.812371 | Other Positive Correlation |
| GPR55 | 0.013258 | 0.812371 | Other Positive Correlation |
| FAM180A | 0.013258 | 0.812371 | Other Positive Correlation |
| CILP | 0.013258 | 0.812371 | Other Positive Correlation |
| LIM2 | 0.013258 | 0.812371 | Other Positive Correlation |
| KLK14 | 0.013258 | 0.812371 | Other Positive Correlation |
| TMF1 | 0.013217 | 0.812949 | Other Positive Correlation |
| SYT10 | 0.013201 | 0.813163 | Other Positive Correlation |
| SALL1 | 0.013201 | 0.813163 | Other Positive Correlation |
| SHISA7 | 0.013146 | 0.81393 | Other Positive Correlation |
| TMED9 | 0.013146 | 0.813932 | Other Positive Correlation |
| TMEM196 | 0.01314 | 0.81402 | Other Positive Correlation |
| ZFP37 | 0.013134 | 0.814098 | Other Positive Correlation |
| TMEM160 | 0.013051 | 0.815249 | Other Positive Correlation |
| VIPAS39 | 0.013043 | 0.815361 | Other Positive Correlation |
| COX15 | 0.013033 | 0.815502 | Other Positive Correlation |
| DNAJB13 | 0.01302 | 0.815677 | Other Positive Correlation |
| PTN | 0.012993 | 0.816062 | Other Positive Correlation |
| UBL7 | 0.01297 | 0.816375 | Other Positive Correlation |
| CTRC | 0.012927 | 0.816977 | Other Positive Correlation |
| POPDC2 | 0.012908 | 0.817241 | Other Positive Correlation |
| GLYR1 | 0.012886 | 0.817553 | Other Positive Correlation |
| COBLL1 | 0.012869 | 0.817788 | Other Positive Correlation |
| ENSG00000260272 | 0.012849 | 0.818056 | Other Positive Correlation |
| GNG8 | 0.012849 | 0.818056 | Other Positive Correlation |
| FCRL2 | 0.012849 | 0.818056 | Other Positive Correlation |
| THRSP | 0.012849 | 0.818056 | Other Positive Correlation |
| FABP1 | 0.012849 | 0.818056 | Other Positive Correlation |
| ENSG00000249240 | 0.012849 | 0.818056 | Other Positive Correlation |
| SCNM1 | 0.012844 | 0.818124 | Other Positive Correlation |
| CD177 | 0.012844 | 0.818133 | Other Positive Correlation |
| RMND5A | 0.012837 | 0.818229 | Other Positive Correlation |
| ZNF182 | 0.012834 | 0.818272 | Other Positive Correlation |
| DISP3 | 0.012826 | 0.818386 | Other Positive Correlation |
| SLN | 0.012812 | 0.818576 | Other Positive Correlation |
| SARNP | 0.012795 | 0.818807 | Other Positive Correlation |
| GPR35 | 0.012766 | 0.819213 | Other Positive Correlation |
| INAFM2 | 0.012743 | 0.819543 | Other Positive Correlation |
| TMEM229B | 0.01272 | 0.81986 | Other Positive Correlation |
| DNAJB4 | 0.012708 | 0.820024 | Other Positive Correlation |
| USP26 | 0.012699 | 0.820143 | Other Positive Correlation |
| CETN3 | 0.012649 | 0.82085 | Other Positive Correlation |
| CPNE9 | 0.012637 | 0.821018 | Other Positive Correlation |
| AADACL3 | 0.012637 | 0.821018 | Other Positive Correlation |
| BTNL8 | 0.012637 | 0.821018 | Other Positive Correlation |
| HORMAD1 | 0.012637 | 0.821018 | Other Positive Correlation |
| OXGR1 | 0.012637 | 0.821018 | Other Positive Correlation |
| ACKR2 | 0.012624 | 0.821191 | Other Positive Correlation |

| | | | |
|---|---|---|---|
| UPB1 | 0.012563 | 0.822043 | Other Positive Correlation |
| ARPC5L | 0.01256 | 0.82208 | Other Positive Correlation |
| DENND10 | 0.012551 | 0.822207 | Other Positive Correlation |
| ALOXE3 | 0.012527 | 0.822539 | Other Positive Correlation |
| CALCOCO1 | 0.012523 | 0.822599 | Other Positive Correlation |
| BSND | 0.012494 | 0.823008 | Other Positive Correlation |
| FBXO11 | 0.012456 | 0.82354 | Other Positive Correlation |
| METTL21C | 0.012453 | 0.823571 | Other Positive Correlation |
| ENSG00000237378 | 0.012395 | 0.824381 | Other Positive Correlation |
| HIGD2B | 0.012332 | 0.825259 | Other Positive Correlation |
| BAHCC1 | 0.012331 | 0.825272 | Other Positive Correlation |
| RAB4B-EGLN2 | 0.012331 | 0.825274 | Other Positive Correlation |
| TMEM151B | 0.012324 | 0.825367 | Other Positive Correlation |
| WWTR1 | 0.012318 | 0.825454 | Other Positive Correlation |
| CDK15 | 0.012295 | 0.825775 | Other Positive Correlation |
| ZNF20 | 0.012264 | 0.826205 | Other Positive Correlation |
| ARID3B | 0.012256 | 0.826321 | Other Positive Correlation |
| CTIF | 0.012225 | 0.82676 | Other Positive Correlation |
| MED27 | 0.012203 | 0.827065 | Other Positive Correlation |
| SLC12A5 | 0.012186 | 0.827303 | Other Positive Correlation |
| NUDT17 | 0.012172 | 0.827496 | Other Positive Correlation |
| BOLA2B | 0.012133 | 0.828032 | Other Positive Correlation |
| TSEN54 | 0.012112 | 0.828333 | Other Positive Correlation |
| NHLRC3 | 0.012101 | 0.82848 | Other Positive Correlation |
| PABPC3 | 0.012083 | 0.828736 | Other Positive Correlation |
| DGCR6L | 0.012064 | 0.828998 | Other Positive Correlation |
| NYAP2 | 0.012053 | 0.829149 | Other Positive Correlation |
| RWDD2A | 0.012032 | 0.829448 | Other Positive Correlation |
| ENSG00000284695 | 0.012016 | 0.829676 | Other Positive Correlation |
| RIMBP3 | 0.012016 | 0.829676 | Other Positive Correlation |
| CCDC201 | 0.012016 | 0.829676 | Other Positive Correlation |
| CALN1 | 0.012016 | 0.829676 | Other Positive Correlation |
| C3orf33 | 0.011985 | 0.830102 | Other Positive Correlation |
| WDR90 | 0.011943 | 0.830692 | Other Positive Correlation |
| ACP3 | 0.011933 | 0.830832 | Other Positive Correlation |
| PHF23 | 0.011916 | 0.831063 | Other Positive Correlation |
| CCER1 | 0.011901 | 0.831272 | Other Positive Correlation |
| TMEM42 | 0.011893 | 0.831383 | Other Positive Correlation |
| SOS2 | 0.011884 | 0.831518 | Other Positive Correlation |
| ZSCAN30 | 0.011847 | 0.832033 | Other Positive Correlation |
| NRARP | 0.011833 | 0.832229 | Other Positive Correlation |
| VPS45 | 0.011829 | 0.832277 | Other Positive Correlation |
| KCNN2 | 0.011828 | 0.832302 | Other Positive Correlation |
| GADD45B | 0.011773 | 0.833059 | Other Positive Correlation |
| SCAP | 0.011735 | 0.833599 | Other Positive Correlation |
| AGBL2 | 0.01172 | 0.833808 | Other Positive Correlation |
| REXO5 | 0.011686 | 0.834282 | Other Positive Correlation |
| IPO9 | 0.011668 | 0.834534 | Other Positive Correlation |

| | | | |
|---|---|---|---|
| AP3M1 | 0.01164 | 0.834927 | Other Positive Correlation |
| KRT79 | 0.011637 | 0.834958 | Other Positive Correlation |
| CSF2RB | 0.011606 | 0.835394 | Other Positive Correlation |
| PCDHGA2 | 0.011594 | 0.835563 | Other Positive Correlation |
| KRBA2 | 0.011593 | 0.835583 | Other Positive Correlation |
| YIPF1 | 0.011577 | 0.835805 | Other Positive Correlation |
| ENSG00000261611 | 0.011571 | 0.835888 | Other Positive Correlation |
| KLHL7 | 0.011546 | 0.836236 | Other Positive Correlation |
| SCARA5 | 0.011537 | 0.836358 | Other Positive Correlation |
| SYT1 | 0.011526 | 0.836518 | Other Positive Correlation |
| RAP2C | 0.011523 | 0.836559 | Other Positive Correlation |
| EPPIN | 0.011521 | 0.83658 | Other Positive Correlation |
| APOL5 | 0.011521 | 0.83658 | Other Positive Correlation |
| CCRL2 | 0.011521 | 0.83658 | Other Positive Correlation |
| SLC9A4 | 0.011521 | 0.83658 | Other Positive Correlation |
| ATF7-NPFF | 0.011521 | 0.83658 | Other Positive Correlation |
| NLRP12 | 0.011521 | 0.83658 | Other Positive Correlation |
| DPEP2 | 0.011521 | 0.83658 | Other Positive Correlation |
| ZMYM3 | 0.011482 | 0.837131 | Other Positive Correlation |
| MCF2 | 0.011477 | 0.837209 | Other Positive Correlation |
| PTGIR | 0.011405 | 0.838214 | Other Positive Correlation |
| LRRC34 | 0.011382 | 0.838525 | Other Positive Correlation |
| CDK18 | 0.011371 | 0.838681 | Other Positive Correlation |
| EPHA4 | 0.011366 | 0.838748 | Other Positive Correlation |
| CPNE4 | 0.011365 | 0.838771 | Other Positive Correlation |
| PNPLA1 | 0.011324 | 0.839347 | Other Positive Correlation |
| TBX6 | 0.01132 | 0.839402 | Other Positive Correlation |
| SLC12A6 | 0.011302 | 0.839649 | Other Positive Correlation |
| GOLGA6L7 | 0.011296 | 0.839736 | Other Positive Correlation |
| ZCCHC4 | 0.011289 | 0.839832 | Other Positive Correlation |
| COX17 | 0.011283 | 0.839916 | Other Positive Correlation |
| PREX2 | 0.011243 | 0.840481 | Other Positive Correlation |
| PAEP | 0.011243 | 0.840481 | Other Positive Correlation |
| PIWIL3 | 0.011243 | 0.840481 | Other Positive Correlation |
| HOXB8 | 0.011243 | 0.840481 | Other Positive Correlation |
| SLC5A12 | 0.011243 | 0.840481 | Other Positive Correlation |
| ITPRID1 | 0.011243 | 0.840481 | Other Positive Correlation |
| BARHL1 | 0.011243 | 0.840481 | Other Positive Correlation |
| SPINK1 | 0.011243 | 0.840481 | Other Positive Correlation |
| MS4A6A | 0.011243 | 0.840481 | Other Positive Correlation |
| TFAP2D | 0.011243 | 0.840481 | Other Positive Correlation |
| TSPAN8 | 0.011243 | 0.840481 | Other Positive Correlation |
| OR5B12 | 0.011243 | 0.840481 | Other Positive Correlation |
| TPTE2 | 0.011243 | 0.840481 | Other Positive Correlation |
| ANKRD55 | 0.011243 | 0.840481 | Other Positive Correlation |
| IRF8 | 0.011243 | 0.840481 | Other Positive Correlation |
| IZUMO1R | 0.011243 | 0.840481 | Other Positive Correlation |
| HBM | 0.011243 | 0.840481 | Other Positive Correlation |

| | | | |
|---|---|---|---|
| ALPG | 0.011243 | 0.840481 | Other Positive Correlation |
| PDZD9 | 0.011243 | 0.840481 | Other Positive Correlation |
| CST4 | 0.011243 | 0.840481 | Other Positive Correlation |
| OR5B2 | 0.011243 | 0.840481 | Other Positive Correlation |
| ABHD17A | 0.01122 | 0.840792 | Other Positive Correlation |
| TTLL5 | 0.011219 | 0.840813 | Other Positive Correlation |
| CSNK2B | 0.011211 | 0.840918 | Other Positive Correlation |
| FGF4 | 0.01121 | 0.840934 | Other Positive Correlation |
| MMP20 | 0.01121 | 0.840934 | Other Positive Correlation |
| EPHA3 | 0.01121 | 0.840934 | Other Positive Correlation |
| ANKRD30A | 0.01121 | 0.840934 | Other Positive Correlation |
| KCNA3 | 0.01121 | 0.840934 | Other Positive Correlation |
| SLITRK3 | 0.01121 | 0.840934 | Other Positive Correlation |
| PAGE5 | 0.01121 | 0.840934 | Other Positive Correlation |
| SI | 0.01121 | 0.840934 | Other Positive Correlation |
| COX6A2 | 0.01121 | 0.840934 | Other Positive Correlation |
| ADGRB3 | 0.01121 | 0.840934 | Other Positive Correlation |
| TUNAR | 0.01121 | 0.840934 | Other Positive Correlation |
| GDF10 | 0.01121 | 0.840934 | Other Positive Correlation |
| NXPH1 | 0.01121 | 0.840934 | Other Positive Correlation |
| ACKR1 | 0.01121 | 0.840934 | Other Positive Correlation |
| SLCO6A1 | 0.01121 | 0.840934 | Other Positive Correlation |
| ADAM7 | 0.01121 | 0.840934 | Other Positive Correlation |
| AMY2A | 0.01121 | 0.840934 | Other Positive Correlation |
| DYDC1 | 0.01121 | 0.840934 | Other Positive Correlation |
| OR10H5 | 0.01121 | 0.840934 | Other Positive Correlation |
| PDZRN4 | 0.01121 | 0.840934 | Other Positive Correlation |
| SYT4 | 0.01121 | 0.840934 | Other Positive Correlation |
| RSPO2 | 0.01121 | 0.840934 | Other Positive Correlation |
| MS4A15 | 0.011207 | 0.840979 | Other Positive Correlation |
| CELA3A | 0.011207 | 0.840979 | Other Positive Correlation |
| LPA | 0.011207 | 0.840979 | Other Positive Correlation |
| ENSG00000259529 | 0.011207 | 0.840979 | Other Positive Correlation |
| GLOD5 | 0.011207 | 0.840979 | Other Positive Correlation |
| ASB10 | 0.011207 | 0.840979 | Other Positive Correlation |
| XRN1 | 0.011206 | 0.84099 | Other Positive Correlation |
| CELF3 | 0.011192 | 0.841191 | Other Positive Correlation |
| MT-CO1 | 0.01119 | 0.841222 | Other Positive Correlation |
| B4GALNT3 | 0.011182 | 0.841329 | Other Positive Correlation |
| HERC6 | 0.011167 | 0.841545 | Other Positive Correlation |
| TTF1 | 0.011119 | 0.84221 | Other Positive Correlation |
| LZTFL1 | 0.011117 | 0.842238 | Other Positive Correlation |
| ZC2HC1C | 0.011111 | 0.842329 | Other Positive Correlation |
| WASF3 | 0.011085 | 0.842684 | Other Positive Correlation |
| RAB43 | 0.011075 | 0.842825 | Other Positive Correlation |
| SLC17A4 | 0.011066 | 0.842953 | Other Positive Correlation |
| NHLH2 | 0.011029 | 0.84348 | Other Positive Correlation |
| CGB7 | 0.011018 | 0.843623 | Other Positive Correlation |

| | | | |
|---|---|---|---|
| BMT2 | 0.010999 | 0.843887 | Other Positive Correlation |
| ZNF488 | 0.010976 | 0.844214 | Other Positive Correlation |
| ZBED3 | 0.010968 | 0.844334 | Other Positive Correlation |
| KREMEN1 | 0.010951 | 0.84456 | Other Positive Correlation |
| ZNF362 | 0.010925 | 0.844933 | Other Positive Correlation |
| SERPINA3 | 0.010896 | 0.84534 | Other Positive Correlation |
| SLC39A2 | 0.010842 | 0.846091 | Other Positive Correlation |
| DTNA | 0.010835 | 0.846197 | Other Positive Correlation |
| RAB12 | 0.010828 | 0.846287 | Other Positive Correlation |
| ULBP3 | 0.010795 | 0.846755 | Other Positive Correlation |
| PKN2 | 0.010763 | 0.847201 | Other Positive Correlation |
| MAPK4 | 0.010761 | 0.847234 | Other Positive Correlation |
| MYCT1 | 0.010739 | 0.847535 | Other Positive Correlation |
| COP1 | 0.010726 | 0.847725 | Other Positive Correlation |
| FMNL1 | 0.010725 | 0.847729 | Other Positive Correlation |
| KLK6 | 0.010724 | 0.847752 | Other Positive Correlation |
| CIDEC | 0.010704 | 0.848026 | Other Positive Correlation |
| SPATA3 | 0.010683 | 0.848328 | Other Positive Correlation |
| FAM167B | 0.010653 | 0.84874 | Other Positive Correlation |
| ADAMTS20 | 0.010643 | 0.848882 | Other Positive Correlation |
| SRD5A2 | 0.010643 | 0.848882 | Other Positive Correlation |
| TEX45 | 0.010614 | 0.849296 | Other Positive Correlation |
| OGFR | 0.01061 | 0.849344 | Other Positive Correlation |
| THSD7B | 0.010608 | 0.849376 | Other Positive Correlation |
| YAF2 | 0.010594 | 0.849573 | Other Positive Correlation |
| ATP6V1D | 0.010549 | 0.850201 | Other Positive Correlation |
| SORBS3 | 0.010542 | 0.850308 | Other Positive Correlation |
| FGF7 | 0.010537 | 0.850374 | Other Positive Correlation |
| PAQR8 | 0.010536 | 0.850391 | Other Positive Correlation |
| GPAT4 | 0.01052 | 0.850607 | Other Positive Correlation |
| MYMX | 0.010518 | 0.850643 | Other Positive Correlation |
| CATSPER3 | 0.010515 | 0.850684 | Other Positive Correlation |
| PRPH2 | 0.010498 | 0.850916 | Other Positive Correlation |
| TMEM81 | 0.010489 | 0.85104 | Other Positive Correlation |
| INSIG2 | 0.010461 | 0.851441 | Other Positive Correlation |
| ROBO1 | 0.010459 | 0.851471 | Other Positive Correlation |
| TNFSF15 | 0.01042 | 0.852018 | Other Positive Correlation |
| NUS1 | 0.010417 | 0.852063 | Other Positive Correlation |
| POU4F3 | 0.010415 | 0.852089 | Other Positive Correlation |
| POLR2J | 0.010412 | 0.85213 | Other Positive Correlation |
| CMTM5 | 0.010395 | 0.85237 | Other Positive Correlation |
| SNORC | 0.01038 | 0.85257 | Other Positive Correlation |
| JAZF1 | 0.010379 | 0.852583 | Other Positive Correlation |
| PHAF1 | 0.010328 | 0.853304 | Other Positive Correlation |
| BMP5 | 0.010327 | 0.853314 | Other Positive Correlation |
| ABHD5 | 0.010291 | 0.853819 | Other Positive Correlation |
| SLC52A3 | 0.010278 | 0.854007 | Other Positive Correlation |
| CALHM2 | 0.010238 | 0.854576 | Other Positive Correlation |

| | | | |
|---|---|---|---|
| ENTREP1 | 0.010194 | 0.855186 | Other Positive Correlation |
| ANGPTL6 | 0.010188 | 0.855271 | Other Positive Correlation |
| ZMAT4 | 0.010173 | 0.855486 | Other Positive Correlation |
| CATSPER4 | 0.010173 | 0.855486 | Other Positive Correlation |
| EIF3C | 0.010115 | 0.856304 | Other Positive Correlation |
| BCO1 | 0.010109 | 0.856382 | Other Positive Correlation |
| RGPD5 | 0.010094 | 0.856591 | Other Positive Correlation |
| AK4P3 | 0.010081 | 0.856773 | Other Positive Correlation |
| ITIH4 | 0.010072 | 0.856904 | Other Positive Correlation |
| ARHGAP32 | 0.010051 | 0.857191 | Other Positive Correlation |
| CLDN8 | 0.010049 | 0.857225 | Other Positive Correlation |
| SIX2 | 0.009993 | 0.858007 | Other Positive Correlation |
| CCDC88B | 0.009986 | 0.858108 | Other Positive Correlation |
| PTPN7 | 0.009979 | 0.858205 | Other Positive Correlation |
| FAHD2B | 0.009969 | 0.858353 | Other Positive Correlation |
| TNFAIP6 | 0.009954 | 0.858554 | Other Positive Correlation |
| PYHIN1 | 0.009942 | 0.858732 | Other Positive Correlation |
| NR2C1 | 0.009925 | 0.858964 | Other Positive Correlation |
| CNTF | 0.009894 | 0.859411 | Other Positive Correlation |
| PDE1A | 0.009866 | 0.859801 | Other Positive Correlation |
| DSC1 | 0.009859 | 0.859894 | Other Positive Correlation |
| GRIN2A | 0.009852 | 0.860002 | Other Positive Correlation |
| ZNF593OS | 0.009845 | 0.860094 | Other Positive Correlation |
| OR1B1 | 0.009837 | 0.860207 | Other Positive Correlation |
| CSF2RA | 0.009829 | 0.860325 | Other Positive Correlation |
| MAP10 | 0.009828 | 0.860328 | Other Positive Correlation |
| UBIAD1 | 0.009827 | 0.860348 | Other Positive Correlation |
| KIF2A | 0.009783 | 0.86097 | Other Positive Correlation |
| DZIP1 | 0.009777 | 0.861051 | Other Positive Correlation |
| ALOX12B | 0.009773 | 0.861101 | Other Positive Correlation |
| FAM13A | 0.009773 | 0.861111 | Other Positive Correlation |
| ZNF646 | 0.009734 | 0.861652 | Other Positive Correlation |
| TMEM114 | 0.009713 | 0.861954 | Other Positive Correlation |
| PNLIPRP3 | 0.009686 | 0.86233 | Other Positive Correlation |
| FETUB | 0.009678 | 0.862445 | Other Positive Correlation |
| RPS6KA6 | 0.009641 | 0.86297 | Other Positive Correlation |
| SKAP2 | 0.009595 | 0.86361 | Other Positive Correlation |
| ENSG00000283599 | 0.009588 | 0.863707 | Other Positive Correlation |
| SERPINB12 | 0.009577 | 0.86387 | Other Positive Correlation |
| TRAF6 | 0.009557 | 0.864151 | Other Positive Correlation |
| CXCL13 | 0.009532 | 0.864502 | Other Positive Correlation |
| DPYS | 0.009532 | 0.864502 | Other Positive Correlation |
| TSPAN16 | 0.009532 | 0.864502 | Other Positive Correlation |
| SH3PXD2B | 0.009529 | 0.864537 | Other Positive Correlation |
| TYK2 | 0.009506 | 0.864863 | Other Positive Correlation |
| BEND5 | 0.009491 | 0.865075 | Other Positive Correlation |
| LRIG1 | 0.00949 | 0.865084 | Other Positive Correlation |
| PIWIL2 | 0.009469 | 0.865384 | Other Positive Correlation |

| | | | |
|---|---|---|---|
| TM4SF4 | 0.00946 | 0.865518 | Other Positive Correlation |
| GNE | 0.009459 | 0.865522 | Other Positive Correlation |
| DDX41 | 0.009442 | 0.865766 | Other Positive Correlation |
| SORCS2 | 0.009419 | 0.866085 | Other Positive Correlation |
| KRBOX4 | 0.009404 | 0.866305 | Other Positive Correlation |
| SUN3 | 0.009366 | 0.866833 | Other Positive Correlation |
| CCDC163 | 0.009322 | 0.867461 | Other Positive Correlation |
| ZNF766 | 0.009313 | 0.867578 | Other Positive Correlation |
| DHX30 | 0.0093 | 0.867761 | Other Positive Correlation |
| DLST | 0.009287 | 0.867944 | Other Positive Correlation |
| ZNF77 | 0.009271 | 0.868168 | Other Positive Correlation |
| PCDHGA7 | 0.00923 | 0.868746 | Other Positive Correlation |
| PI3 | 0.009211 | 0.869019 | Other Positive Correlation |
| RHOH | 0.009208 | 0.869059 | Other Positive Correlation |
| IRAG1 | 0.009185 | 0.869393 | Other Positive Correlation |
| ELOF1 | 0.009181 | 0.869439 | Other Positive Correlation |
| STKLD1 | 0.009173 | 0.869559 | Other Positive Correlation |
| S1PR2 | 0.009152 | 0.869856 | Other Positive Correlation |
| CRTC2 | 0.009132 | 0.870137 | Other Positive Correlation |
| TMEM71 | 0.009111 | 0.870429 | Other Positive Correlation |
| RNF122 | 0.009109 | 0.870451 | Other Positive Correlation |
| TEX26 | 0.009109 | 0.870459 | Other Positive Correlation |
| SEC62 | 0.009104 | 0.870531 | Other Positive Correlation |
| SPRR5 | 0.009078 | 0.870901 | Other Positive Correlation |
| POU6F1 | 0.009072 | 0.87098 | Other Positive Correlation |
| SBNO1 | 0.009057 | 0.871194 | Other Positive Correlation |
| FOXN3 | 0.009046 | 0.871344 | Other Positive Correlation |
| RRH | 0.00895 | 0.872699 | Other Positive Correlation |
| AWAT2 | 0.008946 | 0.872753 | Other Positive Correlation |
| C16orf90 | 0.008936 | 0.872903 | Other Positive Correlation |
| ZDBF2 | 0.008925 | 0.873047 | Other Positive Correlation |
| FER1L6 | 0.008923 | 0.873083 | Other Positive Correlation |
| ZNF626 | 0.008904 | 0.873352 | Other Positive Correlation |
| INTU | 0.008882 | 0.873661 | Other Positive Correlation |
| PCNT | 0.008831 | 0.87438 | Other Positive Correlation |
| PAQR6 | 0.008819 | 0.874552 | Other Positive Correlation |
| GLB1L2 | 0.008817 | 0.874582 | Other Positive Correlation |
| HSD17B13 | 0.008803 | 0.874774 | Other Positive Correlation |
| COG7 | 0.008795 | 0.874891 | Other Positive Correlation |
| SDR9C7 | 0.008794 | 0.874899 | Other Positive Correlation |
| IMPG2 | 0.008782 | 0.875075 | Other Positive Correlation |
| SFTA2 | 0.008731 | 0.875791 | Other Positive Correlation |
| NGRN | 0.008713 | 0.876036 | Other Positive Correlation |
| STAC3 | 0.008668 | 0.876682 | Other Positive Correlation |
| TTC31 | 0.008619 | 0.87737 | Other Positive Correlation |
| TAF5L | 0.008616 | 0.87741 | Other Positive Correlation |
| MSANTD7 | 0.008582 | 0.877892 | Other Positive Correlation |
| LATS2 | 0.008572 | 0.878028 | Other Positive Correlation |

| | | | |
|---|---|---|---|
| MEF2A | 0.00857 | 0.878068 | Other Positive Correlation |
| GNB3 | 0.008548 | 0.878377 | Other Positive Correlation |
| DHX8 | 0.008531 | 0.878614 | Other Positive Correlation |
| SERPINC1 | 0.008531 | 0.878615 | Other Positive Correlation |
| SEMA4G | 0.008516 | 0.878824 | Other Positive Correlation |
| CISH | 0.008505 | 0.878983 | Other Positive Correlation |
| GPD1L | 0.008504 | 0.878996 | Other Positive Correlation |
| ABI2 | 0.00849 | 0.879188 | Other Positive Correlation |
| RUNX2 | 0.008473 | 0.879429 | Other Positive Correlation |
| PTTG1IP | 0.008458 | 0.879649 | Other Positive Correlation |
| KDR | 0.008441 | 0.879887 | Other Positive Correlation |
| TRIR | 0.008436 | 0.879956 | Other Positive Correlation |
| HACD4 | 0.008381 | 0.880735 | Other Positive Correlation |
| NUP188 | 0.008367 | 0.880929 | Other Positive Correlation |
| SLC45A4 | 0.008348 | 0.881194 | Other Positive Correlation |
| BORCS5 | 0.008327 | 0.8815 | Other Positive Correlation |
| NOG | 0.008324 | 0.881538 | Other Positive Correlation |
| VAMP1 | 0.008323 | 0.881547 | Other Positive Correlation |
| ARHGEF37 | 0.008317 | 0.881638 | Other Positive Correlation |
| MPIG6B | 0.008307 | 0.881778 | Other Positive Correlation |
| SLC38A6 | 0.008294 | 0.881959 | Other Positive Correlation |
| PABPN1 | 0.008271 | 0.882283 | Other Positive Correlation |
| EXOC1 | 0.008265 | 0.88237 | Other Positive Correlation |
| TRAFD1 | 0.008259 | 0.882453 | Other Positive Correlation |
| EFHC2 | 0.008222 | 0.88298 | Other Positive Correlation |
| THBD | 0.008174 | 0.883653 | Other Positive Correlation |
| GCOM1 | 0.008122 | 0.884393 | Other Positive Correlation |
| VWA2 | 0.008081 | 0.884976 | Other Positive Correlation |
| PADI4 | 0.008078 | 0.885007 | Other Positive Correlation |
| PTK6 | 0.008076 | 0.885039 | Other Positive Correlation |
| NEURL4 | 0.008068 | 0.885147 | Other Positive Correlation |
| GOLGA6L2 | 0.008058 | 0.8853 | Other Positive Correlation |
| IRAK1BP1 | 0.008026 | 0.885743 | Other Positive Correlation |
| WFDC3 | 0.008025 | 0.885766 | Other Positive Correlation |
| SLAIN2 | 0.008017 | 0.885881 | Other Positive Correlation |
| STK19 | 0.008014 | 0.885912 | Other Positive Correlation |
| USP25 | 0.008 | 0.886109 | Other Positive Correlation |
| RSPH9 | 0.007969 | 0.88655 | Other Positive Correlation |
| TMIGD2 | 0.007962 | 0.886646 | Other Positive Correlation |
| HABP2 | 0.007962 | 0.886646 | Other Positive Correlation |
| CEMIP | 0.007957 | 0.886721 | Other Positive Correlation |
| GATAD2B | 0.007938 | 0.886994 | Other Positive Correlation |
| CYP4F12 | 0.007937 | 0.886999 | Other Positive Correlation |
| APEX2 | 0.00792 | 0.887239 | Other Positive Correlation |
| ZNF517 | 0.007892 | 0.887647 | Other Positive Correlation |
| ADAMTSL2 | 0.007886 | 0.887725 | Other Positive Correlation |
| COPG2 | 0.007885 | 0.887734 | Other Positive Correlation |
| ARL4D | 0.00788 | 0.887815 | Other Positive Correlation |

| | | | |
|---|---|---|---|
| GPR141 | 0.007876 | 0.887873 | Other Positive Correlation |
| FCRLA | 0.007869 | 0.887965 | Other Positive Correlation |
| ARL17A | 0.007865 | 0.88803 | Other Positive Correlation |
| RNF130 | 0.007863 | 0.888051 | Other Positive Correlation |
| XYLB | 0.007842 | 0.888342 | Other Positive Correlation |
| ENSG00000285589 | 0.007831 | 0.888511 | Other Positive Correlation |
| MYO7B | 0.007805 | 0.888876 | Other Positive Correlation |
| KIF6 | 0.007804 | 0.888883 | Other Positive Correlation |
| MT-ND4 | 0.00779 | 0.88909 | Other Positive Correlation |
| RHOT1 | 0.007783 | 0.889187 | Other Positive Correlation |
| GRXCR2 | 0.007778 | 0.88926 | Other Positive Correlation |
| HMCN2 | 0.007766 | 0.889424 | Other Positive Correlation |
| TMEM190 | 0.007743 | 0.889754 | Other Positive Correlation |
| TAF11L11 | 0.007718 | 0.890097 | Other Positive Correlation |
| MAP1LC3B2 | 0.007694 | 0.890438 | Other Positive Correlation |
| PRR29 | 0.007687 | 0.890536 | Other Positive Correlation |
| ZNF572 | 0.00768 | 0.890645 | Other Positive Correlation |
| ZNF385D | 0.00767 | 0.890776 | Other Positive Correlation |
| PSG8 | 0.007657 | 0.890971 | Other Positive Correlation |
| PDS5B | 0.007647 | 0.891103 | Other Positive Correlation |
| TMSB4Y | 0.007647 | 0.89111 | Other Positive Correlation |
| IFNLR1 | 0.007644 | 0.891153 | Other Positive Correlation |
| FBXL17 | 0.007626 | 0.89141 | Other Positive Correlation |
| CRYBA1 | 0.007616 | 0.89154 | Other Positive Correlation |
| PPP1R27 | 0.007615 | 0.891567 | Other Positive Correlation |
| FKBP4 | 0.007609 | 0.891642 | Other Positive Correlation |
| LAT2 | 0.007589 | 0.891926 | Other Positive Correlation |
| SP2 | 0.007568 | 0.892224 | Other Positive Correlation |
| LAMA4 | 0.007559 | 0.892348 | Other Positive Correlation |
| SLF1 | 0.007543 | 0.892578 | Other Positive Correlation |
| CCL2 | 0.007528 | 0.892791 | Other Positive Correlation |
| CCDC120 | 0.007513 | 0.893008 | Other Positive Correlation |
| ACR | 0.007473 | 0.893575 | Other Positive Correlation |
| TMOD2 | 0.007443 | 0.893992 | Other Positive Correlation |
| NTNG2 | 0.007437 | 0.894076 | Other Positive Correlation |
| ITGBL1 | 0.007434 | 0.894118 | Other Positive Correlation |
| EFCAB6 | 0.007417 | 0.894364 | Other Positive Correlation |
| ANKRD27 | 0.007412 | 0.894433 | Other Positive Correlation |
| ZNF227 | 0.007365 | 0.895095 | Other Positive Correlation |
| C17orf97 | 0.007352 | 0.895286 | Other Positive Correlation |
| TNFSF18 | 0.007343 | 0.895409 | Other Positive Correlation |
| SYNE1 | 0.007311 | 0.895865 | Other Positive Correlation |
| METTL7B | 0.0073 | 0.896026 | Other Positive Correlation |
| LVRN | 0.007255 | 0.89666 | Other Positive Correlation |
| RFFL | 0.007253 | 0.896685 | Other Positive Correlation |
| THAP8 | 0.00725 | 0.896725 | Other Positive Correlation |
| OCA2 | 0.007209 | 0.897309 | Other Positive Correlation |
| MGAM | 0.007196 | 0.897498 | Other Positive Correlation |

| | | | |
|---|---|---|---|
| ANKRD50 | 0.007189 | 0.897597 | Other Positive Correlation |
| ZMIZ1 | 0.007181 | 0.897702 | Other Positive Correlation |
| USP42 | 0.007174 | 0.897811 | Other Positive Correlation |
| ARHGAP27 | 0.00716 | 0.897999 | Other Positive Correlation |
| TREX2 | 0.007157 | 0.898041 | Other Positive Correlation |
| SLC24A3 | 0.007155 | 0.898068 | Other Positive Correlation |
| CASP14 | 0.007151 | 0.898129 | Other Positive Correlation |
| ZFP42 | 0.007147 | 0.898193 | Other Positive Correlation |
| TMEM185B | 0.007129 | 0.898449 | Other Positive Correlation |
| NBPF3 | 0.007126 | 0.898485 | Other Positive Correlation |
| COMP | 0.007122 | 0.898547 | Other Positive Correlation |
| FRG1 | 0.007118 | 0.898602 | Other Positive Correlation |
| CALHM5 | 0.0071 | 0.89885 | Other Positive Correlation |
| ZXDC | 0.007082 | 0.899111 | Other Positive Correlation |
| KCP | 0.007076 | 0.899189 | Other Positive Correlation |
| ZNF257 | 0.007057 | 0.899463 | Other Positive Correlation |
| MCPH1 | 0.007026 | 0.899907 | Other Positive Correlation |
| DISP2 | 0.007005 | 0.900206 | Other Positive Correlation |
| HARS2 | 0.006997 | 0.900315 | Other Positive Correlation |
| RNF215 | 0.006996 | 0.900325 | Other Positive Correlation |
| CHCHD7 | 0.006983 | 0.900511 | Other Positive Correlation |
| DNAAF11 | 0.00698 | 0.900559 | Other Positive Correlation |
| GSTT2B | 0.006974 | 0.900634 | Other Positive Correlation |
| KRT222 | 0.006962 | 0.900806 | Other Positive Correlation |
| CDK20 | 0.006948 | 0.901007 | Other Positive Correlation |
| KRTAP19-1 | 0.006935 | 0.901192 | Other Positive Correlation |
| SOX30 | 0.0069 | 0.901684 | Other Positive Correlation |
| MMEL1 | 0.006892 | 0.901807 | Other Positive Correlation |
| PTPN4 | 0.006867 | 0.902158 | Other Positive Correlation |
| MFSD2B | 0.006858 | 0.90229 | Other Positive Correlation |
| RARRES1 | 0.006823 | 0.90278 | Other Positive Correlation |
| LMNTD1 | 0.006775 | 0.903466 | Other Positive Correlation |
| DOP1B | 0.006724 | 0.904183 | Other Positive Correlation |
| LRP12 | 0.006678 | 0.90484 | Other Positive Correlation |
| HSD17B4 | 0.006638 | 0.905409 | Other Positive Correlation |
| NOXO1 | 0.006622 | 0.905635 | Other Positive Correlation |
| ACAP1 | 0.006619 | 0.905675 | Other Positive Correlation |
| STAT3 | 0.006612 | 0.905774 | Other Positive Correlation |
| CEP295 | 0.006589 | 0.906094 | Other Positive Correlation |
| RAPGEF3 | 0.006577 | 0.906265 | Other Positive Correlation |
| METTL23 | 0.006507 | 0.907264 | Other Positive Correlation |
| ATXN7L3 | 0.006492 | 0.907479 | Other Positive Correlation |
| HOATZ | 0.006463 | 0.907885 | Other Positive Correlation |
| KRT86 | 0.006454 | 0.908021 | Other Positive Correlation |
| MIGA1 | 0.006446 | 0.908126 | Other Positive Correlation |
| SLC6A7 | 0.006436 | 0.908263 | Other Positive Correlation |
| BCL2L14 | 0.006378 | 0.90909 | Other Positive Correlation |
| LCN15 | 0.006361 | 0.909337 | Other Positive Correlation |

| | | | |
|---|---|---|---|
| ADGRF5 | 0.006326 | 0.909833 | Other Positive Correlation |
| SLC35A1 | 0.00632 | 0.90991 | Other Positive Correlation |
| CLK3 | 0.00629 | 0.910346 | Other Positive Correlation |
| DNM3 | 0.006253 | 0.910866 | Other Positive Correlation |
| ESCO1 | 0.006217 | 0.911376 | Other Positive Correlation |
| SPRR4 | 0.006204 | 0.911558 | Other Positive Correlation |
| FSD2 | 0.0062 | 0.911612 | Other Positive Correlation |
| CCDC89 | 0.006169 | 0.912056 | Other Positive Correlation |
| ZNF181 | 0.006153 | 0.91229 | Other Positive Correlation |
| UCN | 0.006127 | 0.912651 | Other Positive Correlation |
| SPEM1 | 0.006016 | 0.914235 | Other Positive Correlation |
| SPIN3 | 0.005984 | 0.91468 | Other Positive Correlation |
| SPAG17 | 0.005936 | 0.915369 | Other Positive Correlation |
| ZNF614 | 0.005935 | 0.915386 | Other Positive Correlation |
| SMTNL2 | 0.005931 | 0.915437 | Other Positive Correlation |
| SUPT5H | 0.005925 | 0.915527 | Other Positive Correlation |
| FBXO33 | 0.005902 | 0.915848 | Other Positive Correlation |
| AKAP13 | 0.005902 | 0.91585 | Other Positive Correlation |
| ST3GAL1 | 0.005869 | 0.916315 | Other Positive Correlation |
| SUSD6 | 0.005855 | 0.916524 | Other Positive Correlation |
| MAPK7 | 0.005854 | 0.916528 | Other Positive Correlation |
| ACSF3 | 0.005851 | 0.91657 | Other Positive Correlation |
| GTF2B | 0.005832 | 0.916848 | Other Positive Correlation |
| TNIP2 | 0.005821 | 0.917006 | Other Positive Correlation |
| IL1RL1 | 0.005808 | 0.91719 | Other Positive Correlation |
| KAT7 | 0.005777 | 0.917628 | Other Positive Correlation |
| PEX5 | 0.00577 | 0.917722 | Other Positive Correlation |
| CCDC153 | 0.005752 | 0.917987 | Other Positive Correlation |
| TRABD2B | 0.005726 | 0.918355 | Other Positive Correlation |
| GPR162 | 0.005703 | 0.918679 | Other Positive Correlation |
| ZP3 | 0.005691 | 0.918845 | Other Positive Correlation |
| MOB1B | 0.005684 | 0.918945 | Other Positive Correlation |
| CCR10 | 0.005637 | 0.919615 | Other Positive Correlation |
| GZF1 | 0.005595 | 0.920214 | Other Positive Correlation |
| ZNF649 | 0.005594 | 0.920232 | Other Positive Correlation |
| SSTR4 | 0.005548 | 0.920878 | Other Positive Correlation |
| ENSG00000267618 | 0.005548 | 0.920878 | Other Positive Correlation |
| NDP | 0.005548 | 0.920878 | Other Positive Correlation |
| DNAI2 | 0.005548 | 0.920878 | Other Positive Correlation |
| ENSG00000267314 | 0.005548 | 0.920878 | Other Positive Correlation |
| FGR | 0.005521 | 0.921266 | Other Positive Correlation |
| NHLH1 | 0.00545 | 0.922273 | Other Positive Correlation |
| HOXA3 | 0.005445 | 0.922339 | Other Positive Correlation |
| RAX | 0.005441 | 0.922407 | Other Positive Correlation |
| IHO1 | 0.005362 | 0.92353 | Other Positive Correlation |
| MEGF11 | 0.005334 | 0.92393 | Other Positive Correlation |
| STIMATE-MUSTN1 | 0.005312 | 0.924242 | Other Positive Correlation |
| OR2B8P | 0.005312 | 0.924242 | Other Positive Correlation |

| | | | |
|---|---|---|---|
| TEKT5 | 0.005312 | 0.924242 | Other Positive Correlation |
| GOLGA1 | 0.005307 | 0.924312 | Other Positive Correlation |
| PCMTD1 | 0.005256 | 0.925039 | Other Positive Correlation |
| WBP4 | 0.005235 | 0.925327 | Other Positive Correlation |
| ITK | 0.005226 | 0.925452 | Other Positive Correlation |
| CD84 | 0.005226 | 0.925452 | Other Positive Correlation |
| PLSCR2 | 0.005218 | 0.925579 | Other Positive Correlation |
| DYNLT3 | 0.005182 | 0.926083 | Other Positive Correlation |
| NUP133 | 0.005179 | 0.926122 | Other Positive Correlation |
| ZNF540 | 0.005164 | 0.926343 | Other Positive Correlation |
| SSH2 | 0.005137 | 0.926723 | Other Positive Correlation |
| CDX2 | 0.005121 | 0.926946 | Other Positive Correlation |
| CFAP161 | 0.005111 | 0.92709 | Other Positive Correlation |
| RAC3 | 0.005108 | 0.927132 | Other Positive Correlation |
| INPP4A | 0.005105 | 0.927174 | Other Positive Correlation |
| GARRE1 | 0.005069 | 0.927692 | Other Positive Correlation |
| OR10AC1 | 0.005068 | 0.927713 | Other Positive Correlation |
| CD96 | 0.005056 | 0.927874 | Other Positive Correlation |
| GDF5 | 0.005054 | 0.927911 | Other Positive Correlation |
| NPIPB2 | 0.005052 | 0.927932 | Other Positive Correlation |
| CYP27C1 | 0.005025 | 0.928323 | Other Positive Correlation |
| KLHDC1 | 0.005023 | 0.928341 | Other Positive Correlation |
| ANGPT1 | 0.005021 | 0.928372 | Other Positive Correlation |
| RAB18 | 0.005007 | 0.928577 | Other Positive Correlation |
| NDN | 0.005002 | 0.92865 | Other Positive Correlation |
| OTUD4 | 0.004995 | 0.928743 | Other Positive Correlation |
| USP47 | 0.004974 | 0.929049 | Other Positive Correlation |
| CHN2 | 0.004953 | 0.929347 | Other Positive Correlation |
| ATP6V1A | 0.004939 | 0.929548 | Other Positive Correlation |
| TESMIN | 0.004914 | 0.9299 | Other Positive Correlation |
| TMCO2 | 0.004905 | 0.93003 | Other Positive Correlation |
| LAMP1 | 0.004869 | 0.930542 | Other Positive Correlation |
| IL17RE | 0.004831 | 0.931078 | Other Positive Correlation |
| POT1 | 0.00483 | 0.931087 | Other Positive Correlation |
| NPIPB6 | 0.004828 | 0.931126 | Other Positive Correlation |
| MFSD2A | 0.004819 | 0.931243 | Other Positive Correlation |
| ELN | 0.004801 | 0.931501 | Other Positive Correlation |
| KCNH4 | 0.004801 | 0.931502 | Other Positive Correlation |
| AMHR2 | 0.004801 | 0.931502 | Other Positive Correlation |
| TOMM20L | 0.004757 | 0.932128 | Other Positive Correlation |
| ENTPD1 | 0.004738 | 0.932404 | Other Positive Correlation |
| IFT46 | 0.004699 | 0.932958 | Other Positive Correlation |
| STARD4 | 0.004696 | 0.933006 | Other Positive Correlation |
| ZNF510 | 0.004639 | 0.933805 | Other Positive Correlation |
| DAXX | 0.004615 | 0.934159 | Other Positive Correlation |
| TBC1D31 | 0.0046 | 0.934372 | Other Positive Correlation |
| MLIP | 0.004597 | 0.934403 | Other Positive Correlation |
| ZNF497 | 0.004591 | 0.934502 | Other Positive Correlation |

| | | | |
|---|---|---|---|
| TMEM178A | 0.004588 | 0.934534 | Other Positive Correlation |
| DGKG | 0.004583 | 0.934606 | Other Positive Correlation |
| MUL1 | 0.004556 | 0.934991 | Other Positive Correlation |
| TOR1AIP2 | 0.004548 | 0.935106 | Other Positive Correlation |
| BCL3 | 0.004511 | 0.935639 | Other Positive Correlation |
| TTC22 | 0.004474 | 0.936155 | Other Positive Correlation |
| GALK2 | 0.004467 | 0.936253 | Other Positive Correlation |
| ROS1 | 0.004418 | 0.936959 | Other Positive Correlation |
| NDUFAF7 | 0.004416 | 0.936986 | Other Positive Correlation |
| FOXC1 | 0.004399 | 0.93723 | Other Positive Correlation |
| B4GALNT2 | 0.004391 | 0.937342 | Other Positive Correlation |
| FAM83C | 0.004389 | 0.937375 | Other Positive Correlation |
| PCDHA11 | 0.004383 | 0.937458 | Other Positive Correlation |
| TMEM207 | 0.004383 | 0.937458 | Other Positive Correlation |
| KRT26 | 0.004383 | 0.937458 | Other Positive Correlation |
| KRTAP4-1 | 0.004383 | 0.937458 | Other Positive Correlation |
| SMIM28 | 0.004383 | 0.937458 | Other Positive Correlation |
| KRT27 | 0.004383 | 0.937458 | Other Positive Correlation |
| LCE2B | 0.004383 | 0.937458 | Other Positive Correlation |
| CCDC185 | 0.004383 | 0.937458 | Other Positive Correlation |
| SPINK9 | 0.004383 | 0.937458 | Other Positive Correlation |
| UCN3 | 0.004383 | 0.937458 | Other Positive Correlation |
| DSG4 | 0.004383 | 0.937458 | Other Positive Correlation |
| ASTN1 | 0.004383 | 0.937458 | Other Positive Correlation |
| SLC30A10 | 0.004383 | 0.937458 | Other Positive Correlation |
| CCDC54 | 0.004383 | 0.937458 | Other Positive Correlation |
| PCDHA8 | 0.004383 | 0.937458 | Other Positive Correlation |
| HPGDS | 0.004383 | 0.937458 | Other Positive Correlation |
| CD28 | 0.004383 | 0.937458 | Other Positive Correlation |
| BMP10 | 0.004383 | 0.937458 | Other Positive Correlation |
| CCR5 | 0.004383 | 0.937458 | Other Positive Correlation |
| IL22RA2 | 0.004383 | 0.937458 | Other Positive Correlation |
| TIGIT | 0.004383 | 0.937458 | Other Positive Correlation |
| MUC7 | 0.004383 | 0.937458 | Other Positive Correlation |
| SPRR2B | 0.004383 | 0.937458 | Other Positive Correlation |
| KRTAP17-1 | 0.004383 | 0.937458 | Other Positive Correlation |
| ENSG00000289503 | 0.004383 | 0.937458 | Other Positive Correlation |
| HHLA2 | 0.004383 | 0.937458 | Other Positive Correlation |
| ODAM | 0.004383 | 0.937458 | Other Positive Correlation |
| LIPM | 0.004383 | 0.937458 | Other Positive Correlation |
| LCE2C | 0.004383 | 0.937458 | Other Positive Correlation |
| CD5 | 0.004383 | 0.937458 | Other Positive Correlation |
| KRTAP4-11 | 0.004383 | 0.937458 | Other Positive Correlation |
| BFSP2 | 0.004383 | 0.937458 | Other Positive Correlation |
| KDM4C | 0.00436 | 0.937782 | Other Positive Correlation |
| EBF2 | 0.004313 | 0.938449 | Other Positive Correlation |
| RASGEF1C | 0.004313 | 0.938449 | Other Positive Correlation |
| TLR1 | 0.004306 | 0.93855 | Other Positive Correlation |

| | | | |
|---|---|---|---|
| WDFY1 | 0.004253 | 0.939308 | Other Positive Correlation |
| PLXNA4 | 0.004241 | 0.939485 | Other Positive Correlation |
| OCEL1 | 0.004225 | 0.939709 | Other Positive Correlation |
| GPR176 | 0.00422 | 0.939771 | Other Positive Correlation |
| CYSTM1 | 0.004211 | 0.939908 | Other Positive Correlation |
| SLC9A3R1 | 0.004211 | 0.939912 | Other Positive Correlation |
| LCOR | 0.00421 | 0.939914 | Other Positive Correlation |
| C17orf114 | 0.00421 | 0.939916 | Other Positive Correlation |
| SOGA1 | 0.004204 | 0.94001 | Other Positive Correlation |
| WRAP73 | 0.004161 | 0.940615 | Other Positive Correlation |
| SRP14 | 0.004156 | 0.94069 | Other Positive Correlation |
| RTL9 | 0.004147 | 0.940816 | Other Positive Correlation |
| MMP7 | 0.004134 | 0.941009 | Other Positive Correlation |
| UBQLN2 | 0.004132 | 0.941029 | Other Positive Correlation |
| ARL4A | 0.004052 | 0.942175 | Other Positive Correlation |
| C13orf46 | 0.004016 | 0.942687 | Other Positive Correlation |
| C11orf54 | 0.004012 | 0.942746 | Other Positive Correlation |
| EML4 | 0.003997 | 0.942951 | Other Positive Correlation |
| TTC13 | 0.003985 | 0.943132 | Other Positive Correlation |
| YIPF2 | 0.003956 | 0.943534 | Other Positive Correlation |
| FBXO27 | 0.003949 | 0.943642 | Other Positive Correlation |
| MYO1A | 0.003868 | 0.9448 | Other Positive Correlation |
| KLF8 | 0.003825 | 0.945402 | Other Positive Correlation |
| PANK1 | 0.003815 | 0.945555 | Other Positive Correlation |
| TMEM234 | 0.003776 | 0.946107 | Other Positive Correlation |
| NANOS3 | 0.003764 | 0.946279 | Other Positive Correlation |
| LDOC1 | 0.003755 | 0.9464 | Other Positive Correlation |
| LRRC56 | 0.003751 | 0.946458 | Other Positive Correlation |
| TCN2 | 0.003734 | 0.946696 | Other Positive Correlation |
| SLC10A6 | 0.00373 | 0.946766 | Other Positive Correlation |
| MTRF1 | 0.003692 | 0.947304 | Other Positive Correlation |
| REPS1 | 0.003689 | 0.947348 | Other Positive Correlation |
| ZNF544 | 0.003685 | 0.9474 | Other Positive Correlation |
| ZC3H12D | 0.00364 | 0.948039 | Other Positive Correlation |
| MLC1 | 0.003593 | 0.948712 | Other Positive Correlation |
| HORMAD2 | 0.003593 | 0.948712 | Other Positive Correlation |
| DUX4 | 0.003593 | 0.948712 | Other Positive Correlation |
| MRO | 0.003593 | 0.948712 | Other Positive Correlation |
| IL17REL | 0.00355 | 0.949324 | Other Positive Correlation |
| SSX5 | 0.00355 | 0.949324 | Other Positive Correlation |
| ZFP64 | 0.003526 | 0.94967 | Other Positive Correlation |
| KLHL6 | 0.003475 | 0.950387 | Other Positive Correlation |
| DHRS4 | 0.003473 | 0.950419 | Other Positive Correlation |
| ZDHHC16 | 0.003427 | 0.951076 | Other Positive Correlation |
| TCF20 | 0.003426 | 0.951089 | Other Positive Correlation |
| SYNE4 | 0.003383 | 0.951701 | Other Positive Correlation |
| MITD1 | 0.00337 | 0.951895 | Other Positive Correlation |
| PCDHA3 | 0.003366 | 0.951953 | Other Positive Correlation |

| | | | |
|---|---|---|---|
| CYP3A5 | 0.003334 | 0.952408 | Other Positive Correlation |
| RC3H2 | 0.00333 | 0.952466 | Other Positive Correlation |
| FREM3 | 0.003273 | 0.953268 | Other Positive Correlation |
| LRRC43 | 0.003239 | 0.953765 | Other Positive Correlation |
| REN | 0.003235 | 0.953814 | Other Positive Correlation |
| KLC4 | 0.003207 | 0.954218 | Other Positive Correlation |
| PCDHA6 | 0.003199 | 0.954323 | Other Positive Correlation |
| FAM185A | 0.003195 | 0.954381 | Other Positive Correlation |
| FABP7 | 0.003174 | 0.954688 | Other Positive Correlation |
| POM121L2 | 0.003165 | 0.954822 | Other Positive Correlation |
| TMEM102 | 0.003154 | 0.954975 | Other Positive Correlation |
| ERICH6 | 0.003135 | 0.955248 | Other Positive Correlation |
| CD302 | 0.003096 | 0.955801 | Other Positive Correlation |
| BEST3 | 0.003083 | 0.955977 | Other Positive Correlation |
| GPR89A | 0.003021 | 0.956874 | Other Positive Correlation |
| SART3 | 0.003009 | 0.957041 | Other Positive Correlation |
| MEX3B | 0.003002 | 0.957135 | Other Positive Correlation |
| PABIR2 | 0.002996 | 0.957222 | Other Positive Correlation |
| ENSG00000285942 | 0.002988 | 0.957344 | Other Positive Correlation |
| WWC3 | 0.002987 | 0.957347 | Other Positive Correlation |
| CXCL10 | 0.00294 | 0.958019 | Other Positive Correlation |
| CCDC63 | 0.002933 | 0.95813 | Other Positive Correlation |
| ZNF70 | 0.002927 | 0.958206 | Other Positive Correlation |
| LTC4S | 0.0029 | 0.958589 | Other Positive Correlation |
| FYB1 | 0.002899 | 0.95861 | Other Positive Correlation |
| TOR1B | 0.002893 | 0.958697 | Other Positive Correlation |
| HTATIP2 | 0.002882 | 0.958858 | Other Positive Correlation |
| ADSS1 | 0.002879 | 0.958894 | Other Positive Correlation |
| ERCC4 | 0.002872 | 0.958996 | Other Positive Correlation |
| GP5 | 0.002867 | 0.95907 | Other Positive Correlation |
| ANKRD1 | 0.002841 | 0.959439 | Other Positive Correlation |
| TTC6 | 0.002817 | 0.959782 | Other Positive Correlation |
| POC1B | 0.002806 | 0.959935 | Other Positive Correlation |
| DNAI7 | 0.002783 | 0.960269 | Other Positive Correlation |
| IQCE | 0.002753 | 0.960686 | Other Positive Correlation |
| PRKCD | 0.002724 | 0.961105 | Other Positive Correlation |
| ZC3H7A | 0.002695 | 0.961514 | Other Positive Correlation |
| LINC02210-CRHR1 | 0.002687 | 0.961631 | Other Positive Correlation |
| BPIFA2 | 0.002668 | 0.961906 | Other Positive Correlation |
| ENSG00000285082 | 0.002668 | 0.961906 | Other Positive Correlation |
| SCAF1 | 0.002655 | 0.962085 | Other Positive Correlation |
| PDE7A | 0.002609 | 0.962752 | Other Positive Correlation |
| ADRB3 | 0.002605 | 0.962807 | Other Positive Correlation |
| BIRC3 | 0.002587 | 0.963064 | Other Positive Correlation |
| SLC25A46 | 0.002579 | 0.963174 | Other Positive Correlation |
| KIAA0408 | 0.002575 | 0.963234 | Other Positive Correlation |
| ENSG00000285542 | 0.002575 | 0.963234 | Other Positive Correlation |
| ENSG00000269403 | 0.002575 | 0.963234 | Other Positive Correlation |

| | | | |
|---|---|---|---|
| ARHGAP42 | 0.002567 | 0.96335 | Other Positive Correlation |
| BRD2 | 0.002546 | 0.963638 | Other Positive Correlation |
| DNAH3 | 0.002515 | 0.964085 | Other Positive Correlation |
| SLC2A5 | 0.00249 | 0.96444 | Other Positive Correlation |
| WFDC1 | 0.002467 | 0.964773 | Other Positive Correlation |
| IGFN1 | 0.002446 | 0.965079 | Other Positive Correlation |
| GHITM | 0.002411 | 0.965565 | Other Positive Correlation |
| PPP5C | 0.0024 | 0.965733 | Other Positive Correlation |
| ABHD16A | 0.002385 | 0.965945 | Other Positive Correlation |
| HNRNPH2 | 0.002371 | 0.966147 | Other Positive Correlation |
| DPP6 | 0.002342 | 0.966558 | Other Positive Correlation |
| GJD3 | 0.002341 | 0.966564 | Other Positive Correlation |
| ANXA10 | 0.002338 | 0.966619 | Other Positive Correlation |
| USP51 | 0.002324 | 0.966819 | Other Positive Correlation |
| LHFPL7 | 0.002296 | 0.967216 | Other Positive Correlation |
| EFEMP2 | 0.002288 | 0.967323 | Other Positive Correlation |
| HBQ1 | 0.002285 | 0.967376 | Other Positive Correlation |
| STS | 0.002267 | 0.967633 | Other Positive Correlation |
| STK10 | 0.002261 | 0.967712 | Other Positive Correlation |
| P2RX5-TAX1BP3 | 0.002236 | 0.96807 | Other Positive Correlation |
| VPS13D | 0.002223 | 0.968258 | Other Positive Correlation |
| PTPA | 0.002206 | 0.968502 | Other Positive Correlation |
| CFAP210 | 0.002197 | 0.968623 | Other Positive Correlation |
| SIGLEC15 | 0.002174 | 0.968948 | Other Positive Correlation |
| CNST | 0.002174 | 0.96895 | Other Positive Correlation |
| PPP1R36 | 0.002162 | 0.969126 | Other Positive Correlation |
| ZNF526 | 0.002157 | 0.969199 | Other Positive Correlation |
| ENSG00000284762 | 0.002127 | 0.969631 | Other Positive Correlation |
| PTPN5 | 0.002127 | 0.969631 | Other Positive Correlation |
| CCL3 | 0.002127 | 0.969631 | Other Positive Correlation |
| ZPBP | 0.002127 | 0.969631 | Other Positive Correlation |
| SLC35F5 | 0.00212 | 0.969727 | Other Positive Correlation |
| RAET1G | 0.002098 | 0.970032 | Other Positive Correlation |
| RIGI | 0.002094 | 0.970101 | Other Positive Correlation |
| CDPF1 | 0.00208 | 0.970289 | Other Positive Correlation |
| GEMIN2 | 0.002002 | 0.971413 | Other Positive Correlation |
| PPP6R2 | 0.001991 | 0.971568 | Other Positive Correlation |
| EDN3 | 0.001983 | 0.971674 | Other Positive Correlation |
| GABRP | 0.001974 | 0.971803 | Other Positive Correlation |
| REST | 0.001953 | 0.972109 | Other Positive Correlation |
| SPRYD4 | 0.001952 | 0.972124 | Other Positive Correlation |
| TMEM232 | 0.001887 | 0.973054 | Other Positive Correlation |
| SACM1L | 0.001884 | 0.973095 | Other Positive Correlation |
| MB21D2 | 0.00187 | 0.97329 | Other Positive Correlation |
| TMEM260 | 0.001844 | 0.973671 | Other Positive Correlation |
| PCDH12 | 0.001819 | 0.974018 | Other Positive Correlation |
| TRAPPC13 | 0.001771 | 0.974713 | Other Positive Correlation |
| CCDC186 | 0.001671 | 0.976136 | Other Positive Correlation |

| | | | |
|---|---|---|---|
| TNFAIP8L1 | 0.00166 | 0.976285 | Other Positive Correlation |
| GBP2 | 0.001647 | 0.976473 | Other Positive Correlation |
| LANCL3 | 0.001646 | 0.976491 | Other Positive Correlation |
| CTAGE9 | 0.001641 | 0.976557 | Other Positive Correlation |
| SLC29A2 | 0.001636 | 0.976629 | Other Positive Correlation |
| MYOT | 0.001619 | 0.976881 | Other Positive Correlation |
| SP3 | 0.001618 | 0.976898 | Other Positive Correlation |
| PCDHA7 | 0.001617 | 0.976912 | Other Positive Correlation |
| SLC25A20 | 0.001616 | 0.976922 | Other Positive Correlation |
| MAP1A | 0.001568 | 0.977607 | Other Positive Correlation |
| TUBGCP4 | 0.001533 | 0.978107 | Other Positive Correlation |
| UBE2E1 | 0.001485 | 0.978794 | Other Positive Correlation |
| KDM4A | 0.001474 | 0.978945 | Other Positive Correlation |
| HLA-DRA | 0.001455 | 0.979223 | Other Positive Correlation |
| SLC35E3 | 0.001446 | 0.979349 | Other Positive Correlation |
| MROH6 | 0.001438 | 0.979468 | Other Positive Correlation |
| C18orf32 | 0.001437 | 0.979471 | Other Positive Correlation |
| RIOK2 | 0.001435 | 0.9795 | Other Positive Correlation |
| ATP6V1G1 | 0.001428 | 0.979607 | Other Positive Correlation |
| ANKRD20A1 | 0.001403 | 0.979958 | Other Positive Correlation |
| BTBD19 | 0.001382 | 0.980261 | Other Positive Correlation |
| GDI1 | 0.001373 | 0.98039 | Other Positive Correlation |
| SPATA12 | 0.001321 | 0.981139 | Other Positive Correlation |
| KLK8 | 0.001292 | 0.981541 | Other Positive Correlation |
| TRIB2 | 0.00127 | 0.981859 | Other Positive Correlation |
| WDR31 | 0.001257 | 0.982045 | Other Positive Correlation |
| FOXG1 | 0.001221 | 0.982566 | Other Positive Correlation |
| FAM237B | 0.001215 | 0.982639 | Other Positive Correlation |
| DNAH17 | 0.001202 | 0.982828 | Other Positive Correlation |
| LDLRAP1 | 0.001193 | 0.982959 | Other Positive Correlation |
| ZNG1B | 0.001184 | 0.983087 | Other Positive Correlation |
| TFDP2 | 0.001165 | 0.983358 | Other Positive Correlation |
| FLRT1 | 0.001137 | 0.983761 | Other Positive Correlation |
| ACCS | 0.001125 | 0.983935 | Other Positive Correlation |
| STOML1 | 0.001121 | 0.983982 | Other Positive Correlation |
| TTLL11 | 0.001109 | 0.984161 | Other Positive Correlation |
| HEBP1 | 0.001097 | 0.984331 | Other Positive Correlation |
| UGT2B15 | 0.001086 | 0.984489 | Other Positive Correlation |
| PIK3C2B | 0.001082 | 0.984551 | Other Positive Correlation |
| CDC5L | 0.001067 | 0.984763 | Other Positive Correlation |
| PEX12 | 0.001003 | 0.985672 | Other Positive Correlation |
| MYOZ1 | 0.001 | 0.98572 | Other Positive Correlation |
| TBK1 | 0.000939 | 0.986589 | Other Positive Correlation |
| ZNF776 | 0.000922 | 0.986836 | Other Positive Correlation |
| MTFR1 | 0.000913 | 0.986962 | Other Positive Correlation |
| PDZD8 | 0.000902 | 0.987118 | Other Positive Correlation |
| ENSG00000289791 | 0.000828 | 0.988173 | Other Positive Correlation |
| ZNF354C | 0.000785 | 0.988783 | Other Positive Correlation |

| | | | |
|---|---|---|---|
| POF1B | 0.000669 | 0.990442 | Other Positive Correlation |
| PHETA1 | 0.000629 | 0.991014 | Other Positive Correlation |
| IL2RG | 0.000609 | 0.991302 | Other Positive Correlation |
| CYB561A3 | 0.000561 | 0.991989 | Other Positive Correlation |
| ZNHIT3 | 0.000553 | 0.992108 | Other Positive Correlation |
| CHI3L1 | 0.000531 | 0.99242 | Other Positive Correlation |
| RBM15B | 0.00051 | 0.992713 | Other Positive Correlation |
| FUCA1 | 0.000501 | 0.992849 | Other Positive Correlation |
| CDC42EP2 | 0.000483 | 0.993098 | Other Positive Correlation |
| KIF25 | 0.000479 | 0.993151 | Other Positive Correlation |
| TDRKH | 0.000473 | 0.993244 | Other Positive Correlation |
| PITPNA | 0.000442 | 0.993682 | Other Positive Correlation |
| PTGS1 | 0.000433 | 0.993809 | Other Positive Correlation |
| CRYZL1 | 0.000432 | 0.993834 | Other Positive Correlation |
| ZDHHC15 | 0.000426 | 0.993909 | Other Positive Correlation |
| ZNF687 | 0.000409 | 0.994156 | Other Positive Correlation |
| IQUB | 0.000408 | 0.994171 | Other Positive Correlation |
| CHSY3 | 0.000378 | 0.994599 | Other Positive Correlation |
| MYLK4 | 0.000378 | 0.994608 | Other Positive Correlation |
| IL17RC | 0.000354 | 0.994949 | Other Positive Correlation |
| HCRTR2 | 0.000327 | 0.995323 | Other Positive Correlation |
| PLA2G6 | 0.000319 | 0.995449 | Other Positive Correlation |
| REC114 | 0.000274 | 0.996093 | Other Positive Correlation |
| EPHA5 | 0.000274 | 0.996093 | Other Positive Correlation |
| KRT71 | 0.000274 | 0.996093 | Other Positive Correlation |
| RGPD6 | 0.000274 | 0.996093 | Other Positive Correlation |
| CGA | 0.000274 | 0.996093 | Other Positive Correlation |
| ENSG00000285978 | 0.000274 | 0.996093 | Other Positive Correlation |
| ALG3 | 0.000245 | 0.996504 | Other Positive Correlation |
| MMP3 | 0.000226 | 0.996777 | Other Positive Correlation |
| CNTN2 | 0.000199 | 0.997161 | Other Positive Correlation |
| C15orf40 | 0.000163 | 0.997665 | Other Positive Correlation |
| ADGRE2 | 0.00015 | 0.997861 | Other Positive Correlation |
| ATRN | 0.00014 | 0.998002 | Other Positive Correlation |
| KCNH6 | 0.000135 | 0.998074 | Other Positive Correlation |
| ITLN2 | 0.000135 | 0.998074 | Other Positive Correlation |
| SLC25A2 | 0.000135 | 0.998074 | Other Positive Correlation |
| TPPP2 | 0.000135 | 0.998074 | Other Positive Correlation |
| KLRC3 | 0.000135 | 0.998074 | Other Positive Correlation |
| IMPG1 | 0.000135 | 0.998074 | Other Positive Correlation |
| FOXD4L6 | 0.000103 | 0.998524 | Other Positive Correlation |
| ZNF268 | 9.6E-05 | 0.998629 | Other Positive Correlation |
| RNF128 | 9.49E-05 | 0.998644 | Other Positive Correlation |
| HEPACAM2 | 7.65E-05 | 0.998908 | Other Positive Correlation |
| HAVCR2 | 6.85E-05 | 0.999022 | Other Positive Correlation |
| MRPS11 | 3.46E-05 | 0.999506 | Other Positive Correlation |
| BCL9 | 2.53E-05 | 0.999639 | Other Positive Correlation |
| ENSG00000284691 | -1.8E-05 | 0.99974 | Other Negative Correlation |

| | | | |
|---|---|---|---|
| RPRD1B | -2.8E-05 | 0.999593 | Other Negative Correlation |
| SYCE3 | -4.9E-05 | 0.999296 | Other Negative Correlation |
| MBD3 | -6.3E-05 | 0.999096 | Other Negative Correlation |
| FARP1 | -6.6E-05 | 0.999062 | Other Negative Correlation |
| ABCF3 | -6.7E-05 | 0.999037 | Other Negative Correlation |
| TRIM68 | -0.00011 | 0.998375 | Other Negative Correlation |
| PCSK1 | -0.00012 | 0.99829 | Other Negative Correlation |
| CPT1B | -0.00012 | 0.998233 | Other Negative Correlation |
| PSPN | -0.00012 | 0.998218 | Other Negative Correlation |
| ARL5B | -0.00014 | 0.998062 | Other Negative Correlation |
| CENPVL3 | -0.00015 | 0.997859 | Other Negative Correlation |
| SCARF2 | -0.00016 | 0.997688 | Other Negative Correlation |
| PCDHA12 | -0.00019 | 0.997285 | Other Negative Correlation |
| CCDC38 | -0.0002 | 0.997077 | Other Negative Correlation |
| ARSB | -0.00021 | 0.996936 | Other Negative Correlation |
| NOTCH2NLA | -0.00024 | 0.996637 | Other Negative Correlation |
| POU5F2 | -0.00026 | 0.996353 | Other Negative Correlation |
| CD7 | -0.00026 | 0.996255 | Other Negative Correlation |
| SHKBP1 | -0.00028 | 0.996023 | Other Negative Correlation |
| KDM3B | -0.00032 | 0.995409 | Other Negative Correlation |
| ADHFE1 | -0.00036 | 0.994857 | Other Negative Correlation |
| ZNF786 | -0.00038 | 0.994543 | Other Negative Correlation |
| SLC37A1 | -0.00041 | 0.994135 | Other Negative Correlation |
| ARFGAP1 | -0.00043 | 0.993884 | Other Negative Correlation |
| DDT | -0.00045 | 0.99355 | Other Negative Correlation |
| SEMA5A | -0.00046 | 0.993491 | Other Negative Correlation |
| SPINK8 | -0.00046 | 0.99344 | Other Negative Correlation |
| EXTL1 | -0.00047 | 0.993338 | Other Negative Correlation |
| TM7SF2 | -0.00049 | 0.993 | Other Negative Correlation |
| CDH12 | -0.00053 | 0.992404 | Other Negative Correlation |
| KBTBD6 | -0.00053 | 0.992376 | Other Negative Correlation |
| LATS1 | -0.00055 | 0.992078 | Other Negative Correlation |
| DDX3Y | -0.00057 | 0.991807 | Other Negative Correlation |
| MECP2 | -0.00059 | 0.991524 | Other Negative Correlation |
| MAP3K1 | -0.00059 | 0.991523 | Other Negative Correlation |
| PHTF1 | -0.00061 | 0.991232 | Other Negative Correlation |
| DHRS9 | -0.00065 | 0.99068 | Other Negative Correlation |
| ZSCAN10 | -0.00066 | 0.99055 | Other Negative Correlation |
| C5AR2 | -0.00067 | 0.990391 | Other Negative Correlation |
| APOBEC3H | -0.00068 | 0.990261 | Other Negative Correlation |
| ZFTA | -0.00069 | 0.990186 | Other Negative Correlation |
| ANKRD12 | -0.0007 | 0.989938 | Other Negative Correlation |
| RCBTB2 | -0.00073 | 0.989611 | Other Negative Correlation |
| CSTF2T | -0.00073 | 0.989589 | Other Negative Correlation |
| CEBPE | -0.00074 | 0.989438 | Other Negative Correlation |
| GIT2 | -0.00082 | 0.988283 | Other Negative Correlation |
| POLR2J3.1 | -0.00084 | 0.987995 | Other Negative Correlation |
| DEFB4A | -0.00086 | 0.987664 | Other Negative Correlation |

| | | | |
|---|---|---|---|
| APH1A | -0.00086 | 0.987648 | Other Negative Correlation |
| PARP6 | -0.00088 | 0.98743 | Other Negative Correlation |
| ARF1 | -0.0009 | 0.987202 | Other Negative Correlation |
| PSMD6 | -0.0009 | 0.987197 | Other Negative Correlation |
| SPRR2G | -0.00092 | 0.986816 | Other Negative Correlation |
| SUCLA2 | -0.00096 | 0.98628 | Other Negative Correlation |
| ZNF500 | -0.001 | 0.985726 | Other Negative Correlation |
| LRRC8D | -0.001 | 0.985704 | Other Negative Correlation |
| ZNF641 | -0.00101 | 0.985581 | Other Negative Correlation |
| ZNF324 | -0.00101 | 0.98552 | Other Negative Correlation |
| ZNF579 | -0.00105 | 0.984994 | Other Negative Correlation |
| FOXO4 | -0.00105 | 0.984934 | Other Negative Correlation |
| SMO | -0.00106 | 0.984796 | Other Negative Correlation |
| NPL | -0.00107 | 0.984726 | Other Negative Correlation |
| KASH5 | -0.0011 | 0.984296 | Other Negative Correlation |
| TMED6 | -0.00111 | 0.984078 | Other Negative Correlation |
| CCDC169 | -0.00117 | 0.983295 | Other Negative Correlation |
| GPATCH8 | -0.00117 | 0.983235 | Other Negative Correlation |
| GIMAP8 | -0.00118 | 0.98312 | Other Negative Correlation |
| ACOX2 | -0.00121 | 0.982737 | Other Negative Correlation |
| SLFN12 | -0.00121 | 0.982719 | Other Negative Correlation |
| OXCT2 | -0.00123 | 0.982423 | Other Negative Correlation |
| GAPDHS | -0.00125 | 0.982106 | Other Negative Correlation |
| TRIM64B | -0.00128 | 0.981761 | Other Negative Correlation |
| BOC | -0.00132 | 0.981122 | Other Negative Correlation |
| SLC9A5 | -0.00136 | 0.980549 | Other Negative Correlation |
| RGS22 | -0.00137 | 0.980395 | Other Negative Correlation |
| EXPH5 | -0.00141 | 0.979907 | Other Negative Correlation |
| SLC38A4 | -0.00142 | 0.979699 | Other Negative Correlation |
| MYOCOS | -0.00142 | 0.979655 | Other Negative Correlation |
| CNGA1 | -0.00144 | 0.97945 | Other Negative Correlation |
| WNT16 | -0.00146 | 0.979138 | Other Negative Correlation |
| SYT15B | -0.00147 | 0.979033 | Other Negative Correlation |
| FAM228A | -0.00148 | 0.978809 | Other Negative Correlation |
| MYEOV | -0.00151 | 0.978457 | Other Negative Correlation |
| MGP | -0.00151 | 0.978453 | Other Negative Correlation |
| ANKRD23 | -0.00153 | 0.978096 | Other Negative Correlation |
| ANXA13 | -0.00154 | 0.978068 | Other Negative Correlation |
| ZNF285 | -0.00154 | 0.978041 | Other Negative Correlation |
| BOLA2-SMG1P6 | -0.00154 | 0.977959 | Other Negative Correlation |
| MAN2B1 | -0.00155 | 0.97782 | Other Negative Correlation |
| SRY | -0.00156 | 0.977651 | Other Negative Correlation |
| HAUS3 | -0.0016 | 0.977193 | Other Negative Correlation |
| ALG12 | -0.00161 | 0.97703 | Other Negative Correlation |
| RPP21 | -0.00161 | 0.976939 | Other Negative Correlation |
| OBSL1 | -0.00162 | 0.976814 | Other Negative Correlation |
| TADA1 | -0.00163 | 0.976682 | Other Negative Correlation |
| PEX16 | -0.00163 | 0.976664 | Other Negative Correlation |

| | | | |
|---|---|---|---|
| B3GNT4 | -0.00165 | 0.976439 | Other Negative Correlation |
| MTMR12 | -0.00167 | 0.976192 | Other Negative Correlation |
| TPBGL | -0.00167 | 0.97612 | Other Negative Correlation |
| TRUB2 | -0.00171 | 0.975512 | Other Negative Correlation |
| PSMD3 | -0.00172 | 0.975435 | Other Negative Correlation |
| RRAGB | -0.00177 | 0.974764 | Other Negative Correlation |
| CAPS2 | -0.00177 | 0.974723 | Other Negative Correlation |
| SCYL2 | -0.00177 | 0.974704 | Other Negative Correlation |
| NTN4 | -0.00181 | 0.974155 | Other Negative Correlation |
| PRELP | -0.00185 | 0.973625 | Other Negative Correlation |
| RBPMS | -0.00187 | 0.973255 | Other Negative Correlation |
| DOCK10 | -0.00192 | 0.972566 | Other Negative Correlation |
| AKR1B10 | -0.00199 | 0.971636 | Other Negative Correlation |
| ZNF547 | -0.00199 | 0.971584 | Other Negative Correlation |
| ACTR5 | -0.00203 | 0.970944 | Other Negative Correlation |
| ZDHHC12 | -0.00204 | 0.970816 | Other Negative Correlation |
| IL17B | -0.00208 | 0.970343 | Other Negative Correlation |
| IFI27 | -0.00209 | 0.970101 | Other Negative Correlation |
| C19orf85 | -0.0021 | 0.970006 | Other Negative Correlation |
| ADRA2A | -0.0021 | 0.970006 | Other Negative Correlation |
| CNTN3 | -0.00212 | 0.969795 | Other Negative Correlation |
| CCL27 | -0.00224 | 0.968024 | Other Negative Correlation |
| IFIT5 | -0.00228 | 0.967405 | Other Negative Correlation |
| ALG6 | -0.00228 | 0.967398 | Other Negative Correlation |
| KCNN4 | -0.0023 | 0.967101 | Other Negative Correlation |
| AMTN | -0.0023 | 0.967095 | Other Negative Correlation |
| ZNF627 | -0.00232 | 0.966807 | Other Negative Correlation |
| RBM38 | -0.00233 | 0.966755 | Other Negative Correlation |
| MARS2 | -0.00235 | 0.966455 | Other Negative Correlation |
| HSH2D | -0.00237 | 0.96617 | Other Negative Correlation |
| ROBO3 | -0.00242 | 0.965432 | Other Negative Correlation |
| POU2F1 | -0.00247 | 0.964763 | Other Negative Correlation |
| PIGL | -0.00248 | 0.964634 | Other Negative Correlation |
| MS4A4A | -0.0025 | 0.964247 | Other Negative Correlation |
| ENSG00000284874 | -0.0025 | 0.964247 | Other Negative Correlation |
| TMEM135 | -0.00251 | 0.964205 | Other Negative Correlation |
| FAM8A1 | -0.00252 | 0.963963 | Other Negative Correlation |
| YTHDF1 | -0.00256 | 0.963468 | Other Negative Correlation |
| PAX9 | -0.00259 | 0.963056 | Other Negative Correlation |
| RABGAP1 | -0.00259 | 0.962963 | Other Negative Correlation |
| GTF3C5 | -0.0026 | 0.962909 | Other Negative Correlation |
| SDHAF3 | -0.00262 | 0.962556 | Other Negative Correlation |
| GSDMA | -0.00263 | 0.962398 | Other Negative Correlation |
| HSF5 | -0.00266 | 0.962013 | Other Negative Correlation |
| BTN3A2 | -0.00272 | 0.96121 | Other Negative Correlation |
| P4HA2 | -0.00273 | 0.960965 | Other Negative Correlation |
| UQCC1 | -0.00275 | 0.960757 | Other Negative Correlation |
| NOMO3 | -0.00278 | 0.960309 | Other Negative Correlation |

| | | | |
|---|---|---|---|
| ZNF28 | -0.00284 | 0.959418 | Other Negative Correlation |
| ZNF160 | -0.00287 | 0.959093 | Other Negative Correlation |
| AP1M1 | -0.00287 | 0.959062 | Other Negative Correlation |
| SNTB2 | -0.00287 | 0.958975 | Other Negative Correlation |
| SBNO2 | -0.00291 | 0.958437 | Other Negative Correlation |
| CAMK1 | -0.00292 | 0.958253 | Other Negative Correlation |
| PGAM2 | -0.00296 | 0.957799 | Other Negative Correlation |
| RTN1 | -0.00296 | 0.957694 | Other Negative Correlation |
| KLK13 | -0.00297 | 0.957596 | Other Negative Correlation |
| RNF34 | -0.00297 | 0.957589 | Other Negative Correlation |
| MARCO | -0.00301 | 0.957042 | Other Negative Correlation |
| KCTD17 | -0.00302 | 0.956888 | Other Negative Correlation |
| GLYATL1 | -0.00303 | 0.9567 | Other Negative Correlation |
| ZKSCAN8 | -0.00309 | 0.955921 | Other Negative Correlation |
| PCDHGA6 | -0.0031 | 0.955737 | Other Negative Correlation |
| H2BC8 | -0.00311 | 0.955608 | Other Negative Correlation |
| SLC13A5 | -0.00314 | 0.955105 | Other Negative Correlation |
| ZNF311 | -0.00316 | 0.954858 | Other Negative Correlation |
| C11orf16 | -0.00316 | 0.954842 | Other Negative Correlation |
| HYAL1 | -0.00317 | 0.95475 | Other Negative Correlation |
| NAT1 | -0.00319 | 0.954414 | Other Negative Correlation |
| APOL2 | -0.00322 | 0.953984 | Other Negative Correlation |
| PTPN21 | -0.0033 | 0.952909 | Other Negative Correlation |
| IFT80 | -0.00331 | 0.952715 | Other Negative Correlation |
| VILL | -0.00332 | 0.952573 | Other Negative Correlation |
| MMUT | -0.00336 | 0.952074 | Other Negative Correlation |
| HYAL4 | -0.00337 | 0.95183 | Other Negative Correlation |
| SNRNP200 | -0.00342 | 0.951205 | Other Negative Correlation |
| CHCHD1 | -0.00342 | 0.951175 | Other Negative Correlation |
| ZNF644 | -0.00343 | 0.951075 | Other Negative Correlation |
| PPP6C | -0.00344 | 0.950892 | Other Negative Correlation |
| MCTP2 | -0.00344 | 0.95083 | Other Negative Correlation |
| MMP17 | -0.00347 | 0.950508 | Other Negative Correlation |
| SYNC | -0.00352 | 0.949697 | Other Negative Correlation |
| PLD4 | -0.00354 | 0.949527 | Other Negative Correlation |
| UNC79 | -0.00357 | 0.949098 | Other Negative Correlation |
| KIAA2012 | -0.00358 | 0.948928 | Other Negative Correlation |
| BACE1 | -0.00359 | 0.948791 | Other Negative Correlation |
| SPIN2A | -0.00359 | 0.948743 | Other Negative Correlation |
| PTPN18 | -0.00362 | 0.948332 | Other Negative Correlation |
| E4F1 | -0.00363 | 0.948205 | Other Negative Correlation |
| CEP85 | -0.00363 | 0.948202 | Other Negative Correlation |
| LCE3C | -0.00363 | 0.948176 | Other Negative Correlation |
| PLAA | -0.00368 | 0.947507 | Other Negative Correlation |
| NPRL3 | -0.00377 | 0.946193 | Other Negative Correlation |
| SOST | -0.00382 | 0.945418 | Other Negative Correlation |
| ZNF845 | -0.00385 | 0.945009 | Other Negative Correlation |
| NOD1 | -0.00387 | 0.944773 | Other Negative Correlation |

| | | | |
|---|---|---|---|
| PPP2R2D | -0.00388 | 0.94462 | Other Negative Correlation |
| TMEM139 | -0.00389 | 0.944429 | Other Negative Correlation |
| CFAP299 | -0.0039 | 0.944362 | Other Negative Correlation |
| C9orf43 | -0.00391 | 0.944212 | Other Negative Correlation |
| ETFBKMT | -0.00393 | 0.943869 | Other Negative Correlation |
| CASQ1 | -0.00395 | 0.943646 | Other Negative Correlation |
| PCDHGA4 | -0.00397 | 0.94337 | Other Negative Correlation |
| IGFBP1 | -0.00402 | 0.942575 | Other Negative Correlation |
| FBXL19 | -0.00405 | 0.942168 | Other Negative Correlation |
| TUBGCP6 | -0.00408 | 0.941779 | Other Negative Correlation |
| TVP23B | -0.0041 | 0.941454 | Other Negative Correlation |
| HESX1 | -0.00411 | 0.941316 | Other Negative Correlation |
| SNPH | -0.00415 | 0.940705 | Other Negative Correlation |
| TAS2R20 | -0.00417 | 0.940514 | Other Negative Correlation |
| C5orf63 | -0.00417 | 0.940507 | Other Negative Correlation |
| HSBP1L1 | -0.00419 | 0.940247 | Other Negative Correlation |
| ZNF772 | -0.00419 | 0.940227 | Other Negative Correlation |
| PLXNC1 | -0.00419 | 0.940168 | Other Negative Correlation |
| ZNF830 | -0.0042 | 0.940126 | Other Negative Correlation |
| MTERF1 | -0.0042 | 0.940048 | Other Negative Correlation |
| NEDD4L | -0.0042 | 0.940015 | Other Negative Correlation |
| SERPINB1 | -0.00422 | 0.939763 | Other Negative Correlation |
| MAST3 | -0.00425 | 0.939344 | Other Negative Correlation |
| C7orf31 | -0.00426 | 0.939161 | Other Negative Correlation |
| BCL2L2 | -0.00435 | 0.937905 | Other Negative Correlation |
| GALNT8 | -0.00445 | 0.936518 | Other Negative Correlation |
| HSD11B2 | -0.00446 | 0.93639 | Other Negative Correlation |
| PCDHB7 | -0.00452 | 0.935481 | Other Negative Correlation |
| GABPA | -0.00452 | 0.935462 | Other Negative Correlation |
| ABCF2-H2BK1 | -0.00453 | 0.935292 | Other Negative Correlation |
| CLBA1 | -0.00455 | 0.935068 | Other Negative Correlation |
| ZNF800 | -0.00456 | 0.934954 | Other Negative Correlation |
| SHANK3 | -0.00459 | 0.934503 | Other Negative Correlation |
| PECR | -0.00464 | 0.933745 | Other Negative Correlation |
| RPAP1 | -0.00465 | 0.933678 | Other Negative Correlation |
| DMXL1 | -0.00466 | 0.933481 | Other Negative Correlation |
| SURF4 | -0.00467 | 0.933419 | Other Negative Correlation |
| MED16 | -0.00469 | 0.933019 | Other Negative Correlation |
| CCNYL1 | -0.00472 | 0.932639 | Other Negative Correlation |
| CDKN2AIP | -0.00473 | 0.9325 | Other Negative Correlation |
| ASAH2B | -0.00474 | 0.932304 | Other Negative Correlation |
| MOGS | -0.00475 | 0.932239 | Other Negative Correlation |
| TYW1 | -0.00478 | 0.931852 | Other Negative Correlation |
| PJA1 | -0.00478 | 0.931787 | Other Negative Correlation |
| APPL1 | -0.00479 | 0.931615 | Other Negative Correlation |
| LGALS9 | -0.0048 | 0.931493 | Other Negative Correlation |
| FAM114A2 | -0.00481 | 0.931417 | Other Negative Correlation |
| PPTC7 | -0.00486 | 0.930603 | Other Negative Correlation |

| | | | |
|---|---|---|---|
| CA9 | -0.00487 | 0.930527 | Other Negative Correlation |
| MMAA | -0.00493 | 0.929734 | Other Negative Correlation |
| CD2 | -0.00494 | 0.929541 | Other Negative Correlation |
| ZNF345 | -0.00495 | 0.92939 | Other Negative Correlation |
| MRPS31 | -0.00497 | 0.929136 | Other Negative Correlation |
| EGLN2 | -0.005 | 0.928707 | Other Negative Correlation |
| EFCAB12 | -0.005 | 0.928651 | Other Negative Correlation |
| ACP6 | -0.00505 | 0.927925 | Other Negative Correlation |
| ST6GAL1 | -0.00509 | 0.927374 | Other Negative Correlation |
| TLL2 | -0.00512 | 0.927008 | Other Negative Correlation |
| SEC22A | -0.00512 | 0.926981 | Other Negative Correlation |
| RP2 | -0.00513 | 0.926876 | Other Negative Correlation |
| PAM16 | -0.00513 | 0.926852 | Other Negative Correlation |
| STARD7 | -0.00516 | 0.926449 | Other Negative Correlation |
| MAPK8 | -0.00517 | 0.926284 | Other Negative Correlation |
| MAP3K19 | -0.00519 | 0.925948 | Other Negative Correlation |
| RBP2 | -0.00519 | 0.925948 | Other Negative Correlation |
| ARAP2 | -0.00521 | 0.925682 | Other Negative Correlation |
| PDIA2 | -0.00522 | 0.92558 | Other Negative Correlation |
| STIM1 | -0.00522 | 0.925501 | Other Negative Correlation |
| ARHGAP35 | -0.00526 | 0.92503 | Other Negative Correlation |
| RNF19B | -0.00532 | 0.924167 | Other Negative Correlation |
| FXYD2 | -0.00534 | 0.923879 | Other Negative Correlation |
| C5AR1 | -0.00535 | 0.923715 | Other Negative Correlation |
| DNAH10 | -0.00542 | 0.922764 | Other Negative Correlation |
| C19orf12 | -0.00542 | 0.922694 | Other Negative Correlation |
| VOPP1 | -0.00543 | 0.922518 | Other Negative Correlation |
| TH | -0.00545 | 0.922303 | Other Negative Correlation |
| PIGG | -0.00546 | 0.922119 | Other Negative Correlation |
| APOL3 | -0.00548 | 0.921839 | Other Negative Correlation |
| RAI14 | -0.00551 | 0.921479 | Other Negative Correlation |
| WDPCP | -0.00551 | 0.921403 | Other Negative Correlation |
| ASCL1 | -0.00555 | 0.920834 | Other Negative Correlation |
| ARHGAP17 | -0.00559 | 0.920275 | Other Negative Correlation |
| SLC6A13 | -0.00564 | 0.919614 | Other Negative Correlation |
| IFT81 | -0.00564 | 0.919538 | Other Negative Correlation |
| WDR49 | -0.00571 | 0.918543 | Other Negative Correlation |
| NINL | -0.00574 | 0.918199 | Other Negative Correlation |
| PDE2A | -0.00574 | 0.918191 | Other Negative Correlation |
| TCTN1 | -0.0058 | 0.917298 | Other Negative Correlation |
| STXBP2 | -0.00582 | 0.916945 | Other Negative Correlation |
| PCDHA10 | -0.00585 | 0.916575 | Other Negative Correlation |
| CEP72 | -0.00589 | 0.916068 | Other Negative Correlation |
| VAPB | -0.00594 | 0.915305 | Other Negative Correlation |
| ANAPC10 | -0.00595 | 0.9151 | Other Negative Correlation |
| ZRSR2 | -0.00596 | 0.915056 | Other Negative Correlation |
| MFSD4B | -0.00596 | 0.915012 | Other Negative Correlation |
| HTT | -0.00597 | 0.914838 | Other Negative Correlation |

| | | | |
|---|---|---|---|
| SP1 | -0.00599 | 0.914644 | Other Negative Correlation |
| ADGRG6 | -0.00601 | 0.914297 | Other Negative Correlation |
| PCDHA1 | -0.00602 | 0.914136 | Other Negative Correlation |
| AP5S1 | -0.00607 | 0.913419 | Other Negative Correlation |
| LIPH | -0.00608 | 0.913299 | Other Negative Correlation |
| CPEB2 | -0.00611 | 0.912963 | Other Negative Correlation |
| ENSG00000290146 | -0.00612 | 0.912817 | Other Negative Correlation |
| TTC30B | -0.00612 | 0.912767 | Other Negative Correlation |
| HEXD | -0.00613 | 0.912682 | Other Negative Correlation |
| PADI1 | -0.00617 | 0.912111 | Other Negative Correlation |
| ENSG00000261341 | -0.00617 | 0.91198 | Other Negative Correlation |
| RBM5 | -0.00617 | 0.911976 | Other Negative Correlation |
| KCNN3 | -0.0062 | 0.911612 | Other Negative Correlation |
| CLK4 | -0.00624 | 0.911117 | Other Negative Correlation |
| AP1B1 | -0.00627 | 0.910592 | Other Negative Correlation |
| SPINK13 | -0.00628 | 0.910414 | Other Negative Correlation |
| NSUN4 | -0.00629 | 0.910352 | Other Negative Correlation |
| PHF12 | -0.00631 | 0.91005 | Other Negative Correlation |
| ZPR1 | -0.00633 | 0.909748 | Other Negative Correlation |
| CYP4B1 | -0.00635 | 0.909461 | Other Negative Correlation |
| NPHP4 | -0.00637 | 0.909137 | Other Negative Correlation |
| C9orf152 | -0.00639 | 0.908943 | Other Negative Correlation |
| CYTH1 | -0.0064 | 0.908812 | Other Negative Correlation |
| STK38L | -0.00642 | 0.908539 | Other Negative Correlation |
| CAPS | -0.00644 | 0.908165 | Other Negative Correlation |
| NAAA | -0.00646 | 0.907913 | Other Negative Correlation |
| HAL | -0.00646 | 0.907898 | Other Negative Correlation |
| DAP | -0.00647 | 0.907775 | Other Negative Correlation |
| SP6 | -0.00649 | 0.907525 | Other Negative Correlation |
| ZFP57 | -0.00649 | 0.907438 | Other Negative Correlation |
| TTLL1 | -0.0065 | 0.907299 | Other Negative Correlation |
| FHDC1 | -0.00653 | 0.906892 | Other Negative Correlation |
| S100A9 | -0.00656 | 0.90658 | Other Negative Correlation |
| AP3S2 | -0.00656 | 0.906542 | Other Negative Correlation |
| ICAM1 | -0.00661 | 0.90579 | Other Negative Correlation |
| ABR | -0.00661 | 0.905779 | Other Negative Correlation |
| ZDHHC7 | -0.00667 | 0.90502 | Other Negative Correlation |
| SYCE1 | -0.00677 | 0.90357 | Other Negative Correlation |
| CIBAR2 | -0.00677 | 0.90357 | Other Negative Correlation |
| RFPL4B | -0.00677 | 0.90357 | Other Negative Correlation |
| CD86 | -0.00677 | 0.90357 | Other Negative Correlation |
| GLP2R | -0.00677 | 0.90357 | Other Negative Correlation |
| ALMS1 | -0.00677 | 0.90353 | Other Negative Correlation |
| IQCH | -0.00678 | 0.903449 | Other Negative Correlation |
| HSPBAP1 | -0.0068 | 0.903162 | Other Negative Correlation |
| IST1 | -0.00685 | 0.902435 | Other Negative Correlation |
| PLD1 | -0.00686 | 0.902229 | Other Negative Correlation |
| GPR78 | -0.00687 | 0.902084 | Other Negative Correlation |

| | | | |
|---|---|---|---|
| FAM180B | -0.00687 | 0.902084 | Other Negative Correlation |
| FKBP1C | -0.0069 | 0.901738 | Other Negative Correlation |
| PBXIP1 | -0.0069 | 0.9017 | Other Negative Correlation |
| C3AR1 | -0.00696 | 0.900768 | Other Negative Correlation |
| EPB41L2 | -0.00697 | 0.900766 | Other Negative Correlation |
| SERPINB9 | -0.00698 | 0.900624 | Other Negative Correlation |
| QRICH2 | -0.00699 | 0.900404 | Other Negative Correlation |
| PELI1 | -0.007 | 0.900273 | Other Negative Correlation |
| APOBEC2 | -0.00701 | 0.900109 | Other Negative Correlation |
| COMMD10 | -0.00706 | 0.899451 | Other Negative Correlation |
| C4orf36 | -0.00708 | 0.899108 | Other Negative Correlation |
| ATG4B | -0.00709 | 0.899064 | Other Negative Correlation |
| ENSG00000289701 | -0.00711 | 0.898647 | Other Negative Correlation |
| ATXN7 | -0.00712 | 0.898592 | Other Negative Correlation |
| BTBD16 | -0.00713 | 0.898468 | Other Negative Correlation |
| MED15 | -0.00716 | 0.898048 | Other Negative Correlation |
| RTEL1 | -0.00718 | 0.897705 | Other Negative Correlation |
| RAB8B | -0.00721 | 0.897296 | Other Negative Correlation |
| HYDIN | -0.00722 | 0.897154 | Other Negative Correlation |
| EIF4ENIF1 | -0.00728 | 0.896302 | Other Negative Correlation |
| LIPT2 | -0.00729 | 0.896201 | Other Negative Correlation |
| DISC1 | -0.00731 | 0.89594 | Other Negative Correlation |
| YY1 | -0.00732 | 0.895801 | Other Negative Correlation |
| HEATR4 | -0.00734 | 0.895445 | Other Negative Correlation |
| MEF2D | -0.00734 | 0.895434 | Other Negative Correlation |
| HSD17B3 | -0.00742 | 0.894273 | Other Negative Correlation |
| SH3BP4 | -0.00743 | 0.89416 | Other Negative Correlation |
| NANOG | -0.00746 | 0.893778 | Other Negative Correlation |
| AJM1 | -0.00746 | 0.893714 | Other Negative Correlation |
| ENSG00000251537 | -0.00749 | 0.893343 | Other Negative Correlation |
| RIPPLY2 | -0.0075 | 0.893226 | Other Negative Correlation |
| SOCS5 | -0.0075 | 0.893132 | Other Negative Correlation |
| DCHS2 | -0.00751 | 0.893021 | Other Negative Correlation |
| EPB41L5 | -0.00755 | 0.892528 | Other Negative Correlation |
| BICC1 | -0.00755 | 0.892463 | Other Negative Correlation |
| IMMP1L | -0.00756 | 0.89233 | Other Negative Correlation |
| ZBTB3 | -0.00761 | 0.891689 | Other Negative Correlation |
| ISG20 | -0.0077 | 0.890409 | Other Negative Correlation |
| ARHGEF40 | -0.0077 | 0.890368 | Other Negative Correlation |
| SLC35B3 | -0.0077 | 0.890314 | Other Negative Correlation |
| DMPK | -0.00771 | 0.890261 | Other Negative Correlation |
| ANKRD7 | -0.00771 | 0.890244 | Other Negative Correlation |
| GFAP | -0.00772 | 0.89012 | Other Negative Correlation |
| SGTA | -0.00774 | 0.889823 | Other Negative Correlation |
| VSIG10L2 | -0.00775 | 0.889656 | Other Negative Correlation |
| MGA | -0.00775 | 0.88959 | Other Negative Correlation |
| IFIT1B | -0.00777 | 0.889376 | Other Negative Correlation |
| CFAP58 | -0.00777 | 0.889299 | Other Negative Correlation |

| | | | |
|---|---|---|---|
| ZBTB42 | -0.00779 | 0.889072 | Other Negative Correlation |
| TAB1 | -0.00784 | 0.88833 | Other Negative Correlation |
| LMAN2 | -0.00785 | 0.888179 | Other Negative Correlation |
| KCNK7 | -0.00788 | 0.887881 | Other Negative Correlation |
| RAB3D | -0.00788 | 0.887795 | Other Negative Correlation |
| GAA | -0.00789 | 0.887716 | Other Negative Correlation |
| POLDIP3 | -0.00792 | 0.887175 | Other Negative Correlation |
| POLDIP2 | -0.00797 | 0.886498 | Other Negative Correlation |
| CEACAM19 | -0.00797 | 0.886473 | Other Negative Correlation |
| PCDHA5 | -0.00799 | 0.886318 | Other Negative Correlation |
| SLC5A6 | -0.00802 | 0.885787 | Other Negative Correlation |
| KCNJ1 | -0.00804 | 0.8856 | Other Negative Correlation |
| BST1 | -0.00804 | 0.88559 | Other Negative Correlation |
| NEK5 | -0.00804 | 0.885542 | Other Negative Correlation |
| PCNX3 | -0.00805 | 0.885429 | Other Negative Correlation |
| IGSF9 | -0.0081 | 0.884738 | Other Negative Correlation |
| CHUK | -0.00811 | 0.88455 | Other Negative Correlation |
| DNAJA2 | -0.00811 | 0.884497 | Other Negative Correlation |
| SELPLG | -0.00813 | 0.884346 | Other Negative Correlation |
| SLC16A12 | -0.00813 | 0.88425 | Other Negative Correlation |
| ITPKC | -0.00819 | 0.883364 | Other Negative Correlation |
| ABHD18 | -0.00837 | 0.880835 | Other Negative Correlation |
| TBC1D8B | -0.00842 | 0.880227 | Other Negative Correlation |
| IL36RN | -0.00842 | 0.880212 | Other Negative Correlation |
| ETV3L | -0.00842 | 0.880193 | Other Negative Correlation |
| NEB | -0.00846 | 0.879625 | Other Negative Correlation |
| ADARB2 | -0.00849 | 0.879141 | Other Negative Correlation |
| GMFG | -0.00854 | 0.878555 | Other Negative Correlation |
| ATAD3C | -0.00855 | 0.878364 | Other Negative Correlation |
| CLCF1 | -0.00857 | 0.878119 | Other Negative Correlation |
| ZNF721 | -0.00857 | 0.878116 | Other Negative Correlation |
| ST3GAL6 | -0.00857 | 0.878035 | Other Negative Correlation |
| REC8 | -0.00861 | 0.877552 | Other Negative Correlation |
| CPM | -0.00861 | 0.877511 | Other Negative Correlation |
| COG3 | -0.00863 | 0.877165 | Other Negative Correlation |
| MLLT6 | -0.00864 | 0.877014 | Other Negative Correlation |
| THEGL | -0.00871 | 0.876128 | Other Negative Correlation |
| APOBEC3F | -0.00871 | 0.876085 | Other Negative Correlation |
| PITX2 | -0.00874 | 0.875677 | Other Negative Correlation |
| SUMF1 | -0.0088 | 0.87486 | Other Negative Correlation |
| TULP3 | -0.00882 | 0.874514 | Other Negative Correlation |
| MRAP | -0.00885 | 0.874055 | Other Negative Correlation |
| FAIM | -0.00889 | 0.873499 | Other Negative Correlation |
| ANKRD26 | -0.0089 | 0.873403 | Other Negative Correlation |
| ENSG00000267561 | -0.0089 | 0.87336 | Other Negative Correlation |
| ASAP1 | -0.00891 | 0.873269 | Other Negative Correlation |
| CDC42BPB | -0.00896 | 0.872583 | Other Negative Correlation |
| SULT1C4 | -0.00903 | 0.871621 | Other Negative Correlation |

| | | | |
|---|---|---|---|
| CEP70 | -0.00907 | 0.870978 | Other Negative Correlation |
| PLPP4 | -0.00909 | 0.870696 | Other Negative Correlation |
| IL20 | -0.0091 | 0.870598 | Other Negative Correlation |
| SLC25A43 | -0.0091 | 0.870593 | Other Negative Correlation |
| SMIM10L2A | -0.00914 | 0.870038 | Other Negative Correlation |
| PDP1 | -0.0092 | 0.869204 | Other Negative Correlation |
| ELAC1 | -0.0092 | 0.869168 | Other Negative Correlation |
| TTLL9 | -0.00921 | 0.868964 | Other Negative Correlation |
| ZNF280C | -0.00924 | 0.868628 | Other Negative Correlation |
| SLC22A15 | -0.00924 | 0.868548 | Other Negative Correlation |
| WDR81 | -0.00927 | 0.868234 | Other Negative Correlation |
| CLCA4 | -0.00928 | 0.868117 | Other Negative Correlation |
| C9orf24 | -0.00928 | 0.868053 | Other Negative Correlation |
| ODF2L | -0.00928 | 0.867983 | Other Negative Correlation |
| OPA1 | -0.00938 | 0.866583 | Other Negative Correlation |
| SLC25A38 | -0.00942 | 0.866075 | Other Negative Correlation |
| ABCA5 | -0.00944 | 0.86586 | Other Negative Correlation |
| FAT3 | -0.00945 | 0.865595 | Other Negative Correlation |
| SMYD4 | -0.00946 | 0.865568 | Other Negative Correlation |
| KANSL1L | -0.00948 | 0.865199 | Other Negative Correlation |
| ATPAF2 | -0.00948 | 0.865164 | Other Negative Correlation |
| STK36 | -0.00949 | 0.865052 | Other Negative Correlation |
| GMEB1 | -0.0095 | 0.864945 | Other Negative Correlation |
| PIK3CD | -0.00953 | 0.864597 | Other Negative Correlation |
| SPAG4 | -0.00955 | 0.864257 | Other Negative Correlation |
| ZNF225 | -0.00955 | 0.864249 | Other Negative Correlation |
| TBC1D25 | -0.00957 | 0.863898 | Other Negative Correlation |
| CREG2 | -0.00961 | 0.863458 | Other Negative Correlation |
| CRBN | -0.00964 | 0.863045 | Other Negative Correlation |
| SEC24C | -0.00968 | 0.862405 | Other Negative Correlation |
| MLX | -0.00969 | 0.862337 | Other Negative Correlation |
| PSG6 | -0.00969 | 0.862318 | Other Negative Correlation |
| RWDD2B | -0.00969 | 0.862262 | Other Negative Correlation |
| REM2 | -0.0097 | 0.862175 | Other Negative Correlation |
| RBM4 | -0.0097 | 0.862149 | Other Negative Correlation |
| PCDHGB6 | -0.00972 | 0.861912 | Other Negative Correlation |
| DPF3 | -0.00977 | 0.861155 | Other Negative Correlation |
| CLDN15 | -0.00978 | 0.861051 | Other Negative Correlation |
| PPP1R3B | -0.00979 | 0.860889 | Other Negative Correlation |
| ITGAM | -0.00985 | 0.859997 | Other Negative Correlation |
| F2R | -0.00986 | 0.859822 | Other Negative Correlation |
| NISCH | -0.00987 | 0.859763 | Other Negative Correlation |
| DROSHA | -0.00988 | 0.859655 | Other Negative Correlation |
| AGMAT | -0.00988 | 0.859633 | Other Negative Correlation |
| SEMA3B | -0.00991 | 0.859212 | Other Negative Correlation |
| BIK | -0.00991 | 0.859172 | Other Negative Correlation |
| CALML6 | -0.00991 | 0.859155 | Other Negative Correlation |
| NCF2 | -0.00997 | 0.858302 | Other Negative Correlation |

| | | | |
|---|---|---|---|
| NKX1-2 | -0.01 | 0.857855 | Other Negative Correlation |
| L3MBTL3 | -0.01005 | 0.857143 | Other Negative Correlation |
| CTNNA3 | -0.01006 | 0.857113 | Other Negative Correlation |
| MFSD14A | -0.01007 | 0.856959 | Other Negative Correlation |
| INSL4 | -0.01007 | 0.856888 | Other Negative Correlation |
| TUT4 | -0.01008 | 0.856756 | Other Negative Correlation |
| UGT2B7 | -0.0101 | 0.856496 | Other Negative Correlation |
| C9orf50 | -0.01014 | 0.855959 | Other Negative Correlation |
| ZNF57 | -0.01015 | 0.855764 | Other Negative Correlation |
| DHFR2 | -0.01015 | 0.855742 | Other Negative Correlation |
| MIDN | -0.01018 | 0.855439 | Other Negative Correlation |
| ZNF41 | -0.01024 | 0.85451 | Other Negative Correlation |
| NINJ2 | -0.01028 | 0.854027 | Other Negative Correlation |
| TRMT6 | -0.01029 | 0.853809 | Other Negative Correlation |
| ATP6V1H | -0.0103 | 0.853649 | Other Negative Correlation |
| BAAT | -0.01031 | 0.853607 | Other Negative Correlation |
| TFEC | -0.01032 | 0.853484 | Other Negative Correlation |
| NKPD1 | -0.01038 | 0.852585 | Other Negative Correlation |
| FADS1 | -0.01038 | 0.852515 | Other Negative Correlation |
| CASD1 | -0.01042 | 0.851967 | Other Negative Correlation |
| RNF207 | -0.01044 | 0.851788 | Other Negative Correlation |
| RSPH1 | -0.01044 | 0.851675 | Other Negative Correlation |
| RBM44 | -0.01045 | 0.851562 | Other Negative Correlation |
| ATP2B3 | -0.01048 | 0.851147 | Other Negative Correlation |
| SERPINA5 | -0.01048 | 0.851147 | Other Negative Correlation |
| UNC50 | -0.01057 | 0.849901 | Other Negative Correlation |
| RDH5 | -0.01058 | 0.849736 | Other Negative Correlation |
| GDPD2 | -0.01059 | 0.849626 | Other Negative Correlation |
| PGAP6 | -0.01062 | 0.849141 | Other Negative Correlation |
| ACKR3 | -0.01066 | 0.848663 | Other Negative Correlation |
| PBX2 | -0.01066 | 0.848661 | Other Negative Correlation |
| ASPHD1 | -0.01068 | 0.848423 | Other Negative Correlation |
| CELA3B | -0.01068 | 0.84837 | Other Negative Correlation |
| SALL2 | -0.01071 | 0.847994 | Other Negative Correlation |
| UBE2H | -0.01071 | 0.847896 | Other Negative Correlation |
| SLC38A10 | -0.01072 | 0.847871 | Other Negative Correlation |
| TPD52L1 | -0.01074 | 0.84749 | Other Negative Correlation |
| AKIRIN1 | -0.01076 | 0.847257 | Other Negative Correlation |
| ATP9B | -0.01081 | 0.846582 | Other Negative Correlation |
| PHYKPL | -0.01081 | 0.84657 | Other Negative Correlation |
| ZMAT2 | -0.01085 | 0.845946 | Other Negative Correlation |
| LHB | -0.01086 | 0.845855 | Other Negative Correlation |
| NSG1 | -0.01096 | 0.844402 | Other Negative Correlation |
| ENSG00000266302 | -0.01097 | 0.844276 | Other Negative Correlation |
| PRKCH | -0.01099 | 0.844066 | Other Negative Correlation |
| BTBD6 | -0.01101 | 0.843756 | Other Negative Correlation |
| IL22RA1 | -0.01101 | 0.843707 | Other Negative Correlation |
| GRB14 | -0.01103 | 0.843529 | Other Negative Correlation |

| | | | |
|---|---|---|---|
| EXD2 | -0.01104 | 0.843325 | Other Negative Correlation |
| SLC35F6 | -0.01113 | 0.842022 | Other Negative Correlation |
| AKAP8L | -0.01114 | 0.841903 | Other Negative Correlation |
| NCBP3 | -0.01115 | 0.841769 | Other Negative Correlation |
| IFT122 | -0.01117 | 0.841515 | Other Negative Correlation |
| SLC25A41 | -0.0112 | 0.841072 | Other Negative Correlation |
| STX17 | -0.01123 | 0.840613 | Other Negative Correlation |
| PLEKHG4 | -0.01124 | 0.840537 | Other Negative Correlation |
| SYDE2 | -0.01125 | 0.840437 | Other Negative Correlation |
| TMED7-TICAM2 | -0.01128 | 0.839989 | Other Negative Correlation |
| C1QTNF1 | -0.01128 | 0.839954 | Other Negative Correlation |
| ZNF660 | -0.01129 | 0.839887 | Other Negative Correlation |
| TRIM45 | -0.01131 | 0.839531 | Other Negative Correlation |
| PRKAB1 | -0.01132 | 0.839424 | Other Negative Correlation |
| JAKMIP3 | -0.01138 | 0.838622 | Other Negative Correlation |
| OS9 | -0.01146 | 0.837487 | Other Negative Correlation |
| ENSG00000272916 | -0.01146 | 0.837418 | Other Negative Correlation |
| NYNRIN | -0.01147 | 0.837279 | Other Negative Correlation |
| EHD3 | -0.01148 | 0.837123 | Other Negative Correlation |
| POU6F2 | -0.01149 | 0.836987 | Other Negative Correlation |
| SLC23A2 | -0.01152 | 0.836578 | Other Negative Correlation |
| RGPD8 | -0.01154 | 0.836332 | Other Negative Correlation |
| ZNF667 | -0.01162 | 0.835168 | Other Negative Correlation |
| ABL1 | -0.01167 | 0.834477 | Other Negative Correlation |
| BATF2 | -0.01168 | 0.834415 | Other Negative Correlation |
| SPAST | -0.01169 | 0.834207 | Other Negative Correlation |
| MMP1 | -0.01171 | 0.833949 | Other Negative Correlation |
| ADGRL2 | -0.01172 | 0.833827 | Other Negative Correlation |
| LRRC8E | -0.01173 | 0.833681 | Other Negative Correlation |
| BAG6 | -0.01176 | 0.833296 | Other Negative Correlation |
| PLA2G4F | -0.01176 | 0.833292 | Other Negative Correlation |
| MORN4 | -0.01177 | 0.83312 | Other Negative Correlation |
| ALDH8A1 | -0.01179 | 0.832838 | Other Negative Correlation |
| SCAF4 | -0.0118 | 0.832668 | Other Negative Correlation |
| CD300A | -0.01181 | 0.832599 | Other Negative Correlation |
| CMTR1 | -0.01184 | 0.832197 | Other Negative Correlation |
| PKIA | -0.01186 | 0.831886 | Other Negative Correlation |
| B3GALT5 | -0.01186 | 0.831861 | Other Negative Correlation |
| VPS33B | -0.01187 | 0.831739 | Other Negative Correlation |
| DUSP14 | -0.01187 | 0.831695 | Other Negative Correlation |
| TENT4B | -0.01189 | 0.831484 | Other Negative Correlation |
| ARL13B | -0.01199 | 0.830044 | Other Negative Correlation |
| MCHR1 | -0.012 | 0.829913 | Other Negative Correlation |
| CCL7 | -0.01202 | 0.829577 | Other Negative Correlation |
| LRRIQ3 | -0.01208 | 0.828734 | Other Negative Correlation |
| TRAM2 | -0.01212 | 0.828207 | Other Negative Correlation |
| IRF5 | -0.01218 | 0.827388 | Other Negative Correlation |
| LAMB4 | -0.01223 | 0.826746 | Other Negative Correlation |

| | | | |
|---|---|---|---|
| FGFR1OP2 | -0.01224 | 0.826575 | Other Negative Correlation |
| FBF1 | -0.01225 | 0.826364 | Other Negative Correlation |
| FBXL20 | -0.0123 | 0.825731 | Other Negative Correlation |
| DCAF11 | -0.01234 | 0.825213 | Other Negative Correlation |
| ARMS2 | -0.01234 | 0.825209 | Other Negative Correlation |
| PGPEP1L | -0.01234 | 0.825209 | Other Negative Correlation |
| RAD51D | -0.01238 | 0.824534 | Other Negative Correlation |
| GYG2 | -0.01239 | 0.824495 | Other Negative Correlation |
| SYNGR2 | -0.01243 | 0.823965 | Other Negative Correlation |
| ARMC12 | -0.01243 | 0.823959 | Other Negative Correlation |
| GIPC1 | -0.01244 | 0.823763 | Other Negative Correlation |
| VPS35L | -0.01244 | 0.823755 | Other Negative Correlation |
| ZDHHC9 | -0.01247 | 0.823386 | Other Negative Correlation |
| TEX48 | -0.01249 | 0.823039 | Other Negative Correlation |
| WASHC2C | -0.01251 | 0.822832 | Other Negative Correlation |
| HPS4 | -0.01251 | 0.822813 | Other Negative Correlation |
| CNPY2 | -0.01251 | 0.822803 | Other Negative Correlation |
| ZNF780B | -0.01255 | 0.822277 | Other Negative Correlation |
| UBE2D4 | -0.01259 | 0.821648 | Other Negative Correlation |
| KCNC1 | -0.0126 | 0.821496 | Other Negative Correlation |
| FAM229B | -0.01263 | 0.821041 | Other Negative Correlation |
| GBP4 | -0.01264 | 0.821017 | Other Negative Correlation |
| C12orf71 | -0.01266 | 0.820665 | Other Negative Correlation |
| TCAIM | -0.01267 | 0.820546 | Other Negative Correlation |
| VWA3B | -0.01267 | 0.820529 | Other Negative Correlation |
| ARSK | -0.01268 | 0.820478 | Other Negative Correlation |
| ADA | -0.01268 | 0.820377 | Other Negative Correlation |
| NSMCE2 | -0.0127 | 0.820172 | Other Negative Correlation |
| ADAM28 | -0.01276 | 0.819239 | Other Negative Correlation |
| GALNS | -0.01284 | 0.818202 | Other Negative Correlation |
| NAPB | -0.01284 | 0.818188 | Other Negative Correlation |
| MARVELD2 | -0.01284 | 0.818154 | Other Negative Correlation |
| RFC1 | -0.01292 | 0.817082 | Other Negative Correlation |
| SUDS3 | -0.01298 | 0.816273 | Other Negative Correlation |
| ZNF514 | -0.01298 | 0.816254 | Other Negative Correlation |
| FBXW10B | -0.01298 | 0.816213 | Other Negative Correlation |
| SOX7 | -0.013 | 0.815988 | Other Negative Correlation |
| PRPF18 | -0.01301 | 0.815836 | Other Negative Correlation |
| MYRF | -0.01301 | 0.815827 | Other Negative Correlation |
| STARD9 | -0.01301 | 0.815821 | Other Negative Correlation |
| FCHO1 | -0.01303 | 0.815549 | Other Negative Correlation |
| FOXK2 | -0.01313 | 0.814133 | Other Negative Correlation |
| TRAF4 | -0.01315 | 0.813897 | Other Negative Correlation |
| GRID2IP | -0.01321 | 0.813099 | Other Negative Correlation |
| MC1R | -0.01321 | 0.813092 | Other Negative Correlation |
| UNC5CL | -0.01323 | 0.812825 | Other Negative Correlation |
| ACY3 | -0.01331 | 0.811722 | Other Negative Correlation |
| PHC1 | -0.01332 | 0.811455 | Other Negative Correlation |

| | | | |
|---|---|---|---|
| SLC26A6 | -0.01337 | 0.810782 | Other Negative Correlation |
| ZNF628 | -0.01341 | 0.810274 | Other Negative Correlation |
| RNF187 | -0.01346 | 0.80961 | Other Negative Correlation |
| SENP8 | -0.01351 | 0.808886 | Other Negative Correlation |
| FOXE1 | -0.01354 | 0.808448 | Other Negative Correlation |
| WIPI2 | -0.01354 | 0.808395 | Other Negative Correlation |
| BIVM | -0.01355 | 0.808387 | Other Negative Correlation |
| OSBPL11 | -0.01355 | 0.80826 | Other Negative Correlation |
| SC5D | -0.01355 | 0.808258 | Other Negative Correlation |
| MCM9 | -0.01363 | 0.807196 | Other Negative Correlation |
| CFAP157 | -0.01363 | 0.807167 | Other Negative Correlation |
| DCBLD1 | -0.01367 | 0.806727 | Other Negative Correlation |
| SH3GLB2 | -0.01373 | 0.805803 | Other Negative Correlation |
| BROX | -0.01373 | 0.805775 | Other Negative Correlation |
| HELQ | -0.01376 | 0.80546 | Other Negative Correlation |
| CXorf58 | -0.01379 | 0.804933 | Other Negative Correlation |
| SEC22C | -0.01381 | 0.804651 | Other Negative Correlation |
| AHRR | -0.01384 | 0.80434 | Other Negative Correlation |
| MARCHF10 | -0.01386 | 0.803971 | Other Negative Correlation |
| MAGI1 | -0.01387 | 0.803899 | Other Negative Correlation |
| POLL | -0.01389 | 0.803603 | Other Negative Correlation |
| B3GALNT1 | -0.01397 | 0.802461 | Other Negative Correlation |
| IL34 | -0.01407 | 0.801098 | Other Negative Correlation |
| ZNF425 | -0.01408 | 0.800978 | Other Negative Correlation |
| SETD3 | -0.01409 | 0.80088 | Other Negative Correlation |
| ANG | -0.01409 | 0.800854 | Other Negative Correlation |
| DNLZ | -0.0141 | 0.800744 | Other Negative Correlation |
| DSC2 | -0.0141 | 0.800728 | Other Negative Correlation |
| TJAP1 | -0.0141 | 0.800691 | Other Negative Correlation |
| SLC10A7 | -0.01411 | 0.800546 | Other Negative Correlation |
| LACTB | -0.01413 | 0.80022 | Other Negative Correlation |
| BBOX1 | -0.01414 | 0.800154 | Other Negative Correlation |
| TTC21A | -0.01419 | 0.799447 | Other Negative Correlation |
| CD93 | -0.01421 | 0.799223 | Other Negative Correlation |
| PCDHB16 | -0.01421 | 0.799204 | Other Negative Correlation |
| ENSG00000285943 | -0.01421 | 0.799174 | Other Negative Correlation |
| PCDHA4 | -0.01422 | 0.798997 | Other Negative Correlation |
| MTLN | -0.01425 | 0.798685 | Other Negative Correlation |
| TSHZ2 | -0.01425 | 0.798644 | Other Negative Correlation |
| CASQ2 | -0.01434 | 0.797439 | Other Negative Correlation |
| TAS1R3 | -0.01434 | 0.79736 | Other Negative Correlation |
| KAT6A | -0.01438 | 0.796824 | Other Negative Correlation |
| HDDC3 | -0.01439 | 0.796731 | Other Negative Correlation |
| SGCB | -0.01441 | 0.796408 | Other Negative Correlation |
| MFSD11 | -0.01447 | 0.795605 | Other Negative Correlation |
| ORC4 | -0.01448 | 0.795498 | Other Negative Correlation |
| KRT2 | -0.0145 | 0.795147 | Other Negative Correlation |
| ITPR1 | -0.01452 | 0.794951 | Other Negative Correlation |

| | | | |
|---|---|---|---|
| NDUFA6 | -0.01452 | 0.794935 | Other Negative Correlation |
| PRB3 | -0.01452 | 0.79493 | Other Negative Correlation |
| SLC1A7 | -0.01452 | 0.79493 | Other Negative Correlation |
| GPATCH2 | -0.01452 | 0.794864 | Other Negative Correlation |
| RAB35 | -0.01457 | 0.794249 | Other Negative Correlation |
| ATXN2L | -0.0146 | 0.793794 | Other Negative Correlation |
| RHPN1 | -0.01462 | 0.793491 | Other Negative Correlation |
| ENSG00000285629 | -0.01464 | 0.793251 | Other Negative Correlation |
| FATE1 | -0.01464 | 0.793251 | Other Negative Correlation |
| TPGS2 | -0.01466 | 0.792999 | Other Negative Correlation |
| SPATS1 | -0.01466 | 0.792905 | Other Negative Correlation |
| RFX2 | -0.0147 | 0.792474 | Other Negative Correlation |
| PRSS21 | -0.01471 | 0.792212 | Other Negative Correlation |
| SMPD4 | -0.01474 | 0.791808 | Other Negative Correlation |
| TUBB3 | -0.01476 | 0.791612 | Other Negative Correlation |
| NSG2 | -0.01478 | 0.791325 | Other Negative Correlation |
| LY6D | -0.01482 | 0.790779 | Other Negative Correlation |
| ZW10 | -0.01483 | 0.790553 | Other Negative Correlation |
| EPB41L4B | -0.01485 | 0.790351 | Other Negative Correlation |
| SPEG | -0.01487 | 0.790013 | Other Negative Correlation |
| PCDHB6 | -0.01489 | 0.789724 | Other Negative Correlation |
| TUBA8 | -0.01493 | 0.789212 | Other Negative Correlation |
| SLC4A3 | -0.01496 | 0.788858 | Other Negative Correlation |
| RPS10-NUDT3 | -0.01496 | 0.788832 | Other Negative Correlation |
| C20orf141 | -0.01497 | 0.788675 | Other Negative Correlation |
| FAXDC2 | -0.01498 | 0.788511 | Other Negative Correlation |
| PHLPP2 | -0.01499 | 0.788391 | Other Negative Correlation |
| TMEM167A | -0.015 | 0.78832 | Other Negative Correlation |
| ASB7 | -0.015 | 0.78823 | Other Negative Correlation |
| ALOX12 | -0.01501 | 0.78809 | Other Negative Correlation |
| XRCC1 | -0.01508 | 0.787187 | Other Negative Correlation |
| LMTK2 | -0.0151 | 0.786879 | Other Negative Correlation |
| NPM2 | -0.0151 | 0.786868 | Other Negative Correlation |
| TMEM185A | -0.01513 | 0.786483 | Other Negative Correlation |
| C1QTNF6 | -0.01518 | 0.785756 | Other Negative Correlation |
| RAB1B | -0.01519 | 0.785675 | Other Negative Correlation |
| GINM1 | -0.01521 | 0.785431 | Other Negative Correlation |
| TMEM65 | -0.01522 | 0.785181 | Other Negative Correlation |
| GCN1 | -0.01523 | 0.785155 | Other Negative Correlation |
| SFXN5 | -0.01523 | 0.785146 | Other Negative Correlation |
| RASSF5 | -0.01523 | 0.785112 | Other Negative Correlation |
| H2AC19 | -0.01523 | 0.785045 | Other Negative Correlation |
| KIF7 | -0.01526 | 0.784757 | Other Negative Correlation |
| NDUFA4L2 | -0.01527 | 0.784532 | Other Negative Correlation |
| ZSWIM6 | -0.01528 | 0.784432 | Other Negative Correlation |
| DYRK1A | -0.01528 | 0.784399 | Other Negative Correlation |
| ZNF543 | -0.01535 | 0.783508 | Other Negative Correlation |
| CDC37L1 | -0.01535 | 0.783496 | Other Negative Correlation |

| | | | |
|---|---|---|---|
| ENSG00000276087 | -0.01535 | 0.783407 | Other Negative Correlation |
| TMEM256-PLSCR3 | -0.01535 | 0.783407 | Other Negative Correlation |
| KCNV2 | -0.01535 | 0.783407 | Other Negative Correlation |
| METTL27 | -0.01537 | 0.783127 | Other Negative Correlation |
| DCT | -0.01538 | 0.783077 | Other Negative Correlation |
| AGR3 | -0.01538 | 0.783077 | Other Negative Correlation |
| CCDC175 | -0.01538 | 0.783077 | Other Negative Correlation |
| IL1RAPL1 | -0.01543 | 0.7824 | Other Negative Correlation |
| KCTD19 | -0.01543 | 0.7824 | Other Negative Correlation |
| UGT3A1 | -0.01543 | 0.7824 | Other Negative Correlation |
| AGGF1 | -0.01544 | 0.782212 | Other Negative Correlation |
| ATG2B | -0.01547 | 0.781741 | Other Negative Correlation |
| AKAP3 | -0.01549 | 0.78154 | Other Negative Correlation |
| GIGYF2 | -0.01549 | 0.781525 | Other Negative Correlation |
| NLGN3 | -0.01554 | 0.780852 | Other Negative Correlation |
| YIPF6 | -0.01555 | 0.780763 | Other Negative Correlation |
| TMIGD3 | -0.01555 | 0.78068 | Other Negative Correlation |
| CLIC4 | -0.01558 | 0.780284 | Other Negative Correlation |
| MED12 | -0.0156 | 0.779971 | Other Negative Correlation |
| HHLA1 | -0.01561 | 0.779928 | Other Negative Correlation |
| WDR83OS | -0.01562 | 0.779753 | Other Negative Correlation |
| HPS6 | -0.01571 | 0.778451 | Other Negative Correlation |
| ANXA2R | -0.01572 | 0.778438 | Other Negative Correlation |
| NDUFA3 | -0.01575 | 0.77794 | Other Negative Correlation |
| SMIM6 | -0.01578 | 0.777506 | Other Negative Correlation |
| DIAPH2 | -0.01582 | 0.776929 | Other Negative Correlation |
| PACS1 | -0.01588 | 0.776186 | Other Negative Correlation |
| INPPL1 | -0.01594 | 0.775285 | Other Negative Correlation |
| CAMK2B | -0.01595 | 0.775197 | Other Negative Correlation |
| PPP2R5C | -0.01604 | 0.773968 | Other Negative Correlation |
| NRXN1 | -0.01605 | 0.773797 | Other Negative Correlation |
| BORCS7 | -0.01606 | 0.773754 | Other Negative Correlation |
| FANCD2OS | -0.01607 | 0.773564 | Other Negative Correlation |
| TMEM242 | -0.01608 | 0.77342 | Other Negative Correlation |
| KRT75 | -0.01612 | 0.772835 | Other Negative Correlation |
| CAMKK2 | -0.01614 | 0.77257 | Other Negative Correlation |
| LHX5 | -0.01614 | 0.772546 | Other Negative Correlation |
| GOLGA8A | -0.01615 | 0.772475 | Other Negative Correlation |
| KRT82 | -0.01616 | 0.77233 | Other Negative Correlation |
| PEX7 | -0.01621 | 0.771647 | Other Negative Correlation |
| LIMCH1 | -0.01624 | 0.771273 | Other Negative Correlation |
| ENSG00000285982 | -0.01624 | 0.771243 | Other Negative Correlation |
| PLA1A | -0.01624 | 0.771243 | Other Negative Correlation |
| ENSG00000235007 | -0.01625 | 0.771037 | Other Negative Correlation |
| THTPA | -0.01634 | 0.769857 | Other Negative Correlation |
| VKORC1 | -0.01635 | 0.76968 | Other Negative Correlation |
| DIRAS2 | -0.01637 | 0.769487 | Other Negative Correlation |
| HMGXB4 | -0.01639 | 0.769156 | Other Negative Correlation |

| | | | |
|---|---|---|---|
| MEX3C | -0.01641 | 0.768916 | Other Negative Correlation |
| UBR2 | -0.01643 | 0.7686 | Other Negative Correlation |
| GARIN4 | -0.01646 | 0.768257 | Other Negative Correlation |
| FMR1NB | -0.01649 | 0.767804 | Other Negative Correlation |
| NCOA3 | -0.01649 | 0.767765 | Other Negative Correlation |
| SYT16 | -0.0165 | 0.767742 | Other Negative Correlation |
| DNASE1 | -0.01652 | 0.767421 | Other Negative Correlation |
| CLDND2 | -0.01652 | 0.767418 | Other Negative Correlation |
| HEATR5B | -0.01654 | 0.767119 | Other Negative Correlation |
| OR6B3 | -0.01655 | 0.76706 | Other Negative Correlation |
| WAC | -0.01656 | 0.766907 | Other Negative Correlation |
| ACTR8 | -0.01656 | 0.766876 | Other Negative Correlation |
| NME8 | -0.01659 | 0.766437 | Other Negative Correlation |
| IL12B | -0.01659 | 0.766393 | Other Negative Correlation |
| ABCC5 | -0.01663 | 0.765902 | Other Negative Correlation |
| ZNF143 | -0.01665 | 0.76568 | Other Negative Correlation |
| NOXRED1 | -0.01667 | 0.765301 | Other Negative Correlation |
| SVEP1 | -0.01672 | 0.764615 | Other Negative Correlation |
| TBC1D14 | -0.01675 | 0.764209 | Other Negative Correlation |
| VN1R1 | -0.01676 | 0.764177 | Other Negative Correlation |
| C1orf100 | -0.01677 | 0.764021 | Other Negative Correlation |
| KIAA1586 | -0.01677 | 0.763927 | Other Negative Correlation |
| GORAB | -0.01684 | 0.762982 | Other Negative Correlation |
| ARID4A | -0.01685 | 0.762939 | Other Negative Correlation |
| NXPE2 | -0.01688 | 0.762435 | Other Negative Correlation |
| OPALIN | -0.01688 | 0.762435 | Other Negative Correlation |
| SSTR3 | -0.01688 | 0.762435 | Other Negative Correlation |
| AFDN | -0.01692 | 0.761979 | Other Negative Correlation |
| LHPP | -0.01695 | 0.761594 | Other Negative Correlation |
| IPO11 | -0.01695 | 0.761546 | Other Negative Correlation |
| LDB1 | -0.017 | 0.760879 | Other Negative Correlation |
| ROM1 | -0.01701 | 0.760648 | Other Negative Correlation |
| PPP2R1B | -0.01712 | 0.75919 | Other Negative Correlation |
| RAB24 | -0.01715 | 0.758822 | Other Negative Correlation |
| SLC25A34 | -0.01715 | 0.758767 | Other Negative Correlation |
| B3GAT2 | -0.0172 | 0.758121 | Other Negative Correlation |
| SLC9A9 | -0.01723 | 0.757724 | Other Negative Correlation |
| PSAPL1 | -0.01727 | 0.757237 | Other Negative Correlation |
| GPR157 | -0.01727 | 0.757176 | Other Negative Correlation |
| VPS50 | -0.01728 | 0.756977 | Other Negative Correlation |
| SPRYD3 | -0.01739 | 0.755474 | Other Negative Correlation |
| ZPBP2 | -0.01741 | 0.755309 | Other Negative Correlation |
| SHBG | -0.01741 | 0.755213 | Other Negative Correlation |
| CYP2J2 | -0.01742 | 0.755173 | Other Negative Correlation |
| LMO7 | -0.01747 | 0.754468 | Other Negative Correlation |
| ZFPL1 | -0.01747 | 0.754454 | Other Negative Correlation |
| CMYA5 | -0.01755 | 0.753397 | Other Negative Correlation |
| KANTR | -0.01757 | 0.753024 | Other Negative Correlation |

| | | | |
|---|---|---|---|
| ZC3H3 | -0.01759 | 0.752833 | Other Negative Correlation |
| ENSG00000269755 | -0.01759 | 0.752803 | Other Negative Correlation |
| FBXW5 | -0.01762 | 0.752471 | Other Negative Correlation |
| SPP2 | -0.01764 | 0.752097 | Other Negative Correlation |
| PFKFB2 | -0.01766 | 0.751895 | Other Negative Correlation |
| TATDN3 | -0.01767 | 0.751701 | Other Negative Correlation |
| ERICH1 | -0.01768 | 0.75158 | Other Negative Correlation |
| NDST2 | -0.01771 | 0.751214 | Other Negative Correlation |
| AKAP10 | -0.0178 | 0.749953 | Other Negative Correlation |
| CDH24 | -0.01781 | 0.749768 | Other Negative Correlation |
| FKBP3 | -0.01784 | 0.749366 | Other Negative Correlation |
| ATG16L2 | -0.01789 | 0.748731 | Other Negative Correlation |
| SNX31 | -0.01789 | 0.748694 | Other Negative Correlation |
| OPA3 | -0.01792 | 0.748377 | Other Negative Correlation |
| ZNF14 | -0.01796 | 0.747742 | Other Negative Correlation |
| TXNDC12 | -0.018 | 0.74729 | Other Negative Correlation |
| MRFAP1L1 | -0.018 | 0.747235 | Other Negative Correlation |
| PIN1 | -0.01802 | 0.747027 | Other Negative Correlation |
| FMN1 | -0.01804 | 0.746725 | Other Negative Correlation |
| TRPM6 | -0.01804 | 0.746668 | Other Negative Correlation |
| EXOC8 | -0.01804 | 0.746656 | Other Negative Correlation |
| ATF2 | -0.01805 | 0.746533 | Other Negative Correlation |
| TSPYL1 | -0.01811 | 0.745739 | Other Negative Correlation |
| GUCY2D | -0.01812 | 0.745655 | Other Negative Correlation |
| ABCA12 | -0.01816 | 0.745035 | Other Negative Correlation |
| HDAC11 | -0.0182 | 0.744495 | Other Negative Correlation |
| GARIN2 | -0.0182 | 0.744476 | Other Negative Correlation |
| ULK2 | -0.01822 | 0.74431 | Other Negative Correlation |
| ANKRD34A | -0.01824 | 0.743967 | Other Negative Correlation |
| KLHL18 | -0.01825 | 0.743899 | Other Negative Correlation |
| ZSCAN9 | -0.01825 | 0.743866 | Other Negative Correlation |
| ZC3H6 | -0.01825 | 0.743852 | Other Negative Correlation |
| RPRD1A | -0.01828 | 0.743488 | Other Negative Correlation |
| ZNF669 | -0.01829 | 0.743329 | Other Negative Correlation |
| ATP2B1 | -0.0183 | 0.743123 | Other Negative Correlation |
| H2BC5 | -0.01831 | 0.743019 | Other Negative Correlation |
| ZNF621 | -0.01834 | 0.74268 | Other Negative Correlation |
| TRAPPC8 | -0.01835 | 0.742502 | Other Negative Correlation |
| GOLGA3 | -0.01838 | 0.742075 | Other Negative Correlation |
| HINT3 | -0.01841 | 0.741685 | Other Negative Correlation |
| STON1 | -0.01847 | 0.740916 | Other Negative Correlation |
| KDM5C | -0.01848 | 0.740686 | Other Negative Correlation |
| NOXA1 | -0.01849 | 0.740583 | Other Negative Correlation |
| PPP1R3C | -0.01851 | 0.740353 | Other Negative Correlation |
| M6PR | -0.01853 | 0.740044 | Other Negative Correlation |
| SLC9A3R2 | -0.01856 | 0.739694 | Other Negative Correlation |
| NTAN1 | -0.01858 | 0.739348 | Other Negative Correlation |
| METTL6 | -0.01861 | 0.739048 | Other Negative Correlation |

| | | | |
|---|---|---|---|
| TIMMDC1 | -0.01862 | 0.738848 | Other Negative Correlation |
| CSF1 | -0.01863 | 0.738766 | Other Negative Correlation |
| ZBTB49 | -0.01863 | 0.738703 | Other Negative Correlation |
| MBOAT2 | -0.01865 | 0.738492 | Other Negative Correlation |
| SH3YL1 | -0.01869 | 0.737866 | Other Negative Correlation |
| ZNF195 | -0.01873 | 0.737398 | Other Negative Correlation |
| SRMS | -0.01878 | 0.736757 | Other Negative Correlation |
| CLIP1 | -0.01878 | 0.736633 | Other Negative Correlation |
| ZDHHC4 | -0.01885 | 0.73569 | Other Negative Correlation |
| SEMA3G | -0.01889 | 0.735139 | Other Negative Correlation |
| ZNF24 | -0.01892 | 0.734845 | Other Negative Correlation |
| ENSG00000273291 | -0.01892 | 0.734743 | Other Negative Correlation |
| COPG1 | -0.01893 | 0.734661 | Other Negative Correlation |
| ARHGAP30 | -0.01893 | 0.734626 | Other Negative Correlation |
| ADPGK | -0.01895 | 0.734369 | Other Negative Correlation |
| BGLAP | -0.01896 | 0.734277 | Other Negative Correlation |
| CTU1 | -0.01901 | 0.733627 | Other Negative Correlation |
| KCNE2 | -0.01902 | 0.733483 | Other Negative Correlation |
| MAF1 | -0.01903 | 0.733338 | Other Negative Correlation |
| ZNF888 | -0.01904 | 0.733233 | Other Negative Correlation |
| ZNF487 | -0.01905 | 0.73306 | Other Negative Correlation |
| BDKRB1 | -0.01906 | 0.732956 | Other Negative Correlation |
| ASXL3 | -0.01907 | 0.732838 | Other Negative Correlation |
| HDHD2 | -0.01908 | 0.732618 | Other Negative Correlation |
| TMEM88B | -0.0191 | 0.732393 | Other Negative Correlation |
| HLF | -0.01911 | 0.732235 | Other Negative Correlation |
| MGRN1 | -0.01916 | 0.731544 | Other Negative Correlation |
| CD164L2 | -0.01916 | 0.73151 | Other Negative Correlation |
| PCDHGB1 | -0.0192 | 0.731029 | Other Negative Correlation |
| PSG1 | -0.0192 | 0.731002 | Other Negative Correlation |
| BICRA | -0.01926 | 0.730259 | Other Negative Correlation |
| TECPR2 | -0.01928 | 0.729929 | Other Negative Correlation |
| USP19 | -0.0193 | 0.729734 | Other Negative Correlation |
| WASHC1 | -0.0193 | 0.729704 | Other Negative Correlation |
| TOR4A | -0.01932 | 0.72941 | Other Negative Correlation |
| HPS3 | -0.01939 | 0.728535 | Other Negative Correlation |
| WDR82 | -0.01939 | 0.728527 | Other Negative Correlation |
| NFASC | -0.01939 | 0.728451 | Other Negative Correlation |
| STX16 | -0.0194 | 0.72836 | Other Negative Correlation |
| ZNF823 | -0.01941 | 0.728152 | Other Negative Correlation |
| FCHO2 | -0.01942 | 0.728121 | Other Negative Correlation |
| PCDHGA10 | -0.01942 | 0.728068 | Other Negative Correlation |
| STX1B | -0.01943 | 0.727918 | Other Negative Correlation |
| C1S | -0.01945 | 0.727717 | Other Negative Correlation |
| IQCN | -0.01946 | 0.727553 | Other Negative Correlation |
| ROGDI | -0.0195 | 0.727049 | Other Negative Correlation |
| CHCHD10 | -0.01957 | 0.72606 | Other Negative Correlation |
| RUSF1 | -0.01962 | 0.725402 | Other Negative Correlation |

| | | | |
|---|---|---|---|
| JPH2 | -0.01962 | 0.725376 | Other Negative Correlation |
| TMEM266 | -0.0197 | 0.724318 | Other Negative Correlation |
| XG | -0.01971 | 0.724155 | Other Negative Correlation |
| KPNA3 | -0.01971 | 0.72413 | Other Negative Correlation |
| TGFB1 | -0.0198 | 0.722925 | Other Negative Correlation |
| TMED7 | -0.01981 | 0.722819 | Other Negative Correlation |
| ZBED9 | -0.01982 | 0.722711 | Other Negative Correlation |
| C11orf80 | -0.01988 | 0.721845 | Other Negative Correlation |
| DELE1 | -0.01992 | 0.721375 | Other Negative Correlation |
| C8orf48 | -0.01992 | 0.72135 | Other Negative Correlation |
| GPR183 | -0.02002 | 0.720068 | Other Negative Correlation |
| TIMM17A | -0.02003 | 0.719939 | Other Negative Correlation |
| HMBOX1 | -0.02005 | 0.719627 | Other Negative Correlation |
| IFIT2 | -0.02005 | 0.719582 | Other Negative Correlation |
| ZNF134 | -0.02008 | 0.719256 | Other Negative Correlation |
| SLC34A1 | -0.02015 | 0.718301 | Other Negative Correlation |
| CFAP206 | -0.02017 | 0.717999 | Other Negative Correlation |
| NBPF26 | -0.02017 | 0.717992 | Other Negative Correlation |
| YIPF4 | -0.0202 | 0.717562 | Other Negative Correlation |
| NDUFS7 | -0.02032 | 0.716009 | Other Negative Correlation |
| LRRC47 | -0.02034 | 0.715689 | Other Negative Correlation |
| ZNF707 | -0.02037 | 0.715323 | Other Negative Correlation |
| MKRN2 | -0.0204 | 0.714881 | Other Negative Correlation |
| ATP5MF-PTCD1 | -0.02041 | 0.714808 | Other Negative Correlation |
| ZNF596 | -0.02043 | 0.714509 | Other Negative Correlation |
| BBS2 | -0.02057 | 0.712687 | Other Negative Correlation |
| AANAT | -0.02059 | 0.712425 | Other Negative Correlation |
| ADCY2 | -0.02061 | 0.712166 | Other Negative Correlation |
| SLC25A17 | -0.02065 | 0.711548 | Other Negative Correlation |
| TK2 | -0.02071 | 0.710788 | Other Negative Correlation |
| EXOG | -0.02072 | 0.710712 | Other Negative Correlation |
| GZMB | -0.02076 | 0.710164 | Other Negative Correlation |
| MTX1 | -0.02085 | 0.708866 | Other Negative Correlation |
| CCNJL | -0.02086 | 0.708798 | Other Negative Correlation |
| GPATCH1 | -0.02086 | 0.708728 | Other Negative Correlation |
| OR10H1 | -0.02091 | 0.708165 | Other Negative Correlation |
| DYNLT5 | -0.02091 | 0.708124 | Other Negative Correlation |
| ANKRD65 | -0.02094 | 0.70768 | Other Negative Correlation |
| CLIP2 | -0.02097 | 0.707261 | Other Negative Correlation |
| RFX3 | -0.02109 | 0.705741 | Other Negative Correlation |
| NADK2 | -0.02109 | 0.70567 | Other Negative Correlation |
| LRMDA | -0.02112 | 0.705288 | Other Negative Correlation |
| KCNE3 | -0.02114 | 0.705068 | Other Negative Correlation |
| AMACR | -0.02118 | 0.704581 | Other Negative Correlation |
| XPNPEP1 | -0.02119 | 0.704415 | Other Negative Correlation |
| BST2 | -0.02121 | 0.704178 | Other Negative Correlation |
| RARG | -0.02122 | 0.704046 | Other Negative Correlation |
| ZC2HC1A | -0.02122 | 0.703958 | Other Negative Correlation |

| | | | |
|---|---|---|---|
| C17orf80 | -0.02124 | 0.703751 | Other Negative Correlation |
| HDAC6 | -0.02124 | 0.703742 | Other Negative Correlation |
| FAM20C | -0.02124 | 0.703726 | Other Negative Correlation |
| GSPT2 | -0.02125 | 0.703557 | Other Negative Correlation |
| IFT43 | -0.02131 | 0.702764 | Other Negative Correlation |
| AHSG | -0.02133 | 0.702573 | Other Negative Correlation |
| ADGRG5 | -0.02133 | 0.702495 | Other Negative Correlation |
| ANKRD31 | -0.02134 | 0.702442 | Other Negative Correlation |
| PITPNC1 | -0.02134 | 0.70238 | Other Negative Correlation |
| RANBP9 | -0.02137 | 0.701949 | Other Negative Correlation |
| MTRF1L | -0.02139 | 0.701759 | Other Negative Correlation |
| LRRC28 | -0.02142 | 0.701384 | Other Negative Correlation |
| MTO1 | -0.02142 | 0.701346 | Other Negative Correlation |
| C10orf88 | -0.02143 | 0.70114 | Other Negative Correlation |
| FKTN | -0.02152 | 0.699983 | Other Negative Correlation |
| KHNYN | -0.02157 | 0.699383 | Other Negative Correlation |
| DIPK1A | -0.02158 | 0.699191 | Other Negative Correlation |
| PIK3CA | -0.02158 | 0.699184 | Other Negative Correlation |
| ZAN | -0.0217 | 0.697557 | Other Negative Correlation |
| C14orf93 | -0.02171 | 0.697545 | Other Negative Correlation |
| HERC2 | -0.02174 | 0.697067 | Other Negative Correlation |
| MTARC2 | -0.02177 | 0.69669 | Other Negative Correlation |
| HECTD1 | -0.02177 | 0.696644 | Other Negative Correlation |
| TOMM70 | -0.02187 | 0.695374 | Other Negative Correlation |
| AGO4 | -0.0219 | 0.694943 | Other Negative Correlation |
| RAB30 | -0.02195 | 0.69431 | Other Negative Correlation |
| PIWIL1 | -0.02199 | 0.693783 | Other Negative Correlation |
| KRT74 | -0.02202 | 0.693374 | Other Negative Correlation |
| TRIM74 | -0.02202 | 0.693374 | Other Negative Correlation |
| CLDN19 | -0.02207 | 0.692732 | Other Negative Correlation |
| ZNF322 | -0.02207 | 0.692719 | Other Negative Correlation |
| TMC8 | -0.02207 | 0.692679 | Other Negative Correlation |
| ADGRF4 | -0.02208 | 0.69254 | Other Negative Correlation |
| PCDHB3 | -0.0221 | 0.692284 | Other Negative Correlation |
| PPP1R42 | -0.02214 | 0.691852 | Other Negative Correlation |
| TRIM66 | -0.0222 | 0.690974 | Other Negative Correlation |
| RTN2 | -0.02221 | 0.690847 | Other Negative Correlation |
| DCTN6 | -0.02222 | 0.690807 | Other Negative Correlation |
| PDZD2 | -0.02223 | 0.690584 | Other Negative Correlation |
| SSPN | -0.02227 | 0.690024 | Other Negative Correlation |
| ARG2 | -0.0223 | 0.689701 | Other Negative Correlation |
| EIF3CL | -0.02232 | 0.689408 | Other Negative Correlation |
| PLD5 | -0.02233 | 0.689239 | Other Negative Correlation |
| PAK3 | -0.02234 | 0.689157 | Other Negative Correlation |
| ZFP62 | -0.02238 | 0.688623 | Other Negative Correlation |
| MARCKSL1 | -0.02238 | 0.688589 | Other Negative Correlation |
| LRRC37A3 | -0.02241 | 0.688271 | Other Negative Correlation |
| PIERCE2 | -0.02241 | 0.688204 | Other Negative Correlation |

| | | | |
|---|---|---|---|
| FOXL1 | -0.02242 | 0.68814 | Other Negative Correlation |
| SOD3 | -0.02248 | 0.687373 | Other Negative Correlation |
| C9orf72 | -0.02248 | 0.687336 | Other Negative Correlation |
| USF3 | -0.0225 | 0.687109 | Other Negative Correlation |
| ZSCAN4 | -0.0225 | 0.686997 | Other Negative Correlation |
| PAX8 | -0.02256 | 0.686292 | Other Negative Correlation |
| FASTKD1 | -0.02257 | 0.686126 | Other Negative Correlation |
| MEIS3 | -0.02265 | 0.685079 | Other Negative Correlation |
| CEMIP2 | -0.02267 | 0.684859 | Other Negative Correlation |
| CD72 | -0.02269 | 0.6845 | Other Negative Correlation |
| STEAP3 | -0.02272 | 0.684135 | Other Negative Correlation |
| DTWD2 | -0.02282 | 0.682826 | Other Negative Correlation |
| DHRS2 | -0.02284 | 0.682532 | Other Negative Correlation |
| MSLN | -0.02287 | 0.682156 | Other Negative Correlation |
| USP5 | -0.02288 | 0.682039 | Other Negative Correlation |
| TP53 | -0.0229 | 0.681836 | Other Negative Correlation |
| CES4A | -0.02293 | 0.681454 | Other Negative Correlation |
| CCDC92 | -0.02293 | 0.681352 | Other Negative Correlation |
| CIPC | -0.02295 | 0.681146 | Other Negative Correlation |
| ZNF281 | -0.02296 | 0.681043 | Other Negative Correlation |
| HPS5 | -0.02297 | 0.680929 | Other Negative Correlation |
| PANO1 | -0.02297 | 0.680866 | Other Negative Correlation |
| GTF2E2 | -0.02303 | 0.68011 | Other Negative Correlation |
| SH3BP5 | -0.02305 | 0.679869 | Other Negative Correlation |
| SETD2 | -0.02306 | 0.679734 | Other Negative Correlation |
| ENSG00000277639 | -0.02319 | 0.678003 | Other Negative Correlation |
| MVB12A | -0.02319 | 0.677993 | Other Negative Correlation |
| WHAMM | -0.02322 | 0.677618 | Other Negative Correlation |
| MGAT4B | -0.02325 | 0.677198 | Other Negative Correlation |
| TLE5 | -0.02331 | 0.676473 | Other Negative Correlation |
| CFAP20DC | -0.02331 | 0.676407 | Other Negative Correlation |
| GOLGA8Q | -0.02344 | 0.674712 | Other Negative Correlation |
| KLHDC2 | -0.02345 | 0.674586 | Other Negative Correlation |
| RGL1 | -0.02348 | 0.674238 | Other Negative Correlation |
| PARP15 | -0.0235 | 0.673947 | Other Negative Correlation |
| SANBR | -0.02354 | 0.673352 | Other Negative Correlation |
| CDK16 | -0.02356 | 0.673111 | Other Negative Correlation |
| ADCY7 | -0.02362 | 0.672294 | Other Negative Correlation |
| ACAP3 | -0.02364 | 0.6721 | Other Negative Correlation |
| CYTL1 | -0.02366 | 0.671838 | Other Negative Correlation |
| MAEA | -0.02371 | 0.671151 | Other Negative Correlation |
| LRRC42 | -0.02377 | 0.670397 | Other Negative Correlation |
| LCA5L | -0.02378 | 0.670317 | Other Negative Correlation |
| PSG5 | -0.02382 | 0.669807 | Other Negative Correlation |
| SMARCAL1 | -0.02383 | 0.669655 | Other Negative Correlation |
| ZNF223 | -0.02384 | 0.669457 | Other Negative Correlation |
| C2orf15 | -0.02385 | 0.669357 | Other Negative Correlation |
| PRPF6 | -0.02386 | 0.669229 | Other Negative Correlation |

| | | | |
|---|---|---|---|
| BTG4 | -0.02391 | 0.668617 | Other Negative Correlation |
| PTPMT1 | -0.02394 | 0.668175 | Other Negative Correlation |
| MORN2 | -0.02394 | 0.668126 | Other Negative Correlation |
| TMEM138 | -0.024 | 0.667381 | Other Negative Correlation |
| CHID1 | -0.02401 | 0.667244 | Other Negative Correlation |
| TCEANC | -0.02407 | 0.666469 | Other Negative Correlation |
| ART3 | -0.02408 | 0.666362 | Other Negative Correlation |
| ANO10 | -0.0241 | 0.666106 | Other Negative Correlation |
| CHD2 | -0.0241 | 0.666078 | Other Negative Correlation |
| UBOX5 | -0.02411 | 0.665988 | Other Negative Correlation |
| ATAD2B | -0.02413 | 0.665701 | Other Negative Correlation |
| TRIM3 | -0.0242 | 0.664844 | Other Negative Correlation |
| ITGB5 | -0.02421 | 0.664609 | Other Negative Correlation |
| ZNF384 | -0.02423 | 0.664384 | Other Negative Correlation |
| CHST15 | -0.02427 | 0.663906 | Other Negative Correlation |
| ABCA1 | -0.02428 | 0.66376 | Other Negative Correlation |
| TMEM161B | -0.0243 | 0.66349 | Other Negative Correlation |
| GRIN1 | -0.02434 | 0.663005 | Other Negative Correlation |
| PADI3 | -0.0244 | 0.662173 | Other Negative Correlation |
| C12orf4 | -0.02441 | 0.662127 | Other Negative Correlation |
| MDFIC | -0.02442 | 0.661957 | Other Negative Correlation |
| GATAD2A | -0.02443 | 0.661848 | Other Negative Correlation |
| FGD3 | -0.02445 | 0.661586 | Other Negative Correlation |
| TBC1D23 | -0.02445 | 0.661492 | Other Negative Correlation |
| INF2 | -0.02446 | 0.661388 | Other Negative Correlation |
| ITPKB | -0.02452 | 0.6607 | Other Negative Correlation |
| C17orf99 | -0.02455 | 0.660187 | Other Negative Correlation |
| TMC6 | -0.02459 | 0.659788 | Other Negative Correlation |
| VSTM2L | -0.02463 | 0.659193 | Other Negative Correlation |
| KPNA4 | -0.02466 | 0.658857 | Other Negative Correlation |
| TTC21B | -0.02467 | 0.658739 | Other Negative Correlation |
| MTMR6 | -0.02471 | 0.658129 | Other Negative Correlation |
| RBAK | -0.02471 | 0.658123 | Other Negative Correlation |
| IL1R1 | -0.02472 | 0.658085 | Other Negative Correlation |
| CCDC154 | -0.02479 | 0.65711 | Other Negative Correlation |
| ASPSCR1 | -0.02482 | 0.656731 | Other Negative Correlation |
| PAN3 | -0.02484 | 0.656519 | Other Negative Correlation |
| TPCN2 | -0.02485 | 0.656404 | Other Negative Correlation |
| KCNC4 | -0.02485 | 0.656296 | Other Negative Correlation |
| CIBAR1 | -0.02486 | 0.656264 | Other Negative Correlation |
| NFAT5 | -0.02487 | 0.656087 | Other Negative Correlation |
| ZDHHC1 | -0.02496 | 0.654979 | Other Negative Correlation |
| BIRC2 | -0.025 | 0.654378 | Other Negative Correlation |
| HIPK2 | -0.02506 | 0.653619 | Other Negative Correlation |
| BRAT1 | -0.02509 | 0.653258 | Other Negative Correlation |
| MGAT5B | -0.0251 | 0.653132 | Other Negative Correlation |
| TRAPPC3L | -0.02513 | 0.65277 | Other Negative Correlation |
| PSMC2 | -0.02514 | 0.65262 | Other Negative Correlation |

| | | | |
|---|---|---|---|
| CCDC188 | -0.02515 | 0.652432 | Other Negative Correlation |
| PCDHGC3 | -0.02516 | 0.652409 | Other Negative Correlation |
| B3GNT6 | -0.02517 | 0.652248 | Other Negative Correlation |
| CXCR2 | -0.02517 | 0.652167 | Other Negative Correlation |
| MC4R | -0.02521 | 0.651711 | Other Negative Correlation |
| NOL4 | -0.02526 | 0.651062 | Other Negative Correlation |
| MMP10 | -0.02528 | 0.6508 | Other Negative Correlation |
| SNRPN | -0.02533 | 0.650107 | Other Negative Correlation |
| NR0B1 | -0.02537 | 0.649663 | Other Negative Correlation |
| TRIM37 | -0.02537 | 0.649645 | Other Negative Correlation |
| ITSN1 | -0.02537 | 0.649597 | Other Negative Correlation |
| FAM24B | -0.02539 | 0.649368 | Other Negative Correlation |
| FBXW2 | -0.0254 | 0.649279 | Other Negative Correlation |
| CAMP | -0.02544 | 0.648725 | Other Negative Correlation |
| TMEM19 | -0.02547 | 0.648379 | Other Negative Correlation |
| STAG1 | -0.0255 | 0.647967 | Other Negative Correlation |
| PHLDA3 | -0.02557 | 0.647017 | Other Negative Correlation |
| CCDC180 | -0.02561 | 0.646611 | Other Negative Correlation |
| TSPAN7 | -0.02563 | 0.646329 | Other Negative Correlation |
| L3MBTL1 | -0.02568 | 0.645649 | Other Negative Correlation |
| MEA1 | -0.02568 | 0.645639 | Other Negative Correlation |
| MARCHF6 | -0.0257 | 0.645414 | Other Negative Correlation |
| ENSG00000283886 | -0.02572 | 0.645075 | Other Negative Correlation |
| LGALS7 | -0.02573 | 0.645023 | Other Negative Correlation |
| KATNAL1 | -0.02573 | 0.64499 | Other Negative Correlation |
| MTMR3 | -0.02577 | 0.644467 | Other Negative Correlation |
| SPATA32 | -0.02578 | 0.644345 | Other Negative Correlation |
| NCKAP5 | -0.02582 | 0.643788 | Other Negative Correlation |
| LARS2 | -0.02584 | 0.643579 | Other Negative Correlation |
| SMIM8 | -0.02586 | 0.643345 | Other Negative Correlation |
| SCAF8 | -0.02589 | 0.642904 | Other Negative Correlation |
| VSIG2 | -0.0259 | 0.642804 | Other Negative Correlation |
| LGALSL | -0.0259 | 0.642775 | Other Negative Correlation |
| PDE5A | -0.02602 | 0.641284 | Other Negative Correlation |
| EBLN1 | -0.02604 | 0.640972 | Other Negative Correlation |
| PLCD1 | -0.02607 | 0.640664 | Other Negative Correlation |
| TMEM53 | -0.02611 | 0.640159 | Other Negative Correlation |
| TBXAS1 | -0.02616 | 0.639467 | Other Negative Correlation |
| PWWP3A | -0.02617 | 0.639302 | Other Negative Correlation |
| MRPL55 | -0.02618 | 0.639202 | Other Negative Correlation |
| SLC35A2 | -0.02618 | 0.639188 | Other Negative Correlation |
| TNFSF11 | -0.02621 | 0.638835 | Other Negative Correlation |
| SIN3A | -0.02631 | 0.637593 | Other Negative Correlation |
| EFCAB2 | -0.02631 | 0.637573 | Other Negative Correlation |
| TM2D1 | -0.02637 | 0.636828 | Other Negative Correlation |
| CAB39L | -0.02641 | 0.636312 | Other Negative Correlation |
| ZKSCAN8P1 | -0.02641 | 0.63623 | Other Negative Correlation |
| MORN5 | -0.02644 | 0.635969 | Other Negative Correlation |

| | | | |
|---|---|---|---|
| TMED1 | -0.02649 | 0.635315 | Other Negative Correlation |
| MAPKAPK2 | -0.02661 | 0.633776 | Other Negative Correlation |
| SS18 | -0.02662 | 0.633635 | Other Negative Correlation |
| SEC31B | -0.02663 | 0.633473 | Other Negative Correlation |
| EFL1 | -0.02667 | 0.633008 | Other Negative Correlation |
| CD52 | -0.02667 | 0.632991 | Other Negative Correlation |
| MC5R | -0.0267 | 0.632643 | Other Negative Correlation |
| MAP2K4 | -0.02674 | 0.63207 | Other Negative Correlation |
| RPAP3 | -0.02676 | 0.63182 | Other Negative Correlation |
| ZNF267 | -0.02677 | 0.631749 | Other Negative Correlation |
| HAPSTR1 | -0.02677 | 0.631718 | Other Negative Correlation |
| MAP4K2 | -0.02681 | 0.63116 | Other Negative Correlation |
| SIRT3 | -0.02682 | 0.631089 | Other Negative Correlation |
| POLR2M | -0.02686 | 0.630568 | Other Negative Correlation |
| C1orf127 | -0.02692 | 0.629845 | Other Negative Correlation |
| NVL | -0.02692 | 0.629767 | Other Negative Correlation |
| ERMAP | -0.02693 | 0.629649 | Other Negative Correlation |
| MRPS2 | -0.02697 | 0.629119 | Other Negative Correlation |
| RIN3 | -0.02704 | 0.628301 | Other Negative Correlation |
| USP24 | -0.02705 | 0.628158 | Other Negative Correlation |
| FBLN7 | -0.02708 | 0.627722 | Other Negative Correlation |
| TRIM26 | -0.02708 | 0.627709 | Other Negative Correlation |
| NDUFAF4 | -0.02712 | 0.627292 | Other Negative Correlation |
| MYZAP | -0.02712 | 0.62721 | Other Negative Correlation |
| CXCL8 | -0.02714 | 0.627052 | Other Negative Correlation |
| CASP10 | -0.02715 | 0.626886 | Other Negative Correlation |
| CCNB3 | -0.0272 | 0.62627 | Other Negative Correlation |
| THOP1 | -0.0272 | 0.626262 | Other Negative Correlation |
| PDCD6 | -0.02724 | 0.625707 | Other Negative Correlation |
| BCKDK | -0.02725 | 0.625548 | Other Negative Correlation |
| KCND1 | -0.02726 | 0.625519 | Other Negative Correlation |
| CHM | -0.02727 | 0.625366 | Other Negative Correlation |
| CBY2 | -0.02739 | 0.623793 | Other Negative Correlation |
| LPIN3 | -0.0274 | 0.623744 | Other Negative Correlation |
| TCTN2 | -0.02741 | 0.623603 | Other Negative Correlation |
| BRD3OS | -0.02741 | 0.623519 | Other Negative Correlation |
| MFSD14B | -0.02757 | 0.621531 | Other Negative Correlation |
| MAN2A1 | -0.02758 | 0.621354 | Other Negative Correlation |
| F8A1 | -0.02759 | 0.621272 | Other Negative Correlation |
| PCDHB13 | -0.02764 | 0.62064 | Other Negative Correlation |
| MT-CO2 | -0.02766 | 0.620372 | Other Negative Correlation |
| FGF1 | -0.02767 | 0.620327 | Other Negative Correlation |
| HACE1 | -0.02767 | 0.620266 | Other Negative Correlation |
| PNPLA6 | -0.02771 | 0.619808 | Other Negative Correlation |
| ZKSCAN7 | -0.02771 | 0.619737 | Other Negative Correlation |
| SEC24D | -0.02772 | 0.619669 | Other Negative Correlation |
| PDE12 | -0.02776 | 0.619175 | Other Negative Correlation |
| C12orf57 | -0.02777 | 0.619038 | Other Negative Correlation |

| | | | |
|---|---|---|---|
| SHTN1 | -0.02777 | 0.619008 | Other Negative Correlation |
| SGSM3 | -0.02777 | 0.61897 | Other Negative Correlation |
| HYAL3 | -0.02778 | 0.618937 | Other Negative Correlation |
| FRZB | -0.02778 | 0.618839 | Other Negative Correlation |
| PLXNA2 | -0.0278 | 0.618643 | Other Negative Correlation |
| ABCA6 | -0.0278 | 0.618572 | Other Negative Correlation |
| SLC9A2 | -0.02781 | 0.618542 | Other Negative Correlation |
| TSPAN6 | -0.02781 | 0.618514 | Other Negative Correlation |
| C12orf73 | -0.02782 | 0.618391 | Other Negative Correlation |
| BCLAF3 | -0.02783 | 0.618294 | Other Negative Correlation |
| SLC35G1 | -0.02784 | 0.618129 | Other Negative Correlation |
| TPP1 | -0.02784 | 0.618097 | Other Negative Correlation |
| SMYD3 | -0.02785 | 0.618024 | Other Negative Correlation |
| ZNF746 | -0.0279 | 0.617316 | Other Negative Correlation |
| FOXS1 | -0.02792 | 0.617089 | Other Negative Correlation |
| LRRC1 | -0.02798 | 0.616361 | Other Negative Correlation |
| TMEM64 | -0.02801 | 0.615967 | Other Negative Correlation |
| NEK10 | -0.02802 | 0.615906 | Other Negative Correlation |
| DIPK2A | -0.02803 | 0.615746 | Other Negative Correlation |
| ACYP2 | -0.02804 | 0.615609 | Other Negative Correlation |
| MCTP1 | -0.02813 | 0.614444 | Other Negative Correlation |
| KICS2 | -0.02814 | 0.614411 | Other Negative Correlation |
| SPTBN5 | -0.02814 | 0.614329 | Other Negative Correlation |
| MAPK8IP3 | -0.02814 | 0.614317 | Other Negative Correlation |
| PLAC8 | -0.02814 | 0.614302 | Other Negative Correlation |
| CUL4B | -0.02818 | 0.613865 | Other Negative Correlation |
| TRPM4 | -0.02821 | 0.613489 | Other Negative Correlation |
| BLOC1S1 | -0.02821 | 0.613421 | Other Negative Correlation |
| AAR2 | -0.02826 | 0.612813 | Other Negative Correlation |
| UEVLD | -0.02829 | 0.612504 | Other Negative Correlation |
| C6orf62 | -0.02837 | 0.611494 | Other Negative Correlation |
| MTUS1 | -0.02838 | 0.611283 | Other Negative Correlation |
| RGN | -0.02839 | 0.611167 | Other Negative Correlation |
| MRPS5 | -0.02844 | 0.610572 | Other Negative Correlation |
| DTX2 | -0.02848 | 0.610035 | Other Negative Correlation |
| RALB | -0.0285 | 0.609866 | Other Negative Correlation |
| SLC44A5 | -0.02853 | 0.609446 | Other Negative Correlation |
| IGFBP3 | -0.02855 | 0.609243 | Other Negative Correlation |
| SLC25A19 | -0.02857 | 0.609001 | Other Negative Correlation |
| ENSG00000260643 | -0.02857 | 0.608995 | Other Negative Correlation |
| GPN2 | -0.02858 | 0.608849 | Other Negative Correlation |
| CTSE | -0.02858 | 0.608843 | Other Negative Correlation |
| MED24 | -0.02859 | 0.608712 | Other Negative Correlation |
| KDM2B | -0.02866 | 0.60785 | Other Negative Correlation |
| MFSD3 | -0.02868 | 0.607609 | Other Negative Correlation |
| ACBD4 | -0.02868 | 0.607583 | Other Negative Correlation |
| TBC1D8 | -0.02871 | 0.607149 | Other Negative Correlation |
| ZNF358 | -0.02871 | 0.607133 | Other Negative Correlation |

| | | | |
|---|---|---|---|
| ZNF790 | -0.02872 | 0.607082 | Other Negative Correlation |
| TCEAL9 | -0.02872 | 0.60706 | Other Negative Correlation |
| C19orf54 | -0.02876 | 0.60657 | Other Negative Correlation |
| SIRPB1 | -0.02877 | 0.606459 | Other Negative Correlation |
| PCSK7 | -0.02878 | 0.606369 | Other Negative Correlation |
| EPN3 | -0.02879 | 0.606243 | Other Negative Correlation |
| RPS6KA2 | -0.02882 | 0.605847 | Other Negative Correlation |
| REV1 | -0.02882 | 0.605839 | Other Negative Correlation |
| TTC8 | -0.02882 | 0.605809 | Other Negative Correlation |
| TCAF2 | -0.02883 | 0.605689 | Other Negative Correlation |
| RELL1 | -0.02885 | 0.605501 | Other Negative Correlation |
| TRIM32 | -0.02898 | 0.603821 | Other Negative Correlation |
| RAMAC | -0.02898 | 0.603807 | Other Negative Correlation |
| PLA2G12A | -0.02903 | 0.603214 | Other Negative Correlation |
| RNF44 | -0.02911 | 0.602154 | Other Negative Correlation |
| IGLL1 | -0.02911 | 0.602132 | Other Negative Correlation |
| CLASP2 | -0.02917 | 0.601409 | Other Negative Correlation |
| DDHD1 | -0.0292 | 0.601082 | Other Negative Correlation |
| SPAG1 | -0.02921 | 0.600902 | Other Negative Correlation |
| CASZ1 | -0.02923 | 0.600663 | Other Negative Correlation |
| HACD3 | -0.02924 | 0.600618 | Other Negative Correlation |
| ALX4 | -0.02926 | 0.600289 | Other Negative Correlation |
| ISY1 | -0.02928 | 0.60001 | Other Negative Correlation |
| COG8 | -0.0293 | 0.599778 | Other Negative Correlation |
| IGSF8 | -0.02939 | 0.598734 | Other Negative Correlation |
| TMCC2 | -0.02939 | 0.598643 | Other Negative Correlation |
| KATNBL1 | -0.02941 | 0.598491 | Other Negative Correlation |
| ANKIB1 | -0.02944 | 0.598038 | Other Negative Correlation |
| BRWD1 | -0.02951 | 0.597176 | Other Negative Correlation |
| EBI3 | -0.02954 | 0.596865 | Other Negative Correlation |
| CES1 | -0.02956 | 0.596646 | Other Negative Correlation |
| C12orf60 | -0.02958 | 0.596389 | Other Negative Correlation |
| CLHC1 | -0.02959 | 0.596188 | Other Negative Correlation |
| RPTOR | -0.02962 | 0.59585 | Other Negative Correlation |
| LIPK | -0.02972 | 0.594541 | Other Negative Correlation |
| PCDHGB5 | -0.02974 | 0.594388 | Other Negative Correlation |
| KLHL9 | -0.02974 | 0.594375 | Other Negative Correlation |
| CYBB | -0.02976 | 0.594054 | Other Negative Correlation |
| RFNG | -0.02978 | 0.593859 | Other Negative Correlation |
| ZNF534 | -0.02979 | 0.593722 | Other Negative Correlation |
| EDC3 | -0.02979 | 0.593716 | Other Negative Correlation |
| SEMA7A | -0.02986 | 0.592838 | Other Negative Correlation |
| UGT8 | -0.02987 | 0.59275 | Other Negative Correlation |
| RIMS2 | -0.02987 | 0.592687 | Other Negative Correlation |
| SDCBP | -0.02991 | 0.592286 | Other Negative Correlation |
| PPP1R8 | -0.02994 | 0.591919 | Other Negative Correlation |
| GET1 | -0.02994 | 0.591889 | Other Negative Correlation |
| ADGRE5 | -0.02994 | 0.591839 | Other Negative Correlation |

| | | | |
|---|---|---|---|
| HS2ST1 | -0.02995 | 0.591805 | Other Negative Correlation |
| TCTA | -0.02997 | 0.591495 | Other Negative Correlation |
| RHOXF1 | -0.03003 | 0.590738 | Other Negative Correlation |
| APBB3 | -0.03003 | 0.590706 | Other Negative Correlation |
| SLC30A9 | -0.03005 | 0.590527 | Other Negative Correlation |
| ARHGAP5 | -0.03007 | 0.590235 | Other Negative Correlation |
| FAM135A | -0.03008 | 0.590145 | Other Negative Correlation |
| RAB20 | -0.03009 | 0.59 | Other Negative Correlation |
| FEM1B | -0.03013 | 0.589487 | Other Negative Correlation |
| OR2B11 | -0.03015 | 0.589219 | Other Negative Correlation |
| INHBC | -0.03015 | 0.589219 | Other Negative Correlation |
| ARSG | -0.03017 | 0.589007 | Other Negative Correlation |
| CCR3 | -0.0302 | 0.588714 | Other Negative Correlation |
| SIAE | -0.03022 | 0.588355 | Other Negative Correlation |
| PGLYRP4 | -0.03027 | 0.587851 | Other Negative Correlation |
| KCTD3 | -0.03028 | 0.587714 | Other Negative Correlation |
| ZNF770 | -0.03031 | 0.587316 | Other Negative Correlation |
| ASIC3 | -0.03032 | 0.587217 | Other Negative Correlation |
| UBE2F | -0.03034 | 0.586894 | Other Negative Correlation |
| COL13A1 | -0.03035 | 0.586815 | Other Negative Correlation |
| CFAP95 | -0.03035 | 0.586791 | Other Negative Correlation |
| RPL3L | -0.03037 | 0.586556 | Other Negative Correlation |
| CIC | -0.03057 | 0.584149 | Other Negative Correlation |
| CPD | -0.03057 | 0.58414 | Other Negative Correlation |
| SHROOM2 | -0.03057 | 0.584128 | Other Negative Correlation |
| AMFR | -0.03057 | 0.584108 | Other Negative Correlation |
| NAP1L3 | -0.03061 | 0.583641 | Other Negative Correlation |
| NIPSNAP3A | -0.03061 | 0.583618 | Other Negative Correlation |
| ZSCAN31 | -0.03063 | 0.583357 | Other Negative Correlation |
| STIM2 | -0.03065 | 0.583144 | Other Negative Correlation |
| TMEM30B | -0.03067 | 0.582925 | Other Negative Correlation |
| PAAF1 | -0.03074 | 0.581961 | Other Negative Correlation |
| CPSF7 | -0.03075 | 0.581852 | Other Negative Correlation |
| STRIP1 | -0.03083 | 0.580911 | Other Negative Correlation |
| CLDN16 | -0.03085 | 0.580645 | Other Negative Correlation |
| MAPRE3 | -0.03085 | 0.580629 | Other Negative Correlation |
| KLHDC10 | -0.03088 | 0.580281 | Other Negative Correlation |
| MIER1 | -0.03092 | 0.57981 | Other Negative Correlation |
| MLLT11 | -0.03094 | 0.57957 | Other Negative Correlation |
| USP33 | -0.03103 | 0.578438 | Other Negative Correlation |
| PI4K2A | -0.03104 | 0.578279 | Other Negative Correlation |
| ZNF586 | -0.03105 | 0.578229 | Other Negative Correlation |
| RPN1 | -0.03108 | 0.57778 | Other Negative Correlation |
| OPHN1 | -0.0312 | 0.576319 | Other Negative Correlation |
| LYPLAL1 | -0.03125 | 0.575811 | Other Negative Correlation |
| IGFBP6 | -0.03127 | 0.575546 | Other Negative Correlation |
| AP5B1 | -0.03127 | 0.575537 | Other Negative Correlation |
| ZNF570 | -0.03127 | 0.575498 | Other Negative Correlation |

| | | | |
|---|---|---|---|
| NOTCH2NLR | -0.03129 | 0.575319 | Other Negative Correlation |
| STX7 | -0.0313 | 0.57518 | Other Negative Correlation |
| CORIN | -0.03133 | 0.574803 | Other Negative Correlation |
| SFI1 | -0.03133 | 0.574801 | Other Negative Correlation |
| OFD1 | -0.03138 | 0.574213 | Other Negative Correlation |
| FHIP2B | -0.03139 | 0.574026 | Other Negative Correlation |
| FAHD2A | -0.03146 | 0.573247 | Other Negative Correlation |
| NHEJ1 | -0.03152 | 0.572489 | Other Negative Correlation |
| DUSP22 | -0.03152 | 0.572441 | Other Negative Correlation |
| GRK6 | -0.03154 | 0.572183 | Other Negative Correlation |
| HSDL1 | -0.03157 | 0.571877 | Other Negative Correlation |
| HPCAL1 | -0.03159 | 0.571574 | Other Negative Correlation |
| P4HB | -0.03159 | 0.571557 | Other Negative Correlation |
| ST8SIA6 | -0.0316 | 0.57152 | Other Negative Correlation |
| SMIM27 | -0.0317 | 0.570218 | Other Negative Correlation |
| RGL2 | -0.03173 | 0.569872 | Other Negative Correlation |
| ERCC6L2 | -0.03176 | 0.569517 | Other Negative Correlation |
| HIPK3 | -0.03181 | 0.568985 | Other Negative Correlation |
| LRATD2 | -0.03181 | 0.568888 | Other Negative Correlation |
| TXNDC15 | -0.03182 | 0.568849 | Other Negative Correlation |
| SLC12A9 | -0.03184 | 0.568601 | Other Negative Correlation |
| SERINC1 | -0.03191 | 0.567766 | Other Negative Correlation |
| TECR | -0.03192 | 0.567572 | Other Negative Correlation |
| SSC4D | -0.03196 | 0.567138 | Other Negative Correlation |
| FNDC4 | -0.03198 | 0.566873 | Other Negative Correlation |
| NR6A1 | -0.03205 | 0.566036 | Other Negative Correlation |
| INSYN2B | -0.03208 | 0.565702 | Other Negative Correlation |
| TBC1D24 | -0.03209 | 0.565577 | Other Negative Correlation |
| KLRC2 | -0.03209 | 0.565528 | Other Negative Correlation |
| ENSG00000267001 | -0.03217 | 0.564516 | Other Negative Correlation |
| CYB561D1 | -0.03222 | 0.564015 | Other Negative Correlation |
| RNF167 | -0.03222 | 0.563981 | Other Negative Correlation |
| TMEM94 | -0.03223 | 0.56388 | Other Negative Correlation |
| MRS2 | -0.03225 | 0.563634 | Other Negative Correlation |
| RCOR3 | -0.03226 | 0.563514 | Other Negative Correlation |
| RNF125 | -0.03231 | 0.562859 | Other Negative Correlation |
| CFAP46 | -0.03237 | 0.562174 | Other Negative Correlation |
| TM2D3 | -0.03238 | 0.561984 | Other Negative Correlation |
| MN1 | -0.03241 | 0.561643 | Other Negative Correlation |
| NECAP2 | -0.03241 | 0.561629 | Other Negative Correlation |
| PKD2 | -0.03244 | 0.561302 | Other Negative Correlation |
| PPP3CB | -0.03244 | 0.561271 | Other Negative Correlation |
| AFF1 | -0.03245 | 0.561223 | Other Negative Correlation |
| FZD3 | -0.03247 | 0.560994 | Other Negative Correlation |
| ACBD5 | -0.03257 | 0.559787 | Other Negative Correlation |
| HPSE | -0.03258 | 0.559607 | Other Negative Correlation |
| TRIM9 | -0.0326 | 0.559398 | Other Negative Correlation |
| SGCE | -0.03264 | 0.55883 | Other Negative Correlation |

| | | | |
|---|---|---|---|
| GNA14 | -0.03282 | 0.556694 | Other Negative Correlation |
| DDX59 | -0.03283 | 0.556606 | Other Negative Correlation |
| NOTCH4 | -0.03284 | 0.556429 | Other Negative Correlation |
| MT-ND2 | -0.03285 | 0.556413 | Other Negative Correlation |
| TAFA3 | -0.03285 | 0.556372 | Other Negative Correlation |
| MOSPD2 | -0.0329 | 0.555751 | Other Negative Correlation |
| PRRT3 | -0.03291 | 0.555583 | Other Negative Correlation |
| SGPL1 | -0.03293 | 0.555435 | Other Negative Correlation |
| ABLIM1 | -0.03293 | 0.555348 | Other Negative Correlation |
| KDELR2 | -0.033 | 0.55456 | Other Negative Correlation |
| FUT8 | -0.03308 | 0.553591 | Other Negative Correlation |
| CDK5RAP3 | -0.0331 | 0.553359 | Other Negative Correlation |
| ATXN7L2 | -0.03318 | 0.552368 | Other Negative Correlation |
| SLC25A12 | -0.03321 | 0.552024 | Other Negative Correlation |
| LMBRD1 | -0.03322 | 0.551929 | Other Negative Correlation |
| TRMT11 | -0.03324 | 0.551722 | Other Negative Correlation |
| SELENOO | -0.03325 | 0.551588 | Other Negative Correlation |
| SLX4 | -0.03326 | 0.55144 | Other Negative Correlation |
| CRKL | -0.03328 | 0.551255 | Other Negative Correlation |
| DAPL1 | -0.0333 | 0.550997 | Other Negative Correlation |
| PAFAH1B3 | -0.0333 | 0.550985 | Other Negative Correlation |
| ASXL1 | -0.0333 | 0.550942 | Other Negative Correlation |
| UBAP2 | -0.03332 | 0.550747 | Other Negative Correlation |
| GPR150 | -0.03336 | 0.550196 | Other Negative Correlation |
| CFAP53 | -0.03339 | 0.54993 | Other Negative Correlation |
| SGSM2 | -0.03343 | 0.549368 | Other Negative Correlation |
| CHMP6 | -0.03345 | 0.549178 | Other Negative Correlation |
| IDH1 | -0.03349 | 0.548686 | Other Negative Correlation |
| ADGRG1 | -0.03349 | 0.54865 | Other Negative Correlation |
| NHLRC1 | -0.03356 | 0.547868 | Other Negative Correlation |
| HDAC10 | -0.03357 | 0.54775 | Other Negative Correlation |
| KRTAP4-8 | -0.03358 | 0.54758 | Other Negative Correlation |
| EXOC1L | -0.03358 | 0.54758 | Other Negative Correlation |
| USP28 | -0.0336 | 0.547327 | Other Negative Correlation |
| MFAP3 | -0.03375 | 0.545623 | Other Negative Correlation |
| MUC2 | -0.03377 | 0.545294 | Other Negative Correlation |
| REELD1 | -0.03377 | 0.545294 | Other Negative Correlation |
| ETNK2 | -0.03378 | 0.545225 | Other Negative Correlation |
| CPVL | -0.03379 | 0.545134 | Other Negative Correlation |
| KAT14 | -0.03381 | 0.544909 | Other Negative Correlation |
| PTPRU | -0.03385 | 0.544428 | Other Negative Correlation |
| TUBGCP5 | -0.03387 | 0.544158 | Other Negative Correlation |
| SEPTIN6 | -0.03388 | 0.544096 | Other Negative Correlation |
| ZNF133 | -0.03388 | 0.544047 | Other Negative Correlation |
| OSBPL8 | -0.0339 | 0.543824 | Other Negative Correlation |
| PSMB3 | -0.03391 | 0.543635 | Other Negative Correlation |
| CIAO3 | -0.03392 | 0.543519 | Other Negative Correlation |
| APAF1 | -0.03394 | 0.54329 | Other Negative Correlation |

| | | | |
|---|---|---|---|
| VGLL4 | -0.03396 | 0.543036 | Other Negative Correlation |
| HDAC4 | -0.03398 | 0.542895 | Other Negative Correlation |
| EMP1 | -0.03398 | 0.54287 | Other Negative Correlation |
| SLC50A1 | -0.03399 | 0.542695 | Other Negative Correlation |
| CKM | -0.034 | 0.542674 | Other Negative Correlation |
| ZNF319 | -0.03406 | 0.541904 | Other Negative Correlation |
| BMP8A | -0.0341 | 0.54145 | Other Negative Correlation |
| FBXL7 | -0.0341 | 0.541446 | Other Negative Correlation |
| AP2A1 | -0.03415 | 0.540881 | Other Negative Correlation |
| CHMP7 | -0.03416 | 0.540765 | Other Negative Correlation |
| ABCA13 | -0.03421 | 0.540179 | Other Negative Correlation |
| GNB2 | -0.03425 | 0.539693 | Other Negative Correlation |
| PER3 | -0.03426 | 0.539572 | Other Negative Correlation |
| HIVEP1 | -0.03426 | 0.539518 | Other Negative Correlation |
| ARHGAP4 | -0.03431 | 0.538933 | Other Negative Correlation |
| MAB21L3 | -0.03437 | 0.53818 | Other Negative Correlation |
| SNAPIN | -0.03442 | 0.537668 | Other Negative Correlation |
| ARHGEF10L | -0.03442 | 0.537663 | Other Negative Correlation |
| ZNF573 | -0.03447 | 0.537007 | Other Negative Correlation |
| CEP97 | -0.03453 | 0.536394 | Other Negative Correlation |
| CDIN1 | -0.03455 | 0.536106 | Other Negative Correlation |
| CCNL2 | -0.03455 | 0.53608 | Other Negative Correlation |
| CRNKL1 | -0.03459 | 0.535595 | Other Negative Correlation |
| SPN | -0.03463 | 0.535204 | Other Negative Correlation |
| FAM227B | -0.03463 | 0.535171 | Other Negative Correlation |
| KLRC1 | -0.03473 | 0.533924 | Other Negative Correlation |
| BFAR | -0.03477 | 0.533479 | Other Negative Correlation |
| C14orf28 | -0.03478 | 0.533403 | Other Negative Correlation |
| TMEM199 | -0.03479 | 0.533327 | Other Negative Correlation |
| TXNRD2 | -0.0348 | 0.533193 | Other Negative Correlation |
| USP43 | -0.03483 | 0.532809 | Other Negative Correlation |
| KLHL4 | -0.03484 | 0.532686 | Other Negative Correlation |
| KCNE5 | -0.03486 | 0.532453 | Other Negative Correlation |
| CDNF | -0.03491 | 0.531866 | Other Negative Correlation |
| MAP4K3 | -0.03491 | 0.531833 | Other Negative Correlation |
| THNSL2 | -0.03492 | 0.531801 | Other Negative Correlation |
| ANK2 | -0.03492 | 0.531797 | Other Negative Correlation |
| TCF7 | -0.03495 | 0.531403 | Other Negative Correlation |
| KMO | -0.03498 | 0.530982 | Other Negative Correlation |
| KIAA1328 | -0.03499 | 0.530948 | Other Negative Correlation |
| GPRASP2 | -0.03503 | 0.530396 | Other Negative Correlation |
| ENSG00000188897 | -0.03504 | 0.53029 | Other Negative Correlation |
| CREBZF | -0.03508 | 0.529897 | Other Negative Correlation |
| DYDC2 | -0.03509 | 0.529777 | Other Negative Correlation |
| TBL1Y | -0.03509 | 0.529753 | Other Negative Correlation |
| PTTG1 | -0.03511 | 0.529513 | Other Negative Correlation |
| MAP2K7 | -0.03511 | 0.529463 | Other Negative Correlation |
| NT5C3B | -0.03514 | 0.529197 | Other Negative Correlation |

| | | | |
|---|---|---|---|
| F2RL1 | -0.03519 | 0.528622 | Other Negative Correlation |
| CMC1 | -0.03527 | 0.527653 | Other Negative Correlation |
| TNFRSF10C | -0.03528 | 0.527542 | Other Negative Correlation |
| YIF1B | -0.03531 | 0.527149 | Other Negative Correlation |
| C22orf15 | -0.03533 | 0.526932 | Other Negative Correlation |
| KCTD7 | -0.03535 | 0.526714 | Other Negative Correlation |
| PAGR1 | -0.03536 | 0.526534 | Other Negative Correlation |
| SNAI3 | -0.03537 | 0.526411 | Other Negative Correlation |
| KCNU1 | -0.03539 | 0.526229 | Other Negative Correlation |
| RAB4B | -0.03541 | 0.525989 | Other Negative Correlation |
| WDR27 | -0.0355 | 0.524957 | Other Negative Correlation |
| PEX19 | -0.03551 | 0.524862 | Other Negative Correlation |
| TMPRSS7 | -0.03551 | 0.524808 | Other Negative Correlation |
| NMRAL1 | -0.03552 | 0.524719 | Other Negative Correlation |
| SUN1 | -0.03552 | 0.524676 | Other Negative Correlation |
| TNFRSF10D | -0.03554 | 0.524527 | Other Negative Correlation |
| CLUAP1 | -0.0356 | 0.523744 | Other Negative Correlation |
| KANSL1 | -0.03564 | 0.52335 | Other Negative Correlation |
| ZNF677 | -0.03566 | 0.523034 | Other Negative Correlation |
| MPPE1 | -0.0357 | 0.522613 | Other Negative Correlation |
| CDHR4 | -0.03574 | 0.522123 | Other Negative Correlation |
| ANKRD13A | -0.03575 | 0.521998 | Other Negative Correlation |
| CCDC13 | -0.03576 | 0.521936 | Other Negative Correlation |
| SIK2 | -0.03579 | 0.521606 | Other Negative Correlation |
| SRC | -0.03582 | 0.521243 | Other Negative Correlation |
| ATP8B1 | -0.03586 | 0.520795 | Other Negative Correlation |
| ZNF618 | -0.03586 | 0.52077 | Other Negative Correlation |
| ZNF19 | -0.03589 | 0.520432 | Other Negative Correlation |
| CRYBA4 | -0.03589 | 0.520416 | Other Negative Correlation |
| OCSTAMP | -0.03589 | 0.520416 | Other Negative Correlation |
| JAGN1 | -0.03596 | 0.519587 | Other Negative Correlation |
| EDIL3 | -0.03604 | 0.5187 | Other Negative Correlation |
| ZBTB44 | -0.03605 | 0.51853 | Other Negative Correlation |
| SH3RF1 | -0.03606 | 0.518482 | Other Negative Correlation |
| SIPA1L3 | -0.03632 | 0.515445 | Other Negative Correlation |
| CISD1 | -0.03632 | 0.515388 | Other Negative Correlation |
| PSME3 | -0.03633 | 0.515314 | Other Negative Correlation |
| PEAK3 | -0.03635 | 0.515012 | Other Negative Correlation |
| TSHZ1 | -0.03635 | 0.515008 | Other Negative Correlation |
| TNFSF14 | -0.03636 | 0.514988 | Other Negative Correlation |
| CRTC1 | -0.03639 | 0.51461 | Other Negative Correlation |
| SPAG9 | -0.03648 | 0.513562 | Other Negative Correlation |
| PIK3R5 | -0.03651 | 0.513265 | Other Negative Correlation |
| NAPG | -0.03651 | 0.513209 | Other Negative Correlation |
| MZF1 | -0.03656 | 0.512668 | Other Negative Correlation |
| TLCD4-RWDD3 | -0.03656 | 0.512587 | Other Negative Correlation |
| EVPL | -0.03662 | 0.51196 | Other Negative Correlation |
| OSCP1 | -0.03662 | 0.511959 | Other Negative Correlation |

| | | | |
|---|---|---|---|
| EPC1 | -0.03664 | 0.51168 | Other Negative Correlation |
| ATP5PF | -0.03669 | 0.511116 | Other Negative Correlation |
| PAPPA2 | -0.0367 | 0.511004 | Other Negative Correlation |
| SGMS1 | -0.03675 | 0.510392 | Other Negative Correlation |
| ORMDL1 | -0.03678 | 0.510054 | Other Negative Correlation |
| RPH3AL | -0.0368 | 0.509834 | Other Negative Correlation |
| UNC45B | -0.03686 | 0.509184 | Other Negative Correlation |
| SUPT4H1 | -0.03687 | 0.509025 | Other Negative Correlation |
| MSL1 | -0.03688 | 0.508971 | Other Negative Correlation |
| RAB3C | -0.03692 | 0.50852 | Other Negative Correlation |
| ARID4B | -0.03695 | 0.508174 | Other Negative Correlation |
| COL22A1 | -0.03696 | 0.508002 | Other Negative Correlation |
| ARHGEF17 | -0.03699 | 0.507743 | Other Negative Correlation |
| ZNF443 | -0.03702 | 0.507288 | Other Negative Correlation |
| SLC23A3 | -0.03707 | 0.506737 | Other Negative Correlation |
| TATDN2 | -0.03708 | 0.50666 | Other Negative Correlation |
| DYNLT4 | -0.03708 | 0.506609 | Other Negative Correlation |
| ZNF83 | -0.0371 | 0.50639 | Other Negative Correlation |
| COA8 | -0.03712 | 0.506234 | Other Negative Correlation |
| CALM2 | -0.03714 | 0.505941 | Other Negative Correlation |
| TNK2 | -0.03716 | 0.505764 | Other Negative Correlation |
| ANKRD22 | -0.03716 | 0.505735 | Other Negative Correlation |
| UBR4 | -0.03717 | 0.505571 | Other Negative Correlation |
| HERPUD2 | -0.03723 | 0.504971 | Other Negative Correlation |
| PLCXD1 | -0.03724 | 0.504852 | Other Negative Correlation |
| TRPM3 | -0.03729 | 0.504243 | Other Negative Correlation |
| BBS4 | -0.0373 | 0.504117 | Other Negative Correlation |
| MED23 | -0.03737 | 0.503362 | Other Negative Correlation |
| SLC41A3 | -0.03738 | 0.503255 | Other Negative Correlation |
| PCCA | -0.03741 | 0.502828 | Other Negative Correlation |
| GTPBP8 | -0.03747 | 0.502145 | Other Negative Correlation |
| TAOK1 | -0.03749 | 0.502017 | Other Negative Correlation |
| H3-7 | -0.03756 | 0.50118 | Other Negative Correlation |
| GSDMB | -0.03765 | 0.500153 | Other Negative Correlation |
| ZNF710 | -0.03766 | 0.500079 | Other Negative Correlation |
| COX7A2L | -0.03766 | 0.500019 | Other Negative Correlation |
| ZFYVE26 | -0.03771 | 0.499438 | Other Negative Correlation |
| TASOR | -0.03773 | 0.499229 | Other Negative Correlation |
| TSPAN10 | -0.03774 | 0.499092 | Other Negative Correlation |
| TAF1C | -0.03784 | 0.497986 | Other Negative Correlation |
| NDUFA13 | -0.03788 | 0.497536 | Other Negative Correlation |
| GTF2IRD2B | -0.0379 | 0.497322 | Other Negative Correlation |
| REEP6 | -0.03791 | 0.497234 | Other Negative Correlation |
| CBFA2T2 | -0.03801 | 0.49608 | Other Negative Correlation |
| ZFYVE27 | -0.03802 | 0.495934 | Other Negative Correlation |
| MFSD1 | -0.03804 | 0.495728 | Other Negative Correlation |
| CABCOCO1 | -0.03808 | 0.495219 | Other Negative Correlation |
| ENDOV | -0.0381 | 0.494973 | Other Negative Correlation |

| | | | |
|---|---|---|---|
| BOD1L1 | -0.03812 | 0.494819 | Other Negative Correlation |
| SLC45A3 | -0.03812 | 0.494787 | Other Negative Correlation |
| SULT1A2 | -0.03815 | 0.494486 | Other Negative Correlation |
| DNASE2B | -0.03821 | 0.493787 | Other Negative Correlation |
| NLRX1 | -0.03824 | 0.493466 | Other Negative Correlation |
| HELZ | -0.03825 | 0.49337 | Other Negative Correlation |
| ZNF799 | -0.03825 | 0.493301 | Other Negative Correlation |
| RAB22A | -0.03829 | 0.49288 | Other Negative Correlation |
| ALOX15B | -0.0383 | 0.492711 | Other Negative Correlation |
| GNAT2 | -0.03834 | 0.492339 | Other Negative Correlation |
| GALNT10 | -0.03845 | 0.491098 | Other Negative Correlation |
| TPK1 | -0.03848 | 0.490758 | Other Negative Correlation |
| ARSI | -0.03849 | 0.490624 | Other Negative Correlation |
| CLTC | -0.03851 | 0.490374 | Other Negative Correlation |
| DDA1 | -0.03852 | 0.490239 | Other Negative Correlation |
| ZNF302 | -0.03856 | 0.489808 | Other Negative Correlation |
| DNAH2 | -0.03858 | 0.48966 | Other Negative Correlation |
| IL17RD | -0.0386 | 0.489373 | Other Negative Correlation |
| WIPF2 | -0.0386 | 0.489361 | Other Negative Correlation |
| DPY30 | -0.03864 | 0.488945 | Other Negative Correlation |
| IFTAP | -0.03864 | 0.48893 | Other Negative Correlation |
| PNRC1 | -0.03868 | 0.488461 | Other Negative Correlation |
| PALS1 | -0.03871 | 0.488196 | Other Negative Correlation |
| ADGRD2 | -0.03875 | 0.487744 | Other Negative Correlation |
| PACS2 | -0.03875 | 0.487728 | Other Negative Correlation |
| ZFYVE9 | -0.03877 | 0.487487 | Other Negative Correlation |
| GK5 | -0.03877 | 0.487432 | Other Negative Correlation |
| BLVRA | -0.0388 | 0.487175 | Other Negative Correlation |
| COL24A1 | -0.0388 | 0.487159 | Other Negative Correlation |
| WDR44 | -0.0388 | 0.487122 | Other Negative Correlation |
| GTPBP3 | -0.03885 | 0.486619 | Other Negative Correlation |
| MDH1 | -0.0389 | 0.485962 | Other Negative Correlation |
| TYW1B | -0.03892 | 0.485819 | Other Negative Correlation |
| IL23A | -0.03894 | 0.485558 | Other Negative Correlation |
| ST8SIA4 | -0.03896 | 0.485371 | Other Negative Correlation |
| JHY | -0.03899 | 0.484958 | Other Negative Correlation |
| ZNF300 | -0.039 | 0.484838 | Other Negative Correlation |
| ITGB2 | -0.03903 | 0.484532 | Other Negative Correlation |
| TAF12 | -0.03905 | 0.484341 | Other Negative Correlation |
| FIP1L1 | -0.0391 | 0.483741 | Other Negative Correlation |
| CLTCL1 | -0.03911 | 0.483681 | Other Negative Correlation |
| TAF1 | -0.03913 | 0.483458 | Other Negative Correlation |
| IQANK1 | -0.03916 | 0.483106 | Other Negative Correlation |
| KMT2C | -0.03923 | 0.482286 | Other Negative Correlation |
| MEI1 | -0.03923 | 0.482272 | Other Negative Correlation |
| APCDD1L | -0.03925 | 0.482131 | Other Negative Correlation |
| ACOD1 | -0.03929 | 0.48163 | Other Negative Correlation |
| NAA35 | -0.03932 | 0.481343 | Other Negative Correlation |

| | | | |
|---|---|---|---|
| ATP7B | -0.03933 | 0.48122 | Other Negative Correlation |
| DENND4C | -0.03933 | 0.481171 | Other Negative Correlation |
| GTF2A2 | -0.03934 | 0.481047 | Other Negative Correlation |
| PIP4K2B | -0.03936 | 0.480865 | Other Negative Correlation |
| PIGB | -0.03938 | 0.480665 | Other Negative Correlation |
| HHAT | -0.03943 | 0.480041 | Other Negative Correlation |
| NPAS2 | -0.03944 | 0.48002 | Other Negative Correlation |
| JPH1 | -0.03945 | 0.479816 | Other Negative Correlation |
| ZNF438 | -0.03946 | 0.479714 | Other Negative Correlation |
| DYNC2I1 | -0.03956 | 0.478606 | Other Negative Correlation |
| KBTBD3 | -0.03959 | 0.478276 | Other Negative Correlation |
| C16orf95 | -0.0396 | 0.478182 | Other Negative Correlation |
| ZNF320 | -0.03961 | 0.478071 | Other Negative Correlation |
| PLAG1 | -0.03962 | 0.478006 | Other Negative Correlation |
| KIF16B | -0.03962 | 0.477924 | Other Negative Correlation |
| TOPORS | -0.03963 | 0.477876 | Other Negative Correlation |
| LURAP1 | -0.03966 | 0.47749 | Other Negative Correlation |
| PFDN1 | -0.03972 | 0.476866 | Other Negative Correlation |
| SAR1A | -0.03974 | 0.476632 | Other Negative Correlation |
| SPOCK1 | -0.03977 | 0.476327 | Other Negative Correlation |
| UBE2M | -0.03979 | 0.476047 | Other Negative Correlation |
| PANK2 | -0.03985 | 0.475426 | Other Negative Correlation |
| NMI | -0.03992 | 0.474596 | Other Negative Correlation |
| NSL1 | -0.03994 | 0.474413 | Other Negative Correlation |
| MT-ATP8 | -0.03995 | 0.474261 | Other Negative Correlation |
| OSTF1 | -0.04007 | 0.472962 | Other Negative Correlation |
| RTL8B | -0.04009 | 0.472755 | Other Negative Correlation |
| PNP | -0.04009 | 0.472717 | Other Negative Correlation |
| HMG20B | -0.04013 | 0.47236 | Other Negative Correlation |
| LGALS8 | -0.04013 | 0.472307 | Other Negative Correlation |
| LONP2 | -0.04015 | 0.472052 | Other Negative Correlation |
| MAP3K21 | -0.04017 | 0.471837 | Other Negative Correlation |
| MAN2C1 | -0.04018 | 0.471721 | Other Negative Correlation |
| DHX38 | -0.04018 | 0.471715 | Other Negative Correlation |
| COX11 | -0.04019 | 0.471687 | Other Negative Correlation |
| PAFAH1B2 | -0.0402 | 0.471545 | Other Negative Correlation |
| KANK4 | -0.04026 | 0.470911 | Other Negative Correlation |
| USP46 | -0.04031 | 0.470317 | Other Negative Correlation |
| PPP2R2B | -0.04034 | 0.470041 | Other Negative Correlation |
| MIA | -0.04037 | 0.469678 | Other Negative Correlation |
| JADE3 | -0.04039 | 0.469485 | Other Negative Correlation |
| C1orf216 | -0.04043 | 0.469049 | Other Negative Correlation |
| INTS6L | -0.04048 | 0.4685 | Other Negative Correlation |
| SBSPON | -0.04049 | 0.468354 | Other Negative Correlation |
| NWD1 | -0.0407 | 0.466005 | Other Negative Correlation |
| TBCEL | -0.04072 | 0.465837 | Other Negative Correlation |
| ZNF747 | -0.0408 | 0.46497 | Other Negative Correlation |
| ZSWIM9 | -0.04081 | 0.464873 | Other Negative Correlation |

| | | | |
|---|---|---|---|
| ATP2C1 | -0.04085 | 0.464389 | Other Negative Correlation |
| PSMD5 | -0.04089 | 0.463951 | Other Negative Correlation |
| COPS7B | -0.04089 | 0.463908 | Other Negative Correlation |
| GCNT1 | -0.04091 | 0.463781 | Other Negative Correlation |
| TMEM183A | -0.04091 | 0.463752 | Other Negative Correlation |
| RFX5 | -0.04092 | 0.463625 | Other Negative Correlation |
| SUN2 | -0.04098 | 0.462946 | Other Negative Correlation |
| CPOX | -0.04099 | 0.462865 | Other Negative Correlation |
| ARHGEF18 | -0.04109 | 0.461791 | Other Negative Correlation |
| ATAT1 | -0.04111 | 0.461551 | Other Negative Correlation |
| ZNF420 | -0.04116 | 0.461063 | Other Negative Correlation |
| CREB3 | -0.04116 | 0.461062 | Other Negative Correlation |
| DGAT2 | -0.04125 | 0.460082 | Other Negative Correlation |
| PTPN3 | -0.04126 | 0.459961 | Other Negative Correlation |
| TEAD2 | -0.04126 | 0.459946 | Other Negative Correlation |
| WBP1L | -0.04127 | 0.459816 | Other Negative Correlation |
| FLYWCH1 | -0.04128 | 0.459717 | Other Negative Correlation |
| OR8S1 | -0.04134 | 0.459011 | Other Negative Correlation |
| HHATL | -0.04134 | 0.459011 | Other Negative Correlation |
| WDR37 | -0.04142 | 0.458196 | Other Negative Correlation |
| ADPRHL1 | -0.04143 | 0.458092 | Other Negative Correlation |
| SERPINB10 | -0.04144 | 0.457964 | Other Negative Correlation |
| ZFP69B | -0.04149 | 0.457425 | Other Negative Correlation |
| DDX27 | -0.0415 | 0.457351 | Other Negative Correlation |
| CERS5 | -0.04153 | 0.457024 | Other Negative Correlation |
| RAPGEF6 | -0.04158 | 0.456468 | Other Negative Correlation |
| TFCP2L1 | -0.04158 | 0.456424 | Other Negative Correlation |
| LAPTM4B | -0.04161 | 0.456173 | Other Negative Correlation |
| KIAA1191 | -0.04161 | 0.456132 | Other Negative Correlation |
| NBPF10 | -0.04165 | 0.455695 | Other Negative Correlation |
| REL | -0.04165 | 0.455647 | Other Negative Correlation |
| HCAR2 | -0.04171 | 0.455069 | Other Negative Correlation |
| CASP7 | -0.04173 | 0.454866 | Other Negative Correlation |
| SOX12 | -0.04174 | 0.454673 | Other Negative Correlation |
| GOLGA4 | -0.04175 | 0.454639 | Other Negative Correlation |
| PPIP5K2 | -0.0418 | 0.454064 | Other Negative Correlation |
| BCAN | -0.04184 | 0.453585 | Other Negative Correlation |
| GPR143 | -0.04194 | 0.45256 | Other Negative Correlation |
| ZNF222 | -0.04201 | 0.451844 | Other Negative Correlation |
| CHPF2 | -0.04201 | 0.451811 | Other Negative Correlation |
| CD2BP2 | -0.04207 | 0.451167 | Other Negative Correlation |
| ALKBH1 | -0.0421 | 0.450884 | Other Negative Correlation |
| CCDC68 | -0.0422 | 0.449804 | Other Negative Correlation |
| ZDHHC5 | -0.04223 | 0.449403 | Other Negative Correlation |
| TMEM182 | -0.04225 | 0.449168 | Other Negative Correlation |
| ARHGEF12 | -0.04235 | 0.448151 | Other Negative Correlation |
| DOK7 | -0.04238 | 0.447867 | Other Negative Correlation |
| SLC2A3 | -0.04238 | 0.447774 | Other Negative Correlation |

| | | | |
|---|---|---|---|
| DCLRE1C | -0.04247 | 0.446891 | Other Negative Correlation |
| TCEAL1 | -0.04251 | 0.446391 | Other Negative Correlation |
| PSMB5 | -0.04255 | 0.446 | Other Negative Correlation |
| ACTR10 | -0.04258 | 0.445677 | Other Negative Correlation |
| RAB3B | -0.0426 | 0.445503 | Other Negative Correlation |
| ALDH4A1 | -0.04266 | 0.44479 | Other Negative Correlation |
| VPS16 | -0.04266 | 0.444777 | Other Negative Correlation |
| CUTA | -0.04267 | 0.444725 | Other Negative Correlation |
| FITM2 | -0.04268 | 0.444597 | Other Negative Correlation |
| PCDH18 | -0.04271 | 0.444344 | Other Negative Correlation |
| SEC24B | -0.04273 | 0.444127 | Other Negative Correlation |
| NEIL1 | -0.04275 | 0.443838 | Other Negative Correlation |
| ZFHX4 | -0.04278 | 0.44353 | Other Negative Correlation |
| TOGARAM1 | -0.04281 | 0.44323 | Other Negative Correlation |
| DLG2 | -0.04281 | 0.443221 | Other Negative Correlation |
| PHKA1 | -0.04282 | 0.443096 | Other Negative Correlation |
| ENAH | -0.04286 | 0.442715 | Other Negative Correlation |
| INTS14 | -0.04287 | 0.442562 | Other Negative Correlation |
| PLAUR | -0.04287 | 0.442533 | Other Negative Correlation |
| C2orf66 | -0.04293 | 0.441976 | Other Negative Correlation |
| GOLGA6L10 | -0.04298 | 0.441437 | Other Negative Correlation |
| VANGL1 | -0.04299 | 0.441305 | Other Negative Correlation |
| SPTY2D1 | -0.04306 | 0.440598 | Other Negative Correlation |
| SCX | -0.04316 | 0.439549 | Other Negative Correlation |
| PEBP4 | -0.04321 | 0.439021 | Other Negative Correlation |
| ARHGEF16 | -0.04325 | 0.438576 | Other Negative Correlation |
| EXOC3 | -0.04325 | 0.438568 | Other Negative Correlation |
| C19orf47 | -0.04329 | 0.43809 | Other Negative Correlation |
| ZNF25 | -0.0433 | 0.437977 | Other Negative Correlation |
| ACTN3 | -0.04331 | 0.43788 | Other Negative Correlation |
| UGGT2 | -0.04334 | 0.437591 | Other Negative Correlation |
| GTF3C4 | -0.04335 | 0.437439 | Other Negative Correlation |
| ENSG00000286185 | -0.04339 | 0.437097 | Other Negative Correlation |
| AGO1 | -0.04343 | 0.436623 | Other Negative Correlation |
| CLEC16A | -0.04346 | 0.436285 | Other Negative Correlation |
| SPAG16 | -0.04354 | 0.435463 | Other Negative Correlation |
| NAALADL1 | -0.04356 | 0.435304 | Other Negative Correlation |
| DYM | -0.04358 | 0.435047 | Other Negative Correlation |
| LANCL2 | -0.04361 | 0.434716 | Other Negative Correlation |
| ZNF446 | -0.0437 | 0.43383 | Other Negative Correlation |
| MPP7 | -0.04373 | 0.433522 | Other Negative Correlation |
| SYCE1L | -0.04382 | 0.432525 | Other Negative Correlation |
| PPP1R16A | -0.04387 | 0.431961 | Other Negative Correlation |
| COX8C | -0.04391 | 0.431571 | Other Negative Correlation |
| CROCC | -0.04398 | 0.430877 | Other Negative Correlation |
| TMEM249 | -0.04399 | 0.430789 | Other Negative Correlation |
| PPP2R5D | -0.044 | 0.430598 | Other Negative Correlation |
| PHF3 | -0.04402 | 0.430436 | Other Negative Correlation |

| | | | |
|---|---|---|---|
| AACS | -0.04404 | 0.430255 | Other Negative Correlation |
| TMEM191B | -0.04404 | 0.430194 | Other Negative Correlation |
| PRH2 | -0.04406 | 0.430047 | Other Negative Correlation |
| UACA | -0.04409 | 0.429664 | Other Negative Correlation |
| FTO | -0.0441 | 0.429615 | Other Negative Correlation |
| GP1BA | -0.04411 | 0.42944 | Other Negative Correlation |
| FCER1G | -0.04423 | 0.428252 | Other Negative Correlation |
| MATN2 | -0.04431 | 0.427348 | Other Negative Correlation |
| PPIL4 | -0.04433 | 0.42718 | Other Negative Correlation |
| SERINC3 | -0.04434 | 0.427126 | Other Negative Correlation |
| NDUFAF1 | -0.04441 | 0.426338 | Other Negative Correlation |
| LAX1 | -0.04444 | 0.426046 | Other Negative Correlation |
| NKAPD1 | -0.04445 | 0.425907 | Other Negative Correlation |
| BNIP3L | -0.04446 | 0.425874 | Other Negative Correlation |
| ANKRD61 | -0.04448 | 0.425642 | Other Negative Correlation |
| PLA2G2D | -0.04453 | 0.425133 | Other Negative Correlation |
| HIBCH | -0.04455 | 0.424922 | Other Negative Correlation |
| ICE1 | -0.04456 | 0.424839 | Other Negative Correlation |
| TMEM177 | -0.04463 | 0.424117 | Other Negative Correlation |
| RASGEF1A | -0.04465 | 0.423823 | Other Negative Correlation |
| XAB2 | -0.04466 | 0.42376 | Other Negative Correlation |
| RPUSD1 | -0.04475 | 0.422817 | Other Negative Correlation |
| TLE1 | -0.04475 | 0.422788 | Other Negative Correlation |
| GUCY1A1 | -0.04481 | 0.422231 | Other Negative Correlation |
| TMTC4 | -0.04483 | 0.421994 | Other Negative Correlation |
| ZC3H8 | -0.04484 | 0.421864 | Other Negative Correlation |
| COQ9 | -0.04484 | 0.421856 | Other Negative Correlation |
| VNN1 | -0.04488 | 0.421492 | Other Negative Correlation |
| SHROOM4 | -0.04495 | 0.420698 | Other Negative Correlation |
| RBP1 | -0.04499 | 0.420365 | Other Negative Correlation |
| RPA3 | -0.045 | 0.420261 | Other Negative Correlation |
| HOOK3 | -0.04508 | 0.419382 | Other Negative Correlation |
| DNAH1 | -0.04511 | 0.41914 | Other Negative Correlation |
| MPST | -0.04511 | 0.419119 | Other Negative Correlation |
| MORN1 | -0.04511 | 0.419118 | Other Negative Correlation |
| MINDY1 | -0.04512 | 0.419009 | Other Negative Correlation |
| SYCP2L | -0.04517 | 0.41846 | Other Negative Correlation |
| DDAH2 | -0.04526 | 0.417572 | Other Negative Correlation |
| TMSB15B.1 | -0.04526 | 0.417513 | Other Negative Correlation |
| INTS3 | -0.04529 | 0.417202 | Other Negative Correlation |
| ARHGAP40 | -0.04538 | 0.416267 | Other Negative Correlation |
| SLC33A1 | -0.04544 | 0.415676 | Other Negative Correlation |
| SLC22A18 | -0.04545 | 0.415627 | Other Negative Correlation |
| POMK | -0.04546 | 0.415482 | Other Negative Correlation |
| PIGN | -0.0455 | 0.415033 | Other Negative Correlation |
| TMEM126B | -0.04552 | 0.414826 | Other Negative Correlation |
| AFG3L2 | -0.04557 | 0.414337 | Other Negative Correlation |
| DCUN1D3 | -0.04561 | 0.413944 | Other Negative Correlation |

| | | | |
|---|---|---|---|
| OPLAH | -0.04566 | 0.413414 | Other Negative Correlation |
| TNS2 | -0.04572 | 0.412844 | Other Negative Correlation |
| ZNF346 | -0.04583 | 0.411667 | Other Negative Correlation |
| SSR4 | -0.04589 | 0.41108 | Other Negative Correlation |
| DNAJB14 | -0.04597 | 0.410269 | Other Negative Correlation |
| ERCC8 | -0.04605 | 0.409426 | Other Negative Correlation |
| TMC7 | -0.04608 | 0.4092 | Other Negative Correlation |
| SNX8 | -0.04608 | 0.409179 | Other Negative Correlation |
| PFDN4 | -0.04609 | 0.409058 | Other Negative Correlation |
| IL1RN | -0.04609 | 0.40902 | Other Negative Correlation |
| GAB1 | -0.04624 | 0.407569 | Other Negative Correlation |
| SMIM22 | -0.04625 | 0.407441 | Other Negative Correlation |
| CFAP20 | -0.04628 | 0.40712 | Other Negative Correlation |
| RAB5C | -0.04629 | 0.407034 | Other Negative Correlation |
| UROD | -0.04632 | 0.40667 | Other Negative Correlation |
| WDR45B | -0.04634 | 0.406501 | Other Negative Correlation |
| NDUFB8 | -0.04636 | 0.406349 | Other Negative Correlation |
| CDKL4 | -0.04643 | 0.405631 | Other Negative Correlation |
| FLACC1 | -0.04647 | 0.405211 | Other Negative Correlation |
| PFKFB1 | -0.0465 | 0.404853 | Other Negative Correlation |
| CCN6 | -0.04652 | 0.404694 | Other Negative Correlation |
| ZNF354A | -0.04654 | 0.404486 | Other Negative Correlation |
| SERGEF | -0.04655 | 0.404429 | Other Negative Correlation |
| ZNF789 | -0.04655 | 0.404345 | Other Negative Correlation |
| TMEM128 | -0.04658 | 0.404058 | Other Negative Correlation |
| ACO1 | -0.04659 | 0.404 | Other Negative Correlation |
| IL10RB | -0.04659 | 0.403974 | Other Negative Correlation |
| CSPG4 | -0.04663 | 0.403529 | Other Negative Correlation |
| ERP44 | -0.04667 | 0.403127 | Other Negative Correlation |
| CSNK1G3 | -0.04679 | 0.401945 | Other Negative Correlation |
| PRELID2 | -0.04687 | 0.401126 | Other Negative Correlation |
| PHF21A | -0.04694 | 0.400416 | Other Negative Correlation |
| UBR5 | -0.04697 | 0.400113 | Other Negative Correlation |
| NRDC | -0.04712 | 0.398633 | Other Negative Correlation |
| LINC00672 | -0.04713 | 0.398573 | Other Negative Correlation |
| LRRIQ1 | -0.04716 | 0.398275 | Other Negative Correlation |
| CDK7 | -0.04724 | 0.397493 | Other Negative Correlation |
| IL6 | -0.04726 | 0.397237 | Other Negative Correlation |
| ANAPC4 | -0.04727 | 0.397164 | Other Negative Correlation |
| ARNT | -0.04728 | 0.397005 | Other Negative Correlation |
| PHLDB3 | -0.04731 | 0.3967 | Other Negative Correlation |
| GAL3ST4 | -0.04734 | 0.396482 | Other Negative Correlation |
| LRRC27 | -0.04747 | 0.395192 | Other Negative Correlation |
| MAPK9 | -0.04747 | 0.395122 | Other Negative Correlation |
| PCDHGA8 | -0.04751 | 0.394767 | Other Negative Correlation |
| DGAT1 | -0.04755 | 0.394346 | Other Negative Correlation |
| ZNF654 | -0.04756 | 0.394276 | Other Negative Correlation |
| GMDS | -0.04761 | 0.393806 | Other Negative Correlation |

| | | | |
|---|---|---|---|
| EMILIN1 | -0.04762 | 0.393626 | Other Negative Correlation |
| ABCB6 | -0.04765 | 0.393401 | Other Negative Correlation |
| ENSG00000289281 | -0.04765 | 0.393388 | Other Negative Correlation |
| PGBD2 | -0.04766 | 0.393223 | Other Negative Correlation |
| HOXA1 | -0.04771 | 0.392777 | Other Negative Correlation |
| ZNF852 | -0.04775 | 0.392397 | Other Negative Correlation |
| FBXO44 | -0.04776 | 0.392309 | Other Negative Correlation |
| PCDHB15 | -0.04776 | 0.392264 | Other Negative Correlation |
| CFAP70 | -0.0478 | 0.391867 | Other Negative Correlation |
| SH3TC2 | -0.04781 | 0.391794 | Other Negative Correlation |
| CEP131 | -0.04784 | 0.391427 | Other Negative Correlation |
| CRADD | -0.04788 | 0.391127 | Other Negative Correlation |
| ESYT2 | -0.04789 | 0.390987 | Other Negative Correlation |
| CCDC14 | -0.04791 | 0.390819 | Other Negative Correlation |
| SPRED3 | -0.04793 | 0.390542 | Other Negative Correlation |
| ULK3 | -0.04799 | 0.389998 | Other Negative Correlation |
| KRT81 | -0.048 | 0.389922 | Other Negative Correlation |
| COL23A1 | -0.04803 | 0.389563 | Other Negative Correlation |
| GANAB | -0.04805 | 0.389412 | Other Negative Correlation |
| ZNF837 | -0.04809 | 0.388964 | Other Negative Correlation |
| PSMD9 | -0.04812 | 0.388722 | Other Negative Correlation |
| ATF7IP | -0.04814 | 0.388551 | Other Negative Correlation |
| FBXW12 | -0.04815 | 0.388397 | Other Negative Correlation |
| PHF2 | -0.04817 | 0.388235 | Other Negative Correlation |
| ZKSCAN4 | -0.04819 | 0.388041 | Other Negative Correlation |
| DTX4 | -0.0482 | 0.387946 | Other Negative Correlation |
| MPZL1 | -0.04827 | 0.387252 | Other Negative Correlation |
| SURF1 | -0.04829 | 0.387074 | Other Negative Correlation |
| CALCRL | -0.04829 | 0.387021 | Other Negative Correlation |
| TSPAN3 | -0.0483 | 0.386942 | Other Negative Correlation |
| ZNF16 | -0.0483 | 0.38689 | Other Negative Correlation |
| ODAPH | -0.04832 | 0.386727 | Other Negative Correlation |
| CCNG2 | -0.04832 | 0.386707 | Other Negative Correlation |
| COQ4 | -0.04832 | 0.386685 | Other Negative Correlation |
| METTL22 | -0.04837 | 0.386276 | Other Negative Correlation |
| PYROXD2 | -0.04839 | 0.386022 | Other Negative Correlation |
| PMEL | -0.04841 | 0.385859 | Other Negative Correlation |
| DUSP16 | -0.04846 | 0.385377 | Other Negative Correlation |
| TRMT61B | -0.04854 | 0.384615 | Other Negative Correlation |
| SAMD4B | -0.04856 | 0.384399 | Other Negative Correlation |
| BTN3A3 | -0.04864 | 0.38364 | Other Negative Correlation |
| MRPL35 | -0.04867 | 0.383343 | Other Negative Correlation |
| CDC42EP5 | -0.04871 | 0.382883 | Other Negative Correlation |
| PDCD10 | -0.04875 | 0.382501 | Other Negative Correlation |
| COX6C | -0.04876 | 0.382407 | Other Negative Correlation |
| TMEM184B | -0.04878 | 0.382223 | Other Negative Correlation |
| NNT | -0.04878 | 0.3822 | Other Negative Correlation |
| SEMA4F | -0.04885 | 0.381552 | Other Negative Correlation |

| | | | |
|---|---|---|---|
| COG4 | -0.0489 | 0.381075 | Other Negative Correlation |
| B4GALNT4 | -0.0489 | 0.381036 | Other Negative Correlation |
| NBR1 | -0.04891 | 0.380955 | Other Negative Correlation |
| DNAH11 | -0.04892 | 0.380858 | Other Negative Correlation |
| CDC27 | -0.04892 | 0.380811 | Other Negative Correlation |
| NRBP2 | -0.04894 | 0.380626 | Other Negative Correlation |
| TBCD | -0.04897 | 0.380369 | Other Negative Correlation |
| SZT2 | -0.04899 | 0.380187 | Other Negative Correlation |
| PARP14 | -0.0491 | 0.379106 | Other Negative Correlation |
| POLR2I | -0.0491 | 0.379081 | Other Negative Correlation |
| SCO2 | -0.04913 | 0.3788 | Other Negative Correlation |
| SOD2 | -0.04921 | 0.378086 | Other Negative Correlation |
| RUSC2 | -0.04921 | 0.378061 | Other Negative Correlation |
| CNTN1 | -0.04921 | 0.377992 | Other Negative Correlation |
| RNASET2 | -0.04926 | 0.377591 | Other Negative Correlation |
| RIPK1 | -0.04927 | 0.377487 | Other Negative Correlation |
| RNF165 | -0.04928 | 0.37736 | Other Negative Correlation |
| S1PR5 | -0.04928 | 0.377335 | Other Negative Correlation |
| XPNPEP2 | -0.04941 | 0.376074 | Other Negative Correlation |
| LCP1 | -0.04946 | 0.375615 | Other Negative Correlation |
| DZANK1 | -0.04956 | 0.37468 | Other Negative Correlation |
| BTBD8 | -0.04956 | 0.374629 | Other Negative Correlation |
| AREL1 | -0.0496 | 0.374273 | Other Negative Correlation |
| PLOD3 | -0.04961 | 0.374197 | Other Negative Correlation |
| DNAJC13 | -0.04966 | 0.37368 | Other Negative Correlation |
| PARN | -0.04974 | 0.372925 | Other Negative Correlation |
| MRPL14 | -0.04976 | 0.372712 | Other Negative Correlation |
| ZNF334 | -0.04979 | 0.372432 | Other Negative Correlation |
| LYN | -0.04985 | 0.371882 | Other Negative Correlation |
| BRK1 | -0.04995 | 0.37086 | Other Negative Correlation |
| MICU3 | -0.04998 | 0.370637 | Other Negative Correlation |
| NFKBIE | -0.04998 | 0.370599 | Other Negative Correlation |
| PRODH | -0.05 | 0.370419 | Other Negative Correlation |
| SULF1 | -0.05001 | 0.37035 | Other Negative Correlation |
| LMLN | -0.05003 | 0.370126 | Other Negative Correlation |
| ARMH1 | -0.05004 | 0.369991 | Other Negative Correlation |
| SPATA2L | -0.05005 | 0.369953 | Other Negative Correlation |
| ITPRIPL2 | -0.05015 | 0.368984 | Other Negative Correlation |
| ELP4 | -0.05015 | 0.36894 | Other Negative Correlation |
| GPR68 | -0.05017 | 0.368831 | Other Negative Correlation |
| ENOX2 | -0.05024 | 0.368104 | Other Negative Correlation |
| GSR | -0.05026 | 0.367944 | Other Negative Correlation |
| NCOA7 | -0.05028 | 0.367749 | Other Negative Correlation |
| CX3CL1 | -0.05028 | 0.367732 | Other Negative Correlation |
| RGP1 | -0.05028 | 0.367708 | Other Negative Correlation |
| NSUN5 | -0.05029 | 0.367654 | Other Negative Correlation |
| SHFL | -0.05032 | 0.367401 | Other Negative Correlation |
| BTK | -0.05032 | 0.36736 | Other Negative Correlation |

| | | | |
|---|---|---|---|
| AKAP7 | -0.05036 | 0.36702 | Other Negative Correlation |
| NDUFS1 | -0.0504 | 0.366589 | Other Negative Correlation |
| KLRD1 | -0.05044 | 0.366237 | Other Negative Correlation |
| USO1 | -0.05048 | 0.365862 | Other Negative Correlation |
| CITED2 | -0.05048 | 0.365846 | Other Negative Correlation |
| TMEM191C | -0.05048 | 0.365833 | Other Negative Correlation |
| CCDC115 | -0.05048 | 0.36582 | Other Negative Correlation |
| C2CD3 | -0.05052 | 0.365504 | Other Negative Correlation |
| P2RY2 | -0.05054 | 0.365292 | Other Negative Correlation |
| BAG5 | -0.0506 | 0.36467 | Other Negative Correlation |
| BMPR2 | -0.05065 | 0.364214 | Other Negative Correlation |
| KDELR3 | -0.05068 | 0.36391 | Other Negative Correlation |
| CAT | -0.05071 | 0.363692 | Other Negative Correlation |
| TEX22 | -0.05073 | 0.363446 | Other Negative Correlation |
| NAGLU | -0.05076 | 0.363163 | Other Negative Correlation |
| UBTD1 | -0.05083 | 0.362563 | Other Negative Correlation |
| NR2F6 | -0.05085 | 0.362344 | Other Negative Correlation |
| PRKACB | -0.0509 | 0.361887 | Other Negative Correlation |
| TNK1 | -0.05094 | 0.361513 | Other Negative Correlation |
| ADAMTSL5 | -0.05094 | 0.361494 | Other Negative Correlation |
| DCP1B | -0.05094 | 0.361434 | Other Negative Correlation |
| PSG2 | -0.05098 | 0.361143 | Other Negative Correlation |
| IRGQ | -0.05099 | 0.36105 | Other Negative Correlation |
| FAM83E | -0.05107 | 0.360265 | Other Negative Correlation |
| RABGEF1 | -0.05109 | 0.360077 | Other Negative Correlation |
| SENP2 | -0.05109 | 0.360059 | Other Negative Correlation |
| SAA2 | -0.05118 | 0.359193 | Other Negative Correlation |
| DENND3 | -0.0512 | 0.359047 | Other Negative Correlation |
| CTHRC1 | -0.05122 | 0.358808 | Other Negative Correlation |
| NME3 | -0.05127 | 0.358387 | Other Negative Correlation |
| CA14 | -0.05133 | 0.357803 | Other Negative Correlation |
| PKIG | -0.05135 | 0.35766 | Other Negative Correlation |
| SH3TC1 | -0.05136 | 0.357524 | Other Negative Correlation |
| ASAH1 | -0.05149 | 0.35628 | Other Negative Correlation |
| KLRC4 | -0.05153 | 0.355962 | Other Negative Correlation |
| DAB1 | -0.05153 | 0.355962 | Other Negative Correlation |
| IL6R | -0.05153 | 0.35593 | Other Negative Correlation |
| HMCN1 | -0.05157 | 0.355597 | Other Negative Correlation |
| STRN3 | -0.05159 | 0.355404 | Other Negative Correlation |
| TOB2 | -0.05162 | 0.355101 | Other Negative Correlation |
| THEM6 | -0.05162 | 0.355083 | Other Negative Correlation |
| ZNF652 | -0.05162 | 0.355078 | Other Negative Correlation |
| PRCC | -0.05162 | 0.355062 | Other Negative Correlation |
| APEH | -0.05164 | 0.354882 | Other Negative Correlation |
| RBL2 | -0.05166 | 0.35477 | Other Negative Correlation |
| ARSD | -0.05168 | 0.354502 | Other Negative Correlation |
| NCOR1 | -0.05176 | 0.353825 | Other Negative Correlation |
| SERHL2 | -0.05178 | 0.353595 | Other Negative Correlation |

| | | | |
|---|---|---|---|
| RABL6 | -0.05182 | 0.353245 | Other Negative Correlation |
| KCTD18 | -0.05183 | 0.353151 | Other Negative Correlation |
| CDKN2B | -0.05194 | 0.352148 | Other Negative Correlation |
| TM9SF4 | -0.05197 | 0.3518 | Other Negative Correlation |
| CHMP1A | -0.05198 | 0.351727 | Other Negative Correlation |
| SLC12A4 | -0.05198 | 0.351708 | Other Negative Correlation |
| ZNF784 | -0.05199 | 0.351674 | Other Negative Correlation |
| BPTF | -0.052 | 0.351578 | Other Negative Correlation |
| RAB7A | -0.05202 | 0.35134 | Other Negative Correlation |
| LACC1 | -0.05205 | 0.351088 | Other Negative Correlation |
| SLC16A3 | -0.05206 | 0.350962 | Other Negative Correlation |
| KIAA0825 | -0.0521 | 0.350625 | Other Negative Correlation |
| SLC35G2 | -0.05215 | 0.350137 | Other Negative Correlation |
| ZNF185 | -0.05217 | 0.34997 | Other Negative Correlation |
| TUFT1 | -0.05225 | 0.349287 | Other Negative Correlation |
| CXCL16 | -0.05226 | 0.349112 | Other Negative Correlation |
| SLC30A1 | -0.05229 | 0.348834 | Other Negative Correlation |
| MOGAT1 | -0.0523 | 0.348766 | Other Negative Correlation |
| ANKS1A | -0.05232 | 0.348643 | Other Negative Correlation |
| FBXL8 | -0.05233 | 0.348508 | Other Negative Correlation |
| WRN | -0.05233 | 0.348505 | Other Negative Correlation |
| RTL6 | -0.05234 | 0.348396 | Other Negative Correlation |
| RPP14 | -0.05236 | 0.348196 | Other Negative Correlation |
| CAMLG | -0.05251 | 0.346865 | Other Negative Correlation |
| ISOC1 | -0.05255 | 0.346441 | Other Negative Correlation |
| PLCZ1 | -0.05257 | 0.346268 | Other Negative Correlation |
| KIAA0586 | -0.05258 | 0.346233 | Other Negative Correlation |
| KCTD21 | -0.05259 | 0.346145 | Other Negative Correlation |
| DHDH | -0.05263 | 0.345783 | Other Negative Correlation |
| RASA1 | -0.05277 | 0.344425 | Other Negative Correlation |
| ATL3 | -0.05287 | 0.343519 | Other Negative Correlation |
| GRAPL | -0.05287 | 0.343517 | Other Negative Correlation |
| CDK13 | -0.05289 | 0.343334 | Other Negative Correlation |
| ST3GAL4 | -0.05293 | 0.342994 | Other Negative Correlation |
| DUSP18 | -0.05297 | 0.342612 | Other Negative Correlation |
| TMEM263 | -0.05302 | 0.342206 | Other Negative Correlation |
| RPGRIP1L | -0.05303 | 0.342071 | Other Negative Correlation |
| B3GAT3 | -0.05303 | 0.342056 | Other Negative Correlation |
| JMJD1C | -0.05307 | 0.341773 | Other Negative Correlation |
| TMEM30A | -0.05307 | 0.341713 | Other Negative Correlation |
| CCND3 | -0.05329 | 0.339764 | Other Negative Correlation |
| TMEM14A | -0.05333 | 0.339391 | Other Negative Correlation |
| KIF21A | -0.05334 | 0.339311 | Other Negative Correlation |
| KRIT1 | -0.05334 | 0.33928 | Other Negative Correlation |
| SSR1 | -0.05344 | 0.338401 | Other Negative Correlation |
| TMEM184A | -0.05348 | 0.337997 | Other Negative Correlation |
| KDM7A | -0.05356 | 0.337295 | Other Negative Correlation |
| RTL8C | -0.05358 | 0.337119 | Other Negative Correlation |

| | | | |
|---|---|---|---|
| ZSCAN20 | -0.05375 | 0.335602 | Other Negative Correlation |
| USP50 | -0.05375 | 0.335585 | Other Negative Correlation |
| POFUT2 | -0.0539 | 0.334225 | Other Negative Correlation |
| TEX9 | -0.05392 | 0.334039 | Other Negative Correlation |
| JAK2 | -0.05392 | 0.334016 | Other Negative Correlation |
| DVL3 | -0.05396 | 0.333725 | Other Negative Correlation |
| TTC23 | -0.05396 | 0.333678 | Other Negative Correlation |
| C8orf82 | -0.05398 | 0.333518 | Other Negative Correlation |
| RSPH4A | -0.05399 | 0.33343 | Other Negative Correlation |
| CACNA2D4 | -0.05399 | 0.33337 | Other Negative Correlation |
| MAPK1 | -0.05401 | 0.333203 | Other Negative Correlation |
| NDUFA10 | -0.05403 | 0.333043 | Other Negative Correlation |
| AQP8 | -0.05412 | 0.332257 | Other Negative Correlation |
| TBC1D7 | -0.05417 | 0.331789 | Other Negative Correlation |
| ASPHD2 | -0.0542 | 0.331507 | Other Negative Correlation |
| NCEH1 | -0.05426 | 0.330996 | Other Negative Correlation |
| CCNT2 | -0.05428 | 0.330825 | Other Negative Correlation |
| TMEM140 | -0.05435 | 0.330216 | Other Negative Correlation |
| SEC61G | -0.05437 | 0.330031 | Other Negative Correlation |
| ARSA | -0.05437 | 0.329994 | Other Negative Correlation |
| SLC37A3 | -0.05445 | 0.32934 | Other Negative Correlation |
| CD33 | -0.05449 | 0.328991 | Other Negative Correlation |
| NPHP1 | -0.05456 | 0.328284 | Other Negative Correlation |
| TNFRSF13C | -0.05469 | 0.327167 | Other Negative Correlation |
| LARP4B | -0.05472 | 0.326929 | Other Negative Correlation |
| AHI1 | -0.05473 | 0.32679 | Other Negative Correlation |
| LY6K | -0.05475 | 0.32664 | Other Negative Correlation |
| PDHA1 | -0.05479 | 0.326318 | Other Negative Correlation |
| PYY | -0.05481 | 0.326089 | Other Negative Correlation |
| TARS2 | -0.0549 | 0.325354 | Other Negative Correlation |
| POLG | -0.05493 | 0.325087 | Other Negative Correlation |
| CCDC125 | -0.05494 | 0.324994 | Other Negative Correlation |
| CHRNA9 | -0.05494 | 0.324958 | Other Negative Correlation |
| ZFC3H1 | -0.05496 | 0.324762 | Other Negative Correlation |
| SLC24A4 | -0.05496 | 0.324758 | Other Negative Correlation |
| KLHDC4 | -0.05499 | 0.324517 | Other Negative Correlation |
| TARBP2 | -0.05507 | 0.323853 | Other Negative Correlation |
| MCUR1 | -0.05507 | 0.323786 | Other Negative Correlation |
| LRRC49 | -0.05508 | 0.32376 | Other Negative Correlation |
| QPCT | -0.05508 | 0.323719 | Other Negative Correlation |
| CDC16 | -0.05511 | 0.32348 | Other Negative Correlation |
| GALNT9 | -0.05514 | 0.323182 | Other Negative Correlation |
| TTC4 | -0.05523 | 0.322402 | Other Negative Correlation |
| LITAF | -0.05524 | 0.322366 | Other Negative Correlation |
| DMWD | -0.05524 | 0.3223 | Other Negative Correlation |
| BBS10 | -0.05525 | 0.322278 | Other Negative Correlation |
| SMU1 | -0.05525 | 0.322255 | Other Negative Correlation |
| UBXN2B | -0.05527 | 0.322021 | Other Negative Correlation |

| | | | |
|---|---|---|---|
| ABCG1 | -0.05529 | 0.321879 | Other Negative Correlation |
| PAK1 | -0.05539 | 0.320996 | Other Negative Correlation |
| GPX2 | -0.05541 | 0.320805 | Other Negative Correlation |
| ZNHIT1 | -0.05544 | 0.320615 | Other Negative Correlation |
| MGAT1 | -0.05547 | 0.320278 | Other Negative Correlation |
| ZNF880 | -0.05554 | 0.319657 | Other Negative Correlation |
| APOO | -0.05556 | 0.31952 | Other Negative Correlation |
| ZNF287 | -0.05557 | 0.319441 | Other Negative Correlation |
| SUSD3 | -0.0557 | 0.318295 | Other Negative Correlation |
| UTRN | -0.05571 | 0.318205 | Other Negative Correlation |
| ZNF135 | -0.05574 | 0.317957 | Other Negative Correlation |
| PDZD11 | -0.05575 | 0.317844 | Other Negative Correlation |
| MKLN1 | -0.05581 | 0.317342 | Other Negative Correlation |
| ORAI2 | -0.05595 | 0.316134 | Other Negative Correlation |
| SELENOP | -0.056 | 0.315666 | Other Negative Correlation |
| P2RX1 | -0.05608 | 0.315 | Other Negative Correlation |
| CSF3R | -0.05608 | 0.315 | Other Negative Correlation |
| TVP23C | -0.05616 | 0.314361 | Other Negative Correlation |
| BAIAP2L1 | -0.0562 | 0.313948 | Other Negative Correlation |
| ZNF783 | -0.0562 | 0.313937 | Other Negative Correlation |
| ENSG00000267140 | -0.05622 | 0.31383 | Other Negative Correlation |
| BPNT2 | -0.05624 | 0.313609 | Other Negative Correlation |
| ATN1 | -0.05625 | 0.313541 | Other Negative Correlation |
| CRK | -0.05628 | 0.313288 | Other Negative Correlation |
| DGLUCY | -0.05629 | 0.313196 | Other Negative Correlation |
| CATSPERB | -0.05629 | 0.31316 | Other Negative Correlation |
| RSRC2 | -0.05634 | 0.312759 | Other Negative Correlation |
| RTF2 | -0.05643 | 0.311975 | Other Negative Correlation |
| THAP6 | -0.05645 | 0.311848 | Other Negative Correlation |
| SLC9A7 | -0.05649 | 0.311486 | Other Negative Correlation |
| ST6GALNAC5 | -0.0565 | 0.311383 | Other Negative Correlation |
| CDC73 | -0.0565 | 0.311354 | Other Negative Correlation |
| KDM4D | -0.05653 | 0.311126 | Other Negative Correlation |
| INSL3 | -0.05653 | 0.31112 | Other Negative Correlation |
| ZNF630 | -0.0566 | 0.310575 | Other Negative Correlation |
| GPR153 | -0.0566 | 0.31056 | Other Negative Correlation |
| MON2 | -0.0566 | 0.310558 | Other Negative Correlation |
| STPG1 | -0.05667 | 0.309925 | Other Negative Correlation |
| GLI4 | -0.05669 | 0.309728 | Other Negative Correlation |
| PEX11B | -0.05677 | 0.309092 | Other Negative Correlation |
| MAN1A2 | -0.05678 | 0.309038 | Other Negative Correlation |
| SAV1 | -0.0569 | 0.307995 | Other Negative Correlation |
| SMARCAD1 | -0.05693 | 0.307737 | Other Negative Correlation |
| C14orf119 | -0.05698 | 0.307327 | Other Negative Correlation |
| MRPL22 | -0.05702 | 0.306936 | Other Negative Correlation |
| LST1 | -0.05705 | 0.30669 | Other Negative Correlation |
| ELOVL5 | -0.05708 | 0.306458 | Other Negative Correlation |
| RPS6KB2 | -0.05714 | 0.3059 | Other Negative Correlation |

| | | | |
|---|---|---|---|
| CCDC71 | -0.05716 | 0.305802 | Other Negative Correlation |
| FBXO21 | -0.05717 | 0.305647 | Other Negative Correlation |
| RTP4 | -0.05722 | 0.305244 | Other Negative Correlation |
| ZNF197 | -0.05723 | 0.30518 | Other Negative Correlation |
| CBR3 | -0.05724 | 0.305107 | Other Negative Correlation |
| SPCS3 | -0.05726 | 0.304894 | Other Negative Correlation |
| ZNF169 | -0.05727 | 0.304865 | Other Negative Correlation |
| SAMD8 | -0.05728 | 0.304786 | Other Negative Correlation |
| IKBKB | -0.05733 | 0.304342 | Other Negative Correlation |
| MOB3C | -0.05738 | 0.303875 | Other Negative Correlation |
| DRC3 | -0.05739 | 0.303827 | Other Negative Correlation |
| NDUFS3 | -0.0574 | 0.303741 | Other Negative Correlation |
| ORAI3 | -0.05741 | 0.303626 | Other Negative Correlation |
| PCSK6 | -0.05746 | 0.303233 | Other Negative Correlation |
| WWP2 | -0.0575 | 0.302911 | Other Negative Correlation |
| NHSL1 | -0.05751 | 0.302769 | Other Negative Correlation |
| BLTP1 | -0.05753 | 0.302658 | Other Negative Correlation |
| FOXN1 | -0.05756 | 0.302397 | Other Negative Correlation |
| FRA10AC1 | -0.05757 | 0.302263 | Other Negative Correlation |
| LAMP2 | -0.05763 | 0.301826 | Other Negative Correlation |
| TFPI | -0.05771 | 0.301095 | Other Negative Correlation |
| PIK3IP1 | -0.05773 | 0.300995 | Other Negative Correlation |
| ISL1 | -0.05775 | 0.30083 | Other Negative Correlation |
| RPUSD3 | -0.05776 | 0.300722 | Other Negative Correlation |
| MAGED2 | -0.05807 | 0.298151 | Other Negative Correlation |
| AP4M1 | -0.05809 | 0.297955 | Other Negative Correlation |
| DNASE1L3 | -0.05814 | 0.297499 | Other Negative Correlation |
| TM4SF1 | -0.05817 | 0.297319 | Other Negative Correlation |
| TIMP3 | -0.05819 | 0.297127 | Other Negative Correlation |
| ZNF284 | -0.05823 | 0.296793 | Other Negative Correlation |
| CORO6 | -0.05832 | 0.29602 | Other Negative Correlation |
| AGPAT3 | -0.05837 | 0.295602 | Other Negative Correlation |
| NUP214 | -0.05839 | 0.295426 | Other Negative Correlation |
| NBPF14 | -0.05842 | 0.295173 | Other Negative Correlation |
| NDUFV1 | -0.05852 | 0.294383 | Other Negative Correlation |
| ELL | -0.05854 | 0.294223 | Other Negative Correlation |
| C17orf58 | -0.05855 | 0.294166 | Other Negative Correlation |
| RIC1 | -0.05856 | 0.294016 | Other Negative Correlation |
| TMEM267 | -0.05859 | 0.293839 | Other Negative Correlation |
| COX5B | -0.05862 | 0.293533 | Other Negative Correlation |
| DCAF8 | -0.05863 | 0.293462 | Other Negative Correlation |
| H2BC4 | -0.05863 | 0.293441 | Other Negative Correlation |
| ATP7A | -0.05878 | 0.292242 | Other Negative Correlation |
| TTPAL | -0.05883 | 0.291795 | Other Negative Correlation |
| LNX2 | -0.05885 | 0.291698 | Other Negative Correlation |
| ZBED4 | -0.05892 | 0.291062 | Other Negative Correlation |
| HCAR3 | -0.05892 | 0.29105 | Other Negative Correlation |
| ARRDC3 | -0.05897 | 0.290708 | Other Negative Correlation |

| | | | |
|---|---|---|---|
| ZFYVE16 | -0.05902 | 0.29025 | Other Negative Correlation |
| NT5M | -0.05903 | 0.290213 | Other Negative Correlation |
| TRIP4 | -0.05907 | 0.28986 | Other Negative Correlation |
| PSMG2 | -0.05908 | 0.289776 | Other Negative Correlation |
| OSBP2 | -0.05914 | 0.289315 | Other Negative Correlation |
| MCMDC2 | -0.05915 | 0.289188 | Other Negative Correlation |
| EQTN | -0.05916 | 0.289085 | Other Negative Correlation |
| TNF | -0.05923 | 0.28854 | Other Negative Correlation |
| GHR | -0.05925 | 0.288358 | Other Negative Correlation |
| SRSF8 | -0.05926 | 0.288295 | Other Negative Correlation |
| ZBTB33 | -0.05927 | 0.28826 | Other Negative Correlation |
| LPGAT1 | -0.05928 | 0.288113 | Other Negative Correlation |
| FAM181B | -0.05929 | 0.288038 | Other Negative Correlation |
| ENSG00000289346 | -0.05932 | 0.287848 | Other Negative Correlation |
| ZNF713 | -0.05932 | 0.287847 | Other Negative Correlation |
| SAP18 | -0.05938 | 0.287306 | Other Negative Correlation |
| SVIL | -0.0594 | 0.287185 | Other Negative Correlation |
| L1TD1 | -0.05943 | 0.2869 | Other Negative Correlation |
| RHOJ | -0.05945 | 0.2868 | Other Negative Correlation |
| TIMM17B | -0.05949 | 0.286424 | Other Negative Correlation |
| GAPVD1 | -0.0595 | 0.286372 | Other Negative Correlation |
| REEP5 | -0.05952 | 0.28618 | Other Negative Correlation |
| MICAL1 | -0.0596 | 0.285553 | Other Negative Correlation |
| GAS8 | -0.05963 | 0.285276 | Other Negative Correlation |
| PLA2R1 | -0.05967 | 0.284999 | Other Negative Correlation |
| STARD10 | -0.05978 | 0.284121 | Other Negative Correlation |
| RAD17 | -0.05981 | 0.283888 | Other Negative Correlation |
| OXSM | -0.05983 | 0.283656 | Other Negative Correlation |
| SETBP1 | -0.05984 | 0.283627 | Other Negative Correlation |
| C1orf43 | -0.05986 | 0.283418 | Other Negative Correlation |
| LETM1 | -0.0599 | 0.28309 | Other Negative Correlation |
| GOLGA8N | -0.05994 | 0.282827 | Other Negative Correlation |
| MIEN1 | -0.05995 | 0.282694 | Other Negative Correlation |
| KLHL26 | -0.05997 | 0.282527 | Other Negative Correlation |
| GALC | -0.06 | 0.282286 | Other Negative Correlation |
| AP2A2 | -0.06003 | 0.282052 | Other Negative Correlation |
| FBXO16 | -0.06008 | 0.281675 | Other Negative Correlation |
| FOXQ1 | -0.06009 | 0.281589 | Other Negative Correlation |
| ANAPC13 | -0.06011 | 0.281473 | Other Negative Correlation |
| EIF4G3 | -0.06013 | 0.281242 | Other Negative Correlation |
| TUBG1 | -0.06024 | 0.28041 | Other Negative Correlation |
| SIMC1 | -0.06024 | 0.280406 | Other Negative Correlation |
| ZNF236 | -0.06027 | 0.28013 | Other Negative Correlation |
| CLEC18A | -0.06033 | 0.279708 | Other Negative Correlation |
| MYORG | -0.06043 | 0.278873 | Other Negative Correlation |
| SLC30A7 | -0.06044 | 0.278833 | Other Negative Correlation |
| POGLUT1 | -0.06044 | 0.278825 | Other Negative Correlation |
| DPH1 | -0.06045 | 0.278759 | Other Negative Correlation |

| | | | |
|---|---|---|---|
| PSTPIP1 | -0.06047 | 0.27854 | Other Negative Correlation |
| ZNF205 | -0.06049 | 0.278371 | Other Negative Correlation |
| CD276 | -0.06054 | 0.277972 | Other Negative Correlation |
| USP20 | -0.06057 | 0.277805 | Other Negative Correlation |
| TRIM21 | -0.06059 | 0.277626 | Other Negative Correlation |
| CD2AP | -0.06063 | 0.277277 | Other Negative Correlation |
| IFI30 | -0.06075 | 0.276345 | Other Negative Correlation |
| MT1A | -0.06079 | 0.275982 | Other Negative Correlation |
| GCNT2 | -0.06082 | 0.27577 | Other Negative Correlation |
| DNMT3A | -0.06091 | 0.275065 | Other Negative Correlation |
| OSBPL7 | -0.06095 | 0.274761 | Other Negative Correlation |
| STX6 | -0.06097 | 0.274559 | Other Negative Correlation |
| MAGT1 | -0.06099 | 0.274403 | Other Negative Correlation |
| SULT1B1 | -0.06104 | 0.274074 | Other Negative Correlation |
| IRF2 | -0.06104 | 0.274073 | Other Negative Correlation |
| FRAS1 | -0.06104 | 0.27401 | Other Negative Correlation |
| TRAPPC12 | -0.06106 | 0.273852 | Other Negative Correlation |
| SEPTIN8 | -0.06118 | 0.272938 | Other Negative Correlation |
| SYP | -0.06125 | 0.272374 | Other Negative Correlation |
| FUT6 | -0.06127 | 0.272218 | Other Negative Correlation |
| TRIM27 | -0.06127 | 0.272208 | Other Negative Correlation |
| ASCC2 | -0.06128 | 0.272137 | Other Negative Correlation |
| MBNL2 | -0.06155 | 0.270054 | Other Negative Correlation |
| CYB5D2 | -0.06157 | 0.269921 | Other Negative Correlation |
| WSB1 | -0.06157 | 0.269912 | Other Negative Correlation |
| STXBP5 | -0.0616 | 0.269647 | Other Negative Correlation |
| RNF39 | -0.06166 | 0.26922 | Other Negative Correlation |
| UBQLNL | -0.06181 | 0.268043 | Other Negative Correlation |
| NCKAP1 | -0.06185 | 0.267728 | Other Negative Correlation |
| EME2 | -0.06186 | 0.267677 | Other Negative Correlation |
| RABEP1 | -0.06186 | 0.267612 | Other Negative Correlation |
| SLC27A3 | -0.06188 | 0.26746 | Other Negative Correlation |
| PINLYP | -0.0619 | 0.267368 | Other Negative Correlation |
| NANOGP8 | -0.06192 | 0.267145 | Other Negative Correlation |
| CUBN | -0.06197 | 0.266774 | Other Negative Correlation |
| IL6ST | -0.06198 | 0.266734 | Other Negative Correlation |
| CLDND1 | -0.06212 | 0.26561 | Other Negative Correlation |
| BAALC | -0.06219 | 0.265103 | Other Negative Correlation |
| CYB5R2 | -0.0622 | 0.265033 | Other Negative Correlation |
| ACTR3C | -0.0622 | 0.265 | Other Negative Correlation |
| COG1 | -0.06224 | 0.264706 | Other Negative Correlation |
| TMEM238 | -0.0623 | 0.264276 | Other Negative Correlation |
| DAD1 | -0.06231 | 0.264143 | Other Negative Correlation |
| ILF2 | -0.06232 | 0.264113 | Other Negative Correlation |
| CUL1 | -0.06232 | 0.264094 | Other Negative Correlation |
| DPP3 | -0.06233 | 0.264017 | Other Negative Correlation |
| COPZ1 | -0.06236 | 0.263805 | Other Negative Correlation |
| MYCBP2 | -0.06239 | 0.263579 | Other Negative Correlation |

| | | | |
|---|---|---|---|
| DENND6B | -0.0624 | 0.263482 | Other Negative Correlation |
| MYD88 | -0.06241 | 0.263383 | Other Negative Correlation |
| CHST14 | -0.06243 | 0.263277 | Other Negative Correlation |
| PAIP1 | -0.06254 | 0.262407 | Other Negative Correlation |
| MMP19 | -0.06254 | 0.2624 | Other Negative Correlation |
| ENTPD3 | -0.06257 | 0.262203 | Other Negative Correlation |
| SERPINB13 | -0.06257 | 0.262162 | Other Negative Correlation |
| ZMIZ2 | -0.06267 | 0.261438 | Other Negative Correlation |
| GPBP1L1 | -0.06267 | 0.261428 | Other Negative Correlation |
| ELF4 | -0.06273 | 0.260991 | Other Negative Correlation |
| EFHC1 | -0.06276 | 0.26073 | Other Negative Correlation |
| CYTH3 | -0.06286 | 0.259997 | Other Negative Correlation |
| IQCA1 | -0.06286 | 0.259979 | Other Negative Correlation |
| PCGF2 | -0.06288 | 0.259826 | Other Negative Correlation |
| PPME1 | -0.06289 | 0.259761 | Other Negative Correlation |
| KREMEN2 | -0.0629 | 0.259636 | Other Negative Correlation |
| TIA1 | -0.06292 | 0.259511 | Other Negative Correlation |
| PGGHG | -0.06292 | 0.259505 | Other Negative Correlation |
| UBE3C | -0.06292 | 0.259491 | Other Negative Correlation |
| BEST4 | -0.06293 | 0.259445 | Other Negative Correlation |
| ZNF513 | -0.06298 | 0.259027 | Other Negative Correlation |
| ADIPOR2 | -0.06303 | 0.258698 | Other Negative Correlation |
| MSRA | -0.06309 | 0.258224 | Other Negative Correlation |
| LPCAT4 | -0.06312 | 0.257971 | Other Negative Correlation |
| NPHP3 | -0.06327 | 0.256881 | Other Negative Correlation |
| ZNF691 | -0.06327 | 0.256842 | Other Negative Correlation |
| FANCL | -0.06331 | 0.256588 | Other Negative Correlation |
| TTC23L | -0.06333 | 0.256446 | Other Negative Correlation |
| OAT | -0.06336 | 0.256208 | Other Negative Correlation |
| DNMBP | -0.06343 | 0.255662 | Other Negative Correlation |
| SNX13 | -0.06346 | 0.255448 | Other Negative Correlation |
| WDR47 | -0.06349 | 0.255227 | Other Negative Correlation |
| NFS1 | -0.06359 | 0.254468 | Other Negative Correlation |
| TTBK2 | -0.0636 | 0.25441 | Other Negative Correlation |
| HLCS | -0.06366 | 0.253959 | Other Negative Correlation |
| MFAP1 | -0.06366 | 0.253918 | Other Negative Correlation |
| COLGALT1 | -0.06372 | 0.253526 | Other Negative Correlation |
| STARD13 | -0.06375 | 0.25329 | Other Negative Correlation |
| ANKRA2 | -0.06379 | 0.25299 | Other Negative Correlation |
| PCYOX1L | -0.0638 | 0.252907 | Other Negative Correlation |
| INTS7 | -0.06382 | 0.252725 | Other Negative Correlation |
| SLC25A26 | -0.06385 | 0.252557 | Other Negative Correlation |
| CD226 | -0.06388 | 0.252284 | Other Negative Correlation |
| KLC1 | -0.06395 | 0.25181 | Other Negative Correlation |
| TMEM161A | -0.06395 | 0.251792 | Other Negative Correlation |
| POM121C | -0.06396 | 0.251721 | Other Negative Correlation |
| OTUD5 | -0.064 | 0.2514 | Other Negative Correlation |
| PRDM1 | -0.06401 | 0.251355 | Other Negative Correlation |

| | | | |
|---|---|---|---|
| ZNF282 | -0.06403 | 0.251176 | Other Negative Correlation |
| DCXR | -0.06406 | 0.250997 | Other Negative Correlation |
| SAXO2 | -0.06415 | 0.250304 | Other Negative Correlation |
| TMEM181 | -0.06416 | 0.250235 | Other Negative Correlation |
| PLLP | -0.06422 | 0.249806 | Other Negative Correlation |
| IP6K2 | -0.06424 | 0.249626 | Other Negative Correlation |
| MMP2 | -0.06425 | 0.249564 | Other Negative Correlation |
| KYNU | -0.06426 | 0.249469 | Other Negative Correlation |
| NUDT18 | -0.06427 | 0.249432 | Other Negative Correlation |
| PIK3R2 | -0.06428 | 0.249318 | Other Negative Correlation |
| C12orf76 | -0.0643 | 0.249178 | Other Negative Correlation |
| COA6 | -0.06433 | 0.248959 | Other Negative Correlation |
| TBCCD1 | -0.0644 | 0.248445 | Other Negative Correlation |
| RBBP9 | -0.06446 | 0.248025 | Other Negative Correlation |
| RETREG3 | -0.06447 | 0.247952 | Other Negative Correlation |
| SCAMP1 | -0.06455 | 0.247381 | Other Negative Correlation |
| ZNF18 | -0.06457 | 0.247195 | Other Negative Correlation |
| SHC4 | -0.06464 | 0.246721 | Other Negative Correlation |
| CHST7 | -0.06466 | 0.246512 | Other Negative Correlation |
| STK4 | -0.06472 | 0.246086 | Other Negative Correlation |
| CREBL2 | -0.06478 | 0.245645 | Other Negative Correlation |
| RAE1 | -0.06483 | 0.245267 | Other Negative Correlation |
| TAF13 | -0.06484 | 0.24526 | Other Negative Correlation |
| MYO1D | -0.06493 | 0.244541 | Other Negative Correlation |
| PRPSAP2 | -0.06506 | 0.243615 | Other Negative Correlation |
| SELENOF | -0.06508 | 0.243511 | Other Negative Correlation |
| LRG1 | -0.06512 | 0.243179 | Other Negative Correlation |
| BDH2 | -0.06516 | 0.242885 | Other Negative Correlation |
| PIGV | -0.06518 | 0.24279 | Other Negative Correlation |
| RSBN1 | -0.06523 | 0.242386 | Other Negative Correlation |
| GTF2I | -0.06529 | 0.24196 | Other Negative Correlation |
| ARL14 | -0.06532 | 0.241756 | Other Negative Correlation |
| MAK | -0.06532 | 0.241712 | Other Negative Correlation |
| HEPHL1 | -0.06544 | 0.240909 | Other Negative Correlation |
| TBCB | -0.06548 | 0.24057 | Other Negative Correlation |
| EXTL2 | -0.06557 | 0.239936 | Other Negative Correlation |
| CCNO | -0.06561 | 0.239671 | Other Negative Correlation |
| HLA-DOB | -0.06566 | 0.239328 | Other Negative Correlation |
| TACO1 | -0.06566 | 0.239316 | Other Negative Correlation |
| ZBTB1 | -0.06566 | 0.239277 | Other Negative Correlation |
| ADAM12 | -0.06569 | 0.239098 | Other Negative Correlation |
| TM9SF3 | -0.0657 | 0.238996 | Other Negative Correlation |
| PRDM10 | -0.06571 | 0.238947 | Other Negative Correlation |
| MRPL44 | -0.06573 | 0.238792 | Other Negative Correlation |
| H2BC21 | -0.06574 | 0.238729 | Other Negative Correlation |
| FANCF | -0.06584 | 0.23798 | Other Negative Correlation |
| DDX43 | -0.06588 | 0.237738 | Other Negative Correlation |
| CBFB | -0.06595 | 0.237191 | Other Negative Correlation |

| | | | |
|---|---|---|---|
| LAMTOR5 | -0.06601 | 0.236805 | Other Negative Correlation |
| HOXC4 | -0.06603 | 0.236655 | Other Negative Correlation |
| TTC1 | -0.06607 | 0.236342 | Other Negative Correlation |
| SLC9A1 | -0.0661 | 0.236152 | Other Negative Correlation |
| UBE2L5 | -0.06613 | 0.235951 | Other Negative Correlation |
| PTPN6 | -0.06613 | 0.235926 | Other Negative Correlation |
| GXYLT2 | -0.06613 | 0.235924 | Other Negative Correlation |
| RNF11 | -0.06626 | 0.235033 | Other Negative Correlation |
| ZFAND1 | -0.06628 | 0.234856 | Other Negative Correlation |
| ETAA1 | -0.0663 | 0.234721 | Other Negative Correlation |
| CERS6 | -0.06639 | 0.234096 | Other Negative Correlation |
| KRT9 | -0.06639 | 0.234077 | Other Negative Correlation |
| GLI3 | -0.06643 | 0.233824 | Other Negative Correlation |
| FBXL5 | -0.06649 | 0.233374 | Other Negative Correlation |
| NDUFC2 | -0.0665 | 0.233293 | Other Negative Correlation |
| SPATA6L | -0.06651 | 0.233275 | Other Negative Correlation |
| PLEKHO2 | -0.0666 | 0.232606 | Other Negative Correlation |
| CAPN1 | -0.06667 | 0.232163 | Other Negative Correlation |
| USF2 | -0.0667 | 0.231907 | Other Negative Correlation |
| RBM27 | -0.06676 | 0.231514 | Other Negative Correlation |
| FBXO3 | -0.06679 | 0.231308 | Other Negative Correlation |
| GGT6 | -0.0668 | 0.231224 | Other Negative Correlation |
| SKIC8 | -0.06683 | 0.230992 | Other Negative Correlation |
| MRPS18A | -0.06688 | 0.230624 | Other Negative Correlation |
| LXN | -0.06698 | 0.229988 | Other Negative Correlation |
| ZCWPW2 | -0.06698 | 0.229927 | Other Negative Correlation |
| BRAF | -0.06699 | 0.229886 | Other Negative Correlation |
| HEXIM1 | -0.06704 | 0.229531 | Other Negative Correlation |
| RABGAP1L | -0.06708 | 0.229275 | Other Negative Correlation |
| ANXA11 | -0.06712 | 0.229014 | Other Negative Correlation |
| ZNF408 | -0.06718 | 0.228571 | Other Negative Correlation |
| APOD | -0.06721 | 0.228353 | Other Negative Correlation |
| E2F5 | -0.06724 | 0.228178 | Other Negative Correlation |
| FAM200A | -0.06724 | 0.228143 | Other Negative Correlation |
| TRIM65 | -0.0673 | 0.227722 | Other Negative Correlation |
| GATB | -0.06743 | 0.226838 | Other Negative Correlation |
| CYFIP2 | -0.06748 | 0.226494 | Other Negative Correlation |
| PCSK5 | -0.06756 | 0.225962 | Other Negative Correlation |
| PLEKHA2 | -0.06758 | 0.225772 | Other Negative Correlation |
| CTR9 | -0.06762 | 0.225525 | Other Negative Correlation |
| CLIC3 | -0.06765 | 0.225357 | Other Negative Correlation |
| HRNR | -0.06766 | 0.225274 | Other Negative Correlation |
| SLC5A2 | -0.06776 | 0.2246 | Other Negative Correlation |
| VIL1 | -0.06783 | 0.224071 | Other Negative Correlation |
| CBLN3 | -0.0679 | 0.223642 | Other Negative Correlation |
| STMN3 | -0.0679 | 0.223622 | Other Negative Correlation |
| RNF13 | -0.06792 | 0.223506 | Other Negative Correlation |
| WRNIP1 | -0.06792 | 0.223481 | Other Negative Correlation |

| | | | |
|---|---|---|---|
| NSDHL | -0.06792 | 0.22346 | Other Negative Correlation |
| SIKE1 | -0.06802 | 0.222765 | Other Negative Correlation |
| DZIP3 | -0.06803 | 0.222718 | Other Negative Correlation |
| PDZK1IP1 | -0.06805 | 0.222567 | Other Negative Correlation |
| GNPTAB | -0.0681 | 0.222269 | Other Negative Correlation |
| AFAP1L2 | -0.06819 | 0.221647 | Other Negative Correlation |
| TTLL3 | -0.06821 | 0.221496 | Other Negative Correlation |
| COG5 | -0.06823 | 0.221394 | Other Negative Correlation |
| ST13 | -0.06823 | 0.221338 | Other Negative Correlation |
| TBC1D19 | -0.06835 | 0.220537 | Other Negative Correlation |
| KPNA7 | -0.06843 | 0.220025 | Other Negative Correlation |
| INTS12 | -0.06853 | 0.219314 | Other Negative Correlation |
| KRT23 | -0.06855 | 0.219201 | Other Negative Correlation |
| VANGL2 | -0.06859 | 0.218955 | Other Negative Correlation |
| TREM1 | -0.06863 | 0.218682 | Other Negative Correlation |
| HLA-E | -0.06873 | 0.218 | Other Negative Correlation |
| ACADSB | -0.06886 | 0.217124 | Other Negative Correlation |
| MEIS2 | -0.06894 | 0.216613 | Other Negative Correlation |
| TP53I3 | -0.06894 | 0.216582 | Other Negative Correlation |
| SLC7A6 | -0.06896 | 0.216474 | Other Negative Correlation |
| MYO9A | -0.06907 | 0.215735 | Other Negative Correlation |
| STX19 | -0.06909 | 0.215606 | Other Negative Correlation |
| CFAP69 | -0.06911 | 0.215476 | Other Negative Correlation |
| CAVIN2 | -0.06914 | 0.215253 | Other Negative Correlation |
| KLHL2 | -0.06917 | 0.215069 | Other Negative Correlation |
| CRYBB2 | -0.06918 | 0.214954 | Other Negative Correlation |
| SLC16A13 | -0.06925 | 0.214518 | Other Negative Correlation |
| IMPA1 | -0.06926 | 0.214419 | Other Negative Correlation |
| MED13L | -0.06927 | 0.214387 | Other Negative Correlation |
| IBTK | -0.06933 | 0.214016 | Other Negative Correlation |
| MBP | -0.06941 | 0.21349 | Other Negative Correlation |
| FKBP7 | -0.06945 | 0.213192 | Other Negative Correlation |
| MIPEP | -0.06946 | 0.213143 | Other Negative Correlation |
| SLFN14 | -0.06954 | 0.21263 | Other Negative Correlation |
| PECAM1 | -0.06957 | 0.212401 | Other Negative Correlation |
| PLEKHG2 | -0.06962 | 0.212047 | Other Negative Correlation |
| BACH2 | -0.06971 | 0.211454 | Other Negative Correlation |
| PPP2R3A | -0.06975 | 0.211226 | Other Negative Correlation |
| MYO15B | -0.06978 | 0.211052 | Other Negative Correlation |
| PLCD4 | -0.06978 | 0.211047 | Other Negative Correlation |
| FGFR2 | -0.06981 | 0.210797 | Other Negative Correlation |
| KCTD10 | -0.06982 | 0.210777 | Other Negative Correlation |
| RING1 | -0.06985 | 0.210553 | Other Negative Correlation |
| REEP3 | -0.0699 | 0.210255 | Other Negative Correlation |
| FMOD | -0.06992 | 0.210106 | Other Negative Correlation |
| TMEM179B | -0.07007 | 0.209134 | Other Negative Correlation |
| GRIA1 | -0.07008 | 0.209074 | Other Negative Correlation |
| SLC43A2 | -0.07014 | 0.208683 | Other Negative Correlation |

| | | | |
|---|---|---|---|
| ZNF224 | -0.07015 | 0.208585 | Other Negative Correlation |
| ALG11 | -0.07021 | 0.208223 | Other Negative Correlation |
| CMIP | -0.07026 | 0.207917 | Other Negative Correlation |
| ENSG00000290263 | -0.0703 | 0.207604 | Other Negative Correlation |
| HAAO | -0.07033 | 0.20745 | Other Negative Correlation |
| CARD11 | -0.07033 | 0.20744 | Other Negative Correlation |
| GAMT | -0.07056 | 0.205981 | Other Negative Correlation |
| GPR85 | -0.07059 | 0.205728 | Other Negative Correlation |
| VPS41 | -0.07059 | 0.205726 | Other Negative Correlation |
| FAM241A | -0.0706 | 0.205664 | Other Negative Correlation |
| ADAM17 | -0.0708 | 0.204397 | Other Negative Correlation |
| TMEM44 | -0.07081 | 0.204358 | Other Negative Correlation |
| NDFIP1 | -0.07082 | 0.204304 | Other Negative Correlation |
| NEK11 | -0.07083 | 0.204245 | Other Negative Correlation |
| GPCPD1 | -0.07091 | 0.203714 | Other Negative Correlation |
| SMOC2 | -0.07093 | 0.203583 | Other Negative Correlation |
| STAU2 | -0.07093 | 0.203574 | Other Negative Correlation |
| FABP4 | -0.07096 | 0.203391 | Other Negative Correlation |
| KANK3 | -0.07098 | 0.203255 | Other Negative Correlation |
| CCDC142 | -0.07102 | 0.203024 | Other Negative Correlation |
| NCK2 | -0.07107 | 0.202714 | Other Negative Correlation |
| DCDC2B | -0.07113 | 0.202318 | Other Negative Correlation |
| SCRN2 | -0.07114 | 0.202243 | Other Negative Correlation |
| PRDM6 | -0.07115 | 0.202178 | Other Negative Correlation |
| GOPC | -0.0712 | 0.201885 | Other Negative Correlation |
| SEC16B | -0.07121 | 0.201821 | Other Negative Correlation |
| CCDC102A | -0.07128 | 0.201334 | Other Negative Correlation |
| DMTF1 | -0.0713 | 0.201219 | Other Negative Correlation |
| PCDHGA12 | -0.07138 | 0.200717 | Other Negative Correlation |
| ZNF813 | -0.07139 | 0.200646 | Other Negative Correlation |
| ZNF480 | -0.07145 | 0.200269 | Other Negative Correlation |
| CTSB | -0.0715 | 0.199948 | Other Negative Correlation |
| DNAJC24 | -0.07153 | 0.199784 | Other Negative Correlation |
| PPIL6 | -0.07157 | 0.199494 | Other Negative Correlation |
| LANCL1 | -0.07158 | 0.199437 | Other Negative Correlation |
| RAB11A | -0.07161 | 0.199285 | Other Negative Correlation |
| PHACTR4 | -0.07161 | 0.199284 | Other Negative Correlation |
| TMEM67 | -0.07166 | 0.198963 | Other Negative Correlation |
| KMT5B | -0.07167 | 0.198896 | Other Negative Correlation |
| VPS39 | -0.0717 | 0.198675 | Other Negative Correlation |
| NSMF | -0.07171 | 0.198609 | Other Negative Correlation |
| SIN3B | -0.07176 | 0.198334 | Other Negative Correlation |
| VWA5A | -0.07182 | 0.197958 | Other Negative Correlation |
| RXRB | -0.07187 | 0.197616 | Other Negative Correlation |
| TOP2B | -0.07187 | 0.197609 | Other Negative Correlation |
| WARS1 | -0.07188 | 0.197587 | Other Negative Correlation |
| UBXN4 | -0.07188 | 0.197547 | Other Negative Correlation |
| CEP164 | -0.07189 | 0.197529 | Other Negative Correlation |

| | | | |
|---|---|---|---|
| ARL14EP | -0.07191 | 0.197368 | Other Negative Correlation |
| RIPOR3 | -0.07193 | 0.197232 | Other Negative Correlation |
| PTK2 | -0.07202 | 0.196711 | Other Negative Correlation |
| WNT7A | -0.07202 | 0.196707 | Other Negative Correlation |
| PAQR3 | -0.07209 | 0.19629 | Other Negative Correlation |
| TSPAN9 | -0.0721 | 0.196171 | Other Negative Correlation |
| TCF12 | -0.07228 | 0.195105 | Other Negative Correlation |
| PIGS | -0.07228 | 0.195059 | Other Negative Correlation |
| UBALD1 | -0.07228 | 0.195055 | Other Negative Correlation |
| FKBP8 | -0.07229 | 0.195013 | Other Negative Correlation |
| GFM2 | -0.07241 | 0.194263 | Other Negative Correlation |
| TASP1 | -0.07253 | 0.193509 | Other Negative Correlation |
| PLEKHN1 | -0.07254 | 0.193484 | Other Negative Correlation |
| NBPF1 | -0.07259 | 0.193193 | Other Negative Correlation |
| GRM3 | -0.07262 | 0.192985 | Other Negative Correlation |
| PMPCB | -0.07269 | 0.192572 | Other Negative Correlation |
| SLC16A2 | -0.0727 | 0.192484 | Other Negative Correlation |
| SIRPB2 | -0.07272 | 0.192376 | Other Negative Correlation |
| RDX | -0.07285 | 0.191549 | Other Negative Correlation |
| YME1L1 | -0.07288 | 0.191418 | Other Negative Correlation |
| PRRC2A | -0.07291 | 0.19119 | Other Negative Correlation |
| FAM171A1 | -0.07292 | 0.191143 | Other Negative Correlation |
| CHD6 | -0.07304 | 0.190434 | Other Negative Correlation |
| IRF3 | -0.07304 | 0.190398 | Other Negative Correlation |
| NIPAL3 | -0.07307 | 0.190234 | Other Negative Correlation |
| LPXN | -0.07312 | 0.189913 | Other Negative Correlation |
| ENKUR | -0.07318 | 0.189554 | Other Negative Correlation |
| CCDC191 | -0.07329 | 0.188915 | Other Negative Correlation |
| C1GALT1 | -0.07329 | 0.18891 | Other Negative Correlation |
| CIR1 | -0.07336 | 0.188489 | Other Negative Correlation |
| IZUMO1 | -0.07338 | 0.188373 | Other Negative Correlation |
| LRRC75B | -0.0734 | 0.188205 | Other Negative Correlation |
| MRPS30 | -0.07358 | 0.187136 | Other Negative Correlation |
| SERINC5 | -0.0736 | 0.18704 | Other Negative Correlation |
| TMEM200A | -0.0737 | 0.18641 | Other Negative Correlation |
| G6PC3 | -0.07373 | 0.186278 | Other Negative Correlation |
| SATB1 | -0.07382 | 0.185725 | Other Negative Correlation |
| LMBR1 | -0.07387 | 0.18542 | Other Negative Correlation |
| DLD | -0.07387 | 0.185404 | Other Negative Correlation |
| KCNE1 | -0.07392 | 0.185097 | Other Negative Correlation |
| TRIM41 | -0.07396 | 0.184858 | Other Negative Correlation |
| GRAMD1C | -0.074 | 0.18465 | Other Negative Correlation |
| CDA | -0.07402 | 0.184505 | Other Negative Correlation |
| PLB1 | -0.07424 | 0.183202 | Other Negative Correlation |
| NOX4 | -0.07424 | 0.183194 | Other Negative Correlation |
| ERCC5 | -0.07433 | 0.182691 | Other Negative Correlation |
| NUDT6 | -0.07449 | 0.181714 | Other Negative Correlation |
| ORMDL2 | -0.07452 | 0.181587 | Other Negative Correlation |

| | | | |
|---|---|---|---|
| ENSG00000275674 | -0.07452 | 0.181568 | Other Negative Correlation |
| SULT1E1 | -0.07457 | 0.181286 | Other Negative Correlation |
| RBM33 | -0.07463 | 0.180942 | Other Negative Correlation |
| SSBP4 | -0.07475 | 0.180188 | Other Negative Correlation |
| KCTD14 | -0.07476 | 0.180174 | Other Negative Correlation |
| SHOC1 | -0.07476 | 0.180155 | Other Negative Correlation |
| OSBPL5 | -0.07484 | 0.179705 | Other Negative Correlation |
| KCNRG | -0.07491 | 0.179298 | Other Negative Correlation |
| B9D1 | -0.07494 | 0.179113 | Other Negative Correlation |
| UROS | -0.07497 | 0.178908 | Other Negative Correlation |
| ARMT1 | -0.07504 | 0.178542 | Other Negative Correlation |
| FCHSD1 | -0.07506 | 0.178417 | Other Negative Correlation |
| CCNT1 | -0.07506 | 0.17841 | Other Negative Correlation |
| RB1 | -0.07509 | 0.178224 | Other Negative Correlation |
| AQP3 | -0.07517 | 0.17777 | Other Negative Correlation |
| CHD1L | -0.07531 | 0.176956 | Other Negative Correlation |
| FBXO31 | -0.07532 | 0.17692 | Other Negative Correlation |
| EYA2 | -0.07532 | 0.176911 | Other Negative Correlation |
| BCL2L13 | -0.07536 | 0.176684 | Other Negative Correlation |
| SIL1 | -0.07541 | 0.176409 | Other Negative Correlation |
| DKKL1 | -0.07544 | 0.176206 | Other Negative Correlation |
| MKKS | -0.0755 | 0.175859 | Other Negative Correlation |
| TPTEP2-CSNK1E | -0.07552 | 0.175766 | Other Negative Correlation |
| INTS2 | -0.07553 | 0.17569 | Other Negative Correlation |
| NOSTRIN | -0.07562 | 0.175196 | Other Negative Correlation |
| MT-CO3 | -0.07572 | 0.174619 | Other Negative Correlation |
| WDR6 | -0.07574 | 0.174504 | Other Negative Correlation |
| SMARCA5 | -0.07577 | 0.174351 | Other Negative Correlation |
| SETX | -0.07587 | 0.173736 | Other Negative Correlation |
| TOLLIP | -0.07595 | 0.173315 | Other Negative Correlation |
| ITGA10 | -0.07596 | 0.173257 | Other Negative Correlation |
| GNL1 | -0.07599 | 0.173077 | Other Negative Correlation |
| ROCK2 | -0.07606 | 0.172682 | Other Negative Correlation |
| SLC16A5 | -0.0761 | 0.172458 | Other Negative Correlation |
| SLC35C1 | -0.07611 | 0.172423 | Other Negative Correlation |
| ARPIN | -0.07614 | 0.172231 | Other Negative Correlation |
| UBE2W | -0.07614 | 0.172212 | Other Negative Correlation |
| USB1 | -0.07616 | 0.172131 | Other Negative Correlation |
| MARK4 | -0.07616 | 0.172089 | Other Negative Correlation |
| IQCK | -0.07621 | 0.171814 | Other Negative Correlation |
| HIVEP3 | -0.07625 | 0.171604 | Other Negative Correlation |
| MAPRE1 | -0.07628 | 0.171427 | Other Negative Correlation |
| GGT7 | -0.07636 | 0.170996 | Other Negative Correlation |
| SRRD | -0.07637 | 0.170959 | Other Negative Correlation |
| ZSWIM7 | -0.07642 | 0.170676 | Other Negative Correlation |
| AFG1L | -0.07645 | 0.170473 | Other Negative Correlation |
| POGK | -0.07654 | 0.169973 | Other Negative Correlation |
| BCKDHB | -0.07656 | 0.169849 | Other Negative Correlation |

| | | | |
|---|---|---|---|
| CAMK2D | -0.07656 | 0.169844 | Other Negative Correlation |
| SWAP70 | -0.07666 | 0.169283 | Other Negative Correlation |
| GYG1 | -0.07668 | 0.169203 | Other Negative Correlation |
| ACSF2 | -0.07676 | 0.168759 | Other Negative Correlation |
| ZGLP1 | -0.07677 | 0.168684 | Other Negative Correlation |
| RCAN3 | -0.0768 | 0.168519 | Other Negative Correlation |
| TCEANC2 | -0.07681 | 0.168456 | Other Negative Correlation |
| GBF1 | -0.07682 | 0.168441 | Other Negative Correlation |
| LRFN4 | -0.07689 | 0.168053 | Other Negative Correlation |
| TFEB | -0.07699 | 0.167451 | Other Negative Correlation |
| NEFL | -0.077 | 0.16744 | Other Negative Correlation |
| ECT2L | -0.077 | 0.167394 | Other Negative Correlation |
| CCDC43 | -0.07703 | 0.167241 | Other Negative Correlation |
| LOXL1 | -0.07706 | 0.167103 | Other Negative Correlation |
| ZNF512B | -0.07709 | 0.166908 | Other Negative Correlation |
| ITGA11 | -0.07713 | 0.166724 | Other Negative Correlation |
| MRPS9 | -0.07713 | 0.166722 | Other Negative Correlation |
| TMEM170A | -0.07716 | 0.166547 | Other Negative Correlation |
| OSBPL2 | -0.07718 | 0.166404 | Other Negative Correlation |
| ENSG00000285827 | -0.07723 | 0.166132 | Other Negative Correlation |
| ERC1 | -0.07735 | 0.165485 | Other Negative Correlation |
| COMMD7 | -0.07739 | 0.165296 | Other Negative Correlation |
| HACL1 | -0.07747 | 0.16484 | Other Negative Correlation |
| TMEM101 | -0.07751 | 0.16463 | Other Negative Correlation |
| EVI5 | -0.07753 | 0.164516 | Other Negative Correlation |
| CLINT1 | -0.07753 | 0.164492 | Other Negative Correlation |
| ENTREP3 | -0.07767 | 0.163725 | Other Negative Correlation |
| DHX36 | -0.07769 | 0.163632 | Other Negative Correlation |
| CTSK | -0.07771 | 0.163548 | Other Negative Correlation |
| APOA2 | -0.07771 | 0.163542 | Other Negative Correlation |
| FHIP2A | -0.07774 | 0.163343 | Other Negative Correlation |
| ANGEL2 | -0.07783 | 0.162862 | Other Negative Correlation |
| TAFA5 | -0.07784 | 0.162802 | Other Negative Correlation |
| KANSL3 | -0.07804 | 0.161757 | Other Negative Correlation |
| NLRC4 | -0.07809 | 0.161457 | Other Negative Correlation |
| GABRA1 | -0.07809 | 0.161457 | Other Negative Correlation |
| ZNF493 | -0.07812 | 0.161319 | Other Negative Correlation |
| ZNF512 | -0.07816 | 0.161114 | Other Negative Correlation |
| EP400 | -0.07817 | 0.161053 | Other Negative Correlation |
| SMAD1 | -0.0782 | 0.160891 | Other Negative Correlation |
| WDR19 | -0.07826 | 0.16056 | Other Negative Correlation |
| FAM91A1 | -0.07832 | 0.160251 | Other Negative Correlation |
| GNAQ | -0.07837 | 0.159975 | Other Negative Correlation |
| NDUFA1 | -0.07838 | 0.159896 | Other Negative Correlation |
| PTPN9 | -0.07839 | 0.159851 | Other Negative Correlation |
| NR4A3 | -0.07843 | 0.159656 | Other Negative Correlation |
| KCTD20 | -0.07847 | 0.159449 | Other Negative Correlation |
| TRIP12 | -0.07847 | 0.15941 | Other Negative Correlation |

| | | | |
|---|---|---|---|
| EHHADH | -0.07848 | 0.159355 | Other Negative Correlation |
| NEDD9 | -0.07851 | 0.159227 | Other Negative Correlation |
| NGLY1 | -0.07865 | 0.158495 | Other Negative Correlation |
| WDR41 | -0.07868 | 0.158333 | Other Negative Correlation |
| LRSAM1 | -0.07883 | 0.157531 | Other Negative Correlation |
| VCP | -0.07884 | 0.157462 | Other Negative Correlation |
| ATP5MK | -0.0789 | 0.15713 | Other Negative Correlation |
| PISD | -0.07899 | 0.156674 | Other Negative Correlation |
| MED14 | -0.07902 | 0.1565 | Other Negative Correlation |
| ARHGAP22 | -0.0791 | 0.156112 | Other Negative Correlation |
| DTNB | -0.07928 | 0.155164 | Other Negative Correlation |
| CRLF3 | -0.07935 | 0.154775 | Other Negative Correlation |
| GRIPAP1 | -0.07939 | 0.154582 | Other Negative Correlation |
| GLB1L | -0.07948 | 0.15411 | Other Negative Correlation |
| STRN | -0.0796 | 0.153466 | Other Negative Correlation |
| DAAM1 | -0.07963 | 0.153354 | Other Negative Correlation |
| MST1R | -0.07974 | 0.152766 | Other Negative Correlation |
| ALDH2 | -0.07979 | 0.152492 | Other Negative Correlation |
| PLA2G4C | -0.0798 | 0.152437 | Other Negative Correlation |
| SLAMF7 | -0.07982 | 0.152353 | Other Negative Correlation |
| TRMT2A | -0.07996 | 0.151617 | Other Negative Correlation |
| CASP3 | -0.07998 | 0.151527 | Other Negative Correlation |
| WDFY3 | -0.08 | 0.151437 | Other Negative Correlation |
| WDR11 | -0.08 | 0.151437 | Other Negative Correlation |
| BICRAL | -0.08012 | 0.150803 | Other Negative Correlation |
| TAFAZZIN | -0.08013 | 0.150742 | Other Negative Correlation |
| PLXNB2 | -0.08015 | 0.150676 | Other Negative Correlation |
| ZNF516 | -0.08015 | 0.15064 | Other Negative Correlation |
| PCMTD2 | -0.08017 | 0.150578 | Other Negative Correlation |
| BLTP2 | -0.08017 | 0.150554 | Other Negative Correlation |
| LAMTOR2 | -0.08018 | 0.150512 | Other Negative Correlation |
| SLC22A3 | -0.0802 | 0.150413 | Other Negative Correlation |
| ZMYM6 | -0.08024 | 0.150183 | Other Negative Correlation |
| ETV2 | -0.08027 | 0.150044 | Other Negative Correlation |
| NDUFB9 | -0.08029 | 0.14993 | Other Negative Correlation |
| MBTD1 | -0.0803 | 0.149893 | Other Negative Correlation |
| TPCN1 | -0.08031 | 0.149834 | Other Negative Correlation |
| ENSG00000241962 | -0.08035 | 0.14965 | Other Negative Correlation |
| ISCA1 | -0.08036 | 0.149598 | Other Negative Correlation |
| ZDHHC11 | -0.08042 | 0.1493 | Other Negative Correlation |
| OAF | -0.08042 | 0.149297 | Other Negative Correlation |
| ENSG00000287856 | -0.08043 | 0.149229 | Other Negative Correlation |
| SPIRE1 | -0.08047 | 0.149011 | Other Negative Correlation |
| PAPSS2 | -0.08052 | 0.148778 | Other Negative Correlation |
| B4GALT6 | -0.08063 | 0.148228 | Other Negative Correlation |
| PRPF8 | -0.08072 | 0.147787 | Other Negative Correlation |
| LRRK1 | -0.08072 | 0.147774 | Other Negative Correlation |
| CYSRT1 | -0.08087 | 0.14704 | Other Negative Correlation |

| | | | |
|---|---|---|---|
| TRNAU1AP | -0.08087 | 0.147027 | Other Negative Correlation |
| LPAR5 | -0.08087 | 0.147005 | Other Negative Correlation |
| CCDC22 | -0.08087 | 0.147003 | Other Negative Correlation |
| NTF4 | -0.08101 | 0.146317 | Other Negative Correlation |
| THUMPD1 | -0.08102 | 0.146248 | Other Negative Correlation |
| CCDC57 | -0.08105 | 0.146108 | Other Negative Correlation |
| LRRC57 | -0.08116 | 0.145544 | Other Negative Correlation |
| PRRC1 | -0.08124 | 0.145189 | Other Negative Correlation |
| HRCT1 | -0.08128 | 0.14496 | Other Negative Correlation |
| ANKK1 | -0.08134 | 0.144661 | Other Negative Correlation |
| ZNF277 | -0.08138 | 0.144455 | Other Negative Correlation |
| AP4E1 | -0.08139 | 0.144413 | Other Negative Correlation |
| TMEM17 | -0.0814 | 0.144366 | Other Negative Correlation |
| ZNF30 | -0.08161 | 0.143332 | Other Negative Correlation |
| NUP210L | -0.08177 | 0.142577 | Other Negative Correlation |
| ZFY | -0.08183 | 0.142264 | Other Negative Correlation |
| GJC2 | -0.08191 | 0.141851 | Other Negative Correlation |
| PCNX1 | -0.08197 | 0.141572 | Other Negative Correlation |
| MAP3K7 | -0.08205 | 0.141192 | Other Negative Correlation |
| ALDH16A1 | -0.08207 | 0.141076 | Other Negative Correlation |
| PXDN | -0.08208 | 0.141045 | Other Negative Correlation |
| EDEM3 | -0.0821 | 0.140938 | Other Negative Correlation |
| AGAP4 | -0.08214 | 0.140739 | Other Negative Correlation |
| MYL6B | -0.08218 | 0.140578 | Other Negative Correlation |
| POGLUT2 | -0.08222 | 0.14038 | Other Negative Correlation |
| PPP2R5E | -0.08234 | 0.139784 | Other Negative Correlation |
| PABPC1L | -0.08239 | 0.139562 | Other Negative Correlation |
| ASB13 | -0.08248 | 0.139121 | Other Negative Correlation |
| AIP | -0.08248 | 0.139096 | Other Negative Correlation |
| TMEM156 | -0.08256 | 0.1387 | Other Negative Correlation |
| TCP11L2 | -0.08259 | 0.138603 | Other Negative Correlation |
| MED25 | -0.08275 | 0.137819 | Other Negative Correlation |
| HNRNPLL | -0.08279 | 0.137636 | Other Negative Correlation |
| BPHL | -0.08282 | 0.137484 | Other Negative Correlation |
| ANKFY1 | -0.08283 | 0.137441 | Other Negative Correlation |
| ATAD1 | -0.08286 | 0.137299 | Other Negative Correlation |
| RNASEL | -0.08302 | 0.13653 | Other Negative Correlation |
| CD101 | -0.08303 | 0.136462 | Other Negative Correlation |
| TNS3 | -0.08316 | 0.135871 | Other Negative Correlation |
| AFF4 | -0.08317 | 0.135842 | Other Negative Correlation |
| N4BP2L2 | -0.08326 | 0.135417 | Other Negative Correlation |
| CAMKK1 | -0.08328 | 0.135308 | Other Negative Correlation |
| SLC35E2B | -0.08329 | 0.135256 | Other Negative Correlation |
| PRKAR1B | -0.0833 | 0.135217 | Other Negative Correlation |
| DYNLRB2 | -0.08345 | 0.134486 | Other Negative Correlation |
| TIMM22 | -0.08349 | 0.134298 | Other Negative Correlation |
| STK3 | -0.0835 | 0.134262 | Other Negative Correlation |
| TIGAR | -0.08351 | 0.134238 | Other Negative Correlation |

| | | | |
|---|---|---|---|
| MDP1 | -0.08356 | 0.133968 | Other Negative Correlation |
| GDF11 | -0.08359 | 0.133833 | Other Negative Correlation |
| NOMO1 | -0.08361 | 0.133777 | Other Negative Correlation |
| TNFSF9 | -0.08361 | 0.133772 | Other Negative Correlation |
| ZSCAN5A | -0.08371 | 0.133297 | Other Negative Correlation |
| ACOT13 | -0.08374 | 0.133156 | Other Negative Correlation |
| EFR3A | -0.0838 | 0.132871 | Other Negative Correlation |
| DKK2 | -0.08386 | 0.132573 | Other Negative Correlation |
| PPARD | -0.08391 | 0.132384 | Other Negative Correlation |
| ABAT | -0.08394 | 0.132204 | Other Negative Correlation |
| RP1 | -0.08403 | 0.13179 | Other Negative Correlation |
| MLLT10 | -0.08405 | 0.131714 | Other Negative Correlation |
| FUCA2 | -0.0841 | 0.131462 | Other Negative Correlation |
| DYNC2LI1 | -0.08419 | 0.13105 | Other Negative Correlation |
| EID1 | -0.08426 | 0.130733 | Other Negative Correlation |
| UGGT1 | -0.08428 | 0.130661 | Other Negative Correlation |
| ZBTB4 | -0.0843 | 0.130568 | Other Negative Correlation |
| PIWIL4 | -0.08431 | 0.130513 | Other Negative Correlation |
| GRAMD2B | -0.08432 | 0.130465 | Other Negative Correlation |
| PFKL | -0.08451 | 0.12961 | Other Negative Correlation |
| TLR6 | -0.08455 | 0.129433 | Other Negative Correlation |
| ZNF785 | -0.08459 | 0.129258 | Other Negative Correlation |
| KIF5B | -0.08467 | 0.128891 | Other Negative Correlation |
| TRPM7 | -0.08468 | 0.128814 | Other Negative Correlation |
| TCF7L2 | -0.08473 | 0.1286 | Other Negative Correlation |
| TRIM46 | -0.08489 | 0.127888 | Other Negative Correlation |
| LRRC41 | -0.08492 | 0.127751 | Other Negative Correlation |
| ODF3L1 | -0.08512 | 0.126848 | Other Negative Correlation |
| HCAR1 | -0.08516 | 0.126678 | Other Negative Correlation |
| NBPF15 | -0.0852 | 0.126512 | Other Negative Correlation |
| C6orf226 | -0.08532 | 0.125943 | Other Negative Correlation |
| OR2A7 | -0.08533 | 0.125933 | Other Negative Correlation |
| MRPL2 | -0.08535 | 0.125832 | Other Negative Correlation |
| NMT2 | -0.08536 | 0.12577 | Other Negative Correlation |
| UBE4B | -0.08539 | 0.125651 | Other Negative Correlation |
| ACTA1 | -0.08543 | 0.125465 | Other Negative Correlation |
| DIS3L2 | -0.08548 | 0.125241 | Other Negative Correlation |
| ERGIC1 | -0.08551 | 0.125112 | Other Negative Correlation |
| QSOX2 | -0.08552 | 0.125062 | Other Negative Correlation |
| ZNF22 | -0.08553 | 0.125005 | Other Negative Correlation |
| UBE2D1 | -0.08558 | 0.124806 | Other Negative Correlation |
| LDAF1 | -0.08564 | 0.124545 | Other Negative Correlation |
| KIF17 | -0.08566 | 0.12447 | Other Negative Correlation |
| PATJ | -0.08568 | 0.124375 | Other Negative Correlation |
| MGST2 | -0.08579 | 0.123863 | Other Negative Correlation |
| SUFU | -0.08588 | 0.123483 | Other Negative Correlation |
| TAS2R14 | -0.08593 | 0.123259 | Other Negative Correlation |
| TRABD | -0.08597 | 0.123073 | Other Negative Correlation |

| | | | |
|---|---|---|---|
| ZMYM2 | -0.08634 | 0.121495 | Other Negative Correlation |
| RALBP1 | -0.0864 | 0.121232 | Other Negative Correlation |
| CCDC88C | -0.08644 | 0.12106 | Other Negative Correlation |
| KRTAP19-3 | -0.08647 | 0.120935 | Other Negative Correlation |
| UGT1A5 | -0.08647 | 0.120935 | Other Negative Correlation |
| ATXN2 | -0.08652 | 0.12072 | Other Negative Correlation |
| MAP9 | -0.08653 | 0.120641 | Other Negative Correlation |
| UQCR11 | -0.08656 | 0.120532 | Other Negative Correlation |
| ZNF410 | -0.08663 | 0.120228 | Other Negative Correlation |
| UBR1 | -0.0867 | 0.11993 | Other Negative Correlation |
| CTSS | -0.08673 | 0.119816 | Other Negative Correlation |
| MYSM1 | -0.08676 | 0.119684 | Other Negative Correlation |
| LAGE3 | -0.08677 | 0.119647 | Other Negative Correlation |
| COA3 | -0.08679 | 0.11955 | Other Negative Correlation |
| ZMYM4 | -0.08684 | 0.11933 | Other Negative Correlation |
| NRG4 | -0.08688 | 0.119143 | Other Negative Correlation |
| WASHC4 | -0.08691 | 0.119032 | Other Negative Correlation |
| P3H3 | -0.08698 | 0.118736 | Other Negative Correlation |
| KIF1C | -0.087 | 0.118662 | Other Negative Correlation |
| OTULIN | -0.08706 | 0.118408 | Other Negative Correlation |
| CTTNBP2NL | -0.0871 | 0.118231 | Other Negative Correlation |
| RRAGA | -0.08718 | 0.117894 | Other Negative Correlation |
| PRMT9 | -0.08742 | 0.116872 | Other Negative Correlation |
| SEC14L4 | -0.08745 | 0.116762 | Other Negative Correlation |
| SARAF | -0.08745 | 0.116745 | Other Negative Correlation |
| ELMOD2 | -0.08753 | 0.116428 | Other Negative Correlation |
| PLEKHA4 | -0.08754 | 0.116351 | Other Negative Correlation |
| HOOK2 | -0.08764 | 0.115931 | Other Negative Correlation |
| HMG20A | -0.08769 | 0.115746 | Other Negative Correlation |
| HDGFL2 | -0.08772 | 0.115596 | Other Negative Correlation |
| HFE | -0.08778 | 0.115355 | Other Negative Correlation |
| GOLGA6L4 | -0.08781 | 0.115255 | Other Negative Correlation |
| SDCBP2 | -0.08782 | 0.115198 | Other Negative Correlation |
| AGBL5 | -0.08786 | 0.115036 | Other Negative Correlation |
| RWDD1 | -0.08792 | 0.114782 | Other Negative Correlation |
| FOCAD | -0.08794 | 0.114684 | Other Negative Correlation |
| BOK | -0.08798 | 0.114521 | Other Negative Correlation |
| ACTR1A | -0.08802 | 0.114367 | Other Negative Correlation |
| MBLAC2 | -0.08808 | 0.114134 | Other Negative Correlation |
| FGGY | -0.08818 | 0.11371 | Other Negative Correlation |
| PAPOLG | -0.08821 | 0.113597 | Other Negative Correlation |
| AGFG1 | -0.08824 | 0.113462 | Other Negative Correlation |
| CHTOP | -0.0883 | 0.11321 | Other Negative Correlation |
| FAM168B | -0.08833 | 0.113079 | Other Negative Correlation |
| DDX11 | -0.08834 | 0.113044 | Other Negative Correlation |
| ARMCX1 | -0.08836 | 0.112973 | Other Negative Correlation |
| TLCD5 | -0.08846 | 0.112564 | Other Negative Correlation |
| CTSO | -0.08852 | 0.112317 | Other Negative Correlation |

| | | | |
|---|---|---|---|
| KRBA1 | -0.08854 | 0.112249 | Other Negative Correlation |
| VARS2 | -0.08854 | 0.112244 | Other Negative Correlation |
| VPS53 | -0.08859 | 0.112053 | Other Negative Correlation |
| KDM1B | -0.0887 | 0.111593 | Other Negative Correlation |
| EIF2B2 | -0.08887 | 0.110919 | Other Negative Correlation |
| TANK | -0.08891 | 0.110743 | Other Negative Correlation |
| JPT2 | -0.08895 | 0.110591 | Other Negative Correlation |
| ALDH1B1 | -0.08896 | 0.110531 | Other Negative Correlation |
| SLC25A39 | -0.08902 | 0.110295 | Other Negative Correlation |
| LRP5 | -0.08909 | 0.110016 | Other Negative Correlation |
| UTY | -0.0891 | 0.109996 | Other Negative Correlation |
| CNPY3 | -0.08916 | 0.109756 | Other Negative Correlation |
| ZNF775 | -0.08918 | 0.109669 | Other Negative Correlation |
| ATG4C | -0.08918 | 0.109662 | Other Negative Correlation |
| SELENOW | -0.0892 | 0.10959 | Other Negative Correlation |
| FBXL14 | -0.08921 | 0.109544 | Other Negative Correlation |
| TMEM276 | -0.08924 | 0.109414 | Other Negative Correlation |
| TMTC3 | -0.08933 | 0.109075 | Other Negative Correlation |
| ADNP | -0.0894 | 0.108772 | Other Negative Correlation |
| CNTNAP1 | -0.08946 | 0.108556 | Other Negative Correlation |
| SLC31A2 | -0.08949 | 0.108409 | Other Negative Correlation |
| PRKRA | -0.08957 | 0.108111 | Other Negative Correlation |
| MRTFB | -0.08962 | 0.107927 | Other Negative Correlation |
| SGMS2 | -0.08962 | 0.107922 | Other Negative Correlation |
| EMC2 | -0.08962 | 0.107898 | Other Negative Correlation |
| TUT1 | -0.08963 | 0.107883 | Other Negative Correlation |
| PKP4 | -0.08974 | 0.107427 | Other Negative Correlation |
| VGLL1 | -0.08977 | 0.107299 | Other Negative Correlation |
| TMEM80 | -0.08983 | 0.107099 | Other Negative Correlation |
| SETDB1 | -0.08999 | 0.10645 | Other Negative Correlation |
| NCF1 | -0.09007 | 0.106154 | Other Negative Correlation |
| MID1 | -0.09015 | 0.105844 | Other Negative Correlation |
| CCNH | -0.09018 | 0.105717 | Other Negative Correlation |
| KIAA0319L | -0.09025 | 0.105455 | Other Negative Correlation |
| HDX | -0.09031 | 0.105215 | Other Negative Correlation |
| CDC40 | -0.09036 | 0.105033 | Other Negative Correlation |
| KLC3 | -0.09036 | 0.105028 | Other Negative Correlation |
| TXNRD3 | -0.09042 | 0.104773 | Other Negative Correlation |
| DOCK2 | -0.09047 | 0.104587 | Other Negative Correlation |
| CHURC1 | -0.09063 | 0.103999 | Other Negative Correlation |
| NUDT19 | -0.09067 | 0.103811 | Other Negative Correlation |
| RPS6KA3 | -0.09069 | 0.103752 | Other Negative Correlation |
| ANKS6 | -0.09074 | 0.103555 | Other Negative Correlation |
| KBTBD7 | -0.09075 | 0.103538 | Other Negative Correlation |
| COPS4 | -0.09085 | 0.103159 | Other Negative Correlation |
| TENM4 | -0.09086 | 0.103116 | Other Negative Correlation |
| ALDH6A1 | -0.09087 | 0.103061 | Other Negative Correlation |
| PIGX | -0.09091 | 0.102905 | Other Negative Correlation |

| | | | |
|---|---|---|---|
| MYL5 | -0.09097 | 0.102697 | Other Negative Correlation |
| NUPR1 | -0.09101 | 0.102525 | Other Negative Correlation |
| SETD7 | -0.09104 | 0.102433 | Other Negative Correlation |
| ZNF623 | -0.09109 | 0.102214 | Other Negative Correlation |
| PEX1 | -0.09118 | 0.101877 | Other Negative Correlation |
| PNKP | -0.09119 | 0.101858 | Other Negative Correlation |
| RTF1 | -0.09121 | 0.101764 | Other Negative Correlation |
| LYPLA2 | -0.09124 | 0.101663 | Other Negative Correlation |
| ENTPD6 | -0.09126 | 0.101584 | Other Negative Correlation |
| ITFG1 | -0.09138 | 0.101138 | Other Negative Correlation |
| NDUFA7 | -0.09139 | 0.101109 | Other Negative Correlation |
| CSRNP3 | -0.09139 | 0.101104 | Other Negative Correlation |
| EHBP1L1 | -0.0914 | 0.101059 | Other Negative Correlation |
| GARIN1A | -0.0914 | 0.101054 | Other Negative Correlation |
| C1QTNF12 | -0.09147 | 0.100791 | Other Negative Correlation |
| TMEM106A | -0.09153 | 0.100559 | Other Negative Correlation |
| PLCB3 | -0.09171 | 0.099915 | Other Negative Correlation |
| FUNDC1 | -0.09174 | 0.0998 | Other Negative Correlation |
| NBEAL2 | -0.09175 | 0.099739 | Other Negative Correlation |
| HSPB1 | -0.09181 | 0.099532 | Other Negative Correlation |
| ZBED1 | -0.09186 | 0.099362 | Other Negative Correlation |
| BLZF1 | -0.09191 | 0.099168 | Other Negative Correlation |
| LYRM2 | -0.09206 | 0.098601 | Other Negative Correlation |
| DCTN1 | -0.09208 | 0.098544 | Other Negative Correlation |
| LASP1 | -0.09209 | 0.098505 | Other Negative Correlation |
| ARAF | -0.09218 | 0.098178 | Other Negative Correlation |
| HEG1 | -0.09218 | 0.09816 | Other Negative Correlation |
| ASB9 | -0.09219 | 0.098127 | Other Negative Correlation |
| KIFAP3 | -0.09219 | 0.098119 | Other Negative Correlation |
| SFT2D1 | -0.09225 | 0.097922 | Other Negative Correlation |
| RHBDL2 | -0.09229 | 0.09778 | Other Negative Correlation |
| ELAC2 | -0.09235 | 0.097542 | Other Negative Correlation |
| PSMD4 | -0.09245 | 0.097184 | Other Negative Correlation |
| PPP1R12B | -0.09252 | 0.096923 | Other Negative Correlation |
| POLD4 | -0.09256 | 0.096775 | Other Negative Correlation |
| KCTD11 | -0.09269 | 0.096327 | Other Negative Correlation |
| DEAF1 | -0.09269 | 0.096324 | Other Negative Correlation |
| EXTL3 | -0.09276 | 0.096065 | Other Negative Correlation |
| METTL18 | -0.09285 | 0.095728 | Other Negative Correlation |
| SMCO2 | -0.09288 | 0.095624 | Other Negative Correlation |
| GEMIN8 | -0.09289 | 0.095588 | Other Negative Correlation |
| ETV6 | -0.09301 | 0.095155 | Other Negative Correlation |
| ABLIM3 | -0.09314 | 0.094694 | Other Negative Correlation |
| CNPY4 | -0.09318 | 0.094576 | Other Negative Correlation |
| ZNF875 | -0.09323 | 0.094393 | Other Negative Correlation |
| ANKEF1 | -0.09325 | 0.094303 | Other Negative Correlation |
| ARL17B | -0.09362 | 0.093007 | Other Negative Correlation |
| ARID1B | -0.09368 | 0.092812 | Other Negative Correlation |

| | | | |
|---|---|---|---|
| PRAG1 | -0.09371 | 0.092701 | Other Negative Correlation |
| SNAPC1 | -0.09375 | 0.092541 | Other Negative Correlation |
| ACTR6 | -0.09376 | 0.092523 | Other Negative Correlation |
| TRIM4 | -0.0938 | 0.092375 | Other Negative Correlation |
| CD99 | -0.09386 | 0.092171 | Other Negative Correlation |
| SERPINF2 | -0.09389 | 0.092051 | Other Negative Correlation |
| TEX2 | -0.09391 | 0.092012 | Other Negative Correlation |
| BAG3 | -0.09393 | 0.09192 | Other Negative Correlation |
| PSMD1 | -0.09395 | 0.091857 | Other Negative Correlation |
| ILK | -0.09398 | 0.091771 | Other Negative Correlation |
| EML3 | -0.094 | 0.091699 | Other Negative Correlation |
| NDUFB3 | -0.094 | 0.091674 | Other Negative Correlation |
| GRK2 | -0.09404 | 0.091545 | Other Negative Correlation |
| CRYAB | -0.09404 | 0.091536 | Other Negative Correlation |
| NUDT9 | -0.09406 | 0.091474 | Other Negative Correlation |
| GNPAT | -0.09419 | 0.091024 | Other Negative Correlation |
| AGL | -0.09422 | 0.090921 | Other Negative Correlation |
| MROH1 | -0.09423 | 0.090875 | Other Negative Correlation |
| ENG | -0.09436 | 0.090454 | Other Negative Correlation |
| HYAL2 | -0.09443 | 0.090209 | Other Negative Correlation |
| ZFAND6 | -0.0946 | 0.089608 | Other Negative Correlation |
| AGO3 | -0.09462 | 0.08956 | Other Negative Correlation |
| TRPV1 | -0.09465 | 0.089463 | Other Negative Correlation |
| MED29 | -0.09471 | 0.089257 | Other Negative Correlation |
| APBA3 | -0.09481 | 0.088924 | Other Negative Correlation |
| TMEM106B | -0.09483 | 0.088834 | Other Negative Correlation |
| MT-ND1 | -0.09496 | 0.088402 | Other Negative Correlation |
| IREB2 | -0.09498 | 0.08834 | Other Negative Correlation |
| TANGO2 | -0.09501 | 0.088239 | Other Negative Correlation |
| OSBPL9 | -0.09504 | 0.088135 | Other Negative Correlation |
| AADAT | -0.09505 | 0.088098 | Other Negative Correlation |
| CD74 | -0.09508 | 0.087997 | Other Negative Correlation |
| SKIL | -0.09509 | 0.087953 | Other Negative Correlation |
| ST6GALNAC1 | -0.09512 | 0.087852 | Other Negative Correlation |
| ARF3 | -0.09513 | 0.087821 | Other Negative Correlation |
| SIRT2 | -0.09527 | 0.08737 | Other Negative Correlation |
| TMEM169 | -0.09529 | 0.087311 | Other Negative Correlation |
| HEMK1 | -0.0954 | 0.086934 | Other Negative Correlation |
| NADK | -0.0954 | 0.086918 | Other Negative Correlation |
| PSD4 | -0.09542 | 0.086878 | Other Negative Correlation |
| MT-ND3 | -0.09555 | 0.086446 | Other Negative Correlation |
| MYO9B | -0.09583 | 0.085497 | Other Negative Correlation |
| MTX3 | -0.09588 | 0.085349 | Other Negative Correlation |
| SLC37A2 | -0.09604 | 0.084817 | Other Negative Correlation |
| ARAP3 | -0.09608 | 0.08471 | Other Negative Correlation |
| IKBKE | -0.09619 | 0.084328 | Other Negative Correlation |
| TRAPPC1 | -0.09628 | 0.084035 | Other Negative Correlation |
| ALDOC | -0.0963 | 0.083987 | Other Negative Correlation |

| | | | |
|---|---|---|---|
| HDAC1 | -0.09632 | 0.083932 | Other Negative Correlation |
| SPIDR | -0.09643 | 0.083579 | Other Negative Correlation |
| MACROD1 | -0.09643 | 0.083562 | Other Negative Correlation |
| TWSG1 | -0.09654 | 0.083215 | Other Negative Correlation |
| PLIN3 | -0.09673 | 0.082614 | Other Negative Correlation |
| FAM149B1 | -0.09674 | 0.082558 | Other Negative Correlation |
| RBM6 | -0.09676 | 0.0825 | Other Negative Correlation |
| TMEM9B | -0.09687 | 0.082143 | Other Negative Correlation |
| FAM169A | -0.09689 | 0.082105 | Other Negative Correlation |
| C2orf74 | -0.09693 | 0.081954 | Other Negative Correlation |
| TSPAN14 | -0.09701 | 0.081719 | Other Negative Correlation |
| WWP1 | -0.09704 | 0.081629 | Other Negative Correlation |
| IL31RA | -0.09707 | 0.081523 | Other Negative Correlation |
| NMNAT2 | -0.0972 | 0.081104 | Other Negative Correlation |
| PIN4 | -0.09721 | 0.081082 | Other Negative Correlation |
| UMAD1 | -0.09723 | 0.08102 | Other Negative Correlation |
| SPOP | -0.09729 | 0.080846 | Other Negative Correlation |
| JUND | -0.09729 | 0.080824 | Other Negative Correlation |
| NHSL2 | -0.09734 | 0.080679 | Other Negative Correlation |
| ETFB | -0.09744 | 0.080355 | Other Negative Correlation |
| COQ5 | -0.09745 | 0.080318 | Other Negative Correlation |
| GGPS1 | -0.09755 | 0.080032 | Other Negative Correlation |
| SYK | -0.09757 | 0.07997 | Other Negative Correlation |
| ZNF445 | -0.09757 | 0.079949 | Other Negative Correlation |
| MAMDC2 | -0.09762 | 0.079792 | Other Negative Correlation |
| KDM8 | -0.09788 | 0.078993 | Other Negative Correlation |
| MYO5A | -0.09796 | 0.078751 | Other Negative Correlation |
| IFNAR2 | -0.09802 | 0.078571 | Other Negative Correlation |
| AVIL | -0.09818 | 0.078067 | Other Negative Correlation |
| ATXN10 | -0.09831 | 0.077697 | Other Negative Correlation |
| TAF2 | -0.09836 | 0.077539 | Other Negative Correlation |
| TYSND1 | -0.09846 | 0.077237 | Other Negative Correlation |
| BAD | -0.09853 | 0.077006 | Other Negative Correlation |
| TET2 | -0.09854 | 0.076982 | Other Negative Correlation |
| GOLGB1 | -0.09856 | 0.07693 | Other Negative Correlation |
| TAB3 | -0.09857 | 0.076902 | Other Negative Correlation |
| KIFC3 | -0.09861 | 0.076794 | Other Negative Correlation |
| SFXN3 | -0.09865 | 0.076663 | Other Negative Correlation |
| KLHL42 | -0.09871 | 0.076482 | Other Negative Correlation |
| OXA1L | -0.09872 | 0.076462 | Other Negative Correlation |
| XXYLT1 | -0.09872 | 0.076457 | Other Negative Correlation |
| AKAP9 | -0.09881 | 0.076172 | Other Negative Correlation |
| LAMTOR4 | -0.09884 | 0.076084 | Other Negative Correlation |
| TMEM50A | -0.09908 | 0.075375 | Other Negative Correlation |
| UBXN6 | -0.09913 | 0.075222 | Other Negative Correlation |
| SPACA6 | -0.09914 | 0.075188 | Other Negative Correlation |
| SYT2 | -0.0992 | 0.075016 | Other Negative Correlation |
| ACVR1 | -0.09945 | 0.074303 | Other Negative Correlation |

| | | | |
|---|---|---|---|
| FIZ1 | -0.09952 | 0.074078 | Other Negative Correlation |
| KLHL31 | -0.09957 | 0.073946 | Other Negative Correlation |
| RAB11FIP2 | -0.09964 | 0.073729 | Other Negative Correlation |
| XPA | -0.09964 | 0.073729 | Other Negative Correlation |
| MORC3 | -0.0997 | 0.073556 | Other Negative Correlation |
| ZNF846 | -0.0997 | 0.073554 | Other Negative Correlation |
| DHX40 | -0.09981 | 0.073234 | Other Negative Correlation |
| KLHL17 | -0.09985 | 0.073139 | Other Negative Correlation |
| C3orf52 | -0.09994 | 0.072863 | Other Negative Correlation |
| TAF8 | -0.09995 | 0.072848 | Other Negative Correlation |
| TDP2 | -0.1 | 0.072701 | Other Negative Correlation |
| TBC1D9B | -0.10001 | 0.072651 | Other Negative Correlation |
| COL18A1 | -0.10002 | 0.072648 | Other Negative Correlation |
| ALPK1 | -0.10008 | 0.07246 | Other Negative Correlation |
| CAPZA2 | -0.10012 | 0.07235 | Other Negative Correlation |
| PNCK | -0.10015 | 0.072267 | Other Negative Correlation |
| STAT2 | -0.1002 | 0.072115 | Other Negative Correlation |
| TMEM230 | -0.10038 | 0.071616 | Other Negative Correlation |
| SNRPB2 | -0.10038 | 0.07161 | Other Negative Correlation |
| SFXN1 | -0.10038 | 0.071606 | Other Negative Correlation |
| MYO6 | -0.10039 | 0.071573 | Other Negative Correlation |
| PARP16 | -0.10042 | 0.071493 | Other Negative Correlation |
| DECR2 | -0.10045 | 0.071419 | Other Negative Correlation |
| MAP7 | -0.1005 | 0.071277 | Other Negative Correlation |
| POLI | -0.10067 | 0.070789 | Other Negative Correlation |
| ATXN7L1 | -0.1007 | 0.070716 | Other Negative Correlation |
| PPP2R2A | -0.10073 | 0.070615 | Other Negative Correlation |
| IRAK2 | -0.10075 | 0.070571 | Other Negative Correlation |
| NDUFS8 | -0.10075 | 0.07055 | Other Negative Correlation |
| ATE1 | -0.10075 | 0.070549 | Other Negative Correlation |
| TMEM258 | -0.10076 | 0.070522 | Other Negative Correlation |
| EIPR1 | -0.10086 | 0.070261 | Other Negative Correlation |
| OPN3 | -0.10088 | 0.070185 | Other Negative Correlation |
| USP14 | -0.1009 | 0.070135 | Other Negative Correlation |
| PHACTR2 | -0.10103 | 0.069767 | Other Negative Correlation |
| CATSPER1 | -0.10106 | 0.069708 | Other Negative Correlation |
| FANK1 | -0.10111 | 0.069547 | Other Negative Correlation |
| TAP1 | -0.10112 | 0.069519 | Other Negative Correlation |
| TMED2 | -0.10114 | 0.069483 | Other Negative Correlation |
| TSACC | -0.10116 | 0.069419 | Other Negative Correlation |
| GSTA4 | -0.10122 | 0.069265 | Other Negative Correlation |
| GRK4 | -0.10127 | 0.06911 | Other Negative Correlation |
| HSPG2 | -0.10143 | 0.068683 | Other Negative Correlation |
| BLMH | -0.10145 | 0.06862 | Other Negative Correlation |
| SPECC1 | -0.10147 | 0.068569 | Other Negative Correlation |
| IGSF3 | -0.10155 | 0.06836 | Other Negative Correlation |
| ZBTB20 | -0.10162 | 0.068151 | Other Negative Correlation |
| ANKRD2 | -0.10173 | 0.067847 | Other Negative Correlation |

| | | | |
|---|---|---|---|
| UGT1A6 | -0.1018 | 0.067662 | Other Negative Correlation |
| GHDC | -0.10182 | 0.067619 | Other Negative Correlation |
| MTA3 | -0.10186 | 0.067507 | Other Negative Correlation |
| SPA17 | -0.10193 | 0.067304 | Other Negative Correlation |
| FBXO4 | -0.10211 | 0.066832 | Other Negative Correlation |
| MRPS7 | -0.10216 | 0.066698 | Other Negative Correlation |
| CEP68 | -0.1022 | 0.066599 | Other Negative Correlation |
| EVL | -0.10221 | 0.066555 | Other Negative Correlation |
| C9orf78 | -0.10221 | 0.066549 | Other Negative Correlation |
| PARP8 | -0.10224 | 0.066486 | Other Negative Correlation |
| ENSG00000268655 | -0.10224 | 0.066477 | Other Negative Correlation |
| CD81 | -0.10228 | 0.066363 | Other Negative Correlation |
| THBS3 | -0.10238 | 0.066114 | Other Negative Correlation |
| OTOP2 | -0.10245 | 0.065912 | Other Negative Correlation |
| TRIM35 | -0.10258 | 0.065575 | Other Negative Correlation |
| KLHL20 | -0.10267 | 0.065343 | Other Negative Correlation |
| MSTN | -0.10274 | 0.065153 | Other Negative Correlation |
| PHYHD1 | -0.10283 | 0.064917 | Other Negative Correlation |
| PCYOX1 | -0.10284 | 0.064885 | Other Negative Correlation |
| MSI2 | -0.10286 | 0.06484 | Other Negative Correlation |
| PALMD | -0.10288 | 0.064791 | Other Negative Correlation |
| EVA1A | -0.10298 | 0.064516 | Other Negative Correlation |
| NIPAL4 | -0.10302 | 0.064424 | Other Negative Correlation |
| ZBTB25 | -0.10304 | 0.064375 | Other Negative Correlation |
| ZNF106 | -0.10308 | 0.064255 | Other Negative Correlation |
| TNRC6B | -0.10325 | 0.063839 | Other Negative Correlation |
| GDF9 | -0.10325 | 0.063821 | Other Negative Correlation |
| NEK1 | -0.10329 | 0.063712 | Other Negative Correlation |
| CCDC146 | -0.10331 | 0.063682 | Other Negative Correlation |
| APMAP | -0.10339 | 0.063466 | Other Negative Correlation |
| MRPL10 | -0.10341 | 0.063406 | Other Negative Correlation |
| AGA | -0.10353 | 0.063109 | Other Negative Correlation |
| EPGN | -0.10357 | 0.063012 | Other Negative Correlation |
| FBXL2 | -0.1036 | 0.062938 | Other Negative Correlation |
| UBE2Z | -0.1036 | 0.062926 | Other Negative Correlation |
| VPS26B | -0.10367 | 0.062758 | Other Negative Correlation |
| PCDHGB7 | -0.10368 | 0.062731 | Other Negative Correlation |
| MRPS36 | -0.1037 | 0.062665 | Other Negative Correlation |
| NELFE | -0.1038 | 0.06243 | Other Negative Correlation |
| TSGA10IP | -0.10387 | 0.062241 | Other Negative Correlation |
| DOCK7 | -0.10388 | 0.062205 | Other Negative Correlation |
| B4GAT1 | -0.10392 | 0.062103 | Other Negative Correlation |
| FBXO17 | -0.10396 | 0.062018 | Other Negative Correlation |
| FXYD5 | -0.10399 | 0.061941 | Other Negative Correlation |
| TRANK1 | -0.10404 | 0.061814 | Other Negative Correlation |
| WASHC3 | -0.10406 | 0.061754 | Other Negative Correlation |
| ZNF264 | -0.10431 | 0.061124 | Other Negative Correlation |
| CRELD2 | -0.10434 | 0.061051 | Other Negative Correlation |

| | | | |
|---|---|---|---|
| BRF2 | -0.10435 | 0.061046 | Other Negative Correlation |
| MIB1 | -0.10436 | 0.061021 | Other Negative Correlation |
| OLFML2A | -0.10437 | 0.060974 | Other Negative Correlation |
| GCNA | -0.10439 | 0.06094 | Other Negative Correlation |
| USP53 | -0.10442 | 0.060865 | Other Negative Correlation |
| OARD1 | -0.10444 | 0.0608 | Other Negative Correlation |
| ATG10 | -0.10446 | 0.060755 | Other Negative Correlation |
| BPNT1 | -0.10448 | 0.060706 | Other Negative Correlation |
| IFNAR1 | -0.10452 | 0.060602 | Other Negative Correlation |
| CNGB1 | -0.10468 | 0.060223 | Other Negative Correlation |
| HSD17B10 | -0.10475 | 0.060036 | Other Negative Correlation |
| LYST | -0.10477 | 0.059991 | Other Negative Correlation |
| MUCL3 | -0.10479 | 0.059951 | Other Negative Correlation |
| PRICKLE2 | -0.10498 | 0.059486 | Other Negative Correlation |
| TBC1D2B | -0.10507 | 0.059257 | Other Negative Correlation |
| DNPEP | -0.10511 | 0.059158 | Other Negative Correlation |
| DENND1C | -0.10513 | 0.059107 | Other Negative Correlation |
| ZNF235 | -0.10523 | 0.058873 | Other Negative Correlation |
| KLHL12 | -0.10539 | 0.058485 | Other Negative Correlation |
| ARHGEF7 | -0.10542 | 0.058405 | Other Negative Correlation |
| ANPEP | -0.10549 | 0.058257 | Other Negative Correlation |
| PRXL2C | -0.10557 | 0.058066 | Other Negative Correlation |
| SSR2 | -0.10564 | 0.05788 | Other Negative Correlation |
| NR1H3 | -0.10584 | 0.057415 | Other Negative Correlation |
| FAM227A | -0.10584 | 0.057412 | Other Negative Correlation |
| CARD8 | -0.10585 | 0.057382 | Other Negative Correlation |
| KLHL8 | -0.1059 | 0.057279 | Other Negative Correlation |
| ZNF548 | -0.1059 | 0.057265 | Other Negative Correlation |
| GCC2 | -0.10597 | 0.057118 | Other Negative Correlation |
| SLC39A13 | -0.10598 | 0.057086 | Other Negative Correlation |
| SEC61B | -0.10599 | 0.057067 | Other Negative Correlation |
| NIT1 | -0.10604 | 0.056946 | Other Negative Correlation |
| PIAS3 | -0.10616 | 0.056664 | Other Negative Correlation |
| IDS | -0.10618 | 0.056624 | Other Negative Correlation |
| CC2D1B | -0.10622 | 0.056509 | Other Negative Correlation |
| SERPINA1 | -0.10623 | 0.056503 | Other Negative Correlation |
| STK11IP | -0.10659 | 0.05565 | Other Negative Correlation |
| KCNAB2 | -0.10661 | 0.055622 | Other Negative Correlation |
| NDUFB5 | -0.10677 | 0.055254 | Other Negative Correlation |
| MEN1 | -0.10686 | 0.055042 | Other Negative Correlation |
| USP49 | -0.10696 | 0.054805 | Other Negative Correlation |
| RHBDD1 | -0.10699 | 0.054736 | Other Negative Correlation |
| GNG13 | -0.10702 | 0.054683 | Other Negative Correlation |
| MRC2 | -0.10703 | 0.054646 | Other Negative Correlation |
| TRAF3 | -0.10703 | 0.054642 | Other Negative Correlation |
| RUFY2 | -0.10707 | 0.05456 | Other Negative Correlation |
| LGR4 | -0.1071 | 0.054488 | Other Negative Correlation |
| PIDD1 | -0.10723 | 0.05419 | Other Negative Correlation |

| | | | |
|---|---|---|---|
| MALSU1 | -0.10727 | 0.054119 | Other Negative Correlation |
| AZI2 | -0.10738 | 0.053854 | Other Negative Correlation |
| CNOT2 | -0.10741 | 0.053789 | Other Negative Correlation |
| POGZ | -0.10743 | 0.053759 | Other Negative Correlation |
| SORBS2 | -0.10744 | 0.053734 | Other Negative Correlation |
| WSB2 | -0.10744 | 0.053716 | Other Negative Correlation |
| C19orf53 | -0.10747 | 0.053649 | Other Negative Correlation |
| NCKAP5L | -0.10755 | 0.053472 | Other Negative Correlation |
| RBAK-RBAKDN | -0.10765 | 0.053255 | Other Negative Correlation |
| EFNA1 | -0.10772 | 0.053101 | Other Negative Correlation |
| MAPK14 | -0.10782 | 0.052882 | Other Negative Correlation |
| SNX22 | -0.10786 | 0.052792 | Other Negative Correlation |
| LEPR | -0.10799 | 0.0525 | Other Negative Correlation |
| TP53BP1 | -0.10806 | 0.052351 | Other Negative Correlation |
| UBE2R2 | -0.10811 | 0.05224 | Other Negative Correlation |
| POLR2A | -0.10815 | 0.052164 | Other Negative Correlation |
| PIGT | -0.10815 | 0.052158 | Other Negative Correlation |
| CEP57L1 | -0.10836 | 0.051707 | Other Negative Correlation |
| LRRC37A | -0.10855 | 0.051282 | Other Negative Correlation |
| RNF135 | -0.10856 | 0.051261 | Other Negative Correlation |
| AFAP1 | -0.10867 | 0.051029 | Other Negative Correlation |
| ECE1 | -0.10881 | 0.050727 | Other Negative Correlation |
| POLR2E | -0.10882 | 0.050716 | Other Negative Correlation |
| MT-ATP6 | -0.10891 | 0.05051 | Other Negative Correlation |
| ANKRD17 | -0.10892 | 0.050483 | Other Negative Correlation |
| CMPK1 | -0.10892 | 0.050483 | Other Negative Correlation |
| LGALS4 | -0.10896 | 0.050416 | Other Negative Correlation |
| RABGGTA | -0.10897 | 0.050377 | Other Negative Correlation |
| METTL14 | -0.10901 | 0.050299 | Other Negative Correlation |
| STARD5 | -0.10902 | 0.050288 | Other Negative Correlation |
| TPST2 | -0.10903 | 0.050254 | Other Negative Correlation |
| PNPLA3 | -0.10908 | 0.050158 | Other Negative Correlation |
| F11R | -0.10914 | 0.050018 | Other Negative Correlation |
| MED4 | -0.10917 | 0.049963 | Other Negative Correlation |
| MAP2K2 | -0.1092 | 0.049902 | Other Negative Correlation |
| TRPT1 | -0.10925 | 0.049792 | Other Negative Correlation |
| GDE1 | -0.10931 | 0.049663 | Other Negative Correlation |
| ZBTB38 | -0.10943 | 0.049425 | Other Negative Correlation |
| CCDC32 | -0.10948 | 0.049322 | Other Negative Correlation |
| PSMD7 | -0.10954 | 0.049193 | Other Negative Correlation |
| DIP2B | -0.10965 | 0.04895 | Other Negative Correlation |
| TRAPPC6A | -0.10966 | 0.048941 | Other Negative Correlation |
| GORASP2 | -0.10966 | 0.04894 | Other Negative Correlation |
| C5orf52 | -0.10984 | 0.048565 | Other Negative Correlation |
| IGDCC4 | -0.10987 | 0.048508 | Other Negative Correlation |
| P3H4 | -0.10996 | 0.048327 | Other Negative Correlation |
| LYRM9 | -0.11018 | 0.047879 | Other Negative Correlation |
| NEK3 | -0.11022 | 0.047786 | Other Negative Correlation |

| | | | |
|---|---|---|---|
| AKNAD1 | -0.11023 | 0.047759 | Other Negative Correlation |
| MANBA | -0.11038 | 0.047459 | Other Negative Correlation |
| UBE3B | -0.11045 | 0.047316 | Other Negative Correlation |
| ZSCAN21 | -0.11048 | 0.047269 | Other Negative Correlation |
| RAB3GAP2 | -0.11049 | 0.047239 | Other Negative Correlation |
| DNAJC4 | -0.11059 | 0.047049 | Other Negative Correlation |
| SUGT1 | -0.1106 | 0.047025 | Other Negative Correlation |
| NCOR2 | -0.1107 | 0.046812 | Other Negative Correlation |
| CPA2 | -0.11075 | 0.04673 | Other Negative Correlation |
| STARD6 | -0.11078 | 0.04666 | Other Negative Correlation |
| LRRC4 | -0.11079 | 0.046634 | Other Negative Correlation |
| MAP2K5 | -0.11091 | 0.046394 | Other Negative Correlation |
| GRINA | -0.11093 | 0.04637 | Other Negative Correlation |
| SLC66A2 | -0.11096 | 0.046298 | Other Negative Correlation |
| BMP1 | -0.11097 | 0.04628 | Other Negative Correlation |
| FUBP3 | -0.11099 | 0.046242 | Other Negative Correlation |
| EXOC7 | -0.111 | 0.046215 | Other Negative Correlation |
| TUBGCP3 | -0.11104 | 0.046145 | Other Negative Correlation |
| TTC17 | -0.11105 | 0.046118 | Other Negative Correlation |
| HSBP1 | -0.11112 | 0.045988 | Other Negative Correlation |
| ATG7 | -0.11115 | 0.045927 | Other Negative Correlation |
| NUDT16L1 | -0.11118 | 0.045875 | Other Negative Correlation |
| SPTY2D1OS | -0.11121 | 0.045807 | Other Negative Correlation |
| NEIL2 | -0.11125 | 0.045735 | Other Negative Correlation |
| SPACA9 | -0.11127 | 0.045689 | Other Negative Correlation |
| CEP104 | -0.11128 | 0.045681 | Other Negative Correlation |
| GPR39 | -0.11133 | 0.04557 | Other Negative Correlation |
| PAFAH1B1 | -0.11172 | 0.044821 | Other Negative Correlation |
| ZNF658 | -0.11177 | 0.044725 | Other Negative Correlation |
| STAT4 | -0.11179 | 0.044684 | Other Negative Correlation |
| SLC26A4 | -0.11185 | 0.044578 | Other Negative Correlation |
| ARHGEF9 | -0.11191 | 0.044455 | Other Negative Correlation |
| INAVA | -0.11209 | 0.044109 | Other Negative Correlation |
| TTYH3 | -0.11211 | 0.044072 | Other Negative Correlation |
| UBE2F-SCLY | -0.11231 | 0.0437 | Other Negative Correlation |
| SS18L1 | -0.11253 | 0.043284 | Other Negative Correlation |
| TMCO6 | -0.11263 | 0.043094 | Other Negative Correlation |
| UQCRQ | -0.1127 | 0.042971 | Other Negative Correlation |
| TNFSF10 | -0.11282 | 0.042744 | Other Negative Correlation |
| NUDT22 | -0.1129 | 0.042592 | Other Negative Correlation |
| SEC13 | -0.11297 | 0.042461 | Other Negative Correlation |
| SNX29 | -0.11299 | 0.042433 | Other Negative Correlation |
| ZNF619 | -0.11311 | 0.0422 | Other Negative Correlation |
| IGF2R | -0.11313 | 0.042168 | Other Negative Correlation |
| DDHD2 | -0.11333 | 0.041809 | Other Negative Correlation |
| NUB1 | -0.11333 | 0.041802 | Other Negative Correlation |
| PTPN22 | -0.11341 | 0.041653 | Other Negative Correlation |
| QSOX1 | -0.11348 | 0.041534 | Other Negative Correlation |

| | | | |
|---|---|---|---|
| IVD | -0.11356 | 0.041387 | Other Negative Correlation |
| SPPL2A | -0.11362 | 0.041274 | Other Negative Correlation |
| CFAP44 | -0.11363 | 0.041259 | Other Negative Correlation |
| VPS51 | -0.11388 | 0.040816 | Other Negative Correlation |
| LRBA | -0.11389 | 0.040798 | Other Negative Correlation |
| MRNIP | -0.11391 | 0.04077 | Other Negative Correlation |
| CARD16 | -0.11395 | 0.040685 | Other Negative Correlation |
| TRAPPC3 | -0.11402 | 0.04057 | Other Negative Correlation |
| N4BP1 | -0.11412 | 0.04039 | Other Negative Correlation |
| PIGP | -0.11413 | 0.040372 | Other Negative Correlation |
| BBIP1 | -0.11413 | 0.040366 | Other Negative Correlation |
| SEC23A | -0.11415 | 0.040344 | Other Negative Correlation |
| BDP1 | -0.11424 | 0.040187 | Other Negative Correlation |
| FAM3A | -0.11428 | 0.040102 | Other Negative Correlation |
| ZNF280D | -0.11431 | 0.040049 | Other Negative Correlation |
| NACC1 | -0.11441 | 0.039888 | Other Negative Correlation |
| LSP1 | -0.11442 | 0.039865 | Other Negative Correlation |
| SH3D21 | -0.11445 | 0.039808 | Other Negative Correlation |
| STK39 | -0.11447 | 0.039784 | Other Negative Correlation |
| ROCK1 | -0.11454 | 0.039653 | Other Negative Correlation |
| EIF2B4 | -0.11455 | 0.039641 | Other Negative Correlation |
| PDHB | -0.11459 | 0.039563 | Other Negative Correlation |
| PARP4 | -0.11462 | 0.039523 | Other Negative Correlation |
| CEP126 | -0.11469 | 0.039386 | Other Negative Correlation |
| CERCAM | -0.11471 | 0.039353 | Other Negative Correlation |
| PATZ1 | -0.11473 | 0.039327 | Other Negative Correlation |
| CD82 | -0.11478 | 0.03924 | Other Negative Correlation |
| ARL10 | -0.11488 | 0.039066 | Other Negative Correlation |
| GDPD1 | -0.11491 | 0.039022 | Other Negative Correlation |
| SDHAF4 | -0.11492 | 0.039 | Other Negative Correlation |
| MAP4K5 | -0.11494 | 0.038962 | Other Negative Correlation |
| ST3GAL5 | -0.11525 | 0.038436 | Other Negative Correlation |
| DUS4L | -0.11542 | 0.038146 | Other Negative Correlation |
| HSF1 | -0.11543 | 0.038132 | Other Negative Correlation |
| ZNG1F | -0.11552 | 0.037979 | Other Negative Correlation |
| ZNF217 | -0.11562 | 0.037811 | Other Negative Correlation |
| JARID2 | -0.11566 | 0.037741 | Other Negative Correlation |
| NETO1 | -0.11568 | 0.037718 | Other Negative Correlation |
| SNCG | -0.11574 | 0.037621 | Other Negative Correlation |
| RILPL2 | -0.11585 | 0.037426 | Other Negative Correlation |
| TSTD3 | -0.11594 | 0.037276 | Other Negative Correlation |
| ARHGAP1 | -0.11595 | 0.037266 | Other Negative Correlation |
| AEBP2 | -0.11602 | 0.037149 | Other Negative Correlation |
| IFI6 | -0.11614 | 0.036955 | Other Negative Correlation |
| TRERF1 | -0.1163 | 0.036689 | Other Negative Correlation |
| ZG16B | -0.11634 | 0.036629 | Other Negative Correlation |
| KDM5D | -0.1164 | 0.036526 | Other Negative Correlation |
| IFT140 | -0.11643 | 0.03648 | Other Negative Correlation |

| | | | |
|---|---|---|---|
| POLB | -0.11656 | 0.036273 | Other Negative Correlation |
| FAM193A | -0.11658 | 0.036241 | Other Negative Correlation |
| LRPAP1 | -0.11659 | 0.036219 | Other Negative Correlation |
| SEL1L3 | -0.11661 | 0.036191 | Other Negative Correlation |
| DHRS12 | -0.11681 | 0.035865 | Other Negative Correlation |
| AHCYL1 | -0.11695 | 0.035644 | Other Negative Correlation |
| ECD | -0.11699 | 0.035587 | Other Negative Correlation |
| LAPTM5 | -0.117 | 0.035574 | Other Negative Correlation |
| COPS9 | -0.11702 | 0.035531 | Other Negative Correlation |
| R3HDM1 | -0.11703 | 0.035523 | Other Negative Correlation |
| ODR4 | -0.11711 | 0.035395 | Other Negative Correlation |
| FAM114A1 | -0.11714 | 0.035341 | Other Negative Correlation |
| CRYZ | -0.1173 | 0.035097 | Other Negative Correlation |
| ADAP1 | -0.1174 | 0.034937 | Other Negative Correlation |
| CORO1B | -0.11741 | 0.034924 | Other Negative Correlation |
| NDUFC1 | -0.11745 | 0.034867 | Other Negative Correlation |
| D2HGDH | -0.11749 | 0.034806 | Other Negative Correlation |
| H2AC6 | -0.11749 | 0.034804 | Other Negative Correlation |
| CLIP4 | -0.11759 | 0.034648 | Other Negative Correlation |
| MEAF6 | -0.11762 | 0.034591 | Other Negative Correlation |
| MAVS | -0.11765 | 0.034548 | Other Negative Correlation |
| USP22 | -0.11783 | 0.034281 | Other Negative Correlation |
| CLSTN1 | -0.11806 | 0.033922 | Other Negative Correlation |
| ZNF737 | -0.11816 | 0.033775 | Other Negative Correlation |
| ZNF283 | -0.11836 | 0.033472 | Other Negative Correlation |
| RNF32 | -0.11842 | 0.033379 | Other Negative Correlation |
| UBE2V2 | -0.11849 | 0.033267 | Other Negative Correlation |
| TCAF1 | -0.1185 | 0.033257 | Other Negative Correlation |
| CROT | -0.11853 | 0.033216 | Other Negative Correlation |
| HOXA4 | -0.11856 | 0.033174 | Other Negative Correlation |
| MTRES1 | -0.11856 | 0.033164 | Other Negative Correlation |
| GABRG2 | -0.11858 | 0.033136 | Other Negative Correlation |
| PRDX5 | -0.1186 | 0.033106 | Other Negative Correlation |
| RRN3 | -0.1187 | 0.032966 | Other Negative Correlation |
| TMEM241 | -0.11882 | 0.032788 | Other Negative Correlation |
| SDCCAG8 | -0.11888 | 0.032687 | Other Negative Correlation |
| PROM2 | -0.11896 | 0.03258 | Other Negative Correlation |
| BABAM1 | -0.11903 | 0.032473 | Other Negative Correlation |
| GLRX | -0.11907 | 0.032416 | Other Negative Correlation |
| ZBTB22 | -0.11914 | 0.032313 | Other Negative Correlation |
| RWDD3 | -0.11916 | 0.032279 | Other Negative Correlation |
| OGFRL1 | -0.11947 | 0.031839 | Other Negative Correlation |
| PPCS | -0.11947 | 0.03183 | Other Negative Correlation |
| ENSG00000280571 | -0.11962 | 0.031615 | Other Negative Correlation |
| MTERF2 | -0.11962 | 0.031612 | Other Negative Correlation |
| TCIM | -0.11982 | 0.031325 | Other Negative Correlation |
| LIF | -0.11984 | 0.031307 | Other Negative Correlation |
| NPIPB11 | -0.12013 | 0.030898 | Other Negative Correlation |

| | | | |
|---|---|---|---|
| SRSF4 | -0.12028 | 0.030676 | Other Negative Correlation |
| SRPX | -0.12035 | 0.030588 | Other Negative Correlation |
| CDH1 | -0.12036 | 0.030564 | Other Negative Correlation |
| COMMD2 | -0.12039 | 0.030523 | Other Negative Correlation |
| MAPK13 | -0.1205 | 0.030375 | Other Negative Correlation |
| PSMF1 | -0.12061 | 0.03022 | Other Negative Correlation |
| UBL5 | -0.12065 | 0.030172 | Other Negative Correlation |
| AKAP6 | -0.12067 | 0.030144 | Other Negative Correlation |
| ARL2BP | -0.12078 | 0.02999 | Other Negative Correlation |
| IRAK4 | -0.1208 | 0.029958 | Other Negative Correlation |
| KIDINS220 | -0.12083 | 0.029916 | Other Negative Correlation |
| PRELID3B | -0.12091 | 0.029806 | Other Negative Correlation |
| PJA2 | -0.12103 | 0.029654 | Other Negative Correlation |
| MRPS14 | -0.12111 | 0.029536 | Other Negative Correlation |
| RAET1E | -0.12127 | 0.029324 | Other Negative Correlation |
| TMEM143 | -0.12129 | 0.029296 | Other Negative Correlation |
| MCCC2 | -0.12133 | 0.029238 | Other Negative Correlation |
| DYNC1H1 | -0.12134 | 0.029236 | Other Negative Correlation |
| EPC2 | -0.12142 | 0.029127 | Other Negative Correlation |
| PITRM1 | -0.12148 | 0.029042 | Other Negative Correlation |
| ANKS4B | -0.1215 | 0.029018 | Other Negative Correlation |
| PTPRO | -0.12157 | 0.028925 | Other Negative Correlation |
| IK | -0.12159 | 0.028898 | Other Negative Correlation |
| AMELY | -0.12161 | 0.028874 | Other Negative Correlation |
| SLC4A11 | -0.12163 | 0.028846 | Other Negative Correlation |
| CCDC24 | -0.12173 | 0.028717 | Other Negative Correlation |
| PDPN | -0.12178 | 0.028646 | Other Negative Correlation |
| ZNF490 | -0.12179 | 0.028636 | Other Negative Correlation |
| ARIH1 | -0.12181 | 0.028604 | Other Negative Correlation |
| SDHB | -0.12182 | 0.028592 | Other Negative Correlation |
| FRK | -0.12185 | 0.02856 | Other Negative Correlation |
| PBRM1 | -0.12198 | 0.02839 | Other Negative Correlation |
| P2RX6 | -0.12213 | 0.028184 | Other Negative Correlation |
| MKNK1 | -0.12225 | 0.028031 | Other Negative Correlation |
| PON2 | -0.1225 | 0.027711 | Other Negative Correlation |
| TMEM63B | -0.12264 | 0.027528 | Other Negative Correlation |
| CHMP1B | -0.12269 | 0.027462 | Other Negative Correlation |
| CYP21A2 | -0.12273 | 0.027419 | Other Negative Correlation |
| PSG11 | -0.12273 | 0.027418 | Other Negative Correlation |
| GOLGA8R | -0.12275 | 0.027396 | Other Negative Correlation |
| ZNF791 | -0.12275 | 0.027392 | Other Negative Correlation |
| SGIP1 | -0.1229 | 0.027207 | Other Negative Correlation |
| SND1 | -0.12295 | 0.027144 | Other Negative Correlation |
| BBS12 | -0.12301 | 0.027066 | Other Negative Correlation |
| CRPPA | -0.12302 | 0.027056 | Other Negative Correlation |
| DNAJC16 | -0.12302 | 0.027054 | Other Negative Correlation |
| LY96 | -0.12303 | 0.027045 | Other Negative Correlation |
| BBX | -0.12304 | 0.027026 | Other Negative Correlation |

| | | | |
|---|---|---|---|
| FAM131A | -0.12316 | 0.026876 | Other Negative Correlation |
| CSTB | -0.12318 | 0.026852 | Other Negative Correlation |
| ERAP1 | -0.12334 | 0.02665 | Other Negative Correlation |
| XPR1 | -0.12343 | 0.026537 | Other Negative Correlation |
| TRRAP | -0.12347 | 0.026492 | Other Negative Correlation |
| DHRS7 | -0.12351 | 0.026446 | Other Negative Correlation |
| H3-3A | -0.12351 | 0.026445 | Other Negative Correlation |
| C1RL | -0.12355 | 0.026393 | Other Negative Correlation |
| SUOX | -0.12367 | 0.026244 | Other Negative Correlation |
| CLPP | -0.12369 | 0.026217 | Other Negative Correlation |
| SLC2A6 | -0.12371 | 0.026198 | Other Negative Correlation |
| ARMC9 | -0.12378 | 0.026108 | Other Negative Correlation |
| HDLBP | -0.12389 | 0.025983 | Other Negative Correlation |
| ARHGAP23 | -0.12413 | 0.02569 | Other Negative Correlation |
| SH2D2A | -0.12418 | 0.025628 | Other Negative Correlation |
| ELP1 | -0.12423 | 0.025567 | Other Negative Correlation |
| CKAP4 | -0.12433 | 0.025451 | Other Negative Correlation |
| NIN | -0.12433 | 0.025445 | Other Negative Correlation |
| CARD6 | -0.12437 | 0.025403 | Other Negative Correlation |
| VPS13A | -0.12441 | 0.025359 | Other Negative Correlation |
| NEK9 | -0.12446 | 0.025295 | Other Negative Correlation |
| RDH11 | -0.12446 | 0.025294 | Other Negative Correlation |
| RFX1 | -0.12448 | 0.025274 | Other Negative Correlation |
| SCARB2 | -0.1245 | 0.025243 | Other Negative Correlation |
| DNASE1L1 | -0.12459 | 0.025142 | Other Negative Correlation |
| RHBDF1 | -0.12459 | 0.025138 | Other Negative Correlation |
| SYF2 | -0.12467 | 0.025045 | Other Negative Correlation |
| CD109 | -0.12478 | 0.024922 | Other Negative Correlation |
| ATP1A1 | -0.1248 | 0.024898 | Other Negative Correlation |
| HOMER1 | -0.12485 | 0.024837 | Other Negative Correlation |
| ZSWIM1 | -0.12486 | 0.024825 | Other Negative Correlation |
| METTL25 | -0.12492 | 0.024751 | Other Negative Correlation |
| RSF1 | -0.12495 | 0.024717 | Other Negative Correlation |
| JAK3 | -0.125 | 0.024667 | Other Negative Correlation |
| IDE | -0.12506 | 0.024594 | Other Negative Correlation |
| IDH3B | -0.12509 | 0.02456 | Other Negative Correlation |
| MTNR1A | -0.12538 | 0.02423 | Other Negative Correlation |
| SNX7 | -0.1254 | 0.024205 | Other Negative Correlation |
| VPS13C | -0.1254 | 0.024199 | Other Negative Correlation |
| ARHGEF4 | -0.12543 | 0.024169 | Other Negative Correlation |
| PTPRM | -0.12556 | 0.024019 | Other Negative Correlation |
| VPS25 | -0.12562 | 0.023951 | Other Negative Correlation |
| TGM2 | -0.12567 | 0.023897 | Other Negative Correlation |
| NPIPA9 | -0.12568 | 0.023882 | Other Negative Correlation |
| BRD3 | -0.12571 | 0.023855 | Other Negative Correlation |
| ABHD16B | -0.12572 | 0.023846 | Other Negative Correlation |
| R3HDM4 | -0.1259 | 0.023641 | Other Negative Correlation |
| MKRN2OS | -0.12592 | 0.023615 | Other Negative Correlation |

| | | | |
|---|---|---|---|
| UBXN1 | -0.12593 | 0.02361 | Other Negative Correlation |
| MRPS15 | -0.1261 | 0.023415 | Other Negative Correlation |
| VPS4B | -0.12613 | 0.023388 | Other Negative Correlation |
| CPS1 | -0.12614 | 0.023373 | Other Negative Correlation |
| FXYD3 | -0.12635 | 0.023146 | Other Negative Correlation |
| RAB11FIP3 | -0.1265 | 0.022972 | Other Negative Correlation |
| TUT7 | -0.12654 | 0.022938 | Other Negative Correlation |
| CFAP36 | -0.12655 | 0.022922 | Other Negative Correlation |
| RUFY3 | -0.12664 | 0.022829 | Other Negative Correlation |
| NFIX | -0.12669 | 0.022767 | Other Negative Correlation |
| UPK3B | -0.1267 | 0.02276 | Other Negative Correlation |
| POLM | -0.12673 | 0.02273 | Other Negative Correlation |
| ICAM4 | -0.12682 | 0.022634 | Other Negative Correlation |
| PHACTR1 | -0.12688 | 0.022562 | Other Negative Correlation |
| GYS1 | -0.12695 | 0.022496 | Other Negative Correlation |
| C10orf95 | -0.12696 | 0.022482 | Other Negative Correlation |
| ACOT11 | -0.12699 | 0.022444 | Other Negative Correlation |
| CA12 | -0.127 | 0.022435 | Other Negative Correlation |
| TP73 | -0.12704 | 0.022392 | Other Negative Correlation |
| DNTTIP1 | -0.12708 | 0.022358 | Other Negative Correlation |
| ACACA | -0.12745 | 0.02196 | Other Negative Correlation |
| VWA8 | -0.12752 | 0.021892 | Other Negative Correlation |
| SMDT1 | -0.1276 | 0.021806 | Other Negative Correlation |
| CXXC4 | -0.1276 | 0.021802 | Other Negative Correlation |
| C1D | -0.12787 | 0.02152 | Other Negative Correlation |
| HGSNAT | -0.12793 | 0.021467 | Other Negative Correlation |
| ARHGAP29 | -0.12806 | 0.02133 | Other Negative Correlation |
| ARMCX2 | -0.12819 | 0.021198 | Other Negative Correlation |
| GJB4 | -0.12824 | 0.021148 | Other Negative Correlation |
| LEPROT | -0.12825 | 0.021139 | Other Negative Correlation |
| PFKFB4 | -0.12825 | 0.021134 | Other Negative Correlation |
| PPP2R1A | -0.1283 | 0.021084 | Other Negative Correlation |
| CHMP4B | -0.12833 | 0.021055 | Other Negative Correlation |
| LIPA | -0.12837 | 0.021019 | Other Negative Correlation |
| NDUFB10 | -0.12847 | 0.020912 | Other Negative Correlation |
| GTF2H4 | -0.12855 | 0.020838 | Other Negative Correlation |
| PPP1R35 | -0.12864 | 0.020747 | Other Negative Correlation |
| MR1 | -0.12866 | 0.020721 | Other Negative Correlation |
| WFDC5 | -0.12877 | 0.020617 | Other Negative Correlation |
| STYXL1 | -0.12882 | 0.020569 | Other Negative Correlation |
| RICTOR | -0.12883 | 0.020559 | Other Negative Correlation |
| SPIN4 | -0.12883 | 0.020556 | Other Negative Correlation |
| SPATA20 | -0.12885 | 0.020538 | Other Negative Correlation |
| ARFIP2 | -0.12889 | 0.020498 | Other Negative Correlation |
| DPCD | -0.12916 | 0.020234 | Other Negative Correlation |
| VIRMA | -0.12925 | 0.020147 | Other Negative Correlation |
| GBE1 | -0.1295 | 0.019905 | Other Negative Correlation |
| FOXO3 | -0.12952 | 0.01988 | Other Negative Correlation |

| | | | |
|---|---|---|---|
| TRIM73 | -0.12961 | 0.0198 | Other Negative Correlation |
| FBXO38 | -0.12966 | 0.019751 | Other Negative Correlation |
| RSPH3 | -0.12992 | 0.0195 | Other Negative Correlation |
| CORO1A | -0.1304 | 0.019056 | Other Negative Correlation |
| NECAB3 | -0.13052 | 0.018946 | Other Negative Correlation |
| SMAD3 | -0.13055 | 0.018917 | Other Negative Correlation |
| DOCK6 | -0.13059 | 0.018881 | Other Negative Correlation |
| GIMAP1 | -0.13064 | 0.018831 | Other Negative Correlation |
| MYCBPAP | -0.13077 | 0.018716 | Other Negative Correlation |
| ITGB6 | -0.13097 | 0.018533 | Other Negative Correlation |
| BNIP2 | -0.13098 | 0.018519 | Other Negative Correlation |
| POLR3GL | -0.13104 | 0.018469 | Other Negative Correlation |
| POMZP3 | -0.13105 | 0.018455 | Other Negative Correlation |
| KCMF1 | -0.13111 | 0.018409 | Other Negative Correlation |
| BRWD3 | -0.13128 | 0.018254 | Other Negative Correlation |
| CACUL1 | -0.13128 | 0.018254 | Other Negative Correlation |
| ARFGEF1 | -0.13135 | 0.018192 | Other Negative Correlation |
| ATXN3 | -0.13151 | 0.018047 | Other Negative Correlation |
| PHLDB1 | -0.13154 | 0.018017 | Other Negative Correlation |
| DNAJC18 | -0.13165 | 0.017924 | Other Negative Correlation |
| UCKL1 | -0.13173 | 0.01785 | Other Negative Correlation |
| TARBP1 | -0.13192 | 0.017692 | Other Negative Correlation |
| LPCAT2 | -0.13193 | 0.017676 | Other Negative Correlation |
| COMMD3 | -0.13209 | 0.017542 | Other Negative Correlation |
| AKAP11 | -0.13213 | 0.017511 | Other Negative Correlation |
| SLC35D2 | -0.1323 | 0.017363 | Other Negative Correlation |
| EYS | -0.13238 | 0.017293 | Other Negative Correlation |
| RAB34 | -0.13251 | 0.017183 | Other Negative Correlation |
| TTC3 | -0.13255 | 0.017146 | Other Negative Correlation |
| RGS5 | -0.1326 | 0.017107 | Other Negative Correlation |
| CMAS | -0.1326 | 0.017107 | Other Negative Correlation |
| CORO7 | -0.13263 | 0.017078 | Other Negative Correlation |
| CAND1 | -0.13268 | 0.017041 | Other Negative Correlation |
| PRKG2 | -0.13268 | 0.017036 | Other Negative Correlation |
| FAM177A1 | -0.13276 | 0.016969 | Other Negative Correlation |
| CHD4 | -0.1328 | 0.016938 | Other Negative Correlation |
| DDB1 | -0.13298 | 0.016786 | Other Negative Correlation |
| CTTN | -0.13309 | 0.016701 | Other Negative Correlation |
| C5orf15 | -0.1331 | 0.016692 | Other Negative Correlation |
| YIF1A | -0.1331 | 0.01669 | Other Negative Correlation |
| GLT8D1 | -0.13314 | 0.016657 | Other Negative Correlation |
| USP4 | -0.13327 | 0.016553 | Other Negative Correlation |
| LIPT1 | -0.1333 | 0.016525 | Other Negative Correlation |
| ZCCHC17 | -0.13337 | 0.016469 | Other Negative Correlation |
| PSMB6 | -0.13359 | 0.016289 | Other Negative Correlation |
| UFSP2 | -0.13364 | 0.016248 | Other Negative Correlation |
| CSNK2A1 | -0.13368 | 0.016216 | Other Negative Correlation |
| TMEM79 | -0.13375 | 0.016161 | Other Negative Correlation |

| | | | |
|---|---|---|---|
| ACOX3 | -0.13379 | 0.016132 | Other Negative Correlation |
| ST6GALNAC4 | -0.13379 | 0.016131 | Other Negative Correlation |
| MAML2 | -0.13388 | 0.016055 | Other Negative Correlation |
| TRAM1 | -0.13396 | 0.015989 | Other Negative Correlation |
| RNPC3 | -0.13402 | 0.015945 | Other Negative Correlation |
| PGAP1 | -0.13415 | 0.015844 | Other Negative Correlation |
| ZNF397 | -0.13416 | 0.015835 | Other Negative Correlation |
| TMX3 | -0.13419 | 0.015809 | Other Negative Correlation |
| TRIM8 | -0.13426 | 0.015754 | Other Negative Correlation |
| BCAP29 | -0.13429 | 0.015732 | Other Negative Correlation |
| PSMA4 | -0.13466 | 0.015447 | Other Negative Correlation |
| TMEM254 | -0.13491 | 0.015251 | Other Negative Correlation |
| MT-ND5 | -0.13512 | 0.015092 | Other Negative Correlation |
| INVS | -0.13522 | 0.015019 | Other Negative Correlation |
| TMEM87B | -0.13534 | 0.014927 | Other Negative Correlation |
| RBMS3 | -0.13534 | 0.014924 | Other Negative Correlation |
| FAT4 | -0.13564 | 0.014702 | Other Negative Correlation |
| NSMAF | -0.13586 | 0.014543 | Other Negative Correlation |
| ACAD8 | -0.13586 | 0.014539 | Other Negative Correlation |
| TMEM18 | -0.13594 | 0.014483 | Other Negative Correlation |
| NIPBL | -0.13606 | 0.014397 | Other Negative Correlation |
| KIZ | -0.13629 | 0.014229 | Other Negative Correlation |
| NME4 | -0.13633 | 0.0142 | Other Negative Correlation |
| EXT2 | -0.13637 | 0.014172 | Other Negative Correlation |
| DENR | -0.13648 | 0.014092 | Other Negative Correlation |
| GTF2F1 | -0.1365 | 0.014081 | Other Negative Correlation |
| CAPN12 | -0.13651 | 0.014072 | Other Negative Correlation |
| ABCB7 | -0.1366 | 0.014009 | Other Negative Correlation |
| MBD5 | -0.13664 | 0.013982 | Other Negative Correlation |
| MARVELD1 | -0.13676 | 0.013899 | Other Negative Correlation |
| PSG4 | -0.13686 | 0.013829 | Other Negative Correlation |
| CCNDBP1 | -0.13692 | 0.013786 | Other Negative Correlation |
| CLYBL | -0.137 | 0.013732 | Other Negative Correlation |
| ZNF839 | -0.13704 | 0.013703 | Other Negative Correlation |
| GSAP | -0.13704 | 0.013703 | Other Negative Correlation |
| OGDH | -0.13707 | 0.01368 | Other Negative Correlation |
| ZNF708 | -0.13715 | 0.013627 | Other Negative Correlation |
| ITPR3 | -0.13753 | 0.013366 | Other Negative Correlation |
| IFT52 | -0.13776 | 0.013208 | Other Negative Correlation |
| CETN2 | -0.13785 | 0.01315 | Other Negative Correlation |
| RIDA | -0.13787 | 0.013135 | Other Negative Correlation |
| CTF1 | -0.13789 | 0.013121 | Other Negative Correlation |
| SRPX2 | -0.13793 | 0.013098 | Other Negative Correlation |
| STARD3NL | -0.13801 | 0.013044 | Other Negative Correlation |
| CCBE1 | -0.13801 | 0.013044 | Other Negative Correlation |
| ELFN2 | -0.13801 | 0.013043 | Other Negative Correlation |
| GRHPR | -0.13803 | 0.013028 | Other Negative Correlation |
| ZNF155 | -0.1381 | 0.012983 | Other Negative Correlation |

| | | | |
|---|---|---|---|
| DAW1 | -0.13827 | 0.012873 | Other Negative Correlation |
| RAB32 | -0.13837 | 0.012805 | Other Negative Correlation |
| PDIA4 | -0.13841 | 0.012779 | Other Negative Correlation |
| DHX57 | -0.13854 | 0.012697 | Other Negative Correlation |
| NAB1 | -0.13855 | 0.012688 | Other Negative Correlation |
| TRIM13 | -0.13898 | 0.01241 | Other Negative Correlation |
| SIRT5 | -0.13913 | 0.012313 | Other Negative Correlation |
| SPIRE2 | -0.13926 | 0.012231 | Other Negative Correlation |
| ACOX1 | -0.13934 | 0.012183 | Other Negative Correlation |
| TCF25 | -0.13934 | 0.012183 | Other Negative Correlation |
| KLHL3 | -0.13954 | 0.012062 | Other Negative Correlation |
| SLC16A4 | -0.13961 | 0.012019 | Other Negative Correlation |
| TUBA1A | -0.13962 | 0.012007 | Other Negative Correlation |
| NF1 | -0.13967 | 0.011978 | Other Negative Correlation |
| POMP | -0.13974 | 0.011934 | Other Negative Correlation |
| RBKS | -0.13983 | 0.011879 | Other Negative Correlation |
| PILRB | -0.13995 | 0.011808 | Other Negative Correlation |
| SUGP2 | -0.13997 | 0.011793 | Other Negative Correlation |
| KRCC1 | -0.13997 | 0.011793 | Other Negative Correlation |
| MMGT1 | -0.14013 | 0.011698 | Other Negative Correlation |
| FRRS1 | -0.14013 | 0.011697 | Other Negative Correlation |
| CCDC122 | -0.14027 | 0.011613 | Other Negative Correlation |
| TOB1 | -0.14044 | 0.011512 | Other Negative Correlation |
| CAMK1G | -0.14051 | 0.011469 | Other Negative Correlation |
| AKR1C2 | -0.14056 | 0.011438 | Other Negative Correlation |
| HIBADH | -0.14058 | 0.011428 | Other Negative Correlation |
| TRAF7 | -0.1406 | 0.011415 | Other Negative Correlation |
| TGFBI | -0.14075 | 0.011326 | Other Negative Correlation |
| MRPS25 | -0.14089 | 0.011244 | Other Negative Correlation |
| ACAA1 | -0.14094 | 0.011216 | Other Negative Correlation |
| MNAT1 | -0.14095 | 0.011211 | Other Negative Correlation |
| ERCC2 | -0.14096 | 0.011206 | Other Negative Correlation |
| SGF29 | -0.14098 | 0.011195 | Other Negative Correlation |
| CPSF2 | -0.14103 | 0.011164 | Other Negative Correlation |
| TBCK | -0.14109 | 0.011128 | Other Negative Correlation |
| BCL2L1 | -0.14127 | 0.011025 | Other Negative Correlation |
| SMAD2 | -0.14132 | 0.010998 | Other Negative Correlation |
| ATP5MJ | -0.14138 | 0.010965 | Other Negative Correlation |
| FGD4 | -0.14146 | 0.010919 | Other Negative Correlation |
| CAPNS2 | -0.14149 | 0.010903 | Other Negative Correlation |
| NFKB2 | -0.14161 | 0.010831 | Other Negative Correlation |
| AGK | -0.14167 | 0.010798 | Other Negative Correlation |
| METTL15 | -0.14197 | 0.010633 | Other Negative Correlation |
| AFMID | -0.14218 | 0.010518 | Other Negative Correlation |
| BBLN | -0.14224 | 0.010481 | Other Negative Correlation |
| SMARCA1 | -0.1425 | 0.010341 | Other Negative Correlation |
| GNA12 | -0.14261 | 0.01028 | Other Negative Correlation |
| ENSG00000271741 | -0.14281 | 0.010175 | Other Negative Correlation |

| | | | |
|---|---|---|---|
| EHD1 | -0.14313 | 0.010003 | Other Negative Correlation |
| MAPK10 | -0.14332 | 0.009904 | Other Negative Correlation |
| LRP4 | -0.14334 | 0.009894 | Other Negative Correlation |
| GPALPP1 | -0.14343 | 0.009847 | Other Negative Correlation |
| SH3GLB1 | -0.14364 | 0.00974 | Other Negative Correlation |
| GUCA1B | -0.14398 | 0.009567 | Other Negative Correlation |
| APOC1 | -0.14408 | 0.009517 | Other Negative Correlation |
| CKMT1B | -0.14409 | 0.009512 | Other Negative Correlation |
| RTCA | -0.14419 | 0.009462 | Other Negative Correlation |
| FPGT | -0.14424 | 0.009434 | Other Negative Correlation |
| TNFRSF14 | -0.14434 | 0.009388 | Other Negative Correlation |
| SPG7 | -0.14448 | 0.009314 | Other Negative Correlation |
| TP53INP1 | -0.14453 | 0.00929 | Other Negative Correlation |
| BASP1 | -0.14493 | 0.009095 | Other Negative Correlation |
| UBE2L6 | -0.14496 | 0.00908 | Other Negative Correlation |
| SULT1A1 | -0.14501 | 0.009057 | Other Negative Correlation |
| ADAMTSL4 | -0.14502 | 0.009054 | Other Negative Correlation |
| CNOT6 | -0.14509 | 0.009019 | Other Negative Correlation |
| CBR1 | -0.1452 | 0.008967 | Other Negative Correlation |
| STK16 | -0.14547 | 0.008839 | Other Negative Correlation |
| VDAC3 | -0.1455 | 0.008823 | Other Negative Correlation |
| TRIML2 | -0.14573 | 0.008719 | Other Negative Correlation |
| ARFIP1 | -0.14591 | 0.008632 | Other Negative Correlation |
| YLPM1 | -0.14599 | 0.008598 | Other Negative Correlation |
| CTSZ | -0.14603 | 0.008578 | Other Negative Correlation |
| LENG8 | -0.1461 | 0.008548 | Other Negative Correlation |
| AKT1S1 | -0.14629 | 0.008461 | Other Negative Correlation |
| DAPP1 | -0.1464 | 0.008412 | Other Negative Correlation |
| CAAP1 | -0.14644 | 0.008393 | Other Negative Correlation |
| MPI | -0.14662 | 0.008311 | Other Negative Correlation |
| TEPP | -0.14669 | 0.008282 | Other Negative Correlation |
| RAB6A | -0.14678 | 0.00824 | Other Negative Correlation |
| RSRP1 | -0.14693 | 0.008176 | Other Negative Correlation |
| ENSG00000262304 | -0.14716 | 0.008074 | Other Negative Correlation |
| RIT1 | -0.14737 | 0.007983 | Other Negative Correlation |
| ATP2B4 | -0.14741 | 0.007965 | Other Negative Correlation |
| PLD2 | -0.14752 | 0.007919 | Other Negative Correlation |
| STRIP2 | -0.14782 | 0.007793 | Other Negative Correlation |
| SLC26A7 | -0.14795 | 0.007736 | Other Negative Correlation |
| APOBEC3G | -0.1481 | 0.007676 | Other Negative Correlation |
| SNX32 | -0.14821 | 0.00763 | Other Negative Correlation |
| IFT88 | -0.14847 | 0.007521 | Other Negative Correlation |
| HOPX | -0.14862 | 0.00746 | Other Negative Correlation |
| RELB | -0.14898 | 0.007318 | Other Negative Correlation |
| TMEM98 | -0.14901 | 0.007303 | Other Negative Correlation |
| FEZ2 | -0.14908 | 0.007275 | Other Negative Correlation |
| CDKN1A | -0.1492 | 0.007228 | Other Negative Correlation |
| BAIAP2 | -0.14928 | 0.007199 | Other Negative Correlation |

| | | | |
|---|---|---|---|
| UQCRHL | -0.14944 | 0.007135 | Other Negative Correlation |
| ELAPOR1 | -0.14948 | 0.007121 | Other Negative Correlation |
| ODAD4 | -0.14949 | 0.007114 | Other Negative Correlation |
| WDR48 | -0.14964 | 0.007056 | Other Negative Correlation |
| BAZ2B | -0.14966 | 0.007049 | Other Negative Correlation |
| ENSG00000214265 | -0.14968 | 0.007041 | Other Negative Correlation |
| DHRSX | -0.14971 | 0.00703 | Other Negative Correlation |
| EDF1 | -0.14987 | 0.006971 | Other Negative Correlation |
| HELZ2 | -0.15012 | 0.006873 | Other Negative Correlation |
| SIDT2 | -0.15043 | 0.006757 | Other Negative Correlation |
| NBPF12 | -0.15052 | 0.006727 | Other Negative Correlation |
| OSBPL1A | -0.15066 | 0.006675 | Other Negative Correlation |
| PPP4R2 | -0.15069 | 0.006663 | Other Negative Correlation |
| TBC1D13 | -0.15093 | 0.006575 | Other Negative Correlation |
| CCDC107 | -0.15099 | 0.006553 | Other Negative Correlation |
| BCR | -0.15134 | 0.006429 | Other Negative Correlation |
| RPS6KA1 | -0.15158 | 0.006345 | Other Negative Correlation |
| ASMTL | -0.15164 | 0.006325 | Other Negative Correlation |
| C3 | -0.15191 | 0.006228 | Other Negative Correlation |
| ASH1L | -0.15211 | 0.00616 | Other Negative Correlation |
| ZBED5 | -0.15228 | 0.006104 | Other Negative Correlation |
| PHF6 | -0.15237 | 0.006074 | Other Negative Correlation |
| RNF213 | -0.15257 | 0.006006 | Other Negative Correlation |
| SESN3 | -0.15258 | 0.006001 | Other Negative Correlation |
| SNX33 | -0.15268 | 0.005967 | Other Negative Correlation |
| FKBP14 | -0.15271 | 0.00596 | Other Negative Correlation |
| PSEN1 | -0.15291 | 0.005892 | Other Negative Correlation |
| DDR1 | -0.15315 | 0.005815 | Other Negative Correlation |
| GALNT7 | -0.15324 | 0.005786 | Other Negative Correlation |
| BCO2 | -0.15326 | 0.005778 | Other Negative Correlation |
| DLG1 | -0.15339 | 0.005738 | Other Negative Correlation |
| TNKS1BP1 | -0.15353 | 0.005694 | Other Negative Correlation |
| SLC27A1 | -0.15367 | 0.005648 | Other Negative Correlation |
| DHRS1 | -0.15373 | 0.005629 | Other Negative Correlation |
| RPSAP58 | -0.15386 | 0.00559 | Other Negative Correlation |
| CNOT4 | -0.15399 | 0.005549 | Other Negative Correlation |
| TBC1D22A | -0.15399 | 0.005547 | Other Negative Correlation |
| TPGS1 | -0.15402 | 0.00554 | Other Negative Correlation |
| NUDCD2 | -0.15407 | 0.005522 | Other Negative Correlation |
| PMVK | -0.15431 | 0.00545 | Other Negative Correlation |
| ANO7 | -0.15435 | 0.005436 | Other Negative Correlation |
| COX4I1 | -0.1544 | 0.005421 | Other Negative Correlation |
| SPCS1 | -0.15443 | 0.005413 | Other Negative Correlation |
| OBSCN | -0.15445 | 0.005408 | Other Negative Correlation |
| SNAPC2 | -0.15477 | 0.005312 | Other Negative Correlation |
| EXOC6B | -0.15479 | 0.005305 | Other Negative Correlation |
| BFSP1 | -0.15481 | 0.005298 | Other Negative Correlation |
| CST3 | -0.15493 | 0.005263 | Other Negative Correlation |

| | | | |
|---|---|---|---|
| ZNF558 | -0.15494 | 0.005259 | Other Negative Correlation |
| SP110 | -0.155 | 0.005241 | Other Negative Correlation |
| ZNF827 | -0.15502 | 0.005236 | Other Negative Correlation |
| ANO6 | -0.15514 | 0.005201 | Other Negative Correlation |
| PRSS48 | -0.15522 | 0.005178 | Other Negative Correlation |
| NEPRO | -0.1553 | 0.005154 | Other Negative Correlation |
| COPA | -0.15532 | 0.005148 | Other Negative Correlation |
| PUM1 | -0.15536 | 0.005137 | Other Negative Correlation |
| ENSG00000233757 | -0.15541 | 0.005122 | Other Negative Correlation |
| SCOC | -0.15558 | 0.005072 | Other Negative Correlation |
| USP9X | -0.15566 | 0.00505 | Other Negative Correlation |
| WDR59 | -0.15567 | 0.005046 | Other Negative Correlation |
| PLEKHH1 | -0.1557 | 0.00504 | Other Negative Correlation |
| NFKBIA | -0.1557 | 0.00504 | Other Negative Correlation |
| SDHA | -0.1557 | 0.005038 | Other Negative Correlation |
| CUX1 | -0.15572 | 0.005033 | Other Negative Correlation |
| MAP3K13 | -0.15606 | 0.004938 | Other Negative Correlation |
| SNX12 | -0.15627 | 0.004879 | Other Negative Correlation |
| XRRA1 | -0.15627 | 0.004879 | Other Negative Correlation |
| KLHL13 | -0.1564 | 0.004842 | Other Negative Correlation |
| CABYR | -0.15679 | 0.004737 | Other Negative Correlation |
| TREM2 | -0.15681 | 0.004732 | Other Negative Correlation |
| TMEM92 | -0.15687 | 0.004716 | Other Negative Correlation |
| TTC39B | -0.15703 | 0.004673 | Other Negative Correlation |
| PRKCZ | -0.15713 | 0.004645 | Other Negative Correlation |
| DNAJB5 | -0.15721 | 0.004624 | Other Negative Correlation |
| SCPEP1 | -0.15723 | 0.00462 | Other Negative Correlation |
| FBXO6 | -0.15735 | 0.004588 | Other Negative Correlation |
| COX8A | -0.15737 | 0.004583 | Other Negative Correlation |
| BCAS4 | -0.15739 | 0.004577 | Other Negative Correlation |
| PTMS | -0.15799 | 0.004423 | Other Negative Correlation |
| GABBR1 | -0.15816 | 0.004381 | Other Negative Correlation |
| SLC24A1 | -0.15827 | 0.004353 | Other Negative Correlation |
| BLNK | -0.1583 | 0.004346 | Other Negative Correlation |
| MAGED1 | -0.15832 | 0.004339 | Other Negative Correlation |
| TLK1 | -0.15833 | 0.004337 | Other Negative Correlation |
| RIC8A | -0.15845 | 0.004307 | Other Negative Correlation |
| AKT2 | -0.15886 | 0.004207 | Other Negative Correlation |
| NUMA1 | -0.15888 | 0.004203 | Other Negative Correlation |
| NAXE | -0.15888 | 0.004203 | Other Negative Correlation |
| ZHX3 | -0.15906 | 0.004158 | Other Negative Correlation |
| KANK1 | -0.15911 | 0.004148 | Other Negative Correlation |
| DIXDC1 | -0.15935 | 0.004089 | Other Negative Correlation |
| ATP11A | -0.15942 | 0.004073 | Other Negative Correlation |
| ABHD14A | -0.15959 | 0.004033 | Other Negative Correlation |
| PPIE | -0.15975 | 0.003997 | Other Negative Correlation |
| ACADM | -0.16005 | 0.003927 | Other Negative Correlation |
| KIF3A | -0.16047 | 0.003834 | Other Negative Correlation |

| | | | |
|---|---|---|---|
| QARS1 | -0.16053 | 0.003819 | Other Negative Correlation |
| NUDT7 | -0.16088 | 0.003743 | Other Negative Correlation |
| ECPAS | -0.16112 | 0.00369 | Other Negative Correlation |
| ZDHHC20 | -0.16127 | 0.003658 | Other Negative Correlation |
| SLU7 | -0.16142 | 0.003626 | Other Negative Correlation |
| PTGR3 | -0.16146 | 0.003618 | Other Negative Correlation |
| ZNF451 | -0.16176 | 0.003555 | Other Negative Correlation |
| SOD1 | -0.16183 | 0.00354 | Other Negative Correlation |
| FSIP2 | -0.16186 | 0.003535 | Other Negative Correlation |
| HOMER3 | -0.16192 | 0.003522 | Other Negative Correlation |
| PSD2 | -0.16215 | 0.003475 | Other Negative Correlation |
| COPB2 | -0.16218 | 0.003469 | Other Negative Correlation |
| DNAH9 | -0.16221 | 0.003463 | Other Negative Correlation |
| RERE | -0.16224 | 0.003456 | Other Negative Correlation |
| PRKCA | -0.16233 | 0.003438 | Other Negative Correlation |
| TMEM219 | -0.16237 | 0.00343 | Other Negative Correlation |
| KIAA1522 | -0.16249 | 0.003406 | Other Negative Correlation |
| ECI2 | -0.16254 | 0.003396 | Other Negative Correlation |
| UFC1 | -0.16261 | 0.003382 | Other Negative Correlation |
| ARAP1 | -0.16282 | 0.003341 | Other Negative Correlation |
| CWC22 | -0.16288 | 0.003328 | Other Negative Correlation |
| COL16A1 | -0.16297 | 0.003312 | Other Negative Correlation |
| H6PD | -0.163 | 0.003305 | Other Negative Correlation |
| TSC22D1 | -0.16387 | 0.00314 | Other Negative Correlation |
| SLC66A3 | -0.16388 | 0.003138 | Other Negative Correlation |
| GRN | -0.16397 | 0.003121 | Other Negative Correlation |
| TNFAIP8 | -0.16404 | 0.003109 | Other Negative Correlation |
| CTNNA1 | -0.16418 | 0.003082 | Other Negative Correlation |
| HBEGF | -0.16421 | 0.003077 | Other Negative Correlation |
| TSEN34 | -0.16431 | 0.003059 | Other Negative Correlation |
| FBXW8 | -0.16479 | 0.002973 | Other Negative Correlation |
| PEX11G | -0.16485 | 0.002963 | Other Negative Correlation |
| TXN2 | -0.16486 | 0.00296 | Other Negative Correlation |
| IL27RA | -0.16492 | 0.002951 | Other Negative Correlation |
| ACAA2 | -0.16498 | 0.00294 | Other Negative Correlation |
| SOX6 | -0.16508 | 0.002922 | Other Negative Correlation |
| PSORS1C1 | -0.16521 | 0.002899 | Other Negative Correlation |
| LETMD1 | -0.16525 | 0.002892 | Other Negative Correlation |
| ARL1 | -0.16532 | 0.002881 | Other Negative Correlation |
| CD58 | -0.16553 | 0.002846 | Other Negative Correlation |
| NDUFB7 | -0.16563 | 0.002828 | Other Negative Correlation |
| CDK17 | -0.1659 | 0.002782 | Other Negative Correlation |
| NDUFA11 | -0.16609 | 0.002752 | Other Negative Correlation |
| MMP9 | -0.1661 | 0.002749 | Other Negative Correlation |
| TUBG2 | -0.16646 | 0.002691 | Other Negative Correlation |
| EPS15 | -0.16655 | 0.002676 | Other Negative Correlation |
| CDCP1 | -0.16669 | 0.002654 | Other Negative Correlation |
| COX7B | -0.16675 | 0.002644 | Other Negative Correlation |

| | | | |
|---|---|---|---|
| DLG5 | -0.1671 | 0.002589 | Other Negative Correlation |
| CHP1 | -0.16719 | 0.002575 | Other Negative Correlation |
| DQX1 | -0.16733 | 0.002554 | Other Negative Correlation |
| CCL20 | -0.16756 | 0.002518 | Other Negative Correlation |
| SCEL | -0.16758 | 0.002515 | Other Negative Correlation |
| PABPC4L | -0.16774 | 0.002492 | Other Negative Correlation |
| RAP1GAP2 | -0.1678 | 0.002483 | Other Negative Correlation |
| SYT12 | -0.16807 | 0.002442 | Other Negative Correlation |
| BLOC1S6 | -0.16818 | 0.002426 | Other Negative Correlation |
| GALNT11 | -0.16826 | 0.002414 | Other Negative Correlation |
| MARCHF4 | -0.16856 | 0.00237 | Other Negative Correlation |
| RARS2 | -0.16864 | 0.002359 | Other Negative Correlation |
| STAG2 | -0.16901 | 0.002306 | Other Negative Correlation |
| AAGAB | -0.16908 | 0.002297 | Other Negative Correlation |
| SLX4IP | -0.16916 | 0.002286 | Other Negative Correlation |
| CLTB | -0.16917 | 0.002283 | Other Negative Correlation |
| AVL9 | -0.16938 | 0.002255 | Other Negative Correlation |
| HEXB | -0.16958 | 0.002228 | Other Negative Correlation |
| MLF2 | -0.1698 | 0.002197 | Other Negative Correlation |
| CERS3 | -0.17012 | 0.002155 | Other Negative Correlation |
| SNX1 | -0.17044 | 0.002114 | Other Negative Correlation |
| SGTB | -0.17052 | 0.002102 | Other Negative Correlation |
| BTBD7 | -0.17058 | 0.002095 | Other Negative Correlation |
| USP9Y | -0.17061 | 0.002091 | Other Negative Correlation |
| MYO1C | -0.17078 | 0.00207 | Other Negative Correlation |
| ATP6V0C | -0.17087 | 0.002058 | Other Negative Correlation |
| MGLL | -0.17117 | 0.00202 | Other Negative Correlation |
| GLIPR1L2 | -0.1713 | 0.002005 | Other Negative Correlation |
| IRX2 | -0.1715 | 0.00198 | Other Negative Correlation |
| SEMA4A | -0.17173 | 0.001952 | Other Negative Correlation |
| TWF2 | -0.17174 | 0.00195 | Other Negative Correlation |
| KTN1 | -0.1721 | 0.001908 | Other Negative Correlation |
| ECHS1 | -0.17213 | 0.001904 | Other Negative Correlation |
| SYNPO | -0.17246 | 0.001866 | Other Negative Correlation |
| NUMB | -0.17258 | 0.001852 | Other Negative Correlation |
| GPSM3 | -0.1728 | 0.001826 | Other Negative Correlation |
| ERG28 | -0.17286 | 0.001819 | Other Negative Correlation |
| HMGN4 | -0.17295 | 0.00181 | Other Negative Correlation |
| FYTTD1 | -0.17338 | 0.001762 | Other Negative Correlation |
| UQCR10 | -0.1737 | 0.001727 | Other Negative Correlation |
| PTAFR | -0.1738 | 0.001717 | Other Negative Correlation |
| SHARPIN | -0.17398 | 0.001698 | Other Negative Correlation |
| ATP5IF1 | -0.1741 | 0.001684 | Other Negative Correlation |
| TPR | -0.17415 | 0.001679 | Other Negative Correlation |
| ETNK1 | -0.17416 | 0.001679 | Other Negative Correlation |
| TENT2 | -0.17428 | 0.001666 | Other Negative Correlation |
| TRIM38 | -0.17436 | 0.001657 | Other Negative Correlation |
| S100A13 | -0.17448 | 0.001645 | Other Negative Correlation |

| | | | |
|---|---|---|---|
| RRAS | -0.17457 | 0.001636 | Other Negative Correlation |
| SSC5D | -0.17472 | 0.00162 | Other Negative Correlation |
| RAB2A | -0.17488 | 0.001604 | Other Negative Correlation |
| PNISR | -0.17522 | 0.00157 | Other Negative Correlation |
| HDAC8 | -0.17533 | 0.001559 | Other Negative Correlation |
| ME2 | -0.17536 | 0.001556 | Other Negative Correlation |
| IARS2 | -0.17555 | 0.001538 | Other Negative Correlation |
| PRKAR2A | -0.1756 | 0.001533 | Other Negative Correlation |
| SLC15A4 | -0.1756 | 0.001533 | Other Negative Correlation |
| PARP3 | -0.17563 | 0.00153 | Other Negative Correlation |
| PGM1 | -0.17565 | 0.001528 | Other Negative Correlation |
| MTFMT | -0.17567 | 0.001526 | Other Negative Correlation |
| TIPRL | -0.17585 | 0.001509 | Other Negative Correlation |
| RSU1 | -0.17585 | 0.001509 | Other Negative Correlation |
| PCLO | -0.1759 | 0.001504 | Other Negative Correlation |
| DVL1 | -0.17609 | 0.001487 | Other Negative Correlation |
| MGMT | -0.17628 | 0.001469 | Other Negative Correlation |
| SLC35E1 | -0.17665 | 0.001434 | Other Negative Correlation |
| ARL3 | -0.17679 | 0.001422 | Other Negative Correlation |
| CELSR2 | -0.17691 | 0.001411 | Other Negative Correlation |
| PSD3 | -0.17702 | 0.001401 | Other Negative Correlation |
| STXBP3 | -0.17726 | 0.001381 | Other Negative Correlation |
| ASIC1 | -0.17726 | 0.00138 | Other Negative Correlation |
| CDH4 | -0.17734 | 0.001373 | Other Negative Correlation |
| ALDH1A1 | -0.17739 | 0.001369 | Other Negative Correlation |
| PPP1R10 | -0.17744 | 0.001364 | Other Negative Correlation |
| ANKDD1A | -0.17759 | 0.001351 | Other Negative Correlation |
| CYP2S1 | -0.17765 | 0.001346 | Other Negative Correlation |
| VPS35 | -0.17768 | 0.001344 | Other Negative Correlation |
| AP1M2 | -0.17788 | 0.001326 | Other Negative Correlation |
| CPA4 | -0.17791 | 0.001324 | Other Negative Correlation |
| UBA1 | -0.17828 | 0.001293 | Other Negative Correlation |
| BEX3 | -0.17843 | 0.001281 | Other Negative Correlation |
| SSH3 | -0.17844 | 0.00128 | Other Negative Correlation |
| COMMD6 | -0.17863 | 0.001265 | Other Negative Correlation |
| C1orf159 | -0.17864 | 0.001263 | Other Negative Correlation |
| FXR1 | -0.1787 | 0.001259 | Other Negative Correlation |
| ACTL10 | -0.17874 | 0.001256 | Other Negative Correlation |
| PARD3 | -0.1793 | 0.001211 | Other Negative Correlation |
| FTL | -0.17948 | 0.001197 | Other Negative Correlation |
| DNAJC5 | -0.17951 | 0.001195 | Other Negative Correlation |
| VPS52 | -0.17978 | 0.001174 | Other Negative Correlation |
| DOP1A | -0.17983 | 0.001171 | Other Negative Correlation |
| DHRS4L2 | -0.18018 | 0.001144 | Other Negative Correlation |
| MVP | -0.18025 | 0.00114 | Other Negative Correlation |
| CPNE1 | -0.18047 | 0.001124 | Other Negative Correlation |
| DYRK4 | -0.18047 | 0.001123 | Other Negative Correlation |
| SCG5 | -0.18048 | 0.001123 | Other Negative Correlation |

| | | | |
|---|---|---|---|
| NOL3 | -0.18058 | 0.001116 | Other Negative Correlation |
| EXOC5 | -0.18072 | 0.001105 | Other Negative Correlation |
| POMGNT1 | -0.18137 | 0.00106 | Other Negative Correlation |
| TAFA2 | -0.18176 | 0.001033 | Other Negative Correlation |
| MAP2K6 | -0.18183 | 0.001028 | Other Negative Correlation |
| SPG11 | -0.18194 | 0.001021 | Other Negative Correlation |
| HERC4 | -0.18211 | 0.00101 | Other Negative Correlation |
| RNF7 | -0.1822 | 0.001004 | Other Negative Correlation |
| OTUB1 | -0.18226 | 0.001 | Other Negative Correlation |
| RILPL1 | -0.18254 | 0.000982 | Other Negative Correlation |
| RRBP1 | -0.18255 | 0.000981 | Other Negative Correlation |
| PLEKHG5 | -0.1826 | 0.000978 | Other Negative Correlation |
| ABI1 | -0.18307 | 0.000948 | Other Negative Correlation |
| LNPEP | -0.18322 | 0.000939 | Other Negative Correlation |
| GID8 | -0.18328 | 0.000936 | Other Negative Correlation |
| NBDY | -0.18351 | 0.000922 | Other Negative Correlation |
| PUDP | -0.18362 | 0.000915 | Other Negative Correlation |
| UBA3 | -0.18364 | 0.000913 | Other Negative Correlation |
| FAM210B | -0.18367 | 0.000912 | Other Negative Correlation |
| NEMF | -0.18374 | 0.000908 | Other Negative Correlation |
| GLG1 | -0.18394 | 0.000896 | Other Negative Correlation |
| PSMB7 | -0.18411 | 0.000886 | Other Negative Correlation |
| TMEM131L | -0.18437 | 0.000871 | Other Negative Correlation |
| DLGAP4 | -0.1844 | 0.000869 | Other Negative Correlation |
| TJP1 | -0.18475 | 0.000849 | Other Negative Correlation |
| PITPNM3 | -0.18483 | 0.000844 | Other Negative Correlation |
| ERBIN | -0.18516 | 0.000826 | Other Negative Correlation |
| DNAJC10 | -0.18518 | 0.000825 | Other Negative Correlation |
| TUSC3 | -0.18555 | 0.000805 | Other Negative Correlation |
| PPP2CB | -0.186 | 0.000781 | Other Negative Correlation |
| PIK3C3 | -0.18604 | 0.00078 | Other Negative Correlation |
| FAM120B | -0.18672 | 0.000745 | Other Negative Correlation |
| RNF170 | -0.18679 | 0.000742 | Other Negative Correlation |
| AGR2 | -0.1868 | 0.000741 | Other Negative Correlation |
| QRICH1 | -0.18704 | 0.000729 | Other Negative Correlation |
| ADGRA3 | -0.18715 | 0.000724 | Other Negative Correlation |
| RNPEP | -0.188 | 0.000684 | Other Negative Correlation |
| NDUFAB1 | -0.18832 | 0.000669 | Other Negative Correlation |
| MORF4L1 | -0.18869 | 0.000653 | Other Negative Correlation |
| VPS13B | -0.189 | 0.000639 | Other Negative Correlation |
| NCKIPSD | -0.18951 | 0.000617 | Other Negative Correlation |
| NIPAL2 | -0.18961 | 0.000613 | Other Negative Correlation |
| PHKA2 | -0.18965 | 0.000612 | Other Negative Correlation |
| SMARCD2 | -0.19014 | 0.000592 | Other Negative Correlation |
| MAN1B1 | -0.19019 | 0.00059 | Other Negative Correlation |
| NMRK1 | -0.19028 | 0.000586 | Other Negative Correlation |
| FDXR | -0.19033 | 0.000584 | Other Negative Correlation |
| NCOA4 | -0.19035 | 0.000583 | Other Negative Correlation |

| | | | |
|---|---|---|---|
| COX6B1 | -0.19053 | 0.000576 | Other Negative Correlation |
| IFNGR1 | -0.19067 | 0.000571 | Other Negative Correlation |
| NFRKB | -0.19068 | 0.000571 | Other Negative Correlation |
| SIPA1L1 | -0.19088 | 0.000562 | Other Negative Correlation |
| PRRG4 | -0.19091 | 0.000562 | Other Negative Correlation |
| NENF | -0.19094 | 0.000561 | Other Negative Correlation |
| MRPL47 | -0.19116 | 0.000552 | Other Negative Correlation |
| DPY19L4 | -0.19129 | 0.000547 | Other Negative Correlation |
| ELF1 | -0.19149 | 0.00054 | Other Negative Correlation |
| APPL2 | -0.19156 | 0.000537 | Other Negative Correlation |
| TRIM16 | -0.19167 | 0.000533 | Other Negative Correlation |
| ANAPC5 | -0.19187 | 0.000526 | Other Negative Correlation |
| ODF3B | -0.19191 | 0.000524 | Other Negative Correlation |
| ENSG00000275740 | -0.19266 | 0.000498 | Other Negative Correlation |
| MIB2 | -0.19267 | 0.000498 | Other Negative Correlation |
| ENSG00000258728 | -0.1928 | 0.000493 | Other Negative Correlation |
| TAOK3 | -0.1937 | 0.000464 | Other Negative Correlation |
| MMADHC | -0.19371 | 0.000463 | Other Negative Correlation |
| LIMD1 | -0.19391 | 0.000457 | Other Negative Correlation |
| RASA3 | -0.19399 | 0.000455 | Other Negative Correlation |
| PHF20L1 | -0.19427 | 0.000446 | Other Negative Correlation |
| ADD1 | -0.19457 | 0.000437 | Other Negative Correlation |
| RAB1A | -0.19469 | 0.000433 | Other Negative Correlation |
| NEXN | -0.19547 | 0.00041 | Other Negative Correlation |
| SPRED1 | -0.19645 | 0.000383 | Other Negative Correlation |
| TSTD1 | -0.19647 | 0.000382 | Other Negative Correlation |
| KIAA1671 | -0.19662 | 0.000378 | Other Negative Correlation |
| LHFPL6 | -0.19674 | 0.000375 | Other Negative Correlation |
| TAPBPL | -0.19691 | 0.000371 | Other Negative Correlation |
| MISP | -0.19691 | 0.000371 | Other Negative Correlation |
| GPATCH2L | -0.19751 | 0.000355 | Other Negative Correlation |
| SELENON | -0.19774 | 0.00035 | Other Negative Correlation |
| ARHGEF10 | -0.19822 | 0.000338 | Other Negative Correlation |
| ALOX5AP | -0.19842 | 0.000333 | Other Negative Correlation |
| ENSG00000255639 | -0.19856 | 0.00033 | Other Negative Correlation |
| LIMD2 | -0.19936 | 0.000312 | Other Negative Correlation |
| RAB13 | -0.19942 | 0.00031 | Other Negative Correlation |
| PPP1CA | -0.19943 | 0.00031 | Other Negative Correlation |
| ATP5PD | -0.19965 | 0.000305 | Other Negative Correlation |
| ZKSCAN1 | -0.19972 | 0.000304 | Other Negative Correlation |
| ZNF274 | -0.19995 | 0.000299 | Other Negative Correlation |
| NOP10 | -0.20003 | 0.000297 | Other Negative Correlation |
| PIAS1 | -0.20029 | 0.000292 | Other Negative Correlation |
| ZBTB8A | -0.20034 | 0.000291 | Other Negative Correlation |
| FTH1 | -0.20095 | 0.000278 | Other Negative Correlation |
| ACADVL | -0.2011 | 0.000275 | Other Negative Correlation |
| DCTN2 | -0.20123 | 0.000273 | Other Negative Correlation |
| EDEM1 | -0.20125 | 0.000272 | Other Negative Correlation |

| | | | |
|---|---|---|---|
| AP3B1 | -0.2013 | 0.000271 | Other Negative Correlation |
| PORCN | -0.20145 | 0.000269 | Other Negative Correlation |
| MAP2K3 | -0.20149 | 0.000268 | Other Negative Correlation |
| CRIP2 | -0.20176 | 0.000263 | Other Negative Correlation |
| MXD4 | -0.20254 | 0.000248 | Other Negative Correlation |
| TRAK1 | -0.20267 | 0.000246 | Other Negative Correlation |
| CHMP5 | -0.20269 | 0.000246 | Other Negative Correlation |
| TPBG | -0.20297 | 0.000241 | Other Negative Correlation |
| PRRC2B | -0.2032 | 0.000237 | Other Negative Correlation |
| ATP5MF | -0.20355 | 0.000231 | Other Negative Correlation |
| PSTPIP2 | -0.2039 | 0.000225 | Other Negative Correlation |
| PYM1 | -0.20415 | 0.000221 | Other Negative Correlation |
| SNAP23 | -0.20421 | 0.00022 | Other Negative Correlation |
| ATM | -0.20427 | 0.000219 | Other Negative Correlation |
| RUFY1 | -0.2044 | 0.000217 | Other Negative Correlation |
| MOB1A | -0.20446 | 0.000216 | Other Negative Correlation |
| NDUFA4 | -0.20516 | 0.000205 | Other Negative Correlation |
| TBC1D5 | -0.20524 | 0.000204 | Other Negative Correlation |
| RFXANK | -0.2054 | 0.000202 | Other Negative Correlation |
| AK3 | -0.20554 | 0.0002 | Other Negative Correlation |
| DMD | -0.20559 | 0.000199 | Other Negative Correlation |
| ERMP1 | -0.2064 | 0.000187 | Other Negative Correlation |
| TACSTD2 | -0.20665 | 0.000184 | Other Negative Correlation |
| TMEM164 | -0.20667 | 0.000184 | Other Negative Correlation |
| EPN1 | -0.20686 | 0.000181 | Other Negative Correlation |
| PTRH1 | -0.20686 | 0.000181 | Other Negative Correlation |
| PAPOLA | -0.20761 | 0.000171 | Other Negative Correlation |
| WDR70 | -0.20768 | 0.000171 | Other Negative Correlation |
| DCAF6 | -0.20802 | 0.000166 | Other Negative Correlation |
| PTPRE | -0.20807 | 0.000166 | Other Negative Correlation |
| RCN1 | -0.20808 | 0.000166 | Other Negative Correlation |
| MECR | -0.20834 | 0.000162 | Other Negative Correlation |
| GSDMD | -0.20854 | 0.00016 | Other Negative Correlation |
| CAPZB | -0.20865 | 0.000159 | Other Negative Correlation |
| UBA7 | -0.20879 | 0.000157 | Other Negative Correlation |
| GSTK1 | -0.20893 | 0.000155 | Other Negative Correlation |
| FBXL4 | -0.20894 | 0.000155 | Other Negative Correlation |
| AGTRAP | -0.20903 | 0.000154 | Other Negative Correlation |
| MICALL2 | -0.20909 | 0.000154 | Other Negative Correlation |
| MRPL21 | -0.20939 | 0.00015 | Other Negative Correlation |
| ARPC1B | -0.2099 | 0.000145 | Other Negative Correlation |
| VWDE | -0.21006 | 0.000143 | Other Negative Correlation |
| NT5DC1 | -0.21016 | 0.000142 | Other Negative Correlation |
| NDUFAF3 | -0.2102 | 0.000141 | Other Negative Correlation |
| SPTBN2 | -0.21039 | 0.000139 | Other Negative Correlation |
| MINK1 | -0.21041 | 0.000139 | Other Negative Correlation |
| KDELR1 | -0.211 | 0.000133 | Other Negative Correlation |
| STING1 | -0.21117 | 0.000131 | Other Negative Correlation |

| | | | |
|---|---|---|---|
| SRPK2 | -0.21129 | 0.00013 | Other Negative Correlation |
| SCP2 | -0.21158 | 0.000127 | Other Negative Correlation |
| TKFC | -0.21232 | 0.000121 | Other Negative Correlation |
| NIPSNAP1 | -0.21246 | 0.000119 | Other Negative Correlation |
| TMC4 | -0.21308 | 0.000114 | Other Negative Correlation |
| C4orf3 | -0.21331 | 0.000112 | Other Negative Correlation |
| METTL7A | -0.21354 | 0.00011 | Other Negative Correlation |
| CAST | -0.21361 | 0.000109 | Other Negative Correlation |
| UQCRC2 | -0.21397 | 0.000106 | Other Negative Correlation |
| GBP3 | -0.21403 | 0.000106 | Other Negative Correlation |
| ATRX | -0.21412 | 0.000105 | Other Negative Correlation |
| CDC42EP3 | -0.21435 | 0.000103 | Other Negative Correlation |
| PIR | -0.21441 | 0.000103 | Other Negative Correlation |
| CCDC113 | -0.2145 | 0.000102 | Other Negative Correlation |
| ADD3 | -0.21477 | 0.0001 | Other Negative Correlation |
| PCYT1A | -0.21484 | 9.96E-05 | Other Negative Correlation |
| NAV2 | -0.21507 | 9.78E-05 | Other Negative Correlation |
| NFU1 | -0.21546 | 9.5E-05 | Other Negative Correlation |
| CCDC28B | -0.21548 | 9.48E-05 | Other Negative Correlation |
| ENSG00000203546 | -0.21619 | 8.97E-05 | Other Negative Correlation |
| GLYCTK | -0.21649 | 8.77E-05 | Other Negative Correlation |
| KIAA1217 | -0.21649 | 8.77E-05 | Other Negative Correlation |
| ATP5F1C | -0.21673 | 8.61E-05 | Other Negative Correlation |
| NDUFB4 | -0.21686 | 8.53E-05 | Other Negative Correlation |
| LTB | -0.21702 | 8.42E-05 | Other Negative Correlation |
| CCDC50 | -0.21717 | 8.32E-05 | Other Negative Correlation |
| ARPC3 | -0.21741 | 8.17E-05 | Other Negative Correlation |
| UXS1 | -0.21743 | 8.16E-05 | Other Negative Correlation |
| CAMTA1 | -0.21753 | 8.09E-05 | Other Negative Correlation |
| MATCAP1 | -0.21865 | 7.42E-05 | Other Negative Correlation |
| COX5A | -0.21889 | 7.28E-05 | Other Negative Correlation |
| INTS10 | -0.21942 | 6.99E-05 | Other Negative Correlation |
| MTCH2 | -0.21948 | 6.95E-05 | Other Negative Correlation |
| ATP5PO | -0.21977 | 6.8E-05 | Other Negative Correlation |
| FNDC3B | -0.22009 | 6.63E-05 | Other Negative Correlation |
| FADS3 | -0.22041 | 6.46E-05 | Other Negative Correlation |
| ARL6IP5 | -0.22053 | 6.4E-05 | Other Negative Correlation |
| PRKCI | -0.22071 | 6.32E-05 | Other Negative Correlation |
| AQR | -0.22093 | 6.21E-05 | Other Negative Correlation |
| CPSF1 | -0.22113 | 6.11E-05 | Other Negative Correlation |
| HEXA | -0.22119 | 6.08E-05 | Other Negative Correlation |
| PLEKHA5 | -0.22271 | 5.39E-05 | Other Negative Correlation |
| PRKCSH | -0.22283 | 5.34E-05 | Other Negative Correlation |
| NXPE3 | -0.22333 | 5.13E-05 | Other Negative Correlation |
| ZDHHC24 | -0.22367 | 5E-05 | Other Negative Correlation |
| PAPSS1 | -0.22383 | 4.93E-05 | Other Negative Correlation |
| FAAP20 | -0.2246 | 4.64E-05 | Other Negative Correlation |
| RNF115 | -0.22501 | 4.49E-05 | Other Negative Correlation |

| | | | |
|---|---|---|---|
| FLOT2 | -0.22514 | 4.44E-05 | Other Negative Correlation |
| MDH2 | -0.22534 | 4.37E-05 | Other Negative Correlation |
| CPNE8 | -0.22568 | 4.26E-05 | Other Negative Correlation |
| ZNF343 | -0.22584 | 4.2E-05 | Other Negative Correlation |
| CBY1 | -0.22621 | 4.08E-05 | Other Negative Correlation |
| ALKBH7 | -0.2265 | 3.98E-05 | Other Negative Correlation |
| CPLANE1 | -0.22658 | 3.96E-05 | Other Negative Correlation |
| AP2B1 | -0.2274 | 3.71E-05 | Other Negative Correlation |
| MTHFR | -0.22749 | 3.68E-05 | Other Negative Correlation |
| COX7A2 | -0.22922 | 3.19E-05 | Other Negative Correlation |
| ZNF385A | -0.22926 | 3.18E-05 | Other Negative Correlation |
| MRPL33 | -0.22931 | 3.17E-05 | Other Negative Correlation |
| AKR1C1 | -0.22942 | 3.14E-05 | Other Negative Correlation |
| PHLDA2 | -0.22984 | 3.04E-05 | Other Negative Correlation |
| ITGAV | -0.23024 | 2.94E-05 | Other Negative Correlation |
| MIF4GD | -0.23107 | 2.74E-05 | Other Negative Correlation |
| NDUFB11 | -0.23108 | 2.74E-05 | Other Negative Correlation |
| GATD3 | -0.23118 | 2.72E-05 | Other Negative Correlation |
| ETFA | -0.23265 | 2.41E-05 | Other Negative Correlation |
| CD9 | -0.23273 | 2.39E-05 | Other Negative Correlation |
| ABL2 | -0.23286 | 2.37E-05 | Other Negative Correlation |
| OPTN | -0.23306 | 2.33E-05 | Other Negative Correlation |
| RHOBTB2 | -0.23313 | 2.32E-05 | Other Negative Correlation |
| RAB27B | -0.23376 | 2.2E-05 | Other Negative Correlation |
| LGALS7B | -0.23435 | 2.09E-05 | Other Negative Correlation |
| PDXK | -0.23495 | 1.99E-05 | Other Negative Correlation |
| RAPGEFL1 | -0.23528 | 1.94E-05 | Other Negative Correlation |
| RBFOX2 | -0.23664 | 1.73E-05 | Other Negative Correlation |
| TFG | -0.23708 | 1.66E-05 | Other Negative Correlation |
| PPP1R14A | -0.23723 | 1.64E-05 | Other Negative Correlation |
| CXorf38 | -0.23761 | 1.59E-05 | Other Negative Correlation |
| LAMC1 | -0.23788 | 1.56E-05 | Other Negative Correlation |
| ECH1 | -0.2385 | 1.47E-05 | Other Negative Correlation |
| CEP170 | -0.23878 | 1.44E-05 | Other Negative Correlation |
| ANK3 | -0.23914 | 1.4E-05 | Other Negative Correlation |
| BIRC6 | -0.23919 | 1.39E-05 | Other Negative Correlation |
| DMAC1 | -0.24056 | 1.24E-05 | Other Negative Correlation |
| SMARCA4 | -0.24133 | 1.16E-05 | Other Negative Correlation |
| CUEDC2 | -0.2423 | 1.06E-05 | Other Negative Correlation |
| SERPINB7 | -0.24268 | 1.03E-05 | Other Negative Correlation |
| TOM1L1 | -0.24282 | 1.02E-05 | Other Negative Correlation |
| MYO18A | -0.24332 | 9.75E-06 | Other Negative Correlation |
| PLEKHA1 | -0.24339 | 9.69E-06 | Other Negative Correlation |
| MRPS21 | -0.24364 | 9.48E-06 | Other Negative Correlation |
| ADCY4 | -0.24383 | 9.33E-06 | Other Negative Correlation |
| ACSL3 | -0.24564 | 7.96E-06 | Other Negative Correlation |
| MMP24OS | -0.24606 | 7.67E-06 | Other Negative Correlation |
| CYB5R3 | -0.24655 | 7.35E-06 | Other Negative Correlation |

| | | | |
|---|---|---|---|
| FKBP9 | -0.24752 | 6.75E-06 | Other Negative Correlation |
| NDUFS2 | -0.24764 | 6.68E-06 | Other Negative Correlation |
| CKMT1A | -0.24785 | 6.55E-06 | Other Negative Correlation |
| SMIM30 | -0.24806 | 6.43E-06 | Other Negative Correlation |
| ERGIC3 | -0.24885 | 6E-06 | Other Negative Correlation |
| DBT | -0.24929 | 5.77E-06 | Other Negative Correlation |
| ATP5MC1 | -0.24983 | 5.5E-06 | Other Negative Correlation |
| KRTCAP2 | -0.24986 | 5.48E-06 | Other Negative Correlation |
| UFL1 | -0.2512 | 4.86E-06 | Other Negative Correlation |
| REEP2 | -0.25188 | 4.57E-06 | Other Negative Correlation |
| NDUFB2 | -0.25228 | 4.41E-06 | Other Negative Correlation |
| MSRB2 | -0.25317 | 4.07E-06 | Other Negative Correlation |
| TLN1 | -0.25403 | 3.77E-06 | Other Negative Correlation |
| TBC1D32 | -0.25448 | 3.61E-06 | Other Negative Correlation |
| ATP5MG | -0.25542 | 3.32E-06 | Other Negative Correlation |
| KDM5B | -0.25576 | 3.22E-06 | Other Negative Correlation |
| SH3D19 | -0.25579 | 3.21E-06 | Other Negative Correlation |
| RAB25 | -0.25601 | 3.14E-06 | Other Negative Correlation |
| C5orf24 | -0.25603 | 3.14E-06 | Other Negative Correlation |
| ADAR | -0.25605 | 3.13E-06 | Other Negative Correlation |
| SH3BGRL | -0.25627 | 3.07E-06 | Other Negative Correlation |
| ZMYND11 | -0.25798 | 2.62E-06 | Other Negative Correlation |
| SLC44A1 | -0.25799 | 2.62E-06 | Other Negative Correlation |
| PDCD4 | -0.25814 | 2.59E-06 | Other Negative Correlation |
| CLASP1 | -0.25898 | 2.39E-06 | Other Negative Correlation |
| RNF144B | -0.25922 | 2.34E-06 | Other Negative Correlation |
| SPTLC1 | -0.2594 | 2.3E-06 | Other Negative Correlation |
| RAB11B | -0.25943 | 2.29E-06 | Other Negative Correlation |
| TRIQK | -0.25958 | 2.26E-06 | Other Negative Correlation |
| SDHC | -0.25959 | 2.26E-06 | Other Negative Correlation |
| SPARC | -0.26007 | 2.16E-06 | Other Negative Correlation |
| GGACT | -0.26051 | 2.07E-06 | Other Negative Correlation |
| SERF2 | -0.26069 | 2.04E-06 | Other Negative Correlation |
| R3HDM2 | -0.26275 | 1.68E-06 | Other Negative Correlation |
| ATP5F1B | -0.26335 | 1.59E-06 | Other Negative Correlation |
| EIF4A2 | -0.26415 | 1.47E-06 | Other Negative Correlation |
| ATP1B1 | -0.26451 | 1.42E-06 | Other Negative Correlation |
| CCDC90B | -0.26565 | 1.28E-06 | Other Negative Correlation |
| IL32 | -0.26606 | 1.23E-06 | Other Negative Correlation |
| NAA80 | -0.26682 | 1.14E-06 | Other Negative Correlation |
| TGFB1I1 | -0.26715 | 1.11E-06 | Other Negative Correlation |
| PRKAR1A | -0.26891 | 9.35E-07 | Other Negative Correlation |
| NDUFA5 | -0.26912 | 9.17E-07 | Other Negative Correlation |
| CCDC9B | -0.26959 | 8.76E-07 | Other Negative Correlation |
| STAP2 | -0.27139 | 7.35E-07 | Other Negative Correlation |
| FUOM | -0.27256 | 6.56E-07 | Other Negative Correlation |
| GLTP | -0.27393 | 5.73E-07 | Other Negative Correlation |
| PPA2 | -0.275 | 5.16E-07 | Other Negative Correlation |

| | | | |
|---|---|---|---|
| IFI27L2 | -0.27652 | 4.44E-07 | Other Negative Correlation |
| LY6E | -0.27685 | 4.29E-07 | Other Negative Correlation |
| KRT8 | -0.27814 | 3.78E-07 | Other Negative Correlation |
| TCEAL8 | -0.27821 | 3.75E-07 | Other Negative Correlation |
| PDLIM4 | -0.27985 | 3.18E-07 | Other Negative Correlation |
| ATP5F1A | -0.28042 | 3E-07 | Other Negative Correlation |
| LMAN1 | -0.28166 | 2.65E-07 | Other Negative Correlation |
| TMEM134 | -0.28169 | 2.64E-07 | Other Negative Correlation |
| SF3B6 | -0.28239 | 2.46E-07 | Other Negative Correlation |
| CCNI | -0.28439 | 2E-07 | Other Negative Correlation |
| CYFIP1 | -0.28512 | 1.86E-07 | Other Negative Correlation |
| SEC14L2 | -0.28523 | 1.84E-07 | Other Negative Correlation |
| ARF5 | -0.28724 | 1.49E-07 | Other Negative Correlation |
| PLCB4 | -0.28829 | 1.34E-07 | Other Negative Correlation |
| DSG2 | -0.28994 | 1.13E-07 | Other Negative Correlation |
| SAA1 | -0.29084 | 1.03E-07 | Other Negative Correlation |
| RINL | -0.29151 | 9.55E-08 | Other Negative Correlation |
| NDUFA12 | -0.29198 | 9.08E-08 | Other Negative Correlation |
| VAMP8 | -0.29213 | 8.95E-08 | Other Negative Correlation |
| UQCRC1 | -0.29531 | 6.38E-08 | Other Negative Correlation |
| CASP4 | -0.29648 | 5.63E-08 | Other Negative Correlation |
| COX7C | -0.29719 | 5.21E-08 | Other Negative Correlation |
| TSPAN1 | -0.29742 | 5.09E-08 | Other Negative Correlation |
| ACSS2 | -0.30006 | 3.82E-08 | Moderate Negative Correlation |
| KLHL5 | -0.30029 | 3.72E-08 | Moderate Negative Correlation |
| NDUFS4 | -0.30072 | 3.55E-08 | Moderate Negative Correlation |
| SUCLG2 | -0.30416 | 2.43E-08 | Moderate Negative Correlation |
| CMTM6 | -0.30436 | 2.38E-08 | Moderate Negative Correlation |
| IQGAP1 | -0.30719 | 1.74E-08 | Moderate Negative Correlation |
| ATOX1 | -0.30941 | 1.35E-08 | Moderate Negative Correlation |
| MRPL45 | -0.31449 | 7.59E-09 | Moderate Negative Correlation |
| PPFIBP1 | -0.31489 | 7.25E-09 | Moderate Negative Correlation |
| APLP2 | -0.31877 | 4.62E-09 | Moderate Negative Correlation |
| SORL1 | -0.31896 | 4.51E-09 | Moderate Negative Correlation |
| ATP5ME | -0.3193 | 4.34E-09 | Moderate Negative Correlation |
| PYCARD | -0.32459 | 2.32E-09 | Moderate Negative Correlation |
| CCDC25 | -0.32498 | 2.22E-09 | Moderate Negative Correlation |
| OST4 | -0.32647 | 1.85E-09 | Moderate Negative Correlation |
| FSTL1 | -0.32828 | 1.49E-09 | Moderate Negative Correlation |
| HEBP2 | -0.32978 | 1.24E-09 | Moderate Negative Correlation |
| TAPBP | -0.33509 | 6.47E-10 | Moderate Negative Correlation |
| PSME2 | -0.33787 | 4.58E-10 | Moderate Negative Correlation |
| SNX6 | -0.34297 | 2.4E-10 | Moderate Negative Correlation |
| LGALS1 | -0.34919 | 1.08E-10 | Moderate Negative Correlation |
| UQCRB | -0.35579 | 4.51E-11 | Moderate Negative Correlation |
| DECR1 | -0.36715 | 9.61E-12 | Moderate Negative Correlation |
| PRSS12 | -0.37214 | 4.78E-12 | Moderate Negative Correlation |
| RAC2 | -0.38499 | 7.47E-13 | Moderate Negative Correlation |

| | | | |
|---|---|---|---|
| TMEM256 | -0.38737 | 5.25E-13 | Moderate Negative Correlation |
| LGALS3 | -0.40348 | 4.47E-14 | Moderate Negative Correlation |
| FDPS | -0.4072 | 2.48E-14 | Moderate Negative Correlation |
| EML2 | -0.42187 | 2.27E-15 | Moderate Negative Correlation |
| SAT1 | -0.4455 | 3.74E-17 | Moderate Negative Correlation |

**Table Legend**

| Column Name |
|---|
| Gene name |
| R |
| P |
| Correlation Strength |

| Description |
| --- |
| Official gene symbol standardized by HGNC (HUGO Gene Nomenclature Committee) |
| Pearson correlation coefficient: indicating relationship strength between gene expression among selected cells and selected |
| P-value: statistical significance of correlation (values < 0.05 typically considered statistically significant) |
| Strength of correlation based on absolute R-value (ignores direction). Categories: **Strong** (\|R\| ≥ 0.7), **Moderate** (0.3 ≤ \|R\| < [illegible] |
