## Supplemental Table 5 for "Human Cytomegalovirus Infection of Primary Human Oral Keratinocytes Induces Intermediate Keratinocyte Differentiation and an Altered Innate Immune Response"

| Term | ES | NES | NOM p-val | FDR q-val | FWER p-va | Tag % |
|---|---|---|---|---|---|---|
| **E2F Targets** | 0.735796 | 2.555114 | 0 | 0 | 0 | 43/65 |
| **Oxidative Phosphorylation** | -0.50684 | -2.51674 | 0 | 0 | 0 | 59/149 |
| G2-M Checkpoint | 0.576907 | 2.04058 | 0 | 0 | 0 | 28/68 |
| UV Response Up | 0.545492 | 2.035862 | 0 | 0 | 0 | 57/110 |
| **Adipogenesis** | -0.37742 | -1.85439 | 0 | 0.01214 | 0.01 | 44/142 |
| Spermatogenesis | 0.48954 | 1.776087 | 0 | 0.002514 | 0.014 | 27/81 |
| Hedgehog Signaling | 0.577883 | 1.668692 | 0.002725 | 0.01135 | 0.074 | 14/25 |
| Unfolded Protein Response | 0.48213 | 1.659274 | 0.002484 | 0.010416 | 0.08 | 21/60 |
| **Interferon Alpha Response** | -0.40841 | -1.64952 | 0 | 0.048561 | 0.058 | 20/52 |
| KRAS Signaling Dn | 0.434458 | 1.639683 | 0.001127 | 0.011186 | 0.099 | 43/124 |
| Estrogen Response Early | 0.421015 | 1.615754 | 0.001105 | 0.012751 | 0.123 | 47/142 |
| **Fatty Acid Metabolism** | -0.35126 | -1.61413 | 0 | 0.045526 | 0.07 | 24/95 |
| TGF-beta Signaling | -0.44414 | -1.61342 | 0.007353 | 0.036907 | 0.071 | 11/31 |
| **Interferon Gamma Response** | -0.34228 | -1.59377 | 0 | 0.033588 | 0.078 | 41/112 |
| Protein Secretion | -0.3457 | -1.50083 | 0 | 0.052724 | 0.139 | 15/74 |
| Estrogen Response Late | 0.388796 | 1.493839 | 0.005562 | 0.044856 | 0.415 | 49/139 |
| mTORC1 Signaling | 0.401337 | 1.457901 | 0.012791 | 0.058401 | 0.527 | 28/90 |
| Androgen Response | -0.34277 | -1.42236 | 0.013953 | 0.080733 | 0.232 | 9/55 |
| Hypoxia | 0.369395 | 1.39474 | 0.014428 | 0.098151 | 0.753 | 29/128 |
| IL-2/STAT5 Signaling | 0.361377 | 1.393248 | 0.011186 | 0.091708 | 0.755 | 28/133 |
| Xenobiotic Metabolism | 0.364365 | 1.348591 | 0.039728 | 0.127367 | 0.885 | 28/112 |
| Pperoxisome | -0.30931 | -1.3391 | 0.037736 | 0.12518 | 0.366 | 14/74 |
| Apoptosis | 0.358407 | 1.32627 | 0.048276 | 0.142642 | 0.933 | 28/98 |
| Cholesterol Homeostasis | -0.35953 | -1.32028 | 0.066372 | 0.12456 | 0.391 | 8/37 |
| Myogenesis | 0.346736 | 1.313224 | 0.047404 | 0.14793 | 0.954 | 41/130 |
| Mitotic Spindle | -0.26267 | -1.25793 | 0.033333 | 0.175483 | 0.545 | 35/124 |
| DNA Repair | 0.345852 | 1.247324 | 0.121354 | 0.232787 | 0.993 | 17/86 |
| Glycolysis | 0.328592 | 1.22455 | 0.124722 | 0.258053 | 0.995 | 35/120 |
| Notch Signaling | 0.42697 | 1.202662 | 0.220867 | 0.282427 | 0.998 | 9/22 |
| Reactive Oxygen Species Pathway | -0.31315 | -1.18952 | 0.1625 | 0.24564 | 0.691 | 8/33 |
| Myc Targets V2 | 0.572801 | 1.186466 | 0.256677 | 0.296334 | 1 | 3/7 |
| heme Metabolism | 0.304913 | 1.179286 | 0.169546 | 0.294447 | 1 | 44/151 |
| TNF-alpha Signaling via NF-kB | -0.25518 | -1.17169 | 0.116667 | 0.256441 | 0.733 | 16/105 |
| p53 Pathway | -0.24589 | -1.14898 | 0.15873 | 0.272811 | 0.779 | 35/103 |
| Myc Targets V1 | -0.32429 | -1.09603 | 0.319588 | 0.360325 | 0.883 | 7/24 |
| KRAS Signaling Up | 0.277844 | 1.066516 | 0.330744 | 0.509713 | 1 | 48/138 |
| Coagulation | -0.23904 | -1.05538 | 0.309942 | 0.419449 | 0.937 | 18/78 |
| Inflammatory Response | -0.21729 | -1.05036 | 0.342105 | 0.404059 | 0.942 | 22/129 |
| Angiogenesis | -0.33394 | -1.01405 | 0.414493 | 0.468482 | 0.964 | 3/16 |
| Allograft Rejection | -0.21136 | -0.99277 | 0.471074 | 0.492642 | 0.976 | 27/109 |
| Apical Junction | 0.259768 | 0.977937 | 0.518313 | 0.705115 | 1 | 33/127 |
| Pancreas Beta Cells | 0.341198 | 0.953461 | 0.54073 | 0.734549 | 1 | 13/23 |
| PI3K/AKT/mTOR Signaling | 0.271076 | 0.952855 | 0.551095 | 0.70514 | 1 | 14/70 |

| | | | | | | |
|---|---|---|---|---|---|---|
| Apical Surface | 0.317255 | 0.942788 | 0.555556 | 0.699835 | 1 | 4/29 |
| Complement | 0.241126 | 0.900346 | 0.706621 | 0.761219 | 1 | 49/123 |
| UV Response Dn | 0.242977 | 0.887723 | 0.684211 | 0.759418 | 1 | 34/95 |
| Epithelial Mesenchymal Transitior | 0.229748 | 0.863642 | 0.729025 | 0.777782 | 1 | 20/108 |
| Bile Acid Metabolism | -0.18215 | -0.7934 | 0.903448 | 0.878597 | 1 | 9/79 |
| Wnt-beta Catenin Signaling | 0.236329 | 0.705263 | 0.896317 | 0.967529 | 1 | 9/28 |
| IL-6/JAK/STAT3 Signaling | 0.200871 | 0.675599 | 0.941772 | 0.957019 | 1 | 18/53 |

| Gene % |
| --- |
| 16.24% |
| 11.52% |
| 13.45% |
| 24.05% |
| 12.54% |
| 12.48% |
| 24.94% |
| 12.32% |
| 19.74% |
| 20.37% |
| 16.31% |
| 8.17% |
| 13.56% |
| 22.29% |
| 8.39% |
| 20.32% |
| 15.07% |
| 3.81% |
| 13.74% |
| 12.67% |
| 15.53% |
| 7.69% |
| 15.50% |
| 5.09% |
| 21.11% |
| 15.17% |
| 14.18% |
| 20.41% |
| 21.76% |
| 8.22% |
| 22.28% |
| 22.70% |
| 7.57% |
| 19.00% |
| 11.05% |
| 28.28% |
| 17.67% |
| 11.74% |
| 7.94% |
| 19.58% |
| 20.10% |
| 41.30% |
| 13.74% |

4.21%
33.03%
27.66%
14.88%
7.72%
29.58%
33.92%

| Lead_genes |
| --- |
| UBE2S;UBE2T;TK1;ATAD2;TCF19;BARD1;ORC6;RPA2;POP7;AURKA;MCM5;DSCC1;POLA2;ASF1B;BRCA1;SMC3;KIF |
| DECR1;UQCRB;ATP5ME;NDUFS4;COX7C;UQCRC1;ATP5F1A;NDUFA5;ATP1B1;ATP5F1B;SDHC;ATP5MG;NDUFB2;A |
| UBE2S;E2F1;CDC6;BARD1;ORC6;RPA2;AURKA;MCM5;POLA2;KIF22;SAP30;CHAF1A;SLC7A1;CDKN2C;STIL;MCM3; |
| ENO2;MAPK8IP2;FEN1;SLC25A4;NPTX2;GAL;CCK;HSPA13;KCNH2;ATF3;NPTXR;EPCAM;EIF5;RFC4;MSX1;CEBPG;T |
| DECR1;APLP2;UQCRC1;NDUFA5;SDHC;PDCD4;DBT;ECH1;MDH2;ATP5PO;MTCH2;SCP2;IFNGR1;NDUFAB1;PGM1; |
| YBX2;PCSK1N;AURKA;NEFH;STRBP;MLF1;SHE;TNNI3;RFC4;SEPTIN4;RPL39L;GSTM3;ACE;CLGN;MAST2;CNIH2;LDH |
| VLDLR;AMOT;CNTFR;L1CAM;CRMP1;HEY2;TLE3;THY1;SHH;DPYSL2;GLI1;NKX6-1;PTCH1;ACHE |
| DNAJB9;DNAJC3;CHAC1;CEBPB;STC2;PREB;WFS1;ATF3;MTHFD2;SLC1A4;CEBPG;HSPA9;HERPUD1;WIPI1;DCP2;SI |
| PSME2;LY6E;ADAR;UBA7;ELF1;SP110;HELZ2;UBE2L6;NUB1;TAP1;STAT2;CD74;WARS1;IRF2;IFI30;TRIM21;RTP4;T |
| YBX2;PDK2;BARD1;EFHD1;CALCB;GPRC5C;CAMK1D;TNNI3;SLC38A3;ADRA2C;CPEB3;ARHGDIG;SLC30A3;SNN;YPE |
| TUBB2B;PLAAT3;CLDN7;SYNGR1;INHBB;MYBL1;PMAIP1;STC2;WFS1;OVOL2;ELF3;KCNK15;RAB17;NXT1;SLC1A4;H |
| DECR1;LGALS1;SUCLG2;SDHC;ECH1;MDH2;ACADVL;ALDH1A1;ECHS1;MGLL;ACAA2;ECI2;ACADM;SDHA;CBR1;UBE |
| PPP1CA;TJP1;SMAD3;CDH1;NCOR2;ACVR1;HDAC1;SKIL;ENG;MAP3K7;SMAD1 |
| PSME2;TAPBP;CASP4;VAMP8;LY6E;ADAR;MVP;NFKBIA;SP110;RNF213;HELZ2;CDKN1A;UBE2L6;ST3GAL5;SPPL2A; |
| LMAN1;ERGIC3;TOM1L1;AP2B1;SNAP23;AP3B1;DOP1A;RAB2A;COPB2;SOD1;ARFIP1;ARFGEF1;VPS4B;ATP1A1;IG |
| CKB;PLAAT3;CDC6;GAL;WFS1;OVOL2;NXT1;SLC1A4;HSPB8;TJP3;DNAJC1;LSR;PDLIM3;STIL;HR;KLK11;RNASEH2A; |
| SDF2L1;DDIT3;NIBAN1;VLDLR;AURKA;NFKBIB;TXNRD1;NFIL3;RPA1;MTHFD2;CTH;SLC1A4;HSPA9;TRIB3;SQSTM1; |
| SAT1;LMAN1;KRT8;ACSL3;ITGAV;PIAS1;NCOA4;TNFAIP8;TSC22D1 |
| ENO2;DDIT3;RORA;RRAGD;VLDLR;ADM;KLHL24;STC2;SDC2;ATF3;NFIL3;SAP30;NAGK;GALK1;HMOX1;CDKN1C;PL |
| GABARAPL1;TNFRSF18;RORA;CDC6;RRAGD;APLP1;NFIL3;SHE;COL6A1;BATF3;GUCY1B1;RGS16;CDKN1C;PLIN2;EN |
| HES6;GABARAPL1;APOE;CYP27A1;SAR1B;ETFDH;NDRG2;HMOX1;LPIN2;SMOX;LONP1;MAN1A1;GCH1;RBP4;CYB |
| FDPS;ECH1;SCP2;GSTK1;MVP;ALDH1A1;ECI2;SOD1;ACAA1;ACOX1;VPS4B;IDE;RDH11;PRDX5 |
| ENO2;TIMP2;DDIT3;DNAJC3;ERBB3;PMAIP1;NEFH;BRCA1;ATF3;CTH;HMOX1;SQSTM1;BGN;PPP2R5B;GCH1;CDC |
| FDPS;LGALS3;ACSS2;ECH1;CD9;FBXO6;PMVK;TP53INP1 |
| CKB;ERBB3;DES;STC2;COL6A2;GABARAPL2;KCNH2;HSPB8;REEP1;COX7A1;MB;SGCG;CRAT;ACSL1;NCAM1;AK1;AD |
| CLASP1;CAPZB;ABI1;SYNPO;NUMA1;TLK1;DLG1;BCR;PPP4R2;FGD4;NF1;CTTN;ARFGEF1;ARFIP2;RICTOR;ARHGAP |
| EIF1B;ZWINT;FEN1;STX3;RPA2;RAD51;ARL6IP1;POLA2;RFC4;RFC2;ERCC3;AK1;MPC2;POLD3;GPX4;SEC61A1;SNAP |
| ENO2;RRAGD;VLDLR;AURKA;STC2;GMPPB;SDC2;GFPT1;ELF3;HAX1;SAP30;CLDN3;CTH;GALK1;CHST1;GOT1;LDHC |
| SAP30;NOTCH3;PSENEN;DTX1;FBXW11;ARRB1;FZD5;HEYL;FZD7 |
| ATOX1;NDUFS2;NDUFB4;FTL;SOD1;ERCC2;GLRX;LSP1 |
| MCM5;FARSA;MPHOSPH10 |
| HAGH;GLRX5;GCLM;DMTN;FN3K;TRAK2;LPIN2;YPEL5;SMOX;MFHAS1;ANK1;PPP2R5B;BPGM;NFE2L1;CDR2;GYPC |
| SAT1;PHLDA2;PTPRE;MAP2K3;CCL20;HBEGF;TNFAIP8;TSC22D1;NFKBIA;CDKN1A;RELB;EHD1;NFKB2;SMAD3;SLC |
| SAT1;VAMP8;PTPRE;MXD4;FDXR;HBEGF;TSC22D1;BAIAP2;CDKN1A;TOB1;FOXO3;ELP1;LIF;CD82;PIDD1;CD81;TA |
| COX5A;NDUFAB1;VDAC3;PSMA4;PSMD7;SNRPB2;PSMD1 |
| PCSK1N;GADD45G;SCN1B;SNAP25;IGF2;FGF9;RGS16;GPRC5B;PEG3;BPGM;TNFRSF1B;ACE;RBP4;GYPC;SNAP91;C |
| SPARC;MSRB2;CD9;CRIP2;S100A13;MMP9;C3;APOC1;BMP1;SERPINA1;SIRT2;CTSO;CTSK;CTSB;PECAM1;MMP2;T |
| TAPBP;LY6E;RNF144B;PTPRE;TPBG;ABI1;PTAFR;CCL20;HBEGF;GABBR1;NFKBIA;PSEN1;CDKN1A;ICAM4;PDPN;LIF; |
| FSTL1;ITGAV;LRPAP1 |
| TAPBP;LTB;IFNGR1;ABI1;MMP9;IL27RA;LIF;STAT4;TAP1;IFNAR2;CD74;CTSS;UBE2D1;TLR6;MAP3K7;WARS1;HLA- |
| PCDH1;CLDN7;NRTN;KCNH2;CLDN6;CRB3;PKD1;PPP2R2C;FLNC;SLC30A3;CLDN4;CRAT;PTEN;MADCAM1;PIK3CB; |
| SYT13;PAX6;GCK;LMO2;NKX6-1;AKT3;DPP4;MAFB;STXBP1;NKX2-2;INSM1;NEUROG3;PDX1 |
| DDIT3;E2F1;NFKBIB;PRKAA2;TRIB3;SQSTM1;PTEN;PLCB1;NGF;FGF22;CDKN1B;PAK4;ECSIT;CXCR4 |

BRCA1;NTNG1;MAL;GSTM3
TIMP2;CD55;CEBPB;KLK1;SPOCK2;LRP1;PRSS36;HSPA1A;HPCAL4;SERPING1;RCE1;CLU;CTSD;CTSL;BRPF3;KCNIP3;
MAP1B;VLDLR;SDC2;SLC7A1;NR1D2;SCN8A;SYNJ2;ICA1;FBLN5;PTEN;KALRN;SFMBT1;ATXN1;CDKN1B;CDON;CDC
ENO2;CRLF1;APLP1;COL6A2;ITGA5;MSX1;LRP1;CDH2;SNTB1;BGN;SGCG;FBLN5;FERMT2;MYL9;FBLN2;MATN3;LO
OPTN;SCP2;GSTK1;FDXR;ALDH1A1;PEX11G;SOD1;PRDX5;CROT
WNT6;RBPJ;HEY2;WNT1;FRAT1;WNT5B;PTCH1;AXIN2;FZD8
INHBE;HAX1;HMOX1;TNFRSF1B;CNTFR;IL17RA;CD14;GRB2;ACVRL1;LTBR;MAP3K8;ACVR1B;PIM1;SOCS1;IRF1;IL1

22;RPA1;USP1;MTHFD2;CENPM;RFC2;CDKN2C;PSIP1;MCM3;SLBP;CDC25A;TIMELESS;MRE11;RNASEH2A;DCK;C
TP5MC1;NDUFS2;CYB5R3;ECH1;ETFA;COX7A2;MDH2;ATP5PO;COX5A;NDUFB4;ATP5F1C;UQCRC2;NDUFA4;ATP

YRO3;HMOX1;STARD3;SQSTM1;GGH;NR4A1;CDKN1C;PDLIM3;KLHDC3;AGO2;LHX2;GCH1;OLFM1;RAB27A;HLA-
UQCR10;ECHS1;MGLL;COX7B;NDUFB7;ACAA2;SLC66A3;SOD1;ACADM;COX8A;SLC27A1;C3;SULT1A1;TOB1;ACO

L1;SNCB;KCNN1;MFSD6;CNTFR;OXT;IDUA;PNMT;CCDC106;FGF22;SHOX2;COPZ2;CCNA1;TLX1;HTR1D;FGF16;TE
HSPB8;REEP1;TJP3;TTC39A;PDLIM3;INPP5F;HR;DHRS3;ABCA3;OLFM1;RETREG1;ELOVL2;SLC27A2;XBP1;ADCY9;G

TNFSF10;STAT4;TAP1;STAT2;IFNAR2;CD74;CASP3;SLAMF7;WARS1;PTPN6;MYD88;IRF2;IFI30;TRIM21;RTP4;JAK

ABCA3;OLFM1;SLC27A2;CACNA2D2;CPE;XBP1;SCNN1A;SCUBE2;FAM102A;MAPT;SLC2A8;CCNA1;GLA;LARGE1;S

A;MPP2;FBP1;SLC35B1;PINK1;TMEM176B;GSTM4;GAD1;SLC46A3;GNMT;CYP2E1;ALAS1;EPHX1;CSAD

CY9;CACNA1H;CLU;LARGE1;ATP6AP1;TNNC1;PTP4A3;EFS;TNNC2;SORBS1;FXYD1;BIN1;FABP3;ITGA7;MRAS;EPH
29;NIN;DYNC1H1;ROCK1;PAFAH1B1;TUBGCP3;SORBS2;ARHGEF7;ARAP3;MYO9B;KIFAP3;DOCK2;MID1;RALBP1;

;CYB5A;IDUA;CACNA1H;PMM2;CHPF;EGLN3;SLC37A4;MERTK;ME1;CXCR4;CLDN9;GMPPA;DPYSL4;CAPN5;B3GA

;E2F2;PIGQ;FOXJ2;TAL1;CTNS;FECH;FBXO34;ADD2;EPOR;UCP2;SYNJ1;NARF;LMO2;TENT5C;SLC6A9;MOSPD1;TR

P1;WWP1;NUPR1;ABAT;TNFSF9;PLXNB2;RB1;ERCC5;VWA5A;CYFIP2;HEXIM1;IP6K2;DCXR;ANKRA2;IFI30;TM4SF1

PE;NGF;KIF5C;TMEM176B;TMEM176A;ALDH1A2;EPB41L3;GFPT2;MMP11;CXCR4;ETV5;FLT4;IKZF1;TMEM158;C

GRB7;AMIGO1;MYL9;ATP1A3;CDH15;NECTIN3;THY1;ADRA1B;CLDN9;ARHGEF6;WASL;TAOK2;LAYN;MDK;CADM

;MMP15;GCA;ME1;FDX1;ADRA2B;VCPIP1;CA2;CTSH;MSRB1;GRB2;RASGRP1;LGMN;RNF4;CASP9;CTSV;CPQ;USP[illegible]
[illegible]42BPA;PRKAR2B;DLC1;TGFBR3;MAGI2;FZD2;DDAH1;MIOS;COL1A1;PDGFRB;DBP;PRKCE;AKT3;PIK3R3;PPARG;IG

5MF;ACADVL;ATP5PD;COX6B1;NDUFAB1;UQCR10;ECHS1;ATP6V0C;COX7B;NDUFB7;ACAA2;ACADM;COX8A;SDH

F;DNAJB1;WIZ;SHOX2;NKX2-5;BCL2L11;HSPA2;SPR;CHKA;ATP6V1F;ALAS1;YKT6;EPHX1;CCNE1;PLCL1;CA2;CREG1

FRA1;GAB2;SNX24;SCNN1A;FAM102A;MAPT;AR;MUC1;GLA;SLC22A5;PODXL;DLC1;SIAH2;TFAP2C;SLC1A1;ADCY

LC22A5;CHST8;HOMER2;PRKAR2B;TNNC1;NMU;SIAH2;TFAP2C;EMP2;LLGL2;CHPT1;KCNK5;METTL3;CA2;MDK;M

A2;MMD;CCSER2;PRRX1;ZNF639;BMP2;IL1RL2;WDR33;HOXD11;ADGRA2;SLPI;ETV1;ANKH;CDADC1;TSPAN13;AK

HA;COX4I1;VDAC3;ACAA1;OGDH;ABCB7;MRPS15;IDH3B;SDHB;NDUFC1;PDHB;UQCRQ;NDUFB5;HSD17B10;NDUF

[illegible]S8;OXA1L;ETFB;NDUFB3;NDUFA7;ALDH6A1
