## Supplemental Table 6 for "Human Cytomegalovirus Infection of Primary Human Oral Keratinocytes Induces Intermediate Keratinocyte Differentiation and an Altered Innate Immune Response"

| Gene name | R | P | Correlation Strength |
| --- | --- | --- | --- |
| DDIT3 | 0.641546 | 1.45E-34 | Strong Positive Correlation |
| UCHL1 | 0.633194 | 1.87E-33 | Strong Positive Correlation |
| TUBB2B | 0.628571 | 7.47E-33 | Strong Positive Correlation |
| HES6 | 0.604573 | 6.84E-30 | Strong Positive Correlation |
| RNF157 | 0.60052 | 2.05E-29 | Strong Positive Correlation |
| STC2 | 0.548738 | 6.88E-24 | Strong Positive Correlation |
| KIF1A | 0.526945 | 7.76E-22 | Strong Positive Correlation |
| ENO2 | 0.524591 | 1.27E-21 | Strong Positive Correlation |
| TCF15 | 0.523662 | 1.53E-21 | Strong Positive Correlation |
| GCLM | 0.523374 | 1.63E-21 | Strong Positive Correlation |
| PCK2 | 0.522358 | 2.01E-21 | Strong Positive Correlation |
| ATF3 | 0.511541 | 1.79E-20 | Strong Positive Correlation |
| NIBAN1 | 0.508236 | 3.44E-20 | Strong Positive Correlation |
| ZFR2 | 0.5082 | 3.47E-20 | Strong Positive Correlation |
| TRIB3 | 0.503674 | 8.38E-20 | Strong Positive Correlation |
| CD55 | 0.477446 | 1.09E-17 | Moderate Positive Correlation |
| HERC5 | 0.47 | 4.02E-17 | Moderate Positive Correlation |
| SDF2L1 | 0.469272 | 4.56E-17 | Moderate Positive Correlation |
| EIF1B | 0.466253 | 7.66E-17 | Moderate Positive Correlation |
| NMRK2 | 0.462581 | 1.43E-16 | Moderate Positive Correlation |
| PLAAT3 | 0.456181 | 4.18E-16 | Moderate Positive Correlation |
| RHEBL1 | 0.455711 | 4.52E-16 | Moderate Positive Correlation |
| EEF1A2 | 0.45524 | 4.89E-16 | Moderate Positive Correlation |
| C1QL4 | 0.452375 | 7.83E-16 | Moderate Positive Correlation |
| MEIOC | 0.449783 | 1.19E-15 | Moderate Positive Correlation |
| CALCB | 0.445348 | 2.44E-15 | Moderate Positive Correlation |
| YBX2 | 0.440189 | 5.53E-15 | Moderate Positive Correlation |
| MTHFD2 | 0.436164 | 1.04E-14 | Moderate Positive Correlation |
| HMX1 | 0.435113 | 1.22E-14 | Moderate Positive Correlation |
| CTH | 0.429153 | 3.04E-14 | Moderate Positive Correlation |
| CHAC1 | 0.426275 | 4.69E-14 | Moderate Positive Correlation |
| MAPK8IP2 | 0.420131 | 1.17E-13 | Moderate Positive Correlation |
| MMP23B | 0.417011 | 1.85E-13 | Moderate Positive Correlation |
| RASGRP2 | 0.410682 | 4.6E-13 | Moderate Positive Correlation |
| RBP7 | 0.407835 | 6.89E-13 | Moderate Positive Correlation |
| GARS1 | 0.407358 | 7.37E-13 | Moderate Positive Correlation |
| TAC1 | 0.403522 | 1.26E-12 | Moderate Positive Correlation |
| ASRGL1 | 0.401035 | 1.78E-12 | Moderate Positive Correlation |
| ADM | 0.400605 | 1.89E-12 | Moderate Positive Correlation |
| C1QL1 | 0.397107 | 3.05E-12 | Moderate Positive Correlation |
| CENPF | 0.390024 | 7.93E-12 | Moderate Positive Correlation |
| NRIP3 | 0.386445 | 1.27E-11 | Moderate Positive Correlation |
| ADM2 | 0.385979 | 1.35E-11 | Moderate Positive Correlation |
| NPTX2 | 0.382726 | 2.07E-11 | Moderate Positive Correlation |
| PPP1R1A | 0.379745 | 3.04E-11 | Moderate Positive Correlation |

| | | | |
|---|---|---|---|
| DES | 0.378366 | 3.63E-11 | Moderate Positive Correlation |
| SHISA8 | 0.377053 | 4.29E-11 | Moderate Positive Correlation |
| GATA2 | 0.375848 | 5E-11 | Moderate Positive Correlation |
| FBLL1 | 0.37399 | 6.33E-11 | Moderate Positive Correlation |
| KLF15 | 0.373971 | 6.34E-11 | Moderate Positive Correlation |
| SHE | 0.36922 | 1.15E-10 | Moderate Positive Correlation |
| SKAP1 | 0.368423 | 1.27E-10 | Moderate Positive Correlation |
| SLC10A4 | 0.367568 | 1.41E-10 | Moderate Positive Correlation |
| UBE2S | 0.366133 | 1.68E-10 | Moderate Positive Correlation |
| TNFRSF18 | 0.36607 | 1.69E-10 | Moderate Positive Correlation |
| TXNRD1 | 0.365167 | 1.89E-10 | Moderate Positive Correlation |
| DOC2B | 0.364795 | 1.98E-10 | Moderate Positive Correlation |
| CTSF | 0.362415 | 2.65E-10 | Moderate Positive Correlation |
| CBS | 0.361164 | 3.08E-10 | Moderate Positive Correlation |
| CRLF1 | 0.360993 | 3.14E-10 | Moderate Positive Correlation |
| RAB39B | 0.358034 | 4.48E-10 | Moderate Positive Correlation |
| HSPA13 | 0.356757 | 5.21E-10 | Moderate Positive Correlation |
| ALDH1L2 | 0.356171 | 5.59E-10 | Moderate Positive Correlation |
| PCDH1 | 0.356117 | 5.63E-10 | Moderate Positive Correlation |
| SCN1B | 0.35355 | 7.62E-10 | Moderate Positive Correlation |
| MESP1 | 0.353466 | 7.69E-10 | Moderate Positive Correlation |
| GADD45G | 0.350086 | 1.14E-09 | Moderate Positive Correlation |
| B4GALNT1 | 0.348681 | 1.34E-09 | Moderate Positive Correlation |
| LMO4 | 0.347779 | 1.49E-09 | Moderate Positive Correlation |
| C1QL2 | 0.34661 | 1.7E-09 | Moderate Positive Correlation |
| LPL | 0.346379 | 1.75E-09 | Moderate Positive Correlation |
| METTL4 | 0.346342 | 1.76E-09 | Moderate Positive Correlation |
| TMEM59L | 0.346222 | 1.78E-09 | Moderate Positive Correlation |
| TCEAL2 | 0.345554 | 1.92E-09 | Moderate Positive Correlation |
| SMCO4 | 0.345078 | 2.03E-09 | Moderate Positive Correlation |
| TK1 | 0.337483 | 4.77E-09 | Moderate Positive Correlation |
| ZAR1 | 0.336667 | 5.22E-09 | Moderate Positive Correlation |
| SATL1 | 0.336515 | 5.31E-09 | Moderate Positive Correlation |
| ENHO | 0.336252 | 5.47E-09 | Moderate Positive Correlation |
| ZFAND2A | 0.334596 | 6.56E-09 | Moderate Positive Correlation |
| TNRC6C | 0.332603 | 8.16E-09 | Moderate Positive Correlation |
| NXPH4 | 0.330588 | 1.02E-08 | Moderate Positive Correlation |
| PIEZO2 | 0.330013 | 1.08E-08 | Moderate Positive Correlation |
| MNX1 | 0.328826 | 1.23E-08 | Moderate Positive Correlation |
| RIC3 | 0.328689 | 1.25E-08 | Moderate Positive Correlation |
| DNAJB9 | 0.326922 | 1.51E-08 | Moderate Positive Correlation |
| E2F1 | 0.326723 | 1.54E-08 | Moderate Positive Correlation |
| BMP4 | 0.324608 | 1.93E-08 | Moderate Positive Correlation |
| TLCD4 | 0.32374 | 2.11E-08 | Moderate Positive Correlation |
| HMX3 | 0.320818 | 2.87E-08 | Moderate Positive Correlation |
| HRK | 0.318341 | 3.71E-08 | Moderate Positive Correlation |

| | | | |
|---|---|---|---|
| TLX3 | 0.315881 | 4.79E-08 | Moderate Positive Correlation |
| GARIN5A | 0.315852 | 4.8E-08 | Moderate Positive Correlation |
| WT1 | 0.315493 | 4.98E-08 | Moderate Positive Correlation |
| SEPTIN3 | 0.31398 | 5.81E-08 | Moderate Positive Correlation |
| RCAN2 | 0.312197 | 6.97E-08 | Moderate Positive Correlation |
| TMEM151A | 0.312196 | 6.97E-08 | Moderate Positive Correlation |
| RPP25 | 0.310813 | 8.01E-08 | Moderate Positive Correlation |
| CRYBA2 | 0.310515 | 8.26E-08 | Moderate Positive Correlation |
| SDC2 | 0.309879 | 8.8E-08 | Moderate Positive Correlation |
| DOK5 | 0.308898 | 9.71E-08 | Moderate Positive Correlation |
| INHBE | 0.307471 | 1.12E-07 | Moderate Positive Correlation |
| COLEC12 | 0.307218 | 1.15E-07 | Moderate Positive Correlation |
| C1orf109 | 0.306005 | 1.3E-07 | Moderate Positive Correlation |
| PKN1 | 0.305267 | 1.39E-07 | Moderate Positive Correlation |
| CDK2AP2 | 0.304474 | 1.51E-07 | Moderate Positive Correlation |
| AGT | 0.30377 | 1.61E-07 | Moderate Positive Correlation |
| MAP1B | 0.302517 | 1.82E-07 | Moderate Positive Correlation |
| TESC | 0.301347 | 2.05E-07 | Moderate Positive Correlation |
| RAB3IL1 | 0.30134 | 2.05E-07 | Moderate Positive Correlation |
| SLC7A1 | 0.301068 | 2.1E-07 | Moderate Positive Correlation |
| HOXD8 | 0.300968 | 2.12E-07 | Moderate Positive Correlation |
| KCNK12 | 0.300258 | 2.27E-07 | Moderate Positive Correlation |
| BATF3 | 0.299041 | 2.56E-07 | Other Positive Correlation |
| STAC2 | 0.298992 | 2.57E-07 | Other Positive Correlation |
| DIRAS1 | 0.296417 | 3.29E-07 | Other Positive Correlation |
| RTN4RL2 | 0.295788 | 3.49E-07 | Other Positive Correlation |
| TLX2 | 0.294905 | 3.8E-07 | Other Positive Correlation |
| GK | 0.293851 | 4.2E-07 | Other Positive Correlation |
| RNF175 | 0.292717 | 4.67E-07 | Other Positive Correlation |
| FGF21 | 0.292505 | 4.76E-07 | Other Positive Correlation |
| TMSB15A | 0.291721 | 5.13E-07 | Other Positive Correlation |
| LONRF3 | 0.291508 | 5.23E-07 | Other Positive Correlation |
| SNAP25 | 0.290992 | 5.49E-07 | Other Positive Correlation |
| FLOT1 | 0.28965 | 6.22E-07 | Other Positive Correlation |
| SESN2 | 0.289527 | 6.29E-07 | Other Positive Correlation |
| PDLIM3 | 0.289461 | 6.33E-07 | Other Positive Correlation |
| BARX1 | 0.288582 | 6.87E-07 | Other Positive Correlation |
| RNF212 | 0.288331 | 7.03E-07 | Other Positive Correlation |
| SEC11C | 0.288256 | 7.08E-07 | Other Positive Correlation |
| HAX1 | 0.288145 | 7.15E-07 | Other Positive Correlation |
| RIMKLA | 0.286659 | 8.2E-07 | Other Positive Correlation |
| SCN5A | 0.285638 | 9.01E-07 | Other Positive Correlation |
| NOS1AP | 0.284856 | 9.68E-07 | Other Positive Correlation |
| APLP1 | 0.284275 | 1.02E-06 | Other Positive Correlation |
| C1QTNF4 | 0.284013 | 1.04E-06 | Other Positive Correlation |
| DUSP26 | 0.283564 | 1.09E-06 | Other Positive Correlation |

| | | | |
|---|---|---|---|
| CFAP43 | 0.28187 | 1.27E-06 | Other Positive Correlation |
| SGO2 | 0.28174 | 1.28E-06 | Other Positive Correlation |
| ARHGAP9 | 0.281613 | 1.3E-06 | Other Positive Correlation |
| MARS1 | 0.28119 | 1.35E-06 | Other Positive Correlation |
| COX7A1 | 0.277819 | 1.82E-06 | Other Positive Correlation |
| RHBDL1 | 0.276239 | 2.1E-06 | Other Positive Correlation |
| ECEL1 | 0.276163 | 2.11E-06 | Other Positive Correlation |
| EFHD1 | 0.274933 | 2.35E-06 | Other Positive Correlation |
| CAMK2N2 | 0.274876 | 2.36E-06 | Other Positive Correlation |
| CHST1 | 0.273957 | 2.56E-06 | Other Positive Correlation |
| TIMP4 | 0.272784 | 2.84E-06 | Other Positive Correlation |
| DERL3 | 0.272706 | 2.86E-06 | Other Positive Correlation |
| SLC1A4 | 0.272329 | 2.95E-06 | Other Positive Correlation |
| CRACD | 0.272209 | 2.99E-06 | Other Positive Correlation |
| SLC43A1 | 0.272157 | 3E-06 | Other Positive Correlation |
| RRAGD | 0.27131 | 3.23E-06 | Other Positive Correlation |
| NRGN | 0.271124 | 3.28E-06 | Other Positive Correlation |
| HOXD1 | 0.270684 | 3.41E-06 | Other Positive Correlation |
| COL6A2 | 0.26959 | 3.74E-06 | Other Positive Correlation |
| CPEB1 | 0.269023 | 3.93E-06 | Other Positive Correlation |
| VLDLR | 0.266947 | 4.69E-06 | Other Positive Correlation |
| PRRX2 | 0.266529 | 4.86E-06 | Other Positive Correlation |
| INHBB | 0.266055 | 5.06E-06 | Other Positive Correlation |
| PNMA2 | 0.264754 | 5.65E-06 | Other Positive Correlation |
| ADAMTS2 | 0.264554 | 5.75E-06 | Other Positive Correlation |
| STMND1 | 0.264217 | 5.91E-06 | Other Positive Correlation |
| PELI2 | 0.263509 | 6.27E-06 | Other Positive Correlation |
| PSPH | 0.262827 | 6.64E-06 | Other Positive Correlation |
| PID1 | 0.262642 | 6.75E-06 | Other Positive Correlation |
| TMEM108 | 0.262602 | 6.77E-06 | Other Positive Correlation |
| MYBL1 | 0.262527 | 6.81E-06 | Other Positive Correlation |
| C1QTNF2 | 0.262136 | 7.04E-06 | Other Positive Correlation |
| RFC4 | 0.262016 | 7.11E-06 | Other Positive Correlation |
| NR4A1 | 0.261639 | 7.34E-06 | Other Positive Correlation |
| TMEM145 | 0.261127 | 7.66E-06 | Other Positive Correlation |
| MAL | 0.260248 | 8.23E-06 | Other Positive Correlation |
| FOXL2 | 0.260089 | 8.34E-06 | Other Positive Correlation |
| DUSP15 | 0.260026 | 8.39E-06 | Other Positive Correlation |
| ONECUT2 | 0.259288 | 8.91E-06 | Other Positive Correlation |
| TSC22D3 | 0.259075 | 9.07E-06 | Other Positive Correlation |
| MAFA | 0.258701 | 9.36E-06 | Other Positive Correlation |
| ERICH2 | 0.258638 | 9.4E-06 | Other Positive Correlation |
| POU4F1 | 0.258454 | 9.55E-06 | Other Positive Correlation |
| GOT1 | 0.257165 | 1.06E-05 | Other Positive Correlation |
| CALCA | 0.256761 | 1.1E-05 | Other Positive Correlation |
| CHRD | 0.256086 | 1.16E-05 | Other Positive Correlation |

| | | | |
|---|---|---|---|
| DOK6 | 0.254259 | 1.34E-05 | Other Positive Correlation |
| LY6H | 0.253829 | 1.39E-05 | Other Positive Correlation |
| CYP27A1 | 0.253425 | 1.44E-05 | Other Positive Correlation |
| STXBP6 | 0.253146 | 1.47E-05 | Other Positive Correlation |
| REEP1 | 0.252482 | 1.55E-05 | Other Positive Correlation |
| SLC7A3 | 0.252417 | 1.56E-05 | Other Positive Correlation |
| ERC2 | 0.252399 | 1.56E-05 | Other Positive Correlation |
| PHOX2A | 0.252296 | 1.57E-05 | Other Positive Correlation |
| HOXB7 | 0.252196 | 1.59E-05 | Other Positive Correlation |
| HMX2 | 0.251877 | 1.63E-05 | Other Positive Correlation |
| ZNF483 | 0.251591 | 1.67E-05 | Other Positive Correlation |
| CCK | 0.250501 | 1.82E-05 | Other Positive Correlation |
| NCAM1 | 0.249487 | 1.97E-05 | Other Positive Correlation |
| COX19 | 0.249446 | 1.98E-05 | Other Positive Correlation |
| MSX1 | 0.24917 | 2.02E-05 | Other Positive Correlation |
| LTK | 0.249051 | 2.04E-05 | Other Positive Correlation |
| RND2 | 0.248857 | 2.07E-05 | Other Positive Correlation |
| SGCG | 0.248677 | 2.1E-05 | Other Positive Correlation |
| CDH2 | 0.248424 | 2.14E-05 | Other Positive Correlation |
| RGMB | 0.248209 | 2.18E-05 | Other Positive Correlation |
| GABARAPL1 | 0.247865 | 2.24E-05 | Other Positive Correlation |
| ULBP1 | 0.24731 | 2.34E-05 | Other Positive Correlation |
| FGF9 | 0.247168 | 2.36E-05 | Other Positive Correlation |
| SERTAD4 | 0.246398 | 2.51E-05 | Other Positive Correlation |
| BEND6 | 0.245772 | 2.64E-05 | Other Positive Correlation |
| CTSA | 0.245627 | 2.67E-05 | Other Positive Correlation |
| RORA | 0.243912 | 3.05E-05 | Other Positive Correlation |
| FICD | 0.243784 | 3.08E-05 | Other Positive Correlation |
| KCND3 | 0.242786 | 3.32E-05 | Other Positive Correlation |
| C3orf70 | 0.242697 | 3.35E-05 | Other Positive Correlation |
| NTNG1 | 0.242577 | 3.38E-05 | Other Positive Correlation |
| GAS2L3 | 0.242564 | 3.38E-05 | Other Positive Correlation |
| SYT13 | 0.24228 | 3.45E-05 | Other Positive Correlation |
| EBF3 | 0.242185 | 3.48E-05 | Other Positive Correlation |
| BSPRY | 0.242004 | 3.53E-05 | Other Positive Correlation |
| TMEM200C | 0.240845 | 3.86E-05 | Other Positive Correlation |
| STK40 | 0.240791 | 3.87E-05 | Other Positive Correlation |
| ERO1B | 0.240555 | 3.94E-05 | Other Positive Correlation |
| ZNF367 | 0.240039 | 4.1E-05 | Other Positive Correlation |
| NUP210 | 0.239481 | 4.28E-05 | Other Positive Correlation |
| NEFH | 0.238268 | 4.69E-05 | Other Positive Correlation |
| DCHS1 | 0.238045 | 4.77E-05 | Other Positive Correlation |
| CLGN | 0.237865 | 4.83E-05 | Other Positive Correlation |
| SLC16A11 | 0.237853 | 4.84E-05 | Other Positive Correlation |
| CECR2 | 0.237528 | 4.96E-05 | Other Positive Correlation |
| CENPU | 0.237295 | 5.05E-05 | Other Positive Correlation |

| | | | |
|---|---|---|---|
| TLE2 | 0.237284 | 5.05E-05 | Other Positive Correlation |
| SLC38A3 | 0.237137 | 5.11E-05 | Other Positive Correlation |
| ITPKA | 0.236555 | 5.33E-05 | Other Positive Correlation |
| RBPMS2 | 0.235407 | 5.81E-05 | Other Positive Correlation |
| AVP | 0.235281 | 5.87E-05 | Other Positive Correlation |
| SELENOK | 0.235043 | 5.97E-05 | Other Positive Correlation |
| PRKG1 | 0.23499 | 6E-05 | Other Positive Correlation |
| LHX2 | 0.234712 | 6.12E-05 | Other Positive Correlation |
| IGF2 | 0.233871 | 6.52E-05 | Other Positive Correlation |
| PHF1 | 0.233632 | 6.63E-05 | Other Positive Correlation |
| GAL | 0.233215 | 6.84E-05 | Other Positive Correlation |
| AMN | 0.232687 | 7.11E-05 | Other Positive Correlation |
| DEGS2 | 0.231909 | 7.53E-05 | Other Positive Correlation |
| HERPUD1 | 0.231474 | 7.78E-05 | Other Positive Correlation |
| SLC16A10 | 0.231067 | 8.01E-05 | Other Positive Correlation |
| FGF13 | 0.231001 | 8.05E-05 | Other Positive Correlation |
| RARRES2 | 0.230395 | 8.41E-05 | Other Positive Correlation |
| HID1 | 0.230252 | 8.5E-05 | Other Positive Correlation |
| MORC4 | 0.228827 | 9.43E-05 | Other Positive Correlation |
| RASSF1 | 0.228708 | 9.51E-05 | Other Positive Correlation |
| AIF1L | 0.228441 | 9.7E-05 | Other Positive Correlation |
| HSPA9 | 0.228426 | 9.71E-05 | Other Positive Correlation |
| LINGO1 | 0.228267 | 9.82E-05 | Other Positive Correlation |
| TCF19 | 0.228095 | 9.94E-05 | Other Positive Correlation |
| RUNDC3B | 0.227659 | 0.000103 | Other Positive Correlation |
| HOXB9 | 0.227566 | 0.000103 | Other Positive Correlation |
| WNT6 | 0.227537 | 0.000104 | Other Positive Correlation |
| NOTUM | 0.227479 | 0.000104 | Other Positive Correlation |
| ID4 | 0.227437 | 0.000104 | Other Positive Correlation |
| NPY | 0.227408 | 0.000104 | Other Positive Correlation |
| ZMAT1 | 0.227385 | 0.000105 | Other Positive Correlation |
| SCGB3A1 | 0.22702 | 0.000107 | Other Positive Correlation |
| SAMD11 | 0.226901 | 0.000108 | Other Positive Correlation |
| JPH4 | 0.226853 | 0.000109 | Other Positive Correlation |
| RGS16 | 0.226584 | 0.000111 | Other Positive Correlation |
| MLXIPL | 0.22632 | 0.000113 | Other Positive Correlation |
| DPYSL5 | 0.226266 | 0.000113 | Other Positive Correlation |
| SPTBN4 | 0.22551 | 0.00012 | Other Positive Correlation |
| FSD1 | 0.225499 | 0.00012 | Other Positive Correlation |
| CLCN5 | 0.225023 | 0.000124 | Other Positive Correlation |
| DEUP1 | 0.224467 | 0.000129 | Other Positive Correlation |
| GFRA1 | 0.224397 | 0.00013 | Other Positive Correlation |
| MKX | 0.224067 | 0.000133 | Other Positive Correlation |
| MLEC | 0.223966 | 0.000134 | Other Positive Correlation |
| TIMM44 | 0.223786 | 0.000135 | Other Positive Correlation |
| PIP5K1B | 0.223701 | 0.000136 | Other Positive Correlation |

| | | | |
|---|---|---|---|
| SYCP2 | 0.223203 | 0.000141 | Other Positive Correlation |
| IDUA | 0.222746 | 0.000146 | Other Positive Correlation |
| LONP1 | 0.222631 | 0.000147 | Other Positive Correlation |
| ZBTB18 | 0.221457 | 0.00016 | Other Positive Correlation |
| CENPM | 0.221232 | 0.000162 | Other Positive Correlation |
| ELANE | 0.2211 | 0.000164 | Other Positive Correlation |
| SLC31A1 | 0.221058 | 0.000164 | Other Positive Correlation |
| SFMBT2 | 0.220844 | 0.000167 | Other Positive Correlation |
| CEP128 | 0.220693 | 0.000168 | Other Positive Correlation |
| NKAIN4 | 0.220687 | 0.000168 | Other Positive Correlation |
| DYNC1I1 | 0.220244 | 0.000174 | Other Positive Correlation |
| GPRC5B | 0.219736 | 0.00018 | Other Positive Correlation |
| ILDR2 | 0.219482 | 0.000183 | Other Positive Correlation |
| AKAP12 | 0.219417 | 0.000184 | Other Positive Correlation |
| ZNF333 | 0.218658 | 0.000194 | Other Positive Correlation |
| NES | 0.218555 | 0.000195 | Other Positive Correlation |
| EPRS1 | 0.21836 | 0.000198 | Other Positive Correlation |
| CDKN2C | 0.218258 | 0.000199 | Other Positive Correlation |
| SPOCK2 | 0.217965 | 0.000203 | Other Positive Correlation |
| TFR2 | 0.217849 | 0.000205 | Other Positive Correlation |
| JSRP1 | 0.217592 | 0.000209 | Other Positive Correlation |
| SNAP91 | 0.217465 | 0.000211 | Other Positive Correlation |
| RAB39A | 0.217344 | 0.000212 | Other Positive Correlation |
| SOCS7 | 0.217138 | 0.000215 | Other Positive Correlation |
| HCN4 | 0.21669 | 0.000222 | Other Positive Correlation |
| SNCB | 0.215683 | 0.000238 | Other Positive Correlation |
| PLEKHD1 | 0.215294 | 0.000244 | Other Positive Correlation |
| WNT2B | 0.215278 | 0.000245 | Other Positive Correlation |
| PRR18 | 0.21508 | 0.000248 | Other Positive Correlation |
| C19orf38 | 0.215009 | 0.000249 | Other Positive Correlation |
| RGL3 | 0.21452 | 0.000258 | Other Positive Correlation |
| CDC6 | 0.214337 | 0.000261 | Other Positive Correlation |
| HAGH | 0.21422 | 0.000263 | Other Positive Correlation |
| SCML2 | 0.213682 | 0.000273 | Other Positive Correlation |
| CDK5R2 | 0.213482 | 0.000276 | Other Positive Correlation |
| NR1D2 | 0.213422 | 0.000277 | Other Positive Correlation |
| NFATC2 | 0.213332 | 0.000279 | Other Positive Correlation |
| HUNK | 0.212963 | 0.000286 | Other Positive Correlation |
| BMP7 | 0.212951 | 0.000286 | Other Positive Correlation |
| ART5 | 0.212715 | 0.000291 | Other Positive Correlation |
| ATP1B2 | 0.212449 | 0.000296 | Other Positive Correlation |
| MSI1 | 0.212212 | 0.000301 | Other Positive Correlation |
| STOX1 | 0.212145 | 0.000302 | Other Positive Correlation |
| ENPP5 | 0.211883 | 0.000308 | Other Positive Correlation |
| ADCYAP1 | 0.211469 | 0.000316 | Other Positive Correlation |
| MYDGF | 0.211383 | 0.000318 | Other Positive Correlation |

| | | | |
|---|---|---|---|
| PLEKHG4B | 0.21109 | 0.000325 | Other Positive Correlation |
| PTP4A3 | 0.210912 | 0.000329 | Other Positive Correlation |
| CEBPG | 0.210894 | 0.000329 | Other Positive Correlation |
| YPEL1 | 0.210706 | 0.000333 | Other Positive Correlation |
| RCN3 | 0.210589 | 0.000336 | Other Positive Correlation |
| CDKN2D | 0.210274 | 0.000343 | Other Positive Correlation |
| GNAZ | 0.210232 | 0.000344 | Other Positive Correlation |
| RAI2 | 0.210074 | 0.000347 | Other Positive Correlation |
| KCNK15 | 0.210049 | 0.000348 | Other Positive Correlation |
| FGF19 | 0.20967 | 0.000357 | Other Positive Correlation |
| CLIC6 | 0.209507 | 0.000361 | Other Positive Correlation |
| CCDC110 | 0.209304 | 0.000366 | Other Positive Correlation |
| KIFC2 | 0.209231 | 0.000367 | Other Positive Correlation |
| PTCHD1 | 0.209226 | 0.000368 | Other Positive Correlation |
| SEPTIN4 | 0.209106 | 0.00037 | Other Positive Correlation |
| CHRM3 | 0.208997 | 0.000373 | Other Positive Correlation |
| ZNF85 | 0.208553 | 0.000384 | Other Positive Correlation |
| JAM2 | 0.208445 | 0.000387 | Other Positive Correlation |
| SKOR1 | 0.208247 | 0.000392 | Other Positive Correlation |
| SCAMP5 | 0.208111 | 0.000396 | Other Positive Correlation |
| GPAT3 | 0.207806 | 0.000404 | Other Positive Correlation |
| LYRM1 | 0.207413 | 0.000414 | Other Positive Correlation |
| NMUR1 | 0.207132 | 0.000422 | Other Positive Correlation |
| KCNG3 | 0.207086 | 0.000423 | Other Positive Correlation |
| SCN8A | 0.205787 | 0.000461 | Other Positive Correlation |
| MAFG | 0.205106 | 0.000482 | Other Positive Correlation |
| SLC35F1 | 0.204708 | 0.000494 | Other Positive Correlation |
| ZNF280B | 0.2047 | 0.000495 | Other Positive Correlation |
| KIF5C | 0.203942 | 0.00052 | Other Positive Correlation |
| NOVA1 | 0.203915 | 0.000521 | Other Positive Correlation |
| FGFRL1 | 0.203775 | 0.000525 | Other Positive Correlation |
| GINS4 | 0.20362 | 0.000531 | Other Positive Correlation |
| PPP2R3C | 0.203511 | 0.000534 | Other Positive Correlation |
| EMX1 | 0.203467 | 0.000536 | Other Positive Correlation |
| COLGALT2 | 0.203069 | 0.00055 | Other Positive Correlation |
| CHN1 | 0.202922 | 0.000555 | Other Positive Correlation |
| PKD1 | 0.202826 | 0.000558 | Other Positive Correlation |
| GKAP1 | 0.201973 | 0.00059 | Other Positive Correlation |
| SLC30A3 | 0.201971 | 0.00059 | Other Positive Correlation |
| PDE4A | 0.201807 | 0.000596 | Other Positive Correlation |
| SYN2 | 0.201116 | 0.000623 | Other Positive Correlation |
| ST8SIA2 | 0.200793 | 0.000636 | Other Positive Correlation |
| DMBX1 | 0.200562 | 0.000646 | Other Positive Correlation |
| PPP2R3B | 0.200123 | 0.000664 | Other Positive Correlation |
| CA8 | 0.200004 | 0.000669 | Other Positive Correlation |
| COL6A3 | 0.199918 | 0.000673 | Other Positive Correlation |

| | | | |
|---|---|---|---|
| SELENOM | 0.199793 | 0.000678 | Other Positive Correlation |
| RELL2 | 0.199752 | 0.00068 | Other Positive Correlation |
| LGI2 | 0.199436 | 0.000694 | Other Positive Correlation |
| PPP2R2C | 0.199195 | 0.000704 | Other Positive Correlation |
| CACNA2D2 | 0.19851 | 0.000735 | Other Positive Correlation |
| AARS1 | 0.198053 | 0.000757 | Other Positive Correlation |
| PPM1E | 0.197868 | 0.000766 | Other Positive Correlation |
| ZNF560 | 0.197765 | 0.000771 | Other Positive Correlation |
| FAM162B | 0.197723 | 0.000773 | Other Positive Correlation |
| JADE1 | 0.197578 | 0.00078 | Other Positive Correlation |
| TOX2 | 0.197328 | 0.000792 | Other Positive Correlation |
| PAX2 | 0.19729 | 0.000794 | Other Positive Correlation |
| LRRC4B | 0.197249 | 0.000796 | Other Positive Correlation |
| RAB6B | 0.197229 | 0.000797 | Other Positive Correlation |
| EPHB6 | 0.196956 | 0.000811 | Other Positive Correlation |
| PTH2 | 0.196803 | 0.000819 | Other Positive Correlation |
| PLEKHG4 | 0.19659 | 0.00083 | Other Positive Correlation |
| FOXI2 | 0.196566 | 0.000831 | Other Positive Correlation |
| CCDC167 | 0.196524 | 0.000833 | Other Positive Correlation |
| PRDM16 | 0.19626 | 0.000847 | Other Positive Correlation |
| WFDC2 | 0.196062 | 0.000857 | Other Positive Correlation |
| NR2C2AP | 0.19578 | 0.000872 | Other Positive Correlation |
| OLFM1 | 0.195179 | 0.000906 | Other Positive Correlation |
| DCDC2 | 0.194979 | 0.000917 | Other Positive Correlation |
| AGO2 | 0.194954 | 0.000918 | Other Positive Correlation |
| RADIL | 0.1949 | 0.000922 | Other Positive Correlation |
| HOXD9 | 0.194589 | 0.000939 | Other Positive Correlation |
| BHLHE41 | 0.194547 | 0.000942 | Other Positive Correlation |
| CENPK | 0.194214 | 0.000961 | Other Positive Correlation |
| BEX2 | 0.194203 | 0.000962 | Other Positive Correlation |
| AK5 | 0.194192 | 0.000963 | Other Positive Correlation |
| UTF1 | 0.194183 | 0.000963 | Other Positive Correlation |
| ARX | 0.194044 | 0.000972 | Other Positive Correlation |
| TMOD1 | 0.193432 | 0.001009 | Other Positive Correlation |
| FOXE3 | 0.193293 | 0.001018 | Other Positive Correlation |
| TBX15 | 0.192574 | 0.001064 | Other Positive Correlation |
| FXYD6 | 0.19241 | 0.001074 | Other Positive Correlation |
| HHIPL1 | 0.192236 | 0.001086 | Other Positive Correlation |
| DNASE2 | 0.19134 | 0.001147 | Other Positive Correlation |
| PTPRN | 0.191165 | 0.001159 | Other Positive Correlation |
| HOXA10 | 0.191067 | 0.001166 | Other Positive Correlation |
| PTGER2 | 0.191028 | 0.001169 | Other Positive Correlation |
| WSCD1 | 0.190986 | 0.001172 | Other Positive Correlation |
| DCP2 | 0.190703 | 0.001192 | Other Positive Correlation |
| UBE2T | 0.190372 | 0.001216 | Other Positive Correlation |
| ETFDH | 0.190166 | 0.001232 | Other Positive Correlation |

| | | | |
|---|---|---|---|
| NGF | 0.190106 | 0.001236 | Other Positive Correlation |
| ZNF331 | 0.190102 | 0.001236 | Other Positive Correlation |
| OVOL2 | 0.190011 | 0.001243 | Other Positive Correlation |
| TOX3 | 0.190004 | 0.001244 | Other Positive Correlation |
| UBE2O | 0.189929 | 0.001249 | Other Positive Correlation |
| PTF1A | 0.189681 | 0.001268 | Other Positive Correlation |
| TAL1 | 0.18945 | 0.001286 | Other Positive Correlation |
| IGFBPL1 | 0.189431 | 0.001288 | Other Positive Correlation |
| FRY | 0.189214 | 0.001305 | Other Positive Correlation |
| NEMP1 | 0.189076 | 0.001315 | Other Positive Correlation |
| SNTB1 | 0.18898 | 0.001323 | Other Positive Correlation |
| MTFR2 | 0.188577 | 0.001356 | Other Positive Correlation |
| IFRD1 | 0.188569 | 0.001356 | Other Positive Correlation |
| RPL39L | 0.188311 | 0.001377 | Other Positive Correlation |
| ACOXL | 0.188246 | 0.001383 | Other Positive Correlation |
| ARHGAP45 | 0.188169 | 0.001389 | Other Positive Correlation |
| TAFA4 | 0.187374 | 0.001457 | Other Positive Correlation |
| CFL2 | 0.18718 | 0.001474 | Other Positive Correlation |
| PLCB1 | 0.187082 | 0.001482 | Other Positive Correlation |
| STX3 | 0.186795 | 0.001508 | Other Positive Correlation |
| CHRM4 | 0.186558 | 0.001529 | Other Positive Correlation |
| TBX2 | 0.186315 | 0.001552 | Other Positive Correlation |
| PRADC1 | 0.186174 | 0.001565 | Other Positive Correlation |
| GMPR | 0.186144 | 0.001568 | Other Positive Correlation |
| HSPA5 | 0.185923 | 0.001588 | Other Positive Correlation |
| PSIP1 | 0.185896 | 0.001591 | Other Positive Correlation |
| LDHC | 0.185863 | 0.001594 | Other Positive Correlation |
| DDIT4L | 0.185482 | 0.00163 | Other Positive Correlation |
| SLC30A2 | 0.185146 | 0.001663 | Other Positive Correlation |
| TMEM163 | 0.185117 | 0.001666 | Other Positive Correlation |
| PARM1 | 0.18494 | 0.001683 | Other Positive Correlation |
| ARHGAP20 | 0.184429 | 0.001735 | Other Positive Correlation |
| ACE | 0.184398 | 0.001738 | Other Positive Correlation |
| LRRN1 | 0.184348 | 0.001743 | Other Positive Correlation |
| ANKRD53 | 0.18416 | 0.001762 | Other Positive Correlation |
| FGF16 | 0.183711 | 0.00181 | Other Positive Correlation |
| PGM5 | 0.183689 | 0.001812 | Other Positive Correlation |
| AKR7A3 | 0.18338 | 0.001845 | Other Positive Correlation |
| TCEA3 | 0.183311 | 0.001853 | Other Positive Correlation |
| LONRF2 | 0.183096 | 0.001876 | Other Positive Correlation |
| EMX2 | 0.183064 | 0.00188 | Other Positive Correlation |
| TAF5 | 0.18291 | 0.001896 | Other Positive Correlation |
| ALKAL2 | 0.182355 | 0.001959 | Other Positive Correlation |
| VEGFB | 0.182182 | 0.001979 | Other Positive Correlation |
| C17orf107 | 0.181947 | 0.002006 | Other Positive Correlation |
| BEX5 | 0.181877 | 0.002014 | Other Positive Correlation |

| | | | |
|---|---|---|---|
| TDRD9 | 0.181813 | 0.002022 | Other Positive Correlation |
| ELMOD1 | 0.181654 | 0.00204 | Other Positive Correlation |
| TMEM179 | 0.181559 | 0.002052 | Other Positive Correlation |
| MAN2A2 | 0.181363 | 0.002075 | Other Positive Correlation |
| GATA4 | 0.181275 | 0.002086 | Other Positive Correlation |
| CADM4 | 0.181263 | 0.002087 | Other Positive Correlation |
| ZNF93 | 0.181046 | 0.002113 | Other Positive Correlation |
| VSTM2B | 0.180812 | 0.002142 | Other Positive Correlation |
| TDRD10 | 0.180433 | 0.00219 | Other Positive Correlation |
| GABRQ | 0.18021 | 0.002218 | Other Positive Correlation |
| DLX4 | 0.180002 | 0.002245 | Other Positive Correlation |
| SYVN1 | 0.179937 | 0.002253 | Other Positive Correlation |
| ENSG00000285920 | 0.179663 | 0.002289 | Other Positive Correlation |
| CNTFR | 0.179607 | 0.002296 | Other Positive Correlation |
| ACHE | 0.179456 | 0.002316 | Other Positive Correlation |
| BMP8B | 0.179444 | 0.002318 | Other Positive Correlation |
| KCNN1 | 0.179333 | 0.002333 | Other Positive Correlation |
| CALU | 0.179111 | 0.002363 | Other Positive Correlation |
| DLX6 | 0.179104 | 0.002364 | Other Positive Correlation |
| PAX6 | 0.17888 | 0.002394 | Other Positive Correlation |
| PALM3 | 0.178388 | 0.002462 | Other Positive Correlation |
| CCDC85A | 0.178286 | 0.002477 | Other Positive Correlation |
| BLVRB | 0.178214 | 0.002487 | Other Positive Correlation |
| KHDRBS3 | 0.178164 | 0.002494 | Other Positive Correlation |
| ARHGDIG | 0.177894 | 0.002533 | Other Positive Correlation |
| HHEX | 0.177738 | 0.002555 | Other Positive Correlation |
| RBMXL2 | 0.177546 | 0.002583 | Other Positive Correlation |
| UAP1L1 | 0.177394 | 0.002606 | Other Positive Correlation |
| HLA-B | 0.176708 | 0.002709 | Other Positive Correlation |
| COL11A2 | 0.176225 | 0.002784 | Other Positive Correlation |
| NAT8L | 0.176083 | 0.002806 | Other Positive Correlation |
| GCHFR | 0.175925 | 0.002832 | Other Positive Correlation |
| SP5 | 0.175836 | 0.002846 | Other Positive Correlation |
| TSPAN18 | 0.175814 | 0.002849 | Other Positive Correlation |
| SHOX2 | 0.175447 | 0.002909 | Other Positive Correlation |
| ADRA2C | 0.174821 | 0.003013 | Other Positive Correlation |
| TTLL7 | 0.174703 | 0.003033 | Other Positive Correlation |
| WFS1 | 0.174609 | 0.003049 | Other Positive Correlation |
| FEN1 | 0.17427 | 0.003107 | Other Positive Correlation |
| CLSPN | 0.173606 | 0.003224 | Other Positive Correlation |
| FADS6 | 0.173055 | 0.003325 | Other Positive Correlation |
| FAM3B | 0.173013 | 0.003333 | Other Positive Correlation |
| COL25A1 | 0.172932 | 0.003348 | Other Positive Correlation |
| PLPPR3 | 0.17293 | 0.003348 | Other Positive Correlation |
| NR2F1 | 0.172775 | 0.003377 | Other Positive Correlation |
| ISL2 | 0.172577 | 0.003414 | Other Positive Correlation |

| | | | |
|---|---|---|---|
| ERBB4 | 0.172563 | 0.003417 | Other Positive Correlation |
| ELOVL2 | 0.172412 | 0.003446 | Other Positive Correlation |
| EN2 | 0.172231 | 0.00348 | Other Positive Correlation |
| RAMP1 | 0.171927 | 0.003539 | Other Positive Correlation |
| DTX1 | 0.171881 | 0.003548 | Other Positive Correlation |
| EML5 | 0.171878 | 0.003549 | Other Positive Correlation |
| USP44 | 0.171389 | 0.003646 | Other Positive Correlation |
| DSCC1 | 0.171062 | 0.003712 | Other Positive Correlation |
| CBX7 | 0.170778 | 0.00377 | Other Positive Correlation |
| HTRA3 | 0.170311 | 0.003868 | Other Positive Correlation |
| HCN2 | 0.169756 | 0.003987 | Other Positive Correlation |
| ZNF473 | 0.169654 | 0.004009 | Other Positive Correlation |
| CRTAC1 | 0.169575 | 0.004026 | Other Positive Correlation |
| RHEB | 0.169143 | 0.004122 | Other Positive Correlation |
| FLNC | 0.16912 | 0.004128 | Other Positive Correlation |
| SYNGR3 | 0.169045 | 0.004144 | Other Positive Correlation |
| THRA | 0.169032 | 0.004147 | Other Positive Correlation |
| SDS | 0.168638 | 0.004237 | Other Positive Correlation |
| STAC | 0.168401 | 0.004292 | Other Positive Correlation |
| HPCAL4 | 0.168301 | 0.004315 | Other Positive Correlation |
| MAZ | 0.168266 | 0.004323 | Other Positive Correlation |
| FXYD7 | 0.168208 | 0.004337 | Other Positive Correlation |
| CFAP119 | 0.168106 | 0.004361 | Other Positive Correlation |
| ENTREP2 | 0.167937 | 0.004401 | Other Positive Correlation |
| DHRS13 | 0.167841 | 0.004424 | Other Positive Correlation |
| TRIM71 | 0.167813 | 0.004431 | Other Positive Correlation |
| SELENOV | 0.167756 | 0.004444 | Other Positive Correlation |
| RNF223 | 0.167733 | 0.00445 | Other Positive Correlation |
| EBF4 | 0.167086 | 0.004608 | Other Positive Correlation |
| AR | 0.166713 | 0.004701 | Other Positive Correlation |
| CDT1 | 0.166508 | 0.004753 | Other Positive Correlation |
| PRUNE2 | 0.166442 | 0.00477 | Other Positive Correlation |
| HELT | 0.16631 | 0.004804 | Other Positive Correlation |
| ANK1 | 0.166305 | 0.004805 | Other Positive Correlation |
| SLC26A5 | 0.166041 | 0.004874 | Other Positive Correlation |
| PDE4D | 0.165987 | 0.004888 | Other Positive Correlation |
| SH3GL2 | 0.165936 | 0.004901 | Other Positive Correlation |
| IKZF1 | 0.16575 | 0.00495 | Other Positive Correlation |
| CLDN3 | 0.165735 | 0.004954 | Other Positive Correlation |
| CNIH2 | 0.165634 | 0.004981 | Other Positive Correlation |
| TSEN15 | 0.165627 | 0.004983 | Other Positive Correlation |
| KCNJ10 | 0.16556 | 0.005 | Other Positive Correlation |
| C3orf18 | 0.164676 | 0.005241 | Other Positive Correlation |
| TSPYL2 | 0.164631 | 0.005254 | Other Positive Correlation |
| NGFR | 0.164501 | 0.00529 | Other Positive Correlation |
| MOXD1 | 0.164254 | 0.00536 | Other Positive Correlation |

| | | | |
|---|---|---|---|
| CADM2 | 0.164248 | 0.005362 | Other Positive Correlation |
| DPP10 | 0.164176 | 0.005382 | Other Positive Correlation |
| RTBDN | 0.164169 | 0.005384 | Other Positive Correlation |
| BHMT2 | 0.164088 | 0.005407 | Other Positive Correlation |
| KIRREL2 | 0.164072 | 0.005412 | Other Positive Correlation |
| LYPD1 | 0.163924 | 0.005455 | Other Positive Correlation |
| AMZ1 | 0.163778 | 0.005497 | Other Positive Correlation |
| TUBE1 | 0.163669 | 0.005529 | Other Positive Correlation |
| MYOD1 | 0.163485 | 0.005583 | Other Positive Correlation |
| PEG3 | 0.163471 | 0.005587 | Other Positive Correlation |
| DSCAML1 | 0.163137 | 0.005686 | Other Positive Correlation |
| DHRS3 | 0.162949 | 0.005742 | Other Positive Correlation |
| FREM2 | 0.162818 | 0.005782 | Other Positive Correlation |
| ELAVL3 | 0.162622 | 0.005842 | Other Positive Correlation |
| ARFGEF3 | 0.162602 | 0.005848 | Other Positive Correlation |
| APBB1 | 0.162516 | 0.005875 | Other Positive Correlation |
| SOWAHA | 0.162446 | 0.005896 | Other Positive Correlation |
| TDRD5 | 0.162418 | 0.005905 | Other Positive Correlation |
| SMOX | 0.161923 | 0.00606 | Other Positive Correlation |
| KAZALD1 | 0.161597 | 0.006164 | Other Positive Correlation |
| ASF1B | 0.161481 | 0.006202 | Other Positive Correlation |
| ZNF778 | 0.161398 | 0.006229 | Other Positive Correlation |
| SLC32A1 | 0.160886 | 0.006397 | Other Positive Correlation |
| RIPPLY3 | 0.160513 | 0.006522 | Other Positive Correlation |
| UNC5D | 0.160408 | 0.006558 | Other Positive Correlation |
| RNF168 | 0.160359 | 0.006575 | Other Positive Correlation |
| ENPP1 | 0.160348 | 0.006579 | Other Positive Correlation |
| PPP1R3E | 0.160281 | 0.006601 | Other Positive Correlation |
| CNNM1 | 0.160112 | 0.006659 | Other Positive Correlation |
| SLC6A9 | 0.159842 | 0.006753 | Other Positive Correlation |
| SOX17 | 0.159705 | 0.006801 | Other Positive Correlation |
| UPK3A | 0.159671 | 0.006813 | Other Positive Correlation |
| NEXMIF | 0.159119 | 0.00701 | Other Positive Correlation |
| S1PR1 | 0.159096 | 0.007019 | Other Positive Correlation |
| PIANP | 0.159083 | 0.007023 | Other Positive Correlation |
| GNMT | 0.159006 | 0.007051 | Other Positive Correlation |
| YIPF7 | 0.159 | 0.007053 | Other Positive Correlation |
| ANXA6 | 0.158923 | 0.007081 | Other Positive Correlation |
| NFKBIB | 0.158881 | 0.007096 | Other Positive Correlation |
| ISM1 | 0.158654 | 0.00718 | Other Positive Correlation |
| RNFT2 | 0.157795 | 0.007503 | Other Positive Correlation |
| DOCK3 | 0.157759 | 0.007517 | Other Positive Correlation |
| DPYD | 0.157722 | 0.007531 | Other Positive Correlation |
| CLDN6 | 0.157164 | 0.007749 | Other Positive Correlation |
| ADD2 | 0.157151 | 0.007754 | Other Positive Correlation |
| GRIN2B | 0.157085 | 0.00778 | Other Positive Correlation |

| | | | |
|---|---|---|---|
| ZNF536 | 0.15702 | 0.007806 | Other Positive Correlation |
| NKX2-5 | 0.15614 | 0.008163 | Other Positive Correlation |
| STK32A | 0.156048 | 0.008201 | Other Positive Correlation |
| HYOU1 | 0.155887 | 0.008268 | Other Positive Correlation |
| AGTR1 | 0.15568 | 0.008355 | Other Positive Correlation |
| SP4 | 0.155644 | 0.008371 | Other Positive Correlation |
| CNNM2 | 0.155449 | 0.008454 | Other Positive Correlation |
| DNM1 | 0.154909 | 0.008687 | Other Positive Correlation |
| ENO3 | 0.154754 | 0.008755 | Other Positive Correlation |
| ABHD12B | 0.154717 | 0.008771 | Other Positive Correlation |
| ATRNL1 | 0.154683 | 0.008786 | Other Positive Correlation |
| CENPJ | 0.154378 | 0.008922 | Other Positive Correlation |
| TMEM225B | 0.154284 | 0.008964 | Other Positive Correlation |
| MYH14 | 0.154206 | 0.008999 | Other Positive Correlation |
| CDON | 0.153979 | 0.009102 | Other Positive Correlation |
| HRC | 0.153975 | 0.009104 | Other Positive Correlation |
| SLAIN1 | 0.153868 | 0.009153 | Other Positive Correlation |
| HAT1 | 0.153675 | 0.009242 | Other Positive Correlation |
| NPTXR | 0.153667 | 0.009245 | Other Positive Correlation |
| SIX3 | 0.153614 | 0.00927 | Other Positive Correlation |
| XBP1 | 0.153573 | 0.009289 | Other Positive Correlation |
| RBP4 | 0.153272 | 0.00943 | Other Positive Correlation |
| GALM | 0.15279 | 0.009659 | Other Positive Correlation |
| KCNH2 | 0.152637 | 0.009733 | Other Positive Correlation |
| FGF22 | 0.152555 | 0.009772 | Other Positive Correlation |
| OTUD3 | 0.15233 | 0.009882 | Other Positive Correlation |
| RPA2 | 0.152208 | 0.009942 | Other Positive Correlation |
| SLC7A10 | 0.152147 | 0.009972 | Other Positive Correlation |
| KIAA1958 | 0.151968 | 0.010061 | Other Positive Correlation |
| NRTN | 0.151889 | 0.010101 | Other Positive Correlation |
| BGN | 0.151642 | 0.010225 | Other Positive Correlation |
| C10orf82 | 0.151428 | 0.010333 | Other Positive Correlation |
| CDH15 | 0.151085 | 0.01051 | Other Positive Correlation |
| ITM2C | 0.151042 | 0.010532 | Other Positive Correlation |
| TRAIP | 0.150677 | 0.010723 | Other Positive Correlation |
| PRPH | 0.150574 | 0.010777 | Other Positive Correlation |
| KCNH3 | 0.15049 | 0.010822 | Other Positive Correlation |
| PGP | 0.150477 | 0.010829 | Other Positive Correlation |
| RNASEH2A | 0.150228 | 0.010962 | Other Positive Correlation |
| COL4A4 | 0.150104 | 0.011029 | Other Positive Correlation |
| PENK | 0.15 | 0.011085 | Other Positive Correlation |
| CHST8 | 0.149863 | 0.01116 | Other Positive Correlation |
| PDK2 | 0.149848 | 0.011168 | Other Positive Correlation |
| CDX1 | 0.149718 | 0.011239 | Other Positive Correlation |
| TMEM176A | 0.149534 | 0.01134 | Other Positive Correlation |
| HEY2 | 0.149474 | 0.011374 | Other Positive Correlation |

| | | | |
|---|---|---|---|
| MAP1LC3B | 0.149433 | 0.011397 | Other Positive Correlation |
| ASTN2 | 0.149408 | 0.01141 | Other Positive Correlation |
| HLA-DPA1 | 0.14938 | 0.011426 | Other Positive Correlation |
| CXCR4 | 0.149285 | 0.011479 | Other Positive Correlation |
| CHL1 | 0.14901 | 0.011634 | Other Positive Correlation |
| NKX1-1 | 0.148964 | 0.01166 | Other Positive Correlation |
| SMIM13 | 0.14875 | 0.011782 | Other Positive Correlation |
| SIM1 | 0.148464 | 0.011947 | Other Positive Correlation |
| FAM131C | 0.148287 | 0.01205 | Other Positive Correlation |
| FRMD3 | 0.148205 | 0.012098 | Other Positive Correlation |
| CACNA1H | 0.148188 | 0.012108 | Other Positive Correlation |
| LRRC71 | 0.148183 | 0.012111 | Other Positive Correlation |
| RGS9BP | 0.148058 | 0.012185 | Other Positive Correlation |
| ORC6 | 0.147793 | 0.012342 | Other Positive Correlation |
| CPEB3 | 0.147608 | 0.012452 | Other Positive Correlation |
| CD8A | 0.147584 | 0.012467 | Other Positive Correlation |
| RBFOX3 | 0.14743 | 0.01256 | Other Positive Correlation |
| FRRS1L | 0.147403 | 0.012577 | Other Positive Correlation |
| ZNF695 | 0.14732 | 0.012627 | Other Positive Correlation |
| VIP | 0.147158 | 0.012726 | Other Positive Correlation |
| ABCA3 | 0.147099 | 0.012762 | Other Positive Correlation |
| BRCA1 | 0.146957 | 0.01285 | Other Positive Correlation |
| MGAT4A | 0.146897 | 0.012887 | Other Positive Correlation |
| GATA5 | 0.146877 | 0.012899 | Other Positive Correlation |
| PRKAR2B | 0.146808 | 0.012942 | Other Positive Correlation |
| MAP6 | 0.14677 | 0.012966 | Other Positive Correlation |
| SEMA4D | 0.14667 | 0.013028 | Other Positive Correlation |
| AKTIP | 0.146503 | 0.013133 | Other Positive Correlation |
| PDZD4 | 0.146432 | 0.013178 | Other Positive Correlation |
| RIMS4 | 0.146419 | 0.013186 | Other Positive Correlation |
| TMEM132E | 0.146346 | 0.013233 | Other Positive Correlation |
| REEP4 | 0.146329 | 0.013243 | Other Positive Correlation |
| RAD51 | 0.146258 | 0.013288 | Other Positive Correlation |
| SIGIRR | 0.146013 | 0.013446 | Other Positive Correlation |
| KLK1 | 0.145985 | 0.013463 | Other Positive Correlation |
| CFAP92 | 0.145806 | 0.013579 | Other Positive Correlation |
| FOXD2 | 0.14571 | 0.013642 | Other Positive Correlation |
| SLC27A6 | 0.145676 | 0.013663 | Other Positive Correlation |
| TUB | 0.145621 | 0.0137 | Other Positive Correlation |
| FLT4 | 0.145565 | 0.013736 | Other Positive Correlation |
| SCUBE2 | 0.145498 | 0.013781 | Other Positive Correlation |
| BCL2 | 0.145407 | 0.01384 | Other Positive Correlation |
| ELAVL4 | 0.14521 | 0.013971 | Other Positive Correlation |
| MCOLN3 | 0.144852 | 0.014211 | Other Positive Correlation |
| ZNF726 | 0.144768 | 0.014268 | Other Positive Correlation |
| MATN3 | 0.144733 | 0.014292 | Other Positive Correlation |

| | | | |
|---|---|---|---|
| PRRX1 | 0.144728 | 0.014295 | Other Positive Correlation |
| PITX3 | 0.144301 | 0.014588 | Other Positive Correlation |
| ZIC1 | 0.144273 | 0.014607 | Other Positive Correlation |
| SLC16A6 | 0.144205 | 0.014654 | Other Positive Correlation |
| SRPRA | 0.144174 | 0.014676 | Other Positive Correlation |
| C11orf96 | 0.144121 | 0.014713 | Other Positive Correlation |
| PDRG1 | 0.144062 | 0.014754 | Other Positive Correlation |
| DNAJC3 | 0.14404 | 0.014769 | Other Positive Correlation |
| CHRNA3 | 0.14397 | 0.014818 | Other Positive Correlation |
| CLIC5 | 0.143961 | 0.014824 | Other Positive Correlation |
| SLC25A48 | 0.143864 | 0.014892 | Other Positive Correlation |
| LIN28B | 0.143674 | 0.015026 | Other Positive Correlation |
| ABHD3 | 0.14356 | 0.015107 | Other Positive Correlation |
| DRD2 | 0.143525 | 0.015132 | Other Positive Correlation |
| HSF4 | 0.143512 | 0.015142 | Other Positive Correlation |
| RNF182 | 0.142974 | 0.01553 | Other Positive Correlation |
| TMEM176B | 0.142917 | 0.015572 | Other Positive Correlation |
| NECAB2 | 0.142734 | 0.015707 | Other Positive Correlation |
| DPF1 | 0.142609 | 0.015799 | Other Positive Correlation |
| GPR4 | 0.142549 | 0.015844 | Other Positive Correlation |
| TMEM170B | 0.142501 | 0.01588 | Other Positive Correlation |
| DNAJB2 | 0.142417 | 0.015942 | Other Positive Correlation |
| BTNL9 | 0.142405 | 0.015951 | Other Positive Correlation |
| ATAD2 | 0.14225 | 0.016067 | Other Positive Correlation |
| CNTN4 | 0.141929 | 0.01631 | Other Positive Correlation |
| ZBTB21 | 0.141925 | 0.016313 | Other Positive Correlation |
| FAM149A | 0.141512 | 0.016631 | Other Positive Correlation |
| ENSG00000286022 | 0.14145 | 0.016679 | Other Positive Correlation |
| DAPK1 | 0.141442 | 0.016685 | Other Positive Correlation |
| FHIT | 0.14138 | 0.016733 | Other Positive Correlation |
| CHRDL2 | 0.141372 | 0.01674 | Other Positive Correlation |
| MAPT | 0.141292 | 0.016802 | Other Positive Correlation |
| PCDH8 | 0.141252 | 0.016834 | Other Positive Correlation |
| PSTK | 0.141223 | 0.016857 | Other Positive Correlation |
| POC1A | 0.141136 | 0.016925 | Other Positive Correlation |
| LAMP3 | 0.141022 | 0.017015 | Other Positive Correlation |
| PCDH10 | 0.141014 | 0.017021 | Other Positive Correlation |
| CRELD1 | 0.140951 | 0.017071 | Other Positive Correlation |
| ARHGAP44 | 0.140544 | 0.017396 | Other Positive Correlation |
| SRSF12 | 0.140062 | 0.017789 | Other Positive Correlation |
| MCOLN1 | 0.140058 | 0.017792 | Other Positive Correlation |
| SLC18A3 | 0.140025 | 0.01782 | Other Positive Correlation |
| SOX8 | 0.140001 | 0.017839 | Other Positive Correlation |
| WDSUB1 | 0.139761 | 0.018038 | Other Positive Correlation |
| TYRO3 | 0.139651 | 0.018129 | Other Positive Correlation |
| ANO5 | 0.139568 | 0.018199 | Other Positive Correlation |

| | | | |
|---|---|---|---|
| GYPC | 0.139208 | 0.018503 | Other Positive Correlation |
| LHX3 | 0.13907 | 0.018621 | Other Positive Correlation |
| ALPK3 | 0.138922 | 0.018748 | Other Positive Correlation |
| RIMS1 | 0.138756 | 0.018891 | Other Positive Correlation |
| GRB10 | 0.138456 | 0.019153 | Other Positive Correlation |
| DACT3 | 0.138313 | 0.019278 | Other Positive Correlation |
| CALCOCO2 | 0.13822 | 0.019361 | Other Positive Correlation |
| BARHL2 | 0.137963 | 0.01959 | Other Positive Correlation |
| CACNG5 | 0.137887 | 0.019658 | Other Positive Correlation |
| ZWINT | 0.137879 | 0.019665 | Other Positive Correlation |
| PRKD1 | 0.137824 | 0.019714 | Other Positive Correlation |
| SYT6 | 0.137632 | 0.019888 | Other Positive Correlation |
| KCNA1 | 0.137604 | 0.019913 | Other Positive Correlation |
| GABRB3 | 0.137463 | 0.020041 | Other Positive Correlation |
| ST6GAL2 | 0.137366 | 0.02013 | Other Positive Correlation |
| PRRT1B | 0.137319 | 0.020173 | Other Positive Correlation |
| RASGRP1 | 0.137255 | 0.020232 | Other Positive Correlation |
| GCGR | 0.137093 | 0.020381 | Other Positive Correlation |
| CYP2R1 | 0.136874 | 0.020585 | Other Positive Correlation |
| ADAMTS17 | 0.136858 | 0.020599 | Other Positive Correlation |
| INA | 0.13684 | 0.020616 | Other Positive Correlation |
| GMPPB | 0.13679 | 0.020663 | Other Positive Correlation |
| SCML1 | 0.13669 | 0.020757 | Other Positive Correlation |
| NR3C2 | 0.136652 | 0.020792 | Other Positive Correlation |
| DCLK2 | 0.13659 | 0.020851 | Other Positive Correlation |
| DONSON | 0.136559 | 0.02088 | Other Positive Correlation |
| NRK | 0.136288 | 0.021138 | Other Positive Correlation |
| GUCY1A2 | 0.136267 | 0.021158 | Other Positive Correlation |
| TXNDC16 | 0.136259 | 0.021165 | Other Positive Correlation |
| C14orf132 | 0.136246 | 0.021178 | Other Positive Correlation |
| PLEKHO1 | 0.136109 | 0.02131 | Other Positive Correlation |
| KLHL30 | 0.136105 | 0.021313 | Other Positive Correlation |
| KCNJ4 | 0.135876 | 0.021535 | Other Positive Correlation |
| OXT | 0.135843 | 0.021566 | Other Positive Correlation |
| ITGA7 | 0.135706 | 0.021701 | Other Positive Correlation |
| SLC35D3 | 0.1356 | 0.021804 | Other Positive Correlation |
| LBX1 | 0.135421 | 0.021981 | Other Positive Correlation |
| FAM117A | 0.1351 | 0.022299 | Other Positive Correlation |
| GPR160 | 0.135098 | 0.022301 | Other Positive Correlation |
| ENSG00000287908 | 0.134889 | 0.022512 | Other Positive Correlation |
| WNT1 | 0.134836 | 0.022565 | Other Positive Correlation |
| ARHGAP33 | 0.134827 | 0.022574 | Other Positive Correlation |
| LTBP3 | 0.134728 | 0.022675 | Other Positive Correlation |
| TEDC2 | 0.134683 | 0.02272 | Other Positive Correlation |
| TMEFF1 | 0.13466 | 0.022743 | Other Positive Correlation |
| FBXO48 | 0.134585 | 0.022819 | Other Positive Correlation |

| | | | |
|---|---|---|---|
| ERAS | 0.13438 | 0.02303 | Other Positive Correlation |
| VWA5B2 | 0.13423 | 0.023185 | Other Positive Correlation |
| HOXD4 | 0.13421 | 0.023205 | Other Positive Correlation |
| PODN | 0.134176 | 0.02324 | Other Positive Correlation |
| CACNA1G | 0.133803 | 0.02363 | Other Positive Correlation |
| BMP6 | 0.133757 | 0.023679 | Other Positive Correlation |
| KIAA0319 | 0.133661 | 0.02378 | Other Positive Correlation |
| ZBTB10 | 0.133206 | 0.024266 | Other Positive Correlation |
| KIF22 | 0.133201 | 0.024271 | Other Positive Correlation |
| MAGI2 | 0.132946 | 0.024547 | Other Positive Correlation |
| BNC2 | 0.132885 | 0.024613 | Other Positive Correlation |
| NKAIN1 | 0.132819 | 0.024685 | Other Positive Correlation |
| PCSK1N | 0.132683 | 0.024834 | Other Positive Correlation |
| NKX6-2 | 0.132677 | 0.02484 | Other Positive Correlation |
| CCDC136 | 0.132547 | 0.024984 | Other Positive Correlation |
| ARRB2 | 0.132544 | 0.024987 | Other Positive Correlation |
| SMC1B | 0.132368 | 0.025182 | Other Positive Correlation |
| ARMH4 | 0.132212 | 0.025356 | Other Positive Correlation |
| SORCS1 | 0.132177 | 0.025395 | Other Positive Correlation |
| TPRN | 0.132169 | 0.025404 | Other Positive Correlation |
| SHISAL1 | 0.131907 | 0.025699 | Other Positive Correlation |
| OTUD7B | 0.131899 | 0.025708 | Other Positive Correlation |
| PDXP | 0.131891 | 0.025717 | Other Positive Correlation |
| RAB17 | 0.131804 | 0.025815 | Other Positive Correlation |
| WIPF1 | 0.131727 | 0.025902 | Other Positive Correlation |
| RHBG | 0.131644 | 0.025997 | Other Positive Correlation |
| NRN1L | 0.131551 | 0.026104 | Other Positive Correlation |
| SYPL2 | 0.131545 | 0.026111 | Other Positive Correlation |
| GFPT1 | 0.131511 | 0.02615 | Other Positive Correlation |
| LEFTY2 | 0.131481 | 0.026184 | Other Positive Correlation |
| ZNF439 | 0.131453 | 0.026216 | Other Positive Correlation |
| ENSG00000285526 | 0.131365 | 0.026317 | Other Positive Correlation |
| SLC25A33 | 0.131282 | 0.026414 | Other Positive Correlation |
| KCNS3 | 0.131117 | 0.026605 | Other Positive Correlation |
| CRMP1 | 0.130974 | 0.026773 | Other Positive Correlation |
| CRYM | 0.130972 | 0.026774 | Other Positive Correlation |
| TCF24 | 0.130937 | 0.026816 | Other Positive Correlation |
| ALDH1A2 | 0.130683 | 0.027115 | Other Positive Correlation |
| NKX2-6 | 0.130646 | 0.027159 | Other Positive Correlation |
| SYT3 | 0.130614 | 0.027197 | Other Positive Correlation |
| SHD | 0.130531 | 0.027296 | Other Positive Correlation |
| MIIP | 0.130521 | 0.027307 | Other Positive Correlation |
| CTNND2 | 0.13035 | 0.027512 | Other Positive Correlation |
| MED10 | 0.130218 | 0.02767 | Other Positive Correlation |
| MYRIP | 0.13018 | 0.027717 | Other Positive Correlation |
| NAPB | 0.130116 | 0.027794 | Other Positive Correlation |

| | | | |
|---|---|---|---|
| TFF2 | 0.129952 | 0.027993 | Other Positive Correlation |
| PREB | 0.129949 | 0.027997 | Other Positive Correlation |
| ENSG00000286132 | 0.129748 | 0.028242 | Other Positive Correlation |
| FUZ | 0.129666 | 0.028343 | Other Positive Correlation |
| TEX15 | 0.129617 | 0.028403 | Other Positive Correlation |
| LEFTY1 | 0.129542 | 0.028496 | Other Positive Correlation |
| TRH | 0.129536 | 0.028503 | Other Positive Correlation |
| VMO1 | 0.129517 | 0.028526 | Other Positive Correlation |
| CYP26C1 | 0.12931 | 0.028783 | Other Positive Correlation |
| WDR62 | 0.128749 | 0.029489 | Other Positive Correlation |
| SLC38A8 | 0.128665 | 0.029596 | Other Positive Correlation |
| RUNX1T1 | 0.12861 | 0.029667 | Other Positive Correlation |
| TRPV6 | 0.128602 | 0.029678 | Other Positive Correlation |
| WWOX | 0.128502 | 0.029805 | Other Positive Correlation |
| RSPO4 | 0.128394 | 0.029945 | Other Positive Correlation |
| ADCK2 | 0.128375 | 0.029969 | Other Positive Correlation |
| FOXI3 | 0.128345 | 0.030007 | Other Positive Correlation |
| LCORL | 0.128213 | 0.030178 | Other Positive Correlation |
| ROR2 | 0.128187 | 0.030212 | Other Positive Correlation |
| SEC14L5 | 0.127991 | 0.030468 | Other Positive Correlation |
| PRIMA1 | 0.127954 | 0.030516 | Other Positive Correlation |
| INSR | 0.127737 | 0.030802 | Other Positive Correlation |
| FAM163A | 0.127712 | 0.030835 | Other Positive Correlation |
| MPPED2 | 0.127653 | 0.030913 | Other Positive Correlation |
| BEND4 | 0.127534 | 0.031071 | Other Positive Correlation |
| HOXC9 | 0.127287 | 0.031401 | Other Positive Correlation |
| ANKS1B | 0.127067 | 0.031698 | Other Positive Correlation |
| CSRP2 | 0.127031 | 0.031747 | Other Positive Correlation |
| MYBPC2 | 0.12702 | 0.031761 | Other Positive Correlation |
| NKX6-1 | 0.126885 | 0.031946 | Other Positive Correlation |
| NLGN1 | 0.126679 | 0.032227 | Other Positive Correlation |
| EN1 | 0.126614 | 0.032316 | Other Positive Correlation |
| METTL24 | 0.126422 | 0.032582 | Other Positive Correlation |
| LRRC37B | 0.126327 | 0.032712 | Other Positive Correlation |
| GPX8 | 0.12613 | 0.032988 | Other Positive Correlation |
| OR1R1P | 0.125916 | 0.033289 | Other Positive Correlation |
| AMOT | 0.125682 | 0.03362 | Other Positive Correlation |
| XYLT1 | 0.125549 | 0.033809 | Other Positive Correlation |
| CDH22 | 0.125516 | 0.033857 | Other Positive Correlation |
| NGB | 0.125449 | 0.033953 | Other Positive Correlation |
| HSPA12B | 0.12515 | 0.034385 | Other Positive Correlation |
| C5orf34 | 0.12511 | 0.034443 | Other Positive Correlation |
| PRSS36 | 0.125082 | 0.034483 | Other Positive Correlation |
| ZNF492 | 0.124819 | 0.034868 | Other Positive Correlation |
| NAALAD2 | 0.124791 | 0.034909 | Other Positive Correlation |
| DENND4B | 0.12477 | 0.03494 | Other Positive Correlation |

| | | | |
|---|---|---|---|
| RNF150 | 0.124547 | 0.035269 | Other Positive Correlation |
| LIPE | 0.124395 | 0.035494 | Other Positive Correlation |
| TMEM240 | 0.124293 | 0.035646 | Other Positive Correlation |
| CDK1 | 0.124211 | 0.03577 | Other Positive Correlation |
| NPHS1 | 0.124101 | 0.035935 | Other Positive Correlation |
| MESD | 0.124026 | 0.036049 | Other Positive Correlation |
| PLCL1 | 0.123955 | 0.036156 | Other Positive Correlation |
| ANKRD30BL | 0.123764 | 0.036445 | Other Positive Correlation |
| AKNA | 0.123626 | 0.036656 | Other Positive Correlation |
| KCTD16 | 0.123594 | 0.036706 | Other Positive Correlation |
| HMSD | 0.123542 | 0.036784 | Other Positive Correlation |
| IL18BP | 0.123502 | 0.036847 | Other Positive Correlation |
| PTGIS | 0.123341 | 0.037096 | Other Positive Correlation |
| NR5A1 | 0.123118 | 0.037441 | Other Positive Correlation |
| ATP2A3 | 0.123078 | 0.037504 | Other Positive Correlation |
| COMP | 0.122927 | 0.037741 | Other Positive Correlation |
| C2orf50 | 0.122831 | 0.03789 | Other Positive Correlation |
| SRRM3 | 0.122626 | 0.038215 | Other Positive Correlation |
| DBF4B | 0.122535 | 0.03836 | Other Positive Correlation |
| SLC37A4 | 0.122508 | 0.038402 | Other Positive Correlation |
| ENTPD8 | 0.122505 | 0.038407 | Other Positive Correlation |
| ATOH7 | 0.122486 | 0.038438 | Other Positive Correlation |
| MYO3A | 0.12245 | 0.038495 | Other Positive Correlation |
| KCNK13 | 0.122428 | 0.03853 | Other Positive Correlation |
| CRIP1 | 0.122343 | 0.038666 | Other Positive Correlation |
| TMEM37 | 0.1221 | 0.039057 | Other Positive Correlation |
| MAP3K15 | 0.122018 | 0.03919 | Other Positive Correlation |
| SAPCD2 | 0.121972 | 0.039264 | Other Positive Correlation |
| MRM1 | 0.121811 | 0.039526 | Other Positive Correlation |
| RASGRF2 | 0.121801 | 0.039543 | Other Positive Correlation |
| FOXA1 | 0.12179 | 0.039561 | Other Positive Correlation |
| CYP2E1 | 0.121649 | 0.039792 | Other Positive Correlation |
| DEFB134 | 0.121479 | 0.040072 | Other Positive Correlation |
| CACNA1S | 0.121479 | 0.040072 | Other Positive Correlation |
| NPIPA1 | 0.121472 | 0.040083 | Other Positive Correlation |
| ZNF442 | 0.121438 | 0.040139 | Other Positive Correlation |
| PTH1R | 0.121351 | 0.040283 | Other Positive Correlation |
| UCP1 | 0.121087 | 0.040724 | Other Positive Correlation |
| WDR24 | 0.121073 | 0.040747 | Other Positive Correlation |
| CSN1S1 | 0.121042 | 0.040799 | Other Positive Correlation |
| FCAMR | 0.121042 | 0.040799 | Other Positive Correlation |
| IL18RAP | 0.121042 | 0.040799 | Other Positive Correlation |
| DTNBP1 | 0.121029 | 0.040821 | Other Positive Correlation |
| GXYLT1 | 0.120712 | 0.041356 | Other Positive Correlation |
| EGFL6 | 0.120626 | 0.041501 | Other Positive Correlation |
| HBA1 | 0.120625 | 0.041503 | Other Positive Correlation |

| | | | |
|---|---|---|---|
| FAM110B | 0.120619 | 0.041514 | Other Positive Correlation |
| FN3K | 0.120486 | 0.041739 | Other Positive Correlation |
| JAM3 | 0.120456 | 0.041792 | Other Positive Correlation |
| SPINK2 | 0.120349 | 0.041975 | Other Positive Correlation |
| NRXN2 | 0.120194 | 0.042242 | Other Positive Correlation |
| RBM11 | 0.120188 | 0.042252 | Other Positive Correlation |
| COL6A1 | 0.12009 | 0.042422 | Other Positive Correlation |
| CYB561 | 0.119842 | 0.042853 | Other Positive Correlation |
| CCKBR | 0.119833 | 0.042869 | Other Positive Correlation |
| GMNC | 0.119832 | 0.04287 | Other Positive Correlation |
| ENSG00000257355 | 0.119832 | 0.04287 | Other Positive Correlation |
| HRH3 | 0.119666 | 0.043161 | Other Positive Correlation |
| SEMA6B | 0.119662 | 0.043168 | Other Positive Correlation |
| PHOSPHO1 | 0.119648 | 0.043193 | Other Positive Correlation |
| ATP1A3 | 0.119536 | 0.04339 | Other Positive Correlation |
| PKDCC | 0.119397 | 0.043636 | Other Positive Correlation |
| PBOV1 | 0.119395 | 0.04364 | Other Positive Correlation |
| CHST5 | 0.119395 | 0.04364 | Other Positive Correlation |
| SH3RF3 | 0.119307 | 0.043796 | Other Positive Correlation |
| PPP1R9A | 0.11921 | 0.043969 | Other Positive Correlation |
| IQGAP2 | 0.119082 | 0.044199 | Other Positive Correlation |
| GPC2 | 0.118944 | 0.044447 | Other Positive Correlation |
| EFCC1 | 0.118917 | 0.044495 | Other Positive Correlation |
| PPP1R32 | 0.118857 | 0.044603 | Other Positive Correlation |
| TMIE | 0.118808 | 0.044692 | Other Positive Correlation |
| PDE8B | 0.118739 | 0.044818 | Other Positive Correlation |
| SEC14L6 | 0.118721 | 0.04485 | Other Positive Correlation |
| SPSB4 | 0.118717 | 0.044857 | Other Positive Correlation |
| LPO | 0.118716 | 0.044858 | Other Positive Correlation |
| FEV | 0.118686 | 0.044913 | Other Positive Correlation |
| CFAP107 | 0.118686 | 0.044913 | Other Positive Correlation |
| XAGE2 | 0.118684 | 0.044917 | Other Positive Correlation |
| GPR158 | 0.118684 | 0.044917 | Other Positive Correlation |
| CD300C | 0.118684 | 0.044917 | Other Positive Correlation |
| PFN3 | 0.118642 | 0.044992 | Other Positive Correlation |
| CDKL2 | 0.118612 | 0.045048 | Other Positive Correlation |
| ACVRL1 | 0.118495 | 0.045262 | Other Positive Correlation |
| GLI1 | 0.118429 | 0.045382 | Other Positive Correlation |
| EPCAM | 0.118348 | 0.045531 | Other Positive Correlation |
| PRTN3 | 0.11834 | 0.045545 | Other Positive Correlation |
| C2CD4C | 0.118156 | 0.045885 | Other Positive Correlation |
| TRIM69 | 0.118048 | 0.046086 | Other Positive Correlation |
| DCAF12L1 | 0.11803 | 0.046119 | Other Positive Correlation |
| PSD | 0.118009 | 0.046158 | Other Positive Correlation |
| LRRC10B | 0.117968 | 0.046235 | Other Positive Correlation |
| GPT2 | 0.117945 | 0.046277 | Other Positive Correlation |

| | | | |
|---|---|---|---|
| DEFB124 | 0.117687 | 0.046759 | Other Positive Correlation |
| CLCNKA | 0.117687 | 0.046759 | Other Positive Correlation |
| ENSG00000261915 | 0.117674 | 0.046785 | Other Positive Correlation |
| FAM153A | 0.117616 | 0.046895 | Other Positive Correlation |
| ADRA1D | 0.11743 | 0.047245 | Other Positive Correlation |
| MAGED4B | 0.11743 | 0.047245 | Other Positive Correlation |
| BMP3 | 0.11743 | 0.047245 | Other Positive Correlation |
| SLC8A2 | 0.117406 | 0.047291 | Other Positive Correlation |
| CYP2U1 | 0.117232 | 0.047621 | Other Positive Correlation |
| RNPEPL1 | 0.117122 | 0.047833 | Other Positive Correlation |
| OSBPL6 | 0.116975 | 0.048115 | Other Positive Correlation |
| PRSS16 | 0.116887 | 0.048284 | Other Positive Correlation |
| PRSS8 | 0.11683 | 0.048395 | Other Positive Correlation |
| LRRC46 | 0.116699 | 0.048649 | Other Positive Correlation |
| NXNL2 | 0.116675 | 0.048696 | Other Positive Correlation |
| KBTBD11 | 0.116649 | 0.048746 | Other Positive Correlation |
| VGLL2 | 0.116647 | 0.048749 | Other Positive Correlation |
| SMARCA2 | 0.116542 | 0.048954 | Other Positive Correlation |
| CDKN1C | 0.116505 | 0.049027 | Other Positive Correlation |
| DLC1 | 0.116361 | 0.049309 | Other Positive Correlation |
| FAM241B | 0.116296 | 0.049436 | Other Positive Correlation |
| POMC | 0.116084 | 0.049856 | Other Positive Correlation |
| SV2A | 0.11565 | 0.050724 | Other Positive Correlation |
| TLCD2 | 0.115575 | 0.050875 | Other Positive Correlation |
| GCKR | 0.115553 | 0.050919 | Other Positive Correlation |
| C9orf40 | 0.11554 | 0.050945 | Other Positive Correlation |
| MANF | 0.115467 | 0.051094 | Other Positive Correlation |
| ANKRD45 | 0.115462 | 0.051103 | Other Positive Correlation |
| NDNF | 0.115365 | 0.0513 | Other Positive Correlation |
| ATP6V0E2 | 0.115279 | 0.051475 | Other Positive Correlation |
| RASAL1 | 0.115203 | 0.05163 | Other Positive Correlation |
| MUS81 | 0.115076 | 0.051889 | Other Positive Correlation |
| DLX1 | 0.114884 | 0.052285 | Other Positive Correlation |
| SORCS3 | 0.114794 | 0.05247 | Other Positive Correlation |
| DOCK11 | 0.114718 | 0.052627 | Other Positive Correlation |
| MESP2 | 0.114713 | 0.052638 | Other Positive Correlation |
| PRR35 | 0.114663 | 0.052741 | Other Positive Correlation |
| COL9A2 | 0.114567 | 0.052941 | Other Positive Correlation |
| PREX1 | 0.114554 | 0.052968 | Other Positive Correlation |
| FGF17 | 0.114528 | 0.053023 | Other Positive Correlation |
| CERS1 | 0.114526 | 0.053027 | Other Positive Correlation |
| SLC5A10 | 0.114518 | 0.053045 | Other Positive Correlation |
| SMIM34 | 0.11442 | 0.053249 | Other Positive Correlation |
| SHANK1 | 0.114354 | 0.053388 | Other Positive Correlation |
| MED12L | 0.11428 | 0.053542 | Other Positive Correlation |
| CSTL1 | 0.114255 | 0.053596 | Other Positive Correlation |

| | | | |
|---|---|---|---|
| ENKD1 | 0.114088 | 0.053949 | Other Positive Correlation |
| CHRM1 | 0.114018 | 0.054096 | Other Positive Correlation |
| SMG9 | 0.113967 | 0.054205 | Other Positive Correlation |
| LHX1 | 0.113919 | 0.054308 | Other Positive Correlation |
| TRIM61 | 0.113786 | 0.054591 | Other Positive Correlation |
| DUSP8 | 0.11364 | 0.054903 | Other Positive Correlation |
| ZNF568 | 0.113531 | 0.05514 | Other Positive Correlation |
| APBB1IP | 0.113484 | 0.05524 | Other Positive Correlation |
| LRRC73 | 0.113411 | 0.055398 | Other Positive Correlation |
| PPM1L | 0.113385 | 0.055455 | Other Positive Correlation |
| TIMELESS | 0.11327 | 0.055704 | Other Positive Correlation |
| ALPL | 0.113129 | 0.056012 | Other Positive Correlation |
| EML1 | 0.113083 | 0.056112 | Other Positive Correlation |
| DPYSL4 | 0.11307 | 0.05614 | Other Positive Correlation |
| KLHL35 | 0.112969 | 0.056363 | Other Positive Correlation |
| PLEKHB1 | 0.112966 | 0.056368 | Other Positive Correlation |
| SERP1 | 0.112833 | 0.05666 | Other Positive Correlation |
| GFI1 | 0.112832 | 0.056664 | Other Positive Correlation |
| IFNL4 | 0.112692 | 0.056973 | Other Positive Correlation |
| GAL3ST3 | 0.112679 | 0.057003 | Other Positive Correlation |
| RETREG1 | 0.112676 | 0.057009 | Other Positive Correlation |
| P2RX5 | 0.112628 | 0.057115 | Other Positive Correlation |
| PRKAA2 | 0.112533 | 0.057327 | Other Positive Correlation |
| TBC1D28 | 0.112512 | 0.057374 | Other Positive Correlation |
| CORO2B | 0.112434 | 0.057548 | Other Positive Correlation |
| LKAAEAR1 | 0.112411 | 0.057599 | Other Positive Correlation |
| TP53I11 | 0.112293 | 0.057864 | Other Positive Correlation |
| RNF125 | 0.112224 | 0.058018 | Other Positive Correlation |
| SYDE1 | 0.11221 | 0.058051 | Other Positive Correlation |
| GNG7 | 0.112188 | 0.0581 | Other Positive Correlation |
| COL2A1 | 0.112118 | 0.058258 | Other Positive Correlation |
| CNN1 | 0.1121 | 0.058297 | Other Positive Correlation |
| KCNK9 | 0.112089 | 0.058324 | Other Positive Correlation |
| FBLN5 | 0.112045 | 0.058421 | Other Positive Correlation |
| METRNL | 0.111925 | 0.058695 | Other Positive Correlation |
| LHX9 | 0.111742 | 0.059111 | Other Positive Correlation |
| TAS1R1 | 0.111692 | 0.059225 | Other Positive Correlation |
| DMTN | 0.11166 | 0.059299 | Other Positive Correlation |
| SP8 | 0.111178 | 0.06041 | Other Positive Correlation |
| KISS1R | 0.11117 | 0.06043 | Other Positive Correlation |
| COL15A1 | 0.110717 | 0.06149 | Other Positive Correlation |
| YPEL2 | 0.110699 | 0.061533 | Other Positive Correlation |
| MAT1A | 0.110684 | 0.061569 | Other Positive Correlation |
| KBTBD12 | 0.110671 | 0.061599 | Other Positive Correlation |
| ESAM | 0.11064 | 0.061672 | Other Positive Correlation |
| SLC6A17 | 0.110624 | 0.061712 | Other Positive Correlation |

| | | | |
|---|---|---|---|
| UNC5B | 0.110564 | 0.061853 | Other Positive Correlation |
| NPR1 | 0.110473 | 0.062069 | Other Positive Correlation |
| DOLPP1 | 0.110442 | 0.062142 | Other Positive Correlation |
| RPS6KL1 | 0.110433 | 0.062165 | Other Positive Correlation |
| CCDC180 | 0.110356 | 0.062349 | Other Positive Correlation |
| ENSG00000285641 | 0.110347 | 0.062369 | Other Positive Correlation |
| KRT76 | 0.110293 | 0.062498 | Other Positive Correlation |
| FAM184A | 0.110044 | 0.063097 | Other Positive Correlation |
| MADCAM1 | 0.109978 | 0.063255 | Other Positive Correlation |
| TMEM116 | 0.109974 | 0.063266 | Other Positive Correlation |
| GRIN2C | 0.109969 | 0.063279 | Other Positive Correlation |
| CA7 | 0.109966 | 0.063284 | Other Positive Correlation |
| IGIP | 0.10985 | 0.063565 | Other Positive Correlation |
| PLEKHH2 | 0.10936 | 0.064764 | Other Positive Correlation |
| MAN1C1 | 0.109258 | 0.065015 | Other Positive Correlation |
| GAREM2 | 0.109239 | 0.065062 | Other Positive Correlation |
| RASSF8 | 0.109211 | 0.065132 | Other Positive Correlation |
| PKLR | 0.109155 | 0.065271 | Other Positive Correlation |
| AQP5 | 0.109107 | 0.065389 | Other Positive Correlation |
| EDNRB | 0.1089 | 0.065905 | Other Positive Correlation |
| EXOC3L4 | 0.108888 | 0.065934 | Other Positive Correlation |
| NPAS3 | 0.108857 | 0.066012 | Other Positive Correlation |
| TSPAN7 | 0.108639 | 0.066559 | Other Positive Correlation |
| NKX2-8 | 0.108568 | 0.066738 | Other Positive Correlation |
| RFLNB | 0.108531 | 0.066831 | Other Positive Correlation |
| LDLRAD1 | 0.108528 | 0.066839 | Other Positive Correlation |
| ADCY1 | 0.108494 | 0.066924 | Other Positive Correlation |
| SORBS1 | 0.108471 | 0.066983 | Other Positive Correlation |
| TPPP3 | 0.108455 | 0.067024 | Other Positive Correlation |
| TTC6 | 0.108449 | 0.06704 | Other Positive Correlation |
| MPP2 | 0.108443 | 0.067054 | Other Positive Correlation |
| TMEM121 | 0.108091 | 0.06795 | Other Positive Correlation |
| SOX3 | 0.108063 | 0.068024 | Other Positive Correlation |
| FAM221A | 0.107862 | 0.068539 | Other Positive Correlation |
| CHD5 | 0.107794 | 0.068715 | Other Positive Correlation |
| SCN3B | 0.107793 | 0.068719 | Other Positive Correlation |
| IHH | 0.107758 | 0.068809 | Other Positive Correlation |
| CDKL4 | 0.107756 | 0.068815 | Other Positive Correlation |
| GNG4 | 0.107743 | 0.068847 | Other Positive Correlation |
| MFGE8 | 0.10771 | 0.068933 | Other Positive Correlation |
| MCF2L | 0.107605 | 0.069206 | Other Positive Correlation |
| PKHD1L1 | 0.107603 | 0.069211 | Other Positive Correlation |
| GDF6 | 0.107591 | 0.069242 | Other Positive Correlation |
| POLR2J2 | 0.107567 | 0.069303 | Other Positive Correlation |
| SRPK3 | 0.107292 | 0.070024 | Other Positive Correlation |
| CAMK2A | 0.10725 | 0.070133 | Other Positive Correlation |

| | | | |
|---|---|---|---|
| EPHA7 | 0.107245 | 0.070147 | Other Positive Correlation |
| CAMK1D | 0.107223 | 0.070205 | Other Positive Correlation |
| ZNF76 | 0.107167 | 0.070353 | Other Positive Correlation |
| ECSIT | 0.107101 | 0.070527 | Other Positive Correlation |
| PCDHAC2 | 0.107032 | 0.070709 | Other Positive Correlation |
| HOXD11 | 0.106988 | 0.070826 | Other Positive Correlation |
| FOXA3 | 0.106949 | 0.070929 | Other Positive Correlation |
| ZBTB46 | 0.106945 | 0.07094 | Other Positive Correlation |
| PDE10A | 0.106941 | 0.070949 | Other Positive Correlation |
| NKX2-4 | 0.106925 | 0.070992 | Other Positive Correlation |
| ZNF804A | 0.106925 | 0.070992 | Other Positive Correlation |
| FOXO6 | 0.106868 | 0.071143 | Other Positive Correlation |
| CYP11A1 | 0.10682 | 0.071273 | Other Positive Correlation |
| SHCBP1L | 0.106748 | 0.071463 | Other Positive Correlation |
| TNFAIP8L3 | 0.106726 | 0.071523 | Other Positive Correlation |
| GIPC3 | 0.106689 | 0.07162 | Other Positive Correlation |
| OR51M1 | 0.106641 | 0.07175 | Other Positive Correlation |
| LINGO2 | 0.106568 | 0.071945 | Other Positive Correlation |
| HLA-DRB5 | 0.106516 | 0.072084 | Other Positive Correlation |
| ARHGEF6 | 0.1065 | 0.072128 | Other Positive Correlation |
| C2orf72 | 0.106491 | 0.072151 | Other Positive Correlation |
| OSGIN1 | 0.106201 | 0.072935 | Other Positive Correlation |
| STH | 0.106117 | 0.073165 | Other Positive Correlation |
| GALNT17 | 0.106117 | 0.073165 | Other Positive Correlation |
| SRRM4 | 0.106117 | 0.073165 | Other Positive Correlation |
| GALNTL6 | 0.106117 | 0.073165 | Other Positive Correlation |
| SFRP4 | 0.106117 | 0.073165 | Other Positive Correlation |
| TMEM215 | 0.106117 | 0.073165 | Other Positive Correlation |
| WFIKKN2 | 0.106117 | 0.073165 | Other Positive Correlation |
| TRMT9B | 0.106071 | 0.073289 | Other Positive Correlation |
| IGSF10 | 0.106017 | 0.073437 | Other Positive Correlation |
| ZACN | 0.105972 | 0.073561 | Other Positive Correlation |
| SLC2A4 | 0.105928 | 0.07368 | Other Positive Correlation |
| LVRN | 0.105869 | 0.073842 | Other Positive Correlation |
| STRBP | 0.105855 | 0.073879 | Other Positive Correlation |
| CPXM2 | 0.105755 | 0.074154 | Other Positive Correlation |
| FAAP24 | 0.105741 | 0.074194 | Other Positive Correlation |
| EIF4E1B | 0.105714 | 0.074266 | Other Positive Correlation |
| TAS2R3 | 0.105707 | 0.074287 | Other Positive Correlation |
| KCNA4 | 0.105707 | 0.074287 | Other Positive Correlation |
| ATXN1L | 0.10563 | 0.0745 | Other Positive Correlation |
| ZNF598 | 0.105547 | 0.074728 | Other Positive Correlation |
| FANCI | 0.105486 | 0.074898 | Other Positive Correlation |
| GPAT2 | 0.105339 | 0.075306 | Other Positive Correlation |
| FGF18 | 0.105271 | 0.075497 | Other Positive Correlation |
| GABRB2 | 0.105123 | 0.07591 | Other Positive Correlation |

| | | | |
|---|---|---|---|
| NPBWR1 | 0.105078 | 0.076035 | Other Positive Correlation |
| NHLH1 | 0.105032 | 0.076165 | Other Positive Correlation |
| ODAD2 | 0.105006 | 0.076237 | Other Positive Correlation |
| NDRG4 | 0.104946 | 0.076409 | Other Positive Correlation |
| GAD1 | 0.104779 | 0.076879 | Other Positive Correlation |
| RHOU | 0.104713 | 0.077068 | Other Positive Correlation |
| SELENOS | 0.1045 | 0.077674 | Other Positive Correlation |
| GPR27 | 0.104272 | 0.078327 | Other Positive Correlation |
| ZNF728 | 0.10425 | 0.078389 | Other Positive Correlation |
| KLHL15 | 0.104202 | 0.078528 | Other Positive Correlation |
| PLXDC1 | 0.104111 | 0.07879 | Other Positive Correlation |
| DCUN1D2 | 0.104086 | 0.078862 | Other Positive Correlation |
| DMRT2 | 0.104079 | 0.078885 | Other Positive Correlation |
| ARPP21 | 0.103982 | 0.079164 | Other Positive Correlation |
| HSPB7 | 0.103982 | 0.079164 | Other Positive Correlation |
| SKOR2 | 0.103982 | 0.079164 | Other Positive Correlation |
| LIN7A | 0.103982 | 0.079164 | Other Positive Correlation |
| NXPH2 | 0.103982 | 0.079164 | Other Positive Correlation |
| OTUD6A | 0.103792 | 0.079718 | Other Positive Correlation |
| FNDC5 | 0.10368 | 0.080045 | Other Positive Correlation |
| GLT1D1 | 0.103333 | 0.081064 | Other Positive Correlation |
| PLCXD3 | 0.10324 | 0.081341 | Other Positive Correlation |
| FAM171A2 | 0.103233 | 0.08136 | Other Positive Correlation |
| BRSK2 | 0.103135 | 0.081653 | Other Positive Correlation |
| TNNI3 | 0.103026 | 0.081977 | Other Positive Correlation |
| ISLR2 | 0.102922 | 0.082286 | Other Positive Correlation |
| ANKRD63 | 0.102866 | 0.082456 | Other Positive Correlation |
| LOXL4 | 0.102816 | 0.082606 | Other Positive Correlation |
| SYNE3 | 0.102692 | 0.08298 | Other Positive Correlation |
| KLHL14 | 0.102672 | 0.083038 | Other Positive Correlation |
| PGLYRP2 | 0.102672 | 0.083038 | Other Positive Correlation |
| FAM133A | 0.102672 | 0.083038 | Other Positive Correlation |
| SLC22A10 | 0.102672 | 0.083038 | Other Positive Correlation |
| SARDH | 0.102642 | 0.083128 | Other Positive Correlation |
| RNF144A | 0.102376 | 0.083936 | Other Positive Correlation |
| NPRL3 | 0.10237 | 0.083953 | Other Positive Correlation |
| TUBB1 | 0.102355 | 0.083998 | Other Positive Correlation |
| ZNF98 | 0.102351 | 0.084012 | Other Positive Correlation |
| VIPR2 | 0.10229 | 0.084197 | Other Positive Correlation |
| MIOX | 0.102218 | 0.084415 | Other Positive Correlation |
| HVCN1 | 0.102056 | 0.084914 | Other Positive Correlation |
| LDHD | 0.102048 | 0.084938 | Other Positive Correlation |
| SLC22A17 | 0.101958 | 0.085214 | Other Positive Correlation |
| DNAH6 | 0.101954 | 0.085225 | Other Positive Correlation |
| CCDC160 | 0.101806 | 0.085684 | Other Positive Correlation |
| ANKH | 0.101787 | 0.085741 | Other Positive Correlation |

| | | | |
|---|---|---|---|
| SNCAIP | 0.10161 | 0.086291 | Other Positive Correlation |
| SLC15A1 | 0.10153 | 0.08654 | Other Positive Correlation |
| VASH1 | 0.101294 | 0.087278 | Other Positive Correlation |
| GRM4 | 0.10123 | 0.08748 | Other Positive Correlation |
| THPO | 0.101223 | 0.087502 | Other Positive Correlation |
| TP53TG5 | 0.100969 | 0.088302 | Other Positive Correlation |
| POU2AF3 | 0.100812 | 0.088801 | Other Positive Correlation |
| COL13A1 | 0.100769 | 0.088939 | Other Positive Correlation |
| GFRA4 | 0.100734 | 0.08905 | Other Positive Correlation |
| PI4KA | 0.100696 | 0.089171 | Other Positive Correlation |
| ST7 | 0.100692 | 0.089184 | Other Positive Correlation |
| ALPK2 | 0.100564 | 0.089592 | Other Positive Correlation |
| WNT3 | 0.100516 | 0.089748 | Other Positive Correlation |
| ENSG00000230707 | 0.100446 | 0.089971 | Other Positive Correlation |
| ZNF850 | 0.100408 | 0.090095 | Other Positive Correlation |
| FXYD4 | 0.100358 | 0.090254 | Other Positive Correlation |
| CHST9 | 0.100285 | 0.090489 | Other Positive Correlation |
| HAPLN2 | 0.100244 | 0.090623 | Other Positive Correlation |
| BEX1 | 0.100228 | 0.090674 | Other Positive Correlation |
| CACNG8 | 0.100203 | 0.090755 | Other Positive Correlation |
| CELSR3 | 0.10006 | 0.091218 | Other Positive Correlation |
| MATN4 | 0.100046 | 0.091267 | Other Positive Correlation |
| S100PBP | 0.09995 | 0.091578 | Other Positive Correlation |
| PAQR9 | 0.099672 | 0.092489 | Other Positive Correlation |
| TPTEP2-CSNK1E | 0.099663 | 0.092518 | Other Positive Correlation |
| TM6SF2 | 0.099635 | 0.092611 | Other Positive Correlation |
| SASS6 | 0.099615 | 0.092674 | Other Positive Correlation |
| GVQW3 | 0.099591 | 0.092755 | Other Positive Correlation |
| ENPP4 | 0.099518 | 0.092994 | Other Positive Correlation |
| PCYT1B | 0.099404 | 0.093372 | Other Positive Correlation |
| ZFP41 | 0.09938 | 0.093453 | Other Positive Correlation |
| HAND1 | 0.099243 | 0.093907 | Other Positive Correlation |
| CCDC74A | 0.098967 | 0.09483 | Other Positive Correlation |
| SYNE1 | 0.098914 | 0.095008 | Other Positive Correlation |
| MDGA2 | 0.098905 | 0.095039 | Other Positive Correlation |
| PATE2 | 0.098782 | 0.09545 | Other Positive Correlation |
| PRKCB | 0.098547 | 0.096246 | Other Positive Correlation |
| NEURL1 | 0.098323 | 0.097008 | Other Positive Correlation |
| ZNF92 | 0.098271 | 0.097187 | Other Positive Correlation |
| PAQR8 | 0.098198 | 0.097436 | Other Positive Correlation |
| HLTF | 0.09812 | 0.097702 | Other Positive Correlation |
| ISLR | 0.097851 | 0.098629 | Other Positive Correlation |
| TIMM10 | 0.097826 | 0.098718 | Other Positive Correlation |
| GATM | 0.097791 | 0.098838 | Other Positive Correlation |
| SDK1 | 0.097786 | 0.098855 | Other Positive Correlation |
| CKS2 | 0.097694 | 0.099175 | Other Positive Correlation |

| | | | |
|---|---|---|---|
| JAKMIP2 | 0.097598 | 0.099507 | Other Positive Correlation |
| SEMA6D | 0.097592 | 0.099531 | Other Positive Correlation |
| NELL2 | 0.097577 | 0.099582 | Other Positive Correlation |
| IGSF1 | 0.09757 | 0.099607 | Other Positive Correlation |
| RIMOC1 | 0.097489 | 0.099888 | Other Positive Correlation |
| IGLON5 | 0.097479 | 0.099923 | Other Positive Correlation |
| SLC39A7 | 0.097373 | 0.100297 | Other Positive Correlation |
| ZNF829 | 0.097263 | 0.10068 | Other Positive Correlation |
| RASL10A | 0.09712 | 0.101187 | Other Positive Correlation |
| MTRFR | 0.097048 | 0.10144 | Other Positive Correlation |
| KCNK10 | 0.097045 | 0.10145 | Other Positive Correlation |
| KCNAB3 | 0.096952 | 0.101782 | Other Positive Correlation |
| ANKRD52 | 0.096913 | 0.101919 | Other Positive Correlation |
| FES | 0.096773 | 0.102416 | Other Positive Correlation |
| TUBB4A | 0.096614 | 0.102987 | Other Positive Correlation |
| FOXO3B | 0.096547 | 0.103224 | Other Positive Correlation |
| CFAP74 | 0.096542 | 0.103242 | Other Positive Correlation |
| SHH | 0.096534 | 0.103272 | Other Positive Correlation |
| OTULINL | 0.096494 | 0.103416 | Other Positive Correlation |
| SPAG6 | 0.096469 | 0.103504 | Other Positive Correlation |
| FGF2 | 0.096458 | 0.103545 | Other Positive Correlation |
| HOXB2 | 0.096376 | 0.103841 | Other Positive Correlation |
| SYP | 0.096361 | 0.103892 | Other Positive Correlation |
| DOHH | 0.096338 | 0.103977 | Other Positive Correlation |
| GPRIN2 | 0.096328 | 0.104014 | Other Positive Correlation |
| MT1G | 0.096276 | 0.1042 | Other Positive Correlation |
| C19orf25 | 0.096225 | 0.104384 | Other Positive Correlation |
| MAST1 | 0.095958 | 0.105353 | Other Positive Correlation |
| MTTP | 0.095958 | 0.105355 | Other Positive Correlation |
| GSC | 0.09594 | 0.105419 | Other Positive Correlation |
| SYT9 | 0.095891 | 0.1056 | Other Positive Correlation |
| ERFE | 0.095834 | 0.105808 | Other Positive Correlation |
| SCARF2 | 0.095773 | 0.10603 | Other Positive Correlation |
| SYNM | 0.095416 | 0.107344 | Other Positive Correlation |
| LDLRAD4 | 0.095304 | 0.10776 | Other Positive Correlation |
| ENSG00000284337 | 0.094589 | 0.110441 | Other Positive Correlation |
| PLXND1 | 0.094586 | 0.11045 | Other Positive Correlation |
| MGAT3 | 0.094526 | 0.110678 | Other Positive Correlation |
| TADA2B | 0.094521 | 0.110698 | Other Positive Correlation |
| KANSL1L | 0.094383 | 0.111222 | Other Positive Correlation |
| RFLNA | 0.094378 | 0.11124 | Other Positive Correlation |
| CDO1 | 0.094364 | 0.111295 | Other Positive Correlation |
| FXYD1 | 0.094363 | 0.111299 | Other Positive Correlation |
| ARHGAP36 | 0.094253 | 0.111718 | Other Positive Correlation |
| GUCY1B1 | 0.094233 | 0.111791 | Other Positive Correlation |
| GABRA5 | 0.094126 | 0.112204 | Other Positive Correlation |

| | | | |
|---|---|---|---|
| VAX1 | 0.094026 | 0.112585 | Other Positive Correlation |
| LIN54 | 0.094003 | 0.112674 | Other Positive Correlation |
| FSTL4 | 0.093989 | 0.11273 | Other Positive Correlation |
| HGD | 0.093899 | 0.113073 | Other Positive Correlation |
| ANKRD44 | 0.093821 | 0.113376 | Other Positive Correlation |
| LRRC23 | 0.093804 | 0.113441 | Other Positive Correlation |
| ASCL2 | 0.093625 | 0.114133 | Other Positive Correlation |
| MPPED1 | 0.093554 | 0.11441 | Other Positive Correlation |
| GTPBP2 | 0.093538 | 0.114472 | Other Positive Correlation |
| MACO1 | 0.09349 | 0.114658 | Other Positive Correlation |
| NKX2-3 | 0.093467 | 0.11475 | Other Positive Correlation |
| HS3ST6 | 0.093437 | 0.114865 | Other Positive Correlation |
| ADAM33 | 0.093395 | 0.11503 | Other Positive Correlation |
| HPCA | 0.093301 | 0.115398 | Other Positive Correlation |
| APC2 | 0.093296 | 0.115416 | Other Positive Correlation |
| PNMA3 | 0.093292 | 0.115434 | Other Positive Correlation |
| KCTD13 | 0.093178 | 0.115878 | Other Positive Correlation |
| PCBP3 | 0.093144 | 0.116013 | Other Positive Correlation |
| IRX6 | 0.093119 | 0.116111 | Other Positive Correlation |
| TFAP2B | 0.093114 | 0.116132 | Other Positive Correlation |
| EIF4EBP3 | 0.0931 | 0.116188 | Other Positive Correlation |
| VPS37D | 0.093095 | 0.116205 | Other Positive Correlation |
| SLC25A4 | 0.09306 | 0.116342 | Other Positive Correlation |
| ASTL | 0.092978 | 0.116667 | Other Positive Correlation |
| PRTG | 0.092961 | 0.116734 | Other Positive Correlation |
| DDX25 | 0.092958 | 0.116748 | Other Positive Correlation |
| F7 | 0.092897 | 0.116988 | Other Positive Correlation |
| DGKE | 0.092853 | 0.117161 | Other Positive Correlation |
| HIPK4 | 0.092658 | 0.117938 | Other Positive Correlation |
| ZNF711 | 0.092614 | 0.118114 | Other Positive Correlation |
| SBK1 | 0.092584 | 0.118232 | Other Positive Correlation |
| CA4 | 0.092493 | 0.118594 | Other Positive Correlation |
| CFAP77 | 0.09246 | 0.118729 | Other Positive Correlation |
| HRH2 | 0.092433 | 0.118837 | Other Positive Correlation |
| IL12RB1 | 0.092417 | 0.118901 | Other Positive Correlation |
| PHF21B | 0.092261 | 0.119525 | Other Positive Correlation |
| RNF114 | 0.092244 | 0.119596 | Other Positive Correlation |
| INPP5F | 0.092232 | 0.119644 | Other Positive Correlation |
| CA3 | 0.092173 | 0.11988 | Other Positive Correlation |
| RIIAD1 | 0.092147 | 0.119985 | Other Positive Correlation |
| ADAM11 | 0.092116 | 0.12011 | Other Positive Correlation |
| FAM78A | 0.091973 | 0.12069 | Other Positive Correlation |
| CDC7 | 0.091706 | 0.121778 | Other Positive Correlation |
| ASIC3 | 0.091622 | 0.12212 | Other Positive Correlation |
| PRR19 | 0.091512 | 0.122572 | Other Positive Correlation |
| SEMA3B | 0.091463 | 0.122771 | Other Positive Correlation |

| | | | |
|---|---|---|---|
| TCAP | 0.091427 | 0.122921 | Other Positive Correlation |
| PMP22 | 0.091383 | 0.123101 | Other Positive Correlation |
| TESK1 | 0.091296 | 0.123462 | Other Positive Correlation |
| ZNF678 | 0.091119 | 0.124191 | Other Positive Correlation |
| ICAM5 | 0.091084 | 0.124336 | Other Positive Correlation |
| EOMES | 0.091064 | 0.124419 | Other Positive Correlation |
| L3MBTL4 | 0.090853 | 0.125299 | Other Positive Correlation |
| BEX4 | 0.090837 | 0.125366 | Other Positive Correlation |
| TSPOAP1 | 0.090711 | 0.125893 | Other Positive Correlation |
| VSTM4 | 0.090681 | 0.126017 | Other Positive Correlation |
| PROX1 | 0.090659 | 0.12611 | Other Positive Correlation |
| QRFP | 0.090623 | 0.126262 | Other Positive Correlation |
| LCN15 | 0.090623 | 0.126262 | Other Positive Correlation |
| SYNDIG1 | 0.090623 | 0.126262 | Other Positive Correlation |
| POLA2 | 0.090583 | 0.126432 | Other Positive Correlation |
| CDC25C | 0.090477 | 0.126878 | Other Positive Correlation |
| DCDC2C | 0.09025 | 0.127836 | Other Positive Correlation |
| CLEC20A | 0.090066 | 0.12862 | Other Positive Correlation |
| FAM166C | 0.090062 | 0.128636 | Other Positive Correlation |
| TEX19 | 0.089976 | 0.129004 | Other Positive Correlation |
| FCRLB | 0.089953 | 0.129099 | Other Positive Correlation |
| GPR63 | 0.089885 | 0.129392 | Other Positive Correlation |
| VPS37C | 0.089811 | 0.129708 | Other Positive Correlation |
| TTC28 | 0.089785 | 0.129819 | Other Positive Correlation |
| MFAP4 | 0.089784 | 0.129824 | Other Positive Correlation |
| SLC24A2 | 0.089613 | 0.130559 | Other Positive Correlation |
| RAPH1 | 0.089609 | 0.130578 | Other Positive Correlation |
| RANBP3L | 0.089362 | 0.131645 | Other Positive Correlation |
| NOTO | 0.089173 | 0.132464 | Other Positive Correlation |
| GCC1 | 0.088969 | 0.133358 | Other Positive Correlation |
| TRAPPC9 | 0.088968 | 0.133364 | Other Positive Correlation |
| KCNA7 | 0.088827 | 0.133983 | Other Positive Correlation |
| RAB6D | 0.088824 | 0.133996 | Other Positive Correlation |
| SLIT3 | 0.088725 | 0.13443 | Other Positive Correlation |
| EFNA2 | 0.088712 | 0.134487 | Other Positive Correlation |
| ALX3 | 0.088552 | 0.135192 | Other Positive Correlation |
| BRSK1 | 0.088542 | 0.135237 | Other Positive Correlation |
| IQSEC3 | 0.088468 | 0.135566 | Other Positive Correlation |
| ELFN1 | 0.088295 | 0.136335 | Other Positive Correlation |
| ESX1 | 0.08803 | 0.137521 | Other Positive Correlation |
| TMEM130 | 0.087939 | 0.137931 | Other Positive Correlation |
| SNX11 | 0.087809 | 0.138513 | Other Positive Correlation |
| CCR2 | 0.087795 | 0.138579 | Other Positive Correlation |
| CGB3 | 0.087795 | 0.138579 | Other Positive Correlation |
| PNLIPRP2 | 0.087795 | 0.138579 | Other Positive Correlation |
| PI16 | 0.087795 | 0.138579 | Other Positive Correlation |

| | | | |
|---|---|---|---|
| GPC3 | 0.087435 | 0.140211 | Other Positive Correlation |
| PLPP7 | 0.087409 | 0.140329 | Other Positive Correlation |
| MME | 0.08737 | 0.140505 | Other Positive Correlation |
| BEGAIN | 0.087269 | 0.140969 | Other Positive Correlation |
| AMPH | 0.087264 | 0.14099 | Other Positive Correlation |
| METTL21A | 0.08725 | 0.141054 | Other Positive Correlation |
| ENSG00000286165 | 0.08717 | 0.141422 | Other Positive Correlation |
| CLEC2L | 0.087165 | 0.141441 | Other Positive Correlation |
| SCN11A | 0.087136 | 0.141578 | Other Positive Correlation |
| SORT1 | 0.087049 | 0.141975 | Other Positive Correlation |
| REM1 | 0.086929 | 0.142529 | Other Positive Correlation |
| PRDM12 | 0.086826 | 0.143002 | Other Positive Correlation |
| TRIM36 | 0.086752 | 0.143344 | Other Positive Correlation |
| KCNK4 | 0.086721 | 0.143489 | Other Positive Correlation |
| PSMB11 | 0.086407 | 0.144951 | Other Positive Correlation |
| AVPR1B | 0.086407 | 0.144951 | Other Positive Correlation |
| ERFL | 0.086407 | 0.144951 | Other Positive Correlation |
| ZNF648 | 0.086407 | 0.144951 | Other Positive Correlation |
| ACTN3 | 0.086407 | 0.144951 | Other Positive Correlation |
| LRRC2 | 0.086401 | 0.144978 | Other Positive Correlation |
| SMCO1 | 0.086309 | 0.145408 | Other Positive Correlation |
| TAL2 | 0.086273 | 0.145577 | Other Positive Correlation |
| KIF12 | 0.086227 | 0.145792 | Other Positive Correlation |
| SLC22A16 | 0.08617 | 0.14606 | Other Positive Correlation |
| NEGR1 | 0.085895 | 0.147356 | Other Positive Correlation |
| FAM219A | 0.085883 | 0.147415 | Other Positive Correlation |
| ZBTB14 | 0.085824 | 0.147693 | Other Positive Correlation |
| UBE2QL1 | 0.085745 | 0.148068 | Other Positive Correlation |
| DPPA5 | 0.085745 | 0.148068 | Other Positive Correlation |
| NMBR | 0.085745 | 0.148068 | Other Positive Correlation |
| ENSG00000256206 | 0.085745 | 0.148068 | Other Positive Correlation |
| DLK1 | 0.085745 | 0.148068 | Other Positive Correlation |
| DNAJC27 | 0.085737 | 0.148107 | Other Positive Correlation |
| KCNQ4 | 0.085631 | 0.148608 | Other Positive Correlation |
| ADGRB1 | 0.085584 | 0.148834 | Other Positive Correlation |
| SLC26A1 | 0.085573 | 0.148885 | Other Positive Correlation |
| PHKG2 | 0.085555 | 0.148969 | Other Positive Correlation |
| SNTG2 | 0.085494 | 0.149263 | Other Positive Correlation |
| SLC16A8 | 0.085443 | 0.149504 | Other Positive Correlation |
| CACNB2 | 0.085298 | 0.1502 | Other Positive Correlation |
| OSGIN2 | 0.08499 | 0.151685 | Other Positive Correlation |
| FOXA2 | 0.084954 | 0.151858 | Other Positive Correlation |
| RGCC | 0.084913 | 0.152056 | Other Positive Correlation |
| ACBD7 | 0.084849 | 0.152368 | Other Positive Correlation |
| RAD54L | 0.084757 | 0.152814 | Other Positive Correlation |
| CALY | 0.08469 | 0.15314 | Other Positive Correlation |

| | | | |
|---|---|---|---|
| HS3ST3A1 | 0.084576 | 0.153696 | Other Positive Correlation |
| PPM1H | 0.084545 | 0.153847 | Other Positive Correlation |
| MAP3K9 | 0.084374 | 0.154683 | Other Positive Correlation |
| CLSTN2 | 0.084347 | 0.154815 | Other Positive Correlation |
| NOS2 | 0.08431 | 0.154996 | Other Positive Correlation |
| WDR86 | 0.084123 | 0.155919 | Other Positive Correlation |
| PRM3 | 0.083995 | 0.156553 | Other Positive Correlation |
| PTP4A2 | 0.083706 | 0.157985 | Other Positive Correlation |
| ZNF730 | 0.08369 | 0.158065 | Other Positive Correlation |
| NQO2 | 0.083683 | 0.1581 | Other Positive Correlation |
| ZNF610 | 0.083614 | 0.158448 | Other Positive Correlation |
| EFNB3 | 0.083591 | 0.158562 | Other Positive Correlation |
| EFCAB13 | 0.083535 | 0.158842 | Other Positive Correlation |
| PCP4 | 0.083517 | 0.158932 | Other Positive Correlation |
| FRMPD3 | 0.083517 | 0.158932 | Other Positive Correlation |
| IFNL3 | 0.083517 | 0.158932 | Other Positive Correlation |
| CFAP91 | 0.083517 | 0.158932 | Other Positive Correlation |
| LRRC63 | 0.083517 | 0.158932 | Other Positive Correlation |
| HLX | 0.083508 | 0.158977 | Other Positive Correlation |
| LGI3 | 0.083496 | 0.159035 | Other Positive Correlation |
| DNAJA3 | 0.083398 | 0.159527 | Other Positive Correlation |
| CXCL12 | 0.083388 | 0.159579 | Other Positive Correlation |
| PTGER1 | 0.08337 | 0.159669 | Other Positive Correlation |
| YOD1 | 0.083354 | 0.159746 | Other Positive Correlation |
| UNC5A | 0.083286 | 0.160092 | Other Positive Correlation |
| SMIM24 | 0.083271 | 0.160167 | Other Positive Correlation |
| WNT5A | 0.083171 | 0.160669 | Other Positive Correlation |
| LRRC8B | 0.083113 | 0.160963 | Other Positive Correlation |
| DBF4 | 0.083022 | 0.161425 | Other Positive Correlation |
| WNT11 | 0.082999 | 0.16154 | Other Positive Correlation |
| SEZ6 | 0.082994 | 0.161566 | Other Positive Correlation |
| CCDC77 | 0.082894 | 0.162073 | Other Positive Correlation |
| CH25H | 0.082795 | 0.162579 | Other Positive Correlation |
| INPP5J | 0.082783 | 0.162639 | Other Positive Correlation |
| PIGQ | 0.082747 | 0.162823 | Other Positive Correlation |
| ENSG00000285329 | 0.082694 | 0.163093 | Other Positive Correlation |
| MBD3L2B | 0.082694 | 0.163093 | Other Positive Correlation |
| KL | 0.082632 | 0.163411 | Other Positive Correlation |
| PON3 | 0.082626 | 0.163438 | Other Positive Correlation |
| DACH1 | 0.082505 | 0.16406 | Other Positive Correlation |
| CPT1C | 0.082468 | 0.16425 | Other Positive Correlation |
| ENSG00000285932 | 0.082455 | 0.164317 | Other Positive Correlation |
| IL17D | 0.082453 | 0.164327 | Other Positive Correlation |
| CLEC4A | 0.082426 | 0.164467 | Other Positive Correlation |
| KATNB1 | 0.082253 | 0.165355 | Other Positive Correlation |
| LSM14B | 0.082166 | 0.165802 | Other Positive Correlation |

| | | | |
|---|---|---|---|
| MVB12B | 0.0821 | 0.166147 | Other Positive Correlation |
| TNNC2 | 0.081961 | 0.166864 | Other Positive Correlation |
| GDNF | 0.081949 | 0.166929 | Other Positive Correlation |
| AOC2 | 0.081921 | 0.167076 | Other Positive Correlation |
| DLEU7 | 0.081915 | 0.167107 | Other Positive Correlation |
| SLC8A3 | 0.08189 | 0.167237 | Other Positive Correlation |
| FOXRED1 | 0.081852 | 0.167435 | Other Positive Correlation |
| DLGAP3 | 0.081792 | 0.167746 | Other Positive Correlation |
| STKLD1 | 0.081698 | 0.168239 | Other Positive Correlation |
| ST8SIA5 | 0.081625 | 0.168617 | Other Positive Correlation |
| PRIM2 | 0.081355 | 0.170036 | Other Positive Correlation |
| SBK2 | 0.081285 | 0.170403 | Other Positive Correlation |
| HOXC8 | 0.081247 | 0.170607 | Other Positive Correlation |
| PRR16 | 0.081222 | 0.170738 | Other Positive Correlation |
| ALX1 | 0.081217 | 0.170767 | Other Positive Correlation |
| ESCO2 | 0.081154 | 0.171096 | Other Positive Correlation |
| AMDHD2 | 0.081148 | 0.171128 | Other Positive Correlation |
| ZNF714 | 0.081099 | 0.171388 | Other Positive Correlation |
| SERPING1 | 0.081023 | 0.17179 | Other Positive Correlation |
| VAV3 | 0.080989 | 0.171971 | Other Positive Correlation |
| DOK1 | 0.080987 | 0.171984 | Other Positive Correlation |
| ECRG4 | 0.080906 | 0.172413 | Other Positive Correlation |
| ZNF843 | 0.080841 | 0.172762 | Other Positive Correlation |
| ZNF844 | 0.080834 | 0.172796 | Other Positive Correlation |
| KMT5A | 0.080808 | 0.172936 | Other Positive Correlation |
| SLC34A2 | 0.080708 | 0.173468 | Other Positive Correlation |
| ARHGEF39 | 0.080659 | 0.17373 | Other Positive Correlation |
| GCK | 0.080633 | 0.173873 | Other Positive Correlation |
| MFHAS1 | 0.080605 | 0.174023 | Other Positive Correlation |
| PLA2G10 | 0.080378 | 0.17524 | Other Positive Correlation |
| CENPL | 0.080293 | 0.175696 | Other Positive Correlation |
| ZBTB48 | 0.080148 | 0.176483 | Other Positive Correlation |
| ZNF680 | 0.08014 | 0.176524 | Other Positive Correlation |
| ENSG00000269533 | 0.080119 | 0.176641 | Other Positive Correlation |
| HOXC12 | 0.080119 | 0.176641 | Other Positive Correlation |
| ERICH6 | 0.080119 | 0.176641 | Other Positive Correlation |
| DNAAF6 | 0.080119 | 0.176641 | Other Positive Correlation |
| NR5A2 | 0.080119 | 0.176641 | Other Positive Correlation |
| MAMLD1 | 0.080079 | 0.176855 | Other Positive Correlation |
| KHDC1 | 0.080044 | 0.177048 | Other Positive Correlation |
| CD247 | 0.080023 | 0.177162 | Other Positive Correlation |
| GOLGA6L10 | 0.079951 | 0.17755 | Other Positive Correlation |
| ZBTB7C | 0.079939 | 0.177616 | Other Positive Correlation |
| ESYT3 | 0.079828 | 0.178224 | Other Positive Correlation |
| RSPO1 | 0.079795 | 0.1784 | Other Positive Correlation |
| EPB41L1 | 0.079762 | 0.178582 | Other Positive Correlation |

| | | | |
|---|---|---|---|
| FZD9 | 0.07972 | 0.178811 | Other Positive Correlation |
| DPP7 | 0.079568 | 0.179643 | Other Positive Correlation |
| KLHDC8A | 0.079537 | 0.179814 | Other Positive Correlation |
| RECQL4 | 0.079223 | 0.181545 | Other Positive Correlation |
| CLCC1 | 0.079145 | 0.181977 | Other Positive Correlation |
| C11orf24 | 0.079097 | 0.18224 | Other Positive Correlation |
| LYSMD1 | 0.07905 | 0.182503 | Other Positive Correlation |
| PTCH1 | 0.079006 | 0.182749 | Other Positive Correlation |
| TNFRSF11A | 0.078904 | 0.183317 | Other Positive Correlation |
| NIF3L1 | 0.078887 | 0.183407 | Other Positive Correlation |
| PASK | 0.078763 | 0.184102 | Other Positive Correlation |
| COCH | 0.07871 | 0.184394 | Other Positive Correlation |
| ZFHX3 | 0.078615 | 0.18493 | Other Positive Correlation |
| GSTT2 | 0.078585 | 0.185098 | Other Positive Correlation |
| MIEF1 | 0.078549 | 0.185298 | Other Positive Correlation |
| NTF3 | 0.078541 | 0.185342 | Other Positive Correlation |
| ZSCAN18 | 0.078533 | 0.185391 | Other Positive Correlation |
| NUCB2 | 0.078455 | 0.185828 | Other Positive Correlation |
| KLF2 | 0.078313 | 0.186629 | Other Positive Correlation |
| CD72 | 0.078309 | 0.18665 | Other Positive Correlation |
| UVSSA | 0.07823 | 0.187096 | Other Positive Correlation |
| SAYSD1 | 0.078183 | 0.18736 | Other Positive Correlation |
| HLA-DMB | 0.078178 | 0.187391 | Other Positive Correlation |
| ATP8A1 | 0.078178 | 0.187392 | Other Positive Correlation |
| EPHA10 | 0.078066 | 0.188025 | Other Positive Correlation |
| CLTCL1 | 0.078045 | 0.188146 | Other Positive Correlation |
| CABP1 | 0.078017 | 0.188304 | Other Positive Correlation |
| IL5 | 0.077999 | 0.188403 | Other Positive Correlation |
| APBA1 | 0.077985 | 0.188483 | Other Positive Correlation |
| CABLES1 | 0.07794 | 0.18874 | Other Positive Correlation |
| SMIM45 | 0.077895 | 0.188996 | Other Positive Correlation |
| SLC17A7 | 0.077797 | 0.189556 | Other Positive Correlation |
| PTGDS | 0.077681 | 0.19022 | Other Positive Correlation |
| PNPLA5 | 0.077676 | 0.190246 | Other Positive Correlation |
| PPCDC | 0.077654 | 0.190372 | Other Positive Correlation |
| MAP7D2 | 0.077652 | 0.190387 | Other Positive Correlation |
| RIMBP2 | 0.077624 | 0.190542 | Other Positive Correlation |
| ARHGAP28 | 0.077618 | 0.190582 | Other Positive Correlation |
| GRIN3A | 0.077532 | 0.191073 | Other Positive Correlation |
| TEN1 | 0.077442 | 0.191592 | Other Positive Correlation |
| USP1 | 0.077429 | 0.191666 | Other Positive Correlation |
| GPR162 | 0.077313 | 0.192335 | Other Positive Correlation |
| ZNF189 | 0.077216 | 0.192891 | Other Positive Correlation |
| KBTBD8 | 0.07717 | 0.19316 | Other Positive Correlation |
| KIAA1614 | 0.077163 | 0.193198 | Other Positive Correlation |
| C19orf73 | 0.077163 | 0.193199 | Other Positive Correlation |

| | | | |
|---|---|---|---|
| ADGRB2 | 0.077134 | 0.19337 | Other Positive Correlation |
| ACSL6 | 0.077086 | 0.193649 | Other Positive Correlation |
| MOBP | 0.077057 | 0.193815 | Other Positive Correlation |
| SPX | 0.077017 | 0.194046 | Other Positive Correlation |
| CCSER1 | 0.076991 | 0.194196 | Other Positive Correlation |
| FAM124A | 0.076971 | 0.194315 | Other Positive Correlation |
| BAAT | 0.076864 | 0.194934 | Other Positive Correlation |
| GFPT2 | 0.076854 | 0.194993 | Other Positive Correlation |
| ALKAL1 | 0.076792 | 0.195355 | Other Positive Correlation |
| KCTD2 | 0.076786 | 0.195391 | Other Positive Correlation |
| TMEM200B | 0.076771 | 0.195479 | Other Positive Correlation |
| PPP1R16B | 0.076765 | 0.195516 | Other Positive Correlation |
| CCL3L1 | 0.076719 | 0.195781 | Other Positive Correlation |
| XPO7 | 0.076687 | 0.195971 | Other Positive Correlation |
| GSG1L | 0.076679 | 0.196015 | Other Positive Correlation |
| ITIH3 | 0.076529 | 0.196897 | Other Positive Correlation |
| JAKMIP1 | 0.076529 | 0.196897 | Other Positive Correlation |
| CD8B | 0.076529 | 0.196897 | Other Positive Correlation |
| SFRP2 | 0.076529 | 0.196897 | Other Positive Correlation |
| ADRA1A | 0.076529 | 0.196897 | Other Positive Correlation |
| ZBTB5 | 0.076517 | 0.196968 | Other Positive Correlation |
| SLC7A2 | 0.076431 | 0.197472 | Other Positive Correlation |
| ENSG00000286098 | 0.076426 | 0.197502 | Other Positive Correlation |
| MAPK11 | 0.076347 | 0.197968 | Other Positive Correlation |
| ENSG00000288658 | 0.076325 | 0.198095 | Other Positive Correlation |
| SIM2 | 0.076252 | 0.198526 | Other Positive Correlation |
| IMPG2 | 0.076215 | 0.198744 | Other Positive Correlation |
| RFC2 | 0.076124 | 0.199281 | Other Positive Correlation |
| GFY | 0.076027 | 0.199859 | Other Positive Correlation |
| ADAMTS8 | 0.076027 | 0.199859 | Other Positive Correlation |
| CNTN5 | 0.076027 | 0.199859 | Other Positive Correlation |
| SOX5 | 0.076008 | 0.19997 | Other Positive Correlation |
| CENPP | 0.075982 | 0.200123 | Other Positive Correlation |
| BLTP3A | 0.075936 | 0.200398 | Other Positive Correlation |
| DSN1 | 0.07591 | 0.200555 | Other Positive Correlation |
| PROSER1 | 0.075446 | 0.203328 | Other Positive Correlation |
| PARD3B | 0.075392 | 0.20365 | Other Positive Correlation |
| ATP8B2 | 0.075304 | 0.204178 | Other Positive Correlation |
| GABRR2 | 0.075133 | 0.20521 | Other Positive Correlation |
| PAX5 | 0.075065 | 0.205625 | Other Positive Correlation |
| CDKN1B | 0.07504 | 0.205772 | Other Positive Correlation |
| CCDC74B | 0.074986 | 0.206103 | Other Positive Correlation |
| PANX2 | 0.07486 | 0.206867 | Other Positive Correlation |
| ETNPPL | 0.074841 | 0.206981 | Other Positive Correlation |
| LRRC38 | 0.074841 | 0.206981 | Other Positive Correlation |
| CHAD | 0.074841 | 0.206981 | Other Positive Correlation |

| | | | |
|---|---|---|---|
| ASB18 | 0.074841 | 0.206981 | Other Positive Correlation |
| CCN3 | 0.074726 | 0.207685 | Other Positive Correlation |
| SCT | 0.074705 | 0.207812 | Other Positive Correlation |
| KSR2 | 0.074694 | 0.20788 | Other Positive Correlation |
| ANKS3 | 0.074662 | 0.208078 | Other Positive Correlation |
| ZDHHC18 | 0.074613 | 0.208376 | Other Positive Correlation |
| CDR2 | 0.074569 | 0.208643 | Other Positive Correlation |
| ATP6V0B | 0.074479 | 0.209197 | Other Positive Correlation |
| SDR39U1 | 0.074334 | 0.210083 | Other Positive Correlation |
| TREML2 | 0.074232 | 0.210712 | Other Positive Correlation |
| GRPR | 0.074227 | 0.210745 | Other Positive Correlation |
| FOXC2 | 0.074223 | 0.21077 | Other Positive Correlation |
| SCRG1 | 0.074163 | 0.211139 | Other Positive Correlation |
| GALR3 | 0.074146 | 0.211243 | Other Positive Correlation |
| PABPN1L | 0.074091 | 0.211581 | Other Positive Correlation |
| CAPSL | 0.074091 | 0.211581 | Other Positive Correlation |
| TM6SF1 | 0.074008 | 0.212099 | Other Positive Correlation |
| RAD51AP1 | 0.073962 | 0.21238 | Other Positive Correlation |
| MDGA1 | 0.073856 | 0.213041 | Other Positive Correlation |
| PAQR5 | 0.073764 | 0.213609 | Other Positive Correlation |
| PAIP2B | 0.073739 | 0.213766 | Other Positive Correlation |
| PLIN1 | 0.073399 | 0.215894 | Other Positive Correlation |
| FADS1 | 0.073339 | 0.216268 | Other Positive Correlation |
| TET3 | 0.073326 | 0.216353 | Other Positive Correlation |
| EIF2AK3 | 0.073313 | 0.216433 | Other Positive Correlation |
| ZNF559-ZNF177 | 0.073119 | 0.217655 | Other Positive Correlation |
| CCR9 | 0.073119 | 0.217655 | Other Positive Correlation |
| RAMP3 | 0.073119 | 0.217655 | Other Positive Correlation |
| PTH2R | 0.073119 | 0.217655 | Other Positive Correlation |
| CRYGD | 0.073119 | 0.217655 | Other Positive Correlation |
| WSCD2 | 0.073119 | 0.217655 | Other Positive Correlation |
| SLC5A8 | 0.073119 | 0.217655 | Other Positive Correlation |
| BCL2L12 | 0.07304 | 0.218151 | Other Positive Correlation |
| TMEM74 | 0.072879 | 0.219172 | Other Positive Correlation |
| ZBTB8B | 0.072772 | 0.219852 | Other Positive Correlation |
| SLC1A2 | 0.072704 | 0.220281 | Other Positive Correlation |
| HS3ST5 | 0.072379 | 0.222361 | Other Positive Correlation |
| CACNG7 | 0.072379 | 0.222361 | Other Positive Correlation |
| ADCYAP1R1 | 0.072379 | 0.222361 | Other Positive Correlation |
| GAP43 | 0.072316 | 0.222762 | Other Positive Correlation |
| PRSS56 | 0.072316 | 0.222762 | Other Positive Correlation |
| ACAN | 0.072316 | 0.222762 | Other Positive Correlation |
| LRFN2 | 0.072316 | 0.222762 | Other Positive Correlation |
| HTR3A | 0.072316 | 0.222762 | Other Positive Correlation |
| SLURP1 | 0.072316 | 0.222762 | Other Positive Correlation |
| FTCD | 0.072316 | 0.222762 | Other Positive Correlation |

| | | | |
|---|---|---|---|
| RORB | 0.072287 | 0.22295 | Other Positive Correlation |
| ITGA2B | 0.072198 | 0.223517 | Other Positive Correlation |
| SLC18A2 | 0.072154 | 0.223801 | Other Positive Correlation |
| RASGEF1B | 0.072146 | 0.223856 | Other Positive Correlation |
| NPAS1 | 0.072141 | 0.223888 | Other Positive Correlation |
| LRP1B | 0.072136 | 0.223921 | Other Positive Correlation |
| ZIC5 | 0.0721 | 0.22415 | Other Positive Correlation |
| LIN28A | 0.0721 | 0.22415 | Other Positive Correlation |
| SEL1L | 0.071888 | 0.22552 | Other Positive Correlation |
| PALM | 0.071762 | 0.226338 | Other Positive Correlation |
| COMMD4 | 0.071749 | 0.22642 | Other Positive Correlation |
| LAMC3 | 0.071716 | 0.226635 | Other Positive Correlation |
| KCNJ11 | 0.071713 | 0.226652 | Other Positive Correlation |
| HBZ | 0.071669 | 0.22694 | Other Positive Correlation |
| IFT27 | 0.071642 | 0.227114 | Other Positive Correlation |
| CLEC11A | 0.071638 | 0.227144 | Other Positive Correlation |
| ENSG00000187186 | 0.071634 | 0.22717 | Other Positive Correlation |
| GRB14 | 0.071588 | 0.22747 | Other Positive Correlation |
| CENPQ | 0.071574 | 0.227556 | Other Positive Correlation |
| CD200 | 0.071553 | 0.227693 | Other Positive Correlation |
| KATNAL2 | 0.071459 | 0.22831 | Other Positive Correlation |
| RMI1 | 0.071424 | 0.228538 | Other Positive Correlation |
| UPF2 | 0.071308 | 0.229298 | Other Positive Correlation |
| SLC25A35 | 0.071224 | 0.229846 | Other Positive Correlation |
| STARD9 | 0.071194 | 0.230042 | Other Positive Correlation |
| MMP11 | 0.071176 | 0.230159 | Other Positive Correlation |
| TIGD1 | 0.071049 | 0.230997 | Other Positive Correlation |
| SLC4A4 | 0.07103 | 0.231118 | Other Positive Correlation |
| FAM234B | 0.070921 | 0.231836 | Other Positive Correlation |
| RTEL1 | 0.070865 | 0.232207 | Other Positive Correlation |
| WDR19 | 0.070803 | 0.232618 | Other Positive Correlation |
| ENSG00000159239 | 0.070781 | 0.232763 | Other Positive Correlation |
| PCK1 | 0.070781 | 0.232763 | Other Positive Correlation |
| TERT | 0.070781 | 0.232763 | Other Positive Correlation |
| ENSG00000275778 | 0.070747 | 0.23299 | Other Positive Correlation |
| ACRBP | 0.070643 | 0.23368 | Other Positive Correlation |
| SPRY3 | 0.070632 | 0.233754 | Other Positive Correlation |
| TMDD1 | 0.070588 | 0.234046 | Other Positive Correlation |
| MDM1 | 0.070586 | 0.234057 | Other Positive Correlation |
| TM4SF5 | 0.070583 | 0.234076 | Other Positive Correlation |
| XKR7 | 0.070583 | 0.234076 | Other Positive Correlation |
| LMAN1L | 0.070583 | 0.234076 | Other Positive Correlation |
| CADPS | 0.070583 | 0.234076 | Other Positive Correlation |
| GOLGA8H | 0.070583 | 0.234076 | Other Positive Correlation |
| N4BP2L1 | 0.070408 | 0.23524 | Other Positive Correlation |
| SP9 | 0.070301 | 0.235958 | Other Positive Correlation |

| | | | |
|---|---|---|---|
| CDH7 | 0.070237 | 0.23638 | Other Positive Correlation |
| SLC6A1 | 0.070237 | 0.23638 | Other Positive Correlation |
| MORN5 | 0.070237 | 0.23638 | Other Positive Correlation |
| SYT17 | 0.07014 | 0.23703 | Other Positive Correlation |
| MICA | 0.070052 | 0.237622 | Other Positive Correlation |
| SRCIN1 | 0.070017 | 0.237859 | Other Positive Correlation |
| ANK2 | 0.069989 | 0.238045 | Other Positive Correlation |
| OOSP4A | 0.069933 | 0.238422 | Other Positive Correlation |
| ATG9A | 0.069861 | 0.238909 | Other Positive Correlation |
| PDE1C | 0.069842 | 0.239035 | Other Positive Correlation |
| RASSF2 | 0.069778 | 0.239468 | Other Positive Correlation |
| TTPA | 0.069763 | 0.23957 | Other Positive Correlation |
| PBX4 | 0.069746 | 0.239681 | Other Positive Correlation |
| CTNNA2 | 0.069726 | 0.239819 | Other Positive Correlation |
| MED8 | 0.069722 | 0.239846 | Other Positive Correlation |
| OR2AG1 | 0.069692 | 0.24005 | Other Positive Correlation |
| ZNF550 | 0.069676 | 0.240159 | Other Positive Correlation |
| SEPTIN1 | 0.069646 | 0.240358 | Other Positive Correlation |
| RGS7BP | 0.069591 | 0.240735 | Other Positive Correlation |
| XKR4 | 0.069585 | 0.240773 | Other Positive Correlation |
| ZDHHC11B | 0.069551 | 0.241002 | Other Positive Correlation |
| DACT1 | 0.069518 | 0.241229 | Other Positive Correlation |
| NEU1 | 0.069443 | 0.241735 | Other Positive Correlation |
| SLC16A7 | 0.069414 | 0.241933 | Other Positive Correlation |
| B3GAT1 | 0.069388 | 0.24211 | Other Positive Correlation |
| KCNK5 | 0.06936 | 0.242301 | Other Positive Correlation |
| ANKRD33B | 0.069331 | 0.242501 | Other Positive Correlation |
| FAM219B | 0.069295 | 0.242746 | Other Positive Correlation |
| PRRG3 | 0.069252 | 0.243039 | Other Positive Correlation |
| CDNF | 0.06925 | 0.243047 | Other Positive Correlation |
| RPS19BP1 | 0.06919 | 0.24346 | Other Positive Correlation |
| SPR | 0.069184 | 0.243502 | Other Positive Correlation |
| CARNS1 | 0.069172 | 0.243581 | Other Positive Correlation |
| C1QL3 | 0.06909 | 0.244142 | Other Positive Correlation |
| RSPH10B2 | 0.06909 | 0.244142 | Other Positive Correlation |
| ZNF670 | 0.069036 | 0.24451 | Other Positive Correlation |
| NKD2 | 0.068988 | 0.24484 | Other Positive Correlation |
| ADORA3 | 0.068964 | 0.245003 | Other Positive Correlation |
| ENPP6 | 0.068964 | 0.245003 | Other Positive Correlation |
| ENSG00000258691 | 0.068964 | 0.245003 | Other Positive Correlation |
| ZNF841 | 0.068939 | 0.245178 | Other Positive Correlation |
| RAB27A | 0.068906 | 0.245401 | Other Positive Correlation |
| MZB1 | 0.068905 | 0.24541 | Other Positive Correlation |
| ATOH8 | 0.068865 | 0.245689 | Other Positive Correlation |
| PCOLCE2 | 0.068807 | 0.246087 | Other Positive Correlation |
| SLC9A8 | 0.06879 | 0.246205 | Other Positive Correlation |

| | | | |
|---|---|---|---|
| PLA2G2A | 0.06854 | 0.24793 | Other Positive Correlation |
| RNF5 | 0.068512 | 0.248123 | Other Positive Correlation |
| MMP15 | 0.0685 | 0.248201 | Other Positive Correlation |
| PLCL2 | 0.068442 | 0.248607 | Other Positive Correlation |
| UGGT2 | 0.068437 | 0.248638 | Other Positive Correlation |
| CACNA1B | 0.068402 | 0.248881 | Other Positive Correlation |
| C21orf58 | 0.068378 | 0.249051 | Other Positive Correlation |
| CBLN1 | 0.068276 | 0.24976 | Other Positive Correlation |
| C8orf88 | 0.068168 | 0.250507 | Other Positive Correlation |
| EMILIN1 | 0.068153 | 0.250611 | Other Positive Correlation |
| ZFP69B | 0.068139 | 0.250714 | Other Positive Correlation |
| FOXRED2 | 0.067963 | 0.251941 | Other Positive Correlation |
| ZNF805 | 0.06795 | 0.252034 | Other Positive Correlation |
| SMIM10L2B | 0.067891 | 0.252442 | Other Positive Correlation |
| C2orf81 | 0.067741 | 0.253492 | Other Positive Correlation |
| GREB1L | 0.06774 | 0.253502 | Other Positive Correlation |
| SLC5A5 | 0.067731 | 0.253564 | Other Positive Correlation |
| NIBAN3 | 0.067533 | 0.254963 | Other Positive Correlation |
| TP53I13 | 0.067507 | 0.255142 | Other Positive Correlation |
| UGT3A2 | 0.067481 | 0.255324 | Other Positive Correlation |
| HOXC13 | 0.067364 | 0.25615 | Other Positive Correlation |
| TRIM43 | 0.067295 | 0.256642 | Other Positive Correlation |
| TRIM49C | 0.067295 | 0.256642 | Other Positive Correlation |
| DLL4 | 0.067234 | 0.257073 | Other Positive Correlation |
| LMX1B | 0.067211 | 0.257237 | Other Positive Correlation |
| RASSF3 | 0.067163 | 0.25758 | Other Positive Correlation |
| NEUROD2 | 0.067033 | 0.258504 | Other Positive Correlation |
| CNTN2 | 0.067003 | 0.258719 | Other Positive Correlation |
| CRHR2 | 0.066961 | 0.259013 | Other Positive Correlation |
| NCBP2L | 0.066961 | 0.259013 | Other Positive Correlation |
| GRIK3 | 0.066961 | 0.259013 | Other Positive Correlation |
| CNMD | 0.066961 | 0.259013 | Other Positive Correlation |
| MTUS2 | 0.066939 | 0.259171 | Other Positive Correlation |
| ZKSCAN8P1 | 0.06685 | 0.259809 | Other Positive Correlation |
| CRY1 | 0.066768 | 0.260397 | Other Positive Correlation |
| ACYP1 | 0.06668 | 0.261027 | Other Positive Correlation |
| CCDC184 | 0.066609 | 0.26153 | Other Positive Correlation |
| SHISA3 | 0.066427 | 0.262844 | Other Positive Correlation |
| SMC2 | 0.0664 | 0.263033 | Other Positive Correlation |
| TMEM198 | 0.066138 | 0.264925 | Other Positive Correlation |
| CMTM1 | 0.066081 | 0.265341 | Other Positive Correlation |
| GPAA1 | 0.066068 | 0.265438 | Other Positive Correlation |
| GABRG3 | 0.06606 | 0.265492 | Other Positive Correlation |
| HDAC6 | 0.066058 | 0.265509 | Other Positive Correlation |
| LEMD2 | 0.065888 | 0.266741 | Other Positive Correlation |
| SMIM10 | 0.065824 | 0.267206 | Other Positive Correlation |

| | | | |
|---|---|---|---|
| SLC46A2 | 0.065791 | 0.267453 | Other Positive Correlation |
| ABCB1 | 0.065791 | 0.267453 | Other Positive Correlation |
| GET1-SH3BGR | 0.065791 | 0.267453 | Other Positive Correlation |
| CCNA1 | 0.065786 | 0.267487 | Other Positive Correlation |
| SBSPON | 0.06571 | 0.268037 | Other Positive Correlation |
| ELF5 | 0.065687 | 0.26821 | Other Positive Correlation |
| RNASEK-C17orf49 | 0.065452 | 0.269925 | Other Positive Correlation |
| GRIK5 | 0.065365 | 0.270566 | Other Positive Correlation |
| RFXAP | 0.065279 | 0.271201 | Other Positive Correlation |
| WIPF3 | 0.065204 | 0.271753 | Other Positive Correlation |
| SLC27A2 | 0.065176 | 0.271953 | Other Positive Correlation |
| ROPN1B | 0.065072 | 0.272723 | Other Positive Correlation |
| NKX3-2 | 0.064963 | 0.273531 | Other Positive Correlation |
| GRID1 | 0.064882 | 0.274128 | Other Positive Correlation |
| OSER1 | 0.064809 | 0.274669 | Other Positive Correlation |
| MAGEH1 | 0.064788 | 0.274825 | Other Positive Correlation |
| ENPP2 | 0.064768 | 0.274979 | Other Positive Correlation |
| SCAPER | 0.064665 | 0.275742 | Other Positive Correlation |
| BAMBI | 0.06458 | 0.276373 | Other Positive Correlation |
| DRD5 | 0.064562 | 0.276512 | Other Positive Correlation |
| NLRP14 | 0.064522 | 0.276804 | Other Positive Correlation |
| CYP3A4 | 0.064478 | 0.277136 | Other Positive Correlation |
| HCK | 0.064384 | 0.277837 | Other Positive Correlation |
| AP3M2 | 0.064366 | 0.277973 | Other Positive Correlation |
| ACADL | 0.064324 | 0.278284 | Other Positive Correlation |
| DCAF12L2 | 0.064299 | 0.27847 | Other Positive Correlation |
| SAMD14 | 0.06427 | 0.278694 | Other Positive Correlation |
| INHA | 0.064221 | 0.27906 | Other Positive Correlation |
| LCNL1 | 0.064184 | 0.279335 | Other Positive Correlation |
| TCEAL5 | 0.064184 | 0.279335 | Other Positive Correlation |
| RPH3A | 0.064184 | 0.279335 | Other Positive Correlation |
| MXRA8 | 0.064129 | 0.279753 | Other Positive Correlation |
| TMCO5A | 0.064045 | 0.280381 | Other Positive Correlation |
| PHLPP2 | 0.064033 | 0.280473 | Other Positive Correlation |
| DLX2 | 0.064016 | 0.280602 | Other Positive Correlation |
| STK33 | 0.063799 | 0.282233 | Other Positive Correlation |
| ZFP82 | 0.063733 | 0.282732 | Other Positive Correlation |
| THAP5 | 0.063686 | 0.28309 | Other Positive Correlation |
| GABBR2 | 0.063583 | 0.283876 | Other Positive Correlation |
| NCR3LG1 | 0.063576 | 0.283927 | Other Positive Correlation |
| CFAP73 | 0.063464 | 0.284776 | Other Positive Correlation |
| GREB1 | 0.063397 | 0.285283 | Other Positive Correlation |
| MYEF2 | 0.063331 | 0.285785 | Other Positive Correlation |
| KIF5A | 0.063298 | 0.286044 | Other Positive Correlation |
| TSFM | 0.06322 | 0.286636 | Other Positive Correlation |
| PIP4P1 | 0.063156 | 0.287122 | Other Positive Correlation |

| | | | |
|---|---|---|---|
| RGS3 | 0.063041 | 0.288008 | Other Positive Correlation |
| C3orf80 | 0.062997 | 0.288344 | Other Positive Correlation |
| B3GALT1 | 0.062997 | 0.288344 | Other Positive Correlation |
| DPEP3 | 0.062932 | 0.288837 | Other Positive Correlation |
| BHLHA15 | 0.062914 | 0.28898 | Other Positive Correlation |
| ATG101 | 0.062899 | 0.289092 | Other Positive Correlation |
| EIF3C | 0.062851 | 0.289466 | Other Positive Correlation |
| OR3A3 | 0.062844 | 0.289515 | Other Positive Correlation |
| GAL3ST1 | 0.06275 | 0.290243 | Other Positive Correlation |
| ZNF674 | 0.06275 | 0.290244 | Other Positive Correlation |
| KIT | 0.062716 | 0.290499 | Other Positive Correlation |
| CFTR | 0.062705 | 0.290587 | Other Positive Correlation |
| GNG2 | 0.062695 | 0.290662 | Other Positive Correlation |
| UTS2R | 0.062578 | 0.291569 | Other Positive Correlation |
| CABLES2 | 0.062551 | 0.291773 | Other Positive Correlation |
| TMSB15B | 0.062347 | 0.293356 | Other Positive Correlation |
| ZNF729 | 0.062185 | 0.294613 | Other Positive Correlation |
| ZNF423 | 0.062159 | 0.294816 | Other Positive Correlation |
| GIGYF1 | 0.062157 | 0.294836 | Other Positive Correlation |
| ZNF396 | 0.062138 | 0.294983 | Other Positive Correlation |
| LYNX1 | 0.061915 | 0.296722 | Other Positive Correlation |
| ZNF84 | 0.061857 | 0.297177 | Other Positive Correlation |
| SKIC2 | 0.06185 | 0.297231 | Other Positive Correlation |
| CSMD2 | 0.061846 | 0.29726 | Other Positive Correlation |
| ADGRA2 | 0.061763 | 0.297912 | Other Positive Correlation |
| MUC12 | 0.06173 | 0.298169 | Other Positive Correlation |
| COL4A3 | 0.061671 | 0.298629 | Other Positive Correlation |
| PDGFRB | 0.06165 | 0.298797 | Other Positive Correlation |
| CRYGN | 0.061611 | 0.299101 | Other Positive Correlation |
| KCNJ9 | 0.061611 | 0.299101 | Other Positive Correlation |
| INSM1 | 0.061611 | 0.299101 | Other Positive Correlation |
| LHX8 | 0.061611 | 0.299101 | Other Positive Correlation |
| PTCHD3 | 0.061598 | 0.299204 | Other Positive Correlation |
| ENSG00000284512 | 0.061598 | 0.299204 | Other Positive Correlation |
| MEIOB | 0.061588 | 0.29928 | Other Positive Correlation |
| SDR42E2 | 0.061532 | 0.29972 | Other Positive Correlation |
| TRPC6 | 0.06142 | 0.30061 | Other Positive Correlation |
| COX6B2 | 0.06135 | 0.301158 | Other Positive Correlation |
| SLC2A13 | 0.06132 | 0.301393 | Other Positive Correlation |
| CALHM3 | 0.061307 | 0.3015 | Other Positive Correlation |
| DIS3L | 0.061157 | 0.302683 | Other Positive Correlation |
| COPZ2 | 0.061107 | 0.303078 | Other Positive Correlation |
| SLC4A9 | 0.061092 | 0.303196 | Other Positive Correlation |
| AXIN2 | 0.061079 | 0.303303 | Other Positive Correlation |
| NXPH3 | 0.061006 | 0.303884 | Other Positive Correlation |
| FUT4 | 0.060957 | 0.304267 | Other Positive Correlation |

| | | | |
|---|---|---|---|
| TMC5 | 0.060939 | 0.304414 | Other Positive Correlation |
| PRLR | 0.060855 | 0.305079 | Other Positive Correlation |
| EZH1 | 0.0608 | 0.305521 | Other Positive Correlation |
| DPYSL2 | 0.06069 | 0.3064 | Other Positive Correlation |
| KCNB1 | 0.06064 | 0.306795 | Other Positive Correlation |
| SLCO4A1 | 0.060608 | 0.307051 | Other Positive Correlation |
| EPHA6 | 0.060539 | 0.307606 | Other Positive Correlation |
| KRBOX4 | 0.060447 | 0.308342 | Other Positive Correlation |
| HNF1B | 0.060411 | 0.308634 | Other Positive Correlation |
| TCTE1 | 0.060406 | 0.308672 | Other Positive Correlation |
| C1orf112 | 0.060336 | 0.309231 | Other Positive Correlation |
| DTL | 0.06022 | 0.310168 | Other Positive Correlation |
| SLC15A3 | 0.060166 | 0.310599 | Other Positive Correlation |
| TEX45 | 0.060132 | 0.310876 | Other Positive Correlation |
| PRH2 | 0.060095 | 0.311171 | Other Positive Correlation |
| SOX1 | 0.059976 | 0.312136 | Other Positive Correlation |
| SOBP | 0.05994 | 0.312427 | Other Positive Correlation |
| ZNF530 | 0.059913 | 0.312644 | Other Positive Correlation |
| SH3GL3 | 0.059805 | 0.313518 | Other Positive Correlation |
| CNTF | 0.059763 | 0.313855 | Other Positive Correlation |
| IGSF9B | 0.05973 | 0.314125 | Other Positive Correlation |
| SIPA1L2 | 0.059673 | 0.31459 | Other Positive Correlation |
| SYTL2 | 0.059651 | 0.314771 | Other Positive Correlation |
| CCDC40 | 0.059553 | 0.315564 | Other Positive Correlation |
| TBXA2R | 0.059396 | 0.316842 | Other Positive Correlation |
| PLAAT1 | 0.059378 | 0.316992 | Other Positive Correlation |
| BRME1 | 0.059331 | 0.317377 | Other Positive Correlation |
| CEP162 | 0.059312 | 0.317532 | Other Positive Correlation |
| SAMD10 | 0.059269 | 0.317884 | Other Positive Correlation |
| C19orf81 | 0.059217 | 0.318304 | Other Positive Correlation |
| VSIG8 | 0.059113 | 0.31916 | Other Positive Correlation |
| ZCCHC12 | 0.059059 | 0.319601 | Other Positive Correlation |
| TERB1 | 0.059 | 0.320089 | Other Positive Correlation |
| ANKRD62 | 0.058998 | 0.3201 | Other Positive Correlation |
| REC8 | 0.058989 | 0.320175 | Other Positive Correlation |
| SLC6A3 | 0.058968 | 0.320353 | Other Positive Correlation |
| NEUROG3 | 0.058968 | 0.320353 | Other Positive Correlation |
| CCM2L | 0.058968 | 0.320353 | Other Positive Correlation |
| DEFB126 | 0.058968 | 0.320353 | Other Positive Correlation |
| HNF4G | 0.058968 | 0.320353 | Other Positive Correlation |
| TSPAN33 | 0.058957 | 0.320441 | Other Positive Correlation |
| ZFPM2 | 0.058922 | 0.320733 | Other Positive Correlation |
| RHOQ | 0.058838 | 0.321424 | Other Positive Correlation |
| PRKCG | 0.058823 | 0.321544 | Other Positive Correlation |
| ZMYND12 | 0.058813 | 0.32163 | Other Positive Correlation |
| CGAS | 0.058726 | 0.322343 | Other Positive Correlation |

| | | | |
|---|---|---|---|
| PDE6B | 0.058713 | 0.322452 | Other Positive Correlation |
| MFSD4A | 0.058659 | 0.322898 | Other Positive Correlation |
| ADCY5 | 0.058645 | 0.323017 | Other Positive Correlation |
| RPRM | 0.058627 | 0.323159 | Other Positive Correlation |
| ZNF528 | 0.058513 | 0.324108 | Other Positive Correlation |
| ST6GALNAC3 | 0.058503 | 0.324192 | Other Positive Correlation |
| DLGAP2 | 0.058439 | 0.324722 | Other Positive Correlation |
| ZNF66 | 0.058399 | 0.325054 | Other Positive Correlation |
| FOXP3 | 0.058105 | 0.327497 | Other Positive Correlation |
| FOXJ1 | 0.058042 | 0.32802 | Other Positive Correlation |
| MRPL49 | 0.057977 | 0.328569 | Other Positive Correlation |
| RENBP | 0.057914 | 0.329095 | Other Positive Correlation |
| GIMAP5 | 0.057836 | 0.329745 | Other Positive Correlation |
| FIGNL2 | 0.0578 | 0.330047 | Other Positive Correlation |
| STRA8 | 0.057765 | 0.330339 | Other Positive Correlation |
| GPM6A | 0.057743 | 0.330524 | Other Positive Correlation |
| POU6F1 | 0.057738 | 0.33057 | Other Positive Correlation |
| TCP11 | 0.057699 | 0.33089 | Other Positive Correlation |
| VENTX | 0.057606 | 0.331676 | Other Positive Correlation |
| ZDHHC2 | 0.057526 | 0.332346 | Other Positive Correlation |
| KHDC3L | 0.057421 | 0.33323 | Other Positive Correlation |
| HNF4A | 0.057409 | 0.333333 | Other Positive Correlation |
| OR2L2 | 0.057409 | 0.333333 | Other Positive Correlation |
| ITGA9 | 0.057409 | 0.333333 | Other Positive Correlation |
| GATA1 | 0.057409 | 0.333333 | Other Positive Correlation |
| CACNG1 | 0.057409 | 0.333333 | Other Positive Correlation |
| TMEM74B | 0.057385 | 0.333535 | Other Positive Correlation |
| NUP62CL | 0.057106 | 0.335898 | Other Positive Correlation |
| C5 | 0.057052 | 0.33635 | Other Positive Correlation |
| DMRT3 | 0.05697 | 0.337051 | Other Positive Correlation |
| ENSG00000272104 | 0.05694 | 0.337306 | Other Positive Correlation |
| OR2C3 | 0.05692 | 0.337476 | Other Positive Correlation |
| GLT8D2 | 0.056705 | 0.339304 | Other Positive Correlation |
| AKT3 | 0.056704 | 0.339316 | Other Positive Correlation |
| SLC39A14 | 0.056674 | 0.339569 | Other Positive Correlation |
| RCCD1 | 0.0566 | 0.340203 | Other Positive Correlation |
| NKX2-2 | 0.05649 | 0.341141 | Other Positive Correlation |
| CACNA1F | 0.056436 | 0.341608 | Other Positive Correlation |
| ANGPT2 | 0.056313 | 0.342664 | Other Positive Correlation |
| LAIR1 | 0.056305 | 0.34273 | Other Positive Correlation |
| VASH2 | 0.056253 | 0.343178 | Other Positive Correlation |
| PIFO | 0.056153 | 0.344038 | Other Positive Correlation |
| MEI1 | 0.056113 | 0.344382 | Other Positive Correlation |
| MAGIX | 0.055751 | 0.347502 | Other Positive Correlation |
| HOXC6 | 0.05575 | 0.347511 | Other Positive Correlation |
| RAD9A | 0.055742 | 0.347587 | Other Positive Correlation |

| | | | |
|---|---|---|---|
| C17orf114 | 0.055707 | 0.347886 | Other Positive Correlation |
| ZNF273 | 0.055632 | 0.348536 | Other Positive Correlation |
| ODAD1 | 0.05559 | 0.348906 | Other Positive Correlation |
| FAM184B | 0.055586 | 0.348939 | Other Positive Correlation |
| KLRF1 | 0.055572 | 0.349057 | Other Positive Correlation |
| FGF3 | 0.055302 | 0.351412 | Other Positive Correlation |
| ASB4 | 0.055177 | 0.352497 | Other Positive Correlation |
| TWIST1 | 0.055173 | 0.352537 | Other Positive Correlation |
| ARHGAP31 | 0.055141 | 0.352814 | Other Positive Correlation |
| GARIN1B | 0.055122 | 0.352981 | Other Positive Correlation |
| USP11 | 0.055103 | 0.353149 | Other Positive Correlation |
| GAN | 0.055067 | 0.353465 | Other Positive Correlation |
| ZNF275 | 0.055034 | 0.353755 | Other Positive Correlation |
| CAVIN4 | 0.054989 | 0.354145 | Other Positive Correlation |
| KIF21B | 0.054972 | 0.354297 | Other Positive Correlation |
| ELOVL3 | 0.05491 | 0.354835 | Other Positive Correlation |
| ARL17A | 0.054725 | 0.356464 | Other Positive Correlation |
| RNF148 | 0.054716 | 0.356548 | Other Positive Correlation |
| BAIAP3 | 0.054675 | 0.356906 | Other Positive Correlation |
| CACNA1C | 0.054637 | 0.357237 | Other Positive Correlation |
| GUCA1B | 0.054619 | 0.357398 | Other Positive Correlation |
| ABCG2 | 0.054559 | 0.357932 | Other Positive Correlation |
| LBHD2 | 0.054536 | 0.35813 | Other Positive Correlation |
| KLHDC3 | 0.054376 | 0.359546 | Other Positive Correlation |
| ASGR1 | 0.054337 | 0.359888 | Other Positive Correlation |
| SCN9A | 0.054315 | 0.360089 | Other Positive Correlation |
| GDF15 | 0.054305 | 0.360175 | Other Positive Correlation |
| CGNL1 | 0.054293 | 0.360279 | Other Positive Correlation |
| MCM3 | 0.054251 | 0.360653 | Other Positive Correlation |
| HS6ST3 | 0.054242 | 0.360731 | Other Positive Correlation |
| PDCD1 | 0.054229 | 0.360847 | Other Positive Correlation |
| HAUS6 | 0.054059 | 0.362353 | Other Positive Correlation |
| FAM174B | 0.054055 | 0.362395 | Other Positive Correlation |
| PYDC1 | 0.054043 | 0.362498 | Other Positive Correlation |
| PCSK4 | 0.053969 | 0.363154 | Other Positive Correlation |
| ADRA2B | 0.053921 | 0.363589 | Other Positive Correlation |
| RFX3 | 0.053827 | 0.364419 | Other Positive Correlation |
| KCNQ3 | 0.053811 | 0.364566 | Other Positive Correlation |
| TYW1B | 0.053801 | 0.364654 | Other Positive Correlation |
| DNAL1 | 0.053796 | 0.364704 | Other Positive Correlation |
| BIN1 | 0.053733 | 0.365267 | Other Positive Correlation |
| MFSD6L | 0.053587 | 0.366567 | Other Positive Correlation |
| RNF166 | 0.053533 | 0.367052 | Other Positive Correlation |
| STAG3 | 0.053511 | 0.367247 | Other Positive Correlation |
| N4BP2 | 0.053428 | 0.367997 | Other Positive Correlation |
| SVIP | 0.053383 | 0.368398 | Other Positive Correlation |

| | | | |
|---|---|---|---|
| LYG1 | 0.053368 | 0.368535 | Other Positive Correlation |
| PVALB | 0.053314 | 0.369017 | Other Positive Correlation |
| ENSG00000285713 | 0.05327 | 0.369413 | Other Positive Correlation |
| PTGER3 | 0.05322 | 0.369861 | Other Positive Correlation |
| ODF3L2 | 0.053102 | 0.370928 | Other Positive Correlation |
| KCTD7 | 0.052948 | 0.372322 | Other Positive Correlation |
| LIFR | 0.052926 | 0.372523 | Other Positive Correlation |
| AIFM3 | 0.052863 | 0.373093 | Other Positive Correlation |
| HMOX1 | 0.052843 | 0.373272 | Other Positive Correlation |
| NT5M | 0.05279 | 0.373755 | Other Positive Correlation |
| TNFRSF9 | 0.052768 | 0.373951 | Other Positive Correlation |
| RAB3IP | 0.052751 | 0.374108 | Other Positive Correlation |
| MT3 | 0.052723 | 0.374361 | Other Positive Correlation |
| RAB33A | 0.052702 | 0.374551 | Other Positive Correlation |
| ZNF684 | 0.05266 | 0.37493 | Other Positive Correlation |
| DOCK8 | 0.052654 | 0.374989 | Other Positive Correlation |
| BARD1 | 0.052622 | 0.375277 | Other Positive Correlation |
| HLA-DPB1 | 0.052593 | 0.375535 | Other Positive Correlation |
| ELOVL7 | 0.05256 | 0.375842 | Other Positive Correlation |
| CFAP95 | 0.052507 | 0.376325 | Other Positive Correlation |
| TSHR | 0.052466 | 0.376696 | Other Positive Correlation |
| PYGO1 | 0.052418 | 0.37713 | Other Positive Correlation |
| SLC25A14 | 0.052402 | 0.377278 | Other Positive Correlation |
| BEND7 | 0.052382 | 0.377461 | Other Positive Correlation |
| GPD1 | 0.052381 | 0.377469 | Other Positive Correlation |
| FERMT2 | 0.052352 | 0.377735 | Other Positive Correlation |
| SV2B | 0.052316 | 0.37806 | Other Positive Correlation |
| C9orf24 | 0.052304 | 0.378175 | Other Positive Correlation |
| NAT9 | 0.052202 | 0.3791 | Other Positive Correlation |
| PGAM4 | 0.0522 | 0.379121 | Other Positive Correlation |
| SNTA1 | 0.052191 | 0.379202 | Other Positive Correlation |
| CEP295NL | 0.052169 | 0.379403 | Other Positive Correlation |
| ZNF470 | 0.05214 | 0.37967 | Other Positive Correlation |
| SSTR5 | 0.052051 | 0.380487 | Other Positive Correlation |
| PCDHGA5 | 0.051996 | 0.380992 | Other Positive Correlation |
| RNF208 | 0.051984 | 0.381101 | Other Positive Correlation |
| AQP7 | 0.051967 | 0.381253 | Other Positive Correlation |
| ZNF616 | 0.051913 | 0.381751 | Other Positive Correlation |
| GPR137C | 0.051717 | 0.383556 | Other Positive Correlation |
| TMEM106C | 0.051588 | 0.384744 | Other Positive Correlation |
| PABIR3 | 0.051585 | 0.38477 | Other Positive Correlation |
| KCNV1 | 0.051313 | 0.387284 | Other Positive Correlation |
| SLC44A4 | 0.051308 | 0.387336 | Other Positive Correlation |
| CACNB4 | 0.051294 | 0.387457 | Other Positive Correlation |
| ANKRD34B | 0.051282 | 0.387571 | Other Positive Correlation |
| ANGPTL2 | 0.051221 | 0.388136 | Other Positive Correlation |

| | | | |
|---|---|---|---|
| ETV1 | 0.05121 | 0.388236 | Other Positive Correlation |
| MFAP3L | 0.051035 | 0.38987 | Other Positive Correlation |
| FOXF1 | 0.050944 | 0.390714 | Other Positive Correlation |
| ABHD12 | 0.050808 | 0.391982 | Other Positive Correlation |
| MSL2 | 0.050758 | 0.392444 | Other Positive Correlation |
| ZNF519 | 0.050755 | 0.392475 | Other Positive Correlation |
| ZNF300 | 0.050731 | 0.392695 | Other Positive Correlation |
| C4orf19 | 0.050653 | 0.393428 | Other Positive Correlation |
| TNIK | 0.050638 | 0.393565 | Other Positive Correlation |
| IL19 | 0.05054 | 0.39449 | Other Positive Correlation |
| CCDC163 | 0.050445 | 0.395378 | Other Positive Correlation |
| FRS3 | 0.050337 | 0.396392 | Other Positive Correlation |
| TCF7 | 0.050335 | 0.396413 | Other Positive Correlation |
| ACSS3 | 0.050237 | 0.397326 | Other Positive Correlation |
| TNFRSF8 | 0.050228 | 0.397418 | Other Positive Correlation |
| SLC6A16 | 0.050163 | 0.398022 | Other Positive Correlation |
| ABCG4 | 0.050161 | 0.398049 | Other Positive Correlation |
| PYGM | 0.050145 | 0.398193 | Other Positive Correlation |
| SNED1 | 0.050142 | 0.398221 | Other Positive Correlation |
| TRPM6 | 0.050139 | 0.398249 | Other Positive Correlation |
| ZNF382 | 0.050107 | 0.398558 | Other Positive Correlation |
| SSBP2 | 0.050101 | 0.39861 | Other Positive Correlation |
| GLS2 | 0.05008 | 0.398812 | Other Positive Correlation |
| ZNF691 | 0.050069 | 0.398909 | Other Positive Correlation |
| WDPCP | 0.05005 | 0.399096 | Other Positive Correlation |
| LYPD6 | 0.050046 | 0.399131 | Other Positive Correlation |
| TTC9B | 0.050046 | 0.399134 | Other Positive Correlation |
| GPC5 | 0.049972 | 0.399827 | Other Positive Correlation |
| FBXL16 | 0.049959 | 0.399948 | Other Positive Correlation |
| ATP8A2 | 0.04991 | 0.400417 | Other Positive Correlation |
| DNAJC22 | 0.049878 | 0.400714 | Other Positive Correlation |
| ZNF724 | 0.049832 | 0.401151 | Other Positive Correlation |
| DDTL | 0.049777 | 0.401675 | Other Positive Correlation |
| GBX2 | 0.049737 | 0.402047 | Other Positive Correlation |
| TBKBP1 | 0.049695 | 0.40245 | Other Positive Correlation |
| C1orf87 | 0.049528 | 0.404031 | Other Positive Correlation |
| ZNF177 | 0.049524 | 0.404075 | Other Positive Correlation |
| HYDIN | 0.049434 | 0.404926 | Other Positive Correlation |
| RTL9 | 0.049429 | 0.404973 | Other Positive Correlation |
| NAT16 | 0.049372 | 0.40552 | Other Positive Correlation |
| HOXB3 | 0.049318 | 0.406031 | Other Positive Correlation |
| MERTK | 0.0493 | 0.406199 | Other Positive Correlation |
| DEPDC4 | 0.049257 | 0.406614 | Other Positive Correlation |
| VPS26C | 0.049237 | 0.406805 | Other Positive Correlation |
| DZIP1L | 0.049226 | 0.406908 | Other Positive Correlation |
| HAPLN4 | 0.04919 | 0.407256 | Other Positive Correlation |

| | | | |
|---|---|---|---|
| TMEM208 | 0.049154 | 0.407594 | Other Positive Correlation |
| ZNF41 | 0.049131 | 0.407818 | Other Positive Correlation |
| TOX | 0.049074 | 0.408357 | Other Positive Correlation |
| PTPRU | 0.049073 | 0.408374 | Other Positive Correlation |
| DND1 | 0.048992 | 0.409147 | Other Positive Correlation |
| DPY19L3 | 0.04893 | 0.409743 | Other Positive Correlation |
| ATP6V1FNB | 0.048901 | 0.410021 | Other Positive Correlation |
| MEX3B | 0.048855 | 0.410458 | Other Positive Correlation |
| CHST13 | 0.048854 | 0.410471 | Other Positive Correlation |
| GPRASP1 | 0.048773 | 0.411247 | Other Positive Correlation |
| ARID3A | 0.048768 | 0.411298 | Other Positive Correlation |
| ZFP3 | 0.048713 | 0.411821 | Other Positive Correlation |
| ENSG00000267385 | 0.048617 | 0.412746 | Other Positive Correlation |
| RGS6 | 0.048402 | 0.414821 | Other Positive Correlation |
| RALYL | 0.048342 | 0.415396 | Other Positive Correlation |
| TMEM268 | 0.048339 | 0.415422 | Other Positive Correlation |
| LIN52 | 0.048316 | 0.415646 | Other Positive Correlation |
| KCNH1 | 0.048273 | 0.41606 | Other Positive Correlation |
| NALF2 | 0.048199 | 0.41678 | Other Positive Correlation |
| HMGN5 | 0.048189 | 0.416872 | Other Positive Correlation |
| RMC1 | 0.048172 | 0.417042 | Other Positive Correlation |
| TMPRSS7 | 0.048172 | 0.417046 | Other Positive Correlation |
| LMO1 | 0.048157 | 0.41719 | Other Positive Correlation |
| LRCH2 | 0.048131 | 0.417434 | Other Positive Correlation |
| ZNF853 | 0.048118 | 0.417559 | Other Positive Correlation |
| KCNT2 | 0.048095 | 0.417783 | Other Positive Correlation |
| MPC1 | 0.047882 | 0.419857 | Other Positive Correlation |
| PTER | 0.047805 | 0.420601 | Other Positive Correlation |
| ANKRD10 | 0.047785 | 0.420795 | Other Positive Correlation |
| CSPG5 | 0.047708 | 0.421545 | Other Positive Correlation |
| MRGPRF | 0.047644 | 0.422174 | Other Positive Correlation |
| SLC9A6 | 0.047594 | 0.422655 | Other Positive Correlation |
| RPL17-C18orf32 | 0.04736 | 0.424948 | Other Positive Correlation |
| ZNF285 | 0.047342 | 0.425126 | Other Positive Correlation |
| RNF41 | 0.047338 | 0.425164 | Other Positive Correlation |
| CTTNBP2 | 0.047335 | 0.425193 | Other Positive Correlation |
| ACSS1 | 0.047326 | 0.42528 | Other Positive Correlation |
| LBX2 | 0.047201 | 0.426508 | Other Positive Correlation |
| MPP1 | 0.047177 | 0.426744 | Other Positive Correlation |
| FZD2 | 0.047155 | 0.426953 | Other Positive Correlation |
| SMKR1 | 0.047101 | 0.427488 | Other Positive Correlation |
| FAM167A | 0.046986 | 0.428618 | Other Positive Correlation |
| CDHR3 | 0.046928 | 0.42919 | Other Positive Correlation |
| NLRP9 | 0.046821 | 0.430242 | Other Positive Correlation |
| GALNT16 | 0.046804 | 0.430406 | Other Positive Correlation |
| PRRT4 | 0.046774 | 0.430702 | Other Positive Correlation |

| | | | |
|---|---|---|---|
| BEST1 | 0.046737 | 0.431069 | Other Positive Correlation |
| ANKRD9 | 0.046636 | 0.432064 | Other Positive Correlation |
| MIA2 | 0.046626 | 0.432164 | Other Positive Correlation |
| KCNQ2 | 0.0466 | 0.432424 | Other Positive Correlation |
| POU3F3 | 0.046562 | 0.432796 | Other Positive Correlation |
| CLDN7 | 0.046431 | 0.434092 | Other Positive Correlation |
| ENDOU | 0.046431 | 0.434095 | Other Positive Correlation |
| SRGAP2C | 0.046429 | 0.434116 | Other Positive Correlation |
| HSD17B6 | 0.046427 | 0.434136 | Other Positive Correlation |
| TXLNA | 0.046391 | 0.434488 | Other Positive Correlation |
| ENSG00000285000 | 0.046377 | 0.434634 | Other Positive Correlation |
| PROZ | 0.046377 | 0.434634 | Other Positive Correlation |
| GRPEL2 | 0.046314 | 0.435258 | Other Positive Correlation |
| EPOP | 0.04629 | 0.43549 | Other Positive Correlation |
| PNMA8B | 0.046272 | 0.435675 | Other Positive Correlation |
| TMEM167B | 0.046199 | 0.436396 | Other Positive Correlation |
| NUSAP1 | 0.046156 | 0.436825 | Other Positive Correlation |
| SLC26A11 | 0.04615 | 0.436887 | Other Positive Correlation |
| UCN | 0.046071 | 0.437669 | Other Positive Correlation |
| TEX38 | 0.046028 | 0.438099 | Other Positive Correlation |
| ACER1 | 0.046018 | 0.438199 | Other Positive Correlation |
| PNMA6A | 0.045968 | 0.438703 | Other Positive Correlation |
| RGN | 0.045948 | 0.438893 | Other Positive Correlation |
| SMG5 | 0.045926 | 0.439118 | Other Positive Correlation |
| PIPOX | 0.045864 | 0.439734 | Other Positive Correlation |
| MT1H | 0.045826 | 0.440118 | Other Positive Correlation |
| CPXM1 | 0.045806 | 0.440317 | Other Positive Correlation |
| ARMC7 | 0.045584 | 0.442532 | Other Positive Correlation |
| NOVA2 | 0.045571 | 0.442661 | Other Positive Correlation |
| SAP30 | 0.045567 | 0.442704 | Other Positive Correlation |
| RASA4B | 0.045543 | 0.442949 | Other Positive Correlation |
| GMPPA | 0.045527 | 0.443103 | Other Positive Correlation |
| SYNJ1 | 0.045524 | 0.443136 | Other Positive Correlation |
| CHCT1 | 0.045492 | 0.443462 | Other Positive Correlation |
| PCOLCE | 0.045487 | 0.443511 | Other Positive Correlation |
| FA2H | 0.045483 | 0.443551 | Other Positive Correlation |
| ATP2C2 | 0.045372 | 0.444666 | Other Positive Correlation |
| AQP11 | 0.045277 | 0.445615 | Other Positive Correlation |
| CHRNB4 | 0.045229 | 0.446096 | Other Positive Correlation |
| GOLGA6L9 | 0.045179 | 0.446608 | Other Positive Correlation |
| GPR19 | 0.045121 | 0.447186 | Other Positive Correlation |
| PRTFDC1 | 0.045093 | 0.447474 | Other Positive Correlation |
| CHRNA4 | 0.045022 | 0.448188 | Other Positive Correlation |
| MANEAL | 0.045002 | 0.448389 | Other Positive Correlation |
| SLC6A2 | 0.045001 | 0.448398 | Other Positive Correlation |
| AMIGO1 | 0.04497 | 0.448715 | Other Positive Correlation |

| | | | |
|---|---|---|---|
| HOXA7 | 0.044939 | 0.449029 | Other Positive Correlation |
| SLCO4C1 | 0.044913 | 0.449292 | Other Positive Correlation |
| CYP46A1 | 0.044864 | 0.449781 | Other Positive Correlation |
| C7orf31 | 0.044843 | 0.449999 | Other Positive Correlation |
| RET | 0.044743 | 0.451003 | Other Positive Correlation |
| DIPK1B | 0.044589 | 0.452572 | Other Positive Correlation |
| ZNF14 | 0.044561 | 0.452848 | Other Positive Correlation |
| EIF4E3 | 0.044462 | 0.453862 | Other Positive Correlation |
| TEDC1 | 0.04439 | 0.454587 | Other Positive Correlation |
| DEDD2 | 0.04431 | 0.455404 | Other Positive Correlation |
| PHYHIPL | 0.044296 | 0.455545 | Other Positive Correlation |
| ZNF3 | 0.044288 | 0.455626 | Other Positive Correlation |
| FANCE | 0.044252 | 0.455998 | Other Positive Correlation |
| FABP3 | 0.044226 | 0.456261 | Other Positive Correlation |
| MXD3 | 0.044223 | 0.456289 | Other Positive Correlation |
| TMEM132C | 0.044222 | 0.456303 | Other Positive Correlation |
| RASD1 | 0.044192 | 0.456606 | Other Positive Correlation |
| PPM1N | 0.044096 | 0.457588 | Other Positive Correlation |
| GLIS2 | 0.04409 | 0.45765 | Other Positive Correlation |
| TENT5C | 0.044081 | 0.457744 | Other Positive Correlation |
| DNHD1 | 0.044037 | 0.45819 | Other Positive Correlation |
| CMPK2 | 0.044025 | 0.458311 | Other Positive Correlation |
| STK32B | 0.043962 | 0.458958 | Other Positive Correlation |
| NSUN7 | 0.043794 | 0.460671 | Other Positive Correlation |
| RHBDD1 | 0.043782 | 0.460801 | Other Positive Correlation |
| MPO | 0.04375 | 0.461132 | Other Positive Correlation |
| GPT | 0.043723 | 0.461405 | Other Positive Correlation |
| TMEM47 | 0.043679 | 0.461851 | Other Positive Correlation |
| AGTPBP1 | 0.043672 | 0.461932 | Other Positive Correlation |
| PRKN | 0.04364 | 0.462254 | Other Positive Correlation |
| WNK2 | 0.043619 | 0.46247 | Other Positive Correlation |
| U2AF1L4 | 0.043522 | 0.463472 | Other Positive Correlation |
| CADPS2 | 0.043462 | 0.464086 | Other Positive Correlation |
| BRICD5 | 0.04344 | 0.464312 | Other Positive Correlation |
| SLC25A27 | 0.043394 | 0.464789 | Other Positive Correlation |
| CCDC92B | 0.043256 | 0.466203 | Other Positive Correlation |
| ADAM22 | 0.043196 | 0.466823 | Other Positive Correlation |
| EPHX4 | 0.043041 | 0.468434 | Other Positive Correlation |
| TAF3 | 0.042984 | 0.469021 | Other Positive Correlation |
| PCP2 | 0.0429 | 0.469885 | Other Positive Correlation |
| RDH16 | 0.042895 | 0.469939 | Other Positive Correlation |
| SSTR2 | 0.042824 | 0.470676 | Other Positive Correlation |
| VGF | 0.042801 | 0.470918 | Other Positive Correlation |
| MAPK8IP1 | 0.042787 | 0.471059 | Other Positive Correlation |
| INCA1 | 0.042738 | 0.47157 | Other Positive Correlation |
| NDST3 | 0.042641 | 0.47258 | Other Positive Correlation |

| | | | |
|---|---|---|---|
| PTGES3L | 0.04259 | 0.473103 | Other Positive Correlation |
| KCNJ3 | 0.042565 | 0.473368 | Other Positive Correlation |
| CCDC82 | 0.042543 | 0.473595 | Other Positive Correlation |
| RGS7 | 0.042504 | 0.474 | Other Positive Correlation |
| WDR72 | 0.042501 | 0.474034 | Other Positive Correlation |
| FOXD4L3 | 0.042495 | 0.474096 | Other Positive Correlation |
| LRRC20 | 0.042262 | 0.476525 | Other Positive Correlation |
| IRX3 | 0.042219 | 0.476973 | Other Positive Correlation |
| CRYGC | 0.042161 | 0.477575 | Other Positive Correlation |
| KLHDC1 | 0.042134 | 0.477866 | Other Positive Correlation |
| LRRC51 | 0.042131 | 0.477892 | Other Positive Correlation |
| YJEFN3 | 0.041942 | 0.479869 | Other Positive Correlation |
| PDZD7 | 0.041894 | 0.480376 | Other Positive Correlation |
| ZNF765 | 0.041888 | 0.480442 | Other Positive Correlation |
| GNA13 | 0.041832 | 0.481028 | Other Positive Correlation |
| SMIM43 | 0.041813 | 0.481221 | Other Positive Correlation |
| ASXL3 | 0.041809 | 0.481269 | Other Positive Correlation |
| RELN | 0.041806 | 0.481301 | Other Positive Correlation |
| CSAD | 0.041764 | 0.481739 | Other Positive Correlation |
| LIMS2 | 0.041743 | 0.481958 | Other Positive Correlation |
| FGFR4 | 0.04174 | 0.481997 | Other Positive Correlation |
| LMCD1 | 0.041692 | 0.4825 | Other Positive Correlation |
| PLA2G3 | 0.041661 | 0.48282 | Other Positive Correlation |
| DNASE1L2 | 0.041606 | 0.483396 | Other Positive Correlation |
| DOC2A | 0.041586 | 0.483606 | Other Positive Correlation |
| NOX4 | 0.041578 | 0.483691 | Other Positive Correlation |
| PEX12 | 0.041576 | 0.483718 | Other Positive Correlation |
| LZTS1 | 0.041543 | 0.484061 | Other Positive Correlation |
| KLHL24 | 0.04149 | 0.484624 | Other Positive Correlation |
| BHLHA9 | 0.041457 | 0.484973 | Other Positive Correlation |
| EPM2A | 0.041392 | 0.485659 | Other Positive Correlation |
| UBB | 0.041346 | 0.486136 | Other Positive Correlation |
| ZKSCAN8 | 0.041189 | 0.487802 | Other Positive Correlation |
| BBS4 | 0.040868 | 0.491195 | Other Positive Correlation |
| ENSG00000288637 | 0.040858 | 0.491301 | Other Positive Correlation |
| C16orf86 | 0.040815 | 0.491764 | Other Positive Correlation |
| ENSG00000289720 | 0.040661 | 0.493395 | Other Positive Correlation |
| RBM4B | 0.04055 | 0.494586 | Other Positive Correlation |
| SYNDIG1L | 0.040466 | 0.495476 | Other Positive Correlation |
| MFSD9 | 0.040419 | 0.495976 | Other Positive Correlation |
| TMEM233 | 0.040307 | 0.497171 | Other Positive Correlation |
| GNPTG | 0.040169 | 0.49865 | Other Positive Correlation |
| GPR37L1 | 0.040162 | 0.498723 | Other Positive Correlation |
| MARCHF9 | 0.040153 | 0.498819 | Other Positive Correlation |
| CLCN1 | 0.040072 | 0.499692 | Other Positive Correlation |
| NME1-NME2 | 0.040064 | 0.49977 | Other Positive Correlation |

| | | | |
|---|---|---|---|
| ZNF697 | 0.039987 | 0.500599 | Other Positive Correlation |
| WNT9A | 0.039948 | 0.50102 | Other Positive Correlation |
| ARL17B | 0.039828 | 0.502304 | Other Positive Correlation |
| SEC61A2 | 0.039687 | 0.503815 | Other Positive Correlation |
| ZMYND15 | 0.039632 | 0.504412 | Other Positive Correlation |
| TMEM39A | 0.039504 | 0.505794 | Other Positive Correlation |
| PLA2G2C | 0.039459 | 0.506272 | Other Positive Correlation |
| AFG1L | 0.03922 | 0.508857 | Other Positive Correlation |
| ELMOD3 | 0.039157 | 0.509532 | Other Positive Correlation |
| AFF3 | 0.039155 | 0.509558 | Other Positive Correlation |
| LRP5 | 0.039121 | 0.509927 | Other Positive Correlation |
| DLGAP1 | 0.039046 | 0.510744 | Other Positive Correlation |
| KCNS1 | 0.038983 | 0.511427 | Other Positive Correlation |
| SKIDA1 | 0.038853 | 0.512838 | Other Positive Correlation |
| COX6A1 | 0.038847 | 0.512902 | Other Positive Correlation |
| GRAMD1A | 0.038836 | 0.513021 | Other Positive Correlation |
| TTC21A | 0.038754 | 0.513905 | Other Positive Correlation |
| SHMT1 | 0.038743 | 0.514026 | Other Positive Correlation |
| PDX1 | 0.038692 | 0.514582 | Other Positive Correlation |
| ZNF773 | 0.038658 | 0.514954 | Other Positive Correlation |
| BCKDHA | 0.038626 | 0.515301 | Other Positive Correlation |
| KIF26A | 0.03846 | 0.517104 | Other Positive Correlation |
| P4HB | 0.038449 | 0.517225 | Other Positive Correlation |
| TRIM17 | 0.038393 | 0.517843 | Other Positive Correlation |
| INPP5B | 0.038367 | 0.518127 | Other Positive Correlation |
| SLC1A6 | 0.038211 | 0.519828 | Other Positive Correlation |
| ZFP90 | 0.038201 | 0.519942 | Other Positive Correlation |
| RNF103-CHMP3 | 0.038186 | 0.520097 | Other Positive Correlation |
| TRMU | 0.038178 | 0.520192 | Other Positive Correlation |
| PHYKPL | 0.038037 | 0.521726 | Other Positive Correlation |
| MCM8 | 0.038031 | 0.521797 | Other Positive Correlation |
| SCGB3A2 | 0.03799 | 0.522248 | Other Positive Correlation |
| ZNF239 | 0.037816 | 0.524151 | Other Positive Correlation |
| KCNMB3 | 0.037753 | 0.524847 | Other Positive Correlation |
| CHRNE | 0.037721 | 0.525197 | Other Positive Correlation |
| SLC25A53 | 0.037678 | 0.525672 | Other Positive Correlation |
| BBC3 | 0.03764 | 0.526092 | Other Positive Correlation |
| USP2 | 0.037512 | 0.527497 | Other Positive Correlation |
| KLHL40 | 0.037439 | 0.528305 | Other Positive Correlation |
| TMX4 | 0.037411 | 0.528614 | Other Positive Correlation |
| RAB9B | 0.037244 | 0.530453 | Other Positive Correlation |
| PODXL | 0.03724 | 0.5305 | Other Positive Correlation |
| ZFHX2 | 0.037207 | 0.530861 | Other Positive Correlation |
| IL23R | 0.037207 | 0.530866 | Other Positive Correlation |
| EZH2 | 0.037177 | 0.531197 | Other Positive Correlation |
| INPP5K | 0.03711 | 0.531933 | Other Positive Correlation |

| | | | |
|---|---|---|---|
| GRIP2 | 0.037071 | 0.532369 | Other Positive Correlation |
| ALG10B | 0.037067 | 0.532408 | Other Positive Correlation |
| DTX3 | 0.03703 | 0.532822 | Other Positive Correlation |
| SLC35B1 | 0.037022 | 0.532911 | Other Positive Correlation |
| ADGRD1 | 0.036975 | 0.533432 | Other Positive Correlation |
| C17orf50 | 0.036817 | 0.535183 | Other Positive Correlation |
| TSPO2 | 0.036791 | 0.535468 | Other Positive Correlation |
| HSPB8 | 0.036788 | 0.535503 | Other Positive Correlation |
| ZNF549 | 0.036711 | 0.536354 | Other Positive Correlation |
| USH1C | 0.036667 | 0.536849 | Other Positive Correlation |
| KCNIP3 | 0.036561 | 0.538021 | Other Positive Correlation |
| PPP1R37 | 0.036544 | 0.538214 | Other Positive Correlation |
| CPT1A | 0.0365 | 0.538709 | Other Positive Correlation |
| CCDC17 | 0.036445 | 0.539323 | Other Positive Correlation |
| GGH | 0.036391 | 0.53992 | Other Positive Correlation |
| FRZB | 0.036289 | 0.541061 | Other Positive Correlation |
| ZC3H12B | 0.03624 | 0.541609 | Other Positive Correlation |
| CLCN4 | 0.036145 | 0.542665 | Other Positive Correlation |
| RAMP2 | 0.036143 | 0.542689 | Other Positive Correlation |
| RADX | 0.036023 | 0.544026 | Other Positive Correlation |
| OTUD7A | 0.035977 | 0.544541 | Other Positive Correlation |
| ZNF671 | 0.035972 | 0.544601 | Other Positive Correlation |
| TVP23A | 0.035931 | 0.545054 | Other Positive Correlation |
| CGRRF1 | 0.035921 | 0.545172 | Other Positive Correlation |
| SLF2 | 0.035917 | 0.54521 | Other Positive Correlation |
| ZNF501 | 0.035847 | 0.545999 | Other Positive Correlation |
| DAG1 | 0.035827 | 0.546217 | Other Positive Correlation |
| WDR17 | 0.035746 | 0.547131 | Other Positive Correlation |
| UBE2L5 | 0.035553 | 0.5493 | Other Positive Correlation |
| CARS1 | 0.035494 | 0.549963 | Other Positive Correlation |
| ACTA1 | 0.035489 | 0.550016 | Other Positive Correlation |
| PLAGL2 | 0.035481 | 0.550113 | Other Positive Correlation |
| FUT1 | 0.035477 | 0.550154 | Other Positive Correlation |
| SLC26A8 | 0.035468 | 0.550257 | Other Positive Correlation |
| CDK5RAP2 | 0.035459 | 0.550357 | Other Positive Correlation |
| HOXA6 | 0.035418 | 0.550819 | Other Positive Correlation |
| EPHB1 | 0.035365 | 0.551409 | Other Positive Correlation |
| BEST4 | 0.035349 | 0.551589 | Other Positive Correlation |
| AK7 | 0.035313 | 0.552 | Other Positive Correlation |
| L1CAM | 0.035286 | 0.552301 | Other Positive Correlation |
| AMN1 | 0.035263 | 0.55256 | Other Positive Correlation |
| TTLL3 | 0.035229 | 0.552947 | Other Positive Correlation |
| LAT2 | 0.035162 | 0.553703 | Other Positive Correlation |
| ENSG00000241489 | 0.035108 | 0.554312 | Other Positive Correlation |
| TM4SF19-DYNLT2B | 0.03506 | 0.554854 | Other Positive Correlation |
| ADGRV1 | 0.03505 | 0.554968 | Other Positive Correlation |

| | | | |
|---|---|---|---|
| EPB41L3 | 0.03505 | 0.554972 | Other Positive Correlation |
| HOXA5 | 0.035016 | 0.555346 | Other Positive Correlation |
| EMID1 | 0.035003 | 0.555498 | Other Positive Correlation |
| VEPH1 | 0.034974 | 0.555821 | Other Positive Correlation |
| ZNF700 | 0.034963 | 0.555944 | Other Positive Correlation |
| DMRT1 | 0.034953 | 0.556059 | Other Positive Correlation |
| VSTM5 | 0.034943 | 0.556171 | Other Positive Correlation |
| SATB2 | 0.034849 | 0.557235 | Other Positive Correlation |
| AARD | 0.034838 | 0.557367 | Other Positive Correlation |
| CACNA1A | 0.034832 | 0.55743 | Other Positive Correlation |
| ZNF391 | 0.0348 | 0.557793 | Other Positive Correlation |
| NCALD | 0.034781 | 0.558007 | Other Positive Correlation |
| NOMO2 | 0.034759 | 0.558254 | Other Positive Correlation |
| CREM | 0.034573 | 0.560371 | Other Positive Correlation |
| ZNF555 | 0.034567 | 0.560433 | Other Positive Correlation |
| PNMA8A | 0.034429 | 0.562005 | Other Positive Correlation |
| HOXA9 | 0.034401 | 0.562317 | Other Positive Correlation |
| OLIG2 | 0.034331 | 0.563122 | Other Positive Correlation |
| C1orf141 | 0.034331 | 0.563122 | Other Positive Correlation |
| NPNT | 0.034296 | 0.563512 | Other Positive Correlation |
| ZFP92 | 0.034207 | 0.564534 | Other Positive Correlation |
| GPR12 | 0.034182 | 0.564815 | Other Positive Correlation |
| SPATA6 | 0.034161 | 0.565048 | Other Positive Correlation |
| FAM81A | 0.034075 | 0.566031 | Other Positive Correlation |
| USP6 | 0.033991 | 0.566994 | Other Positive Correlation |
| MMP25 | 0.033866 | 0.568418 | Other Positive Correlation |
| RDH12 | 0.033847 | 0.568637 | Other Positive Correlation |
| TMEM255B | 0.033786 | 0.569337 | Other Positive Correlation |
| STX16-NPEPL1 | 0.033782 | 0.569383 | Other Positive Correlation |
| CLNK | 0.033743 | 0.569828 | Other Positive Correlation |
| LYPD6B | 0.033712 | 0.570175 | Other Positive Correlation |
| SERP2 | 0.033681 | 0.570532 | Other Positive Correlation |
| TMPRSS3 | 0.033677 | 0.57058 | Other Positive Correlation |
| FOXH1 | 0.03363 | 0.571118 | Other Positive Correlation |
| RANBP17 | 0.033627 | 0.571148 | Other Positive Correlation |
| NAGPA | 0.033529 | 0.572282 | Other Positive Correlation |
| CACNB1 | 0.033501 | 0.572597 | Other Positive Correlation |
| SPI1 | 0.033471 | 0.572938 | Other Positive Correlation |
| SLC2A14 | 0.033435 | 0.573358 | Other Positive Correlation |
| AQP1 | 0.033415 | 0.573579 | Other Positive Correlation |
| CALML6 | 0.03336 | 0.574214 | Other Positive Correlation |
| CFAP61 | 0.03336 | 0.574214 | Other Positive Correlation |
| RUSF1 | 0.033314 | 0.574739 | Other Positive Correlation |
| XYLB | 0.033216 | 0.575874 | Other Positive Correlation |
| ATXN1 | 0.033161 | 0.576502 | Other Positive Correlation |
| TRIM34 | 0.033146 | 0.576677 | Other Positive Correlation |

| | | | |
|---|---|---|---|
| KDM4A | 0.033103 | 0.577168 | Other Positive Correlation |
| ENSG00000284292 | 0.033049 | 0.577792 | Other Positive Correlation |
| PURG | 0.033013 | 0.578205 | Other Positive Correlation |
| TTC34 | 0.032973 | 0.57867 | Other Positive Correlation |
| RTN4RL1 | 0.032972 | 0.578683 | Other Positive Correlation |
| EFHC1 | 0.032959 | 0.578824 | Other Positive Correlation |
| IQCB1 | 0.032938 | 0.57907 | Other Positive Correlation |
| GINS3 | 0.032936 | 0.579093 | Other Positive Correlation |
| SPATA13 | 0.032917 | 0.57931 | Other Positive Correlation |
| GNB3 | 0.032907 | 0.579425 | Other Positive Correlation |
| CEP63 | 0.0329 | 0.579505 | Other Positive Correlation |
| LMAN2L | 0.032855 | 0.580025 | Other Positive Correlation |
| JOSD1 | 0.032834 | 0.580269 | Other Positive Correlation |
| ZNF417 | 0.032769 | 0.58102 | Other Positive Correlation |
| AGAP1 | 0.032684 | 0.581996 | Other Positive Correlation |
| CMTM4 | 0.032672 | 0.582136 | Other Positive Correlation |
| HOXC11 | 0.032671 | 0.582152 | Other Positive Correlation |
| TEF | 0.032507 | 0.584051 | Other Positive Correlation |
| RLBP1 | 0.032503 | 0.584094 | Other Positive Correlation |
| LRRN2 | 0.032466 | 0.584524 | Other Positive Correlation |
| LRRC26 | 0.032405 | 0.585226 | Other Positive Correlation |
| DNAJC1 | 0.032229 | 0.587271 | Other Positive Correlation |
| SVOPL | 0.032172 | 0.587927 | Other Positive Correlation |
| SNX16 | 0.032118 | 0.588556 | Other Positive Correlation |
| TBC1D1 | 0.032068 | 0.589144 | Other Positive Correlation |
| CCDC159 | 0.032065 | 0.589178 | Other Positive Correlation |
| CRISPLD2 | 0.032062 | 0.589212 | Other Positive Correlation |
| SOSTDC1 | 0.032054 | 0.589305 | Other Positive Correlation |
| APCDD1 | 0.03203 | 0.589583 | Other Positive Correlation |
| ZNF583 | 0.032017 | 0.589727 | Other Positive Correlation |
| NPPC | 0.032011 | 0.589803 | Other Positive Correlation |
| CDK20 | 0.031974 | 0.590229 | Other Positive Correlation |
| INSC | 0.031942 | 0.590603 | Other Positive Correlation |
| APOBEC3D | 0.031919 | 0.590873 | Other Positive Correlation |
| PTAR1 | 0.031908 | 0.590998 | Other Positive Correlation |
| HPN | 0.031894 | 0.591158 | Other Positive Correlation |
| THAP7 | 0.031867 | 0.591477 | Other Positive Correlation |
| DHRS2 | 0.031675 | 0.59372 | Other Positive Correlation |
| LARP1B | 0.031657 | 0.593928 | Other Positive Correlation |
| STAT5A | 0.031571 | 0.594925 | Other Positive Correlation |
| CCDC158 | 0.031566 | 0.594983 | Other Positive Correlation |
| NPY1R | 0.031555 | 0.595112 | Other Positive Correlation |
| ZNF736 | 0.031466 | 0.596154 | Other Positive Correlation |
| FAM178B | 0.031452 | 0.596312 | Other Positive Correlation |
| PHF10 | 0.031447 | 0.596377 | Other Positive Correlation |
| C1orf226 | 0.03143 | 0.596569 | Other Positive Correlation |

| | | | |
|---|---|---|---|
| SPATA24 | 0.031412 | 0.596791 | Other Positive Correlation |
| CEND1 | 0.031332 | 0.597723 | Other Positive Correlation |
| RASAL3 | 0.031332 | 0.597725 | Other Positive Correlation |
| IZUMO2 | 0.031194 | 0.599338 | Other Positive Correlation |
| ATPAF1 | 0.031189 | 0.59939 | Other Positive Correlation |
| C6orf136 | 0.031186 | 0.599434 | Other Positive Correlation |
| ZNF785 | 0.031168 | 0.599637 | Other Positive Correlation |
| SLC25A42 | 0.031125 | 0.600142 | Other Positive Correlation |
| ZCCHC18 | 0.031122 | 0.600182 | Other Positive Correlation |
| PPP1R15B | 0.031097 | 0.60047 | Other Positive Correlation |
| TXK | 0.031081 | 0.600655 | Other Positive Correlation |
| TRPC3 | 0.031081 | 0.600655 | Other Positive Correlation |
| TUBD1 | 0.03106 | 0.600904 | Other Positive Correlation |
| XKR8 | 0.031034 | 0.601207 | Other Positive Correlation |
| NEFM | 0.031016 | 0.601419 | Other Positive Correlation |
| BBS1 | 0.030946 | 0.602242 | Other Positive Correlation |
| LCK | 0.030933 | 0.602391 | Other Positive Correlation |
| ADPRH | 0.03088 | 0.603016 | Other Positive Correlation |
| C19orf44 | 0.030834 | 0.603556 | Other Positive Correlation |
| ENSG00000257390 | 0.030817 | 0.60376 | Other Positive Correlation |
| SULT4A1 | 0.030771 | 0.604293 | Other Positive Correlation |
| CYP39A1 | 0.030695 | 0.605197 | Other Positive Correlation |
| CAND2 | 0.030568 | 0.606685 | Other Positive Correlation |
| GMEB1 | 0.030511 | 0.607353 | Other Positive Correlation |
| FAM229A | 0.030501 | 0.607471 | Other Positive Correlation |
| ZNF157 | 0.030481 | 0.607711 | Other Positive Correlation |
| GAS7 | 0.030481 | 0.607715 | Other Positive Correlation |
| GNAO1 | 0.030426 | 0.608359 | Other Positive Correlation |
| ENSG00000288695 | 0.030423 | 0.60839 | Other Positive Correlation |
| FGD5 | 0.030423 | 0.6084 | Other Positive Correlation |
| SPOPL | 0.030399 | 0.608679 | Other Positive Correlation |
| TECTA | 0.030378 | 0.608926 | Other Positive Correlation |
| BICDL1 | 0.030361 | 0.609129 | Other Positive Correlation |
| FGF14 | 0.030308 | 0.609755 | Other Positive Correlation |
| RTKN2 | 0.030301 | 0.609838 | Other Positive Correlation |
| GSG1 | 0.030272 | 0.610181 | Other Positive Correlation |
| NDUFA4L2 | 0.030249 | 0.610452 | Other Positive Correlation |
| SOX10 | 0.030218 | 0.610813 | Other Positive Correlation |
| ARC | 0.030062 | 0.612655 | Other Positive Correlation |
| KLK4 | 0.030048 | 0.612819 | Other Positive Correlation |
| SMIM29 | 0.030036 | 0.612971 | Other Positive Correlation |
| PKN3 | 0.029872 | 0.614909 | Other Positive Correlation |
| DLEC1 | 0.029861 | 0.615037 | Other Positive Correlation |
| LARGE1 | 0.029808 | 0.615662 | Other Positive Correlation |
| CPZ | 0.029806 | 0.615691 | Other Positive Correlation |
| RLN1 | 0.029801 | 0.615749 | Other Positive Correlation |

| | | | |
|---|---|---|---|
| TRIM31 | 0.029684 | 0.617136 | Other Positive Correlation |
| MEI4 | 0.029589 | 0.61826 | Other Positive Correlation |
| PNMT | 0.029589 | 0.618268 | Other Positive Correlation |
| GPX3 | 0.029571 | 0.618475 | Other Positive Correlation |
| ARID3C | 0.029561 | 0.618597 | Other Positive Correlation |
| ZNF611 | 0.029494 | 0.619389 | Other Positive Correlation |
| TOR2A | 0.029475 | 0.619619 | Other Positive Correlation |
| LY6L | 0.02936 | 0.620981 | Other Positive Correlation |
| LGR6 | 0.02936 | 0.62099 | Other Positive Correlation |
| ZNF81 | 0.029344 | 0.621179 | Other Positive Correlation |
| NR4A3 | 0.029243 | 0.622377 | Other Positive Correlation |
| CRABP1 | 0.029195 | 0.622946 | Other Positive Correlation |
| TUBB8 | 0.029192 | 0.622981 | Other Positive Correlation |
| SNX30 | 0.029188 | 0.623029 | Other Positive Correlation |
| DHX37 | 0.029186 | 0.623057 | Other Positive Correlation |
| COLEC10 | 0.029161 | 0.623359 | Other Positive Correlation |
| CCNP | 0.029102 | 0.624056 | Other Positive Correlation |
| SLC29A2 | 0.029031 | 0.624902 | Other Positive Correlation |
| SLCO5A1 | 0.028864 | 0.6269 | Other Positive Correlation |
| RND1 | 0.028863 | 0.626915 | Other Positive Correlation |
| CTXN2 | 0.028766 | 0.628068 | Other Positive Correlation |
| ITM2A | 0.028762 | 0.628113 | Other Positive Correlation |
| ZNF682 | 0.028575 | 0.630357 | Other Positive Correlation |
| ZNF10 | 0.028453 | 0.631814 | Other Positive Correlation |
| JPH3 | 0.028427 | 0.632128 | Other Positive Correlation |
| TAFAZZIN | 0.028353 | 0.633017 | Other Positive Correlation |
| EXO5 | 0.028277 | 0.633922 | Other Positive Correlation |
| WAS | 0.028204 | 0.634808 | Other Positive Correlation |
| SCUBE1 | 0.028167 | 0.635243 | Other Positive Correlation |
| NANOG | 0.028059 | 0.636546 | Other Positive Correlation |
| SPATA1 | 0.028049 | 0.636667 | Other Positive Correlation |
| EMILIN3 | 0.027997 | 0.637286 | Other Positive Correlation |
| AANAT | 0.027969 | 0.637621 | Other Positive Correlation |
| CENPT | 0.027946 | 0.637909 | Other Positive Correlation |
| MMP16 | 0.027901 | 0.63844 | Other Positive Correlation |
| WDR25 | 0.0279 | 0.63846 | Other Positive Correlation |
| PUS10 | 0.02787 | 0.638814 | Other Positive Correlation |
| P2RX7 | 0.027813 | 0.639502 | Other Positive Correlation |
| CASP9 | 0.02775 | 0.640262 | Other Positive Correlation |
| AMH | 0.027746 | 0.640316 | Other Positive Correlation |
| CBX4 | 0.027679 | 0.641114 | Other Positive Correlation |
| KDM4D | 0.027654 | 0.641425 | Other Positive Correlation |
| RCOR2 | 0.027641 | 0.641573 | Other Positive Correlation |
| FLT1 | 0.027622 | 0.641806 | Other Positive Correlation |
| MAGI3 | 0.027548 | 0.642695 | Other Positive Correlation |
| ARHGAP24 | 0.027522 | 0.643018 | Other Positive Correlation |

| | | | |
|---|---|---|---|
| CRACR2B | 0.02752 | 0.643035 | Other Positive Correlation |
| FAM171B | 0.0275 | 0.643275 | Other Positive Correlation |
| GABBR1 | 0.027214 | 0.646738 | Other Positive Correlation |
| SPHK2 | 0.027209 | 0.646797 | Other Positive Correlation |
| GPR176 | 0.027172 | 0.647247 | Other Positive Correlation |
| RAB44 | 0.027161 | 0.647376 | Other Positive Correlation |
| NAGA | 0.027133 | 0.647719 | Other Positive Correlation |
| TRIM51 | 0.027077 | 0.648398 | Other Positive Correlation |
| H3Y2 | 0.027077 | 0.648398 | Other Positive Correlation |
| TMEM266 | 0.027062 | 0.648579 | Other Positive Correlation |
| KCNJ2 | 0.02694 | 0.650062 | Other Positive Correlation |
| SMIM10L2A | 0.026937 | 0.650088 | Other Positive Correlation |
| NRROS | 0.026836 | 0.651315 | Other Positive Correlation |
| PCED1A | 0.026825 | 0.651456 | Other Positive Correlation |
| SLC22A15 | 0.026763 | 0.652205 | Other Positive Correlation |
| NECTIN3 | 0.026757 | 0.652281 | Other Positive Correlation |
| IL4I1 | 0.026715 | 0.652786 | Other Positive Correlation |
| HHLA1 | 0.026709 | 0.65286 | Other Positive Correlation |
| TMEM255A | 0.026626 | 0.65387 | Other Positive Correlation |
| MOV10L1 | 0.02659 | 0.654309 | Other Positive Correlation |
| B3GAT2 | 0.026544 | 0.654865 | Other Positive Correlation |
| LRIT3 | 0.026469 | 0.655776 | Other Positive Correlation |
| FOXL2NB | 0.02645 | 0.656011 | Other Positive Correlation |
| PYGO2 | 0.026439 | 0.656138 | Other Positive Correlation |
| ZNF704 | 0.026428 | 0.656283 | Other Positive Correlation |
| SCIMP | 0.026426 | 0.656306 | Other Positive Correlation |
| ZDHHC23 | 0.026422 | 0.656355 | Other Positive Correlation |
| PPARGC1A | 0.026394 | 0.656686 | Other Positive Correlation |
| C17orf75 | 0.026389 | 0.656753 | Other Positive Correlation |
| AP3B2 | 0.026318 | 0.657618 | Other Positive Correlation |
| MSS51 | 0.026309 | 0.657721 | Other Positive Correlation |
| LCTL | 0.026262 | 0.658302 | Other Positive Correlation |
| KLHL22 | 0.026237 | 0.658605 | Other Positive Correlation |
| PTPRC | 0.026196 | 0.659107 | Other Positive Correlation |
| TMPRSS2 | 0.026174 | 0.659377 | Other Positive Correlation |
| PTCRA | 0.026144 | 0.659735 | Other Positive Correlation |
| CITED1 | 0.026144 | 0.659735 | Other Positive Correlation |
| GRM7 | 0.026144 | 0.659735 | Other Positive Correlation |
| HOXA4 | 0.026144 | 0.659735 | Other Positive Correlation |
| ITIH5 | 0.02612 | 0.660028 | Other Positive Correlation |
| FCSK | 0.025967 | 0.661894 | Other Positive Correlation |
| MED22 | 0.025962 | 0.661957 | Other Positive Correlation |
| ZCCHC14 | 0.025958 | 0.662009 | Other Positive Correlation |
| CHRFAM7A | 0.025934 | 0.662302 | Other Positive Correlation |
| SSU72P8 | 0.025917 | 0.662515 | Other Positive Correlation |
| FAM166A | 0.025917 | 0.662515 | Other Positive Correlation |

| | | | |
|---|---|---|---|
| KDM4E | 0.025917 | 0.662515 | Other Positive Correlation |
| SGSM1 | 0.025867 | 0.663124 | Other Positive Correlation |
| CARMIL2 | 0.025832 | 0.663551 | Other Positive Correlation |
| RASL10B | 0.025744 | 0.664624 | Other Positive Correlation |
| ZNF257 | 0.025731 | 0.664788 | Other Positive Correlation |
| LRRIQ4 | 0.025728 | 0.664816 | Other Positive Correlation |
| FOXD3 | 0.025666 | 0.665582 | Other Positive Correlation |
| HYLS1 | 0.025643 | 0.665862 | Other Positive Correlation |
| IL17C | 0.025517 | 0.667402 | Other Positive Correlation |
| PRPF19 | 0.025514 | 0.667438 | Other Positive Correlation |
| NPIPB8 | 0.025499 | 0.667624 | Other Positive Correlation |
| SLC45A2 | 0.025499 | 0.667624 | Other Positive Correlation |
| ENSG00000286221 | 0.025479 | 0.667869 | Other Positive Correlation |
| VSIG2 | 0.025452 | 0.668201 | Other Positive Correlation |
| TMEM52 | 0.025448 | 0.668254 | Other Positive Correlation |
| PITHD1 | 0.025388 | 0.668991 | Other Positive Correlation |
| BCL2L15 | 0.025386 | 0.669011 | Other Positive Correlation |
| ARHGEF17 | 0.025373 | 0.669168 | Other Positive Correlation |
| ABHD8 | 0.025353 | 0.669417 | Other Positive Correlation |
| MRPL46 | 0.025272 | 0.670406 | Other Positive Correlation |
| TBX6 | 0.025197 | 0.671328 | Other Positive Correlation |
| ZNF766 | 0.025123 | 0.672236 | Other Positive Correlation |
| FAM9B | 0.02502 | 0.673505 | Other Positive Correlation |
| TRIM63 | 0.025015 | 0.673567 | Other Positive Correlation |
| ZNF780B | 0.025003 | 0.673722 | Other Positive Correlation |
| KCNB2 | 0.024921 | 0.674722 | Other Positive Correlation |
| LINC02210-CRHR1 | 0.024876 | 0.675273 | Other Positive Correlation |
| MBD6 | 0.024837 | 0.675756 | Other Positive Correlation |
| ZNF19 | 0.024835 | 0.675784 | Other Positive Correlation |
| KHK | 0.024822 | 0.675939 | Other Positive Correlation |
| MUC5B | 0.024759 | 0.676719 | Other Positive Correlation |
| UGT8 | 0.024757 | 0.676751 | Other Positive Correlation |
| KANTR | 0.024679 | 0.677702 | Other Positive Correlation |
| SMG8 | 0.024665 | 0.677877 | Other Positive Correlation |
| TRIM49B | 0.024655 | 0.678002 | Other Positive Correlation |
| TCEAL7 | 0.02458 | 0.678931 | Other Positive Correlation |
| PARP2 | 0.024496 | 0.679967 | Other Positive Correlation |
| ZNF774 | 0.024477 | 0.680198 | Other Positive Correlation |
| PCDHGA8 | 0.024475 | 0.680225 | Other Positive Correlation |
| ACTR3C | 0.024461 | 0.680397 | Other Positive Correlation |
| FAIM2 | 0.024411 | 0.681016 | Other Positive Correlation |
| PKNOX2 | 0.024385 | 0.681331 | Other Positive Correlation |
| OLFML2B | 0.024371 | 0.681509 | Other Positive Correlation |
| CKB | 0.024344 | 0.681835 | Other Positive Correlation |
| TMSB15B.1 | 0.02424 | 0.683125 | Other Positive Correlation |
| EMSY | 0.024168 | 0.684013 | Other Positive Correlation |

| | | | |
|---|---|---|---|
| EPOR | 0.02415 | 0.684244 | Other Positive Correlation |
| ANKRD20A1 | 0.024143 | 0.684328 | Other Positive Correlation |
| PHTF2 | 0.024108 | 0.68476 | Other Positive Correlation |
| ADCY8 | 0.024077 | 0.685145 | Other Positive Correlation |
| TSKS | 0.024017 | 0.685892 | Other Positive Correlation |
| CCSAP | 0.023939 | 0.686847 | Other Positive Correlation |
| CSF1R | 0.023939 | 0.686848 | Other Positive Correlation |
| CPQ | 0.023931 | 0.686958 | Other Positive Correlation |
| CACNA1I | 0.023884 | 0.687535 | Other Positive Correlation |
| SLITRK5 | 0.023849 | 0.687974 | Other Positive Correlation |
| RHBDL3 | 0.023694 | 0.689885 | Other Positive Correlation |
| ZNF529 | 0.023637 | 0.690604 | Other Positive Correlation |
| NPIPB9 | 0.023605 | 0.691001 | Other Positive Correlation |
| OPRK1 | 0.023605 | 0.691001 | Other Positive Correlation |
| NEDD8-MDP1 | 0.023605 | 0.691001 | Other Positive Correlation |
| ENSG00000270249 | 0.023605 | 0.691001 | Other Positive Correlation |
| ATG16L1 | 0.023561 | 0.691548 | Other Positive Correlation |
| SFMBT1 | 0.023546 | 0.691729 | Other Positive Correlation |
| XKR9 | 0.023524 | 0.692003 | Other Positive Correlation |
| LOXL2 | 0.023517 | 0.692094 | Other Positive Correlation |
| RNASE4 | 0.023475 | 0.692617 | Other Positive Correlation |
| ZSCAN1 | 0.023438 | 0.693067 | Other Positive Correlation |
| MSH2 | 0.023389 | 0.693683 | Other Positive Correlation |
| CCDC149 | 0.023278 | 0.695056 | Other Positive Correlation |
| ACOT4 | 0.02327 | 0.695164 | Other Positive Correlation |
| ANO4 | 0.02319 | 0.696151 | Other Positive Correlation |
| NPC2 | 0.023115 | 0.697092 | Other Positive Correlation |
| HOXC5 | 0.023112 | 0.697132 | Other Positive Correlation |
| MSANTD4 | 0.023067 | 0.697693 | Other Positive Correlation |
| SEMA3D | 0.023007 | 0.698437 | Other Positive Correlation |
| ZNF569 | 0.022983 | 0.698731 | Other Positive Correlation |
| MAN1A1 | 0.022904 | 0.699725 | Other Positive Correlation |
| ZNF431 | 0.022902 | 0.69975 | Other Positive Correlation |
| OR52A1 | 0.022843 | 0.700479 | Other Positive Correlation |
| PABPC5 | 0.022751 | 0.701625 | Other Positive Correlation |
| SPICE1 | 0.022734 | 0.701844 | Other Positive Correlation |
| MAJIN | 0.022725 | 0.701953 | Other Positive Correlation |
| HCRTR1 | 0.022707 | 0.702181 | Other Positive Correlation |
| ZMAT5 | 0.02266 | 0.702767 | Other Positive Correlation |
| OR7C1 | 0.022659 | 0.702775 | Other Positive Correlation |
| ENSG00000261832 | 0.022639 | 0.703029 | Other Positive Correlation |
| RAB15 | 0.022614 | 0.703341 | Other Positive Correlation |
| ASPSCR1 | 0.022568 | 0.703924 | Other Positive Correlation |
| STRN4 | 0.022563 | 0.70398 | Other Positive Correlation |
| EYA4 | 0.022561 | 0.704005 | Other Positive Correlation |
| DCLRE1A | 0.022526 | 0.704442 | Other Positive Correlation |

| | | | |
|---|---|---|---|
| LZTR1 | 0.022514 | 0.704588 | Other Positive Correlation |
| GAB3 | 0.022482 | 0.704999 | Other Positive Correlation |
| CLIC2 | 0.022372 | 0.706369 | Other Positive Correlation |
| CENPO | 0.022318 | 0.707043 | Other Positive Correlation |
| PLOD3 | 0.022294 | 0.707343 | Other Positive Correlation |
| GHR | 0.022282 | 0.707494 | Other Positive Correlation |
| ZNF100 | 0.022275 | 0.707587 | Other Positive Correlation |
| CKMT2 | 0.022146 | 0.709206 | Other Positive Correlation |
| TBC1D24 | 0.02214 | 0.709274 | Other Positive Correlation |
| EPDR1 | 0.022074 | 0.710109 | Other Positive Correlation |
| PRR33 | 0.022065 | 0.710214 | Other Positive Correlation |
| SLC23A3 | 0.022036 | 0.710578 | Other Positive Correlation |
| MT1M | 0.022028 | 0.710687 | Other Positive Correlation |
| GCH1 | 0.022001 | 0.711024 | Other Positive Correlation |
| ZNF334 | 0.021964 | 0.711485 | Other Positive Correlation |
| CACNA1D | 0.021933 | 0.711877 | Other Positive Correlation |
| NATD1 | 0.021903 | 0.712256 | Other Positive Correlation |
| GLB1L | 0.021892 | 0.712389 | Other Positive Correlation |
| VEZF1 | 0.021857 | 0.712826 | Other Positive Correlation |
| PBLD | 0.021856 | 0.712837 | Other Positive Correlation |
| CEP44 | 0.021782 | 0.71377 | Other Positive Correlation |
| LRAT | 0.021666 | 0.71523 | Other Positive Correlation |
| SPDYA | 0.021623 | 0.715772 | Other Positive Correlation |
| TBL1X | 0.021615 | 0.715864 | Other Positive Correlation |
| MSX2 | 0.021486 | 0.717489 | Other Positive Correlation |
| LINGO3 | 0.021463 | 0.717786 | Other Positive Correlation |
| TULP2 | 0.021463 | 0.717786 | Other Positive Correlation |
| FOXB1 | 0.021463 | 0.717786 | Other Positive Correlation |
| AQP4 | 0.021463 | 0.717786 | Other Positive Correlation |
| MOG | 0.021369 | 0.718962 | Other Positive Correlation |
| WNT10B | 0.021354 | 0.719152 | Other Positive Correlation |
| LIN9 | 0.021322 | 0.719557 | Other Positive Correlation |
| ABCD2 | 0.021271 | 0.720196 | Other Positive Correlation |
| KRT3 | 0.021271 | 0.720196 | Other Positive Correlation |
| CPLX1 | 0.021248 | 0.720491 | Other Positive Correlation |
| CRB2 | 0.021243 | 0.720555 | Other Positive Correlation |
| MRAP2 | 0.021204 | 0.721042 | Other Positive Correlation |
| TNFRSF1B | 0.021097 | 0.722389 | Other Positive Correlation |
| RBM24 | 0.020957 | 0.724163 | Other Positive Correlation |
| HECW2 | 0.020885 | 0.725072 | Other Positive Correlation |
| BTBD17 | 0.020852 | 0.725488 | Other Positive Correlation |
| NMNAT3 | 0.020831 | 0.725753 | Other Positive Correlation |
| PCDHB7 | 0.020802 | 0.726123 | Other Positive Correlation |
| TRAF5 | 0.020791 | 0.726261 | Other Positive Correlation |
| TRIM49 | 0.020788 | 0.726299 | Other Positive Correlation |
| MYOM2 | 0.020782 | 0.726369 | Other Positive Correlation |

| | | | |
|---|---|---|---|
| CTAGE15 | 0.020722 | 0.727137 | Other Positive Correlation |
| GNA14 | 0.020578 | 0.728954 | Other Positive Correlation |
| DPAGT1 | 0.020543 | 0.729391 | Other Positive Correlation |
| NPIPA2 | 0.020515 | 0.729753 | Other Positive Correlation |
| CADM3 | 0.020513 | 0.729775 | Other Positive Correlation |
| NIM1K | 0.020504 | 0.729896 | Other Positive Correlation |
| HS1BP3 | 0.020449 | 0.730585 | Other Positive Correlation |
| ANOS1 | 0.02043 | 0.730832 | Other Positive Correlation |
| CEP192 | 0.020356 | 0.731769 | Other Positive Correlation |
| KLF14 | 0.020334 | 0.732044 | Other Positive Correlation |
| TNFSF12 | 0.020312 | 0.732321 | Other Positive Correlation |
| ADGRG2 | 0.020304 | 0.732418 | Other Positive Correlation |
| CEBPB | 0.020277 | 0.73276 | Other Positive Correlation |
| TAMALIN | 0.020244 | 0.733185 | Other Positive Correlation |
| MEIG1 | 0.020219 | 0.733498 | Other Positive Correlation |
| MAGEL2 | 0.020211 | 0.733604 | Other Positive Correlation |
| PRRT2 | 0.020172 | 0.734104 | Other Positive Correlation |
| TUBA8 | 0.020009 | 0.736166 | Other Positive Correlation |
| SALL4 | 0.020003 | 0.736237 | Other Positive Correlation |
| SZT2 | 0.019956 | 0.736843 | Other Positive Correlation |
| LRRK2 | 0.019853 | 0.738148 | Other Positive Correlation |
| OR10A2 | 0.019754 | 0.739405 | Other Positive Correlation |
| OIP5 | 0.01975 | 0.739457 | Other Positive Correlation |
| SARNP | 0.019653 | 0.740687 | Other Positive Correlation |
| CYB5RL | 0.019518 | 0.74241 | Other Positive Correlation |
| R3HCC1L | 0.019505 | 0.742577 | Other Positive Correlation |
| RHBDD3 | 0.0195 | 0.742642 | Other Positive Correlation |
| ZNF329 | 0.01948 | 0.742888 | Other Positive Correlation |
| JPH1 | 0.019461 | 0.743136 | Other Positive Correlation |
| RNF222 | 0.019399 | 0.743922 | Other Positive Correlation |
| ZNF540 | 0.019396 | 0.743957 | Other Positive Correlation |
| C19orf84 | 0.01938 | 0.744167 | Other Positive Correlation |
| CCDC187 | 0.019321 | 0.74492 | Other Positive Correlation |
| JMJD7-PLA2G4B | 0.019263 | 0.745659 | Other Positive Correlation |
| KRT32 | 0.019249 | 0.745832 | Other Positive Correlation |
| UNC13A | 0.01922 | 0.746203 | Other Positive Correlation |
| CCDC144A | 0.019127 | 0.747396 | Other Positive Correlation |
| REXO5 | 0.019125 | 0.747415 | Other Positive Correlation |
| PHC1 | 0.019099 | 0.747744 | Other Positive Correlation |
| MDK | 0.019039 | 0.748514 | Other Positive Correlation |
| NEURL2 | 0.019006 | 0.748938 | Other Positive Correlation |
| GALR2 | 0.018981 | 0.749255 | Other Positive Correlation |
| HEATR5A | 0.018872 | 0.750645 | Other Positive Correlation |
| SEMA3G | 0.018832 | 0.751164 | Other Positive Correlation |
| SPNS3 | 0.018798 | 0.751588 | Other Positive Correlation |
| SYNPO2 | 0.01875 | 0.752211 | Other Positive Correlation |

| | | | |
|---|---|---|---|
| ESPN | 0.018684 | 0.753048 | Other Positive Correlation |
| RASA4 | 0.018621 | 0.753863 | Other Positive Correlation |
| RNF149 | 0.018593 | 0.754213 | Other Positive Correlation |
| ZNF48 | 0.018532 | 0.754992 | Other Positive Correlation |
| RAVER2 | 0.018518 | 0.755169 | Other Positive Correlation |
| MSANTD1 | 0.01847 | 0.755789 | Other Positive Correlation |
| PIK3AP1 | 0.018373 | 0.757036 | Other Positive Correlation |
| C12orf56 | 0.018337 | 0.757487 | Other Positive Correlation |
| TUNAR | 0.018315 | 0.757769 | Other Positive Correlation |
| PDZRN4 | 0.018315 | 0.757769 | Other Positive Correlation |
| KCNA3 | 0.018315 | 0.757769 | Other Positive Correlation |
| ADAM7 | 0.018315 | 0.757769 | Other Positive Correlation |
| ANKRD30A | 0.018315 | 0.757769 | Other Positive Correlation |
| OR10H5 | 0.018315 | 0.757769 | Other Positive Correlation |
| PAGE5 | 0.018315 | 0.757769 | Other Positive Correlation |
| NXPH1 | 0.018315 | 0.757769 | Other Positive Correlation |
| ACKR1 | 0.018315 | 0.757769 | Other Positive Correlation |
| AMY2A | 0.018315 | 0.757769 | Other Positive Correlation |
| DYDC1 | 0.018315 | 0.757769 | Other Positive Correlation |
| COX6A2 | 0.018315 | 0.757769 | Other Positive Correlation |
| SLCO6A1 | 0.018315 | 0.757769 | Other Positive Correlation |
| ADGRB3 | 0.018315 | 0.757769 | Other Positive Correlation |
| RSPO2 | 0.018315 | 0.757769 | Other Positive Correlation |
| SYT4 | 0.018315 | 0.757769 | Other Positive Correlation |
| FGF4 | 0.018315 | 0.757769 | Other Positive Correlation |
| SLITRK3 | 0.018315 | 0.757769 | Other Positive Correlation |
| GDF10 | 0.018315 | 0.757769 | Other Positive Correlation |
| SI | 0.018315 | 0.757769 | Other Positive Correlation |
| EPHA3 | 0.018315 | 0.757769 | Other Positive Correlation |
| MMP20 | 0.018315 | 0.757769 | Other Positive Correlation |
| CCDC154 | 0.018289 | 0.758102 | Other Positive Correlation |
| KLLN | 0.018264 | 0.75843 | Other Positive Correlation |
| SYT5 | 0.018238 | 0.758756 | Other Positive Correlation |
| SLC22A31 | 0.018235 | 0.758799 | Other Positive Correlation |
| PLIN5 | 0.01815 | 0.759884 | Other Positive Correlation |
| FBXO30 | 0.018141 | 0.760011 | Other Positive Correlation |
| PLCH1 | 0.018127 | 0.760183 | Other Positive Correlation |
| GPC4 | 0.018064 | 0.760988 | Other Positive Correlation |
| BST1 | 0.018057 | 0.761077 | Other Positive Correlation |
| OR4F17 | 0.018029 | 0.761445 | Other Positive Correlation |
| MAGEE2 | 0.018029 | 0.761445 | Other Positive Correlation |
| NALF1 | 0.018029 | 0.761445 | Other Positive Correlation |
| DMGDH | 0.018023 | 0.761514 | Other Positive Correlation |
| VPREB3 | 0.01799 | 0.761946 | Other Positive Correlation |
| CGREF1 | 0.017913 | 0.762931 | Other Positive Correlation |
| ANKRD7 | 0.017872 | 0.763459 | Other Positive Correlation |

| | | | |
|---|---|---|---|
| IKZF4 | 0.017871 | 0.763475 | Other Positive Correlation |
| MOCS1 | 0.017772 | 0.764737 | Other Positive Correlation |
| KCTD6 | 0.01776 | 0.764901 | Other Positive Correlation |
| XNDC1N | 0.017749 | 0.765041 | Other Positive Correlation |
| RGS11 | 0.017737 | 0.765196 | Other Positive Correlation |
| SNAPC5 | 0.017679 | 0.76594 | Other Positive Correlation |
| CACNA2D1 | 0.01756 | 0.767465 | Other Positive Correlation |
| COQ7 | 0.017557 | 0.767507 | Other Positive Correlation |
| WASH6P | 0.017547 | 0.767629 | Other Positive Correlation |
| ADO | 0.017533 | 0.767811 | Other Positive Correlation |
| MCM5 | 0.017406 | 0.769455 | Other Positive Correlation |
| AUNIP | 0.017387 | 0.769697 | Other Positive Correlation |
| SRGAP2 | 0.017341 | 0.770291 | Other Positive Correlation |
| EYA1 | 0.017337 | 0.770339 | Other Positive Correlation |
| ZNF44 | 0.017303 | 0.770773 | Other Positive Correlation |
| ANKRD35 | 0.017291 | 0.770936 | Other Positive Correlation |
| MEGF9 | 0.01727 | 0.771195 | Other Positive Correlation |
| PCDHB10 | 0.017215 | 0.771914 | Other Positive Correlation |
| AEBP1 | 0.017211 | 0.771963 | Other Positive Correlation |
| MAP1LC3B2 | 0.017145 | 0.772815 | Other Positive Correlation |
| ENSG00000256950 | 0.017034 | 0.774248 | Other Positive Correlation |
| ZNF689 | 0.016974 | 0.775022 | Other Positive Correlation |
| SH3BP2 | 0.016921 | 0.775706 | Other Positive Correlation |
| TMEM169 | 0.016899 | 0.775985 | Other Positive Correlation |
| DPYSL3 | 0.016803 | 0.777224 | Other Positive Correlation |
| NBL1 | 0.016689 | 0.778692 | Other Positive Correlation |
| TLCD3B | 0.016658 | 0.779099 | Other Positive Correlation |
| KCNG2 | 0.016658 | 0.779099 | Other Positive Correlation |
| PELI3 | 0.016652 | 0.779176 | Other Positive Correlation |
| RUBCNL | 0.016594 | 0.779917 | Other Positive Correlation |
| DISP1 | 0.016587 | 0.780007 | Other Positive Correlation |
| ULK1 | 0.016519 | 0.780892 | Other Positive Correlation |
| BUD13 | 0.016434 | 0.78199 | Other Positive Correlation |
| TMEM125 | 0.016408 | 0.782326 | Other Positive Correlation |
| MLF1 | 0.016385 | 0.782624 | Other Positive Correlation |
| KMT5C | 0.016357 | 0.782991 | Other Positive Correlation |
| GRM1 | 0.016336 | 0.783255 | Other Positive Correlation |
| DYNLT5 | 0.016336 | 0.783255 | Other Positive Correlation |
| NMUR2 | 0.016336 | 0.783255 | Other Positive Correlation |
| SCGB1C2 | 0.016336 | 0.783255 | Other Positive Correlation |
| TACR1 | 0.016319 | 0.783477 | Other Positive Correlation |
| NTRK3 | 0.016319 | 0.783477 | Other Positive Correlation |
| TMEM210 | 0.016319 | 0.783477 | Other Positive Correlation |
| IGSF21 | 0.016319 | 0.783477 | Other Positive Correlation |
| CST2 | 0.016319 | 0.783477 | Other Positive Correlation |
| PTPRH | 0.016308 | 0.783625 | Other Positive Correlation |

| | | | |
|---|---|---|---|
| C2orf68 | 0.016091 | 0.786429 | Other Positive Correlation |
| CCDC30 | 0.016077 | 0.78661 | Other Positive Correlation |
| LRFN1 | 0.016036 | 0.787136 | Other Positive Correlation |
| GDPD5 | 0.016027 | 0.787265 | Other Positive Correlation |
| ENTPD2 | 0.015999 | 0.787628 | Other Positive Correlation |
| CCNE2 | 0.015995 | 0.787679 | Other Positive Correlation |
| MAP3K3 | 0.015984 | 0.787821 | Other Positive Correlation |
| ITIH6 | 0.015872 | 0.789274 | Other Positive Correlation |
| OBI1 | 0.015747 | 0.790893 | Other Positive Correlation |
| INSL4 | 0.015672 | 0.791867 | Other Positive Correlation |
| ZNF221 | 0.015496 | 0.79415 | Other Positive Correlation |
| ADCY3 | 0.015468 | 0.794505 | Other Positive Correlation |
| AKAP5 | 0.015443 | 0.794832 | Other Positive Correlation |
| CBX2 | 0.015442 | 0.794847 | Other Positive Correlation |
| LLGL1 | 0.015418 | 0.795163 | Other Positive Correlation |
| GSE1 | 0.015349 | 0.796052 | Other Positive Correlation |
| MT4 | 0.015318 | 0.796459 | Other Positive Correlation |
| GRIN1 | 0.01531 | 0.796567 | Other Positive Correlation |
| CLSTN3 | 0.015287 | 0.796867 | Other Positive Correlation |
| GRAMD1B | 0.015262 | 0.797188 | Other Positive Correlation |
| PCDHB5 | 0.01525 | 0.797345 | Other Positive Correlation |
| CNNM3 | 0.015191 | 0.798114 | Other Positive Correlation |
| TMUB2 | 0.015079 | 0.799566 | Other Positive Correlation |
| NXT1 | 0.015068 | 0.799712 | Other Positive Correlation |
| PPIP5K1 | 0.014977 | 0.800891 | Other Positive Correlation |
| C1orf232 | 0.014938 | 0.801407 | Other Positive Correlation |
| ENSG00000288721 | 0.014937 | 0.801413 | Other Positive Correlation |
| ENSG00000285953 | 0.014908 | 0.801799 | Other Positive Correlation |
| SDR16C5 | 0.014855 | 0.802479 | Other Positive Correlation |
| MATN1 | 0.014854 | 0.802491 | Other Positive Correlation |
| GRIK4 | 0.014841 | 0.802664 | Other Positive Correlation |
| FASTK | 0.014822 | 0.80291 | Other Positive Correlation |
| CYS1 | 0.014736 | 0.804032 | Other Positive Correlation |
| TMEM38B | 0.014722 | 0.804222 | Other Positive Correlation |
| EFR3B | 0.014713 | 0.804327 | Other Positive Correlation |
| RBM20 | 0.014712 | 0.804352 | Other Positive Correlation |
| PROM1 | 0.014704 | 0.804447 | Other Positive Correlation |
| CRISP2 | 0.01467 | 0.804898 | Other Positive Correlation |
| ARIH2 | 0.014662 | 0.805001 | Other Positive Correlation |
| SAP30L | 0.014656 | 0.805072 | Other Positive Correlation |
| INSYN1 | 0.014633 | 0.80537 | Other Positive Correlation |
| ZNF138 | 0.014587 | 0.805977 | Other Positive Correlation |
| CNTD1 | 0.014565 | 0.806257 | Other Positive Correlation |
| MEF2B | 0.014473 | 0.807465 | Other Positive Correlation |
| PKDREJ | 0.014418 | 0.808178 | Other Positive Correlation |
| HARBI1 | 0.014418 | 0.808183 | Other Positive Correlation |

| | | | |
|---|---|---|---|
| TRABD2A | 0.014402 | 0.808385 | Other Positive Correlation |
| EBAG9 | 0.014394 | 0.80849 | Other Positive Correlation |
| TMPRSS5 | 0.014279 | 0.809989 | Other Positive Correlation |
| PODXL2 | 0.014265 | 0.810182 | Other Positive Correlation |
| RASGRP3 | 0.014245 | 0.810441 | Other Positive Correlation |
| SUSD4 | 0.014244 | 0.810451 | Other Positive Correlation |
| LOX | 0.014238 | 0.810526 | Other Positive Correlation |
| COBL | 0.014147 | 0.811722 | Other Positive Correlation |
| CLXN | 0.014049 | 0.812996 | Other Positive Correlation |
| C2CD4D | 0.013991 | 0.81375 | Other Positive Correlation |
| CNOT1 | 0.013967 | 0.814074 | Other Positive Correlation |
| CREBZF | 0.013955 | 0.81422 | Other Positive Correlation |
| EMC9 | 0.01394 | 0.814416 | Other Positive Correlation |
| OOEP | 0.013934 | 0.814503 | Other Positive Correlation |
| ADAL | 0.013886 | 0.815123 | Other Positive Correlation |
| MFSD13A | 0.013791 | 0.816365 | Other Positive Correlation |
| SELPLG | 0.01378 | 0.816513 | Other Positive Correlation |
| EFCAB5 | 0.013755 | 0.816838 | Other Positive Correlation |
| DAAM2 | 0.013587 | 0.819035 | Other Positive Correlation |
| PIGM | 0.013543 | 0.819618 | Other Positive Correlation |
| OR3A2 | 0.01349 | 0.820302 | Other Positive Correlation |
| C11orf54 | 0.013481 | 0.820427 | Other Positive Correlation |
| EDC4 | 0.013468 | 0.820603 | Other Positive Correlation |
| TULP1 | 0.013433 | 0.821051 | Other Positive Correlation |
| NCAN | 0.013431 | 0.821075 | Other Positive Correlation |
| FTCDNL1 | 0.013428 | 0.821114 | Other Positive Correlation |
| CHSY3 | 0.013399 | 0.821502 | Other Positive Correlation |
| RBM46 | 0.013334 | 0.822349 | Other Positive Correlation |
| IKZF3 | 0.013163 | 0.824597 | Other Positive Correlation |
| TMEM232 | 0.013116 | 0.825213 | Other Positive Correlation |
| P4HTM | 0.013094 | 0.825501 | Other Positive Correlation |
| PABPC1L2B | 0.01308 | 0.825679 | Other Positive Correlation |
| AURKC | 0.013066 | 0.825871 | Other Positive Correlation |
| AEN | 0.01305 | 0.826069 | Other Positive Correlation |
| SLC9A3 | 0.01299 | 0.826858 | Other Positive Correlation |
| MARCHF3 | 0.01296 | 0.82725 | Other Positive Correlation |
| PRSS22 | 0.01294 | 0.827524 | Other Positive Correlation |
| OTOS | 0.012915 | 0.827848 | Other Positive Correlation |
| C2CD6 | 0.012905 | 0.82798 | Other Positive Correlation |
| RALGPS1 | 0.012787 | 0.829525 | Other Positive Correlation |
| CFAP45 | 0.01275 | 0.830013 | Other Positive Correlation |
| PGM3 | 0.012743 | 0.830098 | Other Positive Correlation |
| HTR1D | 0.012712 | 0.830508 | Other Positive Correlation |
| ZNF70 | 0.012672 | 0.831039 | Other Positive Correlation |
| GZMB | 0.012663 | 0.831152 | Other Positive Correlation |
| CCHCR1 | 0.012654 | 0.831276 | Other Positive Correlation |

| | | | |
|---|---|---|---|
| SLC17A9 | 0.012626 | 0.831638 | Other Positive Correlation |
| TTC23L | 0.012582 | 0.832221 | Other Positive Correlation |
| ZNF670-ZNF695 | 0.012575 | 0.832304 | Other Positive Correlation |
| ASCL5 | 0.012573 | 0.83234 | Other Positive Correlation |
| CELF5 | 0.012558 | 0.832537 | Other Positive Correlation |
| GPRC5C | 0.012504 | 0.833245 | Other Positive Correlation |
| C17orf67 | 0.012492 | 0.8334 | Other Positive Correlation |
| RTN3 | 0.012465 | 0.833755 | Other Positive Correlation |
| NBPF26 | 0.012448 | 0.833983 | Other Positive Correlation |
| B3GNT9 | 0.012397 | 0.834645 | Other Positive Correlation |
| GANAB | 0.01237 | 0.835001 | Other Positive Correlation |
| HINT2 | 0.012362 | 0.835103 | Other Positive Correlation |
| PIP4K2A | 0.012324 | 0.835605 | Other Positive Correlation |
| MTMR4 | 0.012321 | 0.835652 | Other Positive Correlation |
| TAF11L14 | 0.012309 | 0.835812 | Other Positive Correlation |
| ENSG00000283321 | 0.012309 | 0.835812 | Other Positive Correlation |
| ANKS4B | 0.012309 | 0.835812 | Other Positive Correlation |
| TRIM43B | 0.012309 | 0.835812 | Other Positive Correlation |
| CD8B2 | 0.012309 | 0.835812 | Other Positive Correlation |
| TAF11L13 | 0.012309 | 0.835812 | Other Positive Correlation |
| SLIT2 | 0.012298 | 0.835949 | Other Positive Correlation |
| FZD10 | 0.01226 | 0.836451 | Other Positive Correlation |
| ONECUT3 | 0.012247 | 0.836621 | Other Positive Correlation |
| ANO2 | 0.012207 | 0.837152 | Other Positive Correlation |
| TSNARE1 | 0.01219 | 0.837367 | Other Positive Correlation |
| ZIC2 | 0.012109 | 0.838444 | Other Positive Correlation |
| FBXO15 | 0.012104 | 0.838501 | Other Positive Correlation |
| PBX3 | 0.012001 | 0.83986 | Other Positive Correlation |
| TMEM114 | 0.011984 | 0.840086 | Other Positive Correlation |
| ENO4 | 0.011941 | 0.840648 | Other Positive Correlation |
| ASIP | 0.011927 | 0.840838 | Other Positive Correlation |
| PCDHA13 | 0.011927 | 0.840838 | Other Positive Correlation |
| ENSG00000269476 | 0.011927 | 0.840838 | Other Positive Correlation |
| CNBD2 | 0.011821 | 0.842234 | Other Positive Correlation |
| EIF5 | 0.011774 | 0.842845 | Other Positive Correlation |
| TMEM270 | 0.011697 | 0.843866 | Other Positive Correlation |
| BCOR | 0.011694 | 0.843901 | Other Positive Correlation |
| TUBB8B | 0.011666 | 0.84427 | Other Positive Correlation |
| CDK14 | 0.01161 | 0.845007 | Other Positive Correlation |
| ZNF69 | 0.011606 | 0.845057 | Other Positive Correlation |
| FOXP4 | 0.011566 | 0.845586 | Other Positive Correlation |
| ENSG00000276418 | 0.011415 | 0.847578 | Other Positive Correlation |
| CCDC102B | 0.01141 | 0.847647 | Other Positive Correlation |
| EAF2 | 0.011316 | 0.848883 | Other Positive Correlation |
| SPESP1 | 0.011289 | 0.849243 | Other Positive Correlation |
| CDC25A | 0.011289 | 0.84925 | Other Positive Correlation |

| | | | |
|---|---|---|---|
| PIK3R5 | 0.011264 | 0.849567 | Other Positive Correlation |
| CHDH | 0.011257 | 0.849661 | Other Positive Correlation |
| VAV1 | 0.011254 | 0.849708 | Other Positive Correlation |
| NFE2L1 | 0.011249 | 0.849771 | Other Positive Correlation |
| RTL5 | 0.011195 | 0.850484 | Other Positive Correlation |
| FAM90A27P | 0.01106 | 0.85226 | Other Positive Correlation |
| MYL11 | 0.011026 | 0.852721 | Other Positive Correlation |
| ARMCX4 | 0.010991 | 0.853179 | Other Positive Correlation |
| YPEL4 | 0.01097 | 0.853454 | Other Positive Correlation |
| SIAH3 | 0.010841 | 0.855163 | Other Positive Correlation |
| IL37 | 0.01083 | 0.855308 | Other Positive Correlation |
| SPEF1 | 0.010751 | 0.856353 | Other Positive Correlation |
| KLHL34 | 0.010667 | 0.857461 | Other Positive Correlation |
| PDK4 | 0.010653 | 0.857643 | Other Positive Correlation |
| OPRL1 | 0.010633 | 0.857914 | Other Positive Correlation |
| LEF1 | 0.010579 | 0.858624 | Other Positive Correlation |
| ZIK1 | 0.010564 | 0.858821 | Other Positive Correlation |
| LRRC14B | 0.010564 | 0.858821 | Other Positive Correlation |
| FUNDC2 | 0.010521 | 0.859392 | Other Positive Correlation |
| BRAF | 0.010508 | 0.859565 | Other Positive Correlation |
| FAM102B | 0.010489 | 0.85981 | Other Positive Correlation |
| RASL12 | 0.010428 | 0.86062 | Other Positive Correlation |
| DOCK10 | 0.010419 | 0.860738 | Other Positive Correlation |
| ENSG00000289027 | 0.010409 | 0.860871 | Other Positive Correlation |
| ZNF727 | 0.010344 | 0.861727 | Other Positive Correlation |
| HMGCLL1 | 0.010292 | 0.862414 | Other Positive Correlation |
| GMFG | 0.01026 | 0.862842 | Other Positive Correlation |
| GLI2 | 0.010244 | 0.863051 | Other Positive Correlation |
| HSPA6 | 0.010056 | 0.865541 | Other Positive Correlation |
| KLF12 | 0.010022 | 0.865994 | Other Positive Correlation |
| ZNF500 | 0.010019 | 0.866038 | Other Positive Correlation |
| MIS12 | 0.009908 | 0.867503 | Other Positive Correlation |
| ZNF816 | 0.009669 | 0.870673 | Other Positive Correlation |
| PRRT1 | 0.009596 | 0.871642 | Other Positive Correlation |
| GNS | 0.009563 | 0.872072 | Other Positive Correlation |
| SLC45A1 | 0.009411 | 0.874096 | Other Positive Correlation |
| PAFAH1B3 | 0.009392 | 0.874349 | Other Positive Correlation |
| MYO1A | 0.009258 | 0.876131 | Other Positive Correlation |
| MCOLN2 | 0.009257 | 0.876131 | Other Positive Correlation |
| CFAP65 | 0.009253 | 0.87619 | Other Positive Correlation |
| ZNF534 | 0.009248 | 0.876258 | Other Positive Correlation |
| CACHD1 | 0.009219 | 0.876647 | Other Positive Correlation |
| ZNF26 | 0.009216 | 0.876682 | Other Positive Correlation |
| MPDZ | 0.009209 | 0.87678 | Other Positive Correlation |
| TMEM121B | 0.009187 | 0.877064 | Other Positive Correlation |
| THAP3 | 0.009177 | 0.877198 | Other Positive Correlation |

| | | | |
|---|---|---|---|
| RAPGEF6 | 0.00916 | 0.877426 | Other Positive Correlation |
| DNAAF3 | 0.009138 | 0.877715 | Other Positive Correlation |
| SANBR | 0.009111 | 0.87807 | Other Positive Correlation |
| GBP4 | 0.009105 | 0.878156 | Other Positive Correlation |
| ZNF354C | 0.009046 | 0.878937 | Other Positive Correlation |
| ENSG00000284188 | 0.008973 | 0.879903 | Other Positive Correlation |
| HOXD13 | 0.008968 | 0.879976 | Other Positive Correlation |
| RLN3 | 0.008968 | 0.879976 | Other Positive Correlation |
| ZNF679 | 0.008968 | 0.879976 | Other Positive Correlation |
| HOXC10 | 0.008968 | 0.879976 | Other Positive Correlation |
| DBX2 | 0.008968 | 0.879976 | Other Positive Correlation |
| CALCR | 0.008965 | 0.880018 | Other Positive Correlation |
| NEK5 | 0.008959 | 0.880098 | Other Positive Correlation |
| TEC | 0.00895 | 0.880215 | Other Positive Correlation |
| ZNF347 | 0.008948 | 0.880236 | Other Positive Correlation |
| ROPN1L | 0.008905 | 0.880811 | Other Positive Correlation |
| TMED7-TICAM2 | 0.008871 | 0.881268 | Other Positive Correlation |
| TICRR | 0.008853 | 0.881503 | Other Positive Correlation |
| ZNF878 | 0.008779 | 0.882481 | Other Positive Correlation |
| TFF3 | 0.008771 | 0.88259 | Other Positive Correlation |
| PRR15 | 0.008752 | 0.882842 | Other Positive Correlation |
| PSEN2 | 0.008727 | 0.883183 | Other Positive Correlation |
| WIZ | 0.0086 | 0.884862 | Other Positive Correlation |
| TMEM251 | 0.008599 | 0.884872 | Other Positive Correlation |
| GRK3 | 0.008489 | 0.886344 | Other Positive Correlation |
| GAB2 | 0.008488 | 0.886352 | Other Positive Correlation |
| APOBEC3B | 0.008247 | 0.889563 | Other Positive Correlation |
| ADORA1 | 0.008245 | 0.889583 | Other Positive Correlation |
| IRX5 | 0.008221 | 0.889903 | Other Positive Correlation |
| ATP1A2 | 0.00822 | 0.889926 | Other Positive Correlation |
| PIWIL1 | 0.008206 | 0.890102 | Other Positive Correlation |
| PACS2 | 0.00813 | 0.891111 | Other Positive Correlation |
| ENSG00000268279 | 0.007963 | 0.893337 | Other Positive Correlation |
| OR3A1 | 0.007954 | 0.893466 | Other Positive Correlation |
| SLC51A | 0.007882 | 0.894416 | Other Positive Correlation |
| ENSG00000288698 | 0.007845 | 0.894916 | Other Positive Correlation |
| MXI1 | 0.007772 | 0.89588 | Other Positive Correlation |
| SYN1 | 0.007772 | 0.895889 | Other Positive Correlation |
| ERBB3 | 0.007753 | 0.896132 | Other Positive Correlation |
| APOB | 0.007734 | 0.896384 | Other Positive Correlation |
| MEGF11 | 0.007703 | 0.896809 | Other Positive Correlation |
| PRR36 | 0.00767 | 0.897243 | Other Positive Correlation |
| REM2 | 0.007615 | 0.897972 | Other Positive Correlation |
| ITGAX | 0.007546 | 0.898892 | Other Positive Correlation |
| TMPRSS11A | 0.007543 | 0.898934 | Other Positive Correlation |
| C8orf34 | 0.007508 | 0.899404 | Other Positive Correlation |

| | | | |
|---|---|---|---|
| ABCF2-H2BK1 | 0.007486 | 0.899698 | Other Positive Correlation |
| SYT11 | 0.007471 | 0.899889 | Other Positive Correlation |
| MBOAT1 | 0.007459 | 0.900061 | Other Positive Correlation |
| COMMD3-BMI1 | 0.007427 | 0.90048 | Other Positive Correlation |
| SH2D3C | 0.007412 | 0.900685 | Other Positive Correlation |
| ST7L | 0.0074 | 0.900839 | Other Positive Correlation |
| STOX2 | 0.007364 | 0.901316 | Other Positive Correlation |
| GALNT12 | 0.007304 | 0.902122 | Other Positive Correlation |
| CCDC183 | 0.007266 | 0.902633 | Other Positive Correlation |
| MTO1 | 0.00724 | 0.902968 | Other Positive Correlation |
| MROH8 | 0.007197 | 0.903545 | Other Positive Correlation |
| SP7 | 0.007197 | 0.903547 | Other Positive Correlation |
| F10 | 0.007197 | 0.903547 | Other Positive Correlation |
| ROR1 | 0.007182 | 0.903746 | Other Positive Correlation |
| KCTD8 | 0.00718 | 0.903771 | Other Positive Correlation |
| TBX3 | 0.007168 | 0.903929 | Other Positive Correlation |
| ZNF793 | 0.007152 | 0.904152 | Other Positive Correlation |
| SIX5 | 0.007085 | 0.905039 | Other Positive Correlation |
| FAAH | 0.007035 | 0.90571 | Other Positive Correlation |
| PIK3CB | 0.007029 | 0.905783 | Other Positive Correlation |
| DUOX2 | 0.006948 | 0.906875 | Other Positive Correlation |
| NRN1 | 0.006937 | 0.907023 | Other Positive Correlation |
| NKAIN3 | 0.006922 | 0.907217 | Other Positive Correlation |
| TRABD2B | 0.006905 | 0.907442 | Other Positive Correlation |
| ZNF835 | 0.006873 | 0.907865 | Other Positive Correlation |
| CIAO3 | 0.006871 | 0.907893 | Other Positive Correlation |
| UBAP1L | 0.006843 | 0.90827 | Other Positive Correlation |
| ERVK3-1 | 0.006796 | 0.908895 | Other Positive Correlation |
| AMER2 | 0.006717 | 0.90995 | Other Positive Correlation |
| FAHD1 | 0.006676 | 0.910501 | Other Positive Correlation |
| CCDC148 | 0.006624 | 0.911198 | Other Positive Correlation |
| RGS14 | 0.006583 | 0.911743 | Other Positive Correlation |
| NSUN4 | 0.006567 | 0.911959 | Other Positive Correlation |
| SLFN13 | 0.00656 | 0.912045 | Other Positive Correlation |
| BHMT | 0.006517 | 0.91262 | Other Positive Correlation |
| GNRHR | 0.006506 | 0.912769 | Other Positive Correlation |
| STIMATE | 0.00649 | 0.912985 | Other Positive Correlation |
| FBXL7 | 0.006456 | 0.913439 | Other Positive Correlation |
| RUNX3 | 0.006425 | 0.913858 | Other Positive Correlation |
| TSSK3 | 0.006388 | 0.914342 | Other Positive Correlation |
| PPP1R12C | 0.006377 | 0.914489 | Other Positive Correlation |
| HEPACAM | 0.006353 | 0.914815 | Other Positive Correlation |
| KHDRBS2 | 0.006353 | 0.914815 | Other Positive Correlation |
| SPAAR | 0.006345 | 0.914917 | Other Positive Correlation |
| FAM120AOS | 0.006293 | 0.915613 | Other Positive Correlation |
| CYP3A5 | 0.006276 | 0.915835 | Other Positive Correlation |

| | | | |
|---|---|---|---|
| TBL1Y | 0.006246 | 0.916243 | Other Positive Correlation |
| NFIL3 | 0.006194 | 0.916933 | Other Positive Correlation |
| CLCF1 | 0.006165 | 0.917326 | Other Positive Correlation |
| SFTPC | 0.006124 | 0.917869 | Other Positive Correlation |
| RWDD2A | 0.006085 | 0.918388 | Other Positive Correlation |
| ZSCAN26 | 0.006037 | 0.919038 | Other Positive Correlation |
| FARP1 | 0.005979 | 0.919815 | Other Positive Correlation |
| NPRL2 | 0.00597 | 0.919925 | Other Positive Correlation |
| CDC14B | 0.005957 | 0.920106 | Other Positive Correlation |
| CD79B | 0.005922 | 0.920575 | Other Positive Correlation |
| CHAF1A | 0.005919 | 0.920607 | Other Positive Correlation |
| ALG2 | 0.005918 | 0.920622 | Other Positive Correlation |
| NPIPB3 | 0.005916 | 0.920647 | Other Positive Correlation |
| MYB | 0.0059 | 0.920862 | Other Positive Correlation |
| FLCN | 0.005899 | 0.920878 | Other Positive Correlation |
| TRIM48 | 0.005868 | 0.921296 | Other Positive Correlation |
| ZC3H10 | 0.005829 | 0.921818 | Other Positive Correlation |
| NTRK1 | 0.005765 | 0.922666 | Other Positive Correlation |
| FBXW8 | 0.005744 | 0.922947 | Other Positive Correlation |
| DNAAF8 | 0.005704 | 0.923492 | Other Positive Correlation |
| SHPK | 0.00567 | 0.923945 | Other Positive Correlation |
| RANGRF | 0.005527 | 0.925849 | Other Positive Correlation |
| CES4A | 0.005448 | 0.92691 | Other Positive Correlation |
| HSPA1A | 0.00543 | 0.927147 | Other Positive Correlation |
| PLEKHG1 | 0.00539 | 0.927688 | Other Positive Correlation |
| MT1B | 0.005305 | 0.928829 | Other Positive Correlation |
| POLR2J3.1 | 0.005289 | 0.929033 | Other Positive Correlation |
| PCDH9 | 0.005283 | 0.929124 | Other Positive Correlation |
| UNC13B | 0.005276 | 0.929207 | Other Positive Correlation |
| FCHSD2 | 0.005213 | 0.93005 | Other Positive Correlation |
| BUD31 | 0.005166 | 0.930682 | Other Positive Correlation |
| CFAP298-TCP10L | 0.00515 | 0.930901 | Other Positive Correlation |
| HIP1 | 0.005141 | 0.931017 | Other Positive Correlation |
| HLA-F | 0.005034 | 0.932453 | Other Positive Correlation |
| INKA2 | 0.004917 | 0.934024 | Other Positive Correlation |
| DRD4 | 0.004844 | 0.93499 | Other Positive Correlation |
| SH3BP5L | 0.004762 | 0.936089 | Other Positive Correlation |
| STBD1 | 0.004748 | 0.93628 | Other Positive Correlation |
| ALOX5 | 0.00473 | 0.936522 | Other Positive Correlation |
| ADGRL4 | 0.00469 | 0.937053 | Other Positive Correlation |
| KLF11 | 0.004669 | 0.937342 | Other Positive Correlation |
| SPSB3 | 0.004634 | 0.937807 | Other Positive Correlation |
| CORT | 0.004631 | 0.937845 | Other Positive Correlation |
| SYT7 | 0.004628 | 0.937882 | Other Positive Correlation |
| SIGLEC11 | 0.00452 | 0.939339 | Other Positive Correlation |
| POLN | 0.004456 | 0.940192 | Other Positive Correlation |

| | | | |
|---|---|---|---|
| FKBP7 | 0.004452 | 0.940244 | Other Positive Correlation |
| SALL3 | 0.004259 | 0.942827 | Other Positive Correlation |
| NUP88 | 0.004259 | 0.942837 | Other Positive Correlation |
| AVPR2 | 0.004223 | 0.943313 | Other Positive Correlation |
| PDGFRL | 0.004219 | 0.943363 | Other Positive Correlation |
| GNE | 0.004182 | 0.943867 | Other Positive Correlation |
| F12 | 0.004107 | 0.944866 | Other Positive Correlation |
| ARSH | 0.004099 | 0.944972 | Other Positive Correlation |
| TRAK2 | 0.004023 | 0.946 | Other Positive Correlation |
| TP53TG3D | 0.003984 | 0.946511 | Other Positive Correlation |
| AKR1D1 | 0.003967 | 0.946741 | Other Positive Correlation |
| TREX2 | 0.003954 | 0.946914 | Other Positive Correlation |
| NUDT16 | 0.003926 | 0.947297 | Other Positive Correlation |
| NPR2 | 0.003888 | 0.94781 | Other Positive Correlation |
| L3MBTL1 | 0.003875 | 0.947978 | Other Positive Correlation |
| EDIL3 | 0.003782 | 0.949224 | Other Positive Correlation |
| TMEM221 | 0.003769 | 0.949396 | Other Positive Correlation |
| UPF1 | 0.003713 | 0.950156 | Other Positive Correlation |
| ALG9 | 0.003676 | 0.950645 | Other Positive Correlation |
| PPARA | 0.003675 | 0.950656 | Other Positive Correlation |
| EFCAB2 | 0.003671 | 0.950714 | Other Positive Correlation |
| CNEP1R1 | 0.003628 | 0.951292 | Other Positive Correlation |
| CFAP221 | 0.003624 | 0.951341 | Other Positive Correlation |
| ZNF154 | 0.003509 | 0.952886 | Other Positive Correlation |
| ACADS | 0.003498 | 0.953032 | Other Positive Correlation |
| TAPT1 | 0.00343 | 0.953945 | Other Positive Correlation |
| ACOT1 | 0.003424 | 0.954032 | Other Positive Correlation |
| PRAC1 | 0.003419 | 0.954095 | Other Positive Correlation |
| MELTF | 0.003361 | 0.954869 | Other Positive Correlation |
| KDM8 | 0.003339 | 0.955161 | Other Positive Correlation |
| PLAAT5 | 0.003306 | 0.955612 | Other Positive Correlation |
| TAS2R7 | 0.003306 | 0.955612 | Other Positive Correlation |
| FITM1 | 0.003293 | 0.955782 | Other Positive Correlation |
| ZNF214 | 0.003275 | 0.956025 | Other Positive Correlation |
| ATP2A1 | 0.003249 | 0.956369 | Other Positive Correlation |
| PPP1R12B | 0.003229 | 0.956641 | Other Positive Correlation |
| MT-ND4L | 0.003144 | 0.957783 | Other Positive Correlation |
| CDK11B | 0.003081 | 0.958633 | Other Positive Correlation |
| COX4I2 | 0.003032 | 0.959285 | Other Positive Correlation |
| FNDC1 | 0.002984 | 0.959922 | Other Positive Correlation |
| DDX43 | 0.002978 | 0.960006 | Other Positive Correlation |
| AS3MT | 0.002968 | 0.960147 | Other Positive Correlation |
| ZNF419 | 0.00294 | 0.960515 | Other Positive Correlation |
| CRYBB2 | 0.002918 | 0.960812 | Other Positive Correlation |
| FKRP | 0.002914 | 0.960868 | Other Positive Correlation |
| FCGBP | 0.002866 | 0.961514 | Other Positive Correlation |

| | | | |
|---|---|---|---|
| INAFM1 | 0.002853 | 0.961684 | Other Positive Correlation |
| HGFAC | 0.002794 | 0.962478 | Other Positive Correlation |
| LILRB1 | 0.002782 | 0.962636 | Other Positive Correlation |
| MIGA2 | 0.00264 | 0.964547 | Other Positive Correlation |
| MKS1 | 0.002573 | 0.965441 | Other Positive Correlation |
| ELK1 | 0.002565 | 0.965556 | Other Positive Correlation |
| GPR84 | 0.002513 | 0.966253 | Other Positive Correlation |
| DRC3 | 0.002507 | 0.966334 | Other Positive Correlation |
| TP53BP2 | 0.002489 | 0.966577 | Other Positive Correlation |
| VSIG10L2 | 0.002458 | 0.966987 | Other Positive Correlation |
| CLCNKB | 0.002409 | 0.967647 | Other Positive Correlation |
| ZNF263 | 0.00237 | 0.968164 | Other Positive Correlation |
| F8 | 0.002228 | 0.970079 | Other Positive Correlation |
| ZNF541 | 0.002224 | 0.970133 | Other Positive Correlation |
| SYCP1 | 0.002208 | 0.970349 | Other Positive Correlation |
| CEP85 | 0.002172 | 0.970825 | Other Positive Correlation |
| C5orf49 | 0.00214 | 0.971258 | Other Positive Correlation |
| EXD1 | 0.002103 | 0.971757 | Other Positive Correlation |
| C4orf51 | 0.002046 | 0.972518 | Other Positive Correlation |
| SLC7A7 | 0.001984 | 0.973345 | Other Positive Correlation |
| SLC30A8 | 0.001974 | 0.973481 | Other Positive Correlation |
| MATK | 0.001969 | 0.973557 | Other Positive Correlation |
| ZC3H12D | 0.001931 | 0.974065 | Other Positive Correlation |
| RABL2A | 0.001885 | 0.974678 | Other Positive Correlation |
| SLC22A4 | 0.001859 | 0.975028 | Other Positive Correlation |
| USP27X | 0.001821 | 0.975542 | Other Positive Correlation |
| IRS4 | 0.001787 | 0.975994 | Other Positive Correlation |
| SLC16A14 | 0.001701 | 0.977156 | Other Positive Correlation |
| MAD1L1 | 0.001587 | 0.978687 | Other Positive Correlation |
| EDA | 0.001559 | 0.979061 | Other Positive Correlation |
| HSPA14 | 0.001554 | 0.979128 | Other Positive Correlation |
| OSBP | 0.001504 | 0.979796 | Other Positive Correlation |
| IRF4 | 0.001438 | 0.980685 | Other Positive Correlation |
| SLC29A4 | 0.00142 | 0.980929 | Other Positive Correlation |
| SUV39H2 | 0.001344 | 0.98195 | Other Positive Correlation |
| TMEM50B | 0.001243 | 0.983296 | Other Positive Correlation |
| SGK2 | 0.00118 | 0.984155 | Other Positive Correlation |
| SLC35C2 | 0.001159 | 0.984427 | Other Positive Correlation |
| ZFP69 | 0.001157 | 0.984453 | Other Positive Correlation |
| LMO2 | 0.001128 | 0.984849 | Other Positive Correlation |
| LRTM1 | 0.001124 | 0.984907 | Other Positive Correlation |
| ZSCAN12 | 0.001039 | 0.986042 | Other Positive Correlation |
| CHCHD3 | 0.001006 | 0.986488 | Other Positive Correlation |
| DPH2 | 0.000899 | 0.98792 | Other Positive Correlation |
| IL16 | 0.000889 | 0.988058 | Other Positive Correlation |
| SDHAF1 | 0.000829 | 0.988862 | Other Positive Correlation |

| | | | |
|---|---|---|---|
| FBLN1 | 0.000816 | 0.989034 | Other Positive Correlation |
| B3GAT3 | 0.000796 | 0.989313 | Other Positive Correlation |
| SPN | 0.000751 | 0.989905 | Other Positive Correlation |
| ZSCAN22 | 0.000745 | 0.989993 | Other Positive Correlation |
| BANP | 0.000738 | 0.990083 | Other Positive Correlation |
| FEM1C | 0.000714 | 0.990405 | Other Positive Correlation |
| TRIM58 | 0.000591 | 0.992063 | Other Positive Correlation |
| THSD7A | 0.000572 | 0.992316 | Other Positive Correlation |
| ZFTRAF1 | 0.00057 | 0.992347 | Other Positive Correlation |
| NRSN1 | 0.000548 | 0.992642 | Other Positive Correlation |
| STUM | 0.000548 | 0.992642 | Other Positive Correlation |
| THAP11 | 0.000538 | 0.992766 | Other Positive Correlation |
| DESI2 | 0.000463 | 0.993785 | Other Positive Correlation |
| EDEM2 | 0.000461 | 0.993811 | Other Positive Correlation |
| CALHM6 | 0.000446 | 0.994007 | Other Positive Correlation |
| ADAMTS9 | 0.000416 | 0.994407 | Other Positive Correlation |
| RIBC2 | 0.000355 | 0.995233 | Other Positive Correlation |
| ADAMTS10 | 0.000314 | 0.995775 | Other Positive Correlation |
| ALMS1 | 0.000292 | 0.996083 | Other Positive Correlation |
| PAX1 | 0.000265 | 0.996442 | Other Positive Correlation |
| WNK3 | 0.000229 | 0.996927 | Other Positive Correlation |
| PCDHGB4 | 0.000156 | 0.997899 | Other Positive Correlation |
| FAM107B | 0.000117 | 0.998433 | Other Positive Correlation |
| FAM78B | 0.000104 | 0.998599 | Other Positive Correlation |
| GOLGA8M | 0.000103 | 0.998617 | Other Positive Correlation |
| TTC9 | 8.2E-05 | 0.998899 | Other Positive Correlation |
| ABLIM2 | 3.01E-05 | 0.999595 | Other Positive Correlation |
| CAPN6 | 2.44E-05 | 0.999672 | Other Positive Correlation |
| KLF1 | 2.42E-05 | 0.999675 | Other Positive Correlation |
| TMEM132B | 1.41E-05 | 0.99981 | Other Positive Correlation |
| CHRNA7 | 6.4E-06 | 0.999914 | Other Positive Correlation |
| MIEF2 | -1.8E-06 | 0.999976 | Other Negative Correlation |
| LINS1 | -3.6E-06 | 0.999952 | Other Negative Correlation |
| C6orf120 | -3.9E-05 | 0.999471 | Other Negative Correlation |
| PDE1B | -4.3E-05 | 0.999427 | Other Negative Correlation |
| NGRN | -4.4E-05 | 0.999413 | Other Negative Correlation |
| MRE11 | -0.0002 | 0.997363 | Other Negative Correlation |
| ZNF112 | -0.00026 | 0.996511 | Other Negative Correlation |
| ITIH4 | -0.00035 | 0.995281 | Other Negative Correlation |
| CREB3L3 | -0.00037 | 0.995037 | Other Negative Correlation |
| DGKQ | -0.00042 | 0.99433 | Other Negative Correlation |
| ME1 | -0.00043 | 0.994232 | Other Negative Correlation |
| SOCS2 | -0.00054 | 0.99279 | Other Negative Correlation |
| DUSP9 | -0.00057 | 0.992277 | Other Negative Correlation |
| ASTE1 | -0.00065 | 0.991308 | Other Negative Correlation |
| FASTKD3 | -0.00078 | 0.989508 | Other Negative Correlation |

| | | | |
|---|---|---|---|
| GP1BA | -0.00082 | 0.988976 | Other Negative Correlation |
| GRIK2 | -0.00087 | 0.988267 | Other Negative Correlation |
| MFSD2B | -0.00095 | 0.987214 | Other Negative Correlation |
| KDELR3 | -0.00096 | 0.987132 | Other Negative Correlation |
| TRO | -0.00096 | 0.987102 | Other Negative Correlation |
| ZNF16 | -0.00108 | 0.98546 | Other Negative Correlation |
| SLC41A2 | -0.00111 | 0.985037 | Other Negative Correlation |
| LPIN2 | -0.00112 | 0.984911 | Other Negative Correlation |
| SLFN11 | -0.00116 | 0.984463 | Other Negative Correlation |
| EXOC3L1 | -0.00123 | 0.983423 | Other Negative Correlation |
| IL11 | -0.00138 | 0.981409 | Other Negative Correlation |
| TCERG1L | -0.00139 | 0.98137 | Other Negative Correlation |
| RPRML | -0.00141 | 0.981005 | Other Negative Correlation |
| NEU4 | -0.00141 | 0.981005 | Other Negative Correlation |
| TMEM213 | -0.00141 | 0.981005 | Other Negative Correlation |
| RAB6C | -0.00141 | 0.981005 | Other Negative Correlation |
| HOXB13 | -0.00141 | 0.981005 | Other Negative Correlation |
| CHRM2 | -0.00141 | 0.981005 | Other Negative Correlation |
| HAND2 | -0.00141 | 0.981005 | Other Negative Correlation |
| MAPKAPK5 | -0.00144 | 0.980645 | Other Negative Correlation |
| FER1L6 | -0.00144 | 0.980627 | Other Negative Correlation |
| PEA15 | -0.00146 | 0.980369 | Other Negative Correlation |
| ENSG00000261582 | -0.00148 | 0.980137 | Other Negative Correlation |
| SMIM3 | -0.0016 | 0.978494 | Other Negative Correlation |
| ANKMY1 | -0.00163 | 0.978116 | Other Negative Correlation |
| E2F2 | -0.00165 | 0.977799 | Other Negative Correlation |
| GNAT2 | -0.0017 | 0.977128 | Other Negative Correlation |
| PLD5P1 | -0.0017 | 0.977116 | Other Negative Correlation |
| CRIP3 | -0.00173 | 0.976708 | Other Negative Correlation |
| ITGB3 | -0.00178 | 0.976098 | Other Negative Correlation |
| KCNC3 | -0.00182 | 0.975547 | Other Negative Correlation |
| PACSIN1 | -0.00183 | 0.975428 | Other Negative Correlation |
| NIPSNAP3B | -0.00198 | 0.97344 | Other Negative Correlation |
| ARMC12 | -0.00199 | 0.973249 | Other Negative Correlation |
| ZNF404 | -0.002 | 0.973096 | Other Negative Correlation |
| NOTCH2NLB | -0.00203 | 0.972745 | Other Negative Correlation |
| TEPSIN | -0.00205 | 0.972514 | Other Negative Correlation |
| GABRD | -0.00206 | 0.972358 | Other Negative Correlation |
| WBP2NL | -0.0021 | 0.971828 | Other Negative Correlation |
| B3GLCT | -0.00214 | 0.971264 | Other Negative Correlation |
| GUCA1A | -0.00218 | 0.970694 | Other Negative Correlation |
| ZNF77 | -0.0022 | 0.970389 | Other Negative Correlation |
| STOML1 | -0.00222 | 0.970236 | Other Negative Correlation |
| SPATA5L1 | -0.00229 | 0.969197 | Other Negative Correlation |
| NT5DC4 | -0.00231 | 0.968973 | Other Negative Correlation |
| MYO15A | -0.00239 | 0.967861 | Other Negative Correlation |

| | | | |
|---|---|---|---|
| PRDM6 | -0.00246 | 0.966993 | Other Negative Correlation |
| POMGNT2 | -0.00251 | 0.966298 | Other Negative Correlation |
| DENND2D | -0.00252 | 0.966143 | Other Negative Correlation |
| ZNF763 | -0.00256 | 0.965671 | Other Negative Correlation |
| RELT | -0.00257 | 0.965501 | Other Negative Correlation |
| PAPPA | -0.00262 | 0.96482 | Other Negative Correlation |
| SLC25A18 | -0.00263 | 0.964717 | Other Negative Correlation |
| CCDC9 | -0.00277 | 0.962828 | Other Negative Correlation |
| CEMP1 | -0.00282 | 0.962146 | Other Negative Correlation |
| PRSS2 | -0.00282 | 0.962146 | Other Negative Correlation |
| FAM124B | -0.00282 | 0.962146 | Other Negative Correlation |
| CLCN2 | -0.00288 | 0.961298 | Other Negative Correlation |
| ZNF662 | -0.00298 | 0.960022 | Other Negative Correlation |
| CEP120 | -0.0031 | 0.958382 | Other Negative Correlation |
| OTOP2 | -0.00317 | 0.957476 | Other Negative Correlation |
| ELMO1 | -0.00325 | 0.956308 | Other Negative Correlation |
| CNIH3 | -0.00328 | 0.955976 | Other Negative Correlation |
| STAC3 | -0.00331 | 0.955527 | Other Negative Correlation |
| ARHGAP4 | -0.00334 | 0.955164 | Other Negative Correlation |
| TLE6 | -0.00338 | 0.954552 | Other Negative Correlation |
| ZNF600 | -0.00343 | 0.953987 | Other Negative Correlation |
| RCSD1 | -0.00344 | 0.953852 | Other Negative Correlation |
| HFM1 | -0.00352 | 0.952682 | Other Negative Correlation |
| EFCAB7 | -0.00354 | 0.952444 | Other Negative Correlation |
| HSPA1L | -0.00359 | 0.951855 | Other Negative Correlation |
| ACACB | -0.0037 | 0.950282 | Other Negative Correlation |
| GABRB1 | -0.00374 | 0.949778 | Other Negative Correlation |
| TRIM54 | -0.00376 | 0.949584 | Other Negative Correlation |
| RBP5 | -0.00393 | 0.947244 | Other Negative Correlation |
| PDK3 | -0.00395 | 0.94691 | Other Negative Correlation |
| GSTM4 | -0.00396 | 0.946863 | Other Negative Correlation |
| COL23A1 | -0.00397 | 0.94671 | Other Negative Correlation |
| KCNF1 | -0.00397 | 0.946662 | Other Negative Correlation |
| GET4 | -0.00401 | 0.9462 | Other Negative Correlation |
| SV2C | -0.00402 | 0.946052 | Other Negative Correlation |
| RAB29 | -0.00405 | 0.945693 | Other Negative Correlation |
| SETDB2 | -0.00406 | 0.945504 | Other Negative Correlation |
| MSMP | -0.00411 | 0.944843 | Other Negative Correlation |
| UBE2D4 | -0.00415 | 0.944305 | Other Negative Correlation |
| SLC15A2 | -0.00415 | 0.944292 | Other Negative Correlation |
| CCPG1 | -0.00419 | 0.943807 | Other Negative Correlation |
| SPAG17 | -0.00423 | 0.943175 | Other Negative Correlation |
| ARHGEF33 | -0.00428 | 0.94255 | Other Negative Correlation |
| RTN4R | -0.00439 | 0.94112 | Other Negative Correlation |
| INPP5A | -0.00439 | 0.941067 | Other Negative Correlation |
| TAF11L12 | -0.00443 | 0.940514 | Other Negative Correlation |

| | | | |
|---|---|---|---|
| SCN4A | -0.00445 | 0.940221 | Other Negative Correlation |
| PDIA4 | -0.00455 | 0.938905 | Other Negative Correlation |
| PRAM1 | -0.00459 | 0.938382 | Other Negative Correlation |
| AKR1E2 | -0.00467 | 0.937316 | Other Negative Correlation |
| TMEM143 | -0.00469 | 0.937054 | Other Negative Correlation |
| ZNF35 | -0.0047 | 0.936959 | Other Negative Correlation |
| ARSK | -0.00474 | 0.936397 | Other Negative Correlation |
| OR2A7 | -0.00474 | 0.936324 | Other Negative Correlation |
| ERMAP | -0.00477 | 0.936017 | Other Negative Correlation |
| ENSG00000284956 | -0.00481 | 0.935415 | Other Negative Correlation |
| CDH24 | -0.00483 | 0.935215 | Other Negative Correlation |
| ZNF775 | -0.00483 | 0.935133 | Other Negative Correlation |
| KLK15 | -0.00485 | 0.93488 | Other Negative Correlation |
| ADGB | -0.00485 | 0.93488 | Other Negative Correlation |
| ESPNL | -0.00486 | 0.934845 | Other Negative Correlation |
| ADNP2 | -0.00486 | 0.934808 | Other Negative Correlation |
| SH2D6 | -0.00486 | 0.934763 | Other Negative Correlation |
| TRPM2 | -0.00493 | 0.933834 | Other Negative Correlation |
| GNRH2 | -0.00494 | 0.933677 | Other Negative Correlation |
| JAML | -0.00494 | 0.933677 | Other Negative Correlation |
| ATAD3C | -0.00495 | 0.933624 | Other Negative Correlation |
| DCAF11 | -0.00496 | 0.933391 | Other Negative Correlation |
| KLHDC7B | -0.00498 | 0.93323 | Other Negative Correlation |
| SENP3 | -0.00503 | 0.932508 | Other Negative Correlation |
| PIH1D1 | -0.00504 | 0.932348 | Other Negative Correlation |
| CAPN11 | -0.00505 | 0.932215 | Other Negative Correlation |
| OTUB2 | -0.00506 | 0.932168 | Other Negative Correlation |
| TRIM23 | -0.00507 | 0.932018 | Other Negative Correlation |
| ZNF230 | -0.00512 | 0.931238 | Other Negative Correlation |
| EPO | -0.00513 | 0.931115 | Other Negative Correlation |
| BTBD19 | -0.00514 | 0.931039 | Other Negative Correlation |
| SPECC1L | -0.00514 | 0.931014 | Other Negative Correlation |
| ZNF665 | -0.00515 | 0.930962 | Other Negative Correlation |
| ZNF195 | -0.00519 | 0.930372 | Other Negative Correlation |
| GRM5 | -0.00526 | 0.929477 | Other Negative Correlation |
| NFATC1 | -0.00528 | 0.929176 | Other Negative Correlation |
| ATXN7L3 | -0.00529 | 0.929059 | Other Negative Correlation |
| MBTPS1 | -0.00538 | 0.927814 | Other Negative Correlation |
| ZNF566 | -0.00541 | 0.927447 | Other Negative Correlation |
| ZCCHC4 | -0.00543 | 0.927189 | Other Negative Correlation |
| ZRANB1 | -0.00549 | 0.926396 | Other Negative Correlation |
| NRXN1 | -0.00557 | 0.925318 | Other Negative Correlation |
| RAG1 | -0.00565 | 0.924189 | Other Negative Correlation |
| ENOX1 | -0.0057 | 0.923583 | Other Negative Correlation |
| ZNF641 | -0.00573 | 0.923146 | Other Negative Correlation |
| PWWP2A | -0.00576 | 0.922729 | Other Negative Correlation |

| | | | |
|---|---|---|---|
| BCL2L11 | -0.00589 | 0.921041 | Other Negative Correlation |
| TGFBR3 | -0.00601 | 0.919378 | Other Negative Correlation |
| LCN12 | -0.00604 | 0.918988 | Other Negative Correlation |
| ZNF117 | -0.00605 | 0.918892 | Other Negative Correlation |
| MIS18A | -0.00606 | 0.918662 | Other Negative Correlation |
| SYNGAP1 | -0.00608 | 0.918483 | Other Negative Correlation |
| ZHX1-C8orf76 | -0.00613 | 0.917744 | Other Negative Correlation |
| MLC1 | -0.00625 | 0.916244 | Other Negative Correlation |
| DUX4 | -0.00625 | 0.916244 | Other Negative Correlation |
| HORMAD2 | -0.00625 | 0.916244 | Other Negative Correlation |
| MRO | -0.00625 | 0.916244 | Other Negative Correlation |
| RAX | -0.00625 | 0.916244 | Other Negative Correlation |
| PARD6A | -0.00634 | 0.914943 | Other Negative Correlation |
| BMPER | -0.00636 | 0.914689 | Other Negative Correlation |
| SAT2 | -0.00639 | 0.91431 | Other Negative Correlation |
| NUDT8 | -0.00639 | 0.914288 | Other Negative Correlation |
| ENSG00000260342 | -0.00641 | 0.91411 | Other Negative Correlation |
| C1orf127 | -0.00641 | 0.91411 | Other Negative Correlation |
| AOC1 | -0.00641 | 0.91411 | Other Negative Correlation |
| BLOC1S3 | -0.00643 | 0.913756 | Other Negative Correlation |
| SH2B3 | -0.00651 | 0.912746 | Other Negative Correlation |
| ABCA8 | -0.00651 | 0.912676 | Other Negative Correlation |
| IRF2BP1 | -0.00663 | 0.911065 | Other Negative Correlation |
| MIA3 | -0.00667 | 0.910614 | Other Negative Correlation |
| ZNF232 | -0.00671 | 0.909991 | Other Negative Correlation |
| BOD1 | -0.00675 | 0.90947 | Other Negative Correlation |
| CCDC181 | -0.00681 | 0.908675 | Other Negative Correlation |
| PEPD | -0.00683 | 0.908443 | Other Negative Correlation |
| FIBIN | -0.00685 | 0.908162 | Other Negative Correlation |
| CDKL5 | -0.00687 | 0.907902 | Other Negative Correlation |
| PRSS1 | -0.00687 | 0.907893 | Other Negative Correlation |
| RDH8 | -0.00687 | 0.907893 | Other Negative Correlation |
| PCDHGA6 | -0.00692 | 0.907219 | Other Negative Correlation |
| NR0B1 | -0.00695 | 0.906898 | Other Negative Correlation |
| SLC6A4 | -0.00705 | 0.905512 | Other Negative Correlation |
| KCNK3 | -0.00709 | 0.904949 | Other Negative Correlation |
| LHB | -0.00713 | 0.904388 | Other Negative Correlation |
| MINAR1 | -0.00721 | 0.903309 | Other Negative Correlation |
| DNAI1 | -0.00723 | 0.903167 | Other Negative Correlation |
| ZG16 | -0.00724 | 0.902932 | Other Negative Correlation |
| MAFK | -0.00726 | 0.902707 | Other Negative Correlation |
| C20orf96 | -0.00728 | 0.902451 | Other Negative Correlation |
| ZNF213 | -0.0073 | 0.902235 | Other Negative Correlation |
| NCAPD3 | -0.00732 | 0.901862 | Other Negative Correlation |
| MYL4 | -0.00736 | 0.901387 | Other Negative Correlation |
| AMDHD1 | -0.0074 | 0.900797 | Other Negative Correlation |

| | | | |
|---|---|---|---|
| NGEF | -0.00741 | 0.900669 | Other Negative Correlation |
| RASEF | -0.00742 | 0.900616 | Other Negative Correlation |
| HIVEP3 | -0.00745 | 0.90023 | Other Negative Correlation |
| EIF3CL | -0.00751 | 0.899356 | Other Negative Correlation |
| SLFN12L | -0.00754 | 0.899012 | Other Negative Correlation |
| C3AR1 | -0.00756 | 0.898751 | Other Negative Correlation |
| EML6 | -0.00764 | 0.897639 | Other Negative Correlation |
| UPB1 | -0.0077 | 0.896823 | Other Negative Correlation |
| ENSG00000237378 | -0.00775 | 0.896238 | Other Negative Correlation |
| ANKLE1 | -0.00776 | 0.89605 | Other Negative Correlation |
| PRAMEF14 | -0.00777 | 0.895912 | Other Negative Correlation |
| RPP30 | -0.00778 | 0.895819 | Other Negative Correlation |
| KLF17 | -0.0078 | 0.895531 | Other Negative Correlation |
| PNMA8C | -0.00781 | 0.895375 | Other Negative Correlation |
| ZNF415 | -0.00783 | 0.895093 | Other Negative Correlation |
| ARRDC2 | -0.00784 | 0.894988 | Other Negative Correlation |
| TMEM86B | -0.0079 | 0.894177 | Other Negative Correlation |
| ABHD10 | -0.00792 | 0.89392 | Other Negative Correlation |
| NANP | -0.00796 | 0.893432 | Other Negative Correlation |
| ZNF606 | -0.00804 | 0.89228 | Other Negative Correlation |
| AURKA | -0.00805 | 0.892228 | Other Negative Correlation |
| STX5 | -0.00806 | 0.891984 | Other Negative Correlation |
| SLC22A14 | -0.00813 | 0.891081 | Other Negative Correlation |
| TLE4 | -0.00815 | 0.890892 | Other Negative Correlation |
| C14orf180 | -0.00816 | 0.890678 | Other Negative Correlation |
| OR2AT4 | -0.00816 | 0.890678 | Other Negative Correlation |
| TTF2 | -0.00821 | 0.890094 | Other Negative Correlation |
| CPNE7 | -0.00823 | 0.88979 | Other Negative Correlation |
| LNP1 | -0.00825 | 0.889553 | Other Negative Correlation |
| KCNH8 | -0.00826 | 0.889327 | Other Negative Correlation |
| ITPRIPL1 | -0.00827 | 0.889232 | Other Negative Correlation |
| SLC39A3 | -0.00828 | 0.889083 | Other Negative Correlation |
| PLA2G4D | -0.00833 | 0.888421 | Other Negative Correlation |
| ZNF75D | -0.00833 | 0.8884 | Other Negative Correlation |
| ZNF43 | -0.00834 | 0.888282 | Other Negative Correlation |
| UST | -0.00838 | 0.887814 | Other Negative Correlation |
| NANS | -0.00846 | 0.886733 | Other Negative Correlation |
| POC5 | -0.00851 | 0.886123 | Other Negative Correlation |
| RPUSD2 | -0.00858 | 0.885157 | Other Negative Correlation |
| SCGB2B2 | -0.00867 | 0.883898 | Other Negative Correlation |
| CILP2 | -0.00884 | 0.881674 | Other Negative Correlation |
| RASA2 | -0.00888 | 0.88111 | Other Negative Correlation |
| DLX3 | -0.00891 | 0.880687 | Other Negative Correlation |
| GIP | -0.00898 | 0.879764 | Other Negative Correlation |
| CR2 | -0.009 | 0.879564 | Other Negative Correlation |
| SULT1C2 | -0.00903 | 0.879091 | Other Negative Correlation |

| | | | |
|---|---|---|---|
| TRIM68 | -0.00906 | 0.878748 | Other Negative Correlation |
| C19orf54 | -0.00915 | 0.877618 | Other Negative Correlation |
| OTOF | -0.00916 | 0.8774 | Other Negative Correlation |
| IL3RA | -0.00924 | 0.876384 | Other Negative Correlation |
| CACNA2D4 | -0.00925 | 0.876263 | Other Negative Correlation |
| MYLK3 | -0.00931 | 0.87542 | Other Negative Correlation |
| WFDC10B | -0.00931 | 0.875398 | Other Negative Correlation |
| CNNM4 | -0.00932 | 0.875235 | Other Negative Correlation |
| FLRT1 | -0.00936 | 0.874802 | Other Negative Correlation |
| TFAP2E | -0.00937 | 0.874694 | Other Negative Correlation |
| ABHD15 | -0.00937 | 0.874603 | Other Negative Correlation |
| PDGFRA | -0.00945 | 0.873517 | Other Negative Correlation |
| SRD5A2 | -0.00947 | 0.873277 | Other Negative Correlation |
| ADAMTS20 | -0.00947 | 0.873277 | Other Negative Correlation |
| IGHMBP2 | -0.00953 | 0.872566 | Other Negative Correlation |
| ING4 | -0.00957 | 0.872012 | Other Negative Correlation |
| NOL4 | -0.00958 | 0.871833 | Other Negative Correlation |
| CD34 | -0.0096 | 0.871609 | Other Negative Correlation |
| PCSK6 | -0.0096 | 0.871574 | Other Negative Correlation |
| CFAP99 | -0.00963 | 0.871126 | Other Negative Correlation |
| CCZ1 | -0.00966 | 0.870755 | Other Negative Correlation |
| BOC | -0.00972 | 0.869972 | Other Negative Correlation |
| FGF20 | -0.00977 | 0.8693 | Other Negative Correlation |
| PRMT7 | -0.00983 | 0.868473 | Other Negative Correlation |
| TTC38 | -0.00997 | 0.866702 | Other Negative Correlation |
| BCDIN3D | -0.01005 | 0.865637 | Other Negative Correlation |
| GALK1 | -0.01006 | 0.865518 | Other Negative Correlation |
| PCP4L1 | -0.01007 | 0.865368 | Other Negative Correlation |
| FKBP6 | -0.0101 | 0.864913 | Other Negative Correlation |
| EXTL1 | -0.01014 | 0.864444 | Other Negative Correlation |
| PIGL | -0.01014 | 0.864396 | Other Negative Correlation |
| MTMR7 | -0.01018 | 0.86392 | Other Negative Correlation |
| PLD4 | -0.01021 | 0.863512 | Other Negative Correlation |
| THAP2 | -0.01022 | 0.863407 | Other Negative Correlation |
| TRPM1 | -0.01023 | 0.863295 | Other Negative Correlation |
| CCDC92 | -0.01024 | 0.863045 | Other Negative Correlation |
| PCTP | -0.01026 | 0.862906 | Other Negative Correlation |
| CACTIN | -0.01026 | 0.862781 | Other Negative Correlation |
| CD19 | -0.01033 | 0.861867 | Other Negative Correlation |
| TRIM39 | -0.01034 | 0.861777 | Other Negative Correlation |
| ENSG00000260272 | -0.01037 | 0.861328 | Other Negative Correlation |
| VEGFD | -0.01037 | 0.861328 | Other Negative Correlation |
| THRSP | -0.01037 | 0.861328 | Other Negative Correlation |
| GPR65 | -0.01037 | 0.861328 | Other Negative Correlation |
| GNG8 | -0.01037 | 0.861328 | Other Negative Correlation |
| FCRL2 | -0.01037 | 0.861328 | Other Negative Correlation |

| | | | |
|---|---|---|---|
| CBY2 | -0.01037 | 0.861328 | Other Negative Correlation |
| ENSG00000274322 | -0.01037 | 0.861328 | Other Negative Correlation |
| ENSG00000249240 | -0.01037 | 0.861328 | Other Negative Correlation |
| FABP1 | -0.01037 | 0.861328 | Other Negative Correlation |
| AGMO | -0.01038 | 0.861212 | Other Negative Correlation |
| FGFR1 | -0.0104 | 0.861041 | Other Negative Correlation |
| CD302 | -0.01049 | 0.859798 | Other Negative Correlation |
| CATSPERG | -0.0105 | 0.859726 | Other Negative Correlation |
| CC2D1A | -0.0105 | 0.859716 | Other Negative Correlation |
| TMEM104 | -0.01054 | 0.859155 | Other Negative Correlation |
| FLVCR2 | -0.01054 | 0.859146 | Other Negative Correlation |
| WDR83 | -0.01059 | 0.858508 | Other Negative Correlation |
| TMEM158 | -0.01062 | 0.858086 | Other Negative Correlation |
| RPA1 | -0.01062 | 0.858061 | Other Negative Correlation |
| HOXA1 | -0.01064 | 0.857829 | Other Negative Correlation |
| MLYCD | -0.01067 | 0.857416 | Other Negative Correlation |
| ZNF316 | -0.01073 | 0.856687 | Other Negative Correlation |
| RDH5 | -0.01077 | 0.856148 | Other Negative Correlation |
| PEG10 | -0.01077 | 0.856067 | Other Negative Correlation |
| MAN2B1 | -0.0108 | 0.855644 | Other Negative Correlation |
| CASKIN1 | -0.01092 | 0.854119 | Other Negative Correlation |
| AAK1 | -0.01096 | 0.853606 | Other Negative Correlation |
| LCLAT1 | -0.01096 | 0.853537 | Other Negative Correlation |
| CAMKV | -0.01104 | 0.852578 | Other Negative Correlation |
| C16orf90 | -0.01104 | 0.852578 | Other Negative Correlation |
| PYY | -0.01111 | 0.851637 | Other Negative Correlation |
| ESR1 | -0.01112 | 0.851461 | Other Negative Correlation |
| PAPOLB | -0.01114 | 0.851248 | Other Negative Correlation |
| FAM53B | -0.01118 | 0.850638 | Other Negative Correlation |
| TACR2 | -0.0112 | 0.850401 | Other Negative Correlation |
| CXXC1 | -0.01123 | 0.850064 | Other Negative Correlation |
| CCDC117 | -0.01126 | 0.849611 | Other Negative Correlation |
| SARM1 | -0.01128 | 0.849383 | Other Negative Correlation |
| ZNF605 | -0.01129 | 0.849259 | Other Negative Correlation |
| CCDC33 | -0.0113 | 0.84905 | Other Negative Correlation |
| FLACC1 | -0.01147 | 0.846903 | Other Negative Correlation |
| PCDHB8 | -0.0115 | 0.846478 | Other Negative Correlation |
| CELF3 | -0.01152 | 0.846195 | Other Negative Correlation |
| SPATA17 | -0.0116 | 0.845202 | Other Negative Correlation |
| RYR2 | -0.01164 | 0.844624 | Other Negative Correlation |
| MYO5C | -0.01169 | 0.84399 | Other Negative Correlation |
| ZNF253 | -0.0117 | 0.843815 | Other Negative Correlation |
| AIPL1 | -0.01173 | 0.843489 | Other Negative Correlation |
| ENSG00000267318 | -0.01173 | 0.843395 | Other Negative Correlation |
| C4orf47 | -0.01177 | 0.842867 | Other Negative Correlation |
| PTPRO | -0.01178 | 0.842801 | Other Negative Correlation |

| | | | |
|---|---|---|---|
| HEPN1 | -0.01179 | 0.842619 | Other Negative Correlation |
| CSF2RA | -0.01183 | 0.842075 | Other Negative Correlation |
| NOL4L | -0.01185 | 0.841857 | Other Negative Correlation |
| ENSG00000261611 | -0.01187 | 0.841549 | Other Negative Correlation |
| ABCC6 | -0.01192 | 0.84093 | Other Negative Correlation |
| ZNF420 | -0.01194 | 0.84071 | Other Negative Correlation |
| GOLGA8A | -0.01198 | 0.840129 | Other Negative Correlation |
| THAP1 | -0.012 | 0.839935 | Other Negative Correlation |
| KLHDC9 | -0.01201 | 0.839758 | Other Negative Correlation |
| CYP26B1 | -0.01204 | 0.839307 | Other Negative Correlation |
| SCYL3 | -0.01206 | 0.839079 | Other Negative Correlation |
| TMEM91 | -0.01207 | 0.838983 | Other Negative Correlation |
| PCDHA9 | -0.01211 | 0.838382 | Other Negative Correlation |
| TSPAN13 | -0.01213 | 0.838189 | Other Negative Correlation |
| RAB4B-EGLN2 | -0.01216 | 0.837724 | Other Negative Correlation |
| C1QTNF12 | -0.0122 | 0.837299 | Other Negative Correlation |
| ZNF454 | -0.01221 | 0.837066 | Other Negative Correlation |
| LPA | -0.01221 | 0.837043 | Other Negative Correlation |
| MS4A15 | -0.01221 | 0.837043 | Other Negative Correlation |
| ENSG00000286001 | -0.01221 | 0.837043 | Other Negative Correlation |
| CELA3A | -0.01221 | 0.837043 | Other Negative Correlation |
| COL19A1 | -0.01221 | 0.837043 | Other Negative Correlation |
| ASB10 | -0.01221 | 0.837043 | Other Negative Correlation |
| GLOD5 | -0.01221 | 0.837043 | Other Negative Correlation |
| ENSG00000259529 | -0.01221 | 0.837043 | Other Negative Correlation |
| RNF165 | -0.01223 | 0.836861 | Other Negative Correlation |
| HCN3 | -0.01223 | 0.836849 | Other Negative Correlation |
| LTF | -0.01229 | 0.836098 | Other Negative Correlation |
| ZNF441 | -0.0123 | 0.835969 | Other Negative Correlation |
| WDR88 | -0.01231 | 0.835809 | Other Negative Correlation |
| SLC25A20 | -0.01245 | 0.833905 | Other Negative Correlation |
| PURB | -0.01248 | 0.83357 | Other Negative Correlation |
| EPHA4 | -0.01252 | 0.833095 | Other Negative Correlation |
| CRACR2A | -0.01253 | 0.832896 | Other Negative Correlation |
| ABCC4 | -0.01255 | 0.832702 | Other Negative Correlation |
| PCCA | -0.01258 | 0.832291 | Other Negative Correlation |
| POU5F2 | -0.01263 | 0.831537 | Other Negative Correlation |
| CTRC | -0.01264 | 0.831436 | Other Negative Correlation |
| SLC16A12 | -0.01272 | 0.830387 | Other Negative Correlation |
| CDKL3 | -0.01277 | 0.829688 | Other Negative Correlation |
| CRY2 | -0.0128 | 0.829342 | Other Negative Correlation |
| MYCT1 | -0.01283 | 0.828905 | Other Negative Correlation |
| LRRC56 | -0.013 | 0.826695 | Other Negative Correlation |
| SLC39A11 | -0.01303 | 0.826373 | Other Negative Correlation |
| UBR7 | -0.01305 | 0.826049 | Other Negative Correlation |
| IZUMO4 | -0.01307 | 0.825869 | Other Negative Correlation |

| | | | |
|---|---|---|---|
| SENP1 | -0.01308 | 0.825657 | Other Negative Correlation |
| GALT | -0.01311 | 0.825252 | Other Negative Correlation |
| FOXG1 | -0.01317 | 0.824561 | Other Negative Correlation |
| ZNF385C | -0.01317 | 0.824498 | Other Negative Correlation |
| NBEA | -0.01318 | 0.824391 | Other Negative Correlation |
| RUBCN | -0.01321 | 0.823971 | Other Negative Correlation |
| CFAP161 | -0.01323 | 0.823769 | Other Negative Correlation |
| ACSBG1 | -0.01323 | 0.823679 | Other Negative Correlation |
| SMPD3 | -0.01324 | 0.82357 | Other Negative Correlation |
| PPFIA4 | -0.01328 | 0.823021 | Other Negative Correlation |
| GPLD1 | -0.01333 | 0.822377 | Other Negative Correlation |
| BRINP1 | -0.01344 | 0.821016 | Other Negative Correlation |
| C19orf18 | -0.01346 | 0.820767 | Other Negative Correlation |
| CCDC87 | -0.01347 | 0.82062 | Other Negative Correlation |
| GSDMB | -0.01356 | 0.819346 | Other Negative Correlation |
| ENSG00000260914 | -0.01358 | 0.819089 | Other Negative Correlation |
| NT5C1A | -0.0136 | 0.818917 | Other Negative Correlation |
| SLC52A1 | -0.01362 | 0.818608 | Other Negative Correlation |
| GPR37 | -0.01365 | 0.818241 | Other Negative Correlation |
| OTX1 | -0.01373 | 0.817137 | Other Negative Correlation |
| FAM169A | -0.01374 | 0.816994 | Other Negative Correlation |
| ZNF705A | -0.01378 | 0.81657 | Other Negative Correlation |
| RSPH14 | -0.01378 | 0.816491 | Other Negative Correlation |
| ZSCAN20 | -0.01381 | 0.816056 | Other Negative Correlation |
| CERK | -0.01383 | 0.81582 | Other Negative Correlation |
| VWF | -0.01384 | 0.815695 | Other Negative Correlation |
| FAM53A | -0.01385 | 0.815649 | Other Negative Correlation |
| MATCAP2 | -0.01392 | 0.814742 | Other Negative Correlation |
| AZU1 | -0.01392 | 0.814714 | Other Negative Correlation |
| KLHL25 | -0.01396 | 0.814131 | Other Negative Correlation |
| ZNF317 | -0.01402 | 0.813404 | Other Negative Correlation |
| ADAM21 | -0.01402 | 0.813388 | Other Negative Correlation |
| FAM227B | -0.01409 | 0.8125 | Other Negative Correlation |
| DCAF16 | -0.01413 | 0.811972 | Other Negative Correlation |
| NID2 | -0.01417 | 0.81145 | Other Negative Correlation |
| FAM227A | -0.01417 | 0.811445 | Other Negative Correlation |
| BCL7B | -0.01417 | 0.811429 | Other Negative Correlation |
| FKTN | -0.01418 | 0.811266 | Other Negative Correlation |
| ARHGEF26 | -0.01421 | 0.810836 | Other Negative Correlation |
| PIM1 | -0.0143 | 0.809685 | Other Negative Correlation |
| CSF2RB | -0.01432 | 0.80945 | Other Negative Correlation |
| VN1R1 | -0.01432 | 0.80945 | Other Negative Correlation |
| ZNF181 | -0.01432 | 0.809417 | Other Negative Correlation |
| KLB | -0.01435 | 0.809125 | Other Negative Correlation |
| GLRB | -0.01437 | 0.808842 | Other Negative Correlation |
| ZMAT4 | -0.01439 | 0.808576 | Other Negative Correlation |

| | | | |
|---|---|---|---|
| CATSPER4 | -0.01439 | 0.808576 | Other Negative Correlation |
| OR1I1 | -0.01439 | 0.808576 | Other Negative Correlation |
| TAF7L | -0.0144 | 0.808424 | Other Negative Correlation |
| UNC5CL | -0.0144 | 0.80839 | Other Negative Correlation |
| ZNF771 | -0.01441 | 0.808327 | Other Negative Correlation |
| C6orf163 | -0.01442 | 0.808115 | Other Negative Correlation |
| ISYNA1 | -0.01443 | 0.807998 | Other Negative Correlation |
| RIMBP3C | -0.01447 | 0.807529 | Other Negative Correlation |
| BMAL1 | -0.01448 | 0.807306 | Other Negative Correlation |
| FAM240C | -0.01452 | 0.80684 | Other Negative Correlation |
| ENSG00000249141 | -0.01454 | 0.806642 | Other Negative Correlation |
| ITGA5 | -0.0146 | 0.805787 | Other Negative Correlation |
| CTNS | -0.01461 | 0.805733 | Other Negative Correlation |
| HOXC4 | -0.01471 | 0.80442 | Other Negative Correlation |
| DDN | -0.01472 | 0.804195 | Other Negative Correlation |
| OR1B1 | -0.01473 | 0.804062 | Other Negative Correlation |
| CC2D2B | -0.01474 | 0.804019 | Other Negative Correlation |
| DUSP28 | -0.01476 | 0.803744 | Other Negative Correlation |
| N4BP2L2 | -0.01482 | 0.80295 | Other Negative Correlation |
| ZNF543 | -0.01483 | 0.80279 | Other Negative Correlation |
| RECQL5 | -0.01488 | 0.802219 | Other Negative Correlation |
| APOLD1 | -0.0149 | 0.801853 | Other Negative Correlation |
| SPEM1 | -0.01494 | 0.80133 | Other Negative Correlation |
| PRDM8 | -0.01523 | 0.797584 | Other Negative Correlation |
| SEPTIN5 | -0.01524 | 0.797489 | Other Negative Correlation |
| EFCAB11 | -0.01525 | 0.797381 | Other Negative Correlation |
| TXNDC12 | -0.0153 | 0.796667 | Other Negative Correlation |
| SLC9B1 | -0.01533 | 0.796296 | Other Negative Correlation |
| ADAMTS19 | -0.01533 | 0.796276 | Other Negative Correlation |
| VCPKMT | -0.01535 | 0.796012 | Other Negative Correlation |
| TEK | -0.01536 | 0.795911 | Other Negative Correlation |
| PYCR2 | -0.0154 | 0.795387 | Other Negative Correlation |
| SCAI | -0.01541 | 0.795314 | Other Negative Correlation |
| MRPL18 | -0.01541 | 0.795257 | Other Negative Correlation |
| SLC28A1 | -0.01548 | 0.79432 | Other Negative Correlation |
| LRRN4 | -0.01553 | 0.793734 | Other Negative Correlation |
| FBXO24 | -0.01557 | 0.793172 | Other Negative Correlation |
| GPR132 | -0.01559 | 0.792983 | Other Negative Correlation |
| FAM118A | -0.01562 | 0.792528 | Other Negative Correlation |
| DCLRE1B | -0.01563 | 0.792352 | Other Negative Correlation |
| KCNJ12 | -0.01564 | 0.792262 | Other Negative Correlation |
| LY6G5C | -0.01567 | 0.791945 | Other Negative Correlation |
| LRRC37A3 | -0.01569 | 0.791625 | Other Negative Correlation |
| SIAE | -0.01571 | 0.791305 | Other Negative Correlation |
| CELF4 | -0.01572 | 0.79127 | Other Negative Correlation |
| ZNF594 | -0.01572 | 0.791181 | Other Negative Correlation |

| | | | |
|---|---|---|---|
| FNDC10 | -0.01579 | 0.79034 | Other Negative Correlation |
| CHST6 | -0.01583 | 0.789777 | Other Negative Correlation |
| ANKRD24 | -0.01595 | 0.788209 | Other Negative Correlation |
| ACOT2 | -0.01597 | 0.788001 | Other Negative Correlation |
| GALNT15 | -0.01598 | 0.787832 | Other Negative Correlation |
| TRPV3 | -0.016 | 0.78758 | Other Negative Correlation |
| CLDN10 | -0.016 | 0.787567 | Other Negative Correlation |
| RNF122 | -0.01601 | 0.787531 | Other Negative Correlation |
| PCDHA3 | -0.01604 | 0.787149 | Other Negative Correlation |
| NAPSA | -0.01613 | 0.785899 | Other Negative Correlation |
| WIPI1 | -0.01618 | 0.785293 | Other Negative Correlation |
| SYTL3 | -0.01621 | 0.784944 | Other Negative Correlation |
| EVA1B | -0.01621 | 0.784923 | Other Negative Correlation |
| CCDC153 | -0.01632 | 0.783497 | Other Negative Correlation |
| IL11RA | -0.01634 | 0.783204 | Other Negative Correlation |
| SLC39A1 | -0.01636 | 0.782943 | Other Negative Correlation |
| DNAH7 | -0.0164 | 0.782429 | Other Negative Correlation |
| EXOSC1 | -0.01646 | 0.781704 | Other Negative Correlation |
| ADAMTS14 | -0.01648 | 0.781382 | Other Negative Correlation |
| ZC4H2 | -0.01653 | 0.780708 | Other Negative Correlation |
| ZC2HC1C | -0.01655 | 0.780478 | Other Negative Correlation |
| NAA30 | -0.01664 | 0.779344 | Other Negative Correlation |
| NHLH2 | -0.01671 | 0.778415 | Other Negative Correlation |
| ZNF578 | -0.01673 | 0.778185 | Other Negative Correlation |
| TXNDC11 | -0.01675 | 0.777851 | Other Negative Correlation |
| SCNN1A | -0.01679 | 0.777407 | Other Negative Correlation |
| C8orf44-SGK3 | -0.0168 | 0.777284 | Other Negative Correlation |
| OGDHL | -0.01692 | 0.775696 | Other Negative Correlation |
| AKR1B15 | -0.01694 | 0.775416 | Other Negative Correlation |
| C5orf22 | -0.01702 | 0.774374 | Other Negative Correlation |
| RNF24 | -0.01708 | 0.773618 | Other Negative Correlation |
| SLC4A2 | -0.01713 | 0.773032 | Other Negative Correlation |
| SAMD5 | -0.01714 | 0.772929 | Other Negative Correlation |
| ZBTB20 | -0.0172 | 0.772139 | Other Negative Correlation |
| GTF3C2 | -0.0172 | 0.772053 | Other Negative Correlation |
| MIXL1 | -0.01723 | 0.771779 | Other Negative Correlation |
| ENTPD4 | -0.01727 | 0.771169 | Other Negative Correlation |
| ATF7IP2 | -0.01734 | 0.770323 | Other Negative Correlation |
| OR2B8P | -0.01738 | 0.769825 | Other Negative Correlation |
| STIMATE-MUSTN1 | -0.01738 | 0.769825 | Other Negative Correlation |
| SPEF2 | -0.01742 | 0.769263 | Other Negative Correlation |
| TLX1 | -0.01742 | 0.769225 | Other Negative Correlation |
| PCDHGB5 | -0.01744 | 0.76907 | Other Negative Correlation |
| KCNMB4 | -0.01745 | 0.76887 | Other Negative Correlation |
| CKLF-CMTM1 | -0.01747 | 0.768675 | Other Negative Correlation |
| SLC23A2 | -0.01748 | 0.768519 | Other Negative Correlation |

| | | | |
|---|---|---|---|
| AKR1A1 | -0.01756 | 0.76745 | Other Negative Correlation |
| CEP76 | -0.01758 | 0.76725 | Other Negative Correlation |
| LRGUK | -0.0176 | 0.766997 | Other Negative Correlation |
| SLC6A11 | -0.0176 | 0.766984 | Other Negative Correlation |
| ZFP42 | -0.01763 | 0.766559 | Other Negative Correlation |
| B3GALNT1 | -0.0177 | 0.765717 | Other Negative Correlation |
| HYCC2 | -0.01776 | 0.764951 | Other Negative Correlation |
| CHD1 | -0.01778 | 0.764684 | Other Negative Correlation |
| HAMP | -0.01778 | 0.764626 | Other Negative Correlation |
| PKD1L2 | -0.01778 | 0.764578 | Other Negative Correlation |
| CLPSL1 | -0.01786 | 0.763581 | Other Negative Correlation |
| STAT5B | -0.01787 | 0.763489 | Other Negative Correlation |
| TJAP1 | -0.01788 | 0.763297 | Other Negative Correlation |
| ZNF852 | -0.01789 | 0.76328 | Other Negative Correlation |
| SEC16B | -0.01789 | 0.763203 | Other Negative Correlation |
| AHRR | -0.01789 | 0.763181 | Other Negative Correlation |
| RYBP | -0.01793 | 0.762696 | Other Negative Correlation |
| ZNF701 | -0.01798 | 0.762078 | Other Negative Correlation |
| HECTD2 | -0.01801 | 0.761687 | Other Negative Correlation |
| FAM131B | -0.01813 | 0.760165 | Other Negative Correlation |
| B3GALT6 | -0.01816 | 0.759813 | Other Negative Correlation |
| CHKA | -0.0182 | 0.759298 | Other Negative Correlation |
| CPNE5 | -0.01824 | 0.758722 | Other Negative Correlation |
| ZCWPW1 | -0.01827 | 0.758306 | Other Negative Correlation |
| PTPN4 | -0.01836 | 0.757259 | Other Negative Correlation |
| KDM4C | -0.01841 | 0.756584 | Other Negative Correlation |
| HENMT1 | -0.01842 | 0.756451 | Other Negative Correlation |
| EPHA8 | -0.01843 | 0.756359 | Other Negative Correlation |
| MBD3L2 | -0.01847 | 0.755823 | Other Negative Correlation |
| IQCH | -0.01849 | 0.75554 | Other Negative Correlation |
| ABHD1 | -0.01853 | 0.755066 | Other Negative Correlation |
| TRIM55 | -0.01853 | 0.75504 | Other Negative Correlation |
| DLG4 | -0.01854 | 0.754856 | Other Negative Correlation |
| NKX3-1 | -0.01859 | 0.75423 | Other Negative Correlation |
| H3-5 | -0.01859 | 0.754196 | Other Negative Correlation |
| RPN2 | -0.01866 | 0.753369 | Other Negative Correlation |
| EID2B | -0.01869 | 0.753032 | Other Negative Correlation |
| GPR180 | -0.01871 | 0.752696 | Other Negative Correlation |
| C16orf87 | -0.01873 | 0.752524 | Other Negative Correlation |
| CDH18 | -0.01877 | 0.751934 | Other Negative Correlation |
| GSTA1 | -0.01877 | 0.751934 | Other Negative Correlation |
| METTL7B | -0.01881 | 0.751478 | Other Negative Correlation |
| ERG | -0.01881 | 0.751465 | Other Negative Correlation |
| ZFP62 | -0.01882 | 0.751365 | Other Negative Correlation |
| ZNF681 | -0.01883 | 0.751199 | Other Negative Correlation |
| PDE3B | -0.01892 | 0.750015 | Other Negative Correlation |

| | | | |
|---|---|---|---|
| VSIG10L | -0.01898 | 0.749295 | Other Negative Correlation |
| SLC25A19 | -0.01898 | 0.749254 | Other Negative Correlation |
| PRSS35 | -0.01899 | 0.749148 | Other Negative Correlation |
| HABP2 | -0.01902 | 0.74879 | Other Negative Correlation |
| TMIGD2 | -0.01902 | 0.74879 | Other Negative Correlation |
| CUL4B | -0.01902 | 0.748774 | Other Negative Correlation |
| TTC31 | -0.01902 | 0.748706 | Other Negative Correlation |
| TMEM178A | -0.01903 | 0.748625 | Other Negative Correlation |
| AVIL | -0.0191 | 0.747682 | Other Negative Correlation |
| ACSM3 | -0.01914 | 0.747191 | Other Negative Correlation |
| DHRS11 | -0.01916 | 0.747024 | Other Negative Correlation |
| FBLN7 | -0.01916 | 0.746967 | Other Negative Correlation |
| NEURL4 | -0.01923 | 0.746016 | Other Negative Correlation |
| FAM222A | -0.01924 | 0.74592 | Other Negative Correlation |
| ELAPOR2 | -0.01925 | 0.745874 | Other Negative Correlation |
| PCDH19 | -0.01925 | 0.745816 | Other Negative Correlation |
| SPOCK3 | -0.01926 | 0.745732 | Other Negative Correlation |
| SPATA4 | -0.01929 | 0.745273 | Other Negative Correlation |
| TMEM8B | -0.0193 | 0.745223 | Other Negative Correlation |
| MAP1LC3A | -0.0193 | 0.745191 | Other Negative Correlation |
| YBEY | -0.0193 | 0.745157 | Other Negative Correlation |
| CA5B | -0.01936 | 0.744435 | Other Negative Correlation |
| PARD6B | -0.01937 | 0.744255 | Other Negative Correlation |
| TIGD3 | -0.01949 | 0.742787 | Other Negative Correlation |
| TSNAXIP1 | -0.01949 | 0.742712 | Other Negative Correlation |
| TTC39A | -0.0195 | 0.742605 | Other Negative Correlation |
| SWI5 | -0.01951 | 0.742517 | Other Negative Correlation |
| WBP1 | -0.01952 | 0.742426 | Other Negative Correlation |
| UROC1 | -0.01973 | 0.739707 | Other Negative Correlation |
| ENSG00000275674 | -0.01973 | 0.739707 | Other Negative Correlation |
| ALX4 | -0.01976 | 0.739322 | Other Negative Correlation |
| TESPA1 | -0.01976 | 0.739322 | Other Negative Correlation |
| FRMPD4 | -0.01976 | 0.739322 | Other Negative Correlation |
| PABPC3 | -0.01981 | 0.738695 | Other Negative Correlation |
| FAM43B | -0.01985 | 0.738216 | Other Negative Correlation |
| CPA1 | -0.01988 | 0.737866 | Other Negative Correlation |
| CYRIA | -0.01989 | 0.73771 | Other Negative Correlation |
| ENSG00000225528 | -0.01992 | 0.737275 | Other Negative Correlation |
| SPIB | -0.01993 | 0.737131 | Other Negative Correlation |
| CFAP52 | -0.01997 | 0.736651 | Other Negative Correlation |
| RBM44 | -0.01997 | 0.736606 | Other Negative Correlation |
| ING2 | -0.01999 | 0.73644 | Other Negative Correlation |
| GOSR2 | -0.02005 | 0.735692 | Other Negative Correlation |
| CATSPERB | -0.02008 | 0.735316 | Other Negative Correlation |
| GBA2 | -0.02009 | 0.735078 | Other Negative Correlation |
| ZKSCAN2 | -0.02011 | 0.734905 | Other Negative Correlation |

| | | | |
|---|---|---|---|
| TMEM63C | -0.02012 | 0.734799 | Other Negative Correlation |
| C2CD2 | -0.02013 | 0.734633 | Other Negative Correlation |
| ZNF184 | -0.02019 | 0.733882 | Other Negative Correlation |
| RAB3A | -0.0202 | 0.733754 | Other Negative Correlation |
| GSC2 | -0.02023 | 0.733378 | Other Negative Correlation |
| SNX22 | -0.02025 | 0.733113 | Other Negative Correlation |
| TCEANC | -0.0203 | 0.732489 | Other Negative Correlation |
| MCIDAS | -0.02037 | 0.731594 | Other Negative Correlation |
| NCKAP1L | -0.02038 | 0.731452 | Other Negative Correlation |
| ZFP64 | -0.02045 | 0.730581 | Other Negative Correlation |
| PACRG | -0.02046 | 0.730418 | Other Negative Correlation |
| SERPIND1 | -0.02049 | 0.730049 | Other Negative Correlation |
| TOGARAM2 | -0.0205 | 0.729989 | Other Negative Correlation |
| ATAT1 | -0.02052 | 0.729704 | Other Negative Correlation |
| WWC3 | -0.02053 | 0.72962 | Other Negative Correlation |
| CCNJ | -0.02053 | 0.729521 | Other Negative Correlation |
| ZNF865 | -0.02057 | 0.729025 | Other Negative Correlation |
| EEPD1 | -0.02062 | 0.728372 | Other Negative Correlation |
| RAB26 | -0.02064 | 0.728121 | Other Negative Correlation |
| ZNF208 | -0.0207 | 0.727468 | Other Negative Correlation |
| DKK2 | -0.02075 | 0.726722 | Other Negative Correlation |
| GPAM | -0.02077 | 0.726561 | Other Negative Correlation |
| LRRC74B | -0.02077 | 0.726502 | Other Negative Correlation |
| TMEM150C | -0.02078 | 0.726392 | Other Negative Correlation |
| RP1L1 | -0.0208 | 0.726166 | Other Negative Correlation |
| GLMP | -0.0208 | 0.726152 | Other Negative Correlation |
| CLRN3 | -0.0208 | 0.726126 | Other Negative Correlation |
| ZNF783 | -0.02081 | 0.726061 | Other Negative Correlation |
| LRRC8D | -0.02081 | 0.72601 | Other Negative Correlation |
| LRP1 | -0.02082 | 0.725878 | Other Negative Correlation |
| SPP1 | -0.02084 | 0.725652 | Other Negative Correlation |
| ISL1 | -0.02087 | 0.725281 | Other Negative Correlation |
| ANKZF1 | -0.02092 | 0.724597 | Other Negative Correlation |
| E2F7 | -0.02097 | 0.723946 | Other Negative Correlation |
| PHF7 | -0.02097 | 0.72394 | Other Negative Correlation |
| GJA3 | -0.02102 | 0.723327 | Other Negative Correlation |
| ZNF717 | -0.02109 | 0.722514 | Other Negative Correlation |
| SCNN1B | -0.02109 | 0.722461 | Other Negative Correlation |
| HACE1 | -0.02118 | 0.721354 | Other Negative Correlation |
| RDM1 | -0.02121 | 0.721002 | Other Negative Correlation |
| PET117 | -0.02124 | 0.72054 | Other Negative Correlation |
| SYT15 | -0.02127 | 0.720194 | Other Negative Correlation |
| CNPPD1 | -0.02129 | 0.719938 | Other Negative Correlation |
| CCDC116 | -0.02131 | 0.719756 | Other Negative Correlation |
| CARMIL3 | -0.02134 | 0.719353 | Other Negative Correlation |
| GPM6B | -0.0214 | 0.718515 | Other Negative Correlation |

| | | | |
|---|---|---|---|
| ZNF586 | -0.02142 | 0.718331 | Other Negative Correlation |
| ZNF577 | -0.02144 | 0.718075 | Other Negative Correlation |
| MYLK4 | -0.02146 | 0.717759 | Other Negative Correlation |
| CHRNA5 | -0.02149 | 0.717471 | Other Negative Correlation |
| PCDHAC1 | -0.02149 | 0.717436 | Other Negative Correlation |
| EGFL7 | -0.02151 | 0.717158 | Other Negative Correlation |
| NTN5 | -0.02153 | 0.716956 | Other Negative Correlation |
| H2BC11 | -0.02153 | 0.71692 | Other Negative Correlation |
| GUCA2B | -0.02153 | 0.716911 | Other Negative Correlation |
| RIMS3 | -0.02155 | 0.716743 | Other Negative Correlation |
| BVES | -0.02157 | 0.716498 | Other Negative Correlation |
| CREBBP | -0.0216 | 0.716099 | Other Negative Correlation |
| MED9 | -0.0216 | 0.71609 | Other Negative Correlation |
| BTNL8 | -0.0216 | 0.716058 | Other Negative Correlation |
| AADACL3 | -0.0216 | 0.716058 | Other Negative Correlation |
| HORMAD1 | -0.0216 | 0.716058 | Other Negative Correlation |
| CPNE9 | -0.0216 | 0.716058 | Other Negative Correlation |
| IGSF5 | -0.02162 | 0.715782 | Other Negative Correlation |
| NDRG2 | -0.02165 | 0.715372 | Other Negative Correlation |
| ATPSCKMT | -0.02166 | 0.715268 | Other Negative Correlation |
| ZFAND4 | -0.0217 | 0.714783 | Other Negative Correlation |
| RNF31 | -0.02172 | 0.714591 | Other Negative Correlation |
| GADD45B | -0.02181 | 0.713462 | Other Negative Correlation |
| DSEL | -0.02184 | 0.713089 | Other Negative Correlation |
| ZNF384 | -0.02188 | 0.712503 | Other Negative Correlation |
| NRXN3 | -0.02198 | 0.711321 | Other Negative Correlation |
| SYT10 | -0.02202 | 0.710839 | Other Negative Correlation |
| SALL1 | -0.02202 | 0.710839 | Other Negative Correlation |
| KHDC4 | -0.02202 | 0.710825 | Other Negative Correlation |
| PXDC1 | -0.02205 | 0.710453 | Other Negative Correlation |
| EMILIN2 | -0.02217 | 0.708944 | Other Negative Correlation |
| MTMR8 | -0.02221 | 0.708414 | Other Negative Correlation |
| SEMA6C | -0.02226 | 0.707716 | Other Negative Correlation |
| FBXO10 | -0.02228 | 0.707568 | Other Negative Correlation |
| TLR7 | -0.02232 | 0.707076 | Other Negative Correlation |
| HMGCS2 | -0.02232 | 0.707076 | Other Negative Correlation |
| STOML3 | -0.02232 | 0.707076 | Other Negative Correlation |
| KRTAP3-3 | -0.02232 | 0.707076 | Other Negative Correlation |
| KRTAP2-4 | -0.02232 | 0.707076 | Other Negative Correlation |
| IRAG2 | -0.02232 | 0.707076 | Other Negative Correlation |
| POU2AF2 | -0.02232 | 0.707076 | Other Negative Correlation |
| ZAP70 | -0.02232 | 0.707076 | Other Negative Correlation |
| CLEC2A | -0.02232 | 0.707076 | Other Negative Correlation |
| HOXA11 | -0.02232 | 0.707076 | Other Negative Correlation |
| EVX1 | -0.02232 | 0.707076 | Other Negative Correlation |
| PKD2L1 | -0.02232 | 0.707076 | Other Negative Correlation |

| | | | |
|---|---|---|---|
| LYZ | -0.02232 | 0.707076 | Other Negative Correlation |
| KERA | -0.02232 | 0.707076 | Other Negative Correlation |
| CASP12 | -0.02232 | 0.707076 | Other Negative Correlation |
| HEMGN | -0.02232 | 0.707076 | Other Negative Correlation |
| KRTAP1-4 | -0.02232 | 0.707076 | Other Negative Correlation |
| TBPL2 | -0.02232 | 0.707076 | Other Negative Correlation |
| KRTAP2-1 | -0.02232 | 0.707076 | Other Negative Correlation |
| SAMD7 | -0.02232 | 0.707076 | Other Negative Correlation |
| CLEC12A | -0.02232 | 0.707076 | Other Negative Correlation |
| KCNH7 | -0.02232 | 0.707076 | Other Negative Correlation |
| GTSF1 | -0.02232 | 0.707076 | Other Negative Correlation |
| CD69 | -0.02232 | 0.707076 | Other Negative Correlation |
| PPIAL4H | -0.02232 | 0.707076 | Other Negative Correlation |
| DIO3 | -0.02232 | 0.707076 | Other Negative Correlation |
| KRT38 | -0.02232 | 0.707076 | Other Negative Correlation |
| SIGLEC6 | -0.02232 | 0.707076 | Other Negative Correlation |
| CLEC12B | -0.02232 | 0.707076 | Other Negative Correlation |
| KRIT1 | -0.02234 | 0.706788 | Other Negative Correlation |
| ERN2 | -0.02236 | 0.706534 | Other Negative Correlation |
| AMHR2 | -0.02238 | 0.706289 | Other Negative Correlation |
| KCNH4 | -0.02238 | 0.706289 | Other Negative Correlation |
| LACTB2 | -0.0224 | 0.705989 | Other Negative Correlation |
| LRWD1 | -0.02241 | 0.705932 | Other Negative Correlation |
| ZSCAN5A | -0.02246 | 0.705209 | Other Negative Correlation |
| STIL | -0.02259 | 0.703688 | Other Negative Correlation |
| CDHR1 | -0.02265 | 0.702891 | Other Negative Correlation |
| SESTD1 | -0.02275 | 0.701656 | Other Negative Correlation |
| DNAAF1 | -0.02285 | 0.700381 | Other Negative Correlation |
| TM4SF18 | -0.02288 | 0.700028 | Other Negative Correlation |
| PLCE1 | -0.02292 | 0.699476 | Other Negative Correlation |
| AXDND1 | -0.02292 | 0.699466 | Other Negative Correlation |
| TEKT4 | -0.02296 | 0.699053 | Other Negative Correlation |
| PROB1 | -0.02296 | 0.699012 | Other Negative Correlation |
| CYP4F3 | -0.02299 | 0.698636 | Other Negative Correlation |
| SSUH2 | -0.02307 | 0.697604 | Other Negative Correlation |
| PRDM9 | -0.02317 | 0.696366 | Other Negative Correlation |
| GIMD1 | -0.02317 | 0.696366 | Other Negative Correlation |
| ADGRG7 | -0.02317 | 0.696366 | Other Negative Correlation |
| ARHGEF38 | -0.02317 | 0.696366 | Other Negative Correlation |
| GRID2 | -0.02317 | 0.696366 | Other Negative Correlation |
| SPANXB1 | -0.02317 | 0.696366 | Other Negative Correlation |
| SPANXC | -0.02317 | 0.696366 | Other Negative Correlation |
| C4orf54 | -0.02317 | 0.696366 | Other Negative Correlation |
| EMCN | -0.02317 | 0.696366 | Other Negative Correlation |
| ABHD16B | -0.02317 | 0.696366 | Other Negative Correlation |
| SPANXA1 | -0.02317 | 0.696366 | Other Negative Correlation |

| | | | |
|---|---|---|---|
| CDH20 | -0.02317 | 0.696366 | Other Negative Correlation |
| AMBN | -0.02317 | 0.696366 | Other Negative Correlation |
| DMP1 | -0.02317 | 0.696366 | Other Negative Correlation |
| ZBTB49 | -0.02318 | 0.696262 | Other Negative Correlation |
| BCL6B | -0.02321 | 0.6959 | Other Negative Correlation |
| FAM172A | -0.02328 | 0.695009 | Other Negative Correlation |
| ZBTB2 | -0.02333 | 0.694474 | Other Negative Correlation |
| TVP23C-CDRT4 | -0.02334 | 0.694297 | Other Negative Correlation |
| GLB1L2 | -0.02337 | 0.693882 | Other Negative Correlation |
| GLIPR2 | -0.02339 | 0.693625 | Other Negative Correlation |
| PRR29 | -0.0234 | 0.693584 | Other Negative Correlation |
| RCBTB1 | -0.02341 | 0.693401 | Other Negative Correlation |
| ADCY9 | -0.02344 | 0.693012 | Other Negative Correlation |
| FOXF2 | -0.02357 | 0.691425 | Other Negative Correlation |
| PRKG2 | -0.02362 | 0.690849 | Other Negative Correlation |
| FGFBP2 | -0.02364 | 0.690505 | Other Negative Correlation |
| CST9L | -0.02364 | 0.690505 | Other Negative Correlation |
| OR7D2 | -0.02364 | 0.690505 | Other Negative Correlation |
| COL20A1 | -0.02364 | 0.690505 | Other Negative Correlation |
| MC5R | -0.02364 | 0.690505 | Other Negative Correlation |
| ATP13A2 | -0.02364 | 0.690501 | Other Negative Correlation |
| SIX1 | -0.02368 | 0.690025 | Other Negative Correlation |
| AOAH | -0.02369 | 0.689956 | Other Negative Correlation |
| LDB3 | -0.0237 | 0.68981 | Other Negative Correlation |
| ADRB1 | -0.02373 | 0.689482 | Other Negative Correlation |
| ZSCAN30 | -0.02378 | 0.688805 | Other Negative Correlation |
| RNASEH1 | -0.02379 | 0.688754 | Other Negative Correlation |
| NPFFR2 | -0.02381 | 0.68846 | Other Negative Correlation |
| RGPD5 | -0.02385 | 0.687983 | Other Negative Correlation |
| ZNF862 | -0.02388 | 0.687558 | Other Negative Correlation |
| TNKS | -0.02393 | 0.686948 | Other Negative Correlation |
| CEP152 | -0.02396 | 0.68657 | Other Negative Correlation |
| DDHD1 | -0.02396 | 0.686561 | Other Negative Correlation |
| SHISA6 | -0.02401 | 0.68601 | Other Negative Correlation |
| SLC6A6 | -0.02405 | 0.685507 | Other Negative Correlation |
| TTC22 | -0.02416 | 0.684063 | Other Negative Correlation |
| SLC48A1 | -0.02418 | 0.68391 | Other Negative Correlation |
| NRL | -0.02428 | 0.682621 | Other Negative Correlation |
| OS9 | -0.02429 | 0.682462 | Other Negative Correlation |
| ZNF571 | -0.02433 | 0.682019 | Other Negative Correlation |
| KALRN | -0.02442 | 0.680955 | Other Negative Correlation |
| SGK3 | -0.02444 | 0.680687 | Other Negative Correlation |
| ADAM30 | -0.02446 | 0.680411 | Other Negative Correlation |
| GIMAP2 | -0.02452 | 0.67962 | Other Negative Correlation |
| COL9A3 | -0.02453 | 0.679541 | Other Negative Correlation |
| C6orf58 | -0.02457 | 0.679007 | Other Negative Correlation |

| | | | |
|---|---|---|---|
| SHISA2 | -0.02475 | 0.676886 | Other Negative Correlation |
| LEP | -0.02481 | 0.676074 | Other Negative Correlation |
| PAK5 | -0.02483 | 0.675831 | Other Negative Correlation |
| PCDHB4 | -0.02496 | 0.674259 | Other Negative Correlation |
| MUCL3 | -0.025 | 0.673732 | Other Negative Correlation |
| ZNF821 | -0.02504 | 0.67331 | Other Negative Correlation |
| NHLRC3 | -0.02504 | 0.673242 | Other Negative Correlation |
| SLX1A | -0.02506 | 0.672957 | Other Negative Correlation |
| FBXO25 | -0.02507 | 0.672854 | Other Negative Correlation |
| MAST2 | -0.02509 | 0.67267 | Other Negative Correlation |
| CDSN | -0.02512 | 0.672307 | Other Negative Correlation |
| EIF5A2 | -0.02513 | 0.6722 | Other Negative Correlation |
| CNPY4 | -0.02513 | 0.672145 | Other Negative Correlation |
| ZNF443 | -0.02515 | 0.671949 | Other Negative Correlation |
| TSPAN19 | -0.02516 | 0.671804 | Other Negative Correlation |
| GTDC1 | -0.02517 | 0.671667 | Other Negative Correlation |
| OSR1 | -0.02517 | 0.671631 | Other Negative Correlation |
| SH2B1 | -0.02519 | 0.671382 | Other Negative Correlation |
| SMC3 | -0.0252 | 0.671264 | Other Negative Correlation |
| EXOC6 | -0.02521 | 0.671145 | Other Negative Correlation |
| SPACDR | -0.02521 | 0.671123 | Other Negative Correlation |
| ABCA2 | -0.02525 | 0.670645 | Other Negative Correlation |
| CCDC190 | -0.02528 | 0.670356 | Other Negative Correlation |
| SSMEM1 | -0.02528 | 0.670356 | Other Negative Correlation |
| SEPTIN12 | -0.02528 | 0.670356 | Other Negative Correlation |
| MAP3K10 | -0.02529 | 0.670152 | Other Negative Correlation |
| TBC1D30 | -0.02535 | 0.669504 | Other Negative Correlation |
| ATP6V1E1 | -0.02536 | 0.669327 | Other Negative Correlation |
| COL1A1 | -0.02536 | 0.669296 | Other Negative Correlation |
| NANOS1 | -0.02536 | 0.66928 | Other Negative Correlation |
| KCNA2 | -0.0254 | 0.668866 | Other Negative Correlation |
| IL10RA | -0.02544 | 0.668374 | Other Negative Correlation |
| NKIRAS1 | -0.02544 | 0.668311 | Other Negative Correlation |
| PAX9 | -0.02545 | 0.668214 | Other Negative Correlation |
| PADI2 | -0.02547 | 0.668037 | Other Negative Correlation |
| FUT6 | -0.02549 | 0.667754 | Other Negative Correlation |
| ABCA9 | -0.02553 | 0.667305 | Other Negative Correlation |
| DPEP2NB | -0.02553 | 0.667205 | Other Negative Correlation |
| AZIN2 | -0.02554 | 0.667103 | Other Negative Correlation |
| GRK5 | -0.02554 | 0.667077 | Other Negative Correlation |
| MAS1 | -0.02556 | 0.666872 | Other Negative Correlation |
| MYT1 | -0.02557 | 0.666782 | Other Negative Correlation |
| MCF2L2 | -0.02558 | 0.666689 | Other Negative Correlation |
| SIAH2 | -0.02559 | 0.666537 | Other Negative Correlation |
| LRRN3 | -0.02567 | 0.665534 | Other Negative Correlation |
| GOLGA6L1 | -0.02572 | 0.664891 | Other Negative Correlation |

| | | | |
|---|---|---|---|
| NR1D1 | -0.02574 | 0.664678 | Other Negative Correlation |
| ZNF345 | -0.02575 | 0.664512 | Other Negative Correlation |
| KANSL1 | -0.02576 | 0.664445 | Other Negative Correlation |
| SLC25A29 | -0.02578 | 0.664221 | Other Negative Correlation |
| TMEM151B | -0.02579 | 0.664084 | Other Negative Correlation |
| TSGA10 | -0.02581 | 0.663764 | Other Negative Correlation |
| BRINP2 | -0.02582 | 0.663662 | Other Negative Correlation |
| POLD3 | -0.02585 | 0.66327 | Other Negative Correlation |
| WDR97 | -0.02589 | 0.66278 | Other Negative Correlation |
| ENGASE | -0.02592 | 0.662478 | Other Negative Correlation |
| TIMM23B-AGAP6 | -0.02594 | 0.6622 | Other Negative Correlation |
| MYO16 | -0.026 | 0.661553 | Other Negative Correlation |
| HAVCR2 | -0.02603 | 0.661166 | Other Negative Correlation |
| TRPM3 | -0.02603 | 0.661078 | Other Negative Correlation |
| FGFBP3 | -0.02607 | 0.660637 | Other Negative Correlation |
| ASCL1 | -0.02608 | 0.660529 | Other Negative Correlation |
| TMLHE | -0.02609 | 0.660413 | Other Negative Correlation |
| TNFAIP8L2 | -0.02617 | 0.659364 | Other Negative Correlation |
| CA6 | -0.02617 | 0.659364 | Other Negative Correlation |
| OR2A5 | -0.02617 | 0.659364 | Other Negative Correlation |
| OARD1 | -0.02618 | 0.659289 | Other Negative Correlation |
| CNTLN | -0.02627 | 0.658234 | Other Negative Correlation |
| SHISAL2A | -0.0263 | 0.657819 | Other Negative Correlation |
| C8G | -0.02631 | 0.657775 | Other Negative Correlation |
| CHST10 | -0.02634 | 0.657355 | Other Negative Correlation |
| NDEL1 | -0.02637 | 0.657001 | Other Negative Correlation |
| PIH1D2 | -0.02639 | 0.656681 | Other Negative Correlation |
| LMNTD2 | -0.02644 | 0.656151 | Other Negative Correlation |
| RNFT1 | -0.02648 | 0.655602 | Other Negative Correlation |
| C22orf39 | -0.02651 | 0.655246 | Other Negative Correlation |
| SYT1 | -0.02652 | 0.655101 | Other Negative Correlation |
| ENSG00000283782 | -0.02659 | 0.65432 | Other Negative Correlation |
| PRR22 | -0.02661 | 0.654066 | Other Negative Correlation |
| FSCN2 | -0.02661 | 0.654055 | Other Negative Correlation |
| PLGLB1 | -0.02662 | 0.653989 | Other Negative Correlation |
| SHOX | -0.02662 | 0.653943 | Other Negative Correlation |
| ZNF471 | -0.02663 | 0.653793 | Other Negative Correlation |
| FREM1 | -0.02664 | 0.65364 | Other Negative Correlation |
| CCDC201 | -0.02664 | 0.65364 | Other Negative Correlation |
| ENSG00000284695 | -0.02664 | 0.65364 | Other Negative Correlation |
| CALN1 | -0.02664 | 0.65364 | Other Negative Correlation |
| RIMBP3 | -0.02664 | 0.65364 | Other Negative Correlation |
| TOPBP1 | -0.02668 | 0.653172 | Other Negative Correlation |
| ULK4 | -0.02677 | 0.652137 | Other Negative Correlation |
| AGFG2 | -0.02683 | 0.651381 | Other Negative Correlation |
| AQP10 | -0.0269 | 0.650554 | Other Negative Correlation |

| | | | |
|---|---|---|---|
| SRGAP2B | -0.02697 | 0.649653 | Other Negative Correlation |
| USP32 | -0.02704 | 0.648887 | Other Negative Correlation |
| PARP6 | -0.02705 | 0.648707 | Other Negative Correlation |
| TPST1 | -0.0271 | 0.648164 | Other Negative Correlation |
| TRMT11 | -0.02711 | 0.647958 | Other Negative Correlation |
| PARS2 | -0.02722 | 0.64662 | Other Negative Correlation |
| DGKI | -0.02724 | 0.646477 | Other Negative Correlation |
| CCL24 | -0.02724 | 0.64642 | Other Negative Correlation |
| ZNF585A | -0.02726 | 0.646189 | Other Negative Correlation |
| CCT6B | -0.02727 | 0.646116 | Other Negative Correlation |
| HMCN2 | -0.02728 | 0.645911 | Other Negative Correlation |
| DNAH14 | -0.02739 | 0.644633 | Other Negative Correlation |
| TBCE.1 | -0.02744 | 0.643985 | Other Negative Correlation |
| SLC26A7 | -0.02749 | 0.6434 | Other Negative Correlation |
| TMPRSS6 | -0.0275 | 0.643236 | Other Negative Correlation |
| ADAMTS3 | -0.02757 | 0.642441 | Other Negative Correlation |
| ZNF488 | -0.02761 | 0.641975 | Other Negative Correlation |
| MISP3 | -0.02763 | 0.64177 | Other Negative Correlation |
| NEURL3 | -0.02764 | 0.641572 | Other Negative Correlation |
| ENSG00000283599 | -0.02766 | 0.641403 | Other Negative Correlation |
| ATCAY | -0.02766 | 0.641344 | Other Negative Correlation |
| TTYH2 | -0.02769 | 0.641036 | Other Negative Correlation |
| CACNG2 | -0.02771 | 0.640791 | Other Negative Correlation |
| TSPAN32 | -0.02771 | 0.640791 | Other Negative Correlation |
| SMIM18 | -0.02771 | 0.640791 | Other Negative Correlation |
| CLDN17 | -0.02771 | 0.640791 | Other Negative Correlation |
| GPR17 | -0.02771 | 0.640791 | Other Negative Correlation |
| ENSG00000249319 | -0.02771 | 0.640791 | Other Negative Correlation |
| SPDYC | -0.02771 | 0.640791 | Other Negative Correlation |
| VIL1 | -0.02771 | 0.640745 | Other Negative Correlation |
| PLAGL1 | -0.02777 | 0.640032 | Other Negative Correlation |
| CCDC178 | -0.02778 | 0.639872 | Other Negative Correlation |
| GPR62 | -0.0278 | 0.639613 | Other Negative Correlation |
| RLN2 | -0.02786 | 0.638932 | Other Negative Correlation |
| ATP9A | -0.02787 | 0.638796 | Other Negative Correlation |
| ETNK2 | -0.02789 | 0.638582 | Other Negative Correlation |
| TNRC6A | -0.02791 | 0.638296 | Other Negative Correlation |
| CNKSR2 | -0.02793 | 0.638104 | Other Negative Correlation |
| ZKSCAN5 | -0.02795 | 0.637904 | Other Negative Correlation |
| RBM15 | -0.02797 | 0.637664 | Other Negative Correlation |
| LUM | -0.02801 | 0.637154 | Other Negative Correlation |
| CPNE4 | -0.02804 | 0.636829 | Other Negative Correlation |
| GTPBP1 | -0.02812 | 0.635858 | Other Negative Correlation |
| FAXC | -0.02815 | 0.635442 | Other Negative Correlation |
| TMPRSS12 | -0.02822 | 0.634632 | Other Negative Correlation |
| LPAR1 | -0.02822 | 0.634602 | Other Negative Correlation |

| | | | |
|---|---|---|---|
| MAP3K5 | -0.02825 | 0.634264 | Other Negative Correlation |
| STARD8 | -0.02827 | 0.63407 | Other Negative Correlation |
| MFRP | -0.02827 | 0.633988 | Other Negative Correlation |
| SPACA4 | -0.02827 | 0.633988 | Other Negative Correlation |
| VKORC1L1 | -0.02828 | 0.633924 | Other Negative Correlation |
| SCIN | -0.02829 | 0.633795 | Other Negative Correlation |
| ACKR3 | -0.0283 | 0.633698 | Other Negative Correlation |
| RFX4 | -0.0283 | 0.633633 | Other Negative Correlation |
| DYNLT2 | -0.02832 | 0.633393 | Other Negative Correlation |
| BAIAP2L2 | -0.02832 | 0.633368 | Other Negative Correlation |
| TFE3 | -0.02834 | 0.633198 | Other Negative Correlation |
| NTS | -0.02835 | 0.633072 | Other Negative Correlation |
| CACNA1E | -0.02835 | 0.633027 | Other Negative Correlation |
| GRID2IP | -0.02837 | 0.632851 | Other Negative Correlation |
| GORAB | -0.02837 | 0.63278 | Other Negative Correlation |
| RNF207 | -0.0285 | 0.631302 | Other Negative Correlation |
| GIPC2 | -0.02857 | 0.630458 | Other Negative Correlation |
| CHGB | -0.02858 | 0.630346 | Other Negative Correlation |
| ZHX2 | -0.02862 | 0.629856 | Other Negative Correlation |
| RGPD8 | -0.02864 | 0.629572 | Other Negative Correlation |
| EGR3 | -0.02865 | 0.629438 | Other Negative Correlation |
| CCNI2 | -0.02865 | 0.629425 | Other Negative Correlation |
| TRAM2 | -0.0287 | 0.628824 | Other Negative Correlation |
| COL8A2 | -0.02873 | 0.628532 | Other Negative Correlation |
| ENSG00000285589 | -0.02876 | 0.628097 | Other Negative Correlation |
| DIP2C | -0.02878 | 0.627918 | Other Negative Correlation |
| GARIN1A | -0.02878 | 0.627855 | Other Negative Correlation |
| PMEL | -0.02884 | 0.62721 | Other Negative Correlation |
| FGD1 | -0.02884 | 0.627156 | Other Negative Correlation |
| ZZEF1 | -0.02888 | 0.626671 | Other Negative Correlation |
| PGS1 | -0.02896 | 0.625783 | Other Negative Correlation |
| SPINK1 | -0.02898 | 0.62547 | Other Negative Correlation |
| HOXB8 | -0.02898 | 0.62547 | Other Negative Correlation |
| IZUMO1R | -0.02898 | 0.62547 | Other Negative Correlation |
| HBM | -0.02898 | 0.62547 | Other Negative Correlation |
| TPTE2 | -0.02898 | 0.62547 | Other Negative Correlation |
| IRF8 | -0.02898 | 0.62547 | Other Negative Correlation |
| PDZD9 | -0.02898 | 0.62547 | Other Negative Correlation |
| TSPAN8 | -0.02898 | 0.62547 | Other Negative Correlation |
| CST4 | -0.02898 | 0.62547 | Other Negative Correlation |
| TFAP2D | -0.02898 | 0.62547 | Other Negative Correlation |
| PIWIL3 | -0.02898 | 0.62547 | Other Negative Correlation |
| PAEP | -0.02898 | 0.62547 | Other Negative Correlation |
| PREX2 | -0.02898 | 0.62547 | Other Negative Correlation |
| MS4A6A | -0.02898 | 0.62547 | Other Negative Correlation |
| BARHL1 | -0.02898 | 0.62547 | Other Negative Correlation |

| | | | |
|---|---|---|---|
| ALPG | -0.02898 | 0.62547 | Other Negative Correlation |
| OR5B2 | -0.02898 | 0.62547 | Other Negative Correlation |
| OR5B12 | -0.02898 | 0.62547 | Other Negative Correlation |
| WEE1 | -0.02908 | 0.624321 | Other Negative Correlation |
| SSX5 | -0.0291 | 0.624139 | Other Negative Correlation |
| NLRP8 | -0.0291 | 0.624139 | Other Negative Correlation |
| ERCC3 | -0.02911 | 0.623907 | Other Negative Correlation |
| FILIP1 | -0.02914 | 0.623594 | Other Negative Correlation |
| CREBRF | -0.02916 | 0.623416 | Other Negative Correlation |
| MEFV | -0.0292 | 0.622891 | Other Negative Correlation |
| ANXA13 | -0.0292 | 0.622891 | Other Negative Correlation |
| PRF1 | -0.0292 | 0.622891 | Other Negative Correlation |
| GPRC5D | -0.0292 | 0.622891 | Other Negative Correlation |
| ZDHHC19 | -0.0292 | 0.622891 | Other Negative Correlation |
| GPR55 | -0.0292 | 0.622891 | Other Negative Correlation |
| C19orf67 | -0.0292 | 0.622891 | Other Negative Correlation |
| KLK14 | -0.0292 | 0.622891 | Other Negative Correlation |
| MYOZ2 | -0.0292 | 0.622891 | Other Negative Correlation |
| LIM2 | -0.0292 | 0.622891 | Other Negative Correlation |
| SIGLEC9 | -0.0292 | 0.622891 | Other Negative Correlation |
| GARIN5B | -0.0292 | 0.622891 | Other Negative Correlation |
| KRTAP5-10 | -0.0292 | 0.622891 | Other Negative Correlation |
| TYROBP | -0.0292 | 0.622891 | Other Negative Correlation |
| OR1L8 | -0.0292 | 0.622891 | Other Negative Correlation |
| PLP1 | -0.0292 | 0.622891 | Other Negative Correlation |
| TRPM5 | -0.0292 | 0.622891 | Other Negative Correlation |
| TIE1 | -0.0292 | 0.622891 | Other Negative Correlation |
| FAM180A | -0.0292 | 0.622891 | Other Negative Correlation |
| TRIML1 | -0.0292 | 0.622891 | Other Negative Correlation |
| POC1B-GALNT4 | -0.0292 | 0.622891 | Other Negative Correlation |
| MOGAT2 | -0.0292 | 0.622891 | Other Negative Correlation |
| RNASE1 | -0.0292 | 0.622891 | Other Negative Correlation |
| EPS8L3 | -0.0292 | 0.622891 | Other Negative Correlation |
| LILRA6 | -0.0292 | 0.622891 | Other Negative Correlation |
| MUC21 | -0.0292 | 0.622891 | Other Negative Correlation |
| CAPN9 | -0.02925 | 0.622332 | Other Negative Correlation |
| HSDL1 | -0.02932 | 0.621496 | Other Negative Correlation |
| SOD3 | -0.02932 | 0.621436 | Other Negative Correlation |
| CBX8 | -0.02934 | 0.621258 | Other Negative Correlation |
| ADGRG3 | -0.02936 | 0.62102 | Other Negative Correlation |
| PRDM11 | -0.02936 | 0.62101 | Other Negative Correlation |
| SLC35E4 | -0.02943 | 0.620207 | Other Negative Correlation |
| ZNF593OS | -0.02945 | 0.619923 | Other Negative Correlation |
| GPR141 | -0.02946 | 0.619755 | Other Negative Correlation |
| FBXL14 | -0.02949 | 0.619412 | Other Negative Correlation |
| JMY | -0.02949 | 0.619381 | Other Negative Correlation |

| | | | |
|---|---|---|---|
| XKR5 | -0.02952 | 0.619137 | Other Negative Correlation |
| STX1A | -0.02952 | 0.619056 | Other Negative Correlation |
| FOXD4L6 | -0.02955 | 0.618757 | Other Negative Correlation |
| TRIM11 | -0.02955 | 0.618673 | Other Negative Correlation |
| ACSL5 | -0.02963 | 0.617802 | Other Negative Correlation |
| IGFBP5 | -0.02963 | 0.617747 | Other Negative Correlation |
| RYR1 | -0.02964 | 0.617635 | Other Negative Correlation |
| TEX29 | -0.02966 | 0.617381 | Other Negative Correlation |
| LGALS2 | -0.02967 | 0.617354 | Other Negative Correlation |
| PNPLA2 | -0.02967 | 0.617282 | Other Negative Correlation |
| SLC25A10 | -0.02975 | 0.616354 | Other Negative Correlation |
| GRAP | -0.02979 | 0.615865 | Other Negative Correlation |
| CCR7 | -0.0298 | 0.615705 | Other Negative Correlation |
| MACROD1 | -0.02982 | 0.61553 | Other Negative Correlation |
| ADA2 | -0.02984 | 0.615302 | Other Negative Correlation |
| DLST | -0.02986 | 0.615041 | Other Negative Correlation |
| BCAS2 | -0.02991 | 0.614401 | Other Negative Correlation |
| SCG2 | -0.02993 | 0.614234 | Other Negative Correlation |
| CD93 | -0.02993 | 0.614233 | Other Negative Correlation |
| CD177 | -0.03001 | 0.613291 | Other Negative Correlation |
| PDIK1L | -0.03002 | 0.613172 | Other Negative Correlation |
| PAGE2B | -0.03005 | 0.612804 | Other Negative Correlation |
| CMTM8 | -0.03015 | 0.611568 | Other Negative Correlation |
| SMAD9 | -0.03018 | 0.611277 | Other Negative Correlation |
| GOLT1B | -0.03024 | 0.610496 | Other Negative Correlation |
| GRIN3B | -0.03028 | 0.61007 | Other Negative Correlation |
| ZNF318 | -0.03032 | 0.609561 | Other Negative Correlation |
| H3Y1 | -0.03034 | 0.609412 | Other Negative Correlation |
| ZNF585B | -0.03034 | 0.609375 | Other Negative Correlation |
| BMP8A | -0.03038 | 0.608864 | Other Negative Correlation |
| POP7 | -0.03042 | 0.608443 | Other Negative Correlation |
| COL24A1 | -0.03043 | 0.608265 | Other Negative Correlation |
| ZNF888 | -0.03046 | 0.607909 | Other Negative Correlation |
| PDE7B | -0.03053 | 0.607138 | Other Negative Correlation |
| OTOP3 | -0.03062 | 0.606076 | Other Negative Correlation |
| SHBG | -0.03066 | 0.605593 | Other Negative Correlation |
| PCDHB14 | -0.03067 | 0.605532 | Other Negative Correlation |
| CYP2C18 | -0.03067 | 0.605432 | Other Negative Correlation |
| COL14A1 | -0.03073 | 0.604819 | Other Negative Correlation |
| DNAH11 | -0.03073 | 0.604749 | Other Negative Correlation |
| GLIS3 | -0.03073 | 0.604746 | Other Negative Correlation |
| ECT2 | -0.03081 | 0.603783 | Other Negative Correlation |
| AGAP2 | -0.03085 | 0.60342 | Other Negative Correlation |
| TMEM35A | -0.03088 | 0.603066 | Other Negative Correlation |
| CDH12 | -0.03088 | 0.603028 | Other Negative Correlation |
| C9 | -0.03088 | 0.603028 | Other Negative Correlation |

| | | | |
|---|---|---|---|
| REN | -0.0309 | 0.60279 | Other Negative Correlation |
| RAB33B | -0.03096 | 0.602035 | Other Negative Correlation |
| SNX24 | -0.03098 | 0.601819 | Other Negative Correlation |
| LHX6 | -0.03099 | 0.601741 | Other Negative Correlation |
| ALDH2 | -0.03102 | 0.601399 | Other Negative Correlation |
| FAM72A | -0.03103 | 0.601222 | Other Negative Correlation |
| FAM102A | -0.03103 | 0.601218 | Other Negative Correlation |
| MYRF | -0.03106 | 0.600948 | Other Negative Correlation |
| SPATA3 | -0.03107 | 0.600791 | Other Negative Correlation |
| PNLDC1 | -0.03107 | 0.600791 | Other Negative Correlation |
| ICA1L | -0.03109 | 0.600591 | Other Negative Correlation |
| PAQR6 | -0.0311 | 0.600416 | Other Negative Correlation |
| VWA3B | -0.03114 | 0.600018 | Other Negative Correlation |
| LMNTD1 | -0.03119 | 0.599325 | Other Negative Correlation |
| VAPB | -0.03123 | 0.598925 | Other Negative Correlation |
| AK9 | -0.03125 | 0.598712 | Other Negative Correlation |
| THSD7B | -0.03137 | 0.597324 | Other Negative Correlation |
| NDP | -0.03137 | 0.59727 | Other Negative Correlation |
| DNAI2 | -0.03137 | 0.59727 | Other Negative Correlation |
| SSTR4 | -0.03137 | 0.59727 | Other Negative Correlation |
| ENSG00000267618 | -0.03137 | 0.59727 | Other Negative Correlation |
| ENSG00000267314 | -0.03137 | 0.59727 | Other Negative Correlation |
| DMRTA2 | -0.0314 | 0.5969 | Other Negative Correlation |
| ZNF747 | -0.03142 | 0.596675 | Other Negative Correlation |
| TAF11L11 | -0.03151 | 0.595684 | Other Negative Correlation |
| ZNF676 | -0.03152 | 0.595555 | Other Negative Correlation |
| FZD7 | -0.03154 | 0.595322 | Other Negative Correlation |
| E2F8 | -0.03155 | 0.595209 | Other Negative Correlation |
| PAIP2 | -0.03155 | 0.595172 | Other Negative Correlation |
| ZNF621 | -0.03158 | 0.594843 | Other Negative Correlation |
| RGS5 | -0.0316 | 0.594635 | Other Negative Correlation |
| ATP12A | -0.03163 | 0.594222 | Other Negative Correlation |
| GPANK1 | -0.03167 | 0.593731 | Other Negative Correlation |
| CDYL2 | -0.03168 | 0.593623 | Other Negative Correlation |
| CENPS | -0.03174 | 0.592943 | Other Negative Correlation |
| PPBP | -0.03176 | 0.592699 | Other Negative Correlation |
| FCMR | -0.03177 | 0.592626 | Other Negative Correlation |
| SFTPD | -0.03177 | 0.592616 | Other Negative Correlation |
| ZNF83 | -0.0318 | 0.592274 | Other Negative Correlation |
| HELB | -0.03183 | 0.591922 | Other Negative Correlation |
| MIA | -0.03184 | 0.59182 | Other Negative Correlation |
| OR2B6 | -0.03187 | 0.591437 | Other Negative Correlation |
| ZNF708 | -0.0319 | 0.591135 | Other Negative Correlation |
| SERINC4 | -0.0319 | 0.591125 | Other Negative Correlation |
| RETREG2 | -0.03192 | 0.590898 | Other Negative Correlation |
| LYL1 | -0.03194 | 0.590676 | Other Negative Correlation |

| | | | |
|---|---|---|---|
| TEX30 | -0.03196 | 0.590373 | Other Negative Correlation |
| FHAD1 | -0.03202 | 0.589722 | Other Negative Correlation |
| ZNF114 | -0.03204 | 0.589485 | Other Negative Correlation |
| RASD2 | -0.03205 | 0.589298 | Other Negative Correlation |
| APOM | -0.03206 | 0.589244 | Other Negative Correlation |
| BRF2 | -0.03208 | 0.589033 | Other Negative Correlation |
| FAM161A | -0.03209 | 0.588877 | Other Negative Correlation |
| OGFOD2 | -0.03213 | 0.588424 | Other Negative Correlation |
| NKX1-2 | -0.03213 | 0.588377 | Other Negative Correlation |
| BSN | -0.03214 | 0.58829 | Other Negative Correlation |
| DIRAS3 | -0.0322 | 0.587575 | Other Negative Correlation |
| P4HA3 | -0.03223 | 0.587238 | Other Negative Correlation |
| TRPC4 | -0.03224 | 0.587122 | Other Negative Correlation |
| LRRC45 | -0.03226 | 0.586926 | Other Negative Correlation |
| ENSG00000241962 | -0.03231 | 0.586378 | Other Negative Correlation |
| ANKRD40 | -0.03231 | 0.586373 | Other Negative Correlation |
| ADCK1 | -0.03231 | 0.586282 | Other Negative Correlation |
| SENP8 | -0.03232 | 0.586241 | Other Negative Correlation |
| TMEM196 | -0.03233 | 0.586125 | Other Negative Correlation |
| BIK | -0.03238 | 0.585546 | Other Negative Correlation |
| LRRC24 | -0.0324 | 0.585238 | Other Negative Correlation |
| ATP6V0A2 | -0.03241 | 0.585197 | Other Negative Correlation |
| FOXJ2 | -0.03242 | 0.585079 | Other Negative Correlation |
| STX16 | -0.03243 | 0.584965 | Other Negative Correlation |
| FRAT1 | -0.03247 | 0.5845 | Other Negative Correlation |
| ODAD3 | -0.03247 | 0.584444 | Other Negative Correlation |
| BUD23 | -0.03249 | 0.584222 | Other Negative Correlation |
| TSPAN16 | -0.03257 | 0.583329 | Other Negative Correlation |
| CD33 | -0.03257 | 0.583329 | Other Negative Correlation |
| CXCL13 | -0.03257 | 0.583329 | Other Negative Correlation |
| DPYS | -0.03257 | 0.583329 | Other Negative Correlation |
| ZYG11A | -0.03258 | 0.583252 | Other Negative Correlation |
| GPS1 | -0.03259 | 0.583122 | Other Negative Correlation |
| HKDC1 | -0.03268 | 0.582049 | Other Negative Correlation |
| ATXN7L2 | -0.03269 | 0.581897 | Other Negative Correlation |
| NOG | -0.03271 | 0.581679 | Other Negative Correlation |
| IL21R | -0.03279 | 0.580793 | Other Negative Correlation |
| KIF26B | -0.03282 | 0.580487 | Other Negative Correlation |
| PSORS1C2 | -0.03282 | 0.580392 | Other Negative Correlation |
| COQ10A | -0.03286 | 0.580014 | Other Negative Correlation |
| TBCCD1 | -0.03293 | 0.57911 | Other Negative Correlation |
| ZBTB26 | -0.03296 | 0.578811 | Other Negative Correlation |
| SLC25A25 | -0.03296 | 0.578793 | Other Negative Correlation |
| BRPF3 | -0.03297 | 0.578685 | Other Negative Correlation |
| DLL3 | -0.03298 | 0.578545 | Other Negative Correlation |
| TMEM161A | -0.03299 | 0.578452 | Other Negative Correlation |

| | | | |
|---|---|---|---|
| HLA-G | -0.03301 | 0.578262 | Other Negative Correlation |
| MFSD8 | -0.03302 | 0.578162 | Other Negative Correlation |
| TMEM178B | -0.03303 | 0.57799 | Other Negative Correlation |
| SIRT6 | -0.03305 | 0.577758 | Other Negative Correlation |
| CNOT6L | -0.03309 | 0.577318 | Other Negative Correlation |
| ANKRD2 | -0.0331 | 0.577178 | Other Negative Correlation |
| RAB28 | -0.03317 | 0.576373 | Other Negative Correlation |
| GFOD1 | -0.03319 | 0.576163 | Other Negative Correlation |
| LHFPL7 | -0.03321 | 0.575951 | Other Negative Correlation |
| PIEZO1 | -0.03323 | 0.575749 | Other Negative Correlation |
| MYO7B | -0.03325 | 0.575491 | Other Negative Correlation |
| ZNF624 | -0.03327 | 0.575246 | Other Negative Correlation |
| ELL3 | -0.0333 | 0.574961 | Other Negative Correlation |
| ACR | -0.03334 | 0.574416 | Other Negative Correlation |
| ZNF880 | -0.03335 | 0.574279 | Other Negative Correlation |
| TMEM236 | -0.03336 | 0.574231 | Other Negative Correlation |
| CCN6 | -0.03342 | 0.573517 | Other Negative Correlation |
| LY75 | -0.03346 | 0.573026 | Other Negative Correlation |
| PPP1R3F | -0.03349 | 0.572741 | Other Negative Correlation |
| CMTM5 | -0.0335 | 0.572654 | Other Negative Correlation |
| PRELP | -0.0335 | 0.572615 | Other Negative Correlation |
| ACTRT3 | -0.03352 | 0.572334 | Other Negative Correlation |
| TSSK4 | -0.03356 | 0.571885 | Other Negative Correlation |
| SOHLH2 | -0.0336 | 0.571455 | Other Negative Correlation |
| SHISA7 | -0.03361 | 0.571302 | Other Negative Correlation |
| TMED6 | -0.03365 | 0.570841 | Other Negative Correlation |
| PAK3 | -0.03371 | 0.570236 | Other Negative Correlation |
| GPR3 | -0.03371 | 0.570191 | Other Negative Correlation |
| PCGF5 | -0.03371 | 0.570189 | Other Negative Correlation |
| DAGLA | -0.03375 | 0.569706 | Other Negative Correlation |
| ATP13A4 | -0.03378 | 0.569448 | Other Negative Correlation |
| CA10 | -0.03378 | 0.569433 | Other Negative Correlation |
| PATL1 | -0.03381 | 0.569088 | Other Negative Correlation |
| FXYD2 | -0.03381 | 0.569082 | Other Negative Correlation |
| INTS9 | -0.03384 | 0.568692 | Other Negative Correlation |
| RNF212B | -0.03385 | 0.568624 | Other Negative Correlation |
| ALOX12B | -0.03385 | 0.568553 | Other Negative Correlation |
| CCRL2 | -0.03388 | 0.56824 | Other Negative Correlation |
| DPEP2 | -0.03388 | 0.56824 | Other Negative Correlation |
| APOL5 | -0.03388 | 0.56824 | Other Negative Correlation |
| CNR2 | -0.03388 | 0.56824 | Other Negative Correlation |
| EPPIN | -0.03388 | 0.56824 | Other Negative Correlation |
| ATF7-NPFF | -0.03388 | 0.56824 | Other Negative Correlation |
| NLRP12 | -0.03388 | 0.56824 | Other Negative Correlation |
| GPSM1 | -0.03391 | 0.567966 | Other Negative Correlation |
| PABIR2 | -0.03404 | 0.566436 | Other Negative Correlation |

| | | | |
|---|---|---|---|
| NREP | -0.03413 | 0.565403 | Other Negative Correlation |
| DNMT3B | -0.03419 | 0.564717 | Other Negative Correlation |
| AWAT2 | -0.0342 | 0.564574 | Other Negative Correlation |
| SLC22A1 | -0.03425 | 0.564009 | Other Negative Correlation |
| NUMBL | -0.03427 | 0.563802 | Other Negative Correlation |
| PSMG3 | -0.03433 | 0.563168 | Other Negative Correlation |
| DNAJC14 | -0.03433 | 0.563132 | Other Negative Correlation |
| KANSL2 | -0.03433 | 0.563103 | Other Negative Correlation |
| UNCX | -0.03438 | 0.562584 | Other Negative Correlation |
| KIAA0232 | -0.03442 | 0.56214 | Other Negative Correlation |
| COG1 | -0.03446 | 0.561656 | Other Negative Correlation |
| SCARA5 | -0.03452 | 0.560937 | Other Negative Correlation |
| EVPLL | -0.03455 | 0.560635 | Other Negative Correlation |
| ADGRG5 | -0.03455 | 0.560604 | Other Negative Correlation |
| KICS2 | -0.03456 | 0.560491 | Other Negative Correlation |
| IHO1 | -0.0346 | 0.560081 | Other Negative Correlation |
| DVL3 | -0.03465 | 0.55944 | Other Negative Correlation |
| CCDC88B | -0.03467 | 0.559313 | Other Negative Correlation |
| TPO | -0.03468 | 0.559182 | Other Negative Correlation |
| FRMPD1 | -0.0347 | 0.55892 | Other Negative Correlation |
| ZBTB9 | -0.03472 | 0.5587 | Other Negative Correlation |
| EGFLAM | -0.03475 | 0.558309 | Other Negative Correlation |
| TNFRSF4 | -0.03481 | 0.557675 | Other Negative Correlation |
| POLR3D | -0.03492 | 0.55641 | Other Negative Correlation |
| KCNE3 | -0.03494 | 0.556174 | Other Negative Correlation |
| BCO1 | -0.03495 | 0.556091 | Other Negative Correlation |
| MYO7A | -0.03498 | 0.555745 | Other Negative Correlation |
| ENSG00000285082 | -0.03502 | 0.555305 | Other Negative Correlation |
| BPIFA2 | -0.03502 | 0.555305 | Other Negative Correlation |
| ZFHX4 | -0.03503 | 0.555186 | Other Negative Correlation |
| PLSCR2 | -0.03504 | 0.555105 | Other Negative Correlation |
| TMEM238L | -0.03504 | 0.555057 | Other Negative Correlation |
| CPA6 | -0.03505 | 0.55495 | Other Negative Correlation |
| CALB2 | -0.03517 | 0.553641 | Other Negative Correlation |
| SPRN | -0.03518 | 0.553524 | Other Negative Correlation |
| FAM181A | -0.03519 | 0.553418 | Other Negative Correlation |
| ICAM4 | -0.03519 | 0.553351 | Other Negative Correlation |
| KIAA0408 | -0.03521 | 0.553176 | Other Negative Correlation |
| ENSG00000285542 | -0.03521 | 0.553176 | Other Negative Correlation |
| ENSG00000269403 | -0.03521 | 0.553176 | Other Negative Correlation |
| POLR3H | -0.03524 | 0.552858 | Other Negative Correlation |
| NOS3 | -0.03528 | 0.552319 | Other Negative Correlation |
| CPA2 | -0.03529 | 0.55229 | Other Negative Correlation |
| CSGALNACT1 | -0.0353 | 0.552182 | Other Negative Correlation |
| IP6K3 | -0.03531 | 0.55206 | Other Negative Correlation |
| PHLPP1 | -0.03532 | 0.551898 | Other Negative Correlation |

| | | | |
|---|---|---|---|
| DMXL2 | -0.03535 | 0.551548 | Other Negative Correlation |
| KASH5 | -0.03541 | 0.550908 | Other Negative Correlation |
| KIAA2012 | -0.03543 | 0.550647 | Other Negative Correlation |
| EGLN3 | -0.03544 | 0.550575 | Other Negative Correlation |
| ZFP2 | -0.03547 | 0.550258 | Other Negative Correlation |
| ARMH3 | -0.03547 | 0.550177 | Other Negative Correlation |
| GFRA3 | -0.03555 | 0.549358 | Other Negative Correlation |
| SELENOP | -0.03558 | 0.548996 | Other Negative Correlation |
| PGPEP1 | -0.0356 | 0.54882 | Other Negative Correlation |
| STX11 | -0.0356 | 0.548805 | Other Negative Correlation |
| PTGFR | -0.0356 | 0.548744 | Other Negative Correlation |
| UBXN10 | -0.03564 | 0.548374 | Other Negative Correlation |
| SLC25A11 | -0.03566 | 0.54815 | Other Negative Correlation |
| DYNLRB2 | -0.03566 | 0.548131 | Other Negative Correlation |
| ACAD10 | -0.03569 | 0.547786 | Other Negative Correlation |
| GTF2IRD2 | -0.03571 | 0.547584 | Other Negative Correlation |
| STAB1 | -0.03572 | 0.547425 | Other Negative Correlation |
| TRIM66 | -0.03577 | 0.546846 | Other Negative Correlation |
| FBXO31 | -0.03579 | 0.54659 | Other Negative Correlation |
| NPIPA5 | -0.0358 | 0.546555 | Other Negative Correlation |
| CDC42BPA | -0.03593 | 0.545028 | Other Negative Correlation |
| KIAA0513 | -0.03596 | 0.544759 | Other Negative Correlation |
| SLC16A5 | -0.03603 | 0.543997 | Other Negative Correlation |
| RXRB | -0.03603 | 0.543897 | Other Negative Correlation |
| EXPH5 | -0.03607 | 0.543497 | Other Negative Correlation |
| SERPINF1 | -0.03614 | 0.542767 | Other Negative Correlation |
| LCE5A | -0.03615 | 0.54263 | Other Negative Correlation |
| ZFP37 | -0.0362 | 0.542073 | Other Negative Correlation |
| PCGF6 | -0.03624 | 0.541575 | Other Negative Correlation |
| FGF12 | -0.03625 | 0.54153 | Other Negative Correlation |
| SHC2 | -0.03629 | 0.541063 | Other Negative Correlation |
| TRIL | -0.0363 | 0.540975 | Other Negative Correlation |
| ABHD18 | -0.03632 | 0.540719 | Other Negative Correlation |
| SPNS1 | -0.03633 | 0.540587 | Other Negative Correlation |
| KCND2 | -0.03634 | 0.540512 | Other Negative Correlation |
| LYPD5 | -0.03639 | 0.539901 | Other Negative Correlation |
| BBS9 | -0.0364 | 0.539783 | Other Negative Correlation |
| CEP41 | -0.0365 | 0.538737 | Other Negative Correlation |
| SSX2IP | -0.03651 | 0.538584 | Other Negative Correlation |
| FBN1 | -0.03652 | 0.538501 | Other Negative Correlation |
| PLBD2 | -0.0366 | 0.537572 | Other Negative Correlation |
| SPATC1L | -0.03661 | 0.537527 | Other Negative Correlation |
| NAPEPLD | -0.03661 | 0.537448 | Other Negative Correlation |
| WNT3A | -0.03663 | 0.537291 | Other Negative Correlation |
| TDRD7 | -0.03663 | 0.53726 | Other Negative Correlation |
| MAP4K1 | -0.03666 | 0.536872 | Other Negative Correlation |

| | | | |
|---|---|---|---|
| C2CD2L | -0.0367 | 0.536499 | Other Negative Correlation |
| CD70 | -0.03672 | 0.536292 | Other Negative Correlation |
| EIF4EBP2 | -0.03672 | 0.536242 | Other Negative Correlation |
| PARP16 | -0.03675 | 0.535934 | Other Negative Correlation |
| FRMD5 | -0.0368 | 0.535411 | Other Negative Correlation |
| GYG2 | -0.03682 | 0.535135 | Other Negative Correlation |
| ZFTA | -0.03684 | 0.534886 | Other Negative Correlation |
| ISM2 | -0.03689 | 0.534427 | Other Negative Correlation |
| SPINK8 | -0.03689 | 0.534422 | Other Negative Correlation |
| CCL5 | -0.03693 | 0.533921 | Other Negative Correlation |
| WFDC12 | -0.03693 | 0.533906 | Other Negative Correlation |
| RNF17 | -0.03695 | 0.533759 | Other Negative Correlation |
| ADAD2 | -0.03696 | 0.533597 | Other Negative Correlation |
| BOLA1 | -0.03696 | 0.533549 | Other Negative Correlation |
| PJVK | -0.03703 | 0.532797 | Other Negative Correlation |
| LRRC43 | -0.03707 | 0.532394 | Other Negative Correlation |
| ZNF879 | -0.03712 | 0.531857 | Other Negative Correlation |
| PYHIN1 | -0.03719 | 0.531024 | Other Negative Correlation |
| ADRA2A | -0.03728 | 0.530027 | Other Negative Correlation |
| C19orf85 | -0.03728 | 0.530027 | Other Negative Correlation |
| PTRHD1 | -0.03734 | 0.529371 | Other Negative Correlation |
| KDM7A | -0.03741 | 0.528593 | Other Negative Correlation |
| NLRC3 | -0.03746 | 0.52809 | Other Negative Correlation |
| ZDHHC15 | -0.0375 | 0.527656 | Other Negative Correlation |
| CCDC134 | -0.03755 | 0.527073 | Other Negative Correlation |
| FANCM | -0.03763 | 0.526236 | Other Negative Correlation |
| THBS4 | -0.03764 | 0.526133 | Other Negative Correlation |
| MIER2 | -0.03765 | 0.525995 | Other Negative Correlation |
| VSIG10 | -0.03767 | 0.525751 | Other Negative Correlation |
| HSD17B14 | -0.03778 | 0.524576 | Other Negative Correlation |
| MRGPRE | -0.03782 | 0.524159 | Other Negative Correlation |
| CERS2 | -0.03784 | 0.523868 | Other Negative Correlation |
| ZNF432 | -0.03786 | 0.523631 | Other Negative Correlation |
| HES4 | -0.03786 | 0.523628 | Other Negative Correlation |
| OR6A2 | -0.03788 | 0.523498 | Other Negative Correlation |
| C2orf88 | -0.03788 | 0.523418 | Other Negative Correlation |
| KRT71 | -0.03794 | 0.52283 | Other Negative Correlation |
| REC114 | -0.03794 | 0.52283 | Other Negative Correlation |
| ENSG00000285978 | -0.03794 | 0.52283 | Other Negative Correlation |
| RGPD6 | -0.03794 | 0.52283 | Other Negative Correlation |
| CGA | -0.03794 | 0.52283 | Other Negative Correlation |
| EPHA5 | -0.03794 | 0.52283 | Other Negative Correlation |
| CAMK1 | -0.03798 | 0.522344 | Other Negative Correlation |
| PNMA1 | -0.03799 | 0.522211 | Other Negative Correlation |
| NUP43 | -0.03804 | 0.521664 | Other Negative Correlation |
| CNTNAP1 | -0.03808 | 0.521225 | Other Negative Correlation |

| | | | |
|---|---|---|---|
| HCRTR2 | -0.03808 | 0.521224 | Other Negative Correlation |
| ALOX15 | -0.03813 | 0.520714 | Other Negative Correlation |
| ABI3 | -0.03817 | 0.520312 | Other Negative Correlation |
| ENSG00000235007 | -0.03817 | 0.520225 | Other Negative Correlation |
| NR1H2 | -0.03826 | 0.519339 | Other Negative Correlation |
| DNAI4 | -0.03831 | 0.518695 | Other Negative Correlation |
| JCAD | -0.03832 | 0.518594 | Other Negative Correlation |
| ASAP3 | -0.03833 | 0.518505 | Other Negative Correlation |
| PIK3CA | -0.03835 | 0.518292 | Other Negative Correlation |
| MEN1 | -0.03836 | 0.518203 | Other Negative Correlation |
| PPP1R11 | -0.03842 | 0.517556 | Other Negative Correlation |
| ADGRL1 | -0.03843 | 0.51745 | Other Negative Correlation |
| FMO1 | -0.03844 | 0.51738 | Other Negative Correlation |
| ARF4 | -0.03851 | 0.516585 | Other Negative Correlation |
| GGTA1 | -0.03851 | 0.516582 | Other Negative Correlation |
| NAPA | -0.03853 | 0.516338 | Other Negative Correlation |
| NFKBIL1 | -0.03858 | 0.515831 | Other Negative Correlation |
| MTNR1A | -0.03863 | 0.51526 | Other Negative Correlation |
| SHC3 | -0.03865 | 0.515002 | Other Negative Correlation |
| EDARADD | -0.03869 | 0.514555 | Other Negative Correlation |
| XK | -0.03873 | 0.514153 | Other Negative Correlation |
| MS4A4A | -0.03879 | 0.513566 | Other Negative Correlation |
| ENSG00000284874 | -0.03879 | 0.513566 | Other Negative Correlation |
| ILVBL | -0.0388 | 0.513424 | Other Negative Correlation |
| CLU | -0.03882 | 0.513231 | Other Negative Correlation |
| LYSMD4 | -0.0389 | 0.512326 | Other Negative Correlation |
| ZNF75A | -0.03894 | 0.511901 | Other Negative Correlation |
| TEX37 | -0.03897 | 0.511612 | Other Negative Correlation |
| DTWD2 | -0.03897 | 0.511596 | Other Negative Correlation |
| C2 | -0.03905 | 0.510723 | Other Negative Correlation |
| MTCP1 | -0.03906 | 0.51059 | Other Negative Correlation |
| ANKRD23 | -0.03916 | 0.509496 | Other Negative Correlation |
| MAPK8IP3 | -0.03916 | 0.509475 | Other Negative Correlation |
| SLC25A45 | -0.03917 | 0.509445 | Other Negative Correlation |
| PSENEN | -0.03917 | 0.509435 | Other Negative Correlation |
| CHODL | -0.03918 | 0.509288 | Other Negative Correlation |
| TXLNB | -0.03918 | 0.509238 | Other Negative Correlation |
| SLC9A2 | -0.0392 | 0.509037 | Other Negative Correlation |
| RAB23 | -0.03925 | 0.508578 | Other Negative Correlation |
| DBNDD1 | -0.03931 | 0.50784 | Other Negative Correlation |
| GP5 | -0.03932 | 0.507763 | Other Negative Correlation |
| THEG | -0.03933 | 0.507634 | Other Negative Correlation |
| HOMER2 | -0.03935 | 0.507429 | Other Negative Correlation |
| SVEP1 | -0.03937 | 0.507261 | Other Negative Correlation |
| SPTB | -0.03944 | 0.506531 | Other Negative Correlation |
| TEX26 | -0.03945 | 0.50642 | Other Negative Correlation |

| | | | |
|---|---|---|---|
| ANO7 | -0.03945 | 0.506389 | Other Negative Correlation |
| NUDT10 | -0.03947 | 0.50621 | Other Negative Correlation |
| EEF1AKMT4 | -0.03948 | 0.506091 | Other Negative Correlation |
| BMERB1 | -0.0395 | 0.50584 | Other Negative Correlation |
| DENND1C | -0.03954 | 0.505375 | Other Negative Correlation |
| KIAA1755 | -0.03959 | 0.504881 | Other Negative Correlation |
| MARCHF8 | -0.03961 | 0.504642 | Other Negative Correlation |
| PVR | -0.03962 | 0.504494 | Other Negative Correlation |
| ENSG00000282936 | -0.03963 | 0.504406 | Other Negative Correlation |
| GJA4 | -0.03965 | 0.5042 | Other Negative Correlation |
| MOGAT3 | -0.03965 | 0.504195 | Other Negative Correlation |
| BIN2 | -0.03965 | 0.504195 | Other Negative Correlation |
| MICB | -0.03967 | 0.50399 | Other Negative Correlation |
| NETO1 | -0.03973 | 0.503409 | Other Negative Correlation |
| OVGP1 | -0.03974 | 0.503244 | Other Negative Correlation |
| CDH19 | -0.03981 | 0.502525 | Other Negative Correlation |
| FAHD2B | -0.03985 | 0.502082 | Other Negative Correlation |
| KLC4 | -0.03985 | 0.502053 | Other Negative Correlation |
| NAIF1 | -0.03991 | 0.50143 | Other Negative Correlation |
| ZNF718 | -0.03993 | 0.501181 | Other Negative Correlation |
| RP9 | -0.03996 | 0.500931 | Other Negative Correlation |
| MTMR9 | -0.03996 | 0.500855 | Other Negative Correlation |
| HIF1AN | -0.03998 | 0.500653 | Other Negative Correlation |
| ZMYND19 | -0.03999 | 0.500537 | Other Negative Correlation |
| LAG3 | -0.04003 | 0.500183 | Other Negative Correlation |
| DYNLT4 | -0.04012 | 0.499216 | Other Negative Correlation |
| E4F1 | -0.04014 | 0.498919 | Other Negative Correlation |
| IL17B | -0.04015 | 0.498883 | Other Negative Correlation |
| MUC4 | -0.04015 | 0.498854 | Other Negative Correlation |
| HIC2 | -0.04019 | 0.498396 | Other Negative Correlation |
| SETDB1 | -0.0402 | 0.498362 | Other Negative Correlation |
| ROBO2 | -0.04021 | 0.498263 | Other Negative Correlation |
| CACNG4 | -0.04024 | 0.497884 | Other Negative Correlation |
| NSUN3 | -0.04027 | 0.497525 | Other Negative Correlation |
| PEAK1 | -0.04028 | 0.497494 | Other Negative Correlation |
| CRNN | -0.04029 | 0.497358 | Other Negative Correlation |
| TTC9C | -0.0403 | 0.497195 | Other Negative Correlation |
| C11orf86 | -0.04031 | 0.497146 | Other Negative Correlation |
| USP30 | -0.04037 | 0.49647 | Other Negative Correlation |
| PTN | -0.0404 | 0.496197 | Other Negative Correlation |
| FIBCD1 | -0.04043 | 0.495819 | Other Negative Correlation |
| HSPB3 | -0.04044 | 0.495754 | Other Negative Correlation |
| PNPLA1 | -0.04048 | 0.49538 | Other Negative Correlation |
| GPR137 | -0.04055 | 0.494559 | Other Negative Correlation |
| IL1RL2 | -0.04059 | 0.494198 | Other Negative Correlation |
| KLHDC4 | -0.04064 | 0.493578 | Other Negative Correlation |

| | | | |
|---|---|---|---|
| FIGNL1 | -0.04068 | 0.493193 | Other Negative Correlation |
| CCDC126 | -0.04069 | 0.493067 | Other Negative Correlation |
| BBOF1 | -0.04069 | 0.493061 | Other Negative Correlation |
| CTXND1 | -0.04071 | 0.492877 | Other Negative Correlation |
| TAC3 | -0.04071 | 0.492877 | Other Negative Correlation |
| HIGD2B | -0.04075 | 0.492482 | Other Negative Correlation |
| MYOZ3 | -0.04077 | 0.492279 | Other Negative Correlation |
| KCNE5 | -0.04079 | 0.491979 | Other Negative Correlation |
| TTC30A | -0.04083 | 0.491603 | Other Negative Correlation |
| USP36 | -0.04083 | 0.491598 | Other Negative Correlation |
| TRMT10B | -0.04084 | 0.491476 | Other Negative Correlation |
| H2AL3 | -0.04086 | 0.491239 | Other Negative Correlation |
| AJM1 | -0.0409 | 0.490819 | Other Negative Correlation |
| IL17RB | -0.04092 | 0.49064 | Other Negative Correlation |
| SCX | -0.04093 | 0.49056 | Other Negative Correlation |
| PNPLA7 | -0.04093 | 0.490493 | Other Negative Correlation |
| OAZ3 | -0.04095 | 0.490368 | Other Negative Correlation |
| CD14 | -0.04104 | 0.489376 | Other Negative Correlation |
| CAPS2 | -0.04112 | 0.488546 | Other Negative Correlation |
| CCER1 | -0.04112 | 0.48851 | Other Negative Correlation |
| RAPGEF3 | -0.04118 | 0.487848 | Other Negative Correlation |
| F2RL2 | -0.04121 | 0.487532 | Other Negative Correlation |
| TBC1D20 | -0.04124 | 0.487297 | Other Negative Correlation |
| LRRC69 | -0.04129 | 0.486738 | Other Negative Correlation |
| RNF180 | -0.04129 | 0.48673 | Other Negative Correlation |
| TAF6L | -0.04132 | 0.486385 | Other Negative Correlation |
| ZSCAN16 | -0.04133 | 0.486281 | Other Negative Correlation |
| SEMA3E | -0.04139 | 0.485722 | Other Negative Correlation |
| TMED5 | -0.0414 | 0.485542 | Other Negative Correlation |
| KRBA2 | -0.04145 | 0.485013 | Other Negative Correlation |
| TSEN54 | -0.04146 | 0.484902 | Other Negative Correlation |
| RERG | -0.04153 | 0.484213 | Other Negative Correlation |
| HHAT | -0.04155 | 0.483987 | Other Negative Correlation |
| RAPGEF4 | -0.04157 | 0.483808 | Other Negative Correlation |
| GGN | -0.04174 | 0.481979 | Other Negative Correlation |
| N4BP3 | -0.04175 | 0.481854 | Other Negative Correlation |
| ZC3H11A | -0.04179 | 0.481515 | Other Negative Correlation |
| TELO2 | -0.04181 | 0.481252 | Other Negative Correlation |
| HSF5 | -0.04181 | 0.481241 | Other Negative Correlation |
| TACC1 | -0.04183 | 0.481058 | Other Negative Correlation |
| PCDHA5 | -0.04184 | 0.480934 | Other Negative Correlation |
| PIGA | -0.04185 | 0.480872 | Other Negative Correlation |
| ZNF808 | -0.04187 | 0.480648 | Other Negative Correlation |
| TAFA3 | -0.04189 | 0.480367 | Other Negative Correlation |
| VRTN | -0.04194 | 0.479848 | Other Negative Correlation |
| ENSG00000290146 | -0.04197 | 0.479601 | Other Negative Correlation |

| | | | |
|---|---|---|---|
| FAM237B | -0.04198 | 0.479462 | Other Negative Correlation |
| ENSG00000285085 | -0.04202 | 0.47903 | Other Negative Correlation |
| MRAP | -0.04211 | 0.478067 | Other Negative Correlation |
| CCDC59 | -0.04213 | 0.477894 | Other Negative Correlation |
| CDHR4 | -0.04213 | 0.47788 | Other Negative Correlation |
| DPP6 | -0.04215 | 0.477715 | Other Negative Correlation |
| MYOZ1 | -0.04217 | 0.47752 | Other Negative Correlation |
| MAPK15 | -0.04218 | 0.477343 | Other Negative Correlation |
| MORN4 | -0.04219 | 0.477233 | Other Negative Correlation |
| ZNF461 | -0.04222 | 0.476966 | Other Negative Correlation |
| TPPP | -0.04223 | 0.476827 | Other Negative Correlation |
| ZNF740 | -0.04225 | 0.476599 | Other Negative Correlation |
| NAXD | -0.04227 | 0.476483 | Other Negative Correlation |
| TMEM120B | -0.04227 | 0.476464 | Other Negative Correlation |
| SLX4 | -0.04227 | 0.476426 | Other Negative Correlation |
| H2BC15 | -0.04228 | 0.476354 | Other Negative Correlation |
| F2RL3 | -0.04229 | 0.47622 | Other Negative Correlation |
| SMTNL2 | -0.0423 | 0.476151 | Other Negative Correlation |
| CDC14A | -0.04232 | 0.4759 | Other Negative Correlation |
| CCDC73 | -0.04235 | 0.475626 | Other Negative Correlation |
| DZIP1 | -0.0424 | 0.475094 | Other Negative Correlation |
| MGA | -0.0425 | 0.474052 | Other Negative Correlation |
| SAG | -0.04252 | 0.473872 | Other Negative Correlation |
| ANKDD1B | -0.04256 | 0.473457 | Other Negative Correlation |
| WDR49 | -0.04257 | 0.473365 | Other Negative Correlation |
| GPR146 | -0.04257 | 0.473336 | Other Negative Correlation |
| DUSP19 | -0.04263 | 0.47264 | Other Negative Correlation |
| CFAP210 | -0.04265 | 0.472459 | Other Negative Correlation |
| MYO1G | -0.04266 | 0.472346 | Other Negative Correlation |
| ZNF2 | -0.04268 | 0.47212 | Other Negative Correlation |
| SRP54 | -0.04271 | 0.471862 | Other Negative Correlation |
| CDH23 | -0.04272 | 0.471753 | Other Negative Correlation |
| ZNF599 | -0.04277 | 0.471239 | Other Negative Correlation |
| PCDHA8 | -0.04282 | 0.470726 | Other Negative Correlation |
| TIGIT | -0.04282 | 0.470726 | Other Negative Correlation |
| SLC30A10 | -0.04282 | 0.470726 | Other Negative Correlation |
| KRT27 | -0.04282 | 0.470726 | Other Negative Correlation |
| PCDHA11 | -0.04282 | 0.470726 | Other Negative Correlation |
| CCDC185 | -0.04282 | 0.470726 | Other Negative Correlation |
| KRTAP4-1 | -0.04282 | 0.470726 | Other Negative Correlation |
| ENSG00000289503 | -0.04282 | 0.470726 | Other Negative Correlation |
| MINAR2 | -0.04282 | 0.470726 | Other Negative Correlation |
| HHLA2 | -0.04282 | 0.470726 | Other Negative Correlation |
| CD28 | -0.04282 | 0.470726 | Other Negative Correlation |
| ODAM | -0.04282 | 0.470726 | Other Negative Correlation |
| BFSP2 | -0.04282 | 0.470726 | Other Negative Correlation |

| | | | |
|---|---|---|---|
| SPRR2B | -0.04282 | 0.470726 | Other Negative Correlation |
| DSG4 | -0.04282 | 0.470726 | Other Negative Correlation |
| CD5 | -0.04282 | 0.470726 | Other Negative Correlation |
| LCE2C | -0.04282 | 0.470726 | Other Negative Correlation |
| TMEM207 | -0.04282 | 0.470726 | Other Negative Correlation |
| KRTAP17-1 | -0.04282 | 0.470726 | Other Negative Correlation |
| CCR5 | -0.04282 | 0.470726 | Other Negative Correlation |
| SMIM28 | -0.04282 | 0.470726 | Other Negative Correlation |
| SPINK9 | -0.04282 | 0.470726 | Other Negative Correlation |
| MYL1 | -0.04282 | 0.470726 | Other Negative Correlation |
| HPGDS | -0.04282 | 0.470726 | Other Negative Correlation |
| LCE2B | -0.04282 | 0.470726 | Other Negative Correlation |
| LIPM | -0.04282 | 0.470726 | Other Negative Correlation |
| CCDC54 | -0.04282 | 0.470726 | Other Negative Correlation |
| PSG11 | -0.04282 | 0.470726 | Other Negative Correlation |
| PSG2 | -0.04282 | 0.470726 | Other Negative Correlation |
| KRTAP4-11 | -0.04282 | 0.470726 | Other Negative Correlation |
| KRT26 | -0.04282 | 0.470726 | Other Negative Correlation |
| IL22RA2 | -0.04282 | 0.470726 | Other Negative Correlation |
| MUC7 | -0.04282 | 0.470726 | Other Negative Correlation |
| BMP10 | -0.04282 | 0.470726 | Other Negative Correlation |
| FECH | -0.04291 | 0.469833 | Other Negative Correlation |
| TEX14 | -0.04292 | 0.469722 | Other Negative Correlation |
| BICC1 | -0.04293 | 0.469613 | Other Negative Correlation |
| C7orf57 | -0.04297 | 0.469129 | Other Negative Correlation |
| DEPDC5 | -0.04298 | 0.469047 | Other Negative Correlation |
| POU4F3 | -0.04299 | 0.468986 | Other Negative Correlation |
| RHCE | -0.043 | 0.46883 | Other Negative Correlation |
| MTMR1 | -0.04303 | 0.468576 | Other Negative Correlation |
| RGS9 | -0.04306 | 0.46828 | Other Negative Correlation |
| TRIM9 | -0.04309 | 0.467946 | Other Negative Correlation |
| GPR143 | -0.04309 | 0.46791 | Other Negative Correlation |
| DLG2 | -0.04313 | 0.467486 | Other Negative Correlation |
| WNT5B | -0.04319 | 0.466918 | Other Negative Correlation |
| ZCCHC3 | -0.04324 | 0.466388 | Other Negative Correlation |
| PPARG | -0.04326 | 0.466181 | Other Negative Correlation |
| ZKSCAN7 | -0.04327 | 0.466062 | Other Negative Correlation |
| TSPYL5 | -0.04328 | 0.465909 | Other Negative Correlation |
| GPN2 | -0.0433 | 0.465734 | Other Negative Correlation |
| HAS1 | -0.04331 | 0.465628 | Other Negative Correlation |
| YAF2 | -0.04338 | 0.46496 | Other Negative Correlation |
| ZSCAN23 | -0.04338 | 0.464918 | Other Negative Correlation |
| KLHL6 | -0.0434 | 0.464679 | Other Negative Correlation |
| GRXCR2 | -0.04343 | 0.464436 | Other Negative Correlation |
| HAPLN3 | -0.04351 | 0.46363 | Other Negative Correlation |
| POU2AF1 | -0.04356 | 0.463083 | Other Negative Correlation |

| | | | |
|---|---|---|---|
| KLRC3 | -0.04358 | 0.462895 | Other Negative Correlation |
| IMPG1 | -0.04358 | 0.462895 | Other Negative Correlation |
| KCNH6 | -0.04358 | 0.462895 | Other Negative Correlation |
| ITLN2 | -0.04358 | 0.462895 | Other Negative Correlation |
| SLC25A2 | -0.04358 | 0.462895 | Other Negative Correlation |
| TPPP2 | -0.04358 | 0.462895 | Other Negative Correlation |
| NANOS3 | -0.04361 | 0.462611 | Other Negative Correlation |
| MYO15B | -0.04362 | 0.462479 | Other Negative Correlation |
| CCL3 | -0.04366 | 0.462043 | Other Negative Correlation |
| ENSG00000284762 | -0.04366 | 0.462043 | Other Negative Correlation |
| PTPN5 | -0.04366 | 0.462043 | Other Negative Correlation |
| ZPBP | -0.04366 | 0.462043 | Other Negative Correlation |
| KRT79 | -0.04371 | 0.461488 | Other Negative Correlation |
| SPEG | -0.04372 | 0.461438 | Other Negative Correlation |
| GPR89B | -0.04374 | 0.461226 | Other Negative Correlation |
| TBC1D31 | -0.04376 | 0.461027 | Other Negative Correlation |
| NCDN | -0.04385 | 0.460119 | Other Negative Correlation |
| FAM43A | -0.04388 | 0.45984 | Other Negative Correlation |
| WWP2 | -0.04391 | 0.459539 | Other Negative Correlation |
| HAAO | -0.04393 | 0.459297 | Other Negative Correlation |
| TRAPPC5 | -0.04395 | 0.459114 | Other Negative Correlation |
| CNPY3 | -0.04399 | 0.458655 | Other Negative Correlation |
| ETFBKMT | -0.044 | 0.458584 | Other Negative Correlation |
| CDH4 | -0.04403 | 0.458248 | Other Negative Correlation |
| LRRC3 | -0.04407 | 0.457811 | Other Negative Correlation |
| DTNB | -0.04414 | 0.457124 | Other Negative Correlation |
| OLFM2 | -0.04416 | 0.456921 | Other Negative Correlation |
| STRA6 | -0.04422 | 0.456327 | Other Negative Correlation |
| HSPB9 | -0.04424 | 0.456152 | Other Negative Correlation |
| HHIPL2 | -0.04425 | 0.455974 | Other Negative Correlation |
| BCAS1 | -0.04428 | 0.455669 | Other Negative Correlation |
| ZCWPW2 | -0.0443 | 0.455512 | Other Negative Correlation |
| BANK1 | -0.04431 | 0.455445 | Other Negative Correlation |
| MAP3K2 | -0.04432 | 0.45527 | Other Negative Correlation |
| MYL3 | -0.04436 | 0.454872 | Other Negative Correlation |
| SLTM | -0.04437 | 0.454801 | Other Negative Correlation |
| XKRX | -0.0444 | 0.454504 | Other Negative Correlation |
| FYN | -0.04442 | 0.454267 | Other Negative Correlation |
| RALGAPA1 | -0.04444 | 0.454083 | Other Negative Correlation |
| CLDND2 | -0.04449 | 0.45358 | Other Negative Correlation |
| BTN3A3 | -0.04451 | 0.453366 | Other Negative Correlation |
| NPHP4 | -0.04453 | 0.453129 | Other Negative Correlation |
| NAT2 | -0.04459 | 0.452553 | Other Negative Correlation |
| FIG4 | -0.0446 | 0.452478 | Other Negative Correlation |
| GPR35 | -0.0446 | 0.452425 | Other Negative Correlation |
| ENSG00000263809 | -0.04465 | 0.451985 | Other Negative Correlation |

| | | | |
|---|---|---|---|
| C16orf95 | -0.04468 | 0.451681 | Other Negative Correlation |
| STMN4 | -0.04468 | 0.451606 | Other Negative Correlation |
| CCDC3 | -0.04472 | 0.451198 | Other Negative Correlation |
| MCAM | -0.04473 | 0.45118 | Other Negative Correlation |
| ZNF652 | -0.04477 | 0.45077 | Other Negative Correlation |
| ZFAND3 | -0.04478 | 0.450682 | Other Negative Correlation |
| SLC22A5 | -0.04481 | 0.450354 | Other Negative Correlation |
| RNF32 | -0.04483 | 0.450169 | Other Negative Correlation |
| RSRP1 | -0.04483 | 0.450092 | Other Negative Correlation |
| ASPRV1 | -0.04485 | 0.449949 | Other Negative Correlation |
| GOLGA6L7 | -0.04492 | 0.449171 | Other Negative Correlation |
| DOK3 | -0.04499 | 0.448524 | Other Negative Correlation |
| TMEM117 | -0.04504 | 0.448055 | Other Negative Correlation |
| ECT2L | -0.0451 | 0.4474 | Other Negative Correlation |
| SUPV3L1 | -0.04513 | 0.447067 | Other Negative Correlation |
| ZNF45 | -0.04514 | 0.447002 | Other Negative Correlation |
| BOLA2B | -0.04525 | 0.445931 | Other Negative Correlation |
| MRAS | -0.04525 | 0.445921 | Other Negative Correlation |
| CXCR2 | -0.04528 | 0.445623 | Other Negative Correlation |
| SHISA4 | -0.04529 | 0.44545 | Other Negative Correlation |
| ENTPD1 | -0.04532 | 0.445181 | Other Negative Correlation |
| SCUBE3 | -0.04534 | 0.445024 | Other Negative Correlation |
| LGI4 | -0.04534 | 0.44502 | Other Negative Correlation |
| GRIN2A | -0.04534 | 0.445014 | Other Negative Correlation |
| GTF2A1 | -0.04536 | 0.444823 | Other Negative Correlation |
| ZNF491 | -0.04539 | 0.444469 | Other Negative Correlation |
| FAM86B2 | -0.04544 | 0.444012 | Other Negative Correlation |
| ZNF7 | -0.04546 | 0.443788 | Other Negative Correlation |
| TIMM29 | -0.04549 | 0.443479 | Other Negative Correlation |
| NDUFAF7 | -0.04553 | 0.443111 | Other Negative Correlation |
| CDC42EP2 | -0.04562 | 0.442125 | Other Negative Correlation |
| NYAP2 | -0.04563 | 0.44206 | Other Negative Correlation |
| CCDC157 | -0.04563 | 0.442054 | Other Negative Correlation |
| BCO2 | -0.04568 | 0.441584 | Other Negative Correlation |
| RAD51B | -0.04569 | 0.441465 | Other Negative Correlation |
| PER3 | -0.04572 | 0.441212 | Other Negative Correlation |
| MED1 | -0.04576 | 0.440773 | Other Negative Correlation |
| NKIRAS2 | -0.0458 | 0.440393 | Other Negative Correlation |
| CFDP1 | -0.0458 | 0.440375 | Other Negative Correlation |
| DYNLL2 | -0.04581 | 0.44026 | Other Negative Correlation |
| TYW5 | -0.04581 | 0.440254 | Other Negative Correlation |
| MAPK4 | -0.04585 | 0.439832 | Other Negative Correlation |
| HSPH1 | -0.04588 | 0.439608 | Other Negative Correlation |
| AGBL2 | -0.04589 | 0.439501 | Other Negative Correlation |
| FOXP2 | -0.04592 | 0.43921 | Other Negative Correlation |
| METTL21C | -0.04593 | 0.439119 | Other Negative Correlation |

| | | | |
|---|---|---|---|
| FCRLA | -0.04593 | 0.439073 | Other Negative Correlation |
| SELENBP1 | -0.04602 | 0.438137 | Other Negative Correlation |
| ZNF613 | -0.04611 | 0.437318 | Other Negative Correlation |
| SERPINB12 | -0.04611 | 0.437254 | Other Negative Correlation |
| KRTAP1-1 | -0.04614 | 0.437001 | Other Negative Correlation |
| KRT25 | -0.04614 | 0.437001 | Other Negative Correlation |
| ATP13A5 | -0.04614 | 0.437001 | Other Negative Correlation |
| KRTAP2-2 | -0.04614 | 0.437001 | Other Negative Correlation |
| KRTAP1-5 | -0.04614 | 0.437001 | Other Negative Correlation |
| KRT24 | -0.04614 | 0.437001 | Other Negative Correlation |
| KRTAP1-3 | -0.04614 | 0.437001 | Other Negative Correlation |
| SUPT4H1 | -0.04617 | 0.436649 | Other Negative Correlation |
| NAP1L2 | -0.04624 | 0.436023 | Other Negative Correlation |
| ZNF25 | -0.04626 | 0.435777 | Other Negative Correlation |
| MMP3 | -0.04628 | 0.435562 | Other Negative Correlation |
| ENSG00000272822 | -0.04634 | 0.434978 | Other Negative Correlation |
| KCNN2 | -0.04634 | 0.434956 | Other Negative Correlation |
| TMEM200A | -0.04635 | 0.434894 | Other Negative Correlation |
| ZNF620 | -0.04637 | 0.434723 | Other Negative Correlation |
| FAM89A | -0.04649 | 0.433492 | Other Negative Correlation |
| RBPMS | -0.04651 | 0.433279 | Other Negative Correlation |
| LIMK1 | -0.04652 | 0.433244 | Other Negative Correlation |
| GPR161 | -0.04653 | 0.433141 | Other Negative Correlation |
| ZNF20 | -0.04653 | 0.433081 | Other Negative Correlation |
| CD6 | -0.04654 | 0.433028 | Other Negative Correlation |
| GAPDHS | -0.04658 | 0.432619 | Other Negative Correlation |
| CDK5RAP1 | -0.04659 | 0.432559 | Other Negative Correlation |
| GLRX5 | -0.0466 | 0.432389 | Other Negative Correlation |
| ENSG00000258989 | -0.04662 | 0.432209 | Other Negative Correlation |
| ACP7 | -0.04663 | 0.432102 | Other Negative Correlation |
| SEMA4C | -0.04664 | 0.432014 | Other Negative Correlation |
| KBTBD2 | -0.04666 | 0.431849 | Other Negative Correlation |
| HMGCL | -0.04666 | 0.431783 | Other Negative Correlation |
| CRACDL | -0.04674 | 0.431083 | Other Negative Correlation |
| RTL8A | -0.04674 | 0.431054 | Other Negative Correlation |
| STATH | -0.04675 | 0.430982 | Other Negative Correlation |
| KRT28 | -0.04675 | 0.430982 | Other Negative Correlation |
| TMEM67 | -0.04676 | 0.430885 | Other Negative Correlation |
| ZFAND1 | -0.0469 | 0.429423 | Other Negative Correlation |
| GLIS1 | -0.04691 | 0.429395 | Other Negative Correlation |
| METTL25B | -0.04692 | 0.429311 | Other Negative Correlation |
| GSKIP | -0.04694 | 0.429056 | Other Negative Correlation |
| SLC1A1 | -0.04701 | 0.428347 | Other Negative Correlation |
| OR1F1 | -0.04708 | 0.427698 | Other Negative Correlation |
| PDXDC1 | -0.04709 | 0.427618 | Other Negative Correlation |
| ZSCAN10 | -0.04709 | 0.427581 | Other Negative Correlation |

| | | | |
|---|---|---|---|
| TARBP1 | -0.04714 | 0.427108 | Other Negative Correlation |
| BCAS3 | -0.04716 | 0.426923 | Other Negative Correlation |
| SLC6A12 | -0.04717 | 0.426813 | Other Negative Correlation |
| TM4SF4 | -0.04718 | 0.426687 | Other Negative Correlation |
| S1PR3 | -0.04722 | 0.426352 | Other Negative Correlation |
| CD101 | -0.04724 | 0.426104 | Other Negative Correlation |
| NEO1 | -0.0473 | 0.425508 | Other Negative Correlation |
| EFHC2 | -0.0473 | 0.425493 | Other Negative Correlation |
| POU2F1 | -0.04734 | 0.425102 | Other Negative Correlation |
| GMEB2 | -0.04736 | 0.42493 | Other Negative Correlation |
| MEF2C | -0.0474 | 0.424534 | Other Negative Correlation |
| ANXA10 | -0.0474 | 0.424523 | Other Negative Correlation |
| SGTB | -0.0474 | 0.424509 | Other Negative Correlation |
| LRRC32 | -0.04743 | 0.424304 | Other Negative Correlation |
| BBIP1 | -0.04743 | 0.424257 | Other Negative Correlation |
| FMO5 | -0.04744 | 0.424165 | Other Negative Correlation |
| NBPF10 | -0.04747 | 0.423842 | Other Negative Correlation |
| CIBAR1 | -0.04749 | 0.42365 | Other Negative Correlation |
| ZNF493 | -0.04756 | 0.423004 | Other Negative Correlation |
| ACYP2 | -0.04765 | 0.422147 | Other Negative Correlation |
| TNFSF14 | -0.04766 | 0.422014 | Other Negative Correlation |
| MYOM1 | -0.04767 | 0.421886 | Other Negative Correlation |
| ROS1 | -0.04774 | 0.421207 | Other Negative Correlation |
| FMR1NB | -0.04777 | 0.420917 | Other Negative Correlation |
| FAM228B | -0.04778 | 0.420822 | Other Negative Correlation |
| PACSIN3 | -0.04784 | 0.420217 | Other Negative Correlation |
| KLRC2 | -0.04785 | 0.420143 | Other Negative Correlation |
| SUSD3 | -0.04787 | 0.419999 | Other Negative Correlation |
| ZNF256 | -0.04789 | 0.419793 | Other Negative Correlation |
| TMEM52B | -0.0479 | 0.419701 | Other Negative Correlation |
| PTCD1 | -0.04792 | 0.419522 | Other Negative Correlation |
| PCDHGA11 | -0.04793 | 0.419405 | Other Negative Correlation |
| ZFAT | -0.04799 | 0.418799 | Other Negative Correlation |
| FNIP1 | -0.04802 | 0.418526 | Other Negative Correlation |
| SHQ1 | -0.04802 | 0.418521 | Other Negative Correlation |
| HPF1 | -0.04804 | 0.418352 | Other Negative Correlation |
| ZNF709 | -0.04807 | 0.418062 | Other Negative Correlation |
| SLIT1 | -0.04807 | 0.418052 | Other Negative Correlation |
| KANK3 | -0.04807 | 0.418049 | Other Negative Correlation |
| ZNF502 | -0.0481 | 0.417721 | Other Negative Correlation |
| CRYBA1 | -0.04816 | 0.417182 | Other Negative Correlation |
| RRH | -0.04816 | 0.417109 | Other Negative Correlation |
| MASP1 | -0.04824 | 0.416425 | Other Negative Correlation |
| TEX35 | -0.04824 | 0.416339 | Other Negative Correlation |
| CDK18 | -0.04825 | 0.416319 | Other Negative Correlation |
| SUGP2 | -0.04825 | 0.416301 | Other Negative Correlation |

| | | | |
|---|---|---|---|
| RAD21L1 | -0.04825 | 0.416274 | Other Negative Correlation |
| CELF1 | -0.04827 | 0.416081 | Other Negative Correlation |
| CHAF1B | -0.04828 | 0.416025 | Other Negative Correlation |
| RNF113A | -0.04829 | 0.415888 | Other Negative Correlation |
| ENSG00000267706 | -0.04833 | 0.415491 | Other Negative Correlation |
| NTNG2 | -0.04835 | 0.415313 | Other Negative Correlation |
| ATG9B | -0.04836 | 0.415195 | Other Negative Correlation |
| SCNN1G | -0.04837 | 0.415121 | Other Negative Correlation |
| IPCEF1 | -0.04837 | 0.415092 | Other Negative Correlation |
| SEZ6L2 | -0.04846 | 0.414271 | Other Negative Correlation |
| ANKHD1-EIF4EBP3 | -0.04847 | 0.414143 | Other Negative Correlation |
| COL26A1 | -0.04848 | 0.414102 | Other Negative Correlation |
| ZNF625 | -0.04853 | 0.413561 | Other Negative Correlation |
| RPP21 | -0.04854 | 0.413487 | Other Negative Correlation |
| OSCAR | -0.04862 | 0.412743 | Other Negative Correlation |
| TMSB4Y | -0.0487 | 0.411936 | Other Negative Correlation |
| WTIP | -0.04872 | 0.411769 | Other Negative Correlation |
| SMIM2 | -0.04876 | 0.411339 | Other Negative Correlation |
| XCL1 | -0.04876 | 0.411339 | Other Negative Correlation |
| CX3CR1 | -0.04876 | 0.411339 | Other Negative Correlation |
| C1orf105 | -0.04876 | 0.411339 | Other Negative Correlation |
| VCAM1 | -0.04876 | 0.411339 | Other Negative Correlation |
| ENSG00000281883 | -0.04876 | 0.411339 | Other Negative Correlation |
| TINAG | -0.04876 | 0.411339 | Other Negative Correlation |
| PPEF1 | -0.04876 | 0.411339 | Other Negative Correlation |
| ATP8B4 | -0.04876 | 0.411339 | Other Negative Correlation |
| GSTA2 | -0.04876 | 0.411339 | Other Negative Correlation |
| VWC2 | -0.04876 | 0.411339 | Other Negative Correlation |
| CTXN3 | -0.04876 | 0.411339 | Other Negative Correlation |
| XCL2 | -0.04876 | 0.411339 | Other Negative Correlation |
| TXNDC8 | -0.04876 | 0.411339 | Other Negative Correlation |
| CYP19A1 | -0.04876 | 0.411339 | Other Negative Correlation |
| FASLG | -0.04876 | 0.411339 | Other Negative Correlation |
| HPSE2 | -0.04876 | 0.411339 | Other Negative Correlation |
| H2BW2 | -0.04876 | 0.411339 | Other Negative Correlation |
| SLA | -0.04876 | 0.411339 | Other Negative Correlation |
| DPT | -0.04876 | 0.411339 | Other Negative Correlation |
| H2AC1 | -0.04876 | 0.411339 | Other Negative Correlation |
| HTR2A | -0.04876 | 0.411339 | Other Negative Correlation |
| SIRPG | -0.04876 | 0.411339 | Other Negative Correlation |
| CLEC1B | -0.04876 | 0.411339 | Other Negative Correlation |
| SPAG11B | -0.04876 | 0.411339 | Other Negative Correlation |
| SMR3A | -0.04876 | 0.411339 | Other Negative Correlation |
| SCGN | -0.04876 | 0.411339 | Other Negative Correlation |
| CRH | -0.04876 | 0.411339 | Other Negative Correlation |
| GNGT1 | -0.04876 | 0.411339 | Other Negative Correlation |

| | | | |
|---|---|---|---|
| STYXL2 | -0.04876 | 0.411339 | Other Negative Correlation |
| HAO1 | -0.04876 | 0.411339 | Other Negative Correlation |
| FAM216B | -0.04876 | 0.411339 | Other Negative Correlation |
| CLEC1A | -0.04876 | 0.411339 | Other Negative Correlation |
| PAM16 | -0.04877 | 0.411268 | Other Negative Correlation |
| C16orf91 | -0.0488 | 0.41096 | Other Negative Correlation |
| USP26 | -0.04883 | 0.410683 | Other Negative Correlation |
| EXOG | -0.04884 | 0.410605 | Other Negative Correlation |
| CCDC89 | -0.04885 | 0.410519 | Other Negative Correlation |
| DNM3 | -0.04885 | 0.410472 | Other Negative Correlation |
| CTAGE9 | -0.04888 | 0.410238 | Other Negative Correlation |
| ZNF30 | -0.04889 | 0.410104 | Other Negative Correlation |
| NPIPA8 | -0.04894 | 0.409664 | Other Negative Correlation |
| CPAMD8 | -0.04898 | 0.40925 | Other Negative Correlation |
| NSG2 | -0.04901 | 0.408985 | Other Negative Correlation |
| CRAT | -0.04905 | 0.408566 | Other Negative Correlation |
| SC5D | -0.04907 | 0.408407 | Other Negative Correlation |
| LMBR1L | -0.04909 | 0.408235 | Other Negative Correlation |
| MEIOSIN | -0.04909 | 0.408217 | Other Negative Correlation |
| ADHFE1 | -0.04909 | 0.40819 | Other Negative Correlation |
| ING1 | -0.04921 | 0.407095 | Other Negative Correlation |
| PHF12 | -0.04925 | 0.406658 | Other Negative Correlation |
| NIPA1 | -0.04927 | 0.406521 | Other Negative Correlation |
| CFAP299 | -0.04929 | 0.406333 | Other Negative Correlation |
| SLC25A51 | -0.04934 | 0.405826 | Other Negative Correlation |
| POPDC2 | -0.04937 | 0.405492 | Other Negative Correlation |
| ZNF136 | -0.04938 | 0.405486 | Other Negative Correlation |
| MCEE | -0.04944 | 0.404897 | Other Negative Correlation |
| BCL2L10 | -0.04947 | 0.404621 | Other Negative Correlation |
| MTMR3 | -0.0495 | 0.404338 | Other Negative Correlation |
| PPP2R5B | -0.04951 | 0.40421 | Other Negative Correlation |
| CLDN23 | -0.04959 | 0.403473 | Other Negative Correlation |
| ATP11B | -0.04961 | 0.403257 | Other Negative Correlation |
| FBP1 | -0.04962 | 0.403132 | Other Negative Correlation |
| FZD1 | -0.04966 | 0.402825 | Other Negative Correlation |
| CCDC175 | -0.04968 | 0.402629 | Other Negative Correlation |
| DCT | -0.04968 | 0.402629 | Other Negative Correlation |
| AGR3 | -0.04968 | 0.402629 | Other Negative Correlation |
| TTI1 | -0.04975 | 0.401957 | Other Negative Correlation |
| ESR2 | -0.04977 | 0.401776 | Other Negative Correlation |
| ZNF721 | -0.04977 | 0.401715 | Other Negative Correlation |
| ARFGEF2 | -0.04982 | 0.40131 | Other Negative Correlation |
| PANK4 | -0.04982 | 0.401233 | Other Negative Correlation |
| CFAP70 | -0.04984 | 0.401101 | Other Negative Correlation |
| PTPN21 | -0.04988 | 0.400709 | Other Negative Correlation |
| VAMP2 | -0.0499 | 0.400515 | Other Negative Correlation |

| | | | |
|---|---|---|---|
| BACH2 | -0.04995 | 0.400038 | Other Negative Correlation |
| CRB3 | -0.05 | 0.399531 | Other Negative Correlation |
| METTL3 | -0.05001 | 0.399465 | Other Negative Correlation |
| PTPRB | -0.05008 | 0.398788 | Other Negative Correlation |
| PIAS2 | -0.0501 | 0.398607 | Other Negative Correlation |
| FAT3 | -0.05011 | 0.398538 | Other Negative Correlation |
| GALNT8 | -0.0502 | 0.397688 | Other Negative Correlation |
| HEATR4 | -0.05024 | 0.397322 | Other Negative Correlation |
| SLC35G5 | -0.05031 | 0.396618 | Other Negative Correlation |
| DPM2 | -0.05033 | 0.396413 | Other Negative Correlation |
| CLEC4D | -0.05035 | 0.396279 | Other Negative Correlation |
| MGAM | -0.05043 | 0.395497 | Other Negative Correlation |
| ANKAR | -0.05045 | 0.395364 | Other Negative Correlation |
| CCDC137 | -0.05048 | 0.395073 | Other Negative Correlation |
| TSPAN2 | -0.05049 | 0.394957 | Other Negative Correlation |
| ATG13 | -0.0505 | 0.394889 | Other Negative Correlation |
| ENSG00000266302 | -0.05053 | 0.39458 | Other Negative Correlation |
| ADAMTSL2 | -0.05054 | 0.394487 | Other Negative Correlation |
| PIGZ | -0.05054 | 0.394486 | Other Negative Correlation |
| TDP1 | -0.05056 | 0.394254 | Other Negative Correlation |
| C9orf152 | -0.05058 | 0.394124 | Other Negative Correlation |
| FBXO46 | -0.05068 | 0.393166 | Other Negative Correlation |
| ANKRD49 | -0.0507 | 0.39302 | Other Negative Correlation |
| SULT1E1 | -0.05071 | 0.392928 | Other Negative Correlation |
| MLPH | -0.05073 | 0.392719 | Other Negative Correlation |
| DGKD | -0.05077 | 0.392349 | Other Negative Correlation |
| TLDC2 | -0.05079 | 0.392187 | Other Negative Correlation |
| YIPF5 | -0.05081 | 0.391984 | Other Negative Correlation |
| RSPH9 | -0.05084 | 0.39172 | Other Negative Correlation |
| SLC17A4 | -0.05086 | 0.391481 | Other Negative Correlation |
| PCDHB6 | -0.05086 | 0.391481 | Other Negative Correlation |
| AARS2 | -0.05089 | 0.391256 | Other Negative Correlation |
| SDE2 | -0.05095 | 0.390701 | Other Negative Correlation |
| BEST2 | -0.05095 | 0.390641 | Other Negative Correlation |
| KEL | -0.05096 | 0.390598 | Other Negative Correlation |
| LCA5L | -0.05099 | 0.390304 | Other Negative Correlation |
| BSND | -0.05101 | 0.390054 | Other Negative Correlation |
| C4orf46 | -0.05106 | 0.389664 | Other Negative Correlation |
| STARD3 | -0.05108 | 0.389494 | Other Negative Correlation |
| EFS | -0.0511 | 0.389257 | Other Negative Correlation |
| THAP10 | -0.05112 | 0.389091 | Other Negative Correlation |
| ZNF227 | -0.05116 | 0.388713 | Other Negative Correlation |
| TVP23C | -0.05121 | 0.388224 | Other Negative Correlation |
| SYNPO2L | -0.05122 | 0.388117 | Other Negative Correlation |
| SRGN | -0.05127 | 0.387711 | Other Negative Correlation |
| ADAMTS18 | -0.05129 | 0.387497 | Other Negative Correlation |

| | | | |
|---|---|---|---|
| S100B | -0.05132 | 0.387242 | Other Negative Correlation |
| GOLGA6L2 | -0.0514 | 0.38649 | Other Negative Correlation |
| KLHL11 | -0.05144 | 0.386119 | Other Negative Correlation |
| EFHB | -0.05144 | 0.386078 | Other Negative Correlation |
| ENSG00000269755 | -0.05155 | 0.385074 | Other Negative Correlation |
| CXXC5 | -0.05157 | 0.384936 | Other Negative Correlation |
| ZBTB34 | -0.0516 | 0.384669 | Other Negative Correlation |
| SERPINA3 | -0.05161 | 0.384532 | Other Negative Correlation |
| ARHGEF11 | -0.05162 | 0.384444 | Other Negative Correlation |
| RIGI | -0.05165 | 0.38415 | Other Negative Correlation |
| PDS5B | -0.05166 | 0.384068 | Other Negative Correlation |
| H3C10 | -0.0517 | 0.383738 | Other Negative Correlation |
| CCDC96 | -0.0517 | 0.383682 | Other Negative Correlation |
| ELN | -0.05172 | 0.383527 | Other Negative Correlation |
| TTC39B | -0.05172 | 0.383499 | Other Negative Correlation |
| HLA-C | -0.05174 | 0.383331 | Other Negative Correlation |
| ENSG00000285943 | -0.05179 | 0.382862 | Other Negative Correlation |
| SNAI3 | -0.05182 | 0.382647 | Other Negative Correlation |
| WASF3 | -0.05183 | 0.382491 | Other Negative Correlation |
| SNIP1 | -0.05185 | 0.382375 | Other Negative Correlation |
| SULT1C4 | -0.05185 | 0.382355 | Other Negative Correlation |
| FKBP1C | -0.05188 | 0.382054 | Other Negative Correlation |
| KRTAP3-2 | -0.05188 | 0.38205 | Other Negative Correlation |
| TMEM62 | -0.05194 | 0.381482 | Other Negative Correlation |
| EOLA1 | -0.05201 | 0.380877 | Other Negative Correlation |
| ZDHHC1 | -0.05201 | 0.380835 | Other Negative Correlation |
| SENP7 | -0.05203 | 0.380662 | Other Negative Correlation |
| ARMC5 | -0.05209 | 0.380093 | Other Negative Correlation |
| SIAH1 | -0.0521 | 0.380032 | Other Negative Correlation |
| ZNF248 | -0.05211 | 0.379966 | Other Negative Correlation |
| SPPL2B | -0.05211 | 0.379934 | Other Negative Correlation |
| CHGA | -0.05213 | 0.379771 | Other Negative Correlation |
| PDE1A | -0.05216 | 0.379445 | Other Negative Correlation |
| RBP2 | -0.05221 | 0.379028 | Other Negative Correlation |
| MAP3K19 | -0.05221 | 0.379028 | Other Negative Correlation |
| PPIL6 | -0.05226 | 0.378612 | Other Negative Correlation |
| MYCN | -0.05229 | 0.378335 | Other Negative Correlation |
| DHPS | -0.05231 | 0.378132 | Other Negative Correlation |
| TESMIN | -0.05234 | 0.377825 | Other Negative Correlation |
| MMD | -0.05236 | 0.377648 | Other Negative Correlation |
| POM121L2 | -0.05245 | 0.376805 | Other Negative Correlation |
| LIMCH1 | -0.05248 | 0.376585 | Other Negative Correlation |
| TADA1 | -0.05249 | 0.376477 | Other Negative Correlation |
| SLC10A7 | -0.05249 | 0.37644 | Other Negative Correlation |
| C2CD4B | -0.0525 | 0.376427 | Other Negative Correlation |
| ABCA4 | -0.05253 | 0.376085 | Other Negative Correlation |

| | | | |
|---|---|---|---|
| FAM180B | -0.05253 | 0.376085 | Other Negative Correlation |
| SMCP | -0.05253 | 0.376085 | Other Negative Correlation |
| GPR78 | -0.05253 | 0.376085 | Other Negative Correlation |
| WDR90 | -0.05254 | 0.375978 | Other Negative Correlation |
| PEBP4 | -0.05257 | 0.375751 | Other Negative Correlation |
| C11orf16 | -0.05258 | 0.375636 | Other Negative Correlation |
| CLK4 | -0.0526 | 0.37549 | Other Negative Correlation |
| CGN | -0.05261 | 0.375356 | Other Negative Correlation |
| IRAK1BP1 | -0.05264 | 0.375141 | Other Negative Correlation |
| MCCC1 | -0.05265 | 0.374989 | Other Negative Correlation |
| PPM1B | -0.05266 | 0.374941 | Other Negative Correlation |
| SNX15 | -0.05269 | 0.374674 | Other Negative Correlation |
| ZSCAN31 | -0.05285 | 0.373229 | Other Negative Correlation |
| AMY2B | -0.05287 | 0.373001 | Other Negative Correlation |
| ALPP | -0.05291 | 0.372682 | Other Negative Correlation |
| MYCL | -0.05292 | 0.372599 | Other Negative Correlation |
| KIAA2026 | -0.05299 | 0.371911 | Other Negative Correlation |
| TIRAP | -0.053 | 0.37189 | Other Negative Correlation |
| RIBC1 | -0.05305 | 0.371387 | Other Negative Correlation |
| VWA2 | -0.05306 | 0.371303 | Other Negative Correlation |
| RFESD | -0.05308 | 0.371108 | Other Negative Correlation |
| SLC9A5 | -0.0531 | 0.370988 | Other Negative Correlation |
| ZNF418 | -0.05311 | 0.370867 | Other Negative Correlation |
| INCENP | -0.05312 | 0.370804 | Other Negative Correlation |
| BDNF | -0.05318 | 0.370208 | Other Negative Correlation |
| EXOC3L2 | -0.05318 | 0.370203 | Other Negative Correlation |
| NR1H3 | -0.0532 | 0.370029 | Other Negative Correlation |
| ZNF346 | -0.05323 | 0.369816 | Other Negative Correlation |
| UBQLNL | -0.05323 | 0.369774 | Other Negative Correlation |
| ZNF497 | -0.05324 | 0.369646 | Other Negative Correlation |
| MEIS3 | -0.05335 | 0.368701 | Other Negative Correlation |
| RAPGEF1 | -0.05335 | 0.368671 | Other Negative Correlation |
| GSTM2 | -0.05338 | 0.36847 | Other Negative Correlation |
| SOAT1 | -0.0534 | 0.368269 | Other Negative Correlation |
| C8orf82 | -0.05343 | 0.368021 | Other Negative Correlation |
| HNRNPA0 | -0.05349 | 0.36746 | Other Negative Correlation |
| AP5Z1 | -0.05353 | 0.367042 | Other Negative Correlation |
| CCDC34 | -0.05355 | 0.366912 | Other Negative Correlation |
| SCAMP4 | -0.05358 | 0.366647 | Other Negative Correlation |
| GPR156 | -0.05361 | 0.366356 | Other Negative Correlation |
| NPIPB6 | -0.05364 | 0.366092 | Other Negative Correlation |
| ASB8 | -0.05368 | 0.3657 | Other Negative Correlation |
| EGFL8 | -0.05369 | 0.365626 | Other Negative Correlation |
| NPB | -0.05375 | 0.365105 | Other Negative Correlation |
| VWA7 | -0.05376 | 0.365019 | Other Negative Correlation |
| SPIN2A | -0.05377 | 0.364913 | Other Negative Correlation |

| | | | |
|---|---|---|---|
| CPEB2 | -0.05381 | 0.364615 | Other Negative Correlation |
| FZD5 | -0.05382 | 0.364498 | Other Negative Correlation |
| FIGN | -0.05384 | 0.364284 | Other Negative Correlation |
| PCDHB15 | -0.05384 | 0.364273 | Other Negative Correlation |
| PSG8 | -0.05389 | 0.36384 | Other Negative Correlation |
| ZNF669 | -0.05391 | 0.363722 | Other Negative Correlation |
| RHOXF1 | -0.05393 | 0.363499 | Other Negative Correlation |
| CLHC1 | -0.05395 | 0.36334 | Other Negative Correlation |
| ENSG00000284773 | -0.05397 | 0.363163 | Other Negative Correlation |
| EGLN2 | -0.05404 | 0.362538 | Other Negative Correlation |
| AGO4 | -0.0541 | 0.361956 | Other Negative Correlation |
| MINDY4 | -0.05412 | 0.361807 | Other Negative Correlation |
| IYD | -0.05412 | 0.361794 | Other Negative Correlation |
| TMEM256-PLSCR3 | -0.05412 | 0.361794 | Other Negative Correlation |
| ENSG00000276087 | -0.05412 | 0.361794 | Other Negative Correlation |
| KCNJ1 | -0.05412 | 0.361773 | Other Negative Correlation |
| CATSPER3 | -0.05415 | 0.361556 | Other Negative Correlation |
| CIITA | -0.05422 | 0.360926 | Other Negative Correlation |
| PCNT | -0.05434 | 0.359887 | Other Negative Correlation |
| NLRC5 | -0.05435 | 0.359807 | Other Negative Correlation |
| RNLS | -0.05435 | 0.359804 | Other Negative Correlation |
| C20orf204 | -0.05437 | 0.359592 | Other Negative Correlation |
| KPTN | -0.05443 | 0.359089 | Other Negative Correlation |
| IQCD | -0.05444 | 0.358973 | Other Negative Correlation |
| RMDN3 | -0.05445 | 0.358872 | Other Negative Correlation |
| CRB1 | -0.05449 | 0.358534 | Other Negative Correlation |
| DZANK1 | -0.05451 | 0.3584 | Other Negative Correlation |
| CWH43 | -0.05452 | 0.358275 | Other Negative Correlation |
| MT-ATP8 | -0.05454 | 0.358118 | Other Negative Correlation |
| DNAH9 | -0.05456 | 0.35788 | Other Negative Correlation |
| NOMO3 | -0.05458 | 0.357763 | Other Negative Correlation |
| ZFYVE27 | -0.05459 | 0.357696 | Other Negative Correlation |
| M1AP | -0.05459 | 0.357627 | Other Negative Correlation |
| CACNA2D3 | -0.0546 | 0.35754 | Other Negative Correlation |
| ZXDA | -0.05462 | 0.357413 | Other Negative Correlation |
| S1PR2 | -0.05462 | 0.357397 | Other Negative Correlation |
| HSPA2 | -0.05463 | 0.357268 | Other Negative Correlation |
| FMC1 | -0.05464 | 0.357237 | Other Negative Correlation |
| PHTF1 | -0.05468 | 0.356871 | Other Negative Correlation |
| CEP112 | -0.05471 | 0.356635 | Other Negative Correlation |
| ARSL | -0.05474 | 0.356365 | Other Negative Correlation |
| LYSMD3 | -0.05476 | 0.356195 | Other Negative Correlation |
| ABO | -0.05476 | 0.356187 | Other Negative Correlation |
| PECR | -0.05483 | 0.355575 | Other Negative Correlation |
| RAB19 | -0.05484 | 0.355451 | Other Negative Correlation |
| HBQ1 | -0.05485 | 0.355393 | Other Negative Correlation |

| | | | |
|---|---|---|---|
| TSTD2 | -0.05487 | 0.355211 | Other Negative Correlation |
| VRK3 | -0.05488 | 0.35507 | Other Negative Correlation |
| ABCD4 | -0.05489 | 0.355013 | Other Negative Correlation |
| MEDAG | -0.05497 | 0.354344 | Other Negative Correlation |
| DPY19L1 | -0.05498 | 0.354207 | Other Negative Correlation |
| TTR | -0.05503 | 0.353805 | Other Negative Correlation |
| ENSG00000288635 | -0.05505 | 0.35364 | Other Negative Correlation |
| DCDC1 | -0.05506 | 0.353536 | Other Negative Correlation |
| SLC5A12 | -0.05507 | 0.353418 | Other Negative Correlation |
| ANKRD55 | -0.05507 | 0.353418 | Other Negative Correlation |
| SLC38A11 | -0.05507 | 0.353418 | Other Negative Correlation |
| CPT1B | -0.05509 | 0.353295 | Other Negative Correlation |
| POFUT2 | -0.0551 | 0.353138 | Other Negative Correlation |
| ZNF233 | -0.05511 | 0.353123 | Other Negative Correlation |
| SMPD1 | -0.05518 | 0.352461 | Other Negative Correlation |
| ST20-MTHFS | -0.0552 | 0.352262 | Other Negative Correlation |
| RNF224 | -0.05529 | 0.351505 | Other Negative Correlation |
| CDADC1 | -0.0553 | 0.351439 | Other Negative Correlation |
| DCTN3 | -0.05531 | 0.351344 | Other Negative Correlation |
| TRIM64B | -0.05531 | 0.351333 | Other Negative Correlation |
| CD248 | -0.05531 | 0.351332 | Other Negative Correlation |
| YEATS4 | -0.05532 | 0.351248 | Other Negative Correlation |
| SLC8B1 | -0.05537 | 0.350796 | Other Negative Correlation |
| TMEM214 | -0.05537 | 0.350793 | Other Negative Correlation |
| CASQ1 | -0.05539 | 0.350635 | Other Negative Correlation |
| TCIM | -0.0554 | 0.350574 | Other Negative Correlation |
| GFAP | -0.05544 | 0.350194 | Other Negative Correlation |
| ATF2 | -0.05546 | 0.350003 | Other Negative Correlation |
| ACD | -0.05547 | 0.349963 | Other Negative Correlation |
| CD3D | -0.05548 | 0.349845 | Other Negative Correlation |
| ZNF573 | -0.05557 | 0.349049 | Other Negative Correlation |
| SMAD6 | -0.05561 | 0.3487 | Other Negative Correlation |
| SEC16A | -0.05562 | 0.348629 | Other Negative Correlation |
| CEP295 | -0.05563 | 0.348526 | Other Negative Correlation |
| ANKMY2 | -0.05567 | 0.348203 | Other Negative Correlation |
| BAHD1 | -0.05568 | 0.348152 | Other Negative Correlation |
| SEC22B | -0.0557 | 0.347941 | Other Negative Correlation |
| VMAC | -0.05571 | 0.347889 | Other Negative Correlation |
| ZNF780A | -0.05573 | 0.347673 | Other Negative Correlation |
| LRRC61 | -0.05577 | 0.347309 | Other Negative Correlation |
| MTFR1L | -0.05581 | 0.347025 | Other Negative Correlation |
| POU2F3 | -0.05592 | 0.346054 | Other Negative Correlation |
| FCGR2A | -0.05593 | 0.345987 | Other Negative Correlation |
| CCSER2 | -0.05594 | 0.345881 | Other Negative Correlation |
| HMGB1P1 | -0.05597 | 0.3456 | Other Negative Correlation |
| SIRPB1 | -0.05599 | 0.345419 | Other Negative Correlation |

| | | | |
|---|---|---|---|
| SLC36A1 | -0.05599 | 0.345409 | Other Negative Correlation |
| KNDC1 | -0.05601 | 0.345284 | Other Negative Correlation |
| BCORL1 | -0.0561 | 0.344508 | Other Negative Correlation |
| HEPACAM2 | -0.05613 | 0.3442 | Other Negative Correlation |
| SLC12A5 | -0.05615 | 0.344055 | Other Negative Correlation |
| ZNF660 | -0.05619 | 0.343678 | Other Negative Correlation |
| RAB3B | -0.05621 | 0.343546 | Other Negative Correlation |
| HR | -0.05624 | 0.34325 | Other Negative Correlation |
| HS3ST3B1 | -0.05634 | 0.342386 | Other Negative Correlation |
| DDX11 | -0.05641 | 0.341864 | Other Negative Correlation |
| IGFL2 | -0.05644 | 0.341553 | Other Negative Correlation |
| STON1 | -0.05645 | 0.34151 | Other Negative Correlation |
| NFATC3 | -0.05654 | 0.340723 | Other Negative Correlation |
| NAP1L3 | -0.05656 | 0.340585 | Other Negative Correlation |
| HAUS3 | -0.05656 | 0.340541 | Other Negative Correlation |
| GIPR | -0.0566 | 0.340212 | Other Negative Correlation |
| BTG4 | -0.05662 | 0.340071 | Other Negative Correlation |
| PAAF1 | -0.05662 | 0.339999 | Other Negative Correlation |
| CTSH | -0.05663 | 0.339962 | Other Negative Correlation |
| SOWAHD | -0.05669 | 0.339423 | Other Negative Correlation |
| TIGD6 | -0.05672 | 0.339157 | Other Negative Correlation |
| PCDHA7 | -0.05674 | 0.339005 | Other Negative Correlation |
| SLC39A8 | -0.05675 | 0.338892 | Other Negative Correlation |
| DNAJB12 | -0.05677 | 0.338788 | Other Negative Correlation |
| C20orf141 | -0.05679 | 0.338617 | Other Negative Correlation |
| NUS1 | -0.05682 | 0.338312 | Other Negative Correlation |
| FABP7 | -0.05683 | 0.338243 | Other Negative Correlation |
| KLHL38 | -0.05689 | 0.337771 | Other Negative Correlation |
| HOXA3 | -0.0569 | 0.337664 | Other Negative Correlation |
| CKLF | -0.0569 | 0.337663 | Other Negative Correlation |
| FAM185A | -0.05703 | 0.336574 | Other Negative Correlation |
| CUBN | -0.05704 | 0.336441 | Other Negative Correlation |
| ADRA1B | -0.05706 | 0.336317 | Other Negative Correlation |
| S100A5 | -0.05706 | 0.336278 | Other Negative Correlation |
| AK8 | -0.05708 | 0.336153 | Other Negative Correlation |
| PRSS21 | -0.05709 | 0.336062 | Other Negative Correlation |
| ST3GAL6 | -0.05711 | 0.335878 | Other Negative Correlation |
| ZKSCAN4 | -0.05711 | 0.335878 | Other Negative Correlation |
| CD300A | -0.05711 | 0.335851 | Other Negative Correlation |
| FBXO33 | -0.05712 | 0.335767 | Other Negative Correlation |
| CCDC38 | -0.05718 | 0.335306 | Other Negative Correlation |
| AK4P3 | -0.05718 | 0.335288 | Other Negative Correlation |
| PTPRJ | -0.05724 | 0.334737 | Other Negative Correlation |
| TMEM87A | -0.05725 | 0.334641 | Other Negative Correlation |
| SSTR3 | -0.05727 | 0.334481 | Other Negative Correlation |
| NXPE2 | -0.05727 | 0.334481 | Other Negative Correlation |

| | | | |
|---|---|---|---|
| OPALIN | -0.05727 | 0.334481 | Other Negative Correlation |
| PLK5 | -0.05729 | 0.334377 | Other Negative Correlation |
| RHNO1 | -0.05731 | 0.334143 | Other Negative Correlation |
| KIAA1549 | -0.05734 | 0.333883 | Other Negative Correlation |
| ZNF223 | -0.05735 | 0.333821 | Other Negative Correlation |
| GLP2R | -0.0574 | 0.333388 | Other Negative Correlation |
| CIBAR2 | -0.0574 | 0.333388 | Other Negative Correlation |
| CD86 | -0.0574 | 0.333388 | Other Negative Correlation |
| SYCE1 | -0.0574 | 0.333388 | Other Negative Correlation |
| RFPL4B | -0.0574 | 0.333388 | Other Negative Correlation |
| SLCO2A1 | -0.05746 | 0.332935 | Other Negative Correlation |
| GABARAP | -0.05746 | 0.33287 | Other Negative Correlation |
| KIF27 | -0.05749 | 0.332609 | Other Negative Correlation |
| KCNAB2 | -0.05752 | 0.332437 | Other Negative Correlation |
| FKBP2 | -0.05755 | 0.332163 | Other Negative Correlation |
| EP300 | -0.05756 | 0.332049 | Other Negative Correlation |
| E2F6 | -0.0576 | 0.331763 | Other Negative Correlation |
| PCDHB16 | -0.05769 | 0.33098 | Other Negative Correlation |
| VPS9D1 | -0.05775 | 0.330486 | Other Negative Correlation |
| CLIP3 | -0.05775 | 0.330461 | Other Negative Correlation |
| PDIA2 | -0.05775 | 0.330426 | Other Negative Correlation |
| ENSG00000258465 | -0.0578 | 0.330016 | Other Negative Correlation |
| C1orf100 | -0.0578 | 0.330016 | Other Negative Correlation |
| PHKA1 | -0.05783 | 0.329753 | Other Negative Correlation |
| PLAG1 | -0.05788 | 0.329396 | Other Negative Correlation |
| PIGK | -0.05796 | 0.328717 | Other Negative Correlation |
| KCNN3 | -0.05798 | 0.328503 | Other Negative Correlation |
| CAMTA2 | -0.05801 | 0.328322 | Other Negative Correlation |
| UNC79 | -0.05802 | 0.328197 | Other Negative Correlation |
| PLA2G4A | -0.0581 | 0.327564 | Other Negative Correlation |
| FAM107A | -0.05817 | 0.326963 | Other Negative Correlation |
| ZNF311 | -0.05818 | 0.326891 | Other Negative Correlation |
| FCGRT | -0.0582 | 0.326742 | Other Negative Correlation |
| FRMD1 | -0.0582 | 0.326703 | Other Negative Correlation |
| FYB2 | -0.05823 | 0.326478 | Other Negative Correlation |
| MST1 | -0.05841 | 0.324992 | Other Negative Correlation |
| ZNF160 | -0.05841 | 0.32499 | Other Negative Correlation |
| UPK1A | -0.05844 | 0.324701 | Other Negative Correlation |
| SLC36A4 | -0.05847 | 0.324459 | Other Negative Correlation |
| MUC3A | -0.05848 | 0.324398 | Other Negative Correlation |
| CCDC166 | -0.05853 | 0.323935 | Other Negative Correlation |
| ANAPC1 | -0.05856 | 0.323739 | Other Negative Correlation |
| NIPA2 | -0.05861 | 0.323319 | Other Negative Correlation |
| AP1AR | -0.05865 | 0.322938 | Other Negative Correlation |
| GCA | -0.05866 | 0.322877 | Other Negative Correlation |
| CYP7B1 | -0.05866 | 0.322861 | Other Negative Correlation |

| | | | |
|---|---|---|---|
| ZNF354B | -0.05887 | 0.321163 | Other Negative Correlation |
| IFI30 | -0.05888 | 0.321111 | Other Negative Correlation |
| LHFPL2 | -0.05894 | 0.320562 | Other Negative Correlation |
| FOXE1 | -0.05901 | 0.319981 | Other Negative Correlation |
| SALL2 | -0.05902 | 0.319918 | Other Negative Correlation |
| FAM228A | -0.05902 | 0.319912 | Other Negative Correlation |
| B3GNT7 | -0.05904 | 0.319793 | Other Negative Correlation |
| H3C6 | -0.05905 | 0.319715 | Other Negative Correlation |
| C17orf100 | -0.05906 | 0.319562 | Other Negative Correlation |
| KCNRG | -0.05907 | 0.319474 | Other Negative Correlation |
| MAK | -0.0591 | 0.319264 | Other Negative Correlation |
| GLDC | -0.05911 | 0.319208 | Other Negative Correlation |
| PIK3C2B | -0.05915 | 0.318873 | Other Negative Correlation |
| SHANK3 | -0.05915 | 0.318821 | Other Negative Correlation |
| NBPF11 | -0.05916 | 0.318771 | Other Negative Correlation |
| EPYC | -0.05918 | 0.318581 | Other Negative Correlation |
| PAGR1 | -0.05932 | 0.317466 | Other Negative Correlation |
| ATXN2L | -0.05935 | 0.31725 | Other Negative Correlation |
| CYP4X1 | -0.05941 | 0.316738 | Other Negative Correlation |
| ZNF649 | -0.05945 | 0.31642 | Other Negative Correlation |
| PTPDC1 | -0.05948 | 0.316162 | Other Negative Correlation |
| ETV5 | -0.05951 | 0.315955 | Other Negative Correlation |
| OR10AC1 | -0.05956 | 0.315514 | Other Negative Correlation |
| GNGT2 | -0.05963 | 0.314921 | Other Negative Correlation |
| ZNF570 | -0.05966 | 0.314718 | Other Negative Correlation |
| THTPA | -0.05976 | 0.313877 | Other Negative Correlation |
| CXXC4 | -0.05977 | 0.313807 | Other Negative Correlation |
| OR10H1 | -0.05978 | 0.313726 | Other Negative Correlation |
| CATSPERE | -0.05979 | 0.313626 | Other Negative Correlation |
| ENSG00000285629 | -0.05984 | 0.313199 | Other Negative Correlation |
| FATE1 | -0.05984 | 0.313199 | Other Negative Correlation |
| ZSWIM9 | -0.05991 | 0.312665 | Other Negative Correlation |
| SPTLC2 | -0.05991 | 0.31266 | Other Negative Correlation |
| WDR83OS | -0.05993 | 0.312516 | Other Negative Correlation |
| GIMAP8 | -0.05994 | 0.312398 | Other Negative Correlation |
| ITGA10 | -0.05996 | 0.312292 | Other Negative Correlation |
| ZNF107 | -0.05999 | 0.312007 | Other Negative Correlation |
| ARHGAP26 | -0.06001 | 0.311823 | Other Negative Correlation |
| BTBD9 | -0.06002 | 0.311756 | Other Negative Correlation |
| PXK | -0.06003 | 0.311693 | Other Negative Correlation |
| TESK2 | -0.06007 | 0.31139 | Other Negative Correlation |
| PRR5-ARHGAP8 | -0.06009 | 0.311189 | Other Negative Correlation |
| SLC4A3 | -0.06011 | 0.311055 | Other Negative Correlation |
| KANK4 | -0.06012 | 0.311001 | Other Negative Correlation |
| MYOCOS | -0.0602 | 0.310365 | Other Negative Correlation |
| ZC3H7A | -0.06021 | 0.310209 | Other Negative Correlation |

| | | | |
|---|---|---|---|
| GATAD1 | -0.06027 | 0.309745 | Other Negative Correlation |
| GLYATL1 | -0.06029 | 0.309627 | Other Negative Correlation |
| DDX3Y | -0.06029 | 0.309572 | Other Negative Correlation |
| APLF | -0.0603 | 0.309526 | Other Negative Correlation |
| C14orf39 | -0.06031 | 0.30943 | Other Negative Correlation |
| PPP2R5A | -0.06031 | 0.30941 | Other Negative Correlation |
| SOX30 | -0.06034 | 0.309225 | Other Negative Correlation |
| DYDC2 | -0.06037 | 0.30897 | Other Negative Correlation |
| DNALI1 | -0.06038 | 0.308863 | Other Negative Correlation |
| WFDC1 | -0.06043 | 0.308459 | Other Negative Correlation |
| PLAC9 | -0.06045 | 0.30832 | Other Negative Correlation |
| IL2RG | -0.06052 | 0.307728 | Other Negative Correlation |
| CGGBP1 | -0.06053 | 0.307705 | Other Negative Correlation |
| ISCA2 | -0.06054 | 0.307573 | Other Negative Correlation |
| RBAK-RBAKDN | -0.06056 | 0.307454 | Other Negative Correlation |
| CLEC18A | -0.06062 | 0.306994 | Other Negative Correlation |
| ADAMTS13 | -0.06065 | 0.306756 | Other Negative Correlation |
| ULBP3 | -0.06065 | 0.30669 | Other Negative Correlation |
| IDH2 | -0.06068 | 0.306516 | Other Negative Correlation |
| NPFFR1 | -0.06069 | 0.306432 | Other Negative Correlation |
| ZNF266 | -0.0607 | 0.306337 | Other Negative Correlation |
| ANGPT1 | -0.06075 | 0.305907 | Other Negative Correlation |
| WDR53 | -0.06077 | 0.305744 | Other Negative Correlation |
| ARNT2 | -0.06077 | 0.305734 | Other Negative Correlation |
| PPP1R27 | -0.06083 | 0.305262 | Other Negative Correlation |
| ENSG00000257207 | -0.06084 | 0.305177 | Other Negative Correlation |
| DIPK2A | -0.06087 | 0.304948 | Other Negative Correlation |
| UBIAD1 | -0.0609 | 0.304746 | Other Negative Correlation |
| DOCK2 | -0.06092 | 0.304582 | Other Negative Correlation |
| IGFN1 | -0.06101 | 0.303823 | Other Negative Correlation |
| SIK3 | -0.06103 | 0.30372 | Other Negative Correlation |
| MAOB | -0.06104 | 0.303588 | Other Negative Correlation |
| TOM1 | -0.06106 | 0.303448 | Other Negative Correlation |
| RBM23 | -0.06109 | 0.303203 | Other Negative Correlation |
| GPHA2 | -0.06116 | 0.302627 | Other Negative Correlation |
| TTLL9 | -0.06123 | 0.302139 | Other Negative Correlation |
| SMIM14 | -0.06125 | 0.301964 | Other Negative Correlation |
| NUTM2E | -0.06131 | 0.301464 | Other Negative Correlation |
| H1-8 | -0.06133 | 0.301319 | Other Negative Correlation |
| ENSG00000262633 | -0.06136 | 0.301094 | Other Negative Correlation |
| RPAP1 | -0.06138 | 0.30091 | Other Negative Correlation |
| DENND5B | -0.06139 | 0.300812 | Other Negative Correlation |
| ULK3 | -0.06141 | 0.300664 | Other Negative Correlation |
| FMNL3 | -0.06143 | 0.300542 | Other Negative Correlation |
| OXTR | -0.0615 | 0.299996 | Other Negative Correlation |
| ZNF615 | -0.06153 | 0.29974 | Other Negative Correlation |

| | | | |
|---|---|---|---|
| ENSG00000285253 | -0.06156 | 0.299499 | Other Negative Correlation |
| NTMT1 | -0.0616 | 0.299171 | Other Negative Correlation |
| MRNIP | -0.0616 | 0.299163 | Other Negative Correlation |
| MUC5AC | -0.06163 | 0.298984 | Other Negative Correlation |
| SIGLEC15 | -0.06163 | 0.298983 | Other Negative Correlation |
| SEC61A1 | -0.06167 | 0.298678 | Other Negative Correlation |
| OXGR1 | -0.06168 | 0.298526 | Other Negative Correlation |
| TMEM88 | -0.06178 | 0.297761 | Other Negative Correlation |
| GPR89A | -0.06179 | 0.297714 | Other Negative Correlation |
| YY1AP1 | -0.06182 | 0.297494 | Other Negative Correlation |
| ZNF510 | -0.06185 | 0.297236 | Other Negative Correlation |
| ZBED8 | -0.06185 | 0.297222 | Other Negative Correlation |
| DERL2 | -0.06186 | 0.297139 | Other Negative Correlation |
| CHRNA6 | -0.06187 | 0.297047 | Other Negative Correlation |
| CA11 | -0.06188 | 0.296963 | Other Negative Correlation |
| TRPV1 | -0.06192 | 0.296695 | Other Negative Correlation |
| LIN37 | -0.06192 | 0.296688 | Other Negative Correlation |
| AKAP3 | -0.06194 | 0.296502 | Other Negative Correlation |
| MIS18BP1 | -0.06201 | 0.295954 | Other Negative Correlation |
| BEAN1 | -0.06205 | 0.295661 | Other Negative Correlation |
| TMOD4 | -0.06211 | 0.295227 | Other Negative Correlation |
| SYCE3 | -0.06211 | 0.295172 | Other Negative Correlation |
| SRXN1 | -0.06213 | 0.295055 | Other Negative Correlation |
| CPO | -0.06213 | 0.295031 | Other Negative Correlation |
| EDN2 | -0.06217 | 0.294705 | Other Negative Correlation |
| ZNF776 | -0.06218 | 0.294627 | Other Negative Correlation |
| JADE3 | -0.06221 | 0.294388 | Other Negative Correlation |
| PFN4 | -0.06222 | 0.294341 | Other Negative Correlation |
| POLR2C | -0.06223 | 0.294279 | Other Negative Correlation |
| EDN3 | -0.06224 | 0.294161 | Other Negative Correlation |
| CCNB3 | -0.06225 | 0.294126 | Other Negative Correlation |
| ZNF302 | -0.06225 | 0.294076 | Other Negative Correlation |
| SCGB1A1 | -0.06229 | 0.293828 | Other Negative Correlation |
| C11orf52 | -0.06231 | 0.293647 | Other Negative Correlation |
| CCDC188 | -0.06238 | 0.293104 | Other Negative Correlation |
| PDE6G | -0.06249 | 0.292266 | Other Negative Correlation |
| PRORP | -0.0625 | 0.292193 | Other Negative Correlation |
| CCL7 | -0.0625 | 0.292148 | Other Negative Correlation |
| NSUN5 | -0.06252 | 0.292 | Other Negative Correlation |
| ZNF469 | -0.06253 | 0.291971 | Other Negative Correlation |
| SMYD4 | -0.06253 | 0.291941 | Other Negative Correlation |
| KRT37 | -0.06253 | 0.291909 | Other Negative Correlation |
| SERAC1 | -0.06257 | 0.291647 | Other Negative Correlation |
| POMZP3 | -0.06261 | 0.291314 | Other Negative Correlation |
| SLC35B4 | -0.06261 | 0.291284 | Other Negative Correlation |
| FEM1A | -0.06262 | 0.291213 | Other Negative Correlation |

| | | | |
|---|---|---|---|
| ACY3 | -0.06268 | 0.290751 | Other Negative Correlation |
| MON1A | -0.0627 | 0.290632 | Other Negative Correlation |
| MB | -0.06271 | 0.290576 | Other Negative Correlation |
| ZNF212 | -0.06276 | 0.290187 | Other Negative Correlation |
| DCAF1 | -0.06279 | 0.289933 | Other Negative Correlation |
| TONSL | -0.06281 | 0.289807 | Other Negative Correlation |
| PTPN23 | -0.06282 | 0.289719 | Other Negative Correlation |
| SERPINA5 | -0.06285 | 0.289503 | Other Negative Correlation |
| ATP2B3 | -0.06285 | 0.289503 | Other Negative Correlation |
| PCNX1 | -0.06285 | 0.289502 | Other Negative Correlation |
| HABP4 | -0.06287 | 0.28928 | Other Negative Correlation |
| PLXDC2 | -0.06296 | 0.288657 | Other Negative Correlation |
| KMT2B | -0.06302 | 0.288166 | Other Negative Correlation |
| ARRB1 | -0.06303 | 0.288098 | Other Negative Correlation |
| PRR12 | -0.06304 | 0.28799 | Other Negative Correlation |
| PTPN7 | -0.06305 | 0.28794 | Other Negative Correlation |
| GPR150 | -0.06311 | 0.287477 | Other Negative Correlation |
| BMP5 | -0.06311 | 0.287463 | Other Negative Correlation |
| IQCG | -0.06325 | 0.286369 | Other Negative Correlation |
| FANCD2OS | -0.06327 | 0.286232 | Other Negative Correlation |
| CLCN7 | -0.06341 | 0.285172 | Other Negative Correlation |
| BBS5 | -0.06342 | 0.285077 | Other Negative Correlation |
| CCDC106 | -0.06344 | 0.284969 | Other Negative Correlation |
| FAM90A1 | -0.06349 | 0.284615 | Other Negative Correlation |
| RIPPLY2 | -0.06354 | 0.284222 | Other Negative Correlation |
| C9orf72 | -0.06355 | 0.284153 | Other Negative Correlation |
| PRSS33 | -0.06356 | 0.28402 | Other Negative Correlation |
| CASC3 | -0.06359 | 0.283819 | Other Negative Correlation |
| WDR45 | -0.06361 | 0.28364 | Other Negative Correlation |
| GATA6 | -0.06362 | 0.283591 | Other Negative Correlation |
| CPNE2 | -0.06364 | 0.283465 | Other Negative Correlation |
| CIDEC | -0.06369 | 0.283078 | Other Negative Correlation |
| NMNAT2 | -0.06371 | 0.282907 | Other Negative Correlation |
| CD22 | -0.06373 | 0.28274 | Other Negative Correlation |
| TNFRSF13C | -0.06386 | 0.281807 | Other Negative Correlation |
| PAQR3 | -0.06389 | 0.281524 | Other Negative Correlation |
| GAS2 | -0.0639 | 0.281469 | Other Negative Correlation |
| PRSS27 | -0.06392 | 0.281344 | Other Negative Correlation |
| SMIM27 | -0.06397 | 0.280945 | Other Negative Correlation |
| HSD17B13 | -0.06397 | 0.280922 | Other Negative Correlation |
| GPN3 | -0.06399 | 0.280794 | Other Negative Correlation |
| TNK2 | -0.064 | 0.280708 | Other Negative Correlation |
| NPHP1 | -0.06406 | 0.280304 | Other Negative Correlation |
| TSR2 | -0.06407 | 0.280185 | Other Negative Correlation |
| MED14 | -0.0641 | 0.279994 | Other Negative Correlation |
| RAB37 | -0.06412 | 0.279811 | Other Negative Correlation |

| | | | |
|---|---|---|---|
| NLGN3 | -0.06416 | 0.279513 | Other Negative Correlation |
| UCK1 | -0.0642 | 0.279243 | Other Negative Correlation |
| RPS10-NUDT3 | -0.06424 | 0.278918 | Other Negative Correlation |
| SLC25A44 | -0.06424 | 0.278899 | Other Negative Correlation |
| MOGS | -0.06426 | 0.278797 | Other Negative Correlation |
| GOLPH3L | -0.06427 | 0.278658 | Other Negative Correlation |
| SULT1B1 | -0.06428 | 0.278617 | Other Negative Correlation |
| METTL27 | -0.06429 | 0.27856 | Other Negative Correlation |
| LIPT2 | -0.06431 | 0.278355 | Other Negative Correlation |
| GDF9 | -0.06436 | 0.277988 | Other Negative Correlation |
| OTUD1 | -0.06437 | 0.277963 | Other Negative Correlation |
| SLC46A1 | -0.0644 | 0.277708 | Other Negative Correlation |
| DNAH10 | -0.06441 | 0.277626 | Other Negative Correlation |
| TRPV4 | -0.06443 | 0.277473 | Other Negative Correlation |
| TTLL1 | -0.06445 | 0.277316 | Other Negative Correlation |
| CIB3 | -0.06449 | 0.277028 | Other Negative Correlation |
| KRT74 | -0.06454 | 0.276644 | Other Negative Correlation |
| TRIM74 | -0.06454 | 0.276644 | Other Negative Correlation |
| ZNF506 | -0.06457 | 0.276463 | Other Negative Correlation |
| EOLA2 | -0.06457 | 0.27646 | Other Negative Correlation |
| SPAG4 | -0.06464 | 0.275952 | Other Negative Correlation |
| ZDHHC9 | -0.06466 | 0.275778 | Other Negative Correlation |
| HSF2 | -0.0647 | 0.275472 | Other Negative Correlation |
| ITGAM | -0.06472 | 0.275354 | Other Negative Correlation |
| CLPSL2 | -0.06476 | 0.275015 | Other Negative Correlation |
| CCN5 | -0.06478 | 0.274892 | Other Negative Correlation |
| RNF152 | -0.06479 | 0.274825 | Other Negative Correlation |
| C2orf73 | -0.06483 | 0.274517 | Other Negative Correlation |
| MYPN | -0.06484 | 0.274406 | Other Negative Correlation |
| IFT122 | -0.06485 | 0.27437 | Other Negative Correlation |
| UCN3 | -0.06487 | 0.2742 | Other Negative Correlation |
| VNN2 | -0.06487 | 0.2742 | Other Negative Correlation |
| CCL22 | -0.06488 | 0.274152 | Other Negative Correlation |
| SGSH | -0.06492 | 0.273853 | Other Negative Correlation |
| WNT16 | -0.06493 | 0.273754 | Other Negative Correlation |
| NPTX1 | -0.06496 | 0.273588 | Other Negative Correlation |
| PSAPL1 | -0.06497 | 0.27346 | Other Negative Correlation |
| MBLAC1 | -0.06502 | 0.2731 | Other Negative Correlation |
| PCDH12 | -0.06504 | 0.272993 | Other Negative Correlation |
| STYK1 | -0.06508 | 0.272683 | Other Negative Correlation |
| EXOSC4 | -0.06508 | 0.27266 | Other Negative Correlation |
| CHP2 | -0.06508 | 0.272635 | Other Negative Correlation |
| RCE1 | -0.06509 | 0.272574 | Other Negative Correlation |
| PFKFB1 | -0.0651 | 0.272521 | Other Negative Correlation |
| SLC1A7 | -0.0651 | 0.272521 | Other Negative Correlation |
| PRB3 | -0.0651 | 0.272521 | Other Negative Correlation |

| | | | |
|---|---|---|---|
| ARVCF | -0.06519 | 0.271837 | Other Negative Correlation |
| NTN1 | -0.0652 | 0.27179 | Other Negative Correlation |
| MAST3 | -0.0652 | 0.271783 | Other Negative Correlation |
| CELA3B | -0.06521 | 0.271705 | Other Negative Correlation |
| TAFA2 | -0.06523 | 0.271573 | Other Negative Correlation |
| GOLT1A | -0.06525 | 0.271439 | Other Negative Correlation |
| AKAP17A | -0.06526 | 0.271372 | Other Negative Correlation |
| POMT1 | -0.06528 | 0.271227 | Other Negative Correlation |
| ZNF527 | -0.06528 | 0.271162 | Other Negative Correlation |
| RNF151 | -0.06531 | 0.270963 | Other Negative Correlation |
| ZNF667 | -0.06532 | 0.270866 | Other Negative Correlation |
| SPATA5 | -0.06534 | 0.270777 | Other Negative Correlation |
| EGLN1 | -0.06534 | 0.270756 | Other Negative Correlation |
| HPS4 | -0.06539 | 0.270406 | Other Negative Correlation |
| SCMH1 | -0.0654 | 0.27028 | Other Negative Correlation |
| DENND6B | -0.06541 | 0.270251 | Other Negative Correlation |
| PGPEP1L | -0.06543 | 0.27008 | Other Negative Correlation |
| ARMS2 | -0.06543 | 0.27008 | Other Negative Correlation |
| CTHRC1 | -0.06545 | 0.269919 | Other Negative Correlation |
| KRTAP5-8 | -0.06548 | 0.26971 | Other Negative Correlation |
| TMEM82 | -0.06548 | 0.26971 | Other Negative Correlation |
| ACSM2B | -0.06548 | 0.26971 | Other Negative Correlation |
| TSNAX-DISC1 | -0.06548 | 0.26971 | Other Negative Correlation |
| DDX19A | -0.06549 | 0.269638 | Other Negative Correlation |
| SYT2 | -0.06551 | 0.269481 | Other Negative Correlation |
| CCP110 | -0.06552 | 0.269466 | Other Negative Correlation |
| HNRNPUL2 | -0.06553 | 0.269364 | Other Negative Correlation |
| SLC18B1 | -0.06553 | 0.26932 | Other Negative Correlation |
| EIF2B5 | -0.06554 | 0.269308 | Other Negative Correlation |
| ZBTB39 | -0.06559 | 0.26895 | Other Negative Correlation |
| PPM1M | -0.06559 | 0.268903 | Other Negative Correlation |
| SPRYD3 | -0.06561 | 0.268757 | Other Negative Correlation |
| PLPP2 | -0.06562 | 0.268674 | Other Negative Correlation |
| PROC | -0.0657 | 0.268103 | Other Negative Correlation |
| CCR10 | -0.06571 | 0.268027 | Other Negative Correlation |
| LIN7B | -0.06571 | 0.268021 | Other Negative Correlation |
| C6orf15 | -0.06574 | 0.267805 | Other Negative Correlation |
| RGMA | -0.06574 | 0.267786 | Other Negative Correlation |
| NRG4 | -0.06579 | 0.267477 | Other Negative Correlation |
| DRAXIN | -0.06579 | 0.267427 | Other Negative Correlation |
| ZNF182 | -0.06582 | 0.267259 | Other Negative Correlation |
| HOATZ | -0.06584 | 0.267064 | Other Negative Correlation |
| SUPT7L | -0.06588 | 0.266778 | Other Negative Correlation |
| ZNF572 | -0.06589 | 0.266753 | Other Negative Correlation |
| PPP1R1B | -0.06589 | 0.266747 | Other Negative Correlation |
| SIRPA | -0.0659 | 0.266659 | Other Negative Correlation |

| | | | |
|---|---|---|---|
| TTC13 | -0.0659 | 0.266635 | Other Negative Correlation |
| NOTCH4 | -0.06593 | 0.266469 | Other Negative Correlation |
| CCR1 | -0.06594 | 0.26639 | Other Negative Correlation |
| C18orf54 | -0.06597 | 0.266163 | Other Negative Correlation |
| KRTAP3-1 | -0.06599 | 0.265994 | Other Negative Correlation |
| KRT20 | -0.06599 | 0.265994 | Other Negative Correlation |
| KRT12 | -0.06599 | 0.265994 | Other Negative Correlation |
| CLYBL | -0.06602 | 0.265793 | Other Negative Correlation |
| ARHGEF2 | -0.06602 | 0.265792 | Other Negative Correlation |
| RAB5B | -0.06604 | 0.265663 | Other Negative Correlation |
| ADAT3 | -0.06605 | 0.265591 | Other Negative Correlation |
| RC3H1 | -0.06607 | 0.265446 | Other Negative Correlation |
| CSRNP2 | -0.06612 | 0.265058 | Other Negative Correlation |
| FANCL | -0.06612 | 0.265045 | Other Negative Correlation |
| C2orf69 | -0.06619 | 0.264519 | Other Negative Correlation |
| FBXW11 | -0.06623 | 0.264298 | Other Negative Correlation |
| RGS22 | -0.06625 | 0.264131 | Other Negative Correlation |
| BDH1 | -0.06627 | 0.264003 | Other Negative Correlation |
| C2orf66 | -0.06627 | 0.263992 | Other Negative Correlation |
| PRR20G | -0.06628 | 0.263905 | Other Negative Correlation |
| CFAP97D1 | -0.0663 | 0.263752 | Other Negative Correlation |
| OLAH | -0.06631 | 0.263679 | Other Negative Correlation |
| NPW | -0.06644 | 0.262747 | Other Negative Correlation |
| CENPVL3 | -0.06644 | 0.262732 | Other Negative Correlation |
| HLCS | -0.06646 | 0.262597 | Other Negative Correlation |
| REV3L | -0.06646 | 0.262576 | Other Negative Correlation |
| COA1 | -0.06651 | 0.262225 | Other Negative Correlation |
| RPGRIP1 | -0.06653 | 0.26211 | Other Negative Correlation |
| TPBGL | -0.06654 | 0.262045 | Other Negative Correlation |
| NUDT2 | -0.06655 | 0.261932 | Other Negative Correlation |
| WDR37 | -0.06659 | 0.261699 | Other Negative Correlation |
| MCTP2 | -0.0666 | 0.261616 | Other Negative Correlation |
| HEMK1 | -0.06666 | 0.261186 | Other Negative Correlation |
| DACT2 | -0.06667 | 0.261121 | Other Negative Correlation |
| ZNF365 | -0.06667 | 0.261073 | Other Negative Correlation |
| RNASEK | -0.06669 | 0.260968 | Other Negative Correlation |
| MAPK8 | -0.06671 | 0.260842 | Other Negative Correlation |
| CCIN | -0.06672 | 0.260706 | Other Negative Correlation |
| OLR1 | -0.06674 | 0.260566 | Other Negative Correlation |
| GEMIN2 | -0.06676 | 0.26048 | Other Negative Correlation |
| BRF1 | -0.06676 | 0.260423 | Other Negative Correlation |
| DNAJC28 | -0.06677 | 0.260345 | Other Negative Correlation |
| INTU | -0.0668 | 0.260172 | Other Negative Correlation |
| YPEL3 | -0.06686 | 0.25977 | Other Negative Correlation |
| NYAP1 | -0.0669 | 0.259474 | Other Negative Correlation |
| ABCA6 | -0.0669 | 0.25942 | Other Negative Correlation |

| | | | |
|---|---|---|---|
| ZNF784 | -0.06697 | 0.258983 | Other Negative Correlation |
| RBM48 | -0.06698 | 0.258912 | Other Negative Correlation |
| MRPL38 | -0.06699 | 0.258801 | Other Negative Correlation |
| SLC25A21 | -0.06702 | 0.258583 | Other Negative Correlation |
| FBXO44 | -0.06707 | 0.258265 | Other Negative Correlation |
| TACO1 | -0.06711 | 0.257976 | Other Negative Correlation |
| SYT15B | -0.06712 | 0.257885 | Other Negative Correlation |
| RFX2 | -0.06715 | 0.257698 | Other Negative Correlation |
| TBC1D8B | -0.06715 | 0.257667 | Other Negative Correlation |
| TBXAS1 | -0.06717 | 0.257535 | Other Negative Correlation |
| FBXO42 | -0.06724 | 0.257032 | Other Negative Correlation |
| ZBTB7B | -0.06727 | 0.256829 | Other Negative Correlation |
| ZNF385D | -0.06734 | 0.256319 | Other Negative Correlation |
| GALNT13 | -0.06739 | 0.255988 | Other Negative Correlation |
| ZNF280C | -0.0674 | 0.255899 | Other Negative Correlation |
| FAM53C | -0.06748 | 0.255347 | Other Negative Correlation |
| KRT222 | -0.06749 | 0.255276 | Other Negative Correlation |
| PLEKHA3 | -0.0675 | 0.255213 | Other Negative Correlation |
| ZAN | -0.06751 | 0.255137 | Other Negative Correlation |
| ACOD1 | -0.06751 | 0.255137 | Other Negative Correlation |
| ZNF414 | -0.06751 | 0.255108 | Other Negative Correlation |
| SLC6A13 | -0.06752 | 0.255064 | Other Negative Correlation |
| GPR183 | -0.06754 | 0.254941 | Other Negative Correlation |
| TMEM249 | -0.0676 | 0.25446 | Other Negative Correlation |
| CYP2J2 | -0.06765 | 0.254104 | Other Negative Correlation |
| MAF1 | -0.06769 | 0.253884 | Other Negative Correlation |
| TNK1 | -0.06769 | 0.253859 | Other Negative Correlation |
| KRT2 | -0.06772 | 0.253664 | Other Negative Correlation |
| AZIN1 | -0.06775 | 0.253435 | Other Negative Correlation |
| RCN2 | -0.06775 | 0.253406 | Other Negative Correlation |
| ENSG00000256029 | -0.06778 | 0.253207 | Other Negative Correlation |
| ZNRF3 | -0.0678 | 0.253058 | Other Negative Correlation |
| PCDHA1 | -0.06786 | 0.252629 | Other Negative Correlation |
| ST8SIA1 | -0.06788 | 0.252538 | Other Negative Correlation |
| RTN4IP1 | -0.06789 | 0.252419 | Other Negative Correlation |
| CCDC28A | -0.06789 | 0.252417 | Other Negative Correlation |
| SLC39A2 | -0.06793 | 0.252176 | Other Negative Correlation |
| PRMT9 | -0.06795 | 0.252029 | Other Negative Correlation |
| ZC3H11B | -0.06797 | 0.251903 | Other Negative Correlation |
| BMF | -0.06798 | 0.251851 | Other Negative Correlation |
| RNF38 | -0.068 | 0.251713 | Other Negative Correlation |
| POLDIP3 | -0.06809 | 0.251041 | Other Negative Correlation |
| ALDH1A1 | -0.0681 | 0.250995 | Other Negative Correlation |
| BEND3 | -0.06811 | 0.250893 | Other Negative Correlation |
| IQSEC2 | -0.06818 | 0.250453 | Other Negative Correlation |
| ANKRD39 | -0.06819 | 0.250361 | Other Negative Correlation |

| | | | |
|---|---|---|---|
| TSPAN12 | -0.06822 | 0.250178 | Other Negative Correlation |
| LRRC66 | -0.06825 | 0.24997 | Other Negative Correlation |
| CA2 | -0.06825 | 0.249938 | Other Negative Correlation |
| POU5F1 | -0.06827 | 0.249783 | Other Negative Correlation |
| CLDN8 | -0.06831 | 0.249517 | Other Negative Correlation |
| FAM209A | -0.0684 | 0.248863 | Other Negative Correlation |
| ZNF429 | -0.06841 | 0.248819 | Other Negative Correlation |
| DYRK1A | -0.06842 | 0.248791 | Other Negative Correlation |
| PLD6 | -0.06843 | 0.24866 | Other Negative Correlation |
| NUPR1 | -0.06845 | 0.248579 | Other Negative Correlation |
| LIMD2 | -0.06845 | 0.248531 | Other Negative Correlation |
| SPATS1 | -0.06845 | 0.248521 | Other Negative Correlation |
| ERP44 | -0.06848 | 0.248371 | Other Negative Correlation |
| CCDC142 | -0.0685 | 0.248196 | Other Negative Correlation |
| C4orf36 | -0.06859 | 0.247599 | Other Negative Correlation |
| AK1 | -0.0686 | 0.247544 | Other Negative Correlation |
| ZFYVE9 | -0.06872 | 0.246664 | Other Negative Correlation |
| ENSG00000255339 | -0.06878 | 0.246237 | Other Negative Correlation |
| PIK3C2A | -0.06887 | 0.245674 | Other Negative Correlation |
| PHETA2 | -0.06888 | 0.245611 | Other Negative Correlation |
| TMEM229B | -0.06892 | 0.24534 | Other Negative Correlation |
| EQTN | -0.06892 | 0.245309 | Other Negative Correlation |
| KDR | -0.06895 | 0.245104 | Other Negative Correlation |
| DPM3 | -0.06895 | 0.245096 | Other Negative Correlation |
| KIAA0825 | -0.06907 | 0.244286 | Other Negative Correlation |
| CTNNA3 | -0.06923 | 0.243194 | Other Negative Correlation |
| TNS1 | -0.06924 | 0.243114 | Other Negative Correlation |
| ALG12 | -0.06935 | 0.242401 | Other Negative Correlation |
| SPRR5 | -0.06935 | 0.242341 | Other Negative Correlation |
| LPCAT1 | -0.06937 | 0.242203 | Other Negative Correlation |
| ZNF883 | -0.06942 | 0.2419 | Other Negative Correlation |
| GATAD2B | -0.06943 | 0.241838 | Other Negative Correlation |
| LHX5 | -0.06943 | 0.241838 | Other Negative Correlation |
| TAT | -0.06949 | 0.241415 | Other Negative Correlation |
| STK36 | -0.06949 | 0.241399 | Other Negative Correlation |
| CORIN | -0.06949 | 0.24139 | Other Negative Correlation |
| FAM234A | -0.06951 | 0.241299 | Other Negative Correlation |
| SMIM4 | -0.06956 | 0.24097 | Other Negative Correlation |
| ZBTB44 | -0.06957 | 0.240903 | Other Negative Correlation |
| SLC25A28 | -0.06958 | 0.240822 | Other Negative Correlation |
| ROBO3 | -0.06958 | 0.240816 | Other Negative Correlation |
| ZNF283 | -0.06963 | 0.240471 | Other Negative Correlation |
| CFAP157 | -0.06966 | 0.240287 | Other Negative Correlation |
| ZCCHC7 | -0.06966 | 0.240262 | Other Negative Correlation |
| SGSM3 | -0.06968 | 0.240134 | Other Negative Correlation |
| PADI4 | -0.06973 | 0.239805 | Other Negative Correlation |

| | | | |
|---|---|---|---|
| CSDC2 | -0.06979 | 0.239371 | Other Negative Correlation |
| ERV3-1 | -0.0698 | 0.239286 | Other Negative Correlation |
| RPS6KA5 | -0.06983 | 0.239136 | Other Negative Correlation |
| TNFSF18 | -0.06992 | 0.238512 | Other Negative Correlation |
| CACNB3 | -0.06992 | 0.238499 | Other Negative Correlation |
| FBXL12 | -0.06994 | 0.238345 | Other Negative Correlation |
| MEGF8 | -0.07004 | 0.237732 | Other Negative Correlation |
| GRIP1 | -0.07005 | 0.237656 | Other Negative Correlation |
| NME8 | -0.07007 | 0.23753 | Other Negative Correlation |
| VPS8 | -0.07008 | 0.237463 | Other Negative Correlation |
| ZNF514 | -0.07008 | 0.237409 | Other Negative Correlation |
| CASKIN2 | -0.07014 | 0.23706 | Other Negative Correlation |
| DCAF15 | -0.07017 | 0.236826 | Other Negative Correlation |
| NUPR2 | -0.07019 | 0.236724 | Other Negative Correlation |
| KDF1 | -0.07019 | 0.236712 | Other Negative Correlation |
| KLRG2 | -0.0702 | 0.236604 | Other Negative Correlation |
| DNAH1 | -0.07021 | 0.236554 | Other Negative Correlation |
| FCER1G | -0.07024 | 0.236358 | Other Negative Correlation |
| MYZAP | -0.07026 | 0.236251 | Other Negative Correlation |
| ENSG00000289701 | -0.07035 | 0.235639 | Other Negative Correlation |
| PITPNM2 | -0.07037 | 0.235495 | Other Negative Correlation |
| ATP6V0A4 | -0.07042 | 0.23513 | Other Negative Correlation |
| OGFOD1 | -0.07049 | 0.234707 | Other Negative Correlation |
| HIF3A | -0.0705 | 0.234659 | Other Negative Correlation |
| CIC | -0.07051 | 0.234577 | Other Negative Correlation |
| IL12B | -0.07056 | 0.234255 | Other Negative Correlation |
| HINFP | -0.07057 | 0.234195 | Other Negative Correlation |
| PDE2A | -0.07059 | 0.234032 | Other Negative Correlation |
| PCYOX1L | -0.07062 | 0.233838 | Other Negative Correlation |
| LSP1 | -0.07063 | 0.233753 | Other Negative Correlation |
| SYCP3 | -0.07067 | 0.233523 | Other Negative Correlation |
| GLIPR1L2 | -0.07069 | 0.233394 | Other Negative Correlation |
| CFAP58 | -0.07071 | 0.233242 | Other Negative Correlation |
| PFKFB4 | -0.07071 | 0.233225 | Other Negative Correlation |
| XKR6 | -0.07071 | 0.233217 | Other Negative Correlation |
| PPT2 | -0.0708 | 0.232641 | Other Negative Correlation |
| CCDC171 | -0.0708 | 0.232618 | Other Negative Correlation |
| ALAD | -0.0708 | 0.232611 | Other Negative Correlation |
| PSPN | -0.07085 | 0.232311 | Other Negative Correlation |
| IGFBP1 | -0.07085 | 0.232283 | Other Negative Correlation |
| B4GALNT2 | -0.07092 | 0.231863 | Other Negative Correlation |
| PRKCE | -0.07092 | 0.231813 | Other Negative Correlation |
| THAP6 | -0.07093 | 0.231796 | Other Negative Correlation |
| SEC24A | -0.07095 | 0.231634 | Other Negative Correlation |
| SETD1B | -0.07095 | 0.23163 | Other Negative Correlation |
| IL17REL | -0.07095 | 0.231623 | Other Negative Correlation |

| | | | |
|---|---|---|---|
| INO80B | -0.07097 | 0.231501 | Other Negative Correlation |
| C17orf97 | -0.07098 | 0.23143 | Other Negative Correlation |
| CD2 | -0.07099 | 0.231371 | Other Negative Correlation |
| TFEC | -0.07099 | 0.231369 | Other Negative Correlation |
| RALGDS | -0.07101 | 0.231263 | Other Negative Correlation |
| YY2 | -0.07108 | 0.230806 | Other Negative Correlation |
| RAB43 | -0.07112 | 0.230546 | Other Negative Correlation |
| ARID3B | -0.07113 | 0.230442 | Other Negative Correlation |
| DNAI7 | -0.07119 | 0.230071 | Other Negative Correlation |
| ITGA1 | -0.07121 | 0.229956 | Other Negative Correlation |
| SIRT7 | -0.07126 | 0.229579 | Other Negative Correlation |
| ZPBP2 | -0.07135 | 0.229034 | Other Negative Correlation |
| HMBOX1 | -0.07135 | 0.229025 | Other Negative Correlation |
| BLACAT1 | -0.07137 | 0.228897 | Other Negative Correlation |
| C11orf80 | -0.07138 | 0.228848 | Other Negative Correlation |
| PARVG | -0.07139 | 0.228785 | Other Negative Correlation |
| BICRAL | -0.07139 | 0.228774 | Other Negative Correlation |
| MOAP1 | -0.07143 | 0.228511 | Other Negative Correlation |
| AHSG | -0.07145 | 0.22839 | Other Negative Correlation |
| UBE3D | -0.07148 | 0.228142 | Other Negative Correlation |
| SYN3 | -0.07152 | 0.227903 | Other Negative Correlation |
| PTPN20 | -0.07152 | 0.227889 | Other Negative Correlation |
| MCM3AP | -0.07153 | 0.22786 | Other Negative Correlation |
| DHX16 | -0.07155 | 0.227743 | Other Negative Correlation |
| UBXN11 | -0.07156 | 0.227634 | Other Negative Correlation |
| KLHL12 | -0.07157 | 0.227591 | Other Negative Correlation |
| ST8SIA4 | -0.07158 | 0.227537 | Other Negative Correlation |
| KAT6B | -0.07159 | 0.227466 | Other Negative Correlation |
| GRK6 | -0.07159 | 0.227426 | Other Negative Correlation |
| KIAA1328 | -0.0716 | 0.227409 | Other Negative Correlation |
| KIF6 | -0.07165 | 0.227055 | Other Negative Correlation |
| SLC30A4 | -0.07166 | 0.227024 | Other Negative Correlation |
| PDE3A | -0.07168 | 0.22687 | Other Negative Correlation |
| GDAP2 | -0.0717 | 0.226718 | Other Negative Correlation |
| TMEM191B | -0.07171 | 0.226692 | Other Negative Correlation |
| WASHC1 | -0.07176 | 0.226345 | Other Negative Correlation |
| EVI5L | -0.07177 | 0.226299 | Other Negative Correlation |
| ACVR1C | -0.07178 | 0.226195 | Other Negative Correlation |
| KATNIP | -0.07182 | 0.225952 | Other Negative Correlation |
| ZNF768 | -0.07183 | 0.225924 | Other Negative Correlation |
| TTLL11 | -0.07183 | 0.225889 | Other Negative Correlation |
| NPR3 | -0.07183 | 0.225863 | Other Negative Correlation |
| ADAMTSL3 | -0.07188 | 0.225593 | Other Negative Correlation |
| PRAP1 | -0.07198 | 0.224904 | Other Negative Correlation |
| PPOX | -0.072 | 0.224812 | Other Negative Correlation |
| RFWD3 | -0.07203 | 0.224612 | Other Negative Correlation |

| | | | |
|---|---|---|---|
| CHN2 | -0.07204 | 0.22451 | Other Negative Correlation |
| ABHD4 | -0.07206 | 0.224436 | Other Negative Correlation |
| CCDC191 | -0.07207 | 0.224344 | Other Negative Correlation |
| NLGN4X | -0.07207 | 0.224336 | Other Negative Correlation |
| TLN2 | -0.07209 | 0.22419 | Other Negative Correlation |
| NAALADL1 | -0.07211 | 0.224097 | Other Negative Correlation |
| C9orf64 | -0.07216 | 0.223766 | Other Negative Correlation |
| NT5C3B | -0.07218 | 0.223642 | Other Negative Correlation |
| ALDH8A1 | -0.07218 | 0.223633 | Other Negative Correlation |
| ADAM12 | -0.07227 | 0.223069 | Other Negative Correlation |
| SYT14 | -0.07229 | 0.222944 | Other Negative Correlation |
| PLA1A | -0.07237 | 0.222411 | Other Negative Correlation |
| ENSG00000285982 | -0.07237 | 0.222411 | Other Negative Correlation |
| RAB3C | -0.07237 | 0.222411 | Other Negative Correlation |
| ZNF790 | -0.07245 | 0.221922 | Other Negative Correlation |
| ARL15 | -0.07252 | 0.221469 | Other Negative Correlation |
| GPRIN1 | -0.07261 | 0.220909 | Other Negative Correlation |
| LYPD2 | -0.07263 | 0.22077 | Other Negative Correlation |
| PPM1K | -0.07264 | 0.220663 | Other Negative Correlation |
| CASP2 | -0.07268 | 0.220441 | Other Negative Correlation |
| SFXN5 | -0.07269 | 0.220373 | Other Negative Correlation |
| SEC31B | -0.07272 | 0.220213 | Other Negative Correlation |
| FZD4 | -0.07275 | 0.219973 | Other Negative Correlation |
| BAZ1B | -0.07279 | 0.219732 | Other Negative Correlation |
| HOOK1 | -0.07284 | 0.219427 | Other Negative Correlation |
| POPDC3 | -0.07285 | 0.219352 | Other Negative Correlation |
| ZNF284 | -0.07287 | 0.219237 | Other Negative Correlation |
| LSM11 | -0.07287 | 0.219221 | Other Negative Correlation |
| MXRA5 | -0.07287 | 0.219208 | Other Negative Correlation |
| KCNV2 | -0.07288 | 0.219171 | Other Negative Correlation |
| DNAJC18 | -0.07288 | 0.219153 | Other Negative Correlation |
| STARD6 | -0.07288 | 0.219136 | Other Negative Correlation |
| NCMAP | -0.07293 | 0.21884 | Other Negative Correlation |
| ARHGEF10 | -0.07294 | 0.21878 | Other Negative Correlation |
| PCDHGA4 | -0.07296 | 0.218651 | Other Negative Correlation |
| PPP1R14D | -0.07299 | 0.218445 | Other Negative Correlation |
| FAM110D | -0.073 | 0.218397 | Other Negative Correlation |
| SWT1 | -0.07301 | 0.218348 | Other Negative Correlation |
| EFCAB12 | -0.07305 | 0.218097 | Other Negative Correlation |
| MSANTD7 | -0.0731 | 0.2178 | Other Negative Correlation |
| SRY | -0.0731 | 0.217765 | Other Negative Correlation |
| INTS3 | -0.07316 | 0.217375 | Other Negative Correlation |
| PTK2B | -0.07319 | 0.217227 | Other Negative Correlation |
| RECK | -0.07327 | 0.216719 | Other Negative Correlation |
| LATS1 | -0.07329 | 0.216595 | Other Negative Correlation |
| PPP3CB | -0.07334 | 0.216272 | Other Negative Correlation |

| | | | |
|---|---|---|---|
| USHBP1 | -0.07336 | 0.21614 | Other Negative Correlation |
| MYNN | -0.07338 | 0.21601 | Other Negative Correlation |
| PDZRN3 | -0.0734 | 0.215873 | Other Negative Correlation |
| ZNF57 | -0.07342 | 0.215754 | Other Negative Correlation |
| ZNF304 | -0.07345 | 0.215552 | Other Negative Correlation |
| CHKB | -0.07347 | 0.215433 | Other Negative Correlation |
| PLAAT2 | -0.0735 | 0.215237 | Other Negative Correlation |
| TEKT5 | -0.07351 | 0.215184 | Other Negative Correlation |
| SYTL4 | -0.07351 | 0.215175 | Other Negative Correlation |
| ST8SIA6 | -0.07355 | 0.214964 | Other Negative Correlation |
| MLLT10 | -0.07356 | 0.214871 | Other Negative Correlation |
| APOBR | -0.07356 | 0.21486 | Other Negative Correlation |
| KCNQ1 | -0.07361 | 0.214565 | Other Negative Correlation |
| TIMM8A | -0.07362 | 0.214529 | Other Negative Correlation |
| FAAH2 | -0.07362 | 0.214518 | Other Negative Correlation |
| FAM167B | -0.07362 | 0.214507 | Other Negative Correlation |
| CENPC | -0.07363 | 0.214452 | Other Negative Correlation |
| PCDHGB7 | -0.07366 | 0.214244 | Other Negative Correlation |
| NUP210L | -0.07373 | 0.213804 | Other Negative Correlation |
| ZSWIM1 | -0.07381 | 0.213349 | Other Negative Correlation |
| USP51 | -0.07385 | 0.213093 | Other Negative Correlation |
| CUL7 | -0.07386 | 0.213022 | Other Negative Correlation |
| BEST3 | -0.07393 | 0.212561 | Other Negative Correlation |
| NTRK2 | -0.07402 | 0.212039 | Other Negative Correlation |
| PRDM10 | -0.07407 | 0.211727 | Other Negative Correlation |
| TMCC2 | -0.07408 | 0.211639 | Other Negative Correlation |
| CNKSR1 | -0.07413 | 0.211373 | Other Negative Correlation |
| MMP19 | -0.07414 | 0.211287 | Other Negative Correlation |
| BICDL2 | -0.07418 | 0.211053 | Other Negative Correlation |
| TUG1 | -0.0743 | 0.210282 | Other Negative Correlation |
| COL5A1 | -0.07438 | 0.209793 | Other Negative Correlation |
| PIK3C2G | -0.07442 | 0.209558 | Other Negative Correlation |
| TMEM191C | -0.07443 | 0.209501 | Other Negative Correlation |
| SEC14L4 | -0.07443 | 0.209495 | Other Negative Correlation |
| TJP3 | -0.07443 | 0.209481 | Other Negative Correlation |
| PDE6D | -0.07445 | 0.209392 | Other Negative Correlation |
| ALG6 | -0.07446 | 0.209295 | Other Negative Correlation |
| PLIN2 | -0.07447 | 0.209234 | Other Negative Correlation |
| TSC2 | -0.07448 | 0.209205 | Other Negative Correlation |
| TSHZ1 | -0.0745 | 0.20904 | Other Negative Correlation |
| CDX2 | -0.07454 | 0.208821 | Other Negative Correlation |
| FUT8 | -0.07454 | 0.208802 | Other Negative Correlation |
| TOP3B | -0.07456 | 0.20871 | Other Negative Correlation |
| SIX2 | -0.07456 | 0.208696 | Other Negative Correlation |
| ADAP2 | -0.07457 | 0.208644 | Other Negative Correlation |
| DAPK2 | -0.0746 | 0.208441 | Other Negative Correlation |

| | | | |
|---|---|---|---|
| ZNF445 | -0.07467 | 0.208052 | Other Negative Correlation |
| SPRR4 | -0.07469 | 0.207907 | Other Negative Correlation |
| PIGBOS1 | -0.07472 | 0.207727 | Other Negative Correlation |
| DBP | -0.07473 | 0.207688 | Other Negative Correlation |
| RBBP5 | -0.07473 | 0.207677 | Other Negative Correlation |
| DUSP18 | -0.07473 | 0.207631 | Other Negative Correlation |
| ITGBL1 | -0.07474 | 0.207618 | Other Negative Correlation |
| NR2C2 | -0.07481 | 0.207154 | Other Negative Correlation |
| IFT80 | -0.07485 | 0.206912 | Other Negative Correlation |
| ZNF639 | -0.07497 | 0.206226 | Other Negative Correlation |
| GPR20 | -0.07501 | 0.205934 | Other Negative Correlation |
| UNKL | -0.07503 | 0.205853 | Other Negative Correlation |
| EEFSEC | -0.0751 | 0.205391 | Other Negative Correlation |
| IFT46 | -0.07511 | 0.205358 | Other Negative Correlation |
| PCDHA12 | -0.07512 | 0.205309 | Other Negative Correlation |
| ETV3L | -0.07514 | 0.205156 | Other Negative Correlation |
| AHCYL2 | -0.0752 | 0.204803 | Other Negative Correlation |
| COG3 | -0.07525 | 0.204476 | Other Negative Correlation |
| C9orf50 | -0.07527 | 0.204356 | Other Negative Correlation |
| CNGB1 | -0.07529 | 0.20427 | Other Negative Correlation |
| TIMP2 | -0.0753 | 0.204229 | Other Negative Correlation |
| PCM1 | -0.07533 | 0.204013 | Other Negative Correlation |
| HCFC1 | -0.07539 | 0.203675 | Other Negative Correlation |
| UNK | -0.07541 | 0.203568 | Other Negative Correlation |
| PPP1R3D | -0.07541 | 0.203562 | Other Negative Correlation |
| ARL16 | -0.07541 | 0.203517 | Other Negative Correlation |
| PTOV1 | -0.07546 | 0.203221 | Other Negative Correlation |
| CEP19 | -0.07547 | 0.203183 | Other Negative Correlation |
| C1S | -0.07552 | 0.202877 | Other Negative Correlation |
| PDGFD | -0.07558 | 0.202549 | Other Negative Correlation |
| UBALD2 | -0.0756 | 0.202396 | Other Negative Correlation |
| BRPF1 | -0.07567 | 0.201968 | Other Negative Correlation |
| HLA-DMA | -0.07568 | 0.201924 | Other Negative Correlation |
| NOTCH2NLR | -0.07571 | 0.201757 | Other Negative Correlation |
| TBX1 | -0.07571 | 0.201734 | Other Negative Correlation |
| IDH1 | -0.07573 | 0.201652 | Other Negative Correlation |
| MICU1 | -0.07574 | 0.201557 | Other Negative Correlation |
| AGO1 | -0.07579 | 0.201269 | Other Negative Correlation |
| KRT33B | -0.0758 | 0.201218 | Other Negative Correlation |
| FKBP4 | -0.07582 | 0.20111 | Other Negative Correlation |
| RDH13 | -0.07583 | 0.201018 | Other Negative Correlation |
| SHISA9 | -0.07588 | 0.200715 | Other Negative Correlation |
| PPM1A | -0.07589 | 0.200644 | Other Negative Correlation |
| ZNF383 | -0.07591 | 0.200527 | Other Negative Correlation |
| RGS17 | -0.07594 | 0.200386 | Other Negative Correlation |
| ZNF668 | -0.07597 | 0.20022 | Other Negative Correlation |

| | | | |
|---|---|---|---|
| SEL1L3 | -0.07598 | 0.200158 | Other Negative Correlation |
| CDK2 | -0.076 | 0.200015 | Other Negative Correlation |
| C2orf15 | -0.07602 | 0.199907 | Other Negative Correlation |
| TMCO3 | -0.07603 | 0.199859 | Other Negative Correlation |
| TEX48 | -0.07605 | 0.199717 | Other Negative Correlation |
| TERF2 | -0.07621 | 0.198782 | Other Negative Correlation |
| FABP6 | -0.07624 | 0.198572 | Other Negative Correlation |
| WRAP73 | -0.07625 | 0.19853 | Other Negative Correlation |
| POC1B | -0.07626 | 0.198461 | Other Negative Correlation |
| GGA2 | -0.07632 | 0.198103 | Other Negative Correlation |
| ENSG00000289791 | -0.07633 | 0.198054 | Other Negative Correlation |
| UPF3B | -0.07635 | 0.197974 | Other Negative Correlation |
| SLC25A40 | -0.07635 | 0.197943 | Other Negative Correlation |
| BCL2L14 | -0.0764 | 0.197667 | Other Negative Correlation |
| KLHL3 | -0.07641 | 0.197592 | Other Negative Correlation |
| EIF2AK1 | -0.07641 | 0.19759 | Other Negative Correlation |
| SLC25A23 | -0.07644 | 0.19742 | Other Negative Correlation |
| SCCPDH | -0.07645 | 0.197385 | Other Negative Correlation |
| MTARC2 | -0.07645 | 0.19738 | Other Negative Correlation |
| CAMK1G | -0.07646 | 0.197289 | Other Negative Correlation |
| FOXD1 | -0.07648 | 0.197158 | Other Negative Correlation |
| PPP3CC | -0.07652 | 0.196945 | Other Negative Correlation |
| ZNF200 | -0.07653 | 0.196916 | Other Negative Correlation |
| C9orf43 | -0.07657 | 0.196662 | Other Negative Correlation |
| RRAGB | -0.07658 | 0.196572 | Other Negative Correlation |
| NRG2 | -0.07665 | 0.196188 | Other Negative Correlation |
| ENTREP1 | -0.07668 | 0.196038 | Other Negative Correlation |
| CARF | -0.07668 | 0.195994 | Other Negative Correlation |
| CAPN5 | -0.07669 | 0.195923 | Other Negative Correlation |
| SLC11A1 | -0.0768 | 0.195297 | Other Negative Correlation |
| VAT1 | -0.0768 | 0.195285 | Other Negative Correlation |
| OSBP2 | -0.07681 | 0.195258 | Other Negative Correlation |
| PXDNL | -0.07683 | 0.195152 | Other Negative Correlation |
| USP35 | -0.07685 | 0.195045 | Other Negative Correlation |
| ITGAE | -0.07685 | 0.19499 | Other Negative Correlation |
| TEKT2 | -0.07686 | 0.194955 | Other Negative Correlation |
| PIM3 | -0.07689 | 0.194808 | Other Negative Correlation |
| ZNF341 | -0.0769 | 0.19475 | Other Negative Correlation |
| LMO3 | -0.07696 | 0.194379 | Other Negative Correlation |
| FAM50B | -0.07699 | 0.194223 | Other Negative Correlation |
| CHRNA9 | -0.07707 | 0.193762 | Other Negative Correlation |
| LRRIQ3 | -0.07708 | 0.193675 | Other Negative Correlation |
| MUCL1 | -0.0771 | 0.193576 | Other Negative Correlation |
| PROCA1 | -0.0771 | 0.193538 | Other Negative Correlation |
| TRIM45 | -0.07712 | 0.19345 | Other Negative Correlation |
| PALD1 | -0.07713 | 0.193416 | Other Negative Correlation |

| | | | |
|---|---|---|---|
| TCN1 | -0.07714 | 0.193357 | Other Negative Correlation |
| MC4R | -0.07714 | 0.193349 | Other Negative Correlation |
| SLC6A20 | -0.07716 | 0.193232 | Other Negative Correlation |
| ICA1 | -0.07719 | 0.193073 | Other Negative Correlation |
| MMP24 | -0.07721 | 0.192946 | Other Negative Correlation |
| HSD17B3 | -0.07722 | 0.192898 | Other Negative Correlation |
| STRADB | -0.07728 | 0.192551 | Other Negative Correlation |
| PXMP4 | -0.07728 | 0.192522 | Other Negative Correlation |
| DGCR6L | -0.07728 | 0.192522 | Other Negative Correlation |
| KLHL17 | -0.07729 | 0.192443 | Other Negative Correlation |
| PCDHGA2 | -0.0773 | 0.192436 | Other Negative Correlation |
| C2orf74 | -0.07733 | 0.192257 | Other Negative Correlation |
| RIPOR2 | -0.07734 | 0.19215 | Other Negative Correlation |
| NCF4 | -0.0774 | 0.191824 | Other Negative Correlation |
| SH3BGRL2 | -0.07741 | 0.191793 | Other Negative Correlation |
| EPS8 | -0.07743 | 0.191676 | Other Negative Correlation |
| ZNF823 | -0.07744 | 0.191602 | Other Negative Correlation |
| C19orf47 | -0.07749 | 0.191291 | Other Negative Correlation |
| UGT2B15 | -0.0775 | 0.191255 | Other Negative Correlation |
| SLC40A1 | -0.07752 | 0.19112 | Other Negative Correlation |
| ENSG00000272916 | -0.07755 | 0.190984 | Other Negative Correlation |
| HOGA1 | -0.07757 | 0.190841 | Other Negative Correlation |
| ZNF738 | -0.07767 | 0.190262 | Other Negative Correlation |
| GGA1 | -0.07774 | 0.189853 | Other Negative Correlation |
| GDPGP1 | -0.07775 | 0.189847 | Other Negative Correlation |
| OCRL | -0.07776 | 0.189778 | Other Negative Correlation |
| PNCK | -0.07776 | 0.189758 | Other Negative Correlation |
| PCDHB13 | -0.07787 | 0.189144 | Other Negative Correlation |
| MYOT | -0.07788 | 0.189097 | Other Negative Correlation |
| ZSCAN21 | -0.07791 | 0.188934 | Other Negative Correlation |
| ZNF800 | -0.07791 | 0.188925 | Other Negative Correlation |
| ENSG00000283128 | -0.07792 | 0.188835 | Other Negative Correlation |
| TMIGD3 | -0.07793 | 0.188807 | Other Negative Correlation |
| GID4 | -0.07799 | 0.18848 | Other Negative Correlation |
| CORO7 | -0.07799 | 0.188452 | Other Negative Correlation |
| USP38 | -0.07806 | 0.188045 | Other Negative Correlation |
| ZNF554 | -0.07814 | 0.187585 | Other Negative Correlation |
| ABCB8 | -0.07818 | 0.187398 | Other Negative Correlation |
| PEX14 | -0.07821 | 0.187212 | Other Negative Correlation |
| XYLT2 | -0.07821 | 0.187183 | Other Negative Correlation |
| PDPR | -0.07825 | 0.186989 | Other Negative Correlation |
| WHRN | -0.07828 | 0.186827 | Other Negative Correlation |
| NBAS | -0.0783 | 0.186688 | Other Negative Correlation |
| TSPYL4 | -0.07833 | 0.186551 | Other Negative Correlation |
| USP19 | -0.07838 | 0.186271 | Other Negative Correlation |
| CCDC170 | -0.07844 | 0.185931 | Other Negative Correlation |

| | | | |
|---|---|---|---|
| TUBB3 | -0.07846 | 0.185802 | Other Negative Correlation |
| CTSE | -0.0785 | 0.185581 | Other Negative Correlation |
| GARIN4 | -0.07853 | 0.185422 | Other Negative Correlation |
| GSTM3 | -0.07853 | 0.185415 | Other Negative Correlation |
| PPP1R3G | -0.07855 | 0.185289 | Other Negative Correlation |
| ST3GAL2 | -0.07862 | 0.184904 | Other Negative Correlation |
| GCNT3 | -0.07863 | 0.184858 | Other Negative Correlation |
| TTC30B | -0.07863 | 0.184843 | Other Negative Correlation |
| ZNF132 | -0.07863 | 0.184822 | Other Negative Correlation |
| OXNAD1 | -0.07867 | 0.184631 | Other Negative Correlation |
| ITCH | -0.07869 | 0.184515 | Other Negative Correlation |
| PHLDB3 | -0.0787 | 0.184445 | Other Negative Correlation |
| XPNPEP2 | -0.07874 | 0.184224 | Other Negative Correlation |
| TANGO2 | -0.07879 | 0.183954 | Other Negative Correlation |
| WDR5B | -0.07879 | 0.183952 | Other Negative Correlation |
| KLHL28 | -0.0788 | 0.183895 | Other Negative Correlation |
| CCDC39 | -0.07886 | 0.183579 | Other Negative Correlation |
| DISP2 | -0.07889 | 0.183399 | Other Negative Correlation |
| COP1 | -0.0789 | 0.183336 | Other Negative Correlation |
| PARP3 | -0.07893 | 0.183171 | Other Negative Correlation |
| ZNF664 | -0.07895 | 0.183068 | Other Negative Correlation |
| WDR41 | -0.07896 | 0.183017 | Other Negative Correlation |
| EPC1 | -0.07901 | 0.182705 | Other Negative Correlation |
| PRDM15 | -0.07914 | 0.18203 | Other Negative Correlation |
| RASL11B | -0.07915 | 0.181974 | Other Negative Correlation |
| STAM2 | -0.07916 | 0.181888 | Other Negative Correlation |
| ZNF324B | -0.07917 | 0.181833 | Other Negative Correlation |
| MMEL1 | -0.07917 | 0.181819 | Other Negative Correlation |
| EPB41L2 | -0.07918 | 0.181777 | Other Negative Correlation |
| HEXIM2 | -0.07919 | 0.181731 | Other Negative Correlation |
| CATSPERZ | -0.0792 | 0.181666 | Other Negative Correlation |
| SLC6A7 | -0.07921 | 0.181601 | Other Negative Correlation |
| ZNF584 | -0.07922 | 0.18155 | Other Negative Correlation |
| JAKMIP3 | -0.07925 | 0.181368 | Other Negative Correlation |
| NINL | -0.07929 | 0.181166 | Other Negative Correlation |
| FAM118B | -0.0793 | 0.181127 | Other Negative Correlation |
| ABTB3 | -0.07943 | 0.180415 | Other Negative Correlation |
| ZNF446 | -0.07949 | 0.180083 | Other Negative Correlation |
| ZNF836 | -0.07952 | 0.1799 | Other Negative Correlation |
| ASTN1 | -0.07955 | 0.179718 | Other Negative Correlation |
| COL21A1 | -0.07956 | 0.179691 | Other Negative Correlation |
| NDST2 | -0.07957 | 0.179609 | Other Negative Correlation |
| H2AC19 | -0.0796 | 0.179455 | Other Negative Correlation |
| ZNF251 | -0.07961 | 0.17941 | Other Negative Correlation |
| SYDE2 | -0.07963 | 0.179314 | Other Negative Correlation |
| PTGIR | -0.07965 | 0.179205 | Other Negative Correlation |

| | | | |
|---|---|---|---|
| NFATC4 | -0.07966 | 0.179119 | Other Negative Correlation |
| PRPH2 | -0.07968 | 0.179044 | Other Negative Correlation |
| PCDHGA9 | -0.07975 | 0.178653 | Other Negative Correlation |
| DEDD | -0.07983 | 0.178188 | Other Negative Correlation |
| PHGR1 | -0.07984 | 0.178139 | Other Negative Correlation |
| FBXL15 | -0.07985 | 0.178097 | Other Negative Correlation |
| NAGS | -0.07987 | 0.17799 | Other Negative Correlation |
| DNAH17 | -0.07989 | 0.177861 | Other Negative Correlation |
| LPAR2 | -0.07991 | 0.177793 | Other Negative Correlation |
| ESRRG | -0.07994 | 0.177635 | Other Negative Correlation |
| RFFL | -0.07994 | 0.177618 | Other Negative Correlation |
| LIG4 | -0.07994 | 0.177614 | Other Negative Correlation |
| GOLGA8Q | -0.07997 | 0.177445 | Other Negative Correlation |
| GATB | -0.08001 | 0.177239 | Other Negative Correlation |
| GPR108 | -0.08005 | 0.177023 | Other Negative Correlation |
| C5orf46 | -0.08005 | 0.177018 | Other Negative Correlation |
| HMCN1 | -0.08006 | 0.176957 | Other Negative Correlation |
| MFSD14A | -0.08018 | 0.176296 | Other Negative Correlation |
| ZNF737 | -0.08021 | 0.176122 | Other Negative Correlation |
| ZNF875 | -0.08023 | 0.176028 | Other Negative Correlation |
| SCARF1 | -0.08031 | 0.175631 | Other Negative Correlation |
| ZNF672 | -0.08033 | 0.175515 | Other Negative Correlation |
| ADRB3 | -0.08034 | 0.175466 | Other Negative Correlation |
| PABIR1 | -0.0804 | 0.175116 | Other Negative Correlation |
| PEX5 | -0.08047 | 0.174741 | Other Negative Correlation |
| MTMR10 | -0.08053 | 0.174407 | Other Negative Correlation |
| ORMDL3 | -0.08055 | 0.174338 | Other Negative Correlation |
| SHLD3 | -0.08056 | 0.174252 | Other Negative Correlation |
| GLB1 | -0.0806 | 0.174069 | Other Negative Correlation |
| C15orf62 | -0.08062 | 0.173941 | Other Negative Correlation |
| ILRUN | -0.08063 | 0.173861 | Other Negative Correlation |
| DOK7 | -0.08064 | 0.173823 | Other Negative Correlation |
| P2RY1 | -0.08068 | 0.173636 | Other Negative Correlation |
| C8orf48 | -0.08068 | 0.173602 | Other Negative Correlation |
| LCE3D | -0.08069 | 0.173577 | Other Negative Correlation |
| PIGT | -0.08069 | 0.173552 | Other Negative Correlation |
| ATG16L2 | -0.0807 | 0.173535 | Other Negative Correlation |
| SNX32 | -0.0807 | 0.173513 | Other Negative Correlation |
| NLRP2 | -0.08072 | 0.173423 | Other Negative Correlation |
| ARFGAP3 | -0.08072 | 0.173408 | Other Negative Correlation |
| PTPRN2 | -0.08073 | 0.173371 | Other Negative Correlation |
| MTM1 | -0.08079 | 0.173025 | Other Negative Correlation |
| DNASE2B | -0.08083 | 0.17282 | Other Negative Correlation |
| NNT | -0.08086 | 0.172653 | Other Negative Correlation |
| ANO3 | -0.08091 | 0.172413 | Other Negative Correlation |
| SLC43A3 | -0.08091 | 0.172401 | Other Negative Correlation |

| | | | |
|---|---|---|---|
| CD180 | -0.08093 | 0.172309 | Other Negative Correlation |
| ANKHD1 | -0.081 | 0.171938 | Other Negative Correlation |
| P3H3 | -0.08102 | 0.171803 | Other Negative Correlation |
| PLEKHM3 | -0.08102 | 0.171792 | Other Negative Correlation |
| NUCB1 | -0.0811 | 0.171407 | Other Negative Correlation |
| CD96 | -0.0811 | 0.171407 | Other Negative Correlation |
| ZFP14 | -0.08114 | 0.171175 | Other Negative Correlation |
| TPRA1 | -0.0812 | 0.170871 | Other Negative Correlation |
| APOL4 | -0.08121 | 0.17081 | Other Negative Correlation |
| MORN3 | -0.08123 | 0.17072 | Other Negative Correlation |
| SULT1A3 | -0.08123 | 0.170713 | Other Negative Correlation |
| SERPINB10 | -0.08126 | 0.170561 | Other Negative Correlation |
| RNF145 | -0.0813 | 0.170315 | Other Negative Correlation |
| ZFYVE28 | -0.0813 | 0.170308 | Other Negative Correlation |
| RAB24 | -0.08131 | 0.17028 | Other Negative Correlation |
| REEP6 | -0.08134 | 0.170102 | Other Negative Correlation |
| MINDY3 | -0.08135 | 0.170043 | Other Negative Correlation |
| ZNF607 | -0.08136 | 0.17001 | Other Negative Correlation |
| SLC24A3 | -0.08137 | 0.169981 | Other Negative Correlation |
| ZDHHC11 | -0.0814 | 0.16982 | Other Negative Correlation |
| ERN1 | -0.08144 | 0.169598 | Other Negative Correlation |
| RBPJ | -0.08146 | 0.169467 | Other Negative Correlation |
| PIGV | -0.08147 | 0.169431 | Other Negative Correlation |
| KCNK1 | -0.08152 | 0.169182 | Other Negative Correlation |
| C1orf167 | -0.08153 | 0.169136 | Other Negative Correlation |
| SEMA6A | -0.08153 | 0.169103 | Other Negative Correlation |
| NOXRED1 | -0.08161 | 0.168715 | Other Negative Correlation |
| ANGPTL6 | -0.08162 | 0.168664 | Other Negative Correlation |
| ENSG00000289724 | -0.08173 | 0.168079 | Other Negative Correlation |
| SPANXN5 | -0.08173 | 0.168079 | Other Negative Correlation |
| SERPINA4 | -0.08173 | 0.168079 | Other Negative Correlation |
| P2RY14 | -0.08173 | 0.168079 | Other Negative Correlation |
| STK31 | -0.08173 | 0.168079 | Other Negative Correlation |
| KY | -0.08173 | 0.168079 | Other Negative Correlation |
| PPM1D | -0.08175 | 0.167978 | Other Negative Correlation |
| CTIF | -0.08178 | 0.167809 | Other Negative Correlation |
| BTN2A1 | -0.0818 | 0.167722 | Other Negative Correlation |
| ZXDC | -0.0818 | 0.167694 | Other Negative Correlation |
| PIERCE1 | -0.08184 | 0.167502 | Other Negative Correlation |
| FYB1 | -0.08187 | 0.167363 | Other Negative Correlation |
| IER3IP1 | -0.08187 | 0.167363 | Other Negative Correlation |
| SDR9C7 | -0.08189 | 0.167243 | Other Negative Correlation |
| ZNF222 | -0.08198 | 0.166765 | Other Negative Correlation |
| CA13 | -0.082 | 0.166687 | Other Negative Correlation |
| ATP2B1 | -0.082 | 0.16665 | Other Negative Correlation |
| ADAMTS7 | -0.08201 | 0.166638 | Other Negative Correlation |

| | | | |
|---|---|---|---|
| PLEKHJ1 | -0.08202 | 0.16658 | Other Negative Correlation |
| ASB6 | -0.08204 | 0.166478 | Other Negative Correlation |
| HES5 | -0.08208 | 0.166268 | Other Negative Correlation |
| USH1G | -0.08208 | 0.166233 | Other Negative Correlation |
| CHPT1 | -0.08216 | 0.165848 | Other Negative Correlation |
| CYREN | -0.08222 | 0.165538 | Other Negative Correlation |
| PCSK1 | -0.08225 | 0.165359 | Other Negative Correlation |
| C1orf210 | -0.08225 | 0.165347 | Other Negative Correlation |
| CCNO | -0.08227 | 0.165277 | Other Negative Correlation |
| SLC34A1 | -0.0823 | 0.165092 | Other Negative Correlation |
| HSD11B2 | -0.08231 | 0.165087 | Other Negative Correlation |
| CDC42SE2 | -0.08232 | 0.165019 | Other Negative Correlation |
| SEMA5A | -0.08233 | 0.164969 | Other Negative Correlation |
| EPHX3 | -0.08237 | 0.164767 | Other Negative Correlation |
| MYO1F | -0.08239 | 0.164628 | Other Negative Correlation |
| ETV2 | -0.08244 | 0.164375 | Other Negative Correlation |
| CYP4F12 | -0.08248 | 0.164188 | Other Negative Correlation |
| NDUFA7 | -0.08248 | 0.164184 | Other Negative Correlation |
| GBGT1 | -0.0825 | 0.164084 | Other Negative Correlation |
| DNAJC25 | -0.08251 | 0.164027 | Other Negative Correlation |
| LMTK2 | -0.08253 | 0.163945 | Other Negative Correlation |
| SUSD1 | -0.08254 | 0.163879 | Other Negative Correlation |
| WBP2 | -0.08254 | 0.163869 | Other Negative Correlation |
| TNPO2 | -0.08257 | 0.163736 | Other Negative Correlation |
| SLC66A1 | -0.0826 | 0.163571 | Other Negative Correlation |
| KLRC1 | -0.08269 | 0.163137 | Other Negative Correlation |
| SRMS | -0.08269 | 0.163123 | Other Negative Correlation |
| POP5 | -0.08271 | 0.163035 | Other Negative Correlation |
| ZBTB17 | -0.08273 | 0.162908 | Other Negative Correlation |
| STK19 | -0.08291 | 0.161981 | Other Negative Correlation |
| FNDC11 | -0.08293 | 0.161879 | Other Negative Correlation |
| PIK3R3 | -0.08298 | 0.161659 | Other Negative Correlation |
| DSC1 | -0.08303 | 0.161365 | Other Negative Correlation |
| SERPINA6 | -0.0831 | 0.161032 | Other Negative Correlation |
| DDX59 | -0.08315 | 0.1608 | Other Negative Correlation |
| KDM6B | -0.08317 | 0.160692 | Other Negative Correlation |
| NEBL | -0.0832 | 0.160545 | Other Negative Correlation |
| SETBP1 | -0.08321 | 0.160483 | Other Negative Correlation |
| GRTP1 | -0.08332 | 0.159927 | Other Negative Correlation |
| MACROD2 | -0.08336 | 0.159725 | Other Negative Correlation |
| PEX3 | -0.08336 | 0.159724 | Other Negative Correlation |
| ENSG00000275740 | -0.08341 | 0.15947 | Other Negative Correlation |
| TCEANC2 | -0.08349 | 0.159061 | Other Negative Correlation |
| PLPPR5 | -0.08353 | 0.158853 | Other Negative Correlation |
| MPIG6B | -0.08356 | 0.158729 | Other Negative Correlation |
| SPRY4 | -0.08357 | 0.158653 | Other Negative Correlation |

| | | | |
|---|---|---|---|
| ATG2A | -0.08359 | 0.15855 | Other Negative Correlation |
| C15orf61 | -0.08376 | 0.157728 | Other Negative Correlation |
| ZNF653 | -0.08378 | 0.157624 | Other Negative Correlation |
| IQUB | -0.08379 | 0.15755 | Other Negative Correlation |
| DENND11 | -0.08386 | 0.157232 | Other Negative Correlation |
| ADI1 | -0.08386 | 0.157203 | Other Negative Correlation |
| YIF1B | -0.08389 | 0.15707 | Other Negative Correlation |
| ZNF707 | -0.0839 | 0.157022 | Other Negative Correlation |
| MARF1 | -0.08391 | 0.156986 | Other Negative Correlation |
| TTC32 | -0.08404 | 0.156341 | Other Negative Correlation |
| PAFAH2 | -0.08411 | 0.155975 | Other Negative Correlation |
| RASIP1 | -0.08416 | 0.155731 | Other Negative Correlation |
| ALPK1 | -0.08416 | 0.155715 | Other Negative Correlation |
| ARL8A | -0.08419 | 0.155589 | Other Negative Correlation |
| ACVR2B | -0.08422 | 0.155418 | Other Negative Correlation |
| SIMC1 | -0.08423 | 0.155399 | Other Negative Correlation |
| LY96 | -0.08431 | 0.154987 | Other Negative Correlation |
| FAM83C | -0.08432 | 0.154965 | Other Negative Correlation |
| IFIT1B | -0.08442 | 0.15446 | Other Negative Correlation |
| SMCO2 | -0.08443 | 0.154394 | Other Negative Correlation |
| ID2 | -0.08443 | 0.154387 | Other Negative Correlation |
| ZFYVE1 | -0.08446 | 0.154248 | Other Negative Correlation |
| DVL2 | -0.08447 | 0.154234 | Other Negative Correlation |
| ARMH1 | -0.08451 | 0.154029 | Other Negative Correlation |
| INTS6L | -0.08456 | 0.153763 | Other Negative Correlation |
| GLCCI1 | -0.08461 | 0.153528 | Other Negative Correlation |
| LCE3C | -0.08464 | 0.153406 | Other Negative Correlation |
| HECA | -0.08465 | 0.153345 | Other Negative Correlation |
| FAN1 | -0.08467 | 0.153246 | Other Negative Correlation |
| SCNN1D | -0.08472 | 0.153005 | Other Negative Correlation |
| TRMT2B | -0.08473 | 0.15295 | Other Negative Correlation |
| SNN | -0.08477 | 0.15274 | Other Negative Correlation |
| LRP11 | -0.08479 | 0.152675 | Other Negative Correlation |
| KIF25 | -0.08479 | 0.152637 | Other Negative Correlation |
| SPIN2B | -0.08484 | 0.152385 | Other Negative Correlation |
| TEPP | -0.08485 | 0.152384 | Other Negative Correlation |
| FBXO11 | -0.08486 | 0.152316 | Other Negative Correlation |
| IL2RB | -0.08487 | 0.152284 | Other Negative Correlation |
| SLN | -0.08487 | 0.152262 | Other Negative Correlation |
| CCDC127 | -0.08491 | 0.152078 | Other Negative Correlation |
| MT1A | -0.08499 | 0.151662 | Other Negative Correlation |
| TOMM20L | -0.08501 | 0.151571 | Other Negative Correlation |
| PECAM1 | -0.08503 | 0.151504 | Other Negative Correlation |
| ZNF597 | -0.08509 | 0.151186 | Other Negative Correlation |
| GMCL1 | -0.08517 | 0.150807 | Other Negative Correlation |
| COL5A3 | -0.08517 | 0.150795 | Other Negative Correlation |

| | | | |
|---|---|---|---|
| CLN3 | -0.08521 | 0.150638 | Other Negative Correlation |
| CYP1A2 | -0.08522 | 0.150597 | Other Negative Correlation |
| SPRING1 | -0.08532 | 0.150076 | Other Negative Correlation |
| DPY19L2 | -0.08535 | 0.149954 | Other Negative Correlation |
| ACSF3 | -0.08537 | 0.149873 | Other Negative Correlation |
| APOE | -0.08546 | 0.149447 | Other Negative Correlation |
| ROM1 | -0.08552 | 0.149124 | Other Negative Correlation |
| RPS6KA6 | -0.0856 | 0.148762 | Other Negative Correlation |
| ZNF559 | -0.08561 | 0.148727 | Other Negative Correlation |
| SHANK2 | -0.08568 | 0.148375 | Other Negative Correlation |
| MFSD4B | -0.08569 | 0.148329 | Other Negative Correlation |
| PALB2 | -0.08571 | 0.148244 | Other Negative Correlation |
| WNK4 | -0.08577 | 0.147941 | Other Negative Correlation |
| ABCC2 | -0.0858 | 0.147822 | Other Negative Correlation |
| ZNF226 | -0.08582 | 0.147704 | Other Negative Correlation |
| ORAI2 | -0.08584 | 0.147631 | Other Negative Correlation |
| DCK | -0.08585 | 0.147567 | Other Negative Correlation |
| GJB7 | -0.08586 | 0.147518 | Other Negative Correlation |
| MT1F | -0.08589 | 0.147381 | Other Negative Correlation |
| ADAM23 | -0.08597 | 0.147008 | Other Negative Correlation |
| CHST2 | -0.08601 | 0.146826 | Other Negative Correlation |
| DLX5 | -0.08602 | 0.146754 | Other Negative Correlation |
| FBXO8 | -0.08602 | 0.146751 | Other Negative Correlation |
| PPP1R21 | -0.08608 | 0.146488 | Other Negative Correlation |
| VAMP1 | -0.08609 | 0.146443 | Other Negative Correlation |
| C1orf198 | -0.08613 | 0.146243 | Other Negative Correlation |
| SPIN4 | -0.08613 | 0.146243 | Other Negative Correlation |
| VTI1A | -0.08617 | 0.146062 | Other Negative Correlation |
| TMEM160 | -0.08621 | 0.145884 | Other Negative Correlation |
| CYP21A2 | -0.08631 | 0.145406 | Other Negative Correlation |
| PGC | -0.08631 | 0.145406 | Other Negative Correlation |
| CLPS | -0.08631 | 0.145406 | Other Negative Correlation |
| STC1 | -0.08634 | 0.145247 | Other Negative Correlation |
| SERPINI1 | -0.08634 | 0.145243 | Other Negative Correlation |
| ZSCAN4 | -0.08636 | 0.145162 | Other Negative Correlation |
| CMTR1 | -0.08644 | 0.144802 | Other Negative Correlation |
| NDUFA9 | -0.08645 | 0.144736 | Other Negative Correlation |
| PDPN | -0.08648 | 0.144605 | Other Negative Correlation |
| SNX2 | -0.08658 | 0.144161 | Other Negative Correlation |
| UBQLN4 | -0.08663 | 0.143929 | Other Negative Correlation |
| B4GALT5 | -0.08671 | 0.143554 | Other Negative Correlation |
| TLL2 | -0.08673 | 0.143464 | Other Negative Correlation |
| KIF9 | -0.0868 | 0.143122 | Other Negative Correlation |
| SERTAD2 | -0.08684 | 0.142952 | Other Negative Correlation |
| MPV17L | -0.08685 | 0.142901 | Other Negative Correlation |
| ZNF630 | -0.08685 | 0.142886 | Other Negative Correlation |

| | | | |
|---|---|---|---|
| NCOA6 | -0.08689 | 0.142688 | Other Negative Correlation |
| GJC3 | -0.08697 | 0.142332 | Other Negative Correlation |
| PHACTR1 | -0.08705 | 0.141952 | Other Negative Correlation |
| GON4L | -0.0871 | 0.141737 | Other Negative Correlation |
| KCNE1 | -0.0871 | 0.141728 | Other Negative Correlation |
| IPMK | -0.08717 | 0.141402 | Other Negative Correlation |
| SUSD2 | -0.0872 | 0.141266 | Other Negative Correlation |
| TTC21B | -0.08722 | 0.141211 | Other Negative Correlation |
| ZNG1F | -0.08722 | 0.14117 | Other Negative Correlation |
| SHC4 | -0.08723 | 0.14115 | Other Negative Correlation |
| LRRC34 | -0.08724 | 0.141122 | Other Negative Correlation |
| SATB1 | -0.08724 | 0.141102 | Other Negative Correlation |
| SLC5A6 | -0.08724 | 0.141098 | Other Negative Correlation |
| SOX18 | -0.08732 | 0.140726 | Other Negative Correlation |
| LIF | -0.08735 | 0.140606 | Other Negative Correlation |
| ZNF140 | -0.08737 | 0.140505 | Other Negative Correlation |
| RAPGEF2 | -0.08741 | 0.140313 | Other Negative Correlation |
| NME5 | -0.08741 | 0.140302 | Other Negative Correlation |
| TSLP | -0.08745 | 0.140128 | Other Negative Correlation |
| ZNF267 | -0.08749 | 0.139955 | Other Negative Correlation |
| ZSWIM4 | -0.08755 | 0.139689 | Other Negative Correlation |
| PAK4 | -0.08758 | 0.13957 | Other Negative Correlation |
| GPATCH11 | -0.08764 | 0.139285 | Other Negative Correlation |
| INSYN2B | -0.08769 | 0.139071 | Other Negative Correlation |
| CASTOR2 | -0.08769 | 0.139057 | Other Negative Correlation |
| FAM161B | -0.08774 | 0.138843 | Other Negative Correlation |
| DCUN1D4 | -0.08774 | 0.138838 | Other Negative Correlation |
| C13orf46 | -0.08774 | 0.138823 | Other Negative Correlation |
| SEMA5B | -0.08775 | 0.138775 | Other Negative Correlation |
| SPINDOC | -0.08775 | 0.138762 | Other Negative Correlation |
| CLMN | -0.08783 | 0.138436 | Other Negative Correlation |
| RDH10 | -0.0879 | 0.138113 | Other Negative Correlation |
| DIABLO | -0.0879 | 0.138093 | Other Negative Correlation |
| MEOX1 | -0.08793 | 0.137964 | Other Negative Correlation |
| MAX | -0.08795 | 0.137888 | Other Negative Correlation |
| FBXO28 | -0.08796 | 0.137841 | Other Negative Correlation |
| HYAL4 | -0.08802 | 0.137586 | Other Negative Correlation |
| WDR27 | -0.08805 | 0.13743 | Other Negative Correlation |
| PLCG1 | -0.08805 | 0.137419 | Other Negative Correlation |
| MBTD1 | -0.08806 | 0.137404 | Other Negative Correlation |
| TIGD7 | -0.08808 | 0.137308 | Other Negative Correlation |
| VTCN1 | -0.0881 | 0.13722 | Other Negative Correlation |
| SNX25 | -0.08811 | 0.13717 | Other Negative Correlation |
| SHKBP1 | -0.08815 | 0.136981 | Other Negative Correlation |
| NAALADL2 | -0.08818 | 0.136831 | Other Negative Correlation |
| METTL6 | -0.08823 | 0.136607 | Other Negative Correlation |

| | | | |
|---|---|---|---|
| SLC44A3 | -0.08827 | 0.13646 | Other Negative Correlation |
| RHOH | -0.08832 | 0.136232 | Other Negative Correlation |
| DPF2 | -0.08835 | 0.136072 | Other Negative Correlation |
| CAPN10 | -0.08838 | 0.135977 | Other Negative Correlation |
| SGTA | -0.08838 | 0.135948 | Other Negative Correlation |
| GOLGA8R | -0.08839 | 0.135927 | Other Negative Correlation |
| RAP1GAP | -0.0884 | 0.135859 | Other Negative Correlation |
| DIRAS2 | -0.08848 | 0.135533 | Other Negative Correlation |
| PIK3CD | -0.08849 | 0.13548 | Other Negative Correlation |
| PCDHA6 | -0.0886 | 0.134968 | Other Negative Correlation |
| CFAP20DC | -0.08861 | 0.134927 | Other Negative Correlation |
| SLC25A41 | -0.08862 | 0.134903 | Other Negative Correlation |
| HSD3B7 | -0.08865 | 0.134781 | Other Negative Correlation |
| SETSIP | -0.0887 | 0.134524 | Other Negative Correlation |
| MID2 | -0.08875 | 0.134298 | Other Negative Correlation |
| ABTB1 | -0.08882 | 0.134006 | Other Negative Correlation |
| ADGRL3 | -0.08884 | 0.133909 | Other Negative Correlation |
| C1GALT1C1L | -0.08893 | 0.133545 | Other Negative Correlation |
| PLEKHH1 | -0.08894 | 0.133474 | Other Negative Correlation |
| PF4V1 | -0.08897 | 0.133354 | Other Negative Correlation |
| APBA2 | -0.08902 | 0.133137 | Other Negative Correlation |
| SYNJ2 | -0.08902 | 0.133117 | Other Negative Correlation |
| RARB | -0.08902 | 0.133114 | Other Negative Correlation |
| RIMS2 | -0.08903 | 0.133082 | Other Negative Correlation |
| PIGU | -0.08903 | 0.133075 | Other Negative Correlation |
| TMEM223 | -0.08904 | 0.133028 | Other Negative Correlation |
| CISH | -0.08908 | 0.132865 | Other Negative Correlation |
| ERICH5 | -0.08913 | 0.132645 | Other Negative Correlation |
| RXFP4 | -0.08921 | 0.132306 | Other Negative Correlation |
| C12orf57 | -0.08921 | 0.132289 | Other Negative Correlation |
| ROBO4 | -0.08924 | 0.132166 | Other Negative Correlation |
| ACOX2 | -0.08925 | 0.132153 | Other Negative Correlation |
| TUSC2 | -0.0893 | 0.131928 | Other Negative Correlation |
| SERHL2 | -0.0893 | 0.131913 | Other Negative Correlation |
| FAF1 | -0.0893 | 0.131909 | Other Negative Correlation |
| ALDH4A1 | -0.08933 | 0.131798 | Other Negative Correlation |
| ABCB6 | -0.08936 | 0.131635 | Other Negative Correlation |
| KRT82 | -0.08938 | 0.131582 | Other Negative Correlation |
| HAP1 | -0.0895 | 0.131033 | Other Negative Correlation |
| SAC3D1 | -0.08959 | 0.130665 | Other Negative Correlation |
| ATF7 | -0.08967 | 0.130316 | Other Negative Correlation |
| ARRDC3 | -0.08967 | 0.130308 | Other Negative Correlation |
| CPSF1 | -0.08967 | 0.130307 | Other Negative Correlation |
| HEY1 | -0.08968 | 0.130271 | Other Negative Correlation |
| KANK2 | -0.0897 | 0.130191 | Other Negative Correlation |
| SUPT20H | -0.08971 | 0.130132 | Other Negative Correlation |

| | | | |
|---|---|---|---|
| C14orf93 | -0.08976 | 0.12991 | Other Negative Correlation |
| B3GNT4 | -0.08979 | 0.129781 | Other Negative Correlation |
| C12orf71 | -0.08982 | 0.129655 | Other Negative Correlation |
| DMAP1 | -0.08991 | 0.129305 | Other Negative Correlation |
| INHBC | -0.08993 | 0.12919 | Other Negative Correlation |
| OR2B11 | -0.08993 | 0.12919 | Other Negative Correlation |
| SLC7A6OS | -0.08997 | 0.129035 | Other Negative Correlation |
| ARSG | -0.09 | 0.128897 | Other Negative Correlation |
| SMIM1 | -0.09003 | 0.12879 | Other Negative Correlation |
| TENT4A | -0.09007 | 0.128607 | Other Negative Correlation |
| BMI1 | -0.0901 | 0.128487 | Other Negative Correlation |
| ATP13A1 | -0.09012 | 0.128401 | Other Negative Correlation |
| SAA4 | -0.09016 | 0.128239 | Other Negative Correlation |
| C1QTNF3 | -0.09019 | 0.128103 | Other Negative Correlation |
| LRP4 | -0.0902 | 0.128069 | Other Negative Correlation |
| PRSS54 | -0.0902 | 0.128035 | Other Negative Correlation |
| UBN1 | -0.0903 | 0.127644 | Other Negative Correlation |
| PRX | -0.09032 | 0.127535 | Other Negative Correlation |
| WASHC2C | -0.09032 | 0.127519 | Other Negative Correlation |
| PLEKHG2 | -0.09037 | 0.12732 | Other Negative Correlation |
| PIWIL4 | -0.09049 | 0.126808 | Other Negative Correlation |
| SLC29A3 | -0.0905 | 0.126778 | Other Negative Correlation |
| ECHDC2 | -0.09057 | 0.126484 | Other Negative Correlation |
| TNFRSF19 | -0.09059 | 0.126394 | Other Negative Correlation |
| MIOS | -0.09075 | 0.12575 | Other Negative Correlation |
| ZNF675 | -0.09075 | 0.125711 | Other Negative Correlation |
| CPA5 | -0.09077 | 0.125645 | Other Negative Correlation |
| ZBTB3 | -0.09083 | 0.125409 | Other Negative Correlation |
| PARP15 | -0.09083 | 0.125401 | Other Negative Correlation |
| CCDC146 | -0.0909 | 0.125121 | Other Negative Correlation |
| ZNF22 | -0.0909 | 0.125115 | Other Negative Correlation |
| LETMD1 | -0.09094 | 0.124934 | Other Negative Correlation |
| PTPN9 | -0.09095 | 0.124905 | Other Negative Correlation |
| CENPX | -0.09096 | 0.124872 | Other Negative Correlation |
| POU6F2 | -0.09098 | 0.124752 | Other Negative Correlation |
| UBN2 | -0.091 | 0.124666 | Other Negative Correlation |
| ABCA13 | -0.09101 | 0.124644 | Other Negative Correlation |
| DNLZ | -0.09106 | 0.124431 | Other Negative Correlation |
| ZNF623 | -0.09106 | 0.124428 | Other Negative Correlation |
| PSG1 | -0.09113 | 0.124166 | Other Negative Correlation |
| INSIG2 | -0.09121 | 0.123834 | Other Negative Correlation |
| FMOD | -0.09125 | 0.123631 | Other Negative Correlation |
| ALG11 | -0.09126 | 0.123608 | Other Negative Correlation |
| ANAPC2 | -0.09127 | 0.123577 | Other Negative Correlation |
| ZNF547 | -0.09129 | 0.123493 | Other Negative Correlation |
| CNOT9 | -0.09132 | 0.123349 | Other Negative Correlation |

| | | | |
|---|---|---|---|
| NUBP1 | -0.09133 | 0.123328 | Other Negative Correlation |
| TTC36 | -0.09134 | 0.123286 | Other Negative Correlation |
| CASD1 | -0.0914 | 0.123016 | Other Negative Correlation |
| P2RX4 | -0.09141 | 0.122978 | Other Negative Correlation |
| PMAIP1 | -0.09141 | 0.122971 | Other Negative Correlation |
| CGB7 | -0.09142 | 0.122959 | Other Negative Correlation |
| DCTN4 | -0.09146 | 0.122796 | Other Negative Correlation |
| PCDHB3 | -0.09149 | 0.122669 | Other Negative Correlation |
| LZIC | -0.09155 | 0.122424 | Other Negative Correlation |
| ZNF839 | -0.09155 | 0.122402 | Other Negative Correlation |
| KAZN | -0.09159 | 0.122247 | Other Negative Correlation |
| ZNF436 | -0.09161 | 0.122184 | Other Negative Correlation |
| ELF3 | -0.09161 | 0.122178 | Other Negative Correlation |
| LRRIQ1 | -0.09161 | 0.122157 | Other Negative Correlation |
| ZNG1B | -0.09162 | 0.122139 | Other Negative Correlation |
| SNAI1 | -0.09163 | 0.122083 | Other Negative Correlation |
| TEP1 | -0.09171 | 0.121771 | Other Negative Correlation |
| SNPH | -0.09172 | 0.121711 | Other Negative Correlation |
| USP20 | -0.09178 | 0.121489 | Other Negative Correlation |
| LAMB2 | -0.09178 | 0.121483 | Other Negative Correlation |
| MED16 | -0.09184 | 0.121224 | Other Negative Correlation |
| ICAM1 | -0.09197 | 0.120684 | Other Negative Correlation |
| PLXNC1 | -0.09198 | 0.120677 | Other Negative Correlation |
| ARMC1 | -0.09198 | 0.120659 | Other Negative Correlation |
| NDUFAF5 | -0.092 | 0.12059 | Other Negative Correlation |
| R3HCC1 | -0.09201 | 0.120548 | Other Negative Correlation |
| PSMD11 | -0.09206 | 0.120318 | Other Negative Correlation |
| MAFF | -0.0921 | 0.120162 | Other Negative Correlation |
| SLC16A13 | -0.09211 | 0.120124 | Other Negative Correlation |
| HCST | -0.09215 | 0.119965 | Other Negative Correlation |
| PPP1R3C | -0.09216 | 0.119949 | Other Negative Correlation |
| BICD1 | -0.09217 | 0.119901 | Other Negative Correlation |
| RNF26 | -0.09219 | 0.119818 | Other Negative Correlation |
| SMIM10L1 | -0.0922 | 0.119756 | Other Negative Correlation |
| VPS33A | -0.09223 | 0.119647 | Other Negative Correlation |
| APCDD1L | -0.09225 | 0.119551 | Other Negative Correlation |
| HSPB2 | -0.09226 | 0.119546 | Other Negative Correlation |
| ST3GAL1 | -0.09232 | 0.119291 | Other Negative Correlation |
| RUNX2 | -0.09233 | 0.11924 | Other Negative Correlation |
| RPRD2 | -0.09233 | 0.119237 | Other Negative Correlation |
| PTEN | -0.09233 | 0.119232 | Other Negative Correlation |
| CACFD1 | -0.09234 | 0.119208 | Other Negative Correlation |
| KMT2A | -0.09236 | 0.119109 | Other Negative Correlation |
| MAEA | -0.09242 | 0.1189 | Other Negative Correlation |
| ZFP28 | -0.09245 | 0.118752 | Other Negative Correlation |
| KCNE2 | -0.09246 | 0.118722 | Other Negative Correlation |

| | | | |
|---|---|---|---|
| ILDR1 | -0.09247 | 0.118674 | Other Negative Correlation |
| LEKR1 | -0.09251 | 0.118534 | Other Negative Correlation |
| SH2D2A | -0.09267 | 0.117893 | Other Negative Correlation |
| INAFM2 | -0.09267 | 0.117887 | Other Negative Correlation |
| UTRN | -0.09274 | 0.117624 | Other Negative Correlation |
| NFASC | -0.09275 | 0.117589 | Other Negative Correlation |
| KISS1 | -0.09277 | 0.117493 | Other Negative Correlation |
| CTSK | -0.09277 | 0.117483 | Other Negative Correlation |
| MBIP | -0.09285 | 0.117193 | Other Negative Correlation |
| PLPP3 | -0.09285 | 0.11718 | Other Negative Correlation |
| TMCO4 | -0.09287 | 0.117113 | Other Negative Correlation |
| FNBP1 | -0.09289 | 0.117009 | Other Negative Correlation |
| TEX9 | -0.0929 | 0.116987 | Other Negative Correlation |
| HPGD | -0.09295 | 0.116771 | Other Negative Correlation |
| UPF3A | -0.09297 | 0.116717 | Other Negative Correlation |
| PARP11 | -0.09298 | 0.116662 | Other Negative Correlation |
| VPS45 | -0.09299 | 0.11662 | Other Negative Correlation |
| PLPP1 | -0.09302 | 0.116483 | Other Negative Correlation |
| ATXN7L1 | -0.09305 | 0.116378 | Other Negative Correlation |
| MFSD6 | -0.09306 | 0.116329 | Other Negative Correlation |
| GSR | -0.09307 | 0.116313 | Other Negative Correlation |
| C5AR2 | -0.09309 | 0.116245 | Other Negative Correlation |
| IGFBP3 | -0.09313 | 0.116088 | Other Negative Correlation |
| OR6B3 | -0.09316 | 0.115953 | Other Negative Correlation |
| MCUR1 | -0.09333 | 0.115271 | Other Negative Correlation |
| ECD | -0.09341 | 0.114989 | Other Negative Correlation |
| ZNF713 | -0.09341 | 0.114965 | Other Negative Correlation |
| SPIN3 | -0.09349 | 0.114649 | Other Negative Correlation |
| PLA2G12A | -0.09354 | 0.114459 | Other Negative Correlation |
| POGLUT3 | -0.09357 | 0.114352 | Other Negative Correlation |
| PPFIA3 | -0.09368 | 0.113911 | Other Negative Correlation |
| MARCHF10 | -0.09373 | 0.113741 | Other Negative Correlation |
| RNF4 | -0.09376 | 0.113598 | Other Negative Correlation |
| ZMYM5 | -0.09378 | 0.113516 | Other Negative Correlation |
| TMEM183A | -0.09379 | 0.113481 | Other Negative Correlation |
| NOXA1 | -0.09381 | 0.113405 | Other Negative Correlation |
| HYKK | -0.09384 | 0.113308 | Other Negative Correlation |
| FOLR1 | -0.09389 | 0.113097 | Other Negative Correlation |
| NR2C1 | -0.09391 | 0.113044 | Other Negative Correlation |
| SIRT1 | -0.09392 | 0.112977 | Other Negative Correlation |
| CILP | -0.09393 | 0.112969 | Other Negative Correlation |
| NMB | -0.09394 | 0.112933 | Other Negative Correlation |
| COPZ1 | -0.09394 | 0.112926 | Other Negative Correlation |
| ENSG00000214265 | -0.09396 | 0.112858 | Other Negative Correlation |
| ACVR1B | -0.09396 | 0.112856 | Other Negative Correlation |
| FAM32A | -0.09397 | 0.112802 | Other Negative Correlation |

| | | | |
|---|---|---|---|
| PTPRS | -0.09398 | 0.112755 | Other Negative Correlation |
| GFER | -0.09398 | 0.112753 | Other Negative Correlation |
| GDF11 | -0.09403 | 0.112567 | Other Negative Correlation |
| IPPK | -0.09406 | 0.112456 | Other Negative Correlation |
| C1orf56 | -0.09409 | 0.112325 | Other Negative Correlation |
| FBXO17 | -0.09414 | 0.11216 | Other Negative Correlation |
| PCDHGB1 | -0.09426 | 0.111707 | Other Negative Correlation |
| AMACR | -0.09426 | 0.111705 | Other Negative Correlation |
| OPLAH | -0.09427 | 0.111633 | Other Negative Correlation |
| NECAP1 | -0.09432 | 0.111456 | Other Negative Correlation |
| RAD54L2 | -0.09441 | 0.111126 | Other Negative Correlation |
| HIRA | -0.09442 | 0.111098 | Other Negative Correlation |
| DIAPH2 | -0.09444 | 0.111003 | Other Negative Correlation |
| ME3 | -0.09445 | 0.11096 | Other Negative Correlation |
| TGFBR3L | -0.09455 | 0.110603 | Other Negative Correlation |
| ZCCHC24 | -0.09463 | 0.110272 | Other Negative Correlation |
| CNTNAP2 | -0.0947 | 0.110022 | Other Negative Correlation |
| GCNT2 | -0.09475 | 0.109844 | Other Negative Correlation |
| PRIMPOL | -0.09475 | 0.109834 | Other Negative Correlation |
| MADD | -0.09476 | 0.109803 | Other Negative Correlation |
| CYTL1 | -0.09482 | 0.109572 | Other Negative Correlation |
| AUP1 | -0.09488 | 0.109338 | Other Negative Correlation |
| SLC38A10 | -0.0949 | 0.109276 | Other Negative Correlation |
| PPP1R42 | -0.09494 | 0.109132 | Other Negative Correlation |
| TECPR1 | -0.09498 | 0.108965 | Other Negative Correlation |
| TBC1D15 | -0.09502 | 0.108835 | Other Negative Correlation |
| ZUP1 | -0.09506 | 0.108675 | Other Negative Correlation |
| C11orf91 | -0.09512 | 0.108437 | Other Negative Correlation |
| THEGL | -0.09513 | 0.108415 | Other Negative Correlation |
| TOR1B | -0.09515 | 0.108323 | Other Negative Correlation |
| TMEM35B | -0.09516 | 0.108283 | Other Negative Correlation |
| TDRKH | -0.09522 | 0.10808 | Other Negative Correlation |
| CDC23 | -0.09522 | 0.108071 | Other Negative Correlation |
| STK11IP | -0.09524 | 0.108003 | Other Negative Correlation |
| CHID1 | -0.09527 | 0.107885 | Other Negative Correlation |
| TMEM38A | -0.09532 | 0.107712 | Other Negative Correlation |
| TRPM8 | -0.09535 | 0.10758 | Other Negative Correlation |
| TMEM45B | -0.09539 | 0.107427 | Other Negative Correlation |
| UBR3 | -0.09541 | 0.107375 | Other Negative Correlation |
| SMYD3 | -0.09543 | 0.107296 | Other Negative Correlation |
| CFAP44 | -0.09548 | 0.107106 | Other Negative Correlation |
| NPL | -0.0955 | 0.107031 | Other Negative Correlation |
| CPLANE2 | -0.0955 | 0.107021 | Other Negative Correlation |
| SLC41A1 | -0.09557 | 0.106791 | Other Negative Correlation |
| FAM241A | -0.09558 | 0.106723 | Other Negative Correlation |
| PDE4B | -0.09562 | 0.106596 | Other Negative Correlation |

| | | | |
|---|---|---|---|
| POT1 | -0.09565 | 0.10648 | Other Negative Correlation |
| CNFN | -0.09567 | 0.106425 | Other Negative Correlation |
| EFCAB6 | -0.09569 | 0.106343 | Other Negative Correlation |
| TIAM2 | -0.09569 | 0.106327 | Other Negative Correlation |
| PPP1R26 | -0.09571 | 0.106249 | Other Negative Correlation |
| EIF1AD | -0.09574 | 0.106156 | Other Negative Correlation |
| MRPS5 | -0.09576 | 0.106074 | Other Negative Correlation |
| MON1B | -0.09578 | 0.106022 | Other Negative Correlation |
| ENSG00000273291 | -0.09578 | 0.106003 | Other Negative Correlation |
| IMPA1 | -0.09582 | 0.105871 | Other Negative Correlation |
| COMMD9 | -0.09582 | 0.10585 | Other Negative Correlation |
| ZSCAN29 | -0.09583 | 0.105806 | Other Negative Correlation |
| MAP3K4 | -0.09584 | 0.105774 | Other Negative Correlation |
| FZD8 | -0.09585 | 0.105757 | Other Negative Correlation |
| CSGALNACT2 | -0.09585 | 0.10575 | Other Negative Correlation |
| KIF3C | -0.09586 | 0.105727 | Other Negative Correlation |
| TMTC4 | -0.09589 | 0.105616 | Other Negative Correlation |
| ZBTB40 | -0.09593 | 0.105469 | Other Negative Correlation |
| SLC19A2 | -0.09595 | 0.105388 | Other Negative Correlation |
| TWSG1 | -0.09601 | 0.105176 | Other Negative Correlation |
| EHMT2 | -0.09601 | 0.105157 | Other Negative Correlation |
| PMM2 | -0.09602 | 0.105142 | Other Negative Correlation |
| IL18R1 | -0.0961 | 0.104835 | Other Negative Correlation |
| CIART | -0.0961 | 0.104831 | Other Negative Correlation |
| ACVR1 | -0.09612 | 0.10478 | Other Negative Correlation |
| SMCHD1 | -0.09615 | 0.104644 | Other Negative Correlation |
| DMPK | -0.09619 | 0.104516 | Other Negative Correlation |
| RANBP10 | -0.0962 | 0.104473 | Other Negative Correlation |
| MRPL37 | -0.09625 | 0.104287 | Other Negative Correlation |
| OGT | -0.0963 | 0.104108 | Other Negative Correlation |
| UBE4A | -0.09636 | 0.10391 | Other Negative Correlation |
| METTL22 | -0.09641 | 0.103732 | Other Negative Correlation |
| PKD1L1 | -0.09643 | 0.103641 | Other Negative Correlation |
| RAF1 | -0.09647 | 0.103516 | Other Negative Correlation |
| TTBK2 | -0.09649 | 0.103442 | Other Negative Correlation |
| ANKRD42 | -0.09656 | 0.103175 | Other Negative Correlation |
| SPINK13 | -0.0966 | 0.103035 | Other Negative Correlation |
| MSTO1 | -0.09661 | 0.102983 | Other Negative Correlation |
| SPRED3 | -0.09668 | 0.102756 | Other Negative Correlation |
| PSKH1 | -0.0967 | 0.102691 | Other Negative Correlation |
| RPN1 | -0.09671 | 0.102654 | Other Negative Correlation |
| REPS2 | -0.09677 | 0.10241 | Other Negative Correlation |
| MUL1 | -0.0968 | 0.102316 | Other Negative Correlation |
| PSTPIP1 | -0.09685 | 0.102137 | Other Negative Correlation |
| RPTOR | -0.09687 | 0.102076 | Other Negative Correlation |
| CDH16 | -0.09688 | 0.102046 | Other Negative Correlation |

| | | | |
|---|---|---|---|
| GTF2E1 | -0.09694 | 0.101829 | Other Negative Correlation |
| YJU2B | -0.09694 | 0.101818 | Other Negative Correlation |
| NARS1 | -0.097 | 0.101607 | Other Negative Correlation |
| ZSCAN9 | -0.09703 | 0.101515 | Other Negative Correlation |
| NBPF3 | -0.09704 | 0.101484 | Other Negative Correlation |
| MAP1S | -0.09706 | 0.101384 | Other Negative Correlation |
| PTDSS2 | -0.09709 | 0.101301 | Other Negative Correlation |
| S100A7A | -0.0971 | 0.101241 | Other Negative Correlation |
| ATL3 | -0.09711 | 0.101234 | Other Negative Correlation |
| ZNF563 | -0.09714 | 0.101102 | Other Negative Correlation |
| SPATA2 | -0.09715 | 0.101065 | Other Negative Correlation |
| HCFC2 | -0.09723 | 0.100803 | Other Negative Correlation |
| PCDHB9 | -0.09725 | 0.100727 | Other Negative Correlation |
| B3GALT5 | -0.0973 | 0.10056 | Other Negative Correlation |
| PGBD4 | -0.09733 | 0.100446 | Other Negative Correlation |
| APOBEC3H | -0.09742 | 0.100121 | Other Negative Correlation |
| NOCT | -0.09742 | 0.100117 | Other Negative Correlation |
| C2CD3 | -0.09743 | 0.100113 | Other Negative Correlation |
| IL17RA | -0.0975 | 0.099859 | Other Negative Correlation |
| ZBED9 | -0.09751 | 0.099809 | Other Negative Correlation |
| SPNS2 | -0.09752 | 0.099777 | Other Negative Correlation |
| NEK10 | -0.09757 | 0.099599 | Other Negative Correlation |
| TMEM254 | -0.09757 | 0.099592 | Other Negative Correlation |
| KPNA7 | -0.09758 | 0.099561 | Other Negative Correlation |
| SLC9A4 | -0.09763 | 0.0994 | Other Negative Correlation |
| IFI27 | -0.09767 | 0.099248 | Other Negative Correlation |
| CHIC1 | -0.09773 | 0.099047 | Other Negative Correlation |
| CHCHD4 | -0.09774 | 0.099004 | Other Negative Correlation |
| NKAPL | -0.09774 | 0.099001 | Other Negative Correlation |
| GALK2 | -0.09777 | 0.098901 | Other Negative Correlation |
| CTXN1 | -0.0978 | 0.098802 | Other Negative Correlation |
| KIAA0753 | -0.09782 | 0.098743 | Other Negative Correlation |
| LMF1 | -0.09783 | 0.098688 | Other Negative Correlation |
| ABCD1 | -0.09784 | 0.098672 | Other Negative Correlation |
| SLC19A3 | -0.09794 | 0.098334 | Other Negative Correlation |
| SLC38A7 | -0.09794 | 0.098328 | Other Negative Correlation |
| CRPPA | -0.09797 | 0.098229 | Other Negative Correlation |
| FAM117B | -0.09802 | 0.098041 | Other Negative Correlation |
| HECTD3 | -0.09807 | 0.097884 | Other Negative Correlation |
| ABTB2 | -0.09808 | 0.097846 | Other Negative Correlation |
| USP49 | -0.0981 | 0.097772 | Other Negative Correlation |
| TRPC1 | -0.09821 | 0.097399 | Other Negative Correlation |
| PERM1 | -0.09822 | 0.097375 | Other Negative Correlation |
| TEX101 | -0.09823 | 0.097312 | Other Negative Correlation |
| FSTL5 | -0.09823 | 0.097312 | Other Negative Correlation |
| CST1 | -0.09823 | 0.097312 | Other Negative Correlation |

| | | | |
|---|---|---|---|
| UPRT | -0.09825 | 0.097264 | Other Negative Correlation |
| GAL3ST4 | -0.09835 | 0.096923 | Other Negative Correlation |
| ZNF792 | -0.09837 | 0.096839 | Other Negative Correlation |
| SOST | -0.09838 | 0.096809 | Other Negative Correlation |
| PIGB | -0.0984 | 0.096755 | Other Negative Correlation |
| COL18A1 | -0.09848 | 0.096488 | Other Negative Correlation |
| CLDN5 | -0.09856 | 0.09621 | Other Negative Correlation |
| MPND | -0.09856 | 0.096188 | Other Negative Correlation |
| ATG4C | -0.09869 | 0.095746 | Other Negative Correlation |
| BBS2 | -0.09877 | 0.095481 | Other Negative Correlation |
| USP28 | -0.09882 | 0.095319 | Other Negative Correlation |
| GTPBP3 | -0.09883 | 0.095282 | Other Negative Correlation |
| C2CD4A | -0.09885 | 0.095206 | Other Negative Correlation |
| SORCS2 | -0.09889 | 0.095104 | Other Negative Correlation |
| SLC2A11 | -0.09889 | 0.095082 | Other Negative Correlation |
| PNPLA6 | -0.09892 | 0.094982 | Other Negative Correlation |
| PRKAG2 | -0.09896 | 0.094841 | Other Negative Correlation |
| CISD2 | -0.099 | 0.094716 | Other Negative Correlation |
| CAMKK1 | -0.09903 | 0.094607 | Other Negative Correlation |
| LURAP1 | -0.09904 | 0.094584 | Other Negative Correlation |
| ZDBF2 | -0.09905 | 0.094555 | Other Negative Correlation |
| DHRS9 | -0.09907 | 0.0945 | Other Negative Correlation |
| PTPMT1 | -0.09907 | 0.094469 | Other Negative Correlation |
| DGKG | -0.09911 | 0.09435 | Other Negative Correlation |
| PLBD1 | -0.09913 | 0.094269 | Other Negative Correlation |
| SLC35G2 | -0.09915 | 0.094232 | Other Negative Correlation |
| RINT1 | -0.09916 | 0.094183 | Other Negative Correlation |
| TEX22 | -0.09923 | 0.093951 | Other Negative Correlation |
| IRAG1 | -0.09924 | 0.093909 | Other Negative Correlation |
| SPP2 | -0.0993 | 0.09372 | Other Negative Correlation |
| SLC2A5 | -0.09931 | 0.093687 | Other Negative Correlation |
| CEP78 | -0.09931 | 0.09367 | Other Negative Correlation |
| SIRT3 | -0.09935 | 0.093538 | Other Negative Correlation |
| ENSG00000277639 | -0.09938 | 0.093452 | Other Negative Correlation |
| FARSA | -0.09941 | 0.093366 | Other Negative Correlation |
| VPS54 | -0.09945 | 0.093233 | Other Negative Correlation |
| FAM104A | -0.09949 | 0.093103 | Other Negative Correlation |
| SLC2A8 | -0.09951 | 0.093024 | Other Negative Correlation |
| TLR4 | -0.09952 | 0.092982 | Other Negative Correlation |
| MBD1 | -0.09953 | 0.092956 | Other Negative Correlation |
| HLF | -0.09956 | 0.092851 | Other Negative Correlation |
| CDC34 | -0.09961 | 0.092696 | Other Negative Correlation |
| DCHS2 | -0.09962 | 0.092648 | Other Negative Correlation |
| CDK8 | -0.09964 | 0.092593 | Other Negative Correlation |
| TMOD2 | -0.09965 | 0.09255 | Other Negative Correlation |
| ALDH5A1 | -0.09966 | 0.092529 | Other Negative Correlation |

| | | | |
|---|---|---|---|
| APOBEC2 | -0.09968 | 0.092448 | Other Negative Correlation |
| ICOSLG | -0.09974 | 0.09228 | Other Negative Correlation |
| PGAM2 | -0.09974 | 0.092255 | Other Negative Correlation |
| TBC1D13 | -0.09981 | 0.092043 | Other Negative Correlation |
| KRT40 | -0.09983 | 0.091961 | Other Negative Correlation |
| ASPG | -0.09984 | 0.091922 | Other Negative Correlation |
| POU3F1 | -0.09984 | 0.091922 | Other Negative Correlation |
| FBF1 | -0.0999 | 0.091732 | Other Negative Correlation |
| MAPK7 | -0.09994 | 0.091604 | Other Negative Correlation |
| NINJ2 | -0.10005 | 0.091247 | Other Negative Correlation |
| CLDN15 | -0.1001 | 0.091083 | Other Negative Correlation |
| CAPN7 | -0.10012 | 0.091033 | Other Negative Correlation |
| HEXD | -0.10017 | 0.090876 | Other Negative Correlation |
| ZNF254 | -0.10017 | 0.090869 | Other Negative Correlation |
| TBPL1 | -0.10019 | 0.09081 | Other Negative Correlation |
| TMEM220 | -0.10028 | 0.090493 | Other Negative Correlation |
| RUSC1 | -0.10032 | 0.090391 | Other Negative Correlation |
| KCND1 | -0.10032 | 0.090375 | Other Negative Correlation |
| TEX261 | -0.10034 | 0.090302 | Other Negative Correlation |
| BCL9 | -0.10035 | 0.09028 | Other Negative Correlation |
| RAB8A | -0.10042 | 0.090065 | Other Negative Correlation |
| THAP8 | -0.10042 | 0.090056 | Other Negative Correlation |
| TOMM40L | -0.10044 | 0.089981 | Other Negative Correlation |
| LDLRAD3 | -0.10045 | 0.08995 | Other Negative Correlation |
| GMDS | -0.10046 | 0.089927 | Other Negative Correlation |
| CREB3L4 | -0.1005 | 0.089814 | Other Negative Correlation |
| CSF1 | -0.10056 | 0.089594 | Other Negative Correlation |
| ZNF468 | -0.10058 | 0.089554 | Other Negative Correlation |
| TTF1 | -0.10064 | 0.089337 | Other Negative Correlation |
| PDSS2 | -0.10083 | 0.088746 | Other Negative Correlation |
| TRIM62 | -0.10085 | 0.088689 | Other Negative Correlation |
| YTHDF2 | -0.10088 | 0.088573 | Other Negative Correlation |
| UNC93B1 | -0.1009 | 0.088533 | Other Negative Correlation |
| KLHL31 | -0.10092 | 0.088457 | Other Negative Correlation |
| SPECC1 | -0.10094 | 0.088408 | Other Negative Correlation |
| SHF | -0.10096 | 0.088337 | Other Negative Correlation |
| B3GALT9 | -0.10104 | 0.088067 | Other Negative Correlation |
| MIER3 | -0.10107 | 0.087986 | Other Negative Correlation |
| CPTP | -0.10108 | 0.087958 | Other Negative Correlation |
| ZNF142 | -0.10113 | 0.087779 | Other Negative Correlation |
| ROGDI | -0.10116 | 0.087701 | Other Negative Correlation |
| AKAP10 | -0.10116 | 0.087693 | Other Negative Correlation |
| MTMR14 | -0.10119 | 0.08759 | Other Negative Correlation |
| THY1 | -0.10125 | 0.087403 | Other Negative Correlation |
| CLOCK | -0.1013 | 0.087255 | Other Negative Correlation |
| PKIA | -0.10131 | 0.087215 | Other Negative Correlation |

| | | | |
|---|---|---|---|
| ATP10B | -0.10137 | 0.087036 | Other Negative Correlation |
| PPP1R13B | -0.10139 | 0.086984 | Other Negative Correlation |
| GSTT2B | -0.10141 | 0.086916 | Other Negative Correlation |
| MED26 | -0.10143 | 0.08686 | Other Negative Correlation |
| CBFA2T2 | -0.10146 | 0.08677 | Other Negative Correlation |
| KDM2B | -0.10147 | 0.086713 | Other Negative Correlation |
| TDRD6 | -0.10154 | 0.086497 | Other Negative Correlation |
| TFEB | -0.10158 | 0.086377 | Other Negative Correlation |
| ELMO2 | -0.1017 | 0.086022 | Other Negative Correlation |
| DICER1 | -0.10173 | 0.085921 | Other Negative Correlation |
| GGT7 | -0.10175 | 0.085856 | Other Negative Correlation |
| PHF24 | -0.10182 | 0.085631 | Other Negative Correlation |
| ATP6V1C2 | -0.10186 | 0.085517 | Other Negative Correlation |
| TC2N | -0.10192 | 0.085346 | Other Negative Correlation |
| NDUFS7 | -0.10192 | 0.085341 | Other Negative Correlation |
| STAM | -0.10192 | 0.085323 | Other Negative Correlation |
| EPHB2 | -0.10197 | 0.085192 | Other Negative Correlation |
| RNF2 | -0.10197 | 0.085189 | Other Negative Correlation |
| PLEKHA7 | -0.10198 | 0.08516 | Other Negative Correlation |
| PCDH18 | -0.102 | 0.085086 | Other Negative Correlation |
| FAM20C | -0.102 | 0.085082 | Other Negative Correlation |
| TSACC | -0.10202 | 0.085018 | Other Negative Correlation |
| ENSG00000283201 | -0.10205 | 0.084943 | Other Negative Correlation |
| CDK5R1 | -0.10205 | 0.08494 | Other Negative Correlation |
| HPSE | -0.10205 | 0.084936 | Other Negative Correlation |
| MICU3 | -0.10212 | 0.084703 | Other Negative Correlation |
| RGS19 | -0.10215 | 0.084615 | Other Negative Correlation |
| TLE3 | -0.10231 | 0.084142 | Other Negative Correlation |
| DENND2A | -0.10231 | 0.08414 | Other Negative Correlation |
| USP12 | -0.10232 | 0.084107 | Other Negative Correlation |
| TENT5A | -0.10232 | 0.084094 | Other Negative Correlation |
| EPHX1 | -0.10236 | 0.08397 | Other Negative Correlation |
| FBXO34 | -0.10237 | 0.083944 | Other Negative Correlation |
| ATXN7 | -0.10242 | 0.083795 | Other Negative Correlation |
| CEP85L | -0.10245 | 0.083712 | Other Negative Correlation |
| C17orf113 | -0.10245 | 0.083698 | Other Negative Correlation |
| MMP17 | -0.10246 | 0.083692 | Other Negative Correlation |
| TOP3A | -0.1025 | 0.083552 | Other Negative Correlation |
| CRTC3 | -0.10253 | 0.083468 | Other Negative Correlation |
| ESRP1 | -0.10255 | 0.083408 | Other Negative Correlation |
| INO80E | -0.10258 | 0.083328 | Other Negative Correlation |
| DNAJC12 | -0.10263 | 0.083181 | Other Negative Correlation |
| NHLRC4 | -0.10265 | 0.083098 | Other Negative Correlation |
| CLK1 | -0.10274 | 0.082838 | Other Negative Correlation |
| MTUS1 | -0.10275 | 0.082804 | Other Negative Correlation |
| TRIM2 | -0.10283 | 0.082577 | Other Negative Correlation |

| | | | |
|---|---|---|---|
| AGPAT4 | -0.10285 | 0.082508 | Other Negative Correlation |
| NGDN | -0.10287 | 0.082451 | Other Negative Correlation |
| DMAC2 | -0.10288 | 0.082424 | Other Negative Correlation |
| PHYHD1 | -0.10292 | 0.082293 | Other Negative Correlation |
| MRTFA | -0.10301 | 0.082031 | Other Negative Correlation |
| FASTKD5 | -0.10301 | 0.082021 | Other Negative Correlation |
| CMKLR2 | -0.10302 | 0.081991 | Other Negative Correlation |
| RBSN | -0.10302 | 0.081983 | Other Negative Correlation |
| GLA | -0.10312 | 0.081693 | Other Negative Correlation |
| NSMCE2 | -0.10313 | 0.081657 | Other Negative Correlation |
| CPE | -0.10314 | 0.081633 | Other Negative Correlation |
| TMEM107 | -0.10316 | 0.081589 | Other Negative Correlation |
| RPH3AL | -0.10321 | 0.081416 | Other Negative Correlation |
| GET3 | -0.10324 | 0.081341 | Other Negative Correlation |
| FAM120C | -0.10327 | 0.081253 | Other Negative Correlation |
| RNF19A | -0.10328 | 0.081224 | Other Negative Correlation |
| TPP1 | -0.10338 | 0.080924 | Other Negative Correlation |
| ROPN1 | -0.1034 | 0.08087 | Other Negative Correlation |
| LDB2 | -0.1034 | 0.08087 | Other Negative Correlation |
| ALK | -0.1034 | 0.08087 | Other Negative Correlation |
| DPH1 | -0.10343 | 0.080791 | Other Negative Correlation |
| DNAJC6 | -0.10349 | 0.080594 | Other Negative Correlation |
| BCL6 | -0.1035 | 0.080579 | Other Negative Correlation |
| PGBD1 | -0.10353 | 0.080484 | Other Negative Correlation |
| MXD1 | -0.10357 | 0.080361 | Other Negative Correlation |
| H4C14 | -0.10358 | 0.080343 | Other Negative Correlation |
| RMND5A | -0.10361 | 0.08025 | Other Negative Correlation |
| MAIP1 | -0.10368 | 0.080044 | Other Negative Correlation |
| PGBD2 | -0.10371 | 0.079964 | Other Negative Correlation |
| NPIPB2 | -0.10381 | 0.079663 | Other Negative Correlation |
| KAT2B | -0.10386 | 0.079529 | Other Negative Correlation |
| RAD1 | -0.10386 | 0.079509 | Other Negative Correlation |
| LCMT2 | -0.10389 | 0.079435 | Other Negative Correlation |
| CDKN2AIPNL | -0.10395 | 0.07926 | Other Negative Correlation |
| STX18 | -0.10398 | 0.079164 | Other Negative Correlation |
| PIKFYVE | -0.10405 | 0.078982 | Other Negative Correlation |
| ZNF816-ZNF321P | -0.10406 | 0.078936 | Other Negative Correlation |
| USPL1 | -0.10409 | 0.078845 | Other Negative Correlation |
| AMPD3 | -0.10412 | 0.078776 | Other Negative Correlation |
| PSG5 | -0.10412 | 0.078764 | Other Negative Correlation |
| NHLRC2 | -0.10414 | 0.078696 | Other Negative Correlation |
| CCDC14 | -0.10416 | 0.078639 | Other Negative Correlation |
| DELE1 | -0.10417 | 0.078627 | Other Negative Correlation |
| EFNA3 | -0.10418 | 0.078602 | Other Negative Correlation |
| C17orf99 | -0.10421 | 0.078501 | Other Negative Correlation |
| RMI2 | -0.10423 | 0.078456 | Other Negative Correlation |

| | | | |
|---|---|---|---|
| CLN5 | -0.10424 | 0.078431 | Other Negative Correlation |
| UMAD1 | -0.10426 | 0.078354 | Other Negative Correlation |
| TREM1 | -0.10427 | 0.078326 | Other Negative Correlation |
| HLA-DRB1 | -0.10427 | 0.078321 | Other Negative Correlation |
| BACE1 | -0.10431 | 0.078212 | Other Negative Correlation |
| ENSG00000289281 | -0.10432 | 0.078182 | Other Negative Correlation |
| BAHCC1 | -0.10434 | 0.078144 | Other Negative Correlation |
| NOTCH2NLC | -0.10439 | 0.077999 | Other Negative Correlation |
| ADPRHL1 | -0.10441 | 0.077939 | Other Negative Correlation |
| HEBP1 | -0.10446 | 0.0778 | Other Negative Correlation |
| RYR3 | -0.10447 | 0.077764 | Other Negative Correlation |
| ARFGAP1 | -0.10448 | 0.077743 | Other Negative Correlation |
| CALHM4 | -0.10451 | 0.077644 | Other Negative Correlation |
| KLHL23 | -0.10457 | 0.077468 | Other Negative Correlation |
| ARID5A | -0.10458 | 0.07745 | Other Negative Correlation |
| NRBP2 | -0.10458 | 0.077435 | Other Negative Correlation |
| PPP6R1 | -0.10459 | 0.07743 | Other Negative Correlation |
| UTY | -0.10472 | 0.077055 | Other Negative Correlation |
| KDM2A | -0.10473 | 0.077027 | Other Negative Correlation |
| TENM3 | -0.10476 | 0.076925 | Other Negative Correlation |
| BCLAF3 | -0.10476 | 0.076924 | Other Negative Correlation |
| NVL | -0.10478 | 0.076881 | Other Negative Correlation |
| RPSAP58 | -0.10481 | 0.076781 | Other Negative Correlation |
| RAB2B | -0.10482 | 0.076769 | Other Negative Correlation |
| IQCN | -0.10482 | 0.076765 | Other Negative Correlation |
| CNTN3 | -0.10485 | 0.076681 | Other Negative Correlation |
| CXCL10 | -0.10489 | 0.076555 | Other Negative Correlation |
| CDS1 | -0.1049 | 0.076529 | Other Negative Correlation |
| TAS2R20 | -0.1049 | 0.076528 | Other Negative Correlation |
| NWD1 | -0.10492 | 0.076475 | Other Negative Correlation |
| CTDSPL2 | -0.10493 | 0.07644 | Other Negative Correlation |
| FBXO27 | -0.10495 | 0.076395 | Other Negative Correlation |
| PLCXD1 | -0.10497 | 0.076347 | Other Negative Correlation |
| MUC16 | -0.10498 | 0.076301 | Other Negative Correlation |
| FBXW10B | -0.10505 | 0.076103 | Other Negative Correlation |
| MCRIP2 | -0.10506 | 0.076093 | Other Negative Correlation |
| CDK11A | -0.10506 | 0.076077 | Other Negative Correlation |
| ARF1 | -0.10508 | 0.07603 | Other Negative Correlation |
| PTPRD | -0.10512 | 0.075924 | Other Negative Correlation |
| CCDC169 | -0.10513 | 0.075889 | Other Negative Correlation |
| CALHM2 | -0.10515 | 0.075825 | Other Negative Correlation |
| SIX4 | -0.10518 | 0.075741 | Other Negative Correlation |
| SLC12A7 | -0.10523 | 0.0756 | Other Negative Correlation |
| CDK5RAP3 | -0.10524 | 0.075589 | Other Negative Correlation |
| ITPRID1 | -0.10527 | 0.0755 | Other Negative Correlation |
| FBXO16 | -0.10532 | 0.075353 | Other Negative Correlation |

| | | | |
|---|---|---|---|
| INTS1 | -0.10532 | 0.075346 | Other Negative Correlation |
| FN3KRP | -0.10538 | 0.075191 | Other Negative Correlation |
| NPM2 | -0.1054 | 0.075141 | Other Negative Correlation |
| PIERCE2 | -0.10542 | 0.075081 | Other Negative Correlation |
| SNX21 | -0.10542 | 0.075079 | Other Negative Correlation |
| ICAM2 | -0.10543 | 0.075058 | Other Negative Correlation |
| IKZF5 | -0.1055 | 0.074857 | Other Negative Correlation |
| C1orf216 | -0.10562 | 0.074523 | Other Negative Correlation |
| DNASE1 | -0.10566 | 0.074419 | Other Negative Correlation |
| PLPP4 | -0.10567 | 0.074387 | Other Negative Correlation |
| TAF1C | -0.1057 | 0.07431 | Other Negative Correlation |
| P2RX2 | -0.10571 | 0.074266 | Other Negative Correlation |
| TCTN2 | -0.10572 | 0.074238 | Other Negative Correlation |
| DNAH5 | -0.10575 | 0.074166 | Other Negative Correlation |
| ZC2HC1A | -0.10579 | 0.074065 | Other Negative Correlation |
| NRSN2 | -0.10589 | 0.073781 | Other Negative Correlation |
| PBXIP1 | -0.10589 | 0.073776 | Other Negative Correlation |
| ZNF557 | -0.10603 | 0.073398 | Other Negative Correlation |
| PLA2G2D | -0.10604 | 0.073364 | Other Negative Correlation |
| ARMCX5 | -0.10605 | 0.073335 | Other Negative Correlation |
| MCF2 | -0.10606 | 0.073318 | Other Negative Correlation |
| SYT16 | -0.10607 | 0.07328 | Other Negative Correlation |
| TGDS | -0.10616 | 0.073047 | Other Negative Correlation |
| CEBPE | -0.10621 | 0.072924 | Other Negative Correlation |
| HIP1R | -0.10624 | 0.07284 | Other Negative Correlation |
| RBBP8NL | -0.10628 | 0.072714 | Other Negative Correlation |
| ZNF658 | -0.10632 | 0.072604 | Other Negative Correlation |
| LRRC4 | -0.10633 | 0.07259 | Other Negative Correlation |
| ARHGAP39 | -0.10636 | 0.072502 | Other Negative Correlation |
| FOXN3 | -0.10639 | 0.072435 | Other Negative Correlation |
| ZNF526 | -0.1064 | 0.072389 | Other Negative Correlation |
| ZNF394 | -0.10643 | 0.072323 | Other Negative Correlation |
| AKT1 | -0.10648 | 0.07218 | Other Negative Correlation |
| MEX3D | -0.10649 | 0.072157 | Other Negative Correlation |
| MAPK12 | -0.10649 | 0.072157 | Other Negative Correlation |
| PLXNA4 | -0.10651 | 0.072104 | Other Negative Correlation |
| MT-ND2 | -0.10653 | 0.072058 | Other Negative Correlation |
| NWD2 | -0.10658 | 0.071919 | Other Negative Correlation |
| ARFGAP2 | -0.10663 | 0.071786 | Other Negative Correlation |
| SCYL1 | -0.10664 | 0.071745 | Other Negative Correlation |
| C1GALT1C1 | -0.10668 | 0.071655 | Other Negative Correlation |
| CCDC174 | -0.10669 | 0.071619 | Other Negative Correlation |
| CA14 | -0.10671 | 0.071561 | Other Negative Correlation |
| JAK2 | -0.10672 | 0.071547 | Other Negative Correlation |
| KAT14 | -0.10676 | 0.071422 | Other Negative Correlation |
| TBRG1 | -0.10678 | 0.071374 | Other Negative Correlation |

| | | | |
|---|---|---|---|
| C1orf54 | -0.10681 | 0.071288 | Other Negative Correlation |
| TMEM135 | -0.10684 | 0.071227 | Other Negative Correlation |
| TPK1 | -0.10691 | 0.071031 | Other Negative Correlation |
| PACSIN2 | -0.10692 | 0.07101 | Other Negative Correlation |
| ERVW-1 | -0.10692 | 0.071001 | Other Negative Correlation |
| SH3KBP1 | -0.10698 | 0.070855 | Other Negative Correlation |
| SDC3 | -0.107 | 0.070803 | Other Negative Correlation |
| PCDHGA7 | -0.10702 | 0.070754 | Other Negative Correlation |
| C5AR1 | -0.10702 | 0.070734 | Other Negative Correlation |
| CBLL1 | -0.10703 | 0.070702 | Other Negative Correlation |
| IQSEC1 | -0.10704 | 0.070681 | Other Negative Correlation |
| CCL27 | -0.10706 | 0.070634 | Other Negative Correlation |
| RAB11FIP4 | -0.10708 | 0.070586 | Other Negative Correlation |
| OXSM | -0.10708 | 0.070578 | Other Negative Correlation |
| CUTA | -0.10711 | 0.070505 | Other Negative Correlation |
| ZNF134 | -0.10711 | 0.070493 | Other Negative Correlation |
| ZMIZ2 | -0.10717 | 0.070346 | Other Negative Correlation |
| NME6 | -0.10718 | 0.070321 | Other Negative Correlation |
| PKIG | -0.10719 | 0.070303 | Other Negative Correlation |
| SHROOM4 | -0.10722 | 0.0702 | Other Negative Correlation |
| FETUB | -0.10724 | 0.070169 | Other Negative Correlation |
| NLGN2 | -0.10724 | 0.070163 | Other Negative Correlation |
| ZNF576 | -0.10726 | 0.070117 | Other Negative Correlation |
| GPR173 | -0.10726 | 0.070107 | Other Negative Correlation |
| ALDH3B1 | -0.1073 | 0.06999 | Other Negative Correlation |
| CDC42EP1 | -0.10731 | 0.069979 | Other Negative Correlation |
| ENSG00000188897 | -0.10732 | 0.069944 | Other Negative Correlation |
| BCL3 | -0.10736 | 0.069849 | Other Negative Correlation |
| RPRD1B | -0.10736 | 0.069835 | Other Negative Correlation |
| RAPGEF5 | -0.1074 | 0.069734 | Other Negative Correlation |
| DMRTA1 | -0.10743 | 0.069651 | Other Negative Correlation |
| ZNF229 | -0.10744 | 0.069648 | Other Negative Correlation |
| RHOJ | -0.10751 | 0.06945 | Other Negative Correlation |
| UBOX5 | -0.10752 | 0.069423 | Other Negative Correlation |
| DAP | -0.10758 | 0.069261 | Other Negative Correlation |
| LTO1 | -0.10764 | 0.069107 | Other Negative Correlation |
| LRRC28 | -0.10769 | 0.068992 | Other Negative Correlation |
| ARPIN | -0.10777 | 0.068776 | Other Negative Correlation |
| PTPN22 | -0.1078 | 0.068695 | Other Negative Correlation |
| DCP1A | -0.10783 | 0.068634 | Other Negative Correlation |
| UBE2J1 | -0.10783 | 0.068626 | Other Negative Correlation |
| GULP1 | -0.10789 | 0.068461 | Other Negative Correlation |
| EPG5 | -0.10798 | 0.068231 | Other Negative Correlation |
| SEC24D | -0.10799 | 0.06821 | Other Negative Correlation |
| MAGEE1 | -0.10799 | 0.068207 | Other Negative Correlation |
| DNAAF11 | -0.108 | 0.068183 | Other Negative Correlation |

| | | | |
|---|---|---|---|
| ATP7B | -0.10806 | 0.06802 | Other Negative Correlation |
| CDPF1 | -0.10808 | 0.06799 | Other Negative Correlation |
| MOCS3 | -0.10808 | 0.067981 | Other Negative Correlation |
| FOXS1 | -0.1081 | 0.06792 | Other Negative Correlation |
| TMEM263 | -0.10822 | 0.067627 | Other Negative Correlation |
| TTC7A | -0.10828 | 0.067463 | Other Negative Correlation |
| MTERF1 | -0.10829 | 0.06744 | Other Negative Correlation |
| ZBTB41 | -0.10836 | 0.067269 | Other Negative Correlation |
| PEX6 | -0.10839 | 0.067201 | Other Negative Correlation |
| FOXJ3 | -0.10841 | 0.06715 | Other Negative Correlation |
| BET1L | -0.10854 | 0.066812 | Other Negative Correlation |
| RP1 | -0.10862 | 0.066614 | Other Negative Correlation |
| B3GNT6 | -0.10863 | 0.06658 | Other Negative Correlation |
| CDC16 | -0.10864 | 0.066557 | Other Negative Correlation |
| ADAM19 | -0.10872 | 0.066355 | Other Negative Correlation |
| FLAD1 | -0.10873 | 0.066332 | Other Negative Correlation |
| DAZAP2 | -0.10876 | 0.066255 | Other Negative Correlation |
| GRIN2D | -0.10877 | 0.066234 | Other Negative Correlation |
| NCKIPSD | -0.10881 | 0.066127 | Other Negative Correlation |
| CCZ1B | -0.10883 | 0.066067 | Other Negative Correlation |
| TMEM94 | -0.10884 | 0.066053 | Other Negative Correlation |
| FGF11 | -0.10885 | 0.066037 | Other Negative Correlation |
| KATNAL1 | -0.10886 | 0.066014 | Other Negative Correlation |
| DIPK1A | -0.10888 | 0.065946 | Other Negative Correlation |
| VPS33B | -0.1089 | 0.065902 | Other Negative Correlation |
| SEMA7A | -0.10894 | 0.065816 | Other Negative Correlation |
| GABRP | -0.10899 | 0.065686 | Other Negative Correlation |
| CLTRN | -0.10902 | 0.065616 | Other Negative Correlation |
| TMEM171 | -0.10904 | 0.065543 | Other Negative Correlation |
| PLA2G6 | -0.10909 | 0.065439 | Other Negative Correlation |
| LAMB4 | -0.1092 | 0.065169 | Other Negative Correlation |
| YTHDF3 | -0.10921 | 0.065134 | Other Negative Correlation |
| AGPAT2 | -0.1093 | 0.064913 | Other Negative Correlation |
| DDX20 | -0.10933 | 0.064825 | Other Negative Correlation |
| UHMK1 | -0.10934 | 0.064823 | Other Negative Correlation |
| KLHL7 | -0.10935 | 0.06479 | Other Negative Correlation |
| KCTD21 | -0.10939 | 0.064701 | Other Negative Correlation |
| KHDC1L | -0.1094 | 0.064655 | Other Negative Correlation |
| LEMD3 | -0.10944 | 0.064561 | Other Negative Correlation |
| RMND5B | -0.10948 | 0.064475 | Other Negative Correlation |
| DUSP16 | -0.10954 | 0.064317 | Other Negative Correlation |
| PKP4 | -0.10958 | 0.064217 | Other Negative Correlation |
| SMUG1 | -0.1096 | 0.064171 | Other Negative Correlation |
| H2BC26 | -0.10963 | 0.064112 | Other Negative Correlation |
| KRTAP4-8 | -0.10973 | 0.063865 | Other Negative Correlation |
| EXOC1L | -0.10973 | 0.063865 | Other Negative Correlation |

| | | | |
|---|---|---|---|
| NEIL2 | -0.10974 | 0.063831 | Other Negative Correlation |
| CIB2 | -0.10976 | 0.063783 | Other Negative Correlation |
| UQCRHL | -0.10978 | 0.063732 | Other Negative Correlation |
| ENSG00000203546 | -0.10979 | 0.063714 | Other Negative Correlation |
| NFAM1 | -0.10981 | 0.063655 | Other Negative Correlation |
| SAR1B | -0.10982 | 0.063645 | Other Negative Correlation |
| ING3 | -0.10982 | 0.063633 | Other Negative Correlation |
| P3H1 | -0.10983 | 0.06362 | Other Negative Correlation |
| BMPR1A | -0.10984 | 0.06358 | Other Negative Correlation |
| CDKN2AIP | -0.10987 | 0.063515 | Other Negative Correlation |
| PET100 | -0.10991 | 0.063425 | Other Negative Correlation |
| DENND4A | -0.10992 | 0.063402 | Other Negative Correlation |
| COG8 | -0.10992 | 0.063401 | Other Negative Correlation |
| CDK19 | -0.10997 | 0.063283 | Other Negative Correlation |
| LCP1 | -0.10998 | 0.063259 | Other Negative Correlation |
| REXO1 | -0.10999 | 0.063238 | Other Negative Correlation |
| PIK3R2 | -0.10999 | 0.063216 | Other Negative Correlation |
| TBC1D12 | -0.11001 | 0.063171 | Other Negative Correlation |
| MOB3C | -0.11019 | 0.062753 | Other Negative Correlation |
| ZNF324 | -0.11024 | 0.062616 | Other Negative Correlation |
| ACKR2 | -0.11031 | 0.062464 | Other Negative Correlation |
| FLYWCH1 | -0.11034 | 0.062386 | Other Negative Correlation |
| ST6GAL1 | -0.11036 | 0.062342 | Other Negative Correlation |
| SLC33A1 | -0.11037 | 0.062324 | Other Negative Correlation |
| CCDC93 | -0.11046 | 0.062105 | Other Negative Correlation |
| CNKSR3 | -0.11046 | 0.062089 | Other Negative Correlation |
| TRMT61B | -0.11052 | 0.061967 | Other Negative Correlation |
| CD83 | -0.11055 | 0.061895 | Other Negative Correlation |
| MECP2 | -0.11056 | 0.061866 | Other Negative Correlation |
| CDK15 | -0.11057 | 0.061838 | Other Negative Correlation |
| GALC | -0.11062 | 0.06172 | Other Negative Correlation |
| SEMA4F | -0.11072 | 0.061473 | Other Negative Correlation |
| COG5 | -0.11079 | 0.061317 | Other Negative Correlation |
| ATG4D | -0.11079 | 0.061315 | Other Negative Correlation |
| ORMDL1 | -0.11082 | 0.061258 | Other Negative Correlation |
| BTC | -0.11094 | 0.060968 | Other Negative Correlation |
| ZNF791 | -0.11095 | 0.06094 | Other Negative Correlation |
| VASN | -0.11096 | 0.060929 | Other Negative Correlation |
| PACS1 | -0.11099 | 0.060856 | Other Negative Correlation |
| PNRC2 | -0.1111 | 0.060595 | Other Negative Correlation |
| FDX2 | -0.11111 | 0.060559 | Other Negative Correlation |
| IDNK | -0.11114 | 0.060505 | Other Negative Correlation |
| DNAJA4 | -0.11117 | 0.060431 | Other Negative Correlation |
| MPV17L2 | -0.11123 | 0.0603 | Other Negative Correlation |
| TM4SF19 | -0.11126 | 0.060229 | Other Negative Correlation |
| ZNF490 | -0.11128 | 0.060185 | Other Negative Correlation |

| | | | |
|---|---|---|---|
| PRR5 | -0.11133 | 0.06005 | Other Negative Correlation |
| SLC46A3 | -0.11138 | 0.059941 | Other Negative Correlation |
| RALGPS2 | -0.1114 | 0.059903 | Other Negative Correlation |
| ADCY4 | -0.11145 | 0.059789 | Other Negative Correlation |
| ISOC1 | -0.11148 | 0.059712 | Other Negative Correlation |
| ATG7 | -0.11148 | 0.059701 | Other Negative Correlation |
| ARHGEF40 | -0.11151 | 0.059641 | Other Negative Correlation |
| RNF214 | -0.11154 | 0.059585 | Other Negative Correlation |
| PHF19 | -0.11155 | 0.059556 | Other Negative Correlation |
| FRAS1 | -0.11156 | 0.059522 | Other Negative Correlation |
| PADI1 | -0.11158 | 0.059483 | Other Negative Correlation |
| RFNG | -0.1116 | 0.059429 | Other Negative Correlation |
| SLC5A3 | -0.11163 | 0.059369 | Other Negative Correlation |
| DYNC1LI1 | -0.11165 | 0.059326 | Other Negative Correlation |
| PGAP4 | -0.11167 | 0.059279 | Other Negative Correlation |
| INO80D | -0.11167 | 0.059272 | Other Negative Correlation |
| NBPF15 | -0.11171 | 0.059193 | Other Negative Correlation |
| ERCC8 | -0.11176 | 0.059073 | Other Negative Correlation |
| TM9SF1 | -0.1118 | 0.058968 | Other Negative Correlation |
| SLC35A1 | -0.11181 | 0.058966 | Other Negative Correlation |
| TMEM53 | -0.11182 | 0.058932 | Other Negative Correlation |
| DOT1L | -0.11183 | 0.0589 | Other Negative Correlation |
| UGT2B7 | -0.11184 | 0.058885 | Other Negative Correlation |
| RPP14 | -0.11186 | 0.058847 | Other Negative Correlation |
| ATAD2B | -0.11186 | 0.058835 | Other Negative Correlation |
| SH2B2 | -0.11187 | 0.058822 | Other Negative Correlation |
| HEG1 | -0.11188 | 0.058804 | Other Negative Correlation |
| EAF1 | -0.11195 | 0.058636 | Other Negative Correlation |
| TRIM46 | -0.11205 | 0.058419 | Other Negative Correlation |
| FAM47E | -0.11206 | 0.058386 | Other Negative Correlation |
| HDAC4 | -0.11209 | 0.05831 | Other Negative Correlation |
| RARG | -0.11212 | 0.058257 | Other Negative Correlation |
| PMM1 | -0.11218 | 0.058119 | Other Negative Correlation |
| ALKBH4 | -0.11225 | 0.05796 | Other Negative Correlation |
| RAC3 | -0.11228 | 0.057902 | Other Negative Correlation |
| ZMYM6 | -0.11233 | 0.057782 | Other Negative Correlation |
| ARHGEF19 | -0.11249 | 0.057432 | Other Negative Correlation |
| ZNF619 | -0.1125 | 0.057409 | Other Negative Correlation |
| PDCL | -0.1125 | 0.057402 | Other Negative Correlation |
| TSHZ3 | -0.1125 | 0.0574 | Other Negative Correlation |
| KLHDC2 | -0.11254 | 0.057304 | Other Negative Correlation |
| ZNRF2 | -0.11255 | 0.057283 | Other Negative Correlation |
| AP3S2 | -0.11258 | 0.057214 | Other Negative Correlation |
| ITPR1 | -0.11259 | 0.057202 | Other Negative Correlation |
| CBX1 | -0.11259 | 0.0572 | Other Negative Correlation |
| MPP4 | -0.11261 | 0.057158 | Other Negative Correlation |

| | | | |
|---|---|---|---|
| PPFIBP2 | -0.11266 | 0.057048 | Other Negative Correlation |
| MKRN2 | -0.1127 | 0.056966 | Other Negative Correlation |
| ZNF614 | -0.11275 | 0.056844 | Other Negative Correlation |
| PHF21A | -0.11276 | 0.056812 | Other Negative Correlation |
| SENP5 | -0.11278 | 0.056781 | Other Negative Correlation |
| MPZL3 | -0.1128 | 0.056735 | Other Negative Correlation |
| ATP6V0D2 | -0.11284 | 0.056656 | Other Negative Correlation |
| RHPN1 | -0.11285 | 0.056632 | Other Negative Correlation |
| SAXO2 | -0.11286 | 0.056596 | Other Negative Correlation |
| UACA | -0.11293 | 0.056449 | Other Negative Correlation |
| NSL1 | -0.11295 | 0.056413 | Other Negative Correlation |
| PSG6 | -0.11297 | 0.056353 | Other Negative Correlation |
| MAGI1 | -0.11299 | 0.056325 | Other Negative Correlation |
| INPP5E | -0.11304 | 0.056204 | Other Negative Correlation |
| NOD2 | -0.1131 | 0.056068 | Other Negative Correlation |
| RFX1 | -0.11314 | 0.055977 | Other Negative Correlation |
| TAF4 | -0.11317 | 0.055925 | Other Negative Correlation |
| SLC52A2 | -0.11319 | 0.055877 | Other Negative Correlation |
| RASSF4 | -0.11326 | 0.055727 | Other Negative Correlation |
| HTATSF1 | -0.11328 | 0.055688 | Other Negative Correlation |
| ZNF433 | -0.11332 | 0.055587 | Other Negative Correlation |
| LGMN | -0.11333 | 0.055581 | Other Negative Correlation |
| ABCC10 | -0.11333 | 0.055577 | Other Negative Correlation |
| FRS2 | -0.11338 | 0.055463 | Other Negative Correlation |
| DNAJB1 | -0.11339 | 0.055435 | Other Negative Correlation |
| SAMD12 | -0.11341 | 0.055402 | Other Negative Correlation |
| POLR3K | -0.11348 | 0.055247 | Other Negative Correlation |
| EEF1AKMT3 | -0.11351 | 0.055181 | Other Negative Correlation |
| SMO | -0.11352 | 0.055158 | Other Negative Correlation |
| CCNYL1 | -0.11353 | 0.055146 | Other Negative Correlation |
| SPACA6 | -0.11354 | 0.055128 | Other Negative Correlation |
| GPATCH3 | -0.11359 | 0.055008 | Other Negative Correlation |
| ALDH6A1 | -0.11361 | 0.054959 | Other Negative Correlation |
| IL20RA | -0.11362 | 0.054957 | Other Negative Correlation |
| SPCS2 | -0.11366 | 0.054853 | Other Negative Correlation |
| DHRS4 | -0.11367 | 0.054835 | Other Negative Correlation |
| SLC25A38 | -0.11369 | 0.054793 | Other Negative Correlation |
| LIPT1 | -0.1137 | 0.054781 | Other Negative Correlation |
| ACTBL2 | -0.11371 | 0.054751 | Other Negative Correlation |
| MRM3 | -0.11372 | 0.054739 | Other Negative Correlation |
| YKT6 | -0.11374 | 0.054694 | Other Negative Correlation |
| PATZ1 | -0.11381 | 0.054531 | Other Negative Correlation |
| ZNF687 | -0.11385 | 0.054459 | Other Negative Correlation |
| NHLRC1 | -0.1139 | 0.05435 | Other Negative Correlation |
| RBBP6 | -0.11393 | 0.054286 | Other Negative Correlation |
| RNF215 | -0.11397 | 0.054206 | Other Negative Correlation |

| | | | |
|---|---|---|---|
| HUWE1 | -0.11405 | 0.054037 | Other Negative Correlation |
| GPR85 | -0.1141 | 0.053928 | Other Negative Correlation |
| CCDC13 | -0.11415 | 0.053826 | Other Negative Correlation |
| DUS4L | -0.11415 | 0.053815 | Other Negative Correlation |
| VIPR1 | -0.11421 | 0.053698 | Other Negative Correlation |
| CTC1 | -0.1143 | 0.053492 | Other Negative Correlation |
| ZFYVE26 | -0.11433 | 0.053437 | Other Negative Correlation |
| RCHY1 | -0.1144 | 0.053285 | Other Negative Correlation |
| ZNF135 | -0.1144 | 0.053284 | Other Negative Correlation |
| EXD2 | -0.11447 | 0.053145 | Other Negative Correlation |
| CASTOR1 | -0.11458 | 0.052905 | Other Negative Correlation |
| ZXDB | -0.11463 | 0.052804 | Other Negative Correlation |
| ZNF596 | -0.11464 | 0.052798 | Other Negative Correlation |
| CREG1 | -0.11467 | 0.052719 | Other Negative Correlation |
| NECAB3 | -0.1147 | 0.052663 | Other Negative Correlation |
| ZNF627 | -0.1147 | 0.052658 | Other Negative Correlation |
| CHTOP | -0.11471 | 0.052653 | Other Negative Correlation |
| C12orf60 | -0.11472 | 0.052619 | Other Negative Correlation |
| STX1B | -0.11476 | 0.052542 | Other Negative Correlation |
| C18orf25 | -0.11476 | 0.052539 | Other Negative Correlation |
| XPO6 | -0.1148 | 0.052465 | Other Negative Correlation |
| ZNF408 | -0.11484 | 0.052376 | Other Negative Correlation |
| MTX3 | -0.11501 | 0.052018 | Other Negative Correlation |
| PAOX | -0.11503 | 0.051976 | Other Negative Correlation |
| CLDN14 | -0.11504 | 0.051959 | Other Negative Correlation |
| RNF40 | -0.11508 | 0.051871 | Other Negative Correlation |
| FBXO41 | -0.1152 | 0.051633 | Other Negative Correlation |
| PPP1R36 | -0.1152 | 0.051628 | Other Negative Correlation |
| KBTBD3 | -0.11534 | 0.051343 | Other Negative Correlation |
| FNBP1L | -0.11535 | 0.051332 | Other Negative Correlation |
| NALCN | -0.11537 | 0.051299 | Other Negative Correlation |
| RIPOR1 | -0.1154 | 0.051229 | Other Negative Correlation |
| NFKBID | -0.11541 | 0.051205 | Other Negative Correlation |
| ATXN3 | -0.11547 | 0.05109 | Other Negative Correlation |
| MAPKAPK2 | -0.1155 | 0.051029 | Other Negative Correlation |
| DNAJB11 | -0.11553 | 0.050974 | Other Negative Correlation |
| GBP5 | -0.11554 | 0.050936 | Other Negative Correlation |
| NKD1 | -0.11566 | 0.05071 | Other Negative Correlation |
| QPRT | -0.11571 | 0.050606 | Other Negative Correlation |
| ARSB | -0.11572 | 0.050585 | Other Negative Correlation |
| FAM24B | -0.11573 | 0.05056 | Other Negative Correlation |
| MEST | -0.11574 | 0.05055 | Other Negative Correlation |
| ALKBH1 | -0.11577 | 0.050485 | Other Negative Correlation |
| ZNF264 | -0.11578 | 0.050466 | Other Negative Correlation |
| RCBTB2 | -0.1158 | 0.050422 | Other Negative Correlation |
| GARIN2 | -0.11582 | 0.050386 | Other Negative Correlation |

| | | | |
|---|---|---|---|
| SLC11A2 | -0.11582 | 0.050383 | Other Negative Correlation |
| GK5 | -0.11587 | 0.050282 | Other Negative Correlation |
| RNF20 | -0.11595 | 0.050129 | Other Negative Correlation |
| GON7 | -0.11598 | 0.050058 | Other Negative Correlation |
| PARG | -0.11604 | 0.049943 | Other Negative Correlation |
| SIRT4 | -0.11611 | 0.049814 | Other Negative Correlation |
| ENSG00000284797 | -0.11614 | 0.049751 | Other Negative Correlation |
| PHOSPHO2 | -0.11614 | 0.049741 | Other Negative Correlation |
| CPVL | -0.11616 | 0.04971 | Other Negative Correlation |
| CYP27C1 | -0.11617 | 0.04968 | Other Negative Correlation |
| SNORC | -0.11618 | 0.049673 | Other Negative Correlation |
| CRAMP1 | -0.11626 | 0.049514 | Other Negative Correlation |
| ZC3H3 | -0.11635 | 0.049336 | Other Negative Correlation |
| SORBS3 | -0.11637 | 0.049293 | Other Negative Correlation |
| HSPA12A | -0.11646 | 0.049122 | Other Negative Correlation |
| FGF7 | -0.11646 | 0.049109 | Other Negative Correlation |
| PLEKHG6 | -0.11652 | 0.048996 | Other Negative Correlation |
| TNFSF13B | -0.11655 | 0.04894 | Other Negative Correlation |
| SEC23B | -0.1166 | 0.04884 | Other Negative Correlation |
| PCGF3 | -0.11664 | 0.048755 | Other Negative Correlation |
| RNF185 | -0.11665 | 0.04874 | Other Negative Correlation |
| DUSP23 | -0.11668 | 0.048692 | Other Negative Correlation |
| ZSCAN2 | -0.1167 | 0.048656 | Other Negative Correlation |
| TSPAN15 | -0.11673 | 0.048594 | Other Negative Correlation |
| FAM8A1 | -0.11673 | 0.048588 | Other Negative Correlation |
| DERL1 | -0.11676 | 0.048535 | Other Negative Correlation |
| KPNA5 | -0.11677 | 0.048516 | Other Negative Correlation |
| KCP | -0.11681 | 0.048443 | Other Negative Correlation |
| FAM193A | -0.11682 | 0.048423 | Other Negative Correlation |
| ZNF467 | -0.11682 | 0.048421 | Other Negative Correlation |
| POLD4 | -0.11683 | 0.048395 | Other Negative Correlation |
| CSPG4 | -0.11683 | 0.048393 | Other Negative Correlation |
| ZNF622 | -0.11686 | 0.04834 | Other Negative Correlation |
| INSL3 | -0.11691 | 0.048248 | Other Negative Correlation |
| NACC2 | -0.11697 | 0.048126 | Other Negative Correlation |
| FAAP100 | -0.11701 | 0.048042 | Other Negative Correlation |
| TULP4 | -0.11715 | 0.047788 | Other Negative Correlation |
| PI4KB | -0.11716 | 0.047766 | Other Negative Correlation |
| TRAF6 | -0.11718 | 0.04773 | Other Negative Correlation |
| CAMKMT | -0.11721 | 0.047667 | Other Negative Correlation |
| NOXO1 | -0.11727 | 0.047543 | Other Negative Correlation |
| ZNF513 | -0.11734 | 0.047408 | Other Negative Correlation |
| SPATA7 | -0.11735 | 0.047397 | Other Negative Correlation |
| CLEC7A | -0.11735 | 0.047395 | Other Negative Correlation |
| PREPL | -0.11738 | 0.047348 | Other Negative Correlation |
| FNIP2 | -0.11739 | 0.047327 | Other Negative Correlation |

| | | | |
|---|---|---|---|
| EPC2 | -0.11745 | 0.047204 | Other Negative Correlation |
| CTBP1 | -0.11746 | 0.047185 | Other Negative Correlation |
| USP3 | -0.11755 | 0.04702 | Other Negative Correlation |
| POLR3F | -0.11758 | 0.04696 | Other Negative Correlation |
| SFXN3 | -0.11762 | 0.046895 | Other Negative Correlation |
| ARL5B | -0.11765 | 0.046824 | Other Negative Correlation |
| RPS6KA2 | -0.11769 | 0.046764 | Other Negative Correlation |
| KLF9 | -0.11769 | 0.046752 | Other Negative Correlation |
| DPF3 | -0.11771 | 0.04672 | Other Negative Correlation |
| SELENOH | -0.11771 | 0.046708 | Other Negative Correlation |
| CKM | -0.11782 | 0.046519 | Other Negative Correlation |
| MYRFL | -0.11797 | 0.04624 | Other Negative Correlation |
| CLCA4 | -0.11797 | 0.046235 | Other Negative Correlation |
| SERGEF | -0.11806 | 0.046071 | Other Negative Correlation |
| CCDC97 | -0.11813 | 0.045936 | Other Negative Correlation |
| GABPA | -0.11819 | 0.045819 | Other Negative Correlation |
| C4orf48 | -0.11819 | 0.045814 | Other Negative Correlation |
| MPST | -0.1182 | 0.045812 | Other Negative Correlation |
| GEM | -0.1182 | 0.045806 | Other Negative Correlation |
| ITGB7 | -0.11821 | 0.045793 | Other Negative Correlation |
| ABCA7 | -0.11827 | 0.045668 | Other Negative Correlation |
| B4GAT1 | -0.11831 | 0.045601 | Other Negative Correlation |
| CCNT2 | -0.11841 | 0.045409 | Other Negative Correlation |
| TMEM106A | -0.11848 | 0.045286 | Other Negative Correlation |
| ZNF292 | -0.11849 | 0.045274 | Other Negative Correlation |
| HPDL | -0.1185 | 0.045244 | Other Negative Correlation |
| ATP6V0A1 | -0.11855 | 0.04517 | Other Negative Correlation |
| PRPF8 | -0.1186 | 0.045072 | Other Negative Correlation |
| TMEM65 | -0.11866 | 0.044955 | Other Negative Correlation |
| FAM149B1 | -0.11871 | 0.044861 | Other Negative Correlation |
| CABCOCO1 | -0.11876 | 0.044784 | Other Negative Correlation |
| ARHGAP40 | -0.11877 | 0.044756 | Other Negative Correlation |
| ATOSA | -0.11881 | 0.044684 | Other Negative Correlation |
| TNFAIP6 | -0.11887 | 0.044582 | Other Negative Correlation |
| SCAMP3 | -0.11887 | 0.044571 | Other Negative Correlation |
| GPATCH1 | -0.11899 | 0.044364 | Other Negative Correlation |
| MFSD12 | -0.11903 | 0.044288 | Other Negative Correlation |
| ZNF211 | -0.11904 | 0.044275 | Other Negative Correlation |
| CLPX | -0.11906 | 0.044242 | Other Negative Correlation |
| SPRR2G | -0.11907 | 0.044227 | Other Negative Correlation |
| TMEM41B | -0.11909 | 0.044181 | Other Negative Correlation |
| KRT13 | -0.11913 | 0.044108 | Other Negative Correlation |
| ZFYVE19 | -0.11919 | 0.044006 | Other Negative Correlation |
| ACP6 | -0.11919 | 0.044005 | Other Negative Correlation |
| FOXL1 | -0.11929 | 0.043821 | Other Negative Correlation |
| MYPOP | -0.11932 | 0.043778 | Other Negative Correlation |

| | | | |
|---|---|---|---|
| PPP4R3B | -0.11939 | 0.043654 | Other Negative Correlation |
| ADCY6 | -0.11948 | 0.043495 | Other Negative Correlation |
| MGRN1 | -0.11949 | 0.043476 | Other Negative Correlation |
| BATF | -0.1195 | 0.043446 | Other Negative Correlation |
| AP1M1 | -0.11957 | 0.043331 | Other Negative Correlation |
| FOXO4 | -0.11967 | 0.043163 | Other Negative Correlation |
| NCOA1 | -0.11968 | 0.043135 | Other Negative Correlation |
| TRIM41 | -0.11973 | 0.043047 | Other Negative Correlation |
| NLK | -0.11983 | 0.04288 | Other Negative Correlation |
| LAPTM4A | -0.1199 | 0.042758 | Other Negative Correlation |
| MAPRE3 | -0.1199 | 0.042754 | Other Negative Correlation |
| TUBGCP6 | -0.11993 | 0.042702 | Other Negative Correlation |
| MCMBP | -0.11997 | 0.042623 | Other Negative Correlation |
| TYW1 | -0.11999 | 0.042593 | Other Negative Correlation |
| MMAA | -0.12001 | 0.042562 | Other Negative Correlation |
| ACBD4 | -0.12001 | 0.042562 | Other Negative Correlation |
| ANKRD31 | -0.12009 | 0.042423 | Other Negative Correlation |
| DENND1B | -0.12012 | 0.04237 | Other Negative Correlation |
| ARHGEF5 | -0.12024 | 0.042169 | Other Negative Correlation |
| NOMO1 | -0.12024 | 0.042161 | Other Negative Correlation |
| C17orf80 | -0.1203 | 0.042054 | Other Negative Correlation |
| IL15 | -0.12031 | 0.042041 | Other Negative Correlation |
| SP1 | -0.12032 | 0.04202 | Other Negative Correlation |
| MDP1 | -0.12033 | 0.042001 | Other Negative Correlation |
| GOLM1 | -0.12036 | 0.041962 | Other Negative Correlation |
| TBCD | -0.12038 | 0.041919 | Other Negative Correlation |
| SMAP1 | -0.12041 | 0.041863 | Other Negative Correlation |
| KRTAP19-1 | -0.12046 | 0.041783 | Other Negative Correlation |
| DMD | -0.12049 | 0.041729 | Other Negative Correlation |
| CDC25B | -0.12049 | 0.041728 | Other Negative Correlation |
| NPLOC4 | -0.12055 | 0.041626 | Other Negative Correlation |
| RHOT2 | -0.12061 | 0.041521 | Other Negative Correlation |
| DCAF7 | -0.12062 | 0.041514 | Other Negative Correlation |
| STK38 | -0.1207 | 0.041369 | Other Negative Correlation |
| SLC25A30 | -0.12073 | 0.04132 | Other Negative Correlation |
| KRBOX5 | -0.12074 | 0.041299 | Other Negative Correlation |
| MAML1 | -0.12076 | 0.041277 | Other Negative Correlation |
| BORCS5 | -0.12078 | 0.041248 | Other Negative Correlation |
| PANK1 | -0.12078 | 0.041238 | Other Negative Correlation |
| JMJD1C | -0.12088 | 0.041078 | Other Negative Correlation |
| HLA-DQB1 | -0.12088 | 0.041064 | Other Negative Correlation |
| RAI1 | -0.12091 | 0.041023 | Other Negative Correlation |
| ZNF74 | -0.12093 | 0.040982 | Other Negative Correlation |
| ALKBH3 | -0.12109 | 0.040714 | Other Negative Correlation |
| DYNC2I1 | -0.12113 | 0.04065 | Other Negative Correlation |
| ZNF761 | -0.12115 | 0.040624 | Other Negative Correlation |

| | | | |
|---|---|---|---|
| ARL6 | -0.12115 | 0.040614 | Other Negative Correlation |
| KDM4B | -0.1212 | 0.040531 | Other Negative Correlation |
| COX7A2L | -0.12124 | 0.040465 | Other Negative Correlation |
| DCAF10 | -0.12126 | 0.040433 | Other Negative Correlation |
| GPER1 | -0.1213 | 0.040375 | Other Negative Correlation |
| SOS1 | -0.12139 | 0.040221 | Other Negative Correlation |
| DDHD2 | -0.12139 | 0.040215 | Other Negative Correlation |
| PLAC8 | -0.1216 | 0.03987 | Other Negative Correlation |
| MGP | -0.12167 | 0.039758 | Other Negative Correlation |
| LST1 | -0.12168 | 0.039741 | Other Negative Correlation |
| GPNMB | -0.1217 | 0.039709 | Other Negative Correlation |
| IL20 | -0.12172 | 0.039683 | Other Negative Correlation |
| PLPPR2 | -0.12175 | 0.039633 | Other Negative Correlation |
| SRD5A3 | -0.1218 | 0.03954 | Other Negative Correlation |
| GGCX | -0.12197 | 0.039266 | Other Negative Correlation |
| EDRF1 | -0.12199 | 0.03923 | Other Negative Correlation |
| B4GALT4 | -0.122 | 0.039219 | Other Negative Correlation |
| ABHD17B | -0.12201 | 0.039205 | Other Negative Correlation |
| VWDE | -0.12205 | 0.039145 | Other Negative Correlation |
| UBQLN2 | -0.12205 | 0.039141 | Other Negative Correlation |
| BTBD6 | -0.12207 | 0.039108 | Other Negative Correlation |
| SLC45A4 | -0.12208 | 0.039083 | Other Negative Correlation |
| WWC2 | -0.12217 | 0.038944 | Other Negative Correlation |
| KCMF1 | -0.12217 | 0.038942 | Other Negative Correlation |
| ENSG00000233757 | -0.1222 | 0.038897 | Other Negative Correlation |
| DNAH3 | -0.1222 | 0.038894 | Other Negative Correlation |
| SFT2D3 | -0.12223 | 0.038856 | Other Negative Correlation |
| NPAT | -0.12223 | 0.038842 | Other Negative Correlation |
| TMEM88B | -0.12226 | 0.038805 | Other Negative Correlation |
| SPATA6L | -0.1223 | 0.038734 | Other Negative Correlation |
| STEAP4 | -0.12231 | 0.038712 | Other Negative Correlation |
| IQCE | -0.12233 | 0.038691 | Other Negative Correlation |
| CBFB | -0.12236 | 0.038632 | Other Negative Correlation |
| FAM13A | -0.12248 | 0.038447 | Other Negative Correlation |
| TMEM71 | -0.12249 | 0.038431 | Other Negative Correlation |
| ALOX5AP | -0.12251 | 0.038399 | Other Negative Correlation |
| STXBP1 | -0.12259 | 0.038277 | Other Negative Correlation |
| TMEM81 | -0.1226 | 0.038262 | Other Negative Correlation |
| APOL3 | -0.1226 | 0.038255 | Other Negative Correlation |
| SFI1 | -0.12262 | 0.038231 | Other Negative Correlation |
| ZNF205 | -0.12263 | 0.038208 | Other Negative Correlation |
| TRMT44 | -0.12264 | 0.038194 | Other Negative Correlation |
| TRPM4 | -0.12265 | 0.038174 | Other Negative Correlation |
| SPATA12 | -0.12266 | 0.038159 | Other Negative Correlation |
| LLGL2 | -0.12271 | 0.038081 | Other Negative Correlation |
| TNFAIP8L1 | -0.12273 | 0.038045 | Other Negative Correlation |

| | | | |
|---|---|---|---|
| TRIM35 | -0.12282 | 0.037903 | Other Negative Correlation |
| PTPN6 | -0.12283 | 0.037886 | Other Negative Correlation |
| DDAH1 | -0.1229 | 0.037779 | Other Negative Correlation |
| TMEM60 | -0.12291 | 0.037764 | Other Negative Correlation |
| STAG1 | -0.12298 | 0.037658 | Other Negative Correlation |
| SULF1 | -0.12301 | 0.037608 | Other Negative Correlation |
| CEP70 | -0.12311 | 0.037454 | Other Negative Correlation |
| NAP1L5 | -0.12343 | 0.036957 | Other Negative Correlation |
| ZNF133 | -0.12343 | 0.036955 | Other Negative Correlation |
| PSAP | -0.12344 | 0.036937 | Other Negative Correlation |
| SLC27A3 | -0.12346 | 0.036918 | Other Negative Correlation |
| TBC1D17 | -0.12349 | 0.036866 | Other Negative Correlation |
| ECHDC3 | -0.12355 | 0.036766 | Other Negative Correlation |
| GAS1 | -0.12361 | 0.036686 | Other Negative Correlation |
| DISC1 | -0.12364 | 0.036635 | Other Negative Correlation |
| C1orf159 | -0.12366 | 0.036597 | Other Negative Correlation |
| ABI2 | -0.12367 | 0.036587 | Other Negative Correlation |
| TRNT1 | -0.12367 | 0.036584 | Other Negative Correlation |
| PTPRR | -0.12375 | 0.036472 | Other Negative Correlation |
| PWWP3A | -0.12379 | 0.036409 | Other Negative Correlation |
| PIDD1 | -0.1238 | 0.036385 | Other Negative Correlation |
| KCNC4 | -0.12383 | 0.036348 | Other Negative Correlation |
| IL22RA1 | -0.12384 | 0.036332 | Other Negative Correlation |
| ZSCAN32 | -0.1239 | 0.036241 | Other Negative Correlation |
| B4GALNT4 | -0.12394 | 0.036183 | Other Negative Correlation |
| TAGAP | -0.12395 | 0.036161 | Other Negative Correlation |
| PEX11A | -0.12398 | 0.036123 | Other Negative Correlation |
| PSMD8 | -0.124 | 0.036081 | Other Negative Correlation |
| SLC35A5 | -0.12403 | 0.036048 | Other Negative Correlation |
| GLI4 | -0.12407 | 0.035986 | Other Negative Correlation |
| SPEN | -0.12407 | 0.035983 | Other Negative Correlation |
| TLCD4-RWDD3 | -0.1242 | 0.035782 | Other Negative Correlation |
| DPP4 | -0.1242 | 0.03578 | Other Negative Correlation |
| MLIP | -0.12423 | 0.035744 | Other Negative Correlation |
| MTG2 | -0.12425 | 0.035704 | Other Negative Correlation |
| GOLGA5 | -0.12426 | 0.0357 | Other Negative Correlation |
| ZBTB42 | -0.12426 | 0.035691 | Other Negative Correlation |
| NIPAL2 | -0.12428 | 0.03566 | Other Negative Correlation |
| GDA | -0.12434 | 0.035575 | Other Negative Correlation |
| TBCEL | -0.12439 | 0.035502 | Other Negative Correlation |
| HILPDA | -0.12441 | 0.035477 | Other Negative Correlation |
| ZNF282 | -0.12442 | 0.035462 | Other Negative Correlation |
| MVB12A | -0.1245 | 0.035345 | Other Negative Correlation |
| PAMR1 | -0.1245 | 0.035344 | Other Negative Correlation |
| MYCBP2 | -0.12451 | 0.035328 | Other Negative Correlation |
| TARS3 | -0.12463 | 0.035151 | Other Negative Correlation |

| | | | |
|---|---|---|---|
| NPEPL1 | -0.12463 | 0.035143 | Other Negative Correlation |
| DHDDS | -0.12466 | 0.035095 | Other Negative Correlation |
| JAK3 | -0.12469 | 0.035063 | Other Negative Correlation |
| MAP1A | -0.12471 | 0.035032 | Other Negative Correlation |
| RGL1 | -0.12471 | 0.035025 | Other Negative Correlation |
| KLHL26 | -0.1248 | 0.0349 | Other Negative Correlation |
| SLC35G1 | -0.1248 | 0.034893 | Other Negative Correlation |
| IPO13 | -0.12481 | 0.034884 | Other Negative Correlation |
| PDE8A | -0.12485 | 0.034827 | Other Negative Correlation |
| USP34 | -0.12497 | 0.034644 | Other Negative Correlation |
| NDRG3 | -0.12498 | 0.034632 | Other Negative Correlation |
| LSG1 | -0.12498 | 0.034627 | Other Negative Correlation |
| INTS7 | -0.12501 | 0.034587 | Other Negative Correlation |
| TRIM52 | -0.12501 | 0.034585 | Other Negative Correlation |
| KSR1 | -0.1251 | 0.034459 | Other Negative Correlation |
| RRP8 | -0.12513 | 0.034419 | Other Negative Correlation |
| ATG14 | -0.12514 | 0.034399 | Other Negative Correlation |
| ZNF286A | -0.12518 | 0.034335 | Other Negative Correlation |
| C1orf115 | -0.12519 | 0.034323 | Other Negative Correlation |
| HOMEZ | -0.12531 | 0.034147 | Other Negative Correlation |
| ZNF837 | -0.12533 | 0.034127 | Other Negative Correlation |
| CD7 | -0.12534 | 0.034109 | Other Negative Correlation |
| KDM3A | -0.12536 | 0.034075 | Other Negative Correlation |
| RASSF7 | -0.12538 | 0.03405 | Other Negative Correlation |
| ASPHD2 | -0.12538 | 0.03405 | Other Negative Correlation |
| FBXL3 | -0.12548 | 0.033902 | Other Negative Correlation |
| PLPP6 | -0.12552 | 0.033856 | Other Negative Correlation |
| TMEM175 | -0.12553 | 0.033832 | Other Negative Correlation |
| HSD17B4 | -0.1256 | 0.033733 | Other Negative Correlation |
| RSPH3 | -0.12563 | 0.033688 | Other Negative Correlation |
| AIFM1 | -0.12564 | 0.033683 | Other Negative Correlation |
| RAB4B | -0.12565 | 0.033666 | Other Negative Correlation |
| AHSA1 | -0.12567 | 0.03363 | Other Negative Correlation |
| NFE2 | -0.12574 | 0.033533 | Other Negative Correlation |
| PRMT6 | -0.12575 | 0.033517 | Other Negative Correlation |
| LMF2 | -0.1258 | 0.033448 | Other Negative Correlation |
| TGFB3 | -0.12583 | 0.033414 | Other Negative Correlation |
| NIT1 | -0.12587 | 0.033355 | Other Negative Correlation |
| ZNRF1 | -0.12588 | 0.033337 | Other Negative Correlation |
| MAP2K5 | -0.12591 | 0.0333 | Other Negative Correlation |
| TBC1D16 | -0.12593 | 0.033273 | Other Negative Correlation |
| BET1 | -0.12599 | 0.033189 | Other Negative Correlation |
| BRD1 | -0.12601 | 0.033158 | Other Negative Correlation |
| NCOA2 | -0.12606 | 0.033083 | Other Negative Correlation |
| RIC8B | -0.12608 | 0.033061 | Other Negative Correlation |
| CMYA5 | -0.12611 | 0.033015 | Other Negative Correlation |

| | | | |
|---|---|---|---|
| ANG | -0.12626 | 0.032813 | Other Negative Correlation |
| FBXW4 | -0.12632 | 0.032724 | Other Negative Correlation |
| GPR68 | -0.12632 | 0.032717 | Other Negative Correlation |
| COPRS | -0.12635 | 0.032687 | Other Negative Correlation |
| RBAK | -0.12647 | 0.032509 | Other Negative Correlation |
| CLBA1 | -0.12648 | 0.0325 | Other Negative Correlation |
| CTU1 | -0.12651 | 0.032459 | Other Negative Correlation |
| TMEM161B | -0.12652 | 0.03245 | Other Negative Correlation |
| SECTM1 | -0.12658 | 0.032366 | Other Negative Correlation |
| VPS16 | -0.1266 | 0.032337 | Other Negative Correlation |
| ACOT11 | -0.1266 | 0.03233 | Other Negative Correlation |
| TK2 | -0.12664 | 0.032278 | Other Negative Correlation |
| MBOAT2 | -0.12675 | 0.032135 | Other Negative Correlation |
| SNX19 | -0.12677 | 0.032101 | Other Negative Correlation |
| PIP5K1C | -0.12679 | 0.032071 | Other Negative Correlation |
| ARHGAP10 | -0.12685 | 0.031987 | Other Negative Correlation |
| TRAPPC14 | -0.12686 | 0.031981 | Other Negative Correlation |
| CWF19L2 | -0.12688 | 0.031952 | Other Negative Correlation |
| CYP1A1 | -0.1269 | 0.03193 | Other Negative Correlation |
| SLC7A6 | -0.12691 | 0.031916 | Other Negative Correlation |
| TARS2 | -0.12693 | 0.031882 | Other Negative Correlation |
| ZNF362 | -0.12694 | 0.031873 | Other Negative Correlation |
| JAZF1 | -0.12703 | 0.031742 | Other Negative Correlation |
| ZP3 | -0.12706 | 0.031703 | Other Negative Correlation |
| ZNF582 | -0.1271 | 0.031659 | Other Negative Correlation |
| ZNF496 | -0.1271 | 0.031658 | Other Negative Correlation |
| INIP | -0.12714 | 0.031593 | Other Negative Correlation |
| RUSC2 | -0.12721 | 0.031511 | Other Negative Correlation |
| ACSL1 | -0.12721 | 0.031505 | Other Negative Correlation |
| UBE2B | -0.12722 | 0.031494 | Other Negative Correlation |
| NAGLU | -0.12741 | 0.031234 | Other Negative Correlation |
| PPP1R3B | -0.12745 | 0.031179 | Other Negative Correlation |
| TIGD5 | -0.12747 | 0.031162 | Other Negative Correlation |
| PCMTD2 | -0.12747 | 0.031159 | Other Negative Correlation |
| NUP133 | -0.12748 | 0.03114 | Other Negative Correlation |
| ZNF777 | -0.12752 | 0.031094 | Other Negative Correlation |
| MAPK9 | -0.12752 | 0.031084 | Other Negative Correlation |
| RPS6KC1 | -0.12753 | 0.031082 | Other Negative Correlation |
| PGBD5 | -0.12765 | 0.030916 | Other Negative Correlation |
| ALDH1B1 | -0.12767 | 0.030893 | Other Negative Correlation |
| CHM | -0.12775 | 0.030787 | Other Negative Correlation |
| NUP188 | -0.12777 | 0.030761 | Other Negative Correlation |
| FAM110C | -0.1278 | 0.03072 | Other Negative Correlation |
| USP46 | -0.12786 | 0.030646 | Other Negative Correlation |
| PLA2G15 | -0.12791 | 0.030573 | Other Negative Correlation |
| GDPD1 | -0.12791 | 0.030568 | Other Negative Correlation |

| | | | |
|---|---|---|---|
| TMEM190 | -0.12799 | 0.030468 | Other Negative Correlation |
| EPS15L1 | -0.12799 | 0.030464 | Other Negative Correlation |
| C5orf63 | -0.12802 | 0.030432 | Other Negative Correlation |
| PCSK5 | -0.12802 | 0.030424 | Other Negative Correlation |
| ATP8B1 | -0.12807 | 0.030369 | Other Negative Correlation |
| MTLN | -0.12807 | 0.030363 | Other Negative Correlation |
| ARHGAP42 | -0.12808 | 0.030358 | Other Negative Correlation |
| DNAL4 | -0.12811 | 0.030314 | Other Negative Correlation |
| SDF4 | -0.12814 | 0.03028 | Other Negative Correlation |
| ANKRD12 | -0.12814 | 0.030274 | Other Negative Correlation |
| CASQ2 | -0.12815 | 0.030261 | Other Negative Correlation |
| SLC35A2 | -0.12816 | 0.030244 | Other Negative Correlation |
| TCN2 | -0.12827 | 0.030107 | Other Negative Correlation |
| PPP4R4 | -0.12828 | 0.030087 | Other Negative Correlation |
| XRCC1 | -0.1283 | 0.030068 | Other Negative Correlation |
| CDYL | -0.12831 | 0.030047 | Other Negative Correlation |
| KMT2D | -0.12843 | 0.029903 | Other Negative Correlation |
| B4GALT7 | -0.12844 | 0.029887 | Other Negative Correlation |
| ZNF592 | -0.12854 | 0.029752 | Other Negative Correlation |
| WHAMM | -0.12854 | 0.029751 | Other Negative Correlation |
| ENSG00000262304 | -0.12854 | 0.02975 | Other Negative Correlation |
| USP39 | -0.12859 | 0.029699 | Other Negative Correlation |
| ZNF296 | -0.1286 | 0.029683 | Other Negative Correlation |
| COA7 | -0.12863 | 0.029645 | Other Negative Correlation |
| NOD1 | -0.12868 | 0.029577 | Other Negative Correlation |
| ATG2B | -0.1287 | 0.029556 | Other Negative Correlation |
| MTMR12 | -0.1287 | 0.029555 | Other Negative Correlation |
| ABCB9 | -0.12871 | 0.029534 | Other Negative Correlation |
| ANKRD27 | -0.12877 | 0.029458 | Other Negative Correlation |
| VPS11 | -0.12879 | 0.029434 | Other Negative Correlation |
| ADGRF5 | -0.1288 | 0.029421 | Other Negative Correlation |
| PLA2G4F | -0.12886 | 0.029355 | Other Negative Correlation |
| NBPF1 | -0.12898 | 0.029202 | Other Negative Correlation |
| ST6GALNAC4 | -0.129 | 0.029166 | Other Negative Correlation |
| CLEC2B | -0.12904 | 0.029127 | Other Negative Correlation |
| TCAIM | -0.12905 | 0.029109 | Other Negative Correlation |
| MLLT1 | -0.12914 | 0.029002 | Other Negative Correlation |
| C6orf47 | -0.12918 | 0.028951 | Other Negative Correlation |
| PATJ | -0.12919 | 0.028932 | Other Negative Correlation |
| MTSS2 | -0.12923 | 0.028888 | Other Negative Correlation |
| TAF5L | -0.12924 | 0.028868 | Other Negative Correlation |
| NPDC1 | -0.1293 | 0.028793 | Other Negative Correlation |
| PRKD3 | -0.12935 | 0.028736 | Other Negative Correlation |
| GTF3C4 | -0.12937 | 0.028703 | Other Negative Correlation |
| C17orf58 | -0.12938 | 0.02869 | Other Negative Correlation |
| PGGHG | -0.12941 | 0.028661 | Other Negative Correlation |

| | | | |
|---|---|---|---|
| RWDD3 | -0.12942 | 0.02865 | Other Negative Correlation |
| MKRN1 | -0.12944 | 0.028619 | Other Negative Correlation |
| SLC25A12 | -0.12959 | 0.028437 | Other Negative Correlation |
| BTBD16 | -0.12963 | 0.028389 | Other Negative Correlation |
| EDAR | -0.12964 | 0.028374 | Other Negative Correlation |
| ASXL1 | -0.12966 | 0.028355 | Other Negative Correlation |
| C6orf62 | -0.12967 | 0.028339 | Other Negative Correlation |
| VOPP1 | -0.12968 | 0.028331 | Other Negative Correlation |
| ELOVL5 | -0.1297 | 0.028305 | Other Negative Correlation |
| ITPKB | -0.12975 | 0.028239 | Other Negative Correlation |
| TCTN1 | -0.12981 | 0.028165 | Other Negative Correlation |
| B3GNT8 | -0.12982 | 0.028157 | Other Negative Correlation |
| UBASH3B | -0.12982 | 0.028149 | Other Negative Correlation |
| HSPA1B | -0.12993 | 0.028018 | Other Negative Correlation |
| MED30 | -0.12999 | 0.027947 | Other Negative Correlation |
| YIPF1 | -0.12999 | 0.027942 | Other Negative Correlation |
| TRPM7 | -0.13 | 0.02793 | Other Negative Correlation |
| KRTDAP | -0.13004 | 0.027882 | Other Negative Correlation |
| CAPN14 | -0.13007 | 0.027852 | Other Negative Correlation |
| SPG11 | -0.13008 | 0.027841 | Other Negative Correlation |
| CTSS | -0.13018 | 0.027715 | Other Negative Correlation |
| MGAT2 | -0.13022 | 0.027669 | Other Negative Correlation |
| TMTC1 | -0.13023 | 0.027653 | Other Negative Correlation |
| KLHL36 | -0.1303 | 0.027567 | Other Negative Correlation |
| IRX4 | -0.13037 | 0.027485 | Other Negative Correlation |
| RAB36 | -0.13046 | 0.027384 | Other Negative Correlation |
| SPTBN5 | -0.13057 | 0.027246 | Other Negative Correlation |
| CYP4F11 | -0.13062 | 0.02719 | Other Negative Correlation |
| ZBED4 | -0.13069 | 0.027111 | Other Negative Correlation |
| PILRB | -0.13071 | 0.027085 | Other Negative Correlation |
| SLC9A9 | -0.13077 | 0.027016 | Other Negative Correlation |
| SIDT2 | -0.13078 | 0.026998 | Other Negative Correlation |
| MSI2 | -0.13085 | 0.026919 | Other Negative Correlation |
| CBR3 | -0.13085 | 0.026912 | Other Negative Correlation |
| SOX12 | -0.13098 | 0.026769 | Other Negative Correlation |
| TXNL4B | -0.13098 | 0.02676 | Other Negative Correlation |
| MGARP | -0.13099 | 0.026752 | Other Negative Correlation |
| ACSM1 | -0.131 | 0.026742 | Other Negative Correlation |
| SLC7A8 | -0.13101 | 0.026732 | Other Negative Correlation |
| LTBP1 | -0.13102 | 0.026722 | Other Negative Correlation |
| NSD3 | -0.13104 | 0.026695 | Other Negative Correlation |
| SLC30A5 | -0.13106 | 0.026667 | Other Negative Correlation |
| CMTR2 | -0.13107 | 0.026666 | Other Negative Correlation |
| DENND5A | -0.13108 | 0.026647 | Other Negative Correlation |
| FAM136A | -0.13117 | 0.026538 | Other Negative Correlation |
| TCAF2 | -0.13119 | 0.026523 | Other Negative Correlation |

| | | | |
|---|---|---|---|
| GTF2IRD1 | -0.13119 | 0.02652 | Other Negative Correlation |
| ZNF350 | -0.1312 | 0.026509 | Other Negative Correlation |
| ATG4B | -0.13126 | 0.026444 | Other Negative Correlation |
| SAMD4A | -0.13129 | 0.026407 | Other Negative Correlation |
| ASB3 | -0.13138 | 0.026303 | Other Negative Correlation |
| RPAIN | -0.13139 | 0.02629 | Other Negative Correlation |
| LRP6 | -0.13141 | 0.026266 | Other Negative Correlation |
| D2HGDH | -0.13145 | 0.026218 | Other Negative Correlation |
| ZNF654 | -0.13146 | 0.026212 | Other Negative Correlation |
| ISY1 | -0.13148 | 0.02618 | Other Negative Correlation |
| KCNK7 | -0.13159 | 0.026063 | Other Negative Correlation |
| YIPF2 | -0.13162 | 0.026022 | Other Negative Correlation |
| TAB2 | -0.13164 | 0.026004 | Other Negative Correlation |
| ZNF558 | -0.13169 | 0.025944 | Other Negative Correlation |
| GNB5 | -0.13174 | 0.025883 | Other Negative Correlation |
| SPIDR | -0.13178 | 0.025848 | Other Negative Correlation |
| PCGF2 | -0.13193 | 0.025668 | Other Negative Correlation |
| SETD1A | -0.13196 | 0.025639 | Other Negative Correlation |
| LANCL3 | -0.13198 | 0.025619 | Other Negative Correlation |
| CFAP46 | -0.132 | 0.025588 | Other Negative Correlation |
| NR6A1 | -0.13203 | 0.025565 | Other Negative Correlation |
| GORASP1 | -0.13209 | 0.025497 | Other Negative Correlation |
| ZDHHC21 | -0.13212 | 0.025455 | Other Negative Correlation |
| ANKRD28 | -0.13215 | 0.025428 | Other Negative Correlation |
| IP6K1 | -0.13217 | 0.025402 | Other Negative Correlation |
| TRAPPC6A | -0.13218 | 0.025388 | Other Negative Correlation |
| CASP14 | -0.13224 | 0.02533 | Other Negative Correlation |
| RHBDD2 | -0.13227 | 0.025295 | Other Negative Correlation |
| MYO3B | -0.13228 | 0.025278 | Other Negative Correlation |
| ZRSR2 | -0.13229 | 0.025272 | Other Negative Correlation |
| PYROXD2 | -0.13231 | 0.025245 | Other Negative Correlation |
| TEAD3 | -0.13245 | 0.025093 | Other Negative Correlation |
| SPAG16 | -0.13247 | 0.025071 | Other Negative Correlation |
| CYFIP2 | -0.13248 | 0.025056 | Other Negative Correlation |
| SLC12A8 | -0.1325 | 0.025037 | Other Negative Correlation |
| RNF34 | -0.13251 | 0.025029 | Other Negative Correlation |
| RIC1 | -0.13254 | 0.024987 | Other Negative Correlation |
| PRODH | -0.13256 | 0.024964 | Other Negative Correlation |
| ITPRIP | -0.13258 | 0.024946 | Other Negative Correlation |
| PSME4 | -0.13261 | 0.024919 | Other Negative Correlation |
| BRD8 | -0.13266 | 0.024858 | Other Negative Correlation |
| ANO8 | -0.13272 | 0.024799 | Other Negative Correlation |
| CCDC22 | -0.13277 | 0.024737 | Other Negative Correlation |
| FZR1 | -0.13281 | 0.024699 | Other Negative Correlation |
| SEPSECS | -0.13289 | 0.024612 | Other Negative Correlation |
| GCNA | -0.13289 | 0.024609 | Other Negative Correlation |

| | | | |
|---|---|---|---|
| TTLL5 | -0.13293 | 0.02456 | Other Negative Correlation |
| CABIN1 | -0.13294 | 0.024555 | Other Negative Correlation |
| C12orf4 | -0.13302 | 0.024469 | Other Negative Correlation |
| BICRA | -0.13313 | 0.024343 | Other Negative Correlation |
| SRBD1 | -0.13315 | 0.024327 | Other Negative Correlation |
| COPG1 | -0.1332 | 0.024269 | Other Negative Correlation |
| GXYLT2 | -0.13326 | 0.024204 | Other Negative Correlation |
| SYCP2L | -0.13327 | 0.0242 | Other Negative Correlation |
| KCTD17 | -0.13329 | 0.024177 | Other Negative Correlation |
| WNT7B | -0.1333 | 0.024164 | Other Negative Correlation |
| COPE | -0.13333 | 0.02413 | Other Negative Correlation |
| FAM83F | -0.13337 | 0.024085 | Other Negative Correlation |
| NHSL2 | -0.13341 | 0.024049 | Other Negative Correlation |
| LZTS3 | -0.13342 | 0.024035 | Other Negative Correlation |
| PANK3 | -0.13344 | 0.02401 | Other Negative Correlation |
| PIWIL2 | -0.13345 | 0.024002 | Other Negative Correlation |
| SNAP29 | -0.13346 | 0.023994 | Other Negative Correlation |
| ACADM | -0.13348 | 0.023976 | Other Negative Correlation |
| HDAC11 | -0.13351 | 0.023942 | Other Negative Correlation |
| SLC4A11 | -0.13352 | 0.023929 | Other Negative Correlation |
| TRAF1 | -0.13359 | 0.023859 | Other Negative Correlation |
| MAP2 | -0.13359 | 0.023856 | Other Negative Correlation |
| TMEM241 | -0.13363 | 0.023815 | Other Negative Correlation |
| FBRS | -0.13368 | 0.023756 | Other Negative Correlation |
| C3orf52 | -0.13375 | 0.02369 | Other Negative Correlation |
| ADAMTSL5 | -0.13375 | 0.023688 | Other Negative Correlation |
| XG | -0.13378 | 0.023655 | Other Negative Correlation |
| PLEKHA6 | -0.13386 | 0.023572 | Other Negative Correlation |
| C6orf52 | -0.13386 | 0.02357 | Other Negative Correlation |
| HTRA2 | -0.1339 | 0.023531 | Other Negative Correlation |
| CHSY1 | -0.13391 | 0.023519 | Other Negative Correlation |
| APBB3 | -0.13392 | 0.023511 | Other Negative Correlation |
| TBC1D4 | -0.13392 | 0.023509 | Other Negative Correlation |
| ARL11 | -0.13393 | 0.023498 | Other Negative Correlation |
| CD40 | -0.13398 | 0.023442 | Other Negative Correlation |
| ZNF354A | -0.134 | 0.023423 | Other Negative Correlation |
| CDC42BPG | -0.1341 | 0.02332 | Other Negative Correlation |
| LCOR | -0.13412 | 0.023303 | Other Negative Correlation |
| TGIF2 | -0.13421 | 0.023205 | Other Negative Correlation |
| TNFRSF10C | -0.13423 | 0.023181 | Other Negative Correlation |
| TNNC1 | -0.13431 | 0.023101 | Other Negative Correlation |
| VPS50 | -0.13433 | 0.023085 | Other Negative Correlation |
| ODAPH | -0.13438 | 0.023034 | Other Negative Correlation |
| ARHGEF18 | -0.13438 | 0.023028 | Other Negative Correlation |
| GJA5 | -0.13439 | 0.023022 | Other Negative Correlation |
| DEPTOR | -0.13444 | 0.022968 | Other Negative Correlation |

| | | | |
|---|---|---|---|
| DCAF12 | -0.13447 | 0.022935 | Other Negative Correlation |
| TMPRSS11F | -0.13448 | 0.022926 | Other Negative Correlation |
| ADGRD2 | -0.1345 | 0.022908 | Other Negative Correlation |
| DMWD | -0.13457 | 0.022832 | Other Negative Correlation |
| CRIPT | -0.13464 | 0.022761 | Other Negative Correlation |
| HNMT | -0.1347 | 0.022703 | Other Negative Correlation |
| ZNF416 | -0.13474 | 0.022661 | Other Negative Correlation |
| PISD | -0.13475 | 0.02265 | Other Negative Correlation |
| ATP23 | -0.13477 | 0.022634 | Other Negative Correlation |
| RAB11FIP5 | -0.1348 | 0.022599 | Other Negative Correlation |
| WRN | -0.13481 | 0.022588 | Other Negative Correlation |
| IL1RL1 | -0.13483 | 0.022575 | Other Negative Correlation |
| CYBRD1 | -0.13487 | 0.022535 | Other Negative Correlation |
| COG6 | -0.1349 | 0.022501 | Other Negative Correlation |
| NPHP3 | -0.13492 | 0.02248 | Other Negative Correlation |
| CNOT3 | -0.13506 | 0.022339 | Other Negative Correlation |
| RIPOR3 | -0.13513 | 0.022269 | Other Negative Correlation |
| RHPN2 | -0.13515 | 0.022252 | Other Negative Correlation |
| SLC27A4 | -0.13518 | 0.022215 | Other Negative Correlation |
| OCA2 | -0.13522 | 0.022175 | Other Negative Correlation |
| MKRN3 | -0.13524 | 0.022158 | Other Negative Correlation |
| LRRC27 | -0.13526 | 0.022138 | Other Negative Correlation |
| MOCS2 | -0.13537 | 0.022032 | Other Negative Correlation |
| IFI27L1 | -0.13537 | 0.022028 | Other Negative Correlation |
| RASGEF1A | -0.13538 | 0.022022 | Other Negative Correlation |
| INPPL1 | -0.13541 | 0.021995 | Other Negative Correlation |
| NSMCE3 | -0.13557 | 0.021834 | Other Negative Correlation |
| NRBP1 | -0.13559 | 0.021811 | Other Negative Correlation |
| SLC5A1 | -0.1356 | 0.021808 | Other Negative Correlation |
| TULP3 | -0.1356 | 0.021803 | Other Negative Correlation |
| STARD4 | -0.13562 | 0.021784 | Other Negative Correlation |
| MARS2 | -0.13569 | 0.021719 | Other Negative Correlation |
| WASHC2A | -0.13575 | 0.021654 | Other Negative Correlation |
| ZC3H8 | -0.13582 | 0.02159 | Other Negative Correlation |
| CALB1 | -0.13586 | 0.021548 | Other Negative Correlation |
| A1BG | -0.13586 | 0.021547 | Other Negative Correlation |
| ZNF91 | -0.1359 | 0.02151 | Other Negative Correlation |
| SART3 | -0.13591 | 0.021497 | Other Negative Correlation |
| LRATD2 | -0.13592 | 0.021493 | Other Negative Correlation |
| DCP1B | -0.13593 | 0.021482 | Other Negative Correlation |
| ARID2 | -0.13597 | 0.021439 | Other Negative Correlation |
| CARD8 | -0.13606 | 0.021353 | Other Negative Correlation |
| DISP3 | -0.1361 | 0.021317 | Other Negative Correlation |
| ERI2 | -0.13621 | 0.021215 | Other Negative Correlation |
| DENND1A | -0.13625 | 0.021173 | Other Negative Correlation |
| TRIOBP | -0.13645 | 0.020979 | Other Negative Correlation |

| | | | |
|---|---|---|---|
| DDX24 | -0.13649 | 0.020944 | Other Negative Correlation |
| PLA2G7 | -0.13657 | 0.020867 | Other Negative Correlation |
| ZNF525 | -0.13663 | 0.020811 | Other Negative Correlation |
| AGPAT1 | -0.13663 | 0.02081 | Other Negative Correlation |
| SOCS5 | -0.13669 | 0.020758 | Other Negative Correlation |
| RBM5 | -0.13671 | 0.020736 | Other Negative Correlation |
| LCA5 | -0.13673 | 0.020717 | Other Negative Correlation |
| JRKL | -0.13674 | 0.020712 | Other Negative Correlation |
| PRKCD | -0.13677 | 0.020685 | Other Negative Correlation |
| EPHB3 | -0.13678 | 0.020668 | Other Negative Correlation |
| TMEM204 | -0.13679 | 0.020663 | Other Negative Correlation |
| REL | -0.13684 | 0.020619 | Other Negative Correlation |
| CREG2 | -0.13689 | 0.020573 | Other Negative Correlation |
| SMCR8 | -0.13689 | 0.020566 | Other Negative Correlation |
| ANKRD34A | -0.13693 | 0.020536 | Other Negative Correlation |
| ADAT1 | -0.13693 | 0.020533 | Other Negative Correlation |
| AFTPH | -0.13696 | 0.020501 | Other Negative Correlation |
| RABEP2 | -0.13699 | 0.020476 | Other Negative Correlation |
| MTFMT | -0.1371 | 0.020376 | Other Negative Correlation |
| DYNLRB1 | -0.13711 | 0.020362 | Other Negative Correlation |
| CD52 | -0.13716 | 0.020319 | Other Negative Correlation |
| TGM5 | -0.13717 | 0.020312 | Other Negative Correlation |
| ELAC1 | -0.13723 | 0.020255 | Other Negative Correlation |
| TIGAR | -0.13724 | 0.020241 | Other Negative Correlation |
| ATF7IP | -0.13732 | 0.020174 | Other Negative Correlation |
| FAT4 | -0.13733 | 0.020167 | Other Negative Correlation |
| PPIE | -0.13735 | 0.020147 | Other Negative Correlation |
| ZNF175 | -0.1374 | 0.020099 | Other Negative Correlation |
| BMP2K | -0.13742 | 0.020079 | Other Negative Correlation |
| PRAF2 | -0.13743 | 0.020068 | Other Negative Correlation |
| MSMB | -0.13757 | 0.019948 | Other Negative Correlation |
| BRAP | -0.13758 | 0.019935 | Other Negative Correlation |
| HLA-DRA | -0.13759 | 0.019927 | Other Negative Correlation |
| ART3 | -0.13773 | 0.019799 | Other Negative Correlation |
| SLC28A3 | -0.13774 | 0.01979 | Other Negative Correlation |
| PCDHGB6 | -0.13777 | 0.01976 | Other Negative Correlation |
| OXR1 | -0.13778 | 0.019752 | Other Negative Correlation |
| ZNF517 | -0.13779 | 0.019746 | Other Negative Correlation |
| TMEM25 | -0.13783 | 0.019712 | Other Negative Correlation |
| ZBTB8A | -0.13786 | 0.019679 | Other Negative Correlation |
| PSCA | -0.13789 | 0.019657 | Other Negative Correlation |
| UNG | -0.13795 | 0.019599 | Other Negative Correlation |
| AVEN | -0.13799 | 0.019567 | Other Negative Correlation |
| FHIP1B | -0.13801 | 0.01955 | Other Negative Correlation |
| RNF139 | -0.13816 | 0.01941 | Other Negative Correlation |
| TBC1D22B | -0.13818 | 0.0194 | Other Negative Correlation |

| | | | |
|---|---|---|---|
| AGMAT | -0.13819 | 0.019389 | Other Negative Correlation |
| TOGARAM1 | -0.1382 | 0.019374 | Other Negative Correlation |
| SLC25A43 | -0.13822 | 0.019365 | Other Negative Correlation |
| SLC35C1 | -0.13824 | 0.019341 | Other Negative Correlation |
| MLLT6 | -0.13833 | 0.019261 | Other Negative Correlation |
| MOSMO | -0.13839 | 0.01921 | Other Negative Correlation |
| CUL1 | -0.13841 | 0.019189 | Other Negative Correlation |
| TAF8 | -0.13847 | 0.019141 | Other Negative Correlation |
| ALOXE3 | -0.13852 | 0.019099 | Other Negative Correlation |
| CDK9 | -0.13863 | 0.019003 | Other Negative Correlation |
| ENTPD6 | -0.13863 | 0.018996 | Other Negative Correlation |
| MLLT3 | -0.13865 | 0.018987 | Other Negative Correlation |
| HDAC9 | -0.13867 | 0.018966 | Other Negative Correlation |
| ZNF548 | -0.13868 | 0.018953 | Other Negative Correlation |
| GJC2 | -0.13883 | 0.018829 | Other Negative Correlation |
| SKI | -0.13884 | 0.01882 | Other Negative Correlation |
| ZNF260 | -0.13884 | 0.018817 | Other Negative Correlation |
| FNTB | -0.1389 | 0.018769 | Other Negative Correlation |
| TLR5 | -0.13893 | 0.018744 | Other Negative Correlation |
| CEP57L1 | -0.13894 | 0.018736 | Other Negative Correlation |
| TSSC4 | -0.13908 | 0.018611 | Other Negative Correlation |
| CDKAL1 | -0.13909 | 0.018607 | Other Negative Correlation |
| RCOR3 | -0.13911 | 0.01859 | Other Negative Correlation |
| CRISPLD1 | -0.13914 | 0.018558 | Other Negative Correlation |
| PFDN6 | -0.13917 | 0.018535 | Other Negative Correlation |
| CALHM5 | -0.13931 | 0.018416 | Other Negative Correlation |
| CCDC60 | -0.13932 | 0.018408 | Other Negative Correlation |
| ATP1B3 | -0.13933 | 0.018404 | Other Negative Correlation |
| TCF12 | -0.13934 | 0.01839 | Other Negative Correlation |
| PAN3 | -0.13935 | 0.01838 | Other Negative Correlation |
| DNASE1L1 | -0.13938 | 0.018361 | Other Negative Correlation |
| CLDN19 | -0.13938 | 0.018356 | Other Negative Correlation |
| ADGRG6 | -0.13939 | 0.018347 | Other Negative Correlation |
| OSTM1 | -0.1394 | 0.018339 | Other Negative Correlation |
| ZNF487 | -0.13942 | 0.018327 | Other Negative Correlation |
| CRYZ | -0.13944 | 0.018303 | Other Negative Correlation |
| GZF1 | -0.13945 | 0.018297 | Other Negative Correlation |
| RANBP6 | -0.13949 | 0.018261 | Other Negative Correlation |
| TMEM217 | -0.13952 | 0.018236 | Other Negative Correlation |
| EFEMP2 | -0.13953 | 0.018232 | Other Negative Correlation |
| SNRK | -0.13953 | 0.018229 | Other Negative Correlation |
| CLIP2 | -0.13954 | 0.018227 | Other Negative Correlation |
| VIPAS39 | -0.13959 | 0.018181 | Other Negative Correlation |
| CREB3L1 | -0.13964 | 0.01814 | Other Negative Correlation |
| CCL26 | -0.13969 | 0.018096 | Other Negative Correlation |
| SHROOM1 | -0.13975 | 0.018051 | Other Negative Correlation |

| | | | |
|---|---|---|---|
| ZFPM1 | -0.13975 | 0.018046 | Other Negative Correlation |
| DLG3 | -0.1398 | 0.018004 | Other Negative Correlation |
| TMX2 | -0.13984 | 0.017975 | Other Negative Correlation |
| GJD3 | -0.13985 | 0.017967 | Other Negative Correlation |
| SLC39A13 | -0.13987 | 0.017951 | Other Negative Correlation |
| KLF8 | -0.13994 | 0.017889 | Other Negative Correlation |
| TMEM54 | -0.14007 | 0.017785 | Other Negative Correlation |
| PRUNE1 | -0.14007 | 0.017781 | Other Negative Correlation |
| BLTP3B | -0.14012 | 0.017738 | Other Negative Correlation |
| CEP57 | -0.14018 | 0.017688 | Other Negative Correlation |
| CPSF7 | -0.1402 | 0.017677 | Other Negative Correlation |
| SIDT1 | -0.14022 | 0.017662 | Other Negative Correlation |
| GBF1 | -0.14026 | 0.017626 | Other Negative Correlation |
| PXYLP1 | -0.14027 | 0.017617 | Other Negative Correlation |
| POGK | -0.14028 | 0.01761 | Other Negative Correlation |
| HMG20A | -0.14034 | 0.017565 | Other Negative Correlation |
| STAT2 | -0.14045 | 0.01747 | Other Negative Correlation |
| GPD1L | -0.14051 | 0.017426 | Other Negative Correlation |
| CDAN1 | -0.14051 | 0.017423 | Other Negative Correlation |
| METTL25 | -0.14063 | 0.017327 | Other Negative Correlation |
| APBA3 | -0.14064 | 0.017322 | Other Negative Correlation |
| HCG22 | -0.14067 | 0.017294 | Other Negative Correlation |
| CCS | -0.14075 | 0.017232 | Other Negative Correlation |
| BDKRB1 | -0.14077 | 0.017213 | Other Negative Correlation |
| TRAPPC11 | -0.14084 | 0.01716 | Other Negative Correlation |
| CBLN3 | -0.14084 | 0.017156 | Other Negative Correlation |
| CEP89 | -0.14085 | 0.017148 | Other Negative Correlation |
| STK32C | -0.14104 | 0.017002 | Other Negative Correlation |
| CDIPT | -0.14108 | 0.016965 | Other Negative Correlation |
| DUS2 | -0.14111 | 0.016946 | Other Negative Correlation |
| ANKRD26 | -0.14112 | 0.016939 | Other Negative Correlation |
| CABYR | -0.14114 | 0.016922 | Other Negative Correlation |
| SEC13 | -0.14122 | 0.016857 | Other Negative Correlation |
| RILP | -0.14123 | 0.01685 | Other Negative Correlation |
| TIMM10B | -0.14125 | 0.016834 | Other Negative Correlation |
| CC2D1B | -0.14129 | 0.016805 | Other Negative Correlation |
| STX17 | -0.14132 | 0.016777 | Other Negative Correlation |
| DVL1 | -0.14134 | 0.016768 | Other Negative Correlation |
| IPO11 | -0.14137 | 0.016741 | Other Negative Correlation |
| RNF123 | -0.14137 | 0.01674 | Other Negative Correlation |
| UBR2 | -0.14144 | 0.016687 | Other Negative Correlation |
| PPP3R1 | -0.1415 | 0.016641 | Other Negative Correlation |
| REEP2 | -0.14154 | 0.016612 | Other Negative Correlation |
| SDR42E1 | -0.14156 | 0.016595 | Other Negative Correlation |
| SHLD1 | -0.14164 | 0.016531 | Other Negative Correlation |
| DDX41 | -0.14178 | 0.016424 | Other Negative Correlation |

| | | | |
|---|---|---|---|
| VAC14 | -0.1419 | 0.016333 | Other Negative Correlation |
| MLX | -0.14191 | 0.016324 | Other Negative Correlation |
| ANKRD1 | -0.14195 | 0.016295 | Other Negative Correlation |
| UNC45A | -0.14198 | 0.016274 | Other Negative Correlation |
| SBNO2 | -0.14202 | 0.016241 | Other Negative Correlation |
| LRFN3 | -0.14204 | 0.016227 | Other Negative Correlation |
| RNF146 | -0.14207 | 0.016204 | Other Negative Correlation |
| TLR1 | -0.14215 | 0.016143 | Other Negative Correlation |
| CDK13 | -0.14216 | 0.016136 | Other Negative Correlation |
| ANKRD13D | -0.1422 | 0.016108 | Other Negative Correlation |
| ENG | -0.14257 | 0.015831 | Other Negative Correlation |
| CREBL2 | -0.14259 | 0.015813 | Other Negative Correlation |
| SWSAP1 | -0.14273 | 0.015712 | Other Negative Correlation |
| BTAF1 | -0.14279 | 0.015664 | Other Negative Correlation |
| SNX4 | -0.14279 | 0.015663 | Other Negative Correlation |
| TNS2 | -0.14286 | 0.015613 | Other Negative Correlation |
| TIFA | -0.14287 | 0.015604 | Other Negative Correlation |
| PBX1 | -0.14288 | 0.015602 | Other Negative Correlation |
| ZNF358 | -0.14296 | 0.015544 | Other Negative Correlation |
| ODF3L1 | -0.14298 | 0.015529 | Other Negative Correlation |
| DHTKD1 | -0.14306 | 0.015469 | Other Negative Correlation |
| PCDHB2 | -0.14306 | 0.015466 | Other Negative Correlation |
| COX10 | -0.14308 | 0.015456 | Other Negative Correlation |
| C11orf71 | -0.14308 | 0.015455 | Other Negative Correlation |
| PANO1 | -0.1431 | 0.015438 | Other Negative Correlation |
| BAG6 | -0.14315 | 0.015403 | Other Negative Correlation |
| BTBD3 | -0.14317 | 0.015389 | Other Negative Correlation |
| MBD3 | -0.14318 | 0.01538 | Other Negative Correlation |
| ZNF174 | -0.14325 | 0.015333 | Other Negative Correlation |
| MYMX | -0.14331 | 0.015286 | Other Negative Correlation |
| SMIM6 | -0.14332 | 0.015279 | Other Negative Correlation |
| CCDC192 | -0.14334 | 0.015266 | Other Negative Correlation |
| ZNF165 | -0.14336 | 0.01525 | Other Negative Correlation |
| SEC22A | -0.14336 | 0.015249 | Other Negative Correlation |
| SCARA3 | -0.14339 | 0.015231 | Other Negative Correlation |
| TBC1D8 | -0.14343 | 0.015202 | Other Negative Correlation |
| ANKS6 | -0.14345 | 0.015188 | Other Negative Correlation |
| TMEM185A | -0.14347 | 0.015173 | Other Negative Correlation |
| CLASP2 | -0.14347 | 0.015172 | Other Negative Correlation |
| BATF2 | -0.14348 | 0.015164 | Other Negative Correlation |
| FAM200A | -0.14348 | 0.015163 | Other Negative Correlation |
| PJA1 | -0.1435 | 0.015152 | Other Negative Correlation |
| POGLUT2 | -0.14356 | 0.01511 | Other Negative Correlation |
| PLEKHF1 | -0.14359 | 0.015088 | Other Negative Correlation |
| NBPF14 | -0.14363 | 0.015057 | Other Negative Correlation |
| ERAP1 | -0.14367 | 0.015029 | Other Negative Correlation |

| | | | |
|---|---|---|---|
| OTUD4 | -0.1437 | 0.015007 | Other Negative Correlation |
| C1orf43 | -0.14371 | 0.015004 | Other Negative Correlation |
| PAG1 | -0.14371 | 0.015003 | Other Negative Correlation |
| GNPAT | -0.14371 | 0.015002 | Other Negative Correlation |
| GNPDA2 | -0.14373 | 0.01499 | Other Negative Correlation |
| BRD3OS | -0.14374 | 0.014982 | Other Negative Correlation |
| GPX7 | -0.14375 | 0.014974 | Other Negative Correlation |
| ANKK1 | -0.14375 | 0.014972 | Other Negative Correlation |
| CBL | -0.14375 | 0.014972 | Other Negative Correlation |
| CORO6 | -0.14376 | 0.014965 | Other Negative Correlation |
| MOSPD1 | -0.14378 | 0.014954 | Other Negative Correlation |
| ZNF28 | -0.14384 | 0.014912 | Other Negative Correlation |
| PAXIP1 | -0.14386 | 0.014896 | Other Negative Correlation |
| SLC45A3 | -0.14389 | 0.014875 | Other Negative Correlation |
| UBL7 | -0.14391 | 0.014861 | Other Negative Correlation |
| MAPRE2 | -0.14394 | 0.014836 | Other Negative Correlation |
| SPON2 | -0.14398 | 0.014812 | Other Negative Correlation |
| ALOX12 | -0.14404 | 0.014771 | Other Negative Correlation |
| IGSF22 | -0.14405 | 0.014761 | Other Negative Correlation |
| ANXA9 | -0.14414 | 0.014701 | Other Negative Correlation |
| AMPD2 | -0.14417 | 0.014677 | Other Negative Correlation |
| TMEM234 | -0.14424 | 0.014632 | Other Negative Correlation |
| LSR | -0.14425 | 0.014626 | Other Negative Correlation |
| MIPEP | -0.14427 | 0.014607 | Other Negative Correlation |
| SHROOM2 | -0.14433 | 0.014566 | Other Negative Correlation |
| SPPL2A | -0.14437 | 0.014541 | Other Negative Correlation |
| CRYZL1 | -0.14438 | 0.014536 | Other Negative Correlation |
| USP54 | -0.14439 | 0.014527 | Other Negative Correlation |
| PPP1R9B | -0.1445 | 0.01445 | Other Negative Correlation |
| TMED9 | -0.1445 | 0.014447 | Other Negative Correlation |
| TTC7B | -0.14461 | 0.014376 | Other Negative Correlation |
| EID2 | -0.14463 | 0.014365 | Other Negative Correlation |
| TNFSF9 | -0.14463 | 0.014363 | Other Negative Correlation |
| UBTD2 | -0.1447 | 0.014311 | Other Negative Correlation |
| ARL8B | -0.14479 | 0.014251 | Other Negative Correlation |
| PTPN18 | -0.14492 | 0.014168 | Other Negative Correlation |
| RNGTT | -0.14495 | 0.014146 | Other Negative Correlation |
| GOLGA3 | -0.145 | 0.014114 | Other Negative Correlation |
| NARF | -0.14505 | 0.014079 | Other Negative Correlation |
| TLNRD1 | -0.14506 | 0.014074 | Other Negative Correlation |
| THRB | -0.14506 | 0.014073 | Other Negative Correlation |
| RBM42 | -0.14507 | 0.014067 | Other Negative Correlation |
| GSTCD | -0.14507 | 0.014065 | Other Negative Correlation |
| BTF3L4 | -0.1451 | 0.014043 | Other Negative Correlation |
| RNF39 | -0.14511 | 0.014036 | Other Negative Correlation |
| DRAM2 | -0.14513 | 0.014023 | Other Negative Correlation |

| | | | |
|---|---|---|---|
| TRAPPC2B | -0.14514 | 0.014018 | Other Negative Correlation |
| BMT2 | -0.14514 | 0.014016 | Other Negative Correlation |
| SMAD7 | -0.14521 | 0.013969 | Other Negative Correlation |
| UNC13D | -0.14529 | 0.013915 | Other Negative Correlation |
| GRB7 | -0.14532 | 0.0139 | Other Negative Correlation |
| MBOAT7 | -0.14536 | 0.013872 | Other Negative Correlation |
| PRR14 | -0.1454 | 0.013843 | Other Negative Correlation |
| ELP4 | -0.14549 | 0.013786 | Other Negative Correlation |
| TMEM186 | -0.14558 | 0.013726 | Other Negative Correlation |
| NSMF | -0.14565 | 0.013681 | Other Negative Correlation |
| MOSPD2 | -0.14568 | 0.013659 | Other Negative Correlation |
| TOPORS | -0.1457 | 0.013648 | Other Negative Correlation |
| TRIM24 | -0.14579 | 0.013591 | Other Negative Correlation |
| PLEKHH3 | -0.14583 | 0.013564 | Other Negative Correlation |
| ADAP1 | -0.14594 | 0.013493 | Other Negative Correlation |
| GLI3 | -0.14597 | 0.013473 | Other Negative Correlation |
| MED20 | -0.14597 | 0.01347 | Other Negative Correlation |
| PWWP3B | -0.14604 | 0.013431 | Other Negative Correlation |
| GNG11 | -0.14615 | 0.013358 | Other Negative Correlation |
| UBE4B | -0.14615 | 0.013355 | Other Negative Correlation |
| MYL9 | -0.14631 | 0.013254 | Other Negative Correlation |
| MT-ND4 | -0.14635 | 0.013229 | Other Negative Correlation |
| KAT7 | -0.14636 | 0.013225 | Other Negative Correlation |
| CES1 | -0.14639 | 0.013204 | Other Negative Correlation |
| ANKRD13C | -0.1464 | 0.0132 | Other Negative Correlation |
| PPP1R8 | -0.14643 | 0.013178 | Other Negative Correlation |
| LNX1 | -0.14646 | 0.013159 | Other Negative Correlation |
| BRMS1L | -0.14647 | 0.013152 | Other Negative Correlation |
| CBFA2T3 | -0.14653 | 0.013116 | Other Negative Correlation |
| TBP | -0.14657 | 0.013088 | Other Negative Correlation |
| TEFM | -0.14659 | 0.013081 | Other Negative Correlation |
| ABHD11 | -0.14659 | 0.013077 | Other Negative Correlation |
| ZNF225 | -0.1466 | 0.013071 | Other Negative Correlation |
| TNFSF11 | -0.14667 | 0.013026 | Other Negative Correlation |
| LDAH | -0.14683 | 0.012931 | Other Negative Correlation |
| DCUN1D3 | -0.1469 | 0.012883 | Other Negative Correlation |
| GTPBP8 | -0.147 | 0.012826 | Other Negative Correlation |
| TSPAN17 | -0.14702 | 0.012809 | Other Negative Correlation |
| TRIM27 | -0.14705 | 0.012793 | Other Negative Correlation |
| UBAP1 | -0.14715 | 0.012728 | Other Negative Correlation |
| UBQLN1 | -0.14718 | 0.012711 | Other Negative Correlation |
| ALKBH6 | -0.14721 | 0.012697 | Other Negative Correlation |
| SGCE | -0.14721 | 0.012694 | Other Negative Correlation |
| TPCN2 | -0.14725 | 0.01267 | Other Negative Correlation |
| MAB21L3 | -0.14729 | 0.012647 | Other Negative Correlation |
| ADPRS | -0.14738 | 0.012588 | Other Negative Correlation |

| | | | |
|---|---|---|---|
| ENSG00000260643 | -0.14748 | 0.012529 | Other Negative Correlation |
| MARCHF2 | -0.1475 | 0.012518 | Other Negative Correlation |
| KDM5D | -0.1475 | 0.012515 | Other Negative Correlation |
| IMMP1L | -0.14754 | 0.012491 | Other Negative Correlation |
| C14orf28 | -0.14769 | 0.012404 | Other Negative Correlation |
| GOLGA2 | -0.14769 | 0.012403 | Other Negative Correlation |
| FKBPL | -0.14769 | 0.012401 | Other Negative Correlation |
| PARN | -0.1477 | 0.012397 | Other Negative Correlation |
| PELO | -0.14772 | 0.012384 | Other Negative Correlation |
| CATSPER1 | -0.14776 | 0.01236 | Other Negative Correlation |
| ENOX2 | -0.14778 | 0.012353 | Other Negative Correlation |
| IL1R2 | -0.1478 | 0.012335 | Other Negative Correlation |
| PHETA1 | -0.14782 | 0.012325 | Other Negative Correlation |
| FAM193B | -0.14783 | 0.012322 | Other Negative Correlation |
| TMEM260 | -0.14784 | 0.012314 | Other Negative Correlation |
| MYLIP | -0.1479 | 0.012279 | Other Negative Correlation |
| PAK1 | -0.14797 | 0.012234 | Other Negative Correlation |
| FAM168A | -0.14798 | 0.012233 | Other Negative Correlation |
| APC | -0.14801 | 0.012212 | Other Negative Correlation |
| TCHH | -0.14804 | 0.012197 | Other Negative Correlation |
| TAOK2 | -0.14809 | 0.012164 | Other Negative Correlation |
| SNRNP27 | -0.14812 | 0.012151 | Other Negative Correlation |
| SPTY2D1 | -0.14821 | 0.012097 | Other Negative Correlation |
| SOCS1 | -0.14821 | 0.012094 | Other Negative Correlation |
| ZBTB33 | -0.14822 | 0.01209 | Other Negative Correlation |
| FGFR2 | -0.14827 | 0.012061 | Other Negative Correlation |
| TIMM17B | -0.14832 | 0.012031 | Other Negative Correlation |
| ANGEL2 | -0.14834 | 0.012021 | Other Negative Correlation |
| GATA3 | -0.14836 | 0.012007 | Other Negative Correlation |
| NFE2L3 | -0.14845 | 0.011955 | Other Negative Correlation |
| KTI12 | -0.14849 | 0.011932 | Other Negative Correlation |
| NAA40 | -0.14849 | 0.01193 | Other Negative Correlation |
| CRISP3 | -0.14851 | 0.011919 | Other Negative Correlation |
| ABCF3 | -0.14856 | 0.011894 | Other Negative Correlation |
| RGL2 | -0.14864 | 0.011848 | Other Negative Correlation |
| ENSG00000284691 | -0.14865 | 0.011838 | Other Negative Correlation |
| UQCC1 | -0.14878 | 0.011765 | Other Negative Correlation |
| ACP3 | -0.14883 | 0.011737 | Other Negative Correlation |
| ACO2 | -0.14883 | 0.011736 | Other Negative Correlation |
| KANSL3 | -0.14886 | 0.01172 | Other Negative Correlation |
| CFAP53 | -0.14891 | 0.011693 | Other Negative Correlation |
| BLOC1S1 | -0.14891 | 0.011693 | Other Negative Correlation |
| ZFP91 | -0.14892 | 0.011683 | Other Negative Correlation |
| CWC25 | -0.14894 | 0.011676 | Other Negative Correlation |
| CEMIP | -0.14894 | 0.011673 | Other Negative Correlation |
| ACOT8 | -0.14905 | 0.011614 | Other Negative Correlation |

| | | | |
|---|---|---|---|
| LRRC14 | -0.14908 | 0.011597 | Other Negative Correlation |
| TBX19 | -0.14928 | 0.011482 | Other Negative Correlation |
| CNOT8 | -0.14928 | 0.011481 | Other Negative Correlation |
| GHDC | -0.14939 | 0.011418 | Other Negative Correlation |
| CCL28 | -0.14941 | 0.01141 | Other Negative Correlation |
| MAP7 | -0.14942 | 0.011402 | Other Negative Correlation |
| IKBKG | -0.14955 | 0.011332 | Other Negative Correlation |
| TMEM184C | -0.14965 | 0.011278 | Other Negative Correlation |
| ZNF512B | -0.14968 | 0.011257 | Other Negative Correlation |
| ZNF32 | -0.14976 | 0.011215 | Other Negative Correlation |
| PIP5KL1 | -0.14979 | 0.011198 | Other Negative Correlation |
| SEPTIN6 | -0.14986 | 0.01116 | Other Negative Correlation |
| FGD3 | -0.14988 | 0.011149 | Other Negative Correlation |
| MRPS2 | -0.14994 | 0.011118 | Other Negative Correlation |
| ANO10 | -0.14998 | 0.011096 | Other Negative Correlation |
| NUDCD3 | -0.15001 | 0.011081 | Other Negative Correlation |
| COG7 | -0.15002 | 0.011075 | Other Negative Correlation |
| RDH14 | -0.1501 | 0.011029 | Other Negative Correlation |
| LAPTM5 | -0.15015 | 0.011002 | Other Negative Correlation |
| TUBGCP4 | -0.15019 | 0.010985 | Other Negative Correlation |
| TATDN3 | -0.1502 | 0.010978 | Other Negative Correlation |
| DHFR2 | -0.15021 | 0.01097 | Other Negative Correlation |
| PLEKHA4 | -0.15025 | 0.010953 | Other Negative Correlation |
| MFN1 | -0.15029 | 0.010928 | Other Negative Correlation |
| SKAP2 | -0.15037 | 0.010885 | Other Negative Correlation |
| CNTROB | -0.15053 | 0.0108 | Other Negative Correlation |
| HAL | -0.15059 | 0.010768 | Other Negative Correlation |
| KMT2C | -0.1506 | 0.010765 | Other Negative Correlation |
| PPP6R3 | -0.15062 | 0.010751 | Other Negative Correlation |
| YPEL5 | -0.15067 | 0.010726 | Other Negative Correlation |
| AP3M1 | -0.15068 | 0.010719 | Other Negative Correlation |
| LRG1 | -0.1507 | 0.010712 | Other Negative Correlation |
| SOCS4 | -0.15082 | 0.010648 | Other Negative Correlation |
| NUP50 | -0.15083 | 0.010644 | Other Negative Correlation |
| SNX18 | -0.15088 | 0.010618 | Other Negative Correlation |
| DOP1B | -0.15096 | 0.010573 | Other Negative Correlation |
| GSK3A | -0.15097 | 0.010571 | Other Negative Correlation |
| GDF5 | -0.15101 | 0.010549 | Other Negative Correlation |
| KEAP1 | -0.15108 | 0.010514 | Other Negative Correlation |
| ARL10 | -0.15109 | 0.010505 | Other Negative Correlation |
| PLEKHO2 | -0.15113 | 0.010486 | Other Negative Correlation |
| PADI3 | -0.15115 | 0.010477 | Other Negative Correlation |
| TLK2 | -0.15117 | 0.010465 | Other Negative Correlation |
| KMO | -0.15121 | 0.010445 | Other Negative Correlation |
| STS | -0.15125 | 0.010427 | Other Negative Correlation |
| NOTCH2NLA | -0.15126 | 0.01042 | Other Negative Correlation |

| | | | |
|---|---|---|---|
| FGD4 | -0.15127 | 0.010413 | Other Negative Correlation |
| TSC1 | -0.15132 | 0.01039 | Other Negative Correlation |
| VCPIP1 | -0.15135 | 0.010371 | Other Negative Correlation |
| DAW1 | -0.15136 | 0.010366 | Other Negative Correlation |
| JTB | -0.15137 | 0.010361 | Other Negative Correlation |
| POMT2 | -0.15143 | 0.010331 | Other Negative Correlation |
| PRKACA | -0.15144 | 0.01033 | Other Negative Correlation |
| HM13 | -0.15152 | 0.010288 | Other Negative Correlation |
| TMEM182 | -0.15155 | 0.010269 | Other Negative Correlation |
| PHYHIP | -0.15158 | 0.010257 | Other Negative Correlation |
| VPS41 | -0.1516 | 0.010247 | Other Negative Correlation |
| FRAT2 | -0.15161 | 0.010243 | Other Negative Correlation |
| TMEM61 | -0.15164 | 0.010228 | Other Negative Correlation |
| CCDC71 | -0.1517 | 0.010195 | Other Negative Correlation |
| SIRT2 | -0.15175 | 0.010169 | Other Negative Correlation |
| ROBO1 | -0.15179 | 0.010151 | Other Negative Correlation |
| ZSWIM8 | -0.15181 | 0.010139 | Other Negative Correlation |
| ZNF398 | -0.15182 | 0.010137 | Other Negative Correlation |
| SNAP47 | -0.15195 | 0.010072 | Other Negative Correlation |
| RUNDC3A | -0.15195 | 0.010069 | Other Negative Correlation |
| KCTD14 | -0.15197 | 0.010059 | Other Negative Correlation |
| NACAD | -0.15214 | 0.009974 | Other Negative Correlation |
| FBXL5 | -0.15216 | 0.009968 | Other Negative Correlation |
| ENSG00000170846 | -0.15221 | 0.009942 | Other Negative Correlation |
| HIPK1 | -0.15229 | 0.0099 | Other Negative Correlation |
| BAALC | -0.15231 | 0.009894 | Other Negative Correlation |
| POLM | -0.15234 | 0.009879 | Other Negative Correlation |
| NADK2 | -0.15234 | 0.009878 | Other Negative Correlation |
| ANKRD37 | -0.15241 | 0.009845 | Other Negative Correlation |
| SAMHD1 | -0.15242 | 0.00984 | Other Negative Correlation |
| LRCH4 | -0.15246 | 0.00982 | Other Negative Correlation |
| PFDN1 | -0.15248 | 0.009811 | Other Negative Correlation |
| CFAP69 | -0.15248 | 0.00981 | Other Negative Correlation |
| CYB5R2 | -0.15254 | 0.009777 | Other Negative Correlation |
| BTBD1 | -0.15255 | 0.009774 | Other Negative Correlation |
| POLR3E | -0.1526 | 0.009751 | Other Negative Correlation |
| LIPK | -0.15263 | 0.009737 | Other Negative Correlation |
| ZNF319 | -0.15267 | 0.009717 | Other Negative Correlation |
| CEP97 | -0.15273 | 0.009686 | Other Negative Correlation |
| IPO8 | -0.15274 | 0.009683 | Other Negative Correlation |
| APOBEC3G | -0.15276 | 0.009675 | Other Negative Correlation |
| PAPSS2 | -0.15278 | 0.009666 | Other Negative Correlation |
| GOLGA7B | -0.15282 | 0.009644 | Other Negative Correlation |
| TMEM187 | -0.15285 | 0.009632 | Other Negative Correlation |
| NCOR2 | -0.15286 | 0.009626 | Other Negative Correlation |
| STIM2 | -0.15287 | 0.009621 | Other Negative Correlation |

| | | | |
|---|---|---|---|
| LARS2 | -0.1529 | 0.009604 | Other Negative Correlation |
| MORN1 | -0.15296 | 0.00958 | Other Negative Correlation |
| LENG8 | -0.153 | 0.00956 | Other Negative Correlation |
| PHF8 | -0.153 | 0.009559 | Other Negative Correlation |
| CAPN8 | -0.15303 | 0.009546 | Other Negative Correlation |
| FOXO1 | -0.15303 | 0.009545 | Other Negative Correlation |
| STAU2 | -0.15305 | 0.009533 | Other Negative Correlation |
| ZNF268 | -0.15306 | 0.009531 | Other Negative Correlation |
| PANK2 | -0.15313 | 0.009498 | Other Negative Correlation |
| PCDHA10 | -0.15314 | 0.009491 | Other Negative Correlation |
| USP43 | -0.15317 | 0.00948 | Other Negative Correlation |
| MBTPS2 | -0.15317 | 0.009478 | Other Negative Correlation |
| RAP2A | -0.15318 | 0.009474 | Other Negative Correlation |
| RAB11FIP3 | -0.1532 | 0.009465 | Other Negative Correlation |
| TMEM101 | -0.15322 | 0.009452 | Other Negative Correlation |
| NFATC2IP | -0.15325 | 0.009441 | Other Negative Correlation |
| ACADSB | -0.15332 | 0.009406 | Other Negative Correlation |
| TAB1 | -0.15334 | 0.009398 | Other Negative Correlation |
| NEK11 | -0.15334 | 0.009396 | Other Negative Correlation |
| MAP10 | -0.15341 | 0.009364 | Other Negative Correlation |
| SMPDL3B | -0.15347 | 0.009337 | Other Negative Correlation |
| GTF2IRD2B | -0.15348 | 0.009333 | Other Negative Correlation |
| HACL1 | -0.15355 | 0.009298 | Other Negative Correlation |
| NSD1 | -0.15357 | 0.009292 | Other Negative Correlation |
| CRTC1 | -0.15358 | 0.009286 | Other Negative Correlation |
| ASH2L | -0.15358 | 0.009285 | Other Negative Correlation |
| CLUH | -0.1536 | 0.009275 | Other Negative Correlation |
| PNKP | -0.15368 | 0.00924 | Other Negative Correlation |
| IFT81 | -0.15373 | 0.009215 | Other Negative Correlation |
| TASOR2 | -0.15374 | 0.009212 | Other Negative Correlation |
| DAPL1 | -0.15383 | 0.009173 | Other Negative Correlation |
| IFT20 | -0.15383 | 0.009172 | Other Negative Correlation |
| ZNF224 | -0.15385 | 0.009163 | Other Negative Correlation |
| SUN3 | -0.1539 | 0.009138 | Other Negative Correlation |
| ZNF71 | -0.154 | 0.009095 | Other Negative Correlation |
| CEACAM19 | -0.15403 | 0.009079 | Other Negative Correlation |
| SLC35A4 | -0.15405 | 0.009068 | Other Negative Correlation |
| TNNT3 | -0.15408 | 0.009056 | Other Negative Correlation |
| DPH3 | -0.15412 | 0.009037 | Other Negative Correlation |
| SPTSSB | -0.15414 | 0.009029 | Other Negative Correlation |
| FUT3 | -0.15415 | 0.009026 | Other Negative Correlation |
| NDUFV1 | -0.1542 | 0.009002 | Other Negative Correlation |
| WDR31 | -0.1542 | 0.009 | Other Negative Correlation |
| TTC4 | -0.15421 | 0.008996 | Other Negative Correlation |
| SPRTN | -0.1543 | 0.008957 | Other Negative Correlation |
| ZBTB47 | -0.15432 | 0.008949 | Other Negative Correlation |

| | | | |
|---|---|---|---|
| URGCP | -0.15433 | 0.008942 | Other Negative Correlation |
| WDR20 | -0.15439 | 0.008915 | Other Negative Correlation |
| GPRASP2 | -0.15443 | 0.0089 | Other Negative Correlation |
| TRAPPC2 | -0.15443 | 0.008898 | Other Negative Correlation |
| MAP4K2 | -0.15447 | 0.00888 | Other Negative Correlation |
| LRCH1 | -0.15447 | 0.008879 | Other Negative Correlation |
| DYRK2 | -0.15451 | 0.008863 | Other Negative Correlation |
| PIGN | -0.15452 | 0.008859 | Other Negative Correlation |
| ZNF845 | -0.15455 | 0.008844 | Other Negative Correlation |
| NEK4 | -0.15467 | 0.008791 | Other Negative Correlation |
| ATMIN | -0.15471 | 0.008773 | Other Negative Correlation |
| SLC25A36 | -0.15478 | 0.008745 | Other Negative Correlation |
| ABHD13 | -0.15484 | 0.008719 | Other Negative Correlation |
| ANPEP | -0.15488 | 0.008698 | Other Negative Correlation |
| TMUB1 | -0.15489 | 0.008697 | Other Negative Correlation |
| PLSCR4 | -0.15495 | 0.008669 | Other Negative Correlation |
| TUBG2 | -0.15501 | 0.008642 | Other Negative Correlation |
| RSPH1 | -0.15506 | 0.008621 | Other Negative Correlation |
| SUZ12 | -0.15506 | 0.008619 | Other Negative Correlation |
| TUT1 | -0.15507 | 0.008617 | Other Negative Correlation |
| SAFB2 | -0.15508 | 0.008614 | Other Negative Correlation |
| ADARB2 | -0.15509 | 0.00861 | Other Negative Correlation |
| TRPS1 | -0.1551 | 0.008604 | Other Negative Correlation |
| HEXA | -0.15514 | 0.008586 | Other Negative Correlation |
| NKPD1 | -0.15514 | 0.008585 | Other Negative Correlation |
| ENKUR | -0.15518 | 0.008568 | Other Negative Correlation |
| TRMT12 | -0.15521 | 0.008556 | Other Negative Correlation |
| GPKOW | -0.15521 | 0.008556 | Other Negative Correlation |
| ZNF574 | -0.15524 | 0.008544 | Other Negative Correlation |
| ZW10 | -0.15529 | 0.008522 | Other Negative Correlation |
| RALGAPB | -0.15534 | 0.008501 | Other Negative Correlation |
| GCN1 | -0.15543 | 0.00846 | Other Negative Correlation |
| SAP30BP | -0.15555 | 0.008411 | Other Negative Correlation |
| TCF20 | -0.15559 | 0.008394 | Other Negative Correlation |
| ASB7 | -0.15559 | 0.008393 | Other Negative Correlation |
| UFM1 | -0.1556 | 0.008391 | Other Negative Correlation |
| ADA | -0.15564 | 0.008374 | Other Negative Correlation |
| C3orf33 | -0.15566 | 0.008364 | Other Negative Correlation |
| F8A1 | -0.15567 | 0.008361 | Other Negative Correlation |
| RBM18 | -0.15578 | 0.008315 | Other Negative Correlation |
| NCOA7 | -0.15578 | 0.008315 | Other Negative Correlation |
| ZNF830 | -0.15586 | 0.008281 | Other Negative Correlation |
| GRIPAP1 | -0.15588 | 0.008273 | Other Negative Correlation |
| HYAL3 | -0.15588 | 0.008271 | Other Negative Correlation |
| TRIM3 | -0.15596 | 0.008238 | Other Negative Correlation |
| VWA1 | -0.15599 | 0.008224 | Other Negative Correlation |

| | | | |
|---|---|---|---|
| SMARCD3 | -0.15607 | 0.008192 | Other Negative Correlation |
| DNM2 | -0.15608 | 0.008186 | Other Negative Correlation |
| LRRC75B | -0.15611 | 0.008174 | Other Negative Correlation |
| SNTB2 | -0.15612 | 0.00817 | Other Negative Correlation |
| SUOX | -0.15627 | 0.008109 | Other Negative Correlation |
| C12orf76 | -0.15629 | 0.008101 | Other Negative Correlation |
| ARL4A | -0.15633 | 0.008087 | Other Negative Correlation |
| CCDC63 | -0.15638 | 0.008063 | Other Negative Correlation |
| MMUT | -0.1564 | 0.008056 | Other Negative Correlation |
| STRIP1 | -0.15649 | 0.008019 | Other Negative Correlation |
| G6PC3 | -0.15652 | 0.008006 | Other Negative Correlation |
| SLC50A1 | -0.15654 | 0.007999 | Other Negative Correlation |
| FHIP1A | -0.15655 | 0.007995 | Other Negative Correlation |
| SCAND1 | -0.15656 | 0.007992 | Other Negative Correlation |
| DYM | -0.15662 | 0.007965 | Other Negative Correlation |
| CHRNB1 | -0.15663 | 0.007964 | Other Negative Correlation |
| LRP3 | -0.15666 | 0.00795 | Other Negative Correlation |
| NEDD4L | -0.15677 | 0.007907 | Other Negative Correlation |
| SLC35B2 | -0.15681 | 0.00789 | Other Negative Correlation |
| HDHD2 | -0.15683 | 0.00788 | Other Negative Correlation |
| MPHOSPH10 | -0.15687 | 0.007865 | Other Negative Correlation |
| NEB | -0.15687 | 0.007864 | Other Negative Correlation |
| ERCC4 | -0.1569 | 0.007852 | Other Negative Correlation |
| SLC37A1 | -0.1569 | 0.007852 | Other Negative Correlation |
| RIMKLB | -0.15692 | 0.007845 | Other Negative Correlation |
| AMBRA1 | -0.15694 | 0.00784 | Other Negative Correlation |
| FMN1 | -0.15696 | 0.007829 | Other Negative Correlation |
| RNF14 | -0.15698 | 0.007821 | Other Negative Correlation |
| CD164L2 | -0.15701 | 0.007809 | Other Negative Correlation |
| NEK1 | -0.15705 | 0.007796 | Other Negative Correlation |
| ZC3HC1 | -0.15715 | 0.007756 | Other Negative Correlation |
| TUBGCP5 | -0.1572 | 0.007734 | Other Negative Correlation |
| TMF1 | -0.15733 | 0.007686 | Other Negative Correlation |
| ULK2 | -0.15735 | 0.007677 | Other Negative Correlation |
| PRRG2 | -0.15735 | 0.007676 | Other Negative Correlation |
| IFTAP | -0.15743 | 0.007644 | Other Negative Correlation |
| GNA11 | -0.15744 | 0.00764 | Other Negative Correlation |
| RLF | -0.15745 | 0.007635 | Other Negative Correlation |
| KRT86 | -0.15752 | 0.007611 | Other Negative Correlation |
| RNF121 | -0.15764 | 0.007562 | Other Negative Correlation |
| SP2 | -0.15766 | 0.007555 | Other Negative Correlation |
| CTDSP2 | -0.15767 | 0.00755 | Other Negative Correlation |
| TLR2 | -0.1577 | 0.007538 | Other Negative Correlation |
| MIER1 | -0.15783 | 0.007489 | Other Negative Correlation |
| GOLPH3 | -0.15787 | 0.007475 | Other Negative Correlation |
| FCHO1 | -0.15788 | 0.007469 | Other Negative Correlation |

| | | | |
|---|---|---|---|
| USP25 | -0.15792 | 0.007456 | Other Negative Correlation |
| ARSA | -0.15794 | 0.007447 | Other Negative Correlation |
| SBF1 | -0.15794 | 0.007447 | Other Negative Correlation |
| PPP2R2B | -0.15795 | 0.007444 | Other Negative Correlation |
| HCAR3 | -0.15805 | 0.007406 | Other Negative Correlation |
| PRCD | -0.15807 | 0.007397 | Other Negative Correlation |
| MYO9A | -0.15823 | 0.007336 | Other Negative Correlation |
| EXOC4 | -0.15834 | 0.007297 | Other Negative Correlation |
| CDK7 | -0.15835 | 0.007292 | Other Negative Correlation |
| FZD3 | -0.15839 | 0.007277 | Other Negative Correlation |
| ESCO1 | -0.15847 | 0.007248 | Other Negative Correlation |
| NCBP2AS2 | -0.15853 | 0.007227 | Other Negative Correlation |
| DOCK7 | -0.15854 | 0.007222 | Other Negative Correlation |
| MTPAP | -0.1588 | 0.007126 | Other Negative Correlation |
| WLS | -0.15886 | 0.007103 | Other Negative Correlation |
| PHF13 | -0.15889 | 0.007095 | Other Negative Correlation |
| UBE2J2 | -0.15891 | 0.007085 | Other Negative Correlation |
| TFIP11 | -0.159 | 0.007053 | Other Negative Correlation |
| CYB5R4 | -0.15901 | 0.007051 | Other Negative Correlation |
| ZBTB43 | -0.15903 | 0.007044 | Other Negative Correlation |
| GBE1 | -0.15906 | 0.007033 | Other Negative Correlation |
| FFAR2 | -0.15907 | 0.007029 | Other Negative Correlation |
| LRIF1 | -0.15912 | 0.007011 | Other Negative Correlation |
| DHDH | -0.15915 | 0.007 | Other Negative Correlation |
| ADAM28 | -0.1592 | 0.006981 | Other Negative Correlation |
| ZYG11B | -0.15925 | 0.006962 | Other Negative Correlation |
| C18orf32 | -0.15936 | 0.006925 | Other Negative Correlation |
| EDEM3 | -0.15936 | 0.006924 | Other Negative Correlation |
| LSM8 | -0.15951 | 0.006869 | Other Negative Correlation |
| RSRC1 | -0.15952 | 0.006866 | Other Negative Correlation |
| ALS2 | -0.15953 | 0.006863 | Other Negative Correlation |
| LAMA4 | -0.15957 | 0.00685 | Other Negative Correlation |
| DZIP3 | -0.1596 | 0.00684 | Other Negative Correlation |
| UBAC2 | -0.15961 | 0.006835 | Other Negative Correlation |
| CEP104 | -0.15963 | 0.006829 | Other Negative Correlation |
| TMEM70 | -0.15963 | 0.006827 | Other Negative Correlation |
| AFF1 | -0.15963 | 0.006827 | Other Negative Correlation |
| SAP130 | -0.15966 | 0.006818 | Other Negative Correlation |
| UBXN2B | -0.15967 | 0.006812 | Other Negative Correlation |
| NDUFB9 | -0.15974 | 0.006789 | Other Negative Correlation |
| SOGA1 | -0.15975 | 0.006784 | Other Negative Correlation |
| LENG1 | -0.15976 | 0.006783 | Other Negative Correlation |
| KLHL2 | -0.15978 | 0.006777 | Other Negative Correlation |
| CCDC125 | -0.15985 | 0.00675 | Other Negative Correlation |
| BTN3A1 | -0.1599 | 0.006733 | Other Negative Correlation |
| TRAPPC10 | -0.15997 | 0.006707 | Other Negative Correlation |

| | | | |
|---|---|---|---|
| NUFIP1 | -0.15999 | 0.0067 | Other Negative Correlation |
| KLK7 | -0.16004 | 0.006684 | Other Negative Correlation |
| ZNF593 | -0.16014 | 0.006651 | Other Negative Correlation |
| DIS3L2 | -0.16021 | 0.006627 | Other Negative Correlation |
| GUCD1 | -0.16021 | 0.006625 | Other Negative Correlation |
| LYRM9 | -0.16022 | 0.006621 | Other Negative Correlation |
| SLC38A6 | -0.16024 | 0.006614 | Other Negative Correlation |
| MCTP1 | -0.16027 | 0.006607 | Other Negative Correlation |
| FBXO45 | -0.16027 | 0.006605 | Other Negative Correlation |
| CYTH4 | -0.16028 | 0.006602 | Other Negative Correlation |
| GPR137B | -0.16032 | 0.006588 | Other Negative Correlation |
| SQSTM1 | -0.16036 | 0.006574 | Other Negative Correlation |
| SLC38A4 | -0.1604 | 0.00656 | Other Negative Correlation |
| IL10RB | -0.16045 | 0.006544 | Other Negative Correlation |
| TAS1R3 | -0.16046 | 0.00654 | Other Negative Correlation |
| NBPF12 | -0.16052 | 0.006519 | Other Negative Correlation |
| AGK | -0.16054 | 0.006515 | Other Negative Correlation |
| SRC | -0.16054 | 0.006514 | Other Negative Correlation |
| ATRIP | -0.16054 | 0.006514 | Other Negative Correlation |
| PRR15L | -0.1606 | 0.006494 | Other Negative Correlation |
| USP48 | -0.16063 | 0.006483 | Other Negative Correlation |
| MAP2K4 | -0.16074 | 0.006446 | Other Negative Correlation |
| TRIM6 | -0.16084 | 0.006413 | Other Negative Correlation |
| ZNF524 | -0.16084 | 0.006412 | Other Negative Correlation |
| AFAP1L1 | -0.16088 | 0.0064 | Other Negative Correlation |
| QRICH2 | -0.16098 | 0.006366 | Other Negative Correlation |
| COQ6 | -0.16099 | 0.006362 | Other Negative Correlation |
| HDAC10 | -0.16101 | 0.006356 | Other Negative Correlation |
| HSCB | -0.16102 | 0.006353 | Other Negative Correlation |
| NRDC | -0.16111 | 0.006323 | Other Negative Correlation |
| CCDC57 | -0.16116 | 0.006305 | Other Negative Correlation |
| MYORG | -0.16117 | 0.006302 | Other Negative Correlation |
| MRPL2 | -0.16125 | 0.006276 | Other Negative Correlation |
| ZNF18 | -0.16131 | 0.006256 | Other Negative Correlation |
| ATRN | -0.16131 | 0.006256 | Other Negative Correlation |
| TRAF3IP1 | -0.16136 | 0.00624 | Other Negative Correlation |
| MED21 | -0.16144 | 0.006214 | Other Negative Correlation |
| KCTD18 | -0.16146 | 0.006209 | Other Negative Correlation |
| KBTBD6 | -0.16149 | 0.0062 | Other Negative Correlation |
| ELF4 | -0.16151 | 0.006192 | Other Negative Correlation |
| ZBED3 | -0.16162 | 0.006157 | Other Negative Correlation |
| CCDC71L | -0.16165 | 0.006149 | Other Negative Correlation |
| IFNLR1 | -0.16173 | 0.006122 | Other Negative Correlation |
| STEAP2 | -0.16176 | 0.006111 | Other Negative Correlation |
| HPS3 | -0.16184 | 0.006086 | Other Negative Correlation |
| PDE9A | -0.16186 | 0.006079 | Other Negative Correlation |

| | | | |
|---|---|---|---|
| CFAP300 | -0.16188 | 0.006074 | Other Negative Correlation |
| DHX38 | -0.16189 | 0.00607 | Other Negative Correlation |
| AMMECR1L | -0.1619 | 0.006067 | Other Negative Correlation |
| COASY | -0.16192 | 0.006061 | Other Negative Correlation |
| OPA3 | -0.16192 | 0.006061 | Other Negative Correlation |
| WDR48 | -0.16194 | 0.006055 | Other Negative Correlation |
| PSMB4 | -0.162 | 0.006036 | Other Negative Correlation |
| KLHL13 | -0.16201 | 0.006034 | Other Negative Correlation |
| POM121 | -0.16207 | 0.006012 | Other Negative Correlation |
| FAHD2A | -0.16209 | 0.006006 | Other Negative Correlation |
| ABHD16A | -0.16211 | 0.006002 | Other Negative Correlation |
| PRKAR1B | -0.16225 | 0.005959 | Other Negative Correlation |
| SCRN3 | -0.16226 | 0.005954 | Other Negative Correlation |
| PFKFB2 | -0.16228 | 0.005947 | Other Negative Correlation |
| INVS | -0.16233 | 0.005933 | Other Negative Correlation |
| KIFC3 | -0.16239 | 0.005912 | Other Negative Correlation |
| RPAP3 | -0.16251 | 0.005876 | Other Negative Correlation |
| CNOT6 | -0.16255 | 0.005864 | Other Negative Correlation |
| PRR14L | -0.16255 | 0.005864 | Other Negative Correlation |
| AAMP | -0.16259 | 0.005853 | Other Negative Correlation |
| SLC2A10 | -0.16288 | 0.005762 | Other Negative Correlation |
| TRIM32 | -0.1629 | 0.005758 | Other Negative Correlation |
| CIDEA | -0.16304 | 0.005715 | Other Negative Correlation |
| SLC26A6 | -0.16305 | 0.005711 | Other Negative Correlation |
| LZTFL1 | -0.16305 | 0.005711 | Other Negative Correlation |
| MAU2 | -0.16306 | 0.005708 | Other Negative Correlation |
| SLC25A13 | -0.16307 | 0.005707 | Other Negative Correlation |
| RORC | -0.16332 | 0.005631 | Other Negative Correlation |
| SLC43A2 | -0.16336 | 0.005621 | Other Negative Correlation |
| RTL6 | -0.16339 | 0.00561 | Other Negative Correlation |
| IFT22 | -0.1634 | 0.005609 | Other Negative Correlation |
| SOCS6 | -0.1634 | 0.005607 | Other Negative Correlation |
| CDS2 | -0.1634 | 0.005607 | Other Negative Correlation |
| ASB2 | -0.16344 | 0.005595 | Other Negative Correlation |
| FGF5 | -0.16345 | 0.005593 | Other Negative Correlation |
| SIL1 | -0.1635 | 0.00558 | Other Negative Correlation |
| ANO9 | -0.16359 | 0.005552 | Other Negative Correlation |
| TFCP2L1 | -0.16359 | 0.005551 | Other Negative Correlation |
| ZNF786 | -0.1636 | 0.005549 | Other Negative Correlation |
| PDCD6 | -0.16362 | 0.005544 | Other Negative Correlation |
| PLA2G2F | -0.16363 | 0.005539 | Other Negative Correlation |
| SLC52A3 | -0.16366 | 0.005531 | Other Negative Correlation |
| GLOD4 | -0.16368 | 0.005524 | Other Negative Correlation |
| NETO2 | -0.16371 | 0.005516 | Other Negative Correlation |
| MPC2 | -0.16372 | 0.005513 | Other Negative Correlation |
| CLEC16A | -0.16376 | 0.005501 | Other Negative Correlation |

| | | | |
|---|---|---|---|
| IDH3B | -0.16382 | 0.005486 | Other Negative Correlation |
| TXNRD2 | -0.16382 | 0.005484 | Other Negative Correlation |
| BFSP1 | -0.16389 | 0.005464 | Other Negative Correlation |
| ITPR2 | -0.16403 | 0.005424 | Other Negative Correlation |
| CLPTM1 | -0.16404 | 0.005421 | Other Negative Correlation |
| ATG12 | -0.16406 | 0.005416 | Other Negative Correlation |
| ARID4A | -0.16407 | 0.005413 | Other Negative Correlation |
| MKNK1 | -0.1641 | 0.005403 | Other Negative Correlation |
| SLC49A3 | -0.16412 | 0.005399 | Other Negative Correlation |
| SETD9 | -0.16413 | 0.005394 | Other Negative Correlation |
| ASNSD1 | -0.16415 | 0.005389 | Other Negative Correlation |
| PGLYRP3 | -0.16419 | 0.005378 | Other Negative Correlation |
| GPX2 | -0.16419 | 0.005377 | Other Negative Correlation |
| TDRP | -0.16419 | 0.005377 | Other Negative Correlation |
| SEC22C | -0.16421 | 0.005374 | Other Negative Correlation |
| PFKL | -0.16422 | 0.00537 | Other Negative Correlation |
| EMC1 | -0.16424 | 0.005363 | Other Negative Correlation |
| CMC2 | -0.1643 | 0.005346 | Other Negative Correlation |
| TBC1D25 | -0.16432 | 0.00534 | Other Negative Correlation |
| ETV7 | -0.16433 | 0.00534 | Other Negative Correlation |
| SLC2A3 | -0.16441 | 0.005315 | Other Negative Correlation |
| ZNF626 | -0.16445 | 0.005306 | Other Negative Correlation |
| PPP2R5D | -0.16445 | 0.005305 | Other Negative Correlation |
| TRAM1L1 | -0.16449 | 0.005295 | Other Negative Correlation |
| HELQ | -0.16449 | 0.005294 | Other Negative Correlation |
| BAG4 | -0.16457 | 0.005272 | Other Negative Correlation |
| TNRC6B | -0.1646 | 0.005263 | Other Negative Correlation |
| NISCH | -0.16471 | 0.005233 | Other Negative Correlation |
| SNURF | -0.16475 | 0.005221 | Other Negative Correlation |
| EXTL2 | -0.16487 | 0.005188 | Other Negative Correlation |
| NCBP3 | -0.16491 | 0.005176 | Other Negative Correlation |
| ZFY | -0.16495 | 0.005165 | Other Negative Correlation |
| P2RX6 | -0.16495 | 0.005165 | Other Negative Correlation |
| EFNA5 | -0.16498 | 0.005157 | Other Negative Correlation |
| DGCR6 | -0.165 | 0.005153 | Other Negative Correlation |
| SMIM8 | -0.16503 | 0.005144 | Other Negative Correlation |
| NCKAP5L | -0.16511 | 0.005122 | Other Negative Correlation |
| RIPK2 | -0.16521 | 0.005096 | Other Negative Correlation |
| LURAP1L | -0.16522 | 0.005093 | Other Negative Correlation |
| PANX1 | -0.16522 | 0.005092 | Other Negative Correlation |
| TYK2 | -0.16529 | 0.005072 | Other Negative Correlation |
| USP24 | -0.16531 | 0.005069 | Other Negative Correlation |
| SLC35D1 | -0.16535 | 0.005057 | Other Negative Correlation |
| ACCS | -0.16536 | 0.005055 | Other Negative Correlation |
| MAP4K3 | -0.16539 | 0.005047 | Other Negative Correlation |
| STAT4 | -0.16546 | 0.005027 | Other Negative Correlation |

| | | | |
|---|---|---|---|
| SHOC2 | -0.16555 | 0.005004 | Other Negative Correlation |
| WDR70 | -0.16561 | 0.004988 | Other Negative Correlation |
| TFAP2C | -0.16562 | 0.004985 | Other Negative Correlation |
| TRAPPC3L | -0.16562 | 0.004984 | Other Negative Correlation |
| VPS35L | -0.16563 | 0.004983 | Other Negative Correlation |
| RIOX1 | -0.16563 | 0.004982 | Other Negative Correlation |
| RNF111 | -0.16563 | 0.004982 | Other Negative Correlation |
| SRR | -0.16568 | 0.004968 | Other Negative Correlation |
| SNRNP200 | -0.16571 | 0.00496 | Other Negative Correlation |
| PCNX3 | -0.16573 | 0.004955 | Other Negative Correlation |
| WDR81 | -0.16578 | 0.004941 | Other Negative Correlation |
| ESYT1 | -0.16579 | 0.004941 | Other Negative Correlation |
| MAP3K8 | -0.16586 | 0.00492 | Other Negative Correlation |
| CSTF2T | -0.1659 | 0.00491 | Other Negative Correlation |
| SDHA | -0.16591 | 0.004907 | Other Negative Correlation |
| PLEKHF2 | -0.16606 | 0.004869 | Other Negative Correlation |
| MEGF6 | -0.16618 | 0.004838 | Other Negative Correlation |
| GTF2H4 | -0.16626 | 0.004816 | Other Negative Correlation |
| DEFB4A | -0.16631 | 0.004803 | Other Negative Correlation |
| ZBTB22 | -0.16635 | 0.004792 | Other Negative Correlation |
| IPO9 | -0.16637 | 0.004788 | Other Negative Correlation |
| CEP164 | -0.16641 | 0.004779 | Other Negative Correlation |
| SSR4 | -0.16645 | 0.004768 | Other Negative Correlation |
| CARMIL1 | -0.16646 | 0.004765 | Other Negative Correlation |
| ADCY7 | -0.1665 | 0.004754 | Other Negative Correlation |
| PRXL2B | -0.16654 | 0.004744 | Other Negative Correlation |
| PLA2G4C | -0.16656 | 0.00474 | Other Negative Correlation |
| NAA60 | -0.16657 | 0.004737 | Other Negative Correlation |
| CCDC112 | -0.16659 | 0.004731 | Other Negative Correlation |
| BTBD8 | -0.16668 | 0.004708 | Other Negative Correlation |
| PTPRM | -0.16671 | 0.004701 | Other Negative Correlation |
| HOOK2 | -0.16675 | 0.004691 | Other Negative Correlation |
| COIL | -0.16675 | 0.004691 | Other Negative Correlation |
| PLS1 | -0.16676 | 0.004689 | Other Negative Correlation |
| UBR5 | -0.16679 | 0.004682 | Other Negative Correlation |
| C2orf76 | -0.16682 | 0.004675 | Other Negative Correlation |
| ASB9 | -0.16685 | 0.004667 | Other Negative Correlation |
| HARS2 | -0.16687 | 0.004663 | Other Negative Correlation |
| BST2 | -0.16687 | 0.004662 | Other Negative Correlation |
| PLXNA3 | -0.16687 | 0.004661 | Other Negative Correlation |
| SGPP1 | -0.16688 | 0.00466 | Other Negative Correlation |
| EPHX2 | -0.16694 | 0.004643 | Other Negative Correlation |
| ARMC8 | -0.16698 | 0.004635 | Other Negative Correlation |
| FHIP2A | -0.16699 | 0.004631 | Other Negative Correlation |
| TCEA2 | -0.16707 | 0.004612 | Other Negative Correlation |
| CYP24A1 | -0.16716 | 0.004589 | Other Negative Correlation |

| | | | |
|---|---|---|---|
| NEK3 | -0.16719 | 0.004581 | Other Negative Correlation |
| RRAGC | -0.16721 | 0.004578 | Other Negative Correlation |
| TMTC2 | -0.16731 | 0.004552 | Other Negative Correlation |
| MRC2 | -0.16731 | 0.004552 | Other Negative Correlation |
| HERC1 | -0.16732 | 0.00455 | Other Negative Correlation |
| FIZ1 | -0.16733 | 0.004548 | Other Negative Correlation |
| REPIN1 | -0.16737 | 0.004539 | Other Negative Correlation |
| IL6ST | -0.16752 | 0.004501 | Other Negative Correlation |
| GATC | -0.16755 | 0.004494 | Other Negative Correlation |
| AGBL5 | -0.16757 | 0.00449 | Other Negative Correlation |
| ELF2 | -0.16769 | 0.00446 | Other Negative Correlation |
| CCBE1 | -0.16778 | 0.004438 | Other Negative Correlation |
| PRKRA | -0.1678 | 0.004433 | Other Negative Correlation |
| ENSG00000282988 | -0.16781 | 0.004431 | Other Negative Correlation |
| ICAM3 | -0.16781 | 0.00443 | Other Negative Correlation |
| THBD | -0.16782 | 0.004429 | Other Negative Correlation |
| FHDC1 | -0.16785 | 0.004423 | Other Negative Correlation |
| PIGP | -0.16791 | 0.004406 | Other Negative Correlation |
| PHAF1 | -0.16797 | 0.004394 | Other Negative Correlation |
| EHBP1L1 | -0.16798 | 0.004391 | Other Negative Correlation |
| DPEP1 | -0.16801 | 0.004385 | Other Negative Correlation |
| CIPC | -0.16809 | 0.004364 | Other Negative Correlation |
| GPATCH8 | -0.16815 | 0.004351 | Other Negative Correlation |
| UROD | -0.1682 | 0.004338 | Other Negative Correlation |
| RARS2 | -0.16823 | 0.004332 | Other Negative Correlation |
| DOK4 | -0.16838 | 0.004297 | Other Negative Correlation |
| FAM229B | -0.16838 | 0.004296 | Other Negative Correlation |
| CYB561D1 | -0.16845 | 0.004281 | Other Negative Correlation |
| TAF12 | -0.16849 | 0.00427 | Other Negative Correlation |
| MTERF4 | -0.16852 | 0.004264 | Other Negative Correlation |
| WRNIP1 | -0.16855 | 0.004258 | Other Negative Correlation |
| DHX30 | -0.16855 | 0.004257 | Other Negative Correlation |
| CEBPA | -0.16857 | 0.004254 | Other Negative Correlation |
| SYT12 | -0.16872 | 0.004217 | Other Negative Correlation |
| IFIT2 | -0.16876 | 0.00421 | Other Negative Correlation |
| ACO1 | -0.16882 | 0.004194 | Other Negative Correlation |
| INTS15 | -0.16883 | 0.004194 | Other Negative Correlation |
| CHD2 | -0.16886 | 0.004187 | Other Negative Correlation |
| PSMC4 | -0.16888 | 0.004182 | Other Negative Correlation |
| STYXL1 | -0.16889 | 0.004179 | Other Negative Correlation |
| PEX7 | -0.16893 | 0.00417 | Other Negative Correlation |
| DNMT3A | -0.16896 | 0.004164 | Other Negative Correlation |
| RUNDC1 | -0.16897 | 0.004161 | Other Negative Correlation |
| GARRE1 | -0.16898 | 0.004158 | Other Negative Correlation |
| ALG3 | -0.16899 | 0.004156 | Other Negative Correlation |
| UVRAG | -0.16899 | 0.004156 | Other Negative Correlation |

| | | | |
|---|---|---|---|
| TRMT2A | -0.16901 | 0.004152 | Other Negative Correlation |
| FMNL1 | -0.16902 | 0.004149 | Other Negative Correlation |
| SOX13 | -0.16907 | 0.004138 | Other Negative Correlation |
| SLC6A8 | -0.16924 | 0.004102 | Other Negative Correlation |
| AHI1 | -0.16935 | 0.004076 | Other Negative Correlation |
| MOB1B | -0.16935 | 0.004076 | Other Negative Correlation |
| SPTY2D1OS | -0.16937 | 0.004072 | Other Negative Correlation |
| TTC39C | -0.16939 | 0.004067 | Other Negative Correlation |
| CD276 | -0.16949 | 0.004046 | Other Negative Correlation |
| NSF | -0.16951 | 0.00404 | Other Negative Correlation |
| TP53INP1 | -0.16953 | 0.004036 | Other Negative Correlation |
| FBXO32 | -0.16953 | 0.004035 | Other Negative Correlation |
| C15orf39 | -0.16957 | 0.004027 | Other Negative Correlation |
| ZDHHC16 | -0.16958 | 0.004026 | Other Negative Correlation |
| MTF2 | -0.16958 | 0.004025 | Other Negative Correlation |
| LRRC49 | -0.16962 | 0.004016 | Other Negative Correlation |
| B2M | -0.16976 | 0.003986 | Other Negative Correlation |
| EPB41L4A | -0.16976 | 0.003985 | Other Negative Correlation |
| RHOBTB1 | -0.16979 | 0.003981 | Other Negative Correlation |
| SLC35F3 | -0.16988 | 0.003959 | Other Negative Correlation |
| ZFP57 | -0.16992 | 0.003951 | Other Negative Correlation |
| RTL8B | -0.16995 | 0.003946 | Other Negative Correlation |
| STK11 | -0.17002 | 0.003929 | Other Negative Correlation |
| TRIM65 | -0.17004 | 0.003925 | Other Negative Correlation |
| SUMF1 | -0.17006 | 0.003922 | Other Negative Correlation |
| SLC12A9 | -0.17009 | 0.003915 | Other Negative Correlation |
| GLCE | -0.17011 | 0.00391 | Other Negative Correlation |
| MZF1 | -0.17014 | 0.003905 | Other Negative Correlation |
| EXOC8 | -0.17016 | 0.003901 | Other Negative Correlation |
| CXorf58 | -0.17016 | 0.003901 | Other Negative Correlation |
| ZMIZ1 | -0.17016 | 0.0039 | Other Negative Correlation |
| SFTA2 | -0.17017 | 0.003899 | Other Negative Correlation |
| LNX2 | -0.1702 | 0.003891 | Other Negative Correlation |
| SHB | -0.1703 | 0.00387 | Other Negative Correlation |
| MMP9 | -0.1703 | 0.003869 | Other Negative Correlation |
| CWC27 | -0.17033 | 0.003863 | Other Negative Correlation |
| BCL2L13 | -0.17037 | 0.003855 | Other Negative Correlation |
| MC1R | -0.1704 | 0.003849 | Other Negative Correlation |
| TMEM63B | -0.17042 | 0.003845 | Other Negative Correlation |
| SSBP4 | -0.17043 | 0.003843 | Other Negative Correlation |
| ZNF148 | -0.17046 | 0.003837 | Other Negative Correlation |
| FXR2 | -0.17047 | 0.003835 | Other Negative Correlation |
| MYSM1 | -0.1705 | 0.003828 | Other Negative Correlation |
| WBP4 | -0.17051 | 0.003826 | Other Negative Correlation |
| GCOM1 | -0.17051 | 0.003825 | Other Negative Correlation |
| SASH1 | -0.17056 | 0.003816 | Other Negative Correlation |

| | | | |
|---|---|---|---|
| TIMM23B | -0.17056 | 0.003815 | Other Negative Correlation |
| MEX3A | -0.17056 | 0.003815 | Other Negative Correlation |
| GAST | -0.17059 | 0.003808 | Other Negative Correlation |
| SEMA4G | -0.1706 | 0.003807 | Other Negative Correlation |
| MAP3K1 | -0.17068 | 0.00379 | Other Negative Correlation |
| RNASEL | -0.1707 | 0.003787 | Other Negative Correlation |
| GBA1 | -0.17077 | 0.003772 | Other Negative Correlation |
| TTC14 | -0.17096 | 0.003733 | Other Negative Correlation |
| ARL13B | -0.17105 | 0.003715 | Other Negative Correlation |
| PIP5K1A | -0.17105 | 0.003714 | Other Negative Correlation |
| FCHSD1 | -0.17109 | 0.003707 | Other Negative Correlation |
| MED25 | -0.17109 | 0.003706 | Other Negative Correlation |
| SOS2 | -0.1711 | 0.003704 | Other Negative Correlation |
| MED29 | -0.17113 | 0.003698 | Other Negative Correlation |
| CEP350 | -0.17114 | 0.003695 | Other Negative Correlation |
| KCTD9 | -0.17118 | 0.003688 | Other Negative Correlation |
| CHST12 | -0.17129 | 0.003666 | Other Negative Correlation |
| DNAJB5 | -0.17138 | 0.003647 | Other Negative Correlation |
| LGALS9 | -0.17141 | 0.003642 | Other Negative Correlation |
| RAMAC | -0.17142 | 0.003639 | Other Negative Correlation |
| TRIB2 | -0.17146 | 0.003631 | Other Negative Correlation |
| DNASE1L3 | -0.17148 | 0.003627 | Other Negative Correlation |
| HDAC5 | -0.17152 | 0.00362 | Other Negative Correlation |
| TMPRSS13 | -0.17154 | 0.003616 | Other Negative Correlation |
| PAK6 | -0.17158 | 0.003607 | Other Negative Correlation |
| FBXO9 | -0.17166 | 0.003591 | Other Negative Correlation |
| THOP1 | -0.17174 | 0.003576 | Other Negative Correlation |
| ENSG00000251537 | -0.17177 | 0.00357 | Other Negative Correlation |
| SDHAF2 | -0.1718 | 0.003563 | Other Negative Correlation |
| ZKSCAN3 | -0.17182 | 0.00356 | Other Negative Correlation |
| TRIP4 | -0.17183 | 0.003557 | Other Negative Correlation |
| ABCA5 | -0.17185 | 0.003555 | Other Negative Correlation |
| ATF6B | -0.17186 | 0.003552 | Other Negative Correlation |
| SETD2 | -0.17193 | 0.003538 | Other Negative Correlation |
| MAVS | -0.17202 | 0.003521 | Other Negative Correlation |
| BNIPL | -0.17213 | 0.0035 | Other Negative Correlation |
| TCHP | -0.17223 | 0.003479 | Other Negative Correlation |
| OPHN1 | -0.17224 | 0.003478 | Other Negative Correlation |
| CASP8AP2 | -0.17225 | 0.003477 | Other Negative Correlation |
| TMEM11 | -0.17225 | 0.003477 | Other Negative Correlation |
| CA9 | -0.17231 | 0.003464 | Other Negative Correlation |
| TCF7L2 | -0.17236 | 0.003456 | Other Negative Correlation |
| TCTA | -0.17236 | 0.003455 | Other Negative Correlation |
| ABRAXAS2 | -0.17238 | 0.003452 | Other Negative Correlation |
| TRUB2 | -0.17241 | 0.003445 | Other Negative Correlation |
| RB1CC1 | -0.17242 | 0.003444 | Other Negative Correlation |

| | | | |
|---|---|---|---|
| ZNF236 | -0.17251 | 0.003427 | Other Negative Correlation |
| WDFY3 | -0.17252 | 0.003426 | Other Negative Correlation |
| SH3BP5 | -0.17255 | 0.003418 | Other Negative Correlation |
| CYP4B1 | -0.17265 | 0.0034 | Other Negative Correlation |
| CBARP | -0.1727 | 0.003392 | Other Negative Correlation |
| ADGRE5 | -0.17273 | 0.003385 | Other Negative Correlation |
| PACC1 | -0.17277 | 0.003379 | Other Negative Correlation |
| NDUFA10 | -0.17281 | 0.003371 | Other Negative Correlation |
| PRRT3 | -0.17286 | 0.003361 | Other Negative Correlation |
| ZSWIM3 | -0.17289 | 0.003355 | Other Negative Correlation |
| GEMIN8 | -0.1729 | 0.003354 | Other Negative Correlation |
| CUL5 | -0.17301 | 0.003332 | Other Negative Correlation |
| BRI3 | -0.17302 | 0.003332 | Other Negative Correlation |
| TPGS2 | -0.17302 | 0.003332 | Other Negative Correlation |
| MRPS18A | -0.17304 | 0.003328 | Other Negative Correlation |
| UPK3B | -0.17311 | 0.003315 | Other Negative Correlation |
| MAMSTR | -0.17315 | 0.003308 | Other Negative Correlation |
| ARV1 | -0.17327 | 0.003285 | Other Negative Correlation |
| PEX16 | -0.17331 | 0.003277 | Other Negative Correlation |
| PRR3 | -0.17334 | 0.003273 | Other Negative Correlation |
| CDR2L | -0.17336 | 0.003268 | Other Negative Correlation |
| MED7 | -0.1734 | 0.003262 | Other Negative Correlation |
| TRAPPC6B | -0.17345 | 0.003252 | Other Negative Correlation |
| SSBP3 | -0.17346 | 0.003251 | Other Negative Correlation |
| EFL1 | -0.17352 | 0.003241 | Other Negative Correlation |
| ZNF644 | -0.17353 | 0.003239 | Other Negative Correlation |
| ZNF544 | -0.17353 | 0.003238 | Other Negative Correlation |
| PDE5A | -0.17359 | 0.003228 | Other Negative Correlation |
| PRKACB | -0.17359 | 0.003227 | Other Negative Correlation |
| LY6K | -0.17366 | 0.003215 | Other Negative Correlation |
| PTPRA | -0.17371 | 0.003206 | Other Negative Correlation |
| MICAL1 | -0.17374 | 0.003201 | Other Negative Correlation |
| NELFB | -0.17376 | 0.003198 | Other Negative Correlation |
| ZNF444 | -0.17381 | 0.003188 | Other Negative Correlation |
| ESYT2 | -0.17382 | 0.003186 | Other Negative Correlation |
| FKBP1B | -0.17399 | 0.003156 | Other Negative Correlation |
| SRPX | -0.17406 | 0.003144 | Other Negative Correlation |
| CLN8 | -0.17414 | 0.00313 | Other Negative Correlation |
| SNX8 | -0.17417 | 0.003124 | Other Negative Correlation |
| SNX31 | -0.17424 | 0.003112 | Other Negative Correlation |
| SPIRE2 | -0.17429 | 0.003105 | Other Negative Correlation |
| PTGS1 | -0.17431 | 0.003101 | Other Negative Correlation |
| PFKFB3 | -0.17436 | 0.003092 | Other Negative Correlation |
| ASIC1 | -0.17437 | 0.003089 | Other Negative Correlation |
| DHRS7B | -0.17441 | 0.003082 | Other Negative Correlation |
| COPS7B | -0.17441 | 0.003082 | Other Negative Correlation |

| | | | |
|---|---|---|---|
| VPS13B | -0.17447 | 0.003073 | Other Negative Correlation |
| BRD4 | -0.17467 | 0.003038 | Other Negative Correlation |
| STK4 | -0.17471 | 0.003031 | Other Negative Correlation |
| GDPD2 | -0.17474 | 0.003026 | Other Negative Correlation |
| HEATR5B | -0.17478 | 0.00302 | Other Negative Correlation |
| C12orf43 | -0.17478 | 0.003019 | Other Negative Correlation |
| TTC33 | -0.17479 | 0.003018 | Other Negative Correlation |
| ABL1 | -0.17481 | 0.003015 | Other Negative Correlation |
| BPHL | -0.17487 | 0.003004 | Other Negative Correlation |
| HIPK2 | -0.17489 | 0.003001 | Other Negative Correlation |
| IRAK3 | -0.17495 | 0.002992 | Other Negative Correlation |
| MT-CO1 | -0.175 | 0.002983 | Other Negative Correlation |
| ANKS1A | -0.17502 | 0.002979 | Other Negative Correlation |
| RAB40C | -0.17504 | 0.002977 | Other Negative Correlation |
| ALDH9A1 | -0.17504 | 0.002977 | Other Negative Correlation |
| CRNKL1 | -0.17504 | 0.002976 | Other Negative Correlation |
| AP2A2 | -0.17509 | 0.002968 | Other Negative Correlation |
| AP5M1 | -0.1751 | 0.002965 | Other Negative Correlation |
| ANKRD46 | -0.17518 | 0.002954 | Other Negative Correlation |
| UFSP1 | -0.17521 | 0.002948 | Other Negative Correlation |
| UTP25 | -0.17527 | 0.002939 | Other Negative Correlation |
| MCHR1 | -0.1753 | 0.002933 | Other Negative Correlation |
| KIAA1586 | -0.17538 | 0.002919 | Other Negative Correlation |
| TFCP2 | -0.17541 | 0.002915 | Other Negative Correlation |
| MYH10 | -0.17543 | 0.002911 | Other Negative Correlation |
| CEMIP2 | -0.17545 | 0.002909 | Other Negative Correlation |
| ZCCHC2 | -0.17546 | 0.002907 | Other Negative Correlation |
| IL6 | -0.17551 | 0.002899 | Other Negative Correlation |
| CYSTM1 | -0.17553 | 0.002896 | Other Negative Correlation |
| DKK4 | -0.17553 | 0.002896 | Other Negative Correlation |
| ASAP1 | -0.17555 | 0.002891 | Other Negative Correlation |
| HSF1 | -0.17556 | 0.00289 | Other Negative Correlation |
| ADPGK | -0.17557 | 0.00289 | Other Negative Correlation |
| ATP11A | -0.17559 | 0.002886 | Other Negative Correlation |
| IFIT5 | -0.17567 | 0.002872 | Other Negative Correlation |
| SYNGR1 | -0.17568 | 0.002871 | Other Negative Correlation |
| VPS51 | -0.17572 | 0.002865 | Other Negative Correlation |
| CENATAC | -0.17572 | 0.002864 | Other Negative Correlation |
| CLP1 | -0.17574 | 0.002861 | Other Negative Correlation |
| RNF103 | -0.17578 | 0.002856 | Other Negative Correlation |
| TBCK | -0.17581 | 0.002851 | Other Negative Correlation |
| PDZD8 | -0.17588 | 0.002839 | Other Negative Correlation |
| MED12 | -0.17596 | 0.002826 | Other Negative Correlation |
| PCDHGA12 | -0.17605 | 0.002812 | Other Negative Correlation |
| PITPNC1 | -0.17612 | 0.002801 | Other Negative Correlation |
| MITF | -0.17612 | 0.002801 | Other Negative Correlation |

| | | | |
|---|---|---|---|
| NEIL1 | -0.17619 | 0.00279 | Other Negative Correlation |
| GTF3C5 | -0.1763 | 0.002772 | Other Negative Correlation |
| SHTN1 | -0.17633 | 0.002767 | Other Negative Correlation |
| USF3 | -0.17634 | 0.002766 | Other Negative Correlation |
| TWF2 | -0.17636 | 0.002763 | Other Negative Correlation |
| SNAPIN | -0.17637 | 0.002761 | Other Negative Correlation |
| USP42 | -0.17639 | 0.002759 | Other Negative Correlation |
| SLC13A5 | -0.17643 | 0.002751 | Other Negative Correlation |
| NEDD1 | -0.17645 | 0.002749 | Other Negative Correlation |
| ANKRD54 | -0.1765 | 0.002741 | Other Negative Correlation |
| ELP6 | -0.17651 | 0.00274 | Other Negative Correlation |
| ABHD6 | -0.17658 | 0.002728 | Other Negative Correlation |
| S100A12 | -0.17664 | 0.00272 | Other Negative Correlation |
| TRAF4 | -0.1767 | 0.00271 | Other Negative Correlation |
| TMC7 | -0.17676 | 0.002701 | Other Negative Correlation |
| DDX28 | -0.1768 | 0.002696 | Other Negative Correlation |
| PICALM | -0.1768 | 0.002695 | Other Negative Correlation |
| KDM3B | -0.17681 | 0.002694 | Other Negative Correlation |
| GOLGA7 | -0.17686 | 0.002686 | Other Negative Correlation |
| ATL1 | -0.17692 | 0.002677 | Other Negative Correlation |
| C4orf33 | -0.17695 | 0.002673 | Other Negative Correlation |
| CD53 | -0.17697 | 0.002669 | Other Negative Correlation |
| TRPV2 | -0.17705 | 0.002656 | Other Negative Correlation |
| CSF2 | -0.17706 | 0.002655 | Other Negative Correlation |
| KLK12 | -0.17713 | 0.002645 | Other Negative Correlation |
| ZNF131 | -0.17717 | 0.002639 | Other Negative Correlation |
| NUDT6 | -0.17718 | 0.002638 | Other Negative Correlation |
| TM9SF2 | -0.1772 | 0.002635 | Other Negative Correlation |
| SLC38A9 | -0.17725 | 0.002627 | Other Negative Correlation |
| APOD | -0.17744 | 0.002599 | Other Negative Correlation |
| TM9SF4 | -0.17745 | 0.002597 | Other Negative Correlation |
| IL17RD | -0.17749 | 0.002592 | Other Negative Correlation |
| ULBP2 | -0.17753 | 0.002586 | Other Negative Correlation |
| ZNF813 | -0.17758 | 0.002578 | Other Negative Correlation |
| DMXL1 | -0.17762 | 0.002573 | Other Negative Correlation |
| NUDT18 | -0.1777 | 0.002561 | Other Negative Correlation |
| ACAP1 | -0.17772 | 0.002558 | Other Negative Correlation |
| CCDC69 | -0.17773 | 0.002557 | Other Negative Correlation |
| CKAP2 | -0.17779 | 0.002548 | Other Negative Correlation |
| ALKBH5 | -0.17781 | 0.002545 | Other Negative Correlation |
| TENT5B | -0.17805 | 0.00251 | Other Negative Correlation |
| EIF4ENIF1 | -0.17809 | 0.002504 | Other Negative Correlation |
| PPP5C | -0.1781 | 0.002503 | Other Negative Correlation |
| PRDX2 | -0.17812 | 0.002501 | Other Negative Correlation |
| PRELID2 | -0.17816 | 0.002495 | Other Negative Correlation |
| OCEL1 | -0.17818 | 0.002492 | Other Negative Correlation |

| | | | |
|---|---|---|---|
| PAQR4 | -0.17818 | 0.002491 | Other Negative Correlation |
| IRAK4 | -0.17842 | 0.002458 | Other Negative Correlation |
| FLOT2 | -0.17843 | 0.002457 | Other Negative Correlation |
| CHUK | -0.17845 | 0.002453 | Other Negative Correlation |
| ELL | -0.17846 | 0.002452 | Other Negative Correlation |
| OVOL1 | -0.17856 | 0.002438 | Other Negative Correlation |
| ASS1 | -0.17857 | 0.002436 | Other Negative Correlation |
| RAD17 | -0.17861 | 0.002432 | Other Negative Correlation |
| GTF3C3 | -0.17875 | 0.002412 | Other Negative Correlation |
| SLC25A22 | -0.17876 | 0.002411 | Other Negative Correlation |
| ZFYVE16 | -0.17881 | 0.002403 | Other Negative Correlation |
| AAMDC | -0.17883 | 0.002401 | Other Negative Correlation |
| NDUFA2 | -0.17883 | 0.002401 | Other Negative Correlation |
| ZNF425 | -0.17886 | 0.002397 | Other Negative Correlation |
| PNPLA8 | -0.1789 | 0.002391 | Other Negative Correlation |
| SRSF8 | -0.17895 | 0.002385 | Other Negative Correlation |
| DTNA | -0.17907 | 0.002369 | Other Negative Correlation |
| TCIRG1 | -0.17907 | 0.002369 | Other Negative Correlation |
| KCTD1 | -0.17912 | 0.002362 | Other Negative Correlation |
| MMP2 | -0.17912 | 0.002362 | Other Negative Correlation |
| SLC12A6 | -0.17912 | 0.002361 | Other Negative Correlation |
| PLCB4 | -0.17922 | 0.002348 | Other Negative Correlation |
| CYBC1 | -0.17926 | 0.002342 | Other Negative Correlation |
| TMEM245 | -0.17926 | 0.002342 | Other Negative Correlation |
| CIZ1 | -0.1793 | 0.002338 | Other Negative Correlation |
| ERP27 | -0.17935 | 0.002331 | Other Negative Correlation |
| RELL1 | -0.17935 | 0.00233 | Other Negative Correlation |
| PEX1 | -0.17935 | 0.00233 | Other Negative Correlation |
| CCR3 | -0.17941 | 0.002322 | Other Negative Correlation |
| HGS | -0.17944 | 0.002319 | Other Negative Correlation |
| TMEM30B | -0.17948 | 0.002313 | Other Negative Correlation |
| KIF7 | -0.17954 | 0.002305 | Other Negative Correlation |
| TMEM276 | -0.17955 | 0.002304 | Other Negative Correlation |
| RBBP9 | -0.17957 | 0.002302 | Other Negative Correlation |
| FDX1 | -0.17957 | 0.002301 | Other Negative Correlation |
| ZNF480 | -0.17962 | 0.002295 | Other Negative Correlation |
| PI4K2A | -0.17968 | 0.002287 | Other Negative Correlation |
| MEX3C | -0.17968 | 0.002286 | Other Negative Correlation |
| GOLGA1 | -0.17969 | 0.002285 | Other Negative Correlation |
| AZI2 | -0.17977 | 0.002275 | Other Negative Correlation |
| LPCAT4 | -0.17982 | 0.002269 | Other Negative Correlation |
| HS6ST1 | -0.17989 | 0.002259 | Other Negative Correlation |
| SIRPB2 | -0.17991 | 0.002256 | Other Negative Correlation |
| PLRG1 | -0.17994 | 0.002253 | Other Negative Correlation |
| MTOR | -0.18002 | 0.002243 | Other Negative Correlation |
| LNPK | -0.18004 | 0.00224 | Other Negative Correlation |

| | | | |
|---|---|---|---|
| SP3 | -0.18007 | 0.002235 | Other Negative Correlation |
| BORCS6 | -0.1801 | 0.002232 | Other Negative Correlation |
| POR | -0.18018 | 0.002222 | Other Negative Correlation |
| PINLYP | -0.18021 | 0.002219 | Other Negative Correlation |
| DEPP1 | -0.18023 | 0.002215 | Other Negative Correlation |
| RAB20 | -0.18029 | 0.002208 | Other Negative Correlation |
| KRT80 | -0.18046 | 0.002187 | Other Negative Correlation |
| FANK1 | -0.18061 | 0.002168 | Other Negative Correlation |
| GLT8D1 | -0.18067 | 0.002161 | Other Negative Correlation |
| RALB | -0.18072 | 0.002154 | Other Negative Correlation |
| TMEM45A | -0.18074 | 0.002151 | Other Negative Correlation |
| HBP1 | -0.18077 | 0.002148 | Other Negative Correlation |
| XPR1 | -0.18077 | 0.002147 | Other Negative Correlation |
| AUH | -0.18098 | 0.002122 | Other Negative Correlation |
| RBKS | -0.18104 | 0.002114 | Other Negative Correlation |
| PDE12 | -0.18108 | 0.002109 | Other Negative Correlation |
| FAXDC2 | -0.18113 | 0.002103 | Other Negative Correlation |
| PLD5 | -0.18115 | 0.002101 | Other Negative Correlation |
| DNER | -0.18124 | 0.00209 | Other Negative Correlation |
| EPS15 | -0.18127 | 0.002086 | Other Negative Correlation |
| ZNF846 | -0.18128 | 0.002085 | Other Negative Correlation |
| CENPB | -0.18128 | 0.002085 | Other Negative Correlation |
| TRMT1L | -0.18129 | 0.002083 | Other Negative Correlation |
| CASZ1 | -0.18132 | 0.00208 | Other Negative Correlation |
| XRRA1 | -0.18133 | 0.002079 | Other Negative Correlation |
| CFD | -0.18137 | 0.002074 | Other Negative Correlation |
| GNAL | -0.18139 | 0.002072 | Other Negative Correlation |
| MANSC1 | -0.1814 | 0.00207 | Other Negative Correlation |
| SPATA2L | -0.18143 | 0.002067 | Other Negative Correlation |
| PEX13 | -0.18144 | 0.002066 | Other Negative Correlation |
| CHST14 | -0.18149 | 0.002059 | Other Negative Correlation |
| TRRAP | -0.18156 | 0.002052 | Other Negative Correlation |
| TLCD3A | -0.18157 | 0.00205 | Other Negative Correlation |
| KAT6A | -0.1816 | 0.002047 | Other Negative Correlation |
| GUCY1A1 | -0.18161 | 0.002045 | Other Negative Correlation |
| LRIG1 | -0.18167 | 0.002039 | Other Negative Correlation |
| SEC23IP | -0.18169 | 0.002036 | Other Negative Correlation |
| AP4M1 | -0.18193 | 0.002008 | Other Negative Correlation |
| POLR2M | -0.18204 | 0.001995 | Other Negative Correlation |
| NOP9 | -0.18212 | 0.001986 | Other Negative Correlation |
| FAM83E | -0.18212 | 0.001986 | Other Negative Correlation |
| KMT5B | -0.18212 | 0.001985 | Other Negative Correlation |
| PRKCZ | -0.18215 | 0.001983 | Other Negative Correlation |
| CDK5 | -0.18222 | 0.001975 | Other Negative Correlation |
| CHFR | -0.18227 | 0.001969 | Other Negative Correlation |
| ETV3 | -0.18229 | 0.001967 | Other Negative Correlation |

| | | | |
|---|---|---|---|
| POLI | -0.18229 | 0.001967 | Other Negative Correlation |
| HECTD4 | -0.18234 | 0.001961 | Other Negative Correlation |
| KIAA2013 | -0.18235 | 0.00196 | Other Negative Correlation |
| CROCC | -0.18236 | 0.001958 | Other Negative Correlation |
| PTK6 | -0.18239 | 0.001955 | Other Negative Correlation |
| TGFB1 | -0.18243 | 0.00195 | Other Negative Correlation |
| SMPD4 | -0.18243 | 0.00195 | Other Negative Correlation |
| CORO1A | -0.18254 | 0.001938 | Other Negative Correlation |
| WDTC1 | -0.1826 | 0.001931 | Other Negative Correlation |
| FNDC3A | -0.1827 | 0.00192 | Other Negative Correlation |
| SNAPC3 | -0.18273 | 0.001916 | Other Negative Correlation |
| GNPDA1 | -0.18273 | 0.001916 | Other Negative Correlation |
| CHMP6 | -0.18276 | 0.001913 | Other Negative Correlation |
| MTHFD2L | -0.1828 | 0.001909 | Other Negative Correlation |
| RAP2C | -0.18288 | 0.0019 | Other Negative Correlation |
| NINJ1 | -0.18289 | 0.001898 | Other Negative Correlation |
| NMU | -0.18291 | 0.001897 | Other Negative Correlation |
| MMP1 | -0.18293 | 0.001894 | Other Negative Correlation |
| CSRNP1 | -0.18294 | 0.001893 | Other Negative Correlation |
| TAF1 | -0.18301 | 0.001885 | Other Negative Correlation |
| CPEB4 | -0.18306 | 0.00188 | Other Negative Correlation |
| SLC27A1 | -0.18307 | 0.001879 | Other Negative Correlation |
| SNW1 | -0.18317 | 0.001867 | Other Negative Correlation |
| H2BC8 | -0.18322 | 0.001862 | Other Negative Correlation |
| BCL2L2 | -0.18334 | 0.001849 | Other Negative Correlation |
| KDM5C | -0.18337 | 0.001846 | Other Negative Correlation |
| ANAPC4 | -0.18349 | 0.001833 | Other Negative Correlation |
| KCTD5 | -0.18357 | 0.001824 | Other Negative Correlation |
| SCAF4 | -0.18365 | 0.001817 | Other Negative Correlation |
| ZNF426 | -0.18383 | 0.001797 | Other Negative Correlation |
| LBH | -0.18385 | 0.001795 | Other Negative Correlation |
| EPM2AIP1 | -0.18386 | 0.001794 | Other Negative Correlation |
| LAMTOR3 | -0.18392 | 0.001788 | Other Negative Correlation |
| TNS3 | -0.18399 | 0.00178 | Other Negative Correlation |
| DCAF5 | -0.18404 | 0.001775 | Other Negative Correlation |
| ARAP3 | -0.18407 | 0.001772 | Other Negative Correlation |
| FHIP2B | -0.18407 | 0.001771 | Other Negative Correlation |
| USP5 | -0.1841 | 0.001769 | Other Negative Correlation |
| APOO | -0.18413 | 0.001766 | Other Negative Correlation |
| NMT2 | -0.18425 | 0.001754 | Other Negative Correlation |
| POLRMT | -0.18425 | 0.001753 | Other Negative Correlation |
| S100P | -0.18432 | 0.001746 | Other Negative Correlation |
| CDIN1 | -0.18432 | 0.001746 | Other Negative Correlation |
| COMMD10 | -0.18437 | 0.001741 | Other Negative Correlation |
| SLC6A14 | -0.1845 | 0.001727 | Other Negative Correlation |
| GOLM2 | -0.18451 | 0.001726 | Other Negative Correlation |

| | | | |
|---|---|---|---|
| ATE1 | -0.18461 | 0.001716 | Other Negative Correlation |
| ZNF320 | -0.18466 | 0.001711 | Other Negative Correlation |
| EML4 | -0.18467 | 0.00171 | Other Negative Correlation |
| OCLN | -0.18482 | 0.001696 | Other Negative Correlation |
| BLMH | -0.18482 | 0.001695 | Other Negative Correlation |
| BFAR | -0.1849 | 0.001687 | Other Negative Correlation |
| CYTH2 | -0.18495 | 0.001682 | Other Negative Correlation |
| MOSPD3 | -0.18499 | 0.001678 | Other Negative Correlation |
| SPSB2 | -0.18501 | 0.001676 | Other Negative Correlation |
| TEX264 | -0.18503 | 0.001674 | Other Negative Correlation |
| PHF2 | -0.18511 | 0.001667 | Other Negative Correlation |
| ZNF710 | -0.18514 | 0.001663 | Other Negative Correlation |
| FAM120B | -0.18518 | 0.00166 | Other Negative Correlation |
| PITPNM1 | -0.18521 | 0.001657 | Other Negative Correlation |
| TMEM127 | -0.18523 | 0.001655 | Other Negative Correlation |
| LSM10 | -0.18528 | 0.00165 | Other Negative Correlation |
| ACVR2A | -0.18529 | 0.001649 | Other Negative Correlation |
| RASA3 | -0.18534 | 0.001644 | Other Negative Correlation |
| CETN3 | -0.1854 | 0.001638 | Other Negative Correlation |
| SURF1 | -0.18542 | 0.001636 | Other Negative Correlation |
| APPBP2 | -0.18542 | 0.001636 | Other Negative Correlation |
| LPXN | -0.18543 | 0.001635 | Other Negative Correlation |
| KREMEN1 | -0.1855 | 0.001629 | Other Negative Correlation |
| ACAP3 | -0.18552 | 0.001626 | Other Negative Correlation |
| FOXO3 | -0.18553 | 0.001626 | Other Negative Correlation |
| C18orf21 | -0.18562 | 0.001617 | Other Negative Correlation |
| RNF216 | -0.18571 | 0.001608 | Other Negative Correlation |
| LRSAM1 | -0.18576 | 0.001604 | Other Negative Correlation |
| MKLN1 | -0.1858 | 0.0016 | Other Negative Correlation |
| IGSF9 | -0.18583 | 0.001597 | Other Negative Correlation |
| AP1G1 | -0.1859 | 0.00159 | Other Negative Correlation |
| AIFM2 | -0.18595 | 0.001586 | Other Negative Correlation |
| ZFPL1 | -0.18596 | 0.001585 | Other Negative Correlation |
| RNF44 | -0.186 | 0.001581 | Other Negative Correlation |
| NOTCH3 | -0.186 | 0.001581 | Other Negative Correlation |
| WDFY1 | -0.18601 | 0.001581 | Other Negative Correlation |
| TSTD3 | -0.18601 | 0.00158 | Other Negative Correlation |
| DDX23 | -0.18602 | 0.001579 | Other Negative Correlation |
| MRPL34 | -0.18625 | 0.001557 | Other Negative Correlation |
| DGCR2 | -0.18629 | 0.001554 | Other Negative Correlation |
| SPSB1 | -0.18634 | 0.001549 | Other Negative Correlation |
| PGAP6 | -0.18655 | 0.00153 | Other Negative Correlation |
| DUSP3 | -0.18655 | 0.00153 | Other Negative Correlation |
| SND1 | -0.18661 | 0.001524 | Other Negative Correlation |
| RHOT1 | -0.18662 | 0.001524 | Other Negative Correlation |
| ZNF567 | -0.1867 | 0.001516 | Other Negative Correlation |

| | | | |
|---|---|---|---|
| GPX4 | -0.1869 | 0.001499 | Other Negative Correlation |
| LTV1 | -0.18691 | 0.001498 | Other Negative Correlation |
| AKNAD1 | -0.18691 | 0.001498 | Other Negative Correlation |
| ZBTB1 | -0.18692 | 0.001497 | Other Negative Correlation |
| PUM2 | -0.18693 | 0.001496 | Other Negative Correlation |
| ASB13 | -0.18714 | 0.001478 | Other Negative Correlation |
| HLA-A | -0.18716 | 0.001476 | Other Negative Correlation |
| CCDC102A | -0.18717 | 0.001475 | Other Negative Correlation |
| ARHGAP30 | -0.1872 | 0.001472 | Other Negative Correlation |
| INPP1 | -0.18743 | 0.001452 | Other Negative Correlation |
| GRAMD1C | -0.18745 | 0.00145 | Other Negative Correlation |
| USP4 | -0.18747 | 0.001449 | Other Negative Correlation |
| RANBP3 | -0.1875 | 0.001446 | Other Negative Correlation |
| STX12 | -0.18761 | 0.001437 | Other Negative Correlation |
| SGMS1 | -0.18763 | 0.001435 | Other Negative Correlation |
| B3GALT4 | -0.18765 | 0.001433 | Other Negative Correlation |
| TAF11 | -0.18768 | 0.00143 | Other Negative Correlation |
| FAF2 | -0.1877 | 0.001429 | Other Negative Correlation |
| PUDP | -0.18776 | 0.001424 | Other Negative Correlation |
| PODNL1 | -0.18781 | 0.001419 | Other Negative Correlation |
| SLBP | -0.18783 | 0.001417 | Other Negative Correlation |
| VAMP7 | -0.18791 | 0.001411 | Other Negative Correlation |
| ZNF646 | -0.18791 | 0.001411 | Other Negative Correlation |
| ZC3H6 | -0.18805 | 0.001399 | Other Negative Correlation |
| PPP2R2D | -0.1881 | 0.001395 | Other Negative Correlation |
| HERC6 | -0.18815 | 0.00139 | Other Negative Correlation |
| B3GNT3 | -0.1882 | 0.001387 | Other Negative Correlation |
| RIPK4 | -0.18822 | 0.001385 | Other Negative Correlation |
| SEPTIN8 | -0.18822 | 0.001385 | Other Negative Correlation |
| DYSF | -0.18826 | 0.001382 | Other Negative Correlation |
| CRTC2 | -0.18828 | 0.00138 | Other Negative Correlation |
| STAT1 | -0.18829 | 0.001379 | Other Negative Correlation |
| DNAJC30 | -0.18832 | 0.001377 | Other Negative Correlation |
| USP37 | -0.18845 | 0.001366 | Other Negative Correlation |
| RNF10 | -0.18846 | 0.001365 | Other Negative Correlation |
| SCAP | -0.18848 | 0.001364 | Other Negative Correlation |
| JPH2 | -0.18848 | 0.001364 | Other Negative Correlation |
| VPS26B | -0.1885 | 0.001362 | Other Negative Correlation |
| CTSL | -0.18851 | 0.001361 | Other Negative Correlation |
| MT-ND1 | -0.18851 | 0.001361 | Other Negative Correlation |
| AMTN | -0.18858 | 0.001356 | Other Negative Correlation |
| TSPAN10 | -0.1886 | 0.001354 | Other Negative Correlation |
| DOP1A | -0.18863 | 0.001352 | Other Negative Correlation |
| SGPP2 | -0.18879 | 0.001338 | Other Negative Correlation |
| COG4 | -0.18884 | 0.001334 | Other Negative Correlation |
| SPART | -0.18889 | 0.00133 | Other Negative Correlation |

| | | | |
|---|---|---|---|
| CPLANE1 | -0.18892 | 0.001328 | Other Negative Correlation |
| SIN3A | -0.18909 | 0.001315 | Other Negative Correlation |
| SEC23A | -0.18909 | 0.001314 | Other Negative Correlation |
| STK35 | -0.18914 | 0.00131 | Other Negative Correlation |
| SMAD4 | -0.18915 | 0.00131 | Other Negative Correlation |
| SDHB | -0.18916 | 0.001309 | Other Negative Correlation |
| ENSG00000287856 | -0.18952 | 0.00128 | Other Negative Correlation |
| C1RL | -0.18954 | 0.001279 | Other Negative Correlation |
| GAREM1 | -0.18954 | 0.001279 | Other Negative Correlation |
| ZC3H12C | -0.18954 | 0.001279 | Other Negative Correlation |
| GALNS | -0.18955 | 0.001278 | Other Negative Correlation |
| DDX42 | -0.18966 | 0.00127 | Other Negative Correlation |
| TENT2 | -0.18968 | 0.001268 | Other Negative Correlation |
| TENM4 | -0.18972 | 0.001265 | Other Negative Correlation |
| C1orf52 | -0.18974 | 0.001264 | Other Negative Correlation |
| OBSCN | -0.18978 | 0.001261 | Other Negative Correlation |
| GDI1 | -0.18981 | 0.001259 | Other Negative Correlation |
| AKAP8L | -0.18989 | 0.001253 | Other Negative Correlation |
| MAF | -0.18989 | 0.001252 | Other Negative Correlation |
| TECR | -0.18991 | 0.001251 | Other Negative Correlation |
| RABIF | -0.18991 | 0.001251 | Other Negative Correlation |
| FAM83G | -0.18992 | 0.00125 | Other Negative Correlation |
| FGR | -0.18993 | 0.001249 | Other Negative Correlation |
| STK10 | -0.18999 | 0.001245 | Other Negative Correlation |
| FASTKD1 | -0.19006 | 0.001239 | Other Negative Correlation |
| ELFN2 | -0.19017 | 0.001231 | Other Negative Correlation |
| E2F3 | -0.19017 | 0.001231 | Other Negative Correlation |
| TPRG1L | -0.19022 | 0.001227 | Other Negative Correlation |
| ZNF287 | -0.19036 | 0.001217 | Other Negative Correlation |
| PIAS4 | -0.19041 | 0.001214 | Other Negative Correlation |
| MAGEF1 | -0.19043 | 0.001212 | Other Negative Correlation |
| BIRC2 | -0.19045 | 0.001211 | Other Negative Correlation |
| ZNF703 | -0.19057 | 0.001202 | Other Negative Correlation |
| STX7 | -0.19059 | 0.0012 | Other Negative Correlation |
| NAGK | -0.19062 | 0.001198 | Other Negative Correlation |
| HDX | -0.19067 | 0.001195 | Other Negative Correlation |
| RFK | -0.19068 | 0.001194 | Other Negative Correlation |
| ELOF1 | -0.19071 | 0.001192 | Other Negative Correlation |
| MAP2K7 | -0.19074 | 0.00119 | Other Negative Correlation |
| PWWP2B | -0.19077 | 0.001187 | Other Negative Correlation |
| HLA-DOB | -0.19078 | 0.001187 | Other Negative Correlation |
| MINPP1 | -0.19078 | 0.001186 | Other Negative Correlation |
| DDI2 | -0.19081 | 0.001184 | Other Negative Correlation |
| WFDC3 | -0.19083 | 0.001183 | Other Negative Correlation |
| ZNF281 | -0.19095 | 0.001174 | Other Negative Correlation |
| MGAT5 | -0.19096 | 0.001174 | Other Negative Correlation |

| | | | |
|---|---|---|---|
| ARMCX3 | -0.19099 | 0.001172 | Other Negative Correlation |
| PICK1 | -0.19104 | 0.001168 | Other Negative Correlation |
| C2orf49 | -0.19106 | 0.001166 | Other Negative Correlation |
| CALCRL | -0.1911 | 0.001164 | Other Negative Correlation |
| KLHL29 | -0.19111 | 0.001163 | Other Negative Correlation |
| MEIS2 | -0.19112 | 0.001162 | Other Negative Correlation |
| C3orf38 | -0.19116 | 0.001159 | Other Negative Correlation |
| NFAT5 | -0.19118 | 0.001158 | Other Negative Correlation |
| TMEM19 | -0.19118 | 0.001158 | Other Negative Correlation |
| SNX10 | -0.1912 | 0.001157 | Other Negative Correlation |
| OTUD5 | -0.19139 | 0.001143 | Other Negative Correlation |
| B3GNT2 | -0.19142 | 0.001142 | Other Negative Correlation |
| FHOD3 | -0.19144 | 0.00114 | Other Negative Correlation |
| CYTH1 | -0.19145 | 0.001139 | Other Negative Correlation |
| ASAH2B | -0.19153 | 0.001134 | Other Negative Correlation |
| PDGFB | -0.19172 | 0.001121 | Other Negative Correlation |
| FBXL2 | -0.19175 | 0.001118 | Other Negative Correlation |
| MYO5B | -0.19181 | 0.001114 | Other Negative Correlation |
| RILPL2 | -0.19182 | 0.001114 | Other Negative Correlation |
| DNPEP | -0.19201 | 0.001101 | Other Negative Correlation |
| ITSN2 | -0.19204 | 0.001099 | Other Negative Correlation |
| ASF1A | -0.1921 | 0.001095 | Other Negative Correlation |
| KIF1B | -0.19214 | 0.001092 | Other Negative Correlation |
| RSPRY1 | -0.19222 | 0.001087 | Other Negative Correlation |
| ZNF579 | -0.19231 | 0.001081 | Other Negative Correlation |
| TMEM139 | -0.19231 | 0.001081 | Other Negative Correlation |
| COX16 | -0.19231 | 0.001081 | Other Negative Correlation |
| TMEM17 | -0.19233 | 0.001079 | Other Negative Correlation |
| LEPR | -0.19236 | 0.001078 | Other Negative Correlation |
| STEAP3 | -0.19237 | 0.001077 | Other Negative Correlation |
| MBNL2 | -0.19238 | 0.001076 | Other Negative Correlation |
| SOWAHB | -0.19239 | 0.001076 | Other Negative Correlation |
| ERCC6L2 | -0.19239 | 0.001076 | Other Negative Correlation |
| ISCU | -0.19239 | 0.001075 | Other Negative Correlation |
| SYS1 | -0.19247 | 0.00107 | Other Negative Correlation |
| GPBP1 | -0.19248 | 0.00107 | Other Negative Correlation |
| MUC20 | -0.19249 | 0.001069 | Other Negative Correlation |
| B4GALT6 | -0.19254 | 0.001066 | Other Negative Correlation |
| ZNF274 | -0.19254 | 0.001066 | Other Negative Correlation |
| MAFB | -0.19255 | 0.001065 | Other Negative Correlation |
| FXYD5 | -0.19267 | 0.001057 | Other Negative Correlation |
| SLC7A4 | -0.19268 | 0.001057 | Other Negative Correlation |
| S100A4 | -0.19282 | 0.001048 | Other Negative Correlation |
| DGAT1 | -0.19283 | 0.001047 | Other Negative Correlation |
| STPG1 | -0.19288 | 0.001044 | Other Negative Correlation |
| FAM171A1 | -0.19288 | 0.001044 | Other Negative Correlation |

| | | | |
|---|---|---|---|
| MB21D2 | -0.1929 | 0.001042 | Other Negative Correlation |
| NUP214 | -0.19292 | 0.001041 | Other Negative Correlation |
| TH | -0.19295 | 0.001039 | Other Negative Correlation |
| PLD2 | -0.19303 | 0.001034 | Other Negative Correlation |
| MMP10 | -0.19306 | 0.001032 | Other Negative Correlation |
| WSB2 | -0.19317 | 0.001025 | Other Negative Correlation |
| ARID1B | -0.19317 | 0.001025 | Other Negative Correlation |
| BORCS8 | -0.19318 | 0.001025 | Other Negative Correlation |
| SIPA1L3 | -0.19322 | 0.001022 | Other Negative Correlation |
| ARNT | -0.19322 | 0.001022 | Other Negative Correlation |
| CERS5 | -0.19331 | 0.001017 | Other Negative Correlation |
| LARP6 | -0.19331 | 0.001017 | Other Negative Correlation |
| APPL1 | -0.1934 | 0.001011 | Other Negative Correlation |
| ECM1 | -0.19342 | 0.001009 | Other Negative Correlation |
| DNAJB4 | -0.1935 | 0.001005 | Other Negative Correlation |
| NCOA3 | -0.19351 | 0.001004 | Other Negative Correlation |
| RAB35 | -0.19352 | 0.001004 | Other Negative Correlation |
| CLK3 | -0.19366 | 0.000995 | Other Negative Correlation |
| HNRNPLL | -0.19373 | 0.000991 | Other Negative Correlation |
| TTPAL | -0.19378 | 0.000988 | Other Negative Correlation |
| CYB5D2 | -0.19386 | 0.000982 | Other Negative Correlation |
| SMARCAL1 | -0.1939 | 0.00098 | Other Negative Correlation |
| CYP2S1 | -0.194 | 0.000975 | Other Negative Correlation |
| PTPN3 | -0.19413 | 0.000966 | Other Negative Correlation |
| SLC35F6 | -0.19417 | 0.000964 | Other Negative Correlation |
| ARAF | -0.19419 | 0.000963 | Other Negative Correlation |
| SLC37A3 | -0.19419 | 0.000963 | Other Negative Correlation |
| PDGFC | -0.19423 | 0.000961 | Other Negative Correlation |
| SPRED2 | -0.19424 | 0.00096 | Other Negative Correlation |
| SUN2 | -0.19426 | 0.000959 | Other Negative Correlation |
| WAPL | -0.19426 | 0.000959 | Other Negative Correlation |
| MTIF3 | -0.19428 | 0.000958 | Other Negative Correlation |
| PURA | -0.19429 | 0.000957 | Other Negative Correlation |
| GRAMD4 | -0.19429 | 0.000957 | Other Negative Correlation |
| ATXN2 | -0.19433 | 0.000955 | Other Negative Correlation |
| S100A7 | -0.19446 | 0.000947 | Other Negative Correlation |
| PLA2R1 | -0.19455 | 0.000942 | Other Negative Correlation |
| IRF5 | -0.19456 | 0.000941 | Other Negative Correlation |
| TMEM128 | -0.19466 | 0.000936 | Other Negative Correlation |
| EDC3 | -0.19468 | 0.000934 | Other Negative Correlation |
| GAB1 | -0.19473 | 0.000931 | Other Negative Correlation |
| FKBP15 | -0.19475 | 0.00093 | Other Negative Correlation |
| DAB2IP | -0.19478 | 0.000929 | Other Negative Correlation |
| CAMP | -0.19489 | 0.000922 | Other Negative Correlation |
| TTC8 | -0.1951 | 0.00091 | Other Negative Correlation |
| MAN2B2 | -0.1951 | 0.00091 | Other Negative Correlation |

| | | | |
|---|---|---|---|
| ANKRD6 | -0.19515 | 0.000907 | Other Negative Correlation |
| RPP25L | -0.19525 | 0.000902 | Other Negative Correlation |
| WASHC5 | -0.19529 | 0.0009 | Other Negative Correlation |
| EPS8L1 | -0.19533 | 0.000897 | Other Negative Correlation |
| HSD11B1L | -0.19533 | 0.000897 | Other Negative Correlation |
| GAS2L1 | -0.19541 | 0.000893 | Other Negative Correlation |
| DENND3 | -0.19542 | 0.000892 | Other Negative Correlation |
| HYAL1 | -0.19546 | 0.00089 | Other Negative Correlation |
| ZMYM2 | -0.19552 | 0.000886 | Other Negative Correlation |
| FBXW9 | -0.19572 | 0.000876 | Other Negative Correlation |
| MTX2 | -0.19577 | 0.000873 | Other Negative Correlation |
| OBSL1 | -0.19581 | 0.000871 | Other Negative Correlation |
| PRRG1 | -0.19581 | 0.000871 | Other Negative Correlation |
| S1PR4 | -0.19601 | 0.00086 | Other Negative Correlation |
| VILL | -0.19602 | 0.00086 | Other Negative Correlation |
| NELFA | -0.19603 | 0.000859 | Other Negative Correlation |
| ARL14 | -0.19625 | 0.000847 | Other Negative Correlation |
| SYNC | -0.19627 | 0.000846 | Other Negative Correlation |
| SIK2 | -0.19633 | 0.000843 | Other Negative Correlation |
| SELENOO | -0.19635 | 0.000842 | Other Negative Correlation |
| GMIP | -0.19637 | 0.000841 | Other Negative Correlation |
| TECPR2 | -0.19645 | 0.000837 | Other Negative Correlation |
| TMC1 | -0.19648 | 0.000835 | Other Negative Correlation |
| TRIM33 | -0.19657 | 0.000831 | Other Negative Correlation |
| VPS4A | -0.19658 | 0.00083 | Other Negative Correlation |
| SUFU | -0.19662 | 0.000828 | Other Negative Correlation |
| CYP2W1 | -0.19667 | 0.000825 | Other Negative Correlation |
| SIKE1 | -0.19671 | 0.000823 | Other Negative Correlation |
| RING1 | -0.19674 | 0.000822 | Other Negative Correlation |
| INF2 | -0.19677 | 0.00082 | Other Negative Correlation |
| SAMD4B | -0.19693 | 0.000812 | Other Negative Correlation |
| ARL6IP1 | -0.19694 | 0.000812 | Other Negative Correlation |
| CERS6 | -0.19706 | 0.000805 | Other Negative Correlation |
| LANCL2 | -0.19712 | 0.000802 | Other Negative Correlation |
| MPPE1 | -0.19718 | 0.0008 | Other Negative Correlation |
| GOSR1 | -0.19721 | 0.000798 | Other Negative Correlation |
| CYTH3 | -0.19727 | 0.000795 | Other Negative Correlation |
| LTBR | -0.19738 | 0.000789 | Other Negative Correlation |
| KLHL18 | -0.19747 | 0.000785 | Other Negative Correlation |
| TRAPPC13 | -0.19775 | 0.000771 | Other Negative Correlation |
| BLOC1S4 | -0.19781 | 0.000769 | Other Negative Correlation |
| COG2 | -0.19783 | 0.000767 | Other Negative Correlation |
| ZNF518B | -0.19796 | 0.000761 | Other Negative Correlation |
| TUT4 | -0.198 | 0.000759 | Other Negative Correlation |
| EMC8 | -0.19804 | 0.000757 | Other Negative Correlation |
| GPR155 | -0.19809 | 0.000755 | Other Negative Correlation |

| | | | |
|---|---|---|---|
| TACC2 | -0.19809 | 0.000755 | Other Negative Correlation |
| PALS1 | -0.19817 | 0.000751 | Other Negative Correlation |
| FAM25A | -0.19835 | 0.000743 | Other Negative Correlation |
| LMAN2 | -0.19835 | 0.000743 | Other Negative Correlation |
| KIF16B | -0.19836 | 0.000742 | Other Negative Correlation |
| NXPE3 | -0.19843 | 0.000739 | Other Negative Correlation |
| TRANK1 | -0.19844 | 0.000739 | Other Negative Correlation |
| GRK4 | -0.19846 | 0.000738 | Other Negative Correlation |
| PNKD | -0.19849 | 0.000736 | Other Negative Correlation |
| TMEM267 | -0.1985 | 0.000736 | Other Negative Correlation |
| WASL | -0.19851 | 0.000735 | Other Negative Correlation |
| CDH5 | -0.19851 | 0.000735 | Other Negative Correlation |
| ZNF143 | -0.19854 | 0.000734 | Other Negative Correlation |
| IST1 | -0.19855 | 0.000733 | Other Negative Correlation |
| FBXW5 | -0.19862 | 0.00073 | Other Negative Correlation |
| GAS8 | -0.19866 | 0.000728 | Other Negative Correlation |
| ETFRF1 | -0.19869 | 0.000727 | Other Negative Correlation |
| ACOX3 | -0.19872 | 0.000726 | Other Negative Correlation |
| XAB2 | -0.19873 | 0.000725 | Other Negative Correlation |
| AGGF1 | -0.19889 | 0.000718 | Other Negative Correlation |
| MAMDC2 | -0.19902 | 0.000712 | Other Negative Correlation |
| TMEM192 | -0.19905 | 0.00071 | Other Negative Correlation |
| SPRYD4 | -0.1991 | 0.000709 | Other Negative Correlation |
| SEC24C | -0.19911 | 0.000708 | Other Negative Correlation |
| TMEM120A | -0.19927 | 0.000701 | Other Negative Correlation |
| MROH6 | -0.19927 | 0.000701 | Other Negative Correlation |
| SH2D4A | -0.19931 | 0.000699 | Other Negative Correlation |
| FBXL20 | -0.19943 | 0.000694 | Other Negative Correlation |
| THEMIS2 | -0.19943 | 0.000694 | Other Negative Correlation |
| TARBP2 | -0.19952 | 0.00069 | Other Negative Correlation |
| ZNF688 | -0.19952 | 0.00069 | Other Negative Correlation |
| CTSV | -0.19953 | 0.000689 | Other Negative Correlation |
| LMBRD2 | -0.19957 | 0.000688 | Other Negative Correlation |
| FAM114A2 | -0.19958 | 0.000687 | Other Negative Correlation |
| VPS18 | -0.19959 | 0.000687 | Other Negative Correlation |
| ATL2 | -0.19959 | 0.000687 | Other Negative Correlation |
| GIT2 | -0.19967 | 0.000683 | Other Negative Correlation |
| KRT34 | -0.19972 | 0.000681 | Other Negative Correlation |
| POP4 | -0.19977 | 0.000679 | Other Negative Correlation |
| EPB41L5 | -0.2 | 0.000669 | Other Negative Correlation |
| DEAF1 | -0.20016 | 0.000662 | Other Negative Correlation |
| CILK1 | -0.20025 | 0.000659 | Other Negative Correlation |
| HAPSTR1 | -0.2003 | 0.000657 | Other Negative Correlation |
| LHFPL6 | -0.20031 | 0.000656 | Other Negative Correlation |
| KRT9 | -0.20031 | 0.000656 | Other Negative Correlation |
| ARHGAP12 | -0.20046 | 0.00065 | Other Negative Correlation |

| | | | |
|---|---|---|---|
| GRK2 | -0.20047 | 0.000649 | Other Negative Correlation |
| AP4E1 | -0.20049 | 0.000648 | Other Negative Correlation |
| TMEM14A | -0.2005 | 0.000648 | Other Negative Correlation |
| NEURL1B | -0.20062 | 0.000643 | Other Negative Correlation |
| B4GALNT3 | -0.20064 | 0.000643 | Other Negative Correlation |
| EME2 | -0.20073 | 0.000639 | Other Negative Correlation |
| HIBCH | -0.20075 | 0.000638 | Other Negative Correlation |
| NEK7 | -0.20082 | 0.000635 | Other Negative Correlation |
| MATN2 | -0.20082 | 0.000635 | Other Negative Correlation |
| FOXK2 | -0.20089 | 0.000632 | Other Negative Correlation |
| CYSRT1 | -0.20089 | 0.000632 | Other Negative Correlation |
| ZNF37A | -0.20092 | 0.000631 | Other Negative Correlation |
| MAGT1 | -0.20094 | 0.00063 | Other Negative Correlation |
| ERAL1 | -0.20102 | 0.000627 | Other Negative Correlation |
| RAD21 | -0.20103 | 0.000627 | Other Negative Correlation |
| TERF1 | -0.20118 | 0.000621 | Other Negative Correlation |
| FOXC1 | -0.20125 | 0.000618 | Other Negative Correlation |
| RAI14 | -0.20126 | 0.000617 | Other Negative Correlation |
| TCP11L2 | -0.20131 | 0.000616 | Other Negative Correlation |
| DUOXA1 | -0.20152 | 0.000607 | Other Negative Correlation |
| COA5 | -0.20158 | 0.000605 | Other Negative Correlation |
| ACP2 | -0.20161 | 0.000604 | Other Negative Correlation |
| PCDHGC3 | -0.20161 | 0.000604 | Other Negative Correlation |
| EPB41L4B | -0.20162 | 0.000603 | Other Negative Correlation |
| EMC6 | -0.20169 | 0.000601 | Other Negative Correlation |
| TNFAIP2 | -0.20169 | 0.000601 | Other Negative Correlation |
| PEX11G | -0.20187 | 0.000594 | Other Negative Correlation |
| PRDM1 | -0.20194 | 0.000591 | Other Negative Correlation |
| ZDHHC4 | -0.20197 | 0.00059 | Other Negative Correlation |
| EXOC1 | -0.20204 | 0.000587 | Other Negative Correlation |
| SEC14L1 | -0.2022 | 0.000581 | Other Negative Correlation |
| TMEM9 | -0.20228 | 0.000579 | Other Negative Correlation |
| THNSL2 | -0.20232 | 0.000577 | Other Negative Correlation |
| ZNF772 | -0.20236 | 0.000575 | Other Negative Correlation |
| DBR1 | -0.20246 | 0.000572 | Other Negative Correlation |
| L3MBTL2 | -0.20249 | 0.000571 | Other Negative Correlation |
| COPS5 | -0.20249 | 0.000571 | Other Negative Correlation |
| LATS2 | -0.20252 | 0.000569 | Other Negative Correlation |
| SPRY1 | -0.20255 | 0.000568 | Other Negative Correlation |
| PIP4K2C | -0.20255 | 0.000568 | Other Negative Correlation |
| CLDN16 | -0.20258 | 0.000567 | Other Negative Correlation |
| PLXNA2 | -0.20267 | 0.000564 | Other Negative Correlation |
| TNNT1 | -0.20267 | 0.000564 | Other Negative Correlation |
| MAML3 | -0.20271 | 0.000563 | Other Negative Correlation |
| CSTPP1 | -0.20272 | 0.000562 | Other Negative Correlation |
| BTN2A2 | -0.20272 | 0.000562 | Other Negative Correlation |

| | | | |
|---|---|---|---|
| AACS | -0.20278 | 0.00056 | Other Negative Correlation |
| STIM1 | -0.2028 | 0.00056 | Other Negative Correlation |
| SMG6 | -0.20281 | 0.000559 | Other Negative Correlation |
| CRYL1 | -0.20285 | 0.000558 | Other Negative Correlation |
| CXCL6 | -0.20291 | 0.000556 | Other Negative Correlation |
| MYL5 | -0.20291 | 0.000556 | Other Negative Correlation |
| RB1 | -0.20292 | 0.000555 | Other Negative Correlation |
| DNAJC13 | -0.20293 | 0.000555 | Other Negative Correlation |
| COX11 | -0.20294 | 0.000554 | Other Negative Correlation |
| FAM111A | -0.20295 | 0.000554 | Other Negative Correlation |
| ZDHHC3 | -0.203 | 0.000552 | Other Negative Correlation |
| MIGA1 | -0.20302 | 0.000551 | Other Negative Correlation |
| AZGP1 | -0.20303 | 0.000551 | Other Negative Correlation |
| ABHD5 | -0.20306 | 0.00055 | Other Negative Correlation |
| SLC30A9 | -0.20306 | 0.00055 | Other Negative Correlation |
| DLAT | -0.20306 | 0.00055 | Other Negative Correlation |
| TRIP11 | -0.20314 | 0.000547 | Other Negative Correlation |
| IRF1 | -0.20327 | 0.000543 | Other Negative Correlation |
| ADSS1 | -0.20342 | 0.000538 | Other Negative Correlation |
| ARHGEF10L | -0.20348 | 0.000536 | Other Negative Correlation |
| INTS5 | -0.20355 | 0.000533 | Other Negative Correlation |
| CPM | -0.20356 | 0.000533 | Other Negative Correlation |
| ENDOV | -0.2036 | 0.000531 | Other Negative Correlation |
| TRIM4 | -0.2036 | 0.000531 | Other Negative Correlation |
| ORC4 | -0.2036 | 0.000531 | Other Negative Correlation |
| DHX8 | -0.20368 | 0.000529 | Other Negative Correlation |
| SENP6 | -0.20371 | 0.000528 | Other Negative Correlation |
| DUSP22 | -0.20382 | 0.000524 | Other Negative Correlation |
| CSNK2B | -0.20387 | 0.000522 | Other Negative Correlation |
| TBC1D23 | -0.2039 | 0.000521 | Other Negative Correlation |
| WNT10A | -0.2042 | 0.000511 | Other Negative Correlation |
| INPP4A | -0.20423 | 0.00051 | Other Negative Correlation |
| ANGEL1 | -0.20425 | 0.000509 | Other Negative Correlation |
| METTL18 | -0.20428 | 0.000508 | Other Negative Correlation |
| TMEM199 | -0.20441 | 0.000504 | Other Negative Correlation |
| CERS4 | -0.20444 | 0.000503 | Other Negative Correlation |
| RO60 | -0.20445 | 0.000503 | Other Negative Correlation |
| TMEM248 | -0.20445 | 0.000503 | Other Negative Correlation |
| RNMT | -0.20447 | 0.000502 | Other Negative Correlation |
| JADE2 | -0.20451 | 0.000501 | Other Negative Correlation |
| LMLN | -0.20453 | 0.0005 | Other Negative Correlation |
| TMED1 | -0.20454 | 0.0005 | Other Negative Correlation |
| CHST11 | -0.20454 | 0.0005 | Other Negative Correlation |
| ABCB7 | -0.20455 | 0.0005 | Other Negative Correlation |
| NADK | -0.20456 | 0.000499 | Other Negative Correlation |
| DYNC2LI1 | -0.20458 | 0.000499 | Other Negative Correlation |

| | | | |
|---|---|---|---|
| DNAJC24 | -0.20458 | 0.000498 | Other Negative Correlation |
| VPS13A | -0.20464 | 0.000497 | Other Negative Correlation |
| ZSWIM6 | -0.20466 | 0.000496 | Other Negative Correlation |
| TERF2IP | -0.20473 | 0.000494 | Other Negative Correlation |
| ADAM17 | -0.20475 | 0.000493 | Other Negative Correlation |
| NFKBIE | -0.20495 | 0.000487 | Other Negative Correlation |
| ISOC2 | -0.20495 | 0.000487 | Other Negative Correlation |
| ABT1 | -0.20503 | 0.000484 | Other Negative Correlation |
| TXNDC15 | -0.20504 | 0.000484 | Other Negative Correlation |
| ABCD3 | -0.20506 | 0.000483 | Other Negative Correlation |
| TAF6 | -0.2052 | 0.000479 | Other Negative Correlation |
| MRPL57 | -0.20527 | 0.000477 | Other Negative Correlation |
| RNF19B | -0.20527 | 0.000477 | Other Negative Correlation |
| PTCD3 | -0.20527 | 0.000477 | Other Negative Correlation |
| C6orf226 | -0.20532 | 0.000475 | Other Negative Correlation |
| MYO1D | -0.20534 | 0.000475 | Other Negative Correlation |
| HSPG2 | -0.20545 | 0.000471 | Other Negative Correlation |
| CLDN9 | -0.20552 | 0.000469 | Other Negative Correlation |
| ANKLE2 | -0.20554 | 0.000468 | Other Negative Correlation |
| ZNF197 | -0.20556 | 0.000468 | Other Negative Correlation |
| TIGD2 | -0.20562 | 0.000466 | Other Negative Correlation |
| ARHGEF9 | -0.20573 | 0.000463 | Other Negative Correlation |
| PLB1 | -0.20599 | 0.000455 | Other Negative Correlation |
| CCNJL | -0.20608 | 0.000452 | Other Negative Correlation |
| PDHX | -0.20612 | 0.000451 | Other Negative Correlation |
| RNF25 | -0.2062 | 0.000449 | Other Negative Correlation |
| TVP23B | -0.20624 | 0.000448 | Other Negative Correlation |
| AKT1S1 | -0.20626 | 0.000447 | Other Negative Correlation |
| LACC1 | -0.20629 | 0.000446 | Other Negative Correlation |
| EEF1E1 | -0.20632 | 0.000445 | Other Negative Correlation |
| TRIM26 | -0.20632 | 0.000445 | Other Negative Correlation |
| SLC30A7 | -0.20636 | 0.000444 | Other Negative Correlation |
| CHMP4C | -0.2064 | 0.000443 | Other Negative Correlation |
| FAM50A | -0.20643 | 0.000442 | Other Negative Correlation |
| CHI3L1 | -0.20643 | 0.000442 | Other Negative Correlation |
| RHOB | -0.20656 | 0.000438 | Other Negative Correlation |
| EVL | -0.20662 | 0.000437 | Other Negative Correlation |
| DGLUCY | -0.20664 | 0.000436 | Other Negative Correlation |
| RESF1 | -0.20664 | 0.000436 | Other Negative Correlation |
| WDR7 | -0.20672 | 0.000434 | Other Negative Correlation |
| RTN2 | -0.20687 | 0.000429 | Other Negative Correlation |
| TNF | -0.20688 | 0.000429 | Other Negative Correlation |
| KCTD10 | -0.20691 | 0.000428 | Other Negative Correlation |
| LTC4S | -0.20713 | 0.000422 | Other Negative Correlation |
| NTHL1 | -0.20717 | 0.000421 | Other Negative Correlation |
| TMEM258 | -0.20719 | 0.00042 | Other Negative Correlation |

| | | | |
|---|---|---|---|
| CLPTM1L | -0.20721 | 0.00042 | Other Negative Correlation |
| ARSD | -0.20727 | 0.000418 | Other Negative Correlation |
| EFNA4 | -0.20731 | 0.000417 | Other Negative Correlation |
| XRN1 | -0.20741 | 0.000414 | Other Negative Correlation |
| EPB41 | -0.20742 | 0.000414 | Other Negative Correlation |
| PRKCH | -0.20747 | 0.000413 | Other Negative Correlation |
| CHCHD7 | -0.20749 | 0.000412 | Other Negative Correlation |
| EYA2 | -0.20761 | 0.000409 | Other Negative Correlation |
| POMK | -0.20775 | 0.000405 | Other Negative Correlation |
| FRA10AC1 | -0.20785 | 0.000403 | Other Negative Correlation |
| SELENON | -0.20786 | 0.000402 | Other Negative Correlation |
| RGP1 | -0.20787 | 0.000402 | Other Negative Correlation |
| MTRF1 | -0.20795 | 0.0004 | Other Negative Correlation |
| DNAJB14 | -0.20797 | 0.000399 | Other Negative Correlation |
| COBLL1 | -0.208 | 0.000399 | Other Negative Correlation |
| ZNF628 | -0.2081 | 0.000396 | Other Negative Correlation |
| ACTR5 | -0.20813 | 0.000395 | Other Negative Correlation |
| NOSIP | -0.20827 | 0.000392 | Other Negative Correlation |
| RABGAP1L | -0.20829 | 0.000391 | Other Negative Correlation |
| LTN1 | -0.20841 | 0.000388 | Other Negative Correlation |
| GAK | -0.20841 | 0.000388 | Other Negative Correlation |
| FBXL19 | -0.2085 | 0.000386 | Other Negative Correlation |
| EVI5 | -0.20862 | 0.000383 | Other Negative Correlation |
| TIAM1 | -0.20862 | 0.000383 | Other Negative Correlation |
| EML3 | -0.20867 | 0.000381 | Other Negative Correlation |
| GGA3 | -0.2087 | 0.000381 | Other Negative Correlation |
| CCDC32 | -0.20873 | 0.00038 | Other Negative Correlation |
| CPT2 | -0.20887 | 0.000376 | Other Negative Correlation |
| HSH2D | -0.20891 | 0.000375 | Other Negative Correlation |
| MDH1 | -0.20892 | 0.000375 | Other Negative Correlation |
| RIPK3 | -0.20914 | 0.00037 | Other Negative Correlation |
| H2AC8 | -0.20919 | 0.000368 | Other Negative Correlation |
| ANKRD13A | -0.2093 | 0.000366 | Other Negative Correlation |
| NSMAF | -0.20934 | 0.000365 | Other Negative Correlation |
| FAM91A1 | -0.20937 | 0.000364 | Other Negative Correlation |
| KLK13 | -0.20946 | 0.000362 | Other Negative Correlation |
| SNX27 | -0.2095 | 0.000361 | Other Negative Correlation |
| NT5C2 | -0.20952 | 0.00036 | Other Negative Correlation |
| PRDM4 | -0.20954 | 0.00036 | Other Negative Correlation |
| TMEM64 | -0.20964 | 0.000358 | Other Negative Correlation |
| CHST7 | -0.20967 | 0.000357 | Other Negative Correlation |
| ZBTB8OS | -0.2098 | 0.000354 | Other Negative Correlation |
| RERE | -0.20982 | 0.000353 | Other Negative Correlation |
| PHF3 | -0.20993 | 0.000351 | Other Negative Correlation |
| GFM1 | -0.20998 | 0.00035 | Other Negative Correlation |
| TSC22D2 | -0.21012 | 0.000346 | Other Negative Correlation |

| | | | |
|---|---|---|---|
| SLC49A4 | -0.21026 | 0.000343 | Other Negative Correlation |
| PRR5L | -0.2103 | 0.000342 | Other Negative Correlation |
| WDR82 | -0.2103 | 0.000342 | Other Negative Correlation |
| RAP1GAP2 | -0.21032 | 0.000342 | Other Negative Correlation |
| SERPINB9 | -0.21043 | 0.000339 | Other Negative Correlation |
| SS18L1 | -0.2105 | 0.000338 | Other Negative Correlation |
| MRPS12 | -0.21054 | 0.000337 | Other Negative Correlation |
| LRRK1 | -0.21057 | 0.000336 | Other Negative Correlation |
| SMDT1 | -0.21061 | 0.000335 | Other Negative Correlation |
| LARP4 | -0.2108 | 0.000331 | Other Negative Correlation |
| GGACT | -0.2108 | 0.000331 | Other Negative Correlation |
| TMEM80 | -0.21101 | 0.000326 | Other Negative Correlation |
| AFMID | -0.21103 | 0.000326 | Other Negative Correlation |
| VAMP4 | -0.21103 | 0.000326 | Other Negative Correlation |
| ZCRB1 | -0.2111 | 0.000324 | Other Negative Correlation |
| ANKRD50 | -0.2111 | 0.000324 | Other Negative Correlation |
| EHHADH | -0.21115 | 0.000323 | Other Negative Correlation |
| NPC1 | -0.21116 | 0.000323 | Other Negative Correlation |
| IRF2BPL | -0.21119 | 0.000322 | Other Negative Correlation |
| UBE2A | -0.21127 | 0.000321 | Other Negative Correlation |
| RNF128 | -0.21138 | 0.000318 | Other Negative Correlation |
| DENND4C | -0.21139 | 0.000318 | Other Negative Correlation |
| KHNYN | -0.21149 | 0.000316 | Other Negative Correlation |
| PPP6C | -0.2116 | 0.000314 | Other Negative Correlation |
| RABL6 | -0.21164 | 0.000313 | Other Negative Correlation |
| OSBPL2 | -0.21179 | 0.00031 | Other Negative Correlation |
| CSRNP3 | -0.21184 | 0.000309 | Other Negative Correlation |
| COLGALT1 | -0.21194 | 0.000307 | Other Negative Correlation |
| CRBN | -0.21195 | 0.000306 | Other Negative Correlation |
| ZNF827 | -0.21202 | 0.000305 | Other Negative Correlation |
| TRAPPC8 | -0.21206 | 0.000304 | Other Negative Correlation |
| CASP10 | -0.21208 | 0.000304 | Other Negative Correlation |
| KLHL4 | -0.21215 | 0.000302 | Other Negative Correlation |
| TTC23 | -0.21217 | 0.000302 | Other Negative Correlation |
| STXBP2 | -0.21218 | 0.000302 | Other Negative Correlation |
| DROSHA | -0.21219 | 0.000301 | Other Negative Correlation |
| GDE1 | -0.21223 | 0.000301 | Other Negative Correlation |
| BAP1 | -0.21225 | 0.0003 | Other Negative Correlation |
| PARP8 | -0.21228 | 0.0003 | Other Negative Correlation |
| GPR39 | -0.21229 | 0.000299 | Other Negative Correlation |
| CPPED1 | -0.21248 | 0.000296 | Other Negative Correlation |
| BRAT1 | -0.21252 | 0.000295 | Other Negative Correlation |
| ZNHIT1 | -0.21252 | 0.000295 | Other Negative Correlation |
| CUL2 | -0.21255 | 0.000294 | Other Negative Correlation |
| COMMD5 | -0.21256 | 0.000294 | Other Negative Correlation |
| SPIRE1 | -0.21257 | 0.000294 | Other Negative Correlation |

| | | | |
|---|---|---|---|
| SGCB | -0.21257 | 0.000294 | Other Negative Correlation |
| VSIR | -0.21259 | 0.000293 | Other Negative Correlation |
| BPGM | -0.21261 | 0.000293 | Other Negative Correlation |
| PHRF1 | -0.21262 | 0.000293 | Other Negative Correlation |
| MEF2A | -0.21267 | 0.000292 | Other Negative Correlation |
| FOCAD | -0.21267 | 0.000292 | Other Negative Correlation |
| EHD3 | -0.21288 | 0.000288 | Other Negative Correlation |
| PCDHGA10 | -0.21288 | 0.000288 | Other Negative Correlation |
| HFE | -0.21291 | 0.000287 | Other Negative Correlation |
| PAPOLG | -0.21293 | 0.000287 | Other Negative Correlation |
| RCC1L | -0.21295 | 0.000286 | Other Negative Correlation |
| GRHL1 | -0.21298 | 0.000286 | Other Negative Correlation |
| SLC66A2 | -0.21313 | 0.000283 | Other Negative Correlation |
| CPD | -0.21315 | 0.000283 | Other Negative Correlation |
| USO1 | -0.2132 | 0.000282 | Other Negative Correlation |
| EXOC3 | -0.21335 | 0.000279 | Other Negative Correlation |
| GSAP | -0.21337 | 0.000279 | Other Negative Correlation |
| PITX2 | -0.21347 | 0.000277 | Other Negative Correlation |
| SF3B5 | -0.21359 | 0.000274 | Other Negative Correlation |
| CLCN3 | -0.21364 | 0.000273 | Other Negative Correlation |
| MED13 | -0.21371 | 0.000272 | Other Negative Correlation |
| H2AJ | -0.21373 | 0.000272 | Other Negative Correlation |
| CDK16 | -0.21374 | 0.000272 | Other Negative Correlation |
| C1orf53 | -0.21378 | 0.000271 | Other Negative Correlation |
| TMEM115 | -0.2138 | 0.00027 | Other Negative Correlation |
| ANO1 | -0.21386 | 0.000269 | Other Negative Correlation |
| TET2 | -0.2139 | 0.000269 | Other Negative Correlation |
| RPL3L | -0.21398 | 0.000267 | Other Negative Correlation |
| LRRC47 | -0.21405 | 0.000266 | Other Negative Correlation |
| TLCD5 | -0.21413 | 0.000265 | Other Negative Correlation |
| DYNC2I2 | -0.21418 | 0.000264 | Other Negative Correlation |
| PPIP5K2 | -0.21423 | 0.000263 | Other Negative Correlation |
| SGSM2 | -0.21426 | 0.000262 | Other Negative Correlation |
| FAM222B | -0.21429 | 0.000262 | Other Negative Correlation |
| CTSD | -0.21433 | 0.000261 | Other Negative Correlation |
| HOOK3 | -0.21436 | 0.00026 | Other Negative Correlation |
| ST6GALNAC1 | -0.21437 | 0.00026 | Other Negative Correlation |
| ANKRA2 | -0.21443 | 0.000259 | Other Negative Correlation |
| GGCT | -0.21445 | 0.000259 | Other Negative Correlation |
| SYCE1L | -0.2145 | 0.000258 | Other Negative Correlation |
| S1PR5 | -0.21453 | 0.000257 | Other Negative Correlation |
| TMEM44 | -0.21457 | 0.000257 | Other Negative Correlation |
| MARVELD3 | -0.21462 | 0.000256 | Other Negative Correlation |
| SCRN1 | -0.21476 | 0.000253 | Other Negative Correlation |
| MFSD3 | -0.21478 | 0.000253 | Other Negative Correlation |
| ZDHHC14 | -0.21478 | 0.000253 | Other Negative Correlation |

| | | | |
|---|---|---|---|
| INTS8 | -0.21479 | 0.000253 | Other Negative Correlation |
| ISCA1 | -0.21491 | 0.000251 | Other Negative Correlation |
| GPR157 | -0.21495 | 0.00025 | Other Negative Correlation |
| RAB11FIP2 | -0.21495 | 0.00025 | Other Negative Correlation |
| SLC30A6 | -0.21499 | 0.000249 | Other Negative Correlation |
| AUTS2 | -0.21511 | 0.000247 | Other Negative Correlation |
| DSTYK | -0.21515 | 0.000247 | Other Negative Correlation |
| TMEM63A | -0.21517 | 0.000246 | Other Negative Correlation |
| NFKB2 | -0.21518 | 0.000246 | Other Negative Correlation |
| NMI | -0.21528 | 0.000245 | Other Negative Correlation |
| TBC1D10B | -0.21544 | 0.000242 | Other Negative Correlation |
| DTX4 | -0.21547 | 0.000241 | Other Negative Correlation |
| SNX9 | -0.21548 | 0.000241 | Other Negative Correlation |
| KIAA0319L | -0.21552 | 0.000241 | Other Negative Correlation |
| MSL1 | -0.21558 | 0.00024 | Other Negative Correlation |
| CCDC88C | -0.21559 | 0.000239 | Other Negative Correlation |
| MBD5 | -0.21559 | 0.000239 | Other Negative Correlation |
| PLCD4 | -0.21561 | 0.000239 | Other Negative Correlation |
| R3HDM4 | -0.21563 | 0.000239 | Other Negative Correlation |
| ASCC2 | -0.21566 | 0.000238 | Other Negative Correlation |
| CEP126 | -0.21567 | 0.000238 | Other Negative Correlation |
| POLL | -0.21569 | 0.000238 | Other Negative Correlation |
| B4GALT3 | -0.21571 | 0.000237 | Other Negative Correlation |
| METTL15 | -0.2158 | 0.000236 | Other Negative Correlation |
| ERMP1 | -0.21581 | 0.000236 | Other Negative Correlation |
| TMEM259 | -0.21585 | 0.000235 | Other Negative Correlation |
| UBE2E2 | -0.21594 | 0.000234 | Other Negative Correlation |
| PRDM2 | -0.21599 | 0.000233 | Other Negative Correlation |
| PDE4DIP | -0.21608 | 0.000232 | Other Negative Correlation |
| CRKL | -0.21609 | 0.000231 | Other Negative Correlation |
| CARD11 | -0.21609 | 0.000231 | Other Negative Correlation |
| ARID1A | -0.21614 | 0.000231 | Other Negative Correlation |
| MTFR1 | -0.21614 | 0.000231 | Other Negative Correlation |
| MTF1 | -0.21615 | 0.00023 | Other Negative Correlation |
| PITPNA | -0.21618 | 0.00023 | Other Negative Correlation |
| EMC10 | -0.21629 | 0.000228 | Other Negative Correlation |
| KLF4 | -0.2163 | 0.000228 | Other Negative Correlation |
| CXCL16 | -0.21644 | 0.000226 | Other Negative Correlation |
| UBE2W | -0.21647 | 0.000226 | Other Negative Correlation |
| L3MBTL3 | -0.21651 | 0.000225 | Other Negative Correlation |
| PTPN12 | -0.21653 | 0.000225 | Other Negative Correlation |
| RELA | -0.21659 | 0.000224 | Other Negative Correlation |
| HEPHL1 | -0.21659 | 0.000224 | Other Negative Correlation |
| MFSD11 | -0.21664 | 0.000223 | Other Negative Correlation |
| EBI3 | -0.21665 | 0.000223 | Other Negative Correlation |
| TBC1D14 | -0.21672 | 0.000222 | Other Negative Correlation |

| | | | |
|---|---|---|---|
| COPA | -0.21674 | 0.000221 | Other Negative Correlation |
| OTULIN | -0.21678 | 0.000221 | Other Negative Correlation |
| GNA12 | -0.21678 | 0.000221 | Other Negative Correlation |
| SERPINF2 | -0.2168 | 0.00022 | Other Negative Correlation |
| RBM38 | -0.21691 | 0.000219 | Other Negative Correlation |
| ZNHIT2 | -0.21691 | 0.000219 | Other Negative Correlation |
| IDS | -0.21693 | 0.000218 | Other Negative Correlation |
| CKAP5 | -0.21698 | 0.000218 | Other Negative Correlation |
| KCNG1 | -0.21723 | 0.000214 | Other Negative Correlation |
| SH3D21 | -0.21744 | 0.000211 | Other Negative Correlation |
| COA8 | -0.21746 | 0.000211 | Other Negative Correlation |
| CSNK1G3 | -0.21752 | 0.00021 | Other Negative Correlation |
| RAB5IF | -0.21763 | 0.000208 | Other Negative Correlation |
| NPAS2 | -0.21764 | 0.000208 | Other Negative Correlation |
| USP9Y | -0.21766 | 0.000208 | Other Negative Correlation |
| MUC1 | -0.21779 | 0.000206 | Other Negative Correlation |
| NID1 | -0.21781 | 0.000206 | Other Negative Correlation |
| C1orf74 | -0.21782 | 0.000205 | Other Negative Correlation |
| LHPP | -0.21785 | 0.000205 | Other Negative Correlation |
| MRPS31 | -0.21786 | 0.000205 | Other Negative Correlation |
| BLVRA | -0.21792 | 0.000204 | Other Negative Correlation |
| CEP131 | -0.21794 | 0.000204 | Other Negative Correlation |
| HTATIP2 | -0.21794 | 0.000204 | Other Negative Correlation |
| UCKL1 | -0.21799 | 0.000203 | Other Negative Correlation |
| TDP2 | -0.2181 | 0.000201 | Other Negative Correlation |
| P3H4 | -0.21812 | 0.000201 | Other Negative Correlation |
| P2RY6 | -0.21815 | 0.000201 | Other Negative Correlation |
| TRNAU1AP | -0.21819 | 0.0002 | Other Negative Correlation |
| CAMSAP1 | -0.2182 | 0.0002 | Other Negative Correlation |
| KLHL21 | -0.21821 | 0.0002 | Other Negative Correlation |
| CSTF3 | -0.21832 | 0.000198 | Other Negative Correlation |
| TMEM242 | -0.21834 | 0.000198 | Other Negative Correlation |
| SPAST | -0.21845 | 0.000197 | Other Negative Correlation |
| THBS3 | -0.21845 | 0.000197 | Other Negative Correlation |
| SLC35E2B | -0.21849 | 0.000196 | Other Negative Correlation |
| TNFRSF10D | -0.21863 | 0.000194 | Other Negative Correlation |
| IL23A | -0.21869 | 0.000193 | Other Negative Correlation |
| ZNF428 | -0.21873 | 0.000193 | Other Negative Correlation |
| TMEM156 | -0.21883 | 0.000192 | Other Negative Correlation |
| HCAR2 | -0.21884 | 0.000192 | Other Negative Correlation |
| LRRC57 | -0.21886 | 0.000191 | Other Negative Correlation |
| RASSF5 | -0.21896 | 0.00019 | Other Negative Correlation |
| CLTC | -0.21898 | 0.00019 | Other Negative Correlation |
| TRIM56 | -0.21899 | 0.000189 | Other Negative Correlation |
| PGGT1B | -0.21903 | 0.000189 | Other Negative Correlation |
| ITGB5 | -0.21922 | 0.000186 | Other Negative Correlation |

| | | | |
|---|---|---|---|
| TIMM22 | -0.21925 | 0.000186 | Other Negative Correlation |
| PDE7A | -0.21934 | 0.000185 | Other Negative Correlation |
| SLC2A12 | -0.2194 | 0.000184 | Other Negative Correlation |
| CNOT11 | -0.21945 | 0.000184 | Other Negative Correlation |
| COX15 | -0.21945 | 0.000183 | Other Negative Correlation |
| BSDC1 | -0.21959 | 0.000182 | Other Negative Correlation |
| COA6 | -0.21972 | 0.00018 | Other Negative Correlation |
| LPIN3 | -0.21976 | 0.00018 | Other Negative Correlation |
| SPCS3 | -0.21994 | 0.000177 | Other Negative Correlation |
| ZDHHC7 | -0.22 | 0.000177 | Other Negative Correlation |
| SIPA1 | -0.22003 | 0.000176 | Other Negative Correlation |
| TM2D3 | -0.22004 | 0.000176 | Other Negative Correlation |
| MAML2 | -0.22005 | 0.000176 | Other Negative Correlation |
| UBXN7 | -0.22009 | 0.000176 | Other Negative Correlation |
| TMEM51 | -0.22013 | 0.000175 | Other Negative Correlation |
| PTK2 | -0.22034 | 0.000172 | Other Negative Correlation |
| USP6NL | -0.22035 | 0.000172 | Other Negative Correlation |
| IGSF8 | -0.2205 | 0.000171 | Other Negative Correlation |
| RCN1 | -0.22053 | 0.00017 | Other Negative Correlation |
| WDR59 | -0.22058 | 0.00017 | Other Negative Correlation |
| ELP1 | -0.22065 | 0.000169 | Other Negative Correlation |
| WBP11 | -0.22069 | 0.000168 | Other Negative Correlation |
| POLDIP2 | -0.22069 | 0.000168 | Other Negative Correlation |
| HDAC8 | -0.22071 | 0.000168 | Other Negative Correlation |
| NDUFB8 | -0.22073 | 0.000168 | Other Negative Correlation |
| FLYWCH2 | -0.22075 | 0.000168 | Other Negative Correlation |
| USP33 | -0.22076 | 0.000167 | Other Negative Correlation |
| IKZF2 | -0.22108 | 0.000164 | Other Negative Correlation |
| FAM200B | -0.22112 | 0.000163 | Other Negative Correlation |
| SYNRG | -0.22116 | 0.000163 | Other Negative Correlation |
| ADGRE2 | -0.22119 | 0.000163 | Other Negative Correlation |
| LINC00672 | -0.22119 | 0.000162 | Other Negative Correlation |
| FBLN2 | -0.22138 | 0.00016 | Other Negative Correlation |
| UBE2Q1 | -0.22139 | 0.00016 | Other Negative Correlation |
| TM2D2 | -0.22165 | 0.000157 | Other Negative Correlation |
| REV1 | -0.22166 | 0.000157 | Other Negative Correlation |
| C11orf68 | -0.22169 | 0.000157 | Other Negative Correlation |
| TXNRD3 | -0.22172 | 0.000157 | Other Negative Correlation |
| INTS12 | -0.22185 | 0.000155 | Other Negative Correlation |
| AGA | -0.22193 | 0.000154 | Other Negative Correlation |
| PROS1 | -0.22194 | 0.000154 | Other Negative Correlation |
| ITSN1 | -0.22194 | 0.000154 | Other Negative Correlation |
| SUPT6H | -0.22198 | 0.000154 | Other Negative Correlation |
| NAT14 | -0.22198 | 0.000154 | Other Negative Correlation |
| PKIB | -0.22208 | 0.000153 | Other Negative Correlation |
| GLRX | -0.22219 | 0.000151 | Other Negative Correlation |

| | | | |
|---|---|---|---|
| SPOCK1 | -0.22225 | 0.000151 | Other Negative Correlation |
| WWTR1 | -0.22227 | 0.000151 | Other Negative Correlation |
| SLC39A9 | -0.22233 | 0.00015 | Other Negative Correlation |
| STK39 | -0.22247 | 0.000149 | Other Negative Correlation |
| ACAD8 | -0.22249 | 0.000148 | Other Negative Correlation |
| NEXN | -0.22254 | 0.000148 | Other Negative Correlation |
| ARHGAP35 | -0.22258 | 0.000147 | Other Negative Correlation |
| OXCT2 | -0.22259 | 0.000147 | Other Negative Correlation |
| LRRC8E | -0.22262 | 0.000147 | Other Negative Correlation |
| PSD4 | -0.22264 | 0.000147 | Other Negative Correlation |
| ATF6 | -0.22268 | 0.000146 | Other Negative Correlation |
| SUCLG1 | -0.2227 | 0.000146 | Other Negative Correlation |
| IL34 | -0.22273 | 0.000146 | Other Negative Correlation |
| PEX2 | -0.22286 | 0.000145 | Other Negative Correlation |
| ARHGEF35 | -0.2229 | 0.000144 | Other Negative Correlation |
| UPK1B | -0.22298 | 0.000143 | Other Negative Correlation |
| CRCT1 | -0.223 | 0.000143 | Other Negative Correlation |
| AFAP1 | -0.22301 | 0.000143 | Other Negative Correlation |
| TMEM98 | -0.22301 | 0.000143 | Other Negative Correlation |
| E2F5 | -0.22305 | 0.000143 | Other Negative Correlation |
| POLR3A | -0.22318 | 0.000141 | Other Negative Correlation |
| LCN2 | -0.22321 | 0.000141 | Other Negative Correlation |
| TP73 | -0.22324 | 0.000141 | Other Negative Correlation |
| SNX33 | -0.22346 | 0.000139 | Other Negative Correlation |
| MFSD5 | -0.22346 | 0.000138 | Other Negative Correlation |
| ATP2C1 | -0.22348 | 0.000138 | Other Negative Correlation |
| SMPDL3A | -0.22354 | 0.000138 | Other Negative Correlation |
| SURF4 | -0.22363 | 0.000137 | Other Negative Correlation |
| CSNK1D | -0.22365 | 0.000137 | Other Negative Correlation |
| ILF2 | -0.22375 | 0.000136 | Other Negative Correlation |
| SPRR1A | -0.22378 | 0.000135 | Other Negative Correlation |
| ENSG00000283886 | -0.22382 | 0.000135 | Other Negative Correlation |
| FBXO38 | -0.22386 | 0.000135 | Other Negative Correlation |
| SP6 | -0.2239 | 0.000134 | Other Negative Correlation |
| ALKBH7 | -0.22399 | 0.000133 | Other Negative Correlation |
| PIGG | -0.22401 | 0.000133 | Other Negative Correlation |
| RP2 | -0.22407 | 0.000133 | Other Negative Correlation |
| BTG1 | -0.22408 | 0.000133 | Other Negative Correlation |
| SSH2 | -0.22408 | 0.000133 | Other Negative Correlation |
| AKR1B1 | -0.22409 | 0.000133 | Other Negative Correlation |
| SEPHS2 | -0.22409 | 0.000132 | Other Negative Correlation |
| BBS10 | -0.22409 | 0.000132 | Other Negative Correlation |
| KDELR2 | -0.2241 | 0.000132 | Other Negative Correlation |
| PMPCB | -0.22422 | 0.000131 | Other Negative Correlation |
| LPGAT1 | -0.22427 | 0.000131 | Other Negative Correlation |
| ARHGEF12 | -0.22435 | 0.00013 | Other Negative Correlation |

| | | | |
|---|---|---|---|
| ANAPC7 | -0.22448 | 0.000129 | Other Negative Correlation |
| CUX1 | -0.22463 | 0.000127 | Other Negative Correlation |
| CAT | -0.22472 | 0.000127 | Other Negative Correlation |
| SDHAF4 | -0.22487 | 0.000125 | Other Negative Correlation |
| ATP6AP1 | -0.22497 | 0.000124 | Other Negative Correlation |
| CEACAM1 | -0.22501 | 0.000124 | Other Negative Correlation |
| TNPO3 | -0.22514 | 0.000123 | Other Negative Correlation |
| RNF170 | -0.22514 | 0.000123 | Other Negative Correlation |
| IL31RA | -0.22518 | 0.000123 | Other Negative Correlation |
| RAPGEFL1 | -0.22527 | 0.000122 | Other Negative Correlation |
| SMARCC2 | -0.2253 | 0.000122 | Other Negative Correlation |
| GFOD2 | -0.2253 | 0.000122 | Other Negative Correlation |
| NOTCH2 | -0.22535 | 0.000121 | Other Negative Correlation |
| FRRS1 | -0.22536 | 0.000121 | Other Negative Correlation |
| RAB21 | -0.22549 | 0.00012 | Other Negative Correlation |
| RBM33 | -0.22552 | 0.00012 | Other Negative Correlation |
| ANAPC10 | -0.22558 | 0.000119 | Other Negative Correlation |
| MTCH1 | -0.22562 | 0.000119 | Other Negative Correlation |
| RNF8 | -0.22566 | 0.000118 | Other Negative Correlation |
| KRT75 | -0.22577 | 0.000118 | Other Negative Correlation |
| ZNF618 | -0.226 | 0.000116 | Other Negative Correlation |
| ZNF746 | -0.226 | 0.000116 | Other Negative Correlation |
| STX8 | -0.22601 | 0.000116 | Other Negative Correlation |
| DHX32 | -0.22621 | 0.000114 | Other Negative Correlation |
| MGAT1 | -0.22627 | 0.000113 | Other Negative Correlation |
| ARL3 | -0.22631 | 0.000113 | Other Negative Correlation |
| ZNHIT3 | -0.22632 | 0.000113 | Other Negative Correlation |
| OGFR | -0.22637 | 0.000113 | Other Negative Correlation |
| DOLK | -0.22641 | 0.000112 | Other Negative Correlation |
| PHIP | -0.22643 | 0.000112 | Other Negative Correlation |
| COMMD7 | -0.22647 | 0.000112 | Other Negative Correlation |
| SLC25A17 | -0.2265 | 0.000112 | Other Negative Correlation |
| CHMP7 | -0.22654 | 0.000111 | Other Negative Correlation |
| METTL23 | -0.22662 | 0.000111 | Other Negative Correlation |
| ATXN7L3B | -0.22662 | 0.000111 | Other Negative Correlation |
| VSTM2L | -0.22664 | 0.00011 | Other Negative Correlation |
| TRPC4AP | -0.22667 | 0.00011 | Other Negative Correlation |
| KRBA1 | -0.22673 | 0.00011 | Other Negative Correlation |
| PRICKLE2 | -0.22678 | 0.000109 | Other Negative Correlation |
| PGM2L1 | -0.22678 | 0.000109 | Other Negative Correlation |
| ACTR8 | -0.22697 | 0.000108 | Other Negative Correlation |
| HERC4 | -0.22703 | 0.000107 | Other Negative Correlation |
| CLDN12 | -0.22704 | 0.000107 | Other Negative Correlation |
| PDHA1 | -0.22706 | 0.000107 | Other Negative Correlation |
| AVL9 | -0.22708 | 0.000107 | Other Negative Correlation |
| CNP | -0.22722 | 0.000106 | Other Negative Correlation |

| | | | |
|---|---|---|---|
| TMEM86A | -0.22723 | 0.000106 | Other Negative Correlation |
| BBS12 | -0.22725 | 0.000106 | Other Negative Correlation |
| LY6G6C | -0.22727 | 0.000106 | Other Negative Correlation |
| GRB2 | -0.22741 | 0.000104 | Other Negative Correlation |
| PBX2 | -0.22743 | 0.000104 | Other Negative Correlation |
| HSPB11 | -0.22744 | 0.000104 | Other Negative Correlation |
| FRK | -0.22744 | 0.000104 | Other Negative Correlation |
| GCNT1 | -0.22744 | 0.000104 | Other Negative Correlation |
| ARL14EP | -0.22754 | 0.000104 | Other Negative Correlation |
| PLD1 | -0.22763 | 0.000103 | Other Negative Correlation |
| KIN | -0.22765 | 0.000103 | Other Negative Correlation |
| JHY | -0.2277 | 0.000102 | Other Negative Correlation |
| CNGA1 | -0.22771 | 0.000102 | Other Negative Correlation |
| VARS2 | -0.22776 | 0.000102 | Other Negative Correlation |
| HPS6 | -0.2278 | 0.000102 | Other Negative Correlation |
| CUEDC1 | -0.22781 | 0.000101 | Other Negative Correlation |
| MAGED2 | -0.22782 | 0.000101 | Other Negative Correlation |
| AREL1 | -0.22784 | 0.000101 | Other Negative Correlation |
| CARD14 | -0.22785 | 0.000101 | Other Negative Correlation |
| MDFIC | -0.22799 | 0.0001 | Other Negative Correlation |
| ANXA11 | -0.22807 | 9.96E-05 | Other Negative Correlation |
| PIP4K2B | -0.22808 | 9.96E-05 | Other Negative Correlation |
| DNAJC8 | -0.22808 | 9.95E-05 | Other Negative Correlation |
| RAB12 | -0.22809 | 9.94E-05 | Other Negative Correlation |
| SNX29 | -0.22812 | 9.93E-05 | Other Negative Correlation |
| NFS1 | -0.22816 | 9.9E-05 | Other Negative Correlation |
| SGPL1 | -0.22821 | 9.86E-05 | Other Negative Correlation |
| C10orf88 | -0.22829 | 9.8E-05 | Other Negative Correlation |
| SLC25A26 | -0.22831 | 9.79E-05 | Other Negative Correlation |
| SLC35A3 | -0.22841 | 9.72E-05 | Other Negative Correlation |
| SLC25A46 | -0.22857 | 9.61E-05 | Other Negative Correlation |
| UBR4 | -0.22861 | 9.58E-05 | Other Negative Correlation |
| ZHX3 | -0.22872 | 9.5E-05 | Other Negative Correlation |
| ANXA2R | -0.22872 | 9.5E-05 | Other Negative Correlation |
| SPG7 | -0.22879 | 9.46E-05 | Other Negative Correlation |
| PIP4P2 | -0.22881 | 9.44E-05 | Other Negative Correlation |
| KRT31 | -0.22886 | 9.4E-05 | Other Negative Correlation |
| UBE3B | -0.22887 | 9.4E-05 | Other Negative Correlation |
| ERGIC1 | -0.22888 | 9.39E-05 | Other Negative Correlation |
| EXD3 | -0.22895 | 9.34E-05 | Other Negative Correlation |
| TBC1D7 | -0.22896 | 9.34E-05 | Other Negative Correlation |
| DOCK9 | -0.22909 | 9.25E-05 | Other Negative Correlation |
| ANKRD65 | -0.22911 | 9.24E-05 | Other Negative Correlation |
| ZMYM3 | -0.22913 | 9.23E-05 | Other Negative Correlation |
| FBXL8 | -0.22915 | 9.21E-05 | Other Negative Correlation |
| ARHGEF16 | -0.22915 | 9.21E-05 | Other Negative Correlation |

| | | | |
|---|---|---|---|
| PDCL3 | -0.22928 | 9.13E-05 | Other Negative Correlation |
| BIVM | -0.22931 | 9.11E-05 | Other Negative Correlation |
| SF3A1 | -0.22949 | 8.98E-05 | Other Negative Correlation |
| LZTS2 | -0.22954 | 8.96E-05 | Other Negative Correlation |
| KLHL42 | -0.22955 | 8.95E-05 | Other Negative Correlation |
| OSBPL5 | -0.22956 | 8.94E-05 | Other Negative Correlation |
| TSG101 | -0.22959 | 8.92E-05 | Other Negative Correlation |
| LANCL1 | -0.22959 | 8.92E-05 | Other Negative Correlation |
| SETD3 | -0.22965 | 8.88E-05 | Other Negative Correlation |
| TCTN3 | -0.22971 | 8.85E-05 | Other Negative Correlation |
| SLC25A39 | -0.22975 | 8.82E-05 | Other Negative Correlation |
| SH3GLB2 | -0.22989 | 8.73E-05 | Other Negative Correlation |
| EIF2B2 | -0.22992 | 8.71E-05 | Other Negative Correlation |
| TAPBPL | -0.22999 | 8.66E-05 | Other Negative Correlation |
| PSG4 | -0.23004 | 8.63E-05 | Other Negative Correlation |
| TBK1 | -0.23009 | 8.6E-05 | Other Negative Correlation |
| IL36RN | -0.2301 | 8.59E-05 | Other Negative Correlation |
| MLXIP | -0.23014 | 8.57E-05 | Other Negative Correlation |
| HINT3 | -0.23019 | 8.54E-05 | Other Negative Correlation |
| AAR2 | -0.23029 | 8.48E-05 | Other Negative Correlation |
| MARCHF4 | -0.23032 | 8.46E-05 | Other Negative Correlation |
| TMEM203 | -0.23044 | 8.39E-05 | Other Negative Correlation |
| IDH3A | -0.23049 | 8.36E-05 | Other Negative Correlation |
| MED23 | -0.2305 | 8.35E-05 | Other Negative Correlation |
| LGALS7 | -0.23053 | 8.33E-05 | Other Negative Correlation |
| YAE1 | -0.23065 | 8.26E-05 | Other Negative Correlation |
| FGFR1OP2 | -0.23075 | 8.2E-05 | Other Negative Correlation |
| MIEN1 | -0.23098 | 8.06E-05 | Other Negative Correlation |
| OSBPL7 | -0.23117 | 7.95E-05 | Other Negative Correlation |
| CUL4A | -0.23123 | 7.92E-05 | Other Negative Correlation |
| COQ5 | -0.2313 | 7.88E-05 | Other Negative Correlation |
| H1-10 | -0.23137 | 7.84E-05 | Other Negative Correlation |
| MITD1 | -0.23147 | 7.78E-05 | Other Negative Correlation |
| TLE5 | -0.23151 | 7.76E-05 | Other Negative Correlation |
| BLNK | -0.23152 | 7.75E-05 | Other Negative Correlation |
| RAE1 | -0.23154 | 7.74E-05 | Other Negative Correlation |
| GDPD3 | -0.23154 | 7.74E-05 | Other Negative Correlation |
| USF2 | -0.23164 | 7.68E-05 | Other Negative Correlation |
| CUL3 | -0.23165 | 7.68E-05 | Other Negative Correlation |
| NEDD9 | -0.23166 | 7.67E-05 | Other Negative Correlation |
| TFPI | -0.23167 | 7.67E-05 | Other Negative Correlation |
| IFNAR2 | -0.23167 | 7.66E-05 | Other Negative Correlation |
| KLHL20 | -0.23169 | 7.65E-05 | Other Negative Correlation |
| PRXL2C | -0.2319 | 7.53E-05 | Other Negative Correlation |
| PER1 | -0.23193 | 7.52E-05 | Other Negative Correlation |
| NDUFA13 | -0.23195 | 7.51E-05 | Other Negative Correlation |

| | | | |
|---|---|---|---|
| BMP2 | -0.23195 | 7.51E-05 | Other Negative Correlation |
| SMURF1 | -0.232 | 7.48E-05 | Other Negative Correlation |
| RPS6KB1 | -0.2321 | 7.43E-05 | Other Negative Correlation |
| ANKRD29 | -0.23219 | 7.38E-05 | Other Negative Correlation |
| HACD2 | -0.23225 | 7.35E-05 | Other Negative Correlation |
| NTF4 | -0.23231 | 7.31E-05 | Other Negative Correlation |
| MYO5A | -0.23254 | 7.19E-05 | Other Negative Correlation |
| BTN3A2 | -0.23257 | 7.18E-05 | Other Negative Correlation |
| GRHL3 | -0.23261 | 7.15E-05 | Other Negative Correlation |
| PHYH | -0.23264 | 7.14E-05 | Other Negative Correlation |
| TNIP2 | -0.23277 | 7.07E-05 | Other Negative Correlation |
| PLEKHM1 | -0.23278 | 7.06E-05 | Other Negative Correlation |
| PHKA2 | -0.23281 | 7.05E-05 | Other Negative Correlation |
| RNF181 | -0.23296 | 6.97E-05 | Other Negative Correlation |
| KRT10 | -0.23298 | 6.96E-05 | Other Negative Correlation |
| STAT3 | -0.233 | 6.95E-05 | Other Negative Correlation |
| RIOK2 | -0.23321 | 6.84E-05 | Other Negative Correlation |
| FITM2 | -0.23349 | 6.7E-05 | Other Negative Correlation |
| LRRC58 | -0.23351 | 6.69E-05 | Other Negative Correlation |
| SSPN | -0.23357 | 6.66E-05 | Other Negative Correlation |
| PDPK1 | -0.23359 | 6.65E-05 | Other Negative Correlation |
| LMBRD1 | -0.2338 | 6.55E-05 | Other Negative Correlation |
| CYB561D2 | -0.23381 | 6.55E-05 | Other Negative Correlation |
| VPS25 | -0.23386 | 6.52E-05 | Other Negative Correlation |
| SPRR2E | -0.23397 | 6.47E-05 | Other Negative Correlation |
| FAM174C | -0.23402 | 6.44E-05 | Other Negative Correlation |
| PSMC6 | -0.23404 | 6.44E-05 | Other Negative Correlation |
| GPALPP1 | -0.23431 | 6.3E-05 | Other Negative Correlation |
| ACOX1 | -0.2345 | 6.22E-05 | Other Negative Correlation |
| SCAF8 | -0.23455 | 6.2E-05 | Other Negative Correlation |
| VPS37A | -0.23458 | 6.18E-05 | Other Negative Correlation |
| MSRA | -0.2346 | 6.17E-05 | Other Negative Correlation |
| ST3GAL4 | -0.23462 | 6.16E-05 | Other Negative Correlation |
| ATF5 | -0.23475 | 6.1E-05 | Other Negative Correlation |
| TMEM18 | -0.23476 | 6.1E-05 | Other Negative Correlation |
| CERT1 | -0.23484 | 6.06E-05 | Other Negative Correlation |
| GATD3 | -0.23494 | 6.02E-05 | Other Negative Correlation |
| RTP4 | -0.2351 | 5.95E-05 | Other Negative Correlation |
| MON2 | -0.23515 | 5.92E-05 | Other Negative Correlation |
| DNAJC16 | -0.23522 | 5.89E-05 | Other Negative Correlation |
| SNX13 | -0.23523 | 5.89E-05 | Other Negative Correlation |
| PARP4 | -0.23543 | 5.8E-05 | Other Negative Correlation |
| PHF23 | -0.23547 | 5.78E-05 | Other Negative Correlation |
| HS2ST1 | -0.23554 | 5.75E-05 | Other Negative Correlation |
| TTC17 | -0.23559 | 5.73E-05 | Other Negative Correlation |
| CHPF2 | -0.2356 | 5.73E-05 | Other Negative Correlation |

| | | | |
|---|---|---|---|
| BRWD3 | -0.2357 | 5.69E-05 | Other Negative Correlation |
| NAT1 | -0.23575 | 5.67E-05 | Other Negative Correlation |
| MYL2 | -0.23578 | 5.65E-05 | Other Negative Correlation |
| LARP4B | -0.2358 | 5.65E-05 | Other Negative Correlation |
| KIF13B | -0.23581 | 5.64E-05 | Other Negative Correlation |
| PPTC7 | -0.23585 | 5.62E-05 | Other Negative Correlation |
| SH3YL1 | -0.23588 | 5.61E-05 | Other Negative Correlation |
| RALGAPA2 | -0.23593 | 5.59E-05 | Other Negative Correlation |
| MYO18A | -0.23608 | 5.53E-05 | Other Negative Correlation |
| BNIP2 | -0.23616 | 5.5E-05 | Other Negative Correlation |
| UBE2G1 | -0.23617 | 5.49E-05 | Other Negative Correlation |
| ANKIB1 | -0.23625 | 5.46E-05 | Other Negative Correlation |
| HIVEP2 | -0.23626 | 5.45E-05 | Other Negative Correlation |
| MRTFB | -0.23629 | 5.44E-05 | Other Negative Correlation |
| MAD2L1BP | -0.23631 | 5.43E-05 | Other Negative Correlation |
| TLE1 | -0.23636 | 5.41E-05 | Other Negative Correlation |
| ICE1 | -0.23639 | 5.4E-05 | Other Negative Correlation |
| TMEM230 | -0.23641 | 5.39E-05 | Other Negative Correlation |
| BCAP29 | -0.23652 | 5.35E-05 | Other Negative Correlation |
| POLG | -0.23666 | 5.29E-05 | Other Negative Correlation |
| SECISBP2L | -0.23669 | 5.28E-05 | Other Negative Correlation |
| ZPR1 | -0.2367 | 5.28E-05 | Other Negative Correlation |
| ODF3B | -0.23671 | 5.27E-05 | Other Negative Correlation |
| BRD3 | -0.23675 | 5.26E-05 | Other Negative Correlation |
| DHX57 | -0.23675 | 5.26E-05 | Other Negative Correlation |
| GTF3C1 | -0.23688 | 5.21E-05 | Other Negative Correlation |
| PORCN | -0.23689 | 5.2E-05 | Other Negative Correlation |
| SIRT5 | -0.23689 | 5.2E-05 | Other Negative Correlation |
| NCSTN | -0.23696 | 5.17E-05 | Other Negative Correlation |
| PDHB | -0.23719 | 5.09E-05 | Other Negative Correlation |
| ACTR10 | -0.23729 | 5.05E-05 | Other Negative Correlation |
| RIN3 | -0.23735 | 5.03E-05 | Other Negative Correlation |
| CAVIN2 | -0.23743 | 4.99E-05 | Other Negative Correlation |
| ACAP2 | -0.23758 | 4.94E-05 | Other Negative Correlation |
| ODF2L | -0.23779 | 4.86E-05 | Other Negative Correlation |
| UBR1 | -0.23801 | 4.78E-05 | Other Negative Correlation |
| ARHGAP23 | -0.2381 | 4.75E-05 | Other Negative Correlation |
| HGSNAT | -0.23818 | 4.72E-05 | Other Negative Correlation |
| KCTD3 | -0.23824 | 4.7E-05 | Other Negative Correlation |
| ELAPOR1 | -0.23833 | 4.67E-05 | Other Negative Correlation |
| ANAPC13 | -0.23843 | 4.63E-05 | Other Negative Correlation |
| ATG4A | -0.23856 | 4.59E-05 | Other Negative Correlation |
| MFAP2 | -0.23859 | 4.58E-05 | Other Negative Correlation |
| EP400 | -0.23863 | 4.56E-05 | Other Negative Correlation |
| RABGAP1 | -0.23866 | 4.55E-05 | Other Negative Correlation |
| LRMDA | -0.23867 | 4.55E-05 | Other Negative Correlation |

| | | | |
|---|---|---|---|
| RBMS3 | -0.23868 | 4.55E-05 | Other Negative Correlation |
| PIR | -0.23872 | 4.53E-05 | Other Negative Correlation |
| SRRD | -0.23877 | 4.52E-05 | Other Negative Correlation |
| GPAT4 | -0.23881 | 4.5E-05 | Other Negative Correlation |
| SCAMP1 | -0.23886 | 4.48E-05 | Other Negative Correlation |
| DOCK6 | -0.23898 | 4.44E-05 | Other Negative Correlation |
| A4GALT | -0.23899 | 4.44E-05 | Other Negative Correlation |
| PLAA | -0.239 | 4.44E-05 | Other Negative Correlation |
| GOLGB1 | -0.23913 | 4.39E-05 | Other Negative Correlation |
| HPS1 | -0.23934 | 4.32E-05 | Other Negative Correlation |
| RAB34 | -0.2394 | 4.3E-05 | Other Negative Correlation |
| SMG7 | -0.23946 | 4.28E-05 | Other Negative Correlation |
| XPNPEP1 | -0.23953 | 4.26E-05 | Other Negative Correlation |
| TMEM102 | -0.23959 | 4.24E-05 | Other Negative Correlation |
| TKFC | -0.23967 | 4.22E-05 | Other Negative Correlation |
| ZER1 | -0.23973 | 4.2E-05 | Other Negative Correlation |
| KIF19 | -0.23976 | 4.19E-05 | Other Negative Correlation |
| TRIQK | -0.23977 | 4.18E-05 | Other Negative Correlation |
| CYLD | -0.2398 | 4.18E-05 | Other Negative Correlation |
| C2orf42 | -0.23984 | 4.16E-05 | Other Negative Correlation |
| TP53 | -0.24 | 4.11E-05 | Other Negative Correlation |
| MCPH1 | -0.24001 | 4.11E-05 | Other Negative Correlation |
| NELFE | -0.24002 | 4.1E-05 | Other Negative Correlation |
| AMFR | -0.24004 | 4.1E-05 | Other Negative Correlation |
| SMAD2 | -0.24008 | 4.09E-05 | Other Negative Correlation |
| CCND3 | -0.24013 | 4.07E-05 | Other Negative Correlation |
| DPP3 | -0.24016 | 4.06E-05 | Other Negative Correlation |
| QARS1 | -0.24045 | 3.97E-05 | Other Negative Correlation |
| ASXL2 | -0.2405 | 3.96E-05 | Other Negative Correlation |
| KLC1 | -0.24065 | 3.91E-05 | Other Negative Correlation |
| RNF187 | -0.24074 | 3.89E-05 | Other Negative Correlation |
| SMU1 | -0.24077 | 3.88E-05 | Other Negative Correlation |
| XXYLT1 | -0.24096 | 3.82E-05 | Other Negative Correlation |
| FARS2 | -0.24097 | 3.82E-05 | Other Negative Correlation |
| RAB30 | -0.24098 | 3.82E-05 | Other Negative Correlation |
| TCEAL3 | -0.24106 | 3.79E-05 | Other Negative Correlation |
| SLFN12 | -0.24116 | 3.76E-05 | Other Negative Correlation |
| TOMM70 | -0.24116 | 3.76E-05 | Other Negative Correlation |
| AIP | -0.24118 | 3.76E-05 | Other Negative Correlation |
| TNKS2 | -0.24125 | 3.74E-05 | Other Negative Correlation |
| LDB1 | -0.24132 | 3.72E-05 | Other Negative Correlation |
| SSR1 | -0.24146 | 3.68E-05 | Other Negative Correlation |
| SPRR2D | -0.24146 | 3.68E-05 | Other Negative Correlation |
| SCAF1 | -0.24153 | 3.66E-05 | Other Negative Correlation |
| CAMKK2 | -0.24154 | 3.66E-05 | Other Negative Correlation |
| FTL | -0.2416 | 3.64E-05 | Other Negative Correlation |

| | | | |
|---|---|---|---|
| CD63 | -0.24165 | 3.63E-05 | Other Negative Correlation |
| ITGB2 | -0.24179 | 3.59E-05 | Other Negative Correlation |
| C19orf12 | -0.24179 | 3.59E-05 | Other Negative Correlation |
| OXLD1 | -0.24181 | 3.58E-05 | Other Negative Correlation |
| C8orf76 | -0.24183 | 3.58E-05 | Other Negative Correlation |
| SERTAD3 | -0.24197 | 3.54E-05 | Other Negative Correlation |
| MRPS7 | -0.24206 | 3.51E-05 | Other Negative Correlation |
| CTDP1 | -0.24206 | 3.51E-05 | Other Negative Correlation |
| DDT | -0.24208 | 3.51E-05 | Other Negative Correlation |
| METTL13 | -0.24221 | 3.47E-05 | Other Negative Correlation |
| GYG1 | -0.24222 | 3.47E-05 | Other Negative Correlation |
| MT-ATP6 | -0.24224 | 3.47E-05 | Other Negative Correlation |
| PIGC | -0.24224 | 3.46E-05 | Other Negative Correlation |
| HPCAL1 | -0.24233 | 3.44E-05 | Other Negative Correlation |
| SLC22A18 | -0.24233 | 3.44E-05 | Other Negative Correlation |
| RPP38 | -0.2424 | 3.42E-05 | Other Negative Correlation |
| SPAG1 | -0.24244 | 3.41E-05 | Other Negative Correlation |
| DDX27 | -0.24251 | 3.39E-05 | Other Negative Correlation |
| WWC1 | -0.24252 | 3.39E-05 | Other Negative Correlation |
| GPSM3 | -0.24255 | 3.38E-05 | Other Negative Correlation |
| FAM89B | -0.24259 | 3.37E-05 | Other Negative Correlation |
| BCKDHB | -0.24268 | 3.35E-05 | Other Negative Correlation |
| MYO9B | -0.24276 | 3.33E-05 | Other Negative Correlation |
| SAR1A | -0.24296 | 3.28E-05 | Other Negative Correlation |
| DCAF8 | -0.24306 | 3.25E-05 | Other Negative Correlation |
| RFX5 | -0.24308 | 3.25E-05 | Other Negative Correlation |
| FANCF | -0.24314 | 3.23E-05 | Other Negative Correlation |
| MMP7 | -0.24323 | 3.21E-05 | Other Negative Correlation |
| TBC1D5 | -0.24342 | 3.16E-05 | Other Negative Correlation |
| SPACA9 | -0.24361 | 3.12E-05 | Other Negative Correlation |
| SLC41A3 | -0.24362 | 3.12E-05 | Other Negative Correlation |
| ZBED5 | -0.24371 | 3.09E-05 | Other Negative Correlation |
| WNK1 | -0.24375 | 3.08E-05 | Other Negative Correlation |
| SBNO1 | -0.24377 | 3.08E-05 | Other Negative Correlation |
| RICTOR | -0.2439 | 3.05E-05 | Other Negative Correlation |
| VANGL2 | -0.24392 | 3.04E-05 | Other Negative Correlation |
| FTO | -0.24399 | 3.03E-05 | Other Negative Correlation |
| GLYR1 | -0.24403 | 3.02E-05 | Other Negative Correlation |
| DPP8 | -0.24408 | 3.01E-05 | Other Negative Correlation |
| UEVLD | -0.24414 | 2.99E-05 | Other Negative Correlation |
| HMGXB3 | -0.24415 | 2.99E-05 | Other Negative Correlation |
| ARHGEF7 | -0.24428 | 2.96E-05 | Other Negative Correlation |
| PRPF31 | -0.24428 | 2.96E-05 | Other Negative Correlation |
| UNC119 | -0.24428 | 2.96E-05 | Other Negative Correlation |
| MED31 | -0.24436 | 2.94E-05 | Other Negative Correlation |
| PTMS | -0.24438 | 2.94E-05 | Other Negative Correlation |

| | | | |
|---|---|---|---|
| PDP1 | -0.24438 | 2.94E-05 | Other Negative Correlation |
| KIF3B | -0.24448 | 2.92E-05 | Other Negative Correlation |
| PRICKLE3 | -0.24488 | 2.82E-05 | Other Negative Correlation |
| SLC4A1AP | -0.24503 | 2.79E-05 | Other Negative Correlation |
| TMEM126B | -0.24525 | 2.75E-05 | Other Negative Correlation |
| KLC2 | -0.24529 | 2.74E-05 | Other Negative Correlation |
| TBC1D2B | -0.24536 | 2.72E-05 | Other Negative Correlation |
| WDR44 | -0.2454 | 2.71E-05 | Other Negative Correlation |
| DIP2B | -0.24541 | 2.71E-05 | Other Negative Correlation |
| BAZ2A | -0.24548 | 2.7E-05 | Other Negative Correlation |
| SLC17A5 | -0.24548 | 2.7E-05 | Other Negative Correlation |
| MINDY1 | -0.24554 | 2.69E-05 | Other Negative Correlation |
| LRP12 | -0.24557 | 2.68E-05 | Other Negative Correlation |
| SLC12A4 | -0.24564 | 2.66E-05 | Other Negative Correlation |
| ATP6V0D1 | -0.24572 | 2.65E-05 | Other Negative Correlation |
| ZNF322 | -0.24577 | 2.64E-05 | Other Negative Correlation |
| PDZD2 | -0.24578 | 2.63E-05 | Other Negative Correlation |
| PSMD3 | -0.24582 | 2.63E-05 | Other Negative Correlation |
| GGPS1 | -0.24583 | 2.62E-05 | Other Negative Correlation |
| MTRES1 | -0.24603 | 2.58E-05 | Other Negative Correlation |
| SAV1 | -0.24616 | 2.56E-05 | Other Negative Correlation |
| QPCT | -0.24621 | 2.55E-05 | Other Negative Correlation |
| ZMPSTE24 | -0.24627 | 2.53E-05 | Other Negative Correlation |
| MPI | -0.24645 | 2.5E-05 | Other Negative Correlation |
| PSEN1 | -0.24649 | 2.49E-05 | Other Negative Correlation |
| PRPSAP2 | -0.2465 | 2.49E-05 | Other Negative Correlation |
| TRAPPC12 | -0.2465 | 2.49E-05 | Other Negative Correlation |
| SLC66A3 | -0.24656 | 2.48E-05 | Other Negative Correlation |
| NOSTRIN | -0.24659 | 2.47E-05 | Other Negative Correlation |
| UPK2 | -0.24665 | 2.46E-05 | Other Negative Correlation |
| IWS1 | -0.24669 | 2.45E-05 | Other Negative Correlation |
| OSBPL8 | -0.2467 | 2.45E-05 | Other Negative Correlation |
| WDR6 | -0.2469 | 2.41E-05 | Other Negative Correlation |
| NDUFS1 | -0.24692 | 2.41E-05 | Other Negative Correlation |
| MALT1 | -0.24695 | 2.4E-05 | Other Negative Correlation |
| PRPF6 | -0.24704 | 2.39E-05 | Other Negative Correlation |
| INTS14 | -0.24718 | 2.36E-05 | Other Negative Correlation |
| CRYBG2 | -0.24718 | 2.36E-05 | Other Negative Correlation |
| MEF2D | -0.2473 | 2.34E-05 | Other Negative Correlation |
| SAMD8 | -0.24741 | 2.32E-05 | Other Negative Correlation |
| PKN2 | -0.24741 | 2.32E-05 | Other Negative Correlation |
| PRRC1 | -0.24745 | 2.31E-05 | Other Negative Correlation |
| ATM | -0.24752 | 2.3E-05 | Other Negative Correlation |
| NAAA | -0.24753 | 2.3E-05 | Other Negative Correlation |
| HERC2 | -0.24761 | 2.28E-05 | Other Negative Correlation |
| USP22 | -0.24762 | 2.28E-05 | Other Negative Correlation |

| | | | |
|---|---|---|---|
| NUDT7 | -0.24778 | 2.25E-05 | Other Negative Correlation |
| COX14 | -0.24789 | 2.23E-05 | Other Negative Correlation |
| FAM210A | -0.24816 | 2.19E-05 | Other Negative Correlation |
| UNC119B | -0.24819 | 2.18E-05 | Other Negative Correlation |
| CYB5A | -0.24834 | 2.16E-05 | Other Negative Correlation |
| RUFY3 | -0.24854 | 2.12E-05 | Other Negative Correlation |
| TSPAN6 | -0.24854 | 2.12E-05 | Other Negative Correlation |
| PPP1R14A | -0.24858 | 2.12E-05 | Other Negative Correlation |
| CAMSAP3 | -0.24863 | 2.11E-05 | Other Negative Correlation |
| WSB1 | -0.24863 | 2.11E-05 | Other Negative Correlation |
| ITPKC | -0.24863 | 2.11E-05 | Other Negative Correlation |
| FNDC4 | -0.24865 | 2.1E-05 | Other Negative Correlation |
| COX8A | -0.24886 | 2.07E-05 | Other Negative Correlation |
| DNMBP | -0.24886 | 2.07E-05 | Other Negative Correlation |
| TTC19 | -0.24887 | 2.07E-05 | Other Negative Correlation |
| TNFRSF14 | -0.24897 | 2.05E-05 | Other Negative Correlation |
| FAM199X | -0.24897 | 2.05E-05 | Other Negative Correlation |
| TEX2 | -0.24901 | 2.04E-05 | Other Negative Correlation |
| RIPK1 | -0.24904 | 2.04E-05 | Other Negative Correlation |
| ALDOC | -0.24909 | 2.03E-05 | Other Negative Correlation |
| RNF167 | -0.2491 | 2.03E-05 | Other Negative Correlation |
| HMGXB4 | -0.24914 | 2.02E-05 | Other Negative Correlation |
| MRPL22 | -0.24915 | 2.02E-05 | Other Negative Correlation |
| HEXB | -0.24917 | 2.02E-05 | Other Negative Correlation |
| COL22A1 | -0.24922 | 2.01E-05 | Other Negative Correlation |
| MRPS9 | -0.2493 | 2E-05 | Other Negative Correlation |
| MORC2 | -0.24933 | 1.99E-05 | Other Negative Correlation |
| SPATA20 | -0.24937 | 1.99E-05 | Other Negative Correlation |
| METTL7A | -0.24937 | 1.99E-05 | Other Negative Correlation |
| GSDMA | -0.24942 | 1.98E-05 | Other Negative Correlation |
| EMC4 | -0.24947 | 1.97E-05 | Other Negative Correlation |
| FCHO2 | -0.2495 | 1.97E-05 | Other Negative Correlation |
| TRIP10 | -0.24964 | 1.95E-05 | Other Negative Correlation |
| CCDC120 | -0.24965 | 1.94E-05 | Other Negative Correlation |
| C1orf122 | -0.24972 | 1.93E-05 | Other Negative Correlation |
| PEX19 | -0.24974 | 1.93E-05 | Other Negative Correlation |
| RCAN3 | -0.24978 | 1.92E-05 | Other Negative Correlation |
| HACD4 | -0.24979 | 1.92E-05 | Other Negative Correlation |
| JMJD8 | -0.24994 | 1.9E-05 | Other Negative Correlation |
| TBCC | -0.24997 | 1.89E-05 | Other Negative Correlation |
| PHF6 | -0.25004 | 1.88E-05 | Other Negative Correlation |
| ARMC9 | -0.25004 | 1.88E-05 | Other Negative Correlation |
| HERPUD2 | -0.25017 | 1.86E-05 | Other Negative Correlation |
| BCAR1 | -0.25017 | 1.86E-05 | Other Negative Correlation |
| UBTF | -0.25019 | 1.86E-05 | Other Negative Correlation |
| NKAP | -0.25019 | 1.86E-05 | Other Negative Correlation |

| | | | |
|---|---|---|---|
| H2AC25 | -0.2502 | 1.86E-05 | Other Negative Correlation |
| PPP4C | -0.25028 | 1.85E-05 | Other Negative Correlation |
| ACACA | -0.25039 | 1.83E-05 | Other Negative Correlation |
| NDUFAF4 | -0.25043 | 1.83E-05 | Other Negative Correlation |
| H6PD | -0.25043 | 1.83E-05 | Other Negative Correlation |
| ESRP2 | -0.25046 | 1.82E-05 | Other Negative Correlation |
| ASPHD1 | -0.25049 | 1.82E-05 | Other Negative Correlation |
| FGF1 | -0.25054 | 1.81E-05 | Other Negative Correlation |
| OSCP1 | -0.25071 | 1.79E-05 | Other Negative Correlation |
| RBMX2 | -0.25071 | 1.79E-05 | Other Negative Correlation |
| JPT2 | -0.25087 | 1.76E-05 | Other Negative Correlation |
| PRKRIP1 | -0.25096 | 1.75E-05 | Other Negative Correlation |
| UBL3 | -0.25099 | 1.75E-05 | Other Negative Correlation |
| PCBD1 | -0.25102 | 1.74E-05 | Other Negative Correlation |
| DGKZ | -0.25114 | 1.73E-05 | Other Negative Correlation |
| SACM1L | -0.25137 | 1.69E-05 | Other Negative Correlation |
| R3HDM1 | -0.25139 | 1.69E-05 | Other Negative Correlation |
| CLIC4 | -0.25161 | 1.66E-05 | Other Negative Correlation |
| MST1R | -0.25184 | 1.63E-05 | Other Negative Correlation |
| FBXL17 | -0.25201 | 1.61E-05 | Other Negative Correlation |
| TRAF3 | -0.25209 | 1.6E-05 | Other Negative Correlation |
| TMEM147 | -0.25214 | 1.59E-05 | Other Negative Correlation |
| NGLY1 | -0.25216 | 1.59E-05 | Other Negative Correlation |
| KIF2A | -0.25217 | 1.59E-05 | Other Negative Correlation |
| SMAP2 | -0.25227 | 1.58E-05 | Other Negative Correlation |
| HELZ | -0.25232 | 1.57E-05 | Other Negative Correlation |
| TMC8 | -0.2524 | 1.56E-05 | Other Negative Correlation |
| CLUAP1 | -0.25254 | 1.54E-05 | Other Negative Correlation |
| MYO1C | -0.25262 | 1.53E-05 | Other Negative Correlation |
| VGLL3 | -0.25264 | 1.53E-05 | Other Negative Correlation |
| SMIM12 | -0.25269 | 1.52E-05 | Other Negative Correlation |
| DIXDC1 | -0.2527 | 1.52E-05 | Other Negative Correlation |
| NDUFAF1 | -0.25291 | 1.5E-05 | Other Negative Correlation |
| ALDH16A1 | -0.25291 | 1.5E-05 | Other Negative Correlation |
| ZFAND5 | -0.25299 | 1.49E-05 | Other Negative Correlation |
| PPP1R16A | -0.25304 | 1.48E-05 | Other Negative Correlation |
| MAB21L4 | -0.25313 | 1.47E-05 | Other Negative Correlation |
| AP2B1 | -0.25314 | 1.47E-05 | Other Negative Correlation |
| TMEM184A | -0.25315 | 1.47E-05 | Other Negative Correlation |
| BROX | -0.25315 | 1.47E-05 | Other Negative Correlation |
| KLF16 | -0.25327 | 1.46E-05 | Other Negative Correlation |
| RFC1 | -0.25328 | 1.45E-05 | Other Negative Correlation |
| TPGS1 | -0.25332 | 1.45E-05 | Other Negative Correlation |
| ZNF609 | -0.25336 | 1.44E-05 | Other Negative Correlation |
| STRN3 | -0.25339 | 1.44E-05 | Other Negative Correlation |
| RINL | -0.25341 | 1.44E-05 | Other Negative Correlation |

| | | | |
|---|---|---|---|
| RAB3GAP2 | -0.25347 | 1.43E-05 | Other Negative Correlation |
| TMED7 | -0.25352 | 1.43E-05 | Other Negative Correlation |
| NIN | -0.25361 | 1.42E-05 | Other Negative Correlation |
| BRWD1 | -0.25375 | 1.4E-05 | Other Negative Correlation |
| MTA3 | -0.25382 | 1.39E-05 | Other Negative Correlation |
| SULT1A1 | -0.25385 | 1.39E-05 | Other Negative Correlation |
| MROH1 | -0.25417 | 1.35E-05 | Other Negative Correlation |
| CEP68 | -0.25417 | 1.35E-05 | Other Negative Correlation |
| NUDT16L1 | -0.2543 | 1.34E-05 | Other Negative Correlation |
| ZDHHC13 | -0.25431 | 1.34E-05 | Other Negative Correlation |
| NRARP | -0.2544 | 1.33E-05 | Other Negative Correlation |
| CCM2 | -0.25454 | 1.31E-05 | Other Negative Correlation |
| MRPS30 | -0.25468 | 1.3E-05 | Other Negative Correlation |
| PSMB3 | -0.25475 | 1.29E-05 | Other Negative Correlation |
| STARD7 | -0.25482 | 1.28E-05 | Other Negative Correlation |
| DCTN6 | -0.25482 | 1.28E-05 | Other Negative Correlation |
| CNST | -0.25487 | 1.28E-05 | Other Negative Correlation |
| NTAN1 | -0.25489 | 1.28E-05 | Other Negative Correlation |
| MED24 | -0.2549 | 1.28E-05 | Other Negative Correlation |
| ZC3H4 | -0.25514 | 1.25E-05 | Other Negative Correlation |
| TMEM181 | -0.25517 | 1.25E-05 | Other Negative Correlation |
| APEH | -0.25518 | 1.25E-05 | Other Negative Correlation |
| CRIP2 | -0.25521 | 1.24E-05 | Other Negative Correlation |
| PSMF1 | -0.25524 | 1.24E-05 | Other Negative Correlation |
| ARHGEF3 | -0.25528 | 1.24E-05 | Other Negative Correlation |
| PSD3 | -0.2553 | 1.24E-05 | Other Negative Correlation |
| JKAMP | -0.25546 | 1.22E-05 | Other Negative Correlation |
| SH3BP4 | -0.25558 | 1.21E-05 | Other Negative Correlation |
| ADNP | -0.25563 | 1.2E-05 | Other Negative Correlation |
| PFDN4 | -0.25573 | 1.19E-05 | Other Negative Correlation |
| CALCOCO1 | -0.25574 | 1.19E-05 | Other Negative Correlation |
| KLHL8 | -0.25583 | 1.18E-05 | Other Negative Correlation |
| RBM41 | -0.25583 | 1.18E-05 | Other Negative Correlation |
| CPOX | -0.25588 | 1.18E-05 | Other Negative Correlation |
| KBTBD7 | -0.2559 | 1.18E-05 | Other Negative Correlation |
| TGFB1I1 | -0.25595 | 1.17E-05 | Other Negative Correlation |
| WDR47 | -0.25606 | 1.16E-05 | Other Negative Correlation |
| ELMOD2 | -0.25648 | 1.12E-05 | Other Negative Correlation |
| LYN | -0.2565 | 1.12E-05 | Other Negative Correlation |
| CCNT1 | -0.25653 | 1.12E-05 | Other Negative Correlation |
| RPS6KB2 | -0.25654 | 1.12E-05 | Other Negative Correlation |
| SUCO | -0.25659 | 1.11E-05 | Other Negative Correlation |
| TTC3 | -0.25664 | 1.11E-05 | Other Negative Correlation |
| AVPI1 | -0.25674 | 1.1E-05 | Other Negative Correlation |
| CCDC24 | -0.25707 | 1.07E-05 | Other Negative Correlation |
| TSKU | -0.25716 | 1.06E-05 | Other Negative Correlation |

| | | | |
|---|---|---|---|
| HMG20B | -0.25724 | 1.05E-05 | Other Negative Correlation |
| PBRM1 | -0.25731 | 1.05E-05 | Other Negative Correlation |
| SOX2 | -0.25734 | 1.05E-05 | Other Negative Correlation |
| GPATCH2 | -0.25734 | 1.05E-05 | Other Negative Correlation |
| GOLGA4 | -0.25748 | 1.03E-05 | Other Negative Correlation |
| IQCA1 | -0.25749 | 1.03E-05 | Other Negative Correlation |
| TATDN2 | -0.2575 | 1.03E-05 | Other Negative Correlation |
| SPTSSA | -0.25752 | 1.03E-05 | Other Negative Correlation |
| PNPLA3 | -0.25753 | 1.03E-05 | Other Negative Correlation |
| SERPINB6 | -0.25769 | 1.02E-05 | Other Negative Correlation |
| NCKAP5 | -0.25774 | 1.01E-05 | Other Negative Correlation |
| LMAN1 | -0.2578 | 1.01E-05 | Other Negative Correlation |
| MFSD1 | -0.25788 | 1E-05 | Other Negative Correlation |
| TMED8 | -0.25792 | 9.98E-06 | Other Negative Correlation |
| PIGX | -0.25796 | 9.94E-06 | Other Negative Correlation |
| MUC15 | -0.25799 | 9.92E-06 | Other Negative Correlation |
| APH1A | -0.25804 | 9.87E-06 | Other Negative Correlation |
| ABCA1 | -0.25808 | 9.85E-06 | Other Negative Correlation |
| SETX | -0.25811 | 9.82E-06 | Other Negative Correlation |
| RSBN1 | -0.25812 | 9.81E-06 | Other Negative Correlation |
| FPGT | -0.25814 | 9.8E-06 | Other Negative Correlation |
| ZNF516 | -0.25819 | 9.75E-06 | Other Negative Correlation |
| ZMYM4 | -0.25822 | 9.74E-06 | Other Negative Correlation |
| CHD6 | -0.25823 | 9.72E-06 | Other Negative Correlation |
| SLC16A4 | -0.25824 | 9.72E-06 | Other Negative Correlation |
| ESS2 | -0.25829 | 9.67E-06 | Other Negative Correlation |
| ABHD14A | -0.25831 | 9.66E-06 | Other Negative Correlation |
| RPS6KA3 | -0.25832 | 9.65E-06 | Other Negative Correlation |
| NAA80 | -0.25841 | 9.59E-06 | Other Negative Correlation |
| PLOD1 | -0.25841 | 9.58E-06 | Other Negative Correlation |
| RAET1E | -0.25848 | 9.53E-06 | Other Negative Correlation |
| ETV6 | -0.25855 | 9.47E-06 | Other Negative Correlation |
| FAM114A1 | -0.2586 | 9.43E-06 | Other Negative Correlation |
| TOB2 | -0.2586 | 9.43E-06 | Other Negative Correlation |
| RRAD | -0.2587 | 9.36E-06 | Other Negative Correlation |
| BTRC | -0.25873 | 9.33E-06 | Other Negative Correlation |
| RABAC1 | -0.25883 | 9.26E-06 | Other Negative Correlation |
| TOR1A | -0.25884 | 9.25E-06 | Other Negative Correlation |
| OFD1 | -0.259 | 9.13E-06 | Other Negative Correlation |
| TRAPPC4 | -0.25905 | 9.09E-06 | Other Negative Correlation |
| CHCHD10 | -0.25925 | 8.95E-06 | Other Negative Correlation |
| SRP68 | -0.25929 | 8.91E-06 | Other Negative Correlation |
| RBM27 | -0.25953 | 8.74E-06 | Other Negative Correlation |
| CYBA | -0.25953 | 8.74E-06 | Other Negative Correlation |
| MAPK10 | -0.25961 | 8.68E-06 | Other Negative Correlation |
| VPS39 | -0.25967 | 8.64E-06 | Other Negative Correlation |

| | | | |
|---|---|---|---|
| C1QTNF1 | -0.25967 | 8.63E-06 | Other Negative Correlation |
| LYPLAL1 | -0.25972 | 8.6E-06 | Other Negative Correlation |
| ZNF24 | -0.25984 | 8.51E-06 | Other Negative Correlation |
| GALNT10 | -0.25989 | 8.48E-06 | Other Negative Correlation |
| CSTF1 | -0.2599 | 8.47E-06 | Other Negative Correlation |
| PSMC2 | -0.2599 | 8.47E-06 | Other Negative Correlation |
| MEA1 | -0.26007 | 8.36E-06 | Other Negative Correlation |
| PCSK7 | -0.26007 | 8.35E-06 | Other Negative Correlation |
| GPR153 | -0.26018 | 8.28E-06 | Other Negative Correlation |
| MED13L | -0.26021 | 8.26E-06 | Other Negative Correlation |
| SLX4IP | -0.26024 | 8.24E-06 | Other Negative Correlation |
| RPUSD3 | -0.26033 | 8.18E-06 | Other Negative Correlation |
| RANBP9 | -0.26063 | 7.98E-06 | Other Negative Correlation |
| ZNFX1 | -0.26077 | 7.89E-06 | Other Negative Correlation |
| RNF130 | -0.26082 | 7.85E-06 | Other Negative Correlation |
| FUT2 | -0.26096 | 7.76E-06 | Other Negative Correlation |
| SMARCAD1 | -0.26098 | 7.75E-06 | Other Negative Correlation |
| RAB8B | -0.26105 | 7.7E-06 | Other Negative Correlation |
| WDR45B | -0.26114 | 7.65E-06 | Other Negative Correlation |
| GYS1 | -0.26124 | 7.58E-06 | Other Negative Correlation |
| KIFBP | -0.26125 | 7.58E-06 | Other Negative Correlation |
| LMBR1 | -0.26126 | 7.57E-06 | Other Negative Correlation |
| MTERF2 | -0.26128 | 7.56E-06 | Other Negative Correlation |
| ZNF629 | -0.2613 | 7.54E-06 | Other Negative Correlation |
| PAFAH1B2 | -0.26134 | 7.52E-06 | Other Negative Correlation |
| GRINA | -0.26135 | 7.52E-06 | Other Negative Correlation |
| STARD5 | -0.26177 | 7.26E-06 | Other Negative Correlation |
| NUMB | -0.26179 | 7.24E-06 | Other Negative Correlation |
| MPP7 | -0.26184 | 7.22E-06 | Other Negative Correlation |
| OSBPL11 | -0.26188 | 7.19E-06 | Other Negative Correlation |
| DCXR | -0.26192 | 7.17E-06 | Other Negative Correlation |
| CCNDBP1 | -0.26198 | 7.13E-06 | Other Negative Correlation |
| SRP19 | -0.26198 | 7.13E-06 | Other Negative Correlation |
| CFAP298 | -0.26199 | 7.12E-06 | Other Negative Correlation |
| CKMT1A | -0.262 | 7.12E-06 | Other Negative Correlation |
| PTPRK | -0.26207 | 7.08E-06 | Other Negative Correlation |
| METTL14 | -0.26208 | 7.07E-06 | Other Negative Correlation |
| LEMD1 | -0.26213 | 7.04E-06 | Other Negative Correlation |
| SERINC5 | -0.26214 | 7.04E-06 | Other Negative Correlation |
| MAPK14 | -0.26215 | 7.03E-06 | Other Negative Correlation |
| CSNK2A1 | -0.2622 | 7E-06 | Other Negative Correlation |
| P2RY2 | -0.26223 | 6.98E-06 | Other Negative Correlation |
| ATN1 | -0.26228 | 6.95E-06 | Other Negative Correlation |
| SKIL | -0.26231 | 6.94E-06 | Other Negative Correlation |
| LITAF | -0.26233 | 6.92E-06 | Other Negative Correlation |
| CD74 | -0.26236 | 6.91E-06 | Other Negative Correlation |

| | | | |
|---|---|---|---|
| GPRC5A | -0.26241 | 6.88E-06 | Other Negative Correlation |
| RHOF | -0.26265 | 6.74E-06 | Other Negative Correlation |
| STXBP5 | -0.26273 | 6.7E-06 | Other Negative Correlation |
| KCTD11 | -0.26275 | 6.69E-06 | Other Negative Correlation |
| TRERF1 | -0.26291 | 6.6E-06 | Other Negative Correlation |
| CCNE1 | -0.26316 | 6.46E-06 | Other Negative Correlation |
| MAPK1 | -0.26317 | 6.46E-06 | Other Negative Correlation |
| TMEM33 | -0.26337 | 6.35E-06 | Other Negative Correlation |
| P4HA2 | -0.26338 | 6.34E-06 | Other Negative Correlation |
| ERICH1 | -0.26352 | 6.27E-06 | Other Negative Correlation |
| TTYH1 | -0.26356 | 6.25E-06 | Other Negative Correlation |
| PPP2R2A | -0.26361 | 6.22E-06 | Other Negative Correlation |
| GSPT2 | -0.26364 | 6.21E-06 | Other Negative Correlation |
| ETNK1 | -0.26386 | 6.09E-06 | Other Negative Correlation |
| TP53BP1 | -0.26397 | 6.04E-06 | Other Negative Correlation |
| NF1 | -0.26397 | 6.04E-06 | Other Negative Correlation |
| KIDINS220 | -0.26398 | 6.03E-06 | Other Negative Correlation |
| AKAP11 | -0.26403 | 6E-06 | Other Negative Correlation |
| SLAMF9 | -0.26409 | 5.98E-06 | Other Negative Correlation |
| SLCO3A1 | -0.26412 | 5.96E-06 | Other Negative Correlation |
| GPR107 | -0.26421 | 5.92E-06 | Other Negative Correlation |
| C12orf73 | -0.26422 | 5.91E-06 | Other Negative Correlation |
| ATAD1 | -0.26422 | 5.91E-06 | Other Negative Correlation |
| FOXN1 | -0.26448 | 5.79E-06 | Other Negative Correlation |
| HTT | -0.26469 | 5.68E-06 | Other Negative Correlation |
| VNN1 | -0.26476 | 5.65E-06 | Other Negative Correlation |
| UQCC3 | -0.26481 | 5.62E-06 | Other Negative Correlation |
| TRIM37 | -0.26487 | 5.59E-06 | Other Negative Correlation |
| IL27RA | -0.26492 | 5.57E-06 | Other Negative Correlation |
| ARHGAP32 | -0.26496 | 5.56E-06 | Other Negative Correlation |
| TINCR | -0.26503 | 5.52E-06 | Other Negative Correlation |
| AFG3L2 | -0.26506 | 5.51E-06 | Other Negative Correlation |
| TRAFD1 | -0.26506 | 5.51E-06 | Other Negative Correlation |
| TRIM21 | -0.26556 | 5.28E-06 | Other Negative Correlation |
| CIAO1 | -0.26563 | 5.25E-06 | Other Negative Correlation |
| TEAD2 | -0.26564 | 5.24E-06 | Other Negative Correlation |
| MFAP3 | -0.26565 | 5.24E-06 | Other Negative Correlation |
| SLC35E3 | -0.26579 | 5.18E-06 | Other Negative Correlation |
| ABCA12 | -0.26591 | 5.13E-06 | Other Negative Correlation |
| ARHGAP1 | -0.26604 | 5.07E-06 | Other Negative Correlation |
| DAPK3 | -0.26621 | 5E-06 | Other Negative Correlation |
| CD164 | -0.26632 | 4.95E-06 | Other Negative Correlation |
| FBXO21 | -0.26641 | 4.91E-06 | Other Negative Correlation |
| ENTPD3 | -0.26645 | 4.9E-06 | Other Negative Correlation |
| NCLN | -0.26655 | 4.85E-06 | Other Negative Correlation |
| PLEKHB2 | -0.26657 | 4.84E-06 | Other Negative Correlation |

| | | | |
|---|---|---|---|
| ELAC2 | -0.26665 | 4.81E-06 | Other Negative Correlation |
| MANBA | -0.26665 | 4.81E-06 | Other Negative Correlation |
| MIF4GD | -0.2667 | 4.79E-06 | Other Negative Correlation |
| TP53RK | -0.26698 | 4.68E-06 | Other Negative Correlation |
| AFAP1L2 | -0.26703 | 4.66E-06 | Other Negative Correlation |
| STK16 | -0.2671 | 4.63E-06 | Other Negative Correlation |
| MARCO | -0.26712 | 4.63E-06 | Other Negative Correlation |
| ARCN1 | -0.26724 | 4.58E-06 | Other Negative Correlation |
| CASP3 | -0.26727 | 4.57E-06 | Other Negative Correlation |
| FGGY | -0.26731 | 4.55E-06 | Other Negative Correlation |
| ORAI1 | -0.26734 | 4.54E-06 | Other Negative Correlation |
| RTCB | -0.26736 | 4.53E-06 | Other Negative Correlation |
| FAM135A | -0.26739 | 4.52E-06 | Other Negative Correlation |
| PGRMC2 | -0.2674 | 4.51E-06 | Other Negative Correlation |
| MMGT1 | -0.26751 | 4.48E-06 | Other Negative Correlation |
| SLC10A3 | -0.26753 | 4.47E-06 | Other Negative Correlation |
| TMC6 | -0.26764 | 4.42E-06 | Other Negative Correlation |
| PRCP | -0.26766 | 4.42E-06 | Other Negative Correlation |
| SLC35B3 | -0.26769 | 4.4E-06 | Other Negative Correlation |
| EXTL3 | -0.26772 | 4.39E-06 | Other Negative Correlation |
| RIC8A | -0.2678 | 4.37E-06 | Other Negative Correlation |
| PELI1 | -0.26781 | 4.36E-06 | Other Negative Correlation |
| CAMK2N1 | -0.26781 | 4.36E-06 | Other Negative Correlation |
| FYCO1 | -0.26787 | 4.34E-06 | Other Negative Correlation |
| TMEM185B | -0.26804 | 4.28E-06 | Other Negative Correlation |
| APOL2 | -0.26814 | 4.24E-06 | Other Negative Correlation |
| MARVELD2 | -0.26816 | 4.23E-06 | Other Negative Correlation |
| SORL1 | -0.26821 | 4.21E-06 | Other Negative Correlation |
| TSHZ2 | -0.26844 | 4.13E-06 | Other Negative Correlation |
| DGAT2 | -0.26844 | 4.13E-06 | Other Negative Correlation |
| TENT4B | -0.26856 | 4.09E-06 | Other Negative Correlation |
| GAMT | -0.26865 | 4.06E-06 | Other Negative Correlation |
| WDR33 | -0.26868 | 4.05E-06 | Other Negative Correlation |
| DNAAF10 | -0.26895 | 3.96E-06 | Other Negative Correlation |
| ARAP2 | -0.26907 | 3.92E-06 | Other Negative Correlation |
| KPNA6 | -0.26918 | 3.88E-06 | Other Negative Correlation |
| NBEAL2 | -0.26923 | 3.86E-06 | Other Negative Correlation |
| VPS29 | -0.26939 | 3.81E-06 | Other Negative Correlation |
| TMC4 | -0.2695 | 3.77E-06 | Other Negative Correlation |
| RAB1B | -0.26964 | 3.73E-06 | Other Negative Correlation |
| LRATD1 | -0.26966 | 3.72E-06 | Other Negative Correlation |
| TMEM177 | -0.26968 | 3.72E-06 | Other Negative Correlation |
| VIRMA | -0.26968 | 3.71E-06 | Other Negative Correlation |
| NIPSNAP1 | -0.26972 | 3.7E-06 | Other Negative Correlation |
| RPRD1A | -0.26977 | 3.68E-06 | Other Negative Correlation |
| GEMIN7 | -0.27024 | 3.54E-06 | Other Negative Correlation |

| | | | |
|---|---|---|---|
| KRTCAP2 | -0.27044 | 3.48E-06 | Other Negative Correlation |
| PHACTR4 | -0.27055 | 3.45E-06 | Other Negative Correlation |
| F2R | -0.27067 | 3.41E-06 | Other Negative Correlation |
| WAC | -0.27078 | 3.38E-06 | Other Negative Correlation |
| SBSN | -0.27078 | 3.38E-06 | Other Negative Correlation |
| ABCC5 | -0.27081 | 3.37E-06 | Other Negative Correlation |
| GABARAPL2 | -0.27105 | 3.3E-06 | Other Negative Correlation |
| ATP6V1B2 | -0.27111 | 3.28E-06 | Other Negative Correlation |
| FBXW2 | -0.27134 | 3.22E-06 | Other Negative Correlation |
| CKMT1B | -0.27141 | 3.2E-06 | Other Negative Correlation |
| SCRN2 | -0.27155 | 3.16E-06 | Other Negative Correlation |
| SYNE2 | -0.27156 | 3.16E-06 | Other Negative Correlation |
| TMCC1 | -0.27164 | 3.14E-06 | Other Negative Correlation |
| ZNF677 | -0.27166 | 3.13E-06 | Other Negative Correlation |
| TNFSF15 | -0.27176 | 3.1E-06 | Other Negative Correlation |
| DHRS12 | -0.27177 | 3.1E-06 | Other Negative Correlation |
| LRP10 | -0.27179 | 3.1E-06 | Other Negative Correlation |
| PPP2R5C | -0.27193 | 3.06E-06 | Other Negative Correlation |
| ABHD2 | -0.27197 | 3.05E-06 | Other Negative Correlation |
| SLC44A2 | -0.27197 | 3.05E-06 | Other Negative Correlation |
| NECAP2 | -0.27213 | 3.01E-06 | Other Negative Correlation |
| DDB1 | -0.27219 | 2.99E-06 | Other Negative Correlation |
| ARHGAP27 | -0.27231 | 2.96E-06 | Other Negative Correlation |
| CDC42BPB | -0.27233 | 2.96E-06 | Other Negative Correlation |
| TBC1D22A | -0.27234 | 2.95E-06 | Other Negative Correlation |
| CSK | -0.27241 | 2.93E-06 | Other Negative Correlation |
| DYNC1LI2 | -0.27243 | 2.93E-06 | Other Negative Correlation |
| C1D | -0.27244 | 2.93E-06 | Other Negative Correlation |
| TMEM138 | -0.2725 | 2.91E-06 | Other Negative Correlation |
| GNL1 | -0.27251 | 2.91E-06 | Other Negative Correlation |
| MOB2 | -0.27253 | 2.9E-06 | Other Negative Correlation |
| POF1B | -0.27283 | 2.83E-06 | Other Negative Correlation |
| RBL2 | -0.27289 | 2.81E-06 | Other Negative Correlation |
| ITPR3 | -0.27291 | 2.81E-06 | Other Negative Correlation |
| MT-CYB | -0.27298 | 2.79E-06 | Other Negative Correlation |
| USP8 | -0.27301 | 2.78E-06 | Other Negative Correlation |
| OPA1 | -0.27321 | 2.74E-06 | Other Negative Correlation |
| DYRK4 | -0.27321 | 2.74E-06 | Other Negative Correlation |
| IL1R1 | -0.2733 | 2.71E-06 | Other Negative Correlation |
| NPEPPS | -0.27331 | 2.71E-06 | Other Negative Correlation |
| KLK8 | -0.27338 | 2.7E-06 | Other Negative Correlation |
| PIN1 | -0.27366 | 2.63E-06 | Other Negative Correlation |
| ABLIM3 | -0.27392 | 2.57E-06 | Other Negative Correlation |
| REPS1 | -0.27392 | 2.57E-06 | Other Negative Correlation |
| SKIC8 | -0.274 | 2.55E-06 | Other Negative Correlation |
| CHD1L | -0.27438 | 2.47E-06 | Other Negative Correlation |

| | | | |
|---|---|---|---|
| SCNM1 | -0.27462 | 2.42E-06 | Other Negative Correlation |
| EIF1AY | -0.27468 | 2.4E-06 | Other Negative Correlation |
| VPS13D | -0.27472 | 2.4E-06 | Other Negative Correlation |
| STARD13 | -0.27486 | 2.37E-06 | Other Negative Correlation |
| DTX2 | -0.2749 | 2.36E-06 | Other Negative Correlation |
| NCK2 | -0.275 | 2.34E-06 | Other Negative Correlation |
| ADAMTSL4 | -0.27521 | 2.3E-06 | Other Negative Correlation |
| HDHD3 | -0.27523 | 2.29E-06 | Other Negative Correlation |
| TTYH3 | -0.27525 | 2.29E-06 | Other Negative Correlation |
| PPP2R1B | -0.27528 | 2.28E-06 | Other Negative Correlation |
| EPGN | -0.27539 | 2.26E-06 | Other Negative Correlation |
| SNRPN | -0.27544 | 2.25E-06 | Other Negative Correlation |
| CRLF3 | -0.27545 | 2.25E-06 | Other Negative Correlation |
| MRPL43 | -0.27563 | 2.21E-06 | Other Negative Correlation |
| ATPAF2 | -0.27568 | 2.2E-06 | Other Negative Correlation |
| MT-ND5 | -0.27568 | 2.2E-06 | Other Negative Correlation |
| SLC15A4 | -0.27578 | 2.18E-06 | Other Negative Correlation |
| ABCG1 | -0.27583 | 2.17E-06 | Other Negative Correlation |
| IL17RC | -0.27625 | 2.1E-06 | Other Negative Correlation |
| CMIP | -0.27627 | 2.09E-06 | Other Negative Correlation |
| SLC12A2 | -0.27631 | 2.08E-06 | Other Negative Correlation |
| VGLL4 | -0.27632 | 2.08E-06 | Other Negative Correlation |
| LONP2 | -0.27638 | 2.07E-06 | Other Negative Correlation |
| TP53INP2 | -0.27651 | 2.05E-06 | Other Negative Correlation |
| ENTREP3 | -0.27672 | 2.01E-06 | Other Negative Correlation |
| HIBADH | -0.27673 | 2.01E-06 | Other Negative Correlation |
| KPNA4 | -0.27682 | 1.99E-06 | Other Negative Correlation |
| CCDC122 | -0.27698 | 1.96E-06 | Other Negative Correlation |
| GTF2E2 | -0.27699 | 1.96E-06 | Other Negative Correlation |
| TMEM30A | -0.27699 | 1.96E-06 | Other Negative Correlation |
| VPS53 | -0.27707 | 1.95E-06 | Other Negative Correlation |
| RNF6 | -0.2771 | 1.94E-06 | Other Negative Correlation |
| SEC24B | -0.27711 | 1.94E-06 | Other Negative Correlation |
| TMEM129 | -0.27716 | 1.93E-06 | Other Negative Correlation |
| EMC3 | -0.27728 | 1.91E-06 | Other Negative Correlation |
| POMGNT1 | -0.27731 | 1.91E-06 | Other Negative Correlation |
| SSC4D | -0.27738 | 1.9E-06 | Other Negative Correlation |
| BLTP1 | -0.2774 | 1.89E-06 | Other Negative Correlation |
| DLGAP4 | -0.27753 | 1.87E-06 | Other Negative Correlation |
| BOLA2-SMG1P6 | -0.27756 | 1.87E-06 | Other Negative Correlation |
| DNAJB6 | -0.27757 | 1.86E-06 | Other Negative Correlation |
| LDLRAP1 | -0.27763 | 1.85E-06 | Other Negative Correlation |
| NRIP1 | -0.27766 | 1.85E-06 | Other Negative Correlation |
| COQ4 | -0.27773 | 1.84E-06 | Other Negative Correlation |
| GPCPD1 | -0.27779 | 1.83E-06 | Other Negative Correlation |
| ZDHHC5 | -0.27805 | 1.79E-06 | Other Negative Correlation |

| | | | |
|---|---|---|---|
| NYNRIN | -0.27815 | 1.77E-06 | Other Negative Correlation |
| CCDC186 | -0.27824 | 1.76E-06 | Other Negative Correlation |
| UBE3C | -0.27835 | 1.74E-06 | Other Negative Correlation |
| ERBB2 | -0.27846 | 1.72E-06 | Other Negative Correlation |
| MRFAP1 | -0.27848 | 1.72E-06 | Other Negative Correlation |
| SHFL | -0.27849 | 1.72E-06 | Other Negative Correlation |
| MBD4 | -0.27875 | 1.68E-06 | Other Negative Correlation |
| SUSD6 | -0.27897 | 1.65E-06 | Other Negative Correlation |
| NUP42 | -0.27907 | 1.63E-06 | Other Negative Correlation |
| TMEM41A | -0.27912 | 1.62E-06 | Other Negative Correlation |
| RER1 | -0.27913 | 1.62E-06 | Other Negative Correlation |
| KIZ | -0.27916 | 1.62E-06 | Other Negative Correlation |
| CCDC68 | -0.27923 | 1.61E-06 | Other Negative Correlation |
| EFCAB14 | -0.27938 | 1.59E-06 | Other Negative Correlation |
| GLRX2 | -0.27944 | 1.58E-06 | Other Negative Correlation |
| LRRC42 | -0.27945 | 1.58E-06 | Other Negative Correlation |
| MED27 | -0.27948 | 1.57E-06 | Other Negative Correlation |
| NUDT9 | -0.27953 | 1.57E-06 | Other Negative Correlation |
| LPAR5 | -0.27958 | 1.56E-06 | Other Negative Correlation |
| GRHL2 | -0.27969 | 1.54E-06 | Other Negative Correlation |
| DBT | -0.2797 | 1.54E-06 | Other Negative Correlation |
| CDC37L1 | -0.27972 | 1.54E-06 | Other Negative Correlation |
| CTF1 | -0.27984 | 1.52E-06 | Other Negative Correlation |
| NEPRO | -0.27986 | 1.52E-06 | Other Negative Correlation |
| ASH1L | -0.2799 | 1.51E-06 | Other Negative Correlation |
| ATG10 | -0.28001 | 1.5E-06 | Other Negative Correlation |
| STARD3NL | -0.28004 | 1.5E-06 | Other Negative Correlation |
| DYRK1B | -0.28005 | 1.49E-06 | Other Negative Correlation |
| SLC22A23 | -0.28026 | 1.47E-06 | Other Negative Correlation |
| CRYBG3 | -0.2803 | 1.46E-06 | Other Negative Correlation |
| TAF2 | -0.28045 | 1.44E-06 | Other Negative Correlation |
| EXT2 | -0.28067 | 1.41E-06 | Other Negative Correlation |
| YLPM1 | -0.28068 | 1.41E-06 | Other Negative Correlation |
| C12orf29 | -0.28069 | 1.41E-06 | Other Negative Correlation |
| RPS6KA1 | -0.28074 | 1.4E-06 | Other Negative Correlation |
| AKAP6 | -0.28085 | 1.39E-06 | Other Negative Correlation |
| RITA1 | -0.28092 | 1.38E-06 | Other Negative Correlation |
| UBAP2 | -0.28103 | 1.37E-06 | Other Negative Correlation |
| RUFY1 | -0.28105 | 1.37E-06 | Other Negative Correlation |
| MGST3 | -0.28105 | 1.37E-06 | Other Negative Correlation |
| TUFT1 | -0.28107 | 1.36E-06 | Other Negative Correlation |
| DCBLD1 | -0.28112 | 1.36E-06 | Other Negative Correlation |
| PGLYRP4 | -0.28143 | 1.32E-06 | Other Negative Correlation |
| BAIAP2L1 | -0.2815 | 1.31E-06 | Other Negative Correlation |
| ADGRL2 | -0.28155 | 1.31E-06 | Other Negative Correlation |
| PLD3 | -0.28163 | 1.3E-06 | Other Negative Correlation |

| | | | |
|---|---|---|---|
| GAA | -0.2817 | 1.29E-06 | Other Negative Correlation |
| NAXE | -0.28181 | 1.28E-06 | Other Negative Correlation |
| CIAPIN1 | -0.282 | 1.25E-06 | Other Negative Correlation |
| MTRF1L | -0.28212 | 1.24E-06 | Other Negative Correlation |
| RAB22A | -0.28223 | 1.23E-06 | Other Negative Correlation |
| MAP3K11 | -0.28227 | 1.22E-06 | Other Negative Correlation |
| MAL2 | -0.28229 | 1.22E-06 | Other Negative Correlation |
| CORO1B | -0.28231 | 1.22E-06 | Other Negative Correlation |
| AKR1C2 | -0.28235 | 1.22E-06 | Other Negative Correlation |
| LAGE3 | -0.28259 | 1.19E-06 | Other Negative Correlation |
| RAB11FIP1 | -0.2826 | 1.19E-06 | Other Negative Correlation |
| DNAAF2 | -0.28262 | 1.19E-06 | Other Negative Correlation |
| NDUFA3 | -0.28263 | 1.18E-06 | Other Negative Correlation |
| YIPF3 | -0.28268 | 1.18E-06 | Other Negative Correlation |
| ATP7A | -0.28277 | 1.17E-06 | Other Negative Correlation |
| FAM3A | -0.28305 | 1.14E-06 | Other Negative Correlation |
| NCF1 | -0.28306 | 1.14E-06 | Other Negative Correlation |
| HDAC7 | -0.28315 | 1.13E-06 | Other Negative Correlation |
| PRKD2 | -0.28316 | 1.13E-06 | Other Negative Correlation |
| AEBP2 | -0.28316 | 1.13E-06 | Other Negative Correlation |
| FARP2 | -0.28319 | 1.13E-06 | Other Negative Correlation |
| EPN1 | -0.28326 | 1.12E-06 | Other Negative Correlation |
| SMIM5 | -0.28344 | 1.1E-06 | Other Negative Correlation |
| DEF8 | -0.28359 | 1.09E-06 | Other Negative Correlation |
| ATP6V1H | -0.28363 | 1.08E-06 | Other Negative Correlation |
| RPUSD1 | -0.28381 | 1.06E-06 | Other Negative Correlation |
| DHX40 | -0.28382 | 1.06E-06 | Other Negative Correlation |
| TAB3 | -0.28389 | 1.06E-06 | Other Negative Correlation |
| NEMF | -0.2842 | 1.03E-06 | Other Negative Correlation |
| RRN3 | -0.28435 | 1.01E-06 | Other Negative Correlation |
| NDUFA6 | -0.28439 | 1.01E-06 | Other Negative Correlation |
| BCR | -0.28441 | 1.01E-06 | Other Negative Correlation |
| GCC2 | -0.28454 | 9.96E-07 | Other Negative Correlation |
| TRIM13 | -0.28485 | 9.68E-07 | Other Negative Correlation |
| TMEM238 | -0.28491 | 9.63E-07 | Other Negative Correlation |
| TAF13 | -0.28493 | 9.61E-07 | Other Negative Correlation |
| AKR1B10 | -0.28504 | 9.52E-07 | Other Negative Correlation |
| NKAPD1 | -0.28527 | 9.32E-07 | Other Negative Correlation |
| PNLIPRP3 | -0.28533 | 9.27E-07 | Other Negative Correlation |
| PTPA | -0.28534 | 9.26E-07 | Other Negative Correlation |
| TBC1D9 | -0.28539 | 9.22E-07 | Other Negative Correlation |
| NT5DC1 | -0.28543 | 9.18E-07 | Other Negative Correlation |
| SNAP23 | -0.28548 | 9.14E-07 | Other Negative Correlation |
| STK38L | -0.2855 | 9.12E-07 | Other Negative Correlation |
| C15orf40 | -0.28553 | 9.1E-07 | Other Negative Correlation |
| TSPAN9 | -0.28568 | 8.97E-07 | Other Negative Correlation |

| | | | |
|---|---|---|---|
| ETAA1 | -0.2858 | 8.88E-07 | Other Negative Correlation |
| NSG1 | -0.28584 | 8.85E-07 | Other Negative Correlation |
| KLHDC10 | -0.28586 | 8.83E-07 | Other Negative Correlation |
| RPE | -0.28586 | 8.83E-07 | Other Negative Correlation |
| CRELD2 | -0.28602 | 8.7E-07 | Other Negative Correlation |
| SORBS2 | -0.28604 | 8.68E-07 | Other Negative Correlation |
| KREMEN2 | -0.28619 | 8.57E-07 | Other Negative Correlation |
| SCAMP2 | -0.2862 | 8.55E-07 | Other Negative Correlation |
| LIPH | -0.28634 | 8.45E-07 | Other Negative Correlation |
| L1TD1 | -0.28643 | 8.37E-07 | Other Negative Correlation |
| AP5B1 | -0.28669 | 8.18E-07 | Other Negative Correlation |
| PEX11B | -0.28672 | 8.15E-07 | Other Negative Correlation |
| EXOC2 | -0.28681 | 8.09E-07 | Other Negative Correlation |
| ARMT1 | -0.28693 | 8E-07 | Other Negative Correlation |
| CST6 | -0.28694 | 7.99E-07 | Other Negative Correlation |
| THUMPD3 | -0.28696 | 7.98E-07 | Other Negative Correlation |
| IREB2 | -0.28717 | 7.82E-07 | Other Negative Correlation |
| FKBP8 | -0.28718 | 7.82E-07 | Other Negative Correlation |
| CHD8 | -0.28727 | 7.75E-07 | Other Negative Correlation |
| MGAT4B | -0.28733 | 7.71E-07 | Other Negative Correlation |
| PCLO | -0.28741 | 7.66E-07 | Other Negative Correlation |
| C6orf89 | -0.28746 | 7.62E-07 | Other Negative Correlation |
| GJB4 | -0.28753 | 7.57E-07 | Other Negative Correlation |
| SLC16A2 | -0.28782 | 7.37E-07 | Other Negative Correlation |
| SH3TC1 | -0.28783 | 7.37E-07 | Other Negative Correlation |
| MRPS14 | -0.28788 | 7.33E-07 | Other Negative Correlation |
| CRK | -0.28799 | 7.26E-07 | Other Negative Correlation |
| IRF2 | -0.28801 | 7.25E-07 | Other Negative Correlation |
| TIMM8B | -0.28814 | 7.16E-07 | Other Negative Correlation |
| CBX6 | -0.28815 | 7.15E-07 | Other Negative Correlation |
| TMCC3 | -0.28825 | 7.08E-07 | Other Negative Correlation |
| POLR2I | -0.28829 | 7.06E-07 | Other Negative Correlation |
| UBFD1 | -0.28836 | 7.01E-07 | Other Negative Correlation |
| WDR11 | -0.2885 | 6.92E-07 | Other Negative Correlation |
| EPN3 | -0.28852 | 6.91E-07 | Other Negative Correlation |
| PPP1R35 | -0.28859 | 6.86E-07 | Other Negative Correlation |
| SMOC2 | -0.28859 | 6.86E-07 | Other Negative Correlation |
| PLCB3 | -0.28865 | 6.83E-07 | Other Negative Correlation |
| PRPF18 | -0.28881 | 6.73E-07 | Other Negative Correlation |
| TMED4 | -0.28882 | 6.72E-07 | Other Negative Correlation |
| APEX2 | -0.28888 | 6.68E-07 | Other Negative Correlation |
| ATG3 | -0.28888 | 6.68E-07 | Other Negative Correlation |
| NSDHL | -0.28893 | 6.65E-07 | Other Negative Correlation |
| MT-CO2 | -0.28904 | 6.59E-07 | Other Negative Correlation |
| SEC14L2 | -0.28924 | 6.46E-07 | Other Negative Correlation |
| SYK | -0.28932 | 6.41E-07 | Other Negative Correlation |

| | | | |
|---|---|---|---|
| MFSD14B | -0.28944 | 6.35E-07 | Other Negative Correlation |
| STK17B | -0.28948 | 6.32E-07 | Other Negative Correlation |
| COX17 | -0.28953 | 6.29E-07 | Other Negative Correlation |
| MSRB1 | -0.28957 | 6.27E-07 | Other Negative Correlation |
| PLCD1 | -0.2896 | 6.25E-07 | Other Negative Correlation |
| AHDC1 | -0.28964 | 6.23E-07 | Other Negative Correlation |
| TAX1BP1 | -0.28974 | 6.17E-07 | Other Negative Correlation |
| CIAO2B | -0.2898 | 6.13E-07 | Other Negative Correlation |
| PRCC | -0.28988 | 6.09E-07 | Other Negative Correlation |
| STAU1 | -0.28991 | 6.07E-07 | Other Negative Correlation |
| ACP1 | -0.28994 | 6.05E-07 | Other Negative Correlation |
| MFSD2A | -0.28997 | 6.04E-07 | Other Negative Correlation |
| BAD | -0.29 | 6.02E-07 | Other Negative Correlation |
| TMCO1 | -0.29002 | 6.01E-07 | Other Negative Correlation |
| PSMD4 | -0.29018 | 5.92E-07 | Other Negative Correlation |
| CCL2 | -0.29032 | 5.84E-07 | Other Negative Correlation |
| KANK1 | -0.29042 | 5.79E-07 | Other Negative Correlation |
| ST6GALNAC5 | -0.29047 | 5.76E-07 | Other Negative Correlation |
| HSD17B2 | -0.29048 | 5.76E-07 | Other Negative Correlation |
| SNX7 | -0.29061 | 5.69E-07 | Other Negative Correlation |
| ZNF277 | -0.29075 | 5.62E-07 | Other Negative Correlation |
| HSD11B1 | -0.29085 | 5.56E-07 | Other Negative Correlation |
| NLRX1 | -0.29087 | 5.55E-07 | Other Negative Correlation |
| MRPS15 | -0.29107 | 5.45E-07 | Other Negative Correlation |
| GHITM | -0.29108 | 5.44E-07 | Other Negative Correlation |
| DNAJC19 | -0.29119 | 5.39E-07 | Other Negative Correlation |
| ANKEF1 | -0.29141 | 5.28E-07 | Other Negative Correlation |
| IRGQ | -0.29141 | 5.28E-07 | Other Negative Correlation |
| PPP6R2 | -0.29141 | 5.28E-07 | Other Negative Correlation |
| CCNG2 | -0.29147 | 5.25E-07 | Other Negative Correlation |
| CDC40 | -0.29154 | 5.21E-07 | Other Negative Correlation |
| PRRC2A | -0.29166 | 5.16E-07 | Other Negative Correlation |
| PAF1 | -0.2917 | 5.14E-07 | Other Negative Correlation |
| VAMP3 | -0.29171 | 5.13E-07 | Other Negative Correlation |
| KATNBL1 | -0.29174 | 5.12E-07 | Other Negative Correlation |
| GFM2 | -0.29186 | 5.06E-07 | Other Negative Correlation |
| LSM1 | -0.29186 | 5.06E-07 | Other Negative Correlation |
| UBE2E1 | -0.29211 | 4.94E-07 | Other Negative Correlation |
| IL17RE | -0.29218 | 4.91E-07 | Other Negative Correlation |
| PCYT1A | -0.2922 | 4.9E-07 | Other Negative Correlation |
| C1GALT1 | -0.2923 | 4.86E-07 | Other Negative Correlation |
| TMEM87B | -0.29241 | 4.81E-07 | Other Negative Correlation |
| DAD1 | -0.29262 | 4.71E-07 | Other Negative Correlation |
| CHURC1 | -0.29272 | 4.67E-07 | Other Negative Correlation |
| ZNF217 | -0.29272 | 4.67E-07 | Other Negative Correlation |
| UBE2D1 | -0.29272 | 4.67E-07 | Other Negative Correlation |

| | | | |
|---|---|---|---|
| CX3CL1 | -0.29277 | 4.64E-07 | Other Negative Correlation |
| TRIM47 | -0.29282 | 4.63E-07 | Other Negative Correlation |
| PSMG2 | -0.29282 | 4.63E-07 | Other Negative Correlation |
| RNASET2 | -0.29282 | 4.62E-07 | Other Negative Correlation |
| MFF | -0.29287 | 4.6E-07 | Other Negative Correlation |
| FBH1 | -0.29292 | 4.58E-07 | Other Negative Correlation |
| H1-0 | -0.29304 | 4.53E-07 | Other Negative Correlation |
| TUT7 | -0.29313 | 4.49E-07 | Other Negative Correlation |
| SDSL | -0.29344 | 4.36E-07 | Other Negative Correlation |
| SPA17 | -0.29346 | 4.35E-07 | Other Negative Correlation |
| ZFC3H1 | -0.29349 | 4.34E-07 | Other Negative Correlation |
| CISD1 | -0.29391 | 4.17E-07 | Other Negative Correlation |
| GUCY2D | -0.29431 | 4.02E-07 | Other Negative Correlation |
| PPIL4 | -0.29435 | 4E-07 | Other Negative Correlation |
| LMO7 | -0.29454 | 3.93E-07 | Other Negative Correlation |
| AKIRIN2 | -0.29473 | 3.86E-07 | Other Negative Correlation |
| TMEM42 | -0.29478 | 3.84E-07 | Other Negative Correlation |
| CFAP20 | -0.29499 | 3.77E-07 | Other Negative Correlation |
| FMR1 | -0.29544 | 3.61E-07 | Other Negative Correlation |
| IFI6 | -0.29546 | 3.6E-07 | Other Negative Correlation |
| TYSND1 | -0.29553 | 3.58E-07 | Other Negative Correlation |
| TMPRSS4 | -0.29558 | 3.56E-07 | Other Negative Correlation |
| RELB | -0.29565 | 3.54E-07 | Other Negative Correlation |
| MRPS11 | -0.29576 | 3.5E-07 | Other Negative Correlation |
| PTRH1 | -0.29579 | 3.49E-07 | Other Negative Correlation |
| LAMP1 | -0.29585 | 3.47E-07 | Other Negative Correlation |
| LRRC1 | -0.29602 | 3.42E-07 | Other Negative Correlation |
| SH3RF1 | -0.29608 | 3.4E-07 | Other Negative Correlation |
| KCNN4 | -0.29615 | 3.37E-07 | Other Negative Correlation |
| PI3 | -0.29628 | 3.33E-07 | Other Negative Correlation |
| ERC1 | -0.29639 | 3.3E-07 | Other Negative Correlation |
| CNOT4 | -0.29654 | 3.25E-07 | Other Negative Correlation |
| AGFG1 | -0.29674 | 3.19E-07 | Other Negative Correlation |
| VAMP5 | -0.29683 | 3.16E-07 | Other Negative Correlation |
| LNPEP | -0.2969 | 3.14E-07 | Other Negative Correlation |
| NIPSNAP3A | -0.297 | 3.11E-07 | Other Negative Correlation |
| NCCRP1 | -0.29718 | 3.06E-07 | Other Negative Correlation |
| ATP6V0E1 | -0.29729 | 3.03E-07 | Other Negative Correlation |
| VPS13C | -0.29729 | 3.03E-07 | Other Negative Correlation |
| SLC35E1 | -0.29739 | 3E-07 | Other Negative Correlation |
| CAB39L | -0.29744 | 2.98E-07 | Other Negative Correlation |
| WBP1L | -0.29746 | 2.98E-07 | Other Negative Correlation |
| SUMO1 | -0.29762 | 2.93E-07 | Other Negative Correlation |
| SLAIN2 | -0.29771 | 2.91E-07 | Other Negative Correlation |
| YTHDF1 | -0.29771 | 2.9E-07 | Other Negative Correlation |
| ABHD17A | -0.29772 | 2.9E-07 | Other Negative Correlation |

| | | | |
|---|---|---|---|
| ZNF106 | -0.29772 | 2.9E-07 | Other Negative Correlation |
| PPP2R5E | -0.29782 | 2.87E-07 | Other Negative Correlation |
| TRPT1 | -0.29785 | 2.87E-07 | Other Negative Correlation |
| CHST15 | -0.29793 | 2.84E-07 | Other Negative Correlation |
| EAPP | -0.29794 | 2.84E-07 | Other Negative Correlation |
| DAXX | -0.29812 | 2.79E-07 | Other Negative Correlation |
| ICMT | -0.29816 | 2.78E-07 | Other Negative Correlation |
| PSMD5 | -0.29817 | 2.78E-07 | Other Negative Correlation |
| IFT74 | -0.29825 | 2.76E-07 | Other Negative Correlation |
| BCKDK | -0.29839 | 2.72E-07 | Other Negative Correlation |
| SHARPIN | -0.29843 | 2.71E-07 | Other Negative Correlation |
| SMAD3 | -0.29856 | 2.68E-07 | Other Negative Correlation |
| MRPL44 | -0.29869 | 2.65E-07 | Other Negative Correlation |
| MSLN | -0.2987 | 2.64E-07 | Other Negative Correlation |
| UNC50 | -0.29872 | 2.64E-07 | Other Negative Correlation |
| SS18 | -0.29884 | 2.61E-07 | Other Negative Correlation |
| WNT4 | -0.29886 | 2.6E-07 | Other Negative Correlation |
| CTSO | -0.29903 | 2.56E-07 | Other Negative Correlation |
| CDC42EP4 | -0.29906 | 2.55E-07 | Other Negative Correlation |
| PCMTD1 | -0.2992 | 2.52E-07 | Other Negative Correlation |
| APH1B | -0.29929 | 2.5E-07 | Other Negative Correlation |
| TSNAX | -0.29931 | 2.49E-07 | Other Negative Correlation |
| DLD | -0.29931 | 2.49E-07 | Other Negative Correlation |
| WIPF2 | -0.29931 | 2.49E-07 | Other Negative Correlation |
| PDXK | -0.29945 | 2.46E-07 | Other Negative Correlation |
| AP1B1 | -0.29949 | 2.45E-07 | Other Negative Correlation |
| ATOSB | -0.29963 | 2.42E-07 | Other Negative Correlation |
| OSBPL9 | -0.29965 | 2.41E-07 | Other Negative Correlation |
| SENP2 | -0.29966 | 2.41E-07 | Other Negative Correlation |
| EFNA1 | -0.29978 | 2.38E-07 | Other Negative Correlation |
| PARD3 | -0.29982 | 2.37E-07 | Other Negative Correlation |
| BCAS4 | -0.29989 | 2.36E-07 | Other Negative Correlation |
| EHBP1 | -0.29994 | 2.34E-07 | Other Negative Correlation |
| H2BC21 | -0.30014 | 2.3E-07 | Moderate Negative Correlation |
| SYNE4 | -0.30022 | 2.28E-07 | Moderate Negative Correlation |
| OXA1L | -0.30023 | 2.28E-07 | Moderate Negative Correlation |
| CD2BP2 | -0.30025 | 2.28E-07 | Moderate Negative Correlation |
| MCFD2 | -0.30031 | 2.26E-07 | Moderate Negative Correlation |
| USB1 | -0.30032 | 2.26E-07 | Moderate Negative Correlation |
| ERLEC1 | -0.30033 | 2.26E-07 | Moderate Negative Correlation |
| SDCCAG8 | -0.30072 | 2.17E-07 | Moderate Negative Correlation |
| GBP2 | -0.30081 | 2.16E-07 | Moderate Negative Correlation |
| HMGN3 | -0.30086 | 2.14E-07 | Moderate Negative Correlation |
| AFDN | -0.30095 | 2.12E-07 | Moderate Negative Correlation |
| KIAA1143 | -0.30106 | 2.1E-07 | Moderate Negative Correlation |
| GGT6 | -0.3011 | 2.09E-07 | Moderate Negative Correlation |

| | | | |
|---|---|---|---|
| MAP3K7 | -0.30111 | 2.09E-07 | Moderate Negative Correlation |
| ACBD5 | -0.30122 | 2.07E-07 | Moderate Negative Correlation |
| RASA1 | -0.30127 | 2.06E-07 | Moderate Negative Correlation |
| ELOVL4 | -0.30139 | 2.04E-07 | Moderate Negative Correlation |
| MED15 | -0.30143 | 2.03E-07 | Moderate Negative Correlation |
| SERPINA1 | -0.3015 | 2.01E-07 | Moderate Negative Correlation |
| EDEM1 | -0.3016 | 2E-07 | Moderate Negative Correlation |
| RNF213 | -0.30164 | 1.99E-07 | Moderate Negative Correlation |
| GORASP2 | -0.30168 | 1.98E-07 | Moderate Negative Correlation |
| NIPBL | -0.30168 | 1.98E-07 | Moderate Negative Correlation |
| KRT81 | -0.3017 | 1.98E-07 | Moderate Negative Correlation |
| SFXN1 | -0.30174 | 1.97E-07 | Moderate Negative Correlation |
| ADIPOR2 | -0.30181 | 1.96E-07 | Moderate Negative Correlation |
| GTF2B | -0.30191 | 1.94E-07 | Moderate Negative Correlation |
| DEFB1 | -0.30193 | 1.93E-07 | Moderate Negative Correlation |
| DAB2 | -0.30204 | 1.91E-07 | Moderate Negative Correlation |
| USP14 | -0.30219 | 1.88E-07 | Moderate Negative Correlation |
| TAOK1 | -0.30225 | 1.87E-07 | Moderate Negative Correlation |
| EMC7 | -0.3023 | 1.86E-07 | Moderate Negative Correlation |
| LRPAP1 | -0.3025 | 1.83E-07 | Moderate Negative Correlation |
| ARMCX6 | -0.30251 | 1.83E-07 | Moderate Negative Correlation |
| ARHGEF4 | -0.30279 | 1.78E-07 | Moderate Negative Correlation |
| SAA2 | -0.30285 | 1.77E-07 | Moderate Negative Correlation |
| SWAP70 | -0.30313 | 1.72E-07 | Moderate Negative Correlation |
| SPINT1 | -0.30319 | 1.71E-07 | Moderate Negative Correlation |
| ARFGEF1 | -0.30321 | 1.7E-07 | Moderate Negative Correlation |
| UBAC1 | -0.30323 | 1.7E-07 | Moderate Negative Correlation |
| COX4I1 | -0.30338 | 1.68E-07 | Moderate Negative Correlation |
| MINK1 | -0.3034 | 1.67E-07 | Moderate Negative Correlation |
| DR1 | -0.30344 | 1.67E-07 | Moderate Negative Correlation |
| RAB42 | -0.30352 | 1.65E-07 | Moderate Negative Correlation |
| MPRIP | -0.30353 | 1.65E-07 | Moderate Negative Correlation |
| IGFBP6 | -0.30354 | 1.65E-07 | Moderate Negative Correlation |
| SPTBN2 | -0.30356 | 1.65E-07 | Moderate Negative Correlation |
| YIPF6 | -0.30362 | 1.64E-07 | Moderate Negative Correlation |
| LCMT1 | -0.3038 | 1.61E-07 | Moderate Negative Correlation |
| MCCC2 | -0.30386 | 1.6E-07 | Moderate Negative Correlation |
| SNX17 | -0.3039 | 1.59E-07 | Moderate Negative Correlation |
| ZMAT2 | -0.30393 | 1.59E-07 | Moderate Negative Correlation |
| F2RL1 | -0.30401 | 1.58E-07 | Moderate Negative Correlation |
| SOD1 | -0.30404 | 1.57E-07 | Moderate Negative Correlation |
| RBFOX2 | -0.30409 | 1.56E-07 | Moderate Negative Correlation |
| CAMK2G | -0.30412 | 1.56E-07 | Moderate Negative Correlation |
| ALAS1 | -0.30414 | 1.56E-07 | Moderate Negative Correlation |
| NUDT19 | -0.30417 | 1.55E-07 | Moderate Negative Correlation |
| MARCHF6 | -0.30438 | 1.52E-07 | Moderate Negative Correlation |

| | | | |
|---|---|---|---|
| MRPS6 | -0.30444 | 1.51E-07 | Moderate Negative Correlation |
| TRIM8 | -0.30475 | 1.47E-07 | Moderate Negative Correlation |
| FAM168B | -0.3049 | 1.45E-07 | Moderate Negative Correlation |
| FAM181B | -0.30498 | 1.43E-07 | Moderate Negative Correlation |
| MAP4K5 | -0.30521 | 1.4E-07 | Moderate Negative Correlation |
| PLEKHM2 | -0.30527 | 1.39E-07 | Moderate Negative Correlation |
| SLC35F5 | -0.30527 | 1.39E-07 | Moderate Negative Correlation |
| MRS2 | -0.30538 | 1.38E-07 | Moderate Negative Correlation |
| MRPL55 | -0.30538 | 1.38E-07 | Moderate Negative Correlation |
| USP47 | -0.30545 | 1.37E-07 | Moderate Negative Correlation |
| IBTK | -0.30561 | 1.35E-07 | Moderate Negative Correlation |
| GPATCH2L | -0.30571 | 1.33E-07 | Moderate Negative Correlation |
| SYNJ2BP | -0.30574 | 1.33E-07 | Moderate Negative Correlation |
| NRBF2 | -0.30582 | 1.32E-07 | Moderate Negative Correlation |
| PLEKHA2 | -0.30583 | 1.32E-07 | Moderate Negative Correlation |
| ACTR6 | -0.30599 | 1.3E-07 | Moderate Negative Correlation |
| TMTC3 | -0.30601 | 1.29E-07 | Moderate Negative Correlation |
| CANT1 | -0.30617 | 1.27E-07 | Moderate Negative Correlation |
| SIPA1L1 | -0.30622 | 1.27E-07 | Moderate Negative Correlation |
| CAAP1 | -0.30623 | 1.27E-07 | Moderate Negative Correlation |
| TNFRSF1A | -0.30627 | 1.26E-07 | Moderate Negative Correlation |
| ATXN10 | -0.3066 | 1.22E-07 | Moderate Negative Correlation |
| NDUFB1 | -0.30661 | 1.22E-07 | Moderate Negative Correlation |
| LYST | -0.30665 | 1.21E-07 | Moderate Negative Correlation |
| DAAM1 | -0.30666 | 1.21E-07 | Moderate Negative Correlation |
| DDA1 | -0.30666 | 1.21E-07 | Moderate Negative Correlation |
| MARK2 | -0.30688 | 1.19E-07 | Moderate Negative Correlation |
| RNF43 | -0.30694 | 1.18E-07 | Moderate Negative Correlation |
| ABR | -0.30699 | 1.17E-07 | Moderate Negative Correlation |
| NAP1L4 | -0.30723 | 1.15E-07 | Moderate Negative Correlation |
| ARSI | -0.30727 | 1.14E-07 | Moderate Negative Correlation |
| SLC16A3 | -0.30771 | 1.09E-07 | Moderate Negative Correlation |
| GIGYF2 | -0.30795 | 1.07E-07 | Moderate Negative Correlation |
| MRFAP1L1 | -0.30798 | 1.06E-07 | Moderate Negative Correlation |
| BLTP2 | -0.30807 | 1.06E-07 | Moderate Negative Correlation |
| ALDH3B2 | -0.30812 | 1.05E-07 | Moderate Negative Correlation |
| LGR4 | -0.30818 | 1.04E-07 | Moderate Negative Correlation |
| SNAPC1 | -0.30836 | 1.03E-07 | Moderate Negative Correlation |
| F11R | -0.30845 | 1.02E-07 | Moderate Negative Correlation |
| HSBP1L1 | -0.30852 | 1.01E-07 | Moderate Negative Correlation |
| TICAM1 | -0.30874 | 9.87E-08 | Moderate Negative Correlation |
| NDN | -0.30909 | 9.53E-08 | Moderate Negative Correlation |
| CCDC113 | -0.30918 | 9.44E-08 | Moderate Negative Correlation |
| PPP1R7 | -0.30922 | 9.4E-08 | Moderate Negative Correlation |
| GPBP1L1 | -0.30924 | 9.39E-08 | Moderate Negative Correlation |
| KIAA1191 | -0.30936 | 9.27E-08 | Moderate Negative Correlation |

| | | | |
|---|---|---|---|
| HRCT1 | -0.30943 | 9.21E-08 | Moderate Negative Correlation |
| AP3D1 | -0.3095 | 9.15E-08 | Moderate Negative Correlation |
| ZBTB4 | -0.30952 | 9.12E-08 | Moderate Negative Correlation |
| CBY1 | -0.30954 | 9.11E-08 | Moderate Negative Correlation |
| COMMD6 | -0.30956 | 9.09E-08 | Moderate Negative Correlation |
| UBALD1 | -0.30961 | 9.05E-08 | Moderate Negative Correlation |
| TBC1D10A | -0.30964 | 9.02E-08 | Moderate Negative Correlation |
| GREP1 | -0.3097 | 8.97E-08 | Moderate Negative Correlation |
| TNPO1 | -0.30983 | 8.85E-08 | Moderate Negative Correlation |
| DCTN1 | -0.30995 | 8.74E-08 | Moderate Negative Correlation |
| C14orf119 | -0.30996 | 8.74E-08 | Moderate Negative Correlation |
| RIDA | -0.3101 | 8.61E-08 | Moderate Negative Correlation |
| SNX12 | -0.31013 | 8.59E-08 | Moderate Negative Correlation |
| CARD19 | -0.31039 | 8.37E-08 | Moderate Negative Correlation |
| SPAG9 | -0.31062 | 8.17E-08 | Moderate Negative Correlation |
| FRG1 | -0.31066 | 8.14E-08 | Moderate Negative Correlation |
| BORCS7 | -0.31071 | 8.1E-08 | Moderate Negative Correlation |
| GLG1 | -0.31073 | 8.08E-08 | Moderate Negative Correlation |
| ECI2 | -0.31077 | 8.05E-08 | Moderate Negative Correlation |
| PPME1 | -0.31081 | 8.01E-08 | Moderate Negative Correlation |
| GATAD2A | -0.3111 | 7.78E-08 | Moderate Negative Correlation |
| NMRK1 | -0.31136 | 7.59E-08 | Moderate Negative Correlation |
| MTMR6 | -0.31137 | 7.57E-08 | Moderate Negative Correlation |
| MAN1A2 | -0.31147 | 7.5E-08 | Moderate Negative Correlation |
| SLC9A7 | -0.31162 | 7.39E-08 | Moderate Negative Correlation |
| RC3H2 | -0.31169 | 7.33E-08 | Moderate Negative Correlation |
| PTTG1 | -0.31171 | 7.32E-08 | Moderate Negative Correlation |
| MYEOV | -0.31173 | 7.3E-08 | Moderate Negative Correlation |
| KIFAP3 | -0.31175 | 7.29E-08 | Moderate Negative Correlation |
| NAB1 | -0.31176 | 7.28E-08 | Moderate Negative Correlation |
| FBXO7 | -0.31199 | 7.11E-08 | Moderate Negative Correlation |
| TCAF1 | -0.31202 | 7.09E-08 | Moderate Negative Correlation |
| TMEM205 | -0.31204 | 7.08E-08 | Moderate Negative Correlation |
| ACP5 | -0.3121 | 7.04E-08 | Moderate Negative Correlation |
| GNPTAB | -0.31217 | 6.99E-08 | Moderate Negative Correlation |
| YIPF4 | -0.31224 | 6.94E-08 | Moderate Negative Correlation |
| C1QTNF6 | -0.31226 | 6.92E-08 | Moderate Negative Correlation |
| TAP1 | -0.31242 | 6.82E-08 | Moderate Negative Correlation |
| NME4 | -0.31261 | 6.68E-08 | Moderate Negative Correlation |
| CITED2 | -0.31269 | 6.63E-08 | Moderate Negative Correlation |
| UBE2F | -0.3128 | 6.56E-08 | Moderate Negative Correlation |
| IL6R | -0.31295 | 6.45E-08 | Moderate Negative Correlation |
| RAB6A | -0.313 | 6.42E-08 | Moderate Negative Correlation |
| IRAK2 | -0.31304 | 6.4E-08 | Moderate Negative Correlation |
| MID1 | -0.3131 | 6.36E-08 | Moderate Negative Correlation |
| ABCF1 | -0.31318 | 6.31E-08 | Moderate Negative Correlation |

| | | | |
|---|---|---|---|
| SYF2 | -0.31332 | 6.22E-08 | Moderate Negative Correlation |
| RETREG3 | -0.31345 | 6.14E-08 | Moderate Negative Correlation |
| CHMP1A | -0.31347 | 6.13E-08 | Moderate Negative Correlation |
| HNRNPH2 | -0.3135 | 6.1E-08 | Moderate Negative Correlation |
| CLDN4 | -0.31359 | 6.05E-08 | Moderate Negative Correlation |
| CASP6 | -0.31364 | 6.02E-08 | Moderate Negative Correlation |
| PSMB7 | -0.31365 | 6.01E-08 | Moderate Negative Correlation |
| TOR4A | -0.31366 | 6.01E-08 | Moderate Negative Correlation |
| ITFG1 | -0.31373 | 5.96E-08 | Moderate Negative Correlation |
| NCOR1 | -0.31418 | 5.7E-08 | Moderate Negative Correlation |
| STX19 | -0.31433 | 5.61E-08 | Moderate Negative Correlation |
| TMPRSS11E | -0.31446 | 5.53E-08 | Moderate Negative Correlation |
| TM2D1 | -0.31455 | 5.49E-08 | Moderate Negative Correlation |
| SGMS2 | -0.31459 | 5.46E-08 | Moderate Negative Correlation |
| PPARD | -0.31493 | 5.28E-08 | Moderate Negative Correlation |
| ARIH1 | -0.315 | 5.24E-08 | Moderate Negative Correlation |
| MN1 | -0.31509 | 5.19E-08 | Moderate Negative Correlation |
| ZNF185 | -0.31521 | 5.13E-08 | Moderate Negative Correlation |
| LYRM2 | -0.31538 | 5.04E-08 | Moderate Negative Correlation |
| APOBEC3F | -0.31546 | 5E-08 | Moderate Negative Correlation |
| ERGIC3 | -0.31551 | 4.97E-08 | Moderate Negative Correlation |
| TMEM92 | -0.31554 | 4.96E-08 | Moderate Negative Correlation |
| KLHDC8B | -0.31576 | 4.85E-08 | Moderate Negative Correlation |
| IQANK1 | -0.31605 | 4.7E-08 | Moderate Negative Correlation |
| CNN3 | -0.31609 | 4.68E-08 | Moderate Negative Correlation |
| PIAS1 | -0.31613 | 4.66E-08 | Moderate Negative Correlation |
| MYL6B | -0.31621 | 4.63E-08 | Moderate Negative Correlation |
| PLEKHN1 | -0.31627 | 4.6E-08 | Moderate Negative Correlation |
| PAM | -0.31628 | 4.6E-08 | Moderate Negative Correlation |
| TSEN34 | -0.31633 | 4.57E-08 | Moderate Negative Correlation |
| POLB | -0.31634 | 4.57E-08 | Moderate Negative Correlation |
| HELZ2 | -0.31645 | 4.51E-08 | Moderate Negative Correlation |
| RNF135 | -0.31658 | 4.46E-08 | Moderate Negative Correlation |
| CD2AP | -0.31669 | 4.4E-08 | Moderate Negative Correlation |
| GNAQ | -0.31672 | 4.39E-08 | Moderate Negative Correlation |
| HACD3 | -0.31703 | 4.25E-08 | Moderate Negative Correlation |
| ACAA2 | -0.31728 | 4.15E-08 | Moderate Negative Correlation |
| HBEGF | -0.31737 | 4.1E-08 | Moderate Negative Correlation |
| AGPAT3 | -0.31751 | 4.05E-08 | Moderate Negative Correlation |
| ROCK2 | -0.31758 | 4.02E-08 | Moderate Negative Correlation |
| PTGR1 | -0.31771 | 3.97E-08 | Moderate Negative Correlation |
| CNOT2 | -0.31772 | 3.96E-08 | Moderate Negative Correlation |
| TMEM219 | -0.31779 | 3.93E-08 | Moderate Negative Correlation |
| CARD16 | -0.31795 | 3.87E-08 | Moderate Negative Correlation |
| FEM1B | -0.31799 | 3.85E-08 | Moderate Negative Correlation |
| ORAI3 | -0.31802 | 3.84E-08 | Moderate Negative Correlation |

| | | | |
|---|---|---|---|
| ARAP1 | -0.31811 | 3.8E-08 | Moderate Negative Correlation |
| DUSP14 | -0.31818 | 3.78E-08 | Moderate Negative Correlation |
| KLHL9 | -0.31844 | 3.68E-08 | Moderate Negative Correlation |
| ST3GAL5 | -0.31859 | 3.62E-08 | Moderate Negative Correlation |
| EMC2 | -0.31865 | 3.6E-08 | Moderate Negative Correlation |
| SH3GL1 | -0.31873 | 3.57E-08 | Moderate Negative Correlation |
| SNX1 | -0.31905 | 3.45E-08 | Moderate Negative Correlation |
| TBC1D9B | -0.31909 | 3.44E-08 | Moderate Negative Correlation |
| FBXO3 | -0.31914 | 3.42E-08 | Moderate Negative Correlation |
| SH3PXD2B | -0.3192 | 3.4E-08 | Moderate Negative Correlation |
| IRX2 | -0.31932 | 3.36E-08 | Moderate Negative Correlation |
| RILPL1 | -0.31947 | 3.3E-08 | Moderate Negative Correlation |
| PPFIA1 | -0.31965 | 3.24E-08 | Moderate Negative Correlation |
| SMIM22 | -0.31978 | 3.2E-08 | Moderate Negative Correlation |
| HLA-E | -0.3199 | 3.16E-08 | Moderate Negative Correlation |
| ARHGEF37 | -0.31997 | 3.14E-08 | Moderate Negative Correlation |
| CYB561A3 | -0.31999 | 3.13E-08 | Moderate Negative Correlation |
| RBM47 | -0.32007 | 3.1E-08 | Moderate Negative Correlation |
| TOP2B | -0.32037 | 3.01E-08 | Moderate Negative Correlation |
| KRAS | -0.32055 | 2.95E-08 | Moderate Negative Correlation |
| CERCAM | -0.32057 | 2.95E-08 | Moderate Negative Correlation |
| NTN4 | -0.32061 | 2.93E-08 | Moderate Negative Correlation |
| BOK | -0.32063 | 2.93E-08 | Moderate Negative Correlation |
| ME2 | -0.32071 | 2.9E-08 | Moderate Negative Correlation |
| CALML5 | -0.32072 | 2.9E-08 | Moderate Negative Correlation |
| AKAP13 | -0.32102 | 2.81E-08 | Moderate Negative Correlation |
| SLC9A1 | -0.32119 | 2.76E-08 | Moderate Negative Correlation |
| MKRN2OS | -0.32128 | 2.74E-08 | Moderate Negative Correlation |
| CPA4 | -0.32131 | 2.73E-08 | Moderate Negative Correlation |
| LASP1 | -0.32172 | 2.61E-08 | Moderate Negative Correlation |
| EIPR1 | -0.32175 | 2.6E-08 | Moderate Negative Correlation |
| DHRSX | -0.3218 | 2.59E-08 | Moderate Negative Correlation |
| UCP2 | -0.32182 | 2.58E-08 | Moderate Negative Correlation |
| TSC22D4 | -0.32187 | 2.57E-08 | Moderate Negative Correlation |
| CREB3 | -0.3222 | 2.48E-08 | Moderate Negative Correlation |
| CXCL17 | -0.32261 | 2.38E-08 | Moderate Negative Correlation |
| SEMA4A | -0.32268 | 2.36E-08 | Moderate Negative Correlation |
| DNTTIP1 | -0.32276 | 2.34E-08 | Moderate Negative Correlation |
| TMEM167A | -0.32299 | 2.29E-08 | Moderate Negative Correlation |
| TCEAL1 | -0.32301 | 2.28E-08 | Moderate Negative Correlation |
| SETD7 | -0.32304 | 2.27E-08 | Moderate Negative Correlation |
| CHPF | -0.32329 | 2.22E-08 | Moderate Negative Correlation |
| SPTBN1 | -0.32336 | 2.2E-08 | Moderate Negative Correlation |
| RAET1G | -0.32341 | 2.19E-08 | Moderate Negative Correlation |
| LGALSL | -0.32362 | 2.14E-08 | Moderate Negative Correlation |
| TOM1L2 | -0.32374 | 2.11E-08 | Moderate Negative Correlation |

| | | | |
|---|---|---|---|
| ACBD3 | -0.32391 | 2.08E-08 | Moderate Negative Correlation |
| SPRR2A | -0.32392 | 2.07E-08 | Moderate Negative Correlation |
| ARMCX2 | -0.32418 | 2.02E-08 | Moderate Negative Correlation |
| RHCG | -0.32442 | 1.97E-08 | Moderate Negative Correlation |
| NDFIP1 | -0.32443 | 1.96E-08 | Moderate Negative Correlation |
| PRAG1 | -0.32473 | 1.9E-08 | Moderate Negative Correlation |
| C15orf48 | -0.32477 | 1.89E-08 | Moderate Negative Correlation |
| MARCHF5 | -0.32505 | 1.84E-08 | Moderate Negative Correlation |
| SYNPO | -0.32517 | 1.82E-08 | Moderate Negative Correlation |
| ZCCHC17 | -0.32538 | 1.78E-08 | Moderate Negative Correlation |
| LLPH | -0.32547 | 1.76E-08 | Moderate Negative Correlation |
| TPD52L1 | -0.32558 | 1.74E-08 | Moderate Negative Correlation |
| MTX1 | -0.32562 | 1.73E-08 | Moderate Negative Correlation |
| SULT1A2 | -0.32567 | 1.72E-08 | Moderate Negative Correlation |
| JAK1 | -0.3257 | 1.72E-08 | Moderate Negative Correlation |
| TM9SF3 | -0.32579 | 1.7E-08 | Moderate Negative Correlation |
| WWP1 | -0.32599 | 1.66E-08 | Moderate Negative Correlation |
| NAA35 | -0.32602 | 1.66E-08 | Moderate Negative Correlation |
| NMRAL1 | -0.32626 | 1.62E-08 | Moderate Negative Correlation |
| TCEAL4 | -0.32643 | 1.59E-08 | Moderate Negative Correlation |
| ESRRA | -0.32646 | 1.58E-08 | Moderate Negative Correlation |
| FAM110A | -0.32646 | 1.58E-08 | Moderate Negative Correlation |
| STARD10 | -0.32664 | 1.55E-08 | Moderate Negative Correlation |
| SLMAP | -0.32678 | 1.53E-08 | Moderate Negative Correlation |
| ADIPOR1 | -0.32691 | 1.51E-08 | Moderate Negative Correlation |
| RBM15B | -0.32694 | 1.5E-08 | Moderate Negative Correlation |
| FBXO6 | -0.32702 | 1.49E-08 | Moderate Negative Correlation |
| MAT2B | -0.32726 | 1.45E-08 | Moderate Negative Correlation |
| MAP2K3 | -0.32732 | 1.44E-08 | Moderate Negative Correlation |
| NADSYN1 | -0.32733 | 1.44E-08 | Moderate Negative Correlation |
| POLR2K | -0.3274 | 1.43E-08 | Moderate Negative Correlation |
| MT-ND3 | -0.32745 | 1.42E-08 | Moderate Negative Correlation |
| SUN1 | -0.32747 | 1.42E-08 | Moderate Negative Correlation |
| MT-CO3 | -0.32752 | 1.41E-08 | Moderate Negative Correlation |
| IVD | -0.32772 | 1.38E-08 | Moderate Negative Correlation |
| NDUFS8 | -0.32783 | 1.37E-08 | Moderate Negative Correlation |
| SMAD1 | -0.32788 | 1.36E-08 | Moderate Negative Correlation |
| STK3 | -0.328 | 1.34E-08 | Moderate Negative Correlation |
| NIPAL4 | -0.32813 | 1.32E-08 | Moderate Negative Correlation |
| RRAS | -0.32827 | 1.3E-08 | Moderate Negative Correlation |
| RAB14 | -0.32831 | 1.3E-08 | Moderate Negative Correlation |
| MARK4 | -0.32843 | 1.28E-08 | Moderate Negative Correlation |
| RWDD2B | -0.32864 | 1.25E-08 | Moderate Negative Correlation |
| ACER3 | -0.32875 | 1.24E-08 | Moderate Negative Correlation |
| ZSWIM7 | -0.32876 | 1.24E-08 | Moderate Negative Correlation |
| SCYL2 | -0.32897 | 1.21E-08 | Moderate Negative Correlation |

| | | | |
|---|---|---|---|
| NACC1 | -0.32901 | 1.2E-08 | Moderate Negative Correlation |
| RIT1 | -0.32917 | 1.18E-08 | Moderate Negative Correlation |
| CROT | -0.32946 | 1.15E-08 | Moderate Negative Correlation |
| PRKAR2A | -0.32952 | 1.14E-08 | Moderate Negative Correlation |
| CXCL8 | -0.32967 | 1.12E-08 | Moderate Negative Correlation |
| EXOC5 | -0.3297 | 1.12E-08 | Moderate Negative Correlation |
| TFDP2 | -0.32997 | 1.09E-08 | Moderate Negative Correlation |
| ARHGAP5 | -0.33034 | 1.04E-08 | Moderate Negative Correlation |
| ARL4D | -0.33037 | 1.04E-08 | Moderate Negative Correlation |
| TASOR | -0.3304 | 1.04E-08 | Moderate Negative Correlation |
| AKAP9 | -0.33051 | 1.02E-08 | Moderate Negative Correlation |
| GTF2F1 | -0.33055 | 1.02E-08 | Moderate Negative Correlation |
| HMGN4 | -0.33068 | 1.01E-08 | Moderate Negative Correlation |
| AAGAB | -0.33109 | 9.61E-09 | Moderate Negative Correlation |
| PARP14 | -0.33111 | 9.6E-09 | Moderate Negative Correlation |
| IKBKE | -0.33115 | 9.55E-09 | Moderate Negative Correlation |
| CLIP4 | -0.33124 | 9.46E-09 | Moderate Negative Correlation |
| MPLKIP | -0.33127 | 9.43E-09 | Moderate Negative Correlation |
| XPA | -0.33134 | 9.36E-09 | Moderate Negative Correlation |
| CCDC115 | -0.33169 | 9.01E-09 | Moderate Negative Correlation |
| POLR2J | -0.3319 | 8.8E-09 | Moderate Negative Correlation |
| AGL | -0.33208 | 8.64E-09 | Moderate Negative Correlation |
| MRPL28 | -0.33215 | 8.57E-09 | Moderate Negative Correlation |
| WASHC3 | -0.33267 | 8.1E-09 | Moderate Negative Correlation |
| AP2A1 | -0.33273 | 8.05E-09 | Moderate Negative Correlation |
| HCCS | -0.33285 | 7.94E-09 | Moderate Negative Correlation |
| PPP4R2 | -0.33298 | 7.83E-09 | Moderate Negative Correlation |
| COMMD2 | -0.33299 | 7.82E-09 | Moderate Negative Correlation |
| AURKAIP1 | -0.33306 | 7.76E-09 | Moderate Negative Correlation |
| MAGED1 | -0.33318 | 7.66E-09 | Moderate Negative Correlation |
| WIPI2 | -0.33336 | 7.51E-09 | Moderate Negative Correlation |
| H2BC5 | -0.33344 | 7.45E-09 | Moderate Negative Correlation |
| ZDHHC12 | -0.33351 | 7.39E-09 | Moderate Negative Correlation |
| PLEKHA5 | -0.33362 | 7.3E-09 | Moderate Negative Correlation |
| PGF | -0.33372 | 7.22E-09 | Moderate Negative Correlation |
| TOX4 | -0.33376 | 7.19E-09 | Moderate Negative Correlation |
| FIP1L1 | -0.33381 | 7.15E-09 | Moderate Negative Correlation |
| WNT7A | -0.33406 | 6.96E-09 | Moderate Negative Correlation |
| CUEDC2 | -0.33429 | 6.78E-09 | Moderate Negative Correlation |
| STRN | -0.33434 | 6.75E-09 | Moderate Negative Correlation |
| KLC3 | -0.33448 | 6.64E-09 | Moderate Negative Correlation |
| EPN2 | -0.3345 | 6.63E-09 | Moderate Negative Correlation |
| CNIH4 | -0.33457 | 6.58E-09 | Moderate Negative Correlation |
| SERPINB13 | -0.33459 | 6.57E-09 | Moderate Negative Correlation |
| GNB2 | -0.33484 | 6.39E-09 | Moderate Negative Correlation |
| TRAK1 | -0.33515 | 6.17E-09 | Moderate Negative Correlation |

| | | | |
|---|---|---|---|
| TMX3 | -0.33516 | 6.17E-09 | Moderate Negative Correlation |
| COPS4 | -0.33532 | 6.06E-09 | Moderate Negative Correlation |
| YME1L1 | -0.33534 | 6.04E-09 | Moderate Negative Correlation |
| NFRKB | -0.3354 | 6E-09 | Moderate Negative Correlation |
| CDC73 | -0.33551 | 5.93E-09 | Moderate Negative Correlation |
| DENND10 | -0.336 | 5.62E-09 | Moderate Negative Correlation |
| POLR2E | -0.33611 | 5.55E-09 | Moderate Negative Correlation |
| RAB11B | -0.33614 | 5.54E-09 | Moderate Negative Correlation |
| OGDH | -0.33633 | 5.42E-09 | Moderate Negative Correlation |
| SIN3B | -0.33658 | 5.27E-09 | Moderate Negative Correlation |
| CMC1 | -0.3366 | 5.26E-09 | Moderate Negative Correlation |
| ZDHHC20 | -0.33691 | 5.08E-09 | Moderate Negative Correlation |
| APOC1 | -0.33696 | 5.05E-09 | Moderate Negative Correlation |
| MOB4 | -0.33717 | 4.94E-09 | Moderate Negative Correlation |
| CASP7 | -0.33774 | 4.63E-09 | Moderate Negative Correlation |
| PPP2R3A | -0.33783 | 4.59E-09 | Moderate Negative Correlation |
| DDAH2 | -0.33803 | 4.49E-09 | Moderate Negative Correlation |
| MFAP1 | -0.33832 | 4.35E-09 | Moderate Negative Correlation |
| ATP5MJ | -0.33854 | 4.24E-09 | Moderate Negative Correlation |
| CCDC43 | -0.33875 | 4.14E-09 | Moderate Negative Correlation |
| MRPL41 | -0.3389 | 4.07E-09 | Moderate Negative Correlation |
| PSMD6 | -0.33924 | 3.92E-09 | Moderate Negative Correlation |
| SEC62 | -0.33928 | 3.9E-09 | Moderate Negative Correlation |
| IFT43 | -0.33945 | 3.83E-09 | Moderate Negative Correlation |
| BAZ2B | -0.33959 | 3.77E-09 | Moderate Negative Correlation |
| TIMM17A | -0.33965 | 3.75E-09 | Moderate Negative Correlation |
| PROM2 | -0.34004 | 3.58E-09 | Moderate Negative Correlation |
| LEPROTL1 | -0.34021 | 3.52E-09 | Moderate Negative Correlation |
| ASAH1 | -0.3403 | 3.48E-09 | Moderate Negative Correlation |
| SMARCA1 | -0.3404 | 3.45E-09 | Moderate Negative Correlation |
| BABAM1 | -0.34053 | 3.4E-09 | Moderate Negative Correlation |
| TFPT | -0.34084 | 3.28E-09 | Moderate Negative Correlation |
| GAPVD1 | -0.34088 | 3.26E-09 | Moderate Negative Correlation |
| DCTN5 | -0.34116 | 3.16E-09 | Moderate Negative Correlation |
| ILK | -0.34132 | 3.11E-09 | Moderate Negative Correlation |
| IRF3 | -0.34132 | 3.11E-09 | Moderate Negative Correlation |
| FUCA2 | -0.34143 | 3.07E-09 | Moderate Negative Correlation |
| NECTIN2 | -0.3417 | 2.98E-09 | Moderate Negative Correlation |
| MISP | -0.34184 | 2.93E-09 | Moderate Negative Correlation |
| SLC35D2 | -0.34185 | 2.92E-09 | Moderate Negative Correlation |
| ENDOD1 | -0.34195 | 2.9E-09 | Moderate Negative Correlation |
| ITPRIPL2 | -0.34204 | 2.87E-09 | Moderate Negative Correlation |
| RRBP1 | -0.3421 | 2.84E-09 | Moderate Negative Correlation |
| RHBDF1 | -0.34227 | 2.79E-09 | Moderate Negative Correlation |
| TIMMDC1 | -0.34234 | 2.77E-09 | Moderate Negative Correlation |
| TPCN1 | -0.34266 | 2.67E-09 | Moderate Negative Correlation |

| | | | |
|---|---|---|---|
| RTF2 | -0.34274 | 2.65E-09 | Moderate Negative Correlation |
| UBTD1 | -0.34275 | 2.64E-09 | Moderate Negative Correlation |
| CCDC6 | -0.34277 | 2.64E-09 | Moderate Negative Correlation |
| NCF2 | -0.34283 | 2.62E-09 | Moderate Negative Correlation |
| RNASEH2C | -0.34291 | 2.6E-09 | Moderate Negative Correlation |
| THUMPD1 | -0.34306 | 2.55E-09 | Moderate Negative Correlation |
| DCTN2 | -0.34306 | 2.55E-09 | Moderate Negative Correlation |
| C6orf132 | -0.34313 | 2.53E-09 | Moderate Negative Correlation |
| USP9X | -0.34327 | 2.49E-09 | Moderate Negative Correlation |
| RABEP1 | -0.34351 | 2.43E-09 | Moderate Negative Correlation |
| CNPY2 | -0.34358 | 2.41E-09 | Moderate Negative Correlation |
| GSTA4 | -0.34373 | 2.37E-09 | Moderate Negative Correlation |
| ARHGAP17 | -0.3438 | 2.35E-09 | Moderate Negative Correlation |
| NDUFC1 | -0.34382 | 2.34E-09 | Moderate Negative Correlation |
| ARHGAP29 | -0.34396 | 2.31E-09 | Moderate Negative Correlation |
| CLDND1 | -0.344 | 2.29E-09 | Moderate Negative Correlation |
| RNF115 | -0.34407 | 2.28E-09 | Moderate Negative Correlation |
| FAAP20 | -0.34417 | 2.25E-09 | Moderate Negative Correlation |
| EMP2 | -0.34417 | 2.25E-09 | Moderate Negative Correlation |
| DNAJC4 | -0.34429 | 2.22E-09 | Moderate Negative Correlation |
| DENND2C | -0.34433 | 2.21E-09 | Moderate Negative Correlation |
| KIF3A | -0.34463 | 2.14E-09 | Moderate Negative Correlation |
| DHX29 | -0.34464 | 2.13E-09 | Moderate Negative Correlation |
| COQ10B | -0.34467 | 2.13E-09 | Moderate Negative Correlation |
| UQCRQ | -0.34485 | 2.08E-09 | Moderate Negative Correlation |
| PDZD11 | -0.34488 | 2.08E-09 | Moderate Negative Correlation |
| LDOC1 | -0.34493 | 2.06E-09 | Moderate Negative Correlation |
| PCYOX1 | -0.34496 | 2.06E-09 | Moderate Negative Correlation |
| GALNT11 | -0.34525 | 1.99E-09 | Moderate Negative Correlation |
| PRXL2A | -0.34545 | 1.95E-09 | Moderate Negative Correlation |
| SH3TC2 | -0.34547 | 1.94E-09 | Moderate Negative Correlation |
| ADIRF | -0.34561 | 1.91E-09 | Moderate Negative Correlation |
| CCDC28B | -0.34563 | 1.91E-09 | Moderate Negative Correlation |
| ABAT | -0.34563 | 1.91E-09 | Moderate Negative Correlation |
| COL16A1 | -0.34565 | 1.9E-09 | Moderate Negative Correlation |
| LIPA | -0.34573 | 1.88E-09 | Moderate Negative Correlation |
| PIAS3 | -0.34578 | 1.87E-09 | Moderate Negative Correlation |
| CCNH | -0.34606 | 1.82E-09 | Moderate Negative Correlation |
| RNPEP | -0.34615 | 1.8E-09 | Moderate Negative Correlation |
| STXBP3 | -0.34619 | 1.79E-09 | Moderate Negative Correlation |
| PTGES | -0.34622 | 1.78E-09 | Moderate Negative Correlation |
| TAFA5 | -0.34627 | 1.77E-09 | Moderate Negative Correlation |
| LAMTOR4 | -0.34641 | 1.74E-09 | Moderate Negative Correlation |
| BAG5 | -0.34643 | 1.74E-09 | Moderate Negative Correlation |
| RHOG | -0.34653 | 1.72E-09 | Moderate Negative Correlation |
| KLK6 | -0.34662 | 1.7E-09 | Moderate Negative Correlation |

| | | | |
|---|---|---|---|
| HIPK3 | -0.34676 | 1.67E-09 | Moderate Negative Correlation |
| NENF | -0.34683 | 1.66E-09 | Moderate Negative Correlation |
| PAPOLA | -0.34687 | 1.65E-09 | Moderate Negative Correlation |
| FTH1 | -0.3471 | 1.61E-09 | Moderate Negative Correlation |
| PAPSS1 | -0.34731 | 1.57E-09 | Moderate Negative Correlation |
| RCOR1 | -0.34732 | 1.57E-09 | Moderate Negative Correlation |
| DDRGK1 | -0.34743 | 1.55E-09 | Moderate Negative Correlation |
| QSOX1 | -0.34746 | 1.55E-09 | Moderate Negative Correlation |
| RUFY2 | -0.34765 | 1.51E-09 | Moderate Negative Correlation |
| ZNF750 | -0.34774 | 1.5E-09 | Moderate Negative Correlation |
| ERG28 | -0.34775 | 1.49E-09 | Moderate Negative Correlation |
| DPCD | -0.3478 | 1.49E-09 | Moderate Negative Correlation |
| H2BC4 | -0.34788 | 1.47E-09 | Moderate Negative Correlation |
| FUCA1 | -0.34801 | 1.45E-09 | Moderate Negative Correlation |
| CFAP36 | -0.34803 | 1.45E-09 | Moderate Negative Correlation |
| NUB1 | -0.34804 | 1.45E-09 | Moderate Negative Correlation |
| CETN2 | -0.34806 | 1.44E-09 | Moderate Negative Correlation |
| SHROOM3 | -0.34816 | 1.43E-09 | Moderate Negative Correlation |
| OAF | -0.34827 | 1.41E-09 | Moderate Negative Correlation |
| SDHAF3 | -0.34837 | 1.39E-09 | Moderate Negative Correlation |
| PJA2 | -0.3484 | 1.39E-09 | Moderate Negative Correlation |
| ERBIN | -0.34847 | 1.38E-09 | Moderate Negative Correlation |
| TRIP12 | -0.34871 | 1.34E-09 | Moderate Negative Correlation |
| ST14 | -0.34877 | 1.33E-09 | Moderate Negative Correlation |
| CDC5L | -0.34877 | 1.33E-09 | Moderate Negative Correlation |
| FNDC3B | -0.34882 | 1.32E-09 | Moderate Negative Correlation |
| STX4 | -0.34887 | 1.31E-09 | Moderate Negative Correlation |
| SPCS1 | -0.34891 | 1.31E-09 | Moderate Negative Correlation |
| BIRC6 | -0.34899 | 1.3E-09 | Moderate Negative Correlation |
| DYNC1H1 | -0.34899 | 1.3E-09 | Moderate Negative Correlation |
| ARID4B | -0.34902 | 1.29E-09 | Moderate Negative Correlation |
| FAIM | -0.34912 | 1.28E-09 | Moderate Negative Correlation |
| UFC1 | -0.34947 | 1.23E-09 | Moderate Negative Correlation |
| C19orf53 | -0.3495 | 1.22E-09 | Moderate Negative Correlation |
| DCUN1D1 | -0.34964 | 1.2E-09 | Moderate Negative Correlation |
| ACADVL | -0.34972 | 1.19E-09 | Moderate Negative Correlation |
| PAIP1 | -0.34988 | 1.17E-09 | Moderate Negative Correlation |
| DUSP10 | -0.35028 | 1.12E-09 | Moderate Negative Correlation |
| ATP6V1C1 | -0.35032 | 1.11E-09 | Moderate Negative Correlation |
| ENDOG | -0.35049 | 1.09E-09 | Moderate Negative Correlation |
| ZNF789 | -0.35068 | 1.06E-09 | Moderate Negative Correlation |
| BPNT1 | -0.35097 | 1.03E-09 | Moderate Negative Correlation |
| LRBA | -0.35103 | 1.02E-09 | Moderate Negative Correlation |
| SLC2A6 | -0.35108 | 1.02E-09 | Moderate Negative Correlation |
| CHMP4B | -0.35151 | 9.67E-10 | Moderate Negative Correlation |
| SEC61G | -0.3517 | 9.46E-10 | Moderate Negative Correlation |

| | | | |
|---|---|---|---|
| INKA1 | -0.35176 | 9.39E-10 | Moderate Negative Correlation |
| RNF144B | -0.35179 | 9.36E-10 | Moderate Negative Correlation |
| DLG1 | -0.3518 | 9.35E-10 | Moderate Negative Correlation |
| CHMP2B | -0.35197 | 9.17E-10 | Moderate Negative Correlation |
| PARL | -0.35201 | 9.12E-10 | Moderate Negative Correlation |
| SUGT1 | -0.3522 | 8.92E-10 | Moderate Negative Correlation |
| SCPEP1 | -0.35247 | 8.64E-10 | Moderate Negative Correlation |
| LYPLA2 | -0.35249 | 8.62E-10 | Moderate Negative Correlation |
| SYNGR2 | -0.35262 | 8.49E-10 | Moderate Negative Correlation |
| WFDC5 | -0.35275 | 8.37E-10 | Moderate Negative Correlation |
| MIB1 | -0.35276 | 8.35E-10 | Moderate Negative Correlation |
| CCL20 | -0.35297 | 8.15E-10 | Moderate Negative Correlation |
| ANKFY1 | -0.3531 | 8.03E-10 | Moderate Negative Correlation |
| BMP1 | -0.35327 | 7.87E-10 | Moderate Negative Correlation |
| DLG5 | -0.35335 | 7.8E-10 | Moderate Negative Correlation |
| KLK11 | -0.3536 | 7.57E-10 | Moderate Negative Correlation |
| REEP3 | -0.35404 | 7.19E-10 | Moderate Negative Correlation |
| NAV2 | -0.35432 | 6.96E-10 | Moderate Negative Correlation |
| CDC27 | -0.35439 | 6.9E-10 | Moderate Negative Correlation |
| TST | -0.35441 | 6.88E-10 | Moderate Negative Correlation |
| CBR1 | -0.35475 | 6.61E-10 | Moderate Negative Correlation |
| BRD7 | -0.35504 | 6.39E-10 | Moderate Negative Correlation |
| ST13 | -0.35558 | 5.99E-10 | Moderate Negative Correlation |
| AKIRIN1 | -0.35561 | 5.98E-10 | Moderate Negative Correlation |
| JARID2 | -0.35569 | 5.91E-10 | Moderate Negative Correlation |
| VCP | -0.35573 | 5.89E-10 | Moderate Negative Correlation |
| KPNA3 | -0.35588 | 5.79E-10 | Moderate Negative Correlation |
| HDLBP | -0.35588 | 5.79E-10 | Moderate Negative Correlation |
| KDM5A | -0.35591 | 5.77E-10 | Moderate Negative Correlation |
| TM7SF2 | -0.35627 | 5.52E-10 | Moderate Negative Correlation |
| DHRS4L2 | -0.35644 | 5.41E-10 | Moderate Negative Correlation |
| FDXR | -0.35673 | 5.23E-10 | Moderate Negative Correlation |
| ACOT13 | -0.35697 | 5.09E-10 | Moderate Negative Correlation |
| PINK1 | -0.35705 | 5.03E-10 | Moderate Negative Correlation |
| MBP | -0.35728 | 4.9E-10 | Moderate Negative Correlation |
| MDH2 | -0.35728 | 4.9E-10 | Moderate Negative Correlation |
| DCAF6 | -0.35737 | 4.85E-10 | Moderate Negative Correlation |
| TXN2 | -0.35741 | 4.83E-10 | Moderate Negative Correlation |
| BRD2 | -0.35745 | 4.8E-10 | Moderate Negative Correlation |
| MRPL36 | -0.35771 | 4.66E-10 | Moderate Negative Correlation |
| INAVA | -0.35844 | 4.27E-10 | Moderate Negative Correlation |
| SELENOT | -0.35866 | 4.16E-10 | Moderate Negative Correlation |
| LETM1 | -0.35885 | 4.06E-10 | Moderate Negative Correlation |
| TRAPPC3 | -0.35886 | 4.06E-10 | Moderate Negative Correlation |
| PSMB6 | -0.35895 | 4.01E-10 | Moderate Negative Correlation |
| ARFIP2 | -0.35914 | 3.92E-10 | Moderate Negative Correlation |

| | | | |
|---|---|---|---|
| OLFML2A | -0.35922 | 3.89E-10 | Moderate Negative Correlation |
| FADS3 | -0.3597 | 3.67E-10 | Moderate Negative Correlation |
| PRELID3B | -0.35974 | 3.65E-10 | Moderate Negative Correlation |
| PIK3C3 | -0.35975 | 3.65E-10 | Moderate Negative Correlation |
| VWA5A | -0.3598 | 3.63E-10 | Moderate Negative Correlation |
| RHOV | -0.36011 | 3.49E-10 | Moderate Negative Correlation |
| CLIP1 | -0.36024 | 3.44E-10 | Moderate Negative Correlation |
| DNAJA2 | -0.36026 | 3.43E-10 | Moderate Negative Correlation |
| TRADD | -0.36027 | 3.43E-10 | Moderate Negative Correlation |
| UBA1 | -0.3604 | 3.37E-10 | Moderate Negative Correlation |
| JAGN1 | -0.36067 | 3.27E-10 | Moderate Negative Correlation |
| PSMA3 | -0.36074 | 3.24E-10 | Moderate Negative Correlation |
| S100A8 | -0.36077 | 3.23E-10 | Moderate Negative Correlation |
| CPNE1 | -0.36082 | 3.21E-10 | Moderate Negative Correlation |
| CACUL1 | -0.36097 | 3.15E-10 | Moderate Negative Correlation |
| SNRPA1 | -0.36129 | 3.03E-10 | Moderate Negative Correlation |
| PYM1 | -0.36139 | 2.99E-10 | Moderate Negative Correlation |
| M6PR | -0.36171 | 2.88E-10 | Moderate Negative Correlation |
| PAFAH1B1 | -0.36183 | 2.84E-10 | Moderate Negative Correlation |
| TCF25 | -0.36184 | 2.84E-10 | Moderate Negative Correlation |
| SELENOF | -0.36185 | 2.83E-10 | Moderate Negative Correlation |
| ENAH | -0.36189 | 2.82E-10 | Moderate Negative Correlation |
| TMEM9B | -0.36215 | 2.73E-10 | Moderate Negative Correlation |
| PTGR3 | -0.36219 | 2.72E-10 | Moderate Negative Correlation |
| TUBB2A | -0.36236 | 2.66E-10 | Moderate Negative Correlation |
| KIF21A | -0.36239 | 2.65E-10 | Moderate Negative Correlation |
| KCTD20 | -0.3627 | 2.55E-10 | Moderate Negative Correlation |
| MAPK3 | -0.36274 | 2.54E-10 | Moderate Negative Correlation |
| PNP | -0.36282 | 2.52E-10 | Moderate Negative Correlation |
| NCK1 | -0.36317 | 2.42E-10 | Moderate Negative Correlation |
| UQCRC1 | -0.36319 | 2.41E-10 | Moderate Negative Correlation |
| BIRC3 | -0.36341 | 2.34E-10 | Moderate Negative Correlation |
| RCAN1 | -0.36342 | 2.34E-10 | Moderate Negative Correlation |
| MNAT1 | -0.36374 | 2.25E-10 | Moderate Negative Correlation |
| CLPP | -0.36375 | 2.25E-10 | Moderate Negative Correlation |
| MPHOSPH8 | -0.36403 | 2.17E-10 | Moderate Negative Correlation |
| ZNF770 | -0.3646 | 2.03E-10 | Moderate Negative Correlation |
| TMED2 | -0.36461 | 2.03E-10 | Moderate Negative Correlation |
| MPZL1 | -0.36521 | 1.88E-10 | Moderate Negative Correlation |
| CELSR2 | -0.36533 | 1.86E-10 | Moderate Negative Correlation |
| RAB5A | -0.36534 | 1.85E-10 | Moderate Negative Correlation |
| RAB18 | -0.36541 | 1.84E-10 | Moderate Negative Correlation |
| PPP1R2 | -0.36546 | 1.83E-10 | Moderate Negative Correlation |
| KLK5 | -0.36563 | 1.79E-10 | Moderate Negative Correlation |
| DYNLT1 | -0.36578 | 1.76E-10 | Moderate Negative Correlation |
| ACSF2 | -0.36587 | 1.74E-10 | Moderate Negative Correlation |

| | | | |
|---|---|---|---|
| PUM1 | -0.36621 | 1.67E-10 | Moderate Negative Correlation |
| CCDC91 | -0.36625 | 1.66E-10 | Moderate Negative Correlation |
| RAB5C | -0.36632 | 1.64E-10 | Moderate Negative Correlation |
| CWC15 | -0.36644 | 1.62E-10 | Moderate Negative Correlation |
| SLC26A2 | -0.36684 | 1.54E-10 | Moderate Negative Correlation |
| FAM174A | -0.36689 | 1.53E-10 | Moderate Negative Correlation |
| PIGS | -0.36725 | 1.47E-10 | Moderate Negative Correlation |
| PIN4 | -0.36729 | 1.46E-10 | Moderate Negative Correlation |
| UBA3 | -0.36731 | 1.46E-10 | Moderate Negative Correlation |
| RHOBTB3 | -0.36746 | 1.43E-10 | Moderate Negative Correlation |
| MAN1B1 | -0.36755 | 1.41E-10 | Moderate Negative Correlation |
| PDCD10 | -0.36773 | 1.38E-10 | Moderate Negative Correlation |
| CHIC2 | -0.36798 | 1.34E-10 | Moderate Negative Correlation |
| CD58 | -0.36819 | 1.31E-10 | Moderate Negative Correlation |
| MFN2 | -0.36846 | 1.26E-10 | Moderate Negative Correlation |
| NBR1 | -0.36881 | 1.21E-10 | Moderate Negative Correlation |
| FSTL3 | -0.36886 | 1.2E-10 | Moderate Negative Correlation |
| TMED3 | -0.36899 | 1.18E-10 | Moderate Negative Correlation |
| COQ9 | -0.36916 | 1.16E-10 | Moderate Negative Correlation |
| UBE2L6 | -0.36923 | 1.15E-10 | Moderate Negative Correlation |
| EIF4G3 | -0.36955 | 1.1E-10 | Moderate Negative Correlation |
| OTUB1 | -0.37032 | 1E-10 | Moderate Negative Correlation |
| ZDHHC24 | -0.37051 | 9.78E-11 | Moderate Negative Correlation |
| SPRED1 | -0.37064 | 9.63E-11 | Moderate Negative Correlation |
| BPNT2 | -0.37083 | 9.4E-11 | Moderate Negative Correlation |
| LPCAT2 | -0.37084 | 9.4E-11 | Moderate Negative Correlation |
| TPR | -0.37103 | 9.18E-11 | Moderate Negative Correlation |
| CDKN2B | -0.37104 | 9.16E-11 | Moderate Negative Correlation |
| RALA | -0.37115 | 9.04E-11 | Moderate Negative Correlation |
| PSMD1 | -0.37115 | 9.03E-11 | Moderate Negative Correlation |
| VTA1 | -0.37119 | 8.99E-11 | Moderate Negative Correlation |
| ZG16B | -0.37131 | 8.86E-11 | Moderate Negative Correlation |
| SLC37A2 | -0.37133 | 8.83E-11 | Moderate Negative Correlation |
| TMEM59 | -0.37152 | 8.63E-11 | Moderate Negative Correlation |
| BMPR2 | -0.37157 | 8.57E-11 | Moderate Negative Correlation |
| NDUFB7 | -0.37171 | 8.43E-11 | Moderate Negative Correlation |
| WTAP | -0.37199 | 8.13E-11 | Moderate Negative Correlation |
| BNIP3L | -0.37205 | 8.08E-11 | Moderate Negative Correlation |
| RNF11 | -0.37214 | 7.99E-11 | Moderate Negative Correlation |
| MRPL50 | -0.37217 | 7.96E-11 | Moderate Negative Correlation |
| VANGL1 | -0.37246 | 7.67E-11 | Moderate Negative Correlation |
| ACAA1 | -0.37258 | 7.56E-11 | Moderate Negative Correlation |
| UBXN6 | -0.37266 | 7.48E-11 | Moderate Negative Correlation |
| MEAF6 | -0.37285 | 7.3E-11 | Moderate Negative Correlation |
| TBCB | -0.37299 | 7.17E-11 | Moderate Negative Correlation |
| DNAJC5 | -0.37334 | 6.87E-11 | Moderate Negative Correlation |

| | | | |
|---|---|---|---|
| COPB2 | -0.3736 | 6.65E-11 | Moderate Negative Correlation |
| RRAGA | -0.37377 | 6.51E-11 | Moderate Negative Correlation |
| NDUFS4 | -0.37378 | 6.5E-11 | Moderate Negative Correlation |
| TMEM184B | -0.37393 | 6.38E-11 | Moderate Negative Correlation |
| MRPS25 | -0.37395 | 6.36E-11 | Moderate Negative Correlation |
| PRPSAP1 | -0.37401 | 6.32E-11 | Moderate Negative Correlation |
| BAG1 | -0.37412 | 6.23E-11 | Moderate Negative Correlation |
| MARCKSL1 | -0.37413 | 6.22E-11 | Moderate Negative Correlation |
| RABGGTA | -0.37419 | 6.17E-11 | Moderate Negative Correlation |
| OSBPL1A | -0.37429 | 6.09E-11 | Moderate Negative Correlation |
| LOXL1 | -0.37431 | 6.08E-11 | Moderate Negative Correlation |
| REST | -0.37446 | 5.97E-11 | Moderate Negative Correlation |
| ATP6V1A | -0.3752 | 5.43E-11 | Moderate Negative Correlation |
| ACTR1A | -0.37521 | 5.43E-11 | Moderate Negative Correlation |
| EPHA1 | -0.37567 | 5.12E-11 | Moderate Negative Correlation |
| SCARB2 | -0.3759 | 4.97E-11 | Moderate Negative Correlation |
| NDUFC2 | -0.37611 | 4.84E-11 | Moderate Negative Correlation |
| IGF2R | -0.37619 | 4.79E-11 | Moderate Negative Correlation |
| CMPK1 | -0.37627 | 4.75E-11 | Moderate Negative Correlation |
| JUND | -0.37673 | 4.47E-11 | Moderate Negative Correlation |
| RHBDL2 | -0.37674 | 4.47E-11 | Moderate Negative Correlation |
| NIPAL1 | -0.37683 | 4.41E-11 | Moderate Negative Correlation |
| HDAC1 | -0.37684 | 4.41E-11 | Moderate Negative Correlation |
| HECTD1 | -0.37691 | 4.37E-11 | Moderate Negative Correlation |
| GINM1 | -0.37711 | 4.26E-11 | Moderate Negative Correlation |
| RSF1 | -0.37715 | 4.24E-11 | Moderate Negative Correlation |
| NDUFS3 | -0.37717 | 4.23E-11 | Moderate Negative Correlation |
| MRPL17 | -0.37745 | 4.08E-11 | Moderate Negative Correlation |
| PITRM1 | -0.37754 | 4.03E-11 | Moderate Negative Correlation |
| FYTTD1 | -0.37789 | 3.86E-11 | Moderate Negative Correlation |
| APMAP | -0.37809 | 3.76E-11 | Moderate Negative Correlation |
| TMBIM1 | -0.37826 | 3.68E-11 | Moderate Negative Correlation |
| STX6 | -0.3785 | 3.57E-11 | Moderate Negative Correlation |
| IGFL1 | -0.37866 | 3.5E-11 | Moderate Negative Correlation |
| COX5B | -0.37906 | 3.32E-11 | Moderate Negative Correlation |
| RRP36 | -0.37922 | 3.25E-11 | Moderate Negative Correlation |
| KIAA1217 | -0.37971 | 3.05E-11 | Moderate Negative Correlation |
| PRDX5 | -0.37976 | 3.04E-11 | Moderate Negative Correlation |
| OXSR1 | -0.38005 | 2.93E-11 | Moderate Negative Correlation |
| SMARCA4 | -0.38006 | 2.92E-11 | Moderate Negative Correlation |
| HOPX | -0.38021 | 2.87E-11 | Moderate Negative Correlation |
| NIPAL3 | -0.38029 | 2.84E-11 | Moderate Negative Correlation |
| TNFAIP8 | -0.38031 | 2.83E-11 | Moderate Negative Correlation |
| CAPZA2 | -0.38088 | 2.63E-11 | Moderate Negative Correlation |
| TTC1 | -0.38101 | 2.58E-11 | Moderate Negative Correlation |
| SRPX2 | -0.38134 | 2.48E-11 | Moderate Negative Correlation |

| | | | |
|---|---|---|---|
| UQCRC2 | -0.38135 | 2.47E-11 | Moderate Negative Correlation |
| OPN3 | -0.38139 | 2.46E-11 | Moderate Negative Correlation |
| TLK1 | -0.38168 | 2.37E-11 | Moderate Negative Correlation |
| SMARCD2 | -0.38188 | 2.31E-11 | Moderate Negative Correlation |
| PSME3 | -0.38218 | 2.22E-11 | Moderate Negative Correlation |
| HDGFL2 | -0.38226 | 2.2E-11 | Moderate Negative Correlation |
| TSPAN3 | -0.38245 | 2.15E-11 | Moderate Negative Correlation |
| PPP1R10 | -0.38254 | 2.12E-11 | Moderate Negative Correlation |
| AHCYL1 | -0.3829 | 2.02E-11 | Moderate Negative Correlation |
| UBE2R2 | -0.383 | 2E-11 | Moderate Negative Correlation |
| ABL2 | -0.38301 | 2E-11 | Moderate Negative Correlation |
| STAG2 | -0.38306 | 1.98E-11 | Moderate Negative Correlation |
| IFNAR1 | -0.38306 | 1.98E-11 | Moderate Negative Correlation |
| UQCRFS1 | -0.38312 | 1.97E-11 | Moderate Negative Correlation |
| TANK | -0.38313 | 1.97E-11 | Moderate Negative Correlation |
| MYD88 | -0.38317 | 1.95E-11 | Moderate Negative Correlation |
| CARD6 | -0.38325 | 1.93E-11 | Moderate Negative Correlation |
| FOXQ1 | -0.38327 | 1.93E-11 | Moderate Negative Correlation |
| FBXO2 | -0.38359 | 1.85E-11 | Moderate Negative Correlation |
| CNTN1 | -0.38363 | 1.84E-11 | Moderate Negative Correlation |
| IK | -0.38384 | 1.79E-11 | Moderate Negative Correlation |
| UQCR11 | -0.38395 | 1.76E-11 | Moderate Negative Correlation |
| KYNU | -0.38426 | 1.7E-11 | Moderate Negative Correlation |
| TRIR | -0.38435 | 1.67E-11 | Moderate Negative Correlation |
| YY1 | -0.38454 | 1.63E-11 | Moderate Negative Correlation |
| PPP1CA | -0.38456 | 1.63E-11 | Moderate Negative Correlation |
| MED4 | -0.38457 | 1.63E-11 | Moderate Negative Correlation |
| SMIM26 | -0.38459 | 1.62E-11 | Moderate Negative Correlation |
| FUBP3 | -0.38463 | 1.61E-11 | Moderate Negative Correlation |
| C9orf78 | -0.38478 | 1.58E-11 | Moderate Negative Correlation |
| ITGB8 | -0.38502 | 1.54E-11 | Moderate Negative Correlation |
| SCO2 | -0.38515 | 1.51E-11 | Moderate Negative Correlation |
| MIDN | -0.38546 | 1.45E-11 | Moderate Negative Correlation |
| MKNK2 | -0.38553 | 1.44E-11 | Moderate Negative Correlation |
| CPSF2 | -0.38554 | 1.43E-11 | Moderate Negative Correlation |
| PLXNB2 | -0.38593 | 1.36E-11 | Moderate Negative Correlation |
| EFR3A | -0.38598 | 1.35E-11 | Moderate Negative Correlation |
| RNF13 | -0.38623 | 1.31E-11 | Moderate Negative Correlation |
| PTTG1IP | -0.3865 | 1.26E-11 | Moderate Negative Correlation |
| H2AC6 | -0.38661 | 1.25E-11 | Moderate Negative Correlation |
| SSH3 | -0.38664 | 1.24E-11 | Moderate Negative Correlation |
| LAPTM4B | -0.38673 | 1.23E-11 | Moderate Negative Correlation |
| SDF2 | -0.38687 | 1.2E-11 | Moderate Negative Correlation |
| HOMER1 | -0.38708 | 1.17E-11 | Moderate Negative Correlation |
| AKR1C1 | -0.38715 | 1.16E-11 | Moderate Negative Correlation |
| CXADR | -0.38754 | 1.1E-11 | Moderate Negative Correlation |

| | | | |
|---|---|---|---|
| RAET1L | -0.38794 | 1.04E-11 | Moderate Negative Correlation |
| GIPC1 | -0.38807 | 1.03E-11 | Moderate Negative Correlation |
| COMMD3 | -0.38837 | 9.88E-12 | Moderate Negative Correlation |
| SLC31A2 | -0.38839 | 9.85E-12 | Moderate Negative Correlation |
| BDH2 | -0.38857 | 9.62E-12 | Moderate Negative Correlation |
| CAMK2B | -0.38887 | 9.24E-12 | Moderate Negative Correlation |
| GALNT7 | -0.38914 | 8.92E-12 | Moderate Negative Correlation |
| CHD4 | -0.38918 | 8.88E-12 | Moderate Negative Correlation |
| ABHD17C | -0.3892 | 8.85E-12 | Moderate Negative Correlation |
| N4BP1 | -0.38969 | 8.29E-12 | Moderate Negative Correlation |
| ELOC | -0.38986 | 8.1E-12 | Moderate Negative Correlation |
| PLLP | -0.39001 | 7.94E-12 | Moderate Negative Correlation |
| AFF4 | -0.39016 | 7.79E-12 | Moderate Negative Correlation |
| MALSU1 | -0.39066 | 7.28E-12 | Moderate Negative Correlation |
| IGSF3 | -0.39072 | 7.22E-12 | Moderate Negative Correlation |
| UQCR10 | -0.391 | 6.96E-12 | Moderate Negative Correlation |
| PIK3IP1 | -0.39106 | 6.91E-12 | Moderate Negative Correlation |
| CLASP1 | -0.39111 | 6.86E-12 | Moderate Negative Correlation |
| NFIC | -0.3913 | 6.69E-12 | Moderate Negative Correlation |
| MMP24OS | -0.39133 | 6.66E-12 | Moderate Negative Correlation |
| CAPN12 | -0.39153 | 6.48E-12 | Moderate Negative Correlation |
| ARL2BP | -0.39157 | 6.45E-12 | Moderate Negative Correlation |
| LAYN | -0.39168 | 6.36E-12 | Moderate Negative Correlation |
| FRMD8 | -0.39183 | 6.23E-12 | Moderate Negative Correlation |
| NDUFB5 | -0.39195 | 6.13E-12 | Moderate Negative Correlation |
| SOX7 | -0.39197 | 6.11E-12 | Moderate Negative Correlation |
| BDP1 | -0.39201 | 6.08E-12 | Moderate Negative Correlation |
| RAB9A | -0.39215 | 5.97E-12 | Moderate Negative Correlation |
| OGFRL1 | -0.39249 | 5.7E-12 | Moderate Negative Correlation |
| HSD17B10 | -0.3926 | 5.62E-12 | Moderate Negative Correlation |
| MGMT | -0.39264 | 5.59E-12 | Moderate Negative Correlation |
| ADGRG1 | -0.39287 | 5.42E-12 | Moderate Negative Correlation |
| USP53 | -0.39291 | 5.39E-12 | Moderate Negative Correlation |
| PPA2 | -0.3932 | 5.19E-12 | Moderate Negative Correlation |
| BPTF | -0.3935 | 4.98E-12 | Moderate Negative Correlation |
| MYO6 | -0.3935 | 4.98E-12 | Moderate Negative Correlation |
| TOB1 | -0.39394 | 4.69E-12 | Moderate Negative Correlation |
| UROS | -0.39397 | 4.67E-12 | Moderate Negative Correlation |
| GRHPR | -0.39421 | 4.53E-12 | Moderate Negative Correlation |
| FXR1 | -0.39432 | 4.46E-12 | Moderate Negative Correlation |
| MRPS18C | -0.39433 | 4.45E-12 | Moderate Negative Correlation |
| CD99 | -0.39439 | 4.42E-12 | Moderate Negative Correlation |
| NDUFS2 | -0.39455 | 4.32E-12 | Moderate Negative Correlation |
| LMTK3 | -0.39473 | 4.22E-12 | Moderate Negative Correlation |
| DEXI | -0.39481 | 4.17E-12 | Moderate Negative Correlation |
| PHLDA3 | -0.39483 | 4.16E-12 | Moderate Negative Correlation |

| | | | |
|---|---|---|---|
| C5orf15 | -0.39537 | 3.87E-12 | Moderate Negative Correlation |
| CTSZ | -0.39559 | 3.76E-12 | Moderate Negative Correlation |
| ATP2B4 | -0.39585 | 3.62E-12 | Moderate Negative Correlation |
| RDH11 | -0.39591 | 3.59E-12 | Moderate Negative Correlation |
| PNRC1 | -0.39609 | 3.51E-12 | Moderate Negative Correlation |
| KLK10 | -0.3969 | 3.14E-12 | Moderate Negative Correlation |
| COA3 | -0.3969 | 3.14E-12 | Moderate Negative Correlation |
| ARL1 | -0.39726 | 2.99E-12 | Moderate Negative Correlation |
| CAMTA1 | -0.39808 | 2.67E-12 | Moderate Negative Correlation |
| PPCS | -0.39823 | 2.62E-12 | Moderate Negative Correlation |
| ORMDL2 | -0.39871 | 2.45E-12 | Moderate Negative Correlation |
| CAMK2D | -0.39898 | 2.36E-12 | Moderate Negative Correlation |
| LY6D | -0.39926 | 2.27E-12 | Moderate Negative Correlation |
| WASHC4 | -0.39934 | 2.25E-12 | Moderate Negative Correlation |
| SAMD9 | -0.39956 | 2.18E-12 | Moderate Negative Correlation |
| GRAMD2B | -0.39965 | 2.16E-12 | Moderate Negative Correlation |
| PSORS1C1 | -0.39976 | 2.13E-12 | Moderate Negative Correlation |
| LGALS8 | -0.40032 | 1.97E-12 | Moderate Negative Correlation |
| CTTNBP2NL | -0.40071 | 1.86E-12 | Moderate Negative Correlation |
| AP3B1 | -0.40104 | 1.78E-12 | Moderate Negative Correlation |
| ADD1 | -0.40109 | 1.77E-12 | Moderate Negative Correlation |
| SULT2B1 | -0.40129 | 1.72E-12 | Moderate Negative Correlation |
| NECTIN4 | -0.40134 | 1.71E-12 | Moderate Negative Correlation |
| MLLT11 | -0.40146 | 1.68E-12 | Moderate Negative Correlation |
| ADGRF4 | -0.40169 | 1.63E-12 | Moderate Negative Correlation |
| SP140L | -0.40172 | 1.62E-12 | Moderate Negative Correlation |
| YIF1A | -0.40191 | 1.58E-12 | Moderate Negative Correlation |
| PFN2 | -0.40214 | 1.53E-12 | Moderate Negative Correlation |
| IP6K2 | -0.40248 | 1.46E-12 | Moderate Negative Correlation |
| BLOC1S6 | -0.40256 | 1.44E-12 | Moderate Negative Correlation |
| NUDT22 | -0.4026 | 1.43E-12 | Moderate Negative Correlation |
| MRPL35 | -0.40316 | 1.33E-12 | Moderate Negative Correlation |
| AGTRAP | -0.40321 | 1.32E-12 | Moderate Negative Correlation |
| ZBTB7A | -0.4033 | 1.3E-12 | Moderate Negative Correlation |
| BBOX1 | -0.4034 | 1.28E-12 | Moderate Negative Correlation |
| TMEM106B | -0.40347 | 1.27E-12 | Moderate Negative Correlation |
| SLC30A1 | -0.40395 | 1.19E-12 | Moderate Negative Correlation |
| VDAC3 | -0.40407 | 1.17E-12 | Moderate Negative Correlation |
| OAZ2 | -0.40408 | 1.17E-12 | Moderate Negative Correlation |
| NCOA4 | -0.40413 | 1.16E-12 | Moderate Negative Correlation |
| NDUFA8 | -0.40436 | 1.12E-12 | Moderate Negative Correlation |
| MAP2K2 | -0.40443 | 1.11E-12 | Moderate Negative Correlation |
| ARF5 | -0.40517 | 1E-12 | Moderate Negative Correlation |
| ZKSCAN1 | -0.40518 | 1E-12 | Moderate Negative Correlation |
| EXOC6B | -0.40576 | 9.23E-13 | Moderate Negative Correlation |
| SUPT5H | -0.40577 | 9.22E-13 | Moderate Negative Correlation |

| | | | |
|---|---|---|---|
| BAG3 | -0.40597 | 8.96E-13 | Moderate Negative Correlation |
| CIR1 | -0.40601 | 8.91E-13 | Moderate Negative Correlation |
| HPS5 | -0.40605 | 8.87E-13 | Moderate Negative Correlation |
| RSU1 | -0.40623 | 8.64E-13 | Moderate Negative Correlation |
| EMP1 | -0.40663 | 8.17E-13 | Moderate Negative Correlation |
| ASMTL | -0.40669 | 8.1E-13 | Moderate Negative Correlation |
| KRCC1 | -0.4068 | 7.98E-13 | Moderate Negative Correlation |
| TRIM16 | -0.40706 | 7.68E-13 | Moderate Negative Correlation |
| SUCLA2 | -0.40763 | 7.1E-13 | Moderate Negative Correlation |
| ECH1 | -0.40786 | 6.87E-13 | Moderate Negative Correlation |
| ANK3 | -0.40827 | 6.48E-13 | Moderate Negative Correlation |
| COX7B | -0.40862 | 6.17E-13 | Moderate Negative Correlation |
| SLC9A3R1 | -0.40889 | 5.94E-13 | Moderate Negative Correlation |
| VGLL1 | -0.40896 | 5.87E-13 | Moderate Negative Correlation |
| SUDS3 | -0.40946 | 5.48E-13 | Moderate Negative Correlation |
| EXOC7 | -0.40948 | 5.46E-13 | Moderate Negative Correlation |
| TRMT6 | -0.40952 | 5.43E-13 | Moderate Negative Correlation |
| SAP18 | -0.40981 | 5.21E-13 | Moderate Negative Correlation |
| LAMP2 | -0.41019 | 4.94E-13 | Moderate Negative Correlation |
| NFKBIA | -0.4102 | 4.93E-13 | Moderate Negative Correlation |
| CCDC107 | -0.41028 | 4.87E-13 | Moderate Negative Correlation |
| STING1 | -0.41053 | 4.7E-13 | Moderate Negative Correlation |
| ADGRA3 | -0.41082 | 4.51E-13 | Moderate Negative Correlation |
| CAPZB | -0.41109 | 4.34E-13 | Moderate Negative Correlation |
| TPST2 | -0.41135 | 4.18E-13 | Moderate Negative Correlation |
| SNRPB2 | -0.41146 | 4.12E-13 | Moderate Negative Correlation |
| KIF1C | -0.4115 | 4.09E-13 | Moderate Negative Correlation |
| SELENOW | -0.41178 | 3.93E-13 | Moderate Negative Correlation |
| DHX36 | -0.41194 | 3.84E-13 | Moderate Negative Correlation |
| TGOLN2 | -0.41227 | 3.67E-13 | Moderate Negative Correlation |
| MARCHF7 | -0.41245 | 3.57E-13 | Moderate Negative Correlation |
| UBXN1 | -0.41245 | 3.57E-13 | Moderate Negative Correlation |
| COMMD1 | -0.41278 | 3.41E-13 | Moderate Negative Correlation |
| VPS4B | -0.41297 | 3.31E-13 | Moderate Negative Correlation |
| CTNNA1 | -0.4132 | 3.21E-13 | Moderate Negative Correlation |
| COX5A | -0.41343 | 3.1E-13 | Moderate Negative Correlation |
| TIPRL | -0.41374 | 2.97E-13 | Moderate Negative Correlation |
| CTR9 | -0.41395 | 2.88E-13 | Moderate Negative Correlation |
| MAPK13 | -0.41403 | 2.85E-13 | Moderate Negative Correlation |
| SFT2D1 | -0.41471 | 2.58E-13 | Moderate Negative Correlation |
| CPNE8 | -0.41494 | 2.49E-13 | Moderate Negative Correlation |
| NCEH1 | -0.41547 | 2.31E-13 | Moderate Negative Correlation |
| NDUFAF3 | -0.41566 | 2.25E-13 | Moderate Negative Correlation |
| TOR1AIP2 | -0.41568 | 2.24E-13 | Moderate Negative Correlation |
| PPP2R1A | -0.41623 | 2.07E-13 | Moderate Negative Correlation |
| SDHC | -0.41649 | 1.99E-13 | Moderate Negative Correlation |

| | | | |
|---|---|---|---|
| TSPYL1 | -0.41674 | 1.92E-13 | Moderate Negative Correlation |
| EHD1 | -0.41705 | 1.84E-13 | Moderate Negative Correlation |
| OAT | -0.41748 | 1.72E-13 | Moderate Negative Correlation |
| GTF2A2 | -0.41791 | 1.62E-13 | Moderate Negative Correlation |
| RTL8C | -0.41819 | 1.55E-13 | Moderate Negative Correlation |
| ROCK1 | -0.41841 | 1.51E-13 | Moderate Negative Correlation |
| RFXANK | -0.41872 | 1.44E-13 | Moderate Negative Correlation |
| CAPNS2 | -0.41986 | 1.22E-13 | Moderate Negative Correlation |
| CD47 | -0.41992 | 1.21E-13 | Moderate Negative Correlation |
| SPINT2 | -0.42019 | 1.16E-13 | Moderate Negative Correlation |
| TRAPPC1 | -0.42027 | 1.15E-13 | Moderate Negative Correlation |
| PEDS1 | -0.42028 | 1.14E-13 | Moderate Negative Correlation |
| NAPG | -0.42034 | 1.13E-13 | Moderate Negative Correlation |
| VPS35 | -0.42049 | 1.11E-13 | Moderate Negative Correlation |
| WDR26 | -0.42127 | 9.89E-14 | Moderate Negative Correlation |
| C3 | -0.42127 | 9.89E-14 | Moderate Negative Correlation |
| SOD2 | -0.42174 | 9.23E-14 | Moderate Negative Correlation |
| EVPL | -0.42197 | 8.91E-14 | Moderate Negative Correlation |
| SP110 | -0.4221 | 8.75E-14 | Moderate Negative Correlation |
| EDF1 | -0.42311 | 7.53E-14 | Moderate Negative Correlation |
| TCEAL9 | -0.42372 | 6.88E-14 | Moderate Negative Correlation |
| PALMD | -0.42376 | 6.83E-14 | Moderate Negative Correlation |
| COX6B1 | -0.42398 | 6.62E-14 | Moderate Negative Correlation |
| ATP5F1C | -0.42438 | 6.23E-14 | Moderate Negative Correlation |
| ATP5F1A | -0.42483 | 5.83E-14 | Moderate Negative Correlation |
| MARVELD1 | -0.42505 | 5.64E-14 | Moderate Negative Correlation |
| TUSC3 | -0.42555 | 5.23E-14 | Moderate Negative Correlation |
| ANAPC5 | -0.42572 | 5.1E-14 | Moderate Negative Correlation |
| SPPL3 | -0.4258 | 5.04E-14 | Moderate Negative Correlation |
| MR1 | -0.42591 | 4.96E-14 | Moderate Negative Correlation |
| CD109 | -0.42636 | 4.63E-14 | Moderate Negative Correlation |
| KIAA1671 | -0.42668 | 4.41E-14 | Moderate Negative Correlation |
| ATP6V1F | -0.42675 | 4.37E-14 | Moderate Negative Correlation |
| HMCES | -0.42687 | 4.29E-14 | Moderate Negative Correlation |
| HYAL2 | -0.42694 | 4.25E-14 | Moderate Negative Correlation |
| CCDC9B | -0.42719 | 4.09E-14 | Moderate Negative Correlation |
| SERINC1 | -0.42731 | 4.02E-14 | Moderate Negative Correlation |
| PLCD3 | -0.42733 | 4E-14 | Moderate Negative Correlation |
| TNFSF10 | -0.42746 | 3.93E-14 | Moderate Negative Correlation |
| NDUFB3 | -0.42747 | 3.92E-14 | Moderate Negative Correlation |
| FAM3C | -0.42822 | 3.5E-14 | Moderate Negative Correlation |
| POLR3GL | -0.42898 | 3.12E-14 | Moderate Negative Correlation |
| SH3GLB1 | -0.42903 | 3.1E-14 | Moderate Negative Correlation |
| PMVK | -0.43018 | 2.6E-14 | Moderate Negative Correlation |
| AGR2 | -0.43033 | 2.54E-14 | Moderate Negative Correlation |
| ANKRD17 | -0.431 | 2.3E-14 | Moderate Negative Correlation |

| | | | |
|---|---|---|---|
| SLC9A3R2 | -0.43116 | 2.24E-14 | Moderate Negative Correlation |
| TUBA1A | -0.4312 | 2.23E-14 | Moderate Negative Correlation |
| KLHL5 | -0.43152 | 2.12E-14 | Moderate Negative Correlation |
| ABI1 | -0.43184 | 2.02E-14 | Moderate Negative Correlation |
| TJP1 | -0.432 | 1.97E-14 | Moderate Negative Correlation |
| GSN | -0.43225 | 1.9E-14 | Moderate Negative Correlation |
| LDAF1 | -0.43277 | 1.75E-14 | Moderate Negative Correlation |
| ELOB | -0.4328 | 1.74E-14 | Moderate Negative Correlation |
| ATRX | -0.43289 | 1.72E-14 | Moderate Negative Correlation |
| PHF20L1 | -0.43326 | 1.62E-14 | Moderate Negative Correlation |
| NDUFB4 | -0.434 | 1.45E-14 | Moderate Negative Correlation |
| ACSL3 | -0.43424 | 1.4E-14 | Moderate Negative Correlation |
| FKBP3 | -0.43476 | 1.29E-14 | Moderate Negative Correlation |
| TAOK3 | -0.43518 | 1.21E-14 | Moderate Negative Correlation |
| ARPC5L | -0.43518 | 1.21E-14 | Moderate Negative Correlation |
| UBE2M | -0.43526 | 1.19E-14 | Moderate Negative Correlation |
| FKBP14 | -0.43575 | 1.11E-14 | Moderate Negative Correlation |
| CALM2 | -0.43616 | 1.04E-14 | Moderate Negative Correlation |
| SMIM30 | -0.43656 | 9.74E-15 | Moderate Negative Correlation |
| SESN3 | -0.4367 | 9.53E-15 | Moderate Negative Correlation |
| COPS9 | -0.43677 | 9.44E-15 | Moderate Negative Correlation |
| BAIAP2 | -0.43712 | 8.93E-15 | Moderate Negative Correlation |
| AP1M2 | -0.43741 | 8.54E-15 | Moderate Negative Correlation |
| SERINC3 | -0.43804 | 7.75E-15 | Moderate Negative Correlation |
| CLINT1 | -0.43848 | 7.23E-15 | Moderate Negative Correlation |
| COX6C | -0.43867 | 7.01E-15 | Moderate Negative Correlation |
| PPIG | -0.43872 | 6.96E-15 | Moderate Negative Correlation |
| NME3 | -0.43908 | 6.58E-15 | Moderate Negative Correlation |
| SNX3 | -0.43921 | 6.44E-15 | Moderate Negative Correlation |
| ATP5MC1 | -0.43934 | 6.32E-15 | Moderate Negative Correlation |
| SH3D19 | -0.44001 | 5.69E-15 | Moderate Negative Correlation |
| UBE2H | -0.44005 | 5.65E-15 | Moderate Negative Correlation |
| UBE2V2 | -0.44191 | 4.21E-15 | Moderate Negative Correlation |
| TSPAN14 | -0.44249 | 3.85E-15 | Moderate Negative Correlation |
| ADD3 | -0.44278 | 3.67E-15 | Moderate Negative Correlation |
| CYB5R3 | -0.44287 | 3.62E-15 | Moderate Negative Correlation |
| UFL1 | -0.44296 | 3.57E-15 | Moderate Negative Correlation |
| CHCHD1 | -0.44312 | 3.48E-15 | Moderate Negative Correlation |
| NUMA1 | -0.44327 | 3.39E-15 | Moderate Negative Correlation |
| ARFIP1 | -0.44386 | 3.09E-15 | Moderate Negative Correlation |
| NDUFA11 | -0.44397 | 3.04E-15 | Moderate Negative Correlation |
| NCKAP1 | -0.44423 | 2.92E-15 | Moderate Negative Correlation |
| TIMP3 | -0.44427 | 2.9E-15 | Moderate Negative Correlation |
| ANO6 | -0.44479 | 2.66E-15 | Moderate Negative Correlation |
| ADGRF1 | -0.44483 | 2.65E-15 | Moderate Negative Correlation |
| TRIM38 | -0.44509 | 2.54E-15 | Moderate Negative Correlation |

| | | | |
|---|---|---|---|
| SLPI | -0.44528 | 2.47E-15 | Moderate Negative Correlation |
| KRT23 | -0.44544 | 2.41E-15 | Moderate Negative Correlation |
| TUBG1 | -0.44558 | 2.35E-15 | Moderate Negative Correlation |
| DDX52 | -0.44584 | 2.25E-15 | Moderate Negative Correlation |
| CAPS | -0.44587 | 2.25E-15 | Moderate Negative Correlation |
| CDC42EP5 | -0.44624 | 2.11E-15 | Moderate Negative Correlation |
| RWDD1 | -0.44628 | 2.1E-15 | Moderate Negative Correlation |
| FUNDC1 | -0.44664 | 1.98E-15 | Moderate Negative Correlation |
| ABLIM1 | -0.44689 | 1.91E-15 | Moderate Negative Correlation |
| LGALS1 | -0.44707 | 1.85E-15 | Moderate Negative Correlation |
| PPL | -0.4471 | 1.84E-15 | Moderate Negative Correlation |
| FSTL1 | -0.44775 | 1.66E-15 | Moderate Negative Correlation |
| IFNGR1 | -0.44861 | 1.44E-15 | Moderate Negative Correlation |
| ADAR | -0.4487 | 1.42E-15 | Moderate Negative Correlation |
| CHP1 | -0.44875 | 1.41E-15 | Moderate Negative Correlation |
| MAP9 | -0.44883 | 1.39E-15 | Moderate Negative Correlation |
| SCD5 | -0.44886 | 1.39E-15 | Moderate Negative Correlation |
| MTCH2 | -0.44887 | 1.38E-15 | Moderate Negative Correlation |
| LAMTOR2 | -0.44906 | 1.34E-15 | Moderate Negative Correlation |
| SARAF | -0.44942 | 1.27E-15 | Moderate Negative Correlation |
| TAF7 | -0.44977 | 1.2E-15 | Moderate Negative Correlation |
| NFU1 | -0.45014 | 1.13E-15 | Moderate Negative Correlation |
| SLU7 | -0.45143 | 9.13E-16 | Moderate Negative Correlation |
| PDZK1IP1 | -0.45158 | 8.91E-16 | Moderate Negative Correlation |
| CCNL2 | -0.45205 | 8.25E-16 | Moderate Negative Correlation |
| FURIN | -0.45217 | 8.09E-16 | Moderate Negative Correlation |
| SCP2 | -0.45233 | 7.89E-16 | Moderate Negative Correlation |
| ATP5PD | -0.45277 | 7.34E-16 | Moderate Negative Correlation |
| REEP5 | -0.45286 | 7.23E-16 | Moderate Negative Correlation |
| RDX | -0.45373 | 6.27E-16 | Moderate Negative Correlation |
| DYNLT3 | -0.45405 | 5.94E-16 | Moderate Negative Correlation |
| TPBG | -0.45411 | 5.89E-16 | Moderate Negative Correlation |
| ARMCX1 | -0.45475 | 5.3E-16 | Moderate Negative Correlation |
| DSC2 | -0.45529 | 4.84E-16 | Moderate Negative Correlation |
| ATP1A1 | -0.45555 | 4.64E-16 | Moderate Negative Correlation |
| ATP5F1B | -0.45628 | 4.11E-16 | Moderate Negative Correlation |
| PSMD7 | -0.45726 | 3.5E-16 | Moderate Negative Correlation |
| UTP3 | -0.45785 | 3.17E-16 | Moderate Negative Correlation |
| RAB3D | -0.45794 | 3.12E-16 | Moderate Negative Correlation |
| CDH1 | -0.45855 | 2.82E-16 | Moderate Negative Correlation |
| MORN2 | -0.45995 | 2.23E-16 | Moderate Negative Correlation |
| RAB13 | -0.46021 | 2.13E-16 | Moderate Negative Correlation |
| CKAP4 | -0.46036 | 2.08E-16 | Moderate Negative Correlation |
| HEXIM1 | -0.46038 | 2.07E-16 | Moderate Negative Correlation |
| GRN | -0.46093 | 1.89E-16 | Moderate Negative Correlation |
| MRPS36 | -0.46108 | 1.84E-16 | Moderate Negative Correlation |

| | | | |
|---|---|---|---|
| PRKAR1A | -0.46117 | 1.82E-16 | Moderate Negative Correlation |
| MAP3K13 | -0.4613 | 1.78E-16 | Moderate Negative Correlation |
| MOB1A | -0.46218 | 1.53E-16 | Moderate Negative Correlation |
| TMEM132A | -0.46247 | 1.46E-16 | Moderate Negative Correlation |
| ATP6AP2 | -0.46301 | 1.33E-16 | Moderate Negative Correlation |
| LEPROT | -0.46306 | 1.32E-16 | Moderate Negative Correlation |
| BID | -0.46474 | 9.91E-17 | Moderate Negative Correlation |
| SERPINB1 | -0.46486 | 9.71E-17 | Moderate Negative Correlation |
| MXD4 | -0.46499 | 9.5E-17 | Moderate Negative Correlation |
| HSBP1 | -0.46504 | 9.43E-17 | Moderate Negative Correlation |
| SDCBP2 | -0.46577 | 8.32E-17 | Moderate Negative Correlation |
| CLSTN1 | -0.46588 | 8.17E-17 | Moderate Negative Correlation |
| TGM2 | -0.46595 | 8.07E-17 | Moderate Negative Correlation |
| SRP14 | -0.46596 | 8.05E-17 | Moderate Negative Correlation |
| CAPN1 | -0.46628 | 7.63E-17 | Moderate Negative Correlation |
| PGM1 | -0.46645 | 7.41E-17 | Moderate Negative Correlation |
| TFG | -0.4669 | 6.85E-17 | Moderate Negative Correlation |
| NDUFA4 | -0.4674 | 6.29E-17 | Moderate Negative Correlation |
| CDC42EP3 | -0.46776 | 5.92E-17 | Moderate Negative Correlation |
| BOD1L1 | -0.46795 | 5.72E-17 | Moderate Negative Correlation |
| GID8 | -0.46838 | 5.32E-17 | Moderate Negative Correlation |
| LRRC41 | -0.46867 | 5.06E-17 | Moderate Negative Correlation |
| NDUFA1 | -0.46895 | 4.82E-17 | Moderate Negative Correlation |
| ZNF385A | -0.46904 | 4.74E-17 | Moderate Negative Correlation |
| SRPK2 | -0.46918 | 4.63E-17 | Moderate Negative Correlation |
| TRAM1 | -0.46932 | 4.52E-17 | Moderate Negative Correlation |
| ATP6V1D | -0.46969 | 4.24E-17 | Moderate Negative Correlation |
| CRYAB | -0.47038 | 3.76E-17 | Moderate Negative Correlation |
| ETFA | -0.47046 | 3.71E-17 | Moderate Negative Correlation |
| MALL | -0.4705 | 3.68E-17 | Moderate Negative Correlation |
| MGLL | -0.47065 | 3.59E-17 | Moderate Negative Correlation |
| MORF4L1 | -0.47076 | 3.52E-17 | Moderate Negative Correlation |
| EVA1A | -0.47176 | 2.96E-17 | Moderate Negative Correlation |
| UBXN4 | -0.47192 | 2.87E-17 | Moderate Negative Correlation |
| SVIL | -0.47209 | 2.79E-17 | Moderate Negative Correlation |
| SEC61B | -0.47213 | 2.77E-17 | Moderate Negative Correlation |
| FEZ2 | -0.47341 | 2.21E-17 | Moderate Negative Correlation |
| NDUFB10 | -0.4735 | 2.18E-17 | Moderate Negative Correlation |
| ARF3 | -0.47351 | 2.18E-17 | Moderate Negative Correlation |
| ISG20 | -0.474 | 2E-17 | Moderate Negative Correlation |
| ELF1 | -0.47443 | 1.85E-17 | Moderate Negative Correlation |
| DPY30 | -0.47485 | 1.72E-17 | Moderate Negative Correlation |
| PPIC | -0.47494 | 1.69E-17 | Moderate Negative Correlation |
| PLEKHG5 | -0.47572 | 1.48E-17 | Moderate Negative Correlation |
| PHACTR2 | -0.47625 | 1.34E-17 | Moderate Negative Correlation |
| PSMB5 | -0.47634 | 1.32E-17 | Moderate Negative Correlation |

| | | | |
|---|---|---|---|
| MIB2 | -0.47739 | 1.1E-17 | Moderate Negative Correlation |
| GSTK1 | -0.4776 | 1.06E-17 | Moderate Negative Correlation |
| RAB7A | -0.47802 | 9.81E-18 | Moderate Negative Correlation |
| BASP1 | -0.47818 | 9.54E-18 | Moderate Negative Correlation |
| NDUFAB1 | -0.47821 | 9.49E-18 | Moderate Negative Correlation |
| OSTF1 | -0.47999 | 6.9E-18 | Moderate Negative Correlation |
| OPTN | -0.48065 | 6.13E-18 | Moderate Negative Correlation |
| EIF4G1 | -0.48067 | 6.11E-18 | Moderate Negative Correlation |
| ECE1 | -0.48111 | 5.64E-18 | Moderate Negative Correlation |
| NOP10 | -0.48123 | 5.53E-18 | Moderate Negative Correlation |
| NDUFV3 | -0.48183 | 4.96E-18 | Moderate Negative Correlation |
| TMEM179B | -0.48197 | 4.83E-18 | Moderate Negative Correlation |
| LRRFIP2 | -0.4823 | 4.56E-18 | Moderate Negative Correlation |
| RAP1A | -0.48249 | 4.4E-18 | Moderate Negative Correlation |
| ATP5PO | -0.48393 | 3.39E-18 | Moderate Negative Correlation |
| CCDC50 | -0.48415 | 3.26E-18 | Moderate Negative Correlation |
| ISG15 | -0.48437 | 3.13E-18 | Moderate Negative Correlation |
| ANKRD22 | -0.48443 | 3.1E-18 | Moderate Negative Correlation |
| CD81 | -0.48459 | 3.01E-18 | Moderate Negative Correlation |
| LACTB | -0.48494 | 2.82E-18 | Moderate Negative Correlation |
| UXS1 | -0.485 | 2.79E-18 | Moderate Negative Correlation |
| ATP5MF | -0.48543 | 2.58E-18 | Moderate Negative Correlation |
| BBLN | -0.48545 | 2.57E-18 | Moderate Negative Correlation |
| PPP2CB | -0.48557 | 2.52E-18 | Moderate Negative Correlation |
| MRPL47 | -0.48596 | 2.35E-18 | Moderate Negative Correlation |
| PTPRE | -0.48635 | 2.18E-18 | Moderate Negative Correlation |
| KTN1 | -0.48657 | 2.1E-18 | Moderate Negative Correlation |
| PTPN1 | -0.48662 | 2.08E-18 | Moderate Negative Correlation |
| KIAA1522 | -0.48694 | 1.96E-18 | Moderate Negative Correlation |
| IARS2 | -0.49013 | 1.09E-18 | Moderate Negative Correlation |
| CTTN | -0.49053 | 1.01E-18 | Moderate Negative Correlation |
| NR2F6 | -0.4906 | 9.98E-19 | Moderate Negative Correlation |
| ATP5MK | -0.49089 | 9.46E-19 | Moderate Negative Correlation |
| ZBED1 | -0.49096 | 9.34E-19 | Moderate Negative Correlation |
| COX7C | -0.4918 | 7.98E-19 | Moderate Negative Correlation |
| SERF2 | -0.49351 | 5.81E-19 | Moderate Negative Correlation |
| PSMA4 | -0.49379 | 5.51E-19 | Moderate Negative Correlation |
| HIF1A | -0.49477 | 4.58E-19 | Moderate Negative Correlation |
| CRABP2 | -0.49499 | 4.4E-19 | Moderate Negative Correlation |
| KDM5B | -0.495 | 4.38E-19 | Moderate Negative Correlation |
| PTAFR | -0.49533 | 4.12E-19 | Moderate Negative Correlation |
| MMADHC | -0.49569 | 3.86E-19 | Moderate Negative Correlation |
| ZBTB38 | -0.4957 | 3.85E-19 | Moderate Negative Correlation |
| C5orf24 | -0.49628 | 3.44E-19 | Moderate Negative Correlation |
| LYPD3 | -0.49653 | 3.29E-19 | Moderate Negative Correlation |
| DHRS7 | -0.49675 | 3.15E-19 | Moderate Negative Correlation |

| | | | |
|---|---|---|---|
| GTF2I | -0.49778 | 2.59E-19 | Moderate Negative Correlation |
| RTF1 | -0.49877 | 2.15E-19 | Moderate Negative Correlation |
| GOPC | -0.49945 | 1.89E-19 | Moderate Negative Correlation |
| IRF2BP2 | -0.50038 | 1.58E-19 | Strong Negative Correlation |
| MRPL14 | -0.50077 | 1.47E-19 | Strong Negative Correlation |
| NDUFB11 | -0.50109 | 1.38E-19 | Strong Negative Correlation |
| MLF2 | -0.50197 | 1.17E-19 | Strong Negative Correlation |
| ZFAND6 | -0.50242 | 1.07E-19 | Strong Negative Correlation |
| SCEL | -0.50265 | 1.02E-19 | Strong Negative Correlation |
| CTSB | -0.50273 | 1.01E-19 | Strong Negative Correlation |
| LGALS7B | -0.50389 | 8.04E-20 | Strong Negative Correlation |
| H3-3A | -0.50403 | 7.82E-20 | Strong Negative Correlation |
| PLIN3 | -0.50418 | 7.59E-20 | Strong Negative Correlation |
| RAB27B | -0.50494 | 6.55E-20 | Strong Negative Correlation |
| MRPL33 | -0.50556 | 5.81E-20 | Strong Negative Correlation |
| PXDN | -0.50569 | 5.67E-20 | Strong Negative Correlation |
| BRK1 | -0.50606 | 5.27E-20 | Strong Negative Correlation |
| POMP | -0.5064 | 4.93E-20 | Strong Negative Correlation |
| FAM177A1 | -0.50722 | 4.2E-20 | Strong Negative Correlation |
| MRPS21 | -0.50761 | 3.89E-20 | Strong Negative Correlation |
| MRPL21 | -0.51069 | 2.12E-20 | Strong Negative Correlation |
| CAMLG | -0.51076 | 2.09E-20 | Strong Negative Correlation |
| YWHAG | -0.51089 | 2.04E-20 | Strong Negative Correlation |
| C11orf58 | -0.51271 | 1.42E-20 | Strong Negative Correlation |
| IL1RN | -0.5145 | 9.92E-21 | Strong Negative Correlation |
| RAB1A | -0.51451 | 9.89E-21 | Strong Negative Correlation |
| CLIC3 | -0.51477 | 9.4E-21 | Strong Negative Correlation |
| CHMP1B | -0.51532 | 8.41E-21 | Strong Negative Correlation |
| BCL2L1 | -0.51614 | 7.14E-21 | Strong Negative Correlation |
| FUOM | -0.51648 | 6.67E-21 | Strong Negative Correlation |
| PON2 | -0.51913 | 3.89E-21 | Strong Negative Correlation |
| ATP5IF1 | -0.51922 | 3.82E-21 | Strong Negative Correlation |
| BEX3 | -0.52001 | 3.25E-21 | Strong Negative Correlation |
| RBP1 | -0.52135 | 2.47E-21 | Strong Negative Correlation |
| SF3B6 | -0.52175 | 2.28E-21 | Strong Negative Correlation |
| NDUFB2 | -0.52219 | 2.08E-21 | Strong Negative Correlation |
| ETFB | -0.52248 | 1.96E-21 | Strong Negative Correlation |
| ITGAV | -0.52363 | 1.55E-21 | Strong Negative Correlation |
| PBDC1 | -0.52381 | 1.49E-21 | Strong Negative Correlation |
| TSTD1 | -0.52401 | 1.43E-21 | Strong Negative Correlation |
| PRRG4 | -0.52432 | 1.34E-21 | Strong Negative Correlation |
| ITGB6 | -0.52438 | 1.32E-21 | Strong Negative Correlation |
| CDKN1A | -0.529 | 5.03E-22 | Strong Negative Correlation |
| TNKS1BP1 | -0.53006 | 4.03E-22 | Strong Negative Correlation |
| MSRB2 | -0.5303 | 3.83E-22 | Strong Negative Correlation |
| RAB2A | -0.53045 | 3.71E-22 | Strong Negative Correlation |

| | | | |
|---|---|---|---|
| RAB11A | -0.5313 | 3.1E-22 | Strong Negative Correlation |
| GNB1 | -0.53183 | 2.77E-22 | Strong Negative Correlation |
| KIF5B | -0.53254 | 2.38E-22 | Strong Negative Correlation |
| RALBP1 | -0.5331 | 2.11E-22 | Strong Negative Correlation |
| FAM210B | -0.53406 | 1.72E-22 | Strong Negative Correlation |
| RNF7 | -0.53473 | 1.49E-22 | Strong Negative Correlation |
| LAMTOR5 | -0.53489 | 1.44E-22 | Strong Negative Correlation |
| CMAS | -0.53572 | 1.21E-22 | Strong Negative Correlation |
| S100A9 | -0.53623 | 1.08E-22 | Strong Negative Correlation |
| SAA1 | -0.53626 | 1.07E-22 | Strong Negative Correlation |
| HOMER3 | -0.53682 | 9.52E-23 | Strong Negative Correlation |
| ALOX15B | -0.53696 | 9.23E-23 | Strong Negative Correlation |
| ECHS1 | -0.537 | 9.16E-23 | Strong Negative Correlation |
| UBE2Z | -0.53713 | 8.9E-23 | Strong Negative Correlation |
| ATP5PF | -0.53721 | 8.75E-23 | Strong Negative Correlation |
| FDPS | -0.53723 | 8.7E-23 | Strong Negative Correlation |
| COX7A2 | -0.53746 | 8.29E-23 | Strong Negative Correlation |
| AK3 | -0.53758 | 8.07E-23 | Strong Negative Correlation |
| ATP5MG | -0.53761 | 8.02E-23 | Strong Negative Correlation |
| DHRS1 | -0.5397 | 5.09E-23 | Strong Negative Correlation |
| EIF4A2 | -0.53975 | 5.04E-23 | Strong Negative Correlation |
| DSG2 | -0.54028 | 4.49E-23 | Strong Negative Correlation |
| SDCBP | -0.54033 | 4.44E-23 | Strong Negative Correlation |
| TMEM50A | -0.54108 | 3.77E-23 | Strong Negative Correlation |
| BBX | -0.54468 | 1.7E-23 | Strong Negative Correlation |
| CST3 | -0.54504 | 1.57E-23 | Strong Negative Correlation |
| CAST | -0.54633 | 1.18E-23 | Strong Negative Correlation |
| CMTM6 | -0.54874 | 6.88E-24 | Strong Negative Correlation |
| CD82 | -0.55283 | 2.73E-24 | Strong Negative Correlation |
| TOLLIP | -0.55399 | 2.09E-24 | Strong Negative Correlation |
| UBL5 | -0.55577 | 1.39E-24 | Strong Negative Correlation |
| NBDY | -0.55645 | 1.19E-24 | Strong Negative Correlation |
| TSC22D1 | -0.55651 | 1.17E-24 | Strong Negative Correlation |
| IFI27L2 | -0.55764 | 9.02E-25 | Strong Negative Correlation |
| TMEM134 | -0.55965 | 5.66E-25 | Strong Negative Correlation |
| IRF6 | -0.56052 | 4.62E-25 | Strong Negative Correlation |
| GBP3 | -0.56147 | 3.7E-25 | Strong Negative Correlation |
| NDUFA12 | -0.56288 | 2.66E-25 | Strong Negative Correlation |
| KDELR1 | -0.56502 | 1.6E-25 | Strong Negative Correlation |
| MGST2 | -0.56799 | 7.87E-26 | Strong Negative Correlation |
| PLAUR | -0.56884 | 6.41E-26 | Strong Negative Correlation |
| DAPP1 | -0.57078 | 4.02E-26 | Strong Negative Correlation |
| DYNC1I2 | -0.57281 | 2.45E-26 | Strong Negative Correlation |
| POLR2A | -0.57335 | 2.15E-26 | Strong Negative Correlation |
| TP53I3 | -0.57398 | 1.84E-26 | Strong Negative Correlation |
| PLEKHA1 | -0.57539 | 1.3E-26 | Strong Negative Correlation |

| | | | |
|---|---|---|---|
| STAP2 | -0.57575 | 1.19E-26 | Strong Negative Correlation |
| ACOT7 | -0.57631 | 1.04E-26 | Strong Negative Correlation |
| NDUFA5 | -0.57632 | 1.04E-26 | Strong Negative Correlation |
| PRKCI | -0.57848 | 6.08E-27 | Strong Negative Correlation |
| TAPBP | -0.57903 | 5.3E-27 | Strong Negative Correlation |
| EID1 | -0.58112 | 3.14E-27 | Strong Negative Correlation |
| SCOC | -0.58157 | 2.8E-27 | Strong Negative Correlation |
| SERPINB7 | -0.58232 | 2.32E-27 | Strong Negative Correlation |
| SNCG | -0.58347 | 1.74E-27 | Strong Negative Correlation |
| ATP1B1 | -0.58599 | 9.17E-28 | Strong Negative Correlation |
| PSME2 | -0.58885 | 4.41E-28 | Strong Negative Correlation |
| KRT6B | -0.59116 | 2.42E-28 | Strong Negative Correlation |
| TMEM79 | -0.59186 | 2.02E-28 | Strong Negative Correlation |
| ARPC3 | -0.59277 | 1.59E-28 | Strong Negative Correlation |
| CLDN1 | -0.59361 | 1.28E-28 | Strong Negative Correlation |
| CCDC25 | -0.59405 | 1.14E-28 | Strong Negative Correlation |
| CA12 | -0.59429 | 1.07E-28 | Strong Negative Correlation |
| DECR1 | -0.59493 | 9.05E-29 | Strong Negative Correlation |
| ATP6V1G1 | -0.59514 | 8.55E-29 | Strong Negative Correlation |
| PDCD4 | -0.59761 | 4.46E-29 | Strong Negative Correlation |
| PHLDA2 | -0.59862 | 3.4E-29 | Strong Negative Correlation |
| KRT8 | -0.59981 | 2.48E-29 | Strong Negative Correlation |
| PDLIM4 | -0.59992 | 2.4E-29 | Strong Negative Correlation |
| TSPAN1 | -0.60552 | 5.28E-30 | Strong Negative Correlation |
| TM4SF1 | -0.60814 | 2.57E-30 | Strong Negative Correlation |
| ATP5MC3 | -0.60868 | 2.22E-30 | Strong Negative Correlation |
| AQP3 | -0.60994 | 1.57E-30 | Strong Negative Correlation |
| CLTB | -0.613 | 6.67E-31 | Strong Negative Correlation |
| HEBP2 | -0.61356 | 5.7E-31 | Strong Negative Correlation |
| PYCARD | -0.61418 | 4.79E-31 | Strong Negative Correlation |
| UQCRB | -0.61492 | 3.88E-31 | Strong Negative Correlation |
| ATOX1 | -0.61858 | 1.37E-31 | Strong Negative Correlation |
| CDCP1 | -0.61868 | 1.33E-31 | Strong Negative Correlation |
| TCEAL8 | -0.61956 | 1.04E-31 | Strong Negative Correlation |
| MVP | -0.62742 | 1.05E-32 | Strong Negative Correlation |
| CDA | -0.62764 | 9.82E-33 | Strong Negative Correlation |
| LGALS3 | -0.6301 | 4.73E-33 | Strong Negative Correlation |
| S100A13 | -0.63302 | 1.97E-33 | Strong Negative Correlation |
| CHMP5 | -0.63719 | 5.56E-34 | Strong Negative Correlation |
| OST4 | -0.63774 | 4.7E-34 | Strong Negative Correlation |
| SNX6 | -0.63928 | 2.92E-34 | Strong Negative Correlation |
| CSTB | -0.63965 | 2.61E-34 | Strong Negative Correlation |
| CASP4 | -0.63994 | 2.38E-34 | Strong Negative Correlation |
| DDR1 | -0.64476 | 5.31E-35 | Strong Negative Correlation |
| CD9 | -0.65193 | 5.37E-36 | Strong Negative Correlation |
| CD24 | -0.65589 | 1.48E-36 | Strong Negative Correlation |

| | | | |
|---|---|---|---|
| IL32 | -0.65951 | 4.46E-37 | Strong Negative Correlation |
| FXYD3 | -0.66655 | 4.15E-38 | Strong Negative Correlation |
| COMT | -0.67003 | 1.25E-38 | Strong Negative Correlation |
| VAMP8 | -0.67012 | 1.21E-38 | Strong Negative Correlation |
| RAB25 | -0.68531 | 5.32E-41 | Strong Negative Correlation |
| ARL6IP5 | -0.69301 | 2.98E-42 | Strong Negative Correlation |
| C4orf3 | -0.72369 | 1.17E-47 | Very Strong Negative Correlation |
| HSPB1 | -0.75236 | 2.07E-53 | Very Strong Negative Correlation |
| SAT1 | -0.75563 | 4.06E-54 | Very Strong Negative Correlation |
| TACSTD2 | -0.76549 | 2.55E-56 | Very Strong Negative Correlation |

**Table Legend**

| Column Name |
| --- |
| Gene name |
| R |
| P |
| Correlation Strength |

| Description |
|---|
| Official gene symbol standardized by HGNC (HUGO Gene Nomenclature Committee) |
| Pearson correlation coefficient: indicating relationship strength between gene expression among selected cells and selecte |
| P-value: statistical significance of correlation (values < 0.05 typically considered statistically significant) |
| Strength of correlation based on absolute R-value (ignores direction). Categories: **Strong** (\|R\| ≥ 0.7), **Moderate** (0.3 ≤ \|R\| < |
