## Supplemental Table 7 for "Human Cytomegalovirus Infection of Primary Human Oral Keratinocytes Induces Intermediate Keratinocyte Differentiation and an Altered Innate Immune Response"

| Term | ES | NES | NOM p-val | FDR q-val | FWER p-va | Tag % |
|---|---|---|---|---|---|---|
| **Oxidative Phosphorylation** | -0.66895 | -2.12298 | 0 | 0 | 0 | 116/152 |
| Spermatogenesis | 0.380574 | 1.915681 | 0 | 0.011321 | 0.003 | 32/81 |
| Protein Secretion | -0.62291 | -1.89216 | 0 | 0 | 0 | 50/74 |
| Adipogenesis | -0.5535 | -1.74675 | 0 | 0.000313 | 0.001 | 88/145 |
| E2F Targets | 0.365383 | 1.69588 | 0 | 0.018868 | 0.01 | 27/69 |
| Myc Targets V1 | -0.61494 | -1.68031 | 0.001068 | 0.003525 | 0.015 | 18/28 |
| Mitotic Spindle | -0.53165 | -1.66264 | 0 | 0.003384 | 0.018 | 62/130 |
| **Interferon Alpha Response** | -0.54428 | -1.58878 | 0.001016 | 0.009556 | 0.059 | 30/51 |
| **Interferon Gamma Response** | -0.51023 | -1.58557 | 0 | 0.008191 | 0.059 | 59/116 |
| **TNF-alpha Signaling via NF-kB** | -0.50067 | -1.56451 | 0 | 0.009634 | 0.079 | 60/107 |
| TGF-beta Signaling | -0.56082 | -1.55771 | 0.001046 | 0.009608 | 0.089 | 24/32 |
| DNA Repair | -0.49726 | -1.52509 | 0.001006 | 0.014851 | 0.148 | 48/83 |
| Fatty Acid Metabolism | -0.48765 | -1.51928 | 0 | 0.014184 | 0.155 | 39/94 |
| Apoptosis | -0.48627 | -1.50817 | 0 | 0.014804 | 0.172 | 41/101 |
| **Complement** | -0.47145 | -1.4871 | 0 | 0.018365 | 0.225 | 49/124 |
| Coagulation | -0.47665 | -1.4664 | 0.005025 | 0.022558 | 0.29 | 29/79 |
| Androgen Response | -0.48977 | -1.44386 | 0.010215 | 0.027321 | 0.35 | 26/55 |
| p53 Pathway | -0.46166 | -1.43986 | 0.002006 | 0.027082 | 0.361 | 55/104 |
| Estrogen Response Late | -0.44216 | -1.39392 | 0.001 | 0.044951 | 0.553 | 32/143 |
| heme Metabolism | -0.43267 | -1.38735 | 0 | 0.045795 | 0.586 | 52/151 |
| Pperoxisome | -0.45375 | -1.38699 | 0.024267 | 0.043583 | 0.587 | 38/73 |
| Reactive Oxygen Species Pathway | -0.46816 | -1.30531 | 0.090147 | 0.104661 | 0.898 | 15/33 |
| IL-6/JAK/STAT3 Signaling | -0.42808 | -1.26924 | 0.102041 | 0.145689 | 0.959 | 28/54 |
| Xenobiotic Metabolism | -0.39376 | -1.23421 | 0.063063 | 0.198538 | 0.987 | 44/119 |
| KRAS Signaling Dn | 0.246908 | 1.227442 | 0 | 0.179874 | 0.135 | 45/129 |
| mTORC1 Signaling | -0.38905 | -1.21613 | 0.101508 | 0.227054 | 0.994 | 26/93 |
| Cholesterol Homeostasis | -0.42402 | -1.20666 | 0.184874 | 0.237762 | 0.996 | 11/37 |
| PI3K/AKT/mTOR Signaling | -0.38759 | -1.17787 | 0.178894 | 0.296566 | 1 | 30/73 |
| Inflammatory Response | -0.37408 | -1.17523 | 0.113113 | 0.290871 | 1 | 35/129 |
| Apical Junction | -0.37064 | -1.16687 | 0.14343 | 0.299071 | 1 | 30/128 |
| Estrogen Response Early | -0.36894 | -1.16639 | 0.112112 | 0.289766 | 1 | 45/147 |
| Apical Surface | -0.40895 | -1.13851 | 0.295699 | 0.348421 | 1 | 8/30 |
| Hedgehog Signaling | 0.276276 | 1.125666 | 0.174603 | 0.229245 | 0.22 | 10/24 |
| Glycolysis | -0.35508 | -1.11189 | 0.218218 | 0.411562 | 1 | 38/124 |
| Notch Signaling | -0.42638 | -1.09929 | 0.343646 | 0.435065 | 1 | 8/20 |
| Epithelial Mesenchymal Transitior | -0.35227 | -1.09905 | 0.269 | 0.422174 | 1 | 18/110 |
| UV Response Dn | -0.34821 | -1.07593 | 0.349749 | 0.475119 | 1 | 29/93 |
| Bile Acid Metabolism | -0.34433 | -1.05615 | 0.381288 | 0.518922 | 1 | 30/81 |
| Angiogenesis | -0.40801 | -1.03562 | 0.45618 | 0.564788 | 1 | 7/16 |
| Unfolded Protein Response | -0.34485 | -1.02485 | 0.458883 | 0.580744 | 1 | 21/59 |
| Allograft Rejection | -0.33042 | -1.01886 | 0.451904 | 0.583034 | 1 | 47/108 |
| G2-M Checkpoint | -0.32083 | -0.97456 | 0.554326 | 0.684266 | 1 | 30/79 |
| UV Response Up | -0.30973 | -0.96728 | 0.585 | 0.686869 | 1 | 31/110 |

| | | | | | | |
|---|---|---|---|---|---|---|
| IL-2/STAT5 Signaling | -0.30669 | -0.96538 | 0.597598 | 0.674162 | 1 | 36/134 |
| KRAS Signaling Up | -0.29267 | -0.92741 | 0.692 | 0.745499 | 1 | 45/138 |
| Myogenesis | -0.29418 | -0.92342 | 0.700701 | 0.736164 | 1 | 41/134 |
| Myc Targets V2 | -0.44189 | -0.90722 | 0.618454 | 0.751351 | 1 | 4/7 |
| Hypoxia | -0.28644 | -0.89123 | 0.767768 | 0.763541 | 1 | 45/124 |
| Wnt-beta Catenin Signaling | -0.26174 | -0.71789 | 0.874598 | 0.953816 | 1 | 16/28 |
| Pancreas Beta Cells | -0.21984 | -0.57864 | 0.953564 | 0.986248 | 1 | 2/24 |

| Gene % |
| --- |
| 25.91% |
| 16.43% |
| 24.64% |
| 30.47% |
| 13.29% |
| 18.64% |
| 21.09% |
| 28.71% |
| 28.71% |
| 30.18% |
| 34.20% |
| 30.60% |
| 22.20% |
| 21.74% |
| 26.54% |
| 28.14% |
| 27.76% |
| 26.66% |
| 7.89% |
| 22.20% |
| 37.44% |
| 18.88% |
| 33.77% |
| 26.70% |
| 16.66% |
| 9.81% |
| 11.69% |
| 22.25% |
| 19.93% |
| 13.67% |
| 19.71% |
| 14.43% |
| 12.45% |
| 19.48% |
| 27.18% |
| 7.57% |
| 22.47% |
| 31.09% |
| 28.14% |
| 19.18% |
| 38.24% |
| 28.97% |
| 18.72% |

23.27%
27.61%
27.86%
43.39%
30.71%
48.00%
6.26%

| Lead_genes |
| --- |
| UQCRB;ATP5MC3;ATP6V1G1;DECR1;ATP1B1;NDUFA5;ATP5MG;COX7A2;ATP5PF;ECHS1;ETFB;NDUFB2;COX7C;AT |
| YBX2;SHE;RFC4;NEFH;CLGN;SNAP91;ADCYAP1;SEPTIN4;RPL39L;CHRM4;LDHC;ACE;CNIH2;PCSK1N;CDK1;JAM3;PC |
| RAB2A;ATP1A1;ARFIP1;NAPG;VPS4B;LAMP2;AP3B1;RAB9A;IGF2R;COPB2;RAB5A;TMED2;M6PR;RAB14;PAM;ERG |
| PDCD4;DECR1;SNCG;NDUFA5;ECHS1;ETFB;YWHAG;DHRS7;ATP5PO;NDUFAB1;MGLL;PGM1;REEP5;SCP2;MTCH2; |
| MTHFD2;UBE2S;TK1;TCF19;CENPM;CDKN2C;GINS4;UBE2T;PSIP1;DSCC1;ASF1B;RPA2;RNASEH2A;ORC6;BRCA1;AT |
| PSMA4;NDUFAB1;PSMD7;COX5A;SNRPB2;VDAC3;PSMD1;SNRPA1;UBE2E1;ACP1;STARD7;PSMB3;PSMD3;PRPF31 |
| RALBP1;KIF5B;CTTN;NUMA1;GSN;ABI1;ROCK1;CAPZB;CLASP1;TLK1;NCK1;PAFAH1B1;CLIP1;ARFIP2;CDC27;DLG1; |
| PSME2;ISG15;ELF1;ISG20;ADAR;SP110;CD47;SAMD9;UBE2L6;PSMA3;NUB1;PARP14;HELZ2;TAP1;GBP2;IRF2;TRIM |
| VAMP8;CASP4;MVP;PSME2;TAPBP;CDKN1A;HIF1A;PTPN1;ISG15;ISG20;ADAR;TNFSF10;SP110;SOD2;NFKBIA;MYD |
| SAT1;PHLDA2;PLAUR;TSC22D1;CDKN1A;PTPRE;SOD2;EHD1;NFKBIA;PNRC1;KYNU;TANK;TNFAIP8;RCAN1;BIRC3;T |
| CDH1;FURIN;TJP1;PPP1CA;HDAC1;BMPR2;ARID4B;SMAD1;SPTBN1;MAP3K7;SMAD3;SKIL;SMURF1;BMP2;WWTR |
| CDA;POLR2A;SDCBP;AK3;NME3;POLR3GL;EDF1;GTF2A2;SUPT5H;RALA;TMED2;PNP;CETN2;POLR2E;POLR2J;GTF2 |
| DECR1;ECHS1;MGLL;LGALS1;SERINC1;SDHC;ECH1;SUCLA2;RDH11;GRHPR;UROS;HSD17B10;ACAA1;UBE2L6;MDH |
| SAT1;HSPB1;CASP4;LGALS3;PDCD4;CDKN1A;BCL2L1;ISG20;BID;IFNGR1;TIMP3;GSN;TNFSF10;SOD2;ROCK1;EMP1 |
| CASP4;S100A13;LGALS3;CDA;ATOX1;PLAUR;S100A9;CTSB;PPP2CB;C3;EHD1;LAMP2;KYNU;STX4;RHOG;LIPA;CASP |
| CD9;S100A13;MSRB2;CTSB;FURIN;TIMP3;GSN;C3;LAMP2;BMP1;APOC1;GNB2;DUSP14;SERPINA1;CTSO;KLK8;CRI |
| SAT1;KRT8;TSC22D1;TMEM50A;ITGAV;ACSL3;NCOA4;TNFAIP8;SLC26A2;PIAS1;H1-0;ABHD2;SRP19;RPS6KA3;LMA |
| SAT1;VAMP8;TM4SF1;TSC22D1;CD82;ALOX15B;CDKN1A;PTPRE;CD81;MXD4;HEXIM1;BAIAP2;IP6K2;PHLDA3;TOB |
| CD9;PDCD4;CA12;S100A9;ETFB;CLIC3;ISG20;CDH1;TPBG;DYNLT3;ADD3;COX6C;AGR2;MAPK13;SLC9A3R1;SULT2B |
| AQP3;CAST;SDCBP;CTSB;OPTN;PDZK1IP1;C3;LAMP2;CIR1;NCOA4;SLC30A1;ADD1;UROS;ARL2BP;IGSF3;BNIP3L;TM |
| MVP;FDPS;CRABP2;GSTK1;SCP2;SOD2;VPS4B;ECH1;SULT2B1;RDH11;PRDX5;ACAA1;ECI2;NUDT19;SOD1;PEX11B; |
| ATOX1;LAMTOR5;NDUFB4;SOD2;NDUFS2;OXSR1;MBP;SOD1;PTPA;NDUFA6;GLRX2;FTL;MSRA;CAT;GLRX |
| CD9;PTPN1;IFNGR1;MYD88;IFNAR1;TNFRSF1A;IL1R1;STAT3;GRB2;EBI3;TNF;IRF1;LTBR;PDGFC;LEPR;STAT1;TGFB1 |
| COMT;CDA;DHRS1;DHRS7;SPINT2;ECH1;KYNU;PINK1;CBR1;PTGES;DDAH2;CROT;PTGR1;PPARD;CASP6;TNFRSF1A |
| CALCB;YBX2;EFHD1;SLC38A3;IDUA;SNCB;YPEL1;SLC30A3;FGF16;CNTFR;KCNN1;ARHGDIG;SHOX2;ADRA2C;FGF22 |
| CD9;DAPP1;CDKN1A;RAB1A;PSMA4;PSMB5;ATP6V1D;PGM1;TUBG1;ADD3;ATP5MC1;ARPC5L;ACSL3;SLC9A3R1;M |
| CD9;LGALS3;PLAUR;FDPS;PMVK;ECH1;PNRC1;TM7SF2;FBXO6;NSDHL;MAL2 |
| ARPC3;DAPP1;CDKN1A;MKNK2;PPP1CA;MYD88;NCK1;PRKAR2A;RIT1;MAP2K3;TNFRSF1A;MAP3K7;CAB39L;RPS6 |
| TAPBP;PLAUR;CD82;CDKN1A;PTAFR;HIF1A;PTPRE;TPBG;ABI1;TNFSF10;NFKBIA;SLC31A2;ITGB8;IFNAR1;CCL20;RN |
| ALOX15B;CDH1;TUBG1;BAIAP2;TJP1;MAPK13;CTNNA1;RSU1;NECTIN4;CD99;LAYN;CNTN1;MPZL1;BMP1;DLG1;ST |
| AQP3;KRT8;CA12;CLIC3;ELF1;SVIL;TGM2;TPBG;DYNLT3;ABLIM1;ADD3;SLC9A3R1;SULT2B1;KLK10;TOB1;OPN3;BA |
| HSPB1;PLAUR;LYPD3;RHCG;ADIPOR2;CX3CL1;AFAP1L2;LYN |
| VLDLR;CNTFR;ACHE;HEY2;CRMP1;NKX6-1;AMOT;GLI1;SHH;PTCH1 |
| AK3;PPP2CB;NDUFV3;ISG20;DSC2;TPBG;SDHC;HOMER1;COPB2;MDH2;HDLBP;BPNT1;QSOX1;AGL;CHPF;ME2;PA |
| PPARD;DTX2;APH1A;NOTCH2;MAML2;DTX4;NOTCH3;TCF7L2 |
| SAT1;IL32;PDLIM4;PLAUR;ITGAV;BASP1;TGM2;FSTL1;LGALS1;TIMP3;PFN2;LOXL1;FSTL3;BMP1;FUCA1;QSOX1;CC |
| MGLL;ADD3;ATRX;TJP1;ATP2B4;MGMT;DLG1;PIAS3;CITED2;SIPA1L1;DAB2;NIPBL;SMAD3;ADGRL2;ERBB2;MAPK1 |
| RBP1;OPTN;GSTK1;SCP2;BBOX1;SULT2B1;PRDX5;FDXR;CROT;SOD1;LONP2;ABCA1;PEX19;SLC22A18;PHYH;CAT;N |
| ITGAV;FSTL1;LRPAP1;PTK2;CXCL6;S100A4;THBD |
| EIF4A2;KIF5B;EIF4G1;FKBP14;BAG3;YIF1A;TUBB2A;PAIP1;SPCS1;CNOT2;DCTN1;EDEM1;CNOT4;LSM1;CCL2;EXOC |
| TAPBP;HIF1A;IFNGR1;ABI1;CD47;EIF4G3;NCK1;HLA-E;TAP1;MAP3K7;GBP2;UBE2D1;CCL2;CSK;F2R;IL27RA;CD74;I |
| KIF5B;HIF1A;NUMA1;ATRX;MNAT1;PAFAH1B1;CDC27;CHMP1A;PTTG1;DR1;SS18;SMAD3;SLC12A2;MAPK14;CCN |
| CLTB;AQP3;BID;FURIN;HYAL2;ATP6V1F;SOD2;SELENOW;NFKBIA;ACAA1;CDKN2B;PPP1R2;TST;ATP6V1C1;CDC5L; |

CDCP1;IRF6;ITGAV;BCL2L1;CD81;TGM2;CKAP4;FURIN;IFNGR1;TNFSF10;EMP1;CTSZ;HOPX;IGF2R;BMPR2;PNP;SY[illegible]
TSPAN1;PLAUR;MALL;SLPI;TOR1AIP2;EMP1;BIRC3;VWA5A;PRELID3B;CCL20;FUCA1;STRN;WNT7A;CROT;HBEGF;[illegible]
EIF4A2;FDPS;CDKN1A;SVIL;CRYAB;ABLIM1;GSN;CAMK2B;PLXNB2;BAG1;NAV2;SYNGR2;AGL;RIT1;TPD52L1;HBEG[illegible]
NDUFAF4;MPHOSPH10;UNG;FARSA
CA12;PLAUR;CDKN1A;ISG20;PGM1;TGM2;TPBG;TPST2;SULT2B1;PNRC1;GRHPR;PRDX5;BNIP3L;HDLBP;PGF;PAM[illegible]
PPARD;NUMB;TP53;NCSTN;ADAM17;DKK4;HDAC5;NCOR2;CUL1;HDAC11;MAML1;NKD1;FZD8;HEY1;DVL2;RBPJ
SRP14;SPCS1

TP5MF;ATP5PO;NDUFAB1;ETFA;ATP6V1D;NDUFA1;NDUFA4;ATP5F1B;ATP5PD;CYB5R3;ATP5MC1;COX6C;NDUFE
OMC;GFI1;STRBP;GAD1;TNNI3;DDX25;PHKG2;DBF4;HBZ;ACRBP;CCNA1;DPEP3;CFTR;TCP11;ELOVL3;PCSK4
GIC3;SOD1;YIPF6;ARFGEF1;VAMP3;SNAP23;ATP6V1H;ATP7A;RAB22A;RER1;ARCN1;MAPK1;RPS6KA3;ABCA1;LMA
IFNGR1;C3;SDHC;COX7B;ECH1;TOB1;UQCR10;UQCR11;TANK;NDUFS3;CMPK1;RNF11;NDUFB7;COQ9;UQCRC1;JA

DYNC1H1;ARHGAP29;PPP4R2;ARHGAP5;MARK4;SYNPO;SPTBN1;AKAP13;CD2AP;MID1;KIFAP3;ABR;ARFGEF1;RA

D88;UBE2L6;PNP;PSMA3;CASP7;PARP14;ST3GAL5;HELZ2;TAP1;RNF213;VAMP5;CCL2;IRF2;PELI1;CASP3;TRIM21;
UBB2A;CCL20;SLC2A6;MAP2K3;HBEGF;TAP1;F2RL1;EFNA1;SMAD3;RELB;CCL2;LITAF;ABCA1;TRIP10;TNIP2;BMP2

F1;POLR2K;POLB;NME4;CANT1;GTF2B;DAD1;COX17;POLR2I;TAF13;NUDT9;DDB1;NELFE;TP53;RAE1;TSG101;CST
2;CBR1;ACADVL;HCCS;ACAA2;ECI2;ADIPOR2;DLD;NSDHL;TP53INP2;VNN1;CPOX;PCBD1;BCKDHB;PDHB;ACOX1;F
;ADD1;MGMT;IGF2R;BNIP3L;BIRC3;FDXR;CASP7;CASP6;TAP1;SOD1;IGFBP6;GUCY2D;H1-0;ERBB2;F2R;CASP3;PS
7;APOC1;GNB2;APOBEC3F;USP14;SERPINA1;CTSO;MSRB1;IRF2;PCLO;USP8;PRCP;CASP3;LYN;KIF2A;PPP4C;PSEN

31;PLXNB2;MKNK2;CDKN2B;VWA5A;FDXR;ST14;FUCA1;ABAT;WWP1;TPD52L1;HBEGF;TAP1;KLK8;ABCC5;F2R;TR
31;KLK10;ATP2B4;TOB1;OPN3;BAG1;SLC26A2;CELSR2;TST;KLK11;DLG5;ST14;PTGES;EMP2;RABEP1;JAK1;TPD52L
MEM9B;CDC27;DCUN1D1;ENDOD1;GAPVD1;TFDP2;MAP2K3;ADIPOR1;UCP2;ACP5;FBXO7;DAAM1;UBAC1;H1-0;
LONP2;ABCC5;ACOX1;SLC25A17;CAT;PEX2;EHHADH;ABCD3;PEX13;ALDH9A1;CLN8;EPHX2;CDK7;SLC35B2;STS;A

;MARCHF6;ALAS1;MCCC2;HSD11B1;HSD17B2;ACP1;AKR1C2;IL1R1;VNN1;DCXR;BCAR1;CYB5A;SLC12A4;DDT;TK
;PDK2;CPEB3;OXT;TFF2;TEX15;NGB;NPHS1;COL2A1;CAMK1D;ARPP21;TNNI3;TFAP2B;GDNF;KLHDC8A;NTF3;IL5;

KA1;PPP2R1B;PIN1;MAPK1;MAPK10;RPS6KA3;ACACA;RIPK1;SMAD2;TBK1;GRB2;CLTC;PITX2;TIAM1;AKT1S1;CSN
JF144B;RHOG;CXCL8;HBEGF;SGMS2;IRAK2;CX3CL1;CCL2;IL1R1;TNFSF15;MARCO;P2RY2;ABCA1;LYN;PSEN1;RTP4
TX4;COL16A1;NECTIN2;GTF2F1;RRAS;CERCAM;CLDN4;RASA1;CX3CL1;SYK;LDLRAP1;GAMT;NF1;RHOF;MAPK14
AG1;RHOBTB3;SLC26A2;CELSR2;NAV2;PTGES;ABAT;ENDOD1;NADSYN1;TPD52L1;ZNF185;CANT1;SEC14L2;NRIP1;

M;CASP6;CITED2;SLC16A3;SOD1;DLD;NSDHL;RPE;EXT2;P4HA2;GYS1;PLOD1;MED24;KIF2A;CYB5A;MPI;CHPF2;HS

14;BCKDHB;SLC22A18;SCAF8;TFPI;GCNT1;NOTCH2;ATP2C1;MRPS31;PRDM2;PHF3;ARHGEF9;NEK7;AGGF1
IPC1;ABCD3;PEX11G;PEX13;PEX1;PNPLA8;ALDH9A1;PEX16;ABCA5;PEX7;EPHX2;TFCP2L1;SLC35B2;HACL1

LYN;ITGB2;CCND3;IFNAR2;GCNT1;RPL3L;TNF;HLA-DOB;FGR;STAT1;HLA-A;ACVR2A;TGFB1;IL6;MMP9;B2M;STAT4
T1;ATF5;CUL3;CUL4A;NOTCH2;RAD21;PURA;MEIS2;E2F3;TGFB1;ABL1;CUL5;CASP8AP2;MTF2;EFNA5;ARID4A
TAP1;MARK2;ALAS1;CASP3;CCNE1;GRINA;RRAD;LYN;SULT1A1;CCND3;NAT1;BMP2;CNP;MGAT1;BTG1;E2F5

NGR2;SWAP70;CASP3;CCNE1;SERPINB6;MYO1C;CCND3;ST3GAL4;BMP2;SMPDL3A;IKZF2;MUC1;SNX9;TIAM1;RA
F2RL1;SDCCAG8;CAB39L;KCNN4;HSD11B1;ZNF277;ATG10;WDR33;NIN;DNMBP;ITGB2;BMP2;TFPI;AVL9;GLRX;KLI
F;MYL6B;CNN3;GAA;CTF1;GABARAPL2;MYO1C;MEF2D;MYL2;SSPN;ATP6AP1;WWTR1;ITGB5;PDE4DIP;MEF2A;RE

;CASP6;CITED2;EFNA1;CCNG2;ATP7A;GAA;P4HA2;GYS1;ALDOC;WSB1;XPNPEP1;BTG1;GLRX;IDS;S100A4;PDGFB;

34;NDUFB3;ATP6V1F;ATP5F1A;ATP5F1C;COX6B1;OAT;SDHC;COX5A;COX7B;ECH1;SUCLA2;NDUFA8;VDAC3;MRPL

AN1;GOLGA4;AP2B1;CD63;SCAMP1;MON2;SSPN;TSG101;CLTC;SCRN1;CLCN3;USO1;VAMP4;COG2;KIF1B;STX7;D
AGN1;MDH2;TST;UQCRQ;ESRRA;UCP2;AGPAT3;ACAA2;SOD1;ADIPOR2;DLD;CCNG2;GHITM;MGST3;DBT;CHCHD1

ASA1;CDC42EP4;STAU1;SORBS2;STK38L;BCR;MAP3K11;NCK2;ARHGAP27;NF1;RHOF;RANBP9;ARHGEF3;NIN;RFC

;TRAFD1;CD74;ZNFX1;RIPK1;RTP4;SSPN;STAT3;IFNAR2;OGFR;BTG1;NMI;AUTS2;BPGM;TRIM26;IRF1;LATS2;TNFA
2;PER1;BTG1;IL23A;RELA;KLF4;NFKB2;TNF;RHOB;RNF19B;NFKBIE;IRF1;CXCL6;TNFAIP2;DNAJB4;NFAT5;BIRC2;SPS

TF3;POLL;TAF6;RNMT;TARBP2;ARL6IP1;GPX4;SURF1;ELL;GTF3C5;CLP1;NELFB;TAF12;MPC2;POM121;ERCC4;ADA

N1;PDP1;XPNPEP1;DOCK9;GRB2;CTSD;CASP10;PRDM4;CPM;IRF1;CTSV;PDGFB;RABIF;CTSL;FDX1;S100A12;IL6

AFD1;DCXR;RRAD;SERTAD3;CCND3;TP53;KIF13B;BMP2;BTG1;ELP1;KLF4;ANKRA2;CTSD;H2AJ;RNF19B;RB1;S100

MGST3;LRP10;SLC10A3;P4HA2;BTRC;CPOX;CCND3;ATG4A;CAT;ARHGEF12;HTATIP2;BLVRA;RBM38;ISCA1;CLCN3

ITIH3;SLC16A7;SLC5A5;CCNA1;COPZ2;SLC6A3;PDE6B;CACNG1;C5;CACNA1F;MAGIX;PDCD1;STAG3;BARD1

;ABHD2;P2RY2;MED13L;TSKU;MED24;ESRP2;MINDY1;UNC119;WWC1;AMFR;FRK;ARL3;RAPGEFL1;MUC1;KLF4 [illegible]4;ANO1;BPGM;EVI5;RABGAP1L;ADAM17;PRDM1;MMP10;MAFB;GUCY1A1;APOD;CSF2;TRIB2;MAP3K1;MMP9

L35;NDUFS2;HSD17B10;NDUFB5;UQCR10;BDH2;UQCR11;UQCRFS1;UQCRC2;COX5B;NDUFS3;NDUFC2;NDUFS4;[illegible]

L0;ABCA1;SULT1A1;COX8A;SLC66A3;BAZ2A;DDT;RAB34;GPAT4;CAVIN2;ACOX1;SSPN;PHYH;COQ5;IDH3A;CAT;S[illegible]

1;BCAR1;KIF3B;ARHGEF7;RICTOR;MYO9B;KLC1;RABGAP1;NEDD9;NOTCH2;ARHGEF12;SSH2;CSNK1D;ITSN1;CEP1

ACAA1;NDUFB7;MFN2;UQCRC1;MDH2;ATP6V1C1;ACADVL;UQCRQ;NDUFC1;TIMM17A;CASP7;OGDH;HCCS;NDU

JCLG1;ITSN1;CPT2;LTC4S;PTCD3;DLAT;HIBCH;ATL2;ARAF;DGAT1;SDHB;GPX4;SLC27A1;POR;CHUK;DHRS7B;PFKFI

FS8;ACAA2;NDUFB1;ALAS1;COX4I1;OXA1L;DLD;ATP6V0E1;MRPS11;MRPS15;COX17;TIMM8B;NDUFA6;ATP6V1H

I;NDUFA3;MGST3;OPA1;AFG3L2;MRPS30;COX8A;CYB5A;NDUFS1;PDP1;TOMM70;PDHB;PHYH;IDH3A;PDHA1;AT

P6AP1;SUCLG1;NDUFB8;COX15;ISCA1;MRPS12;MDH1;MTRF1;PDHX;ABCB7;DLAT;COX11;MTX2;ISCU;SDHB;GPX

[illegible]4;RHOT1;MRPL34;SURF1;POR;FDX1;TCIRG1;NDUFA2
